# ADAM protease inhibition overcomes resistance of breast cancer stem-like cells to γδ T cell immunotherapy

**DOI:** 10.1101/2020.07.17.207472

**Authors:** Indrani Dutta, Dylan Dieters-Castator, James W. Papatzimas, Anais Medina, Julia Schueler, Darren J. Derksen, Gilles Lajoie, Lynne-Marie Postovit, Gabrielle M. Siegers

## Abstract

Breast cancer stem cells (BCSC) are highly resistant to current therapies, and are responsible for metastatic burden and relapse. Gamma delta T cells (γδTc) are immunosurveillance cells with tremendous anti-tumoral activity, and a growing number of clinical trials have confirmed the safety of γδTc immunotherapy for various malignancies. Herein, we demonstrate that γδTc can kill BCSC, but to a lesser extent than non-cancer stem cells (NSC). Immune evasion was orchestrated by several mechanisms. The BCSC secretome rendered γδTc hypo-responsive by reducing proliferation, cytotoxicity and IFN-γ production, while increasing expression of co-inhibitory receptors on γδTc. BCSC and target cells surviving γδTc cytotoxicity had higher PD-L1 co-inhibitory ligand expression, and blocking PD-1 on γδTc significantly overcame BCSC resistance to γδTc killing. Fas/FasL signaling was dysfunctional in BCSC due to upregulation of the anti-apoptotic protein MCL-1, which could be partially overcome using dMCL1-2, an MCL-1 degrader. Moreover, the BCSC fraction shed higher levels of the NKG2D ligand MICA compared to NSC. Inhibiting MICA shedding using the ADAM inhibitor GW280264X overcame BCSC resistance to γδTc killing, rendering BCSC as sensitive to γδTc cytotoxicity as NSC. Collectively, our data unravel multiple mechanisms exploited by BCSC to evade γδTc killing, which may also come into play in BCSC resistance to other cytotoxic lymphocytes. Developing strategies to overcome this resistance will increase the efficacy of cancer immunotherapy and lead to improved outcomes for cancer patients.

**One Sentence Summary:** Breast cancer stem-like cells are resistant to γδ T cell targeting, which can be overcome by inhibiting ADAM proteases that facilitate MICA/B shedding.

## Introduction

Breast cancer is the most frequently diagnosed neoplasia and the leading cause of cancer-related mortality in females worldwide(*1*). This high rate of mortality is often attributed to drug resistance, recurrence and metastatic spread predominantly owing to breast cancer stem-like cells (BCSC), a subpopulation of cells which seem to drive tumour initiation and progression. The CD44^+^/CD24^-/low^ phenotype has been used extensively to successfully enrich for BCSC in both cell lines and tumour samples(*2*). The presence of CD44^+^/CD24^-/low^ cells correlates with the presence of distant metastases(*2*) and predicts poor prognosis in triple-negative breast cancer (TNBC)(*3*). Hence, by virtue of their high tumorigenic potential and drug resistance, BCSC have emerged as one of the most crucial therapeutic targets for breast cancer treatment.

Immunotherapy has tremendous promise for cancer treatment. While the majority of immunotherapies have focused on harnessing the potential of alpha beta T cells (αβTc), γδTc also play an important role in cancer immunity and have potent anti-tumoral activity(*4*). In γδTc, target recognition is primarily mediated by the γδ T-cell antigen receptor (TCR) and/or Natural Killer receptors, such as NKG2D(*5, 6*) that recognizes NKG2D ligands upregulated on transformed cells(*7*). Upon recognition, γδTc lyse tumours *via* perforin, granzymes, Fas Ligand (FasL) and TNF-related apoptosis-inducing ligand (TRAIL)(*6*). Activated γδTc also secrete high levels of pro-inflammatory cytokines such as interferon-γ (IFN-γ) that help stimulate and regulate the function of other immune cells(*8, 9*).

Several studies have reported that γδTc can target breast cancer cell *lines*(*10–14*) and phase I clinical trials have confirmed the safety of γδTc, either *via* adoptive transfer or *in vivo* stimulation with aminobisphosphonates such as Zoledronate(*15–17*). In one such study treating breast cancer patients with advanced-stage refractory disease, three of ten patients survived past 12 months and showed durable maintenance of robust γδTc numbers(*18*). For a recent review of γδTc in breast cancer, please see (*19*).

In this study, we used primary human γδTc derived and expanded from the blood of healthy donors(*20*) with the goal of determining whether γδTc can target BCSC. We demonstrate that while γδTc could kill BCSC, some were resistant to γδTc cytotoxicity; this was orchestrated by several immune evasion mechanisms, which we specifically targeted to overcome resistance and restore sensitivity to γδTc immunotherapy.

## Results

### Breast cancer stem-like cells are more resistant to γδ T cell targeting than non-stem-like cells

To address whether γδTc can kill BCSC as well as NSC, we used three cell line models and two different techniques to identify and isolate BCSC. One approach was to sort BCSC and NSC populations based on their expression of CD44 and CD24 on the cell surface. In a panel of human breast cancer cell lines, we found a range of BCSC percentages, from 0.5-88.6% (Fig. S1A-G). We chose to use SUM149 TNBC breast cancer cells as targets, since SUM149 comprise ~10% CD44^+^CD24^-^ BCSC on average (Fig. S1A), allowing us to isolate these cells, and CD44^+^CD24^+^ NSC, using fluorescence activated cell sorting (FACS). BCSC prevalence in SUM149 is much higher than that in TNBC-derived PDX401, or the luminal A cell lines MCF-7 and T47D (Fig. S1B-D). In contrast, the TNBC lines MDA-MB-231 and SUM159 were composed almost entirely of CD44^+^CD24^-^ cells; we did not use these (Fig. S1E-F), as it would be difficult to isolate sufficient BCSC and NSC for comparison. After sorting SUM149 into CD44^+^CD24^-^ BCSC and CD44^+^CD24^+^ NSC (Fig. S1I), we put them into culture, and found that the CD44^+^CD24^-^ population reverted back to the heterogeneity of the original population after 24h (Fig. S1I, J); hence, sorted cells were used directly in experiments. We performed serial dilution and sphere formation assays with sorted populations, confirming that SUM149 CD44^+^CD24^-^ cells have much higher sphere forming ability than CD44^+^CD24^+^ cells, which is a functional indicator of BCSC (Fig. S1K). We also confirmed that PDX401 mammospheres (3D) were enriched for CD44^+^CD24^-^ cells compared to adherent monolayer cultures (2D) (Fig. S1L), and used mammosphere assays to enrich for BCSC in the patient-derived xenograft (PDX) cell line PDX401 that did not have a distinct CD44^+^CD24^-^ population (Fig. S1B).

For experiments in this study, we derived primary γδTc cell cultures from twenty different healthy donors; culture subset compositions and purities are listed in Table S1. In cytotoxicity assays (n=3 independent experiments conducted with different donor-derived cultures), we determined that SUM149 BCSC are 10 to 30% less susceptible to γδTc killing than NSC (Fig. 1A, BCSC versus NSC p=0.0054, cumulative results from n=3 independent experiments shown in Fig. S1M; here and elsewhere experiments conducted with different donor cultures are indicated by different symbols in graphs. Statistical analyses employed and p-values are listed in Tables S2 and S3. The decreased susceptibility observed for SUM149 CD24^-^ cells also held true for SUM149-derived 3D versus 2D target cells (Fig. 1B, p=0.0261 at 10:1 effector:target (E:T) ratio, n=4 biological replicates shown in Fig. S1N). As a corollary, we wanted to determine if breast cancer cells resistant to γδTc targeting are enriched for stem-like cells; thus, we cocultured breast cancer cells with γδTc at 1:1 for 24h and assessed mammosphere forming capacity over two generations using the surviving resistant cells. Resistant cells exhibited significantly higher mammosphere forming ability compared to untreated cells, indicating that they were enriched for BCSC (Fig. 1C, p=0.0093, n=3 shown in Fig. S1O). As was the case for SUM149 cells, 3D PDX401 cells were also more resistant to γδTc cytotoxicity than 2D (Fig. 1D, n=4 shown in Fig. S1P), and 24h 1:1 co-cultures of γδTc with PDX401 yielded cells with a significantly greater capacity for mammosphere formation compared to target cells cultured alone (Fig. 1E, p<0.0001); data from two other biological replicates are shown in supplemental Fig. S1Q together with a representative image of mammospheres formed by PDX401 (Fig. S1R).

**Figure 1.**
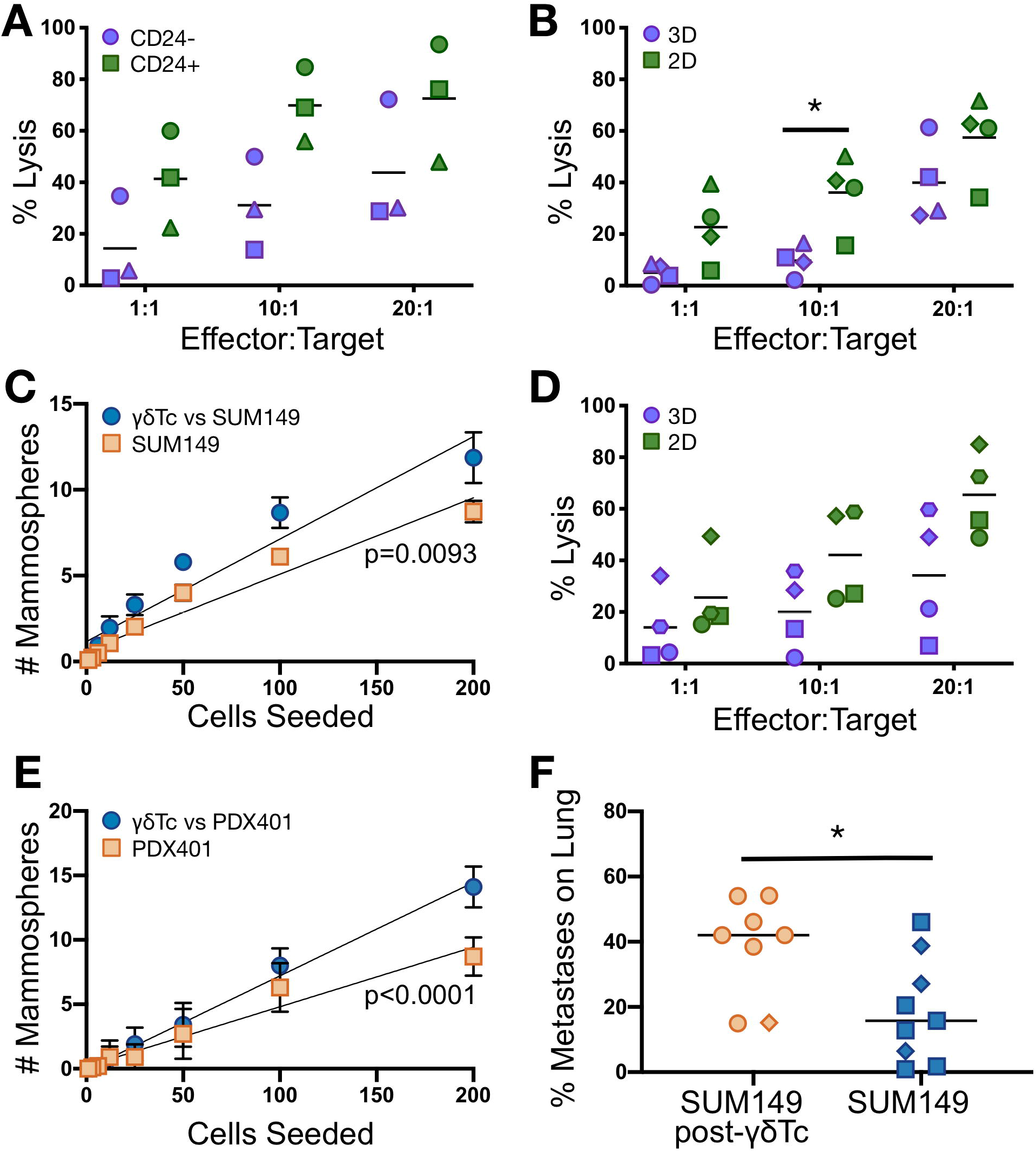
Breast cancer stem-like cells are resistant to γδ T cell targeting. **(A)** SUM149 cells were sorted and used in cytotoxicity assays (cumulative, n=3). **(B)** Second generation mammospheres (3D) and adherent (2D) SUM149 were dissociated, filtered into single cell suspensions and used as targets (cumulative, n=4) in Calcein AM cytotoxicity assays. **(C)** SUM149 cells were co-cultured with γδTc at 1:1 for 24 h, γδTc were removed, and mammosphere forming potential of targets determined over two generations (cumulative, n=3). **(D)** The patient derived xenograft cell line PDX 401 was used as a target in cytotoxicity assays as in (A) (n=4). **(E)** PDX401 were co-cultured with γδTc and mammospheres quantified as in (C); a representative example is shown. **(F)** Quantification of lung metastasis in mice injected with SUM149 alone or SUM149 treated with γδTc (n=17); mice injected with 90μl cell suspension are denoted with diamond shape, the rest who received 100μl are all denoted by circles. Data are presented as means in A, B and D, mean ± SEM in C and E, and medians in F. Statistical tests employed were: (A, B, D) Two-way ANOVA followed by Sidak’s post hoc test for multiple comparisons between groups; (C, E) simple linear regression; (F) Unpaired twotailed t-test, *p<0.05.

We used a similar approach to test the functionality of SUM149 resistant to γδTc *in vivo*, by determining their ability to develop experimental lung metastases in mice. Untreated SUM149 cells or those that had been treated with γδTc 1:1 for 24h were injected into the tail veins of NSG mice. After 18 weeks, mice were sacrificed and lungs injected with India ink to indicate metastases macroscopically, presenting as white spots on black-stained lungs (Fig. S1S and S1T). Calculation of the percent metastases, indicating the percentage of “white area” in the black lung, revealed that γδTc-treated SUM149 generated significantly more metastatic lesions (median white area 42%) compared to untreated SUM149 cells (median white area 15.7%), indicating the presence of a larger stem-like cell population in SUM149 cells treated with γδTc (Fig. 1F, p=0.0110).

### Gamma delta T cell degranulation and IFN-γ secretion are impaired in the presence of breast cancer stem-like cells

We performed degranulation assays by incubating γδTc with BCSC or NSC from SUM149 as well as PDX401 cells for 3h and measuring surface CD107a expression using flow cytometry; PMA/ionomycin-activated γδTc were our positive control. We found that γδTc incubated with BCSC degranulate significantly less compared to those incubated with NSC (SUM149 Fig. 2A, p=0.0361; PDX401 Fig. 2B; biological replicates Fig. S2A-G).

**Figure 2.**
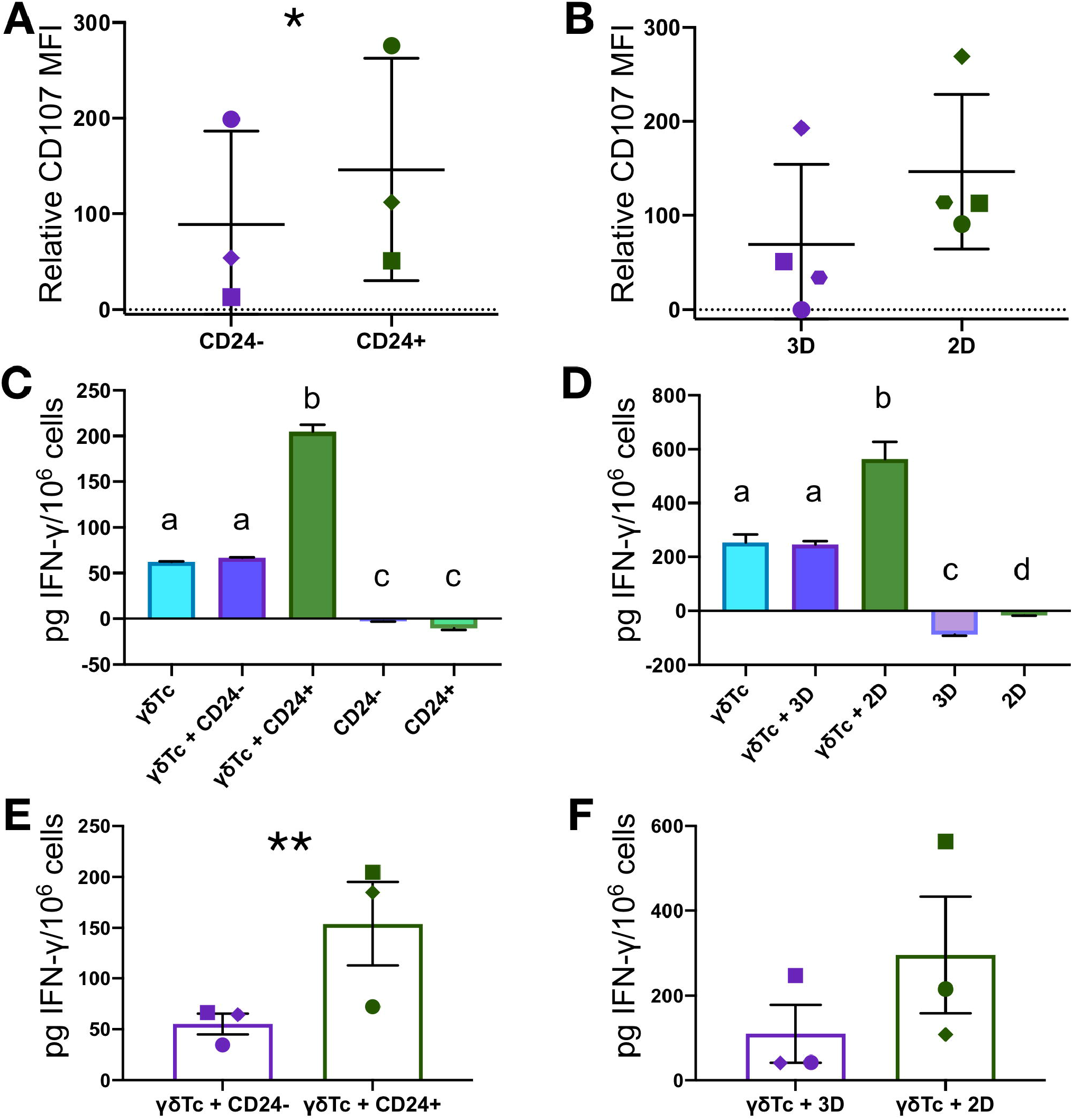
Gamma delta T cell degranulation and IFN-γ secretion are impaired in the presence of breast cancer stem-like cells. **(A)** Cumulative results of flow cytometric analysis of CD107a expression (degranulation) on γδ T cell cultures derived from different donors co-incubated with SUM149 (n=3); MFIs of CD107a were normalized over “no target” controls. **(B)** As in A but with PDX401 cells (n=4). **(C)** IFN-γ ELISA was performed on conditioned media (CM) from γδTc co-incubated with CD24^-^ or CD24^+^ SUM149 or **(D)** 3D or 2D PDX401 for 24h. **(E)** Cumulative results for IFN-γ secretion by γδTc incubated with CD24^-^ or CD24^+^ SUM149 including the experiment in C (n=3) and **(F)** 3D or 2D PDX401 targets including the experiment in D are shown (n=3). Data are presented as mean ± SD in A-D, and mean ± SEM in E and F. Statistical tests employed were: (A) Two-tailed paired t-test, (B, F) Wilcoxon two-tailed test. (C, D) One-way ANOVA followed by Tukey’s post hoc test for multiple comparisons between groups. a-b, p<0.0001 in C and a-b, p=0.0010 in D. (E) One-tailed ratio paired t-test. *p<0.05, **p<0.01.

We also measured IFN-γ secretion in conditioned medium (CM) from γδTc and co-cultures of γδTc with SUM149 BCSC or NSC (Fig. 2C, p<0.0001 for γδTc+CD24^+^ versus γδTc+CD24’) and PDX401 (Fig. 2D, p=0.0010 for γδTc+2D versus γδTc+3D). When co-incubated with BCSC, γδTc release significantly less IFN-γ than those co-cultured with NSC. Cumulative results also show significantly lower IFN-γ release by CD24^-^ SUM149 co-incubated γδTc (Fig. 2E, n=3, ratio paired one-tailed t-test, p=0.0071). This trend was also seen in cumulative analysis of experiments using 3D and 2D PDX401 (Fig. 2F, one-tailed Wilcoxon test, p=0.1250); however, in all individual biological replicates done with both cell lines, significant differences in IFN-γ secretion were observed between γδTc co-incubated with BCSC versus NSC (Fig. S2H-K).

### Breast cancer stem-like cells secrete factors that inhibit γδ T cell function

Impairment of γδTc function in the presence of BCSC prompted our investigation into the impact of secreted factors produced by BCSC on γδTc. To this end, we seeded equal numbers of SUM149 and PDX401 BCSC and NSC, and collected their CM after 24h. We labeled γδTc with CellTrace Violet (CTV) and incubated them with different CM for 24h and then replaced CM with fresh media. Six days later, flow cytometric assessment of CTV-labeled γδTc revealed greater CTV dilution by γδTc incubated with NSC CM compared to BCSC CM (Fig. 3A). Proliferation modelling indicated a greater proportion of cells in earlier proliferation divisions in BCSC CM compared to NSC CM (compare peaks 0 and 1 in Fig. 3B and C, respectively). This is depicted graphically in Fig. 3D. Compiled analyses of replication indices showed reduced proliferation of γδTc incubated with BCSC media compared to NSC media, but the difference was not significant when experiments were combined (Fig. 3E). Proliferation modelling outputs of other experiments are shown in supplemental Fig. S3A-F (n=3) as well as graphical representation of proliferation indices for combined experiments (Fig. S3G). To assess whether CM impacted γδTc viability, we incubated γδTc with different CM for 24, 48 and 72h and stained them with Zombie Aqua and AnnexinV; the viability of γδTc incubated with BCSC CM was slightly but not significantly reduced compared to γδTc incubated with NSC CM (Fig. S3H; Fig. S3I and J, cumulative results for SUM149 and PDX401, respectively).

**Figure 3.**
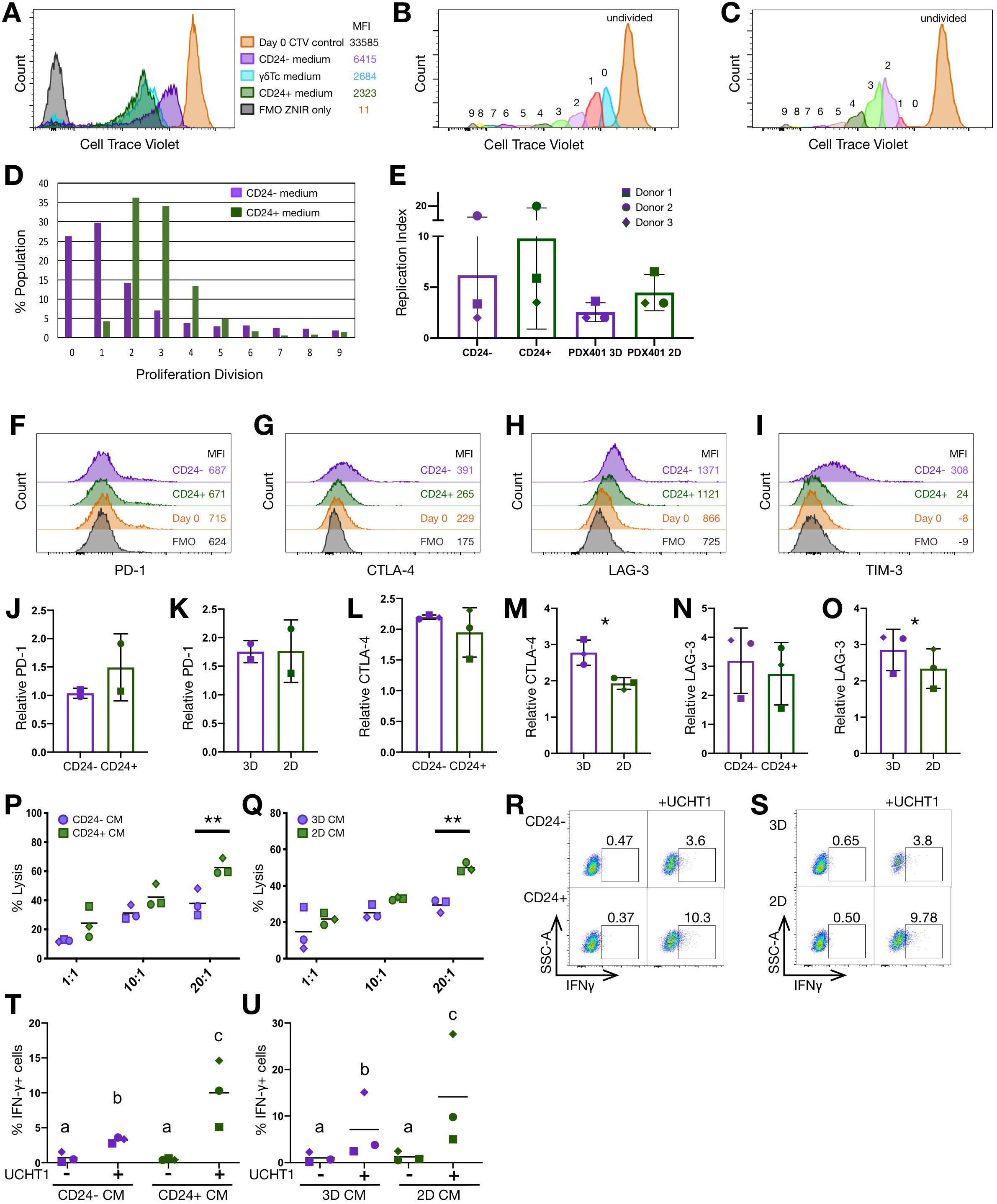
Breast cancer stem-like cells secrete factors that inhibit gamma delta T cell function. **(A)** To assess proliferation, γδTc were labelled with CellTrace Violet (CTV) and incubated with SUM149 CD24^+^ and CD24^-^ conditioned medium (CM) for 24h, washed, then further cultured in fresh medium. Six days later, the CTV signal was measured using flow cytometry. **(B)** Output from the FlowJo 10.5.3 Proliferation Modeling Tool, showing numbered peaks corresponding to divisions of the populations within the cultures that proliferated to different extents, for γδTc that had been incubated in CD24-CM, or **(C)** CD24^+^ CM. **(D)** Graphical representation of replication divisions undergone by γδTc in the proliferation experiment shown in A-C. **(E)** Graph of cumulative replication indices of γδTc treated with SUM149 or PDX401 CM including experiment in A-D (n=3). **(F-I)** γδTc were incubated with SUM149 CD24^+^ or CD24^-^ CM for 24h; representative histograms for expression of the indicated co-inhibitory receptors on γδTc seven days after incubation with CM are depicted. Ctrl were γδTc incubated with regular γδTc media instead of conditioned media. **(J-O)** Cumulative results including the experiment in F-I (n=2 J and K, n=3 for L-O; see symbols unique to biological replicates) for MFI normalized to corresponding FMOs for the indicated co-inhibitory receptors on γδTc incubated with SUM149 CD24^+^ and CD24^-^ CM or PDX401 2D and 3D CM. **(P)** After incubation with SUM149 CD24^+^ or CD24^-^ CM or **(Q)** PDX401 2D or 3D CM, γδTc were used to target the unsorted population of SUM149 and PDX401, respectively, in Calcein AM cytotoxicity assays at the indicated effector: target (E:T) ratios (n=3). **(R)** γδTc incubated with SUM149 CD24^-^ or CD24^+^ CM or **(S)** PDX 2D or 3D CM were treated with anti-CD3 antibody (UCHT1) for 6h and IFN-γ expression was determined *via* intracellular flow. **(T)** Cumulative results for % IFN-γ producing γδTc are depicted for γδTc incubated with SUM149 CD24^+^ and CD24^-^ CM including the experiment in R (n=3) and **(U)** PDX401 2D and 3D CM including the experiment in S (n=3). Data are presented as mean ± SD in E and J-O, and mean values are represented by lines in P, Q, T and U. Statistical tests employed were: (E) One-way ANOVA followed by Si dak’s post hoc test for multiple comparisons between groups; (L-O) One-tailed paired t-test, (P, Q) Two-way ANOVA followed by Sidak’s post hoc test, (T) One-tailed paired ratio t-test, a-b p=0.0482, b-c p=0.0255, a-c p=0.0107. (U) One-tailed paired ratio t-test, a-b p=0.0094, b-c p=0.0085, a-c p=0.0079. *p<0.05, **p<0.01.

We then incubated γδTc for 24h in different CM, replaced this media with complete medium and measured the status of inhibitory receptors on γδTc seven days after treatment. In a representative example from one experiment, PD-1 expression was similar (Fig. 3F), but CTLA-4, LAG-3 and TIM-3 were markedly increased on γδTc treated with SUM149 BCSC (Fig. 3G-I). Compiling data from different biological replicates, these differences did not always achieve significance; however, this is likely due to inherent variability among γδTc cells obtained from different donors. This was the case for PD-1 expression on γδTc treated with SUM149 CD24-versus CD24^+^ CM (Fig. 3J) as well as PDX401 3D versus 2D CM (Fig. 3K). CTLA-4 trended higher on γδTc treated with CD24-CM compared to CD24^+^ CM (Fig. 3L), and was significantly higher on 3D CM^-^ compared to 2D CM-treated γδTc (Fig. 3M, p=0.0371). The same was true for LAG-3 (Fig. 3N, p=0.0797 and Fig. 3O, p=0.0282 for SUM149 and PDX401, respectively). Histogram overlays of inhibitory receptor expression from different biological replicates are shown in supplemental Fig. S3K-N.

Since chronic upregulation of these receptors can render cells dysfunctional(*21*), we assessed the cytotoxic ability of γδTc that had been incubated with BCSC versus NSC CM. After incubation with BCSC CM, γδTc displayed only half of the cytotoxicity exhibited by CD24^+^ CM-treated γδTc at 20:1 against SUM149 (Fig. 3P, n=3, p=0.0043 at 20:1, Fig. S3O-Q). PDX 2D CM-treated γδTc were almost 1.5 times more cytotoxic than PDX 3D-CM treated γδTc at 20:1 (Fig. 3Q, n=3, p=0.0022; Fig. S3R-T).

We then stimulated γδTc with anti-CD3 antibody, incubated them overnight with BCSC or NSC CM and then performed intracellular staining for IFN-γ. Flow cytometric analysis revealed reduced percentages of IFN-γ-producing γδTc treated with CD24^-^ CM from SUM149 (Fig. 3R) and PDX401 3D CM compared to their respective NSC CM (Fig. 3S). Significant differences were observed in cumulative results (SUM149, Fig. 3T, n=3, p=0.0255, Fig. S3U-W; PDX401, Fig. 3U, p=0.0085, Fig. S3X,Y).

### The PD-1/PD-L1 pathway contributes to resistance of breast cancer stem-like cells to γδ T cell killing

Programmed cell death-ligand 1 (PD-L1) is often expressed by tumor cells, enabling binding to PD-1 on immune cells and thereby attenuating their responses(*22*). PD-L1 expression has been associated with poor prognosis in different solid tumors, including breast cancer(*23*). Hence, to compare expression of PD-L1 and PD-L2 - another ligand of PD-1 - on BCSC and NSC, we co-stained SUM149 with antibodies against CD24, CD44 and PD-L1 or PD-L1 and PD-L2 simultaneously and acquired samples by flow cytometry. Gating controls are shown in Fig. S4A and B. Unexpectedly, PD-L1 expression was slightly higher on SUM149 NSC compared to BCSC (Fig.4A and S4C). In fact, plotting PD-L1 versus CD24 shows a positive correlation (Fig. 4B). However, PDX401 3D cells expressed significantly higher PD-L1 than 2D adherent cells (Fig. 4C, p=0.0288; S4D-F), and PD-L2 trended higher in PDX401 3D compared to 2D cells (Fig. 4D, S4G-H).

**Figure 4.**
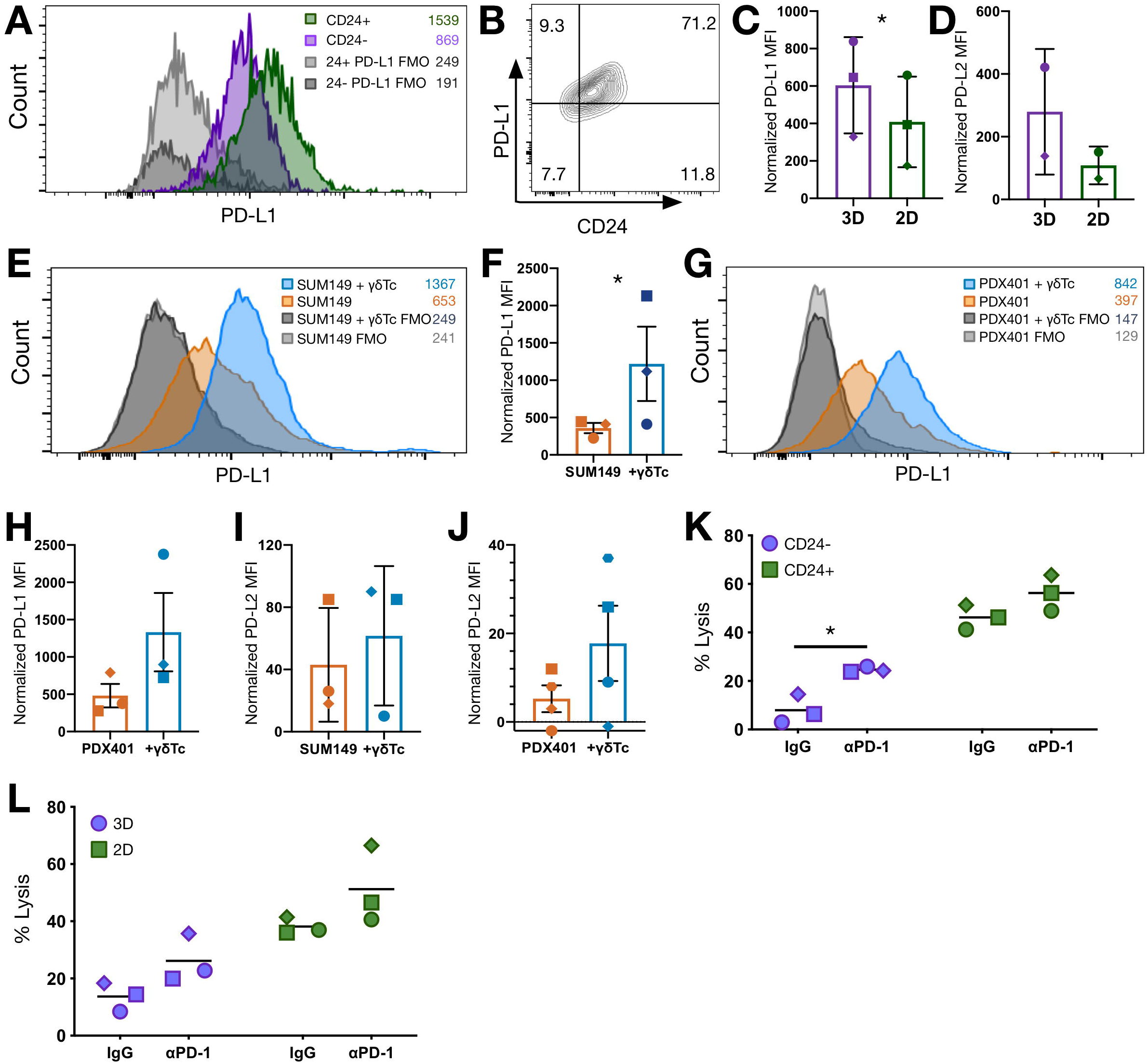
Inhibitory ligands are expressed on breast cancer stem-like cells resistant to γδ T cell killing. **(A)** PD-L1 expression was detected by flow cytometry on CD24^-^ and CD24^+^ SUM149 and (**B)** CD24 vs PD-L1 is plotted. **(C)** MFI (normalized to FMO) of PD-L1 (n=3) and **(D)** PD-L2 (n=2). on 2D and 3D PDX401 cells were measured **(E)** Target cells were co-incubated with γδTc at a 1:1 ratio overnight and surface expression of PD-L1 was detected. **(F)** Cumulative results of PD-L1 MFI normalized to FMO including the experiment in E are shown (n=3). **(G)** Experiment done as in E but with PDX401 3D and 2D cells. **(H)** Cumulative results of three biological replicates including G. **(I)** PD-L2 was compared between co-incubated targets and targets alone on SUM149 (n= 3) and **(J)** PDX401 cells (n=4). **(K)** γδTc were incubated with anti-PD-1 blocking antibody and then co-incubated with SUM149 CD24^-^ or CD24^+^ cells or **(L)** PDX401 3D or 2D cells at E: T 20:1 for 4h. Cumulative results are shown (n=3). Data are presented as mean ± SD in C, D, K and L, and as mean ± SEM in F and H-J. Mean values are represented by lines in K and L. Statistical tests employed were: (C, F, H-J) One-tailed paired ratio t-test (K, L) Two-way ANOVA followed by Tukey’s post hoc test. *p<0.05.

We then treated SUM149 with γδTc overnight, washed off γδTc and determined PD-L1 expression on the remaining resistant target cells. PD-L1 expression was indeed significantly elevated on target cells resistant to γδTc killing compared to target cells alone (Fig. 4E, 4F, n=3, p=0.0310, S4I-J). Similar results were obtained using PDX401 cells as targets (Fig. 4G, S4K-L); however, when cumulative results were analyzed, the difference was not significant (Fig. 4H, n=3, p=0.0939). PD-L2 expression also trended higher in γδTc-resistant targets (Fig.4I-J, S4M-S). To determine if blocking the PD-1/PD-L1 pathway increases sensitivity of BCSC to γδTc killing, we added monoclonal anti-PD-1 antibody in cytotoxicity assays against BCSC and NSC. Anti-PD1 treatment significantly increased γδTc lysis of SUM149 CD24^-^ cells but not to the level of CD24^+^ SUM149 lysis (Fig. 4K, n=3, p=0.0222, S4T-V, all p<0.05). Anti-PD-1 antibody also increased lysis of CD24^+^ cells, significantly so in one of three biological replicates (Fig. S4V, p=0.0045). While cumulative results trended toward increased lysis of PDX401 3D (Fig. 4L), of three biological replicates (Fig. S4W-Y), two showed a significant increase in lysis of 3D targets in the presence of anti-PD-1 antibody (Fig. S4X, p=0.0141; S4Y, p=0.0267) and in one case, 2D lysis was also significantly enhanced (Fig. S4Y, p=0.0033).

### The anti-apoptotic protein MCL-1 is upregulated in breast cancer stem-like cells

We next sought to characterize the contribution of receptors and ligands to the cytotoxicity of γδTc against BCSC and NSC using blocking antibodies. Blocking NKG2D, FasL or TRAIL on γδTc reduced cytotoxicity against SUM149, NKG2D blocking significantly so (Fig. S5A, p=0.0472 NKG2D versus IgG). In a separate experiment using γδTc from a different donor, blocking MICA/B or CD54 on SUM149 significantly reduced lysis (Fig. S5B, p=0.0286 and 0.0240, respectively). Significant reductions in lysis were also evident when blocking NKG2D and FasL in cytotoxicity assays against PDX401 (Fig. S5C, p=0.0117 and 0.0042, respectively). Blocking MICA/B or CD54 also lead to reduction of cytotoxicity against PDX401, but this did not achieve significance (Fig. S5C). We then did blocking assays with individual populations of SUM149 CD24^+^, CD24^-^, PDX401 2D and 3D (Fig. 5A-E) and found similar significant reductions in cytotoxicity against CD24^-^ SUM149 when sTRAIL, MICA/B or CD54 were blocked (Fig. 5A; p=0.0216, 0.0006, <0.0001, respectively); however, blocking NKG2D or FasL on γδTc did not significantly reduce cytotoxicity against CD24^-^ targets (Fig. 5A). In contrast, blocking any of these receptors in cytotoxicity assays against SUM149 CD24^+^ cells significantly reduced target cell lysis (Fig. 5B, all p<0.001 versus Ig). Cumulative results for log2 fold change in cytotoxicity on blocking FasL versus IgG control approached significance (Fig. 5C, n=4, p=0.0591). Cytotoxicity of γδTc against PDX401 3D remained unchanged despite blocking (Fig. 5D); however, γδTc cytotoxicity against PDX401 2D cells was reduced significantly on blocking any of the above-mentioned factors (Fig. 5E, all p<0.001). Blocking FasL versus IgG was significantly different between PDX401 2D and PDX401 3D cells (Fig. 5F, n=3, p<0.0037). All of the individual blocking cytotoxicity experiments are shown in supplemental Fig. S5D-J.

**Figure 5.**
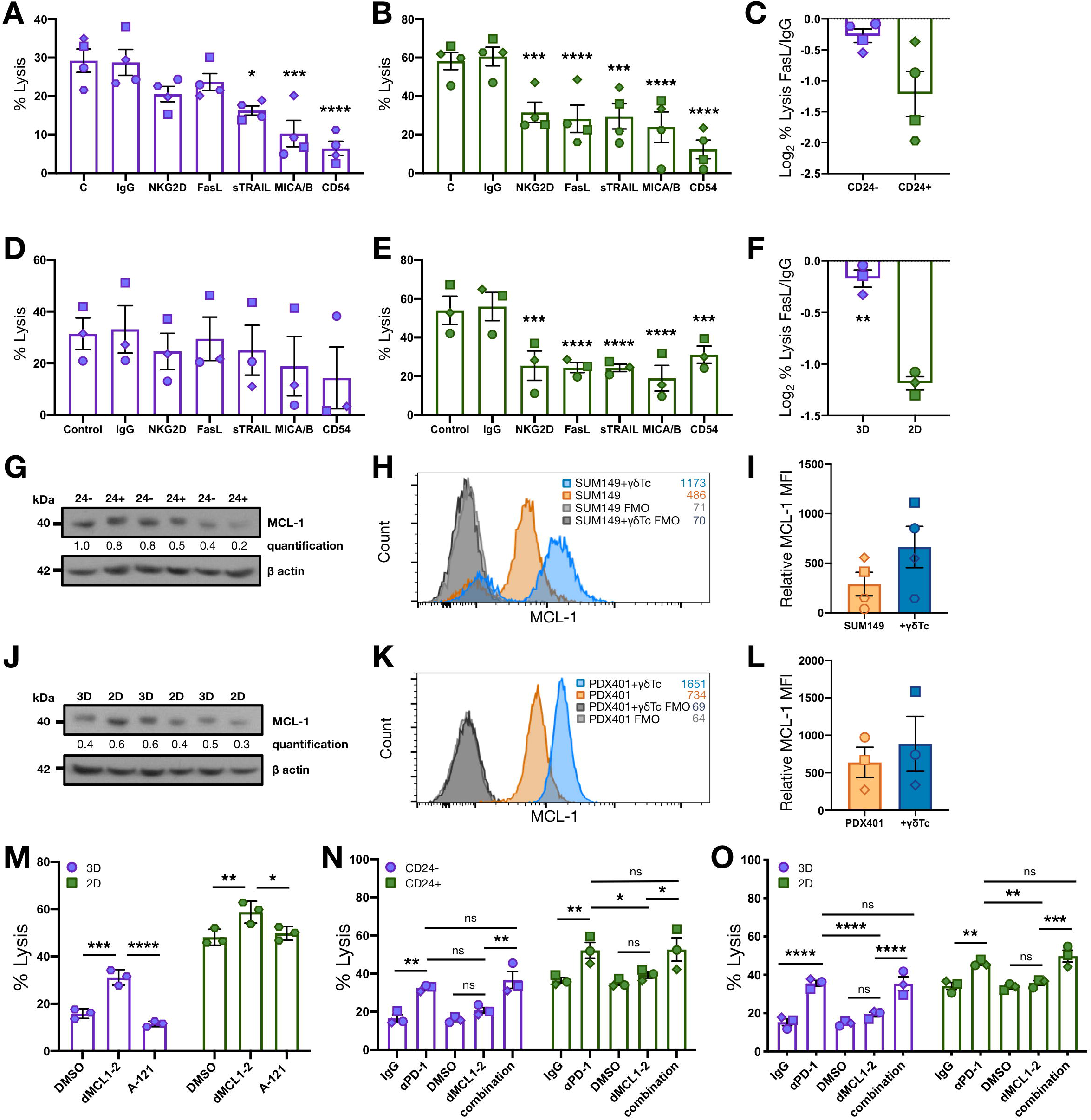
The Fas-FasL pathway is dysfunctional in breast cancer stem-like cells. **(A)** SUM149 were sorted into CD24^+^ and **(B)** CD24^-^ fractions, incubated with the indicated antibodies for 30 mins and then co-incubated with γδTc at E: T 20:1 for 4h. Cumulative results are shown (n=4). **(C)** The log2 fold decrease in target cell lysis upon FasL blocking compared to IgG control was calculated for the experiment shown in A and B (n=4). **(D)** Similarly, PDX401 3D and **(E)** 2D cells were incubated with antibodies followed by γδTc (n=3) and **(F)** the log2 fold decrease in target lysis upon FasL blocking compared to IgG control was calculated (n=3). **(G)** Expression of the anti-apoptotic protein MCL-1 was determined using western blot analysis of lysates from three sets of sorted SUM149 CD24^-^ and CD24^+^ cells as indicated. Molecular weight markers are shown on the left in kDa; β-actin loading controls are below corresponding panels and quantification of MCL-1 band intensities normalized to β-actin controls is shown. **(H)** SUM149 cells were subjected to a γδ T cell cytotoxicity assay at 1:1 (E: T) overnight. Target cells were then stained for intracellular MCL-1 protein and acquired *via* flow cytometry. **(I)** Cumulative results for SUM149 are shown including the experiment in H (n=3). **(J)** Experiments done as in G-I but with **(J-L)** PDX401 3D and 2D cells (n=3). **(M)** Sorted SUM149 CD24^+^ and CD24^-^ cells were co-incubated with γδTc at E: T 20:1 in a 4h Calcein AM cytotoxicity assay (n=4); where indicated 500nM dMCL1-2 or A-1210477 (A-121) were added to the coincubation, with the same volume of DMSO as a vehicle control. **(N)** Cumulative results of cytotoxicity assays done with dMCL1-2 or anti-PD-1 against SUM149 (n=3) or **(O)** PDX401 cells (n=3). Data are presented as mean ± SEM in all graphs except M, which shows mean ± SD. Statistical tests employed were: (A, B, D, E) RM One-way ANOVA followed by Bonferroni’s post hoc test; (C, F, I, L) One-tailed paired t-test; (M) Two-way ANOVA followed by Sidak’s post hoc test. (N-O) Two-way ANOVA followed by Tukey’s post hoc test. *p<0.05, **p<0.01, ***p<0.001, ****p<0.0001.

Given that the Fas-FasL pathway, which induces apoptosis in target cells, was dysregulated in BCSC of both cell lines, we explored surface expression of Fas(CD95) on CD24^-^ and CD24^+^ cells, finding them similar, as were expression of TRAIL R1 and TRAIL R2 (Fig. S5K). This was also true for PDX401 3D and 2D cells (Fig. S5L). Since CSC can upregulate anti-apoptotic proteins to resist induction of apoptosis(*24*), we determined the expression of several different anti-apoptotic proteins *via* western blotting (Fig. S5M) and found that MCL-1 was upregulated in CD24^-^ cells (Fig. 5G) as was Bcl-XL in two out of three replicates (Fig. S5M, left panels). Flow cytometric analysis revealed that surface MCL-1 expression was also higher in SUM 149 cells resistant to γδTc killing compared to targets alone (Fig. 5H; 5I, n=4, p=0.09; S5N). PDX401 3D also expressed higher MCL-1 (Fig. 5J), Bcl-XL and Survivin levels than 2D cells (Fig. S5M, right panels). MCL-1 was also increased in resistant compared to untreated PDX401 cells (Fig. 5K; 5L, n=3; S5O).

To target the overexpression of MCL-1 in resistant BCSC, we employed a novel MCL-1 protein degrader, dMCL1-2, and the failed inhibitor A-1210477 (A-121), from which dMCL1-2 was derived, as a control(*25*). Degradation of MCL-1 on addition of dMCL1-2 was confirmed in both SUM149 and PDX401 cell lines *via* western blotting (Fig. S5P-Q).

Addition of dMCL1-2 in a cytotoxicity assay with γδTc derived from one particular donor showed significant upregulation of cell lysis of PDX 2D cells (Fig. 5M, p=0.0073) and 3D cells even more so compared to the vehicle control DMSO (Fig. 5M, p=0.0003). Assays done with two other γδTc cultures derived from this donor yielded significant increases in cytotoxicity against CD24^-^ SUM149 (Fig. S5R, p=0.0254, and S5S, p=0.0239); however, dMCL1-2 had no effect in assays done with γδTc derived from five other donors (Fig. S5T-X). Using CD24^-^ and CD24^+^ SUM149 as targets, we performed cytotoxicity assays using a combination of the anti-PD1 antibody and dMCL1-2; however, we did not see any synergistic effect in cumulative analysis of experiments using BCSC and NSC SUM149 (Fig. 5N, n=3; S5V-X left panels) or PDX401 targets (Fig. 5O, n=3; S5V-X right panels). Since MCL-1 is an anti-apoptotic protein, we also tested the viability of γδTc after 4hr treatment with the degrader dMCL1-2, mimicking the conditions of the cytotoxicity assays, and saw no impact on γδTc viability (Fig. S5Y).

### Breast cancer stem-like cells display lower expression of MICA/B on their surface

We then turned our attention to the expression of the NKG2D ligands, MICA/B and UL16 binding proteins (ULBP)-2-6, on BCSC and NSC (Fig. 6, S6). Histogram overlays from a representative experiment show greater expression of these NKG2D ligands on CD24^+^ NSC compared to CD24^-^ BCSC (Fig. 6A, S6A). After normalization of median fluorescence intensity (MFI) values and compiling data from multiple experiments, we observed that MICA/B, ULBP-3 and ULBP-4 expression were significantly lower in CD24^-^ SUM149 cells compared to CD24^+^ SUM149 cells (Fig. 6B, n=3, p=0.0428, 0.0467, and p=0.0430 respectively; S6B-E), while ULBP-2,5,6 was also down-regulated in SUM149 CD24^-^ compared to CD24^+^, nearly reaching significance (Fig. 6B, p=0.0518; S6B-E). MICA/B surface expression was also significantly lower on PDX 3D cells compared to 2D (Fig.6C and D left panels, p=0.0088; S6F,G), but expression of the other NKG2D ligands was variable (Fig. 6C and D, S6F,G).

**Figure 6.**
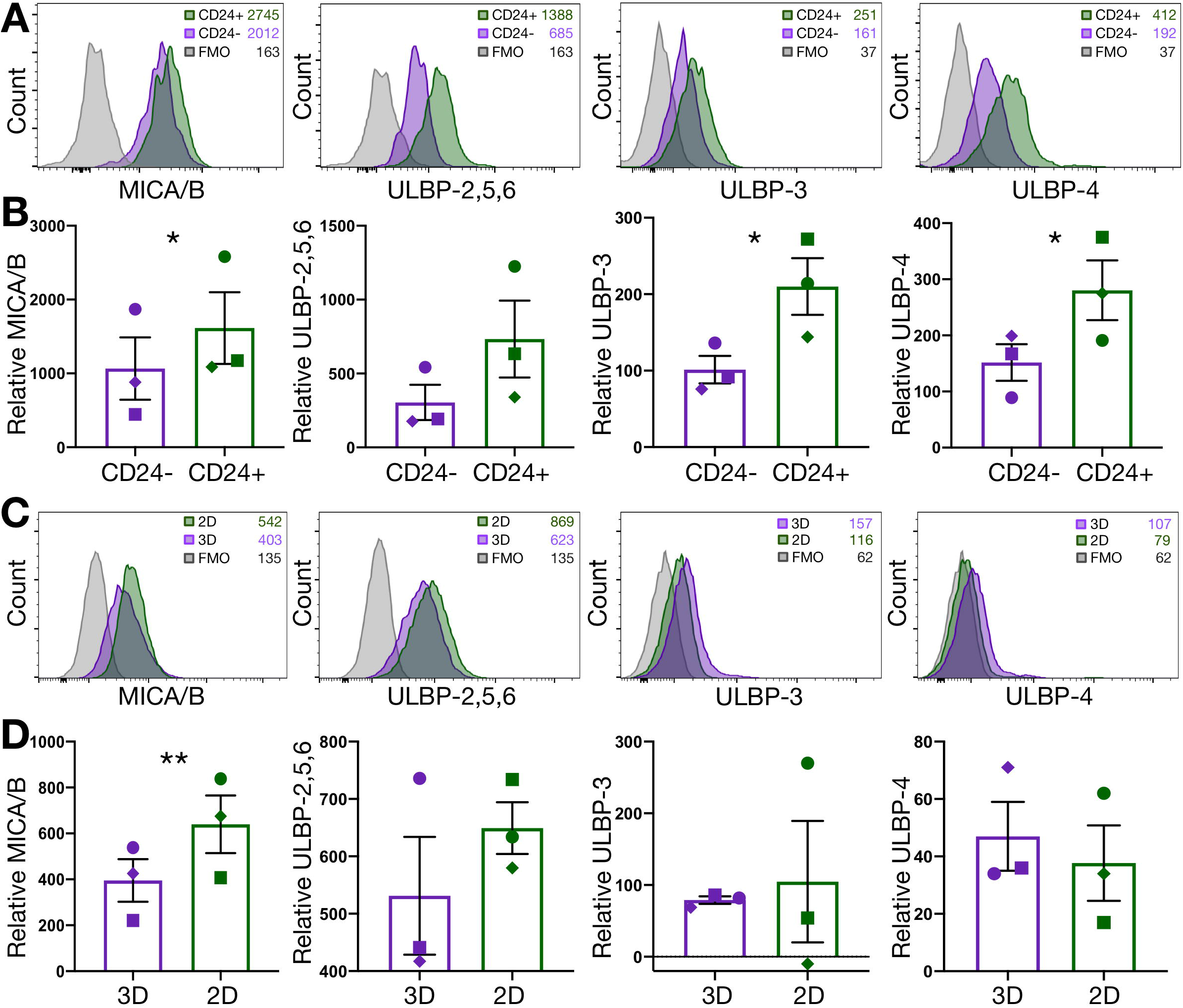
Breast cancer stem-like cells have lower surface expression of MICA/B. **(A)** NKG2D ligand expression was determined on SUM149 CD24^-^ and CD24^+^ cells by flow cytometry and **(B)** cumulative results for normalized values including those from the experiment in A are shown (n=3). **(C)** NKG2D ligands on 2D and 3D PDX401 cells and **(D)** cumulative data comprising normalized values including those from the experiment in C are shown (n=3). Data in graphs are presented as mean ± SEM. For statistics shown in B and D, one-tailed paired t-tests were employed. *p<0.05, **p<0.01.

### Breast cancer stem-like cells shed higher levels of MICA conferring resistance to γδ T cell killing that can be overcome with ADAM inhibition

One mechanism regulating NKG2D ligands on the surface of cancer cells is NKG2D ligand shedding, which can occur by proteolytic activity or exosomes(*26*). Strikingly, proteomic analysis of CM from BCSC and NSC revealed that MICA was one of the top upregulated proteins secreted by both SUM149 and PDX401 BCSC. A heat map and scatter plot depicting the top differentially secreted proteins between the BCSC and NSC, common between both cells lines, is shown (Fig. 7A-B). Increased MICA in BCSC compared to NSC supernatants from both SUM149 and PDX401 cell lines was also confirmed by ELISAs (Fig.7C and D, n=4, p=0.0033 and p=0.0038, respectively; Fig. S7E-F). We then targeted “a disintegrin and metalloproteinase” (ADAM)10 and 17 that are known to be involved in tumor-associated proteolytic release of soluble MICA(*27, 28*). Proteomic analysis showed that ADAM 10 and 17 were also upregulated in BCSC CM of both cell lines (Fig 7B, S7C-D). We then used GW280264X (ADAMi), an inhibitor specific for both ADAM10 and ADAM17, and found that the surface level of MICA/B, as measured by flow cytometry, increased significantly on both SUM149 (Fig.7E top panel and 7F, n=3, p=0.0008; Fig. S7G) and PDX401 (Fig.7E bottom panel and 7G, n=3, p=0.0011; Fig. S7H). In addition, MICA shedding, as evidenced by soluble MICA levels measured by ELISA, was also significantly abrogated in both cell lines upon addition of ADAMi (Fig.7H, n=3, p=0.0332, S7I; and 7I, n=3, p=0.0058, S7J). Adding this inhibitor to cytotoxicity assays lead to significantly enhanced γδTc cytotoxicity against both BCSC and NSC SUM149 (Fig.7J, CD24-DMSO versus ADAMi p<0.0001; CD24^+^ DMSO versus ADAMi, p=0.0007; S7K-M) and PDX401 (Fig.7K, 3D DMSO versus ADAMi, p<0.0001; 2D DMSO versus ADAMi, p<0.0001; S7N-P). Notably, combining ADAMi with γδTc brought the levels of BCSC lysis to those of untreated NSC (Fig.7J, CD24-ADAMi versus CD24^+^ DMSO, p=0.6781; S7K-M; and Fig.7K 3D ADAMi versus 2D DMSO, p=0.2139, S7N-P), overcoming the resistance of BCSC to γδTc cytotoxicity.

**Figure 7.**
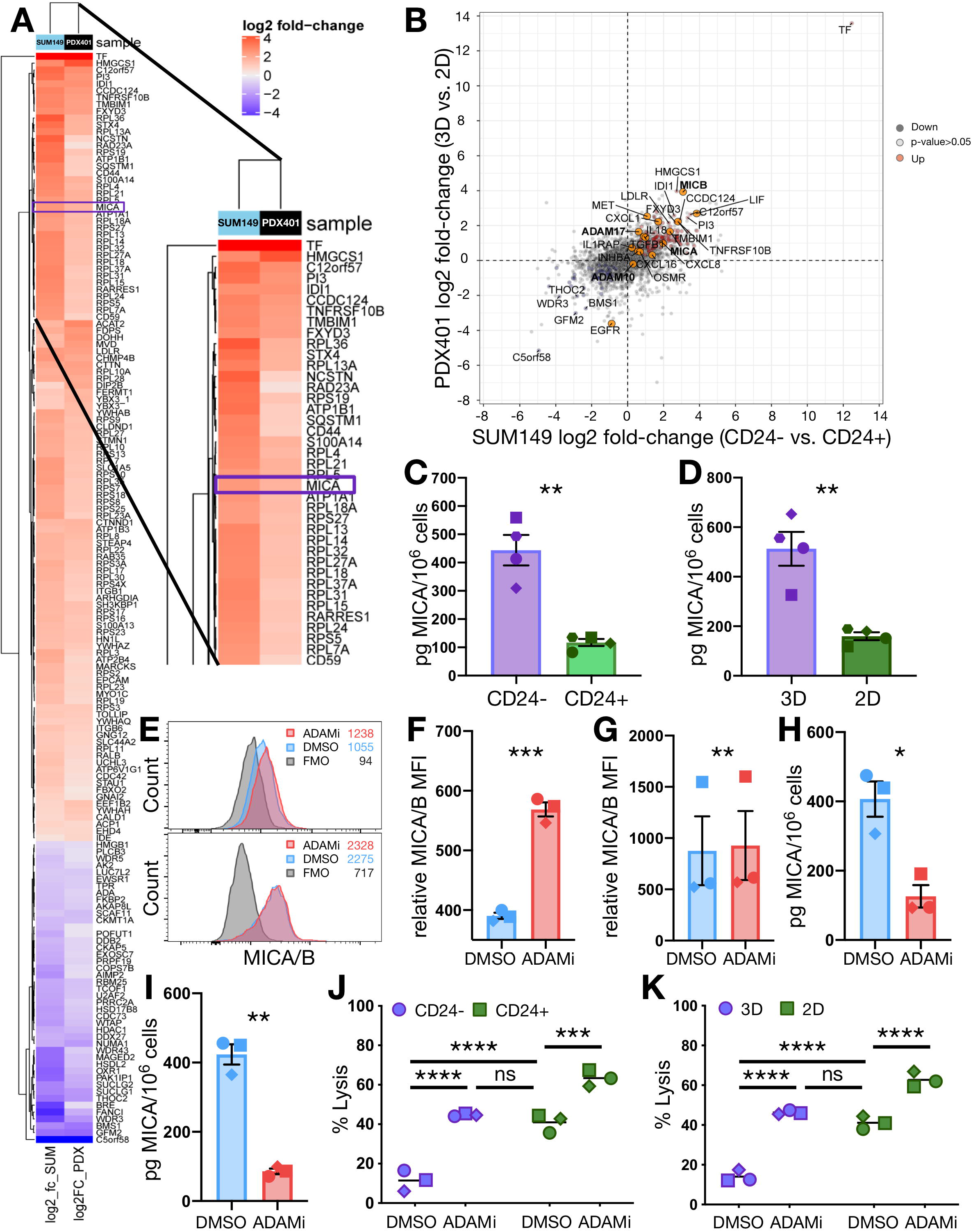
Breast cancer stem-like cells shed MICA rendering them resistant to γδ T cell killing, which can be reversed by inhibiting ADAM proteases. **(A)** Secretome analysis of SUM149 and PDX BCSC was performed with mass spectrometry; the log2 fold changes of the most differentially secreted proteins are depicted as a heat map; a magnified version of the top 39 proteins, with MICA boxed in purple is provided. **(B)** Scatter plot depiction of secretome analysis shown in A. **(C)** ELISA quantification of MICA shed by sorted CD24-versus CD24+ SUM149 cells (n=4) and **(D)** 2D and 3D PDX401 (n=4). **(E)** Targets were treated with the ADAM protease inhibitor GW280264X (ADAMi) for 4h and MICA/B surface expression was determined *via* flow cytometry. Representative overlays of SUM149 (top) and PDX401 cells (bottom) are shown and **(F)** cumulative results for MICA MFI (normalized to FMO controls) on SUM149 (n=3) and **(G)** PDX401 are shown (n=3). **(H)** Soluble MICA secreted by SUM149 in the presence of ADAMi or vehicle control was measured by ELISA (n=3) and **(I)** the same was done with supernatants from PDX401 cells treated in the same way (n=3). (**J**) SUM149 CD24^-^ and CD24^+^ cells (n=3) or **(K)** PDX401 3D and 2D cells (n=3) were incubated with ADAMi and then co-incubated with γδTc at E:T 20:1 for 4h. Data are presented as mean ± SEM in C, D and F-I. Mean values are represented by lines in J and K. Statistical tests employed were: (C, D, F-I) One-tailed paired t-test; (J and K) Two-way ANOVA followed by Tukey’s post hoc test. *p<0.05, **p<0.01, ***p<0.001, ****p<0.0001.

We also repeated many of these experiments with a luminal A breast cancer cell line, MCF-7, and found the results similar to those in experiments done with SUM149 and PDX401 cells. MCF-7 cells do not have a distinct CD44^+^CD24^-^ population (Fig. S1C,G) and so we generated MCF-7 mammospheres to enrich for CD44^+^CD24^-^ BCSC (Fig. S8A). MCF-7 3D cells were more resistant to γδTc cytotoxicity than their 2D counterparts (Fig. S8B, n=4, p=0.0239 at 20:1 E:T), and cells resistant to γδTc killing had higher sphere forming ability (Fig.S8C, n=3, p=0.0016). In the presence of MCF-7 3D compared to 2D cells, γδTc degranulated less (Fig. S8G, n=3), and produced significantly less IFN-γ (Fig. SH, n=4, p=0.0262). Proliferation of γδTc was also lower when incubated with MCF-7 3D CM compared to 2D CM (Fig. S8I, n=3, proliferation index, p=0.0216, and replication index, p=0.0020). There were no differences in viability (Fig. S8J, n=4) and expression of co-inhibitory receptors on γδTc incubated with 3D or 2D CM (PD-1 Fig. S8K, n=2; CTLA-4 S8L, n=3; LAG-3 S8M, n=3), but γδTc incubated with 3D CM released less IFN-γ (Fig. S8N, n=3, p=0.0452). MCF-7 3D cells had higher PD-L1 and PD-L2 expression on their surface compared to 2D cells (Fig. S8O, n=2), yet co-incubation with γδTc did not increase PD-L1 expression on MCF-7 (Fig. S8P, n=1). Blocking receptors and performing cytotoxicity assays yielded varying results: blocking NKG2D or FasL did not reduce cytotoxicity of γδTc against 3D cells (Fig. S8Q left panel, n=3), but significantly reduced cytotoxicity against 2D MCF-7 (Fig. S8Q right panel, n=3, p≤0.0005 for all). The Log2 fold change in cytotoxicity on blocking FasL versus IgG was lower on MCF-7 3D compared to 2D cells, but not significantly so (Fig. S8R, n=3, p=0.1614). In line with this, while MCF-7 3D cells expressed higher levels of MCL-1 compared to 2D cells (Fig. S8S), unlike in SUM149 and PDX401, there was no difference in MCL-1 expression before and after treatment with γδTc (Fig. S8T). Yet, similar to SUM149 and PDX401 BCSC, MCF-7 3D cells expressed less surface MICA/B compared to 2D MCF-7 (Fig. S8U), while ULBP expression was variable (Fig. S8V-X).

## Discussion

Gamma delta T cells are a promising alternative to αβTc for T cell immunotherapy, offering several notable advantages: Clinical trials have established their safety; their activation is typically HLA-independent thus they do not cause graft-versus-host disease and are not impacted by peptide-MHC antigen loss, which diminishes the effectiveness of αβTc therapy; and their kinetics in response to stress signals are faster(*6*).

Clinical trials with γδTc have thus far been early phase, designed to test safety and conducted in standard-treatment-refractory advanced stage cancer patients; unsurprisingly in this setting, they have shown variable efficacy(*29*), but later phase clinical trials are still lacking. Despite this, discussion regarding lack of patient response to γδTc therapy has included the possibility of activation-induced anergy or unresponsive cells(*17*), and it is prudent to investigate such potential obstacles. Several immune evasion tactics are exploited by cancer cells to overcome recognition and attack, and the immunosuppressive nature of the TME is a major hurdle facing successful immunotherapy.

Few thus far have investigated CSC targeting by γδTc, and those that have - in colon cancer, ovarian and neuroblastoma(*30–32*) - did not compare their cytotoxicity against CSCs and NSC in parallel to get a complete picture as we have done. However, it was recently shown that pretreatment of tumor cells with Zoledronate, and subsequent treatment with primary autologous γδTc and CD8^+^ T cells in combination was more effective than either cell type alone(*33*). These studies used aminobisphosphonates such as Zoledronate that selectively expand the Vδ2 γδTc subset; however, several pre-clinical cancer studies have suggested that Vδ1 may be superior to Vδ2 cells in terms of both cytotoxicity and durability(*34, 35*). Vδ1 cells are more resistant to activation-induced cell death (AICD)(*36*), which has previously posed significant problems in clinical trials due to chronic stimulation of Vδ2 cells(*17*). Hence, both subsets should be taken into consideration for the development of next-generation γδTc-based immunotherapies. Here, we used polyclonal primary human γδTc, and many of these cultures contained significant proportions of Vδ1 γδTc, which likely contributed to lysis of BCSC and NSC (Table S1).

We found that while γδTc can kill BCSC, BCSC employed multiple strategies to evade γδTc attack. Both CD44^+^CD24^-^ and mammosphere-enriched 3D BCSC were less susceptible to γδTc cytotoxicity than CD44^+^CD24^+^or adherent 2D NSC (Fig. 1A,B,D; 4K,L; 5A,B,D,E,M-O; 7J,K). *In vivo*, mice injected with SUM149 cells previously treated with γδTc exhibited significantly more metastatic lesions compared to mice injected with untreated SUM149 (Fig. 1F). Since BCSC have a greater ability to form metastatic lesions(*2*), this result suggests enrichment of BCSC during co-incubation with γδTc, which we saw in our *in vitro* cytotoxicity assays (Fig. 1C,E). Activated γδTc perform their cytotoxic function by degranulation to release perforins and granzymes, and they also produce IFN-γ (*6, 8*). These functions were inhibited in the presence of BCSC from both SUM149 and PDX401 TNBC-derived cell lines, as well as the luminal A cell line, MCF-7 (Fig. 2A-F, S8B). Interestingly, co-incubation with PDX401 lead to two- to three times more IFN-γ production by γδTc compared to incubation with SUM149 or MCF-7 cells, suggesting that PDX401 cells activate γδTc to a greater extent (Fig. 2E-F, S8G-H). In addition to decreasing degranulation and IFN-γ production, BCSC CM rendered γδTc dysfunctional by chronically increasing their expression of co-inhibitory receptors (Fig. 3) while reducing their proliferation and cytotoxicity, similar to a study in which incubation with CSC CM from colorectal cancer patients significantly reduced proliferation and IFN-γ production by γδTc (*37*).

Since checkpoint inhibition has become a widely used and effective tool in the clinic(*38*), we tested the differential expression of PD-L1 and found it not only to be higher on 3D compared to 2D PDX401 cells (Fig. 4C), but also higher on target cells resistant to γδTc (Fig. 4E-H). Hence, it is possible that ligands on BCSC, and resistant target cells in general, engage PD-1 on γδTc to suppress γδTc activity. It is important to note, however, that PD-1 is also a marker of T cell activation; PD-1 is transiently upregulated in activated T cells, but chronically expressed on dysfunctional T cells (*22*). We did not see differences in PD-1 expression on γδTc upon exposure to BCSC CM (Fig. 3F,J), yet blocking the PD-1/PD-L1 pathway was able to partly surmount resistance to γδTc cytotoxicity, specifically in BCSC (Fig. 4K,L). Our results are also in agreement with studies showing enhanced γδTc cytotoxicity against various cancers such as lymphoma, multiple myeloma, different types of leukemia, B cell lymphoma and neuroblastoma upon blocking PD-1/PD-L1 pathway(*39–42*).

Blocking assays demonstrated that Fas/FasL signaling, an important pathway that γδTc exploit to trigger apoptosis in their targets, is dysfunctional in BCSC (Fig. 5A,C,D,F). In line with this, the anti-apoptotic protein MCL-1 was overexpressed in BCSC in general (Fig.5G,J), and also in cells resistant to γδTc targeting (Fig.5H,I,K,L). MCL-1 overexpression renders cancer cells resistant to several drugs(*43*) and is associated with poor outcome in breast cancer(*44*). While we verified that the failed MCL-1 degrader A-121 leads to accumulation of this protein and that the novel dMCL1-2 degrader indeed reduced MCL-1 levels (Fig. S5P,Q)(*25*), the success of the degrader in sensitizing BCSC to γδTc targeting (Fig. 5M-O, Fig. S5R-X) appeared to be donor dependent: experiments in Fig. 5M, Fig. S5R and S5S were done with different γδTc cultures derived from the same donor. Interestingly, this was the same donor whose cells were used in the only experiment in which MCL-1 expression was not upregulated after treatment with γδTc (Fig. S5O, top panel). If this was also the case in cytotoxicity assays, and MCL-1 was not upregulated in response to this donor’s γδTc, it could explain why dMCL1-2 worked so well in those particular experiments. We would likely need to use greater quantities of dMCL1-2 to elicit the same effect in experiments in which MCL-1 was upregulated in the presence of γδTc. This also raises the intriguing question as to why γδTc from this particular donor did not incite MCL-1 upregulation while those from other donors did. Apart from various intrinsic factors, MCL-1 expression can be modulated by several extrinsic factors like IL-6 and IFN-γ (*45, 46*), which were perhaps secreted differentially by this specific donor’s γδTc and played a role in regulating MCL-1 in targets.

Several soluble factors are released by tumour cells or stromal cells that can exert suppressive effects on immune cells (*47*). Mass spectrometry analysis indicating global differences in protein expression between BCSC and NSC secretomes identified potential “hits” that could translate into therapeutic targets (Tables S5 and S6). KEGG analysis identified cytokines and chemokines such as TGF-β1/2, IL-18, CXCL16 and CXCL8 upregulated in BCSC from both cell lines (Fig. 7B, S7C,D). IL-18 was upregulated in BCSC, but its impact on γδTc is unclear: Vδ2 expansion by Zoledronate and IL-2 can be strongly promoted by addition of IL-18(*48*); yet, in contrast, a study on malaria suggested that IL-18, in synergy with IL-12, can induce expression of the co-inhibitory receptor TIM-3 on the surface of γδTc, rendering them functionally impaired(*49*). In our study, incubation with CM from SUM149 CD24^-^ also lead to TIM-3 upregulation (Fig.3I). Serotransferrin (TF), an abundant protein involved in iron metabolism and associated with poor prognosis in Cholangiocarcicoma (*50*), is also dramatically over-expressed in BCSCs of both SUM149 and PDX401 (Fig. 7B).

Amongst several others, TGF-β1 was a prominent soluble factor upregulated in both SUM149 and PDX401 BCSC, and its impact on γδTc is unclear. One study reported that TGF-β1 can enhance cytotoxic activity of Vδ2 T cell (*10*), while another demonstrated that TGF-β1 and IL-15 can polarize Vγ9Vδ2 cells (Vδ2) to FOXP3+ regulatory γδTc that suppress proliferation of activated PBMC including αβTc (*51*). Yet another study suggested that Vδ2 can be polarized into γδT17 cells (producing IL-17) in the presence of TGF-β1, IL-6 and IL-1 β (*52*). These γδT17 cells are associated with the immune-suppressive TME, which further aids development of cancer(*53*). Using gain- and loss-of-function models, we recently found that the TGF-β family member NODAL inversely correlated with MICA/B expression on breast cancer cells, which impacted the ability of γδTc to target them(*54*). Compared to NSC, BCSC consistently displayed lower expression of cell surface MICA/B (Fig. 6A-D, S6), likely rendering BCSC invisible to γδTc, similar to what we observed in our study of NODAL and γδTc(*54*).

Soluble MICA shed from the cell surface can downregulate NKG2D on T and NK cells(*55*), and elevated levels of circulating soluble MICA have been associated with poor clinical prognosis and metastasis in multiple cancer types, including breast cancer(*56, 57*). Proteomic analysis followed by ELISA verification demonstrated significantly enhanced MICA shedding by BCSC compared to NSC (Fig. 7C-D). Proteomic analysis also revealed increased expression of ADAM10 and 17 in the BCSC CM (Fig. 7B, Fig. S7C and D). ADAMs are proteases whose key role is proteolytic release of ectodomains of trans-membranous proteins, such as cytokines, growth factors, and cell adhesion molecules, including the NKG2D ligand MICA(*27*). They are closely associated with cancer progression, inflammation, tissue damage and repair, and preclinical studies with ADAM inhibitors against cancer and autoimmune diseases have shown encouraging results. However, thus far clinical trials have not been successful(*58*); for example, a trial with one such inhibitor for rheumatoid arthritis was halted in phase I/II due to hepatotoxicity(*59*). Typically, toxicity is correlated with dose; we predict that administering γδTc in combination with an inhibitor would enable lower doses to be employed, which could alleviate this issue. The inhibitor we used in this study, GW280264X, has not yet been tested in clinical trials; however, using GW280264X to block MICA shedding lead to significantly enhanced γδTc cytotoxicity against BCSC (Fig. 7J, K). Interestingly, this increased lysis of BCSC to the level of cytotoxicity observed against NSC in the absence of the inhibitor, suggesting that MICA shedding is the main mechanism underlying BCSC resistance to γδTc cytotoxicity. Since using this inhibitor in combination with γδTc also increased cytotoxicity against NSC, this approach could be considered to generally enhance γδTc targeting of cancer types using MICA shedding as an immune evasion mechanism. This mechanism was also employed under hypoxia, a biophysical property of the TME associated with poor patient outcome, in which we observed increased MICA shedding that correlated with reduced cytotoxicity of γδTc against breast cancer cell lines(*14*).

Since the GW280264X ADAM inhibitor overcame BCSC resistance to γδTc targeting, and also enhanced the cytotoxicity of γδTc against NSC, such inhibitors should be further developed for therapeutic applications. Our study provides a strong rationale for additional pre-clinical and clinical validation of ADAM inhibitors in combination with γδTc immunotherapy to improve patient outcomes.

## Materials and Methods

### Study Design

The overall objective of this study was to determine whether human γδTc are able to kill BCSC and, if not, how BCSC resistance can be overcome. We hypothesized that, similar to other treatment modalities, a fraction of BCSC would be resistant to γδTc cytotoxicity and that, once mechanisms were uncovered, this resistance could be overcome. In controlled laboratory experiments to test his hypothesis and determine mechanisms at play, we used 57 different primary human γδTc derived from the blood of 20 healthy donors. This study was conducted in compliance with the recommendations of the Research Ethics Guidelines, Health Research Ethics Board of Alberta—Cancer Committee with written informed consent from all participants according to the Declaration of Helsinki. The γδTc cultures were monitored for subset composition and purity by flow cytometry (Table S1). TNBC breast cancer lines SUM149, patient derived cell line PDX401 and luminal A breast cancer line, MCF-7 were used as targets. BCSC were enriched either by fluorescence activated cell sorting for CD44^+^CD24^-^ cells or by mammosphere assays (3D), and were compared to non-stem cells (NSC) that were CD44^+^CD24^-^ or adherent (2D), respectively. We used Calcein AM cytotoxicity assays, blocking assays, degranulation assays, mammosphere assays, tumor surface antigen profiling *via* flow cytometry, ELISAs, Western blotting and mass spectrometry, details of which are given in Materials and Methods, as well as the Extended Materials and Methods in Supplementary Materials. Due to limitations with respect to BCSC modeling *in vivo*, we were unable to perform extensive *in vivo* studies; however, we did explore lung metastasis formation in female NOD.Cg-Prkdcscid Il2rgtm1Wjl/SzJ (NSG) mice, which had received tail vein injections of SUM149 cells treated with γδTc or SUM149 cells alone; further details are provided below under “Mice”. The Animal Use Subcommittee at the University of Alberta approved the *in vivo* experiment protocol used in this study (AUP00001288). This experiment was randomized; mice were numbered with ear markings and treatments assigned using the randomization function in Excel. Animals from both treatment groups were housed together in mixed cages. After injections were performed, treatments were not listed on monitoring sheets to ensure observations henceforth (including outcomes) were blinded. Health monitoring, including weighing and clinical observation, was performed weekly. At the experimental endpoint, metastasis formation was quantified blindly by two investigators. Detailed descriptions of all experiments done are provided in the Materials and Methods below or in the Supplementary Materials. Sample sizes were chosen empirically to ensure sufficient statistical power and were in the line with field standards for the techniques used in this study. Data were only excluded if identifiable technical failure occurred in the course of the experiment (i.e. failure of positive or negative controls) or if primary cell viability was poor at the outset of an experiment. No outliers were excluded. The number of biological replicates is specified in the figure legends and in cases where cumulative figures are shown in the main figures, the results for the individual biological replicates are shown in the corresponding supplementary figures; technical replicates are also indicated.

### Primary Human γδ T Cells

Primary human γδTc cultures were established and maintained as described (*20*). Briefly, Peripheral blood mononuclear cells were extracted from blood drawn from healthy donors using density gradient separation (Lymphoprep, Stem Cell Technologies, Vancouver, BC, Canada) and cultured at 1 × 10^6^ cells/ml in RPMI complete medium containing 1μg/ml Concanavalin A (Sigma-Aldrich, Oakville, ON, Canada), 10% fetal bovine serum (FBS), 1× MEM NEAA, 10mM HEPES, 1 mM sodium pyruvate (all Invitrogen, Burlington, ON, Canada) plus 10ng/ml recombinant human IL-2 and IL-4 (Miltenyi Biotec, Auburn, CA, USA). Cells were maintained in a humidified incubator at 37°C with 5% CO_2_. Cells were counted every two to three days and densities were adjusted back to 1 × 10^6^ cells/ml by adding fresh medium and cytokines. After seven days, αβTc were depleted by labeling with anti-TCRαβ PE antibodies (BioLegend, San Diego, CA, USA) followed by anti-PE microbeads (Miltenyi Biotec), filtering through a 50μm Cell Trics filter (Partec, Görlitz, Germany) and passing through an LD depletion column on a MidiMACS magnet (both from Miltenyi Biotec). The flow-through contained γδTc, which were collected and further cultured in RPMI complete medium plus cytokines (as above) at 37°C with 5% CO_2_. Subset percentages and purities of different donor-derived γδTc cultures are listed in Table S1. As γδTc were most differentiated and therefore most cytotoxic by the end of the culture period (days 19-21), most experiments including cytotoxicity assays were done on those days. Exceptions were proliferation and inhibitory receptor assays, which were begun earlier in the culturing period.

### Breast Cancer Cell Lines

Inflammatory TNBC line SUM149 cells were cultured in Ham’s F-12 medium with 5% heat-inactivated FBS, 10mM HEPES, 1μg/mL hydrocortisone and 5μg/mL insulin. The patient derived xenograft cell line PDX401, obtained *via* collaboration with Oncotest (Charles River Discovery, Freiburg, Germany), was derived from a well-differentiated basal-like TNBC tumour. PDX401 were cultured in RPMI-1640 containing 10% FBS and 1% Gentamycin. Luminal A breast cancer cell lines, MCF-7 and T47D, were cultured in RPMI with 10% FBS. All breast cancer cells were authenticated at the Sick Kids Research Institute, Toronto, ON, Canada.

### Mice

8-11-week-old NOD.Cg-Prkdcscid Il2rgtm1Wjl/SzJ (NSG) mice were injected with SUM149 cells alone or SUM149 cells previously treated with γδTc, into their tail veins. 500,000 cells were suspended in 100μl PBS and injected into the tail vein, however, some mice only received 90μl cell suspension, as indicated in the lung quantification. 18 weeks later, the mice were sacrificed and their lungs stained with India Ink as per Wexler to identify macro metastasis(*60*); lungs were then stored in Fekete’s solution. The metastases appear white on the black stained lungs.

### Image Analysis

Lung images were taken with an iPhone 11, which has a 12 mega-pixels wide-angle camera with the lungs submerged in Fekete’s solution. The images were opened with ImageJ (Wayne Rasband National Institute of Health, USA, version 1.52q), converted to 8-bits and inverted. Regions of Interest (ROI) were traced around the “white metastases” (that appear black when inverted) and around total lung, and the areas of these regions were noted. Percent metastases were calculated as 100 x (area of white metastasis/area of total lung).

### Calcein AM (CalAM) Cytotoxicity/Blocking Assays

Cytotoxicity and blocking assays were done as described (*13*); details are in the supplemental materials. For degrading MCL-1 during cytotoxicity assays, 500nM dMCL1-2 or DMSO was added directly to the co-incubation. The MCL-1 degrader, dMCL1-2, was synthesized by the Dersken lab, University of Calgary(*25*). For ADAM inhibition experiments, 3μM ADAM inhibitor (GW280264X, Aobious Inc., Massachusetts, USA) or DMSO was added to the co-incubation.

### Mammosphere Assays

Target cells were trypsinized, passed through a 50μm cell strainer (CellTrics) and seeded into ultra-low adherent plates (Corning, NY, USA) in MammoCult media (StemCell Technologies, Vancouver, BC, Canada) as per manufacturer’s instructions. Mammospheres formed after 7 days were collected by centrifugation at 1000 rpm. For serial passaging, obtained mammospheres were dissociated into single cells and seeded again into ultra-low adherent plates (Corning).

## Flow Cytometry

### Surface Marker Staining

Gamma delta T cells and/or breast cancer cell lines were stained as described in the supplemental materials; the detailed list of antibodies used can be found in Supplemental Table S4. Antibody-stained cells were analyzed using a FACS CANTO II (Becton Dickinson, Mississauga, ON, Canada). Cell sorting was done using a FACSAria III cell sorter (Becton Dickinson). For details on the flow cytometers, please see “Flow Cytometer Specifications” in the supplemental materials.

### CD107 Assays

CD107 assays were performed as previously published(*35*); details are in the supplemental materials.

### Detection of Intracellular IFN-γ

Conditioned media (CM) treated γδTc were incubated with 5μg Anti-CD3 (UCHT1, BioLegend) per million cells followed by addition of 6μg/ml of monensin (Golgi-Stop, BD) after an hour. Subsequently after another 5h, cells were washed, fixed and permeabilized and stained with anti-IFN-γ antibody according to the manufacturer’s instructions (Cytoperm/Cytofix™ solution kit, BD Biosciences). PMA/ionomycin stimulated cells were used as a positive control.

### Detection of Apoptosis

Apoptosis analyses were performed as previously published (*13*); details are in the supplemental materials.

### CellTrace Violet Proliferation Assay

Gamma delta T cells were incubated with the indicated CM for 24h, washed, labeled with 1μM CellTrace Violet (Invitrogen) as per manufacturer’s instructions, and cultured in fresh media for six additional days, then washed and fixed in FACS buffer containing 2% paraformaldehyde prior to flow acquisition. Proliferation modeling was performed using FlowJo™ software, version 10.5.3.

### ELISA

Culture supernatants (1-2 ml) were stored at −80°C and upon thaw, Halt™ Protease and Phosphatase Inhibitor Cocktail (PIC, Thermo Fisher Scientific) was added to samples prior to use in ELISAs or further storage at 4°C. The following ELISA kits were used: Human MICA DuoSet ELISA (R&D systems) and ELISA MAX™ Deluxe Set Human IFN-γ (Biolegend). For MICA ELISAs, supernatants were concentrated using Amicon Ultra-4 10 K spin columns (Merck-Millipore, Carrigtwohill, Ireland). The columns were centrifuged at 3,000 g for 2h at 12°C. The final volume of concentrated media was adjusted to 200μl, and 100μl per well were plated in duplicate. All ELISAs were done in accordance to the manufacturer’s instructions. Absorbance was measured at both 450 and 550 nm using a FLUOstar Omega plate reader (BMG Labtech, Offenburg, Germany) with Omega Software version 5.11. For calculating concentrations, the difference linear regression fit of the standard curve was used. ELISA data were normalized to cell numbers.

### Immunoblotting

Immunoblotting was performed as described in the Supplementary Materials and Methods; antibodies used are listed in Table S4.

### Mass spectrometry

Details of mass spectrometry sample preparation and data analyses are included in the Supplementary Materials and Methods.

## Statistics

All statistical tests were performed as indicated in the figure legends, using Prism 7.0 for Mac OSX (GraphPad Software, San Diego, CA, USA). The Shapiro-Wilk test was done to determine normality for all data; in cases where normality was not achieved, non-parametric tests were employed. Depending on the experiment design, the following methods were used to determine significance: Student’s t-test, Wilcoxon test, one-way analysis of variance (ANOVA), or twoway ANOVA. ANOVA tests were followed by Tukey’s, Bonferroni’s, or Sidak’s post hoc test for multiple comparisons between groups as indicated. All data tested for differences were considered significant when p<0.05; levels of significance are denoted by asterisks or letters as defined in the figure legends. All statistical analysis done and p-values are listed in Table S2 for main figures and Table S3 for supplemental.

## Supplementary Materials

### Supplementary Materials and Methods

Calcein AM (CalAM) Cytotoxicity/Blocking Assays; Flow Cytometry (Surface Marker Staining, CD107 Assays, Detection of Apoptosis, Flow Cytometer Specifications); Immunoblotting; Mass Spectrometry (Sample preparation, LC-MS/MS, Data analysis).

**Fig. S1.** Breast cancer stem-like cells in CD44+CD24-SUM149 and PDX401 3D mammospheres are further enriched upon co-incubation with γδ T cells.

**Fig. S2.** Gamma delta T cell degranulation and IFN-γ secretion are impaired in the presence of breast cancer stem-like cells.

**Fig. S3.** Breast cancer stem-like cells secrete factors that inhibit γδ T cell function but do not impact their viability.

**Fig S4.** Inhibitory ligands are expressed on breast cancer stem-like cells resistant to γδ T cell killing.

**Fig S5.** The Fas-FasL pathway is dysfunctional in breast cancer stem-like cells.

**Fig S6.** MICA/B is downregulated on the surface of breast cancer stem-like cells.

**Fig S7.** The ADAM inhibitor GW280264X prevents MICA shedding and enhances γδ T cell cytotoxicity against breast cancer stem-like cells.

**Table S1**. Subset percentages and purities of donor-derived γδ T cell cultures.

**Table S2**. Statistical tests employed and resulting p-values for experiments shown in Figures 1–7.

**Table S3.** Statistical tests employed and resulting p-values for experiments shown in Supplemental Figures S1-S8.

**Table S4.** Flow cytometry, western blot and blocking assay antibodies.

**Table S5.** Proteins acquired by mass spectrometry that were differentially expressed in conditioned medium from CD24-versus CD24+ SUM149 cells.

**Table S6.** Proteins acquired by mass spectrometry that were differentially expressed in conditioned medium from 3D versus 2D PDX401 cells.

