## Supplementary Materials for "ADAM protease inhibition overcomes resistance of breast cancer stem-like cells to γδ T cell immunotherapy"

### Supplementary Materials and Methods

#### Calcein AM (CalAM) Cytotoxicity/Blocking Assays

Target cells were labeled with 5µM CalAM in PBS (Invitrogen/Thermo Fisher Scientific, Waltham, MA, USA). Target cells were diluted to a density of 30,000 cells/100µl medium (RPMI 1640 plus 10% heat-inactivated FBS; 10mM HEPES; 1× MEM NEAA; 1 mM sodium pyruvate; 50U/ml penicillin–streptomycin; and 2mM l-glutamine, all from Invitrogen) and γδTc were re-suspended to 600,000 cells/100µl in medium. An aliquot of the diluted γδTc were further diluted down to 300,000 cells/100µl in medium and 30,000 cells/100µl to obtain effector:target ratios of 20:1, 10:1 and 1:1, respectively. 100µl CalAM-labeled targets were added to 100µl γδTc in 96-well round-bottom plates in triplicate followed by incubation at 37°C for 4 hours (h). For blocking experiments (including PD-1 blocking), 4µg of blocking antibody were added to 400µl cell suspension of γδTc or targets (as indicated) for each test in Eppendorf tubes, then 100µl were plated per well in a 96-well round-bottomed plate in triplicate. For blocking assays, untreated and IgG-treated cells were used as experimental controls. After incubation with blocking antibody at 37°C for 30 min, CalAM-labeled targets were added to γδTc followed by incubation at 37°C for 4h. Plates were then centrifuged and supernatants transferred to fresh 96-well plates (Corning 96-well, black cell culture-treated flat-bottom microplate with clear bottom, Fisher Scientific) for CalAM fluorescence detection on a fluorimeter (FLUOstar Omega, BMG labtech). Controls were CalAM-labeled target cells incubated alone (spon = spontaneous release) or with 0.05% Triton-X 100 (Thermo Fisher Scientific) (max = maximum release). Percent lysis was calculated: [(test − spon)/(max − spon)] × 100%.

**Flow Cytometry**

**Surface Marker Staining**

Gamma delta T cells and/or breast cancer cell lines were re-suspended to a dilution of $1 \times 10^6$ cells/100µl and stained with 1µl/$10^6$ cells Zombie Aqua (ZA) or Zombie Near Infrared (ZNIR) fixable viability dye in PBS (ZA or ZNIR, BioLegend) for 15–20 min at room temperature in the dark, washed and then fluorochrome-conjugated antibodies diluted in FACS buffer [PBS containing 1% FBS and 2mM EDTA (Invitrogen)] were used to stain γδTc. Breast cancer cell lines were blocked at $10 \times 10^6$ cells/ml in FACS buffer containing 50µl/ml Trustain FcX (BioLegend) on ice for 30 min prior to antibody incubation. After blocking, cells were centrifuged and supernatants were removed, leaving 10µl FACS buffer plus block/$10^6$ cells. Antibodies and FACS buffer were added to 20µl, and cells were then incubated on ice for 10–15 min followed by washing. All cells were fixed in FACS buffer containing 2% paraformaldehyde (Sigma-Aldrich), stored at 4°C and acquired within one week. For flow sorting, cells were stained as described above in sorting buffer (dPBS with 2% FBS, 1mM EDTA, 10mM HEPES and 5% penicillin/streptamycin); live cells were gated according to their forward scatter and side scatter profiles. A detailed list of antibodies used can be found in Table S4.

**CD107 Assays**

CD107 assays were performed as previously published (*35*). Briefly, 1 x $10^5$ γδTc with 5µL anti-CD107 AF647 (Biolegend) were plated alone or mixed with 5 x $10^5$ target cells in 200µl complete media in a 96-well round-bottomed plate in triplicates. To the positive control wells, Phorbol-12-myristate-13 acetate (PMA)/ionomycin (0.15nM/0.3µg/mL final concentrations) was added. Cells were incubated for 1h at 37°C and then 6µg/ml monensin (Golgi-stop, BD) were added. Cells were further incubated for another 2h at 37°C, after which cells were covered and put on ice. Further staining was performed as described in surface marker staining above.

**Detection of Apoptosis**

Apoptosis analysis were performed as previously published (*13*). Cultured γδTc or cancer cells were stained with 5ng/µl ZA fixable viability dye (BioLegend) first for 15–20 min after which they were washed with 1× Annexin V (AnnV) binding buffer (BioLegend), and stained with AnnV FITC (BioLegend, 1:20) on ice for 15 min in the dark. Cells were then washed and re-suspended in 200µl AnnV binding buffer plus 2% paraformaldehyde and stored at 4°C until analyzed, within one week.

**Flow Cytometer Specifications**

Antibody-stained cells were analyzed using a FACS CANTO II (Becton Dickinson, Mississauga, ON, Canada) that has an air-cooled 405-nm solid state diode, 30 mW fiber power output violet laser, with 450/50 and 510/50 band pass (BP) [502 long pass (LP) detector]; a 488-nm solid state, 20-mW blue laser with 530/30 BP (502 LP), 585/42 BP (556 LP), 670 LP (655 LP), and 780/60 BP (735 LP) filters; and a 633-nm HeNe, 17-mW red laser with 660/20 BP and 780/60 BO (735 LP) filters. For calibration, CS&T beads (Becton Dickenson, Mississauga, ON, Canada) were used. Forward and side-scatter properties were used to discriminate live and dead cells and gate singlets. Fluorescence minus one (FMO) controls were used to set gates. Cell sorting was done using a FACSAria III cell sorter (Becton Dickinson, Mississauga, ON, Canada), that had fiber-launched fixed-wavelength air cooled lasers: 488 nm blue laser with 530/30nm, 695/40 nm filters, 561nm Yellow/Green laser with 582/21 nm, 610/20 nm, 670/14 nm,710/50 nm, 780/60 nm filters,

633 nm red laser with 660/20 nm, 730/45 nm, 780/60 nm filters, 405 or 375 nm Violet or near UV laser with 450/40 nm, 510/50 nm, 610/20 nm, 660/20 nm, 710/50 nm and 780/60 nm filters. Analysis of acquired data was performed using FlowJo© software (Tree Star, Ashland, OR, USA, Version 10.0.8r1).

**Immunoblotting**

Cell lysates were prepared by treating γδTc or breast cancer cell lines with M-PER Mammalian Protein Extraction Reagent (Thermo Fisher Scientific) containing PIC at 10µl lysis buffer per million γδTc or 50µl per million tumour cells followed by incubation at room temperature for 10 min. Lysates were then centrifuged at 13,000 rpm for 20 min at 4°C, after which supernatants were transferred to fresh tubes and 5× reducing sample buffer [0.0625 M Tris/HCl pH6.8, 2% SDS, 20% glycerol, 0.05% β-mercaptoethanol, 0.025% (w/v) Bromophenol Blue] was added. Samples were boiled 5 min, cooled, and briefly centrifuged in a benchtop centrifuge prior to running on 10% or 12% SDS-PAGE gels. Proteins were transferred onto Immobilon-FL PVDF membranes (Millipore) using the Trans-Blot Turbo Transfer System (Bio-Rad, Mississauga, ON, Canada). The "high MW" program was used for ADAM 10 and ADAM 17 detection; "mixed MW" program was used for all other proteins. Membranes were blocked for 40 min in 3% milk in TBST, followed by overnight incubation in primary antibody baths at 4°C. The next day, after washing, membranes were incubated with the corresponding species-specific HRP-labeled secondary antibody for 1h, followed by further washing and then detection using Clarity™ Western ECL Substrate (Bio-Rad). Primary antibodies baths were comprised of PBS containing 2% bovine serum albumin and 0.05% sodium azide. Antibodies used are listed in Table S4.

**Mass Spectrometry**

**Sample preparation**

To prepare CM for LC-MS, concentrated CM were lyophilized and re-suspended in 8M urea, 50mM ammonium bicarbonate (ABC), 10mM dithiothreitol (DTT) and 2% sodium dodecyl sulfate. CM protein were quantified using a Pierce™ 660nm Protein Assay with Ionic Detergent Compatibility Reagent (Thermo Scientific™) and ~10-50 µg was reduced in 10mM DTT for 30 minutes and alkylated in 100mM iodoacetamide for 30 minutes at room temperature in the dark. Proteins were immediately precipitated in methanol/chloroform according to Wessel and Flügge (*61*). Briefly, samples were topped up to 150µL with 50mM ABC, mixed with 600µL ice cold methanol and 150µL of ice cold chloroform, and vortexed thoroughly. An additional 450µL of 4°C distilled water was added followed by vortexing and centrifugation at 14,000 xg for 5 min. The upper aqueous/methanol phase was carefully removed to avoid disturbing the precipitated protein interphase. A second 450µL of cold methanol was added to each sample before the tube was inverted several times and centrifugaged at 14,000 xg for 5 min. The remaining methanol/chloroform was discarded and the precipitated protein pellet was left to air dry in a fume hood. On-pellet in-solution protein digestion was performed similarly to Duan *et al. (62)*. Briefly, precipitated protein pellets were reconstituted in 100µL of 50mM ABC (pH 8) and sonicated (~3 x 0.5s pulses) with a probe sonicator (Fisher Scientific, Waltham, MA) to the break up the pellet. LysC (Wako Chemicals, USA) and mass spec grade trypsin/LysC mix (Promega, Madison, WI, USA) were added to protein samples at 1:100 and 1:50 ratios, respectively. Protein digestion was carried out at 37°C on a ThermoMixer C (Eppendorf) at 400 rpm overnight (~18h). The next day an additional volume of trypsin/LysC mix (1:100 ratio) was added to each sample and mixed at

1400 rpm. After 3-4h, digests were acidified to pH 3-4 with 10µL of 10% formic acid (FA) and centrifuged at 14,000xg to pellet insoluble material prior to LC-MS/MS.

**LC-MS/MS**

Digests were analyzed on Q Exactive Plus mass spectrometer (Thermo Scientific) connected to an Waters ACQUITY M-Class UPLC. Solvent A consisted of water/0.1% FA and solvent B consisted of acetonitrile/0.1% FA. Peptides (~1µg estimated using a Pierce™ bicinchoninic acid assay) were initially loaded onto an ACQUITY UPLC M-Class Symmetry C18 Trap Column (5µm, 180µm x 20mm) and trapped for 6 minutes at a flow rate of 5µl/min at 99% A/1% B. Peptides were separated on an ACQUITY UPLC M-Class Peptide BEH C18 Column (130Å, 1.7µm, 75µm X 250mm) operating at a flow rate of 300nL/min at 35°C using a non-linear gradient consisting of 1-7% B over 1 minute, 7-23% B over 179 minutes and 23-35% B over 60 minutes, followed by washing and re-equilibration. The MS acquisition instrument settings are same as used in (*63*).

**Data analysis**

MS files were searched in MaxQuant (version 1.6.3.4) against the Human Uniprot database (reviewed entries plus isoforms) (*64*). Missed cleavages were set to 3 and cysteine carbamidomethylation was set as a fixed modification. Oxidation (M), N-terminal acetylation (protein), and deamidation (NQ) were set as variable amino acid modifications (max. number of modifications per peptide = 5). LFQ Min. ratio count was set to 1 and all other settings were left default. Protein and peptide FDR was left at 0.01 (1%) and the decoy database was set to revert. The 'match between runs' feature was utilized to maximize proteome coverage and label-free quantification (*65*). Search results were loaded into Perseus or R, and proteins labeled as 'only identified by site', 'matched to reverse' or 'potential contaminant' were removed (*66*). Protein identifications with label free quantification (LFQ) values in ≥2 biological replicates were retained for downstream analysis and missing values were imputed using a width of 0.3 and down shift of 1.8. Gene ontology cellular component (GOCC) annotation were performed using Metascape (version 3.0). Gene set enrichment analysis (GSEA) were performed in GSEA v3.0 (Broad Institute) with a minimum gene set size of 10 and permutation type set to gene_set. Molecular Signature Database (MSigDB) v6.2 collections: canonical pathways, hallmark, KEGG gene sets, C2 curated sets (*67*). Heatmaps were produced using the Bioconductor package complexHeatmap (*68*).

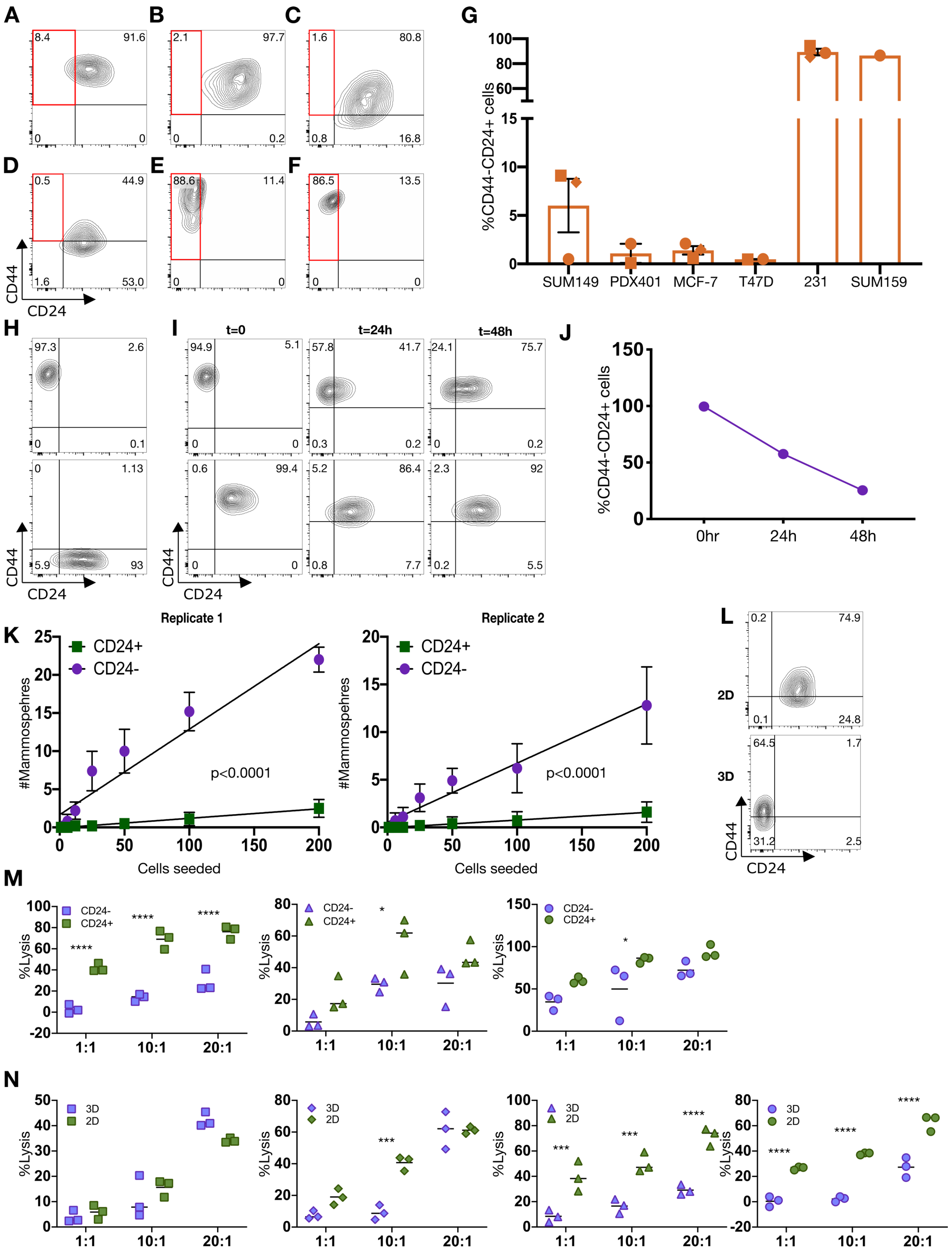

A
B
C
D
E
F
G
H
I
J
K
L
M
N
CD44
CD24
%CD44-CD24+ cells
SUM149
PDX401
MCF-7
T47D
231
SUM159
t=0
t=24h
t=48h
0hr
24h
48h
Replicate 1
Replicate 2
CD24+
CD24-
#Mammospehres
p<0.0001
Cells seeded
2D
3D
%Lysis
1:1
10:1
20:1

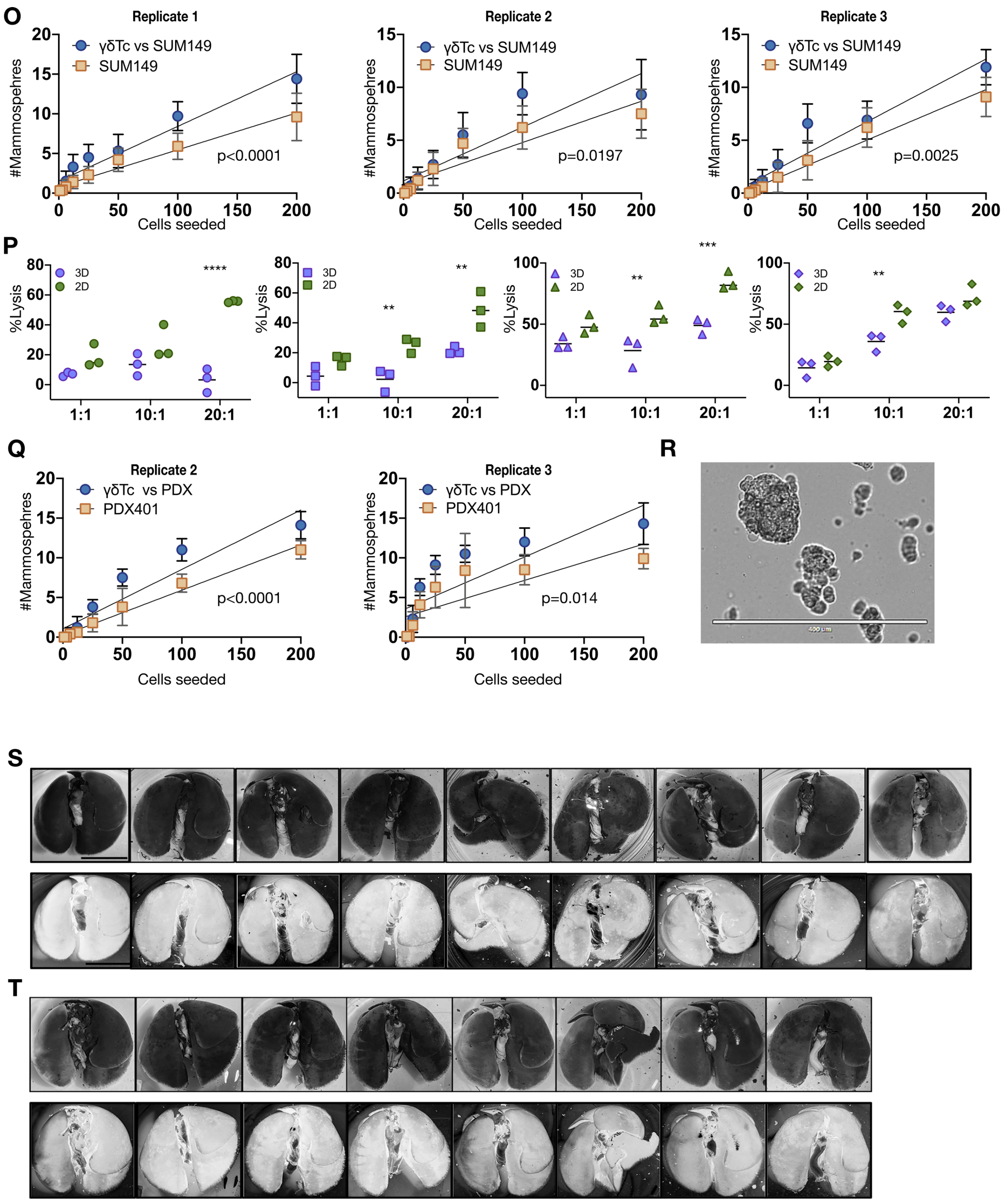


**Figure S1. Breast cancer stem-like cells in CD44+CD24- SUM149 and PDX401 3D mammospheres are further enriched upon co-incubation with γδ T cells.** Representative examples from a panel of breast cancer cell lines **(A)** SUM149 (n=3), **(B)** PDX401 (n=2), **(C)** MCF-7 (n=3), **(D)** T47D (n=2), **(E)** MDA-MB-231 (n=3), and **(F)** SUM159 (n=1) that were stained for surface expression of CD44 and CD24 followed by flow cytometric acquisition, which is graphically represented in **(G)**. **(H)** Quadrant gates were set for analyses in A-F using fluorescence minus one (FMO) controls as shown in this representative example for **(I)** SUM149 cells sorted into CD44+CD24- (top left panel) and CD44+CD24+ (bottom left panel) fractions; sorted populations were maintained in culture and stained with CD24 and CD44 after the indicated times to determine the kinetics of differentiation, depicted graphically in **(J)**. **(K)** Sorted SUM149 cells were seeded as indicated and cultured for seven days under low-serum and low attachment conditions. The spheres that formed were counted (n=2). **(L)** Second-generation PDX401 adherent cells (2D) and mammospheres (3D) were dissociated, then CD44 and CD24 expression were determined by flow cytometry (n=1). **(M)** SUM149 cells were sorted and used in Calcein AM cytotoxicity assays with γδTc derived from three different donors. **(N)** Second generation mammospheres (3D) and adherent (2D) SUM149 were dissociated, filtered into single cell suspensions and used as targets in cytotoxicity assays with γδTc from four different donors. **(O)** SUM149 alone or SUM149 treated with γδTc, from which γδTc were removed, were seeded and the number of spheres formed were counted. In each experiment, ten technical replicates were counted. Three biological replicates of this experiment, in which γδTc were derived from different donors, are shown. **(P)** Experiment done as in **(N)** but with PDX401 3D and 2D cells, and with γδTc derived from four different donors. The symbols used here match the donor symbols from the cumulative figure. **(Q)** Experiments done as in O but with PDX401 target cells. **(R)** Image of second generation PDX401 mammospheres. **(S)** SUM149 cells alone or those previously treated with γδTc **(T)** were injected into the tail veins of NSG mice and, 18 weeks later, mice were sacrificed and lungs stained with India ink. White spots in the black stained lungs indicate macroscopic metastases; scale = 1 cm. Corresponding inverted images of the lungs are below each set of original lung images. are Data are presented as mean ± SEM in G; mean ± SD in K, O and Q; and mean in M, N and P. Statistical tests employed were: (K, O, Q) Simple linear regression, p values are indicated in figure; (M, N, P) Two-way ANOVA followed by Sidak's post hoc test for multiple comparisons between groups. *$p<0.05$, **$p<0.01$, ***$p<0.001$, ****$p<0.0001$.

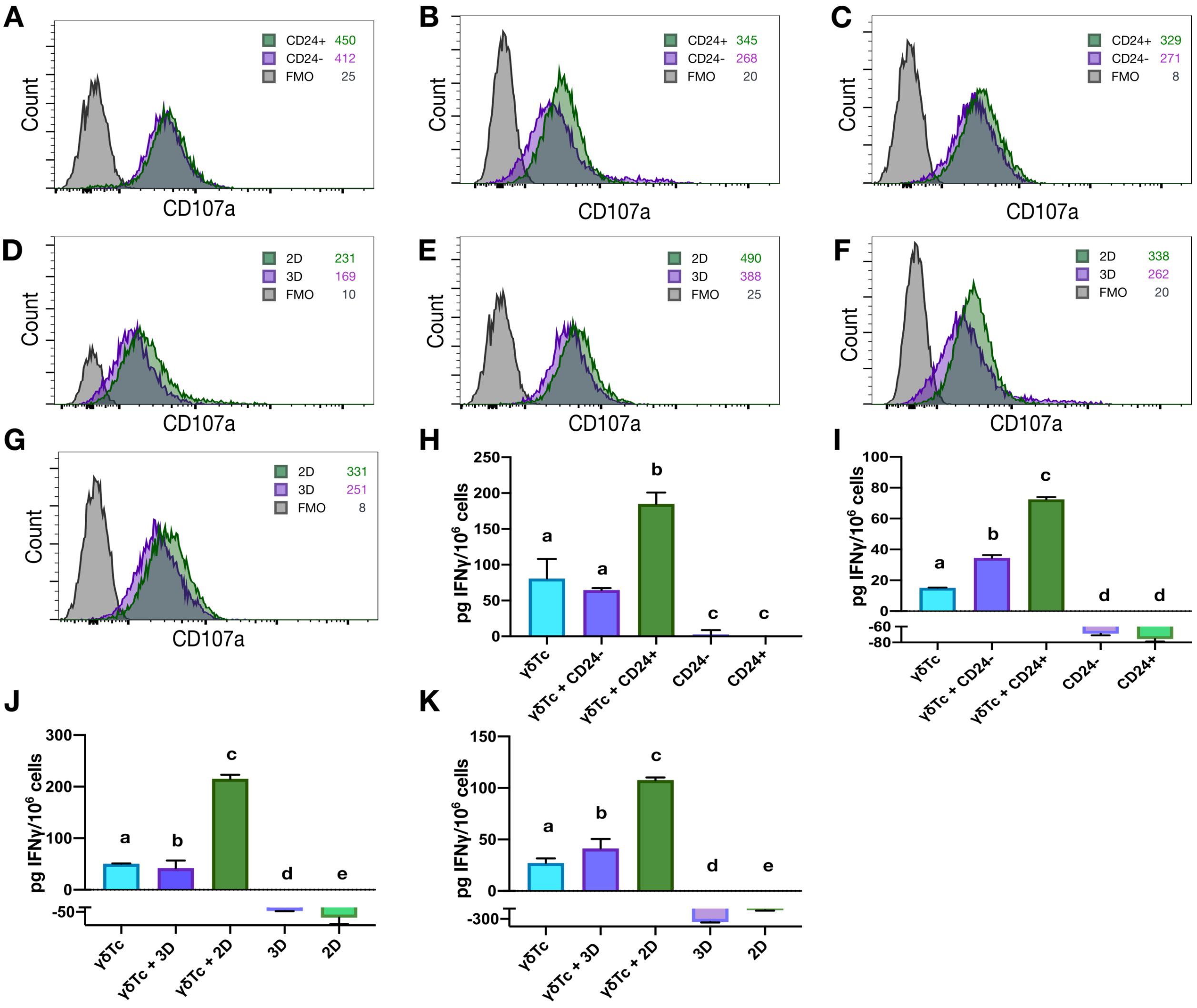


**Figure S2. Gamma delta T cell degranulation and IFN-γ secretion are impaired in the presence of Breast Cancer Stem-Like Cells. (A)** Overlay of of CD107a expression (degranulation) is shown for γδTc incubated with SUM149 CD24- and incubated with CD24+ cells is shown for γδTc derived from donor 1, **(B)** donor 2 and **(C)** donor 3. **(D)** Experiments done as in A-C but with PDX401 3D (BCSC) or 2D (NSC) cells and γδTc from donor 1 **(E)** donor 2, **(F)** donor 3 and **(G)** donor 4. **(H)** INF-γ ELISA was performed on conditioned media (CM) from $CD24^{+}$ or $CD24^{-}$ SUM149 co-incubated with γδTc derived from donor 2 and **(I)** donor 3 for 24h. **(J)** INF-γ ELISA on conditioned media (CM) from 2D or 3D PDX401 co-incubated with γδTc derived from donor 2 and **(K)** donor 3 for 24h. Donor numbers were reset for each set of experiments. Data are presented as mean ± SD. Statistical tests employed were: (G-J) One-way ANOVA followed by Tukey's post hoc test for multiple comparisons. In H, a-b, $p<0.01$; in I and J, b-c, $p<0.0001$; in K, b-c, $p=0.0038$.

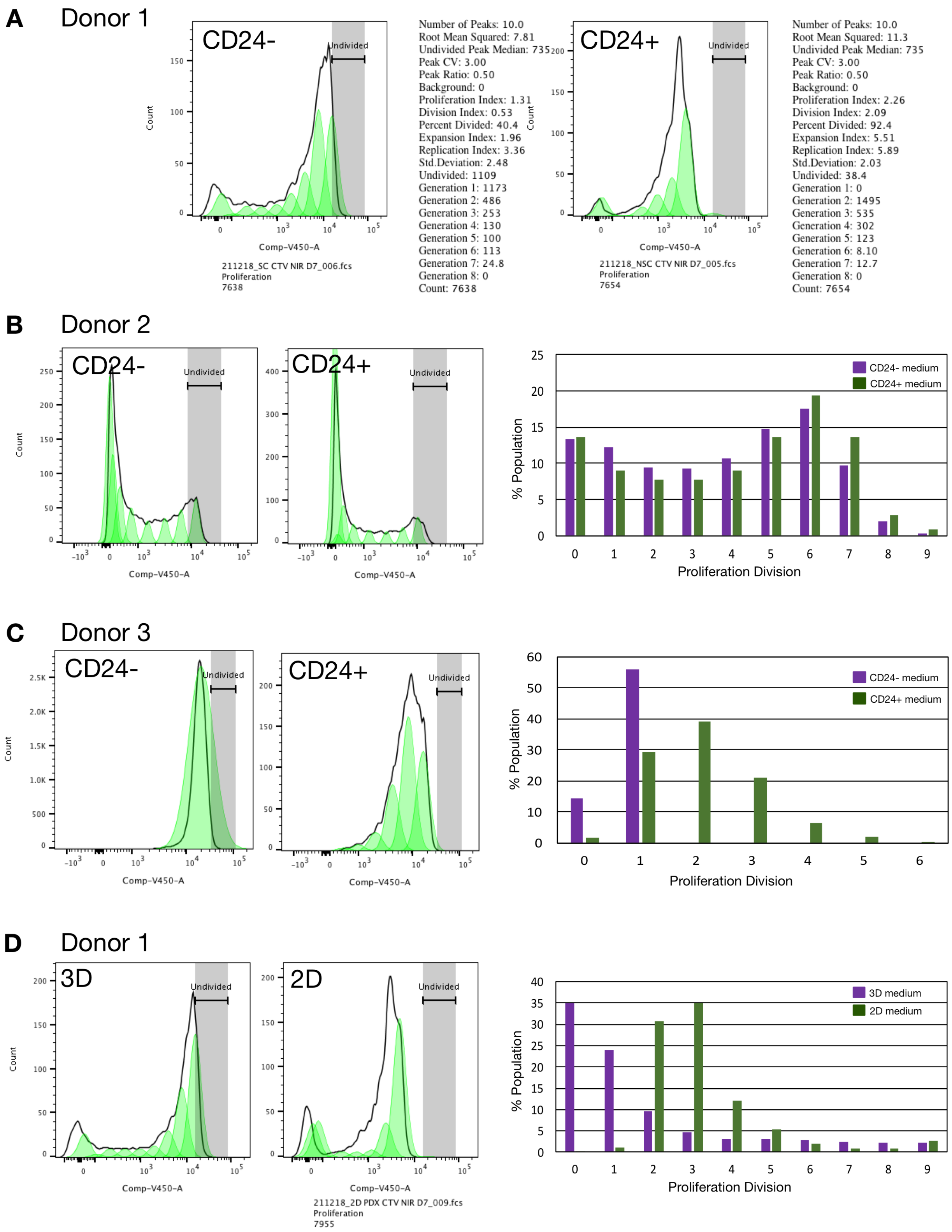

A
Donor 1
CD24-
Undivided
Count
Comp-V450-A
211218_SC CTV NIR D7_006.fcs
Proliferation
7638
Number of Peaks: 10.0
Root Mean Squared: 7.81
Undivided Peak Median: 735
Peak CV: 3.00
Peak Ratio: 0.50
Background: 0
Proliferation Index: 1.31
Division Index: 0.53
Percent Divided: 40.4
Expansion Index: 1.96
Replication Index: 3.36
Std.Deviation: 2.48
Undivided: 1109
Generation 1: 1173
Generation 2: 486
Generation 3: 253
Generation 4: 130
Generation 5: 100
Generation 6: 113
Generation 7: 24.8
Generation 8: 0
Count: 7638
CD24+
Undivided
Count
Comp-V450-A
211218_NSC CTV NIR D7_005.fcs
Proliferation
7654
Number of Peaks: 10.0
Root Mean Squared: 11.3
Undivided Peak Median: 735
Peak CV: 3.00
Peak Ratio: 0.50
Background: 0
Proliferation Index: 2.26
Division Index: 2.09
Percent Divided: 92.4
Expansion Index: 5.51
Replication Index: 5.89
Std.Deviation: 2.03
Undivided: 38.4
Generation 1: 0
Generation 2: 1495
Generation 3: 535
Generation 4: 302
Generation 5: 123
Generation 6: 8.10
Generation 7: 12.7
Generation 8: 0
Count: 7654
B
Donor 2
CD24-
CD24+
Undivided
Count
Comp-V450-A
CD24- medium
CD24+ medium
% Population
Proliferation Division
C
Donor 3
CD24-
CD24+
Undivided
Count
Comp-V450-A
CD24- medium
CD24+ medium
% Population
Proliferation Division
D
Donor 1
3D
2D
Undivided
Count
Comp-V450-A
211218_2D PDX CTV NIR D7_009.fcs
Proliferation
7955
3D medium
2D medium
% Population
Proliferation Division

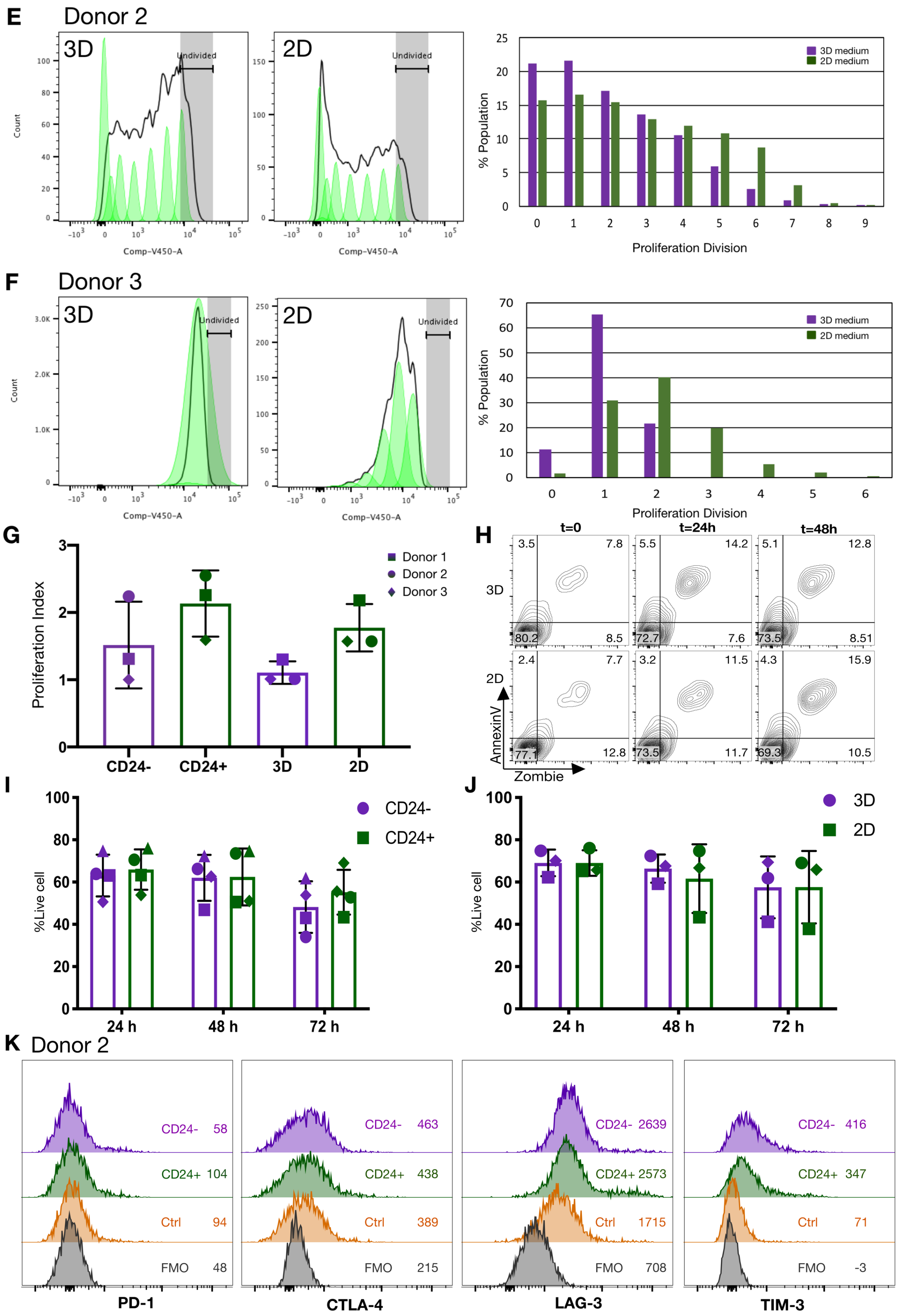
E
Donor 2
3D
2D
Undivided
Count
Comp-V450-A
% Population
3D medium
2D medium
Proliferation Division
F
Donor 3
3D
2D
Undivided
Count
Comp-V450-A
% Population
3D medium
2D medium
Proliferation Division
G
Proliferation Index
Donor 1
Donor 2
Donor 3
CD24-
CD24+
3D
2D
H
t=0
t=24h
t=48h
3D
2D
3.5
7.8
80.2
8.5
5.5
14.2
72.7
7.6
5.1
12.8
73.5
8.51
2.4
7.7
77.1
12.8
3.2
11.5
73.5
11.7
4.3
15.9
69.3
10.5
AnnexinV
Zombie
I
%Live cell
CD24-
CD24+
24 h
48 h
72 h
J
%Live cell
3D
2D
24 h
48 h
72 h
K
Donor 2
CD24- 58
CD24+ 104
Ctrl 94
FMO 48
PD-1
CD24- 463
CD24+ 438
Ctrl 389
FMO 215
CTLA-4
CD24- 2639
CD24+ 2573
Ctrl 1715
FMO 708
LAG-3
CD24- 416
CD24+ 347
Ctrl 71
FMO -3
TIM-3

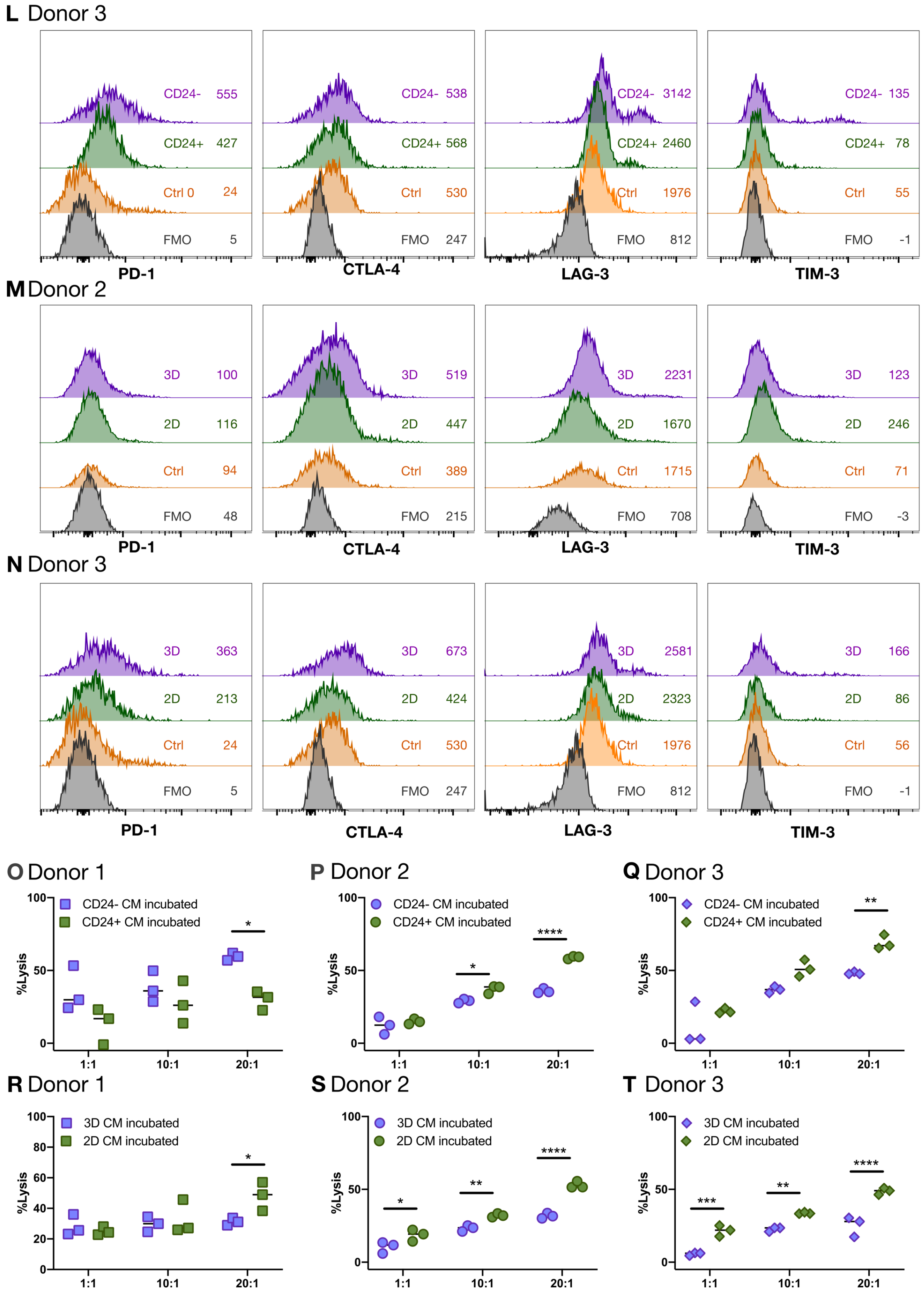
L Donor 3
CD24- 555
CD24+ 427
Ctrl 0 24
FMO 5
PD-1
CD24- 538
CD24+ 568
Ctrl 530
FMO 247
CTLA-4
CD24- 3142
CD24+ 2460
Ctrl 1976
FMO 812
LAG-3
CD24- 135
CD24+ 78
Ctrl 55
FMO -1
TIM-3
M Donor 2
3D 100
2D 116
Ctrl 94
FMO 48
PD-1
3D 519
2D 447
Ctrl 389
FMO 215
CTLA-4
3D 2231
2D 1670
Ctrl 1715
FMO 708
LAG-3
3D 123
2D 246
Ctrl 71
FMO -3
TIM-3
N Donor 3
3D 363
2D 213
Ctrl 24
FMO 5
PD-1
3D 673
2D 424
Ctrl 530
FMO 247
CTLA-4
3D 2581
2D 2323
Ctrl 1976
FMO 812
LAG-3
3D 166
2D 86
Ctrl 56
FMO -1
TIM-3
O Donor 1
P Donor 2
Q Donor 3
CD24- CM incubated
CD24+ CM incubated
%Lysis
1:1
10:1
20:1
R Donor 1
S Donor 2
T Donor 3
3D CM incubated
2D CM incubated
%Lysis
1:1
10:1
20:1

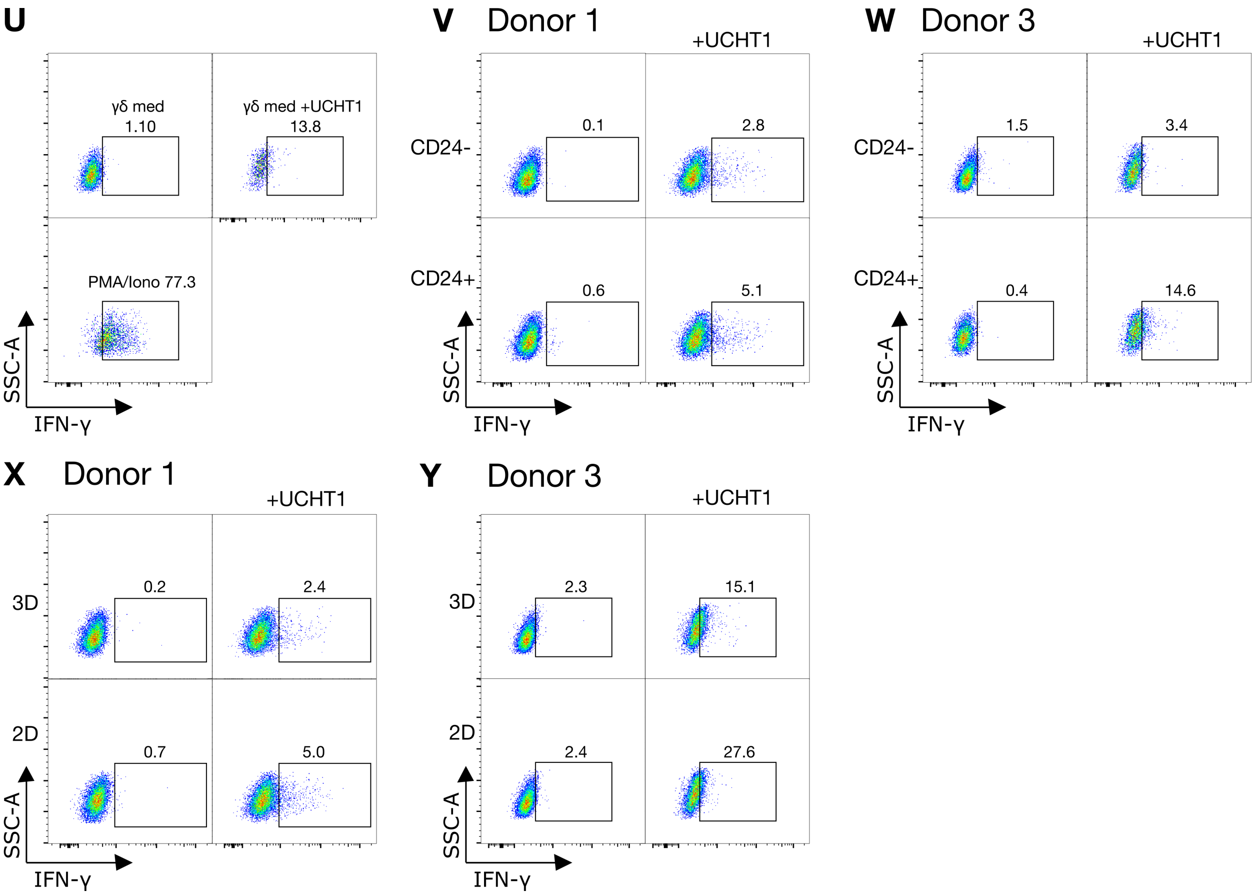


**Figure S3 Breast cancer stem-like cells secrete factors that inhibit gamma delta T cell function but do not impact their viability. (A)** γδTc were labelled with Cell Trace Violet (CTV) and incubated with SUM149 SC (CD24-) or NSC (CD24+) conditioned medium (CM) for 24h, washed and their proliferation measured six days later using flow cytometry. Output from the FlowJo 10.5.3 Proliferation Modeling Tool for γδTc derived from donor 1, **(B)** donor 2 and **(C)** donor 3. **(D)** Experiments done as in A-C but with PDX401 3D and 2D media treatment of γδTc from donor 1, **(E)** donor 2 and **(F)** donor 3. **(G)** Graph of cumulative proliferation index values from the three donor cultures treated with SUM149 or PDX401 CM. **(H)** γδTc were incubated with PDX 2D and 3D CM for 24 h, washed and viability was accessed using Zombie Aqua and AnnexinV staining after indicated duration. **(I)** Cumulative results for percent live cells of γδTc incubated with SUM149 CD24- and CD24+ CM (n=4) or **(J)** PDX401 3D and 2D CM (n=3). **(K)** γδTc were incubated with SUM149 CD24- and CD24+ CM for 24h; histograms for expression of the indicated co-inhibitory receptors on γδTc seven days after incubation with SUM149 CD24- and CD24+ are depicted for donor 2 with median fluorescence intensities indicated. **(L)** The same experiment was done with γδTc from donor 3. **(M)** Experiments done as in K-L but with PDX401 3D and 2D media treating γδTc from donor 2 and **(N)** donor 3. Ctrl stands for cells that were incubated with regular media instead of conditioned media. After incubation with SUM149 CD24- and CD24+ CM, γδTc were used to target unsorted SUM149 in Calcein AM cytotoxicity assays at the indicated effector: target (E:T) ratios. **(O)** Percent lysis of SUM149 cells using γδTc from donor 1, **(P)** donor 2 and **(Q)** donor 3. **(R)** Experiments done as in O-Q but with PDX401 target cells and PDX401 3D and 2D media-treated γδTc from donor 1, **(S)** donor 2 and **(T)** donor 3. **(U)**

γδTc incubated with SUM149 CD24- and CD24+ CM were treated with anti-CD3 antibody (UCHT1) for 6h and IFN-γ expression was determined via intracellular flow cytometry. Unstimulated cells were used to set gates and γδTc treated with PMA/Ionomycin were used as positive control. Control gates from donor 3 are shown as an example. **(V)** Percentage of IFN-γ-producing γδTc is depicted for γδTc derived from donor 1 and **(W)** donor 3. **(X)** Similarly, results from PDX401 3D and 2D CM-treated γδTc derived from donor 1 and **(Y)** donor 3 are shown. Donor numbers are reset for each set of experiments. Data are presented as mean ± SD in G, I-J, and median in O-T. Statistical tests employed were: (G) One-way ANOVA followed by Sidak's post hoc test for multiple comparisons between groups. (I-J, O-T) One-way ANOVA followed by Sidak's post hoc test for multiple comparisons between group. No significance was achieved for G, I-J. *P<0.05, **P<0.01, ***P<0.001, ****P<0.0001.

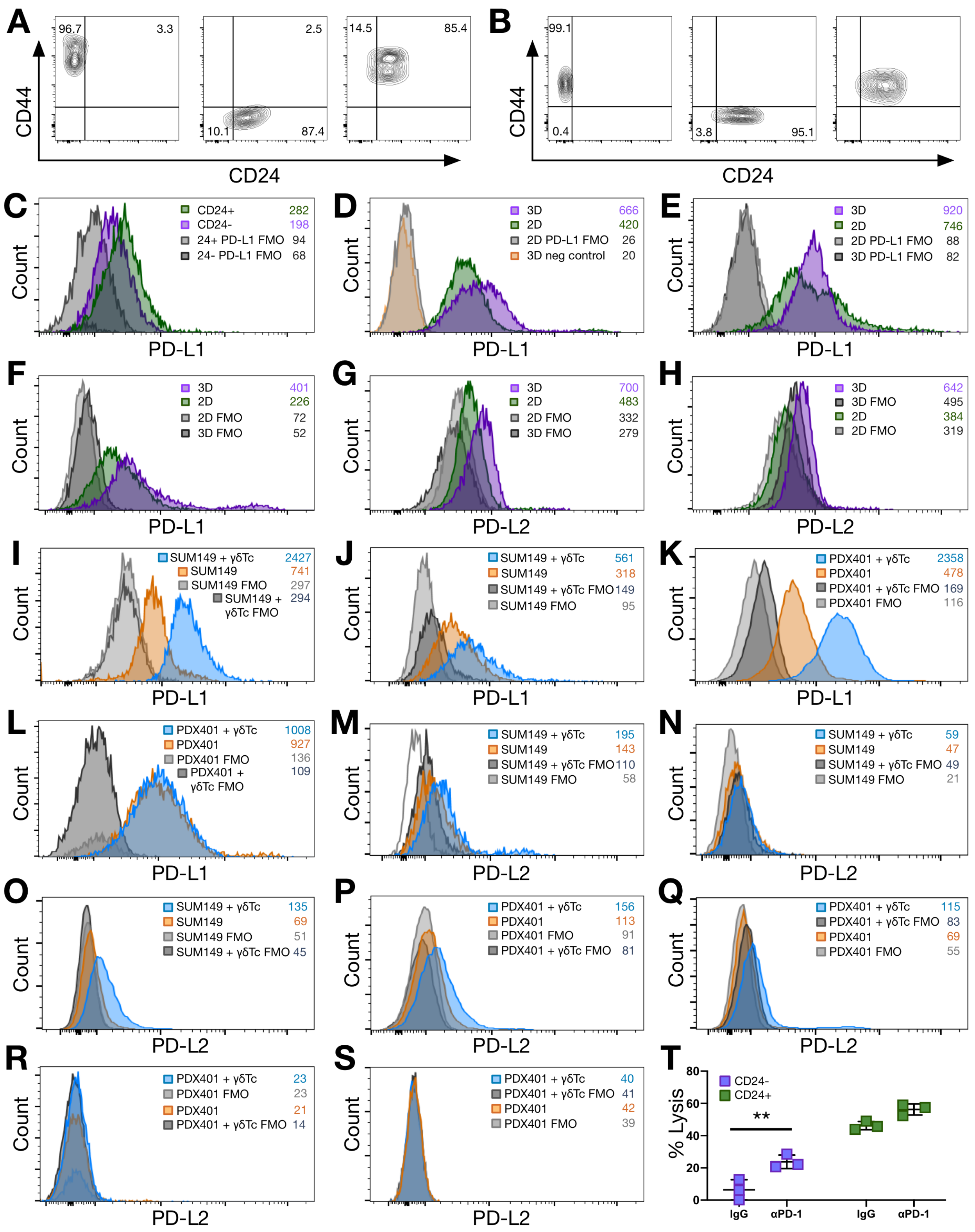

A
96.7
3.3
2.5
14.5
85.4
10.1
87.4
CD44
CD24
B
99.1
0.4
3.8
95.1
CD44
CD24
C
CD24+ 282
CD24- 198
24+ PD-L1 FMO 94
24- PD-L1 FMO 68
Count
PD-L1
D
3D 666
2D 420
2D PD-L1 FMO 26
3D neg control 20
Count
PD-L1
E
3D 920
2D 746
2D PD-L1 FMO 88
3D PD-L1 FMO 82
Count
PD-L1
F
3D 401
2D 226
2D FMO 72
3D FMO 52
Count
PD-L1
G
3D 700
2D 483
2D FMO 332
3D FMO 279
Count
PD-L2
H
3D 642
3D FMO 495
2D 384
2D FMO 319
Count
PD-L2
I
SUM149 + γδTc 2427
SUM149 741
SUM149 FMO 297
SUM149 + γδTc FMO 294
Count
PD-L1
J
SUM149 + γδTc 561
SUM149 318
SUM149 + γδTc FMO 149
SUM149 FMO 95
Count
PD-L1
K
PDX401 + γδTc 2358
PDX401 478
PDX401 + γδTc FMO 169
PDX401 FMO 116
Count
PD-L1
L
PDX401 + γδTc 1008
PDX401 927
PDX401 FMO 136
PDX401 + γδTc FMO 109
Count
PD-L1
M
SUM149 + γδTc 195
SUM149 143
SUM149 + γδTc FMO 110
SUM149 FMO 58
Count
PD-L2
N
SUM149 + γδTc 59
SUM149 47
SUM149 + γδTc FMO 49
SUM149 FMO 21
Count
PD-L2
O
SUM149 + γδTc 135
SUM149 69
SUM149 FMO 51
SUM149 + γδTc FMO 45
Count
PD-L2
P
PDX401 + γδTc 156
PDX401 113
PDX401 FMO 91
PDX401 + γδTc FMO 81
Count
PD-L2
Q
PDX401 + γδTc 115
PDX401 + γδTc FMO 83
PDX401 69
PDX401 FMO 55
Count
PD-L2
R
PDX401 + γδTc 23
PDX401 FMO 23
PDX401 21
PDX401 + γδTc FMO 14
Count
PD-L2
S
PDX401 + γδTc 40
PDX401 + γδTc FMO 41
PDX401 42
PDX401 FMO 39
Count
PD-L2
T
CD24-
CD24+
**
% Lysis
0
20
40
60
80
IgG
αPD-1
IgG
αPD-1

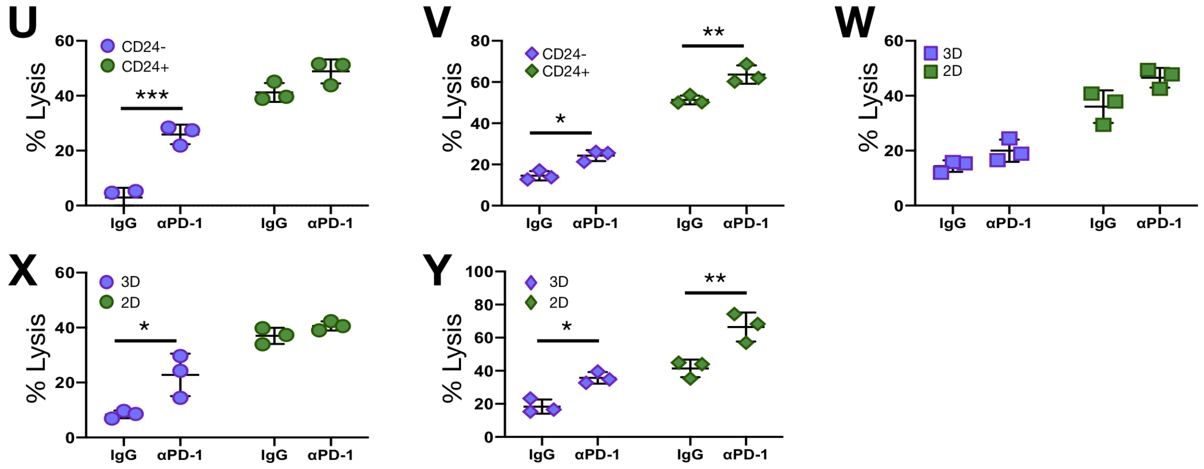


**Figure S4. Inhibitory ligands are expressed on breast cancer stem-like cells resistant to gamma delta T cell killing.** PD-L1 expression on SUM149 CD24- and CD24+ cells were detected by flow cytometry (A) Florescence minus one (FMO) controls are shown for replicate 1 and (B) replicate 2. (C) Histogram overlays of PD-L1 expression on SUM149 replicate 2. (D) Histogram overlays for PD-L1 expression on PDX 401 3D and 2D cells, replicate 1, (E) replicate 2 and (F) replicate 3 is shown. (G) Histogram overlays for PD-L2 expression on PDX 401 3D and 3D cells, replicate 1 and (H) replicate 2. Target cells were co-incubated with γδTc at a 1:1 ratio overnight and surface expression of PD-L1 was determined using flow cytometry. (I) Histogram overlays PD-L1 expression are shown for replicate 2, and (J) replicate 3. (K) Experiments done as in I-J but with PDX401 cells, replicate 2 and (L) replicate 3. (M) PD-L2 was also compared between co-incubated targets and targets alone on SUM149 replicate 1, (N) replicate 2, (O) replicate 3, and (P) PDX401 cells replicate 1, (Q) replicate 2, (R) replicate 3 and (S) replicate 4. γδTc were incubated with blocking antibody for PD-1 and then co-incubated with (T) SUM149 CD24-, CD24+ cells replicate 1, (U) replicate 2 and (V) replicate 3, or (W) PDX401 2D, PDX 3D cells replicate 1, (X) replicate 2 and (Y) replicate 3; at E: T 20:1 for 4 h. Donor numbers were reset for each set of experiments. Data are presented as mean ± SD in T-Y. Statistical tests employed were: (T-Y) One-way ANOVA followed by Tukey's post hoc test. *P<0.05, **P<0.01.

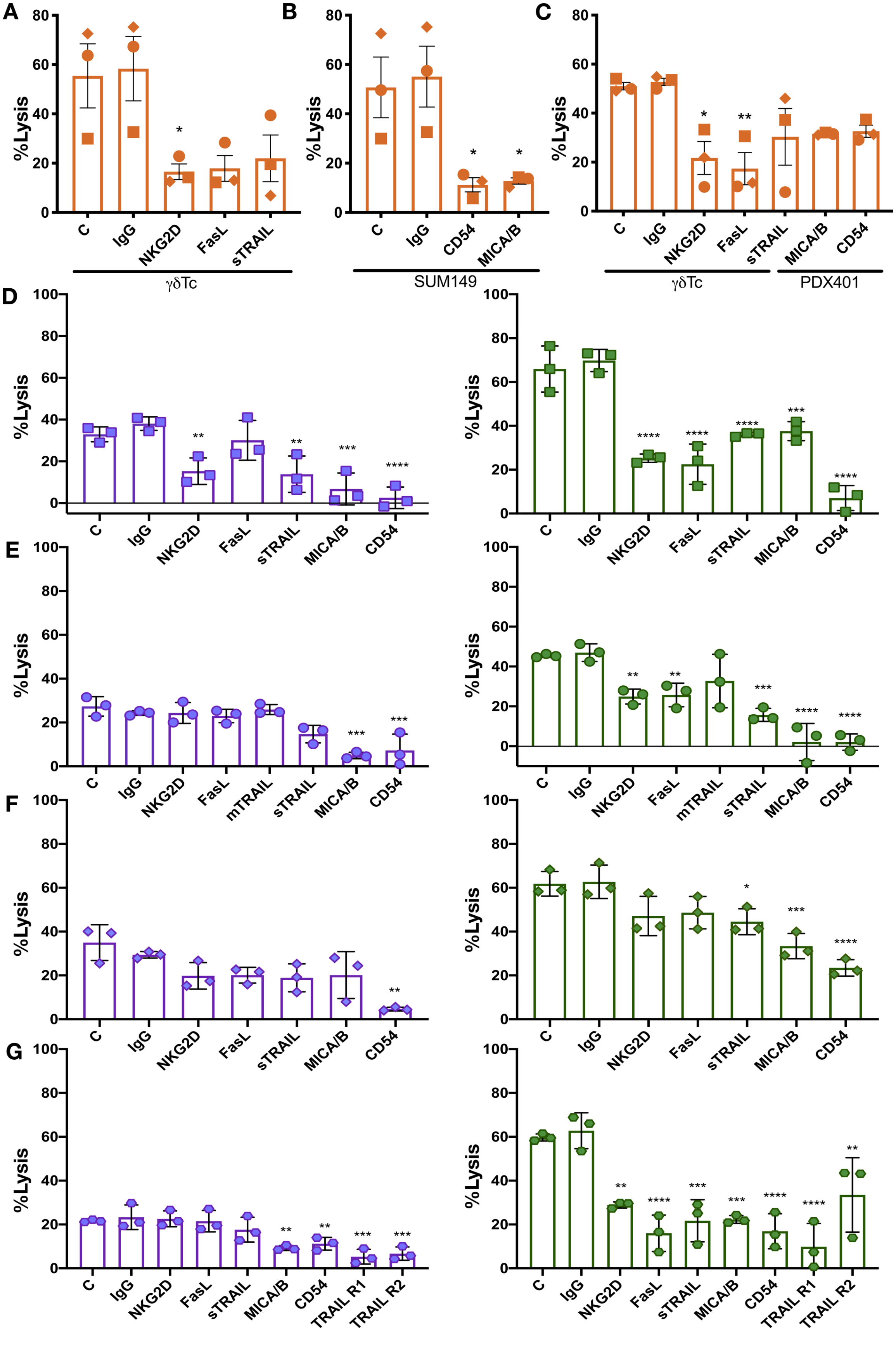
A
B
C
D
E
F
G
%Lysis
C
IgG
NKG2D
FasL
sTRAIL
mTRAIL
MICA/B
CD54
TRAIL R1
TRAIL R2
γδTc
SUM149
PDX401

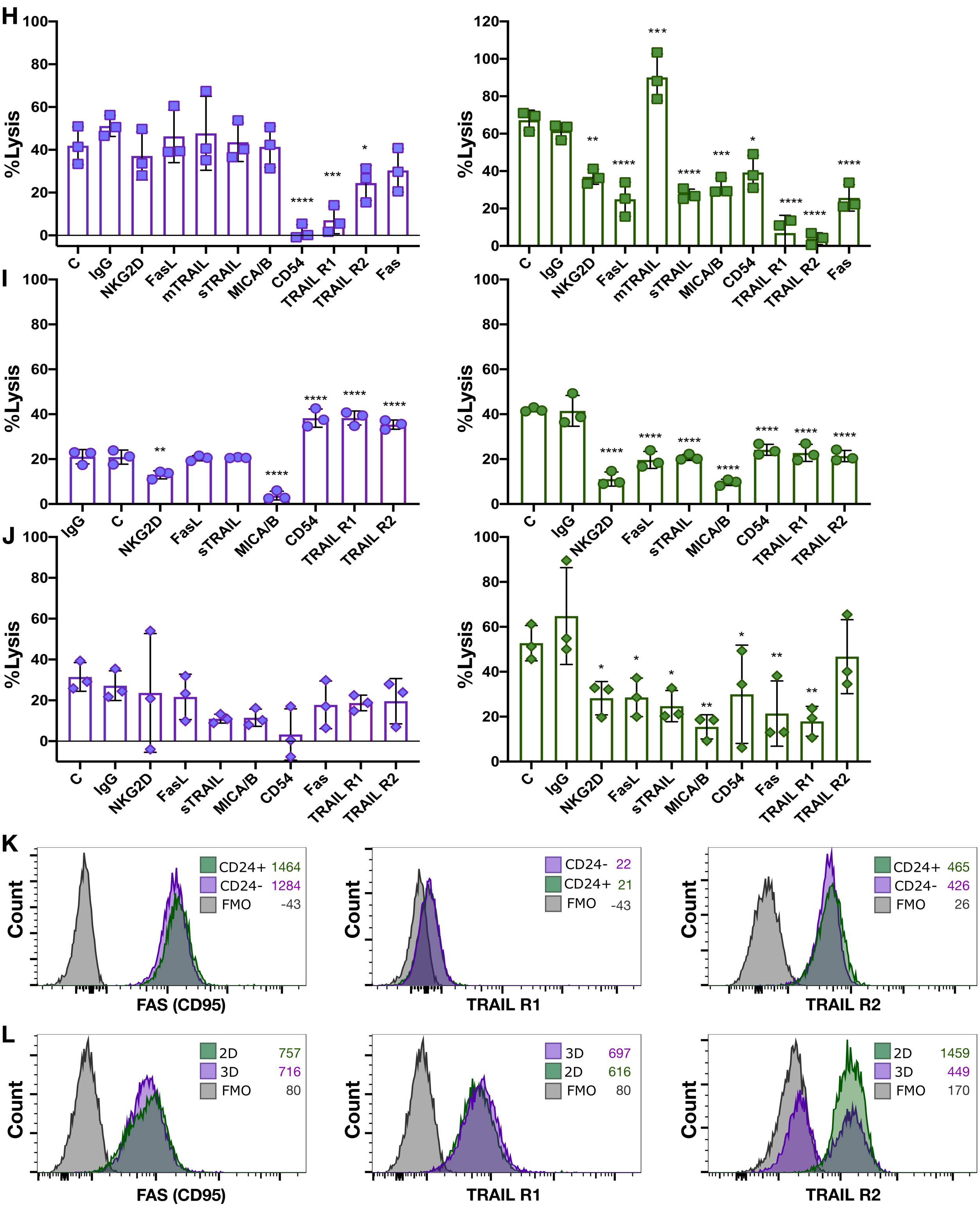
H
I
J
K
L
%Lysis
C
IgG
NKG2D
FasL
mTRAIL
sTRAIL
MICA/B
CD54
TRAIL R1
TRAIL R2
Fas
Count
FAS (CD95)
TRAIL R1
TRAIL R2
CD24+ 1464
CD24- 1284
FMO -43
CD24- 22
CD24+ 21
FMO -43
CD24+ 465
CD24- 426
FMO 26
2D 757
3D 716
FMO 80
3D 697
2D 616
FMO 80
2D 1459
3D 449
FMO 170

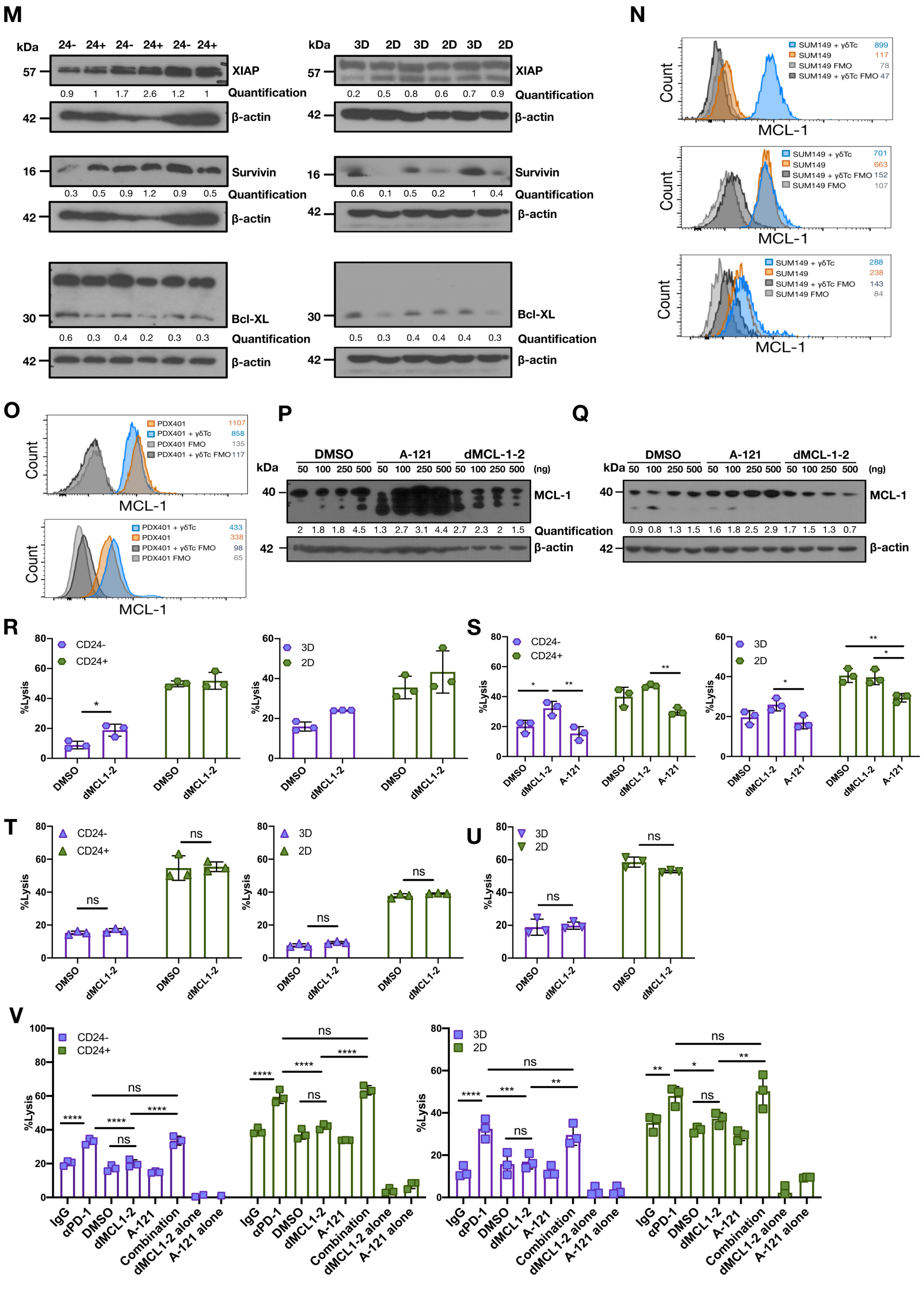
M
kDa 24- 24+ 24- 24+ 24- 24+
57 XIAP
0.9 1 1.7 2.6 1.2 1 Quantification
42 β-actin
16 Survivin
0.3 0.5 0.9 1.2 0.9 0.5 Quantification
42 β-actin
30 Bcl-XL
0.6 0.3 0.4 0.2 0.3 0.3 Quantification
42 β-actin
kDa 3D 2D 3D 2D 3D 2D
57 XIAP
0.2 0.5 0.8 0.6 0.7 0.9 Quantification
42 β-actin
16 Survivin
0.6 0.1 0.5 0.2 1 0.4 Quantification
42 β-actin
30 Bcl-XL
0.5 0.3 0.4 0.4 0.4 0.3 Quantification
42 β-actin
N
Count
MCL-1
SUM149 + γδTc 899
SUM149 117
SUM149 FMO 78
SUM149 + γδTc FMO 47
SUM149 + γδTc 701
SUM149 663
SUM149 + γδTc FMO 152
SUM149 FMO 107
SUM149 + γδTc 288
SUM149 238
SUM149 + γδTc FMO 143
SUM149 FMO 84
O
Count
MCL-1
PDX401 1107
PDX401 + γδTc 858
PDX401 FMO 135
PDX401 + γδTc FMO 117
PDX401 + γδTc 433
PDX401 338
PDX401 + γδTc FMO 98
PDX401 FMO 65
P
kDa DMSO A-121 dMCL-1-2
50 100 250 500 50 100 250 500 50 100 250 500 (ng)
40 MCL-1
2 1.8 1.8 4.5 1.3 2.7 3.1 4.4 2.7 2.3 2 1.5 Quantification
42 β-actin
Q
kDa DMSO A-121 dMCL-1-2
50 100 250 500 50 100 250 500 50 100 250 500 (ng)
40 MCL-1
0.9 0.8 1.3 1.5 1.6 1.8 2.5 2.9 1.7 1.5 1.3 0.7
42 β-actin
R
%Lysis
CD24-
CD24+
3D
2D
DMSO dMCL1-2 DMSO dMCL1-2
S
%Lysis
CD24-
CD24+
3D
2D
DMSO dMCL1-2 A-121 DMSO dMCL1-2 A-121
T
%Lysis
CD24-
CD24+
3D
2D
ns
DMSO dMCL1-2 DMSO dMCL1-2
U
%Lysis
3D
2D
ns
DMSO dMCL1-2 DMSO dMCL1-2
V
%Lysis
CD24-
CD24+
3D
2D
ns
IgG αPD-1 DMSO dMCL1-2 A-121 Combination dMCL1-2 alone A-121 alone

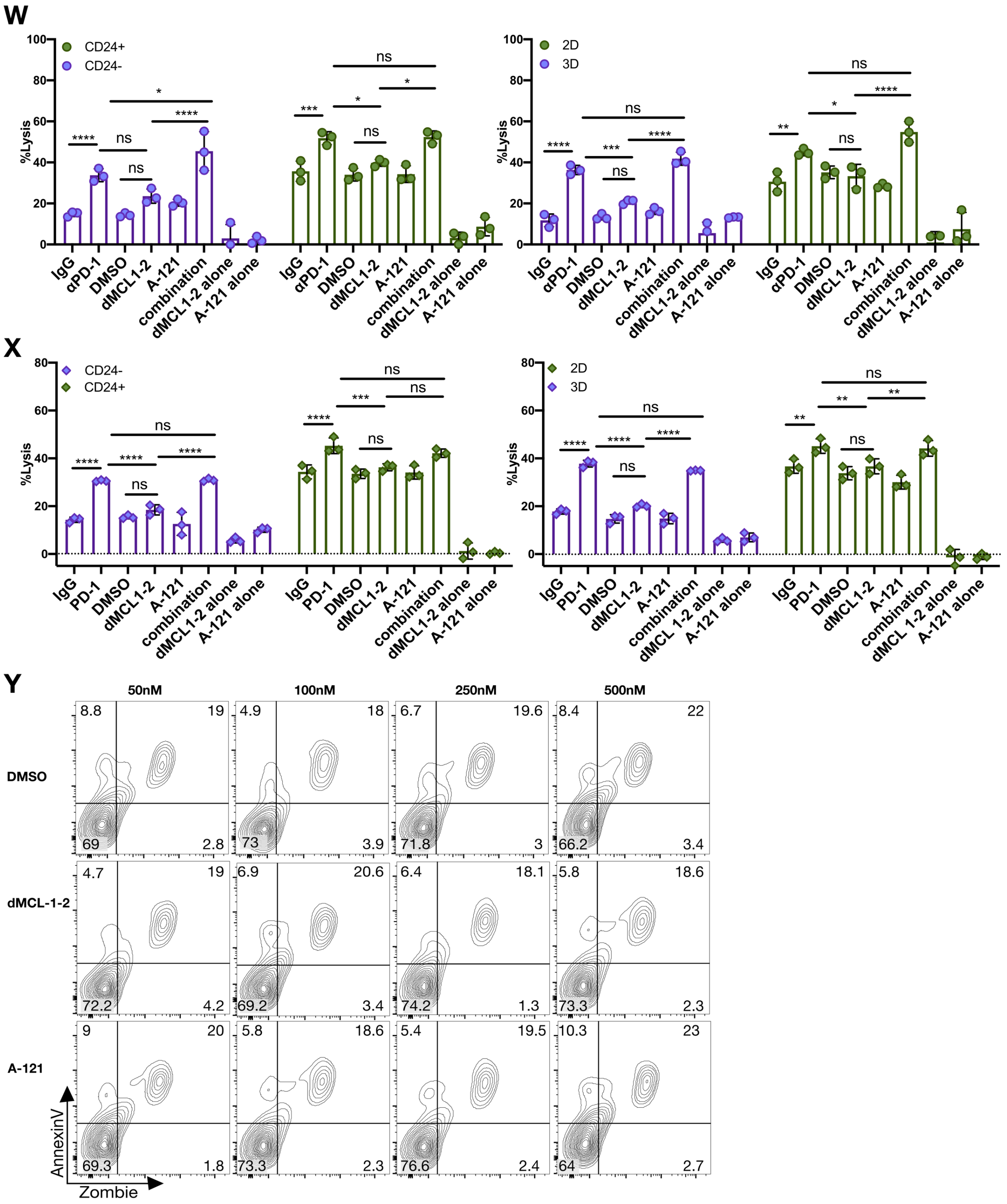


**Figure S5. Fas-FasL pathway dysfunctional in breast cancer ste m-like cells. (A)** γδTc or **(B)** SUM149 cells were incubated with the indicated antibodies for 30 mins and then co-incubated at E:T 20:1 for 4h in a Calcein AM release assay to assess cytotoxicity **(C)** Blocking and cytotoxicity assay as in A and B, but with γδTc and PDX401 target cells. **(D)** SUM149 were sorted into $CD24^-$ (left panel) and $CD24^+$ (right panel) fractions, incubated with the indicated antibodies and then co-incubated with γδTc at E: T 20:1 for 4h from donor 1, **(E)** donor 2, **(F)** donor 3 and **(G)** donor 4. **(H)** Similarly, PDX401 3D (left panel) and 2D cells (right panel) were incubated with antibodies followed by γδTc from donor 1, **(I)** donor 2 and **(J)** donor 3. mTRAIL stands for membrane bound TRAIL and sTRAIL stands for soluble TRAIL. **(K)** Histogram overlays of indicated receptors on SUM149 CD24- and CD24+ cells and **(L)** PDX 3D and 2D cells are shown. **(M)** Expression of indicated anti-apoptotic proteins was determined using western blot analysis of lysates from three sets of sorted SUM149 $CD24^-$ and $CD24^+$ cells (left panel) and PDX401 3D and 2D cells (right panel). Molecular weight markers are shown on the left in kDa; β-actin loading controls and quantifications are shown below corresponding panels. For SUM149 cells, XIAP and Survivin were probed on the same blot and for PDX401 cells, Survivin and Bcl-XL were probed on the same blot. **(N)** SUM149 cells were subject to a γδTc cytotoxicity assay at 1:1 (E: T) ratio overnight. Target cells were then stained for intracellular MCL-1 protein that was detected *via* flow cytometry. Histogram overlays of experiments done with γδTc from donor 2 (top), donor 3 (middle) and donor 4 (bottom) are shown. **(O)** Experiment done as in N with PDX401 cells and γδTc from donor 2 (top) and 3 (bottom) are shown. **(P)** SUM149 and **(Q)** PDX401 that were treated with the MCL-1 degrader dMCL1-2 and failed inhibitor A-1210477 (A-121) are depicted. Molecular weight markers are shown on the left in kDa; β -actin loading controls and quantifications are shown below corresponding panels. **(R)** Sorted SUM149 CD24+ and CD24- cells (left) were co-incubated with γδTc derived from donor 1 at E: T 20:1 in a 4-hour Calcein AM cytotoxicity assay; indicated treatments were added to the co-incubation. The same experiment was performed in parallel with PDX401 3D and 2D cells (right). Experiments done with γδTc derived from **(S)** donor 2, **(T)** donor 3, **(U)** donor 4, **(V)** donor 5, **(W)** donor 6 and **(X)** donor 7 are shown. **(Y)** γδTc were incubated with the indicated concentration of DMSO, dMCL1-2 and A-121 for 4h and viability was accessed using Zombie Aqua and AnnexinV staining. Donor numbers were reset for each set of experiments. Data are presented as mean ± SEM in A-C and mean ± SD in D-J and R-X. Statistical tests employed were: (A-J, R-U) One-way ANOVA followed by Sidak's post hoc test, (V-X) Two-way ANOVA followed by Tukey's post hoc test. $*P<0.05$, $**P<0.01$, $***P<0.001$, $****P<0.0001$.

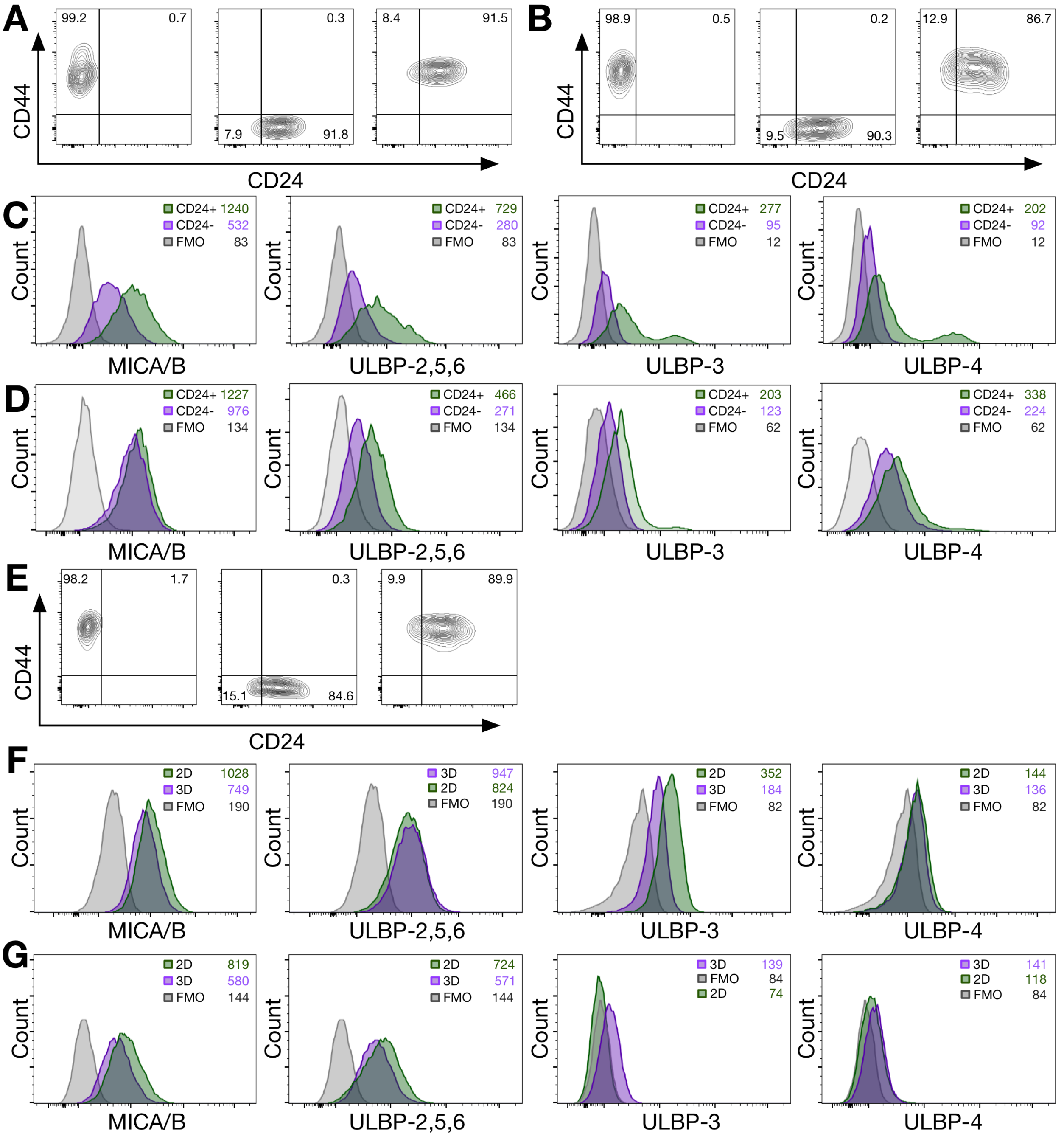


**Figure S6. MICA/B is downregulated on the surface of breast cancer stem-like cells. (A)** NKG2D ligand expression was determined on SUM149 CD24- and CD24+ cells by flow cytometry. Florescence minus one (FMO) controls used to set gates for CD44 and CD24 for replicate 1 and **(B)** replicate 2. **(C)** Histogram overlays of NKG2D ligand expression on SUM149 CD24- and CD24+ cells for replicate 2 and **(D)** replicate 3. **(E)** FMO controls are also shown for replicate 3. **(F)** Histogram overlays of NKG2D ligand expression on PDX401 3D and 2D cells for replicate 2 and **(G)** replicate 3.

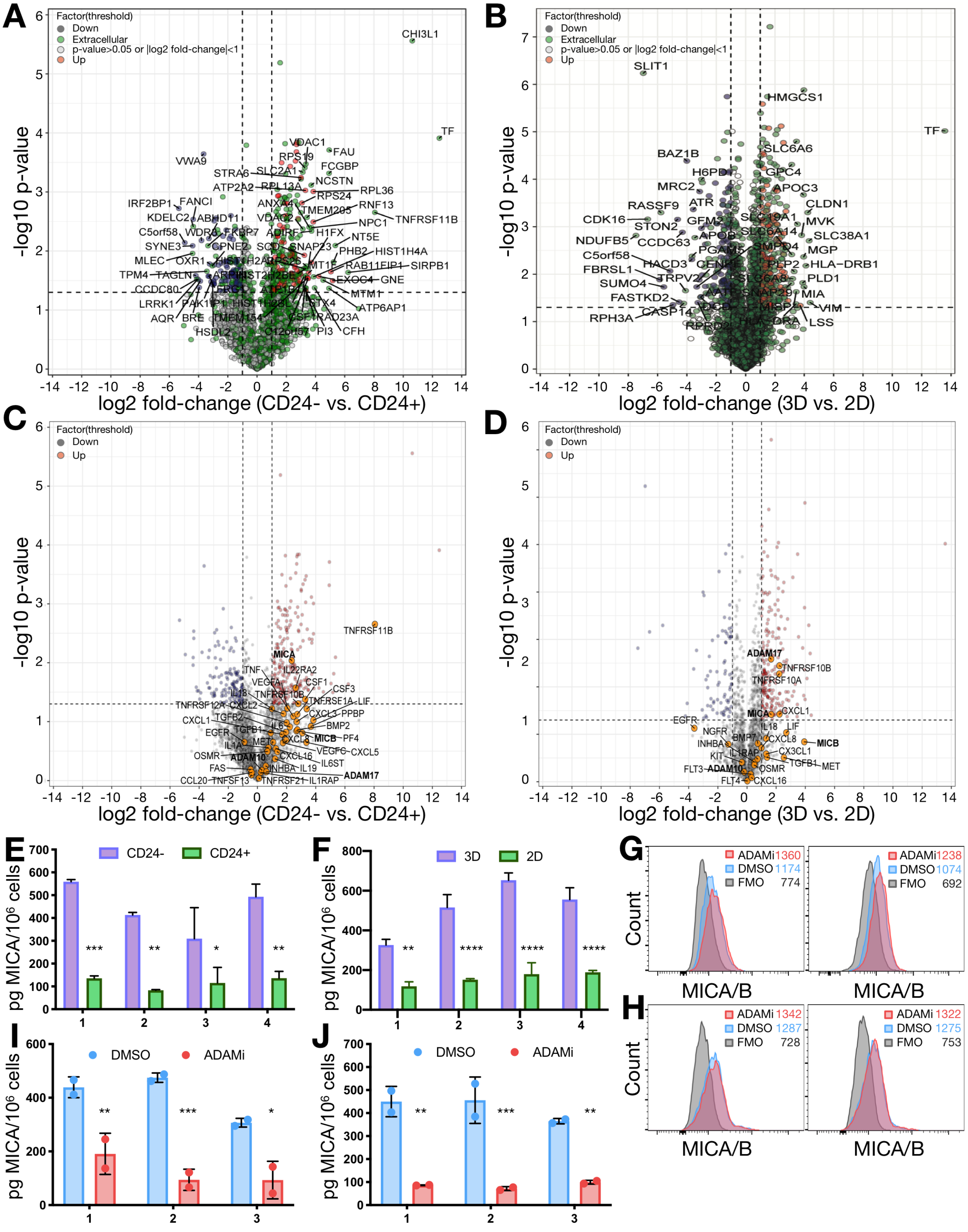
A
Factor(threshold)
Down
Extracellular
p-value>0.05 or |log2 fold-change|<1
Up
-log10 p-value
log2 fold-change (CD24- vs. CD24+)
CHI3L1
TF
VWA9
TNFRSF11B
B
log2 fold-change (3D vs. 2D)
SLIT1
HMGCS1
TF
BAZ1B
SLC6A6
C
MICA
TNFRSF11B
MICB
ADAM10
ADAM17
D
ADAM17
TNFRSF10B
TNFRSF10A
MICA
MICB
EGFR
ADAM10
E
CD24-
CD24+
pg MICA/10^6 cells
F
3D
2D
G
ADAMi 1360
DMSO 1174
FMO 774
ADAMi 1238
DMSO 1074
FMO 692
Count
MICA/B
H
ADAMi 1342
DMSO 1287
FMO 728
ADAMi 1322
DMSO 1275
FMO 753
I
DMSO
ADAMi
J

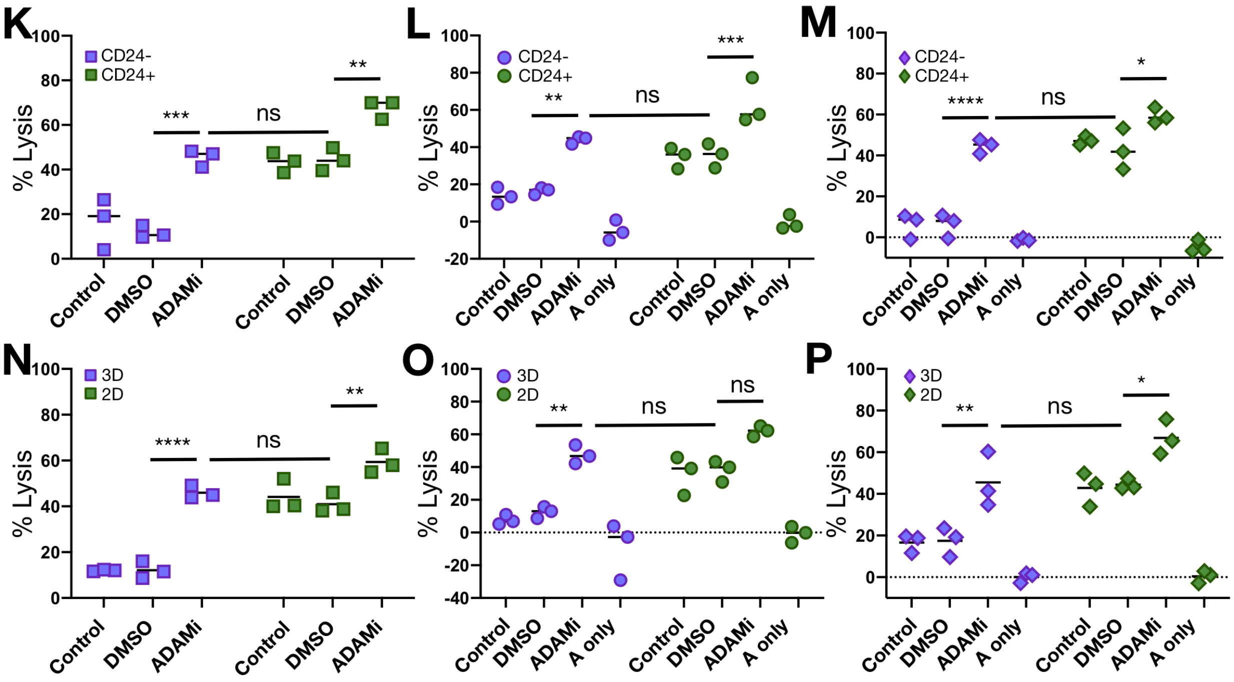


**Figure S7. The ADAM inhibitor GW280264X prevents MICA shedding and enhances γδ T cell cytotoxicity against breast cancer stem-like cells.** Mass spectrometry analysis of SUM149 $CD24^-$ and $CD24^+$ cells (n=3) and PDX401 3D (n=3) and 2D (n=6) conditioned media. **(A)** Volcano plot of quantified "extracellular" proteins. Log2fold-change indicates proteins that showed the most significant differential expression between SUM149 $CD24^-$ and $CD24^+$CM, and **(B)** PDX401 3D and 2D CM. **(C)** Several cytokines, chemokines and other relevant factors differentially secreted by sorted SUM149 cells and **(D)** PDX401 3D and 2D cells. Vertical and horizontal dotted lines indicate Log2 fold-changes ≥2 and the −Log10 p-value cut off for p<0 respectively. **(E)** MICA shed by SUM149 sorted cells and **(F)** 2D and 3D PDX401 was determined by ELISA. Replicate number is indicated on x-axis. **(G)** Targets were treated with ADAM protease inhibitor GW280264X (ADAMi) for 4h, and MICA/B surface expression was determined *via* flow cytometry for SUM149 replicate 2 (left) and replicate 3 (right), as well as **(H)** PDX401 replicate 2 (left) and (right). **(I)** Soluble MICA in media after treatment was determined using ELISA for SUM149 and **(J)** PDX401 cells; replicate numbers are indicated on x-axis. (**K**) SUM149 $CD24^-$ and $CD24^+$ cells were incubated with ADAMi and then co-incubated with γδTc at E:T 20:1 for 4h. Results from experiments done with γδTc from donor 1, **(L)** donor 2 and **(M)** donor 3. **(N)** Experiments done as in K-M but with PDX401 2D and 3D cells and γδTc from donor 1, **(O)** donor 2 and **(P)** donor 3. "A only" stands for target cells treated with ADAMi only. Donor numbers were reset for each set of experiments. Data are presented as mean ± SD (E,F,I,J) or median (K-P). Statistical tests employed were: (E,F,I,J) Two-way ANOVA followed by Bonferroni's post hoc test; (K-P) Two-way ANOVA followed by Tukey's post hoc test; *P<0.05, **P<0.01, ***P<0.001, ****P<0.0001.

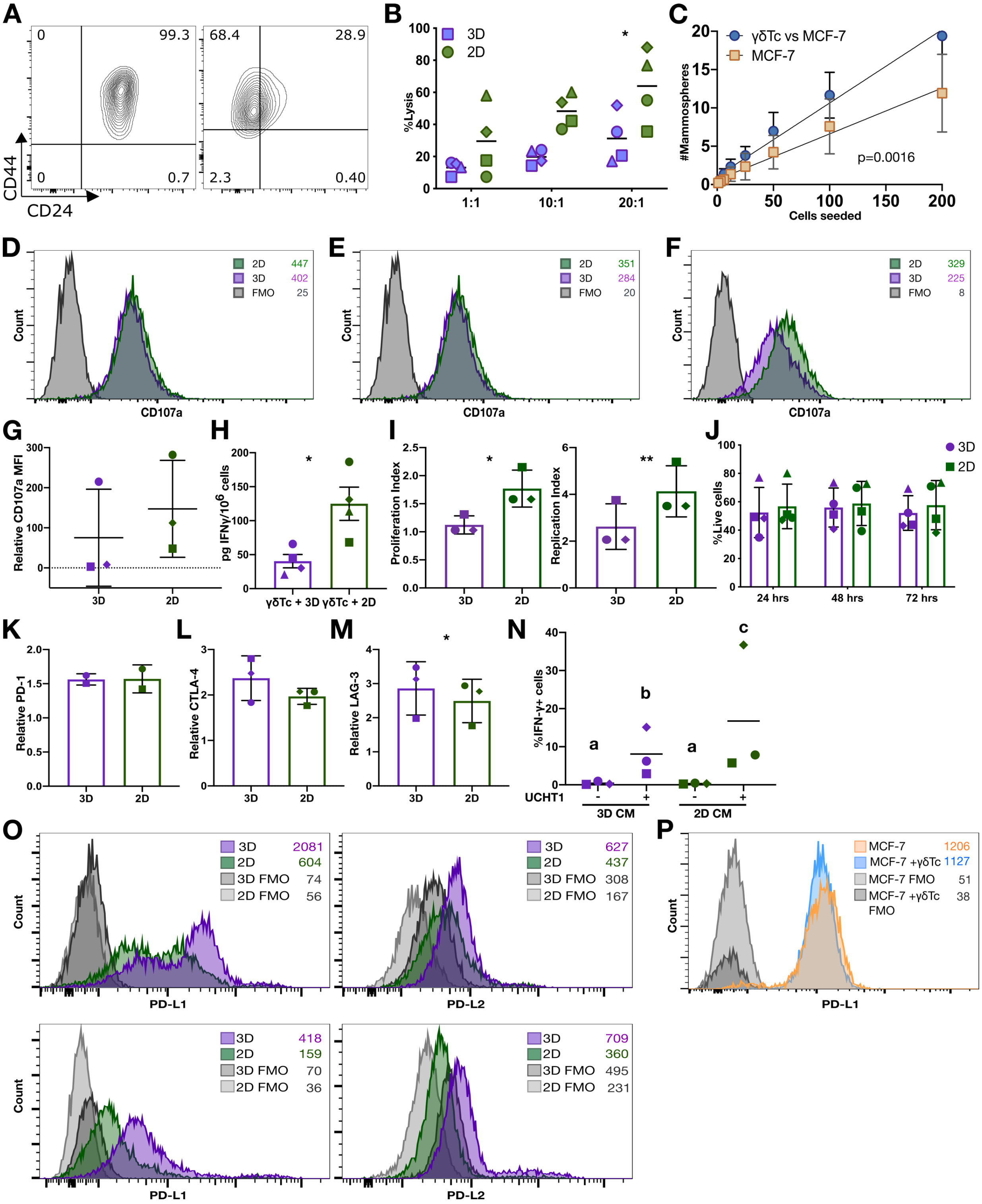
A
0
99.3
0
0.7
68.4
28.9
2.3
0.40
CD44
CD24
B
%Lysis
3D
2D
1:1
10:1
20:1
*
C
γδTc vs MCF-7
MCF-7
#Mammospheres
Cells seeded
p=0.0016
D
2D 447
3D 402
FMO 25
Count
CD107a
E
2D 351
3D 284
FMO 20
Count
CD107a
F
2D 329
3D 225
FMO 8
Count
CD107a
G
Relative CD107a MFI
3D
2D
H
pg IFNγ/10^6 cells
γδTc + 3D
γδTc + 2D
*
I
Proliferation Index
Replication Index
3D
2D
*
**
J
%Live cells
3D
2D
24 hrs
48 hrs
72 hrs
K
Relative PD-1
3D
2D
L
Relative CTLA-4
3D
2D
M
Relative LAG-3
3D
2D
*
N
%IFN-γ+ cells
a
b
a
c
UCHT1
3D CM
2D CM
O
3D 2081
2D 604
3D FMO 74
2D FMO 56
Count
PD-L1
3D 627
2D 437
3D FMO 308
2D FMO 167
PD-L2
3D 418
2D 159
3D FMO 70
2D FMO 36
Count
PD-L1
3D 709
2D 360
3D FMO 495
2D FMO 231
PD-L2
P
MCF-7 1206
MCF-7 +γδTc 1127
MCF-7 FMO 51
MCF-7 +γδTc FMO 38
Count
PD-L1

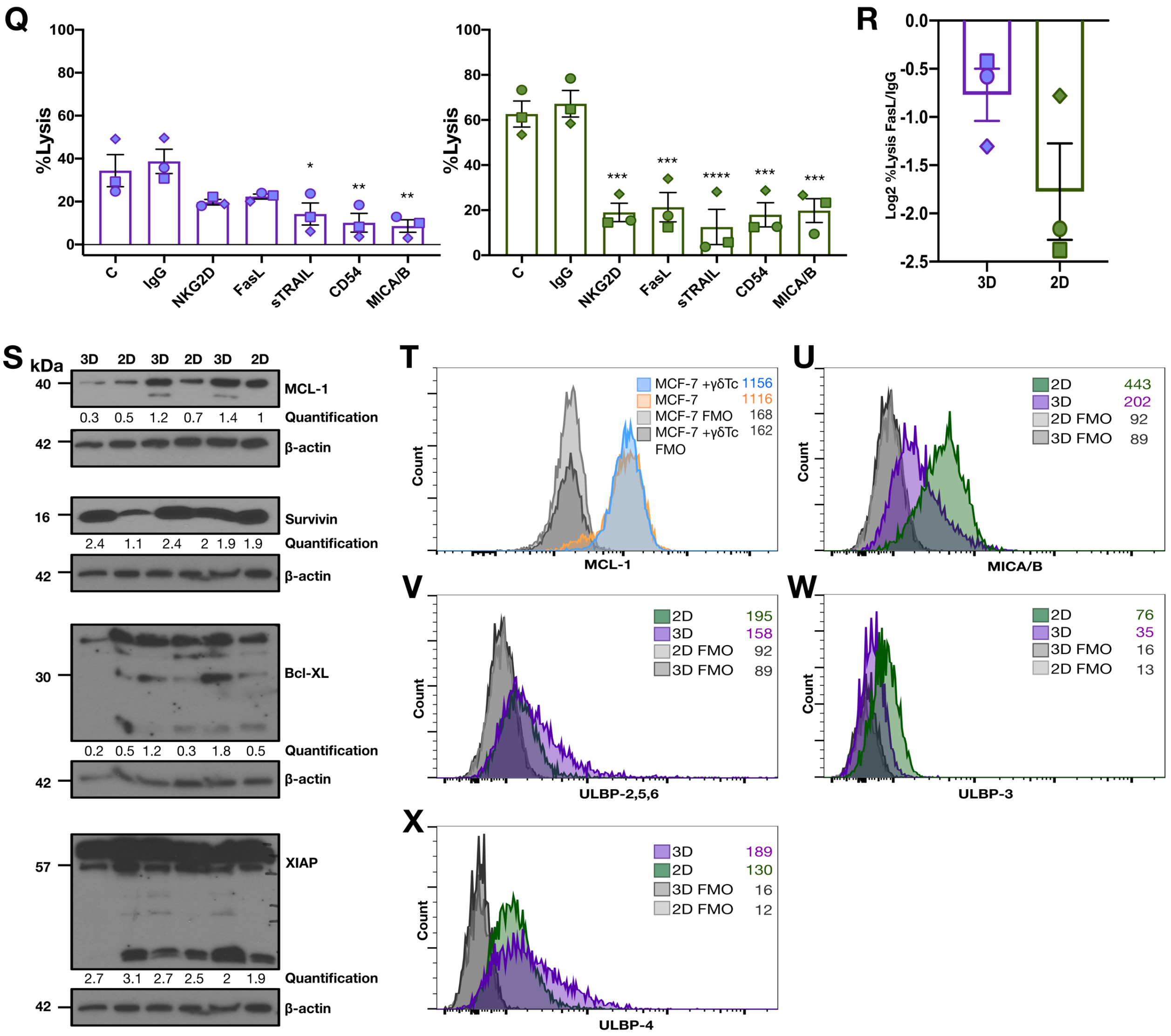


**Figure S8. The MCF-7 stem-like cell population behaves very similar to SUM149 and PDX401 stem-like cells.** **(A)** Second generation mammospheres (3D) and adherent (2D) MCF-7 cells were dissociated, then CD44 and CD24 expression were determined by flow cytometry (n=1). **(B)** Second generation mammospheres (3D) and adherent (2D) MCF-7 were dissociated, filtered into single cell suspensions and used as targets in Calcein AM cytotoxicity assays (n=4) **(C)** MCF-7 cells were co-cultured with γδTc at 1:1 overnight, γδTc were then removed, and mammosphere forming potential of targets determined over two generations (cumulative, n=3). **(D)** Overlay of CD107a expression (degranulation) for γδTc incubated with MCF-7 3D and 2D cells for γδTc derived from donor 1, **(E)** donor 2 and **(F)** donor 3. **(G)** Cumulative results (n=3) for MFI of CD107a. **(H)** IFN-γ ELISA was performed on conditioned media (CM) from γδTc co-incubated with 3D or 2D MCF-7 for 24h (n=4). **(I)** Using the FlowJo 10.5.3 Proliferation Modeling Tool, proliferation index values and replication index values were calculated for three donor cultures treated with MCF-7 3D and 2D CM; cumulative data are shown (n=3). **(J)** γδTc viability after incubation with MCF-7 3D or 2D conditioned medium (CM) was assessed using AnnexinV and Zombie Aqua at specified time points; cumulative results are shown (n=4). **(K)** γδTc were incubated with MCF-7 3D and 2D cells for 24h; cumulative results for expression of co-inhibitory receptors PD-1 (n=2), **(L)** CTLA-4 (n=3), and **(M)** LAG-3 (n=4) on γδTc seven days later. **(N)** γδTc incubated with MCF-7 3D and 2D CM were treated with anti-CD3 antibody overnight and IFN-γ expression was determined *via* intracellular flow cytometry; cumulative results for % IFN-γ producing cells (n=3). **(O)** Histogram overlays showing PD-L1 expression (top left) and PD-L2 expression (top right) on second generation 3D and adherent 2D MCF-7 cells. The same experiment with a different biological replicate is shown in the bottom panels. **(P)** MCF-7 cells were treated with γδTc at 1:1 overnight, and surface expression of PD-L1 was compared between treated and untreated cells. **(Q)** 3D and 2D cells were incubated with the indicated antibodies and then co-incubated with γδTc at E:T 20:1 for 4h (n=3). Cumulative results for three experiments are depicted. **(R)** The decrease in target cell lysis upon FasL blocking (n=3) is indicated for the experiment. **(S)** Expression of anti-apoptotic proteins was determined using western blot analysis of lysates from three different sets of MCF-7 3D and 2D cells as indicated. Molecular weight (MW) markers are shown on the left; β-actin loading controls and quantification of band intensities normalized to β-actin controls are shown below corresponding panels. Survivin and XIAP were probed on the same blot **(T)** MCF-7 were subject to a γδTc cytotoxicity assay at 1:1 (E:T) ratio overnight. Post-cytotoxicity, target cells were stained for intracellular MCL-1 expression, detected *via* flow cytometry. (**U**) Histogram overlays of MICA/B (**V**) ULBP-2,5,6 **(W)** ULBP-3 and **(X)** ULBP-4 expression on MCF-7 3D and 2D cells is shown. Data are presented as: mean in B and N; mean ± SD in C, G, and I-M; mean ± SEM in H, Q, and R. Statistical tests employed were: (B, J); Two-way ANOVA followed Sidak's post hoc test for multiple comparisons between groups; (C) simple linear regression; (G, L) One-tailed Wilcoxon test; (H) Two-tailed ratio paired t-test; (I) Two-tailed paired t-test; (K, M, R) One-tailed paired t-test; (N) One-tailed ratio paired t-test, b-c, p= 0.0452; (Q) One-way ANOVA followed by Bonferroni's post hoc test. $^*p<0.05$, $^{**}p<0.01$, $^{***}p<0.001$, $^{****}p<0.0001$.

**Table S1** Subset percentages and purities of donor-derived γδ T cell cultures. Donors are listed in order of appearance of the first γδ T cell culture from that donor in the figures. Flow refers to the day cells were harvested, stained with fixable viability dye followed by the following antibodies: pan γδ TCR, Vδ1 TCR and Vδ2 TCR, after which they were washed and fixed; samples were acquired by flow cytometry within one week. The purity is calculated as the sum of %Vδ1, %Vδ2 and %γδTCR+Vδ1-Vδ2-. Day indicates the day on which the experiment(s) listed in the figure panel ended and cells were harvested for analysis. Their use in experiments for figures in this manuscript are listed in the order in which they appear.

| ID | Flow | %Vδ1 | %Vδ2 | % γδ+Vδ1-Vδ2- | % Purity | Day | Figure(s) |
|---|---|---|---|---|---|---|---|
| 1-1 | 19 | 24.1 | 34.1 | 7.2 | 65.4 | 20<br>21 | 1A, S1M-1<br>5A-C, S5G |
| 1-2 | 19 | 28.5 | 50.5 | 6.0 | 85.0 | 21 | 1D, S1P-2, S3I, S8J |
| 1-3 | 21 | 43.3 | 43.6 | 7.3 | 94.3 | 18 | 1E |
| 1-4 | 21 | 69.4 | 12.8 | 8.4 | 90.6 | 19<br>21 | S3J<br>5N, S5W |
| 1-5 | 22 | 69.0 | 6.6 | 16.9 | 92.5 | 20 | S8H |
| 1-6 | 21 | 26.3 | 50.6 | 3.5 | 80.4 | 19 | S8Q,R |
| 2-1 | 22 | 1.8 | 92.4 | 1.6 | 95.8 | 22 | 1A, S1M-2 |
| 2-2 | 21 | 0.9 | 94.3 | 1.4 | 96.6 | 19<br>20 | S1Q-1<br>S5C |
| 2-3 | 21 | 0.8 | 94 | 1.6 | 96.4 | 18<br>21 | 2A,B, S2C, S2G, S8F,G<br>3L-O, S3B,E,G,L,N, S8L,M, S8I |
| 2-4 | 21 | 1.3 | 88.1 | 2.7 | 92.1 | 21<br>20 | 3P,Q, 5D-F, S3O,R,S5J<br>4E-J, S4O,S |
| 2-5 | 21 | 1.1 | 94.4 | 1.3 | 96.8 | 21 | S8H |
| 3-1 | 21 | 11.0 | 77.9 | 3.7 | 92.6 | 20 | 1A, S1M-3 |
| 3-2 | 21 | 53.5 | 23.8 | 13.2 | 90.5 | 21 | 1D, S1P-1 |
| 3-3 | 24 | 29.0 | 50.5 | 13.4 | 92.9 | 20<br>21<br>19 | 2B, S2D<br>S3C,F,G, S8I<br>S5C |
| 4-1 | 21 | 52.2 | 4.0 | 14.7 | 70.9 | 21 | 1B, S1N-1, S8Q,R |
| 4-2 | 21 | 8.0 | 77.1 | 1.5 | 86.6 | 21<br>20<br>22 | 2C,E,F<br>S8H<br>S8J |
| 4-3 | 21 | 30.0 | 29.8 | 22.0 | 81.8 | 20 | 2E,F, S2I |
| 5-1 | 22 | 41.2 | 47.1 | 7.0 | 95.3 | 22 | 1B, S1N-2 |
| 5-2 | 21 | 23.2 | 69.4 | 5.5 | 98.1 | 20<br>21 | S5A,B, S5A,B<br>5A-C, S5E |
| 6-1 | 21 | 29.2 | 56.4 | 6.1 | 91.7 | 20 | 1B,C, S1N-3,O-1, S8C |
| 6-2 | 22 | 13.4 | 71.5 | 5.2 | 90.1 | 22 | S3I |
| 7-1 | 21 | 21.9 | 73.4 | 4.2 | 99.5 | 20<br>18 | 1B, S1N-4, S8Q,R<br>1C, S1O-2 |
| 7-2 | 21 | 40.0 | 44.2 | 7.7 | 91.9 | 18 | S1Q-2 |
| 7-3 | 21 | 4.9 | 1.2 | 91.1 | 97.2 | 19 | 2E,F, S2H, S8H |
| 7-4 | 21 | 0.6 | 81.3 | 8.6 | 90.5 | 18<br>21 | 3T,U, S3U,W,Y, S8N<br>4I,J, 5I,L, S4M,Q, S5N-3,O-2 |
| 7-5 | 20 | 14.0 | 74.4 | 2.1 | 90.5 | 21 | S5A |

| ID | Flow | %Vδ1 | %Vδ2 | % γδ+Vδ1-Vδ2- | % Purity | Day | Figure(s) |
|---|---|---|---|---|---|---|---|
| 7-6 | 18 | 14.8 | 67.8 | 2.4 | 85.0 | 22 | S8B |
| 8-1 | 20 | 28.0 | 52.3 | 7.2 | 87.5 | 20 | 1C, S1O-3 |
| 8-2 | 19 | 16.4 | 74.6 | 2.3 | 93.3 | 20<br>21 | S3H-J, S8J<br>1D, S1P-4 |
| 8-3 | 20 | 32.3 | 27.4 | 12.3 | 72.0 | 22 | 4J, S4P |
| 8-4 | 20 | 45.1 | 33.3 | 6.9 | 85.3 | 19 | 7J,K, S7L,O |
| 9-1 | 20 | 17.4 | 71.7 | 2.3 | 91.4 | 20 | 1D, S1P-3, S3I, S8J |
| 9-2 | 21 | 1.6 | 87.2 | 9.0 | 97.8 | 20 | 1F |
| 9-3 | 22 | 8.4 | 85.2 | 3.7 | 97.3 | 19<br>22 | 2D, 5D-F, S5H<br>4F, S4I, S8P, S8T |
| 9-4 | 21 | 7.0 | 84.5 | 5.1 | 96.6 | 18<br>20<br>22 | 3J-O, S3K,M, S8K-M<br>S5C<br>S5U |
| 9-5 | 22 | 4.0 | 87.2 | 2.8 | 94.0 | 22 | 4L, S4Y |
| 9-6 | 21 | 2.5 | 92.4 | 3.3 | 98.2 | 20 | 7J,K, S7K,N |
| 10-1 | 21 | 1.9 | 94.1 | 1.7 | 97.7 | 21 | 2A,B, S2A, S2E, S8D,G |
| 11-1 | 21 | 10.3 | 75.9 | 3.3 | 89.5 | 21 | 2A,B, S2B, S2F, S8E,G |
| 11-2 | 18 | 11.1 | 76.9 | 2.5 | 90.5 | 20 | 4K,L, S2J, S4V,X, S5T |
| 11-3 | 21 | 16.5 | 60.8 | 11.3 | 88.6 | 20 | 5D-F, S5I |
| 12 | 21 | 60.2 | 6.0 | 20.3 | 86.5 | 19 | S2K |
| 13-1 | 21 | 9.4 | 81.7 | 4.8 | 95.9 | 21 | 3A-O, S3A,D,G, S8I,K-M |
| 13-2 | 20 | 17.6 | 72.0 | 2.1 | 91.7 | 19<br>21 | S3J<br>5M |
| 13-3 | 21 | 4.0 | 87.6 | 3.5 | 95.1 | 18<br>20 | 3R-U, S8N<br>4F,H-L, 5I,L, S4J,L,N,R,U,W, S5N-2,O-1,R |
| 13-4 | 21 | 21.1 | 64.3 | 1.8 | 87.2 | 20 | 4H, S4K |
| 13-5 | 21 | 46.0 | 42.2 | 7.4 | 95.6 | 20 | S5S |
| 13-6 | 20 | 16.2 | 79.2 | 2.2 | 97.6 | 20 | 7J,K, S7M,P |
| 14 | 20 | 4.5 | 88.0 | 3.9 | 96.4 | 20 | 3P,Q, 5H,I,K,L, S3P,S |
| 15 | 21 | 0.1 | 87.9 | 2.6 | 90.6 | 20<br>21 | 3P,Q, 5I,N, S3Q,T, S5N-1<br>S5V |
| 16 | 20 | 17.8 | 68.5 | 3.4 | 89.7 | 20 | 5A-C, S5A,B,D |
| 17 | 20 | 16.4 | 62.5 | 4.8 | 83.7 | 19<br>20 | 5A-C, S5B, S5F<br>S8C |
| 18 | 23 | 9.9 | 65.9 | 9.4 | 85.2 | 20 | 5N, S5X |
| 19-1 | 20 | 70.0 | 14.9 | 11.5 | 96.4 | 21 | S8B |
| 19-2 | 21 | 49.1 | 32.2 | 8.7 | 90.0 | 20 | S8B |
| 20-1 | 21 | 49.4 | 32.0 | 7.3 | 88.7 | 21 | S8B |
| 20-2 | 20 | 24.4 | 64.6 | 7.5 | 96.5 | 19 | S8C |

**Table S2.** Statistical tests employed and resulting p-values for experiments shown in Figures 1-7. Bonf=Bonferroni's multiple comparisons test; E:T=effector:target ratio; F=failed; L=left panel; R=right panel; RM=repeated measures; S-W=Shapiro-Wilk Normality test; Tukey=Tukey's multiple comparisons test.

| Figure | S-W | Test | Sample Comparison | p-value |
|---|---|---|---|---|
| 1A | All passed | Two-way ANOVA | CD24-/CD24+ | 0.0054 |
| 1B | All passed | Two-way ANOVA, Sidak | 3D/2D 10:1 | 0.0261 |
| 1C | All passed | Simple Linear Regression | γδTc vs. SUM149/SUM149 | 0.0093 |
| 1D | All passed | Two-way ANOVA, Sidak | 3D/2D 20:1 | 0.0666 |
| 1E | All failed | Simple Linear Regression | γδTc vs. PDX401/PDX401 | <0.0001 |
| 1F | All passed | Unpaired 1-tailed t-test | SUM149 post-γδTc/SUM149 | 0.0110 |
| 2A | All passed | Paired 2-tailed t-test | CD24-/CD24+ | 0.0361 |
| 2B | F: 2D | Wilcoxon | 3D/2D | 0.1250 |
| 2C | n=2, too small | One-way ANOVA, Tukey | γδTc + CD24-/γδTc, a-a | 0.6871 |
| | | | γδTc + CD24+/γδTc, a-b | <0.0001 |
| | | | γδTc + CD24-/ γδTc + CD24+, a-b | <0.0001 |
| 2D | n=2, too small | One-way ANOVA, Tukey | γδTc + 3D/γδTc, a-a | 0.9994 |
| | | | γδTc + 2D/γδTc, a-b | 0.0011 |
| | | | γδTc + 3D/ γδTc + 2D, a-b | 0.0010 |
| 2E | All passed | 1-tail ratio paired t-test | γδTc + CD24-/ γδTc + CD24+ | 0.0071 |
| 2F | F: γδTc + 3D | Wilcoxon | γδTc + 3D/ γδTc + 2D | 0.1250 |
| 3E | All passed | One-way ANOVA, Sidak | CD24-/CD24+ | 0.7216 |
| | | | 3D/2D | 0.9414 |
| 3J | n=2, too small | No test performed | | |
| 3K | n=2, too small | No test performed | | |
| 3L | All passed | Paired 1-tailed t-test | CD24-/CD24+ | 0.2105 |
| 3M | All passed | Paired 1-tailed t-test | 3D/2D | 0.0371 |
| 3N | All passed | Paired 1-tailed t-test | CD24-/CD24+ | 0.0797 |
| 3O | All passed | Paired 1-tailed t-test | 3D/2D | 0.0282 |
| 3P | All passed | Two-way ANOVA, Sidak | CD24-/CD24+ at 1:1 | 0.1865 |
| | | | CD24-/CD24+ at 10:1 | 0.2411 |
| | | | CD24-/CD24+ at 20:1 | 0.0043 |
| 3Q | All passed | Two-way ANOVA, Sidak | 3D/2D at 1:1 | 0.3949 |
| | | | 3D/2D at 10:1 | 0.3165 |
| | | | 3D/2D at 20:1 | 0.0022 |
| 3T | All passed | 1-tail ratio paired t-test | CD24- stim/CD24- unstim, a-b | 0.0482 |
| | | | CD24+ stim/CD24+ unstim, a-c | 0.0107 |
| | | | CD24- stim/CD24+ stim, b-c | 0.0255 |
| 3U | All passed | 1-tail ratio paired t-test | 3D stim/3D unstim, a-b | 0.0094 |
| | | | 2D stim/2D unstim, a-c | 0.0079 |

| Figure | S-W | Test | Sample Comparison | p-value |
|---|---|---|---|---|
| 3U | | | 3D stim/2D stim, b-c | 0.0085 |
| 4C | All passed | | 3D/2D | 0.0288 |
| 4D | n=2, too small | No test performed | | |
| 4F | All passed | 1-tail ratio paired t-test | γδTc + SUM149/SUM149 | 0.0310 |
| 4H | All passed | 1-tail ratio paired t-test | γδTc + PDX401/ PDX401 | 0.0939 |
| 4I | All passed | 1-tail ratio paired t-test | γδTc + SUM149/SUM149 | 0.2808 |
| 4J | All passed | 1-tail ratio paired t-test | γδTc + PDX401/ PDX401 | 0.0809 |
| 4K | All passed | One-way ANOVA, Tukey | CD24- αPD-1 vs. IgG | 0.0222 |
| | | | CD24+ αPD-1 vs. IgG | 0.1847 |
| 4L | All passed | One-way ANOVA, Tukey | αPD-1 3D vs. IgG | 0.3405 |
| | | | αPD-1 2D vs. IgG | 0.3032 |
| 5A | All passed | RM 1-way ANOVA, Bonf | NKG2D, FasL vs. IgG | >0.05 |
| | | | sTRAIL vs. IgG | 0.0216 |
| | | | MICA/B vs. IgG | 0.0006 |
| | | | CD54 vs. IgG | <0.0001 |
| 5B | F: NKG2D | RM 1-way ANOVA, Bonf | NKG2D vs. IgG | 0.0004 |
| | | | FasL vs. IgG | <0.0001 |
| | | | sTRAIL vs. IgG | 0.0002 |
| | | | MICA/B, CD54 vs. IgG | <0.0001 |
| 5C | All passed | Paired 1-tailed t-test | CD24-/CD24+ | 0.0591 |
| 5D | All passed | RM 1-way ANOVA, Bonf | Any vs. IgG | >0.9999 |
| 5E | All passed | RM 1-way ANOVA, Bonf | NKG2D vs. IgG | 0.0001 |
| | | | FasL, sTRAIL, MICA/B | <0.0001 |
| | | | CD54 vs. IgG | 0.0008 |
| 5F | All passed | Paired 1-tailed t-test | 3D/2D | 0.0037 |
| 5I | All passed | Paired 1-tailed t-test | γδTc + SUM149/SUM149 | 0.0950 |
| 5L | All passed | Paired 1-tailed t-test | γδTc + PDX401/ PDX401 | 0.2722 |
| 5M | All passed | One-way ANOVA, Sidak | 3D dMCL1-2 vs. DMSO | 0.0003 |
| | | | 3D A-121 vs. DMSO | 0.5805 |
| | | | 3D dMCL1-2 vs. A-121 | <0.0001 |
| | | | 2D dMCL1-2 vs. DMSO | 0.0073 |
| | | | 2D A-121 vs. DMSO | 0.9886 |
| | | | 2D dMCL1-2 vs. A-121 | 0.0237 |
| 5N | All passed | Two-way ANOVA, Tukey | CD24- αPD-1 vs. IgG | 0.0086 |
| | | | CD24- DMSO vs. dMCL1-2 | 0.7553 |
| | | | CD24- αPD-1 vs. dMCL1-2 | 0.0728 |
| | | | CD24- αPD-1 vs. Combination | 0.8545 |
| | | | CD24- dMCL1-2 vs. Combination | 0.0088 |
| | | | CD24+ αPD-1 vs. IgG | 0.0096 |
| | | | CD24+ DMSO vs. dMCL1-2 | 0.8346 |
| | | | CD24+ αPD-1 vs. dMCL1-2 | 0.0430 |

| Figure | S-W | Test | Sample Comparison | p-value |
|---|---|---|---|---|
| 5N | | | CD24+ αPD-1 vs. Combination | >0.9999 |
| | | | CD24+ dMCL1-2 vs. Combination | 0.0347 |
| 5O | All passed | Two-way ANOVA, Tukey | 3D αPD-1 vs. IgG | <0.0001 |
| | | | 3D DMSO vs. dMCL1-2 | 0.4344 |
| | | | 3D αPD-1 vs. dMCL1-2 | <0.0001 |
| | | | 3D αPD-1 vs. Combination | >0.9999 |
| | | | 3D dMCL1-2 vs. Combination | <0.0001 |
| | | | 2D αPD-1 vs. IgG | 0.0024 |
| | | | 2D DMSO vs. dMCL1-2 | 0.9352 |
| | | | 2D αPD-1 vs. dMCL1-2 | 0.0088 |
| | | | 2D αPD-1 vs. Combination | 0.6505 |
| | | | 2D dMCL1-2 vs. Combination | 0.0004 |
| 6B | All passed | Paired 1-tailed t-test | CD24-/CD24+ MICA/B | 0.0428 |
| | All passed | Paired 1-tailed t-test | CD24-/CD24+ ULBP-2,5,6 | 0.0518 |
| | All passed | Paired 1-tailed t-test | CD24-/CD24+ ULBP-3 | 0.0467 |
| | All passed | Paired 1-tailed t-test | CD24-/CD24+ ULBP-4 | 0.0430 |
| 6D | All passed | Paired 1-tailed t-test | 3D/2D MICA/B | 0.0088 |
| | All passed | Paired 1-tailed t-test | 3D/2D ULBP-2,5,6 | 0.2084 |
| | All passed | Paired 1-tailed t-test | 3D/2D ULBP-3 | 0.3923 |
| | All passed | Paired 1-tailed t-test | 3D/2D ULBP-4 | 0.3388 |
| 7C | All passed | Paired 1-tailed t-test | CD24-/CD24+ | 0.0033 |
| 7D | All passed | Paired 1-tailed t-test | 3D/2D | 0.0038 |
| 7F | All passed | Paired 1-tailed t-test | ADAMi vs. DMSO | 0.0008 |
| 7G | All passed | Paired 1-tailed t-test | ADAMi vs. DMSO | 0.0011 |
| 7H | F: ADAMi | Ratio paired t-test | ADAMi vs. DMSO | 0.0332 |
| 7I | All passed | Paired 1-tailed t-test | ADAMi vs. DMSO | 0.0058 |
| 7J | All passed | Two-way ANOVA, Tukey | CD24- ADAMi vs. DMSO | <0.0001 |
| | | | CD24+ ADAMi vs. DMSO | 0.0007 |
| | | | CD24- ADAMi vs. CD24+ DMSO | 0.6781 |
| | | | CD24- DMSO vs. CD24+ DMSO | <0.0001 |
| 7K | All passed | Two-way ANOVA, Tukey | 3D ADAMi vs. DMSO | <0.0001 |
| | | | 2D ADAMi vs. DMSO | <0.0001 |
| | | | 3D ADAMi vs. 2D DMSO | 0.2139 |
| | | | 3D DMSO vs. 2D DMSO | <0.0001 |

**Table S3.** Statistical tests employed and resulting p-values for experiments shown in Supplemental Figures S1-S8. Biological replicates are listed in order from left to right as indicated by numbers 1-4; Bonf=Bonferroni's multiple comparisons test; E:T=effector:target ratio; F=failed; L=left panel; R=right panel; S-W=Shapiro-Wilk Normality test; Tukey=Tukey's multiple comparisons test.

| Figure | S-W | Test | Sample Comparison | p-value |
|---|---|---|---|---|
| S1K-1 | CD24- passed | Simple Linear Regression | CD24-/CD24+ | <0.0001 |
| S1K-2 | CD24- passed | Simple Linear Regression | CD24-/CD24+ | <0.0001 |
| S1M-1 | All passed | Two-way ANOVA | CD24-/CD24+ at all E:T | <0.0001 |
| S1M-2 | F: CD24+ 20:1 | Two-way ANOVA | CD24-/CD24+ at 10:1 | 0.0349 |
| S1M-3 | All passed | Two-way ANOVA | CD24-/CD24+ at 10:1 | 0.0433 |
| S1N-1 | All passed | Two-way ANOVA | 3D/2D at all E:T | >0.05 |
| S1N-2 | All passed | Two-way ANOVA | 3D/2D at 10:1 | 0.0001 |
| S1N-3 | All passed | Two-way ANOVA | 3D/2D at 1:1 | 0.0007 |
| | | | 3D/2D at 10:1 | 0.0003 |
| | | | 3D/2D at 20:1 | <0.0001 |
| S1N-4 | All passed | Two-way ANOVA | 3D/2D at all E:T | <0.0001 |
| S1O-1 | All passed | Simple Linear Regression | γδTc vs. SUM149/SUM149 | <0.0001 |
| S1O-2 | All passed | Simple Linear Regression | γδTc vs. SUM149/SUM149 | 0.0197 |
| S1O-3 | F:SUM149 | Simple Linear Regression | γδTc vs. SUM149/SUM149 | 0.0025 |
| S1P-1 | All passed | Two-way ANOVA | 3D/2D at 20:1 | <0.0001 |
| S1P-2 | All passed | Two-way ANOVA | 3D/2D at 10:1 | 0.0045 |
| | | | 3D/2D at 20:1 | 0.0011 |
| S1P-3 | All passed | Two-way ANOVA | 3D/2D at 10:1 | 0.0026 |
| | | | 3D/2D at 20:1 | 0.0004 |
| S1P-4 | All passed | Two-way ANOVA | 3D/2D at 10:1 | 0.0066 |
| S1Q-1 | F:PDX401 | Simple Linear Regression | γδTc vs. PDX401/PDX401 | <0.0001 |
| S1Q-2 | All passed | Simple Linear Regression | γδTc vs. PDX401/PDX401 | 0.0140 |
| S2H | n=2, too small | One-way ANOVA, Tukey | γδTc + CD24-/γδTc, a-a | 0.7916 |
| | | | γδTc + CD24+/γδTc, a-b | 0.0043 |
| | | | γδTc/CD24- or CD24+, a-c | <0.05 |
| | | | γδTc + CD24-/CD24- or CD24+, a-c | <0.05 |
| | | | γδTc + CD24+/CD24- or CD24+, b-c | <0.001 |
| | | | γδTc + CD24-/ γδTc vs. CD24+, a-b | 0.0022 |
| S2I | n=2, too small | One-way ANOVA, Tukey | γδTc + CD24-/γδTc, a-b | 0.0011 |
| | | | γδTc + CD24+/γδTc, a-c | <0.0001 |
| | | | γδTc/CD24- or CD24+, a-d | <0.0001 |
| | | | γδTc + CD24-/CD24- or CD24+, b-d | <0.0001 |
| | | | γδTc + CD24+/CD24- or CD24+, c-d | <0.0001 |
| | | | γδTc + CD24-/ γδTc + CD24+, b-c | <0.0001 |

| Figure | S-W | Test | Sample Comparison | p-value |
|---|---|---|---|---|
| S2J | n=2, too small | One-way ANOVA, Tukey | γδTc + 3D/γδTc, a-b | 0.9011 |
| | | | γδTc + 2D/γδTc, a-c | <0.0001 |
| | | | γδTc/3D a-d | 0.0009 |
| | | | γδTc/2D a-e | 0.0005 |
| | | | γδTc + 3D/3D b-d | 0.0014 |
| | | | γδTc + 3D/2D b-e | 0.0007 |
| | | | γδTc + 2D/3D, c-d | <0.0001 |
| | | | γδTc + 2D/2D, c-e | <0.0001 |
| | | | γδTc + 3D/ γδTc + 2D, b-c | <0.0001 |
| S2K | n=2, too small | One-way ANOVA, Tukey | γδTc + 3D/γδTc, a-b | 0.5693 |
| | | | γδTc + 2D/γδTc, a-c | 0.0016 |
| | | | γδTc/3D a-d | <0.0001 |
| | | | γδTc/2D a-e | 0.0007 |
| | | | γδTc + 3D/3D b-d | <0.0001 |
| | | | γδTc + 3D/2D b-e | 0.0004 |
| | | | γδTc + 2D/3D, c-d | <0.0001 |
| | | | γδTc + 2D/2D, c-e | <0.0001 |
| | | | γδTc + 3D/ γδTc + 2D, b-c | 0.0038 |
| S3G | All passed | One-way ANOVA, Sidak | CD24-/CD24+ | 0.2247 |
| | | | 3D/2D | 0.1750 |
| S3I | All passed | Two-way ANOVA, Sidak | CD24-/CD24+ all time points | >0.05 |
| S3J | All passed | Two-way ANOVA, Sidak | 3D/2D | >0.05 |
| S3O | All passed | Two-way ANOVA, Sidak | CD24-/CD24+ at 20:1 | 0.0227 |
| S3P | All passed | Two-way ANOVA, Sidak | CD24-/CD24+ at 10:1 | 0.0170 |
| | | | CD24-/CD24+ at 20:1 | <0.0001 |
| S3Q | All passed | Two-way ANOVA, Sidak | CD24-/CD24+ at 20:1 | 0.0089 |
| S3R | All passed | Two-way ANOVA, Sidak | 3D/2D at 20:1 | 0.0355 |
| S3S | All passed | Two-way ANOVA, Sidak | 3D/2D at 1:1 | 0.0100 |
| | | Two-way ANOVA, Sidak | 3D/2D at 10:1 | 0.0060 |
| | | Two-way ANOVA, Sidak | 3D/2D at 20:1 | <0.0001 |
| S3T | All passed | Two-way ANOVA, Sidak | 3D/2D at 1:1 | 0.0003 |
| | | | 3D/2D at 10:1 | 0.0063 |
| | | | 3D/2D at 20:1 | <0.0001 |
| S4T | All passed | One-way ANOVA, Tukey | CD24- αPD-1 vs. IgG | 0.0051 |
| | | | CD24+ αPD-1 vs. IgG | 0.0838 |
| S4U | All passed | One-way ANOVA, Tukey | CD24- αPD-1 vs. IgG | 0.0003 |
| | | | CD24+ αPD-1 vs. IgG | 0.1317 |
| S4V | All passed | One-way ANOVA, Tukey | CD24- αPD-1 vs. IgG | 0.0172 |
| | | | CD24+ αPD-1 vs. IgG | 0.0045 |
| S4W | All passed | One-way ANOVA, Tukey | 3D αPD-1 vs. IgG | 0.4082 |

| Figure | S-W | Test | Sample Comparison | p-value |
|---|---|---|---|---|
| S4W | | | 2D αPD-1 vs. IgG | 0.0570 |
| S4X | All passed | One-way ANOVA, Tukey | 3D αPD-1 vs. IgG | 0.0141 |
| | | | 2D αPD-1 vs. IgG | 0.7328 |
| S4Y | All passed | One-way ANOVA, Tukey | 3D αPD-1 vs. IgG | 0.0267 |
| | | | 2D αPD-1 vs. IgG | 0.0033 |
| S5A | All passed | One-way ANOVA, Sidak | NKG2D vs. IgG | 0.0472 |
| | | | FasL, sTRAIL vs. IgG | >0.05 |
| S5B | All passed | One-way ANOVA, Sidak | CD54 vs. IgG | 0.0240 |
| | | | MICA/B vs. IgG | 0.0286 |
| S5C | All passed | One-way ANOVA, Sidak | NKG2D vs. IgG | 0.0117 |
| | | | FasL vs. IgG | 0.0042 |
| | | | sTRAIL, MICA/B, CD54 vs. IgG | >0.05 |
| S5D(L) | All passed | One-way ANOVA, Sidak | NKG2D vs. IgG | 0.0059 |
| | | | FasL vs. IgG | >0.05 |
| | | | sTRAIL vs. IgG | 0.0035 |
| | | | MICA/B vs. IgG | 0.0003 |
| | | | CD54 vs. IgG | <0.0001 |
| S5D(R) | All passed | One-way ANOVA, Sidak | NKG2D, FasL, sTRAIL, CD54 vs. IgG | <0.0001 |
| | | | MICA/B vs. IgG | 0.0001 |
| S5E(L) | All passed | One-way ANOVA, Sidak | NKG2D, FasL, m or sTRAIL vs. IgG | >0.05 |
| | | | MICA/B vs. IgG | 0.0002 |
| | | | CD54 vs. IgG | 0.0007 |
| S5E(R) | All passed | One-way ANOVA, Sidak | NKG2D vs. IgG | 0.0072 |
| | | | FasL vs. IgG | 0.0097 |
| | | | mTRAIL vs. IgG | 0.1311 |
| | | | sTRAIL vs. IgG | 0.0002 |
| | | | MICA/B, CD54 vs. IgG | <0.0001 |
| S5F(L) | All passed | One-way ANOVA, Sidak | NKG2D, FasL, sTRAIL vs. IgG | >0.05 |
| | | | MICA/B vs. IgG | 0.4403 |
| | | | CD54 vs. IgG | 0.0016 |
| S5F(R) | All passed | One-way ANOVA, Sidak | NKG2D, FasL vs. IgG | >0.05 |
| | | | sTRAIL vs. IgG | 0.0273 |
| | | | MICA/B vs. IgG | 0.0005 |
| | | | CD54 vs. IgG | <0.0001 |
| S5G(L) | All passed | One-way ANOVA, Sidak | NKG2D, FasL, sTRAIL vs. IgG | >0.05 |
| | | | MICA/B vs. IgG | 0.0022 |
| | | | CD54 vs. IgG | 0.0086 |
| | | | TRAIL R1 vs. IgG | 0.0001 |
| | | | TRAIL R2 vs. IgG | 0.0004 |
| S5G(R) | F: TRAIL R2 | One-way ANOVA, Sidak | NKG2D vs. IgG | 0.0014 |

| Figure | S-W | Test | Sample Comparison | p-value |
|---|---|---|---|---|
| S5G(R) | | | FasL, CD54, TRAILR1 vs. IgG | <0.0001 |
| | | | sTRAIL, MICA/B vs. IgG | 0.0002 |
| | | | TRAIL R2 vs. IgG | 0.0058 |
| S5H(L) | F: FasL | One-way ANOVA, Sidak | NKG2D, FasL, m or sTRAIL vs. IgG | >0.05 |
| | | | MICA/B vs. IgG | >0.05 |
| | | | CD54 vs. IgG | <0.0001 |
| | | | TRAIL R1 vs. IgG | 0.0002 |
| | | | TRAIL R2 vs. IgG | 0.0320 |
| | | | Fas vs. IgG | 0.1606 |
| S5H(R) | F: MICA/B | One-way ANOVA, Sidak | NKG2D vs. IgG | 0.0038 |
| | | | FasL, sTRAIL, Fas vs. IgG | <0.0001 |
| | | | mTRAIL vs. IgG | 0.0007 |
| | | | MICA/B vs. IgG | 0.0004 |
| | | | CD54 vs. IgG | 0.0100 |
| | | | TRAIL R1, TRAIL R2 vs. IgG | <0.0001 |
| S5I(L) | All passed | One-way ANOVA, Sidak | NKG2D vs. IgG | 0.0092 |
| | | | FasL, sTRAIL vs. IgG | >0.05 |
| | | | MICA/B, CD54 vs. IgG | <0.0001 |
| | | | TRAIL R1, TRAIL R2 vs. IgG | <0.0001 |
| S5I(R) | All passed | One-way ANOVA, Sidak | NKG2D, FasL, sTRAIL vs. IgG | <0.0001 |
| | | | MICA/B, CD54, TRAILR1/R2 vs. IgG | <0.0001 |
| S5J(L) | All passed | One-way ANOVA, Sidak | NKG2D, FasL, sTRAIL vs. IgG | >0.05 |
| | | | MICA/B, CD54 vs. IgG | >0.05 |
| | | | Fas, TRAIL R1, TRAIL R2 vs. IgG | >0.05 |
| S5J(R) | F: MICA/B, Fas | One-way ANOVA, Sidak | NKG2D vs. IgG | 0.0255 |
| | | | FasL vs. IgG | 0.0275 |
| | | | sTRAIL vs. IgG | 0.0119 |
| | | | MICA/B vs. IgG | 0.0016 |
| | | | CD54 vs. IgG | 0.0366 |
| | | | Fas vs. IgG | 0.0059 |
| | | | TRAIL R1 vs. IgG | 0.0028 |
| | | | TRAIL R2 vs. IgG | 0.6449 |
| S5R(L) | All passed | One-way ANOVA, Sidak | CD24- dMCL1-2 vs. DMSO | 0.0254 |
| | | | CD24+ dMCL1-2 vs. DMSO | 0.7828 |
| S5R(R) | All passed | One-way ANOVA, Sidak | CD24- dMCL1-2 vs. DMSO | 0.2639 |
| | | | CD24+ dMCL1-2 vs. DMSO | 0.2875 |
| S5S(L) | F:CD24-DMSO | One-way ANOVA, Sidak | CD24- dMCL1-2 vs. DMSO | 0.0239 |
| | | | CD24- A-121 vs. DMSO | 0.7440 |
| | | | CD24- dMCL1-2 vs. A-121 | 0.0022 |
| | | | CD24+ dMCL1-2 vs. DMSO | 0.2884 |
| | | | CD24+ A-121 vs. DMSO | 0.0824 |

| Figure | S-W | Test | Sample Comparison | p-value |
|---|---|---|---|---|
| S5S(L) | | | CD24+ dMCL1-2 vs. A-121 | 0.0019 |
| S5S(R) | All passed | One-way ANOVA, Sidak | 3D dMCL1-2 vs. DMSO | 0.1791 |
| | | | 3D A-121 vs. DMSO | 0.9254 |
| | | | 3D dMCL1-2 vs. A-121 | 0.0318 |
| | | | 2D dMCL1-2 vs. DMSO | 0.9998 |
| | | | 2D A-121 vs. DMSO | 0.0068 |
| | | | 2D dMCL1-2 vs. A-121 | 0.0120 |
| S5T(L) | All passed | One-way ANOVA, Sidak | CD24- dMCL1-2 vs. DMSO | 0.9681 |
| | | | CD24+ dMCL1-2 vs. DMSO | 0.9934 |
| S5T(R) | All passed | One-way ANOVA, Sidak | 3D dMCL1-2 vs. DMSO | 0.1065 |
| | | | 2D dMCL1-2 vs. DMSO | 0.1669 |
| S5U | All passed | One-way ANOVA, Sidak | 3D dMCL1-2 vs. DMSO | 0.9782 |
| | | | 2D dMCL1-2 vs. DMSO | 0.1526 |
| S5V(L) | All passed | Two-way ANOVA, Tukey | CD24- αPD-1 vs. IgG | <0.0001 |
| | | | CD24- αPD-1 vs. dMCL1-2 | <0.0001 |
| | | | CD24- αPD-1 vs. Combination | >0.9999 |
| | | | CD24- DMSO vs. dMCL1-2 | 0.7125 |
| | | | CD24- DMSO vs. A-121 | 0.8731 |
| | | | CD24- dMCL1-2 vs. A-121 | 0.0845 |
| | | | CD24- dMCL1-2 vs. Combination | <0.0001 |
| | | | CD24- dMCL1-2 only vs. A-121 only | >0.9999 |
| | | | CD24+ αPD-1 vs. IgG | <0.0001 |
| | | | CD24+ αPD-1 vs. dMCL1-2 | <0.0001 |
| | | | CD24+ αPD-1 vs. Combination | 0.3429 |
| | | | CD24+ DMSO vs. dMCL1-2 | 0.1558 |
| | | | CD24+ DMSO vs. A-121 | 0.4577 |
| | | | CD24+ dMCL1-2 vs. A-121 | 0.0009 |
| | | | CD24+ dMCL1-2 vs. Combination | <0.0001 |
| | | | CD24+ dMCL1-2 only vs. A-121only | 0.5036 |
| S5V(R) | All passed | Two-way ANOVA, Tukey | 3D αPD-1 vs. IgG | <0.0001 |
| | | | 3D αPD-1 vs. dMCL1-2 | 0.0005 |
| | | | 3D αPD-1 vs. Combination | 0.9792 |
| | | | 3D DMSO vs. dMCL1-2 | >0.9999 |
| | | | 3D DMSO vs. A-121 | 0.9549 |
| | | | 3D dMCL1-2 vs. A-121 | 0.8200 |
| | | | 3D dMCL1-2 vs. Combination | 0.0065 |
| | | | 3D dMCL1-2 only vs. A-121 only | >0.9999 |
| | | | 2D αPD-1 vs. IgG | 0.0057 |
| | | | 2D αPD-1 vs. dMCL1-2 | 0.0293 |
| | | | 2D αPD-1 vs. Combination | 0.9962 |
| | | | 2D DMSO vs. dMCL1-2 | 0.6859 |

| Figure | S-W | Test | Sample Comparison | p-value |
|---|---|---|---|---|
| S5V(R) | | | 2D DMSO vs. A-121 | 0.9891 |
| | | | 2D dMCL1-2 vs. A-121 | 0.2141 |
| | | | 2D dMCL1-2 vs. Combination | 0.0047 |
| | | | 2D dMCL1-2 only vs. A-121 only | 0.3152 |
| S5W(L) | All passed | Two-way ANOVA, Tukey | CD24- αPD-1 vs. IgG | <0.0001 |
| | | | CD24- αPD-1 vs. dMCL1-2 | 0.0761 |
| | | | CD24- αPD-1 vs. Combination | 0.0248 |
| | | | CD24- DMSO vs. dMCL1-2 | 0.1365 |
| | | | CD24- DMSO vs. A-121 | 0.6204 |
| | | | CD24- dMCL1-2 vs. A-121 | 0.9754 |
| | | | CD24- dMCL1-2 vs. Combination | <0.0001 |
| | | | CD24- dMCL1-2 only vs. A-121 only | >0.9999 |
| | | | CD24+ αPD-1 vs. IgG | 0.0007 |
| | | | CD24+ αPD-1 vs. dMCL1-2 | 0.0199 |
| | | | CD24+ αPD-1 vs. Combination | >0.9999 |
| | | | CD24+ DMSO vs. dMCL1-2 | 0.6657 |
| | | | CD24+ DMSO vs. A-121 | >0.9999 |
| | | | CD24+ dMCL1-2 vs. A-121 | 0.7131 |
| | | | CD24+ dMCL1-2 vs. Combination | 0.0129 |
| | | | CD24+ dMCL1-2 only vs. A-121only | 0.7111 |
| S5W(R) | All passed | Two-way ANOVA, Tukey | 3D αPD-1 vs. IgG | <0.0001 |
| | | | 3D αPD-1 vs. dMCL1-2 | 0.0008 |
| | | | 3D αPD-1 vs. Combination | 0.6862 |
| | | | 3D DMSO vs. dMCL1-2 | 0.3164 |
| | | | 3D DMSO vs. A-121 | 0.9841 |
| | | | 3D dMCL1-2 vs. A-121 | 0.8425 |
| | | | 3D dMCL1-2 vs. Combination | <0.0001 |
| | | | 3D dMCL1-2 only vs. A-121 only | 0.2801 |
| | | | 2D αPD-1 vs. IgG | 0.0022 |
| | | | 2D αPD-1 vs. dMCL1-2 | 0.0200 |
| | | | 2D αPD-1 vs. Combination | 0.0721 |
| | | | 2D DMSO vs. dMCL1-2 | 0.9989 |
| | | | 2D DMSO vs. A-121 | 0.4506 |
| | | | 2D dMCL1-2 vs. A-121 | 0.8059 |
| | | | 2D dMCL1-2 vs. Combination | <0.0001 |
| | | | 2D dMCL1-2 only vs. A-121 only | 0.4860 |
| S5X(L) | All passed | Two-way ANOVA, Tukey | CD24- αPD-1 vs. IgG | <0.0001 |
| | | | CD24- αPD-1 vs. dMCL1-2 | <0.0001 |
| | | | CD24- αPD-1 vs. Combination | >0.9999 |
| | | | CD24- DMSO vs. dMCL1-2 | 0.7247 |
| | | | CD24- DMSO vs. A-121 | 0.7564 |

| Figure | S-W | Test | Sample Comparison | p-value |
|---|---|---|---|---|
| S5X(L) | | | CD24- dMCL1-2 vs. A-121 | 0.0522 |
| | | | CD24- dMCL1-2 vs. Combination | <0.0001 |
| | | | CD24- dMCL1-2 only vs. A-121 only | 0.2634 |
| | | | CD24+ αPD-1 vs. IgG | <0.0001 |
| | | | CD24+ αPD-1 vs. dMCL1-2 | 0.0006 |
| | | | CD24+ αPD-1 vs. Combination | 0.6840 |
| | | | CD24+ DMSO vs. dMCL1-2 | 0.8042 |
| | | | CD24+ DMSO vs. A-121 | >0.9999 |
| | | | CD24+ dMCL1-2 vs. A-121 | 0.9258 |
| | | | CD24+ dMCL1-2 vs. Combination | 0.0521 |
| | | | CD24+ dMCL1-2 only vs. A-121only | 0.9998 |
| S5X(R) | All passed | Two-way ANOVA, Tukey | 3D αPD-1 vs. IgG | <0.0001 |
| | | | 3D αPD-1 vs. dMCL1-2 | <0.0001 |
| | | | 3D αPD-1 vs. Combination | 0.7728 |
| | | | 3D DMSO vs. dMCL1-2 | 0.0929 |
| | | | 3D DMSO vs. A-121 | >0.9999 |
| | | | 3D dMCL1-2 vs. A-121 | 0.1101 |
| | | | 3D dMCL1-2 vs. Combination | <0.0001 |
| | | | 3D dMCL1-2 only vs. A-121 only | 0.9983 |
| | | | 2D αPD-1 vs. IgG | 0.0017 |
| | | | 2D αPD-1 vs. dMCL1-2 | 0.0017 |
| | | | 2D αPD-1 vs. Combination | 0.9995 |
| | | | 2D DMSO vs. dMCL1-2 | 0.7790 |
| | | | 2D DMSO vs. A-121 | 0.4842 |
| | | | 2D dMCL1-2 vs. A-121 | 0.0229 |
| | | | 2D dMCL1-2 vs. Combination | 0.0067 |
| | | | 2D dMCL1-2 only vs. A-121 only | >0.9999 |
| S7E | All passed | Two-way ANOVA, Bonf | Replicate 1 CD24- vs. CD24+ | 0.0004 |
| | | | Replicate 2 CD24- vs. CD24+ | 0.0019 |
| | | | Replicate 3 CD24- vs. CD24+ | 0.0420 |
| | | | Replicate 4 CD24- vs. CD24+ | 0.0011 |
| S7F | All passed | Two-way ANOVA, Bonf | Replicate 1 3D vs. 2D | 0.0041 |
| | | | Replicate 2 3D vs. 2D | <0.0001 |
| | | | Replicate 3 3D vs. 2D | <0.0001 |
| | | | Replicate 4 3D vs. 2D | <0.0001 |
| S7I | DMSO passed | Two-way ANOVA, Bonf | Replicate 1 ADAMi vs. DMSO | 0.0069 |
| | | | Replicate 2 ADAMi vs. DMSO | 0.0007 |
| | | | Replicate 3 ADAMi vs. DMSO | 0.0143 |
| S7J | All passed | Two-way ANOVA, Bonf | Replicate 1 ADAMi vs. DMSO | 0.0010 |
| | | | Replicate 2 ADAMi vs. DMSO | 0.0007 |
| | | | Replicate 3 ADAMi vs. DMSO | 0.0053 |

| Figure | S-W | Test | Sample Comparison | p-value |
|---|---|---|---|---|
| S7K | F: 24+ ADAMi | Two-way ANOVA, Tukey | CD24- ADAMi vs. DMSO | 0.0002 |
| | | | CD24- ADAMi vs. CD24+ DMSO | >0.9999 |
| | | | CD24+ ADAMi vs. DMSO | 0.0054 |
| S7L | All passed | Two-way ANOVA, Tukey | CD24- ADAMi vs. DMSO | 0.0010 |
| | | | CD24- ADAMi vs. CD24+ DMSO | 0.6988 |
| | | | CD24+ ADAMi vs. DMSO | 0.0009 |
| S7M | All passed | Two-way ANOVA, Tukey | CD24- ADAMi vs. DMSO | <0.0001 |
| | | | CD24- ADAMi vs. CD24+ DMSO | 0.9998 |
| | | | CD24+ ADAMi vs. DMSO | 0.0208 |
| S7N | F:2D Control | Two-way ANOVA, Tukey | 3D ADAMi vs. DMSO | <0.0001 |
| | | | 3D ADAMi vs. 2D DMSO | 0.7278 |
| | | | 2D ADAMi vs. DMSO | 0.0025 |
| S7O | All passed | Two-way ANOVA, Tukey | 3D ADAMi vs. DMSO | 0.0024 |
| | | | 3D ADAMi vs. 2D DMSO | 0.8584 |
| | | | 2D ADAMi vs. DMSO | 0.0501 |
| S7P | All passed | Two-way ANOVA, Tukey | 3D ADAMi vs. DMSO | 0.0035 |
| | | | 3D ADAMi vs. 2D DMSO | >0.9999 |
| | | | 2D ADAMi vs. DMSO | 0.0226 |
| S8B | All passed | Two-way ANOVA, Sidak | 3D/2D at 1:1 | 0.3845 |
| | | | 3D/2D at 10:1 | 0.0532 |
| | | | 3D/2D at 20:1 | 0.0239 |
| S8C | F: MCF-7 | Simple Linear Regression | γδTc vs. MCF-7/MCF-7 | 0.0016 |
| S8G | F: 3D | Wilcoxon | 3D vs. 2D | 0.2500 |
| S8H | All passed | 2-tail ratio paired t-test | γδTc + 3D/ γδTc + 2D | 0.0262 |
| S8I(L) | All passed | 2-tailed paired t-test | 3D vs. 2D | 0.0216 |
| S8I(R) | All passed | 2-tailed paired t-test | 3D vs. 2D | 0.0020 |
| S8J | All passed | Two-way ANOVA, Sidak | 3D vs. 2D, all time points | >0.05 |
| S8K | n=2, too small | No test performed | | |
| S8L | F: 2D | Wilcoxon | 3D vs. 2D | 0.2500 |
| S8M | All passed | 1-tailed paired t-test | 3D vs. 2D | 0.0282 |
| S8N | All passed | 1-tail ratio paired t-test | 3D stim vs. 2D stim, b-c | 0.0452 |
| S8Q | F:2D CD54 | One-way ANOVA, Sidak | 3D NKG2D, FasL vs. IgG | >0.05 |
| | | | 3D sTRAIL vs. IgG | 0.0115 |
| | | | 3D CD54 vs. IgG | 0.0033 |
| | | | 3D MICA/B vs. IgG | 0.0021 |
| | | | 2D NKG2D vs. IgG | 0.0003 |
| | | | 2D FasL vs. IgG | 0.0005 |
| | | | 2D sTRAIL vs. IgG | <0.0001 |
| | | | 2D CD54 vs. IgG | 0.0002 |
| | | | 2D MICA/B vs. IgG | 0.0003 |
| S8R | All passed | 1-tailed paired t-test | 3D vs. 2D | 0.1614 |

**Table S4.** Flow cytometry, western blot and blocking assay antibodies.

| **Flow Cytometry Antibody** | **Dilution or Concentration** | **Clone** | **Company** |
|---|---|---|---|
| CD107a AlexaFluor (AF) 647 | 1:40 | H4A3 | BioLegend |
| CD24 APC | 1:50 | REA832 | Miltenyi |
| CD44 PerCP Cy5.5 | 1:50 | BJ18 | BioLegend |
| CD44 Vioblue | 1:20 | REA690 | Miltenyi |
| CTLA-4(CD152) APC | 1:10 | L3D10 | BioLegend |
| Fas (CD95) APC | 1:100 | DX2 | BioLegend |
| IFN-γ VioBlue | 1:50 | REA600 | Miltenyi |
| LAG-3 FITC | 1:20 | 11C3C65 | BioLegend |
| MCL-1 AF488 | 1:20 | 22 | Santa Cruz |
| MICA/B PE | 0.1µg | 6D4, | BioLegend |
| PD-1(CD279) BV421 | 1:10 | EH12.2H7 | BioLegend |
| PD-L1 APC | 1:10 | 29E.2A3 | BioLegend |
| PD-L2 PE | 1:10 | 24F.10C12 | BioLegend |
| TCR Vδ1 FITC | 1:10 | REA173 | Miltenyi |
| TCR Vδ2 PerCP | 1:25 | B6 | BioLegend |
| TCRγδ PE | 1:25 | B1 | BioLegend |
| TIM-3 APC/Fire750 | 1:20 | F38-2E2 | BioLegend |
| TRAIL R1 APC | 1:20 | DJR1 | BioLegend |
| TRAIL R2 APC | 1:5 | DJR2-4 | BioLegend |
| ULBP-2,5,6 PE | 0.2µg | 165903 | R&D systems |
| ULBP-3 APC | 0.04µg | 166510 | R&D systems |
| ULBP-4 APC | 0.1µg | 709116 | R&D systems |
| **Western Blot Antibodies** | **Dilution or Concentration** | **Clone** | **Company** |
| rabbit anti-human ADAM10 | 1:500 | ab1997 | Abcam |
| rabbit anti-human ADAM17 | 1:1000 | ab39162 | Abcam |
| mouse anti-human BCL-2 | 1:1000 | C2 | Santa Cruz |
| mouse anti-human BCL-XL | 1:1000 | H5 | Santa Cruz |
| mouse anti-human MCL-1 | 1:1000 | 22 | Santa Cruz |
| rabbit anti-human Survivin | 1:1000 | 71G4B7E | Cell Signaling |
| mouse anti-human XIAP | 1:500 | 48/hILP/XIAP(RUO) | BD Biosciences |
| mouse anti-human β-Actin | 1:3000 | C4 | Santa Cruz |
| rabbit anti-human β-Actin | 1:2000 | 13E5 | Cell Signaling |

| **Blocking Antibodies** | **Concentration** | **Clone** | **Company** |
| --- | --- | --- | --- |
| CD54 | 1µg/100ul | HCD54 | BioLegend |
| Fas | 1µg/100ul | A16086F | BioLegend |
| FasL | 1µg/100ul | NOK-1 | BioLegend |
| IgG1k | 1µg/100ul | MOPC-21 | BioLegend |
| MICA/B | 1µg/100ul | 6D4 | BioLegend |
| NKG2D | 1µg/100ul | 1D11 | BioLegend |
| PD-1 | 1µg/100ul | A17188B | BioLegend |
| TRAIL | 1µg/100ul | RIK-2 | BioLegend |
| TRAIL R1 | 1µg/100ul | DJR1 | BioLegend |
| TRAIL R2 | 1µg/100ul | DJR2-4 | BioLegend |

**Table S5.** Proteins acquired by mass spectrometry that were differentially expressed in conditioned medium from CD24- versus CD24+ SUM149 cells.

| Protein.names | Majority.protein.IDs | Gene.names | Extracellular or secreted | log2_fold-change | -log10_p-value | NSC_CM1 | NSC_CM2 | NSC_CM3 | SC_CM1 | SC_CM2 | SC_CM3 |
|---|---|---|---|---|---|---|---|---|---|---|---|
| Alpha-2-macroglobulin-like protein 1 | A8K2U0;A8K2U0-2 | A2ML1 | Extracellular or secreted | -0.121333333 | 0.073457712 | 26.3336 | 27.4951 | 27.0295 | 27.7596 | 26.4519 | 26.2827 |
| Acetoacetyl-CoA synthetase | Q86V21;Q86V21-3 | AACS | Other | 1.093633333 | 0.408213968 | 22.3305 | 22.9403 | 24.5981 | 22.7101 | 24.643 | 25.7967 |
| Mth938 domain-containing protein | Q9H7C9 | AAMDC | Other | 0.394366667 | 0.261045249 | 26.6917 | 26.3919 | 25.1269 | 26.8417 | 26.8006 | 25.7513 |
| Angio-associated migratory cell protein | Q13685 | AAMP | Extracellular or secreted | -0.043833333 | 0.014032932 | 26.1017 | 26.5418 | 24.7767 | 27.3211 | 25.6611 | 24.3065 |
| Protein AAR2 homolog | Q9Y312 | AAR2 | Other | 0.342033333 | 0.194239637 | 22.19 | 22.8063 | 23.7864 | 22.7089 | 22.8532 | 24.2467 |
| Alanine--tRNA ligase, cytoplasmic | P49588;P49588-2 | AARS | Extracellular or secreted | 0.8794 | 0.525490263 | 28.3602 | 28.6808 | 30.135 | 29.0108 | 30.1173 | 30.6861 |
| L-aminoadipate-semialdehyde dehydrogenase-phosphop | Q9NRN7 | AASDHPPT | Extracellular or secreted | -0.616433333 | 0.545093064 | 24.2088 | 24.6924 | 24.8266 | 23.1081 | 24.3461 | 24.4243 |
| ATP-binding cassette sub-family E member 1 | P61221 | ABCE1 | Other | -0.2675 | 0.339284073 | 26.8658 | 27.1817 | 27.6172 | 26.4708 | 27.176 | 27.2154 |
| ATP-binding cassette sub-family F member 1 | Q8NE71;Q8NE71-2 | ABCF1 | Other | -0.246833333 | 0.20276495 | 28.2021 | 27.5742 | 29.0616 | 28.1059 | 27.8719 | 28.1196 |
| ATP-binding cassette sub-family F member 2 | Q9UG63;Q9UG63-2 | ABCF2 | Extracellular or secreted | -0.735933333 | 0.735244329 | 22.5592 | 21.4116 | 22.4897 | 21.1601 | 21.1897 | 21.9029 |
| Mycophenolic acid acyl-glucuronide esterase, mitochond | Q9NUJ1;Q9NUJ1-2 | ABHD10 | Extracellular or secreted | -1.2533 | 1.765061685 | 29.4396 | 29.5557 | 30.2233 | 28.3649 | 28.8262 | 28.2676 |
| Alpha/beta hydrolase domain-containing protein 11 | Q8NFV4;Q8NFV4-4;Q8NFV4- | ABHD11 | Other | -3.296 | 2.211169833 | 26.418 | 26.3581 | 27.5934 | 22.9072 | 23.1647 | 24.4096 |
| Alpha/beta hydrolase domain-containing protein 14B | Q96IU4 | ABHD14B | Extracellular or secreted | -0.3051 | 0.321124272 | 28.9395 | 29.1632 | 28.4719 | 28.107 | 29.1824 | 28.3699 |
| Abl interactor 1 | Q8IZP0-10;Q8IZP0-8;Q8IZP0- | ABI1 | Extracellular or secreted | 0.1096 | 0.078012607 | 25.7864 | 26.7096 | 27.1998 | 26.2498 | 26.6294 | 27.1454 |
| Costars family protein ABRACL | Q9P1F3 | ABRACL | Other | 0.203366667 | 0.091129204 | 27.3455 | 27.8184 | 27.0376 | 27.0071 | 29.0498 | 26.7547 |
| 3-ketoacyl-CoA thiolase, peroxisomal | P09110;P09110-2 | ACAA1 | Other | 0.0205 | 0.018018034 | 25.4526 | 25.5156 | 26.4287 | 25.5324 | 26.2069 | 25.7191 |
| 3-ketoacyl-CoA thiolase, mitochondrial | P42765 | ACAA2 | Extracellular or secreted | -0.344933333 | 0.233052035 | 27.6113 | 28.9679 | 29.2651 | 27.9211 | 28.72 | 28.1684 |
| Acetyl-CoA carboxylase 1;Biotin carboxylase | Q13085;Q13085-4;Q13085-3 | ACACA | Extracellular or secreted | 2.076233333 | 0.334905809 | 19.782 | 22.4584 | 27.0427 | 23.8399 | 23.8126 | 27.8593 |
| Medium-chain specific acyl-CoA dehydrogenase, mitocho | P11310;P11310-2 | ACADM | Extracellular or secreted | 0.6266 | 0.443957345 | 25.2366 | 25.3541 | 26.8476 | 26.747 | 25.9316 | 26.6395 |
| Very long-chain specific acyl-CoA dehydrogenase, mitoch | P49748-2;P49748;P49748-3 | ACADVL | Other | -0.290066667 | 0.172468594 | 26.9745 | 26.9096 | 28.7083 | 27.0852 | 27.3164 | 27.3206 |
| Acetyl-CoA acetyltransferase, mitochondrial | P24752 | ACAT1 | Extracellular or secreted | 0.1719 | 0.087669952 | 25.479 | 26.9862 | 27.5856 | 26.8638 | 27.2753 | 26.4274 |
| Acetyl-CoA acetyltransferase, cytosolic | Q9BWD1;Q9BWD1-2 | ACAT2 | Extracellular or secreted | 1.602366667 | 1.322150736 | 28.6136 | 28.8905 | 29.037 | 29.6445 | 30.8934 | 30.8103 |
| Golgi resident protein GCP60 | Q9H3P7 | ACBD3 | Other | -0.2099 | 0.217390634 | 28.3605 | 28.2889 | 28.344 | 27.6028 | 27.983 | 28.7779 |
| Apoptotic chromatin condensation inducer in the nucleu | Q9UKV3-5;Q9UKV3 | ACIN1 | Extracellular or secreted | -1.093233333 | 1.069964472 | 29.6954 | 28.6235 | 29.689 | 28.4045 | 28.1704 | 28.1533 |
| ATP-citrate synthase | P53396;P53396-2;P53396-3 | ACLY | Extracellular or secreted | 0.724666667 | 1.081716685 | 30.788 | 30.8552 | 31.4347 | 31.2876 | 31.9004 | 32.0639 |
| Cytoplasmic aconitate hydratase | P21399 | ACO1 | Extracellular or secreted | -0.495266667 | 0.496126492 | 27.6866 | 28.4605 | 28.4557 | 27.0524 | 27.8376 | 28.227 |
| Aconitate hydratase, mitochondrial | Q99798 | ACO2 | Other | -0.409733333 | 0.394655473 | 29.5242 | 30.4243 | 30.8277 | 29.8505 | 30.0809 | 29.6156 |
| Acyl-coenzyme A thioesterase 1;Acyl-coenzyme A thioest | Q86TX2;P49753;P49753-2 | ACOT1;ACOT2 | Other | -2.343833333 | 2.916439642 | 26.5359 | 26.5956 | 27.0973 | 24.3061 | 24.2986 | 24.5926 |
| Acyl-coenzyme A thioesterase 13;Acyl-coenzyme A thioe | Q9NPJ3-2;Q9NPJ3 | ACOT13 | Extracellular or secreted | -1.044733333 | 0.665854915 | 24.5288 | 24.6229 | 25.0017 | 22.5032 | 24.4298 | 24.0862 |
| Cytosolic acyl coenzyme A thioester hydrolase | O00154-4;O00154-6;O00154 | ACOT7 | Extracellular or secreted | -0.1876 | 0.124419633 | 27.071 | 26.7323 | 27.8101 | 26.1361 | 27.3381 | 27.5764 |
| Peroxisomal acyl-coenzyme A oxidase 1 | Q15067-2;Q15067 | ACOX1 | Extracellular or secreted | -0.2388 | 0.048058166 | 21.2398 | 21.2806 | 25.9972 | 22.416 | 23.3126 | 22.0726 |
| Low molecular weight phosphotyrosine protein phospha | P24666;P24666-4 | ACP1 | Extracellular or secreted | 0.789633333 | 1.571765935 | 26.7019 | 27.2968 | 27.0308 | 27.5961 | 28.095 | 27.7073 |
| Lysosomal acid phosphatase | P11117 | ACP2 | Extracellular or secreted | 0.097166667 | 0.162638392 | 26.7079 | 26.3783 | 26.3036 | 26.7546 | 26.7349 | 26.1918 |
| Lysophosphatidic acid phosphatase type 6 | Q9NPH0;Q9NPH0-2 | ACP6 | Extracellular or secreted | 0.226566667 | 0.094346536 | 23.258 | 21.2382 | 21.0649 | 22.1252 | 22.8825 | 21.2331 |
| Prostatic acid phosphatase;PAPf39 | P15309;P15309-2;P15309-3 | ACPP | Extracellular or secreted | -0.179733333 | 0.091123855 | 23.4176 | 25.1325 | 23.4655 | 23.6031 | 24.6243 | 23.249 |
| Acyl-CoA synthetase family member 3, mitochondrial | Q4G176 | ACSF3 | Other | -0.743066667 | 0.226697088 | 23.4911 | 20.7868 | 24.3685 | 23.4376 | 21.3886 | 21.591 |
| Long-chain-fatty-acid--CoA ligase 1 | P33121-3;P33121;P33121-2 | ACSL1 | Extracellular or secreted | -1.803233333 | 0.695480394 | 25.4933 | 25.7807 | 26.2902 | 23.7126 | 22.5534 | 25.8885 |
| Long-chain-fatty-acid--CoA ligase 4 | O60488-2;O60488 | ACSL4 | Extracellular or secreted | 1.327766667 | 0.385308787 | 21.2482 | 21.6253 | 23.5927 | 21.0843 | 24.3672 | 24.998 |
| Acetyl-coenzyme A synthetase, cytoplasmic | Q9NR19;Q9NR19-2 | ACSS2 | Other | 3.314633333 | 1.083955061 | 22.8667 | 24.6209 | 26.672 | 26.263 | 28.6691 | 29.1714 |
| Actin, alpha skeletal muscle;Actin, alpha cardiac muscle 1 | P68133;P68032;P63267;P62 | ACTA1;ACTC1; | Extracellular or secreted | 0.7075 | 0.121011975 | 30.8714 | 26.8761 | 32.1897 | 27.889 | 31.7696 | 32.4011 |
| Actin, cytoplasmic 1;Actin, cytoplasmic 1, N-terminally pr | P60709 | ACTB | Extracellular or secreted | 1.233533333 | 0.524462901 | 25.3221 | 25.7214 | 28.0651 | 26.9893 | 27.1749 | 28.645 |
| Beta-actin-like protein 2 | Q562R1 | ACTBL2 | Extracellular or secreted | -4.516166667 | 0.498709728 | 22.2501 | 32.0275 | 32.6072 | 25.5976 | 21.1271 | 26.6116 |
| Actin, cytoplasmic 2;Actin, cytoplasmic 2, N-terminally pr | P63261 | ACTG1 | Extracellular or secreted | 0.6569 | 0.542364386 | 34.4163 | 34.5942 | 35.8292 | 35.3164 | 35.364 | 36.13 |
| Actin-like protein 6A | O96019;O96019-2 | ACTL6A | Extracellular or secreted | -0.828833333 | 1.05597445 | 29.685 | 29.8536 | 30.2323 | 28.5628 | 29.139 | 29.5826 |
| Alpha-actinin-1 | P12814;P12814-3;P12814-2; | ACTN1 | Extracellular or secreted | -0.025333333 | 0.087889654 | 34.4254 | 34.4096 | 34.162 | 34.404 | 34.2152 | 34.3018 |
| Alpha-actinin-2 | P35609-2;P35609 | ACTN2 | Extracellular or secreted | 3.814166667 | 0.676517831 | 21.7483 | 28.369 | 22.4385 | 27.7115 | 28.0415 | 28.2453 |
| Alpha-actinin-3 | Q08043 | ACTN3 | Extracellular or secreted | 2.226766667 | 2.641879926 | 25.6451 | 25.4599 | 25.6554 | 27.7601 | 27.5554 | 28.1252 |
| Alpha-actinin-4 | O43707;O43707-2;O43707-3 | ACTN4 | Extracellular or secreted | -0.252166667 | 0.518875632 | 35.7644 | 35.5746 | 35.3486 | 35.6158 | 35.2944 | 35.0209 |
| Actin-related protein 10 | Q9NZ32 | ACTR10 | Other | -0.0312 | 0.018626903 | 21.0319 | 21.2726 | 22.2215 | 22.3028 | 20.9171 | 21.2125 |
| Alpha-centractin | P61163 | ACTR1A | Extracellular or secreted | -0.402066667 | 0.379653615 | 28.0876 | 28.5802 | 29.1289 | 27.568 | 28.3537 | 28.6688 |
| Beta-centractin | P42025 | ACTR1B | Extracellular or secreted | -2.123033333 | 1.091555787 | 24.416 | 24.9556 | 25.344 | 22.1486 | 22.002 | 24.1959 |
| Actin-related protein 2 | P61160;P61160-2 | ACTR2 | Extracellular or secreted | 0.0786 | 0.060173605 | 29.7747 | 29.9735 | 30.2226 | 29.3239 | 30.1263 | 30.7564 |
| Actin-related protein 3 | P61158 | ACTR3 | Extracellular or secreted | 0.036566667 | 0.058522923 | 30.7603 | 30.7432 | 31.1399 | 30.6474 | 30.8688 | 31.2369 |
| Aminoacylase-1 | Q03154;Q03154-4;Q03154-2 | ACY1 | Extracellular or secreted | -0.297533333 | 0.199568794 | 31.3037 | 30.7527 | 30.221 | 31.263 | 30.497 | 29.6248 |
| Acylphosphatase-1 | P07311 | ACYP1 | Extracellular or secreted | 0.035566667 | 0.025601069 | 27.4139 | 27.512 | 26.615 | 27.5666 | 27.5978 | 26.4832 |
| Acylphosphatase-2 | P14621 | ACYP2 | Extracellular or secreted | 0.896333333 | 0.993234802 | 25.6341 | 26.2045 | 25.185 | 26.8461 | 26.9014 | 25.9651 |
| Adenosine deaminase | P00813 | ADA | Extracellular or secreted | -1.0706 | 1.381120469 | 28.5586 | 28.4265 | 27.7073 | 27.1702 | 27.3359 | 26.9745 |
| Disintegrin and metalloproteinase domain-containing pro | O14672;O14672-2 | ADAM10 | Extracellular or secreted | 0.304266667 | 0.173875266 | 26.6351 | 27.0421 | 25.6544 | 27.7714 | 26.3569 | 26.1161 |
| Disintegrin and metalloproteinase domain-containing pro | Q13444-10;Q13444-9;Q1344 | ADAM15 | Extracellular or secreted | 2.103933333 | 0.747755481 | 24.7187 | 24.1992 | 21.3139 | 25.1091 | 26.0679 | 25.3666 |
| Disintegrin and metalloproteinase domain-containing pro | P78536-2;P78536 | ADAM17 | Extracellular or secreted | 0.613 | 0.149864969 | 23.2964 | 23.4438 | 22.9987 | 21.0608 | 25.6557 | 24.8614 |
| Disintegrin and metalloproteinase domain-containing pro | Q13443-2;Q13443 | ADAM9 | Extracellular or secreted | 1.751933333 | 0.94061524 | 23.1201 | 25.3386 | 24.5751 | 26.0832 | 26.0859 | 26.1205 |
| A disintegrin and metalloproteinase with thrombospond | Q9UHI8 | ADAMTS1 | Extracellular or secreted | -0.905833333 | 0.452168445 | 26.6041 | 25.5788 | 24.0741 | 24.2681 | 24.0205 | 25.2509 |
| A disintegrin and metalloproteinase with thrombospond | O75173;O75173-2 | ADAMTS4 | Extracellular or secreted | -1.006366667 | 0.413777817 | 22.9401 | 23.9668 | 20.9947 | 22.4026 | 20.708 | 21.7719 |
| ADAMTS-like protein 4 | Q6UY14;Q6UY14-3;Q6UY14- | ADAMTSL4 | Extracellular or secreted | -1.459866667 | 1.674577751 | 23.4241 | 23.0667 | 23.9788 | 22.5203 | 22.0593 | 21.5104 |
| Double-stranded RNA-specific adenosine deaminase | P55265-5;P55265;P55265-4; | ADAR | Other | -1.3989 | 0.555982362 | 22.2656 | 23.1178 | 25.4588 | 21.8042 | 22.2138 | 22.6275 |
| Alpha-adducin | P35611-2;P35611-6;P35611; | ADD1 | Extracellular or secreted | -0.365466667 | 0.366319425 | 25.1792 | 25.5449 | 26.1116 | 25.1471 | 24.7568 | 25.8354 |
| Beta-adducin | P35612 | ADD2 | Extracellular or secreted | -0.751733333 | 0.579091426 | 22.9678 | 23.0257 | 23.579 | 23.3117 | 21.6065 | 22.3991 |
| Gamma-adducin | Q9UEY8-2;Q9UEY8 | ADD3 | Extracellular or secreted | -1.598266667 | 0.796549017 | 24.2599 | 23.8077 | 25.103 | 21.6196 | 22.5303 | 24.2259 |
| Alcohol dehydrogenase class-3 | P11766 | ADH5 | Extracellular or secreted | 0.555566667 | 1.245487883 | 29.8237 | 29.8864 | 29.5906 | 30.573 | 30.3812 | 30.0132 |
| 1,2-dihydroxy-3-keto-5-methylthiopentene dioxygenase | Q9BV57;Q9BV57-2 | ADI1 | Extracellular or secreted | 0.6159 | 0.630839424 | 27.9859 | 27.8737 | 27.1417 | 28.7507 | 28.488 | 27.6103 |
| Adipogenesis regulatory factor | Q15847 | ADIRF | Extracellular or secreted | 3.728166667 | 2.337068793 | 20.2175 | 20.1056 | 21.2036 | 23.2744 | 24.8204 | 24.6164 |
| Adenosine kinase | P55263;P55263-2;P55263-3; | ADK | Other | 0.000466667 | 0.000464579 | 28.8513 | 29.6909 | 28.7271 | 29.2038 | 29.1295 | 28.9374 |
| Activity-dependent neuroprotector homeobox protein | Q9H2P0 | ADNP | Extracellular or secreted | -1.449766667 | 0.317109285 | 25.3941 | 22.7244 | 28.1267 | 25.5232 | 22.192 | 24.1807 |
| 2-aminoethanethiol dioxygenase | Q96SZ5 | ADO | Other | 0.0058 | 0.014607452 | 23.6639 | 23.2631 | 23.5377 | 23.5187 | 23.5606 | 23.4028 |
| Poly(ADP-ribose) glycohydrolase ARH3 | Q9NX46 | ADPRHL2 | Extracellular or secreted | -0.2371 | 0.272120379 | 29.5298 | 28.811 | 28.6931 | 29.1992 | 28.7174 | 28.406 |
| Beta-adrenergic receptor kinase 1 | P25098 | ADRBK1 | Extracellular or secreted | 0.031566667 | 0.042345985 | 21.668 | 21.7331 | 21.2148 | 21.1868 | 21.708 | 21.8158 |
| Proteasomal ubiquitin receptor ADRM1 | Q16186 | ADRM1 | Extracellular or secreted | -0.0483 | 0.039891174 | 26.2817 | 25.699 | 26.6842 | 25.6842 | 26.1292 | 26.7066 |
| Adenylosuccinate lyase | P30566;P30566-2 | ADSL | Other | -0.9066 | 0.956719819 | 27.7154 | 28.2845 | 28.018 | 26.3859 | 27.4185 | 27.4937 |
| Adenylosuccinate synthetase isozyme 2 | P30520 | ADSS | Extracellular or secreted | 0.394266667 | 0.432443894 | 28.8605 | 29.5843 | 29.5936 | 29.1513 | 30.1546 | 29.9153 |
| AF4/FMR2 family member 4 | Q9UHB7;Q9UHB7-2 | AFF4 | Other | -0.8462 | 0.382383422 | 23.0962 | 20.6055 | 22.9464 | 22.0329 | 21.3076 | 20.769 |
| N(4)-(beta-N-acetylglucosaminyl)-L-asparaginase;Glycosy | P20933 | AGA | Extracellular or secreted | 1.3716 | 0.865588871 | 27.3451 | 29.407 | 28.3282 | 29.9018 | 29.9836 | 29.3097 |
| Arf-GAP domain and FG repeat-containing protein 1 | P52594-2;P52594-3;P52594; | AGFG1 | Other | 0.6097 | 0.505131938 | 25.6184 | 25.9892 | 24.4708 | 26.1464 | 26.0155 | 25.7456 |
| Glycogen debranching enzyme;4-alpha-glucanotransfera | P35573;P35573-2;P35573-3 | AGL | Other | -1.201666667 | 1.023731578 | 27.2309 | 27.8707 | 27.8741 | 25.6031 | 26.693 | 27.0746 |
| Protein argonaute-1 | Q9UL18 | AGO1 | Other | -0.671966667 | 0.664211317 | 21.7721 | 22.2653 | 22.1986 | 21.729 | 20.6392 | 21.8519 |
| Protein argonaute-2 | Q9UKV8;Q9UKV8-2 | AGO2 | Other | -0.9199 | 0.809089385 | 27.0352 | 26.0725 | 27.2138 | 26.2176 | 25.0797 | 26.2645 |
| Protein argonaute-3;Protein argonaute-4 | Q9H9G7;Q9H9G7-2;Q9HCK5 | AGO3;AGO4 | Other | 1.211233333 | 0.578342896 | 22.3084 | 22.5493 | 20.4633 | 22.7269 | 21.9916 | 24.2362 |
| Agrin;Agrin N-terminal 110 kDa subunit;Agrin C-terminal | O00468-6;O00468-3;O00468 | AGRN | Extracellular or secreted | 2.234633333 | 1.269609399 | 31.8349 | 32.4491 | 30.5681 | 33.9767 | 33.7102 | 33.8691 |
| Type-1 angiotensin II receptor-associated protein | Q6RW13-2;Q6RW13 | AGTRAP | Extracellular or secreted | 1.3694 | 0.556451556 | 24.4869 | 21.6343 | 24.3476 | 25.0267 | 24.7288 | 24.8215 |
| Protein ELYS | Q8WYP5;Q8WYP5-3;Q8WYP | AHCTF1 | Extracellular or secreted | 0.633633333 | 0.365873277 | 22.6289 | 22.1059 | 24.2301 | 23.4107 | 24.0559 | 23.3992 |
| Adenosylhomocysteinase | P23526;P23526-2 | AHCY | Extracellular or secreted | -0.2305 | 0.776388744 | 31.7657 | 32.0284 | 31.7785 | 31.4205 | 31.6953 | 31.7653 |
| Putative adenosylhomocysteinase 2;Putative adenosylho | O43865-2;O43865;Q96HN2- | AHCYL1;AHCYL | Extracellular or secreted | -0.1449 | 0.100005463 | 24.3661 | 25.0973 | 25.6039 | 24.2904 | 24.7647 | 25.5775 |
| Neuroblast differentiation-associated protein AHNAK | Q09666 | AHNAK | Extracellular or secreted | 1.395533333 | 1.117407567 | 29.6669 | 29.8229 | 31.1147 | 31.2112 | 31.7097 | 31.8702 |
| Activator of 90 kDa heat shock protein ATPase homolog 1 | O95433;O95433-2 | AHSA1 | Extracellular or secreted | -1.0348 | 1.626974976 | 29.385 | 28.9858 | 29.5242 | 27.8204 | 28.4317 | 28.5385 |
| Axin interactor, dorsalization-associated protein | Q96BJ3 | AIDA | Other | 0.0254 | 0.035420559 | 25.5634 | 25.4997 | 24.935 | 25.0901 | 25.5019 | 25.4823 |
| Apoptosis-inducing factor 1, mitochondrial | O95831-3;O95831;O95831-2 | AIFM1 | Other | -1.892266667 | 2.208772183 | 31.8614 | 31.1005 | 31.9496 | 29.8857 | 29.9985 | 29.3505 |
| Absent in melanoma 1 protein | Q9Y4K1 | AIM1 | Extracellular or secreted | -0.341666667 | 0.257047153 | 26.4701 | 25.6044 | 26.1991 | 26.4633 | 24.9208 | 25.8645 |
| Aminoacyl tRNA synthase complex-interacting multifunct | Q12904;Q12904-2 | AIMP1 | Extracellular or secreted | -0.5996 | 1.114195469 | 30.364 | 29.7902 | 30.4014 | 29.455 | 29.5554 | 29.7464 |
| Aminoacyl tRNA synthase complex-interacting multifunct | Q13155 | AIMP2 | Other | -2.157966667 | 1.496005607 | 26.6443 | 26.0076 | 27.7381 | 24.1138 | 24.8242 | 24.9781 |
| AH receptor-interacting protein | O00170 | AIP | Extracellular or secreted | -0.7327 | 0.510061919 | 26.4004 | 26.5281 | 27.9114 | 25.6376 | 26.029 | 26.9752 |
| Adenylate kinase isoenzyme 1 | P00568 | AK1 | Extracellular or secreted | -1.277933333 | 1.034708635 | 28.2501 | 29.0564 | 29.4578 | 26.8744 | 28.4238 | 27.6323 |
| Adenylate kinase 2, mitochondrial;Adenylate kinase 2, m | P54819-2;P54819;P54819-5; | AK2 | Extracellular or secreted | -0.987666667 | 1.311660002 | 32.1998 | 31.6941 | 32.0532 | 30.8588 | 31.5233 | 30.602 |
| GTP:AMP phosphotransferase AK3, mitochondrial | Q9UIJ7;Q9UIJ7-3;Q9UIJ7-2 | AK3 | Other | -1.415133333 | 1.79136194 | 29.415 | 28.9236 | 29.8246 | 27.7038 | 28.1965 | 28.0175 |
| Adenylate kinase 4, mitochondrial | P27144 | AK4 | Extracellular or secreted | -0.792 | 0.236938177 | 22.6942 | 24.06 | 25.3517 | 21.1591 | 24.5727 | 23.9981 |
| A-kinase anchor protein 12 | Q02952-3;Q02952-2;Q02952 | AKAP12 | Extracellular or secreted | -1.504066667 | 0.874172979 | 27.146 | 26.8696 | 26.6406 | 26.4367 | 25.4526 | 24.2547 |
| A-kinase anchor protein 13 | Q12802-4;Q12802;Q12802-2 | AKAP13 | Other | -0.3573 | 0.247414527 | 21.4601 | 20.3661 | 21.9921 | 21.4617 | 20.8277 | 20.457 |
| A-kinase anchor protein 2 | Q9Y2D5-6;Q9Y2D5-4;Q9Y2D | AKAP2 | Other | 0.658866667 | 0.258526401 | 24.7445 | 21.699 | 23.9875 | 23.6968 | 24.1225 | 24.5883 |
| A-kinase anchor protein 8 | O43823 | AKAP8 | Other | -0.334933333 | 0.13075287 | 22.9663 | 22.1172 | 25.0633 | 23.2347 | 22.8149 | 23.0924 |
| A-kinase anchor protein 8-like | Q9ULX6;Q9ULX6-2 | AKAP8L | Other | -1.1733 | 1.433475161 | 27.0954 | 26.6751 | 27.6269 | 25.8318 | 25.869 | 26.1767 |
| Alcohol dehydrogenase [NADP(+)] | P14550 | AKR1A1 | Extracellular or secreted | -0.214333333 | 0.201501741 | 31.4549 | 31.0165 | 30.8957 | 31.434 | 31.0686 | 30.2215 |
| Aldose reductase | P15121 | AKR1B1 | Extracellular or secreted | -0.1883 | 0.255100667 | 34.5503 | 34.9149 | 34.0603 | 34.3059 | 34.5794 | 34.0753 |
| Aldo-keto reductase family 1 member B10 | O60218 | AKR1B10 | Extracellular or secreted | -0.1342 | 0.10318339 | 25.197 | 24.3051 | 24.1468 | 24.9524 | 24.4935 | 23.8004 |
| Aldo-keto reductase family 1 member C1 | Q04828 | AKR1C1 | Extracellular or secreted | 0.231233333 | 0.396456795 | 29.6192 | 29.0186 | 29.2348 | 29.4445 | 29.8529 | 29.2689 |
| Aldo-keto reductase family 1 member C2 | P52895 | AKR1C2 | Other | -0.014966667 | 0.014254813 | 32.569 | 31.4852 | 31.9462 | 32.1893 | 31.9918 | 31.7744 |
| Aldo-keto reductase family 1 member C3 | P42330 | AKR1C3 | Extracellular or secreted | -1.325733333 | 1.099476857 | 29.5107 | 29.5569 | 29.7803 | 28.6964 | 28.7102 | 27.4641 |
| 3-oxo-5-beta-steroid 4-dehydrogenase | P51857-2;P51857 | AKR1D1 | Extracellular or secreted | 0.521333333 | 0.60107942 | 26.894 | 27.3172 | 27.7382 | 27.9709 | 27.2664 | 28.2761 |

| | | | | | | | | | | | |
|---|---|---|---|---|---|---|---|---|---|---|---|
| Aflatoxin B1 aldehyde reductase member 2 | O43488 | AKR7A2 | Extracellular or secreted | -0.344433333 | 0.235005968 | 25.5842 | 26.6147 | 26.4839 | 24.953 | 26.4534 | 26.2431 |
| RAC-beta serine/threonine-protein kinase | P31751-2;P31751 | AKT2 | Extracellular or secreted | -0.0106 | 0.00513775 | 21.6766 | 21.7748 | 20.4385 | 22.2731 | 21.0745 | 20.5105 |
| RAC-gamma serine/threonine-protein kinase | Q9Y243-2;Q9Y243 | AKT3 | Extracellular or secreted | 0.664633333 | 0.663908334 | 23.6658 | 23.1368 | 22.818 | 23.1452 | 24.1491 | 24.3202 |
| Delta-aminolevulinic acid dehydratase | P13716;P13716-2 | ALAD | Extracellular or secreted | 0.1334 | 0.29131807 | 26.5071 | 26.0936 | 26.1186 | 26.6292 | 26.2443 | 26.246 |
| CD166 antigen | Q13740-2;Q13740 | ALCAM | Extracellular or secreted | 0.581333333 | 0.604163451 | 30.7155 | 30.6102 | 29.7886 | 31.4405 | 31.0561 | 30.3617 |
| Aldehyde dehydrogenase family 16 member A1 | Q8IZ83-3;Q8IZ83 | ALDH16A1 | Extracellular or secreted | -1.518466667 | 2.418597197 | 27.1631 | 27.1916 | 27.439 | 25.8538 | 25.4198 | 25.9647 |
| Delta-1-pyrroline-5-carboxylate synthase;Glutamate 5-ki | P54886-2;P54886 | ALDH18A1 | Other | -0.326666667 | 0.07848357 | 21.914 | 23.5956 | 25.7592 | 21.5999 | 23.8883 | 24.8006 |
| Aldehyde dehydrogenase family 1 member A3 | P47895 | ALDH1A3 | Extracellular or secreted | -0.4849 | 0.436888164 | 30.3882 | 31.0705 | 29.6235 | 29.9134 | 29.8868 | 29.8273 |
| Aldehyde dehydrogenase X, mitochondrial | P30837 | ALDH1B1 | Other | -0.929866667 | 0.398561015 | 22.055 | 22.8734 | 24.8978 | 21.4897 | 22.7544 | 22.7925 |
| Aldehyde dehydrogenase, mitochondrial | P05091;P05091-2 | ALDH2 | Extracellular or secreted | -0.703933333 | 0.417542041 | 27.5258 | 27.8783 | 29.5786 | 27.576 | 27.808 | 27.4869 |
| Succinate-semialdehyde dehydrogenase, mitochondrial | P51649;P51649-2 | ALDH5A1 | Other | -1.424233333 | 0.440030314 | 21.9534 | 23.6419 | 24.8886 | 20.6276 | 21.3877 | 24.1959 |
| Alpha-aminoadipic semialdehyde dehydrogenase | P49419-2;P49419;P49419-4 | ALDH7A1 | Extracellular or secreted | -0.028566667 | 0.022095494 | 27.7486 | 28.0905 | 29.0252 | 28.573 | 27.9579 | 28.2477 |
| 4-trimethylaminobutyraldehyde dehydrogenase | P49189;P49189-2 | ALDH9A1 | Extracellular or secreted | -0.020433333 | 0.129326855 | 27.7755 | 27.8091 | 27.8163 | 27.6773 | 27.8049 | 27.8574 |
| Fructose-bisphosphate aldolase A | P04075;P04075-2 | ALDOA | Extracellular or secreted | 0.331533333 | 0.691259302 | 35.1151 | 35.1219 | 34.8758 | 35.5642 | 35.5391 | 35.0041 |
| Fructose-bisphosphate aldolase C | P09972 | ALDOC | Extracellular or secreted | 1.495833333 | 1.969601992 | 30.5373 | 30.5668 | 30.6152 | 32.2503 | 32.2156 | 31.7409 |
| THO complex subunit 4 | Q86V81 | ALYREF | Other | 0.0019 | 0.000979942 | 29.1786 | 27.9772 | 29.973 | 28.692 | 29.1703 | 29.2722 |
| Putative N-acetylglucosamine-6-phosphate deacetylase | Q9Y303;Q9Y303-2;Q9Y303-3 | AMDHD2 | Other | -0.199633333 | 0.232313187 | 24.4002 | 24.0308 | 23.5438 | 24.1072 | 23.9203 | 23.3484 |
| Amphoterin-induced protein 2 | Q86SJ2 | AMIGO2 | Extracellular or secreted | 0.852066667 | 0.563090811 | 28.0322 | 27.6431 | 26.1467 | 28.2898 | 28.3038 | 27.7846 |
| AMP deaminase 2 | Q01433-3;Q01433-5;Q01433 | AMPD2 | Other | -1.2135 | 1.03529384 | 27.2464 | 26.3161 | 27.8256 | 26.1137 | 25.3627 | 26.2712 |
| Alpha-amylase 1;Alpha-amylase 2B;Pancreatic alpha-amy | P04745;P19961;P04746;P19 | AMY1A;AMY2E | Extracellular or secreted | 0.932866667 | 0.404276863 | 29.6412 | 28.9606 | 26.773 | 29.6346 | 29.49 | 29.0488 |
| Anaphase-promoting complex subunit 1 | Q9H1A4 | ANAPC1 | Other | -0.359933333 | 0.147337995 | 20.584 | 20.636 | 22.6697 | 22.0902 | 20.1195 | 20.6002 |
| Anaphase-promoting complex subunit 10 | Q9UM13 | ANAPC10 | Other | -0.1478 | 0.113387161 | 22.6171 | 22.4077 | 21.9583 | 21.3694 | 22.4157 | 22.7546 |
| Angiogenin | P03950 | ANG | Extracellular or secreted | -2.451266667 | 0.843644377 | 24.7309 | 25.713 | 26.7127 | 21.0953 | 24.7017 | 24.0058 |
| Angiopoietin-1 | Q15389-2;Q15389 | ANGPT1 | Extracellular or secreted | -0.018866667 | 0.007390104 | 21.9037 | 22.942 | 20.6313 | 20.8448 | 22.0516 | 22.524 |
| Angiopoietin-related protein 4 | Q9BY76;Q9BY76-3;Q9BY76-2 | ANGPTL4 | Extracellular or secreted | -0.937166667 | 0.122751116 | 29.3967 | 22.3232 | 22.0184 | 26.1017 | 23.0572 | 21.7679 |
| Ankyrin-2 | Q01484-5;Q01484-2;Q01484 | ANK2 | Extracellular or secreted | -0.107633333 | 0.072458177 | 25.0001 | 25.6567 | 25.5904 | 26.1185 | 25.2859 | 24.5199 |
| Rabankyrin-5 | Q9P2R3;Q9P2R3-2;Q9P2R3-4 | ANKFY1 | Extracellular or secreted | -0.0488 | 0.018551319 | 24.0388 | 24.6212 | 26.4064 | 24.4182 | 24.5374 | 25.9644 |
| Ankyrin repeat domain-containing protein 18A;Ankyrin r | Q8IVF6;A2A2Z9 | ANKRD18A;AN | Other | 1.758 | 0.268634394 | 22.3519 | 29.2531 | 29.7627 | 28.6083 | 29.1047 | 28.9287 |
| Actin-binding protein anillin | Q9NQW6-2;Q9NQW6 | ANLN | Other | -0.9456 | 0.291122053 | 22.8795 | 21.0104 | 24.6146 | 23.2515 | 21.8913 | 20.5249 |
| Acidic leucine-rich nuclear phosphoprotein 32 family me | P39687 | ANP32A | Other | 0.350633333 | 0.22343733 | 32.2238 | 31.3185 | 31.1908 | 32.7654 | 32.0114 | 31.0082 |
| Acidic leucine-rich nuclear phosphoprotein 32 family me | Q92688-2;Q92688 | ANP32B | Extracellular or secreted | -0.254866667 | 0.215106588 | 31.987 | 31.4742 | 30.7364 | 31.5047 | 31.3339 | 30.5944 |
| Acidic leucine-rich nuclear phosphoprotein 32 family me | Q9BTT0-3;Q9BTT0;Q9BTT0-2 | ANP32E | Other | -0.091766667 | 0.089431369 | 27.6398 | 27.2306 | 27.7232 | 28.0579 | 26.9779 | 27.2825 |
| Aminopeptidase N | P15144 | ANPEP | Extracellular or secreted | 2.303266667 | 1.882956624 | 31.3211 | 30.7114 | 30.112 | 33.6986 | 33.0554 | 32.3003 |
| Anthrax toxin receptor 1 | Q9H6X2-4;Q9H6X2-6;Q9H6X | ANTXR1 | Extracellular or secreted | -0.0317 | 0.006732071 | 24.5681 | 24.7256 | 20.8357 | 21.6335 | 24.2062 | 24.1946 |
| Annexin A1 | P04083 | ANXA1 | Extracellular or secreted | 1.7936 | 1.341437096 | 31.1542 | 31.9036 | 30.3994 | 32.8243 | 33.1843 | 32.8294 |
| Annexin A11 | P50995-2;P50995 | ANXA11 | Extracellular or secreted | 2.156 | 1.426056222 | 21.3622 | 21.6645 | 22.882 | 24.3097 | 23.9336 | 24.1334 |
| Annexin A2;Putative annexin A2-like protein | P07355;P07355-2;A6NMY6 | ANXA2;ANXA2 | Extracellular or secreted | 2.502366667 | 1.977293472 | 31.9623 | 31.4507 | 30.6673 | 34.2194 | 33.7263 | 33.6417 |
| Annexin A3 | P12429 | ANXA3 | Extracellular or secreted | 1.1591 | 0.939153373 | 31.7947 | 32.32 | 30.8403 | 33.546 | 32.5889 | 32.2974 |
| Annexin A4 | P09525;P09525-2 | ANXA4 | Extracellular or secreted | 3.562466667 | 2.403832137 | 24.3148 | 23.0211 | 23.8615 | 27.6072 | 27.8586 | 26.419 |
| Annexin A5 | P08758 | ANXA5 | Extracellular or secreted | 1.886033333 | 1.408624637 | 30.4504 | 30.8808 | 29.2873 | 32.4522 | 32.1998 | 31.6246 |
| Annexin A6 | P08133-2;P08133 | ANXA6 | Extracellular or secreted | 1.1278 | 0.865691025 | 27.8364 | 28.3926 | 26.7979 | 29.4896 | 28.7219 | 28.1988 |
| Annexin A7 | P20073-2;P20073 | ANXA7 | Extracellular or secreted | 1.7032 | 0.489388761 | 24.6154 | 20.3381 | 23.8284 | 24.7576 | 24.3441 | 24.7898 |
| Annexin A8 | P13928;P13928-3 | ANXA8 | Extracellular or secreted | 0.220933333 | 0.153205026 | 28.3076 | 28.7504 | 27.0921 | 28.2829 | 28.4605 | 28.0695 |
| Annexin A8-like protein 2 | Q5VT79;Q5VT79-2 | ANXA8L2 | Extracellular or secreted | -0.970733333 | 1.359367562 | 22.5955 | 23.2429 | 22.7987 | 22.4219 | 21.7074 | 21.5956 |
| Amiloride-sensitive amine oxidase [copper-containing] | P19801;P19801-2 | AOC1 | Extracellular or secreted | -0.470733333 | 0.266312757 | 22.6184 | 24.0032 | 22.4842 | 21.6362 | 23.4062 | 22.6512 |
| AP-1 complex subunit beta-1 | Q10567-3;Q10567-2;Q10567 | AP1B1 | Other | -0.550166667 | 0.549314007 | 28.8136 | 28.8133 | 29.3463 | 27.9681 | 28.1701 | 29.1845 |
| AP-1 complex subunit gamma-1 | O43747;O43747-2 | AP1G1 | Other | -0.4582 | 0.242311559 | 26.0639 | 26.3018 | 27.265 | 24.9009 | 26.2965 | 27.0587 |
| AP-1 complex subunit gamma-like 2 | O75843 | AP1G2 | Other | -0.645666667 | 0.265065174 | 23.3307 | 23.8986 | 26.1808 | 23.4413 | 23.6319 | 24.3999 |
| AP-1 complex subunit mu-1 | Q9BXS5;Q9BXS5-2 | AP1M1 | Extracellular or secreted | -0.372333333 | 0.230248457 | 27.2066 | 27.0665 | 28.272 | 26.4604 | 26.8458 | 28.1219 |
| AP-1 complex subunit mu-2 | Q9Y6Q5;Q9Y6Q5-2 | AP1M2 | Other | -0.232866667 | 0.101752856 | 26.0424 | 25.8721 | 27.0038 | 24.7615 | 26.2109 | 27.2473 |
| AP-1 complex subunit sigma-1A | P61966;P61966-2 | AP1S1 | Extracellular or secreted | -0.548633333 | 0.275168231 | 26.0152 | 26.4096 | 27.1057 | 24.7246 | 26.0114 | 27.1486 |
| AP-2 complex subunit alpha-1 | O95782-2;O95782 | AP2A1 | Extracellular or secreted | -1.197633333 | 0.465641446 | 28.4364 | 27.8273 | 28.7355 | 25.2981 | 27.5133 | 28.5949 |
| AP-2 complex subunit alpha-2 | O94973;O94973-2;O94973-3 | AP2A2 | Extracellular or secreted | -1.3821 | 0.550019367 | 28.1211 | 28.631 | 29.2167 | 25.5356 | 27.4641 | 28.8228 |
| AP-2 complex subunit beta | P63010;P63010-2;P63010-3 | AP2B1 | Extracellular or secreted | -0.7615 | 0.510539765 | 29.6837 | 30.109 | 30.568 | 28.4443 | 29.2607 | 30.3712 |
| AP-2 complex subunit mu | Q96CW1-2;Q96CW1 | AP2M1 | Extracellular or secreted | -0.585733333 | 0.395288297 | 27.7221 | 28.1032 | 28.856 | 26.8019 | 27.5423 | 28.5799 |
| AP-2 complex subunit sigma | P53680-2;P53680 | AP2S1 | Extracellular or secreted | -0.365133333 | 0.193926674 | 24.376 | 24.6364 | 25.7386 | 23.787 | 24.1734 | 25.6952 |
| AP-3 complex subunit beta-1 | O00203-3;O00203 | AP3B1 | Other | -0.7843 | 0.689754855 | 25.3572 | 25.1137 | 26.2251 | 24.0763 | 24.8442 | 25.4226 |
| AP-3 complex subunit delta-1 | O14617-5;O14617-2;O14617 | AP3D1 | Other | -1.5458 | 0.688069051 | 26.2294 | 25.6323 | 27.2333 | 23.1666 | 25.3217 | 25.9693 |
| AP-3 complex subunit mu-1 | Q9Y2T2 | AP3M1 | Other | -0.1209 | 0.062782631 | 22.2179 | 22.4194 | 23.4636 | 21.5396 | 22.8396 | 23.359 |
| AP-3 complex subunit sigma-1 | Q92572 | AP3S1 | Other | -0.633766667 | 0.173819757 | 21.7175 | 23.122 | 24.3651 | 22.8247 | 20.3064 | 24.1722 |
| Acylamino-acid-releasing enzyme | P13798 | APEH | Extracellular or secreted | -0.111133333 | 0.073090418 | 32.2773 | 31.5874 | 31.052 | 32.2379 | 31.4829 | 30.8625 |
| DNA-(apurinic or apyrimidinic site) lyase;DNA-(apurinic o | P27695 | APEX1 | Other | -0.313233333 | 0.247032803 | 32.5284 | 31.9823 | 31.8219 | 32.57 | 31.7622 | 31.0607 |
| Apoptosis inhibitor 5 | Q9BZZ5-2;Q9BZZ5;Q9BZZ5-3 | API5 | Other | 0.271666667 | 0.1020798 | 26.5441 | 26.5737 | 28.2923 | 26.07 | 27.4647 | 28.6904 |
| Methylthioribulose-1-phosphate dehydratase | Q96GX9;Q96GX9-3 | APIP | Other | 0.630366667 | 0.858554381 | 27.9542 | 27.1567 | 27.1012 | 28.2053 | 28.0187 | 27.8792 |
| Aprataxin and PNK-like factor | Q8IW19 | APLF | Other | 0.025 | 0.0110615 | 21.8278 | 23.6455 | 23.3187 | 23.7016 | 22.0284 | 23.137 |
| Amyloid-like protein 2 | Q06481-3;Q06481-6;Q06481 | APLP2 | Extracellular or secreted | 1.005 | 0.584597957 | 29.6487 | 28.9862 | 27.4426 | 29.9856 | 29.8512 | 29.2557 |
| Adipocyte plasma membrane-associated protein | Q9HDC9-2;Q9HDC9 | APMAP | Extracellular or secreted | 2.170133333 | 1.03109977 | 21.4175 | 22.1935 | 23.9853 | 25.1718 | 24.4593 | 24.4756 |
| NAD(P)H-hydrate epimerase | Q8NCW5;Q8NCW5-2 | APOA1BP | Extracellular or secreted | -0.1555 | 0.092539177 | 30.4698 | 30.1643 | 29.4888 | 30.6785 | 30.0523 | 28.9256 |
| Apolipoprotein B-100;Apolipoprotein B-48 | P04114 | APOB | Extracellular or secreted | -1.295433333 | 1.29130378 | 22.6943 | 22.8371 | 22.0535 | 21.7065 | 20.5054 | 21.4867 |
| DNA dC->dU-editing enzyme APOBEC-3C | Q9NRW3 | APOBEC3C | Other | 2.145566667 | 0.640955313 | 23.0875 | 21.6248 | 20.1586 | 24.5698 | 21.3851 | 25.3527 |
| Amyloid beta A4 protein;N-APP;Soluble APP-alpha;Solub | P05067-11;P05067-8;P05067 | APP | Extracellular or secreted | 0.4762 | 0.346449426 | 31.9572 | 31.6664 | 30.332 | 32.0962 | 31.8727 | 31.4153 |
| DCC-interacting protein 13-alpha | Q9UKG1 | APPL1 | Extracellular or secreted | -1.141166667 | 0.353723438 | 23.8143 | 24.267 | 24.7628 | 20.749 | 24.449 | 24.2226 |
| Adenine phosphoribosyltransferase | P07741;P07741-2 | APRT | Extracellular or secreted | 0.7227 | 1.042636339 | 26.7754 | 26.8722 | 27.5983 | 27.483 | 27.8832 | 28.0478 |
| Intron-binding protein aquarius | O60306 | AQR | Other | -3.887033333 | 1.48902835 | 28.637 | 28.6329 | 31.1394 | 26.0366 | 25.6512 | 25.0604 |
| Arf-GAP with Rho-GAP domain, ANK repeat and PH doma | Q8WZ64 | ARAP2 | Other | -0.5416 | 0.115800717 | 23.5819 | 26.0626 | 24.3045 | 24.0462 | 21.5659 | 26.7121 |
| Coatomer subunit delta | P48444;P48444-2 | ARCN1 | Extracellular or secreted | -0.5685 | 0.867048076 | 30.166 | 30.196 | 30.3366 | 29.2698 | 29.6244 | 30.0989 |
| Amphiregulin | P15514 | AREG | Extracellular or secreted | 1.756 | 0.33998215 | 27.1016 | 25.3182 | 20.8801 | 27.9416 | 25.1635 | 25.4628 |
| ADP-ribosylation factor 1;ADP-ribosylation factor 3 | P84077;P61204;P61204-2 | ARF1;ARF3 | Extracellular or secreted | -0.219933333 | 0.183313329 | 28.0832 | 28.9837 | 29.2048 | 27.9393 | 28.7685 | 28.9041 |
| ADP-ribosylation factor 4 | P18085 | ARF4 | Extracellular or secreted | 0.349 | 0.523798273 | 24.3974 | 23.6927 | 24.3034 | 24.6708 | 24.6716 | 24.0981 |
| ADP-ribosylation factor 5 | P84085 | ARF5 | Extracellular or secreted | -0.507733333 | 0.327023744 | 23.8242 | 23.1798 | 23.3361 | 21.8127 | 23.6103 | 23.3939 |
| ADP-ribosylation factor 6 | P62330 | ARF6 | Extracellular or secreted | 0.736866667 | 0.645989199 | 23.9407 | 23.1584 | 24.6531 | 24.2493 | 24.7298 | 24.9837 |
| ADP-ribosylation factor GTPase-activating protein 1 | Q8N6T3-4;Q8N6T3-5;Q8N6T | ARFGAP1 | Other | 0.531166667 | 0.255494288 | 21.076 | 23.4183 | 22.324 | 22.0529 | 22.7309 | 23.628 |
| ADP-ribosylation factor GTPase-activating protein 2 | Q8N6H7 | ARFGAP2 | Extracellular or secreted | 0.064166667 | 0.051475912 | 23.5658 | 23.6273 | 23.332 | 22.9937 | 23.3893 | 24.3346 |
| ADP-ribosylation factor GTPase-activating protein 3 | Q9NP61-2;Q9NP61 | ARFGAP3 | Other | 0.393533333 | 0.606424985 | 21.769 | 21.9646 | 22.0246 | 21.8776 | 22.732 | 22.3292 |
| Brefeldin A-inhibited guanine nucleotide-exchange prote | Q9Y6D6 | ARFGEF1 | Other | -2.3482 | 0.997379556 | 25.4837 | 24.8096 | 24.2205 | 22.5815 | 23.9629 | 20.9248 |
| Brefeldin A-inhibited guanine nucleotide-exchange prote | Q9Y6D5 | ARFGEF2 | Other | 1.4068 | 0.545130251 | 23.5234 | 23.9742 | 26.6451 | 25.5761 | 25.8515 | 26.9355 |
| Arfaptin-1 | P53367;P53367-2 | ARFIP1 | Other | 0.868766667 | 0.565845516 | 23.1164 | 24.5728 | 25.035 | 25.1206 | 25.0903 | 25.1196 |
| Arginase-1 | P05089;P05089-2;P05089-3 | ARG1 | Extracellular or secreted | 2.7015 | 1.101546126 | 22.0896 | 21.6996 | 23.9707 | 27.02 | 24.0669 | 24.7775 |
| Rho GTPase-activating protein 1 | Q07960 | ARHGAP1 | Extracellular or secreted | -0.784 | 0.550788268 | 26.7362 | 27.1245 | 28.1124 | 25.745 | 26.4852 | 27.3909 |
| Rho GTPase-activating protein 12 | Q8IWW6-3;Q8IWW6-2;Q8IW | ARHGAP12 | Other | 2.188566667 | 0.631076089 | 24.9125 | 26.1065 | 21.6865 | 26.0482 | 27.4385 | 25.7845 |
| Rho GDP-dissociation inhibitor 1 | P52565;P52565-2 | ARHGDIA | Extracellular or secreted | 1.5895 | 2.580347075 | 30.2704 | 30.843 | 30.65 | 32.143 | 32.3761 | 32.0128 |
| Rho GDP-dissociation inhibitor 2 | P52566 | ARHGDIB | Extracellular or secreted | 0.381766667 | 0.335487663 | 27.1674 | 25.9162 | 26.9206 | 27.5124 | 27.0284 | 26.6087 |
| Rho guanine nucleotide exchange factor 1 | Q92888;Q92888-3;Q92888-2 | ARHGEF1 | Extracellular or secreted | -0.935533333 | 0.184282592 | 23.3461 | 23.3659 | 25.6897 | 20.3382 | 23.0208 | 26.2361 |
| Rho guanine nucleotide exchange factor 6;Rho guanine n | Q15052-2;Q14155-1;Q14155 | ARHGEF6;ARH( | Other | 0.344833333 | 0.112395353 | 21.1672 | 22.3366 | 23.0849 | 20.7427 | 22.9531 | 23.9274 |
| AT-rich interactive domain-containing protein 1A | O14497;O14497-2;O14497-3 | ARID1A | Other | -2.1048 | 0.823300392 | 26.2571 | 26.6594 | 28.3939 | 24.1269 | 23.9803 | 26.8888 |
| AT-rich interactive domain-containing protein 4A | P29374-3;P29374-2;P29374 | ARID4A | Other | -2.367466667 | 0.584252127 | 25.7296 | 25.7162 | 21.3012 | 23.2472 | 22.3758 | 20.0216 |
| AT-rich interactive domain-containing protein 4B | Q4LE39-3;Q4LE39-2;Q4LE39 | ARID4B | Other | -0.1759 | 0.033580955 | 23.9659 | 19.3753 | 24.1027 | 22.4407 | 20.9275 | 23.548 |
| E3 ubiquitin-protein ligase ARIH1 | Q9Y4X5 | ARIH1 | Other | -1.126033333 | 0.666151369 | 26.4324 | 26.1935 | 26.6711 | 24.0341 | 25.8653 | 26.0195 |
| E3 ubiquitin-protein ligase ARIH2 | O95376 | ARIH2 | Other | -0.3202 | 0.355212603 | 23.5212 | 23.8305 | 24.483 | 23.2086 | 23.6112 | 24.0543 |
| ADP-ribosylation factor-like protein 2 | P36404;P36404-2 | ARL2 | Extracellular or secreted | -0.246133333 | 0.148616131 | 21.7651 | 22.7763 | 23.632 | 23.0007 | 22.1599 | 22.2744 |
| ADP-ribosylation factor-like protein 3 | P36405 | ARL3 | Extracellular or secreted | 0.111733333 | 0.059842633 | 22.0097 | 21.6963 | 23.6308 | 22.637 | 22.2818 | 22.7532 |
| ADP-ribosylation factor-like protein 6-interacting protein | Q66PJ3-7;Q66PJ3-4;Q66PJ3- | ARL6IP4 | Other | -1.072633333 | 0.477132445 | 22.4087 | 23.4997 | 23.3325 | 22.3149 | 23.2891 | 20.419 |
| PRA1 family protein 3 | O75915 | ARL6IP5 | Extracellular or secreted | 1.664733333 | 1.027471417 | 22.5017 | 21.589 | 23.6891 | 23.6937 | 24.0814 | 24.9989 |
| ADP-ribosylation factor-like protein 8B | Q9NVJ2 | ARL8B | Extracellular or secreted | 0.047433333 | 0.019612548 | 23.6574 | 23.7005 | 25.8089 | 23.8079 | 24.9603 | 24.5409 |
| Protein-glutamate O-methyltransferase | Q9H993 | ARMT1 | Other | -0.123066667 | 0.248842342 | 28.2907 | 28.0632 | 27.7629 | 28.1576 | 27.8056 | 27.7844 |
| Actin-related protein 2/3 complex subunit 1A | Q92747;Q92747-2 | ARPC1A | Extracellular or secreted | 0.1187 | 0.110016604 | 25.2191 | 25.806 | 25.4477 | 24.9385 | 25.8427 | 26.0477 |
| Actin-related protein 2/3 complex subunit 1B | O15143 | ARPC1B | Extracellular or secreted | 0.369433333 | 0.609286579 | 30.2327 | 30.0215 | 30.4677 | 30.2906 | 30.4929 | 31.0467 |
| Actin-related protein 2/3 complex subunit 2 | O15144 | ARPC2 | Extracellular or secreted | -0.032666667 | 0.033621526 | 30.3337 | 30.3389 | 30.3879 | 29.7686 | 30.3569 | 30.837 |
| Actin-related protein 2/3 complex subunit 3 | O15145 | ARPC3 | Extracellular or secreted | 0.2111 | 0.360403242 | 29.2327 | 29.3818 | 29.6118 | 29.229 | 29.6891 | 29.9415 |
| Actin-related protein 2/3 complex subunit 4 | P59998;P59998-3;P59998-4; | ARPC4 | Extracellular or secreted | 0.241133333 | 0.396842396 | 29.1546 | 29.2683 | 29.4411 | 29.1333 | 29.5414 | 29.9127 |
| Actin-related protein 2/3 complex subunit 5 | O15511;O15511-2 | ARPC5 | Extracellular or secreted | 0.245533333 | 0.388741717 | 28.6903 | 28.5652 | 28.956 | 28.566 | 29.0299 | 29.3522 |
| Actin-related protein 2/3 complex subunit 5-like protein | Q9BPX5 | ARPC5L | Extracellular or secreted | 0.3053 | 0.286740286 | 25.5032 | 25.7528 | 25.961 | 25.3402 | 26.6589 | 26.1338 |
| Arpin | Q7Z6K5;Q7Z6K5-2 | ARPIN | Other | -3.336 | 1.573250217 | 24.4921 | 24.5875 | 25.1029 | 20.7574 | 20.7496 | 22.6675 |
| cAMP-regulated phosphoprotein 19 | P56211;P56211-2 | ARPP19 | Other | 0.205933333 | 0.074061818 | 23.2454 | 21.6076 | 24.705 | 23.1905 | 23.9271 | 23.0582 |
| Arrestin domain-containing protein 1 | Q8N5I2 | ARRDC1 | Extracellular or secreted | -0.542 | 0.212748847 | 23.4898 | 25.1047 | 22.4291 | 24.1675 | 22.0819 | 23.1482 |
| Arylsulfatase A;Arylsulfatase A component B;Arylsulfatas | P15289-2;P15289 | ARSA | Extracellular or secreted | 1.270766667 | 1.57877224 | 25.1776 | 26.0627 | 25.2461 | 26.8204 | 27.0467 | 26.4316 |
| Arylsulfatase B | P15848;P15848-2 | ARSB | Extracellular or secreted | -0.656866667 | 0.494597907 | 24.7061 | 24.3244 | 23.775 | 22.9077 | 24.5466 | 23.3806 |
| Acid ceramidase;Acid ceramidase subunit alpha;Acid cera | Q13510;Q13510-2;Q13510-3 | ASAH1 | Extracellular or secreted | 0.843066667 | 1.359916384 | 28.4007 | 28.3674 | 28.0192 | 28.752 | 29.5208 | 29.0437 |
| Activating signal cointegrator 1 complex subunit 1 | Q8N9N2-2;Q8N9N2 | ASCC1 | Other | 0.160433333 | 0.133896391 | 23.1156 | 23.5036 | 23.9799 | 23.2539 | 23.4207 | 24.4058 |

| | | | | | | | | | | | |
|---|---|---|---|---|---|---|---|---|---|---|---|
| Histone chaperone ASF1A | Q9Y294 | ASF1A | Other | -0.056266667 | 0.050202593 | 26.4382 | 25.2786 | 25.456 | 25.5738 | 25.7103 | 25.7199 |
| Set1/Ash2 histone methyltransferase complex subunit AS | Q9UBL3-3;Q9UBL3;Q9UBL3- | ASH2L | Other | -1.5583 | 0.791663787 | 23.9635 | 23.2741 | 25.3962 | 21.4509 | 22.7858 | 23.7222 |
| Argininosuccinate lyase | P04424;P04424-3;P04424-2 | ASL | Extracellular or secreted | 0.646566667 | 1.91373533 | 26.7229 | 26.5204 | 26.4472 | 27.2339 | 27.1705 | 27.2258 |
| ATPase ASNA1 | O43681 | ASNA1 | Extracellular or secreted | -0.6666 | 0.435944162 | 26.9192 | 26.7085 | 27.5592 | 25.3246 | 26.6204 | 27.2421 |
| Asparagine synthetase [glutamine-hydrolyzing] | P08243-2;P08243;P08243-3 | ASNS | Other | -0.336266667 | 0.219427723 | 27.2573 | 27.7445 | 28.3992 | 26.4912 | 27.8648 | 28.0362 |
| Aspartyl/asparaginyl beta-hydroxylase | Q12797-10;Q12797 | ASPH | Extracellular or secreted | -0.7438 | 0.222256426 | 22.4359 | 23.4401 | 23.9984 | 23.7981 | 23.6212 | 20.2237 |
| Tether containing UBX domain for GLUT4 | Q9BZE9;Q9BZE9-2;Q9BZE9-3 | ASPSCR1 | Extracellular or secreted | -1.279933333 | 0.692473431 | 24.8175 | 25.2338 | 26.9159 | 23.455 | 24.356 | 25.3164 |
| Argininosuccinate synthase | P00966 | ASS1 | Extracellular or secreted | -0.2722 | 0.481858784 | 28.6082 | 28.8673 | 29.196 | 28.2656 | 28.7689 | 28.8204 |
| Protein asunder homolog | Q9NVM9;Q9NVM9-2 | ASUN | Other | -2.155466667 | 0.70970617 | 25.5191 | 25.486 | 27.5441 | 22.4217 | 23.433 | 26.2281 |
| Arginyl-tRNA--protein transferase 1 | O95260-2;O95260 | ATE1 | Other | -2.967033333 | 1.223152939 | 26.3008 | 26.1475 | 26.3623 | 21.7868 | 23.8628 | 24.2599 |
| Cyclic AMP-dependent transcription factor ATF-1;Cyclic A | P18846-2;P18846;Q03060-2 | ATF1;CREB1 | Other | 0.690666667 | 0.345919995 | 24.4875 | 23.5983 | 26.1108 | 25.0527 | 25.3882 | 25.8277 |
| Activating transcription factor 7-interacting protein 1 | Q6VMQ6-2;Q6VMQ6-5;Q6V | ATF7IP | Other | -2.133966667 | 0.821447897 | 24.5202 | 24.0184 | 24.7089 | 20.725 | 22.0891 | 24.0315 |
| Autophagy-related protein 16-1 | Q676U5-3;Q676U5-2;Q676U | ATG16L1 | Other | 0.581466667 | 0.159767574 | 20.9389 | 24.2869 | 25.0854 | 23.979 | 23.9264 | 24.1502 |
| Ubiquitin-like-conjugating enzyme ATG3 | Q9NT62;Q9NT62-2 | ATG3 | Other | -0.206766667 | 0.151704974 | 24.2884 | 23.8486 | 25.1629 | 23.8571 | 23.9386 | 24.8839 |
| Autophagy protein 5 | Q9H1Y0-2;Q9H1Y0 | ATG5 | Extracellular or secreted | -0.380266667 | 0.407777596 | 23.7258 | 24.1667 | 24.434 | 23.2063 | 23.6475 | 24.3319 |
| Ubiquitin-like modifier-activating enzyme ATG7 | O95352-2;O95352;O95352-3 | ATG7 | Other | -0.164633333 | 0.0536834 | 24.1638 | 24.7259 | 25.0688 | 23.2411 | 23.8148 | 26.4087 |
| Bifunctional purine biosynthesis protein PURH;Phosphor | P31939;P31939-2 | ATIC | Extracellular or secreted | 0.031333333 | 0.042237974 | 30.1649 | 30.2385 | 30.8579 | 30.4925 | 30.2366 | 30.6262 |
| Atlastin-2 | Q8NHH9-3;Q8NHH9-5;Q8NH | ATL2 | Other | -0.035466667 | 0.013060883 | 22.8528 | 22.9788 | 25.3432 | 23.4176 | 23.335 | 24.3158 |
| Atlastin-3 | Q6DD88 | ATL3 | Other | 0.694366667 | 0.397902989 | 25.2269 | 25.513 | 27.231 | 26.4005 | 26.2877 | 27.3658 |
| Copper transport protein ATOX1 | O00244 | ATOX1 | Other | 1.2277 | 1.178786887 | 25.2638 | 25.8427 | 25.451 | 26.4327 | 27.5033 | 26.3046 |
| Sodium/potassium-transporting ATPase subunit alpha-1 | P05023-4;P05023;P05023-3; | ATP1A1 | Extracellular or secreted | 2.4478 | 1.855692441 | 30.7248 | 29.3237 | 29.5543 | 32.6037 | 32.533 | 31.8095 |
| Sodium/potassium-transporting ATPase subunit beta-1 | P05026-2;P05026 | ATP1B1 | Extracellular or secreted | 3.333733333 | 1.583545546 | 27.0365 | 25.3703 | 24.9567 | 29.2552 | 29.3737 | 28.7358 |
| Sodium/potassium-transporting ATPase subunit beta-3 | P54709 | ATP1B3 | Extracellular or secreted | 1.422266667 | 1.81873539 | 27.5048 | 26.6738 | 26.7443 | 28.5926 | 28.6066 | 27.9905 |
| Sarcoplasmic/endoplasmic reticulum calcium ATPase 2 | P16615-5;P16615-2;P16615- | ATP2A2 | Extracellular or secreted | 3.308333333 | 3.026215772 | 20.8747 | 21.0791 | 21.7873 | 24.5111 | 25.0138 | 24.1412 |
| Plasma membrane calcium-transporting ATPase 1 | P20020-6;P20020;P20020-4; | ATP2B1 | Extracellular or secreted | 1.3511 | 1.04988411 | 26.0312 | 25.8989 | 24.5209 | 26.9379 | 27.1939 | 26.3725 |
| Plasma membrane calcium-transporting ATPase 4 | P23634-7;P23634-6;P23634- | ATP2B4 | Extracellular or secreted | 1.847433333 | 2.146913101 | 26.5677 | 25.6456 | 25.9578 | 28.1277 | 28.0388 | 27.5469 |
| ATP synthase subunit alpha, mitochondrial | P25705;P25705-2;P25705-3 | ATP5A1 | Extracellular or secreted | 2.075933333 | 0.434300771 | 27.5638 | 27.8536 | 29.3605 | 33.9165 | 28.6972 | 28.392 |
| ATP synthase subunit beta, mitochondrial | P06576 | ATP5B | Extracellular or secreted | 0.387466667 | 0.106130345 | 30.2988 | 29.9738 | 32.1522 | 33.3861 | 30.3864 | 29.8147 |
| ATP synthase subunit gamma, mitochondrial | P36542;P36542-2 | ATP5C1 | Extracellular or secreted | 3.268133333 | 0.671414294 | 21.4691 | 21.2847 | 22.0215 | 28.4533 | 23.574 | 22.5524 |
| ATP synthase subunit delta, mitochondrial | P30049 | ATP5D | Other | 1.139833333 | 1.170820275 | 24.8351 | 25.4071 | 25.5266 | 26.4416 | 27.0005 | 25.7462 |
| ATP synthase F(0) complex subunit B1, mitochondrial | P24539 | ATP5F1 | Extracellular or secreted | 2.926166667 | 1.62086003 | 20.9893 | 20.4414 | 19.4673 | 24.0984 | 23.5902 | 21.9879 |
| ATP synthase subunit d, mitochondrial | O75947-2;O75947 | ATP5H | Extracellular or secreted | 2.374566667 | 1.051290852 | 20.1056 | 22.0451 | 22.9412 | 24.7033 | 24.0288 | 23.4835 |
| ATP synthase subunit O, mitochondrial | P48047 | ATP5O | Extracellular or secreted | 3.453033333 | 0.57124862 | 21.0506 | 22.2011 | 20.9025 | 29.4095 | 22.6845 | 22.4193 |
| V-type proton ATPase subunit S1 | Q15904 | ATP6AP1 | Extracellular or secreted | 4.921533333 | 1.367355358 | 22.1883 | 25.725 | 22.0956 | 27.4931 | 28.4551 | 28.8253 |
| Renin receptor | O75787-2;O75787 | ATP6AP2 | Extracellular or secreted | 1.086833333 | 0.402583667 | 24.3535 | 26.0054 | 25.276 | 24.4925 | 26.5285 | 27.8744 |
| V-type proton ATPase subunit d 1 | P61421 | ATP6V0D1 | Extracellular or secreted | 0.9451 | 1.387166527 | 24.4541 | 24.3173 | 24.9023 | 25.0404 | 25.5836 | 25.885 |
| V-type proton ATPase catalytic subunit A | P38606;P38606-2 | ATP6V1A | Extracellular or secreted | -0.050166667 | 0.038133717 | 28.6783 | 28.6425 | 29.3258 | 28.0726 | 29.219 | 29.2045 |
| V-type proton ATPase subunit B, brain isoform | P21281 | ATP6V1B2 | Extracellular or secreted | -0.286933333 | 0.218050552 | 27.7439 | 27.5745 | 28.468 | 26.919 | 27.6177 | 28.3889 |
| V-type proton ATPase subunit C 1 | P21283 | ATP6V1C1 | Extracellular or secreted | 0.765233333 | 0.623445511 | 23.7726 | 23.6093 | 24.8228 | 24.2722 | 24.6205 | 25.6077 |
| V-type proton ATPase subunit D | Q9Y5K8 | ATP6V1D | Extracellular or secreted | 0.653533333 | 0.594798507 | 23.4166 | 23.5677 | 24.6297 | 23.9584 | 24.6235 | 24.9927 |
| V-type proton ATPase subunit E 1 | P36543-2;P36543;P36543-3 | ATP6V1E1 | Extracellular or secreted | 0.9852 | 0.792525785 | 24.1843 | 23.6991 | 24.3541 | 24.6749 | 26.0053 | 24.5129 |
| V-type proton ATPase subunit F | Q16864;Q16864-2 | ATP6V1F | Extracellular or secreted | 2.038 | 2.513808588 | 24.5186 | 24.7524 | 25.3099 | 26.5681 | 27.1089 | 27.0179 |
| V-type proton ATPase subunit G 1 | O75348 | ATP6V1G1 | Extracellular or secreted | 1.162366667 | 1.524861963 | 24.3257 | 24.639 | 25.215 | 25.4279 | 26.1878 | 26.0511 |
| V-type proton ATPase subunit H | Q9UI12-2;Q9UI12 | ATP6V1H | Extracellular or secreted | -0.565466667 | 0.158173664 | 24.6039 | 24.527 | 26.1608 | 22.2057 | 25.3105 | 26.0791 |
| Attractin | O75882-2;O75882-3;O75882 | ATRN | Extracellular or secreted | 0.962166667 | 0.522167221 | 29.8919 | 30.3514 | 28.145 | 31.0036 | 30.6224 | 29.6488 |
| Transcriptional regulator ATRX | P46100-2;P46100-5;P46100- | ATRX | Other | -1.932766667 | 0.704352703 | 23.7772 | 23.4081 | 23.3629 | 23.2917 | 21.7019 | 19.7563 |
| Ataxin-10 | Q9UBB4-2;Q9UBB4 | ATXN10 | Extracellular or secreted | -2.050733333 | 0.589500462 | 25.6901 | 25.7512 | 26.6558 | 21.46 | 24.5699 | 25.915 |
| Ataxin-2 | Q99700-2;Q99700-4;Q99700 | ATXN2 | Other | -0.398366667 | 0.061376553 | 25.5759 | 19.7161 | 25.4866 | 22.8013 | 21.5761 | 25.2061 |
| Ataxin-2-like protein | Q8WWM7-6;Q8WWM7-8;Q | ATXN2L | Other | 0.9592 | 1.483934461 | 25.8741 | 25.6937 | 25.8355 | 26.5334 | 26.5909 | 27.1566 |
| Aurora kinase A | O14965 | AURKA | Other | -1.287433333 | 0.268347314 | 25.9636 | 21.1969 | 26.0875 | 24.9919 | 21.6946 | 22.6992 |
| Tyrosine-protein kinase receptor UFO | P30530-2;P30530 | AXL | Extracellular or secreted | 2.563666667 | 0.696347352 | 25.0711 | 25.0901 | 21.5806 | 28.8308 | 25.7526 | 24.8494 |
| Zinc-alpha-2-glycoprotein | P25311 | AZGP1 | Extracellular or secreted | 0.8182 | 0.742112771 | 27.255 | 26.4912 | 26.108 | 28.0998 | 27.4092 | 26.7998 |
| Beta-2-microglobulin;Beta-2-microglobulin form pI 5.3 | P61769 | B2M | Extracellular or secreted | 0.6857 | 0.192590289 | 29.4651 | 31.8074 | 27.9414 | 30.7504 | 31.5204 | 29.0002 |
| Galactosylgalactosylxylosylprotein 3-beta-glucuronosyltr | O94766-2;O94766 | B3GAT3 | Extracellular or secreted | 0.528933333 | 0.896094706 | 23.3504 | 23.9511 | 23.3483 | 23.7089 | 24.2051 | 24.3226 |
| N-acetyllactosaminide beta-1,3-N-acetylglucosaminyltra | Q9NY97-2;Q9NY97 | B3GNT2 | Extracellular or secreted | -0.271133333 | 0.13379348 | 23.3077 | 24.5053 | 23.4774 | 22.2896 | 24.4321 | 23.7553 |
| Beta-1,4-galactosyltransferase 1;Lactose synthase A prot | P15291-2;P15291 | B4GALT1 | Extracellular or secreted | -0.561366667 | 0.425119597 | 29.122 | 29.4741 | 27.9904 | 27.6948 | 28.3626 | 28.845 |
| Beta-1,4-galactosyltransferase 4;N-acetyllactosamine syn | O60513 | B4GALT4 | Extracellular or secreted | -0.159233333 | 0.087104726 | 26.2988 | 27.321 | 26.2721 | 25.5198 | 26.5018 | 27.3926 |
| Beta-1,4-galactosyltransferase 5 | O43286 | B4GALT5 | Extracellular or secreted | 0.599766667 | 0.533233339 | 26.1873 | 27.3614 | 25.9984 | 26.9625 | 27.3002 | 27.0837 |
| Beta-1,4-glucuronyltransferase 1 | O43505 | B4GAT1 | Extracellular or secreted | 0.123466667 | 0.044934121 | 27.1564 | 26.5891 | 24.4177 | 25.7309 | 26.914 | 25.8887 |
| BRISC and BRCA1-A complex member 1 | Q9NWV8 | BABAM1 | Other | 0.2251 | 0.14273606 | 25.1529 | 24.3067 | 25.9389 | 25.7244 | 24.6803 | 25.6691 |
| BAG family molecular chaperone regulator 2 | O95816;O95816-2 | BAG2 | Other | -0.706766667 | 0.750954796 | 26.8146 | 26.7871 | 26.9366 | 25.9092 | 25.6854 | 26.8234 |
| BAG family molecular chaperone regulator 3 | O95817 | BAG3 | Extracellular or secreted | 0.836833333 | 1.051175604 | 25.5975 | 25.464 | 25.3521 | 26.8593 | 26.1293 | 25.9355 |
| BAG family molecular chaperone regulator 5 | Q9UL15;Q9UL15-2 | BAG5 | Other | -0.865666667 | 0.455207024 | 19.3547 | 20.2909 | 21.6602 | 19.7601 | 20.2282 | 18.7205 |
| Large proline-rich protein BAG6 | P46379-2;P46379;P46379-3; | BAG6 | Other | -1.219233333 | 0.667849122 | 25.7345 | 25.9545 | 26.23 | 23.4422 | 25.0486 | 25.7705 |
| Brain-specific angiogenesis inhibitor 1-associated protein | Q9UQB8-3;Q9UQB8-5;Q9UQ | BAIAP2 | Extracellular or secreted | -0.049266667 | 0.069629906 | 26.73 | 26.3756 | 26.7322 | 26.1434 | 26.7189 | 26.8277 |
| Brain-specific angiogenesis inhibitor 1-associated protein | Q9UHR4 | BAIAP2L1 | Extracellular or secreted | 0.337566667 | 0.479451752 | 25.0064 | 24.2428 | 24.219 | 25.0945 | 24.6352 | 24.7512 |
| Barrier-to-autointegration factor;Barrier-to-autointegrat | O75531 | BANF1 | Extracellular or secreted | 2.143966667 | 1.130551495 | 28.1567 | 28.4014 | 28.5201 | 31.7669 | 29.9993 | 29.7439 |
| Chromatin complexes subunit BAP18 | Q8IXM2;Q8IXM2-3;Q8IXM2- | BAP18 | Other | -1.7894 | 0.52935422 | 25.7107 | 25.554 | 26.184 | 21.4864 | 25.5463 | 25.0478 |
| Brain acid soluble protein 1 | P80723;P80723-2 | BASP1 | Extracellular or secreted | 1.6272 | 0.444532631 | 25.309 | 22.6814 | 26.9929 | 28.4334 | 25.9754 | 25.4561 |
| Apoptosis regulator BAX | Q07812-5;Q07812-8;Q07812 | BAX | Extracellular or secreted | 0.0526 | 0.045848208 | 26.1184 | 25.7991 | 26.7642 | 25.7991 | 26.3046 | 26.7358 |
| Bromodomain adjacent to zinc finger domain protein 1A | Q9NRL2-2;Q9NRL2 | BAZ1A | Other | -1.380033333 | 0.211030655 | 28.1145 | 21.1143 | 21.2002 | 21.4197 | 23.0636 | 21.8056 |
| Tyrosine-protein kinase BAZ1B | Q9UIG0-2;Q9UIG0 | BAZ1B | Other | -1.3643 | 0.571941604 | 28.6257 | 27.6573 | 25.5955 | 25.8667 | 25.1649 | 26.754 |
| B-cell receptor-associated protein 31 | P51572;P51572-2 | BCAP31 | Extracellular or secreted | 0.798266667 | 0.69313085 | 25.7656 | 25.3355 | 26.8115 | 26.4442 | 26.7205 | 27.1427 |
| Pre-mRNA-splicing factor SPF27 | O75934 | BCAS2 | Other | -0.919866667 | 0.411081719 | 23.3772 | 23.9564 | 24.7162 | 21.4715 | 23.7374 | 24.0813 |
| Branched-chain-amino-acid aminotransferase, cytosolic | P54687-4;P54687;P54687-5; | BCAT1 | Other | 0.252433333 | 0.144556121 | 30.8662 | 30.5646 | 29.6336 | 31.4893 | 30.6612 | 29.6712 |
| Branched-chain-amino-acid aminotransferase, mitochon | O15382;O15382-2 | BCAT2 | Other | -1.484266667 | 0.948057772 | 29.2259 | 28.7309 | 29.2093 | 28.4667 | 27.7398 | 26.5068 |
| BRCA2 and CDKN1A-interacting protein | Q9P287;Q9P287-4;Q9P287-3 | BCCIP | Other | -0.9521 | 1.853624346 | 28.3493 | 28.1113 | 28.404 | 27.0501 | 27.3342 | 27.624 |
| Bcl-2-like protein 13 | Q9BXK5;Q9BXK5-2 | BCL2L13 | Other | -1.475666667 | 0.555132771 | 25.6792 | 25.8197 | 26.6264 | 22.5476 | 25.591 | 25.5597 |
| Bcl-2-associated transcription factor 1 | Q9NYF8-2;Q9NYF8;Q9NYF8- | BCLAF1 | Other | -0.022266667 | 0.014706221 | 26.6165 | 25.538 | 26.8572 | 26.8798 | 26.1402 | 25.9249 |
| Biglycan | P21810 | BGN | Extracellular or secreted | 1.003466667 | 0.55408428 | 26.1569 | 27.1576 | 24.8489 | 26.4295 | 26.986 | 27.7583 |
| S-methylmethionine--homocysteine S-methyltransferase | Q9H2M3-2;Q9H2M3;Q9308 | BHMT2;BHMT | Extracellular or secreted | -0.151833333 | 0.522539245 | 26.1254 | 25.7682 | 26.0317 | 25.7061 | 25.9067 | 25.857 |
| Protein bicaudal D homolog 2 | Q8TD16;Q8TD16-2 | BICD2 | Extracellular or secreted | 1.690166667 | 0.488788611 | 23.2691 | 20.4896 | 24.9738 | 24.2209 | 24.7005 | 24.8816 |
| BH3-interacting domain death agonist;BH3-interacting do | P55957;P55957-2 | BID | Extracellular or secreted | -0.1635 | 0.318536283 | 26.0435 | 25.4823 | 25.6869 | 25.3147 | 25.6915 | 25.716 |
| Myc box-dependent-interacting protein 1 | O00499-9;O00499-10;O0049 | BIN1 | Extracellular or secreted | -1.097433333 | 1.503461458 | 25.2721 | 25.829 | 25.0051 | 24.0719 | 24.3505 | 24.3915 |
| Baculoviral IAP repeat-containing protein 6 | Q9NR09 | BIRC6 | Other | -0.747866667 | 0.38399657 | 23.3416 | 23.939 | 23.3465 | 22.7572 | 21.5568 | 24.0695 |
| Bleomycin hydrolase | Q13867 | BLMH | Extracellular or secreted | 0.329933333 | 0.19198213 | 30.2284 | 29.6727 | 29.0854 | 31.0534 | 29.752 | 29.1709 |
| Biliverdin reductase A | P53004 | BLVRA | Extracellular or secreted | -0.928366667 | 1.836774654 | 27.7796 | 28.1046 | 27.5473 | 26.5785 | 26.9683 | 27.0996 |
| Flavin reductase (NADPH) | P30043 | BLVRB | Extracellular or secreted | -0.337266667 | 0.282349886 | 28.6267 | 28.5476 | 29.4044 | 27.804 | 29.1435 | 28.6194 |
| Bone morphogenetic protein 1 | P13497;P13497-5;P13497-2; | BMP1 | Extracellular or secreted | 0.576966667 | 0.335825108 | 28.1179 | 28.6668 | 26.6047 | 27.835 | 28.4898 | 28.7955 |
| Bone morphogenetic protein 2 | P12643 | BMP2 | Extracellular or secreted | 2.739366667 | 0.861898578 | 21.9269 | 21.3399 | 20.4216 | 21.6066 | 25.4808 | 24.8191 |
| Bone morphogenetic protein 5 | P22003-2;P22003 | BMP5 | Extracellular or secreted | -0.616533333 | 0.237137889 | 22.7032 | 23.8264 | 20.6352 | 22.0904 | 21.7135 | 21.5113 |
| Ribosome biogenesis protein BMS1 homolog | Q14692 | BMS1 | Other | -2.329833333 | 2.249528724 | 24.1513 | 23.3056 | 24.35 | 21.163 | 21.8742 | 21.7802 |
| Biorientation of chromosomes in cell division protein 1-li | Q8NFC6 | BOD1L1 | Other | -0.952366667 | 2.025690166 | 25.7187 | 25.3007 | 25.7648 | 24.5457 | 24.8141 | 24.5673 |
| BolA-like protein 2 | Q9H3K6;Q9H3K6-2 | BOLA2 | Extracellular or secreted | 0.235433333 | 0.274295999 | 25.0173 | 25.8488 | 26.0368 | 25.8977 | 25.93 | 25.7815 |
| Ribosome biogenesis protein BOP1 | Q14137;Q14137-2 | BOP1 | Other | -0.534266667 | 0.140450921 | 23.9409 | 20.6729 | 24.4436 | 23.7239 | 22.5287 | 21.202 |
| Bisphosphoglycerate mutase | P07738 | BPGM | Extracellular or secreted | 0.589533333 | 0.256893189 | 21.8044 | 22.4845 | 21.2136 | 22.4724 | 23.7823 | 21.0164 |
| 3(2),5-bisphosphate nucleotidase 1 | O95861;O95861-2;O95861-4 | BPNT1 | Extracellular or secreted | 0.093266667 | 0.367860879 | 29.9328 | 29.6213 | 29.7064 | 29.8181 | 29.9194 | 29.8028 |
| Nucleosome-remodeling factor subunit BPTF | Q12830-4;Q12830-2;Q12830 | BPTF | Extracellular or secreted | 2.005933333 | 1.168381071 | 24.7439 | 23.8571 | 26.0553 | 27.5819 | 26.6428 | 26.4494 |
| Lys-63-specific deubiquitinase BRCC36 | P46736-2;P46736;P46736-3; | BRCC3 | Other | -2.700566667 | 1.201408438 | 25.1155 | 25.4216 | 25.609 | 21.7279 | 22.1753 | 24.1412 |
| Bromodomain-containing protein 2 | P25440-4;P25440-3;P25440; | BRD2 | Other | -2.513466667 | 0.774621402 | 25.2343 | 24.7906 | 25.5778 | 20.7936 | 24.921 | 22.3477 |
| Bromodomain-containing protein 4 | O60885;O60885-2;O60885-3 | BRD4 | Other | -0.841866667 | 0.541602968 | 26.1495 | 24.7682 | 26.33 | 24.3255 | 24.5429 | 25.8537 |
| BRCA1-A complex subunit BRE | Q9NXR7;Q9NXR7-4;Q9NXR7 | BRE | Other | -3.957733333 | 1.382348236 | 25.6567 | 25.3392 | 25.3019 | 23.1333 | 20.9989 | 20.2924 |
| Protein BRICK1 | Q8WUW1;Q8WUW1-2 | BRK1 | Extracellular or secreted | 0.438266667 | 0.389939296 | 25.0656 | 25.97 | 24.677 | 26.021 | 25.8589 | 25.1475 |
| BRO1 domain-containing protein BROX | Q5VW32;Q5VW32-2 | BROX | Extracellular or secreted | -0.692233333 | 0.484816079 | 27.7505 | 28.6941 | 28.7004 | 26.6767 | 28.036 | 28.3556 |
| BSD domain-containing protein 1 | Q9NW68-8;Q9NW68-2;Q9N | BSDC1 | Other | 0.622466667 | 0.609696651 | 23.5112 | 22.5773 | 23.0626 | 23.1686 | 24.3755 | 23.4744 |
| Basigin | P35613-2;P35613;P35613-4 | BSG | Extracellular or secreted | 1.1936 | 1.21758128 | 30.694 | 29.838 | 30.3704 | 31.9881 | 31.706 | 30.7891 |
| Basigin | P35613-3 | BSG | Extracellular or secreted | 0.2914 | 0.234402791 | 26.8055 | 26.3979 | 26.5806 | 27.5461 | 27.068 | 26.0441 |
| Bone marrow stromal antigen 2 | Q10589-2;Q10589 | BST2 | Extracellular or secreted | 1.8302 | 1.544232902 | 25.6746 | 24.9022 | 24.6741 | 27.2399 | 27.4247 | 26.0769 |
| Biotinidase | P43251-4;P43251;P43251-3; | BTD | Extracellular or secreted | 1.751366667 | 0.734741647 | 27.3917 | 27.5452 | 24.7471 | 29.0096 | 28.3432 | 27.5853 |
| Transcription factor BTF3 | P20290-2;P20290 | BTF3 | Other | -1.1683 | 1.002948455 | 26.8251 | 28.2998 | 27.8888 | 26.1897 | 26.9411 | 26.378 |
| Transcription factor BTF3 homolog 4 | Q96K17;Q96K17-2 | BTF3L4 | Other | -0.220366667 | 0.227353911 | 27.7736 | 28.0614 | 28.7583 | 27.4986 | 28.191 | 28.2426 |
| Mitotic checkpoint protein BUB3 | O43684-2;O43684 | BUB3 | Other | -0.342866667 | 0.452537925 | 28.6022 | 28.852 | 28.9038 | 27.97 | 28.4054 | 28.954 |
| BUD13 homolog | Q9BRD0 | BUD13 | Other | -0.370533333 | 0.07940819 | 20.2044 | 24.0547 | 25.1509 | 23.4919 | 21.8141 | 22.9924 |
| Protein BUD31 homolog | P41223;P41223-2 | BUD31 | Other | -0.124633333 | 0.100102311 | 21.099 | 22.1299 | 21.3899 | 20.7659 | 21.758 | 21.721 |
| Basic leucine zipper and W2 domain-containing protein 1 | Q7L1Q6;Q7L1Q6-4;Q7L1Q6- | BZW1 | Other | -1.311433333 | 0.84258323 | 28.2229 | 28.6777 | 28.4369 | 26.0265 | 27.4282 | 27.9485 |
| Basic leucine zipper and W2 domain-containing protein 2 | Q9Y6E2 | BZW2 | Other | -0.7075 | 0.94300202 | 26.3451 | 26.7761 | 27.2968 | 25.9464 | 26.1257 | 26.2234 |
| UPF0693 protein C10orf32 | Q96B45 | C10orf32 | Other | 0.309466667 | 0.469748799 | 22.7889 | 22.116 | 22.1829 | 22.5596 | 22.4182 | 23.0384 |
| Ester hydrolase C11orf54 | Q9H0W9-2;Q9H0W9;Q9H0W | C11orf54 | Extracellular or secreted | 0.4807 | 0.406625492 | 28.3877 | 28.0536 | 27.3495 | 29.0934 | 28.407 | 27.7325 |
| UPF0696 protein C11orf68 | Q9H3H3;Q9H3H3-2;Q9H3H3 | C11orf68 | Other | -1.9962 | 0.847353004 | 25.2374 | 24.9393 | 25.4624 | 21.5918 | 23.5422 | 24.5165 |

| | | | | | | | | | | | |
|---|---|---|---|---|---|---|---|---|---|---|---|
| Protein Hikeshi | Q53FT3 | C11orf73 | Extracellular or secreted | -0.0322 | 0.054821406 | 25.6687 | 25.2695 | 25.0159 | 25.326 | 25.2533 | 25.2782 |
| UPF0160 protein MYG1, mitochondrial | Q9HB07 | C12orf10 | Extracellular or secreted | 0.2286 | 0.51838057 | 25.9755 | 25.6718 | 26.2504 | 26.2757 | 26.1538 | 26.154 |
| Uncharacterized protein C12orf43 | Q96C57 | C12orf43 | Other | -1.332133333 | 0.469535088 | 23.6232 | 20.2837 | 23.3572 | 20.6587 | 21.3006 | 21.3084 |
| Protein C10 | Q99622 | C12orf57 | Other | 3.497733333 | 1.533578764 | 20.3127 | 22.5968 | 22.5879 | 25.217 | 25.9492 | 24.8244 |
| Uncharacterized protein C14orf142 | Q9BXV9 | C14orf142 | Other | 0.2585 | 0.089253042 | 21.0088 | 22.7559 | 23.575 | 23.4664 | 23.3384 | 21.3104 |
| UPF0568 protein C14orf166 | Q9Y224 | C14orf166 | Other | -1.2693 | 1.160348154 | 28.1019 | 29.0244 | 28.4901 | 26.4788 | 27.498 | 27.8317 |
| Uncharacterized protein C15orf52 | Q6ZUT6-4;Q6ZUT6;Q6ZUT6- | C15orf52 | Other | 0.5154 | 0.616011541 | 24.9457 | 24.5565 | 25.644 | 25.5912 | 25.7182 | 25.383 |
| Protein Njmu-R1 | Q9HAS0 | C17orf75 | Extracellular or secreted | -1.420766667 | 0.877980304 | 22.7627 | 22.6302 | 22.6278 | 22.407 | 20.7352 | 20.6162 |
| Uncharacterized protein C19orf43 | Q9BQ61 | C19orf43 | Other | 0.053233333 | 0.068676727 | 25.6933 | 25.714 | 26.3742 | 25.7611 | 25.9176 | 26.2625 |
| UPF0587 protein C1orf123 | Q9NWV4 | C1orf123 | Extracellular or secreted | 0.0404 | 0.038193025 | 28.4568 | 28.8483 | 28.4435 | 29.0217 | 28.8619 | 27.9862 |
| Uncharacterized protein C1orf50 | Q9BV19 | C1orf50 | Other | 1.733933333 | 1.677844446 | 22.2154 | 22.0574 | 21.1127 | 24.088 | 23.5091 | 22.9902 |
| Complement component 1 Q subcomponent-binding pro | Q07021 | C1QBP | Extracellular or secreted | -0.7387 | 0.535432726 | 32.1787 | 31.913 | 32.3782 | 32.126 | 31.7312 | 30.3966 |
| Complement C1r subcomponent;Complement C1r subco | P00736 | C1R | Extracellular or secreted | 0.594066667 | 0.327435607 | 30.9734 | 31.6423 | 29.549 | 30.5817 | 31.9374 | 31.4278 |
| Complement C1r subcomponent-like protein | Q9NZP8 | C1RL | Extracellular or secreted | 1.837333333 | 1.133680742 | 23.3901 | 23.6706 | 21.7313 | 24.4487 | 25.4316 | 24.4237 |
| Complement C1s subcomponent;Complement C1s subco | P09871 | C1S | Extracellular or secreted | 0.9867 | 0.842464597 | 30.0565 | 30.7018 | 29.1846 | 30.4574 | 31.4157 | 31.0299 |
| UPF0687 protein C20orf27 | Q9GZN8;Q9GZN8-2 | C20orf27 | Other | -1.4381 | 1.466244826 | 25.4595 | 24.2928 | 25.1964 | 23.0836 | 23.7713 | 23.7795 |
| UPF0769 protein C21orf59 | P57076 | C21orf59 | Other | -0.093666667 | 0.040876501 | 23.128 | 22.4653 | 21.7185 | 21.1772 | 22.4139 | 23.4397 |
| C2 domain-containing protein 5 | Q86YS7;Q86YS7-2;Q86YS7-3 | C2CD5 | Extracellular or secreted | 1.491933333 | 0.644101259 | 19.4732 | 21.1333 | 22.4901 | 22.263 | 21.8805 | 23.4289 |
| Complement C3;Complement C3 beta chain;C3-beta-c;C | P01024 | C3 | Extracellular or secreted | 0.876566667 | 0.563010039 | 36.4091 | 37.2382 | 35.2065 | 36.7276 | 37.5499 | 37.206 |
| UPF0609 protein C4orf27 | Q9NWY4 | C4orf27 | Other | -0.9003 | 1.097138521 | 25.8733 | 26.8061 | 25.9207 | 25.632 | 25.3759 | 24.8913 |
| UPF0489 protein C5orf22 | Q49AR2;Q49AR2-2 | C5orf22 | Other | 0.014166667 | 0.008848956 | 24.1144 | 23.1396 | 22.3999 | 23.2359 | 23.2509 | 23.2096 |
| Putative uncharacterized protein C5orf58 | C9J3I9 | C5orf58 | Other | -4.9463 | 2.139267081 | 28.2125 | 28.2289 | 27.1692 | 21.8309 | 22.79 | 24.1508 |
| Uncharacterized protein C6orf203 | Q9P0P8 | C6orf203 | Other | -1.902766667 | 0.694896218 | 25.7674 | 25.1024 | 25.7562 | 21.6249 | 25.0187 | 24.2741 |
| Protein PAXX | Q9BUH6;Q9BUH6-2 | C9orf142 | Extracellular or secreted | -0.709333333 | 0.327493927 | 23.0243 | 22.8526 | 22.8778 | 23.1214 | 22.8975 | 20.6078 |
| UPF0553 protein C9orf64 | Q5T6V5 | C9orf64 | Other | -1.002933333 | 0.805256083 | 28.5853 | 28.7061 | 27.3559 | 27.5822 | 27.6 | 26.4563 |
| Carbonic anhydrase 12 | O43570-2;O43570 | CA12 | Extracellular or secreted | -0.0529 | 0.087349938 | 24.6057 | 23.9362 | 24.2046 | 24.0548 | 24.2848 | 24.2482 |
| Carbonic anhydrase 13 | Q8N1Q1 | CA13 | Other | 0.331066667 | 0.327146745 | 24.6442 | 24.3985 | 23.6107 | 24.7594 | 24.883 | 24.0042 |
| Carbonic anhydrase 2 | P00918 | CA2 | Extracellular or secreted | 0.2665 | 0.160934912 | 30.9666 | 30.382 | 29.5635 | 31.437 | 30.449 | 29.8256 |
| Carbonic anhydrase 9 | Q16790 | CA9 | Extracellular or secreted | 0.046166667 | 0.054974211 | 26.5527 | 25.6654 | 26.2394 | 25.9919 | 26.2582 | 26.3459 |
| Calcium-binding protein 39 | Q9Y376 | CAB39 | Extracellular or secreted | -0.787233333 | 0.881745797 | 27.1257 | 27.6675 | 26.8974 | 25.785 | 26.846 | 26.6979 |
| Voltage-dependent R-type calcium channel subunit alpha | Q15878-2;Q15878-3;Q15878 | CACNA1E | Extracellular or secreted | 1.748133333 | 1.912600238 | 27.4565 | 27.2795 | 26.4703 | 29.3087 | 28.6429 | 28.4991 |
| Calcyclin-binding protein | Q9HB71;Q9HB71-3 | CACYBP | Extracellular or secreted | -0.5238 | 1.172732247 | 30.6244 | 30.1215 | 30.32 | 29.5934 | 29.7907 | 30.1104 |
| CAD protein;Glutamine-dependent carbamoyl-phosphat | P27708 | CAD | Extracellular or secreted | -0.361233333 | 0.300063254 | 28.866 | 28.6743 | 29.8988 | 28.6293 | 28.3543 | 29.3718 |
| Cell adhesion molecule 4 | Q8NFZ8 | CADM4 | Extracellular or secreted | 0.164833333 | 0.120168893 | 24.7711 | 24.0068 | 23.7718 | 24.7871 | 24.6994 | 23.5577 |
| Calretinin | P22676 | CALB2 | Other | 3.303166667 | 1.187129265 | 22.3148 | 19.2579 | 21.362 | 24.4903 | 24.2691 | 24.0848 |
| Caldesmon | Q05682-5;Q05682-4;Q05682 | CALD1 | Extracellular or secreted | 0.6794 | 1.884684725 | 28.001 | 27.9311 | 27.9463 | 28.6827 | 28.7665 | 28.4674 |
| Calmodulin-like protein 5 | Q9NZT1 | CALML5 | Extracellular or secreted | 1.857 | 0.822582823 | 22.0869 | 22.0719 | 24.6559 | 24.8104 | 24.0379 | 25.5374 |
| Calreticulin | P27797 | CALR | Extracellular or secreted | 0.185033333 | 0.09117087 | 34.8159 | 34.2758 | 34.7574 | 36.0968 | 34.4277 | 33.8797 |
| Calreticulin-3 | Q96L12 | CALR3 | Extracellular or secreted | -1.745466667 | 0.665613992 | 24.4887 | 23.7215 | 25.4531 | 20.9832 | 23.0442 | 24.3995 |
| Calumenin | O43852;O43852-3;O43852-6 | CALU | Extracellular or secreted | 1.621933333 | 0.741930933 | 30.343 | 29.1938 | 31.3296 | 33.3951 | 31.5679 | 30.7692 |
| Calumenin | O43852-2;O43852-4;O43852 | CALU | Extracellular or secreted | 1.721266667 | 0.400329541 | 25.0601 | 20.8112 | 25.832 | 27.1247 | 25.2132 | 24.5292 |
| Calcium/calmodulin-dependent protein kinase type II sub | Q13555-10;Q13555-3;Q1355 | CAMK2G | Extracellular or secreted | -0.128933333 | 0.034924908 | 20.7385 | 22.777 | 24.6146 | 22.9032 | 23.1715 | 21.6686 |
| Cullin-associated NEDD8-dissociated protein 1 | Q86VP6;Q86VP6-2 | CAND1 | Extracellular or secreted | -0.368966667 | 0.350764547 | 30.0154 | 30.6449 | 30.8151 | 29.4644 | 30.2193 | 30.6848 |
| Soluble calcium-activated nucleotidase 1 | Q8WVQ1-3;Q8WVQ1;Q8WV | CANT1 | Extracellular or secreted | 1.257433333 | 0.347570618 | 27.0937 | 28.4715 | 23.9258 | 28.027 | 27.8042 | 27.4321 |
| Calnexin | P27824;P27824-2;P27824-3 | CANX | Extracellular or secreted | -0.224766667 | 0.130539555 | 30.3485 | 29.0262 | 30.9603 | 30.2687 | 29.8478 | 29.5442 |
| Adenylyl cyclase-associated protein 1 | Q01518 | CAP1 | Extracellular or secreted | 0.0765 | 0.09294769 | 31.6928 | 32.2613 | 31.696 | 31.5285 | 32.0704 | 32.2807 |
| Adenylyl cyclase-associated protein 1 | Q01518-2 | CAP1 | Extracellular or secreted | -1.1785 | 0.799473701 | 23.7791 | 23.801 | 22.366 | 21.2077 | 22.3406 | 22.8623 |
| Adenylyl cyclase-associated protein 2 | P40123-2;P40123 | CAP2 | Extracellular or secreted | 0.368266667 | 0.337146493 | 24.4097 | 23.3115 | 24.3377 | 23.8838 | 24.4543 | 24.8256 |
| Macrophage-capping protein | P40121;P40121-2 | CAPG | Extracellular or secreted | -0.641866667 | 0.547803666 | 30.4644 | 30.7165 | 31.1306 | 29.249 | 30.4949 | 30.642 |
| Calpain-1 catalytic subunit | P07384 | CAPN1 | Extracellular or secreted | -0.316533333 | 0.942052244 | 29.1755 | 29.2879 | 29.1105 | 28.6456 | 28.898 | 29.0807 |
| Calpain-2 catalytic subunit | P17655;P17655-2 | CAPN2 | Extracellular or secreted | -0.139866667 | 0.108906147 | 29.0209 | 29.471 | 29.9086 | 28.6775 | 29.304 | 29.9994 |
| Calpain small subunit 1 | P04632 | CAPNS1 | Extracellular or secreted | -0.405066667 | 0.458951198 | 28.7972 | 29.4083 | 29.1347 | 28.0702 | 29.0951 | 28.9597 |
| Caprin-1 | Q14444-2;Q14444 | CAPRIN1 | Extracellular or secreted | 0.050333333 | 0.048969995 | 30.4159 | 29.6987 | 29.3528 | 29.8648 | 29.6339 | 30.1197 |
| Calcyphosin | Q13938-4;Q13938;Q13938-3 | CAPS | Extracellular or secreted | -0.391133333 | 0.160159864 | 21.2623 | 23.5017 | 22.7234 | 22.279 | 20.9196 | 23.1154 |
| F-actin-capping protein subunit alpha-1 | P52907 | CAPZA1 | Extracellular or secreted | -0.390666667 | 0.468949382 | 30.8515 | 31.2006 | 31.0908 | 30.0301 | 30.9473 | 30.9935 |
| F-actin-capping protein subunit alpha-2 | P47755 | CAPZA2 | Extracellular or secreted | -0.5378 | 0.53875669 | 28.56 | 29.0546 | 28.929 | 27.5792 | 28.5113 | 28.8397 |
| F-actin-capping protein subunit beta | P47756-2;P47756 | CAPZB | Extracellular or secreted | -0.533433333 | 0.699672813 | 31.2414 | 31.3704 | 31.4082 | 30.2464 | 30.9906 | 31.1827 |
| Calcium-regulated heat stable protein 1 | Q9Y2V2 | CARHSP1 | Extracellular or secreted | -0.0667 | 0.046106527 | 25.7849 | 25.5785 | 26.7861 | 25.4108 | 26.5335 | 26.0051 |
| ATP-dependent (S)-NAD(P)H-hydrate dehydratase | Q8IW45;Q8IW45-2;Q8IW45- | CARKD | Other | -0.263466667 | 0.695107177 | 24.5089 | 24.9766 | 24.8965 | 24.4266 | 24.6219 | 24.5431 |
| Histone-arginine methyltransferase CARM1 | Q86X55-2;Q86X55-1;Q86X55 | CARM1 | Extracellular or secreted | -1.257233333 | 1.367511357 | 26.4888 | 26.3748 | 27.3472 | 24.9671 | 25.4653 | 26.0067 |
| Cysteine--tRNA ligase, cytoplasmic | P49589-3;P49589;P49589-2 | CARS | Other | -0.145266667 | 0.239079049 | 28.2518 | 28.2017 | 28.2164 | 27.6421 | 28.2488 | 28.3432 |
| Probable cysteine--tRNA ligase, mitochondrial | Q9HA77 | CARS2 | Other | -0.7691 | 0.232101758 | 22.2992 | 24.7992 | 25.4196 | 24.1651 | 21.6454 | 24.3938 |
| Protein CASC5 | Q8NG31-3;Q8NG31-4;Q8NG | CASC5 | Extracellular or secreted | 0.0336 | 0.014931584 | 21.3511 | 23.6416 | 22.5496 | 22.7879 | 22.9026 | 21.9526 |
| Peripheral plasma membrane protein CASK | O14936-3;O14936-4;O14936 | CASK | Extracellular or secreted | 0.865533333 | 1.307791538 | 25.7 | 25.4253 | 26.2497 | 26.4188 | 26.652 | 26.9008 |
| Caspase-14;Caspase-14 subunit p17, mature form;Caspa | P31944 | CASP14 | Extracellular or secreted | 0.832133333 | 0.955570147 | 28.8468 | 28.9226 | 29.8482 | 29.9244 | 29.9331 | 30.2565 |
| Caspase-3;Caspase-3 subunit p17;Caspase-3 subunit p12 | P42574 | CASP3 | Extracellular or secreted | 0.206766667 | 0.425274692 | 28.218 | 28.0878 | 27.6673 | 28.4379 | 28.0895 | 28.066 |
| Caspase-6;Caspase-6 subunit p18;Caspase-6 subunit p11 | P55212 | CASP6 | Other | -0.419466667 | 0.325508178 | 22.6169 | 23.5255 | 23.5438 | 22.7314 | 22.1142 | 23.5822 |
| Calpastatin | P20810-9;P20810-10;P20810 | CAST | Other | 1.1415 | 1.108399935 | 29.4564 | 28.6687 | 29.3663 | 31.0215 | 30.1905 | 29.7039 |
| Catalase | P04040 | CAT | Extracellular or secreted | -0.044433333 | 0.03067181 | 30.865 | 30.141 | 31.1366 | 31.3087 | 30.7211 | 29.9795 |
| Caveolin-1 | Q03135;Q03135-2 | CAV1 | Extracellular or secreted | 1.633433333 | 1.082040174 | 24.6507 | 24.2167 | 24.7185 | 27.0763 | 26.1899 | 25.22 |
| Core-binding factor subunit beta | Q13951-2;Q13951 | CBFB | Other | -0.5317 | 0.79312163 | 27.2227 | 27.6041 | 27.598 | 26.4558 | 27.056 | 27.3179 |
| Carbonyl reductase [NADPH] 1 | P16152;P16152-2 | CBR1 | Extracellular or secreted | -0.2528 | 0.268617886 | 29.0912 | 30.0234 | 29.8606 | 28.9542 | 29.7807 | 29.4819 |
| Carbonyl reductase [NADPH] 3 | O75828 | CBR3 | Extracellular or secreted | -0.568 | 0.586817156 | 25.8626 | 26.4453 | 27.0079 | 25.5403 | 26.4153 | 25.6562 |
| Carbonyl reductase family member 4 | Q8N4T8-2;Q8N4T8 | CBR4 | Other | -1.724833333 | 1.148310233 | 24.4929 | 25.6588 | 26.197 | 22.8397 | 24.5551 | 23.7794 |
| Chromobox protein homolog 1 | P83916 | CBX1 | Other | 0.1049 | 0.138542621 | 26.1704 | 26.0584 | 26.8088 | 26.4024 | 26.7268 | 26.2231 |
| Chromobox protein homolog 3 | Q13185 | CBX3 | Other | -0.223 | 0.744382629 | 30.1587 | 29.8862 | 30.2031 | 29.8945 | 30.0045 | 29.68 |
| Chromobox protein homolog 5 | P45973 | CBX5 | Other | -0.144766667 | 0.138505382 | 26.6656 | 25.6699 | 26.7856 | 26.0614 | 26.2188 | 26.4066 |
| Coiled-coil and C2 domain-containing protein 1A | Q6P1N0-2;Q6P1N0 | CC2D1A | Extracellular or secreted | -0.6648 | 0.798376788 | 22.7181 | 22.3807 | 22.5611 | 22.2185 | 22.1848 | 21.2622 |
| Cell division cycle and apoptosis regulator protein 1 | Q8IX12-2;Q8IX12 | CCAR1 | Other | -1.935766667 | 0.707783919 | 27.8984 | 27.9737 | 29.1607 | 24.4853 | 26.6753 | 28.0649 |
| Cell cycle and apoptosis regulator protein 2 | Q8N163-2;Q8N163 | CCAR2 | Other | -1.715833333 | 1.076733297 | 28.8049 | 28.6843 | 30.1183 | 26.4367 | 27.5976 | 28.4257 |
| Kynurenine--oxoglutarate transaminase 1 | Q16773-2;Q16773 | CCBL1 | Other | -0.856066667 | 0.763318017 | 26.8376 | 26.4722 | 25.634 | 26.0283 | 25.5906 | 24.7567 |
| Kynurenine--oxoglutarate transaminase 3 | Q6YP21-3;Q6YP21 | CCBL2 | Other | 0.5164 | 0.624951009 | 28.6521 | 28.3757 | 28.225 | 29.5077 | 28.8701 | 28.4242 |
| Coiled-coil domain-containing protein 12 | Q8WUD4 | CCDC12 | Other | 0.646466667 | 0.357875289 | 23.4839 | 21.6617 | 23.6896 | 23.5213 | 24.2016 | 23.0517 |
| Coiled-coil domain-containing protein 124 | Q96CT7 | CCDC124 | Extracellular or secreted | 2.913866667 | 1.483888587 | 21.0428 | 22.8223 | 23.1386 | 24.1521 | 26.3475 | 25.2457 |
| Coiled-coil domain-containing protein 132 | Q96JG6-2;Q96JG6-3;Q96JG6 | CCDC132 | Extracellular or secreted | 0.750466667 | 0.299238846 | 22.9357 | 19.7905 | 20.9964 | 22.4156 | 21.6197 | 21.9387 |
| Coiled-coil domain-containing protein 25 | Q86WR0;Q86WR0-2 | CCDC25 | Extracellular or secreted | -0.5676 | 1.007537885 | 27.7969 | 27.1327 | 27.6938 | 26.8598 | 26.9583 | 27.1025 |
| Coiled-coil domain-containing protein 43 | Q96MW1-2;Q96MW1 | CCDC43 | Other | 2.492766667 | 1.596152592 | 21.1422 | 20.1753 | 21.9422 | 24.4996 | 22.7721 | 23.4663 |
| Coiled-coil domain-containing protein 58 | Q4VC31 | CCDC58 | Other | -1.3516 | 1.703218972 | 28.9831 | 28.7749 | 29.0843 | 27.6007 | 28.0106 | 27.1762 |
| Coiled-coil domain-containing protein 6 | Q16204 | CCDC6 | Other | 0.5244 | 1.871549761 | 26.1274 | 25.8987 | 26.2103 | 26.4813 | 26.7507 | 26.5776 |
| Coiled-coil domain-containing protein 68 | Q9H2F9 | CCDC68 | Other | 0.920833333 | 0.203779467 | 22.0239 | 22.6997 | 22.0959 | 26.0928 | 22.945 | 20.5442 |
| Coiled-coil domain-containing protein 80 | Q76M96;Q76M96-2 | CCDC80 | Extracellular or secreted | -4.132033333 | 1.521574348 | 27.7946 | 28.6714 | 27.7956 | 22.6669 | 23.6164 | 25.5822 |
| Coiled-coil domain-containing protein 86 | Q9H6F5 | CCDC86 | Other | -2.968366667 | 1.497294912 | 28.6964 | 27.3249 | 26.4859 | 25.7509 | 23.514 | 24.3372 |
| Girdin | Q3V6T2-5;Q3V6T2-4;Q3V6T2 | CCDC88A | Extracellular or secreted | -2.5667 | 0.74685703 | 26.6167 | 25.1512 | 24.8341 | 20.3539 | 24.133 | 24.415 |
| Coiled-coil domain-containing protein 9 | Q9Y3X0 | CCDC9 | Other | -1.436033333 | 0.674130291 | 22.9773 | 22.742 | 22.7205 | 22.9681 | 20.6052 | 20.5584 |
| C-C motif chemokine 20;CCL20(1-67);CCL20(1-64);CCL20 | P78556-2;P78556 | CCL20 | Extracellular or secreted | -0.389033333 | 0.137503878 | 25.2632 | 27.5039 | 24.9272 | 24.5326 | 25.2352 | 26.7594 |
| Cyclin-H | P51946 | CCNH | Other | 0.299233333 | 0.341512417 | 22.9728 | 22.4084 | 23.5369 | 23.2611 | 23.23 | 23.3247 |
| Cyclin-K | O75909-1;O75909;O75909-4 | CCNK | Other | -1.302333333 | 0.448128489 | 23.4769 | 24.1067 | 25.3287 | 20.8945 | 23.7861 | 24.3247 |
| Copper chaperone for superoxide dismutase | O14618 | CCS | Other | -0.298433333 | 0.317591573 | 24.3893 | 24.8573 | 23.9094 | 23.5799 | 24.5039 | 24.1769 |
| T-complex protein 1 subunit beta | P78371;P78371-2 | CCT2 | Extracellular or secreted | -1.093666667 | 1.029766545 | 31.121 | 31.498 | 32.0608 | 29.6937 | 30.7087 | 30.9964 |
| T-complex protein 1 subunit gamma | P49368;P49368-2 | CCT3 | Extracellular or secreted | -0.846166667 | 0.684855607 | 31.337 | 31.5946 | 32.3093 | 30.0325 | 31.0807 | 31.5892 |
| T-complex protein 1 subunit delta | P50991;P50991-2 | CCT4 | Extracellular or secreted | -0.7877 | 0.661285474 | 30.6323 | 31.2147 | 31.7392 | 29.6129 | 30.5429 | 31.0673 |
| T-complex protein 1 subunit epsilon | P48643;P48643-2 | CCT5 | Extracellular or secreted | -0.967166667 | 0.968689605 | 31.3376 | 31.6647 | 32.3066 | 30.1685 | 30.8166 | 31.4223 |
| T-complex protein 1 subunit zeta | P40227;P40227-2 | CCT6A | Extracellular or secreted | -0.7102 | 0.627103535 | 30.522 | 31.1528 | 31.6078 | 29.709 | 30.365 | 31.078 |
| T-complex protein 1 subunit zeta-2 | Q92526;Q92526-2;Q92526-3 | CCT6B | Other | 2.4052 | 0.398265153 | 21.0904 | 27.0013 | 20.5512 | 26.2888 | 22.4444 | 27.1253 |
| T-complex protein 1 subunit eta | Q99832;Q99832-3;Q99832-4 | CCT7 | Extracellular or secreted | -0.7394 | 0.621529773 | 30.5258 | 30.974 | 31.7003 | 29.5924 | 30.3895 | 31 |
| T-complex protein 1 subunit theta | P50990;P50990-2;P50990-3 | CCT8 | Extracellular or secreted | -0.778366667 | 0.817921504 | 31.7132 | 31.975 | 32.4738 | 30.7087 | 31.182 | 31.9362 |
| CD109 antigen | Q6YHK3;Q6YHK3-4;Q6YHK3- | CD109 | Extracellular or secreted | -0.566833333 | 0.4883441 | 29.5885 | 29.7869 | 28.3968 | 28.3257 | 28.9652 | 28.7808 |
| Monocyte differentiation antigen CD14;Monocyte differe | P08571 | CD14 | Extracellular or secreted | 0.828266667 | 0.536095951 | 27.6263 | 28.8929 | 27.2077 | 27.8294 | 29.1958 | 29.1865 |
| CD151 antigen | P48509 | CD151 | Other | 1.8581 | 0.475339625 | 26.8034 | 26.7689 | 22.3612 | 27.0758 | 27.3867 | 27.0453 |
| CD276 antigen | Q5ZPR3-3;Q5ZPR3-4;Q5ZPR3 | CD276 | Extracellular or secreted | 0.2414 | 0.236416982 | 28.4993 | 28.0303 | 27.4942 | 28.3151 | 28.6935 | 27.7394 |
| CD2-associated protein | Q9Y5K6 | CD2AP | Extracellular or secreted | 0.1802 | 0.3279712 | 27.4454 | 27.4857 | 27.8871 | 27.4546 | 27.8537 | 28.0505 |
| DNA-directed RNA polymerase I subunit RPA34 | O15446;O15446-2 | CD3EAP | Other | -0.108166667 | 0.037660106 | 24.0368 | 23.1096 | 25.4103 | 25.4882 | 23.4869 | 23.2571 |
| CD44 antigen | P16070-8;P16070-4;P16070- | CD44 | Extracellular or secreted | 2.9415 | 2.267282027 | 30.6405 | 30.3104 | 29.3699 | 33.7316 | 32.9899 | 32.4238 |
| Membrane cofactor protein | P15529-7;P15529-4;P15529- | CD46 | Extracellular or secreted | 0.6082 | 0.630109273 | 27.9091 | 27.1484 | 27.2133 | 28.5672 | 28.1525 | 27.3757 |
| Leukocyte surface antigen CD47 | Q08722-2;Q08722-3;Q08722 | CD47 | Extracellular or secreted | -0.3911 | 0.235974533 | 27.4835 | 26.2173 | 26.7442 | 27.3119 | 26.482 | 25.4778 |
| Complement decay-accelerating factor | P08174-4;P08174;P08174-3 | CD55 | Extracellular or secreted | 0.1352 | 0.125713168 | 29.3668 | 29.2492 | 28.4132 | 29.5668 | 29.1784 | 28.6896 |
| Lymphocyte function-associated antigen 3 | P19256-2;P19256-3;P19256 | CD58 | Extracellular or secreted | 1.339766667 | 1.643937084 | 26.0665 | 25.3852 | 25.1856 | 27.0392 | 27.237 | 26.3804 |
| CD59 glycoprotein | P13987 | CD59 | Extracellular or secreted | 2.406933333 | 1.991034347 | 28.7506 | 29.3085 | 28.1014 | 31.7548 | 31.2089 | 30.4176 |
| CD63 antigen | P08962-3;P08962-2;P08962 | CD63 | Extracellular or secreted | 1.559933333 | 0.371479547 | 25.3723 | 25.1498 | 21.1071 | 26.8419 | 23.3585 | 26.1086 |

| | | | | | | | | | | | |
|---|---|---|---|---|---|---|---|---|---|---|---|
| CD81 antigen | P60033 | CD81 | Extracellular or secreted | 0.190566667 | 0.119783399 | 27.8601 | 28.0254 | 26.8154 | 28.5155 | 27.7592 | 26.9979 |
| CD82 antigen | P27701-2;P27701 | CD82 | Extracellular or secreted | 2.552466667 | 1.89888901 | 26.2652 | 26.6917 | 25.2029 | 29.3506 | 28.3905 | 28.0761 |
| CD9 antigen | P21926 | CD9 | Extracellular or secreted | 1.0919 | 1.625327943 | 29.7474 | 29.5566 | 28.9637 | 30.7814 | 30.5779 | 30.1841 |
| CD97 antigen;CD97 antigen subunit alpha;CD97 antigen | P48960-2;P48960-3;P48960 | CD97 | Extracellular or secreted | 0.961933333 | 0.881461765 | 25.8812 | 25.4825 | 24.5833 | 26.768 | 26.4114 | 25.6534 |
| Cytidine deaminase | P32320 | CDA | Extracellular or secreted | 1.310266667 | 1.248167567 | 28.9052 | 30.1311 | 29.8256 | 30.6603 | 31.1216 | 31.0108 |
| Cell division cycle protein 123 homolog | O75794 | CDC123 | Other | 0.370866667 | 0.344204765 | 22.5567 | 21.5324 | 22.6116 | 23.1328 | 22.4434 | 22.2371 |
| Hsp90 co-chaperone Cdc37;Hsp90 co-chaperone Cdc37, | Q16543 | CDC37 | Extracellular or secreted | -0.113066667 | 0.177449544 | 29.349 | 29.5447 | 29.4938 | 29.032 | 29.2417 | 29.7746 |
| Pre-mRNA-processing factor 17 | O60508 | CDC40 | Extracellular or secreted | -1.278933333 | 0.589274251 | 23.4054 | 23.7461 | 24.722 | 21.1795 | 22.8598 | 23.9974 |
| Cell division control protein 42 homolog | P60953;P60953-1 | CDC42 | Extracellular or secreted | 1.2014 | 1.735622726 | 28.2028 | 27.993 | 28.0132 | 29.0723 | 29.6714 | 29.0695 |
| Serine/threonine-protein kinase MRCK beta | Q9Y5S2 | CDC42BPB | Extracellular or secreted | -0.250233333 | 0.288186742 | 23.8866 | 23.6573 | 23.632 | 23.9989 | 22.907 | 23.5193 |
| Cell division cycle 5-like protein | Q99459 | CDC5L | Other | 0.0453 | 0.0179779 | 23.9986 | 26.4976 | 26.2016 | 25.6034 | 25.7064 | 25.5239 |
| Parafibromin | Q6P1J9 | CDC73 | Other | -1.845 | 1.567463468 | 25.7969 | 25.5688 | 25.6559 | 23.3566 | 23.66 | 24.47 |
| CUB domain-containing protein 1 | Q9H5V8;Q9H5V8-2 | CDCP1 | Extracellular or secreted | 0.998133333 | 0.698477251 | 27.661 | 28.0751 | 26.2824 | 28.5262 | 28.4766 | 28.0101 |
| Cadherin-1;E-Cad/CTF1;E-Cad/CTF2;E-Cad/CTF3 | P12830;P12830-2 | CDH1 | Extracellular or secreted | 0.632166667 | 0.792200226 | 31.844 | 31.7203 | 31.1841 | 32.6989 | 32.2658 | 31.6802 |
| Cadherin-13 | P55290;P55290-4;P55290-5; | CDH13 | Extracellular or secreted | 1.445666667 | 0.977375326 | 25.5594 | 26.652 | 24.7293 | 27.415 | 27.2768 | 26.5859 |
| Cadherin-2 | P19022;P19022-2 | CDH2 | Extracellular or secreted | 4.199066667 | 1.190352696 | 23.9774 | 24.8435 | 20.8444 | 26.8289 | 28.0554 | 27.3782 |
| Cadherin-3 | P22223;P22223-2 | CDH3 | Extracellular or secreted | 0.620766667 | 0.393400316 | 29.0766 | 29.7373 | 27.7602 | 29.8321 | 29.5763 | 29.028 |
| Cyclin-dependent kinase 1 | P06493;P06493-2 | CDK1 | Extracellular or secreted | -2.052633333 | 0.914787593 | 26.0745 | 24.51 | 25.8338 | 21.743 | 24.4939 | 24.0235 |
| Cyclin-dependent kinase 11B;Cyclin-dependent kinase 11 | P21127-5;P21127-10;P21127 | CDK11B;CDK11 | Other | 0.683933333 | 0.799931795 | 23.6729 | 23.5536 | 23.834 | 24.6383 | 24.7385 | 23.7355 |
| Cyclin-dependent kinase 2 | P24941;P24941-2 | CDK2 | Other | -1.391266667 | 0.707087228 | 25.8514 | 25.5801 | 27.0674 | 23.3214 | 25.332 | 25.6717 |
| CDK5 regulatory subunit-associated protein 3 | Q96JB5;Q96JB5-4;Q96JB5-2 | CDK5RAP3 | Other | -0.6132 | 0.137721766 | 23.6953 | 24.4194 | 26.5512 | 21.6524 | 24.8548 | 26.3191 |
| Cyclin-dependent kinase 6 | Q00534 | CDK6 | Other | 0.931066667 | 0.667557467 | 22.0766 | 20.9728 | 22.7905 | 22.3715 | 23.0091 | 23.2525 |
| CDKN2A-interacting protein | Q9NXV6 | CDKN2AIP | Other | -0.363733333 | 0.078608337 | 23.1435 | 21.9424 | 25.8379 | 21.0249 | 23.9641 | 24.8436 |
| CDKN2AIP N-terminal-like protein | Q96HQ2;Q96HQ2-2 | CDKN2AIPNL | Other | -1.3116 | 1.023333427 | 24.4772 | 24.4001 | 24.1837 | 22.1473 | 23.5525 | 23.4264 |
| Corneodesmosin | Q15517 | CDSN | Extracellular or secreted | 0.395 | 0.06408382 | 22.5014 | 21.8068 | 22.9601 | 26.7565 | 21.4685 | 20.2283 |
| Protein CDV3 homolog | Q9UKY7;Q9UKY7-2 | CDV3 | Extracellular or secreted | 0.1419 | 0.104591171 | 28.6741 | 27.8584 | 28.0781 | 29.1511 | 28.1142 | 27.771 |
| CCAAT/enhancer-binding protein beta | P17676-2;P17676;P17676-3 | CEBPB | Other | -0.856 | 0.455254785 | 25.723 | 24.5739 | 26.4275 | 23.6739 | 24.7206 | 25.7619 |
| Cat eye syndrome critical region protein 5 | Q9BXW7-2;Q9BXW7 | CECR5 | Extracellular or secreted | -0.198466667 | 0.042395357 | 20.0863 | 24.0751 | 24.9525 | 23.1577 | 22.7611 | 22.5997 |
| CUGBP Elav-like family member 1 | Q92879-2;Q92879-3;Q92879 | CELF1 | Other | -1.1358 | 0.140911775 | 25.1472 | 20.0369 | 27.1765 | 20.4348 | 21.3747 | 27.1437 |
| Cadherin EGF LAG seven-pass G-type receptor 2 | Q9HCU4 | CELSR2 | Extracellular or secreted | 0.884066667 | 0.402204425 | 24.0878 | 25.75 | 22.9081 | 24.8329 | 25.2586 | 25.3066 |
| Major centromere autoantigen B | P07199 | CENPB | Other | -1.687666667 | 0.991436056 | 21.4113 | 22.5181 | 22.7933 | 20.738 | 19.3696 | 21.5521 |
| Centromere protein F | P49454 | CENPF | Other | -1.0698 | 0.818674329 | 22.925 | 22.7366 | 22.2249 | 21.3961 | 20.7922 | 22.4888 |
| Centrosomal protein of 164 kDa | Q9UPV0-2;Q9UPV0 | CEP164 | Extracellular or secreted | -2.911466667 | 0.623807001 | 27.4168 | 26.3072 | 24.1066 | 21.2581 | 26.5265 | 21.3116 |
| Centrosomal protein of 170 kDa | Q5SW79-2;Q5SW79-3;Q5SW | CEP170 | Other | 1.972366667 | 0.949534787 | 20.1859 | 21.1275 | 22.7029 | 23.3342 | 23.1191 | 23.4801 |
| Centrosomal protein of 63 kDa | Q96MT8-4;Q96MT8-3;Q96M | CEP63 | Other | -4.172533333 | 1.090592398 | 27.0851 | 26.9973 | 27.5388 | 22.6662 | 21.0102 | 25.4272 |
| Liver carboxylesterase 1 | P23141-3;P23141;P23141-2 | CES1 | Extracellular or secreted | -2.3353 | 1.532871375 | 30.3229 | 30.871 | 31.6633 | 29.2237 | 29.0942 | 27.5334 |
| Cilia- and flagella-associated protein 20 | Q9Y6A4 | CFAP20 | Extracellular or secreted | -1.0841 | 0.515121571 | 24.0396 | 24.2116 | 25.6006 | 22.1152 | 23.7944 | 24.6899 |
| Complement factor B;Complement factor B Ba fragment; | P00751;P00751-2 | CFB | Extracellular or secreted | 1.847333333 | 0.75394238 | 32.6816 | 34.23 | 31.0345 | 34.7084 | 34.7965 | 33.9832 |
| Complement factor D | P00746 | CFD | Extracellular or secreted | -1.7751 | 0.580011224 | 21.4998 | 25.4242 | 24.1961 | 21.8639 | 22.8276 | 21.1033 |
| Craniofacial development protein 1 | Q9UEE9-2;Q9UEE9 | CFDP1 | Extracellular or secreted | -0.080033333 | 0.063602227 | 24.5226 | 23.6424 | 24.8184 | 24.0085 | 23.9802 | 24.7546 |
| Complement factor H | P08603;P08603-2 | CFH | Extracellular or secreted | 3.939966667 | 1.37591812 | 27.2309 | 27.9549 | 24.7721 | 29.9185 | 31.091 | 30.7683 |
| Complement factor I;Complement factor I heavy chain;C | P05156 | CFI | Extracellular or secreted | 0.669733333 | 0.289775105 | 21.4415 | 23.2562 | 22.56 | 22.7176 | 22.0021 | 24.5472 |
| Cofilin-1 | P23528 | CFL1 | Extracellular or secreted | -0.2261 | 0.242351522 | 33.495 | 34.0125 | 33.2759 | 32.8494 | 33.8643 | 33.3914 |
| Cofilin-2 | Q9Y281;Q9Y281-3 | CFL2 | Extracellular or secreted | -0.2504 | 0.417631392 | 26.0762 | 26.6732 | 26.0462 | 26.0292 | 26.2643 | 25.7509 |
| Cingulin | Q9P2M7;Q9P2M7-2 | CGN | Other | -2.935833333 | 2.508977915 | 24.5304 | 24.8006 | 25.1294 | 22.0702 | 21.2804 | 22.3023 |
| Chromatin assembly factor 1 subunit B | Q13112 | CHAF1B | Other | -1.210566667 | 1.415621854 | 22.9421 | 23.6568 | 23.8325 | 21.7049 | 22.4244 | 22.6704 |
| Coiled-coil-helix-coiled-coil-helix domain-containing prot | Q9Y6H1;Q5T1J5 | CHCHD2;CHCH | Other | 1.6305 | 0.860108239 | 24.1746 | 23.5392 | 25.9255 | 25.6594 | 26.4048 | 26.4666 |
| Mitochondrial intermembrane space import and assemb | Q8N4Q1;Q8N4Q1-2 | CHCHD4 | Other | -0.7807 | 0.529375302 | 27.9208 | 27.0522 | 28.3494 | 27.9616 | 26.8153 | 26.2034 |
| Coiled-coil-helix-coiled-coil-helix domain-containing prot | Q9BSY4;Q9BSY4-2 | CHCHD5 | Other | 0.613966667 | 0.113672317 | 23.3236 | 22.6631 | 23.3199 | 25.4722 | 25.6153 | 20.061 |
| Chromodomain-helicase-DNA-binding protein 2 | O14647-3;O14647-2;O14647 | CHD2 | Extracellular or secreted | -1.136866667 | 0.720747872 | 23.9975 | 23.2216 | 24.8447 | 23.6378 | 23.1883 | 21.8271 |
| Chromodomain-helicase-DNA-binding protein 3 | Q12873-2;Q12873;Q12873-3 | CHD3 | Other | -1.8611 | 1.150736061 | 24.3937 | 24.228 | 25.0713 | 22.1606 | 22.0647 | 23.8844 |
| Chromodomain-helicase-DNA-binding protein 4 | Q14839;Q14839-2 | CHD4 | Other | -2.9634 | 1.281404616 | 28.0269 | 27.7031 | 29.7175 | 24.2153 | 25.2452 | 27.0968 |
| Calcium homeostasis endoplasmic reticulum protein | Q8IWX8 | CHERP | Other | -0.963466667 | 0.412957776 | 25.131 | 27.0244 | 27.107 | 23.9592 | 26.2661 | 26.1467 |
| Chitinase-3-like protein 1 | P36222 | CHI3L1 | Extracellular or secreted | 10.62146667 | 5.558759331 | 19.8196 | 19.6819 | 19.2013 | 29.9502 | 30.4547 | 30.1623 |
| Chitinase-3-like protein 2 | Q15782-6;Q15782;Q15782-5 | CHI3L2 | Extracellular or secreted | -0.239566667 | 0.163760001 | 28.3862 | 27.2395 | 26.7967 | 27.0826 | 27.7348 | 26.8863 |
| Chitinase domain-containing protein 1 | Q9BWS9-3;Q9BWS9;Q9BWS | CHID1 | Extracellular or secreted | -1.159033333 | 0.593428829 | 24.1395 | 24.4406 | 24.3558 | 21.6966 | 24.0688 | 23.6934 |
| Rab proteins geranylgeranyltransferase component A 1 | P24386 | CHM | Other | -0.3501 | 0.220489524 | 23.154 | 24.0367 | 24.075 | 22.3613 | 23.8112 | 24.0429 |
| Charged multivesicular body protein 1a | Q9HD42 | CHMP1A | Extracellular or secreted | 0.4192 | 0.646587066 | 24.3376 | 25.0823 | 24.8465 | 25.0431 | 25.5534 | 24.9275 |
| Charged multivesicular body protein 3 | Q9Y3E7;Q9Y3E7-4;Q9Y3E7-3 | CHMP3 | Extracellular or secreted | 1.739733333 | 0.476400194 | 24.7537 | 25.5582 | 21.0749 | 24.9517 | 25.9119 | 25.7424 |
| Charged multivesicular body protein 4b | Q9H444 | CHMP4B | Extracellular or secreted | 2.379866667 | 2.210391294 | 24.2845 | 24.6783 | 23.8557 | 26.6417 | 26.7797 | 26.5367 |
| Charged multivesicular body protein 5 | Q9NZZ3 | CHMP5 | Extracellular or secreted | 0.827966667 | 0.310799132 | 22.1344 | 21.8366 | 21.4727 | 23.6177 | 23.6277 | 20.6822 |
| Cysteine and histidine-rich domain-containing protein 1 | Q9UHD1;Q9UHD1-2 | CHORDC1 | Other | -0.3906 | 0.524758814 | 28.5059 | 29.0162 | 29.007 | 27.95 | 28.8743 | 28.533 |
| Calcineurin B homologous protein 1 | Q99653 | CHP1 | Extracellular or secreted | 2.435933333 | 1.273537597 | 21.1977 | 23.1424 | 23.3915 | 24.9551 | 25.9634 | 24.1209 |
| Chromatin accessibility complex protein 1 | Q9NRG0 | CHRAC1 | Other | -1.6163 | 1.063452863 | 22.6637 | 23.1352 | 23.3574 | 20.9892 | 20.7796 | 22.5386 |
| Carbohydrate sulfotransferase 11 | Q9NPF2-2;Q9NPF2 | CHST11 | Extracellular or secreted | 1.7141 | 0.477541703 | 22.5772 | 25.0423 | 20.426 | 23.2463 | 24.5395 | 25.402 |
| Carbohydrate sulfotransferase 14 | Q8NCH0 | CHST14 | Extracellular or secreted | 0.782233333 | 0.21621367 | 23.3051 | 26.6399 | 23.1514 | 23.5366 | 25.9439 | 25.9626 |
| Probable cytosolic iron-sulfur protein assembly protein C | O76071 | CIAO1 | Other | 0.086533333 | 0.10284439 | 26.9537 | 26.1015 | 26.8377 | 26.9671 | 26.5865 | 26.5989 |
| Anamorsin | Q6FI81-3;Q6FI81 | CIAPIN1 | Other | -0.143833333 | 0.161449799 | 25.9071 | 25.1228 | 25.8765 | 25.8328 | 25.0978 | 25.5443 |
| Calcium and integrin-binding protein 1 | Q99828;Q99828-2 | CIB1 | Extracellular or secreted | 1.930433333 | 1.221234879 | 22.2737 | 22.5298 | 22.112 | 23.7634 | 25.2845 | 23.6589 |
| Cold-inducible RNA-binding protein | Q14011;Q14011-2 | CIRBP | Other | 1.025 | 0.435078662 | 23.1923 | 23.6108 | 26.0382 | 25.478 | 25.2691 | 25.1692 |
| Cirhin | Q969X6-2;Q969X6-3;Q969X6 | CIRH1A | Extracellular or secreted | -0.108266667 | 0.038121044 | 23.5409 | 23.4196 | 23.2362 | 25.1015 | 22.3898 | 22.3806 |
| Cytoskeleton-associated protein 4 | Q07065 | CKAP4 | Extracellular or secreted | 0.7826 | 1.355236927 | 24.0172 | 23.8687 | 23.3488 | 24.8232 | 24.5403 | 24.219 |
| Cytoskeleton-associated protein 5 | Q14008-2;Q14008;Q14008-3 | CKAP5 | Other | -1.417933333 | 1.404600509 | 27.8714 | 28.8304 | 28.7539 | 26.3652 | 27.4348 | 27.4019 |
| Creatine kinase U-type, mitochondrial | P12532;P12532-2 | CKMT1A | Extracellular or secreted | -1.224 | 1.430153855 | 31.0068 | 30.5098 | 30.8465 | 29.9035 | 29.8224 | 28.9652 |
| Claudin-4;Claudin-9;Claudin-6;Claudin-3 | O14493;O95484;P56747;O15 | CLDN4;CLDN9; | Extracellular or secreted | 2.273933333 | 0.796635696 | 23.9359 | 24.5452 | 21.1053 | 25.1269 | 25.9176 | 25.3637 |
| Claudin-7 | O95471-2;O95471 | CLDN7 | Extracellular or secreted | 2.306 | 1.034423514 | 21.8674 | 24.2344 | 21.5861 | 25.5741 | 25.0197 | 24.0121 |
| Claudin domain-containing protein 1 | Q9NY35-2;Q9NY35 | CLDND1 | Extracellular or secreted | 1.850866667 | 1.514334459 | 21.6652 | 20.9767 | 20.1815 | 23.3769 | 22.8391 | 22.16 |
| Protein CLEC16A | Q2KHT3-2;Q2KHT3 | CLEC16A | Extracellular or secreted | 0.196766667 | 0.159036841 | 30.3436 | 29.4781 | 29.0954 | 30.2559 | 29.9379 | 29.3136 |
| Calmegin | O14967-2;O14967 | CLGN | Extracellular or secreted | -0.769733333 | 0.224075255 | 23.0003 | 21.5457 | 23.0431 | 21.2195 | 20.0231 | 24.0373 |
| Chloride intracellular channel protein 1 | O00299 | CLIC1 | Extracellular or secreted | 0.016133333 | 0.01016457 | 30.9105 | 32.3281 | 31.8585 | 31.0924 | 32.0149 | 32.0382 |
| Chloride intracellular channel protein 4 | Q9Y696 | CLIC4 | Extracellular or secreted | -0.4299 | 0.475939024 | 29.3351 | 29.9556 | 29.5605 | 28.5278 | 29.4808 | 29.5529 |
| Chloride intracellular channel protein 5;Chloride intracel | Q9NZA1-3;Q9NZA1-2;Q9NZA | CLIC5;CLIC6 | Extracellular or secreted | -0.405866667 | 0.639445794 | 23.775 | 24.3462 | 23.5536 | 23.227 | 23.7133 | 23.5169 |
| Clathrin interactor 1 | Q14677;Q14677-3;Q14677-2 | CLINT1 | Other | -0.1023 | 0.066674712 | 26.8836 | 25.1862 | 25.9676 | 25.883 | 25.6674 | 26.1801 |
| CAP-Gly domain-containing linker protein 1 | P30622-2;P30622-1;P30622 | CLIP1 | Extracellular or secreted | 1.100566667 | 0.720583747 | 24.462 | 25.6536 | 26.4649 | 26.3344 | 27.0717 | 26.4761 |
| Ceroid-lipofuscinosis neuronal protein 5 | O75503 | CLN5 | Extracellular or secreted | 1.271166667 | 2.251299146 | 25.5877 | 25.7304 | 25.612 | 26.9911 | 27.0899 | 26.6626 |
| Methylosome subunit pICln | P54105 | CLNS1A | Extracellular or secreted | -1.681133333 | 0.582465923 | 24.6916 | 25.0269 | 25.093 | 24.5595 | 21.0947 | 24.1139 |
| Caseinolytic peptidase B protein homolog | Q9H078-3;Q9H078-2;Q9H07 | CLPB | Other | -2.393266667 | 0.679252475 | 25.0432 | 23.8888 | 25.9523 | 19.887 | 23.832 | 23.9855 |
| ATP-dependent Clp protease proteolytic subunit, mitoch | Q16740 | CLPP | Other | -0.090066667 | 0.050079402 | 25.778 | 25.3255 | 27.2333 | 26.1239 | 26.1401 | 25.8026 |
| Calsyntenin-1;Soluble Alc-alpha;CTF1-alpha | O94985-2;O94985 | CLSTN1 | Extracellular or secreted | 1.1127 | 0.645056721 | 31.8658 | 32.4702 | 30.2866 | 32.4746 | 32.9069 | 32.5792 |
| Calsyntenin-3 | Q9BQT9;Q9BQT9-2 | CLSTN3 | Extracellular or secreted | 3.516466667 | 0.863465344 | 25.6944 | 27.669 | 22.6562 | 28.8871 | 28.7075 | 28.9744 |
| Clathrin light chain A | P09496-5;P09496-2;P09496- | CLTA | Extracellular or secreted | 0.352466667 | 0.385039534 | 24.4044 | 25.2342 | 25.4381 | 25.0096 | 25.7304 | 25.3941 |
| Clathrin light chain B | P09497-2;P09497 | CLTB | Extracellular or secreted | 0.331833333 | 0.269900352 | 26.1732 | 26.6752 | 27.5118 | 26.9658 | 27.687 | 26.7029 |
| Clathrin heavy chain 1 | Q00610-2;Q00610 | CLTC | Extracellular or secreted | -0.633066667 | 0.602627612 | 32.313 | 32.9152 | 33.3143 | 31.6059 | 32.1702 | 32.8672 |
| Clusterin;Clusterin beta chain;Clusterin alpha chain | P10909-4;P10909;P10909-5; | CLU | Extracellular or secreted | -0.2132 | 0.194023523 | 27.15 | 27.8053 | 26.5657 | 26.5671 | 27.2692 | 27.0451 |
| Clustered mitochondria protein homolog | O75153 | CLUH | Other | -2.621866667 | 0.940604093 | 28.0241 | 27.6301 | 28.6311 | 23.6313 | 25.6332 | 27.1552 |
| N-acylneuraminate cytidylyltransferase | Q8NFW8;Q8NFW8-2 | CMAS | Other | -0.092633333 | 0.028102119 | 26.7818 | 27.1564 | 29.041 | 25.9677 | 27.8422 | 28.8914 |
| COX assembly mitochondrial protein homolog | Q7Z7K0 | CMC1 | Other | -1.385766667 | 1.541063052 | 23.0093 | 22.4391 | 22.8457 | 22.0057 | 21.0641 | 21.067 |
| UMP-CMP kinase | P30085;P30085-2 | CMPK1 | Extracellular or secreted | -0.050366667 | 0.087623833 | 29.0813 | 29.0939 | 29.0399 | 28.749 | 29.3902 | 28.9248 |
| Cellular nucleic acid-binding protein | P62633-2;P62633;P62633-7; | CNBP | Other | 0.0262 | 0.016043853 | 28.8753 | 29.2859 | 27.8809 | 28.3024 | 29.39 | 28.4283 |
| Cellular nucleic acid-binding protein | P62633-8;P62633-4 | CNBP | Other | -0.106 | 0.055533887 | 22.699 | 22.2097 | 21.4505 | 20.929 | 22.5751 | 22.5371 |
| Cytosolic non-specific dipeptidase | Q96KP4;Q96KP4-2 | CNDP2 | Extracellular or secreted | 0.2789 | 0.822644786 | 30.3232 | 30.6114 | 30.3134 | 30.5371 | 30.9325 | 30.6151 |
| Protein cornichon homolog 4 | Q9P003 | CNIH4 | Other | 1.672166667 | 2.151208104 | 21.8423 | 22.551 | 22.6142 | 24.0032 | 24.3565 | 23.6643 |
| Calponin-2 | Q99439;Q99439-2 | CNN2 | Extracellular or secreted | -0.1098 | 0.07122515 | 25.4653 | 26.9419 | 26.4213 | 25.7111 | 26.0093 | 26.7787 |
| Calponin-3 | Q15417;Q15417-3;Q15417-2 | CNN3 | Other | 2.9137 | 0.644682352 | 19.8647 | 25.3464 | 20.7128 | 24.3791 | 24.8237 | 25.4622 |
| CCR4-NOT transcription complex subunit 1 | A5YKK6-2;A5YKK6;A5YKK6-3 | CNOT1 | Extracellular or secreted | -1.893166667 | 0.399540902 | 24.7395 | 24.9664 | 26.2684 | 19.9457 | 24.631 | 25.7181 |
| CCR4-NOT transcription complex subunit 3 | O75175 | CNOT3 | Other | -0.511666667 | 0.272656565 | 22.2996 | 22.8469 | 22.9417 | 21.698 | 21.3342 | 23.521 |
| 2,3-cyclic-nucleotide 3-phosphodiesterase | P09543-2;P09543 | CNP | Extracellular or secreted | 0.231133333 | 0.283429724 | 24.5619 | 24.9814 | 25.5654 | 25.4943 | 25.1776 | 25.1302 |
| Protein canopy homolog 2 | Q9Y2B0 | CNPY2 | Extracellular or secreted | -0.909933333 | 0.694700476 | 29.5894 | 29.1818 | 29.8923 | 29.58 | 28.4787 | 27.875 |
| Protein canopy homolog 3 | Q9BT09 | CNPY3 | Extracellular or secreted | -0.8824 | 1.198318261 | 26.0642 | 25.9748 | 26.5774 | 25.2595 | 24.8866 | 25.8231 |
| Contactin-1 | Q12860-2;Q12860;Q12860-3 | CNTN1 | Extracellular or secreted | 1.535733333 | 0.98404654 | 28.7266 | 28.4128 | 26.921 | 29.7733 | 29.5554 | 29.3389 |
| Cytochrome c oxidase assembly factor 6 homolog | Q5JTJ3-3;Q5JTJ3;Q5JTJ3-2 | COA6 | Other | -1.143733333 | 0.791297254 | 27.7798 | 27.2443 | 28.2675 | 27.1496 | 27.1878 | 25.523 |
| Cytochrome c oxidase assembly factor 7 | Q96BR5 | COA7 | Other | -0.6999 | 1.150986081 | 25.3216 | 25.6334 | 26.0866 | 25.2655 | 24.9959 | 24.6805 |
| Cochlin | O43405;O43405-2 | COCH | Extracellular or secreted | 0.3887 | 0.180210281 | 26.9514 | 27.5402 | 25.0341 | 26.7501 | 27.1524 | 26.7893 |
| Conserved oligomeric Golgi complex subunit 4 | Q9H9E3;Q9H9E3-2;Q9H9E3- | COG4 | Other | -0.4284 | 0.117992122 | 22.0298 | 23.8007 | 25.5459 | 21.6838 | 24.106 | 24.3014 |
| Conserved oligomeric Golgi complex subunit 6 | Q9Y2V7;Q9Y2V7-2 | COG6 | Other | -0.801333333 | 0.381278401 | 21.5506 | 21.5126 | 23.3006 | 22.6288 | 20.7478 | 20.5832 |
| Conserved oligomeric Golgi complex subunit 7 | P83436 | COG7 | Other | -1.066033333 | 0.854176733 | 24.7346 | 24.5757 | 23.6771 | 22.9535 | 22.6679 | 24.1679 |
| Conserved oligomeric Golgi complex subunit 8 | Q96MW5 | COG8 | Other | -0.4519 | 0.265685593 | 21.8635 | 22.1736 | 23.1278 | 20.8294 | 22.5084 | 22.4714 |
| Collagen alpha-1(XII) chain | Q99715;Q99715-4;Q99715-2 | COL12A1 | Extracellular or secreted | 1.311366667 | 0.980314563 | 32.2548 | 32.2665 | 30.7435 | 32.5305 | 33.3819 | 33.2865 |
| Collagen alpha-1(XVI) chain | Q07092-2;Q07092 | COL16A1 | Extracellular or secreted | 1.175366667 | 1.383387155 | 26.7572 | 26.2866 | 25.7351 | 26.9953 | 27.4128 | 27.8969 |

| | | | | | | | | | | | |
|---|---|---|---|---|---|---|---|---|---|---|---|
| Collagen alpha-1(XVIII) chain;Endostatin | P39060-2;P39060-1;P39060 | COL18A1 | Extracellular or secreted | 1.18 | 0.543859222 | 23.4306 | 20.6029 | 22.2067 | 23.2446 | 23.1514 | 23.3842 |
| Collagen alpha-1(IV) chain;Arresten | P02462 | COL4A1 | Extracellular or secreted | 0.4222 | 0.714770168 | 25.1453 | 24.748 | 24.6224 | 25.4507 | 25.4969 | 24.8347 |
| Collagen alpha-2(IV) chain;Canstatin | P08572 | COL4A2 | Extracellular or secreted | -0.041033333 | 0.030537584 | 28.5411 | 28.8952 | 27.5024 | 28.2259 | 28.4833 | 28.1064 |
| Collagen type IV alpha-3-binding protein | Q9Y5P4-2;Q9Y5P4;Q9Y5P4-3 | COL4A3BP | Other | -0.2648 | 0.150894458 | 22.5183 | 22.8292 | 22.7172 | 21.6094 | 22.0511 | 23.6098 |
| Collagen alpha-1(V) chain | P20908-2;P20908 | COL5A1 | Extracellular or secreted | 0.9939 | 0.196462744 | 23.4241 | 24.2013 | 19.7089 | 20.7159 | 24.6828 | 24.9173 |
| Collagen alpha-1(VI) chain | P12109 | COL6A1 | Extracellular or secreted | 0.695666667 | 0.37430223 | 30.1643 | 29.9854 | 27.9961 | 30.0743 | 30.21 | 29.9485 |
| Collagen alpha-2(VI) chain | P12110-3;P12110-2;P12110 | COL6A2 | Extracellular or secreted | 1.274066667 | 0.942558813 | 23.6092 | 23.2829 | 21.9885 | 23.4986 | 24.7013 | 24.5029 |
| Collagen alpha-1(VII) chain | Q02388-2;Q02388 | COL7A1 | Extracellular or secreted | 0.891566667 | 1.365902993 | 26.3215 | 26.6422 | 25.8233 | 26.8526 | 27.2304 | 27.3787 |
| Procollagen galactosyltransferase 1 | Q8NBJ5 | COLGALT1 | Extracellular or secreted | 0.035366667 | 0.018114946 | 26.4852 | 26.0977 | 28.0721 | 27.0003 | 26.6694 | 27.0914 |
| COMM domain-containing protein 3 | Q9UBI1 | COMMD3 | Other | 1.1456 | 0.691390388 | 22.8023 | 23.1531 | 21.2844 | 22.9519 | 23.2093 | 24.5154 |
| COMM domain-containing protein 4 | Q9H0A8-3;Q9H0A8-2;Q9H0A | COMMD4 | Other | 0.3692 | 0.160192235 | 20.6144 | 21.2887 | 22.5716 | 23.0822 | 21.6153 | 20.8848 |
| COMM domain-containing protein 8 | Q9NX08 | COMMD8 | Other | -0.230066667 | 0.10648715 | 22.4282 | 22.8979 | 23.7207 | 21.4529 | 23.4464 | 23.4573 |
| COMM domain-containing protein 9 | Q9P000;Q9P000-2 | COMMD9 | Other | 0.253033333 | 0.194692728 | 21.5351 | 22.3546 | 21.2812 | 21.2254 | 22.275 | 22.4296 |
| Catechol O-methyltransferase | P21964-2;P21964 | COMT | Extracellular or secreted | 0.215033333 | 0.144823218 | 26.7366 | 26.7895 | 27.8359 | 26.5004 | 27.7155 | 27.7912 |
| Coatomer subunit alpha;Xenin;Proxenin | P53621;P53621-2 | COPA | Extracellular or secreted | -0.155766667 | 0.101494102 | 29.6723 | 29.7099 | 30.6955 | 29.2573 | 29.6416 | 30.7115 |
| Coatomer subunit beta | P53618 | COPB1 | Extracellular or secreted | -0.178733333 | 0.107482531 | 28.5377 | 28.5622 | 29.6472 | 28.2743 | 28.2539 | 29.6827 |
| Coatomer subunit beta | P35606;P35606-2 | COPB2 | Other | -0.3496 | 0.24135459 | 29.3857 | 29.267 | 30.0729 | 28.5918 | 28.8851 | 30.1999 |
| Coatomer subunit epsilon | O14579;O14579-2;O14579-3 | COPE | Other | 0.023533333 | 0.067818047 | 29.0721 | 29.3147 | 29.0587 | 28.9963 | 29.2687 | 29.2511 |
| Coatomer subunit gamma-1 | Q9Y678 | COPG1 | Other | -0.437633333 | 0.275674856 | 29.0579 | 28.9766 | 29.9908 | 28.1079 | 28.686 | 29.9185 |
| Coatomer subunit gamma-2 | Q9UBF2;Q9UBF2-2 | COPG2 | Other | -2.546466667 | 0.776210683 | 25.1592 | 25.3538 | 26.5879 | 23.0833 | 21.0356 | 25.3426 |
| COP9 signalosome complex subunit 2 | P61201;P61201-2 | COPS2 | Other | -0.7978 | 1.002220902 | 28.4989 | 28.4777 | 29.0566 | 27.3494 | 27.9062 | 28.3842 |
| COP9 signalosome complex subunit 3 | Q9UNS2;Q9UNS2-2 | COPS3 | Other | -1.297366667 | 1.109013765 | 28.1764 | 28.2194 | 29.0598 | 26.449 | 27.155 | 27.9595 |
| COP9 signalosome complex subunit 4 | Q9BT78;Q9BT78-2 | COPS4 | Extracellular or secreted | -0.964233333 | 0.835960589 | 28.6381 | 28.799 | 28.8184 | 26.9625 | 28.0689 | 28.3314 |
| COP9 signalosome complex subunit 5 | Q92905 | COPS5 | Other | -1.095366667 | 1.203997998 | 27.8187 | 27.9663 | 28.1078 | 26.2543 | 27.2339 | 27.1185 |
| COP9 signalosome complex subunit 6 | Q7L5N1 | COPS6 | Extracellular or secreted | -0.607666667 | 0.555490773 | 27.7524 | 28.1674 | 28.5421 | 26.771 | 27.7306 | 28.1373 |
| COP9 signalosome complex subunit 7a | Q9UBW8 | COPS7A | Other | -2.123266667 | 0.623163057 | 25.5938 | 25.8417 | 26.2463 | 21.4189 | 24.0563 | 25.8368 |
| COP9 signalosome complex subunit 7b | Q9H9Q2;Q9H9Q2-3;Q9H9Q2 | COPS7B | Other | -1.980833333 | 1.816105961 | 25.6771 | 26.0442 | 26.3777 | 23.4727 | 23.9747 | 24.7091 |
| COP9 signalosome complex subunit 8 | Q99627-2;Q99627 | COPS8 | Extracellular or secreted | -0.3766 | 0.300035672 | 26.6204 | 27.1524 | 27.5448 | 26.0165 | 26.6898 | 27.4815 |
| Coatomer subunit zeta-1 | P61923;P61923-4;P61923-5 | COPZ1 | Other | 0.3354 | 0.228556074 | 26.5493 | 26.603 | 27.7339 | 26.6944 | 27.0803 | 28.1177 |
| Coronin-1A | P31146 | CORO1A | Extracellular or secreted | 0.214833333 | 0.244580538 | 25.6037 | 26.1958 | 25.9321 | 25.8776 | 25.7882 | 26.7103 |
| Coronin-1B | Q9BR76 | CORO1B | Extracellular or secreted | 0.416733333 | 0.527390845 | 29.1787 | 29.2006 | 29.8901 | 29.3809 | 29.869 | 30.2697 |
| Coronin-1C | Q9ULV4;Q9ULV4-2;Q9ULV4- | CORO1C | Extracellular or secreted | -0.4147 | 0.731711431 | 29.0838 | 29.5463 | 29.3932 | 28.531 | 28.998 | 29.2502 |
| Coronin-7 | P57737-2;P57737-4;P57737; | CORO7 | Other | -1.2448 | 0.457250805 | 20.3199 | 22.5152 | 23.7916 | 20.9157 | 20.2386 | 21.738 |
| Coactosin-like protein | Q14019 | COTL1 | Extracellular or secreted | -0.2778 | 0.320052619 | 31.5084 | 31.4548 | 30.6187 | 31.0523 | 31.1771 | 30.5191 |
| Cytochrome c oxidase copper chaperone | Q14061 | COX17 | Other | 1.481533333 | 0.612615226 | 25.4273 | 23.6628 | 24.4434 | 27.7728 | 25.3624 | 24.8429 |
| Cytochrome c oxidase subunit 6B1 | P14854 | COX6B1 | Other | 0.1322 | 0.05649398 | 24.4088 | 24.9194 | 25.2661 | 25.2888 | 26.1016 | 23.6005 |
| Carboxypeptidase A4 | Q9UI42;Q9UI42-2 | CPA4 | Extracellular or secreted | -1.446 | 0.677628318 | 28.8514 | 28.9499 | 26.546 | 27.1155 | 27.2602 | 25.6336 |
| Carboxypeptidase D | O75976;O75976-2 | CPD | Extracellular or secreted | -0.7961 | 0.812100078 | 22.6841 | 22.2348 | 22.7519 | 22.1629 | 22.1037 | 21.0159 |
| Carboxypeptidase E | P16870-2;P16870 | CPE | Extracellular or secreted | 0.036133333 | 0.026138301 | 29.1337 | 28.5752 | 28.2403 | 28.2074 | 28.4164 | 29.4338 |
| Carboxypeptidase M | P14384 | CPM | Extracellular or secreted | -1.2157 | 1.284941823 | 24.4387 | 24.6917 | 25.5343 | 23.4994 | 24.24 | 23.2782 |
| Copine-1 | Q99829 | CPNE1 | Extracellular or secreted | -0.153333333 | 0.107545845 | 27.4808 | 27.3324 | 28.5123 | 26.9901 | 27.6504 | 28.225 |
| Copine-2;Copine-5;Copine-9;Copine-4;Copine-6;Copine-7 | Q96FN4;Q9HCH3-2;Q96FN4- | CPNE2;CPNE5; | Extracellular or secreted | -3.060466667 | 1.748384897 | 25.7729 | 25.1254 | 24.0577 | 22.7991 | 20.7928 | 22.1827 |
| Copine-3 | O75131 | CPNE3 | Extracellular or secreted | 1.1885 | 1.336839868 | 25.8278 | 25.0509 | 25.95 | 26.7563 | 26.9209 | 26.717 |
| Oxygen-dependent coproporphyrinogen-III oxidase, mito | P36551 | CPOX | Other | -1.453466667 | 2.369366741 | 30.1178 | 29.6646 | 29.7437 | 28.4788 | 28.6559 | 28.031 |
| Serine/threonine-protein phosphatase CPPED1 | Q9BRF8;Q9BRF8-2 | CPPED1 | Extracellular or secreted | 0.465333333 | 0.23201221 | 25.4401 | 24.2862 | 23.6322 | 25.9722 | 24.82 | 23.9623 |
| Carbamoyl-phosphate synthase [ammonia], mitochondri | P31327-2;P31327;P31327-3 | CPS1 | Other | 0.9473 | 0.642682834 | 26.113 | 26.8054 | 25.8854 | 28.1855 | 27.2069 | 26.2533 |
| Cleavage and polyadenylation specificity factor subunit 1 | Q10570 | CPSF1 | Other | -2.1366 | 0.888302447 | 25.9686 | 25.6389 | 27.3988 | 22.4782 | 24.5513 | 25.567 |
| Cleavage and polyadenylation specificity factor subunit 3 | Q9UKF6 | CPSF3 | Other | -1.8396 | 0.593201697 | 24.5385 | 24.8424 | 27.2468 | 21.8379 | 23.7081 | 25.5629 |
| Cleavage and polyadenylation specificity factor subunit 6 | Q16630-3;Q16630;Q16630-2 | CPSF6 | Other | -0.6226 | 0.397009389 | 26.8541 | 25.9486 | 27.3815 | 25.1875 | 26.1775 | 26.9514 |
| Cleavage and polyadenylation specificity factor subunit 7 | Q8N684-2;Q8N684;Q8N684- | CPSF7 | Other | -0.540766667 | 0.216838867 | 23.2663 | 23.2936 | 25.5296 | 22.4558 | 23.4429 | 24.5685 |
| Probable serine carboxypeptidase CPVL | Q9H3G5 | CPVL | Extracellular or secreted | -0.0177 | 0.008685217 | 29.248 | 29.5767 | 27.7615 | 29.3145 | 29.0771 | 28.1415 |
| Cellular retinoic acid-binding protein 2 | P29373 | CRABP2 | Extracellular or secreted | -0.910466667 | 0.310991921 | 23.8117 | 24.1908 | 21.2189 | 20.9505 | 23.4913 | 22.0482 |
| Carnitine O-acetyltransferase | P43155-2;P43155;P43155-3 | CRAT | Other | -1.2608 | 0.706104594 | 24.1575 | 25.5254 | 26.4857 | 23.6297 | 24.8347 | 23.9218 |
| Protein CREG1 | O75629 | CREG1 | Extracellular or secreted | 0.1513 | 0.082391264 | 26.7806 | 26.2664 | 26.1222 | 25.4644 | 27.5017 | 26.657 |
| Cysteine-rich with EGF-like domain protein 2 | Q6UXH1-4;Q6UXH1;Q6UXH1 | CRELD2 | Extracellular or secreted | -0.424266667 | 0.210822142 | 26.8178 | 25.803 | 26.4206 | 27.2718 | 25.5035 | 24.9933 |
| Cysteine-rich motor neuron 1 protein;Processed cysteine | Q9NZV1 | CRIM1 | Extracellular or secreted | 1.781766667 | 2.184577813 | 23.7957 | 24.137 | 23.6158 | 25.6624 | 26.0457 | 25.1857 |
| Cysteine-rich protein 1 | P50238 | CRIP1 | Other | 0.093766667 | 0.032149721 | 25.7953 | 28.4552 | 26.196 | 25.9407 | 27.6604 | 27.1267 |
| Adapter molecule crk | P46108 | CRK | Extracellular or secreted | -0.467833333 | 0.772764249 | 29.4127 | 29.0895 | 28.6678 | 28.7211 | 28.798 | 28.2474 |
| Crk-like protein | P46109 | CRKL | Extracellular or secreted | -0.380066667 | 1.120731439 | 27.5266 | 27.2793 | 27.2035 | 27.0387 | 27.1156 | 26.7149 |
| Cytokine receptor-like factor 3 | Q8IUI8-2;Q8IUI8 | CRLF3 | Extracellular or secreted | -0.239466667 | 0.188291175 | 23.321 | 23.3023 | 24.447 | 22.8741 | 23.5807 | 23.8971 |
| Rootletin | Q5TZA2-2;Q5TZA2 | CROCC | Extracellular or secreted | 0.562033333 | 0.453333483 | 28.5075 | 28.952 | 27.3847 | 29.1317 | 28.6829 | 28.7157 |
| Cartilage-associated protein | O75718 | CRTAP | Extracellular or secreted | 0.2453 | 0.121934333 | 21.5161 | 22.744 | 23.8487 | 23.2192 | 23.0362 | 22.5893 |
| Quinone oxidoreductase | Q08257-3;Q08257 | CRYZ | Extracellular or secreted | -0.239 | 0.291460999 | 30.9933 | 30.6495 | 30.6347 | 30.9011 | 30.7156 | 29.9438 |
| Quinone oxidoreductase-like protein 1 | O95825;O95825-2 | CRYZL1 | Other | -0.295666667 | 0.847512477 | 26.6808 | 26.7116 | 26.5523 | 26.4407 | 26.5233 | 26.0937 |
| Citrate synthase, mitochondrial | O75390 | CS | Extracellular or secreted | -1.164366667 | 1.060403495 | 31.6896 | 31.3296 | 31.7355 | 31.0883 | 30.455 | 29.7183 |
| Cold shock domain-containing protein E1 | O75534-2;O75534;O75534-3 | CSDE1 | Extracellular or secreted | 0.117433333 | 0.070479461 | 26.9703 | 26.7919 | 27.8747 | 26.5019 | 27.3502 | 28.1371 |
| Exportin-2 | P55060-3;P55060;P55060-4 | CSE1L | Extracellular or secreted | -0.820333333 | 0.64406621 | 31.2542 | 32.5103 | 32.512 | 30.718 | 31.0646 | 32.0329 |
| Macrophage colony-stimulating factor 1;Processed macr | P09603;P09603-2;P09603-3 | CSF1 | Extracellular or secreted | 3.278766667 | 1.378508054 | 26.4218 | 26.871 | 24.328 | 29.3727 | 29.4405 | 28.6439 |
| Granulocyte colony-stimulating factor | P09919;P09919-3;P09919-2 | CSF3 | Extracellular or secreted | 3.3825 | 1.217290349 | 26.94 | 26.6568 | 23.8951 | 29.1789 | 29.8133 | 28.6472 |
| Tyrosine-protein kinase CSK | P41240 | CSK | Extracellular or secreted | -0.590766667 | 0.250564194 | 22.4117 | 22.1111 | 23.3778 | 21.7466 | 20.7996 | 23.5821 |
| Casein kinase I isoform alpha | P48729-3;P48729;P48729-2 | CSNK1A1 | Other | -0.475566667 | 0.684426541 | 25.4003 | 25.2732 | 25.9769 | 24.7804 | 24.9156 | 25.5277 |
| Casein kinase II subunit alpha;Casein kinase II subunit alp | P68400;P68400-2;Q8NEV1 | CSNK2A1;CSNK | Extracellular or secreted | 1.134233333 | 0.632012303 | 24.8355 | 22.6266 | 24.4616 | 25.0528 | 24.7656 | 25.508 |
| Casein kinase II subunit alpha | P19784 | CSNK2A2 | Other | 1.005966667 | 0.373105282 | 23.8413 | 21.9755 | 25.3833 | 24.1345 | 24.5426 | 25.5409 |
| Casein kinase II subunit beta | P67870 | CSNK2B | Extracellular or secreted | 0.684566667 | 0.531803585 | 25.3333 | 24.8834 | 26.5191 | 26.051 | 26.1529 | 26.5856 |
| Chondroitin sulfate proteoglycan 4 | Q6UVK1 | CSPG4 | Extracellular or secreted | -1.339666667 | 0.993541101 | 28.845 | 29.4104 | 27.8117 | 27.2393 | 27.4021 | 27.4067 |
| Cysteine and glycine-rich protein 1 | P21291 | CSRP1 | Extracellular or secreted | 0.184233333 | 0.109670361 | 28.8711 | 30.0483 | 28.4929 | 28.5784 | 29.8652 | 29.5214 |
| Cysteine and glycine-rich protein 2 | Q16527 | CSRP2 | Other | -2.1743 | 1.125313612 | 24.7886 | 24.4312 | 25.3905 | 21.5587 | 23.9789 | 22.5498 |
| Cystatin-C | P01034 | CST3 | Extracellular or secreted | 0.084433333 | 0.049438554 | 31.3041 | 31.883 | 30.188 | 30.6792 | 31.6931 | 31.2561 |
| Cystatin-M | Q15828 | CST6 | Extracellular or secreted | 0.761733333 | 0.235064329 | 28.3001 | 30.2277 | 26.2394 | 29.2626 | 29.4377 | 28.3521 |
| Cystatin-F | O76096 | CST7 | Extracellular or secreted | 3.892466667 | 1.223109926 | 25.216 | 22.4673 | 21.9225 | 26.8484 | 27.1133 | 27.3215 |
| Cystatin-A;Cystatin-A, N-terminally processed | P01040 | CSTA | Extracellular or secreted | 0.9686 | 0.627048272 | 28.5046 | 28.6794 | 28.6088 | 30.3947 | 29.8542 | 28.4497 |
| Cystatin-B | P04080 | CSTB | Extracellular or secreted | 1.027733333 | 0.863130454 | 32.0009 | 32.3828 | 30.9529 | 32.6555 | 33.4594 | 32.3049 |
| Cleavage stimulation factor subunit 1 | Q05048 | CSTF1 | Other | -0.276933333 | 0.1942325 | 27.0354 | 27.0484 | 28.184 | 26.4377 | 27.2042 | 27.7951 |
| Cleavage stimulation factor subunit 2 | P33240-2;P33240 | CSTF2 | Other | -1.0059 | 0.320453647 | 26.2773 | 26.5177 | 27.9963 | 23.7488 | 26.5888 | 27.436 |
| Cleavage stimulation factor subunit 2 tau variant | Q9H0L4 | CSTF2T | Other | -0.944266667 | 0.279825849 | 22.9287 | 22.0361 | 25.5702 | 22.5167 | 21.1454 | 24.0401 |
| Cleavage stimulation factor subunit 3 | Q12996 | CSTF3 | Other | -1.0626 | 0.601376456 | 27.1111 | 27.1224 | 27.7676 | 25.2336 | 26.0499 | 27.5298 |
| C-terminal-binding protein 1 | Q13363-2;Q13363 | CTBP1 | Other | -0.210466667 | 0.037081778 | 24.7322 | 23.3601 | 27.5927 | 23.1763 | 23.9729 | 27.9044 |
| C-terminal-binding protein 2 | P56545;P56545-3;P56545-2 | CTBP2 | Other | -0.213733333 | 0.060304941 | 22.963 | 21.061 | 24.1089 | 23.7828 | 20.8962 | 22.8127 |
| Di-N-acetylchitobiase | Q01459 | CTBS | Extracellular or secreted | 1.079466667 | 1.505888055 | 27.8874 | 27.5039 | 27.1202 | 28.9458 | 28.6866 | 28.1175 |
| Transcriptional repressor CTCF | P49711;P49711-2 | CTCF | Other | 0.7976 | 0.227212046 | 22.9207 | 19.2963 | 22.7867 | 21.2656 | 22.8297 | 23.3012 |
| RNA polymerase II subunit A C-terminal domain phospha | Q9Y5B0-4;Q9Y5B0 | CTDP1 | Other | -2.489766667 | 1.661421975 | 27.6651 | 27.0963 | 26.3013 | 25.5637 | 24.1582 | 23.8715 |
| Connective tissue growth factor | P29279;P29279-2 | CTGF | Extracellular or secreted | -1.213333333 | 0.551066122 | 27.0637 | 26.4446 | 24.4257 | 25.6229 | 24.8399 | 23.8312 |
| Cystathionine gamma-lyase | P32929;P32929-2;P32929-3 | CTH | Extracellular or secreted | -0.263466667 | 0.343310844 | 26.967 | 27.4116 | 27.8582 | 26.8055 | 27.4128 | 27.2281 |
| Catenin alpha-1 | P35221;P35221-2;P35221-3 | CTNNA1 | Extracellular or secreted | -0.175833333 | 0.204056439 | 31.0223 | 30.7941 | 31.2058 | 30.4706 | 30.6052 | 31.4189 |
| Catenin alpha-2 | P26232-2;P26232-5;P26232; | CTNNA2 | Extracellular or secreted | -1.115733333 | 1.165199764 | 28.7018 | 29.297 | 28.2018 | 27.7015 | 27.524 | 27.6279 |
| Catenin beta-1 | P35222 | CTNNB1 | Extracellular or secreted | 0.667966667 | 0.919348439 | 29.5699 | 29.4369 | 29.862 | 29.8887 | 30.1675 | 30.8165 |
| Beta-catenin-like protein 1 | Q8WYA6-4;Q8WYA6;Q8WYA | CTNNBL1 | Other | -1.316533333 | 0.294803525 | 23.4546 | 23.4605 | 26.3616 | 21.0899 | 22.2085 | 26.0287 |
| Catenin delta-1 | O60716-21;O60716-19;O607 | CTNND1 | Extracellular or secreted | 1.5286 | 1.911183131 | 28.0699 | 27.2947 | 27.8983 | 29.1171 | 29.2625 | 29.4691 |
| CTP synthase 1 | P17812;P17812-2 | CTPS1 | Other | -0.697466667 | 0.67802292 | 27.6264 | 27.8666 | 28.7081 | 26.8353 | 27.2925 | 27.9809 |
| CTP synthase 2 | Q9NRF8 | CTPS2 | Other | -0.168033333 | 0.128472043 | 21.3668 | 21.5418 | 22.7813 | 21.8578 | 21.5884 | 21.7396 |
| RNA polymerase-associated protein CTR9 homolog | Q6PD62 | CTR9 | Other | 0.8121 | 0.250860491 | 23.3799 | 23.057 | 21.5848 | 25.2711 | 21.44 | 23.7469 |
| Lysosomal protective protein;Lysosomal protective prote | P10619-2;P10619 | CTSA | Extracellular or secreted | 0.515866667 | 0.717801517 | 28.0043 | 28.5917 | 27.6538 | 28.6789 | 28.725 | 28.3935 |
| Cathepsin B;Cathepsin B light chain;Cathepsin B heavy ch | P07858 | CTSB | Extracellular or secreted | 0.7964 | 0.907576683 | 28.9126 | 29.61 | 28.4744 | 29.6605 | 30.0664 | 29.6593 |
| Dipeptidyl peptidase 1;Dipeptidyl peptidase 1 exclusion d | P53634 | CTSC | Extracellular or secreted | 1.201333333 | 0.778196525 | 31.9088 | 32.4886 | 30.5122 | 33.1993 | 33.0879 | 32.2264 |
| Cathepsin D;Cathepsin D light chain;Cathepsin D heavy c | P07339 | CTSD | Extracellular or secreted | 1.0628 | 1.120754698 | 32.044 | 32.0644 | 30.991 | 32.526 | 33.1448 | 32.617 |
| Cathepsin F | Q9UBX1 | CTSF | Extracellular or secreted | 0.4629 | 0.170104501 | 23.1782 | 23.7976 | 22.2777 | 21.7527 | 24.4925 | 24.397 |
| Pro-cathepsin H;Cathepsin H mini chain;Cathepsin H;Cath | P09668 | CTSH | Extracellular or secreted | 1.676366667 | 0.948072638 | 28.3043 | 29.5511 | 27.2994 | 29.679 | 30.3891 | 30.1158 |
| Cathepsin L1;Cathepsin L1 heavy chain;Cathepsin L1 light | P07711 | CTSL | Extracellular or secreted | 0.431266667 | 0.327439355 | 29.3894 | 29.4602 | 28.1765 | 29.9747 | 29.5448 | 28.8004 |
| Cathepsin O | P43234 | CTSO | Extracellular or secreted | 1.024866667 | 0.319871284 | 20.8986 | 23.8443 | 22.0862 | 21.3164 | 24.3242 | 24.2631 |
| Cathepsin S | P25774-2;P25774 | CTSS | Extracellular or secreted | 2.090266667 | 1.579950533 | 23.6733 | 24.1069 | 23.2123 | 25.1632 | 26.6608 | 25.4393 |
| Cathepsin L2 | O60911 | CTSV | Extracellular or secreted | -0.329233333 | 0.125047897 | 28.8669 | 29.7803 | 26.9538 | 28.8394 | 28.4227 | 27.3512 |
| Cathepsin Z | Q9UBR2 | CTSZ | Extracellular or secreted | 1.338466667 | 0.762463883 | 31.7123 | 32.3059 | 30.0748 | 33.0624 | 33.0322 | 32.0138 |
| Src substrate cortactin | Q14247;Q14247-3;Q14247-2 | CTTN | Extracellular or secreted | 1.947733333 | 1.678214594 | 27.6285 | 27.4673 | 28.417 | 29.7359 | 29.7781 | 29.842 |
| Cullin-1 | Q13616 | CUL1 | Other | -1.0769 | 0.685846721 | 27.5399 | 27.9786 | 28.6756 | 25.8414 | 27.3096 | 27.8124 |
| Cullin-2 | Q13617;Q13617-2 | CUL2 | Other | -0.746533333 | 0.510813799 | 27.6789 | 28.0604 | 28.6454 | 26.3823 | 27.5191 | 28.2437 |
| Cullin-3 | Q13618-2;Q13618;Q13618-3 | CUL3 | Extracellular or secreted | -2.7973 | 0.961622277 | 25.6414 | 26.0474 | 27.0296 | 22.3811 | 22.3364 | 25.609 |
| Cullin-4A | Q13619;Q13619-2 | CUL4A | Other | -0.628666667 | 0.548561569 | 26.9594 | 27.2533 | 27.4457 | 25.7831 | 26.7149 | 27.2744 |
| Cullin-4B | Q13620-3;Q13620-1;Q13620 | CUL4B | Extracellular or secreted | -1.3522 | 0.599921509 | 24.7697 | 24.458 | 24.7794 | 21.6221 | 24.0648 | 24.2636 |
| Cullin-5 | Q93034 | CUL5 | Other | -1.747066667 | 1.639838611 | 27.6625 | 27.1679 | 26.5669 | 25.2844 | 25.2845 | 25.5872 |

| | | | | | | | | | | | |
|---|---|---|---|---|---|---|---|---|---|---|---|
| Protein CutA | O60888-3;O60888;O60888-2 | CUTA | Extracellular or secreted | 0.0334 | 0.017976346 | 29.5117 | 29.8969 | 28.0571 | 29.4424 | 29.3375 | 28.786 |
| Copper homeostasis protein cutC homolog | Q9NTM9 | CUTC | Other | -0.412433333 | 0.2916551 | 25.9076 | 25.1814 | 24.4631 | 24.9152 | 25.3664 | 24.0332 |
| Homeobox protein cut-like 1 | P39880-5;P39880-2;P39880; | CUX1 | Other | -2.9292 | 0.773409147 | 26.4111 | 25.7588 | 26.5635 | 20.5263 | 24.4499 | 24.9696 |
| Spliceosome-associated protein CWC15 homolog | Q9P013 | CWC15 | Other | 1.223633333 | 0.546756629 | 21.2604 | 20.2835 | 23.1757 | 22.3649 | 23.2644 | 22.7612 |
| Pre-mRNA-splicing factor CWC22 homolog | Q9HCG8 | CWC22 | Other | 0.488066667 | 0.159598396 | 22.2143 | 19.9127 | 23.4169 | 22.9492 | 21.5826 | 22.4763 |
| Peptidyl-prolyl cis-trans isomerase CWC27 homolog | Q6UX04-2;Q6UX04 | CWC27 | Other | -1.117 | 0.462337132 | 24.4273 | 23.6162 | 26.6417 | 23.6185 | 23.2562 | 24.4595 |
| CWF19-like protein 1 | Q69YN2;Q69YN2-3;Q69YN2- | CWF19L1 | Other | 0.2902 | 0.114069593 | 21.4322 | 23.2648 | 23.9142 | 22.5644 | 23.1377 | 24.1397 |
| Coxsackievirus and adenovirus receptor | P78310-2;P78310-6;P78310; | CXADR | Extracellular or secreted | 0.100666667 | 0.062354214 | 24.1618 | 23.1278 | 22.4558 | 23.5972 | 23.5571 | 22.8931 |
| Growth-regulated alpha protein;GRO-alpha(4-73);GRO-a | P09341 | CXCL1 | Extracellular or secreted | 1.705333333 | 0.783185461 | 33.9337 | 34.5563 | 31.9079 | 35.0674 | 35.366 | 35.0805 |
| C-X-C motif chemokine 16 | Q9H2A7 | CXCL16 | Extracellular or secreted | 1.3666 | 0.488999528 | 25.731 | 27.3755 | 23.7387 | 26.9364 | 27.6986 | 26.31 |
| C-X-C motif chemokine 2;GRO-beta(5-73) | P19875 | CXCL2 | Extracellular or secreted | 1.789066667 | 1.133610651 | 25.8679 | 26.5892 | 24.5881 | 26.7071 | 28.0902 | 27.6151 |
| C-X-C motif chemokine 3;GRO-gamma(5-73) | P19876 | CXCL3 | Extracellular or secreted | 2.6903 | 0.998846006 | 31.2539 | 32.3186 | 29.0881 | 33.3708 | 33.952 | 33.4087 |
| C-X-C motif chemokine 5;ENA-78(8-78);ENA-78(9-78) | P42830 | CXCL5 | Extracellular or secreted | 3.358533333 | 0.648706737 | 22.2101 | 27.9929 | 21.9775 | 25.7529 | 28.7935 | 27.7097 |
| Interleukin-8;MDNCF-a;Interleukin-8;IL-8(5-77);IL-8(6-77 | P10145 | CXCL8 | Extracellular or secreted | 1.968566667 | 0.688824777 | 31.4874 | 31.7915 | 28.4223 | 32.4831 | 32.9649 | 32.1589 |
| Cytochrome b5 | P00167-2;P00167;P00167-3 | CYB5A | Extracellular or secreted | 1.7203 | 0.477585144 | 23.1081 | 23.5329 | 27.3853 | 26.0516 | 26.4236 | 26.712 |
| Cytochrome b5 type B | O43169 | CYB5B | Other | 1.303966667 | 0.712111282 | 21.6162 | 22.0182 | 23.8744 | 24.1716 | 23.8731 | 23.376 |
| NADH-cytochrome b5 reductase 2 | Q6BCY4;Q6BCY4-2 | CYB5R2 | Other | 1.865833333 | 1.484774955 | 27.2928 | 27.4974 | 26.0998 | 29.1219 | 28.9538 | 28.4118 |
| NADH-cytochrome b5 reductase 3;NADH-cytochrome b5 | P00387-2;P00387;P00387-3 | CYB5R3 | Extracellular or secreted | 1.471233333 | 0.935372579 | 20.8445 | 22.6091 | 22.6065 | 23.0414 | 23.1581 | 24.2743 |
| Cytochrome c | P99999 | CYCS | Other | -1.614766667 | 1.655693967 | 32.6823 | 32.782 | 32.2113 | 30.8775 | 31.5456 | 30.4082 |
| Cytoplasmic FMR1-interacting protein 1 | Q7L576;Q7L576-2 | CYFIP1 | Extracellular or secreted | -0.055533333 | 0.042564947 | 27.7048 | 28.0193 | 28.8947 | 27.6816 | 28.1813 | 28.5893 |
| Cytoplasmic FMR1-interacting protein 2 | Q96F07-2;Q96F07 | CYFIP2 | Extracellular or secreted | -0.588833333 | 0.350016547 | 22.3288 | 22.8902 | 23.7262 | 21.3914 | 22.4609 | 23.3264 |
| Cysteine and histidine-rich protein 1 | Q6ZMK1-2;Q6ZMK1 | CYHR1 | Extracellular or secreted | 0.190133333 | 0.055528107 | 21.2166 | 23.4794 | 24.2074 | 21.6359 | 23.8454 | 23.9925 |
| Cytochrome P450 2B6 | P20813-2;P20813 | CYP2B6 | Extracellular or secreted | -2.440333333 | 0.90034442 | 24.8936 | 24.3441 | 23.3797 | 23.7381 | 20.2984 | 21.2599 |
| Lanosterol 14-alpha demethylase | Q16850-2;Q16850 | CYP51A1 | Extracellular or secreted | 1.462733333 | 0.9752858 | 21.3954 | 20.7571 | 21.6209 | 21.744 | 23.6995 | 22.7181 |
| Protein CYR61 | O00622 | CYR61 | Extracellular or secreted | -0.4117 | 0.210992297 | 29.4624 | 29.8946 | 27.6788 | 29.0073 | 28.6393 | 28.1541 |
| D-2-hydroxyglutarate dehydrogenase, mitochondrial | Q8N465;Q8N465-3 | D2HGDH | Extracellular or secreted | -0.6211 | 0.224059786 | 23.0819 | 20.4535 | 23.6282 | 22.3286 | 21.3666 | 21.6051 |
| Dystroglycan;Alpha-dystroglycan;Beta-dystroglycan | Q14118 | DAG1 | Extracellular or secreted | 0.316166667 | 0.171295257 | 31.189 | 31.4172 | 29.4438 | 31.014 | 31.4253 | 30.5592 |
| Bifunctional ATP-dependent dihydroxyacetone kinase/FA | Q3LXA3;Q3LXA3-2 | DAK | Extracellular or secreted | -0.508033333 | 0.563369037 | 26.8249 | 27.3714 | 27.5719 | 26.1024 | 27.0541 | 27.0876 |
| Aspartate--tRNA ligase, cytoplasmic | P14868;P14868-2 | DARS | Extracellular or secreted | -0.483333333 | 0.28069941 | 29.1646 | 28.9755 | 30.5785 | 28.4457 | 28.8102 | 30.0127 |
| Aspartate--tRNA ligase, mitochondrial | Q6PI48 | DARS2 | Other | -1.4385 | 0.315819143 | 21.3544 | 21.2601 | 26.3177 | 22.4206 | 21.0576 | 21.1385 |
| DAZ-associated protein 1 | Q96EP5-2;Q96EP5 | DAZAP1 | Other | -0.660933333 | 0.68337048 | 27.5924 | 27.847 | 27.884 | 26.512 | 27.0547 | 27.7739 |
| Acyl-CoA-binding protein | P07108;P07108-3;P07108-2; | DBI | Extracellular or secreted | 1.073833333 | 0.624822282 | 27.0481 | 27.1093 | 26.2592 | 28.9844 | 27.9311 | 26.7226 |
| Drebrin | Q16643;Q16643-2;Q16643-3 | DBN1 | Extracellular or secreted | 3.357333333 | 0.752571791 | 24.5789 | 21.2364 | 26.9357 | 27.2119 | 27.529 | 28.0821 |
| Drebrin-like protein | Q9UJU6;Q9UJU6-5 | DBNL | Extracellular or secreted | 0.887266667 | 0.167722537 | 22.8789 | 24.0678 | 25.5293 | 21.5288 | 26.4639 | 27.1451 |
| Drebrin-like protein | Q9UJU6-2;Q9UJU6-3;Q9UJU | DBNL | Extracellular or secreted | 0.5112 | 0.316316308 | 21.907 | 21.2296 | 22.2098 | 21.1976 | 22.572 | 23.1104 |
| Lariat debranching enzyme | Q9UK59;Q9UK59-2 | DBR1 | Other | -4.176666667 | 0.688720262 | 28.4212 | 30.1374 | 28.2519 | 26.8593 | 20.1793 | 27.2419 |
| Lipoamide acyltransferase component of branched-chain | P11182 | DBT | Other | 0.7024 | 0.254055526 | 22.4151 | 19.7335 | 22.9455 | 22.5957 | 22.7097 | 21.8959 |
| DDB1- and CUL4-associated factor 8 | Q5TAQ9;Q5TAQ9-2 | DCAF8 | Other | -0.2989 | 0.093005839 | 23.7542 | 22.732 | 24.4003 | 21.3566 | 23.9653 | 24.6679 |
| Discoidin, CUB and LCCL domain-containing protein 1 | Q8N8Z6-2;Q8N8Z6 | DCBLD1 | Other | 0.2367 | 0.212691525 | 26.1844 | 26.276 | 25.0486 | 25.919 | 26.0804 | 26.2197 |
| Discoidin, CUB and LCCL domain-containing protein 2 | Q96PD2;Q96PD2-2 | DCBLD2 | Extracellular or secreted | 1.106 | 0.509781131 | 23.7345 | 24.7329 | 22.311 | 25.8521 | 24.5945 | 23.6498 |
| Dermcidin;Survival-promoting peptide;DCD-1 | P81605;P81605-2 | DCD | Extracellular or secreted | 1.950066667 | 0.758721209 | 26.2217 | 25.1815 | 27.7422 | 30.0355 | 26.9434 | 28.0167 |
| Deoxycytidine kinase | P27707 | DCK | Other | -2.796733333 | 2.55896308 | 27.4249 | 27.821 | 26.9408 | 25.1187 | 24.0246 | 24.6532 |
| Decorin | P07585;P07585-4 | DCN | Extracellular or secreted | -0.539333333 | 0.2911741 | 26.8585 | 25.4677 | 24.5167 | 25.1512 | 25.161 | 24.9127 |
| m7GpppX diphosphatase | Q96C86 | DCPS | Other | -0.122833333 | 0.125131644 | 28.2186 | 28.3163 | 29.1589 | 28.115 | 28.4311 | 28.7722 |
| Deoxycytidylate deaminase | P32321;P32321-2 | DCTD | Extracellular or secreted | -0.609466667 | 0.668710319 | 28.4743 | 28.6062 | 28.3931 | 27.2439 | 28.4178 | 27.9835 |
| Dynactin subunit 1 | Q14203-3;Q14203-4;Q14203 | DCTN1 | Other | -0.660233333 | 0.409318982 | 28.6942 | 28.5708 | 29.6867 | 27.3812 | 28.2448 | 29.345 |
| Dynactin subunit 2 | Q13561;Q13561-3;Q13561-2 | DCTN2 | Extracellular or secreted | 0.229766667 | 0.181566168 | 27.8325 | 28.3948 | 28.5038 | 27.6345 | 28.8601 | 28.9258 |
| Dynactin subunit 3 | O75935-3;O75935;O75935-2 | DCTN3 | Other | -0.5855 | 0.160878998 | 24.0109 | 25.1919 | 24.3523 | 21.4834 | 24.7402 | 25.575 |
| Dynactin subunit 4 | Q9UJW0-2;Q9UJW0;Q9UJW | DCTN4 | Other | -0.7925 | 0.477450065 | 24.7523 | 24.6726 | 25.6618 | 23.0146 | 24.8794 | 24.8152 |
| Dynactin subunit 5 | Q9BTE1-3;Q9BTE1-2;Q9BTE1 | DCTN5 | Other | 0.156833333 | 0.208666806 | 22.0886 | 21.4471 | 22.2273 | 21.9078 | 21.9333 | 22.3924 |
| dCTP pyrophosphatase 1 | Q9H773 | DCTPP1 | Other | -0.071233333 | 0.048252933 | 29.155 | 28.4069 | 28.1853 | 29.2493 | 28.4447 | 27.8395 |
| DCN1-like protein 1 | Q96GG9 | DCUN1D1 | Other | 0.256866667 | 0.267440701 | 25.8541 | 25.5232 | 25.367 | 25.1749 | 26.2848 | 26.0546 |
| DCN1-like protein 5 | Q9BTE7 | DCUN1D5 | Other | -0.028266667 | 0.007441261 | 20.7287 | 19.6861 | 22.5339 | 22.4474 | 21.1658 | 19.2507 |
| L-xylulose reductase | Q7Z4W1 | DCXR | Extracellular or secreted | -1.2225 | 0.756010259 | 25.6235 | 25.662 | 27.4904 | 25.0224 | 24.5901 | 25.4959 |
| N(G),N(G)-dimethylarginine dimethylaminohydrolase 1 | O94760;O94760-2 | DDAH1 | Extracellular or secreted | 0.9435 | 0.543931026 | 29.1277 | 28.2826 | 27.2882 | 30.1799 | 29.0731 | 28.276 |
| N(G),N(G)-dimethylarginine dimethylaminohydrolase 2 | O95865 | DDAH2 | Extracellular or secreted | 0.176866667 | 0.150595376 | 27.0518 | 26.849 | 25.8669 | 27.0313 | 26.9587 | 26.3083 |
| DNA damage-binding protein 1 | Q16531 | DDB1 | Extracellular or secreted | -0.6471 | 0.679347774 | 33.3866 | 32.6774 | 32.3999 | 32.7412 | 32.1373 | 31.6441 |
| DNA damage-binding protein 2 | Q92466;Q92466-4;Q92466-5 | DDB2 | Other | -1.312433333 | 1.851335586 | 28.3155 | 28.1578 | 28.0147 | 26.7246 | 27.2593 | 26.5668 |
| Protein DDI1 homolog 2 | Q5TDH0;Q5TDH0-3;Q5TDH0 | DDI2 | Other | -0.1077 | 0.092675033 | 26.4385 | 25.8081 | 27.0538 | 26.0311 | 26.2701 | 26.6761 |
| Dolichyl-diphosphooligosaccharide--protein glycosyltran | P39656-3;P39656;P39656-2 | DDOST | Extracellular or secreted | 0.537433333 | 0.385596407 | 20.4688 | 21.5811 | 20.6043 | 21.8229 | 20.4961 | 21.9475 |
| Epithelial discoidin domain-containing receptor 1 | Q08345-2;Q08345-6;Q08345 | DDR1 | Extracellular or secreted | 0.1927 | 0.525032754 | 28.3311 | 28.1994 | 28.0605 | 28.3507 | 28.6399 | 28.1785 |
| D-dopachrome decarboxylase;D-dopachrome decarboxy | P30046;P30046-2;A6NHG4 | DDT;DDTL | Extracellular or secreted | -0.0214 | 0.017014528 | 30.3336 | 30.3752 | 29.6252 | 30.4712 | 30.379 | 29.4196 |
| ATP-dependent RNA helicase DDX1 | Q92499;Q92499-3;Q92499-2 | DDX1 | Other | -0.802966667 | 0.339906694 | 28.3585 | 28.8034 | 29.4786 | 26.5159 | 28.2279 | 29.4878 |
| Probable ATP-dependent RNA helicase DDX17 | Q92841;Q92841-1;Q92841-3 | DDX17 | Extracellular or secreted | -0.991433333 | 0.669127402 | 28.1544 | 28.9547 | 29.6907 | 27.3131 | 27.5774 | 28.935 |
| ATP-dependent RNA helicase DDX19B;ATP-dependent R | Q9UMR2-3;Q9NUU7-2;Q9U | DDX19B;DDX1 | Extracellular or secreted | -0.181633333 | 0.103436438 | 22.4555 | 22.3905 | 24.0435 | 22.492 | 22.4598 | 23.3928 |
| Probable ATP-dependent RNA helicase DDX20 | Q9UHI6 | DDX20 | Extracellular or secreted | -0.095 | 0.039234237 | 22.3591 | 20.3222 | 21.2067 | 22.1205 | 20.1477 | 21.3348 |
| Nucleolar RNA helicase 2 | Q9NR30;Q9NR30-2 | DDX21 | Other | -0.608766667 | 0.294840335 | 28.4359 | 27.7792 | 29.1238 | 29.2662 | 27.0769 | 27.1695 |
| Probable ATP-dependent RNA helicase DDX23 | Q9BUQ8 | DDX23 | Extracellular or secreted | -0.9945 | 0.471275538 | 25.1673 | 24.7293 | 27.3009 | 25.1758 | 24.2424 | 24.7958 |
| ATP-dependent RNA helicase DDX24 | Q9GZR7-2;Q9GZR7 | DDX24 | Other | -0.482033333 | 0.274154361 | 22.1213 | 20.8096 | 22.7069 | 22.2336 | 20.9273 | 21.0308 |
| Probable ATP-dependent RNA helicase DDX27 | Q96GQ7 | DDX27 | Other | -1.424733333 | 1.437133206 | 24.9892 | 25.7889 | 26.1253 | 24.762 | 23.6785 | 24.1887 |
| ATP-dependent RNA helicase DDX39A | O00148;O00148-2;O00148-3 | DDX39A | Other | -2.5892 | 0.680029301 | 26.0906 | 25.5558 | 28.2934 | 22.6454 | 22.6175 | 26.9093 |
| Spliceosome RNA helicase DDX39B | Q13838;Q13838-2 | DDX39B | Other | -1.038066667 | 0.754739946 | 31.1994 | 31.189 | 32.1541 | 29.5188 | 30.6257 | 31.2838 |
| ATP-dependent RNA helicase DDX3X;ATP-dependent RN | O00571-2;O00571;O15523-2 | DDX3X;DDX3Y | Extracellular or secreted | 0.411666667 | 0.279199331 | 25.3495 | 25.078 | 26.5931 | 25.4096 | 26.2042 | 26.6418 |
| ATP-dependent RNA helicase DDX42 | Q86XP3-2;Q86XP3 | DDX42 | Other | -1.207166667 | 0.47836901 | 26.6527 | 26.6667 | 28.3395 | 24.4243 | 26.0467 | 27.5664 |
| Probable ATP-dependent RNA helicase DDX46 | Q7L014 | DDX46 | Other | -1.0089 | 0.375059668 | 24.0519 | 23.6456 | 26.4274 | 22.9374 | 23.0353 | 25.1255 |
| Probable ATP-dependent RNA helicase DDX47 | Q9H0S4-2;Q9H0S4 | DDX47 | Other | -0.4819 | 0.704739196 | 21.9645 | 22.3958 | 22.2334 | 22.236 | 21.4511 | 21.4609 |
| Probable ATP-dependent RNA helicase DDX5 | P17844;P17844-2 | DDX5 | Extracellular or secreted | 0.018 | 0.012882094 | 25.4128 | 25.6989 | 26.7269 | 26.0718 | 25.549 | 26.2718 |
| ATP-dependent RNA helicase DDX51 | Q8N8A6 | DDX51 | Other | -1.0993 | 0.676580473 | 22.8911 | 21.682 | 23.7772 | 22.069 | 22.0232 | 20.9602 |
| Probable ATP-dependent RNA helicase DDX6 | P26196 | DDX6 | Other | -0.209933333 | 0.107294448 | 28.2191 | 27.8991 | 29.3538 | 27.4733 | 28.0408 | 29.3281 |
| 2,4-dienoyl-CoA reductase, mitochondrial | Q16698-2;Q16698 | DECR1 | Extracellular or secreted | 0.559566667 | 0.217870239 | 21.8835 | 22.5431 | 24.9021 | 23.8373 | 23.8262 | 23.3439 |
| Protein DEK | P35659;P35659-2 | DEK | Other | -0.1438 | 0.134917552 | 29.1217 | 29.2154 | 29.6213 | 28.6277 | 29.0765 | 29.8228 |
| DENN domain-containing protein 4C | Q5VZ89-5;Q5VZ89-3;Q5VZ89 | DENND4C | Extracellular or secreted | -0.176633333 | 0.101994989 | 22.6244 | 22.2574 | 23.6442 | 21.8278 | 22.7403 | 23.428 |
| Density-regulated protein | O43583 | DENR | Other | 0.010133333 | 0.176072575 | 27.0556 | 27.0512 | 26.9983 | 27.0237 | 27.0605 | 27.0513 |
| Deoxyribose-phosphate aldolase | Q9Y315 | DERA | Extracellular or secreted | -0.803366667 | 0.415924844 | 26.6718 | 26.674 | 27.6484 | 24.7769 | 26.9539 | 26.8533 |
| DNA fragmentation factor subunit alpha | O00273;O00273-2 | DFFA | Other | 0.5802 | 0.585997394 | 25.0862 | 24.9981 | 26.0398 | 26.097 | 26.3663 | 25.4014 |
| Non-syndromic hearing impairment protein 5 | O60443;O60443-3 | DFNA5 | Extracellular or secreted | -0.4096 | 0.219881731 | 20.893 | 20.9109 | 22.2342 | 22.0729 | 20.4406 | 20.2958 |
| Deoxyguanosine kinase, mitochondrial | Q16854-2;Q16854;Q16854-4 | DGUOK | Other | -1.783366667 | 0.867309638 | 22.8487 | 23.0834 | 25.1591 | 21.9136 | 21.8888 | 21.9387 |
| Delta(24)-sterol reductase | Q15392-2;Q15392 | DHCR24 | Other | 2.586033333 | 1.747170879 | 21.5592 | 20.7712 | 22.2142 | 24.2662 | 23.8466 | 24.1899 |
| 7-dehydrocholesterol reductase | Q9UBM7 | DHCR7 | Other | 0.616233333 | 0.26118908 | 20.8133 | 22.5735 | 22.2388 | 23.4884 | 22.9846 | 21.0013 |
| Dihydrofolate reductase | P00374-2;P00374 | DHFR | Other | 0.627966667 | 0.389428225 | 23.0025 | 22.5511 | 21.8714 | 24.2419 | 22.4301 | 22.6369 |
| Deoxyhypusine synthase | P49366;P49366-2;P49366-3 | DHPS | Other | -0.274333333 | 0.651337877 | 29.2103 | 28.9716 | 28.6819 | 28.5252 | 28.6304 | 28.8852 |
| Dehydrogenase/reductase SDR family member 7 | Q9Y394-2;Q9Y394 | DHRS7 | Extracellular or secreted | 2.5877 | 1.229527294 | 23.0858 | 21.7873 | 20.4606 | 24.857 | 23.7353 | 24.5045 |
| Probable 2-oxoglutarate dehydrogenase E1 component | Q96HY7 | DHTKD1 | Other | -2.139633333 | 0.378684122 | 21.2094 | 20.3218 | 27.0617 | 21.0465 | 20.5228 | 20.6047 |
| Pre-mRNA-splicing factor ATP-dependent RNA helicase D | O43143 | DHX15 | Other | -1.378433333 | 1.131841887 | 28.3864 | 28.2037 | 29.5834 | 26.774 | 27.232 | 28.0322 |
| Putative pre-mRNA-splicing factor ATP-dependent RNA h | O60231 | DHX16 | Other | 1.566266667 | 0.403738808 | 20.2276 | 19.5424 | 22.3808 | 23.875 | 19.5738 | 23.4008 |
| ATP-dependent RNA helicase DHX36 | Q9H2U1-3;Q9H2U1-2;Q9H2 | DHX36 | Extracellular or secreted | -0.3386 | 0.188469266 | 22.8733 | 24.221 | 24.9129 | 23.1041 | 23.7791 | 24.1082 |
| Pre-mRNA-splicing factor ATP-dependent RNA helicase P | Q92620;Q92620-2 | DHX38 | Other | -0.209466667 | 0.087185452 | 22.6207 | 22.4185 | 24.3173 | 24.0869 | 22.5566 | 22.0846 |
| ATP-dependent RNA helicase DHX8 | Q14562 | DHX8 | Extracellular or secreted | 0.962833333 | 0.537606396 | 21.4738 | 23.433 | 23.5816 | 24.104 | 23.7497 | 23.5232 |
| ATP-dependent RNA helicase A | Q08211 | DHX9 | Other | -0.713933333 | 0.342966115 | 29.6737 | 29.7435 | 31.7663 | 28.872 | 29.5501 | 30.6196 |
| Diablo homolog, mitochondrial | Q9NR28-2;Q9NR28;Q9NR28 | DIABLO | Extracellular or secreted | -0.785833333 | 0.405609095 | 26.3879 | 25.4049 | 27.2226 | 24.335 | 26.4269 | 25.896 |
| Protein diaphanous homolog 1 | O60610-2;O60610-3;O60610 | DIAPH1 | Extracellular or secreted | -1.070433333 | 0.676928587 | 28.1948 | 28.1964 | 28.6033 | 26.1218 | 27.5236 | 28.1378 |
| Death-inducer obliterator 1 | Q9BTC0 | DIDO1 | Other | -1.4135 | 0.570029343 | 23.6035 | 24.3262 | 24.8955 | 20.9813 | 23.8327 | 23.7707 |
| Digestive organ expansion factor homolog | Q68CQ4 | DIEXF | Other | -0.141066667 | 0.109285218 | 22.1423 | 21.753 | 21.8194 | 22.0856 | 20.9155 | 22.2904 |
| Disco-interacting protein 2 homolog B | Q9P265 | DIP2B | Extracellular or secreted | 0.774433333 | 2.613833743 | 22.8094 | 23.0352 | 23.0595 | 23.8411 | 23.6819 | 23.7044 |
| Exosome complex exonuclease RRP44 | Q9Y2L1-2;Q9Y2L1 | DIS3 | Other | -1.420766667 | 0.559114429 | 24.3736 | 23.9972 | 26.5159 | 22.9848 | 22.5039 | 25.1357 |
| DIS3-like exonuclease 2 | Q8IYB7-3;Q8IYB7-2;Q8IYB7; | DIS3L2 | Other | -1.528833333 | 1.328756552 | 23.1037 | 21.8248 | 22.9623 | 21.7499 | 20.9937 | 20.5607 |
| H/ACA ribonucleoprotein complex subunit 4 | O60832;O60832-2 | DKC1 | Other | -0.593233333 | 0.355245385 | 28.1861 | 26.7366 | 28.1954 | 27.995 | 27.071 | 26.2724 |
| Dickkopf-related protein 1 | O94907 | DKK1 | Extracellular or secreted | 1.5388 | 1.380164732 | 25.8448 | 26.2297 | 25.0684 | 27.1993 | 27.3829 | 27.1771 |
| Dihydrolipoyllysine-residue acetyltransferase componen | P10515 | DLAT | Other | -0.433433333 | 0.383940199 | 25.7656 | 24.9104 | 26.3391 | 25.3655 | 25.4423 | 24.907 |
| Dihydrolipoyl dehydrogenase, mitochondrial | P09622;P09622-2;P09622-3 | DLD | Other | -0.539433333 | 0.335391824 | 32.4562 | 31.5475 | 31.8653 | 32.3709 | 31.5019 | 30.3779 |
| Disks large homolog 1 | Q12959-5;Q12959-3;Q12959 | DLG1 | Extracellular or secreted | 0.384766667 | 0.555840689 | 27.0511 | 26.4818 | 27.1737 | 26.8504 | 27.4504 | 27.5601 |
| Disks large-associated protein 5 | Q15398-3;Q15398-1;Q15398 | DLGAP5 | Other | -0.999166667 | 0.900971763 | 23.0628 | 23.7312 | 24.5133 | 22.4593 | 23.0844 | 22.7661 |
| Dihydrolipoyllysine-residue succinyltransferase compone | P36957;P36957-2 | DLST | Extracellular or secreted | 1.075733333 | 0.727237181 | 25.2429 | 24.8188 | 26.6251 | 27.2579 | 26.6749 | 25.9812 |
| Deleted in malignant brain tumors 1 protein | Q9UGM3-5;Q9UGM3-3;Q9U | DMBT1 | Extracellular or secreted | 3.7991 | 1.223085365 | 28.8881 | 28.9225 | 25.6304 | 30.9232 | 31.6128 | 32.3023 |
| DnaJ homolog subfamily A member 1 | P31689;P31689-2 | DNAJA1 | Extracellular or secreted | -0.3637 | 0.327230947 | 27.1969 | 26.9168 | 27.7598 | 26.3806 | 26.7551 | 27.6467 |
| DnaJ homolog subfamily A member 2 | O60884 | DNAJA2 | Extracellular or secreted | -0.521766667 | 0.315020224 | 26.0393 | 26.114 | 27.8483 | 25.8846 | 25.8707 | 26.681 |
| DnaJ homolog subfamily A member 3, mitochondrial | Q96EY1-2;Q96EY1;Q96EY1-3 | DNAJA3 | Extracellular or secreted | 0.117833333 | 0.114443715 | 24.4767 | 24.409 | 24.0174 | 24.6848 | 24.8141 | 23.7577 |
| DnaJ homolog subfamily B member 1 | P25685;P25685-2 | DNAJB1 | Extracellular or secreted | -0.268966667 | 0.134582842 | 25.1126 | 24.7575 | 26.644 | 24.5104 | 25.1346 | 26.0622 |
| DnaJ homolog subfamily B member 11 | Q9UBS4 | DNAJB11 | Extracellular or secreted | 0.420133333 | 0.43718809 | 26.6252 | 26.4814 | 27.5858 | 27.1308 | 27.1118 | 27.7102 |

| | | | | | | | | | | | |
|---|---|---|---|---|---|---|---|---|---|---|---|
| DnaJ homolog subfamily B member 12 | Q9NXW2;Q9NXW2-2 | DNAJB12 | Other | -0.6334 | 0.650237774 | 23.2536 | 22.9501 | 22.3184 | 21.8328 | 22.8847 | 21.9044 |
| DnaJ homolog subfamily C member 1 | Q96KC8 | DNAJC1 | Extracellular or secreted | -1.236966667 | 0.841075417 | 23.0123 | 21.9272 | 23.7709 | 22.4121 | 21.0223 | 21.5651 |
| DnaJ homolog subfamily C member 10 | Q8IXB1-2;Q8IXB1;Q8IXB1-3 | DNAJC10 | Other | -1.61 | 1.678987155 | 25.7649 | 26.2675 | 26.9124 | 24.211 | 25.1048 | 24.799 |
| DnaJ homolog subfamily C member 2;DnaJ homolog subf | Q99543;Q99543-2 | DNAJC2 | Other | -1.732333333 | 0.97720608 | 26.0096 | 25.288 | 25.8655 | 24.1428 | 25.034 | 22.7893 |
| DnaJ homolog subfamily C member 5 | Q9H3Z4-2;Q9H3Z4 | DNAJC5 | Extracellular or secreted | 0.971633333 | 1.193978518 | 24.415 | 24.2329 | 24.9085 | 25.8695 | 25.7074 | 24.8944 |
| DnaJ homolog subfamily C member 8 | O75937 | DNAJC8 | Extracellular or secreted | -0.172633333 | 0.264262727 | 29.775 | 29.8753 | 29.3652 | 29.4567 | 29.8767 | 29.1642 |
| DnaJ homolog subfamily C member 9 | Q8WXX5 | DNAJC9 | Extracellular or secreted | -0.830166667 | 2.055625772 | 28.0932 | 28.1763 | 27.8333 | 27.2845 | 27.3804 | 26.9474 |
| Deoxyribonuclease-2-alpha | O00115-2;O00115 | DNASE2 | Extracellular or secreted | 0.766533333 | 0.951545332 | 30.7063 | 30.2382 | 29.652 | 31.1692 | 31.0699 | 30.657 |
| Dynamin-1-like protein | O00429-2;O00429-4;O00429 | DNM1L | Other | -1.699933333 | 0.926192915 | 27.7906 | 28.0037 | 29.3386 | 25.4145 | 26.8785 | 27.7401 |
| Dynamin-2 | P50570-5;P50570;P50570-2; | DNM2 | Extracellular or secreted | -0.616466667 | 0.224885531 | 25.6876 | 25.8173 | 27.5875 | 24.3896 | 25.4936 | 27.3598 |
| DNA (cytosine-5)-methyltransferase 1 | P26358;P26358-2 | DNMT1 | Other | -0.782233333 | 0.544924438 | 28.0204 | 26.7991 | 27.5192 | 27.5452 | 25.7719 | 26.6749 |
| Aspartyl aminopeptidase | Q9ULA0 | DNPEP | Extracellular or secreted | -0.3096 | 0.292504651 | 29.1129 | 29.3892 | 28.87 | 29.5015 | 28.7471 | 28.1947 |
| 2-deoxynucleoside 5-phosphate N-hydrolase 1 | O43598;O43598-2 | DNPH1 | Extracellular or secreted | -0.197233333 | 0.110612441 | 26.5527 | 27.071 | 27.4519 | 25.6872 | 27.3725 | 27.4242 |
| Deoxynucleotidyltransferase terminal-interacting protein | Q5QJE6 | DNTTIP2 | Other | -0.305566667 | 0.144245301 | 23.0029 | 21.2886 | 23.0273 | 23.2037 | 21.6433 | 21.5551 |
| Dedicator of cytokinesis protein 7 | Q96N67-4;Q96N67-3;Q96N6 | DOCK7 | Other | -0.230066667 | 0.074272742 | 23.6138 | 20.6524 | 23.7938 | 22.5642 | 22.2208 | 22.5848 |
| Deoxyhypusine hydroxylase | Q9BU89 | DOHH | Other | 1.4886 | 2.93992287 | 23.8311 | 24.1861 | 23.7579 | 25.6026 | 25.4347 | 25.2036 |
| Zinc finger protein ubi-d4 | Q92785 | DPF2 | Other | -1.4275 | 0.558459488 | 25.5698 | 26.0293 | 26.8391 | 22.857 | 25.182 | 26.1167 |
| Diphthamide biosynthesis protein 2 | Q9BQC3-2;Q9BQC3 | DPH2 | Other | -1.4689 | 1.841901428 | 22.4081 | 22.2859 | 22.2546 | 20.4459 | 21.0437 | 21.0523 |
| Diphthine synthase | Q9H2P9-3;Q9H2P9-2;Q9H2P | DPH5 | Other | -1.466833333 | 1.55859227 | 22.7072 | 23.7579 | 23.6497 | 21.6207 | 22.3965 | 21.6971 |
| Dipeptidyl peptidase 3 | Q9NY33;Q9NY33-4 | DPP3 | Extracellular or secreted | 0.379 | 0.236740244 | 32.9342 | 32.2792 | 31.5699 | 33.4917 | 32.6285 | 31.8001 |
| Dipeptidyl peptidase 4;Dipeptidyl peptidase 4 membrane | P27487 | DPP4 | Extracellular or secreted | 1.269233333 | 1.083885488 | 27.444 | 28.3481 | 26.9258 | 28.7489 | 29.0833 | 28.6934 |
| Dipeptidyl peptidase 2 | Q9UHL4 | DPP7 | Extracellular or secreted | 0.060566667 | 0.084360186 | 28.3475 | 27.9322 | 28.0201 | 28.2626 | 28.4733 | 27.7456 |
| Dipeptidyl peptidase 9 | Q86TI2;Q86TI2-4;Q86TI2-2 | DPP9 | Other | -0.249533333 | 0.345981504 | 27.5735 | 28.1153 | 27.5102 | 27.0526 | 27.5699 | 27.8279 |
| Protein dpy-30 homolog | Q9C005 | DPY30 | Other | 0.5231 | 0.513446942 | 24.3956 | 25.5754 | 24.7161 | 25.7432 | 24.889 | 25.6242 |
| Dihydropyrimidinase-related protein 2 | Q16555;Q16555-2 | DPYSL2 | Extracellular or secreted | -0.2734 | 0.300630597 | 30.4659 | 30.2695 | 31.0423 | 29.7918 | 30.392 | 30.7737 |
| Protein Dr1 | Q01658 | DR1 | Other | -0.173366667 | 0.12482165 | 25.9173 | 25.8556 | 25.9758 | 25.236 | 25.3013 | 26.6913 |
| Dr1-associated corepressor | Q14919;Q14919-2 | DRAP1 | Other | 0.3492 | 0.080222437 | 20.6561 | 24.0188 | 25.2128 | 22.5887 | 23.6545 | 24.6921 |
| Developmentally-regulated GTP-binding protein 1 | Q9Y295 | DRG1 | Other | -1.1526 | 1.2052141 | 26.7243 | 26.8313 | 27.7021 | 25.2974 | 26.1327 | 26.3698 |
| Desmocollin-1 | Q08554-2;Q08554 | DSC1 | Extracellular or secreted | 1.960566667 | 0.54007111 | 26.8844 | 23.4392 | 27.7984 | 29.4685 | 26.7031 | 27.8321 |
| Desmocollin-2 | Q02487-2;Q02487 | DSC2 | Extracellular or secreted | 0.245066667 | 0.129553635 | 28.412 | 29.2232 | 27.3235 | 29.3018 | 28.5388 | 27.8533 |
| Desmocollin-3 | Q14574-2;Q14574 | DSC3 | Extracellular or secreted | 0.214466667 | 0.089909069 | 27.8913 | 28.9158 | 26.2387 | 28.2915 | 27.8675 | 27.5302 |
| Desmoglein-1 | Q02413 | DSG1 | Extracellular or secreted | 2.428166667 | 0.661528182 | 26.9374 | 23.2883 | 27.8339 | 29.7088 | 27.2974 | 28.3379 |
| Desmoglein-2 | Q14126 | DSG2 | Extracellular or secreted | 0.6099 | 1.002534957 | 30.2227 | 30.3052 | 29.6498 | 30.9859 | 30.7142 | 30.3073 |
| Desmoglein-3 | P32926 | DSG3 | Extracellular or secreted | -0.507533333 | 0.336460277 | 25.7895 | 26.4089 | 25.7942 | 26.4973 | 25.3677 | 24.605 |
| Desmoplakin | P15924;P15924-3;P15924-2 | DSP | Extracellular or secreted | 0.214566667 | 0.146169001 | 29.1173 | 29.6903 | 30.2738 | 30.5113 | 29.079 | 30.1348 |
| Destrin | P60981;P60981-2 | DSTN | Extracellular or secreted | -0.051833333 | 0.100677427 | 29.8863 | 30.1778 | 30.1565 | 29.7119 | 30.2114 | 30.1418 |
| D-tyrosyl-tRNA(Tyr) deacylase 1 | Q8TEA8 | DTD1 | Other | -0.351033333 | 0.485315195 | 27.494 | 27.0156 | 26.8815 | 26.9967 | 27.0597 | 26.2816 |
| Thymidylate kinase | P23919-2;P23919 | DTYMK | Other | -0.471066667 | 0.392291794 | 25.398 | 24.5958 | 26.155 | 24.9799 | 24.9039 | 24.8518 |
| Deoxyuridine 5-triphosphate nucleotidohydrolase, mitoc | P33316-2;P33316 | DUT | Extracellular or secreted | -0.5194 | 0.630144246 | 29.6238 | 28.8772 | 28.5695 | 28.7372 | 28.5206 | 28.2545 |
| Decapping and exoribonuclease protein | O77932 | DXO | Extracellular or secreted | -1.1975 | 0.524264897 | 24.7705 | 23.892 | 23.7035 | 23.9699 | 23.5917 | 21.2119 |
| Cytoplasmic dynein 1 heavy chain 1 | Q14204 | DYNC1H1 | Extracellular or secreted | -0.219766667 | 0.105714397 | 29.7189 | 29.9117 | 31.5327 | 29.4707 | 29.9503 | 31.083 |
| Cytoplasmic dynein 1 intermediate chain 2 | Q13409-3;Q13409-2;Q13409 | DYNC1I2 | Other | -0.051366667 | 0.046415793 | 27.1956 | 27.3628 | 28.3298 | 27.6953 | 27.5546 | 27.4842 |
| Cytoplasmic dynein 1 light intermediate chain 1 | Q9Y6G9 | DYNC1LI1 | Extracellular or secreted | -0.992266667 | 0.186477806 | 21.6504 | 21.9936 | 27.1618 | 20.9933 | 23.0293 | 23.8064 |
| Cytoplasmic dynein 1 light intermediate chain 2 | O43237;O43237-2 | DYNC1LI2 | Other | 0.461066667 | 0.161944356 | 23.9174 | 24.4114 | 26.5639 | 24.1673 | 25.5492 | 26.5594 |
| Dynein light chain 1, cytoplasmic | P63167 | DYNLL1 | Extracellular or secreted | -0.044533333 | 0.038021226 | 27.6814 | 28.3373 | 28.6003 | 27.5851 | 28.4658 | 28.4345 |
| Dynein light chain 2, cytoplasmic | Q96FJ2 | DYNLL2 | Extracellular or secreted | -0.928766667 | 0.505542556 | 25.2182 | 25.6252 | 25.4175 | 23.1019 | 25.1652 | 25.2075 |
| Dynein light chain roadblock-type 1 | Q9NP97 | DYNLRB1 | Other | 0.5533 | 0.6938383 | 26.3878 | 27.0799 | 26.5992 | 27.0145 | 27.819 | 26.8933 |
| Dynein light chain Tctex-type 1 | P63172 | DYNLT1 | Extracellular or secreted | -0.1401 | 0.09695315 | 24.9235 | 24.5946 | 24.8692 | 23.6986 | 25.0816 | 25.1868 |
| Probable rRNA-processing protein EBP2 | Q99848 | EBNA1BP2 | Other | -1.243166667 | 0.662960998 | 24.8294 | 23.9381 | 24.1287 | 21.6317 | 23.868 | 23.667 |
| 3-beta-hydroxysteroid-Delta(8),Delta(7)-isomerase | Q15125 | EBP | Extracellular or secreted | 1.940133333 | 1.219749039 | 21.0635 | 22.5492 | 22.7986 | 24.3852 | 23.9592 | 23.8873 |
| Endothelin-converting enzyme 1 | P42892-3;P42892-2;P42892- | ECE1 | Extracellular or secreted | 0.073266667 | 0.108261352 | 23.9757 | 24.1635 | 23.4227 | 24.0696 | 23.8926 | 23.8195 |
| Delta(3,5)-Delta(2,4)-dienoyl-CoA isomerase, mitochond | Q13011 | ECH1 | Extracellular or secreted | -1.174166667 | 1.306699098 | 29.8913 | 29.5355 | 30.6537 | 28.9594 | 29.1279 | 28.4707 |
| Ethylmalonyl-CoA decarboxylase | Q9NTX5-3;Q9NTX5-6;Q9NTX | ECHDC1 | Extracellular or secreted | 1.763566667 | 0.555592128 | 23.7722 | 20.5097 | 24.3973 | 24.1284 | 24.4403 | 25.4012 |
| Enoyl-CoA hydratase domain-containing protein 3, mitoc | Q96DC8;Q96DC8-2 | ECHDC3 | Other | -2.948033333 | 1.222682378 | 24.8718 | 25.5826 | 26.8974 | 22.9033 | 21.2579 | 24.3465 |
| Enoyl-CoA hydratase, mitochondrial | P30084 | ECHS1 | Extracellular or secreted | -1.230733333 | 1.019114632 | 27.7046 | 28.4455 | 29.2751 | 27.0791 | 27.6765 | 26.9774 |
| Enoyl-CoA delta isomerase 1, mitochondrial | P42126;P42126-2 | ECI1 | Extracellular or secreted | -1.1828 | 0.977235439 | 28.8289 | 28.4404 | 29.1249 | 28.0405 | 28.0978 | 26.7075 |
| Extracellular matrix protein 1 | Q16610;Q16610-4;Q16610-2 | ECM1 | Extracellular or secreted | 0.8319 | 0.456493253 | 28.6644 | 29.5471 | 27.2431 | 29.7319 | 29.559 | 28.6594 |
| Proteasome-associated protein ECM29 homolog | Q5VYK3 | ECM29 | Other | -0.888533333 | 0.465903521 | 27.2322 | 28.4273 | 28.207 | 25.7926 | 27.2037 | 28.2046 |
| Enhancer of mRNA-decapping protein 4 | Q6P2E9;Q6P2E9-2 | EDC4 | Other | -1.753833333 | 0.65254044 | 24.2389 | 23.4658 | 26.313 | 22.023 | 22.077 | 24.6562 |
| Endothelial differentiation-related factor 1 | O60869-2;O60869-3;O60869 | EDF1 | Extracellular or secreted | 0.614933333 | 0.58532411 | 27.4095 | 27.2621 | 26.1621 | 27.9393 | 27.5113 | 27.2279 |
| Early endosome antigen 1 | Q15075 | EEA1 | Extracellular or secreted | -0.824833333 | 0.991906892 | 28.3094 | 27.6828 | 28.2159 | 26.6359 | 27.4235 | 27.6742 |
| Elongation factor 1-alpha 1;Putative elongation factor 1- | P68104;Q5VTE0;P68104-2 | EEF1A1;EEF1A1 | Extracellular or secreted | -0.399933333 | 0.270988517 | 33.1306 | 33.6245 | 33.8752 | 32.2335 | 33.1576 | 34.0394 |
| Elongation factor 1-alpha 2 | Q05639 | EEF1A2 | Other | -1.708033333 | 1.635885009 | 28.2715 | 29.5226 | 28.9984 | 26.7714 | 27.7674 | 27.1296 |
| Elongation factor 1-beta | P24534 | EEF1B2 | Other | 0.761933333 | 1.552043596 | 29.3279 | 29.3111 | 29.4459 | 29.8724 | 30.3845 | 30.1138 |
| Elongation factor 1-delta | P29692;P29692-2 | EEF1D | Other | 0.171233333 | 0.276377403 | 31.3854 | 31.3352 | 31.1938 | 31.0402 | 31.7881 | 31.5998 |
| Elongation factor 1-delta | P29692-3 | EEF1D | Other | -1.740166667 | 0.432089337 | 25.0201 | 25.5008 | 25.1673 | 20.5299 | 25.5414 | 24.3964 |
| Eukaryotic translation elongation factor 1 epsilon-1 | O43324;O43324-2 | EEF1E1 | Extracellular or secreted | -0.945066667 | 0.847764148 | 26.8334 | 26.4801 | 27.8913 | 25.7154 | 26.2526 | 26.4016 |
| Elongation factor 1-gamma | P26641;P26641-2 | EEF1G | Extracellular or secreted | -1.0082 | 1.535897023 | 33.104 | 32.6184 | 32.8047 | 31.6168 | 31.5863 | 32.2994 |
| Elongation factor 2 | P13639 | EEF2 | Extracellular or secreted | -0.8541 | 0.822107131 | 33.0238 | 33.7518 | 33.5477 | 31.8678 | 32.667 | 33.2262 |
| EGF-containing fibulin-like extracellular matrix protein 1 | Q12805-2;Q12805-4;Q12805 | EFEMP1 | Extracellular or secreted | -1.654533333 | 1.496323643 | 29.856 | 30.8958 | 29.6802 | 27.8224 | 28.7166 | 28.9294 |
| EGF-containing fibulin-like extracellular matrix protein 1 | Q12805-5 | EFEMP1 | Extracellular or secreted | 2.106366667 | 0.727895258 | 21.349 | 21.4657 | 24.6513 | 24.1974 | 24.9167 | 24.671 |
| EF-hand domain-containing protein D2 | Q96C19 | EFHD2 | Other | -0.0461 | 0.084686789 | 30.1634 | 30.5987 | 30.0366 | 30.1019 | 30.3712 | 30.1873 |
| Ephrin-A1;Ephrin-A1, secreted form | P20827;P20827-2 | EFNA1 | Extracellular or secreted | 0.598833333 | 0.283614649 | 27.8739 | 29.2752 | 26.7406 | 28.579 | 29.2086 | 27.8986 |
| Ephrin-A5 | P52803 | EFNA5 | Extracellular or secreted | -0.068066667 | 0.036859147 | 26.0449 | 26.1715 | 24.3485 | 25.5796 | 25.3846 | 25.3965 |
| Ephrin-B1 | P98172 | EFNB1 | Extracellular or secreted | -0.2118 | 0.105000358 | 27.1561 | 27.0729 | 25.3535 | 26.6721 | 26.7949 | 25.4801 |
| Ephrin-B2 | P52799 | EFNB2 | Extracellular or secreted | 1.1962 | 0.553719881 | 26.6497 | 27.3347 | 24.6909 | 28.0189 | 27.7684 | 26.4766 |
| Elongation factor Tu GTP-binding domain-containing pro | Q7Z2Z2-2;Q7Z2Z2 | EFTUD1 | Other | -0.2177 | 0.062508165 | 23.1017 | 23.8758 | 25.0859 | 21.9184 | 23.9873 | 25.5046 |
| 116 kDa U5 small nuclear ribonucleoprotein component | Q15029;Q15029-3;Q15029-2 | EFTUD2 | Other | -0.379266667 | 0.361534487 | 28.9793 | 28.7195 | 29.9168 | 28.5437 | 28.66 | 29.2741 |
| Epidermal growth factor receptor | P00533;P00533-4;P00533-3; | EGFR | Extracellular or secreted | -0.889233333 | 0.653395659 | 28.4612 | 27.9118 | 27.3994 | 27.7272 | 27.3288 | 26.0487 |
| Egl nine homolog 1 | Q9GZT9-3;Q9GZT9-2;Q9GZT | EGLN1 | Other | -0.785066667 | 0.488545312 | 20.9488 | 21.8075 | 21.5306 | 21.8064 | 19.7894 | 20.3359 |
| EH domain-binding protein 1-like protein 1 | Q8N3D4 | EHBP1L1 | Other | 0.2835 | 0.14665903 | 23.8477 | 23.4339 | 24.3987 | 23.2 | 23.9564 | 25.3744 |
| EH domain-containing protein 1 | Q9H4M9 | EHD1 | Extracellular or secreted | 1.149166667 | 2.022999861 | 27.8373 | 28.1564 | 28.4614 | 29.5181 | 29.1921 | 29.1924 |
| EH domain-containing protein 2 | Q9NZN4 | EHD2 | Extracellular or secreted | -1.734233333 | 1.460534158 | 26.6151 | 27.4947 | 28.0359 | 25.1963 | 26.3478 | 25.3989 |
| EH domain-containing protein 4 | Q9H223 | EHD4 | Extracellular or secreted | 0.6632 | 1.542124542 | 25.2716 | 25.6435 | 25.6922 | 26.4347 | 26.2304 | 25.9318 |
| Histone-lysine N-methyltransferase EHMT2 | Q96KQ7-2;Q96KQ7;Q96KQ7 | EHMT2 | Extracellular or secreted | 0.150666667 | 0.192372922 | 22.2979 | 22.1016 | 22.4844 | 21.9191 | 22.7859 | 22.6309 |
| Eukaryotic translation initiation factor 1;Eukaryotic trans | P41567;O60739 | EIF1;EIF1B | Other | 0.829733333 | 0.685887071 | 23.942 | 24.9543 | 23.5262 | 24.5499 | 25.6517 | 24.7101 |
| Probable RNA-binding protein EIF1AD | Q8N9N8 | EIF1AD | Other | 2.347433333 | 1.240587105 | 22.3222 | 20.2186 | 22.3179 | 24.041 | 24.677 | 23.183 |
| Eukaryotic translation initiation factor 1A, X-chromosom | P47813;O14602 | EIF1AX;EIF1AY | Other | -0.499766667 | 0.587324523 | 26.6706 | 27.221 | 27.2263 | 26.4923 | 26.0188 | 27.1075 |
| Eukaryotic translation initiation factor 2A;Eukaryotic tran | Q9BY44-3;Q9BY44;Q9BY44-4 | EIF2A | Extracellular or secreted | -0.0541 | 0.02855488 | 26.8657 | 27.2898 | 28.2889 | 26.7095 | 27.2446 | 28.328 |
| Interferon-induced, double-stranded RNA-activated prot | P19525-2;P19525 | EIF2AK2 | Other | 1.246833333 | 0.574437563 | 21.5644 | 20.4334 | 23.2071 | 22.0812 | 23.5735 | 23.2907 |
| Translation initiation factor eIF-2B subunit alpha | Q14232;Q14232-2 | EIF2B1 | Extracellular or secreted | -0.2457 | 0.152293959 | 24.7762 | 25.6528 | 25.6033 | 24.1081 | 25.3356 | 25.8515 |
| Translation initiation factor eIF-2B subunit beta | P49770 | EIF2B2 | Other | -0.472966667 | 0.239836523 | 24.7588 | 24.6616 | 26.0841 | 23.5644 | 24.8189 | 25.7023 |
| Translation initiation factor eIF-2B subunit gamma | Q9NR50-3;Q9NR50-2;Q9NR5 | EIF2B3 | Other | -0.827066667 | 0.525687783 | 24.6947 | 24.8638 | 26.4364 | 23.865 | 24.4198 | 25.2289 |
| Translation initiation factor eIF-2B subunit delta | Q9UI10;Q9UI10-3;Q9UI10-2 | EIF2B4 | Other | -1.263966667 | 0.690050312 | 23.7354 | 23.9795 | 25.5169 | 21.9889 | 23.3519 | 24.0991 |
| Translation initiation factor eIF-2B subunit epsilon | Q13144 | EIF2B5 | Other | -1.187066667 | 0.517068323 | 24.9756 | 24.244 | 25.661 | 22.2149 | 23.9299 | 25.1746 |
| Eukaryotic translation initiation factor 2D | P41214;P41214-2 | EIF2D | Other | -0.442233333 | 0.182472516 | 22.6844 | 21.2193 | 22.8372 | 23.0417 | 21.9352 | 20.4373 |
| Eukaryotic translation initiation factor 2 subunit 1 | P05198 | EIF2S1 | Extracellular or secreted | -0.762866667 | 0.542390671 | 29.897 | 30.5214 | 30.6733 | 28.6452 | 29.684 | 30.4739 |
| Eukaryotic translation initiation factor 2 subunit 2 | P20042 | EIF2S2 | Other | 0.4129 | 0.258690667 | 29.9894 | 28.7599 | 30.3596 | 30.9056 | 29.5307 | 29.9113 |
| Eukaryotic translation initiation factor 2 subunit 3;Putati | P41091;Q2VIR3;Q2VIR3-2 | EIF2S3;EIF2S3L | Extracellular or secreted | -0.578366667 | 0.374286585 | 30.071 | 30.8144 | 31.2602 | 29.1796 | 30.2087 | 31.0222 |
| Eukaryotic translation initiation factor 3 subunit A | Q14152;Q14152-2 | EIF3A | Other | -0.3579 | 0.414520561 | 30.4766 | 30.4775 | 31.0923 | 29.8945 | 30.1816 | 30.8966 |
| Eukaryotic translation initiation factor 3 subunit B | P55884;P55884-2 | EIF3B | Extracellular or secreted | -0.5807 | 2.064283806 | 30.4985 | 30.395 | 30.6816 | 29.9631 | 29.7852 | 30.0847 |
| Eukaryotic translation initiation factor 3 subunit C;Eukary | Q99613;Q99613-2;B5ME19 | EIF3C;EIF3CL | Other | -0.491266667 | 0.599350137 | 29.5338 | 29.5017 | 30.0678 | 28.8235 | 28.9995 | 29.8065 |
| Eukaryotic translation initiation factor 3 subunit D | O15371-2;O15371-3;O15371 | EIF3D | Other | -0.538866667 | 0.530520319 | 28.4311 | 29.1013 | 29.3023 | 27.9259 | 28.1937 | 29.0985 |
| Eukaryotic translation initiation factor 3 subunit E | P60228 | EIF3E | Extracellular or secreted | -0.391266667 | 0.312436833 | 28.3989 | 28.3852 | 29.4393 | 27.8568 | 28.11 | 29.0828 |
| Eukaryotic translation initiation factor 3 subunit F | O00303 | EIF3F | Other | -0.4126 | 0.4160764 | 28.6452 | 28.7293 | 28.8205 | 27.6921 | 28.2833 | 28.9818 |
| Eukaryotic translation initiation factor 3 subunit G | O75821 | EIF3G | Other | 0.1081 | 0.23196586 | 28.6455 | 28.4176 | 28.6268 | 28.5162 | 28.5056 | 28.9924 |
| Eukaryotic translation initiation factor 3 subunit H | O15372 | EIF3H | Extracellular or secreted | -0.616533333 | 0.651340312 | 28.203 | 28.3077 | 28.5458 | 27.3617 | 27.3913 | 28.4539 |
| Eukaryotic translation initiation factor 3 subunit I | Q13347 | EIF3I | Extracellular or secreted | -0.094966667 | 0.113407963 | 29.2521 | 28.4715 | 29.1859 | 28.6991 | 28.7158 | 29.2097 |
| Eukaryotic translation initiation factor 3 subunit J | O75822;O75822-2;O75822-3 | EIF3J | Other | 0.098233333 | 0.141399609 | 28.8234 | 28.1386 | 28.8821 | 28.6904 | 28.788 | 28.6604 |
| Eukaryotic translation initiation factor 3 subunit K | Q9UBQ5-2;Q9UBQ5 | EIF3K | Extracellular or secreted | -0.247766667 | 0.323361737 | 27.5968 | 28.4405 | 28.2899 | 27.5319 | 27.971 | 28.081 |
| Eukaryotic translation initiation factor 3 subunit L | Q9Y262;Q9Y262-2 | EIF3L | Other | -0.421066667 | 0.518145692 | 28.6112 | 28.7724 | 29.5253 | 28.3247 | 28.3476 | 28.9734 |
| Eukaryotic translation initiation factor 3 subunit M | Q7L2H7;Q7L2H7-2 | EIF3M | Other | -2.021366667 | 0.702626836 | 28.0555 | 28.4531 | 28.6061 | 24.2187 | 27.1799 | 27.652 |
| Eukaryotic initiation factor 4A-I | P60842;P60842-2 | EIF4A1 | Extracellular or secreted | -0.345833333 | 0.197070084 | 30.3093 | 30.9568 | 31.6178 | 29.7426 | 30.47 | 31.6338 |
| Eukaryotic initiation factor 4A-II;Eukaryotic initiation fact | Q14240;Q14240-2 | EIF4A2 | Other | 0.022066667 | 0.018755617 | 27.1033 | 27.6255 | 27.3906 | 27.2434 | 26.8808 | 28.0614 |
| Eukaryotic initiation factor 4A-III;Eukaryotic initiation fac | P38919 | EIF4A3 | Other | -1.3258 | 0.351792288 | 27.4859 | 26.4351 | 29.2527 | 24.0509 | 26.6169 | 28.5285 |
| Eukaryotic translation initiation factor 4B | P23588;P23588-2 | EIF4B | Other | 0.770733333 | 0.711534727 | 29.6005 | 28.5853 | 29.9205 | 30.6005 | 30.1071 | 29.7109 |
| Eukaryotic translation initiation factor 4E | P06730;P06730-2;P06730-3 | EIF4E | Extracellular or secreted | -0.291066667 | 0.225873819 | 27.1318 | 27.545 | 28.1036 | 26.5164 | 27.4851 | 27.9057 |
| Eukaryotic translation initiation factor 4E type 2 | O60573;O60573-2 | EIF4E2 | Other | -0.714866667 | 0.225883337 | 23.5594 | 23.5527 | 23.7874 | 23.4842 | 20.7254 | 24.5453 |
| Eukaryotic translation initiation factor 4E-binding protein | Q13541 | EIF4EBP1 | Other | 2.251366667 | 0.525932158 | 20.5089 | 20.9726 | 21.5056 | 25.6904 | 23.8687 | 20.1821 |

| | | | | | | | | | | | |
|---|---|---|---|---|---|---|---|---|---|---|---|
| Eukaryotic translation initiation factor 4E-binding protein | Q13542 | EIF4EBP2 | Other | -0.901866667 | 0.584324847 | 22.7654 | 22.6924 | 21.8671 | 22.0045 | 22.2291 | 20.3857 |
| Eukaryotic translation initiation factor 4 gamma 1 | Q04637-4 | EIF4G1 | Other | -0.041033333 | 0.049095328 | 29.5112 | 29.2096 | 30.0128 | 29.5037 | 29.2754 | 29.8314 |
| Eukaryotic translation initiation factor 4 gamma 2 | P78344-2;P78344 | EIF4G2 | Other | -1.4625 | 1.520713966 | 25.8333 | 25.0237 | 25.7455 | 23.5813 | 23.9035 | 24.7302 |
| Eukaryotic translation initiation factor 4 gamma 3 | O43432-4;O43432;O43432-3 | EIF4G3 | Other | 0.5486 | 0.504662262 | 26.0154 | 25.5056 | 25.9598 | 26.8876 | 25.5661 | 26.6729 |
| Eukaryotic translation initiation factor 4H | Q15056-2;Q15056 | EIF4H | Other | 1.5326 | 1.132870457 | 26.2673 | 26.4542 | 27.8187 | 28.0137 | 28.4877 | 28.6366 |
| Eukaryotic translation initiation factor 5 | P55010 | EIF5 | Extracellular or secreted | -0.642733333 | 0.736914435 | 27.6972 | 28.6233 | 28.0652 | 27.0119 | 27.4191 | 28.0265 |
| Eukaryotic translation initiation factor 5A-1;Eukaryotic tr | P63241;P63241-2;Q6IS14 | EIF5A;EIF5AL1 | Extracellular or secreted | -0.469033333 | 1.647117884 | 32.2007 | 31.9994 | 32.2873 | 31.5007 | 31.7789 | 31.8007 |
| Eukaryotic translation initiation factor 5B | O60841 | EIF5B | Other | -0.0092 | 0.010564523 | 29.8143 | 29.4796 | 29.5124 | 30.0718 | 29.517 | 29.1899 |
| Eukaryotic translation initiation factor 6 | P56537;P56537-2 | EIF6 | Extracellular or secreted | -0.2698 | 1.338143341 | 30.3036 | 30.3468 | 30.3759 | 30.1619 | 29.9414 | 30.1136 |
| Zinc phosphodiesterase ELAC protein 2 | Q9BQ52-4;Q9BQ52;Q9BQ52 | ELAC2 | Other | -3.1251 | 0.846987429 | 26.1881 | 26.2855 | 26.3393 | 20.7493 | 23.364 | 25.3243 |
| ELAV-like protein 1 | Q15717;Q15717-2 | ELAVL1 | Other | -1.3849 | 0.811989972 | 29.3865 | 29.0762 | 30.227 | 27.0367 | 28.2178 | 29.2805 |
| Engulfment and cell motility protein 3 | Q96BJ8;Q96BJ8-3;Q96BJ8-2 | ELMO3 | Other | 0.5697 | 0.165751579 | 22.4703 | 21.4977 | 25.1781 | 23.92 | 22.3976 | 24.5376 |
| Elongator complex protein 2 | Q6IA86-2;Q6IA86-3;Q6IA86- | ELP2 | Extracellular or secreted | -0.262933333 | 0.271371514 | 23.9712 | 24.0647 | 24.075 | 23.0822 | 23.9906 | 24.2493 |
| Elongator complex protein 3 | Q9H9T3-4;Q9H9T3-5;Q9H9T | ELP3 | Other | -0.3206 | 0.320147532 | 23.343 | 23.4497 | 24.4748 | 23.7116 | 23.3922 | 23.2019 |
| Emerin | P50402 | EMD | Other | 0.094 | 0.038718321 | 24.8319 | 23.9344 | 26.5769 | 25.242 | 25.2609 | 25.1223 |
| Ribosomal RNA small subunit methyltransferase NEP1 | Q92979 | EMG1 | Other | -0.369933333 | 0.156541004 | 22.1414 | 21.9213 | 24.3834 | 23.1138 | 22.0884 | 22.1341 |
| Echinoderm microtubule-associated protein-like 2 | O95834;O95834-2;O95834-3 | EML2 | Other | -0.178533333 | 0.312952827 | 30.3607 | 30.5196 | 30.1591 | 30.0948 | 30.5453 | 29.8637 |
| Echinoderm microtubule-associated protein-like 3 | Q32P44;Q32P44-2 | EML3 | Other | -0.8449 | 1.676265953 | 25.5374 | 25.6462 | 26.0709 | 24.6519 | 24.8625 | 25.2054 |
| Echinoderm microtubule-associated protein-like 4 | Q9HC35;Q9HC35-2 | EML4 | Other | 0.065033333 | 0.120043218 | 28.9639 | 28.4141 | 28.9133 | 28.9647 | 28.6991 | 28.8226 |
| Protein enabled homolog | Q8N8S7-3;Q8N8S7-2;Q8N8S | ENAH | Extracellular or secreted | -0.577366667 | 0.910891834 | 28.736 | 28.4024 | 28.2893 | 27.7278 | 27.595 | 28.3728 |
| Endonuclease domain-containing 1 protein | O94919 | ENDOD1 | Extracellular or secreted | -0.546733333 | 0.267320883 | 24.3546 | 23.21 | 23.308 | 24.4827 | 22.4585 | 22.2912 |
| Endoglin | P17813-2;P17813 | ENG | Extracellular or secreted | 1.1591 | 0.739067238 | 21.3792 | 21.509 | 23.2335 | 23.7244 | 22.9351 | 22.9395 |
| Alpha-enolase | P06733 | ENO1 | Extracellular or secreted | 0.242966667 | 0.22559786 | 36.4218 | 36.2873 | 35.8876 | 37.1029 | 36.4066 | 35.8161 |
| Gamma-enolase | P09104;P09104-2 | ENO2 | Extracellular or secreted | 1.004266667 | 0.792364973 | 30.422 | 30.1758 | 29.4059 | 31.8577 | 30.9361 | 30.2227 |
| Beta-enolase | P13929;P13929-3;P13929-2 | ENO3 | Extracellular or secreted | 0.660466667 | 0.440853942 | 26.2129 | 25.8619 | 25.4488 | 27.6099 | 26.0673 | 25.8278 |
| Enolase-phosphatase E1 | Q9UHY7;Q9UHY7-2 | ENOPH1 | Extracellular or secreted | -0.0752 | 0.038496245 | 29.431 | 28.7157 | 27.7357 | 29.2357 | 28.7034 | 27.7177 |
| Alpha-endosulfine | O43768-2;O43768;O43768-9 | ENSA | Other | 0.122233333 | 0.072665047 | 25.8028 | 25.661 | 26.2593 | 26.9251 | 26.097 | 25.0677 |
| Ectonucleoside triphosphate diphosphohydrolase 3 | O75355-2;O75355 | ENTPD3 | Extracellular or secreted | 2.0735 | 1.285841384 | 20.7217 | 21.1983 | 21.8013 | 23.5139 | 24.2185 | 22.2094 |
| Transcription and mRNA export factor ENY2 | Q9NPA8-2;Q9NPA8 | ENY2 | Other | -0.7231 | 0.304059077 | 22.6194 | 23.3534 | 23.7201 | 22.8822 | 23.7695 | 20.8719 |
| Band 4.1-like protein 1 | Q9H4G0-2;Q9H4G0;Q9H4G0 | EPB41L1 | Extracellular or secreted | 0.502866667 | 0.215797921 | 24.8649 | 23.8135 | 26.3771 | 24.5905 | 25.6795 | 26.2941 |
| Band 4.1-like protein 2 | O43491;O43491-3;O43491-2 | EPB41L2 | Extracellular or secreted | 0.2941 | 0.300123119 | 28.2751 | 27.0905 | 27.9917 | 28.0524 | 27.9214 | 28.2658 |
| Epithelial cell adhesion molecule | P16422 | EPCAM | Extracellular or secreted | 1.583166667 | 1.490868464 | 29.3714 | 29.1236 | 28.2875 | 31.1851 | 30.4073 | 29.9396 |
| Mammalian ependymin-related protein 1 | Q9UM22-2;Q9UM22 | EPDR1 | Extracellular or secreted | 0.693566667 | 1.099600862 | 23.6112 | 22.9608 | 22.8657 | 24.014 | 23.5161 | 23.9883 |
| Ephrin type-A receptor 2 | P29317 | EPHA2 | Extracellular or secreted | 0.637766667 | 0.971197171 | 29.6509 | 29.2109 | 28.8724 | 30.2893 | 29.7598 | 29.5984 |
| Ephrin type-B receptor 2 | P29323-2;P29323-3;P29323 | EPHB2 | Extracellular or secreted | 0.902333333 | 0.569172947 | 26.2379 | 26.1123 | 24.4024 | 27.0987 | 26.3023 | 26.0586 |
| Ephrin type-B receptor 3 | P54753 | EPHB3 | Extracellular or secreted | 0.658833333 | 0.79817286 | 22.0294 | 23.1028 | 22.5493 | 23.3969 | 23.1704 | 23.0907 |
| Ephrin type-B receptor 4 | P54760;P54760-2;P54760-4 | EPHB4 | Extracellular or secreted | 0.492166667 | 0.390571083 | 26.754 | 27.374 | 25.7544 | 27.1165 | 27.2664 | 26.976 |
| Epoxide hydrolase 1 | P07099 | EPHX1 | Other | -0.2569 | 0.23443869 | 26.711 | 26.2238 | 27.5502 | 26.327 | 26.6338 | 26.7535 |
| Epsin-2 | O95208-2;O95208-3;O95208 | EPN2 | Other | 0.846766667 | 0.529886938 | 21.8256 | 22.6015 | 20.5345 | 22.4692 | 22.4789 | 22.5538 |
| Epiplakin | P58107 | EPPK1 | Other | -1.952 | 0.963867506 | 29.1479 | 29.2056 | 29.1319 | 28.5317 | 26.1359 | 26.9618 |
| Bifunctional glutamate/proline--tRNA ligase;Glutamate-- | P07814 | EPRS | Other | -0.482766667 | 0.464620689 | 31.2534 | 31.7299 | 32.3779 | 30.755 | 31.3336 | 31.8243 |
| Epidermal growth factor receptor substrate 15 | P42566 | EPS15 | Extracellular or secreted | -0.159833333 | 0.387479341 | 26.8904 | 26.6755 | 26.4149 | 26.4901 | 26.3259 | 26.6853 |
| Epidermal growth factor receptor substrate 15-like 1 | Q9UBC2-4;Q9UBC2-3;Q9UB( | EPS15L1 | Extracellular or secreted | -0.495666667 | 0.875936546 | 27.5229 | 26.8339 | 26.9754 | 26.5936 | 26.7318 | 26.5198 |
| Epidermal growth factor receptor kinase substrate 8 | Q12929;Q12929-2 | EPS8 | Extracellular or secreted | 0.895666667 | 1.421550311 | 24.0669 | 23.8827 | 23.5674 | 24.605 | 24.4204 | 25.1786 |
| Endoplasmic reticulum aminopeptidase 1 | Q9NZ08;Q9NZ08-2 | ERAP1 | Extracellular or secreted | -0.0411 | 0.063072541 | 26.2039 | 25.8646 | 26.0765 | 26.2687 | 26.1343 | 25.6187 |
| Endoplasmic reticulum aminopeptidase 2 | Q6P179-3;Q6P179 | ERAP2 | Extracellular or secreted | -1.036766667 | 1.383390172 | 23.078 | 23.4765 | 23.4509 | 21.7965 | 22.6987 | 22.3999 |
| Protein LAP2 | Q96RT1-7;Q96RT1-5;Q96RT1 | ERBB2IP | Extracellular or secreted | -0.876533333 | 0.335887525 | 23.0338 | 23.4896 | 23.2925 | 23.4727 | 20.4633 | 23.2503 |
| ELKS/Rab6-interacting/CAST family member 1 | Q8IUD2;Q8IUD2-2;Q8IUD2-3 | ERC1 | Extracellular or secreted | 0.828266667 | 0.687664728 | 25.2783 | 26.1407 | 26.8662 | 27.137 | 27.0776 | 26.5554 |
| Enhancer of rudimentary homolog | P84090 | ERH | Other | -0.149766667 | 0.251613325 | 30.5237 | 30.919 | 30.8249 | 30.6912 | 30.8994 | 30.2277 |
| ERI1 exoribonuclease 3 | O43414-3;O43414-2;O43414 | ERI3 | Other | -0.513666667 | 0.318612849 | 22.5904 | 24.3133 | 24.2716 | 23.7573 | 23.1237 | 22.7533 |
| ERO1-like protein alpha | Q96HE7 | ERO1L | Extracellular or secreted | -0.116133333 | 0.109681129 | 30.3964 | 30.2651 | 31.3729 | 30.713 | 30.625 | 30.348 |
| Endoplasmic reticulum resident protein 29 | P30040 | ERP29 | Extracellular or secreted | -0.243166667 | 0.275149155 | 28.6085 | 28.4661 | 29.3655 | 28.8345 | 28.7331 | 28.143 |
| Endoplasmic reticulum resident protein 44 | Q9BS26 | ERP44 | Extracellular or secreted | -0.432866667 | 0.490470599 | 29.7079 | 28.8734 | 29.9534 | 29.3391 | 29.1222 | 28.7748 |
| S-formylglutathione hydrolase | P10768 | ESD | Extracellular or secreted | -0.364066667 | 0.310737733 | 29.1145 | 30.2784 | 29.0344 | 29.0956 | 29.5336 | 28.7059 |
| ESF1 homolog | Q9H501 | ESF1 | Extracellular or secreted | 0.1257 | 0.045676268 | 22.7959 | 21.3704 | 24.1351 | 22.1675 | 22.7313 | 23.7797 |
| Epithelial splicing regulatory protein 1 | Q6NXG1-2;Q6NXG1-5;Q6NX( | ESRP1 | Extracellular or secreted | -1.115 | 0.392094163 | 24.2926 | 23.3769 | 24.4407 | 21.1918 | 22.7312 | 24.8422 |
| Extended synaptotagmin-1 | Q9BSJ8;Q9BSJ8-2 | ESYT1 | Extracellular or secreted | -0.971033333 | 1.909528459 | 26.5962 | 26.0563 | 26.124 | 25.355 | 25.0603 | 25.4481 |
| Eukaryotic peptide chain release factor subunit 1 | P62495;P62495-2 | ETF1 | Other | -0.5156 | 0.520485361 | 28.9665 | 29.1924 | 29.2103 | 27.8921 | 28.769 | 29.1613 |
| Electron transfer flavoprotein subunit alpha, mitochondr | P13804-2;P13804 | ETFA | Extracellular or secreted | -0.167733333 | 0.071447139 | 25.4096 | 25.297 | 27.6189 | 25.5104 | 26.1638 | 26.1481 |
| Electron transfer flavoprotein subunit beta | P38117;P38117-2 | ETFB | Extracellular or secreted | -0.202033333 | 0.087677878 | 24.7459 | 25.3593 | 27.1397 | 24.8473 | 25.8523 | 25.9392 |
| Persulfide dioxygenase ETHE1, mitochondrial | O95571 | ETHE1 | Other | -1.2751 | 0.627726047 | 24.1225 | 23.8387 | 26.0418 | 22.2111 | 24.0514 | 23.9152 |
| Ellis-van Creveld syndrome protein | P57679 | EVC | Extracellular or secreted | -2.455366667 | 0.637267565 | 26.0068 | 25.4418 | 21.3671 | 22.3023 | 21.0486 | 22.0987 |
| RNA-binding protein EWS | Q01844-6;Q01844-3;Q01844 | EWSR1 | Extracellular or secreted | -0.931766667 | 1.97079619 | 28.5164 | 28.6309 | 28.1452 | 27.2307 | 27.5371 | 27.7294 |
| Exocyst complex component 4 | Q96A65;Q96A65-2 | EXOC4 | Extracellular or secreted | 4.220033333 | 1.570243876 | 23.3127 | 22.0791 | 24.9987 | 27.874 | 28.0908 | 27.0858 |
| Exosome complex component CSL4 | Q9Y3B2 | EXOSC1 | Other | -0.649266667 | 0.451349455 | 21.6167 | 23.0375 | 22.5336 | 22.3404 | 20.8456 | 22.054 |
| Exosome component 10 | Q01780-2;Q01780 | EXOSC10 | Other | -1.041666667 | 0.353842128 | 21.4943 | 23.5884 | 25.1038 | 21.255 | 22.9868 | 22.8197 |
| Exosome complex component RRP4 | Q13868;Q13868-3;Q13868-2 | EXOSC2 | Other | -2.4605 | 1.792004236 | 27.3278 | 26.541 | 27.3351 | 23.7348 | 24.8077 | 25.2799 |
| Exosome complex component RRP41 | Q9NPD3 | EXOSC4 | Other | -2.5071 | 0.725388469 | 26.6621 | 26.2657 | 26.4785 | 21.406 | 25.3463 | 25.1327 |
| Exosome complex component RRP46 | Q9NQT4 | EXOSC5 | Other | -1.726 | 1.209968108 | 23.6015 | 23.2281 | 24.6302 | 22.61 | 22.6002 | 21.0716 |
| Exosome complex component MTR3 | Q5RKV6 | EXOSC6 | Other | -2.0139 | 0.59902449 | 25.0625 | 24.9792 | 26.0912 | 20.8199 | 24.4353 | 24.836 |
| Exosome complex component RRP42 | Q15024 | EXOSC7 | Other | -1.516533333 | 1.342561695 | 27.0237 | 26.504 | 27.2185 | 24.5843 | 25.8343 | 25.778 |
| Exosome complex component RRP43 | Q96B26 | EXOSC8 | Other | -1.625066667 | 1.207931269 | 25.7769 | 24.9199 | 26.4389 | 24.027 | 23.9925 | 24.241 |
| Exosome complex component RRP45 | Q06265;Q06265-2;Q06265-3 | EXOSC9 | Extracellular or secreted | -1.466366667 | 1.258309049 | 26.7451 | 26.2987 | 27.1866 | 24.6731 | 25.0482 | 26.11 |
| Exostosin-1 | Q16394 | EXT1 | Other | 1.661833333 | 0.595652394 | 27.2027 | 28.426 | 24.8342 | 28.1303 | 28.7402 | 28.5779 |
| Exostosin-2 | Q93063;Q93063-2;Q93063-3 | EXT2 | Extracellular or secreted | 1.833033333 | 0.784361256 | 27.8056 | 29.0192 | 26.0062 | 28.9781 | 29.6489 | 29.7031 |
| Exostosin-like 2;Processed exostosin-like 2 | Q9UBQ6 | EXTL2 | Extracellular or secreted | 0.4664 | 0.149153589 | 26.0119 | 27.0239 | 23.3832 | 25.9786 | 26.0094 | 25.8302 |
| Ezrin | P15311 | EZR | Extracellular or secreted | -0.176833333 | 0.202913221 | 32.7329 | 32.5888 | 31.9217 | 32.6321 | 32.2262 | 31.8546 |
| Junctional adhesion molecule A | Q9Y624;Q9Y624-2 | F11R | Extracellular or secreted | 1.065866667 | 1.481793024 | 27.7584 | 27.4031 | 27.0281 | 28.9217 | 28.4123 | 28.0532 |
| Tissue factor | P13726;P13726-2 | F3 | Extracellular or secreted | 0.065066667 | 0.025433006 | 26.5258 | 27.3555 | 24.9195 | 27.2344 | 25.9853 | 25.7763 |
| Fatty acid-binding protein, epidermal | Q01469 | FABP5 | Extracellular or secreted | -1.391933333 | 1.824807507 | 31.3717 | 31.6637 | 31.5267 | 30.1802 | 30.4839 | 29.7222 |
| Gastrotropin | P51161;P51161-2 | FABP6 | Extracellular or secreted | 2.473733333 | 1.211707697 | 22.2636 | 22.805 | 21.2688 | 23.8534 | 26.1014 | 23.8038 |
| FAS-associated death domain protein | Q13158 | FADD | Extracellular or secreted | -1.155033333 | 0.88076095 | 24.498 | 25.2346 | 23.5168 | 23.463 | 23.5162 | 22.8051 |
| Fatty acid desaturase 2 | O95864-3;O95864-4;O95864 | FADS2 | Extracellular or secreted | 2.555633333 | 1.51597041 | 19.9784 | 20.951 | 21.4578 | 22.1745 | 24.1608 | 23.7188 |
| FAS-associated factor 1 | Q9UNN5;Q9UNN5-2 | FAF1 | Extracellular or secreted | -0.7727 | 2.150042265 | 27.593 | 27.3939 | 27.6162 | 26.5672 | 26.7641 | 26.9537 |
| Fumarylacetoacetase | P16930;P16930-2 | FAH | Extracellular or secreted | -0.305833333 | 0.168212446 | 30.403 | 29.9847 | 29.0448 | 30.2547 | 29.8286 | 28.4317 |
| Acylpyruvase FAHD1, mitochondrial | Q6P587;Q6P587-2;Q6P587-3 | FAHD1 | Other | -0.423733333 | 0.374108099 | 28.6116 | 28.4512 | 28.3052 | 28.6202 | 28.2639 | 27.2127 |
| Fumarylacetoacetate hydrolase domain-containing prote | Q96GK7;Q6P2I3 | FAHD2A;FAHD | Other | -1.7048 | 0.640565598 | 24.7364 | 24.539 | 24.4183 | 21.1587 | 24.6129 | 22.8077 |
| Protein FAM107B | Q9H098;Q9H098-2 | FAM107B | Extracellular or secreted | 1.1191 | 0.866211021 | 25.9943 | 25.3108 | 24.3713 | 26.8238 | 26.5624 | 25.6475 |
| Constitutive coactivator of PPAR-gamma-like protein 1 | Q9NZB2-4;Q9NZB2;Q9NZB2- | FAM120A | Extracellular or secreted | -1.094833333 | 1.014281814 | 27.832 | 27.7948 | 27.7725 | 27.2243 | 25.9923 | 26.8982 |
| Protein Niban | Q9BZQ8 | FAM129A | Extracellular or secreted | 1.994466667 | 0.475442125 | 20.3577 | 24.2372 | 25.6511 | 25.3627 | 25.4956 | 25.3711 |
| Niban-like protein 1 | Q96TA1-2;Q96TA1 | FAM129B | Extracellular or secreted | 0.205266667 | 0.17259273 | 28.2373 | 29.1462 | 29.3467 | 28.5787 | 29.1829 | 29.5844 |
| Protein FAM136A | Q96C01 | FAM136A | Other | 1.624766667 | 0.762473105 | 21.4263 | 22.1087 | 24.0895 | 24.0532 | 24.5565 | 23.8891 |
| Protein FAM160B1 | Q5W0V3-2;Q5W0V3 | FAM160B1 | Other | -1.4918 | 0.68899427 | 21.4436 | 21.7616 | 24.0396 | 21.0352 | 20.531 | 21.2032 |
| Soluble lamin-associated protein of 75 kDa | Q9Y6X4 | FAM169A | Other | -0.8939 | 0.680199697 | 21.3764 | 23.0949 | 22.0934 | 21.0052 | 21.4331 | 21.4447 |
| BRISC complex subunit Abro1 | Q15018 | FAM175B | Other | -1.8921 | 1.047426982 | 24.0372 | 23.2587 | 23.7723 | 21.5234 | 23.0482 | 20.8203 |
| Protein FAM188A | Q9H8M7 | FAM188A | Other | -0.9955 | 0.309094845 | 24.4409 | 22.8736 | 24.0602 | 20.5142 | 24.2684 | 23.6056 |
| Glycosaminoglycan xylosylkinase | O75063 | FAM20B | Other | 0.205666667 | 0.087433131 | 20.6045 | 23.2103 | 22.2185 | 22.5321 | 22.4181 | 21.7001 |
| Extracellular serine/threonine protein kinase FAM20C | Q8IXL6;Q8IXL6-2 | FAM20C | Extracellular or secreted | 2.1598 | 0.953235018 | 23.3155 | 24.3414 | 21.5146 | 24.7755 | 25.3181 | 25.5573 |
| Protein FAM3A | P98173-3;P98173-2;P98173 | FAM3A | Extracellular or secreted | 0.4859 | 0.36571806 | 24.7498 | 26.1933 | 24.7098 | 25.342 | 25.8252 | 25.9434 |
| Protein FAM3C | Q92520 | FAM3C | Extracellular or secreted | 1.4226 | 0.717943623 | 29.9481 | 30.1619 | 27.7984 | 31.1356 | 31.0019 | 30.0387 |
| Protein FAM49B | Q9NUQ9;Q9NUQ9-2 | FAM49B | Extracellular or secreted | 0.219766667 | 0.261362418 | 27.7169 | 28.5406 | 28.3644 | 27.9829 | 28.6574 | 28.6409 |
| Protein FAM50A | Q14320 | FAM50A | Other | -0.6615 | 0.832388546 | 26.7688 | 26.5783 | 26.6988 | 25.4388 | 26.3497 | 26.2729 |
| Protein FAM83B | Q5T0W9 | FAM83B | Other | -1.154333333 | 0.515452098 | 24.0492 | 22.9586 | 23.8225 | 22.1792 | 24.0374 | 21.1507 |
| MIP18 family protein FAM96A | Q9H5X1;Q9H5X1-2 | FAM96A | Extracellular or secreted | 0.345133333 | 0.408462217 | 22.713 | 22.8151 | 22.7741 | 23.2087 | 23.6078 | 22.5211 |
| Protein FAM98A | Q8NCA5-2;Q8NCA5 | FAM98A | Other | -0.324366667 | 0.085054575 | 20.712 | 24.0016 | 24.0389 | 21.2336 | 22.6741 | 23.8717 |
| Protein FAM98B | Q52LJ0-2;Q52LJ0 | FAM98B | Other | 0.192866667 | 0.118211364 | 25.1204 | 25.5636 | 26.663 | 25.3228 | 25.9803 | 26.6225 |
| Fanconi anemia group I protein | Q9NVI1-2;Q9NVI1-1;Q9NVI1 | FANCI | Other | -4.370566667 | 2.534719849 | 27.557 | 26.8861 | 26.8784 | 23.4699 | 21.9703 | 22.7696 |
| Prolyl endopeptidase FAP;Antiplasmin-cleaving enzyme F | Q12884 | FAP | Extracellular or secreted | 1.470466667 | 1.644723095 | 27.3947 | 26.987 | 26.4553 | 28.5858 | 28.8316 | 27.831 |
| Phenylalanine--tRNA ligase alpha subunit | Q9Y285;Q9Y285-2 | FARSA | Other | 0.252233333 | 0.505696488 | 26.1607 | 26.2677 | 26.6592 | 26.3089 | 26.8293 | 26.7061 |
| Phenylalanine--tRNA ligase beta subunit | Q9NSD9;Q9NSD9-2 | FARSB | Other | 0.399966667 | 0.414997624 | 27.0107 | 27.4069 | 28.236 | 27.8504 | 27.8508 | 28.1523 |
| Tumor necrosis factor receptor superfamily member 6 | P25445-6;P25445 | FAS | Extracellular or secreted | -0.468966667 | 0.190297633 | 23.4534 | 21.7487 | 21.9532 | 20.5991 | 21.8994 | 23.2499 |
| Fatty acid synthase;[Acyl-carrier-protein] S-acetyltransfe | P49327 | FASN | Extracellular or secreted | 0.526666667 | 0.312574584 | 33.5079 | 34.1181 | 34.8355 | 33.6213 | 34.8901 | 35.5301 |
| Protocadherin Fat 1;Protocadherin Fat 1, nuclear form | Q14517 | FAT1 | Extracellular or secreted | -1.6147 | 0.999640279 | 29.4595 | 29.9005 | 28.0162 | 27.7085 | 27.4917 | 27.3319 |
| Protocadherin Fat 4 | Q6V0I7-2;Q6V0I7-3;Q6V0I7 | FAT4 | Extracellular or secreted | 0.3694 | 0.052380629 | 21.0136 | 25.8251 | 25.9814 | 21.0524 | 26.3919 | 26.484 |
| 40S ribosomal protein S30 | P62861 | FAU | Extracellular or secreted | 4.936666667 | 3.710266662 | 21.6449 | 21.7165 | 22.4042 | 27.2583 | 26.9871 | 26.3302 |
| rRNA 2-O-methyltransferase fibrillarin | P22087 | FBL | Extracellular or secreted | -2.392433333 | 0.724511431 | 25.0302 | 21.5248 | 25.5216 | 22.5124 | 20.6751 | 21.7118 |
| Filamin-binding LIM protein 1 | Q8WUP2-3;Q8WUP2 | FBLIM1 | Other | 0.7203 | 0.401629157 | 23.1501 | 24.2102 | 21.8833 | 23.7615 | 23.619 | 24.024 |
| Fibulin-1 | P23142;P23142-2;P23142-3; | FBLN1 | Extracellular or secreted | -1.528966667 | 1.835772142 | 23.8022 | 24.106 | 23.4033 | 22.2602 | 22.7219 | 21.7425 |
| Fibrillin-1 | P35555 | FBN1 | Extracellular or secreted | 0.250866667 | 0.06815917 | 23.2727 | 23.4847 | 20.9784 | 20.872 | 23.4608 | 24.1556 |

| | | | | | | | | | | | |
|---|---|---|---|---|---|---|---|---|---|---|---|
| Fructose-1,6-bisphosphatase 1 | P09467 | FBP1 | Extracellular or secreted | 1.4466 | 1.285562516 | 28.4107 | 27.6974 | 26.9772 | 29.5028 | 29.3305 | 28.5918 |
| Fructose-1,6-bisphosphatase isozyme 2 | O00757 | FBP2 | Extracellular or secreted | 1.503166667 | 1.924758164 | 25.1504 | 24.6483 | 24.3445 | 26.5058 | 26.4257 | 25.7212 |
| F-box only protein 2 | Q9UK22 | FBXO2 | Extracellular or secreted | 0.988233333 | 1.759295142 | 29.0991 | 28.9335 | 29.4461 | 30.0697 | 30.1961 | 30.1776 |
| F-box only protein 22 | Q8NEZ5 | FBXO22 | Other | -1.745733333 | 1.425577027 | 23.9072 | 24.3133 | 23.7298 | 21.6388 | 23.0389 | 22.0354 |
| High affinity immunoglobulin epsilon receptor subunit ga | P30273 | FCER1G | Extracellular or secreted | -1.0324 | 0.330097388 | 23.9765 | 23.5686 | 20.3487 | 22.2968 | 21.2729 | 21.2269 |
| IgGFc-binding protein | Q9Y6R7 | FCGBP | Extracellular or secreted | 4.9239 | 3.315548186 | 20.3615 | 21.4875 | 21.2697 | 25.9795 | 26.5084 | 25.4025 |
| Squalene synthase | P37268-5;P37268;P37268-4; | FDFT1 | Other | 2.343533333 | 1.11797068 | 19.7867 | 19.7242 | 22.1092 | 23.544 | 21.9716 | 23.1351 |
| Farnesyl pyrophosphate synthase | P14324-2;P14324 | FDPS | Other | 1.8391 | 2.403453164 | 31.0566 | 30.5846 | 30.5989 | 32.9305 | 32.6675 | 32.1594 |
| Adrenodoxin, mitochondrial | P10109 | FDX1 | Other | -0.9846 | 0.490471157 | 23.2138 | 23.6604 | 25.6789 | 23.1708 | 23.3693 | 23.0592 |
| Flap endonuclease 1 | P39748;P39748-2 | FEN1 | Extracellular or secreted | -0.409566667 | 0.270298616 | 26.4443 | 27.7155 | 28.0263 | 26.3804 | 26.9567 | 27.6203 |
| Fermitin family homolog 1 | Q9BQL6;Q9BQL6-2;Q9BQL6- | FERMT1 | Extracellular or secreted | 1.1151 | 1.350449455 | 26.4901 | 26.489 | 26.0505 | 27.267 | 27.07 | 28.0379 |
| Fermitin family homolog 2 | Q96AC1;Q96AC1-3;Q96AC1- | FERMT2 | Extracellular or secreted | -1.095766667 | 0.790628544 | 29.1043 | 29.3615 | 29.9109 | 27.3959 | 28.4807 | 29.2128 |
| Fibroblast growth factor 2 | P09038-2;P09038-3;P09038- | FGF2 | Extracellular or secreted | 1.616566667 | 0.954270772 | 22.5483 | 22.5207 | 20.6163 | 23.0849 | 24.0861 | 23.364 |
| Fibroblast growth factor-binding protein 1 | Q14512 | FGFBP1 | Extracellular or secreted | 0.4119 | 0.343264748 | 29.6128 | 29.2852 | 30.6228 | 29.7383 | 30.3142 | 30.704 |
| Fibroblast growth factor receptor-like 1 | Q8N441 | FGFRL1 | Extracellular or secreted | 0.7982 | 0.353064417 | 23.7268 | 24.0272 | 21.5196 | 24.7617 | 23.7205 | 23.186 |
| Fumarate hydratase, mitochondrial | P07954-2;P07954 | FH | Extracellular or secreted | -0.464633333 | 0.552119153 | 31.6756 | 31.5178 | 32.0836 | 31.7496 | 31.4388 | 30.6947 |
| Four and a half LIM domains protein 2 | Q14192 | FHL2 | Other | 0.942133333 | 0.476222166 | 23.6367 | 24.0075 | 21.6024 | 23.7077 | 24.2843 | 24.081 |
| Four and a half LIM domains protein 3 | Q13643 | FHL3 | Other | -1.131833333 | 0.389669117 | 23.5857 | 23.9741 | 23.1269 | 20.3311 | 23.9616 | 22.9985 |
| FH1/FH2 domain-containing protein 1 | Q9Y613 | FHOD1 | Other | -1.545 | 0.177898369 | 29.9215 | 30.4084 | 28.2911 | 30.7999 | 31.2488 | 21.9373 |
| Filamin A-interacting protein 1-like | Q4L180-2;Q4L180;Q4L180-5 | FILIP1L | Other | -1.806366667 | 0.375776458 | 26.7301 | 26.1898 | 21.9175 | 22.0095 | 25.7611 | 21.6477 |
| Pre-mRNA 3-end-processing factor FIP1 | Q6UN15-4;Q6UN15-3;Q6UN | FIP1L1 | Other | -1.1734 | 1.111199141 | 23.3781 | 22.8967 | 24.234 | 22.3488 | 21.8428 | 22.797 |
| Mitochondrial fission 1 protein | Q9Y3D6 | FIS1 | Other | -1.158266667 | 0.859011957 | 23.5432 | 24.7597 | 25.133 | 22.5658 | 23.4992 | 23.8961 |
| Peptidyl-prolyl cis-trans isomerase FKBP10 | Q96AY3 | FKBP10 | Extracellular or secreted | -0.721633333 | 0.949714979 | 28.0853 | 27.5094 | 28.4854 | 27.4694 | 27.3401 | 27.1057 |
| Peptidyl-prolyl cis-trans isomerase FKBP14 | Q9NWM8 | FKBP14 | Extracellular or secreted | -1.382433333 | 0.533454079 | 23.3926 | 23.6034 | 24.7135 | 23.7759 | 23.1912 | 20.5951 |
| FK506-binding protein 15 | Q5T1M5-2;Q5T1M5 | FKBP15 | Other | 0.5087 | 0.235304985 | 24.4134 | 21.7962 | 23.591 | 23.9852 | 23.403 | 23.9385 |
| Peptidyl-prolyl cis-trans isomerase FKBP1A | P62942 | FKBP1A | Extracellular or secreted | -0.191566667 | 0.189689097 | 29.1558 | 29.2831 | 28.3112 | 28.906 | 29.0088 | 28.2606 |
| Peptidyl-prolyl cis-trans isomerase FKBP2 | P26885 | FKBP2 | Extracellular or secreted | -1.114933333 | 1.572841178 | 29.6415 | 29.7511 | 30.0137 | 28.6906 | 29.0967 | 28.2742 |
| Peptidyl-prolyl cis-trans isomerase FKBP3 | Q00688 | FKBP3 | Other | 0.410666667 | 0.656310725 | 29.4658 | 29.6396 | 29.5071 | 29.5827 | 30.3905 | 29.8713 |
| Peptidyl-prolyl cis-trans isomerase FKBP4;Peptidyl-prolyl | Q02790 | FKBP4 | Extracellular or secreted | -1.035366667 | 1.584711469 | 32.6593 | 32.4276 | 32.311 | 31.004 | 31.726 | 31.5618 |
| Peptidyl-prolyl cis-trans isomerase FKBP5;Peptidyl-prolyl | Q13451;Q13451-2 | FKBP5 | Extracellular or secreted | -2.320233333 | 1.3147749 | 28.8022 | 28.6599 | 30.5371 | 26.6146 | 27.3604 | 27.0635 |
| Peptidyl-prolyl cis-trans isomerase FKBP7 | Q9Y680-3;Q9Y680;Q9Y680-1 | FKBP7 | Extracellular or secreted | -3.356766667 | 2.08952167 | 24.9145 | 24.6726 | 24.6183 | 20.6637 | 21.6601 | 21.8113 |
| Peptidyl-prolyl cis-trans isomerase FKBP9 | O95302;O95302-3 | FKBP9 | Extracellular or secreted | -0.196933333 | 0.174031558 | 26.5295 | 25.4664 | 26.0949 | 26.4152 | 25.6505 | 25.4343 |
| FAD synthase;Molybdenum cofactor biosynthesis protein | Q8NFF5-3;Q8NFF5-2;Q8NFF | FLAD1 | Extracellular or secreted | -0.807566667 | 0.408502787 | 21.1442 | 21.6514 | 23.5784 | 21.0591 | 21.2781 | 21.6141 |
| Protein flightless-1 homolog | Q13045-2;Q13045;Q13045-3 | FLII | Other | -0.591366667 | 0.280329826 | 26.1652 | 26.4535 | 27.7799 | 25.1387 | 26.0173 | 27.4685 |
| Filamin-A | P21333-2;P21333 | FLNA | Extracellular or secreted | -0.9745 | 2.277432228 | 35.7389 | 35.8574 | 35.7462 | 34.6644 | 34.752 | 35.0026 |
| Filamin-B | O75369-8;O75369-2;O75369 | FLNB | Extracellular or secreted | -0.776233333 | 2.173786478 | 34.5262 | 34.8981 | 34.6232 | 33.7687 | 33.9296 | 34.0205 |
| Filamin-C | Q14315-2;Q14315 | FLNC | Extracellular or secreted | -0.9395 | 0.180630887 | 23.1267 | 26.9807 | 26.5249 | 26.4374 | 25.8549 | 21.5215 |
| Flotillin-1 | O75955-2;O75955 | FLOT1 | Extracellular or secreted | 2.3369 | 0.741044759 | 24.5127 | 20.7299 | 24.0274 | 25.8779 | 25.6095 | 24.7933 |
| Flotillin-2 | Q14254 | FLOT2 | Extracellular or secreted | 1.430266667 | 0.816948191 | 22.8484 | 20.7636 | 22.133 | 23.6877 | 24.0347 | 22.3134 |
| Leucine-rich repeat transmembrane protein FLRT3 | Q9NZU0 | FLRT3 | Extracellular or secreted | 0.554233333 | 0.993909615 | 26.6034 | 27.2607 | 26.6588 | 27.5632 | 27.1941 | 27.4283 |
| Fragile X mental retardation protein 1 | Q06787-6;Q06787-2;Q06787 | FMR1 | Other | -0.697433333 | 0.584006252 | 26.401 | 25.7472 | 27.2195 | 25.6659 | 25.2827 | 26.3268 |
| Fibronectin;Anastellin;Ugl-Y1;Ugl-Y2;Ugl-Y3 | P02751;P02751-3;P02751-8; | FN1 | Extracellular or secreted | 1.9389 | 0.786635155 | 36.014 | 36.9733 | 33.9272 | 37.4816 | 37.627 | 37.6226 |
| Fibronectin;Anastellin;Ugl-Y1;Ugl-Y2;Ugl-Y3 | P02751-15;P02751-11;P0275 | FN1 | Extracellular or secreted | -0.275866667 | 0.066339477 | 23.156 | 25.0901 | 21.9794 | 21.4153 | 22.7305 | 25.2521 |
| Fibronectin;Anastellin;Ugl-Y1;Ugl-Y2;Ugl-Y3 | P02751-5 | FN1 | Extracellular or secreted | -0.927433333 | 0.209436626 | 27.0459 | 26.7822 | 23.7641 | 22.8834 | 27.4659 | 24.4606 |
| Fructosamine-3-kinase | Q9H479 | FN3K | Other | 0.207533333 | 0.122683599 | 22.638 | 23.9885 | 24.4499 | 23.3896 | 24.0965 | 24.2129 |
| Ketosamine-3-kinase | Q9HA64 | FN3KRP | Other | -0.441333333 | 0.402455973 | 25.4184 | 26.1681 | 26.538 | 24.9651 | 25.7783 | 26.0571 |
| Formin-binding protein 1-like | Q5T0N5-3;Q5T0N5-4;Q5T0N | FNBP1L | Extracellular or secreted | -0.0551 | 0.036476115 | 23.0012 | 23.9711 | 22.968 | 22.5556 | 23.3134 | 23.906 |
| Formin-binding protein 4 | Q8N3X1;Q8N3X1-2 | FNBP4 | Other | -2.2405 | 1.026772334 | 25.268 | 24.1186 | 25.1763 | 23.3447 | 20.9851 | 23.5116 |
| Protein farnesyltransferase/geranylgeranyltransferase ty | P49354;P49354-2 | FNTA | Other | -0.335266667 | 2.304015032 | 26.5706 | 26.7021 | 26.5648 | 26.2109 | 26.3357 | 26.2851 |
| Protein farnesyltransferase subunit beta | P49356;P49356-2 | FNTB | Other | 0.254833333 | 0.466076956 | 24.5751 | 24.1052 | 23.9071 | 24.232 | 24.5009 | 24.619 |
| Focadhesin | Q5VW36 | FOCAD | Extracellular or secreted | -1.358233333 | 0.564555085 | 22.7748 | 23.4966 | 23.5852 | 20.4078 | 21.8082 | 23.5659 |
| Forkhead box protein K2 | Q01167-2;Q01167 | FOXK2 | Other | -0.2873 | 0.092219967 | 22.8373 | 22.7204 | 24.0882 | 21.0308 | 23.4091 | 24.3441 |
| Extracellular matrix protein FRAS1 | Q86XX4;Q86XX4-2;Q86XX4-5 | FRAS1 | Extracellular or secreted | -0.881833333 | 0.220557297 | 24.0088 | 27.7932 | 23.1077 | 24.1193 | 23.9223 | 24.2226 |
| Protein FRG1 | Q14331 | FRG1 | Other | -3.2695 | 1.57136924 | 25.7033 | 27.1512 | 26.4056 | 21.7132 | 24.0667 | 23.6717 |
| Fascin | Q16658 | FSCN1 | Extracellular or secreted | -0.516 | 0.517465239 | 31.4414 | 32.1905 | 30.9119 | 31.1481 | 31.2222 | 30.6255 |
| Follistatin | P19883-2;P19883 | FST | Extracellular or secreted | 2.226933333 | 0.682214514 | 25.3689 | 26.6819 | 22.479 | 27.8177 | 27.0138 | 26.3791 |
| Follistatin-related protein 1 | Q12841;Q12841-2 | FSTL1 | Extracellular or secreted | 1.477933333 | 0.949565781 | 32.2355 | 32.819 | 30.8513 | 33.7845 | 33.5677 | 32.9874 |
| Follistatin-related protein 3 | O95633-2;O95633 | FSTL3 | Extracellular or secreted | 0.527033333 | 0.260947192 | 29.8659 | 30.5224 | 28.1493 | 30.51 | 30.1571 | 29.4516 |
| Ferritin heavy chain;Ferritin heavy chain, N-terminally pr | P02794 | FTH1 | Extracellular or secreted | 0.556566667 | 0.471966704 | 32.1847 | 31.817 | 31.5676 | 33.1291 | 32.5023 | 31.6076 |
| Ferritin light chain | P02792 | FTL | Extracellular or secreted | -0.235933333 | 0.142729698 | 28.1307 | 28.6733 | 28.1837 | 28.7727 | 28.5211 | 26.9861 |
| Alpha-ketoglutarate-dependent dioxygenase FTO | Q9C0B1 | FTO | Other | -2.7093 | 1.111738414 | 25.5828 | 25.6505 | 25.5028 | 21.2675 | 23.7674 | 23.5733 |
| pre-rRNA processing protein FTSJ3 | Q8IY81 | FTSJ3 | Other | -2.2607 | 0.813018399 | 23.7196 | 22.3822 | 25.7322 | 22.5009 | 22.5278 | 20.0232 |
| Far upstream element-binding protein 1 | Q96AE4 | FUBP1 | Other | -0.410333333 | 0.347452907 | 30.0194 | 29.5898 | 30.0198 | 28.6516 | 29.6125 | 30.1339 |
| Far upstream element-binding protein 1 | Q96AE4-2 | FUBP1 | Other | 1.583 | 0.453877232 | 19.6967 | 21.2408 | 24.1275 | 22.1683 | 23.797 | 23.8487 |
| Far upstream element-binding protein 3 | Q96I24;Q96I24-2 | FUBP3 | Other | 1.736733333 | 0.3673445 | 21.169 | 20.0442 | 25.2323 | 21.7119 | 24.2601 | 25.6837 |
| Tissue alpha-L-fucosidase | P04066 | FUCA1 | Extracellular or secreted | 1.425233333 | 2.61107431 | 27.4244 | 27.5156 | 27.0087 | 28.6015 | 29.003 | 28.6199 |
| Plasma alpha-L-fucosidase | Q9BTY2 | FUCA2 | Extracellular or secreted | 0.512966667 | 0.50885919 | 27.2153 | 27.4237 | 26.2167 | 27.607 | 27.7148 | 27.0728 |
| FUN14 domain-containing protein 2 | Q9BWH2 | FUNDC2 | Other | -0.7134 | 1.047482083 | 23.8388 | 23.2433 | 23.2081 | 22.8292 | 22.252 | 23.0688 |
| Furin | P09958 | FURIN | Extracellular or secreted | 0.8987 | 0.304899401 | 26.6281 | 28.1789 | 24.4354 | 27.2396 | 27.1258 | 27.5731 |
| RNA-binding protein FUS | P35637-2;P35637 | FUS | Other | -0.3336 | 0.719830825 | 31.0483 | 30.625 | 31.0649 | 30.3879 | 30.4629 | 30.8866 |
| Frataxin, mitochondrial;Frataxin intermediate form;Frata | Q16595;Q16595-2;Q16595-3 | FXN | Extracellular or secreted | -0.45 | 0.346794543 | 27.5574 | 27.1568 | 27.0337 | 27.4448 | 27.0845 | 25.8686 |
| Fragile X mental retardation syndrome-related protein 1 | P51114;P51114-3;P51114-2 | FXR1 | Extracellular or secreted | -0.902366667 | 0.452363632 | 24.7776 | 23.8414 | 26.4127 | 23.7115 | 24.4061 | 24.207 |
| Fragile X mental retardation syndrome-related protein 2 | P51116 | FXR2 | Extracellular or secreted | 0.204666667 | 0.073321742 | 23.1269 | 20.7525 | 23.6794 | 22.4244 | 22.486 | 23.2624 |
| FXYD domain-containing ion transport regulator 3 | Q14802;Q14802-2;Q14802-3 | FXYD3 | Extracellular or secreted | 2.649933333 | 1.312734334 | 21.7254 | 23.5412 | 21.0632 | 23.7919 | 25.4308 | 25.0569 |
| Frizzled-6 | O60353-2;O60353 | FZD6 | Extracellular or secreted | 0.5158 | 0.156725833 | 22.6541 | 23.4322 | 22.7797 | 25.6558 | 21.8413 | 22.9163 |
| Ras GTPase-activating protein-binding protein 1 | Q13283 | G3BP1 | Extracellular or secreted | -0.136033333 | 0.120108538 | 29.557 | 30.2614 | 30.3057 | 29.2614 | 30.0933 | 30.3613 |
| Ras GTPase-activating protein-binding protein 2 | Q9UN86-2;Q9UN86 | G3BP2 | Other | -0.6883 | 1.120351874 | 29.3526 | 29.4021 | 29.7505 | 28.3789 | 28.9018 | 29.1596 |
| Glucose-6-phosphate 1-dehydrogenase | P11413;P11413-2;P11413-3 | G6PD | Extracellular or secreted | 0.703866667 | 0.640757231 | 30.105 | 30.358 | 31.0211 | 30.3991 | 31.5852 | 31.6114 |
| Lysosomal alpha-glucosidase;76 kDa lysosomal alpha-glu | P10253 | GAA | Extracellular or secreted | -0.5091 | 0.631537754 | 31.0946 | 30.987 | 30.4955 | 30.8003 | 30.4633 | 29.7862 |
| Gamma-aminobutyric acid receptor-associated protein-li | P60520 | GABARAPL2 | Other | -0.132666667 | 0.151659437 | 24.3224 | 24.2817 | 23.6878 | 23.7993 | 24.4609 | 23.6337 |
| GA-binding protein alpha chain | Q06546 | GABPA | Other | -0.897366667 | 0.590526635 | 23.6814 | 22.7605 | 24.4174 | 22.0372 | 22.4809 | 23.6491 |
| Galactocerebrosidase | P54803-4;P54803-3;P54803 | GALC | Extracellular or secreted | 0.700033333 | 1.76966551 | 21.601 | 22.0018 | 21.9452 | 22.2971 | 22.6695 | 22.6815 |
| UDP-glucose 4-epimerase | Q14376;Q14376-2 | GALE | Extracellular or secreted | 0.084566667 | 0.021407629 | 23.819 | 23.7075 | 26.8162 | 23.3485 | 25.0353 | 26.2126 |
| N-acetylgalactosamine kinase | Q01415-2;Q01415 | GALK2 | Other | -1.267833333 | 0.693456174 | 25.6054 | 26.5271 | 26.2241 | 23.4582 | 25.5712 | 25.5237 |
| N-acetylgalactosamine-6-sulfatase | P34059 | GALNS | Extracellular or secreted | 1.4467 | 1.30015083 | 27.4904 | 27.4504 | 26.7638 | 29.1566 | 29.0188 | 27.8693 |
| Polypeptide N-acetylgalactosaminyltransferase 12 | Q8IXK2 | GALNT12 | Extracellular or secreted | 2.7637 | 0.755444937 | 24.539 | 24.9197 | 20.6853 | 25.8921 | 26.5023 | 26.0407 |
| Polypeptide N-acetylgalactosaminyltransferase 2;Polype | Q10471 | GALNT2 | Extracellular or secreted | -0.3564 | 0.277223204 | 28.9372 | 29.1234 | 27.6435 | 28.257 | 28.4053 | 27.9726 |
| Polypeptide N-acetylgalactosaminyltransferase 3 | Q14435 | GALNT3 | Extracellular or secreted | -1.2694 | 1.436430713 | 27.2324 | 26.8734 | 26.1459 | 25.1706 | 25.8721 | 25.4008 |
| Polypeptide N-acetylgalactosaminyltransferase 6 | Q8NCL4 | GALNT6 | Extracellular or secreted | 0.457033333 | 0.432424818 | 26.576 | 27.4069 | 26.0393 | 27.1205 | 27.2697 | 27.0031 |
| N-acetylgalactosaminyltransferase 7 | Q86SF2 | GALNT7 | Extracellular or secreted | 0.0381 | 0.030347878 | 27.1096 | 27.3355 | 26.0786 | 26.6948 | 27.1141 | 26.8291 |
| Neutral alpha-glucosidase AB | Q14697 | GANAB | Extracellular or secreted | 0.0335 | 0.027645089 | 30.9373 | 30.9296 | 32.0798 | 31.2745 | 31.419 | 31.3537 |
| Glyceraldehyde-3-phosphate dehydrogenase | P04406 | GAPDH | Extracellular or secreted | 1.025566667 | 0.877683642 | 35.6965 | 35.3099 | 35.2801 | 37.2405 | 36.3832 | 35.7395 |
| Glyceraldehyde-3-phosphate dehydrogenase | P04406-2 | GAPDH | Extracellular or secreted | 0.139166667 | 0.102205254 | 27.2728 | 26.5227 | 26.4921 | 27.7221 | 26.5109 | 26.4721 |
| GTPase-activating protein and VPS9 domain-containing p | Q14C86-3;Q14C86-4;Q14C86 | GAPVD1 | Other | 0.009233333 | 0.005468137 | 24.2064 | 24.1693 | 25.5988 | 24.1251 | 25.0304 | 24.8467 |
| H/ACA ribonucleoprotein complex subunit 1 | Q9NY12-2;Q9NY12 | GAR1 | Other | -0.200533333 | 0.036626634 | 23.3294 | 21.5091 | 25.3176 | 20.3436 | 25.337 | 23.8739 |
| Glycine--tRNA ligase | P41250 | GARS | Extracellular or secreted | -0.336533333 | 0.400799447 | 29.6073 | 30.3768 | 30.4561 | 29.3684 | 29.9292 | 30.133 |
| Trifunctional purine biosynthetic protein adenosine-3;Ph | P22102;P22102-2 | GART | Extracellular or secreted | -1.1414 | 1.735583916 | 30.1636 | 29.9505 | 30.6547 | 28.7862 | 29.0492 | 29.5092 |
| Growth arrest-specific protein 6 | Q14393;Q14393-3;Q14393-1 | GAS6 | Extracellular or secreted | 1.302466667 | 2.666245693 | 28.7473 | 28.9533 | 28.52 | 29.8158 | 30.2833 | 30.0289 |
| Transcriptional repressor p66-alpha | Q86YP4-2;Q86YP4;Q86YP4-3 | GATAD2A | Other | 0.178733333 | 0.052554518 | 21.5829 | 23.4665 | 25.2441 | 22.8097 | 24.0838 | 23.9362 |
| Glucosylceramidase | P04062-4;P04062-2;P04062; | GBA | Extracellular or secreted | -0.065233333 | 0.038956884 | 25.8846 | 26.9325 | 25.9957 | 25.2919 | 26.6986 | 26.6266 |
| 1,4-alpha-glucan-branching enzyme | Q04446 | GBE1 | Extracellular or secreted | -0.2984 | 0.861652037 | 28.8087 | 28.6205 | 29.0743 | 28.6668 | 28.4446 | 28.4969 |
| Interferon-induced guanylate-binding protein 1;Interfero | P32455;P32456 | GBP1;GBP2 | Extracellular or secreted | -1.183366667 | 0.770179241 | 24.5853 | 24.6912 | 24.6525 | 23.8811 | 24.1524 | 22.3454 |
| GRIP and coiled-coil domain-containing protein 2 | Q8IWJ2 | GCC2 | Other | -0.268666667 | 0.136637657 | 25.0909 | 23.2677 | 24.9003 | 23.3785 | 24.2037 | 24.8707 |
| GTP cyclohydrolase 1 feedback regulatory protein | P30047 | GCHFR | Other | -1.3322 | 0.708793123 | 23.4733 | 23.18 | 22.7235 | 23.2014 | 20.8998 | 21.279 |
| Glutamate--cysteine ligase catalytic subunit | P48506 | GCLC | Other | -0.260066667 | 0.386696461 | 28.1382 | 27.7038 | 28.3119 | 27.9473 | 28.0633 | 27.3631 |
| Glutamate--cysteine ligase regulatory subunit | P48507;P48507-2 | GCLM | Other | -0.728533333 | 0.637817913 | 28.0986 | 27.0708 | 28.5136 | 27.1958 | 27.0666 | 27.235 |
| Translational activator GCN1 | Q92616 | GCN1L1 | Other | 0.6922 | 0.301062864 | 25.3916 | 24.7929 | 27.3251 | 27.0938 | 25.4984 | 26.994 |
| N-acetyllactosaminide beta-1,6-N-acetylglucosaminyl-tra | Q8N0V5-2;Q8N0V5-3;Q8N0V | GCNT2 | Other | 0.601133333 | 0.631794306 | 22.397 | 23.1524 | 21.9181 | 23.2065 | 23.1004 | 22.964 |
| Glycine cleavage system H protein, mitochondrial | P23434 | GCSH | Extracellular or secreted | -1.0852 | 0.673138576 | 31.4734 | 31.1074 | 31.6099 | 31.482 | 30.0214 | 29.4317 |
| Guanine deaminase | Q9Y2T3;Q9Y2T3-3;Q9Y2T3-2 | GDA | Extracellular or secreted | 0.000333333 | 0.000185197 | 31.9025 | 31.011 | 30.4245 | 31.6161 | 31.3973 | 30.3256 |
| Rab GDP dissociation inhibitor alpha | P31150 | GDI1 | Other | 0.2897 | 0.276399045 | 31.68 | 31.4797 | 30.6794 | 32.0644 | 31.5813 | 31.0625 |
| Rab GDP dissociation inhibitor beta | P50395;P50395-2 | GDI2 | Extracellular or secreted | -0.3442 | 0.705691133 | 33.3784 | 33.587 | 33.0425 | 32.784 | 33.2976 | 32.8937 |
| Gem-associated protein 5 | Q8TEQ6 | GEMIN5 | Other | -1.1506 | 1.288128111 | 27.4404 | 27.6446 | 27.5993 | 25.8367 | 26.7253 | 26.6705 |
| FAD-linked sulfhydryl oxidase ALR | P55789;P55789-2 | GFER | Extracellular or secreted | -0.9033 | 0.290691221 | 20.8028 | 23.3852 | 24.1836 | 23.2723 | 20.9303 | 21.4591 |
| Ribosome-releasing factor 2, mitochondrial | Q969S9-5;Q969S9-4;Q969S9 | GFM2 | Other | -2.890333333 | 2.28387007 | 24.9218 | 24.6887 | 23.6954 | 22.2551 | 21.3038 | 21.076 |
| Glutamine--fructose-6-phosphate aminotransferase [isor | Q06210-2;Q06210 | GFPT1 | Extracellular or secreted | 0.448666667 | 0.252223933 | 27.0473 | 27.795 | 29.0074 | 27.7445 | 28.3187 | 29.1325 |
| ADP-ribosylation factor-binding protein GGA1 | Q9UJY5-4;Q9UJY5;Q9UJY5-3 | GGA1 | Other | -0.9857 | 0.79166996 | 24.6335 | 23.1608 | 23.2753 | 22.3847 | 22.5291 | 23.1987 |
| Gamma-glutamylcyclotransferase | O75223;O75223-3 | GGCT | Extracellular or secreted | -0.385066667 | 0.354526756 | 30.1333 | 29.6834 | 28.9989 | 29.7133 | 29.2945 | 28.6526 |
| Gamma-glutamyl hydrolase | Q92820 | GGH | Extracellular or secreted | -0.178566667 | 0.116961587 | 32.5521 | 32.3686 | 31.3198 | 32.2848 | 32.3206 | 31.0994 |

| | | | | | | | | | | | |
|---|---|---|---|---|---|---|---|---|---|---|---|
| Geranylgeranyl pyrophosphate synthase | O95749;O95749-2 | GGPS1 | Other | 0.618366667 | 1.698328388 | 26.869 | 26.7655 | 27.1666 | 27.4798 | 27.6726 | 27.5038 |
| Gamma-glutamyltransferase 5;Gamma-glutamyltransfera | P36269-2;P36269;P36269-3 | GGT5 | Extracellular or secreted | 1.755566667 | 1.12164823 | 19.7689 | 21.2429 | 21.2188 | 23.38 | 22.6213 | 21.496 |
| Glucose-induced degradation protein 4 homolog | Q8IVV7 | GID4 | Other | -1.009566667 | 0.472742096 | 23.2491 | 20.9626 | 22.4211 | 21.4667 | 22.1532 | 19.9842 |
| Glucose-induced degradation protein 8 homolog | Q9NWU2 | GID8 | Other | -0.0893 | 0.166334459 | 26.0523 | 25.8463 | 26.1963 | 25.6482 | 25.9379 | 26.2409 |
| PERQ amino acid-rich with GYF domain-containing prote | Q6Y7W6-4;Q6Y7W6-5;Q6Y7 | GIGYF2 | Other | 0.091933333 | 0.051334624 | 24.8855 | 24.5372 | 23.755 | 25.2081 | 24.7447 | 23.5007 |
| DNA replication complex GINS protein PSF1 | Q14691 | GINS1 | Other | -1.160633333 | 0.514498179 | 23.6968 | 23.3592 | 23.45 | 22.3154 | 23.8306 | 20.8781 |
| DNA replication complex GINS protein PSF2 | Q9Y248 | GINS2 | Other | -0.7754 | 0.982038069 | 25.7074 | 26.0309 | 25.0258 | 24.5871 | 25.0373 | 24.8135 |
| DNA replication complex GINS protein PSF3 | Q9BRX5;Q9BRX5-3;Q9BRX5- | GINS3 | Other | -0.794566667 | 1.370656238 | 26.4004 | 26.5309 | 26.0565 | 25.1774 | 25.9091 | 25.5176 |
| DNA replication complex GINS protein SLD5;DNA replicat | Q9BRT9;Q9BRT9-2 | GINS4 | Other | -2.618566667 | 0.648967886 | 26.1934 | 25.8263 | 25.5806 | 24.4229 | 25.0759 | 20.2458 |
| PDZ domain-containing protein GIPC1 | O14908;O14908-2 | GIPC1 | Extracellular or secreted | 1.2017 | 1.988882636 | 25.1375 | 24.6128 | 25.1778 | 25.8091 | 26.2866 | 26.4375 |
| ARF GTPase-activating protein GIT1 | Q9Y2X7;Q9Y2X7-3 | GIT1 | Other | 0.389666667 | 0.304438887 | 22.5675 | 23.2229 | 23.7164 | 24.312 | 23.4087 | 22.9551 |
| Alpha-galactosidase A | P06280 | GLA | Extracellular or secreted | -0.457466667 | 0.84804426 | 26.7246 | 26.979 | 26.3288 | 26.0665 | 26.5441 | 26.0494 |
| Beta-galactosidase | P16278-3;P16278;P16278-2 | GLB1 | Extracellular or secreted | -0.023233333 | 0.04041047 | 27.4587 | 27.6743 | 27.8838 | 27.3459 | 27.8 | 27.8012 |
| D-glucuronyl C5-epimerase | O94923 | GLCE | Extracellular or secreted | 2.052433333 | 0.691467648 | 20.2762 | 24.0152 | 21.3891 | 23.8301 | 24.2413 | 23.7664 |
| Golgi apparatus protein 1 | Q92896;Q92896-2;Q92896-3 | GLG1 | Extracellular or secreted | -0.242766667 | 0.205135922 | 28.7503 | 28.2076 | 27.5376 | 28.1293 | 28.2908 | 27.3471 |
| Glomulin | Q92990;Q92990-2 | GLMN | Other | -1.0942 | 0.248862897 | 23.7418 | 23.5724 | 24.8304 | 19.8038 | 24.3419 | 24.7163 |
| Lactoylglutathione lyase | Q04760-2;Q04760 | GLO1 | Extracellular or secreted | -0.009 | 0.005567033 | 31.4949 | 31.9513 | 30.3687 | 31.2942 | 31.6147 | 30.879 |
| Glyoxalase domain-containing protein 4 | Q9HC38-2;Q9HC38;Q9HC38- | GLOD4 | Extracellular or secreted | -0.406066667 | 0.373380182 | 31.2373 | 30.7455 | 30.3491 | 30.9361 | 30.5003 | 29.6773 |
| Glutaredoxin-1 | P35754 | GLRX | Extracellular or secreted | 0.786733333 | 1.050879958 | 24.7142 | 24.9044 | 24.0063 | 25.0495 | 25.7401 | 25.1955 |
| Glutaredoxin-3 | O76003 | GLRX3 | Extracellular or secreted | -0.782233333 | 1.151292506 | 29.4205 | 29.7128 | 29.1408 | 28.1374 | 28.8684 | 28.9216 |
| Glutaminase kidney isoform, mitochondrial | O94925-3;O94925 | GLS | Extracellular or secreted | 0.105333333 | 0.021340974 | 21.4635 | 21.5256 | 26.1416 | 22.9245 | 23.3119 | 23.2103 |
| Glycolipid transfer protein | Q9NZD2 | GLTP | Extracellular or secreted | 0.391233333 | 0.174927017 | 20.2222 | 21.8676 | 22.1157 | 22.9829 | 21.0095 | 21.3868 |
| Glutamate dehydrogenase 1, mitochondrial;Glutamate d | P00367;P00367-3;P00367-2; | GLUD1;GLUD2 | Extracellular or secreted | -0.2162 | 0.149104717 | 29.3756 | 29.1177 | 30.3013 | 29.9621 | 29.5755 | 28.6084 |
| Glutamine synthetase | P15104 | GLUL | Extracellular or secreted | 0.974033333 | 0.309492026 | 21.7077 | 21.1024 | 23.4419 | 20.9526 | 24.04 | 24.1815 |
| Putative oxidoreductase GLYR1 | Q49A26-5;Q49A26-4;Q49A2 | GLYR1 | Other | -1.4422 | 0.714269457 | 21.7518 | 23.4757 | 24.3443 | 22.2832 | 20.9041 | 22.0579 |
| Ganglioside GM2 activator;Ganglioside GM2 activator iso | P17900 | GM2A | Extracellular or secreted | 0.569766667 | 0.537941723 | 29.2042 | 30.0563 | 29.3642 | 29.4484 | 30.7539 | 30.1317 |
| GDP-mannose 4,6 dehydratase | O60547-2;O60547 | GMDS | Extracellular or secreted | 0.677766667 | 1.166726042 | 26.2395 | 26.4424 | 26.96 | 27.3239 | 27.0134 | 27.3379 |
| Glia maturation factor beta | P60983 | GMFB | Other | 0.144166667 | 0.255493017 | 28.6964 | 29.0868 | 28.6067 | 28.647 | 29.2308 | 28.9446 |
| GEM-interacting protein | Q9P107-2;Q9P107 | GMIP | Other | -1.477733333 | 0.766505207 | 25.5892 | 24.7052 | 23.8195 | 22.0761 | 24.5031 | 23.1015 |
| Mannose-1-phosphate guanyltransferase alpha | Q96IJ6;Q96IJ6-2 | GMPPA | Extracellular or secreted | 0.4136 | 0.206807771 | 24.2104 | 22.496 | 23.0752 | 24.8297 | 23.2577 | 22.935 |
| Mannose-1-phosphate guanyltransferase beta | Q9Y5P6;Q9Y5P6-2 | GMPPB | Extracellular or secreted | 0.6168 | 1.1319549 | 25.8299 | 26.4351 | 25.8506 | 26.741 | 26.4936 | 26.7314 |
| GMP reductase 2 | Q9P2T1;Q9P2T1-2;Q9P2T1-3 | GMPR2 | Other | -0.298933333 | 0.565859003 | 27.0236 | 26.8501 | 26.9476 | 27.0144 | 26.5826 | 26.3275 |
| GMP synthase [glutamine-hydrolyzing] | P49915;P49915-2 | GMPS | Other | -0.5434 | 0.577546284 | 29.4971 | 29.8449 | 30.0647 | 28.6078 | 29.3388 | 29.8299 |
| Guanine nucleotide-binding protein subunit alpha-11 | P29992 | GNA11 | Extracellular or secreted | 1.0449 | 0.68536751 | 25.6816 | 23.6819 | 24.8732 | 25.2922 | 26.0553 | 26.0239 |
| Guanine nucleotide-binding protein G(i) subunit alpha-1 | P63096;P63096-2 | GNAI1 | Extracellular or secreted | 0.723566667 | 0.554619706 | 25.0799 | 23.3685 | 24.1491 | 24.9313 | 25.1141 | 24.7228 |
| Guanine nucleotide-binding protein G(i) subunit alpha-2 | P04899;P04899-4;P04899-6; | GNAI2 | Extracellular or secreted | 0.856933333 | 1.774128455 | 28.655 | 28.1599 | 28.228 | 29.1494 | 29.4907 | 28.9736 |
| Guanine nucleotide-binding protein G(k) subunit alpha | P08754 | GNAI3 | Extracellular or secreted | 1.038566667 | 1.059487615 | 26.8263 | 25.8079 | 25.7236 | 27.6812 | 26.7069 | 27.0854 |
| Guanine nucleotide-binding protein G(q) subunit alpha | P50148 | GNAQ | Extracellular or secreted | -0.5284 | 0.290995862 | 21.2125 | 22.3534 | 21.3222 | 20.7539 | 22.3045 | 20.2445 |
| Guanine nucleotide-binding protein G(s) subunit alpha is | Q5JWF2;P63092;P63092-4;P | GNAS | Extracellular or secreted | 1.076133333 | 1.28993044 | 26.1826 | 25.6789 | 26.7128 | 27.4914 | 27.0209 | 27.2904 |
| Guanine nucleotide-binding protein G(I)/G(S)/G(T) subun | P62873;P62873-2 | GNB1 | Extracellular or secreted | 0.4437 | 0.527906245 | 27.8805 | 27.0171 | 28.0395 | 28.0194 | 28.1566 | 28.0922 |
| Guanine nucleotide-binding protein G(I)/G(S)/G(T) subun | P62879 | GNB2 | Extracellular or secreted | 0.242766667 | 0.403893568 | 28.9573 | 28.5902 | 29.3677 | 29.3381 | 29.1177 | 29.1877 |
| Guanine nucleotide-binding protein subunit beta-2-like 1 | P63244 | GNB2L1 | Extracellular or secreted | 0.415033333 | 1.158163871 | 30.3795 | 30.6456 | 30.823 | 31.0404 | 31.2078 | 30.845 |
| Bifunctional UDP-N-acetylglucosamine 2-epimerase/N-ac | Q9Y223-2;Q9Y223-3;Q9Y223 | GNE | Other | 5.173066667 | 1.496323936 | 20.0024 | 20.8474 | 22.739 | 24.1011 | 26.6355 | 28.3714 |
| Guanine nucleotide-binding protein G(I)/G(S)/G(O) subun | Q9UBI6 | GNG12 | Extracellular or secreted | 1.107733333 | 1.999597457 | 27.2785 | 27.3524 | 26.7737 | 28.3654 | 28.3754 | 27.987 |
| Guanine nucleotide-binding protein G(I)/G(S)/G(O) subun | P63218 | GNG5 | Extracellular or secreted | 1.562 | 0.360016557 | 21.1512 | 25.7076 | 25.297 | 27.6619 | 24.2831 | 24.8968 |
| Guanine nucleotide-binding protein-like 1 | P36915;P36915-2 | GNL1 | Extracellular or secreted | 0.9101 | 0.169946007 | 21.2812 | 20.7826 | 25.3315 | 24.7444 | 20.5337 | 24.8475 |
| Glucosamine-6-phosphate isomerase 1 | P46926;P46926-2 | GNPDA1 | Extracellular or secreted | -0.2739 | 0.254009639 | 30.1014 | 29.5379 | 29.2347 | 29.9211 | 29.3893 | 28.7419 |
| Glucosamine-6-phosphate isomerase 2 | Q8TDQ7;Q8TDQ7-3;Q8TDQ7 | GNPDA2 | Other | -0.209866667 | 0.223324009 | 26.4507 | 25.992 | 26.0819 | 26.3749 | 26.1947 | 25.3254 |
| Glucosamine 6-phosphate N-acetyltransferase | Q96EK6 | GNPNAT1 | Other | -0.8706 | 0.851682737 | 28.8913 | 29.611 | 28.2675 | 27.9852 | 28.4443 | 27.7285 |
| N-acetylglucosamine-6-sulfatase | P15586;P15586-2 | GNS | Extracellular or secreted | 0.878433333 | 1.015953932 | 29.1831 | 28.7316 | 28.5038 | 30.273 | 29.631 | 29.1498 |
| Golgin subfamily A member 2 | Q08379-2;Q08379;Q9NVV4- | GOLGA2 | Other | 0.821833333 | 0.376045117 | 22.5899 | 20.7674 | 23.536 | 22.7972 | 23.0004 | 23.5612 |
| Golgin subfamily A member 3 | Q08378;Q08378-2;Q08378-4 | GOLGA3 | Other | 0.205333333 | 0.216527839 | 25.2786 | 24.4355 | 25.4305 | 24.9478 | 25.2137 | 25.5991 |
| Golgin subfamily A member 4 | Q13439-3;Q13439-4;Q13439 | GOLGA4 | Extracellular or secreted | -1.055633333 | 0.998922786 | 28.5155 | 27.3921 | 28.6461 | 26.8316 | 26.9456 | 27.6096 |
| Golgin subfamily A member 5 | Q8TBA6-2;Q8TBA6 | GOLGA5 | Other | -0.7131 | 0.459186516 | 24.5396 | 23.2207 | 24.598 | 22.5151 | 24.2293 | 23.4746 |
| Golgin subfamily B member 1 | Q14789-4;Q14789-3;Q14789 | GOLGB1 | Other | -0.307133333 | 0.465912095 | 28.7074 | 28.0107 | 28.726 | 28.1268 | 28.4513 | 27.9446 |
| Golgi integral membrane protein 4 | O00461 | GOLIM4 | Other | 1.861766667 | 1.837919698 | 20.1956 | 21.3415 | 20.9033 | 23.2524 | 22.257 | 22.5163 |
| Golgi membrane protein 1 | Q8NBJ4-2;Q8NBJ4 | GOLM1 | Extracellular or secreted | 1.0629 | 0.699151643 | 28.6376 | 28.1488 | 27.3009 | 30.0847 | 29.0227 | 28.1686 |
| Golgi reassembly-stacking protein 2 | Q9H8Y8-2;Q9H8Y8;Q9H8Y8- | GORASP2 | Other | -0.01 | 0.004730969 | 26.0804 | 26.3307 | 26.8025 | 25.3237 | 26.3487 | 27.5112 |
| Golgi SNAP receptor complex member 1 | O95249;O95249-2 | GOSR1 | Other | 0.698166667 | 0.175996422 | 24.2595 | 21.3598 | 20.093 | 20.9414 | 23.5883 | 23.2771 |
| Aspartate aminotransferase, cytoplasmic | P17174;P17174-2 | GOT1 | Extracellular or secreted | -0.4412 | 0.288474895 | 33.4082 | 32.727 | 32.3677 | 33.3085 | 32.3681 | 31.5027 |
| Aspartate aminotransferase, mitochondrial | P00505;P00505-2 | GOT2 | Extracellular or secreted | -1.1078 | 0.950276394 | 34.2936 | 33.8036 | 33.5414 | 33.4415 | 32.9343 | 31.9394 |
| Glypican-1;Secreted glypican-1 | P35052;P35052-2 | GPC1 | Extracellular or secreted | 1.217533333 | 1.027198253 | 29.5685 | 28.9611 | 28.1142 | 30.7265 | 30.076 | 29.4939 |
| Glypican-6;Secreted glypican-6 | Q9Y625 | GPC6 | Extracellular or secreted | 2.356733333 | 1.584976996 | 23.6348 | 24.9769 | 23.4545 | 26.717 | 26.3806 | 26.0388 |
| Glycerol-3-phosphate dehydrogenase, mitochondrial | P43304;P43304-2 | GPD2 | Extracellular or secreted | 0.829366667 | 0.749171783 | 23.4676 | 24.2016 | 22.8462 | 24.0577 | 24.9665 | 23.9793 |
| Gephyrin;Molybdopterin adenylyltransferase;Molybdopt | Q9NQX3;Q9NQX3-2 | GPHN | Extracellular or secreted | -0.4195 | 0.310900083 | 27.1894 | 26.5279 | 27.5001 | 25.99 | 26.4362 | 27.5327 |
| Glucose-6-phosphate isomerase | P06744;P06744-2 | GPI | Extracellular or secreted | -0.296066667 | 0.222329218 | 34.6417 | 34.3105 | 33.9139 | 34.7574 | 34.04 | 33.1805 |
| G patch domain and KOW motifs-containing protein | Q92917 | GPKOW | Other | -1.736133333 | 0.414638791 | 24.7558 | 25.2549 | 25.1105 | 20.159 | 24.7757 | 24.9781 |
| G-protein coupled receptor 126 | Q86SQ4-2;Q86SQ4;Q86SQ4- | GPR126 | Extracellular or secreted | -0.1156 | 0.042262434 | 25.5859 | 26.1978 | 23.5529 | 25.1815 | 25.6642 | 24.1441 |
| G-protein coupled receptor 56;GPR56 N-terminal fragme | Q9Y653-5;Q9Y653-2;Q9Y653 | GPR56 | Extracellular or secreted | 0.0657 | 0.050086531 | 22.3548 | 23.4447 | 22.4562 | 22.4212 | 23.3696 | 22.662 |
| Retinoic acid-induced protein 3 | Q8NFJ5 | GPRC5A | Extracellular or secreted | 0.808833333 | 0.936809606 | 27.579 | 28.5151 | 27.5752 | 29.1888 | 28.4869 | 28.4201 |
| COP9 signalosome complex subunit 1 | Q13098-5;Q13098;Q13098-7 | GPS1 | Other | -1.4069 | 0.779857151 | 27.8644 | 28.2294 | 28.6187 | 25.5809 | 26.987 | 27.9239 |
| Glutathione peroxidase 1 | P07203 | GPX1 | Extracellular or secreted | 1.194133333 | 0.78962525 | 24.567 | 24.3836 | 26.1307 | 25.5005 | 26.289 | 26.8742 |
| Phospholipid hydroperoxide glutathione peroxidase, mit | P36969-2;P36969 | GPX4 | Extracellular or secreted | 0.0686 | 0.067336059 | 23.84 | 23.3741 | 24.1673 | 23.5412 | 23.6467 | 24.3993 |
| Growth factor receptor-bound protein 2 | P62993;P62993-2 | GRB2 | Extracellular or secreted | -1.087066667 | 1.304285337 | 27.8276 | 27.861 | 27.1295 | 25.9232 | 26.764 | 26.8697 |
| Grainyhead-like protein 2 homolog | Q6ISB3-2;Q6ISB3 | GRHL2 | Extracellular or secreted | -0.8983 | 0.588699033 | 23.6932 | 23.7029 | 24.0014 | 21.7471 | 23.4756 | 23.4799 |
| Glyoxylate reductase/hydroxypyruvate reductase | Q9UBQ7 | GRHPR | Extracellular or secreted | -0.387833333 | 0.652882201 | 26.4474 | 27.1969 | 26.908 | 26.5127 | 26.6824 | 26.1937 |
| GRIP1-associated protein 1 | Q4V328;Q4V328-2;Q4V328- | GRIPAP1 | Extracellular or secreted | 2.204033333 | 0.818735792 | 21.7087 | 21.8752 | 21.3821 | 22.0473 | 24.0996 | 25.4312 |
| Granulins;Acrogranin;Paragranulin;Granulin-1;Granulin-2 | P28799;P28799-2;P28799-3 | GRN | Extracellular or secreted | 0.945466667 | 0.847078535 | 29.2928 | 30.4514 | 29.0887 | 30.5287 | 30.9105 | 30.2301 |
| GrpE protein homolog 1, mitochondrial | Q9HAV7 | GRPEL1 | Other | -0.359833333 | 0.239961506 | 28.0742 | 28.3711 | 29.7117 | 28.1868 | 28.9112 | 27.9795 |
| G-rich sequence factor 1 | Q12849-5;Q12849 | GRSF1 | Other | -2.9358 | 1.762556231 | 26.1043 | 25.4315 | 26.4583 | 22.883 | 22.2092 | 24.0945 |
| Glutamate-rich WD repeat-containing protein 1 | Q9BQ67 | GRWD1 | Other | -0.411066667 | 0.290431787 | 25.8903 | 26.0386 | 27.4159 | 25.5343 | 26.0945 | 26.4828 |
| Gasdermin-A | Q96QA5 | GSDMA | Other | 2.715933333 | 0.733620009 | 22.6605 | 21.4318 | 23.2502 | 27.6938 | 22.8247 | 24.9718 |
| Glycogen synthase kinase-3 beta | P49841;P49841-2 | GSK3B | Extracellular or secreted | 0.052733333 | 0.077609824 | 24.1971 | 24.5659 | 24.5801 | 24.2347 | 24.3735 | 24.8931 |
| Gelsolin | P06396-2;P06396-4;P06396- | GSN | Extracellular or secreted | 0.340833333 | 1.434585381 | 29.9586 | 30.0837 | 29.7963 | 30.2686 | 30.4178 | 30.1747 |
| Eukaryotic peptide chain release factor GTP-binding subu | P15170-2;P15170-3;P15170 | GSPT1 | Other | -0.012633333 | 0.011690179 | 27.8345 | 27.6549 | 28.4928 | 27.7459 | 27.7179 | 28.4805 |
| Glutathione reductase, mitochondrial | P00390-4;P00390-2;P00390; | GSR | Extracellular or secreted | -0.132833333 | 0.094597999 | 30.1564 | 29.5561 | 29.3818 | 30.3161 | 29.562 | 28.8177 |
| Glutathione synthetase | P48637;P48637-2 | GSS | Extracellular or secreted | 0.724533333 | 0.568005096 | 30.6735 | 30.0178 | 29.3226 | 31.4299 | 30.7514 | 30.0062 |
| Glutathione S-transferase kappa 1 | Q9Y2Q3;Q9Y2Q3-3;Q9Y2Q3- | GSTK1 | Extracellular or secreted | -1.487833333 | 0.710981224 | 23.1629 | 24.3409 | 23.8262 | 23.8641 | 21.2849 | 21.7175 |
| Glutathione S-transferase Mu 3 | P21266 | GSTM3 | Extracellular or secreted | 0.200633333 | 0.344495259 | 24.5978 | 24.2516 | 24.923 | 24.5578 | 24.7769 | 25.0396 |
| Glutathione S-transferase omega-1 | P78417;P78417-3;P78417-2 | GSTO1 | Extracellular or secreted | 0.569333333 | 0.63028442 | 32.0159 | 32.2406 | 31.1629 | 32.7296 | 32.4374 | 31.9604 |
| Glutathione S-transferase P | P09211 | GSTP1 | Extracellular or secreted | -0.391733333 | 0.881939417 | 33.8539 | 33.7925 | 33.5297 | 33.474 | 33.5276 | 32.9993 |
| Glutathione S-transferase theta-1 | P30711;P30711-2 | GSTT1 | Extracellular or secreted | 0.0291 | 0.04300449 | 25.152 | 25.1197 | 24.6873 | 25.0258 | 25.3133 | 24.7072 |
| Glutathione S-transferase theta-2B;Glutathione S-transfe | P0CG30;P0CG29 | GSTT2B;GSTT2 | Extracellular or secreted | 0.181033333 | 0.069413405 | 23.4316 | 23.4862 | 23.2481 | 21.9149 | 24.0201 | 24.774 |
| Maleylacetoacetate isomerase | O43708;O43708-2;O43708-3 | GSTZ1 | Other | -0.776833333 | 0.612557227 | 24.4355 | 24.2548 | 25.6828 | 23.4218 | 24.5915 | 24.0293 |
| Transcription initiation factor IIA subunit 1;Transcription | P52655 | GTF2A1 | Other | -0.821366667 | 0.396774308 | 23.0051 | 22.9729 | 22.8751 | 20.5779 | 22.8501 | 22.961 |
| General transcription factor IIE subunit 1 | P29083 | GTF2E1 | Other | -0.345466667 | 0.242940707 | 21.8654 | 22.6913 | 21.9623 | 21.0144 | 21.7885 | 22.6797 |
| General transcription factor IIF subunit 1 | P35269 | GTF2F1 | Other | -0.664033333 | 0.763993343 | 25.9561 | 26.732 | 26.5231 | 25.1558 | 25.8203 | 26.243 |
| General transcription factor IIF subunit 2 | P13984 | GTF2F2 | Other | -1.032533333 | 0.729537189 | 24.1541 | 24.4404 | 24.2755 | 22.2657 | 23.4456 | 24.0611 |
| General transcription factor II-I | P78347-2;P78347-4;P78347- | GTF2I | Other | -0.551566667 | 0.255206696 | 27.3135 | 27.3962 | 28.417 | 25.8509 | 27.175 | 28.4461 |
| General transcription factor 3C polypeptide 1 | Q12789-3;Q12789 | GTF3C1 | Other | -2.217833333 | 0.733798115 | 24.6494 | 24.5561 | 23.9921 | 24.4182 | 21.3072 | 20.8187 |
| Guanylate kinase | Q16774;Q16774-2;Q16774-3 | GUK1 | Other | -0.2805 | 0.247102898 | 25.5768 | 24.8173 | 26.1829 | 24.9999 | 25.5827 | 25.1529 |
| Beta-glucuronidase | P08236-2;P08236;P08236-3 | GUSB | Extracellular or secreted | -0.3432 | 0.403656607 | 27.3725 | 27.1946 | 27.4679 | 27.5134 | 27.074 | 26.418 |
| Glycogenin-1 | P46976-2;P46976;P46976-3 | GYG1 | Extracellular or secreted | -0.904266667 | 0.520450528 | 25.5092 | 24.91 | 26.8841 | 24.428 | 24.3304 | 25.8321 |
| Glycogen [starch] synthase, muscle | P13807-2;P13807 | GYS1 | Other | 1.621866667 | 0.509932445 | 20.9611 | 21.4453 | 24.7071 | 22.9713 | 23.8871 | 25.1207 |
| Histone H1x | Q92522 | H1FX | Other | 3.7304 | 1.805762439 | 22.1683 | 20.1785 | 21.6587 | 24.6497 | 25.3916 | 25.1554 |
| Histone H2A.V;Histone H2A.Z | Q71UI9;P0C0S5;Q71UI9-3;Q | H2AFV;H2AFZ | Extracellular or secreted | 2.790066667 | 1.461947567 | 26.5306 | 26.2368 | 28.3826 | 29.4956 | 29.0977 | 30.9269 |
| Histone H2AX;Histone H2A type 1-A | P16104;Q96QV6 | H2AFX;HIST1H | Extracellular or secreted | 2.551433333 | 1.924167013 | 30.04 | 31.0635 | 31.4133 | 33.3979 | 33.0197 | 33.7535 |
| Core histone macro-H2A.1 | O75367-3;O75367;O75367-2 | H2AFY | Extracellular or secreted | 0.789766667 | 0.583456606 | 29.5673 | 28.598 | 29.9871 | 29.4927 | 30.0285 | 31.0005 |
| Histone H3.3 | P84243 | H3F3A | Extracellular or secreted | 2.935933333 | 1.445962332 | 29.8599 | 31.1797 | 31.8903 | 34.0567 | 33.7318 | 33.9492 |
| GDH/6PGL endoplasmic bifunctional protein;Glucose 1-d | O95479 | H6PD | Extracellular or secreted | -0.451933333 | 0.292394887 | 23.1198 | 23.4606 | 24.2045 | 22.099 | 23.6022 | 23.7279 |
| Hyaluronan-binding protein 2;Hyaluronan-binding protei | Q14520-2;Q14520 | HABP2 | Extracellular or secreted | -1.483233333 | 0.498885816 | 26.4706 | 26.6127 | 26.2766 | 22.7304 | 25.9532 | 26.2266 |
| Very-long-chain (3R)-3-hydroxyacyl-CoA dehydratase 3 | Q9P035 | HACD3 | Other | -4.7936 | 1.258246994 | 28.9497 | 27.5448 | 28.7096 | 21.0078 | 25.1547 | 24.6608 |
| Hydroxyacyl-coenzyme A dehydrogenase, mitochondrial | Q16836;Q16836-3;Q16836-2 | HADH | Other | -1.789966667 | 2.598560895 | 30.9163 | 30.717 | 31.2862 | 29.2385 | 29.494 | 28.8171 |
| Trifunctional enzyme subunit alpha, mitochondrial;Long- | P40939 | HADHA | Other | 1.1209 | 0.416708524 | 21.3372 | 23.3941 | 24.8189 | 24.4542 | 24.5404 | 23.9183 |
| Hydroxyacylglutathione hydrolase, mitochondrial | Q16775-2;Q16775 | HAGH | Extracellular or secreted | 0.1675 | 0.126049446 | 28.9274 | 28.5143 | 27.7792 | 29.0785 | 28.7488 | 27.8961 |
| Histidine ammonia-lyase | P42357-3;P42357;P42357-2 | HAL | Other | -1.275766667 | 0.34343848 | 23.9118 | 25.5857 | 23.602 | 25.6921 | 22.3189 | 21.2612 |
| Hyaluronan and proteoglycan link protein 3 | Q96S86 | HAPLN3 | Extracellular or secreted | 2.5381 | 0.667149955 | 24.4029 | 25.1725 | 20.5528 | 26.0761 | 26.2247 | 25.4417 |
| Histidine--tRNA ligase, cytoplasmic | P12081;P12081-4;P12081-2; | HARS | Other | 0.040766667 | 0.043679978 | 30.0307 | 29.7023 | 29.0333 | 29.7029 | 29.7192 | 29.4665 |
| Histone acetyltransferase type B catalytic subunit | O14929;O14929-2 | HAT1 | Other | -1.092033333 | 1.144422683 | 28.0372 | 27.3437 | 27.8291 | 26.1208 | 26.4879 | 27.3252 |

| | | | | | | | | | | | |
|---|---|---|---|---|---|---|---|---|---|---|---|
| Hemoglobin subunit beta;LVV-hemorphin-7;Spinorphin | P68871 | HBB | Extracellular or secreted | 1.1158 | 0.454099362 | 22.7977 | 21.1197 | 24.1539 | 23.9773 | 24.6494 | 22.792 |
| Host cell factor 1;HCF N-terminal chain 1;HCF N-terminal | P51610;P51610-2 | HCFC1 | Other | -1.957533333 | 0.584774775 | 28.6585 | 29.1139 | 29.9937 | 24.8483 | 27.9252 | 29.12 |
| Histone deacetylase 1 | Q13547 | HDAC1 | Other | -1.555466667 | 1.575756812 | 26.7703 | 26.8 | 27.8088 | 25.8126 | 24.9943 | 25.9058 |
| Histone deacetylase 2 | Q92769;Q92769-3 | HDAC2 | Other | -1.4214 | 0.986534741 | 28.4547 | 28.8511 | 29.5415 | 26.5536 | 27.5953 | 28.4342 |
| Histone deacetylase 6 | Q9UBN7;Q9UBN7-2 | HDAC6 | Extracellular or secreted | -0.7529 | 0.650031665 | 22.3812 | 22.324 | 23.2687 | 21.0753 | 22.3244 | 22.3155 |
| HD domain-containing protein 2 | Q7Z4H3;Q7Z4H3-2 | HDDC2 | Extracellular or secreted | -0.3679 | 0.300727652 | 24.6932 | 23.5291 | 24.8557 | 24.4876 | 23.7845 | 23.7022 |
| Guanosine-3,5-bis(diphosphate) 3-pyrophosphohydrolas | Q8N4P3;Q8N4P3-2 | HDDC3 | Other | -0.1708 | 0.121829042 | 27.8699 | 27.5053 | 27.2075 | 27.9469 | 27.6653 | 26.4581 |
| Hepatoma-derived growth factor | P51858;P51858-2;P51858-3 | HDGF | Extracellular or secreted | 0.239233333 | 0.168438091 | 33.1998 | 32.3099 | 32.5084 | 33.7519 | 32.786 | 32.1979 |
| Hepatoma-derived growth factor-related protein 2 | Q7Z4V5;Q7Z4V5-2;Q7Z4V5-3 | HDGFRP2 | Other | -0.018233333 | 0.017059516 | 27.6384 | 26.5747 | 27.4575 | 27.0404 | 27.2862 | 27.2893 |
| Hepatoma-derived growth factor-related protein 3 | Q9Y3E1 | HDGFRP3 | Other | 0.184333333 | 0.201771148 | 27.8809 | 27.4052 | 27.0124 | 27.7652 | 27.9513 | 27.135 |
| Pseudouridine-5-phosphatase | Q08623;Q08623-3;Q08623-4 | HDHD1 | Extracellular or secreted | -0.2913 | 0.222345779 | 27.2628 | 26.6839 | 25.6583 | 26.1999 | 26.3802 | 26.151 |
| Haloacid dehalogenase-like hydrolase domain-containing | Q9H0R4;Q9H0R4-2 | HDHD2 | Extracellular or secreted | -0.113133333 | 0.067979502 | 26.0886 | 25.9205 | 25.3276 | 26.6521 | 25.4344 | 24.9108 |
| Haloacid dehalogenase-like hydrolase domain-containing | Q9BSH5 | HDHD3 | Other | -2.032633333 | 0.911741155 | 25.9133 | 26.1079 | 26.9001 | 22.6075 | 25.1191 | 25.0968 |
| Vigilin | Q00341;Q00341-2 | HDLBP | Extracellular or secreted | -0.120266667 | 0.04234219 | 25.701 | 25.6514 | 27.756 | 25.0751 | 26.2585 | 27.414 |
| HEAT repeat-containing protein 1;HEAT repeat-containin | Q9H583 | HEATR1 | Other | -0.924166667 | 0.133121115 | 23.1155 | 26.5343 | 24.6444 | 22.8855 | 28.1617 | 20.4745 |
| HEAT repeat-containing protein 3 | Q7Z4Q2-2;Q7Z4Q2 | HEATR3 | Other | 0.596433333 | 0.147258645 | 21.1077 | 20.8017 | 23.9449 | 20.8341 | 22.1433 | 24.6662 |
| Heme-binding protein 1 | Q9NRV9 | HEBP1 | Extracellular or secreted | -1.803833333 | 1.1432293 | 28.7588 | 28.55 | 28.8643 | 25.9411 | 27.7518 | 27.0687 |
| Heme-binding protein 2 | Q9Y5Z4;Q9Y5Z4-2 | HEBP2 | Extracellular or secreted | 0.751966667 | 0.650460869 | 27.9587 | 28.0603 | 27.6748 | 29.392 | 28.6826 | 27.8751 |
| E3 ubiquitin-protein ligase HECTD1 | Q9ULT8 | HECTD1 | Other | -0.761366667 | 0.248973498 | 21.5804 | 22.3255 | 21.667 | 23.2994 | 19.9697 | 20.0197 |
| E3 ubiquitin-protein ligase HECTD3 | Q5T447;Q5T447-2 | HECTD3 | Other | 0.127166667 | 0.043218118 | 20.8001 | 22.2376 | 23.9039 | 21.8477 | 22.3229 | 23.1525 |
| Probable E3 ubiquitin-protein ligase HERC4 | Q5GLZ8-3;Q5GLZ8-2;Q5GLZ8 | HERC4 | Other | 0.779366667 | 0.253094699 | 21.5131 | 24.4922 | 25.1286 | 24.2293 | 24.3834 | 24.8593 |
| Beta-hexosaminidase subunit alpha | P06865 | HEXA | Extracellular or secreted | 0.8544 | 1.189658255 | 30.2273 | 30.3815 | 29.6649 | 31.2064 | 31.1934 | 30.4371 |
| Beta-hexosaminidase subunit beta;Beta-hexosaminidase | P07686 | HEXB | Extracellular or secreted | 0.710166667 | 0.762455581 | 30.9401 | 30.81 | 30.4178 | 31.9636 | 31.5754 | 30.7594 |
| Protein HEXIM1 | O94992 | HEXIM1 | Other | 1.624766667 | 0.73191074 | 23.061 | 21.5089 | 23.8022 | 24.3386 | 23.145 | 25.7628 |
| Hepatocyte growth factor-regulated tyrosine kinase subs | O14964-2;O14964 | HGS | Extracellular or secreted | 0.995433333 | 0.812126518 | 25.7752 | 27.171 | 26.0143 | 26.6112 | 27.8203 | 27.5153 |
| 3-hydroxyisobutyrate dehydrogenase, mitochondrial | P31937 | HIBADH | Extracellular or secreted | -0.9976 | 1.132861548 | 28.0948 | 28.435 | 28.9489 | 27.5458 | 28.0237 | 26.9164 |
| 3-hydroxyisobutyryl-CoA hydrolase, mitochondrial | Q6NVY1;Q6NVY1-2 | HIBCH | Extracellular or secreted | -1.064333333 | 1.020844601 | 29.0021 | 29.0418 | 29.1557 | 28.4203 | 28.2996 | 27.2867 |
| Histidine triad nucleotide-binding protein 1 | P49773 | HINT1 | Extracellular or secreted | 0.442866667 | 0.351322502 | 26.5743 | 27.4926 | 27.9239 | 27.1275 | 28.2701 | 27.9218 |
| Histidine triad nucleotide-binding protein 2, mitochondr | Q9BX68 | HINT2 | Extracellular or secreted | 0.062966667 | 0.022270772 | 22.6717 | 23.737 | 25.4238 | 24.0521 | 24.8038 | 23.1655 |
| Huntingtin-interacting protein 1 | O00291-3;O00291-4;O00291 | HIP1 | Other | -0.719166667 | 0.187112166 | 26.0049 | 23.8556 | 26.4066 | 22.5238 | 24.8622 | 26.7236 |
| HIRA-interacting protein 3 | Q9BW71 | HIRIP3 | Other | -0.965033333 | 0.765208606 | 23.9164 | 23.2302 | 22.6492 | 21.9947 | 23.1759 | 21.7301 |
| Histone H1.1 | Q02539 | HIST1H1A | Other | 1.7262 | 0.666393847 | 22.2289 | 22.3531 | 25.1107 | 23.9709 | 24.6867 | 26.2137 |
| Histone H1.5 | P16401 | HIST1H1B | Extracellular or secreted | 1.937866667 | 1.419241934 | 29.0709 | 28.8217 | 30.1256 | 30.456 | 31.2507 | 32.1251 |
| Histone H1.2 | P16403 | HIST1H1C | Other | 0.479566667 | 0.274223471 | 27.3105 | 25.6744 | 26.6039 | 26.0608 | 27.127 | 27.8397 |
| Histone H1.3 | P16402 | HIST1H1D | Other | 0.944466667 | 0.286755064 | 24.7015 | 21.3683 | 24.8288 | 23.4145 | 24.7503 | 25.5672 |
| Histone H1.4 | P10412 | HIST1H1E | Extracellular or secreted | 1.3278 | 1.060597359 | 29.3285 | 29.0279 | 30.2731 | 30.0255 | 31.0461 | 31.5413 |
| Histone H2A type 1-J;Histone H2A type 1-H;Histone H2A | Q99878;Q96KK5;Q9BTM1;Q | HIST1H2AJ;HIS | Extracellular or secreted | 3.212866667 | 1.704571162 | 28.8005 | 30.2802 | 30.8032 | 32.668 | 33.2818 | 33.5727 |
| Histone H2B type 1-L;Histone H2B type 1-M;Histone H2B | Q99880;Q99879;Q99877;Q9 | HIST1H2BL;HIS | Extracellular or secreted | 3.192933333 | 1.558633988 | 30.4726 | 31.9508 | 32.6309 | 34.6408 | 34.6868 | 35.3055 |
| Histone H4 | P62805 | HIST1H4A | Extracellular or secreted | 3.864766667 | 1.562829263 | 29.8453 | 31.2731 | 32.466 | 34.7794 | 34.862 | 35.5373 |
| Histone H2B type 2-E;Histone H2B type 1-B;Histone H2B | Q16778;P33778;P23527;P06 | HIST2H2BE;HIS | Extracellular or secreted | 3.328166667 | 1.702204686 | 26.6546 | 28.2017 | 28.5962 | 30.8836 | 30.9654 | 31.588 |
| Histone H3.2 | Q71DI3 | HIST2H3A | Extracellular or secreted | -1.420533333 | 0.507771842 | 25.5452 | 22.5507 | 22.2637 | 22.3958 | 22.5854 | 21.1168 |
| Hexokinase-1 | P19367;P19367-4;P19367-2; | HK1 | Other | -0.4404 | 0.285726994 | 30.4919 | 30.5842 | 31.7601 | 29.6474 | 30.6101 | 31.2575 |
| Hexokinase-2 | P52789 | HK2 | Other | -1.161033333 | 0.229928697 | 24.517 | 26.0602 | 29.262 | 23.8657 | 24.2441 | 28.2463 |
| HLA class I histocompatibility antigen, A-68 alpha chain | P01891 | HLA-A | Extracellular or secreted | 1.5647 | 1.540251759 | 29.6436 | 29.1182 | 28.6444 | 31.0871 | 31.025 | 29.9882 |
| HLA class I histocompatibility antigen, A-34 alpha chain;H | P30453;P30457;P30450;P18 | HLA-A | Extracellular or secreted | 1.336666667 | 1.352143124 | 25.3071 | 24.7582 | 24.0863 | 26.2758 | 26.3973 | 25.4885 |
| HLA class I histocompatibility antigen, B-42 alpha chain;H | P30480;P30462;P01889;Q95 | HLA-B | Extracellular or secreted | 1.799466667 | 1.683161868 | 24.4295 | 23.6875 | 23.2034 | 25.6695 | 26.0845 | 24.9648 |
| HLA class I histocompatibility antigen, Cw-12 alpha chain | P30508;Q07000;P30510;P30 | HLA-C | Extracellular or secreted | 1.492133333 | 1.129535001 | 27.6861 | 27.2665 | 26.5434 | 29.1072 | 29.1884 | 27.6768 |
| HLA class I histocompatibility antigen, alpha chain F | P30511;P30511-3 | HLA-F | Extracellular or secreted | -0.820533333 | 0.419159559 | 22.4219 | 22.2794 | 22.6052 | 21.0908 | 23.0684 | 20.6857 |
| Porphobilinogen deaminase | P08397-2;P08397;P08397-4; | HMBS | Other | -0.8986 | 1.075630025 | 28.2101 | 28.0496 | 28.1304 | 26.6645 | 27.5683 | 27.4615 |
| Hemicentin-1 | Q96RW7-2;Q96RW7 | HMCN1 | Extracellular or secreted | 1.462866667 | 0.381333022 | 25.7403 | 23.7199 | 21.3733 | 26.2905 | 25.812 | 23.1196 |
| High mobility group protein HMG-I/HMG-Y | P17096;P17096-3 | HMGA1 | Other | 1.2389 | 0.785458218 | 30.9498 | 29.2348 | 30.7017 | 32.4596 | 31.359 | 30.7844 |
| High mobility group protein HMG-I/HMG-Y | P17096-2 | HMGA1 | Other | 1.028066667 | 0.363060562 | 27.0085 | 25.4873 | 27.6605 | 29.4343 | 27.7257 | 26.0805 |
| High mobility group protein HMGI-C | P52926;P52926-3;P52926-6; | HMGA2 | Other | 2.024233333 | 0.995214054 | 24.1774 | 21.6002 | 23.1633 | 25.9476 | 25.0208 | 24.0452 |
| High mobility group protein B1;Putative high mobility gro | P09429;B2RPK0 | HMGB1;HMGB | Extracellular or secreted | -0.634333333 | 1.516861452 | 32.7576 | 32.6211 | 32.5805 | 32.1365 | 32.1736 | 31.7461 |
| High mobility group protein B2 | P26583 | HMGB2 | Extracellular or secreted | -0.479333333 | 0.667260794 | 32.3229 | 31.5062 | 31.73 | 31.5229 | 31.648 | 30.9502 |
| High mobility group protein B3 | O15347 | HMGB3 | Other | -0.3811 | 0.513885855 | 29.9523 | 29.1876 | 30.0707 | 29.1737 | 29.6163 | 29.2773 |
| Hydroxymethylglutaryl-CoA lyase, mitochondrial | P35914 | HMGCL | Extracellular or secreted | -1.1125 | 0.394448523 | 24.1624 | 24.8265 | 26.1164 | 22.335 | 25.7805 | 23.6523 |
| Hydroxymethylglutaryl-CoA synthase, cytoplasmic | Q01581 | HMGCS1 | Extracellular or secreted | 2.7303 | 1.673482014 | 27.023 | 25.6539 | 25.9115 | 28.8488 | 29.0391 | 28.8914 |
| Non-histone chromosomal protein HMG-14 | P05114 | HMGN1 | Other | 1.279233333 | 0.346416064 | 29.4955 | 26.1447 | 30.5643 | 31.1954 | 29.1991 | 29.6477 |
| Non-histone chromosomal protein HMG-17 | P05204 | HMGN2 | Other | 1.4353 | 0.525429505 | 20.5031 | 21.9056 | 20.9906 | 24.5631 | 22.0248 | 21.1173 |
| High mobility group nucleosome-binding domain-contain | Q15651 | HMGN3 | Other | 0.947166667 | 0.321605397 | 24.2122 | 22.3919 | 25.6493 | 26.5212 | 24.2277 | 24.346 |
| High mobility group nucleosome-binding domain-contain | P82970 | HMGN5 | Other | 1.836466667 | 0.845361227 | 22.8997 | 23.4534 | 23.5154 | 26.6995 | 24.0621 | 24.6163 |
| Heme oxygenase 1 | P09601 | HMOX1 | Extracellular or secreted | -2.182266667 | 1.104722403 | 23.4307 | 24.4893 | 23.7795 | 22.2474 | 22.6246 | 20.2807 |
| Heme oxygenase 2 | P30519-2;P30519 | HMOX2 | Extracellular or secreted | -1.568733333 | 0.752479215 | 27.1793 | 27.4995 | 27.0158 | 24.1223 | 26.5882 | 26.2779 |
| Hematological and neurological expressed 1 protein;Hem | Q9UK76;Q9UK76-3;Q9UK76- | HN1 | Extracellular or secreted | 2.800166667 | 0.801494145 | 20.9411 | 23.9286 | 25.2893 | 26.4713 | 26.3004 | 25.7878 |
| Hematological and neurological expressed 1-like protein | Q9H910;Q9H910-3;Q9H910- | HN1L | Extracellular or secreted | 1.421233333 | 2.931411318 | 26.4369 | 26.3892 | 26.7879 | 27.8182 | 28.1878 | 27.8717 |
| Heterogeneous nuclear ribonucleoprotein A0 | Q13151 | HNRNPA0 | Other | 0.857633333 | 1.121670298 | 23.9487 | 24.2174 | 24.9094 | 24.8878 | 25.4699 | 25.2907 |
| Heterogeneous nuclear ribonucleoprotein A1;Heterogen | P09651-2;P09651;P09651-3; | HNRNPA1;HNR | Extracellular or secreted | -0.6767 | 0.819221874 | 32.9543 | 33.1754 | 33.5238 | 31.9232 | 32.9462 | 32.754 |
| Heterogeneous nuclear ribonucleoproteins A2/B1 | P22626;P22626-2 | HNRNPA2B1 | Other | -0.191066667 | 0.212504757 | 33.5922 | 33.0145 | 34.065 | 33.0914 | 33.4008 | 33.6063 |
| Heterogeneous nuclear ribonucleoprotein A3 | P51991 | HNRNPA3 | Other | 0.2988 | 0.153687981 | 29.1844 | 29.5121 | 30.9983 | 29.3225 | 30.3742 | 30.8945 |
| Heterogeneous nuclear ribonucleoprotein A3 | P51991-2 | HNRNPA3 | Other | 0.460566667 | 0.089043177 | 22.4009 | 20.516 | 24.6193 | 23.4403 | 20.3447 | 25.1329 |
| Heterogeneous nuclear ribonucleoprotein A/B | Q99729-3;Q99729-4;Q99729 | HNRNPAB | Other | -0.742966667 | 1.367619135 | 31.0856 | 31.0207 | 31.0671 | 30.1326 | 30.6388 | 30.1731 |
| Heterogeneous nuclear ribonucleoproteins C1/C2 | P07910-2;P07910;P07910-4; | HNRNPC | Extracellular or secreted | -1.461633333 | 1.466681551 | 31.3367 | 30.8439 | 32.0432 | 29.5563 | 29.7554 | 30.5272 |
| Heterogeneous nuclear ribonucleoprotein D0 | Q14103-3;Q14103;Q14103-4 | HNRNPD | Extracellular or secreted | -0.631333333 | 0.906707014 | 31.7456 | 32.1676 | 32.0244 | 30.8272 | 31.5335 | 31.6829 |
| Heterogeneous nuclear ribonucleoprotein D-like | O14979-3;O14979-2;O14979 | HNRNPDL | Extracellular or secreted | 0.157633333 | 0.084996083 | 28.1767 | 28.1639 | 29.2908 | 27.6351 | 29.1637 | 29.3055 |
| Heterogeneous nuclear ribonucleoprotein F;Heterogene | P52597 | HNRNPF | Other | -0.550366667 | 0.330722356 | 30.1366 | 29.9193 | 31.0834 | 28.9484 | 29.645 | 30.8948 |
| Heterogeneous nuclear ribonucleoprotein H;Heterogene | P31943 | HNRNPH1 | Other | 0.1595 | 0.083596037 | 28.9539 | 28.9066 | 30.4789 | 28.9657 | 29.4205 | 30.4317 |
| Heterogeneous nuclear ribonucleoprotein H2 | P55795 | HNRNPH2 | Other | 0.167266667 | 0.041289344 | 24.5084 | 23.6809 | 26.2462 | 23.3704 | 24.3887 | 27.1782 |
| Heterogeneous nuclear ribonucleoprotein H3 | P31942-2;P31942;P31942-3; | HNRNPH3 | Other | -0.954566667 | 0.613529006 | 28.0817 | 27.7638 | 29.46 | 26.8338 | 27.2054 | 28.3396 |
| Heterogeneous nuclear ribonucleoprotein K | P61978;P61978-2 | HNRNPK | Extracellular or secreted | 0.155866667 | 0.073817264 | 30.1987 | 30.8116 | 31.8055 | 30.1257 | 31.0517 | 32.106 |
| Heterogeneous nuclear ribonucleoprotein K | P61978-3 | HNRNPK | Extracellular or secreted | 0.2441 | 0.079582752 | 23.1903 | 25.2772 | 25.5723 | 23.4356 | 25.2638 | 26.0727 |
| Heterogeneous nuclear ribonucleoprotein L | P14866;P14866-2 | HNRNPL | Extracellular or secreted | -0.483533333 | 0.186122794 | 28.8203 | 29.2586 | 31.2487 | 28.5826 | 28.7035 | 30.5909 |
| Heterogeneous nuclear ribonucleoprotein L-like | Q8WVV9-5;Q8WVV9-4;Q8W | HNRNPLL | Other | -1.0794 | 0.405138341 | 22.4162 | 23.3605 | 25.4653 | 22.1149 | 21.9043 | 23.9846 |
| Heterogeneous nuclear ribonucleoprotein M | P52272-2;P52272 | HNRNPM | Extracellular or secreted | -0.015233333 | 0.004938907 | 27.6828 | 26.358 | 29.2516 | 26.9266 | 27.5283 | 28.7918 |
| Heterogeneous nuclear ribonucleoprotein R | O43390;O43390-3 | HNRNPR | Other | -1.438533333 | 1.318711991 | 30.5129 | 30.9945 | 31.5633 | 28.8521 | 29.7002 | 30.2028 |
| Heterogeneous nuclear ribonucleoprotein R | O43390-2;O43390-4 | HNRNPR | Other | -1.660766667 | 0.785929109 | 23.0803 | 23.8062 | 23.8341 | 22.8631 | 22.5438 | 20.3314 |
| Heterogeneous nuclear ribonucleoprotein U | Q00839;Q00839-2 | HNRNPU | Other | -0.7903 | 0.502871566 | 30.3528 | 29.6124 | 31.6473 | 29.906 | 29.3858 | 29.9498 |
| Heterogeneous nuclear ribonucleoprotein U-like protein | Q9BUJ2-4;Q9BUJ2-2;Q9BUJ2 | HNRNPUL1 | Other | 0.6163 | 0.405227821 | 26.1503 | 26.247 | 27.711 | 26.5403 | 27.8119 | 27.605 |
| Heterogeneous nuclear ribonucleoprotein U-like protein | Q1KMD3 | HNRNPUL2 | Other | -0.931233333 | 0.663888383 | 29.2089 | 29.1201 | 30.7414 | 28.7904 | 28.61 | 28.8763 |
| Homer protein homolog 3 | Q9NSC5-5;Q9NSC5-2;Q9NSC | HOMER3 | Extracellular or secreted | 2.6116 | 1.277298476 | 20.745 | 22.7122 | 21.4606 | 22.7986 | 24.735 | 25.219 |
| Protein Hook homolog 2 | Q96ED9-2;Q96ED9 | HOOK2 | Other | 0.755766667 | 0.877607391 | 22.8187 | 23.081 | 22.7677 | 24.2719 | 23.2194 | 23.4434 |
| Protein Hook homolog 3 | Q86VS8 | HOOK3 | Other | -0.470166667 | 0.90779648 | 25.7472 | 25.164 | 25.7578 | 25.0854 | 25.279 | 24.8941 |
| Heterochromatin protein 1-binding protein 3 | Q5SSJ5;Q5SSJ5-2;Q5SSJ5-3 | HP1BP3 | Other | 1.403966667 | 0.498819187 | 28.1238 | 26.4568 | 29.6621 | 28.3593 | 29.0539 | 31.0414 |
| Hippocalcin-like protein 1;Neuron-specific calcium-bindir | P37235;P84074 | HPCAL1;HPCA | Extracellular or secreted | 2.026666667 | 1.936527488 | 27.208 | 27.3989 | 26.2833 | 28.5478 | 29.1667 | 29.2557 |
| 4-hydroxyphenylpyruvate dioxygenase-like protein | Q96IR7 | HPDL | Other | -2.183033333 | 1.824553027 | 28.1551 | 27.7574 | 29.1135 | 25.7025 | 26.5819 | 26.1925 |
| Hypoxanthine-guanine phosphoribosyltransferase | P00492 | HPRT1 | Extracellular or secreted | -0.733433333 | 1.251122902 | 31.091 | 30.8186 | 30.4306 | 30.2604 | 30.2272 | 29.6523 |
| Hemopexin | P02790 | HPX | Extracellular or secreted | 2.1508 | 1.533005364 | 23.2825 | 22.2852 | 21.5558 | 24.377 | 24.0458 | 25.1531 |
| Ribonuclease UK114 | P52758 | HRSP12 | Extracellular or secreted | -2.491933333 | 0.914190427 | 26.3228 | 25.3994 | 25.5634 | 21.3148 | 24.6337 | 23.8613 |
| Heparan-sulfate 6-O-sulfotransferase 2 | Q96MM7-2;Q96MM7-3;Q96 | HS6ST2 | Extracellular or secreted | -1.266333333 | 0.528633146 | 24.8306 | 25.1429 | 24.0211 | 21.5946 | 24.4551 | 24.1459 |
| Heat shock factor-binding protein 1 | O75506 | HSBP1 | Other | -1.808066667 | 0.963737395 | 22.3863 | 22.4764 | 21.6766 | 21.5453 | 20.4337 | 19.1361 |
| 3-hydroxyacyl-CoA dehydrogenase type-2 | Q99714;Q99714-2 | HSD17B10 | Extracellular or secreted | -0.852366667 | 0.386371403 | 27.5013 | 26.6098 | 29.279 | 26.0584 | 27.4186 | 27.356 |
| Very-long-chain 3-oxoacyl-CoA reductase | Q53GQ0 | HSD17B12 | Extracellular or secreted | 1.1407 | 0.313993635 | 20.2422 | 24.7984 | 23.3173 | 23.9361 | 24.1881 | 23.6558 |
| Peroxisomal multifunctional enzyme type 2;(3R)-hydroxy | P51659;P51659-3;P51659-2 | HSD17B4 | Other | -1.8753 | 1.143320086 | 26.168 | 26.6634 | 28.185 | 24.2809 | 25.4833 | 25.6263 |
| Estradiol 17-beta-dehydrogenase 8 | Q92506 | HSD17B8 | Extracellular or secreted | -1.889566667 | 1.390985177 | 25.9663 | 25.6167 | 26.7946 | 23.997 | 25.1841 | 23.5278 |
| Hydroxysteroid dehydrogenase-like protein 2 | Q6YN16;Q6YN16-2 | HSDL2 | Other | -3.043066667 | 1.428796473 | 23.661 | 25.5859 | 25.9716 | 22.4086 | 22.9811 | 20.6996 |
| Heat shock protein HSP 90-alpha | P07900;P07900-2 | HSP90AA1 | Extracellular or secreted | -0.6376 | 1.162986789 | 33.3683 | 33.1799 | 33.1806 | 32.2854 | 32.579 | 32.9516 |
| Heat shock protein HSP 90-beta | P08238 | HSP90AB1 | Extracellular or secreted | -0.433366667 | 0.52828049 | 34.9349 | 34.7996 | 35.0997 | 34.0131 | 34.4379 | 35.0831 |
| Putative heat shock protein HSP 90-beta 2 | Q58FF8 | HSP90AB2P | Other | -5.012333333 | 0.7356313 | 30.1066 | 29.732 | 30.036 | 25.6886 | 28.8763 | 20.2727 |
| Putative heat shock protein HSP 90-beta 4 | Q58FF6 | HSP90AB4P | Other | -3.0091 | 0.49111761 | 28.9799 | 29.5843 | 28.889 | 21.5617 | 27.8637 | 29.0005 |
| Endoplasmin | P14625 | HSP90B1 | Extracellular or secreted | 0.397166667 | 0.253526578 | 31.5112 | 31.5614 | 33.2282 | 32.7548 | 32.4349 | 32.3026 |
| Heat shock 70 kDa protein 13 | P48723 | HSPA13 | Extracellular or secreted | -0.219966667 | 0.227617578 | 26.8926 | 27.4735 | 26.5881 | 26.2158 | 27.014 | 27.0645 |
| Heat shock 70 kDa protein 14 | Q0VDF9 | HSPA14 | Other | -1.308833333 | 0.924861533 | 22.2146 | 22.0614 | 23.6529 | 21.4426 | 21.1582 | 21.4016 |
| Heat shock 70 kDa protein 1B;Heat shock 70 kDa protein | P0DMV9;P0DMV8;P0DMV8- | HSPA1B;HSPA1 | Extracellular or secreted | -0.545566667 | 1.784016871 | 34.2718 | 34.0319 | 33.9161 | 33.5273 | 33.6538 | 33.402 |
| Heat shock-related 70 kDa protein 2 | P54652 | HSPA2 | Extracellular or secreted | -0.844733333 | 1.242445139 | 27.0253 | 26.4 | 27.0165 | 25.4925 | 26.163 | 26.2521 |
| Heat shock 70 kDa protein 4 | P34932 | HSPA4 | Extracellular or secreted | -0.364566667 | 0.55097776 | 33.6446 | 33.077 | 32.9152 | 33.2152 | 32.7514 | 32.5765 |
| Heat shock 70 kDa protein 4L | O95757 | HSPA4L | Other | -0.101666667 | 0.135099606 | 27.1619 | 26.8597 | 27.5404 | 26.7074 | 27.1888 | 27.3608 |
| 78 kDa glucose-regulated protein | P11021 | HSPA5 | Extracellular or secreted | -0.034233333 | 0.022464781 | 33.9066 | 33.535 | 34.1799 | 34.6727 | 33.7475 | 33.0986 |
| Heat shock 70 kDa protein 6;Putative heat shock 70 kDa | P17066;P48741 | HSPA6;HSPA7 | Extracellular or secreted | -0.199666667 | 0.357671582 | 32.0656 | 31.9809 | 31.6699 | 31.9757 | 31.8105 | 31.3312 |
| Heat shock cognate 71 kDa protein | P11142;P11142-2 | HSPA8 | Extracellular or secreted | 0.024933333 | 0.04926344 | 35.1143 | 34.9502 | 34.5699 | 34.9548 | 34.9207 | 34.8337 |

| | | | | | | | | | | | |
|---|---|---|---|---|---|---|---|---|---|---|---|
| Stress-70 protein, mitochondrial | P38646 | HSPA9 | Extracellular or secreted | -0.921066667 | 0.783899625 | 30.6117 | 30.8128 | 32.04 | 29.9127 | 30.511 | 30.2776 |
| Heat shock protein beta-1 | P04792 | HSPB1 | Extracellular or secreted | 0.179833333 | 0.08519031 | 30.1488 | 29.9188 | 32.0588 | 30.459 | 30.9093 | 31.2976 |
| Intraflagellar transport protein 25 homolog | Q9Y547 | HSPB11 | Extracellular or secreted | -0.166966667 | 0.106494962 | 28.7582 | 28.687 | 27.8481 | 28.73 | 28.749 | 27.3134 |
| Hsp70-binding protein 1 | Q9NZL4;Q9NZL4-3;Q9NZL4-2 | HSPBP1 | Other | -2.1259 | 1.344449402 | 26.0213 | 26.5986 | 26.8583 | 23.3439 | 24.4983 | 25.2583 |
| 60 kDa heat shock protein, mitochondrial | P10809 | HSPD1 | Extracellular or secreted | -0.8659 | 0.722684575 | 33.0926 | 32.3518 | 33.931 | 32.5222 | 32.4227 | 31.8328 |
| 10 kDa heat shock protein, mitochondrial | P61604 | HSPE1 | Extracellular or secreted | -0.4717 | 0.451312634 | 31.5135 | 31.6755 | 32.2129 | 32.0292 | 31.2313 | 30.7263 |
| Basement membrane-specific heparan sulfate proteogly | P98160 | HSPG2 | Extracellular or secreted | 1.404866667 | 2.31209497 | 31.0515 | 31.1403 | 30.5948 | 32.3406 | 32.1729 | 32.4877 |
| Heat shock protein 105 kDa | Q92598;Q92598-2;Q92598-3 | HSPH1 | Extracellular or secreted | 0.154 | 0.317820653 | 31.5336 | 31.0196 | 31.2172 | 31.6548 | 31.2128 | 31.3648 |
| HIV Tat-specific factor 1 | O43719 | HTATSF1 | Other | -0.228966667 | 0.147426737 | 27.9308 | 26.8655 | 26.6398 | 27.5908 | 26.1493 | 27.0091 |
| Serine protease HTRA1 | Q92743 | HTRA1 | Extracellular or secreted | 1.1189 | 0.604820839 | 28.4849 | 28.6035 | 27.7343 | 28.208 | 29.3291 | 30.6423 |
| Serine protease HTRA2, mitochondrial | O43464-3;O43464;O43464-2 | HTRA2 | Extracellular or secreted | -2.138633333 | 1.622478316 | 25.5291 | 25.4409 | 26.8436 | 23.3707 | 24.2456 | 23.7814 |
| Huntingtin | P42858 | HTT | Other | -1.263033333 | 0.312093315 | 21.6707 | 23.1189 | 26.5453 | 22.2756 | 21.5443 | 23.7259 |
| E3 ubiquitin-protein ligase HUWE1 | Q7Z6Z7;Q7Z6Z7-2;Q7Z6Z7-3 | HUWE1 | Extracellular or secreted | -0.540966667 | 0.619676884 | 30.3295 | 30.1755 | 31.0411 | 29.625 | 29.7539 | 30.5443 |
| Hydrocephalus-inducing protein homolog | Q4G0P3;Q4G0P3-6;Q4G0P3- | HYDIN | Other | 1.4423 | 0.380650481 | 26.5499 | 27.5549 | 22.9305 | 26.208 | 27.7257 | 27.4285 |
| Putative hydroxypyruvate isomerase | Q5T013;Q5T013-3;Q5T013-4 | HYI | Other | 0.310366667 | 0.274262963 | 27.7894 | 27.2363 | 26.7864 | 28.1616 | 27.622 | 26.9596 |
| Hypoxia up-regulated protein 1 | Q9Y4L1 | HYOU1 | Extracellular or secreted | 0.268866667 | 0.257894433 | 28.8646 | 28.5507 | 29.7967 | 29.4699 | 29.1266 | 29.4221 |
| Huntingtin-interacting protein K | Q9NX55 | HYPK | Other | -0.568566667 | 0.414557396 | 24.7937 | 24.7429 | 26.3141 | 24.7376 | 24.6327 | 24.7747 |
| Isoamyl acetate-hydrolyzing esterase 1 homolog | Q2TAA2;Q2TAA2-2 | IAH1 | Extracellular or secreted | -0.584466667 | 1.461931529 | 28.7713 | 28.5764 | 28.7792 | 27.8483 | 28.2842 | 28.241 |
| Isoleucine--tRNA ligase, cytoplasmic | P41252 | IARS | Extracellular or secreted | -0.5918 | 0.43807126 | 29.42 | 29.3599 | 30.8085 | 28.7565 | 29.2129 | 29.8436 |
| Isoleucine--tRNA ligase, mitochondrial | Q9NSE4 | IARS2 | Other | -0.835566667 | 0.267517007 | 26.9142 | 28.6116 | 30.3356 | 26.2916 | 28.3865 | 28.6766 |
| Putative transferase CAF17, mitochondrial | Q5T440 | IBA57 | Other | -0.918 | 0.74211718 | 24.8396 | 26.1609 | 26.2554 | 24.2193 | 25.1812 | 25.1014 |
| Intercellular adhesion molecule 1 | P05362 | ICAM1 | Extracellular or secreted | 1.1034 | 2.105563432 | 28.1532 | 28.1418 | 27.754 | 29.3888 | 29.1617 | 28.8087 |
| Insulin-degrading enzyme | P14735 | IDE | Extracellular or secreted | 0.3954 | 1.467681864 | 27.5452 | 27.458 | 27.715 | 27.9642 | 27.9779 | 27.9623 |
| Isocitrate dehydrogenase [NADP] cytoplasmic | O75874 | IDH1 | Extracellular or secreted | 0.587266667 | 0.71260126 | 30.6041 | 31.5738 | 30.8887 | 31.324 | 32.0839 | 31.4205 |
| Isocitrate dehydrogenase [NADP], mitochondrial | P48735-2;P48735 | IDH2 | Extracellular or secreted | -1.165666667 | 0.319133138 | 22.1515 | 23.5693 | 25.5918 | 23.7882 | 23.6464 | 20.381 |
| Isocitrate dehydrogenase [NAD] subunit alpha, mitochon | P50213-2;P50213 | IDH3A | Other | -2.894733333 | 2.08268193 | 26.5551 | 26.5086 | 27.8566 | 23.8558 | 23.5452 | 24.8351 |
| Isocitrate dehydrogenase [NAD] subunit beta, mitochond | O43837-2;O43837 | IDH3B | Other | -1.144 | 0.604373745 | 22.3845 | 21.5052 | 23.9225 | 22.2138 | 20.8731 | 21.2933 |
| Isopentenyl-diphosphate Delta-isomerase 1 | Q13907;Q13907-2 | IDI1 | Other | 2.587433333 | 1.718170168 | 28.5785 | 28.5595 | 28.5683 | 31.1843 | 31.771 | 30.5133 |
| Iduronate 2-sulfatase;Iduronate 2-sulfatase 42 kDa chain | P22304 | IDS | Extracellular or secreted | 1.360466667 | 0.698811934 | 28.0862 | 28.7592 | 26.3237 | 29.7367 | 29.2877 | 28.2261 |
| Alpha-L-iduronidase | P35475 | IDUA | Extracellular or secreted | 1.081566667 | 1.212354757 | 23.161 | 23.8528 | 23.0441 | 24.1029 | 25.0811 | 24.1186 |
| Gamma-interferon-inducible lysosomal thiol reductase | P13284 | IFI30 | Extracellular or secreted | -0.583433333 | 0.34362044 | 29.7592 | 29.6575 | 27.9809 | 28.0256 | 29.2947 | 28.327 |
| Interferon-induced 35 kDa protein | P80217;P80217-2 | IFI35 | Other | 1.650833333 | 1.549876518 | 24.0853 | 24.1009 | 23.3079 | 26.2331 | 25.0138 | 25.1997 |
| Interferon-induced transmembrane protein 3 | Q01628 | IFITM3 | Extracellular or secreted | 1.435633333 | 2.713828026 | 25.6136 | 25.4785 | 25.1465 | 27.0188 | 26.8049 | 26.7218 |
| Immunoglobulin-binding protein 1 | P78318 | IGBP1 | Other | -0.189666667 | 0.244974053 | 24.7332 | 24.5917 | 24.5469 | 24.3004 | 24.0325 | 24.9699 |
| Insulin-like growth factor II;Insulin-like growth factor II;In | P01344;P01344-2;P01344-3 | IGF2 | Extracellular or secreted | -4.542133333 | 1.262759316 | 27.9115 | 27.8249 | 25.8029 | 20.5587 | 25.0746 | 22.2796 |
| Cation-independent mannose-6-phosphate receptor | P11717 | IGF2R | Extracellular or secreted | 1.2098 | 1.786298705 | 28.3297 | 28.2148 | 27.5978 | 29.5176 | 29.3795 | 28.8746 |
| Insulin-like growth factor-binding protein 2 | P18065 | IGFBP2 | Extracellular or secreted | 1.027633333 | 0.926870912 | 27.706 | 27.6413 | 26.8059 | 27.8648 | 29.2166 | 28.1547 |
| Insulin-like growth factor-binding protein 3 | P17936;P17936-2 | IGFBP3 | Extracellular or secreted | 3.040666667 | 0.8743091 | 26.8071 | 29.2768 | 24.9295 | 28.2202 | 30.3418 | 31.5734 |
| Insulin-like growth factor-binding protein 4 | P22692;P22692-2 | IGFBP4 | Extracellular or secreted | 0.792633333 | 0.795326142 | 29.3874 | 29.3113 | 28.2996 | 29.2963 | 30.3118 | 29.7681 |
| Insulin-like growth factor-binding protein 6 | P24592 | IGFBP6 | Extracellular or secreted | 0.739766667 | 0.412952443 | 30.6829 | 31.5 | 29.2396 | 31.404 | 31.5808 | 30.657 |
| Insulin-like growth factor-binding protein 7 | Q16270-2;Q16270 | IGFBP7 | Extracellular or secreted | 2.1889 | 1.091445473 | 29.121 | 28.9967 | 26.8265 | 29.5112 | 31.2399 | 30.7598 |
| Ig alpha-1 chain C region | P01876 | IGHA1 | Other | 1.585833333 | 0.496519867 | 22.9718 | 21.7255 | 24.3058 | 25.2239 | 26.166 | 22.3707 |
| Immunoglobulin superfamily member 3 | O75054;O75054-2 | IGSF3 | Other | -0.658033333 | 0.767904786 | 24.1506 | 23.5101 | 23.3892 | 22.6903 | 23.642 | 22.7435 |
| Immunoglobulin superfamily member 8 | Q969P0;Q969P0-3 | IGSF8 | Extracellular or secreted | -0.252233333 | 0.452833401 | 29.2365 | 29.1941 | 28.6337 | 29.0371 | 28.6203 | 28.6502 |
| Protein Red | Q13123 | IK | Extracellular or secreted | -0.8651 | 0.402248303 | 22.3459 | 22.3785 | 23.3803 | 21.0469 | 23.4189 | 21.0436 |
| Elongator complex protein 1 | O95163 | IKBKAP | Other | -0.396766667 | 0.310125783 | 24.4837 | 24.6467 | 25.6043 | 24.2949 | 23.9818 | 25.2677 |
| NF-kappa-B essential modulator | Q9Y6K9;Q9Y6K9-2 | IKBKG | Other | -0.103133333 | 0.427319923 | 25.3857 | 25.1291 | 25.1437 | 25.2111 | 25.0098 | 25.1282 |
| Interleukin-18 | Q14116-2;Q14116 | IL18 | Extracellular or secreted | 0.9938 | 1.219258795 | 27.6598 | 27.1456 | 28.1353 | 28.8234 | 28.4717 | 28.627 |
| Interleukin-19 | Q9UHD0;Q9UHD0-2;Q9UHD | IL19 | Extracellular or secreted | 1.212733333 | 0.368454532 | 20.5682 | 23.9542 | 22.2659 | 24.0375 | 24.8014 | 21.5876 |
| Interleukin-1 alpha | P01583 | IL1A | Extracellular or secreted | 0.75 | 0.558412717 | 26.7909 | 25.6079 | 25.2029 | 27.2898 | 26.3834 | 26.1785 |
| Interleukin-1 receptor accessory protein | Q9NPH3-2;Q9NPH3;Q9NPH3 | IL1RAP | Extracellular or secreted | 0.2337 | 0.139841842 | 27.1582 | 26.818 | 25.3002 | 26.7724 | 26.7833 | 26.4218 |
| Interleukin-1 receptor antagonist protein | P18510-2;P18510-4;P18510; | IL1RN | Extracellular or secreted | -0.642 | 0.59533663 | 27.8088 | 28.0957 | 27.6685 | 26.9755 | 28.0147 | 26.6568 |
| Interleukin-22 receptor subunit alpha-2 | Q969J5-3;Q969J5-2;Q969J5 | IL22RA2 | Extracellular or secreted | 2.6226 | 1.57147421 | 21.8106 | 22.8224 | 21.2811 | 24.0043 | 25.7707 | 24.0069 |
| Interleukin-27 receptor subunit alpha | Q6UWB1 | IL27RA | Extracellular or secreted | 0.029666667 | 0.029008659 | 25.6427 | 25.405 | 24.6779 | 25.34 | 25.5362 | 24.9384 |
| Interleukin-36 gamma | Q9NZH8;Q9NZH8-2 | IL36G | Extracellular or secreted | 2.551333333 | 1.331190801 | 23.8011 | 24.3471 | 22.0662 | 26.3912 | 26.6074 | 24.8698 |
| Interleukin-6 | P05231 | IL6 | Extracellular or secreted | 2.2244 | 0.911012015 | 28.1782 | 25.6999 | 25.1801 | 28.2844 | 29.4291 | 28.0179 |
| Interleukin-6 receptor subunit beta | P40189-2;P40189-3;P40189 | IL6ST | Extracellular or secreted | 1.264433333 | 0.542709568 | 20.3463 | 23.1037 | 22.1154 | 22.5065 | 24.3665 | 22.4857 |
| Interleukin enhancer-binding factor 2 | Q12905 | ILF2 | Other | -1.096833333 | 0.873224702 | 30.2564 | 30.5173 | 31.2318 | 28.7086 | 29.6669 | 30.3395 |
| Interleukin enhancer-binding factor 3 | Q12906;Q12906-5;Q12906-2 | ILF3 | Other | -0.398966667 | 0.24384914 | 25.6477 | 25.5106 | 27.0359 | 25.0452 | 25.4786 | 26.4735 |
| Interleukin enhancer-binding factor 3 | Q12906-7;Q12906-4;Q12906 | ILF3 | Other | -1.041866667 | 0.79147317 | 31.531 | 31.398 | 32.3674 | 29.7927 | 30.8985 | 31.4796 |
| Integrin-linked kinase-associated serine/threonine phosp | Q9H0C8 | ILKAP | Other | -2.250633333 | 0.775926081 | 24.1174 | 24.7622 | 24.8627 | 21.2145 | 21.2867 | 24.4892 |
| MICOS complex subunit MIC60 | Q16891-3;Q16891-2;Q16891 | IMMT | Other | -0.9563 | 0.301117636 | 24.2307 | 21.9774 | 26.0154 | 23.3661 | 22.8106 | 23.1779 |
| Inositol monophosphatase 1 | P29218;P29218-3;P29218-2 | IMPA1 | Extracellular or secreted | 0.036266667 | 0.053955401 | 27.7486 | 27.5665 | 27.6304 | 28.0213 | 27.7464 | 27.2866 |
| Inosine-5-monophosphate dehydrogenase 1 | P20839;P20839-3;P20839-7; | IMPDH1 | Other | 0.6019 | 0.127760991 | 23.4759 | 22.3419 | 26.428 | 22.3553 | 26.5119 | 25.1843 |
| Inosine-5-monophosphate dehydrogenase 2 | P12268 | IMPDH2 | Extracellular or secreted | 0.2445 | 0.162527444 | 29.0492 | 28.6308 | 30.2173 | 29.3049 | 29.2017 | 30.1242 |
| Immortalization up-regulated protein | Q9GZP8-2;Q9GZP8 | IMUP | Other | 1.103633333 | 0.191387773 | 25.0879 | 20.6804 | 27.0621 | 27.0579 | 25.6624 | 23.421 |
| Inhibitor of growth protein 1 | Q9UK53-3;Q9UK53-4;Q9UK5 | ING1 | Other | 0.4195 | 0.199238222 | 20.4551 | 21.052 | 22.0651 | 20.3258 | 22.4531 | 22.0518 |
| Inhibin alpha chain | P05111 | INHA | Extracellular or secreted | -2.401933333 | 0.686089433 | 26.7617 | 27.16 | 25.9963 | 21.6264 | 25.9195 | 25.1663 |
| Inhibin beta A chain | P08476 | INHBA | Extracellular or secreted | 0.571133333 | 0.246058481 | 30.9911 | 32.1033 | 29.2281 | 31.3217 | 31.6238 | 31.0904 |
| Importin-11 | Q9UI26;Q9UI26-2 | IPO11 | Extracellular or secreted | -2.0547 | 0.845214648 | 25.262 | 25.0966 | 25.7095 | 21.5594 | 23.8375 | 24.5071 |
| Importin-4 | Q8TEX9;Q8TEX9-2 | IPO4 | Other | -0.972366667 | 0.257879633 | 21.5684 | 24.1792 | 25.3973 | 21.584 | 21.9447 | 24.6991 |
| Importin-5 | O00410;O00410-3;O00410-2 | IPO5 | Other | -0.8658 | 1.11181065 | 30.3636 | 30.7359 | 30.3989 | 29.0643 | 29.8885 | 29.9482 |
| Importin-7 | O95373 | IPO7 | Other | -1.028233333 | 0.943274553 | 28.5989 | 29.5579 | 29.5392 | 27.5013 | 28.2536 | 28.8564 |
| Importin-9 | Q96P70 | IPO9 | Extracellular or secreted | -1.208366667 | 0.712677128 | 27.5199 | 27.9718 | 28.733 | 25.6447 | 27.1447 | 27.8102 |
| Ras GTPase-activating-like protein IQGAP1 | P46940 | IQGAP1 | Extracellular or secreted | -0.773933333 | 0.635321179 | 32.5611 | 32.7648 | 32.8477 | 31.1221 | 32.0203 | 32.7094 |
| Interferon regulatory factor 2-binding protein 1 | Q8IU81 | IRF2BP1 | Other | -5.3493 | 2.721627288 | 28.8207 | 28.8768 | 27.2321 | 23.8431 | 22.1491 | 22.8895 |
| Interferon regulatory factor 2-binding protein 2 | Q7Z5L9-2;Q7Z5L9 | IRF2BP2 | Other | 0.1959 | 0.112971949 | 25.6236 | 26.9176 | 26.936 | 25.9069 | 26.6768 | 27.4812 |
| Ubiquitin-like protein ISG15 | P05161 | ISG15 | Extracellular or secreted | 0.215566667 | 0.173411296 | 27.4641 | 26.8804 | 26.9146 | 26.4952 | 27.8734 | 27.5372 |
| Isochorismatase domain-containing protein 1 | Q96CN7 | ISOC1 | Extracellular or secreted | -1.198333333 | 1.261473086 | 27.4261 | 27.4352 | 28.4746 | 26.1133 | 26.9734 | 26.6542 |
| Isochorismatase domain-containing protein 2, mitochond | Q96AB3;Q96AB3-2;Q96AB3- | ISOC2 | Other | -0.729466667 | 0.535408842 | 25.3505 | 25.9047 | 27.0226 | 24.8548 | 25.9499 | 25.2847 |
| IST1 homolog | P53990-2;P53990;P53990-4; | IST1 | Extracellular or secreted | -0.568866667 | 1.027242378 | 26.628 | 26.6319 | 26.9958 | 26.0376 | 25.917 | 26.5945 |
| Inositol-3-phosphate synthase 1 | Q9NPH2-2;Q9NPH2-3;Q9NP | ISYNA1 | Other | -0.8765 | 1.151379626 | 27.6316 | 27.5342 | 26.7349 | 26.6207 | 26.5475 | 26.103 |
| E3 ubiquitin-protein ligase Itchy homolog | Q96J02-3;Q96J02-2;Q96J02 | ITCH | Extracellular or secreted | 0.146433333 | 0.105729366 | 22.494 | 22.0454 | 22.0634 | 22.3095 | 21.5786 | 23.154 |
| Integrin-alpha FG-GAP repeat-containing protein 2 | Q969R8 | ITFG2 | Other | -0.563466667 | 0.465216883 | 22.321 | 22.2582 | 22.3372 | 22.2364 | 20.8295 | 22.1601 |
| Protein ITFG3 | Q9H0X4-2;Q9H0X4 | ITFG3 | Extracellular or secreted | -0.359933333 | 0.080419953 | 24.0781 | 23.7162 | 23.8514 | 24.5927 | 25.3802 | 20.593 |
| Integrin alpha-1 | P56199 | ITGA1 | Extracellular or secreted | 2.993966667 | 2.650710206 | 23.8434 | 23.773 | 23.0107 | 25.9911 | 27.0984 | 26.5195 |
| Integrin alpha-2 | P17301 | ITGA2 | Extracellular or secreted | 2.304 | 1.174270619 | 28.9732 | 28.449 | 26.66 | 30.682 | 30.4926 | 29.8196 |
| Integrin alpha-3;Integrin alpha-3 heavy chain;Integrin alp | P26006;P26006-1 | ITGA3 | Extracellular or secreted | 1.094033333 | 0.82701889 | 27.6874 | 26.9797 | 26.2323 | 28.8886 | 27.9331 | 27.3598 |
| Integrin alpha-5;Integrin alpha-5 heavy chain;Integrin alp | P08648 | ITGA5 | Extracellular or secreted | 0.070066667 | 0.066590898 | 26.5936 | 25.4258 | 25.7938 | 25.9704 | 26.02 | 26.033 |
| Integrin alpha-6;Integrin alpha-6 heavy chain;Integrin alp | P23229-4;P23229-2;P23229- | ITGA6 | Extracellular or secreted | 0.865866667 | 0.687734198 | 27.9082 | 27.9527 | 26.6033 | 29.053 | 28.1205 | 27.8883 |
| Integrin alpha-V;Integrin alpha-V heavy chain;Integrin alp | P06756-3;P06756;P06756-2 | ITGAV | Extracellular or secreted | 0.8256 | 0.728218752 | 26.7349 | 26.1438 | 25.2703 | 26.9633 | 26.9798 | 26.6827 |
| Integrin beta-1 | P05556;P05556-2;P05556-5; | ITGB1 | Extracellular or secreted | 1.581466667 | 1.565648305 | 29.5984 | 29.3248 | 28.726 | 31.4645 | 30.693 | 30.2361 |
| Integrin beta-4 | P16144-4;P16144-2;P16144; | ITGB4 | Extracellular or secreted | 0.614866667 | 0.928030727 | 27.0896 | 26.8409 | 26.9418 | 28.0185 | 27.5193 | 27.1791 |
| Integrin beta-5 | P18084 | ITGB5 | Extracellular or secreted | 0.7402 | 0.781655922 | 24.9141 | 23.9794 | 24.2057 | 25.66 | 25.1435 | 24.5163 |
| Integrin beta-6 | P18564-2;P18564 | ITGB6 | Extracellular or secreted | 1.2385 | 1.452324126 | 25.4483 | 24.738 | 24.4087 | 26.4763 | 25.9068 | 25.9274 |
| Integrin beta-like protein 1 | O95965-2;O95965-3;O95965 | ITGBL1 | Extracellular or secreted | 2.280133333 | 0.480829965 | 24.1007 | 27.1536 | 20.9441 | 27.1156 | 26.2326 | 25.6906 |
| Integral membrane protein 2B;BRI2, membrane form;BR | Q9Y287-2;Q9Y287 | ITM2B | Extracellular or secreted | -1.1121 | 0.494731437 | 23.1022 | 25.5286 | 23.1924 | 23.4166 | 23.3292 | 21.7411 |
| Inosine triphosphate pyrophosphatase | Q9BY32;Q9BY32-2;Q9BY32-3 | ITPA | Other | -0.202033333 | 0.332973022 | 26.8088 | 27.4902 | 27.1135 | 26.6783 | 26.9261 | 27.202 |
| Intersectin-1 | Q15811-13;Q15811-6;Q1581 | ITSN1 | Extracellular or secreted | -0.938933333 | 1.257187713 | 22.9591 | 22.3048 | 22.8181 | 22.1786 | 21.8428 | 21.2438 |
| Isovaleryl-CoA dehydrogenase, mitochondrial | P26440-2;P26440 | IVD | Other | 1.237866667 | 1.894422197 | 23.4091 | 23.7059 | 24.1088 | 25.1383 | 24.7919 | 25.0072 |
| Protein IWS1 homolog | Q96ST2;Q96ST2-3;Q96ST2-2 | IWS1 | Other | -0.4174 | 0.304055252 | 26.2511 | 24.7274 | 25.3413 | 25.5337 | 24.3937 | 25.1402 |
| Protein jagged-1 | P78504;P78504-2 | JAG1 | Extracellular or secreted | -0.351766667 | 0.20190313 | 27.4492 | 27.6933 | 25.9289 | 27.1176 | 26.9594 | 25.9391 |
| JmjC domain-containing protein 7 | P0C870;P0C869-8;P0C869-7; | JMJD7 | Other | -0.136333333 | 0.093363333 | 23.7916 | 23.1044 | 22.3611 | 23.5173 | 22.8968 | 22.434 |
| Junction plakoglobin | P14923 | JUP | Extracellular or secreted | 0.5284 | 0.503481519 | 29.6956 | 29.6993 | 30.8231 | 30.743 | 30.1221 | 30.9381 |
| Lysine--tRNA ligase | Q15046;Q15046-2 | KARS | Extracellular or secreted | -0.373933333 | 0.415320552 | 30.6332 | 30.5284 | 31.184 | 29.9124 | 30.3175 | 30.9939 |
| Histone acetyltransferase KAT7 | O95251-2;O95251-4;O95251 | KAT7 | Other | 2.476033333 | 0.943513334 | 21.2441 | 23.4223 | 20.281 | 24.384 | 23.9476 | 24.0439 |
| E3 ubiquitin-protein ligase KCMF1 | Q9P0J7 | KCMF1 | Other | -0.348 | 0.246015615 | 24.3699 | 25.237 | 25.7623 | 24.1119 | 24.7694 | 25.4439 |
| Kv channel-interacting protein 2;Kv channel-interacting p | Q9NS61-6;Q9NZI2-2;Q9NS61 | KCNIP2;KCNIP1 | Extracellular or secreted | 0.8608 | 0.239350649 | 23.2825 | 22.8857 | 22.2943 | 24.8199 | 25.1173 | 21.1077 |
| Intermediate conductance calcium-activated potassium c | O15554 | KCNN4 | Extracellular or secreted | 0.642333333 | 0.586151174 | 21.066 | 20.1058 | 19.8126 | 21.4106 | 21.1132 | 20.3876 |
| BTB/POZ domain-containing protein KCTD12 | Q96CX2 | KCTD12 | Extracellular or secreted | -0.341866667 | 0.367111183 | 25.5659 | 24.5862 | 25.6799 | 25.0253 | 24.806 | 24.9751 |
| BTB/POZ domain-containing protein KCTD3 | Q9Y597-2;Q9Y597 | KCTD3 | Other | -0.675466667 | 0.303030053 | 24.1874 | 22.1482 | 24.0375 | 23.2603 | 21.542 | 23.5444 |
| KDEL motif-containing protein 2 | Q7Z4H8;Q7Z4H8-2;Q7Z4H8- | KDELC2 | Extracellular or secreted | -4.365933333 | 2.414262695 | 25.7059 | 26.017 | 26.6436 | 22.0315 | 22.4399 | 20.7973 |
| Keratinocyte differentiation factor 1 | Q8NAX2 | KDF1 | Other | 0.377833333 | 0.284360774 | 22.7673 | 21.5348 | 22.7219 | 22.7916 | 23.2877 | 22.0782 |
| Lysine-specific histone demethylase 1A | O60341;O60341-2 | KDM1A | Extracellular or secreted | -1.5685 | 0.593372207 | 23.7413 | 24.2334 | 26.4741 | 22.1359 | 22.7364 | 24.871 |
| Lysine-specific demethylase 3B | Q7LBC6;Q7LBC6-2 | KDM3B | Extracellular or secreted | 0.5595 | 0.485457087 | 22.1613 | 21.5643 | 21.5876 | 23.1858 | 21.8271 | 21.9788 |
| KH domain-containing, RNA-binding, signal transduction- | Q07666;Q07666-3;Q07666-2 | KHDRBS1 | Extracellular or secreted | 0.7719 | 0.504193571 | 27.3547 | 27.3435 | 28.8451 | 27.7978 | 28.7348 | 29.3264 |
| Far upstream element-binding protein 2 | Q92945 | KHSRP | Other | -0.674966667 | 0.470637633 | 28.7774 | 28.7236 | 29.504 | 27.4019 | 28.334 | 29.2442 |
| Pumilio domain-containing protein KIAA0020 | Q15397 | KIAA0020 | Other | 1.6255 | 0.99131344 | 20.5933 | 21.1413 | 21.0886 | 21.8158 | 22.1422 | 23.7417 |
| WASH complex subunit strumpellin | Q12768 | KIAA0196 | Other | -0.610433333 | 0.235048022 | 21.7916 | 21.3659 | 24.0234 | 21.331 | 21.0716 | 22.947 |

| | | | | | | | | | | | |
|---|---|---|---|---|---|---|---|---|---|---|---|
| Dyslexia-associated protein KIAA0319-like protein | Q8IZA0-2;Q8IZA0 | KIAA0319L | Extracellular or secreted | -0.361033333 | 0.13354677 | 22.219 | 23.8101 | 23.0414 | 21.9721 | 21.6439 | 24.3714 |
| KIF1-binding protein | Q96EK5 | KIAA1279 | Other | 0.430766667 | 0.41223541 | 24.1733 | 24.822 | 25.0426 | 24.7703 | 24.7425 | 25.8174 |
| Protein virilizer homolog | Q69YN4-3;Q69YN4;Q69YN4- | KIAA1429 | Other | -1.589166667 | 0.339030904 | 22.4683 | 20.1974 | 26.0214 | 22.6799 | 20.2256 | 21.0141 |
| LisH domain and HEAT repeat-containing protein KIAA14 | Q9P260;Q9P260-2 | KIAA1468 | Other | -0.5388 | 0.248218402 | 20.9465 | 20.5619 | 23.0209 | 21.3596 | 21.1701 | 20.3832 |
| Uncharacterized protein KIAA1522 | Q9P206;Q9P206-3;Q9P206-2 | KIAA1522 | Other | 0.5579 | 0.460705026 | 24.2118 | 23.5425 | 23.889 | 25.0412 | 23.5558 | 24.72 |
| Shootin-1 | A0MZ66-8;A0MZ66-4;A0MZ( | KIAA1598 | Other | -0.454 | 1.014051682 | 24.543 | 24.2857 | 24.0497 | 23.5919 | 24.1216 | 23.8029 |
| Kinesin-like protein KIF11 | P52732 | KIF11 | Other | 2.122766667 | 0.844108807 | 24.9195 | 21.8255 | 23.8656 | 25.6987 | 25.7872 | 25.493 |
| Kinesin-like protein KIF20B | Q96Q89-4;Q96Q89-3;Q96Q8 | KIF20B | Other | -1.7392 | 0.200769869 | 28.4996 | 27.7244 | 28.36 | 20.5079 | 28.0216 | 30.8369 |
| Kinesin-like protein KIF23 | Q02241-2;Q02241;Q02241-3 | KIF23 | Extracellular or secreted | -0.306433333 | 0.726758116 | 25.04 | 24.7743 | 25.1359 | 24.3841 | 24.9082 | 24.7386 |
| Kinesin-like protein KIF2C | Q99661-2;Q99661 | KIF2C | Other | 0.0388 | 0.009874076 | 22.9745 | 20.3037 | 23.2758 | 23.7174 | 22.2967 | 20.6563 |
| Kinesin-1 heavy chain | P33176 | KIF5B | Extracellular or secreted | -0.947233333 | 1.201736939 | 29.8442 | 29.7962 | 30.2068 | 28.5676 | 28.8948 | 29.5431 |
| Kin of IRRE-like protein 1 | Q96J84;Q96J84-2;Q96J84-3 | KIRREL | Extracellular or secreted | 0.101633333 | 0.053143501 | 25.4615 | 24.7493 | 23.4947 | 24.924 | 24.9971 | 24.0893 |
| Kinesin light chain 1 | Q07866-8;Q07866-2;Q07866 | KLC1 | Other | -0.6337 | 1.250674161 | 27.1677 | 27.4649 | 27.638 | 26.4201 | 26.9878 | 26.9616 |
| Kinesin light chain 2 | Q9H0B6;Q9H0B6-2 | KLC2 | Other | -1.9478 | 0.692415311 | 22.8596 | 24.2238 | 25.0035 | 20.3119 | 21.9755 | 23.9561 |
| Kelch domain-containing protein 7B | Q96G42 | KLHDC7B | Other | -1.0015 | 0.433937313 | 21.5692 | 23.4438 | 24.482 | 21.4957 | 22.8286 | 22.1662 |
| Kallikrein-6 | Q92876-2;Q92876 | KLK6 | Extracellular or secreted | 0.6979 | 0.271883129 | 21.3323 | 21.2508 | 22.7212 | 20.7057 | 23.3665 | 23.3258 |
| Kinetochore-associated protein 1 | P50748 | KNTC1 | Extracellular or secreted | -0.328266667 | 0.089685564 | 20.4517 | 22.5306 | 24.6119 | 22.5211 | 21.585 | 22.5033 |
| Importin subunit alpha-5;Importin subunit alpha-5, N-ter | P52294 | KPNA1 | Other | -0.0226 | 0.022650698 | 26.9053 | 26.8853 | 27.8115 | 27.2662 | 26.9875 | 27.2806 |
| Importin subunit alpha-1 | P52292 | KPNA2 | Other | -0.5602 | 1.325985496 | 27.5195 | 27.9588 | 27.987 | 27.0201 | 27.4034 | 27.3612 |
| Importin subunit alpha-4 | O00505 | KPNA3 | Other | -1.3197 | 0.855667138 | 26.5742 | 26.7542 | 27.6413 | 25.2781 | 24.9124 | 26.8201 |
| Importin subunit alpha-3 | O00629 | KPNA4 | Extracellular or secreted | -0.273633333 | 0.320766604 | 27.2737 | 27.2548 | 27.9598 | 26.8498 | 27.0917 | 27.7259 |
| Importin subunit alpha-7 | O60684 | KPNA6 | Other | 1.560866667 | 0.542025562 | 21.7406 | 20.52 | 23.6005 | 21.7497 | 24.0636 | 24.7304 |
| Importin subunit beta-1 | Q14974;Q14974-2 | KPNB1 | Extracellular or secreted | -0.859466667 | 0.750407676 | 31.2075 | 31.8862 | 32.1728 | 30.1011 | 31.0415 | 31.5455 |
| Keratinocyte proline-rich protein | Q5T749 | KPRP | Extracellular or secreted | 1.854466667 | 0.805799828 | 25.6974 | 24.7013 | 27.6824 | 27.5835 | 27.3611 | 28.6999 |
| GTPase KRas;GTPase KRas, N-terminally processed | P01116-2;P01116 | KRAS | Extracellular or secreted | 0.9255 | 2.057330226 | 25.1599 | 25.625 | 25.5049 | 26.5729 | 26.1112 | 26.3822 |
| Keratin, type I cytoskeletal 18 | P05783 | KRT18 | Extracellular or secreted | -0.003733333 | 0.001833987 | 31.7699 | 29.6982 | 31.0756 | 30.8964 | 30.4879 | 31.1482 |
| Keratin, type II cytoskeletal 6B | P04259 | KRT6B | Extracellular or secreted | -1.953033333 | 0.531815454 | 27.1334 | 22.634 | 24.6766 | 21.9439 | 21.9475 | 24.6935 |
| Kinectin | Q86UP2;Q86UP2-2;Q86UP2- | KTN1 | Extracellular or secreted | 0.269433333 | 0.189842137 | 30.0623 | 29.4851 | 31.1979 | 30.5175 | 30.5026 | 30.5335 |
| Kynureninase | Q16719;Q16719-2 | KYNU | Other | 0.7147 | 0.913274318 | 33.7728 | 33.57 | 32.8851 | 34.3008 | 34.4346 | 33.6366 |
| Neural cell adhesion molecule L1 | P32004-3;P32004-2;P32004 | L1CAM | Extracellular or secreted | -0.3692 | 0.238259514 | 32.5076 | 32.4171 | 31.2523 | 32.5217 | 31.5977 | 30.95 |
| Beta-lactamase-like protein 2 | Q53H82 | LACTB2 | Other | -0.529733333 | 0.641082593 | 29.3571 | 28.9168 | 28.5948 | 28.5801 | 28.8414 | 27.858 |
| Ladinin-1 | O00515 | LAD1 | Extracellular or secreted | 1.734333333 | 1.015695302 | 25.3651 | 24.8862 | 26.8785 | 27.554 | 27.2017 | 27.5771 |
| EKC/KEOPS complex subunit LAGE3 | Q14657 | LAGE3 | Other | 0.738833333 | 0.480859034 | 22.2575 | 21.5527 | 21.5387 | 21.8162 | 22.0867 | 23.6625 |
| Laminin subunit alpha-4 | Q16363-2;Q16363 | LAMA4 | Extracellular or secreted | 0.6691 | 0.336594361 | 31.5273 | 29.8687 | 29.0565 | 31.2896 | 30.7445 | 30.4257 |
| Laminin subunit alpha-5 | O15230 | LAMA5 | Extracellular or secreted | 0.7818 | 1.191406404 | 31.7627 | 31.9422 | 31.137 | 32.1058 | 32.4605 | 32.621 |
| Laminin subunit beta-1 | P07942 | LAMB1 | Extracellular or secreted | 0.615966667 | 0.317387425 | 32.6892 | 32.4022 | 30.4146 | 32.7182 | 32.4022 | 32.2335 |
| Laminin subunit beta-2 | P55268 | LAMB2 | Extracellular or secreted | 0.099933333 | 0.08756572 | 29.745 | 30.6814 | 29.7138 | 29.7333 | 30.1107 | 30.596 |
| Laminin subunit beta-3 | Q13751 | LAMB3 | Extracellular or secreted | 0.218566667 | 0.084676397 | 31.8367 | 33.4373 | 31.0094 | 31.2052 | 32.6452 | 33.0887 |
| Laminin subunit gamma-1 | P11047 | LAMC1 | Extracellular or secreted | 0.5704 | 0.385833664 | 32.7336 | 32.862 | 31.1416 | 32.9911 | 32.7334 | 32.7239 |
| Laminin subunit gamma-2 | Q13753;Q13753-2 | LAMC2 | Extracellular or secreted | 0.8599 | 0.373391725 | 31.3079 | 33.1118 | 30.2057 | 31.7728 | 32.589 | 32.8433 |
| Lysosome-associated membrane glycoprotein 1 | P11279;P11279-2 | LAMP1 | Extracellular or secreted | 0.087866667 | 0.090469503 | 29.6475 | 29.6182 | 29.492 | 29.9208 | 30.0666 | 29.0339 |
| Lysosome-associated membrane glycoprotein 2 | P13473-2;P13473;P13473-3 | LAMP2 | Extracellular or secreted | 0.7738 | 1.122091471 | 26.7222 | 26.5084 | 26.5506 | 27.8378 | 27.2192 | 27.0456 |
| Ragulator complex protein LAMTOR1 | Q6IAA8 | LAMTOR1 | Extracellular or secreted | -0.898666667 | 0.27324054 | 23.891 | 24.4896 | 25.2325 | 21.2922 | 25.043 | 24.5819 |
| Ragulator complex protein LAMTOR2 | Q9Y2Q5;Q9Y2Q5-2 | LAMTOR2 | Extracellular or secreted | 0.070966667 | 0.068003984 | 24.2646 | 24.6172 | 25.162 | 24.2451 | 24.9757 | 25.0359 |
| Ragulator complex protein LAMTOR3 | Q9UHA4;Q9UHA4-2 | LAMTOR3 | Extracellular or secreted | -1.224566667 | 0.442226118 | 24.9612 | 25.2182 | 26.1634 | 22.1815 | 25.5586 | 24.929 |
| LanC-like protein 1 | O43813 | LANCL1 | Extracellular or secreted | -1.624366667 | 0.879010388 | 25.1733 | 26.7727 | 25.05 | 22.8778 | 25.1286 | 24.1165 |
| LanC-like protein 2 | Q9NS86 | LANCL2 | Extracellular or secreted | -0.494966667 | 0.316795941 | 22.9455 | 21.9731 | 22.4399 | 20.8573 | 22.4818 | 22.5345 |
| Cytosol aminopeptidase | P28838-2;P28838 | LAP3 | Extracellular or secreted | 0.4195 | 0.279065922 | 31.4475 | 30.7971 | 30.4141 | 32.1677 | 31.3445 | 30.405 |
| La-related protein 1 | Q6PKG0;Q6PKG0-3 | LARP1 | Other | 0.2602 | 0.638495298 | 25.4321 | 25.2622 | 25.7381 | 25.8358 | 25.5012 | 25.876 |
| La-related protein 7 | Q4G0J3;Q4G0J3-3;Q4G0J3-2 | LARP7 | Other | -2.106933333 | 1.832573994 | 25.4558 | 25.2464 | 24.2199 | 22.5822 | 22.4887 | 23.5304 |
| Leucine--tRNA ligase, cytoplasmic | Q9P2J5;Q9P2J5-2 | LARS | Other | -0.6491 | 0.341267046 | 27.5203 | 27.0371 | 29.0265 | 26.263 | 27.3796 | 27.994 |
| LIM and SH3 domain protein 1 | Q14847;Q14847-2;Q14847-3 | LASP1 | Extracellular or secreted | 0.649233333 | 1.195726247 | 28.3025 | 28.8448 | 28.9528 | 29.3273 | 29.127 | 29.5935 |
| Lamin-B receptor | Q14739 | LBR | Other | 1.254733333 | 0.558258566 | 25.2776 | 24.8168 | 27.5576 | 27.2522 | 27.0876 | 27.0764 |
| Leucine carboxyl methyltransferase 1 | Q9UIC8-3;Q9UIC8;Q9UIC8-2 | LCMT1 | Other | -0.492166667 | 0.393619937 | 21.9942 | 22.9281 | 21.3264 | 21.8848 | 21.5344 | 21.353 |
| Neutrophil gelatinase-associated lipocalin | P80188 | LCN2 | Extracellular or secreted | 2.125566667 | 0.968070937 | 34.9776 | 35.7923 | 33.0376 | 36.9907 | 37.0845 | 36.109 |
| Neutrophil gelatinase-associated lipocalin | P80188-2 | LCN2 | Extracellular or secreted | 1.851966667 | 1.891366481 | 21.1123 | 21.2194 | 22.1055 | 23.579 | 23.6711 | 22.743 |
| Plastin-2 | P13796 | LCP1 | Extracellular or secreted | 0.721833333 | 1.206014052 | 27.5866 | 28.1737 | 27.4535 | 28.384 | 28.7523 | 28.243 |
| L-lactate dehydrogenase A chain | P00338;P00338-3;P00338-5; | LDHA | Extracellular or secreted | -0.097 | 0.09339633 | 35.5298 | 35.1063 | 34.8068 | 35.5554 | 35.0858 | 34.5107 |
| L-lactate dehydrogenase B chain | P07195 | LDHB | Extracellular or secreted | -0.033066667 | 0.026051895 | 35.5952 | 35.2623 | 34.8136 | 35.8409 | 35.1112 | 34.6198 |
| Low-density lipoprotein receptor | P01130-3;P01130-5;P01130; | LDLR | Extracellular or secreted | 2.283833333 | 2.711322925 | 29.2352 | 28.5368 | 28.6124 | 31.4055 | 31.1841 | 30.6463 |
| Leukocyte cell-derived chemotaxin-2 | O14960 | LECT2 | Extracellular or secreted | -0.5539 | 0.180094535 | 23.0009 | 20.0062 | 23.2811 | 22.1397 | 20.7817 | 21.7051 |
| RNA polymerase-associated protein LEO1 | Q8WVC0;Q8WVC0-2 | LEO1 | Other | -0.487633333 | 0.595942952 | 27.55 | 26.6697 | 26.6838 | 26.234 | 26.3035 | 26.9031 |
| Prolyl 3-hydroxylase 1 | Q32P28-4;Q32P28;Q32P28-3 | LEPRE1 | Extracellular or secreted | -0.4498 | 1.252556966 | 26.5869 | 26.3427 | 26.8002 | 25.9484 | 26.253 | 26.179 |
| LETM1 and EF-hand domain-containing protein 1, mitoch | O95202 | LETM1 | Other | 1.068533333 | 0.40160163 | 22.0283 | 20.5024 | 23.892 | 23.0943 | 22.5614 | 23.9726 |
| Beta-1,3-N-acetylglucosaminyltransferase lunatic fringe | Q8NES3-3;Q8NES3 | LFNG | Extracellular or secreted | 1.6916 | 0.392853475 | 24.5355 | 25.9609 | 20.5677 | 24.8998 | 25.6377 | 25.6014 |
| Galectin-1 | P09382 | LGALS1 | Extracellular or secreted | 0.536333333 | 1.017721531 | 32.268 | 32.2183 | 31.6632 | 32.3442 | 32.841 | 32.5733 |
| Galectin-3 | P17931 | LGALS3 | Extracellular or secreted | -0.862333333 | 1.477362594 | 30.7992 | 30.6196 | 30.1519 | 29.5113 | 29.7405 | 29.7319 |
| Galectin-3-binding protein | Q08380 | LGALS3BP | Extracellular or secreted | 0.9437 | 0.562784319 | 31.2668 | 31.6942 | 29.632 | 31.611 | 32.4276 | 31.3855 |
| Galectin-8 | O00214;O00214-2 | LGALS8 | Extracellular or secreted | -0.1728 | 0.185283304 | 25.1542 | 25.3673 | 24.755 | 24.3229 | 25.156 | 25.2792 |
| Legumain | Q99538;Q99538-2;Q99538-3 | LGMN | Extracellular or secreted | 0.513 | 0.358955249 | 30.0697 | 30.2976 | 28.7472 | 30.5662 | 30.5374 | 29.5499 |
| Phospholysine phosphohistidine inorganic pyrophosphat | Q9H008;Q9H008-2 | LHPP | Other | -2.534 | 2.055699675 | 25.429 | 25.2005 | 24.2374 | 22.0967 | 22.7376 | 22.4306 |
| Leukemia inhibitory factor | P15018 | LIF | Extracellular or secreted | 3.837333333 | 1.025018153 | 26.9068 | 26.7335 | 22.7156 | 29.8427 | 29.6846 | 28.3406 |
| DNA ligase 1 | P18858-2;P18858-3;P18858 | LIG1 | Other | -1.1757 | 0.580890699 | 23.5 | 22.6089 | 24.3463 | 23.3766 | 20.9185 | 22.633 |
| LIM domain and actin-binding protein 1 | Q9UHB6-3;Q9UHB6-2;Q9UH | LIMA1 | Extracellular or secreted | 0.606 | 0.120914165 | 21.7063 | 26.0009 | 25.0461 | 22.3248 | 25.7528 | 26.4937 |
| LIM and calponin homology domains-containing protein | Q9UPQ0-9;Q9UPQ0-6 | LIMCH1 | Other | 2.921433333 | 0.582725338 | 21.5858 | 22.1413 | 22.6225 | 21.2831 | 27.351 | 26.4798 |
| LIM and senescent cell antigen-like-containing domain p | P48059;P48059-4;P48059-2; | LIMS1 | Extracellular or secreted | -0.746833333 | 0.380824283 | 21.9823 | 22.6657 | 22.8547 | 21.3583 | 20.7456 | 23.1583 |
| Protein lin-7 homolog C;Protein lin-7 homolog A | Q9NUP9;O14910 | LIN7C;LIN7A | Extracellular or secreted | 0.195133333 | 0.165046911 | 26.2897 | 26.5534 | 26.9643 | 26.0235 | 27.1958 | 27.1735 |
| Lysosomal acid lipase/cholesteryl ester hydrolase | P38571-2;P38571 | LIPA | Extracellular or secreted | 0.587233333 | 0.526782593 | 24.6374 | 25.0636 | 25.0782 | 24.7532 | 25.5759 | 26.2118 |
| Protein ERGIC-53 | P49257 | LMAN1 | Extracellular or secreted | -1.8561 | 1.891636786 | 27.8584 | 28.7211 | 27.6647 | 26.383 | 25.6869 | 26.606 |
| Vesicular integral-membrane protein VIP36 | Q12907 | LMAN2 | Extracellular or secreted | -0.343866667 | 0.35383035 | 29.1721 | 28.3371 | 28.0013 | 28.1031 | 28.4828 | 27.893 |
| Prelamin-A/C;Lamin-A/C | P02545;P02545-6;P02545-3; | LMNA | Other | 0.110833333 | 0.0742202 | 32.397 | 32.1172 | 33.4594 | 32.1248 | 33.1742 | 33.0071 |
| Prelamin-A/C;Lamin-A/C | P02545-2 | LMNA | Other | -0.714933333 | 0.594565547 | 29.4636 | 28.9776 | 29.5809 | 27.7427 | 28.8655 | 29.2691 |
| Lamin-B1 | P20700 | LMNB1 | Other | -0.2923 | 0.167833248 | 30.5397 | 29.4631 | 31.5066 | 30.0062 | 30.6528 | 29.9735 |
| Lamin-B2 | Q03252 | LMNB2 | Other | -0.509033333 | 0.330878867 | 28.597 | 27.7444 | 29.6918 | 27.8539 | 28.4769 | 28.1753 |
| LIM domain only protein 7 | Q8WWI1-3;Q8WWI1-4;Q8W | LMO7 | Other | -0.1568 | 0.106383327 | 26.2119 | 25.6544 | 26.1511 | 25.5074 | 25.2422 | 26.7974 |
| Lon protease homolog, mitochondrial | P36776-2;P36776;P36776-3 | LONP1 | Other | -2.823666667 | 1.142440772 | 26.7638 | 26.1191 | 29.0288 | 24.2179 | 24.1226 | 25.1002 |
| Protein-lysine 6-oxidase | P28300 | LOX | Extracellular or secreted | 0.606233333 | 0.290443223 | 28.7966 | 29.1135 | 26.6838 | 28.4974 | 28.8585 | 29.0567 |
| Lysyl oxidase homolog 1 | Q08397 | LOXL1 | Extracellular or secreted | 1.4846 | 0.290499152 | 26.1014 | 26.3359 | 20.6089 | 26.1709 | 26.1665 | 25.1626 |
| Lysyl oxidase homolog 2 | Q9Y4K0 | LOXL2 | Extracellular or secreted | 1.7475 | 1.488190554 | 31.357 | 31.5923 | 30.3019 | 32.5171 | 32.8629 | 33.1137 |
| Lipoma-preferred partner | Q93052 | LPP | Extracellular or secreted | -0.3722 | 0.162067929 | 23.7093 | 24.725 | 23.0526 | 22.8547 | 22.6564 | 24.8592 |
| Lipopolysaccharide-responsive and beige-like anchor pro | P50851;P50851-2 | LRBA | Extracellular or secreted | 0.1347 | 0.078540049 | 26.2428 | 25.7351 | 27.4896 | 26.4229 | 26.2661 | 27.1826 |
| Leucine-rich alpha-2-glycoprotein | P02750 | LRG1 | Extracellular or secreted | 1.833333333 | 0.869882725 | 27.5548 | 28.3217 | 25.6305 | 29.3392 | 29.4145 | 28.2533 |
| Prolow-density lipoprotein receptor-related protein 1;Lo | Q07954 | LRP1 | Extracellular or secreted | 0.145966667 | 0.244760776 | 27.1961 | 26.7955 | 26.5688 | 27.2882 | 26.8132 | 26.8969 |
| Low-density lipoprotein receptor-related protein 6 | O75581 | LRP6 | Extracellular or secreted | 0.1194 | 0.024810048 | 24.8305 | 25.3137 | 20.6096 | 24.0544 | 24.069 | 22.9886 |
| Low-density lipoprotein receptor-related protein 8 | Q14114-3;Q14114;Q14114-2 | LRP8 | Extracellular or secreted | 1.8715 | 1.572576841 | 26.7478 | 26.7993 | 25.6275 | 28.4135 | 27.9628 | 28.4128 |
| Alpha-2-macroglobulin receptor-associated protein | P30533 | LRPAP1 | Extracellular or secreted | 0.2065 | 0.148720096 | 25.0157 | 24.834 | 25.9277 | 26.09 | 25.5584 | 24.7485 |
| Leucine-rich PPR motif-containing protein, mitochondria | P42704 | LRPPRC | Other | -0.620866667 | 0.261619381 | 27.4895 | 27.7835 | 30.2077 | 28.1117 | 27.9055 | 27.6009 |
| Leucine-rich repeat-containing protein 1 | Q9BTT6;Q9BTT6-2 | LRRC1 | Other | 1.032333333 | 0.812484751 | 21.7841 | 21.1831 | 21.1245 | 21.431 | 22.9228 | 22.8349 |
| Leucine-rich repeat-containing protein 16A | Q5VZK9-2;Q5VZK9 | LRRC16A | Extracellular or secreted | -1.067966667 | 0.648159636 | 25.6063 | 24.1959 | 26.2343 | 24.196 | 23.6227 | 25.0139 |
| Leucine-rich repeat-containing protein 40 | Q9H9A6 | LRRC40 | Other | -0.415066667 | 0.251924259 | 22.32 | 22.2947 | 23.5989 | 23.2934 | 21.716 | 21.959 |
| Leucine-rich repeat-containing protein 47 | Q8N1G4 | LRRC47 | Other | -0.3252 | 0.152853983 | 25.6474 | 26.6946 | 27.2185 | 24.9187 | 26.7567 | 26.9095 |
| Leucine-rich repeat-containing protein 59 | Q96AG4 | LRRC59 | Other | -1.364766667 | 0.975869376 | 27.4506 | 27.9812 | 29.2186 | 26.6051 | 26.6572 | 27.2938 |
| Leucine-rich repeat flightless-interacting protein 1 | Q32MZ4-3;Q32MZ4-2;Q32M | LRRFIP1 | Extracellular or secreted | 0.396633333 | 0.297130043 | 28.764 | 27.6467 | 29.0137 | 29.52 | 28.3839 | 28.7104 |
| Leucine-rich repeat flightless-interacting protein 2 | Q9Y608-2;Q9Y608-4;Q9Y608 | LRRFIP2 | Other | 0.477266667 | 1.023134778 | 25.0173 | 24.4436 | 24.5536 | 25.2001 | 25.2777 | 24.9685 |
| Leucine-rich repeat serine/threonine-protein kinase 1 | Q38SD2 | LRRK1 | Other | -4.041833333 | 1.523717568 | 26.5587 | 24.7697 | 25.487 | 21.0413 | 23.3748 | 20.2738 |
| U6 snRNA-associated Sm-like protein LSm1 | O15116 | LSM1 | Other | 0.527066667 | 0.220696376 | 20.8798 | 22.3583 | 22.3448 | 22.4517 | 23.6975 | 21.0149 |
| U6 snRNA-associated Sm-like protein LSm2 | Q9Y333 | LSM2 | Other | -0.167566667 | 0.217951822 | 26.537 | 26.8126 | 26.334 | 25.9029 | 26.7714 | 26.5066 |
| U6 snRNA-associated Sm-like protein LSm4 | Q9Y4Z0 | LSM4 | Other | -0.3715 | 0.896125154 | 26.7698 | 26.7307 | 26.9545 | 26.155 | 26.6926 | 26.4929 |
| U6 snRNA-associated Sm-like protein LSm5 | Q9Y4Y9 | LSM5 | Other | 0.493833333 | 0.322964565 | 23.3373 | 25.1342 | 24.6516 | 25.4265 | 24.6935 | 24.4846 |
| U6 snRNA-associated Sm-like protein LSm6 | P62312 | LSM6 | Extracellular or secreted | -0.2969 | 0.467034895 | 26.5082 | 26.352 | 26.5925 | 25.7321 | 26.2847 | 26.5452 |
| U6 snRNA-associated Sm-like protein LSm7 | Q9UK45 | LSM7 | Other | -0.246366667 | 0.39421727 | 26.0954 | 26.4759 | 26.8118 | 25.9849 | 26.5245 | 26.1346 |
| U6 snRNA-associated Sm-like protein LSm8 | O95777 | LSM8 | Other | 0.4353 | 0.521448767 | 27.7393 | 27.9434 | 27.8073 | 28.6214 | 28.5395 | 27.635 |
| Lipolysis-stimulated lipoprotein receptor | Q86X29-6;Q86X29-5;Q86X29 | LSR | Extracellular or secreted | 1.260233333 | 1.035162214 | 29.4992 | 29.4545 | 28.1106 | 30.6925 | 30.1693 | 29.9832 |
| Lipolysis-stimulated lipoprotein receptor | Q86X29-3 | LSR | Extracellular or secreted | 0.460133333 | 0.129615475 | 24.5095 | 21.4305 | 23.6859 | 23.8044 | 25.1963 | 22.0056 |
| Lanosterol synthase | P48449-3;P48449;P48449-2 | LSS | Other | 1.919033333 | 1.077531216 | 21.5044 | 21.9299 | 23.6082 | 23.787 | 23.7093 | 25.3033 |
| Leukotriene A-4 hydrolase | P09960;P09960-4;P09960-2; | LTA4H | Extracellular or secreted | -0.1029 | 0.08305991 | 32.2266 | 31.8018 | 31.1967 | 32.1999 | 31.6266 | 31.0899 |
| Latent-transforming growth factor beta-binding protein | Q14766-2;Q14766-5;Q14766 | LTBP1 | Extracellular or secreted | 0.912066667 | 0.497068512 | 26.8722 | 27.0446 | 25.358 | 26.3064 | 27.3414 | 28.3632 |
| Latent-transforming growth factor beta-binding protein | Q14767 | LTBP2 | Extracellular or secreted | 1.668733333 | 0.660851705 | 27.0739 | 27.9991 | 24.827 | 28.3378 | 28.3225 | 28.2459 |
| Latent-transforming growth factor beta-binding protein | Q9NS15;Q9NS15-2 | LTBP3 | Extracellular or secreted | 2.2496 | 1.946354737 | 29.2016 | 29.3103 | 28.2214 | 30.5551 | 31.1017 | 31.8253 |

| | | | | | | | | | | | |
|---|---|---|---|---|---|---|---|---|---|---|---|
| Latent-transforming growth factor beta-binding protein | Q8N2S1-2;Q8N2S1-3;Q8N2S | LTBP4 | Extracellular or secreted | -0.400633333 | 0.55455051 | 28.7727 | 29.2804 | 28.3348 | 28.393 | 28.4713 | 28.3217 |
| Putative RNA-binding protein Luc7-like 2 | Q9Y383;Q96HJ9-2;Q9Y383-3 | LUC7L2 | Other | -0.860166667 | 1.996477112 | 27.9505 | 27.9752 | 27.5392 | 27.0577 | 26.7349 | 27.0918 |
| Luc7-like protein 3 | O95232 | LUC7L3 | Other | -0.9257 | 0.668206551 | 25.5414 | 26.1381 | 26.1725 | 24.0174 | 25.2563 | 25.8012 |
| Leucine zipper protein 1 | Q86V48-2;Q86V48-3;Q86V4 | LUZP1 | Extracellular or secreted | -1.134766667 | 0.875978168 | 25.8561 | 25.0184 | 26.6258 | 24.8615 | 23.9838 | 25.2507 |
| Latexin | Q9BS40 | LXN | Extracellular or secreted | -0.415633333 | 0.39448178 | 28.7944 | 28.7655 | 28.6056 | 27.5614 | 28.451 | 28.9062 |
| Cell growth-regulating nucleolar protein | Q9NX58 | LYAR | Other | -1.420666667 | 1.128314529 | 25.814 | 25.1669 | 26.7838 | 24.2852 | 24.1181 | 25.0994 |
| Ly6/PLAUR domain-containing protein 3 | O95274 | LYPD3 | Extracellular or secreted | 1.394966667 | 1.215601348 | 28.3787 | 29.1632 | 27.6897 | 30.2606 | 29.8347 | 29.3212 |
| Acyl-protein thioesterase 1 | O75608-2;O75608 | LYPLA1 | Extracellular or secreted | -0.925866667 | 0.290487427 | 24.9286 | 27.0751 | 26.436 | 23.0473 | 26.3863 | 26.2285 |
| Acyl-protein thioesterase 2 | O95372 | LYPLA2 | Extracellular or secreted | 0.140366667 | 0.076784498 | 25.3461 | 26.7117 | 27.0851 | 25.8687 | 26.599 | 27.0963 |
| Lysophospholipase-like protein 1 | Q5VWZ2;Q5VWZ2-2 | LYPLAL1 | Extracellular or secreted | -2.4134 | 0.628693009 | 24.6043 | 26.295 | 25.1932 | 24.6307 | 24.1812 | 20.0404 |
| Lysozyme C | P61626 | LYZ | Extracellular or secreted | 0.726733333 | 0.49606742 | 25.1152 | 23.661 | 24.9119 | 26.1686 | 25.0116 | 24.6881 |
| Protein LZIC | Q8WZA0;Q8WZA0-2 | LZIC | Other | -0.174333333 | 0.142039714 | 26.2105 | 24.8559 | 25.7738 | 25.5298 | 25.715 | 25.0724 |
| Cation-dependent mannose-6-phosphate receptor | P20645 | M6PR | Extracellular or secreted | 0.369533333 | 0.298991146 | 24.5986 | 24.5972 | 25.2089 | 26.0173 | 24.9383 | 24.5577 |
| Mitotic spindle assembly checkpoint protein MAD1 | Q9Y6D9;Q9Y6D9-3 | MAD1L1 | Extracellular or secreted | -1.535433333 | 0.587645177 | 25.7657 | 24.3693 | 27.2911 | 23.0434 | 23.9887 | 25.7877 |
| Mitotic spindle assembly checkpoint protein MAD2A | Q13257 | MAD2L1 | Other | -1.3439 | 1.484622932 | 27.8 | 26.8483 | 27.8379 | 25.7708 | 26.263 | 26.4207 |
| Macrophage erythroblast attacher | Q7L5Y9-3;Q7L5Y9-2;Q7L5Y9 | MAEA | Extracellular or secreted | -1.162233333 | 0.586574959 | 22.5345 | 24.7931 | 24.7932 | 22.7073 | 22.4947 | 23.4321 |
| Melanoma-associated antigen D2 | Q9UNF1-2;Q9UNF1 | MAGED2 | Other | -2.952566667 | 1.444619923 | 24.5721 | 23.1341 | 24.9919 | 22.7337 | 20.6835 | 20.4232 |
| Protein mago nashi homolog | P61326 | MAGOH | Other | -0.6094 | 0.733657629 | 27.8549 | 27.8273 | 28.6834 | 27.0328 | 27.907 | 27.5976 |
| Protein MAK16 homolog | Q9BXY0 | MAK16 | Other | -1.398533333 | 0.748407963 | 23.4541 | 21.8453 | 24.2348 | 21.5499 | 22.1602 | 21.6285 |
| Mannosyl-oligosaccharide 1,2-alpha-mannosidase IA | P33908;P33908-2 | MAN1A1 | Extracellular or secreted | 0.2183 | 0.089795838 | 25.6971 | 27.2389 | 24.508 | 25.6738 | 26.5094 | 25.9157 |
| Endoplasmic reticulum mannosyl-oligosaccharide 1,2-alp | Q9UKM7 | MAN1B1 | Extracellular or secreted | -0.469066667 | 0.407317572 | 25.764 | 25.0646 | 24.395 | 24.4527 | 25.1403 | 24.2234 |
| Alpha-mannosidase 2 | Q16706 | MAN2A1 | Extracellular or secreted | 0.99 | 0.229395853 | 27.3259 | 27.8748 | 22.9911 | 27.4769 | 27.23 | 26.4549 |
| Lysosomal alpha-mannosidase;Lysosomal alpha-mannos | O00754-2;O00754 | MAN2B1 | Extracellular or secreted | 1.1053 | 1.689992737 | 28.9551 | 29.1313 | 28.4211 | 30.2262 | 30.0561 | 29.5411 |
| Epididymis-specific alpha-mannosidase | Q9Y2E5;Q9Y2E5-2 | MAN2B2 | Extracellular or secreted | 0.7967 | 0.79933147 | 27.0455 | 26.6665 | 26.32 | 27.7519 | 27.9437 | 26.7265 |
| Alpha-mannosidase 2C1 | Q9NTJ4-2;Q9NTJ4;Q9NTJ4-4 | MAN2C1 | Other | -0.654733333 | 0.351582524 | 26.3478 | 25.8711 | 25.6785 | 26.1352 | 25.8533 | 23.9447 |
| Beta-mannosidase | O00462 | MANBA | Extracellular or secreted | 1.103833333 | 0.945346314 | 26.7353 | 26.8621 | 25.4819 | 27.6707 | 27.6538 | 27.0663 |
| Mesencephalic astrocyte-derived neurotrophic factor | P55145 | MANF | Extracellular or secreted | -0.531666667 | 0.370193211 | 30.597 | 30.397 | 30.0723 | 30.7731 | 29.7691 | 28.9291 |
| MANSC domain-containing protein 1 | Q9H8J5;Q9H8J5-2 | MANSC1 | Extracellular or secreted | 0.576233333 | 0.188953855 | 23.3158 | 20.2469 | 21.8134 | 23.0756 | 23.1706 | 20.8586 |
| Microtubule-associated protein 1B;MAP1B heavy chain; | P46821 | MAP1B | Extracellular or secreted | 0.730566667 | 1.118415154 | 22.6133 | 22.5887 | 22.7557 | 23.3209 | 23.0288 | 23.7997 |
| Microtubule-associated proteins 1A/1B light chain 3B;Mi | Q9GZQ8;A6NCE7;Q9H492;Q | MAP1LC3B;MA | Other | 0.394466667 | 0.371471122 | 27.0441 | 26.4211 | 26.2503 | 27.6137 | 26.9333 | 26.3519 |
| Dual specificity mitogen-activated protein kinase kinase | Q02750;Q02750-2 | MAP2K1 | Extracellular or secreted | -1.275933333 | 0.276702623 | 29.5024 | 28.9796 | 24.8241 | 25.7103 | 28.644 | 25.124 |
| Dual specificity mitogen-activated protein kinase kinase | P46734-2;P46734;P46734-3 | MAP2K3 | Other | -1.6395 | 0.37220082 | 26.0736 | 25.6716 | 25.5249 | 24.3913 | 21.1384 | 26.8219 |
| Microtubule-associated protein 4 | P27816;P27816-6;P27816-2 | MAP4 | Extracellular or secreted | -0.081566667 | 0.130346794 | 30.0037 | 29.3824 | 30.0355 | 29.6399 | 29.7504 | 29.7866 |
| Mitogen-activated protein kinase kinase kinase kinase 4 | O95819-2;O95819-6;O95819 | MAP4K4 | Other | 1.0752 | 0.510841239 | 28.8392 | 26.7021 | 26.4685 | 29.3138 | 27.6165 | 28.3051 |
| Ensconsin | Q14244-5;Q14244-3;Q14244 | MAP7 | Extracellular or secreted | 0.367133333 | 0.076194237 | 23.4463 | 22.8456 | 25.0745 | 21.1437 | 25.6399 | 25.6842 |
| MAP7 domain-containing protein 1 | Q3KQU3-4;Q3KQU3;Q3KQU | MAP7D1 | Other | 0.0178 | 0.025438956 | 26.6378 | 26.3875 | 26.2898 | 26.7261 | 26.5878 | 26.0546 |
| Mitogen-activated protein kinase 1 | P28482-2;P28482 | MAPK1 | Extracellular or secreted | -0.6571 | 1.369163299 | 27.5233 | 28.025 | 27.9189 | 26.8615 | 27.422 | 27.2124 |
| Mitogen-activated protein kinase 14 | Q16539;Q16539-2;Q16539-5 | MAPK14 | Extracellular or secreted | -0.7125 | 0.902659816 | 26.8758 | 26.8884 | 26.6063 | 25.4989 | 26.2894 | 26.4447 |
| MAPK-interacting and spindle-stabilizing protein-like | Q8NDC0 | MAPK1IP1L | Other | 2.383266667 | 1.533824834 | 21.2939 | 22.0178 | 21.346 | 24.7141 | 22.9458 | 24.1476 |
| Mitogen-activated protein kinase 9 | P45984-3;P45984-2;P45984- | MAPK9 | Other | -0.511166667 | 0.805933688 | 23.6555 | 23.453 | 23.7584 | 22.6412 | 23.2593 | 23.4329 |
| MAP kinase-activated protein kinase 3 | Q16644 | MAPKAPK3 | Other | -0.511533333 | 0.684749542 | 22.0283 | 22.0448 | 21.9319 | 22.0421 | 21.1428 | 21.2855 |
| Microtubule-associated protein RP/EB family member 1 | Q15691 | MAPRE1 | Extracellular or secreted | -0.310466667 | 0.474998965 | 30.5579 | 30.3566 | 30.1585 | 29.5724 | 30.1989 | 30.3703 |
| Myristoylated alanine-rich C-kinase substrate | P29966 | MARCKS | Extracellular or secreted | 1.558066667 | 1.850784312 | 28.6138 | 28.0367 | 28.6418 | 30.5488 | 29.7977 | 29.62 |
| MARCKS-related protein | P49006 | MARCKSL1 | Extracellular or secreted | 3.654266667 | 0.984807653 | 21.4501 | 24.9865 | 20.4778 | 26.9179 | 25.6367 | 25.3226 |
| Methionine--tRNA ligase, cytoplasmic | P56192;P56192-2 | MARS | Extracellular or secreted | -1.004433333 | 0.595075947 | 27.8848 | 27.5004 | 29.5588 | 26.6918 | 27.3785 | 27.8604 |
| S-adenosylmethionine synthase isoform type-2 | P31153;P31153-2 | MAT2A | Other | 0.181433333 | 0.11891139 | 30.744 | 30.3845 | 29.6336 | 31.27 | 30.2873 | 29.7491 |
| Methionine adenosyltransferase 2 subunit beta | Q9NZL9;Q9NZL9-2;Q9NZL9-4 | MAT2B | Extracellular or secreted | -0.047866667 | 0.049876741 | 29.7421 | 29.3146 | 28.7665 | 29.1581 | 29.5296 | 28.9919 |
| Matrin-3 | P43243;P43243-2 | MATR3 | Extracellular or secreted | -1.513166667 | 0.863140419 | 27.8766 | 27.4192 | 29.5978 | 26.3371 | 26.4627 | 27.5543 |
| Mitochondrial antiviral-signaling protein | Q7Z434;Q7Z434-4 | MAVS | Other | -0.300466667 | 0.553241783 | 26.7927 | 26.6002 | 26.9722 | 26.2378 | 26.8884 | 26.3375 |
| Protein max | P61244-3;P61244-4;P61244- | MAX | Other | -0.6437 | 0.641306157 | 23.3087 | 23.3615 | 24.1163 | 22.7085 | 22.4852 | 23.6617 |
| Myoglobin | P02144 | MB | Extracellular or secreted | -0.7229 | 0.357731644 | 23.0412 | 23.2042 | 23.0534 | 21.0297 | 23.6362 | 22.4642 |
| Methyl-CpG-binding domain protein 2 | Q9UBB5 | MBD2 | Other | -1.768833333 | 0.810051589 | 23.2012 | 22.4439 | 24.2648 | 22.3448 | 19.9022 | 22.3564 |
| Metallo-beta-lactamase domain-containing protein 1 | A4D2B0 | MBLAC1 | Other | 1.008333333 | 0.79984022 | 21.2661 | 22.1993 | 20.575 | 22.9504 | 22.1793 | 21.9357 |
| Membrane-bound transcription factor site-1 protease | Q14703 | MBTPS1 | Other | 1.048966667 | 0.69659623 | 25.5434 | 26.2602 | 24.3342 | 26.4942 | 26.5147 | 26.2758 |
| Cell surface glycoprotein MUC18 | P43121;P43121-2 | MCAM | Extracellular or secreted | -0.2798 | 0.219278735 | 27.4358 | 26.1874 | 26.1217 | 26.5595 | 26.4998 | 25.8462 |
| DNA replication licensing factor MCM2 | P49736 | MCM2 | Other | -2.335466667 | 0.425110011 | 26.3175 | 26.2114 | 27.5248 | 20.297 | 25.6764 | 27.0739 |
| DNA replication licensing factor MCM3 | P25205;P25205-2 | MCM3 | Other | 0.2 | 0.101483084 | 26.9889 | 26.9805 | 27.7876 | 28.7149 | 26.7818 | 26.8603 |
| DNA replication licensing factor MCM4 | P33991 | MCM4 | Other | -1.043933333 | 0.681924513 | 26.371 | 26.0744 | 27.6973 | 24.8123 | 25.7072 | 26.4914 |
| DNA replication licensing factor MCM5 | P33992 | MCM5 | Other | -0.160633333 | 0.125266967 | 26.5093 | 26.107 | 27.3654 | 25.9672 | 26.5959 | 26.9367 |
| DNA replication licensing factor MCM6 | Q14566 | MCM6 | Other | -0.687733333 | 0.817245438 | 27.7799 | 27.6072 | 28.3118 | 26.7184 | 27.1268 | 27.7905 |
| DNA replication licensing factor MCM7 | P33993;P33993-3 | MCM7 | Other | -0.629566667 | 0.120542688 | 24.7297 | 23.9368 | 27.6657 | 22.114 | 24.9642 | 27.3653 |
| Mini-chromosome maintenance complex-binding protei | Q9BTE3-2;Q9BTE3;Q9BTE3-3 | MCMBP | Extracellular or secreted | -2.106733333 | 2.35493073 | 27.0073 | 26.9222 | 27.5029 | 24.8935 | 24.6633 | 25.5554 |
| Malignant T-cell-amplified sequence 1 | Q9ULC4;Q9ULC4-3;Q9ULC4- | MCTS1 | Extracellular or secreted | -0.5045 | 0.335008395 | 26.4685 | 28.0138 | 27.4199 | 25.9431 | 27.1589 | 27.2867 |
| Mediator of DNA damage checkpoint protein 1 | Q14676-2;Q14676;Q14676-3 | MDC1 | Other | -1.534 | 0.50091436 | 24.816 | 22.6344 | 26.6337 | 23.0245 | 23.1446 | 23.313 |
| Malate dehydrogenase, cytoplasmic | P40925;P40925-3;P40925-2 | MDH1 | Extracellular or secreted | -0.034166667 | 0.025875881 | 32.9364 | 32.5852 | 32.2548 | 33.173 | 32.6547 | 31.8462 |
| Malate dehydrogenase, mitochondrial | P40926;P40926-2 | MDH2 | Extracellular or secreted | -0.944933333 | 0.835026197 | 34.5478 | 34.3308 | 34.5405 | 33.9019 | 33.9807 | 32.7017 |
| Midkine | P21741;P21741-2 | MDK | Extracellular or secreted | 0.706033333 | 0.428384202 | 25.1687 | 24.8542 | 23.4127 | 25.8959 | 25.2974 | 24.3604 |
| NADP-dependent malic enzyme | P48163;P48163-2 | ME1 | Other | 0.537966667 | 0.795403278 | 30.2579 | 30.0792 | 29.6599 | 30.7085 | 30.8549 | 30.0475 |
| NAD-dependent malic enzyme, mitochondrial | P23368;P23368-2 | ME2 | Other | -0.454833333 | 0.371490057 | 28.2383 | 27.4439 | 28.4221 | 28.2098 | 27.7179 | 26.8121 |
| Methyl-CpG-binding protein 2 | P51608-2;P51608 | MECP2 | Extracellular or secreted | 0.6802 | 0.301089345 | 25.0556 | 24.4976 | 26.8434 | 26.8491 | 24.9955 | 26.5926 |
| Mediator of RNA polymerase II transcription subunit 23 | Q9ULK4-2;Q9ULK4-4;Q9ULK | MED23 | Other | -0.860566667 | 0.438326886 | 22.9754 | 21.909 | 21.309 | 22.5403 | 20.385 | 20.6864 |
| Multiple epidermal growth factor-like domains protein 8 | Q7Z7M0-2;Q7Z7M0 | MEGF8 | Extracellular or secreted | 1.263866667 | 0.713171751 | 23.7756 | 21.863 | 21.7885 | 23.6621 | 24.5503 | 23.0063 |
| Protein MEMO1 | Q9Y316;Q9Y316-3;Q9Y316-2 | MEMO1 | Other | -0.195166667 | 0.10291078 | 25.8148 | 27.848 | 26.1711 | 26.2558 | 26.7608 | 26.2318 |
| 7SK snRNA methylphosphate capping enzyme | Q7L2J0 | MEPCE | Other | -0.2999 | 0.182846335 | 22.0713 | 21.8936 | 21.9169 | 22.3996 | 22.0622 | 20.5203 |
| LDLR chaperone MESD | Q14696 | MESDC2 | Extracellular or secreted | -1.0256 | 2.236865105 | 26.6486 | 26.8811 | 26.4251 | 25.6412 | 25.8567 | 25.3801 |
| Hepatocyte growth factor receptor | P08581;P08581-2;P08581-3 | MET | Extracellular or secreted | 1.081133333 | 0.650312192 | 28.0444 | 28.391 | 26.3428 | 28.5791 | 29.0966 | 28.3459 |
| Methionine aminopeptidase 1 | P53582 | METAP1 | Other | -0.423833333 | 1.562874778 | 28.2618 | 27.9592 | 28.0616 | 27.608 | 27.5598 | 27.8433 |
| Methionine aminopeptidase 2 | P50579;P50579-3;P50579-2 | METAP2 | Extracellular or secreted | -0.6473 | 1.074987746 | 28.1423 | 27.5071 | 27.6128 | 27.5132 | 26.9242 | 26.8829 |
| Meteorin-like protein | Q641Q3-2;Q641Q3 | METRNL | Extracellular or secreted | 0.418866667 | 0.146390334 | 24.9186 | 25.4103 | 24.7909 | 23.5026 | 26.2622 | 26.6116 |
| tRNA (guanine-N(7)-)-methyltransferase | Q9UBP6 | METTL1 | Other | -0.0905 | 0.032036935 | 22.449 | 20.8347 | 23.8761 | 22.1236 | 21.9892 | 22.7755 |
| Microfibrillar-associated protein 1 | P55081 | MFAP1 | Extracellular or secreted | -0.695433333 | 0.377801324 | 22.4885 | 21.8879 | 23.1952 | 23.1103 | 21.3734 | 21.0016 |
| Microfibrillar-associated protein 2 | P55001-3;P55001-2;P55001 | MFAP2 | Extracellular or secreted | 1.428333333 | 0.355510164 | 26.2078 | 27.546 | 22.5357 | 26.8367 | 27.0828 | 26.655 |
| Lactadherin;Lactadherin short form;Medin | Q08431-3;Q08431;Q08431-4 | MFGE8 | Extracellular or secreted | 1.717 | 1.497204054 | 26.1154 | 26.3129 | 25.1913 | 27.0576 | 28.3656 | 27.3474 |
| Melanotransferrin | P08582 | MFI2 | Extracellular or secreted | -0.519233333 | 0.247936641 | 27.2493 | 27.2287 | 25.1033 | 26.6371 | 26.0626 | 25.3239 |
| Alpha-1,6-mannosyl-glycoprotein 2-beta-N-acetylglucosa | Q10469 | MGAT2 | Extracellular or secreted | 0.797433333 | 0.37515639 | 21.5255 | 23.3521 | 20.9038 | 21.7756 | 23.2161 | 23.182 |
| Alpha-1,3-mannosyl-glycoprotein 4-beta-N-acetylglucosa | Q9UQ53-2;Q9UQ53;Q9UQ5 | MGAT4B | Extracellular or secreted | -0.945366667 | 0.219842328 | 25.8567 | 25.9844 | 25.7157 | 21.8843 | 26.9634 | 25.873 |
| Alpha-1,6-mannosylglycoprotein 6-beta-N-acetylglucosa | Q09328 | MGAT5 | Extracellular or secreted | -0.7416 | 0.934795623 | 26.5254 | 26.1894 | 25.6077 | 25.4741 | 25.7453 | 24.8783 |
| Protein O-GlcNAcase | O60502-4;O60502;O60502-2 | MGEA5 | Other | -0.969866667 | 0.614015982 | 26.7827 | 26.5279 | 26.5928 | 24.4926 | 26.4259 | 26.0753 |
| Mitochondrial genome maintenance exonuclease 1 | Q9BQP7 | MGME1 | Other | -2.222633333 | 1.003475385 | 25.1356 | 25.0075 | 25.582 | 22.0396 | 24.5729 | 22.4447 |
| Microsomal glutathione S-transferase 1 | P10620;P10620-2 | MGST1 | Other | 2.974 | 1.276664705 | 25.464 | 22.588 | 23.493 | 27.817 | 26.998 | 25.652 |
| Melanoma inhibitory activity protein 3 | Q5JRA6-2;Q5JRA6;Q5JRA6-3 | MIA3 | Extracellular or secreted | 0.125366667 | 0.274926785 | 25.4654 | 25.1882 | 25.3427 | 25.7554 | 25.4002 | 25.2168 |
| MHC class I polypeptide-related sequence A | Q29983 | MICA | Extracellular or secreted | 2.352133333 | 2.037455729 | 24.857 | 23.6297 | 24.5507 | 26.362 | 27.1436 | 26.5882 |
| MICAL-like protein 1 | Q8N3F8 | MICALL1 | Extracellular or secreted | -0.549533333 | 0.290849031 | 22.0434 | 23.198 | 24.4364 | 22.8134 | 22.3968 | 22.819 |
| MHC class I polypeptide-related sequence B | Q29980;Q29980-2 | MICB | Extracellular or secreted | 3.083266667 | 0.807569816 | 25.5261 | 24.0556 | 20.811 | 27.671 | 27.4785 | 24.493 |
| Mesoderm induction early response protein 1 | Q8N108-16;Q8N108-14;Q8N | MIER1 | Extracellular or secreted | -0.8704 | 0.521832853 | 22.4052 | 24.1774 | 24.0835 | 22.0441 | 22.4779 | 23.5329 |
| Macrophage migration inhibitory factor | P14174 | MIF | Extracellular or secreted | 0.1066 | 0.115101293 | 32.1445 | 32.886 | 31.8939 | 32.5579 | 32.5396 | 32.1467 |
| Multiple inositol polyphosphate phosphatase 1 | Q9UNW1;Q9UNW1-4;Q9UN | MINPP1 | Extracellular or secreted | 0.253966667 | 0.455368266 | 27.0314 | 26.5006 | 26.436 | 26.6737 | 27.1766 | 26.8796 |
| Mitochondrial intermediate peptidase | Q99797 | MIPEP | Other | -1.857633333 | 0.457683084 | 21.3759 | 21.2529 | 25.9031 | 21.0024 | 20.9342 | 21.0224 |
| Antigen KI-67 | P46013;P46013-2 | MKI67 | Other | -0.6413 | 0.315056304 | 27.1523 | 25.5636 | 27.5464 | 26.5766 | 26.7836 | 24.9782 |
| Muskelin | Q9UL63-2;Q9UL63 | MKLN1 | Other | -0.050766667 | 0.013062756 | 23.2558 | 23.0467 | 20.6182 | 23.482 | 20.3534 | 22.933 |
| Malectin | Q14165 | MLEC | Extracellular or secreted | -4.413166667 | 1.958975939 | 26.1007 | 25.1248 | 25.6938 | 22.0086 | 19.9348 | 21.7364 |
| Target of rapamycin complex subunit LST8 | Q9BVC4-4;Q9BVC4;Q9BVC4- | MLST8 | Other | 1.018266667 | 0.911997922 | 21.9683 | 21.2475 | 22.6998 | 22.7509 | 22.9183 | 23.3012 |
| Collagenase 3 | P45452 | MMP13 | Extracellular or secreted | 0.759666667 | 0.142523582 | 29.0997 | 30.2115 | 24.2616 | 29.0649 | 28.8852 | 27.9017 |
| Matrix metalloproteinase-14 | P50281 | MMP14 | Extracellular or secreted | 0.025 | 0.01761687 | 26.7534 | 26.2161 | 25.3364 | 26.4077 | 26.2455 | 25.7277 |
| Matrix metalloproteinase-15 | P51511 | MMP15 | Extracellular or secreted | 0.117166667 | 0.106111089 | 25.1191 | 24.3673 | 24.1654 | 24.9295 | 24.9511 | 24.1227 |
| 72 kDa type IV collagenase;PEX | P08253;P08253-3;P08253-2 | MMP2 | Extracellular or secreted | 0.667333333 | 0.400308075 | 30.3586 | 30.9847 | 28.9337 | 30.1842 | 31.1524 | 30.9424 |
| Matrilysin | P09237 | MMP7 | Extracellular or secreted | 1.1377 | 0.320843414 | 25.5669 | 28.422 | 24.6896 | 25.5505 | 28.3796 | 28.1615 |
| Matrix metalloproteinase-9;67 kDa matrix metalloprotei | P14780 | MMP9 | Extracellular or secreted | 0.0882 | 0.022128849 | 31.7528 | 31.4888 | 27.8707 | 30.4287 | 30.3308 | 30.6174 |
| MMS19 nucleotide excision repair protein homolog | Q96T76;Q96T76-8;Q96T76-5 | MMS19 | Other | -1.137066667 | 0.483403644 | 22.3554 | 21.9775 | 24.7244 | 22.3133 | 20.9222 | 22.4106 |
| MOB kinase activator 1A;MOB kinase activator 1B | Q9H8S9;Q7L9L4;Q7L9L4-2;Q | MOB1A;MOB1 | Extracellular or secreted | 0.114033333 | 0.128397356 | 26.0817 | 26.6611 | 26.4406 | 26.1301 | 27.0398 | 26.3556 |
| MOB-like protein phocein | Q9Y3A3-2;Q9Y3A3-3;Q9Y3A | MOB4 | Other | -0.389266667 | 0.886224031 | 25.8304 | 26.0056 | 26.1989 | 25.2935 | 25.8028 | 25.7708 |
| Molybdenum cofactor sulfurase | Q96EN8 | MOCOS | Other | -1.708433333 | 0.580275121 | 27.2211 | 27.3503 | 27.5244 | 23.5223 | 26.203 | 27.2452 |
| Mannosyl-oligosaccharide glucosidase | Q13724-2;Q13724 | MOGS | Extracellular or secreted | 0.274633333 | 0.069603277 | 21.6075 | 22.9689 | 23.4649 | 20.6601 | 24.8324 | 23.3727 |
| Mortality factor 4-like protein 1 | Q9UBU8-2;Q9UBU8;Q9UBU | MORF4L1 | Other | -1.770966667 | 0.606606038 | 25.4469 | 25.9736 | 25.2864 | 21.6678 | 24.3628 | 25.3634 |
| Putative helicase MOV-10 | Q9HCE1;Q9HCE1-2 | MOV10 | Extracellular or secreted | -0.2262 | 0.152808974 | 26.0128 | 25.68 | 26.8147 | 25.268 | 25.8043 | 26.7566 |
| Mannose-6-phosphate isomerase | P34949;P34949-2 | MPI | Extracellular or secreted | 0.141 | 0.301818426 | 28.2123 | 27.9092 | 27.6388 | 28.0944 | 28.1731 | 27.9158 |
| Myosin phosphatase Rho-interacting protein | Q6WCQ1;Q6WCQ1-2;Q6WC | MPRIP | Other | -1.9069 | 1.846554399 | 25.9117 | 25.7193 | 26.5606 | 24.5777 | 23.4619 | 24.4313 |
| 3-mercaptopyruvate sulfurtransferase | P25325;P25325-2 | MPST | Extracellular or secreted | 1.529966667 | 0.507625974 | 21.4227 | 22.3572 | 25.2092 | 24.2599 | 24.6943 | 24.6248 |
| Myelin protein zero-like protein 1 | O95297-2;O95297;O95297-4 | MPZL1 | Extracellular or secreted | 1.548866667 | 0.938067611 | 24.6508 | 22.6043 | 23.433 | 25.5963 | 25.5827 | 24.1557 |

| | | | | | | | | | | | |
|---|---|---|---|---|---|---|---|---|---|---|---|
| C-type mannose receptor 2 | Q9UBG0 | MRC2 | Other | 0.773566667 | 0.549662198 | 25.5151 | 24.0438 | 23.8212 | 25.1153 | 25.276 | 25.3095 |
| Double-strand break repair protein MRE11A | P49959;P49959-3;P49959-2 | MRE11A | Other | -1.6573 | 0.612307679 | 25.5176 | 24.2804 | 26.638 | 21.8696 | 24.7494 | 24.8451 |
| MRG/MORF4L-binding protein | Q9NV56 | MRGBP | Other | 0.1696 | 0.106951089 | 25.1673 | 24.6366 | 25.9411 | 26.2195 | 25.2787 | 24.7556 |
| Methylthioribose-1-phosphate isomerase | Q9BV20;Q9BV20-2 | MRI1 | Other | -0.044766667 | 0.030766122 | 24.5704 | 23.9774 | 25.088 | 24.0077 | 24.2696 | 25.2242 |
| 39S ribosomal protein L1, mitochondrial | Q9BYD6 | MRPL1 | Other | -0.480466667 | 0.211755152 | 21.7188 | 22.2859 | 24.2054 | 23.0854 | 21.9277 | 21.7556 |
| 39S ribosomal protein L12, mitochondrial | P52815 | MRPL12 | Other | -0.351233333 | 0.238558633 | 27.2485 | 28.2429 | 28.8711 | 27.4951 | 28.4179 | 27.3958 |
| 28S ribosomal protein S27, mitochondrial | Q92552;Q92552-2 | MRPS27 | Other | 1.4374 | 0.443553265 | 21.3539 | 24.2087 | 20.1016 | 23.9756 | 23.4241 | 22.5767 |
| Ribosome-recycling factor, mitochondrial | Q96E11-2;Q96E11-8;Q96E11 | MRRF | Other | -1.417633333 | 0.484319247 | 24.5033 | 24.2723 | 24.2943 | 20.7359 | 24.1843 | 23.8968 |
| mRNA turnover protein 4 homolog | Q9UKD2 | MRTO4 | Other | -0.090966667 | 0.058748276 | 26.3132 | 27.9207 | 27.6336 | 26.9611 | 27.2283 | 27.4052 |
| DNA mismatch repair protein Msh2 | P43246 | MSH2 | Other | -1.446333333 | 0.64764561 | 25.8286 | 25.4258 | 27.7358 | 23.7205 | 24.7458 | 26.1849 |
| DNA mismatch repair protein Msh3 | P20585 | MSH3 | Extracellular or secreted | 0.5182 | 0.1522563 | 21.4564 | 20.1315 | 24.046 | 23.1232 | 21.7645 | 22.3008 |
| DNA mismatch repair protein Msh6 | P52701-4;P52701-3;P52701; | MSH6 | Extracellular or secreted | -0.7162 | 0.147729888 | 20.4621 | 20.5317 | 25.3797 | 20.7632 | 20.8833 | 22.5784 |
| Mesothelin;Megakaryocyte-potentiating factor;Mesothe | Q13421-4;Q13421-3;Q13421 | MSLN | Extracellular or secreted | 0.057133333 | 0.028847198 | 32.466 | 33.5937 | 31.4576 | 32.3262 | 32.8429 | 32.5196 |
| Moesin | P26038 | MSN | Extracellular or secreted | -0.062833333 | 0.111996447 | 34.3781 | 34.3902 | 33.9502 | 34.458 | 34.0785 | 33.9935 |
| Mitochondrial peptide methionine sulfoxide reductase | Q9UJ68-2;Q9UJ68-3;Q9UJ68 | MSRA | Extracellular or secreted | 0.2119 | 1.144535603 | 24.7331 | 24.5294 | 24.6788 | 24.8731 | 24.8502 | 24.8537 |
| Hepatocyte growth factor-like protein;Hepatocyte growt | P26927 | MST1 | Extracellular or secreted | -0.816 | 0.675060911 | 23.619 | 22.4833 | 22.2239 | 21.639 | 21.6136 | 22.6256 |
| Metallothionein-1F | P04733 | MT1F | Other | 3.518133333 | 1.625633143 | 22.3735 | 24.2594 | 21.7189 | 25.9936 | 27.3604 | 25.5522 |
| Metallothionein-1X | P80297 | MT1X | Other | -0.6082 | 0.493065903 | 25.8062 | 26.9874 | 25.7167 | 25.8311 | 25.9783 | 24.8763 |
| Metallothionein-2 | P02795 | MT2A | Other | 0.7777 | 0.252764339 | 30.1546 | 31.1979 | 29.0047 | 32.8392 | 30.4625 | 29.3886 |
| Metastasis-associated protein MTA1 | Q13330-3;Q13330;Q13330-2 | MTA1 | Other | -2.648733333 | 1.293528229 | 23.9189 | 24.3842 | 25.1576 | 20.7387 | 21.5321 | 23.2437 |
| Metastasis-associated protein MTA2 | O94776;O94776-2 | MTA2 | Other | -0.8238 | 0.431916186 | 26.7104 | 26.6016 | 28.2327 | 25.4306 | 26.1079 | 27.5348 |
| Metastasis-associated protein MTA3 | Q9BTC8-2;Q9BTC8 | MTA3 | Other | -0.256966667 | 0.100805563 | 22.542 | 20.197 | 22.5835 | 21.9013 | 20.6604 | 21.9899 |
| Mitochondrial carrier homolog 2 | Q9Y6C9 | MTCH2 | Extracellular or secreted | 2.709566667 | 1.819319658 | 21.1662 | 20.1936 | 21.7619 | 23.8814 | 23.3489 | 24.0201 |
| Protein LYRIC | Q86UE4 | MTDH | Extracellular or secreted | -0.472633333 | 0.309599266 | 30.2892 | 28.6623 | 29.9628 | 29.89 | 28.9025 | 28.7039 |
| C-1-tetrahydrofolate synthase, cytoplasmic;Methylenete | P11586 | MTHFD1 | Extracellular or secreted | -0.1454 | 0.05848379 | 27.7389 | 27.9921 | 29.9271 | 27.601 | 28.2861 | 29.3348 |
| Bifunctional methylenetetrahydrofolate dehydrogenase/ | P13995;P13995-2 | MTHFD2 | Extracellular or secreted | -0.532 | 0.332917588 | 24.999 | 24.8314 | 26.6873 | 25.0606 | 25.0328 | 24.8283 |
| Myotubularin | Q13496-2;Q13496 | MTM1 | Extracellular or secreted | 4.522 | 1.504962318 | 21.7538 | 19.1622 | 22.3005 | 25.177 | 25.3178 | 26.2877 |
| Myotrophin | P58546 | MTPN | Extracellular or secreted | 0.4426 | 0.141932901 | 27.5336 | 28.1546 | 26.9512 | 29.8056 | 27.9892 | 26.1724 |
| Methionine synthase | Q99707;Q99707-2 | MTR | Other | -1.9458 | 1.56494912 | 28.1943 | 27.2974 | 28.4617 | 25.4903 | 25.7278 | 26.8979 |
| Methylmalonyl-CoA mutase, mitochondrial | P22033 | MUT | Other | -2.9864 | 1.113726877 | 25.1001 | 26.1905 | 27.6153 | 21.3902 | 24.6459 | 23.9106 |
| Multivesicular body subunit 12A | Q96EY5-2;Q96EY5-3;Q96EY5 | MVB12A | Extracellular or secreted | 0.554633333 | 0.924598421 | 22.3585 | 22.9158 | 22.156 | 22.8748 | 23.24 | 22.9794 |
| Diphosphomevalonate decarboxylase | P53602 | MVD | Other | 1.5819 | 1.438559098 | 24.4695 | 24.7623 | 25.7545 | 26.1796 | 26.8297 | 26.7227 |
| Major vault protein | Q14764 | MVP | Extracellular or secreted | 1.2312 | 1.165901461 | 24.2695 | 24.7669 | 25.4794 | 26.012 | 26.1965 | 26.0009 |
| Matrix-remodeling-associated protein 7 | P84157-2;P84157-3;P84157 | MXRA7 | Extracellular or secreted | -0.029833333 | 0.006193787 | 21.1954 | 19.2539 | 24.271 | 22.3134 | 21.1816 | 21.1358 |
| Myb-binding protein 1A | Q9BQG0;Q9BQG0-2 | MYBBP1A | Other | -1.5815 | 0.632771538 | 25.2397 | 23.3342 | 24.9841 | 22.4987 | 21.6155 | 24.6993 |
| C-Myc-binding protein | Q99417 | MYCBP | Other | 0.137 | 0.023238504 | 20.2517 | 25.4603 | 25.4134 | 24.3542 | 24.9963 | 22.1859 |
| Myeloid differentiation primary response protein MyD88 | Q99836-5;Q99836;Q99836-6 | MYD88 | Extracellular or secreted | -1.015033333 | 0.325634242 | 23.241 | 22.4837 | 22.613 | 24.0658 | 20.5995 | 20.6273 |
| Myeloid-derived growth factor | Q969H8 | MYDGF | Extracellular or secreted | -1.246166667 | 0.985746032 | 29.7677 | 29.3127 | 29.4314 | 28.8689 | 28.5471 | 27.3573 |
| Myosin-10 | P35580;P35580-5;P35580-2; | MYH10 | Extracellular or secreted | -1.734366667 | 1.217150135 | 28.2992 | 27.2271 | 29.0383 | 25.7269 | 26.8734 | 26.7612 |
| Myosin-14 | Q7Z406;Q7Z406-6;Q7Z406-2 | MYH14 | Extracellular or secreted | 0.203933333 | 0.162377428 | 24.6448 | 24.9088 | 25.5227 | 24.454 | 25.6488 | 25.5853 |
| Myosin-9 | P35579;P35579-2 | MYH9 | Extracellular or secreted | -0.916233333 | 1.0077785 | 33.3832 | 32.3153 | 33.161 | 31.932 | 32.2019 | 31.9769 |
| Myosin regulatory light chain 12A;Myosin regulatory ligh | P19105;O14950;P24844 | MYL12A;MYL1 | Extracellular or secreted | -0.1127 | 0.08829153 | 28.2721 | 29.1537 | 29.3623 | 28.2877 | 29.3496 | 28.8127 |
| Myosin light polypeptide 6 | P60660-2;P60660 | MYL6 | Extracellular or secreted | -0.429866667 | 0.22016759 | 28.5848 | 28.8949 | 30.7352 | 29.5647 | 28.6597 | 28.7009 |
| Myosin light chain kinase, smooth muscle;Myosin light ch | Q15746-11;Q15746-6;Q1574 | MYLK | Extracellular or secreted | 0.389666667 | 0.663111619 | 26.5769 | 26.7834 | 26.7332 | 27.5327 | 26.8832 | 26.8466 |
| Unconventional myosin-XVIIIa | Q92614-3;Q92614-4;Q92614 | MYO18A | Other | -1.0221 | 0.340876246 | 26.5433 | 26.6126 | 27.9898 | 24.0536 | 26.238 | 27.7878 |
| Unconventional myosin-Ib | O43795-2;O43795 | MYO1B | Extracellular or secreted | 0.8157 | 0.697619934 | 26.2285 | 25.3394 | 26.875 | 26.8825 | 27.3503 | 26.6572 |
| Unconventional myosin-Ic | O00159-3 | MYO1C | Extracellular or secreted | 1.436533333 | 1.588444875 | 27.8594 | 26.9371 | 27.8585 | 29.0577 | 29.2147 | 28.6922 |
| Unconventional myosin-VI | Q9UM54-5;Q9UM54-2;Q9UI | MYO6 | Extracellular or secreted | 0.0316 | 0.025254465 | 23.2819 | 22.8073 | 23.7216 | 22.9381 | 23.0192 | 23.9483 |
| Myoferlin | Q9NZM1-6;Q9NZM1-3;Q9NZ | MYOF | Extracellular or secreted | 0.320233333 | 0.304759936 | 26.1788 | 26.2773 | 25.243 | 26.3055 | 26.6393 | 25.715 |
| N-alpha-acetyltransferase 10 | P41227;P41227-2 | NAA10 | Other | -0.883466667 | 1.058364019 | 27.4539 | 28.0884 | 28.2021 | 26.4686 | 27.1078 | 27.5176 |
| N-alpha-acetyltransferase 15, NatA auxiliary subunit | Q9BXJ9;Q9BXJ9-4 | NAA15 | Other | -0.804266667 | 1.365974614 | 28.5793 | 28.4225 | 29.0748 | 27.5931 | 27.8216 | 28.2491 |
| N-alpha-acetyltransferase 20 | P61599;P61599-2 | NAA20 | Other | 0.0561 | 0.035980416 | 21.8242 | 22.5275 | 22.0089 | 22.5476 | 22.7364 | 21.2449 |
| N-alpha-acetyltransferase 25, NatB auxiliary subunit | Q14CX7;Q14CX7-2 | NAA25 | Other | -0.6941 | 2.390096526 | 27.8234 | 27.8123 | 27.6268 | 26.9694 | 27.2346 | 26.9762 |
| N-alpha-acetyltransferase 50 | Q9GZZ1;Q9GZZ1-2 | NAA50 | Extracellular or secreted | -0.660866667 | 0.285344877 | 24.725 | 26.2721 | 26.1949 | 23.7128 | 25.1158 | 26.3808 |
| Nascent polypeptide-associated complex subunit alpha;N | Q13765;E9PAV3-2;E9PAV3 | NACA | Extracellular or secreted | 0.122733333 | 0.636229257 | 29.6074 | 29.735 | 29.7681 | 29.9185 | 29.8707 | 29.6895 |
| Glutamine-dependent NAD(+) synthetase | Q6IA69-2;Q6IA69 | NADSYN1 | Other | 0.4997 | 0.207310017 | 20.9152 | 23.6244 | 22.7129 | 22.044 | 23.5132 | 23.1944 |
| NEDD8-activating enzyme E1 regulatory subunit | Q13564-2;Q13564;Q13564-4 | NAE1 | Extracellular or secreted | -0.255666667 | 0.490873547 | 28.5965 | 29.0453 | 28.9762 | 28.2792 | 28.6951 | 28.8767 |
| H/ACA ribonucleoprotein complex non-core subunit NAF | Q96HR8-2;Q96HR8 | NAF1 | Other | 0.2343 | 0.230582488 | 22.9091 | 22.8852 | 22.3075 | 22.3848 | 22.8719 | 23.548 |
| Alpha-N-acetylgalactosaminidase | P17050 | NAGA | Extracellular or secreted | -0.302633333 | 0.243047101 | 26.8975 | 26.5567 | 26.8957 | 27.1143 | 26.7073 | 25.6204 |
| N-acetyl-D-glucosamine kinase | Q9UJ70;Q9UJ70-2 | NAGK | Extracellular or secreted | 0.386266667 | 0.731864349 | 25.4854 | 26.1365 | 26.0125 | 26.1919 | 26.2183 | 26.383 |
| Alpha-N-acetylglucosaminidase;Alpha-N-acetylglucosami | P54802 | NAGLU | Extracellular or secreted | 1.278933333 | 1.120139634 | 26.8775 | 27.3337 | 25.9814 | 28.117 | 28.1127 | 27.7997 |
| Nicotinamide phosphoribosyltransferase | P43490 | NAMPT | Extracellular or secreted | 0.4852 | 0.833347642 | 33.4809 | 33.3746 | 32.7724 | 33.8663 | 33.7656 | 33.4516 |
| N-acylneuraminate-9-phosphatase | Q8TBE9 | NANP | Other | -0.808 | 0.322808954 | 21.5188 | 21.2984 | 23.9343 | 22.2984 | 21.6319 | 20.3972 |
| Sialic acid synthase | Q9NR45 | NANS | Extracellular or secreted | -0.4587 | 0.893448355 | 28.7887 | 29.4533 | 29.1255 | 28.5469 | 28.7761 | 28.6684 |
| Nucleosome assembly protein 1-like 1 | P55209-2;P55209;P55209-3 | NAP1L1 | Other | -0.688166667 | 0.999193648 | 29.2171 | 29.0939 | 29.6998 | 28.2686 | 28.5466 | 29.1311 |
| Nucleosome assembly protein 1-like 4 | Q99733;Q99733-2 | NAP1L4 | Other | -0.393933333 | 0.383059153 | 28.6566 | 28.9854 | 29.4228 | 28.0088 | 28.619 | 29.2552 |
| Alpha-soluble NSF attachment protein | P54920 | NAPA | Extracellular or secreted | -0.376266667 | 0.225764241 | 27.5619 | 26.5307 | 27.8835 | 25.939 | 27.4606 | 27.4477 |
| Nicotinate phosphoribosyltransferase | Q6XQN6;Q6XQN6-2;Q6XQN | NAPRT | Extracellular or secreted | -0.7532 | 1.629522171 | 28.9234 | 28.8191 | 28.395 | 28.0245 | 28.136 | 27.7174 |
| Asparagine--tRNA ligase, cytoplasmic | O43776 | NARS | Extracellular or secreted | -0.384066667 | 0.435408771 | 29.9229 | 30.4186 | 30.1096 | 29.1147 | 30.1235 | 30.0607 |
| Nuclear autoantigenic sperm protein | P49321;P49321-3;P49321-2; | NASP | Other | -0.350433333 | 0.470882348 | 31.2272 | 30.4426 | 30.5877 | 30.7911 | 30.3585 | 30.0566 |
| Neuroblastoma suppressor of tumorigenicity 1 | P41271;P41271-2 | NBL1 | Extracellular or secreted | 3.8595 | 0.960686742 | 24.5632 | 24.6554 | 23.0752 | 30.5787 | 25.4356 | 27.858 |
| Condensin complex subunit 1 | Q15021 | NCAPD2 | Other | -1.1045 | 0.370892484 | 25.3531 | 24.8096 | 27.4766 | 23.6126 | 24.0754 | 26.6378 |
| Condensin complex subunit 3 | Q9BPX3 | NCAPG | Other | 0.258666667 | 0.373755457 | 25.9556 | 25.5128 | 26.0016 | 26.3947 | 26.2376 | 25.6137 |
| Condensin complex subunit 2 | Q15003;Q15003-2 | NCAPH | Other | -1.3719 | 0.586532261 | 23.0878 | 23.7513 | 25.6673 | 21.7222 | 22.5607 | 24.1078 |
| Nuclear cap-binding protein subunit 1 | Q09161 | NCBP1 | Other | -0.682966667 | 0.364044308 | 26.1761 | 26.8368 | 27.7618 | 25.3485 | 25.932 | 27.4453 |
| F-box only protein 50 | Q6ZVX7 | NCCRP1 | Extracellular or secreted | 2.238 | 0.853063015 | 20.949 | 22.161 | 22.1106 | 25.9157 | 22.6358 | 23.3831 |
| Neurochondrin | Q9UBB6-2;Q9UBB6;Q9UBB6 | NCDN | Other | 0.266566667 | 0.154865368 | 22.3189 | 22.4484 | 22.889 | 21.7552 | 22.919 | 23.7818 |
| Cytoplasmic protein NCK1 | P16333;P16333-2 | NCK1 | Extracellular or secreted | 1.2644 | 0.412662814 | 20.4687 | 23.9242 | 23.9213 | 23.9753 | 24.0985 | 24.0336 |
| Nck-associated protein 1 | Q9Y2A7;Q9Y2A7-2 | NCKAP1 | Extracellular or secreted | -0.1469 | 0.109016086 | 27.0533 | 27.2913 | 28.1383 | 26.8693 | 27.1233 | 28.0496 |
| Nucleolin | P19338 | NCL | Extracellular or secreted | -0.236466667 | 0.692126377 | 34.0595 | 33.8573 | 33.7838 | 33.8624 | 33.6994 | 33.4294 |
| Nuclear receptor coactivator 7 | Q8NI08-5;Q8NI08-7;Q8NI08- | NCOA7 | Other | -0.1059 | 0.042646446 | 23.1101 | 21.1821 | 22.148 | 20.8059 | 22.9258 | 22.3908 |
| Nicastrin | Q92542-2;Q92542 | NCSTN | Extracellular or secreted | 3.703333333 | 3.11474856 | 21.839 | 22.605 | 22.3606 | 26.197 | 25.8623 | 25.8553 |
| Protein NDRG1 | Q92597;Q92597-3;Q92597-2 | NDRG1 | Extracellular or secreted | 0.8076 | 0.526521316 | 26.2659 | 26.6003 | 28.061 | 27.2455 | 27.6333 | 28.4712 |
| Protein NDRG3 | Q9UGV2-2;Q9UGV2;Q9UGV | NDRG3 | Extracellular or secreted | 0.574766667 | 0.629692267 | 24.4682 | 24.6068 | 25.5666 | 25.1488 | 25.4876 | 25.7295 |
| NADH dehydrogenase [ubiquinone] 1 alpha subcomplex | P51970 | NDUFA8 | Other | -1.0973 | 0.924089951 | 26.4439 | 26.3672 | 26.1278 | 24.8134 | 26.0746 | 24.759 |
| Acyl carrier protein, mitochondrial | O14561 | NDUFAB1 | Other | -0.025566667 | 0.005804654 | 25.1063 | 23.9499 | 27.9627 | 27.2241 | 24.9697 | 24.7484 |
| NADH-ubiquinone oxidoreductase 75 kDa subunit, mitoc | P28331;P28331-2;P28331-4; | NDUFS1 | Other | -0.5925 | 0.12293179 | 22.4698 | 22.5549 | 27.4372 | 23.4095 | 23.3388 | 23.9361 |
| NADH dehydrogenase [ubiquinone] flavoprotein 2, mitoc | P19404 | NDUFV2 | Other | -1.2031 | 1.133596493 | 24.0617 | 24.3938 | 25.3058 | 22.8471 | 23.9824 | 23.3225 |
| Nebulin | P20929-4;P20929;P20929-3; | NEB | Extracellular or secreted | -2.177733333 | 1.443903661 | 25.5445 | 25.2653 | 25.183 | 22.2886 | 23.3091 | 23.8619 |
| Adaptin ear-binding coat-associated protein 1 | Q8NC96-2;Q8NC96 | NECAP1 | Extracellular or secreted | 0.145333333 | 0.156348377 | 22.8847 | 23.2171 | 22.9015 | 22.5819 | 23.6699 | 23.1875 |
| Adaptin ear-binding coat-associated protein 2 | Q9NVZ3;Q9NVZ3-2;Q9NVZ3- | NECAP2 | Extracellular or secreted | -0.154833333 | 0.028928148 | 24.8589 | 21.486 | 25.2337 | 21.0793 | 24.5025 | 25.5323 |
| NEDD8 | Q15843 | NEDD8 | Extracellular or secreted | 0.292166667 | 0.49149193 | 26.7827 | 27.2293 | 27.0718 | 27.212 | 27.7344 | 27.0139 |
| Neuronal growth regulator 1 | Q7Z3B1;Q7Z3B1-2 | NEGR1 | Extracellular or secreted | 1.804733333 | 2.238980024 | 24.5517 | 25.4038 | 24.9186 | 26.979 | 26.9931 | 26.3162 |
| Serine/threonine-protein kinase Nek9 | Q8TD19 | NEK9 | Other | -0.4332 | 0.136919269 | 22.409 | 23.4121 | 23.4061 | 24.2729 | 20.6429 | 23.0118 |
| Negative elongation factor B | Q8WX92 | NELFB | Other | -1.293433333 | 0.299103425 | 22.2584 | 20.9056 | 25.5972 | 20.4492 | 20.7191 | 23.7126 |
| Negative elongation factor C/D | Q8IXH7-4;Q8IXH7 | NELFCD | Other | -0.776733333 | 0.375926043 | 23.158 | 21.9827 | 22.8721 | 21.3431 | 20.9637 | 23.3758 |
| Nuclear export mediator factor NEMF | O60524-4;O60524-5;O60524 | NEMF | Other | -1.1746 | 0.363745487 | 22.9965 | 23.4664 | 24.2652 | 23.3222 | 20.0322 | 23.8499 |
| Neudesin | Q9UMX5 | NENF | Extracellular or secreted | 0.363766667 | 0.086687677 | 24.327 | 24.6633 | 27.4309 | 27.9553 | 25.3914 | 24.1658 |
| Neogenin | Q92859-3;Q92859-2;Q92859 | NEO1 | Extracellular or secreted | 3.245766667 | 1.276106783 | 23.5594 | 21.1233 | 20.9887 | 25.5432 | 24.8562 | 25.0093 |
| Sialidase-1 | Q99519 | NEU1 | Extracellular or secreted | 1.816066667 | 1.203779241 | 23.7581 | 24.5361 | 24.6335 | 25.0458 | 26.8852 | 26.4449 |
| Nuclear factor 1 B-type | O00712;O00712-4;O00712-5 | NFIB | Other | -2.033933333 | 0.737271016 | 24.1387 | 23.9249 | 25.847 | 21.8631 | 21.2922 | 24.6535 |
| Nuclear factor NF-kappa-B p105 subunit;Nuclear factor N | P19838;P19838-2;P19838-3 | NFKB1 | Other | -0.553266667 | 0.079713396 | 25.4049 | 21.0622 | 27.2084 | 20.7833 | 25.1792 | 26.0532 |
| Nuclear factor NF-kappa-B p100 subunit;Nuclear factor N | Q00653-4;Q00653;Q00653-3 | NFKB2 | Extracellular or secreted | 1.115066667 | 0.273396797 | 22.5232 | 20.1088 | 24.6214 | 22.2393 | 22.9345 | 25.4248 |
| NFU1 iron-sulfur cluster scaffold homolog, mitochondria | Q9UMS0-3;Q9UMS0;Q9UMS | NFU1 | Other | 0.052966667 | 0.03501389 | 25.7554 | 25.7629 | 26.1916 | 26.2045 | 26.6162 | 25.0481 |
| Nuclear transcription factor Y subunit beta | P25208 | NFYB | Other | -1.276333333 | 0.636516488 | 22.4678 | 23.7864 | 24.2887 | 21.5979 | 21.4613 | 23.6547 |
| Nuclear transcription factor Y subunit gamma | Q13952-6;Q13952-7;Q13952 | NFYC | Other | -0.0874 | 0.117063394 | 24.1219 | 24.6191 | 24.8474 | 24.5843 | 24.629 | 24.1129 |
| Neuroguidin | Q8NEJ9-2;Q8NEJ9 | NGDN | Other | -1.631366667 | 0.598695552 | 21.1631 | 20.7704 | 24.0346 | 20.6689 | 20.0998 | 20.3053 |
| NHL repeat-containing protein 2 | Q8NBF2;Q8NBF2-2 | NHLRC2 | Other | -1.118166667 | 0.635731227 | 25.2476 | 26.213 | 26.1351 | 23.4241 | 25.5608 | 25.2563 |
| NHL repeat-containing protein 3 | Q5JS37;Q5JS37-2 | NHLRC3 | Extracellular or secreted | 1.505 | 1.06051373 | 25.8989 | 25.7588 | 25.9579 | 28.0141 | 27.685 | 26.4315 |
| H/ACA ribonucleoprotein complex subunit 2 | Q9NX24 | NHP2 | Other | 0.266666667 | 0.227881411 | 23.5411 | 23.965 | 24.925 | 24.2623 | 24.2533 | 24.7155 |
| NHP2-like protein 1;NHP2-like protein 1, N-terminally pr | P55769 | NHP2L1 | Other | -0.758033333 | 0.518369101 | 25.9253 | 26.2376 | 27.7217 | 25.6449 | 26.1211 | 25.8445 |
| Nidogen-1 | P14543-2;P14543 | NID1 | Extracellular or secreted | 0.7284 | 0.442936991 | 23.0488 | 22.7081 | 21.7148 | 22.0936 | 23.9336 | 23.6297 |
| Nidogen-2 | Q14112 | NID2 | Extracellular or secreted | 1.7113 | 1.251323665 | 31.4271 | 31.502 | 30.2577 | 31.8476 | 32.9662 | 33.5069 |
| NIF3-like protein 1 | Q9GZT8-2;Q9GZT8;Q9GZT8- | NIF3L1 | Other | -0.0773 | 0.046756112 | 29.6498 | 28.9946 | 28.519 | 29.7722 | 28.9694 | 28.1899 |
| MKI67 FHA domain-interacting nucleolar phosphoprotei | Q9BYG3 | NIFK | Other | -0.445833333 | 0.335918103 | 24.794 | 23.6178 | 25.0966 | 24.646 | 23.7458 | 23.7791 |
| Ninjurin-1 | Q92982 | NINJ1 | Other | -0.1878 | 0.059300095 | 20.9846 | 21.5871 | 23.1514 | 22.641 | 22.5563 | 19.9624 |
| 60S ribosome subunit biogenesis protein NIP7 homolog | Q9Y221-2;Q9Y221 | NIP7 | Other | -1.8677 | 0.942451244 | 22.8346 | 23.732 | 24.7654 | 22.3753 | 20.4954 | 22.8582 |
| Protein NipSnap homolog 3A | Q9UFN0 | NIPSNAP3A | Other | -0.7414 | 0.939035571 | 27.2688 | 27.5806 | 27.927 | 26.9567 | 27.303 | 26.2925 |
| Nitrilase homolog 1 | Q86X76-2;Q86X76-4;Q86X76 | NIT1 | Extracellular or secreted | -0.387966667 | 0.133890874 | 22.8277 | 23.0874 | 23.1461 | 20.7009 | 24.011 | 23.1854 |

| | | | | | | | | | | | |
|---|---|---|---|---|---|---|---|---|---|---|---|
| Omega-amidase NIT2 | Q9NQR4 | NIT2 | Extracellular or secreted | -0.326766667 | 1.308062373 | 29.6739 | 29.7336 | 29.5529 | 29.3786 | 29.4528 | 29.1487 |
| Notchless protein homolog 1 | Q9NVX2;Q9NVX2-2 | NLE1 | Other | -0.7107 | 0.717250833 | 22.6045 | 23.5752 | 23.34 | 21.8334 | 22.54 | 23.0142 |
| Neuroligin-4, X-linked;Neuroligin-4, Y-linked;Neuroligin-1 | Q8N0W4;Q8N0W4-2;Q8NFZ | NLGN4X;NLGN | Extracellular or secreted | 0.072533333 | 0.032796291 | 23.5071 | 23.1369 | 22.3039 | 21.9184 | 24.1397 | 23.1074 |
| Neurolysin, mitochondrial | Q9BYT8 | NLN | Extracellular or secreted | -0.536133333 | 1.303559796 | 28.4318 | 28.0707 | 28.2991 | 28.015 | 27.6763 | 27.5019 |
| Nucleoside diphosphate kinase A | P15531;P15531-2 | NME1 | Extracellular or secreted | -1.054 | 1.736775855 | 30.54 | 29.9928 | 30.0559 | 29.4186 | 29.2694 | 28.7387 |
| Nucleoside diphosphate kinase B | P22392-2;P22392 | NME2 | Other | -0.936266667 | 1.307606929 | 33.8271 | 33.913 | 33.1212 | 32.7166 | 32.8439 | 32.492 |
| Putative nucleoside diphosphate kinase | O60361 | NME2P1 | Other | -0.1076 | 0.087442379 | 24.7511 | 24.8724 | 23.9547 | 24.8256 | 24.6608 | 23.769 |
| Nucleoside diphosphate kinase 7 | Q9Y5B8-2;Q9Y5B8 | NME7 | Other | 0.303566667 | 0.302765353 | 21.0786 | 21.7286 | 22.2294 | 21.7113 | 21.8052 | 22.4308 |
| NmrA-like family domain-containing protein 1 | Q9HBL8 | NMRAL1 | Other | 0.428766667 | 0.426347375 | 22.5818 | 22.5549 | 22.7617 | 23.605 | 23.2459 | 22.3338 |
| Glycylpeptide N-tetradecanoyltransferase 1 | P30419;P30419-2 | NMT1 | Extracellular or secreted | -0.506533333 | 1.111501188 | 28.8282 | 29.2806 | 29.098 | 28.2328 | 28.7363 | 28.7181 |
| Nicotinamide N-methyltransferase | P40261 | NNMT | Other | -2.384133333 | 0.748804937 | 28.0641 | 28.5076 | 29.1838 | 23.8592 | 26.9299 | 27.814 |
| Nucleolar protein 11 | Q9H8H0;Q9H8H0-2 | NOL11 | Other | -0.960033333 | 0.374477463 | 22.2411 | 23.1029 | 25.4351 | 22.3002 | 22.4805 | 23.1183 |
| Nucleolar protein 3 | O60936;O60936-3;O60936-1 | NOL3 | Other | 0.327866667 | 0.327656962 | 26.4333 | 25.9732 | 27.23 | 27.0821 | 26.8226 | 26.7154 |
| Nucleolar protein 7 | Q9UMY1-2;Q9UMY1 | NOL7 | Other | 1.4903 | 1.232803613 | 21.7008 | 22.0953 | 22.2507 | 23.0723 | 24.3597 | 23.0857 |
| Nucleolar and coiled-body phosphoprotein 1 | Q14978-2;Q14978;Q14978-3 | NOLC1 | Other | -1.306066667 | 0.841202533 | 30.4017 | 29.241 | 28.7152 | 28.9112 | 28.3753 | 27.1532 |
| Nodal modulator 2;Nodal modulator 1;Nodal modulator | Q5JPE7-2;Q15155;P69849;Q | NOMO2;NOMO | Other | -0.0865 | 0.077500328 | 25.8263 | 25.9019 | 25.0703 | 24.9343 | 25.7867 | 25.818 |
| Non-POU domain-containing octamer-binding protein | Q15233;Q15233-2 | NONO | Other | 1.208766667 | 0.504989745 | 26.0868 | 26.2956 | 28.2029 | 26.6901 | 28.0719 | 29.4496 |
| H/ACA ribonucleoprotein complex subunit 3 | Q9NPE3 | NOP10 | Other | -1.3182 | 0.461133913 | 22.2991 | 22.653 | 24.6389 | 20.1101 | 22.03 | 23.4963 |
| Probable 28S rRNA (cytosine(4447)-C(5))-methyltransfera | P46087-2;P46087;P46087-4; | NOP2 | Other | 0.426666667 | 0.160788439 | 24.6588 | 24.2843 | 27.0674 | 26.4325 | 25.8771 | 24.9809 |
| Nucleolar protein 56 | O00567 | NOP56 | Other | -1.7027 | 0.808287145 | 27.9664 | 26.3778 | 29.135 | 27.0036 | 25.6558 | 25.7117 |
| Nucleolar protein 58 | Q9Y2X3 | NOP58 | Other | -1.979866667 | 0.991202516 | 26.9936 | 26.0469 | 28.5457 | 26.2019 | 24.2598 | 25.1849 |
| Neurogenic locus notch homolog protein 2;Notch 2 extra | Q04721 | NOTCH2 | Extracellular or secreted | 0.2343 | 0.335533006 | 23.9998 | 24.0901 | 23.3393 | 23.7321 | 24.219 | 24.181 |
| Neurogenic locus notch homolog protein 3;Notch 3 extra | Q9UM47 | NOTCH3 | Extracellular or secreted | 0.4009 | 0.292791055 | 26.438 | 27.4147 | 26.4292 | 26.3024 | 27.4105 | 27.7717 |
| Protein NOV homolog | P48745 | NOV | Extracellular or secreted | 1.873666667 | 0.978507591 | 27.7482 | 27.6555 | 25.7217 | 28.917 | 28.9131 | 28.9163 |
| Niemann-Pick C1 protein | O15118 | NPC1 | Extracellular or secreted | 3.744666667 | 2.393918791 | 22.3904 | 21.1561 | 21.0298 | 25.9366 | 25.4758 | 24.3979 |
| Epididymal secretory protein E1 | P61916;P61916-2 | NPC2 | Extracellular or secreted | 1.319533333 | 0.846695638 | 31.9674 | 32.5455 | 30.5538 | 32.9527 | 33.477 | 32.5956 |
| Probable aminopeptidase NPEPL1 | Q8NDH3;Q8NDH3-5;Q8NDH | NPEPL1 | Other | 0.028433333 | 0.014790999 | 27.3682 | 27.1418 | 26.2741 | 27.8118 | 27.0816 | 25.976 |
| Puromycin-sensitive aminopeptidase | P55786;P55786-2 | NPEPPS | Extracellular or secreted | -0.098033333 | 0.076638307 | 32.5296 | 32.4515 | 31.5838 | 32.6769 | 32.0677 | 31.5262 |
| Nuclear protein localization protein 4 homolog | Q8TAT6;Q8TAT6-2 | NPLOC4 | Other | -0.478833333 | 0.348758702 | 27.8542 | 27.9757 | 28.5552 | 26.7226 | 27.8013 | 28.4247 |
| Nucleophosmin | P06748-2;P06748;P06748-3 | NPM1 | Other | -0.575166667 | 1.022181445 | 34.3459 | 33.7097 | 33.7777 | 33.2643 | 33.4528 | 33.3907 |
| Nucleoplasmin-3 | O75607 | NPM3 | Other | -0.882166667 | 1.308610769 | 28.5255 | 28.1983 | 27.8214 | 26.8243 | 27.5359 | 27.5385 |
| Nephronectin | Q6UXI9;Q6UXI9-6;Q6UXI9-3 | NPNT | Extracellular or secreted | 0.707033333 | 0.421424529 | 21.1423 | 22.9082 | 20.9345 | 22.0548 | 22.2658 | 22.7855 |
| Neuroplastin | Q9Y639-3;Q9Y639-1;Q9Y639 | NPTN | Extracellular or secreted | 1.217966667 | 1.337298899 | 26.6276 | 26.0751 | 25.9046 | 27.807 | 27.6979 | 26.7563 |
| NAD(P)H dehydrogenase [quinone] 1 | P15559-3;P15559-2;P15559 | NQO1 | Extracellular or secreted | 0.5175 | 0.328473167 | 27.1697 | 27.681 | 29.0552 | 28.2521 | 28.1942 | 29.0121 |
| Ribosyldihydronicotinamide dehydrogenase [quinone] | P16083 | NQO2 | Extracellular or secreted | -0.561833333 | 0.354217539 | 30.9889 | 30.3821 | 29.798 | 30.7623 | 29.8548 | 28.8664 |
| Nuclear receptor 2C2-associated protein | Q86WQ0;Q86WQ0-2 | NR2C2AP | Extracellular or secreted | -0.960666667 | 0.586371452 | 22.7894 | 23.8108 | 22.6341 | 22.3557 | 23.0218 | 20.9748 |
| GTPase NRas | P01111 | NRAS | Extracellular or secreted | 2.101466667 | 1.2989981 | 21.4498 | 22.3771 | 20.441 | 23.2956 | 23.9402 | 23.3365 |
| Nuclear receptor-binding protein | Q9UHY1 | NRBP1 | Other | 0.173066667 | 0.122310133 | 24.5505 | 25.4014 | 25.1437 | 24.3429 | 25.4746 | 25.7973 |
| Nardilysin | O43847;O43847-2 | NRD1 | Other | 0.058766667 | 0.116273869 | 29.5643 | 29.6025 | 29.0738 | 29.511 | 29.5021 | 29.4038 |
| Nuclear respiratory factor 1 | Q16656;Q16656-4;Q16656-2 | NRF1 | Extracellular or secreted | -1.010533333 | 0.761510822 | 23.4981 | 22.6338 | 24.1592 | 21.6053 | 22.6426 | 23.0116 |
| Neuropilin-1 | O14786;O14786-3;O14786-2 | NRP1 | Extracellular or secreted | 2.637566667 | 0.561788749 | 28.6579 | 28.979 | 23.4836 | 30.4742 | 29.0477 | 29.5113 |
| Neuropilin-2 | O60462-4;O60462-5;O60462 | NRP2 | Extracellular or secreted | -1.189066667 | 0.31720617 | 26.1123 | 25.3047 | 24.8956 | 21.497 | 25.6332 | 25.6152 |
| Sterol-4-alpha-carboxylate 3-dehydrogenase, decarboxyl | Q15738 | NSDHL | Other | 2.4075 | 1.948106213 | 20.624 | 20.5122 | 21.3161 | 22.4239 | 23.5492 | 23.7017 |
| Vesicle-fusing ATPase | P46459;P46459-2 | NSF | Extracellular or secreted | -0.606833333 | 0.149497961 | 26.7688 | 24.9764 | 27.5461 | 23.7781 | 25.5164 | 28.1763 |
| NSFL1 cofactor p47 | Q9UNZ2;Q9UNZ2-5;Q9UNZ2 | NSFL1C | Extracellular or secreted | 0.676 | 1.339240023 | 29.7278 | 29.403 | 29.9897 | 30.2785 | 30.3926 | 30.4774 |
| Kinetochore-associated protein NSL1 homolog | Q96IY1-2;Q96IY1 | NSL1 | Other | 0.0825 | 0.035249727 | 21.8227 | 21.3047 | 23.1845 | 23.1351 | 22.2256 | 21.1987 |
| tRNA (cytosine(34)-C(5))-methyltransferase | Q08J23;Q08J23-3;Q08J23-2 | NSUN2 | Other | -0.6925 | 0.202289472 | 21.2031 | 22.9666 | 24.6024 | 20.9059 | 21.8886 | 23.9001 |
| 5(3)-deoxyribonucleotidase, cytosolic type | Q8TCD5 | NT5C | Extracellular or secreted | -0.974033333 | 0.399466947 | 24.1899 | 26.5045 | 25.8484 | 23.0188 | 25.4622 | 25.1397 |
| Cytosolic purine 5-nucleotidase | P49902;P49902-2 | NT5C2 | Other | -0.2252 | 0.255734347 | 25.1682 | 25.9004 | 25.5627 | 24.7692 | 25.5434 | 25.6431 |
| 5-nucleotidase domain-containing protein 1 | Q5TFE4;Q5TFE4-2 | NT5DC1 | Other | -0.166666667 | 0.751317379 | 29.0874 | 28.8501 | 28.8214 | 28.7919 | 28.7753 | 28.6917 |
| 5-nucleotidase | P21589-2;P21589 | NT5E | Extracellular or secreted | 5.354733333 | 2.092618307 | 23.2197 | 22.2441 | 21.6476 | 26.1652 | 28.3162 | 28.6942 |
| Nucleobindin-1 | Q02818 | NUCB1 | Extracellular or secreted | 2.1086 | 1.490155161 | 27.2194 | 27.8335 | 28.3508 | 30.9208 | 29.4868 | 29.3219 |
| Nucleobindin-2;Nesfatin-1 | P80303-2;P80303 | NUCB2 | Extracellular or secreted | 1.1458 | 1.315921276 | 22.641 | 22.8839 | 22.5549 | 23.8378 | 24.3492 | 23.3302 |
| Nuclear ubiquitous casein and cyclin-dependent kinase s | Q9H1E3;Q9H1E3-2 | NUCKS1 | Other | 0.0924 | 0.052389888 | 31.2482 | 30.1067 | 30.1971 | 31.5114 | 30.4469 | 29.8709 |
| Nuclear migration protein nudC | Q9Y266 | NUDC | Other | -0.9499 | 1.723351706 | 31.4783 | 30.9793 | 31.0105 | 29.8581 | 30.4947 | 30.2656 |
| NudC domain-containing protein 1 | Q96RS6;Q96RS6-2;Q96RS6-3 | NUDCD1 | Other | -0.9284 | 1.598900045 | 28.8794 | 28.6189 | 29.1166 | 27.5363 | 28.1002 | 28.1932 |
| NudC domain-containing protein 2 | Q8WVJ2 | NUDCD2 | Extracellular or secreted | -0.579066667 | 0.329328917 | 24.4762 | 23.9961 | 23.4629 | 22.3479 | 24.5228 | 23.3273 |
| 7,8-dihydro-8-oxoguanine triphosphatase | P36639-4;P36639-3;P36639- | NUDT1 | Extracellular or secreted | -0.403366667 | 0.203455931 | 28.1981 | 26.9802 | 26.6207 | 27.6887 | 27.1908 | 25.7094 |
| Probable 8-oxo-dGTP diphosphatase NUDT15 | Q9NV35 | NUDT15 | Other | 0.1028 | 0.107345913 | 23.1726 | 22.5614 | 22.5922 | 22.8481 | 23.3772 | 22.4093 |
| Nucleoside diphosphate-linked moiety X motif 19, mitocl | A8MXV4 | NUDT19 | Extracellular or secreted | -1.8097 | 1.103257588 | 22.9549 | 23.1888 | 23.5519 | 20.5618 | 22.5273 | 21.1774 |
| Cleavage and polyadenylation specificity factor subunit 5 | O43809 | NUDT21 | Other | -1.603166667 | 1.261626686 | 27.7403 | 27.9762 | 27.9406 | 25.8333 | 25.9189 | 27.0954 |
| Diphosphoinositol polyphosphate phosphohydrolase 1 | O95989 | NUDT3 | Extracellular or secreted | -0.280166667 | 0.870889209 | 26.2801 | 26.3421 | 25.9494 | 25.9738 | 25.772 | 25.9853 |
| ADP-sugar pyrophosphatase | Q9UKK9 | NUDT5 | Extracellular or secreted | -0.355333333 | 0.45144672 | 31.8368 | 31.1615 | 30.9001 | 31.2551 | 30.9411 | 30.6362 |
| ADP-ribose pyrophosphatase, mitochondrial | Q9BW91-2;Q9BW91 | NUDT9 | Extracellular or secreted | 0.524733333 | 0.101518264 | 24.0766 | 20.4769 | 22.4293 | 24.251 | 24.4891 | 19.8169 |
| Nuclear mitotic apparatus protein 1 | Q14980;Q14980-2;Q14980-4 | NUMA1 | Extracellular or secreted | -1.4718 | 1.807140134 | 31.6128 | 30.9745 | 31.9054 | 29.7194 | 30.0396 | 30.3183 |
| Nuclear pore complex protein Nup153 | P49790-2;P49790;P49790-3 | NUP153 | Other | -1.756566667 | 0.788765026 | 24.1366 | 23.8904 | 26.482 | 23.0924 | 22.7804 | 23.3665 |
| Nuclear pore complex protein Nup160 | Q12769 | NUP160 | Other | 0.295433333 | 0.851069113 | 23.5314 | 23.4493 | 23.62 | 23.5845 | 24.0328 | 23.8697 |
| Nuclear pore complex protein Nup214 | P35658-2;P35658-4;P35658; | NUP214 | Extracellular or secreted | -1.116 | 0.494194015 | 21.7549 | 21.4693 | 24.0316 | 22.2417 | 21.1985 | 20.4676 |
| Nucleoporin Nup37 | Q8NFH4 | NUP37 | Other | -0.646866667 | 0.607610603 | 26.7879 | 25.7213 | 27.0308 | 25.9446 | 25.7229 | 25.9319 |
| Nucleoporin Nup43 | Q8NFH3 | NUP43 | Other | -2.231166667 | 0.781362113 | 25.8214 | 23.1264 | 26.4909 | 21.6386 | 22.7393 | 24.3673 |
| Nuclear pore complex protein Nup50 | Q9UKX7-2;Q9UKX7 | NUP50 | Other | -0.830633333 | 1.133315793 | 23.7842 | 24.6398 | 24.1672 | 22.9543 | 23.7849 | 23.3601 |
| Nucleoporin p54 | Q7Z3B4-2;Q7Z3B4-3;Q7Z3B4 | NUP54 | Other | -0.6654 | 0.57360984 | 23.1593 | 22.7088 | 24.0439 | 22.2346 | 22.3834 | 23.2978 |
| Nuclear pore glycoprotein p62 | P37198 | NUP62 | Other | -1.521266667 | 0.925546326 | 21.7618 | 23.3931 | 23.7871 | 21.7632 | 21.6907 | 20.9243 |
| Nuclear pore complex protein Nup88 | Q99567 | NUP88 | Other | -0.1457 | 0.051921082 | 22.0438 | 21.4613 | 23.9099 | 23.2255 | 22.6117 | 21.1407 |
| Nuclear pore complex protein Nup93 | Q8N1F7-2;Q8N1F7 | NUP93 | Other | -1.190166667 | 0.995130715 | 24.4715 | 24.8009 | 25.8785 | 23.2011 | 23.9411 | 24.4382 |
| Nuclear pore complex protein Nup98-Nup96;Nuclear por | P52948-4;P52948-3;P52948- | NUP98 | Other | -2.433366667 | 1.287619033 | 24.3676 | 23.848 | 25.2394 | 21.6817 | 21.0768 | 23.3964 |
| Nucleolar and spindle-associated protein 1 | Q9BXS6-2;Q9BXS6;Q9BXS6-7 | NUSAP1 | Other | -1.460733333 | 0.946304697 | 25.6489 | 26.85 | 27.6594 | 24.7736 | 25.2387 | 25.7638 |
| Nuclear transport factor 2 | P61970 | NUTF2 | Extracellular or secreted | -0.2181 | 0.141799212 | 32.3593 | 32.5476 | 31.2261 | 31.6261 | 32.5845 | 31.2681 |
| Nuclear RNA export factor 1 | Q9UBU9;Q9UBU9-2 | NXF1 | Other | -0.876833333 | 0.413271511 | 21.8112 | 23.1973 | 24.4108 | 21.355 | 22.6092 | 22.8246 |
| Nucleoredoxin | Q6DKJ4;Q6DKJ4-3 | NXN | Other | -0.599466667 | 0.771987531 | 25.7306 | 25.9675 | 25.7909 | 24.8028 | 25.1033 | 25.7845 |
| NTF2-related export protein 1;NTF2-related export prote | Q9UKK6;Q9NPJ8;Q9NPJ8-2;( | NXT1;NXT2 | Other | -3.000266667 | 1.135234485 | 25.5816 | 24.1879 | 25.1514 | 20.3772 | 23.7153 | 21.8276 |
| Out at first protein homolog | Q86UD1 | OAF | Extracellular or secreted | 1.5199 | 1.006109071 | 25.4519 | 25.8526 | 23.9789 | 26.3621 | 27.2797 | 26.2013 |
| Ornithine aminotransferase, mitochondrial;Ornithine am | P04181;P04181-2 | OAT | Other | -0.6064 | 0.374259848 | 27.8131 | 27.8555 | 29.4103 | 28.0838 | 28.2664 | 26.9095 |
| Occludin | Q16625-5;Q16625-4;Q16625 | OCLN | Extracellular or secreted | 2.6963 | 1.420661155 | 24.1651 | 21.9495 | 22.9286 | 26.6835 | 25.8511 | 24.5975 |
| 2-oxoglutarate dehydrogenase, mitochondrial | Q02218;Q02218-2 | OGDH | Other | -0.280633333 | 0.135431697 | 25.8795 | 26.2787 | 28.1666 | 26.4311 | 26.7891 | 26.2627 |
| Prolyl 3-hydroxylase OGFOD1 | Q8N543;Q8N543-2 | OGFOD1 | Other | -0.691033333 | 0.415177442 | 26.6836 | 26.179 | 24.6061 | 25.077 | 25.1445 | 25.1741 |
| Opioid growth factor receptor | Q9NZT2-2;Q9NZT2 | OGFR | Other | -0.033466667 | 0.024876096 | 27.4756 | 26.2698 | 27.4529 | 27.4021 | 26.8225 | 26.8733 |
| UDP-N-acetylglucosamine--peptide N-acetylglucosaminy | O15294;O15294-2;O15294-3 | OGT | Extracellular or secreted | -1.482333333 | 0.512620808 | 23.3982 | 24.832 | 24.7327 | 22.7852 | 20.983 | 24.7477 |
| Obg-like ATPase 1 | Q9NTK5;Q9NTK5-3;Q9NTK5- | OLA1 | Extracellular or secreted | -0.553533333 | 0.913269467 | 30.0003 | 30.6454 | 30.372 | 29.365 | 30.0252 | 29.9669 |
| Olfactomedin-4 | Q6UX06 | OLFM4 | Extracellular or secreted | 0.455166667 | 0.160757331 | 21.9113 | 23.2524 | 21.1049 | 20.9372 | 22.8654 | 23.8315 |
| Olfactomedin-like protein 3 | Q9NRN5-2;Q9NRN5;Q9NRN5 | OLFML3 | Extracellular or secreted | 0.567966667 | 0.261464225 | 20.7517 | 20.6333 | 22.8358 | 21.5068 | 21.5119 | 22.906 |
| Dynamin-like 120 kDa protein, mitochondrial;Dynamin-li | O60313;O60313-11;O60313- | OPA1 | Other | 0.3739 | 0.277638686 | 24.0252 | 25.099 | 25.6867 | 25.1061 | 25.2398 | 25.5867 |
| 5-oxoprolinase | O14841 | OPLAH | Other | 1.5108 | 0.404763883 | 21.2327 | 21.5677 | 25.1396 | 22.6341 | 23.9423 | 25.896 |
| Optineurin | Q96CV9-2;Q96CV9;Q96CV9- | OPTN | Extracellular or secreted | 1.510466667 | 1.061947204 | 24.3257 | 23.6541 | 25.3029 | 25.8699 | 25.9805 | 25.9637 |
| Oxysterol-binding protein 1 | P22059 | OSBP | Other | -0.3016 | 0.190069044 | 25.9056 | 26.455 | 27.531 | 25.6915 | 26.3189 | 26.9764 |
| Oxysterol-binding protein-related protein 9 | Q96SU4-5;Q96SU4-4;Q96SU | OSBPL9 | Other | -0.276866667 | 0.081743417 | 22.2257 | 20.5977 | 22.729 | 19.6073 | 22.3022 | 22.8123 |
| Probable tRNA N6-adenosine threonylcarbamoyltransfer | Q9NPF4 | OSGEP | Other | -0.168866667 | 0.332407959 | 25.9825 | 25.6602 | 25.6997 | 25.5085 | 25.3706 | 25.9567 |
| Oncostatin-M-specific receptor subunit beta | Q99650-2;Q99650 | OSMR | Extracellular or secreted | 0.688366667 | 0.49292294 | 27.3313 | 26.4626 | 25.5727 | 27.6006 | 27.2498 | 26.5813 |
| Osteoclast-stimulating factor 1 | Q92882 | OSTF1 | Extracellular or secreted | -0.198866667 | 0.139055323 | 25.7485 | 26.9717 | 26.4487 | 25.4608 | 26.8042 | 26.3073 |
| Ubiquitin thioesterase OTUB1 | Q96FW1 | OTUB1 | Extracellular or secreted | 0.1305 | 0.43813637 | 30.0923 | 29.8505 | 30.1442 | 29.9811 | 30.2265 | 30.2709 |
| OTU domain-containing protein 6B | Q8N6M0 | OTUD6B | Other | -0.864566667 | 0.719138087 | 24.6614 | 24.1793 | 24.7762 | 22.9146 | 24.4974 | 23.6112 |
| Ovarian cancer-associated gene 2 protein | Q8WZ82 | OVCA2 | Other | -1.172466667 | 0.267686664 | 24.0277 | 24.6037 | 24.5667 | 20.0599 | 24.4543 | 25.1665 |
| Succinyl-CoA:3-ketoacid coenzyme A transferase 1, mito | P55809 | OXCT1 | Extracellular or secreted | -0.747466667 | 0.665002335 | 28.1322 | 27.685 | 28.3661 | 27.9199 | 27.5316 | 26.4894 |
| Oxidation resistance protein 1 | Q8N573-4;Q8N573-8;Q8N57 | OXR1 | Other | -3.240866667 | 1.779782836 | 29.4414 | 29.0446 | 28.5423 | 26.3538 | 24.6289 | 26.323 |
| 3-oxoacyl-[acyl-carrier-protein] synthase, mitochondrial | Q9NWU1;Q9NWU1-2 | OXSM | Other | -1.200766667 | 0.893461821 | 24.9178 | 24.5808 | 26.2231 | 23.7037 | 24.7186 | 23.6971 |
| Serine/threonine-protein kinase OSR1 | O95747 | OXSR1 | Extracellular or secreted | -1.144 | 1.515971947 | 28.2027 | 27.3131 | 27.9825 | 26.7019 | 26.411 | 26.9534 |
| Prolyl 4-hydroxylase subunit alpha-1 | P13674-3;P13674-2;P13674 | P4HA1 | Extracellular or secreted | 0.562566667 | 0.661255258 | 27.7363 | 26.9778 | 27.9956 | 28.5701 | 27.8242 | 28.0031 |
| Prolyl 4-hydroxylase subunit alpha-2 | O15460-2;O15460 | P4HA2 | Extracellular or secreted | -0.9779 | 0.705495829 | 25.5604 | 25.2458 | 26.7447 | 24.0175 | 25.4634 | 25.1363 |
| Protein disulfide-isomerase | P07237 | P4HB | Extracellular or secreted | -0.117433333 | 0.155178232 | 33.4387 | 33.2219 | 33.9505 | 33.7192 | 33.4491 | 33.0905 |
| Proliferation-associated protein 2G4 | Q9UQ80;Q9UQ80-2 | PA2G4 | Extracellular or secreted | -0.471666667 | 0.433392576 | 30.5815 | 31.8021 | 31.2866 | 30.1518 | 31.0931 | 31.0103 |
| Proteasomal ATPase-associated factor 1 | Q9BRP4-2;Q9BRP4;Q9BRP4- | PAAF1 | Other | -1.762666667 | 0.718417594 | 23.6322 | 21.5501 | 24.7326 | 22.1373 | 20.9777 | 21.5119 |
| Polyadenylate-binding protein 1 | P11940;P11940-2 | PABPC1 | Extracellular or secreted | 0.0219 | 0.023999207 | 30.2367 | 30.3302 | 30.8696 | 30.077 | 30.5478 | 30.8774 |
| Polyadenylate-binding protein 4 | Q13310-2;Q13310;Q13310-3 | PABPC4 | Other | -0.008966667 | 0.005821451 | 27.6959 | 27.2796 | 28.8102 | 27.7756 | 27.7206 | 28.2626 |
| Polyadenylate-binding protein 2 | Q86U42-2;Q86U42;Q92843- | PABPN1 | Other | 0.5898 | 0.504308417 | 26.1739 | 26.46 | 27.6097 | 27.7167 | 27.1707 | 27.1256 |
| Protein kinase C and casein kinase substrate in neurons p | Q9UNF0-2;Q9UNF0 | PACSIN2 | Extracellular or secreted | -0.0564 | 0.118279349 | 27.3512 | 27.8784 | 27.5861 | 27.4201 | 27.5532 | 27.6732 |
| Protein kinase C and casein kinase substrate in neurons p | Q9UKS6 | PACSIN3 | Extracellular or secreted | -0.0426 | 0.041896393 | 25.0656 | 26.0839 | 25.538 | 25.4753 | 25.8395 | 25.2449 |
| RNA polymerase II-associated factor 1 homolog | Q8N7H5;Q8N7H5-3;Q8N7H5 | PAF1 | Other | -0.412733333 | 0.180873381 | 26.6734 | 24.737 | 25.5361 | 24.8538 | 24.3332 | 26.5213 |
| Platelet-activating factor acetylhydrolase IB subunit alph | P43034 | PAFAH1B1 | Extracellular or secreted | -0.1967 | 0.325940504 | 30.1446 | 30.5843 | 29.8296 | 29.8297 | 30.1152 | 30.0235 |
| Platelet-activating factor acetylhydrolase IB subunit beta | P68402;P68402-3;P68402-2; | PAFAH1B2 | Extracellular or secreted | -0.442633333 | 0.355154686 | 27.1911 | 27.6294 | 27.7987 | 26.1871 | 27.5616 | 27.5426 |

| | | | | | | | | | | | |
|---|---|---|---|---|---|---|---|---|---|---|---|
| Platelet-activating factor acetylhydrolase IB subunit gam | Q15102 | PAFAH1B3 | Extracellular or secreted | 0.2353 | 0.223662383 | 27.1105 | 27.4384 | 28.0617 | 27.2356 | 27.8027 | 28.2782 |
| Multifunctional protein ADE2;Phosphoribosylaminoimida | P22234;P22234-2 | PAICS | Extracellular or secreted | -1.2114 | 1.490075789 | 29.4844 | 29.6673 | 29.9163 | 28.1345 | 28.2737 | 29.0256 |
| Polyadenylate-binding protein-interacting protein 1 | Q9H074-3;Q9H074-2;Q9H07 | PAIP1 | Other | -0.0848 | 0.165891691 | 25.4728 | 25.6112 | 25.2203 | 25.4948 | 25.0431 | 25.512 |
| Polyadenylate-binding protein-interacting protein 2 | Q9BPZ3 | PAIP2 | Other | -1.997266667 | 1.231588685 | 24.396 | 25.2393 | 24.5665 | 22.7995 | 21.6946 | 23.7159 |
| Serine/threonine-protein kinase PAK 1 | Q13153;Q13153-2 | PAK1 | Extracellular or secreted | -0.722033333 | 0.959166291 | 24.2698 | 23.8636 | 23.304 | 23.074 | 23.4343 | 22.763 |
| p21-activated protein kinase-interacting protein 1 | Q9NWT1 | PAK1IP1 | Extracellular or secreted | -3.149566667 | 1.478702406 | 24.5836 | 26.1895 | 27.1996 | 21.9405 | 23.1797 | 23.4038 |
| Serine/threonine-protein kinase PAK 2;PAK-2p27;PAK-2p | Q13177 | PAK2 | Extracellular or secreted | 0.391133333 | 0.36869175 | 26.8742 | 26.6217 | 27.1073 | 26.5736 | 27.2758 | 27.9272 |
| Serine/threonine-protein kinase PAK 4 | O96013 | PAK4 | Other | -0.900733333 | 0.452884619 | 23.0665 | 21.8392 | 23.8098 | 23.2228 | 21.0897 | 21.7008 |
| Palladin | Q8WX93-3;Q8WX93;Q8WX9 | PALLD | Extracellular or secreted | -0.2284 | 0.472368067 | 27.648 | 27.819 | 28.008 | 27.4155 | 27.4285 | 27.9458 |
| Peptidyl-glycine alpha-amidating monooxygenase;Peptid | P19021-2;P19021-4;P19021- | PAM | Extracellular or secreted | 0.3813 | 0.211852493 | 26.8649 | 27.9236 | 25.795 | 26.7304 | 27.3165 | 27.6805 |
| Iron/zinc purple acid phosphatase-like protein | Q6ZNF0 | PAPL | Extracellular or secreted | 3.907833333 | 0.880573411 | 26.6854 | 26.4442 | 21.7786 | 28.8373 | 29.2772 | 28.5172 |
| Poly(A) polymerase alpha | P51003 | PAPOLA | Other | 0.871866667 | 0.419600788 | 24.5479 | 24.2891 | 26.7491 | 25.9281 | 26.4138 | 25.8598 |
| Pappalysin-1 | Q13219 | PAPPA | Extracellular or secreted | 2.782466667 | 0.858761106 | 24.0928 | 27.1042 | 22.9809 | 28.5929 | 26.5541 | 27.3783 |
| Bifunctional 3-phosphoadenosine 5-phosphosulfate synt | O43252 | PAPSS1 | Other | -0.1192 | 0.043711852 | 27.6623 | 28.3427 | 28.5314 | 26.4776 | 28.298 | 29.4032 |
| Protein deglycase DJ-1 | Q99497 | PARK7 | Extracellular or secreted | 0.036966667 | 0.025160236 | 33.9429 | 33.5359 | 33.0175 | 34.2619 | 33.4917 | 32.8536 |
| Poly(A)-specific ribonuclease PARN | O95453-2;O95453;O95453-3 | PARN | Other | -1.579733333 | 0.712582448 | 23.2215 | 23.3811 | 25.2037 | 20.8121 | 22.9556 | 23.2994 |
| Poly [ADP-ribose] polymerase 1 | P09874 | PARP1 | Other | -1.4825 | 0.980499908 | 31.6186 | 31.6655 | 32.0812 | 29.4348 | 30.1699 | 31.3131 |
| Alpha-parvin | Q9NVD7 | PARVA | Other | -1.291233333 | 0.906970201 | 25.6016 | 25.25 | 25.0125 | 23.2575 | 23.7115 | 25.0214 |
| PRKC apoptosis WT1 regulator protein | Q96IZ0 | PAWR | Extracellular or secreted | -0.5463 | 0.239764953 | 23.6508 | 22.8248 | 23.9094 | 21.3431 | 23.4908 | 23.9122 |
| Protein PBDC1 | Q9BVG4 | PBDC1 | Other | -0.0051 | 0.008760593 | 27.5759 | 27.2645 | 27.2192 | 27.1105 | 27.6364 | 27.2974 |
| Lymphokine-activated killer T-cell-originated protein kina | Q96KB5;Q96KB5-2 | PBK | Other | -2.100733333 | 1.751534405 | 27.8157 | 26.741 | 27.3831 | 25.2637 | 25.0785 | 25.2954 |
| Pterin-4-alpha-carbinolamine dehydratase | P61457 | PCBD1 | Extracellular or secreted | 0.203733333 | 0.168756918 | 27.1746 | 27.2644 | 25.9594 | 26.9441 | 27.1479 | 26.9176 |
| Poly(rC)-binding protein 1 | Q15365 | PCBP1 | Extracellular or secreted | -0.616933333 | 0.535955885 | 30.0457 | 30.1161 | 30.7538 | 28.8912 | 29.8167 | 30.3569 |
| Poly(rC)-binding protein 2 | Q15366-6;Q15366-3;Q1536 | PCBP2 | Extracellular or secreted | -0.818866667 | 0.490505157 | 27.6071 | 27.9144 | 28.3907 | 25.9347 | 27.4841 | 28.0368 |
| Protocadherin-20 | Q8N6Y1 | PCDH20 | Extracellular or secreted | 2.653166667 | 0.851616727 | 24.6444 | 24.6926 | 21.2498 | 25.739 | 26.4139 | 26.3934 |
| Pre-mRNA cleavage complex 2 protein Pcf11 | O94913 | PCF11 | Extracellular or secreted | -0.334033333 | 0.165141352 | 21.1518 | 20.8617 | 21.575 | 20.4893 | 22.2058 | 19.8913 |
| Polycomb group RING finger protein 6 | Q9BYE7-3;Q9BYE7 | PCGF6 | Other | 0.829933333 | 0.812160823 | 26.0667 | 26.8593 | 26.1253 | 27.2889 | 27.7774 | 26.4748 |
| PCI domain-containing protein 2 | Q5JVF3-3;Q5JVF3-2;Q5JVF3; | PCID2 | Other | -0.010133333 | 0.001677423 | 19.4547 | 25.2839 | 24.2966 | 21.9739 | 22.9931 | 24.0378 |
| Phosphoenolpyruvate carboxykinase [GTP], mitochondri | Q16822;Q16822-3;Q16822-2 | PCK2 | Extracellular or secreted | -1.477033333 | 0.523077277 | 25.3678 | 26.1313 | 28.8913 | 24.8027 | 25.6449 | 25.5117 |
| Protein piccolo | Q9Y6V0-6;Q9Y6V0 | PCLO | Extracellular or secreted | 0.455666667 | 0.195680906 | 26.8797 | 27.5233 | 26.3154 | 28.93 | 26.7424 | 26.413 |
| Pericentriolar material 1 protein | Q15154-5;Q15154-4;Q15154 | PCM1 | Other | 0.823866667 | 0.399851593 | 25.4186 | 27.7037 | 27.7305 | 27.5013 | 28.2944 | 27.5287 |
| Protein-L-isoaspartate(D-aspartate) O-methyltransferase | P22061;P22061-2 | PCMT1 | Extracellular or secreted | -1.168033333 | 0.851098804 | 29.3771 | 29.7492 | 29.5272 | 27.3784 | 28.9538 | 28.8172 |
| Proliferating cell nuclear antigen | P12004 | PCNA | Extracellular or secreted | 0.308066667 | 0.328549651 | 32.6862 | 32.799 | 31.7479 | 33.0081 | 32.7101 | 32.4391 |
| PEST proteolytic signal-containing nuclear protein | Q8WW12;Q8WW12-2 | PCNP | Other | -0.623233333 | 1.952519606 | 27.6736 | 27.6881 | 27.9809 | 26.9951 | 27.3318 | 27.146 |
| Pecanex-like protein 2 | A6NKB5 | PCNXL2 | Other | -0.924933333 | 0.683027847 | 24.8622 | 23.6797 | 23.1442 | 22.4831 | 22.92 | 23.5082 |
| Procollagen C-endopeptidase enhancer 2 | Q9UKZ9 | PCOLCE2 | Extracellular or secreted | -1.0471 | 0.530668214 | 24.2005 | 24.0654 | 23.5261 | 21.4036 | 23.431 | 23.8161 |
| Proprotein convertase subtilisin/kexin type 6 | P29122-6;P29122-5;P29122- | PCSK6 | Extracellular or secreted | -0.0994 | 0.071751901 | 22.2829 | 22.8381 | 21.3196 | 21.785 | 22.0889 | 22.2685 |
| Proprotein convertase subtilisin/kexin type 9 | Q8NBP7 | PCSK9 | Extracellular or secreted | 1.5495 | 1.588810793 | 31.7973 | 31.2054 | 31.7235 | 32.5229 | 33.1614 | 33.6904 |
| Choline-phosphate cytidylyltransferase A | P49585 | PCYT1A | Other | -1.7935 | 2.253049045 | 23.2294 | 22.5557 | 22.9107 | 21.1584 | 21.5061 | 20.6508 |
| Ethanolamine-phosphate cytidylyltransferase | Q99447;Q99447-3;Q99447-2 | PCYT2 | Other | 1.561633333 | 0.425968945 | 22.6558 | 22.1196 | 23.8852 | 22.1888 | 24.2535 | 26.9032 |
| 28 kDa heat- and acid-stable phosphoprotein | Q13442 | PDAP1 | Other | 1.289566667 | 0.712954305 | 28.2112 | 26.3037 | 28.4499 | 29.7078 | 28.3518 | 28.7739 |
| Programmed cell death protein 10 | Q9BUL8 | PDCD10 | Extracellular or secreted | -0.5406 | 1.071863236 | 25.1818 | 25.6134 | 25.5 | 24.6768 | 24.7317 | 25.2649 |
| Programmed cell death protein 5 | O14737;O14737-2 | PDCD5 | Extracellular or secreted | -0.4779 | 0.561086124 | 28.9748 | 29.3507 | 28.3648 | 28.3151 | 28.8822 | 28.0593 |
| Programmed cell death protein 6 | O75340;O75340-2 | PDCD6 | Extracellular or secreted | 0.364433333 | 0.722037 | 25.5305 | 25.3345 | 25.9805 | 26.0227 | 25.8622 | 26.0539 |
| Programmed cell death 6-interacting protein | Q8WUM4;Q8WUM4-2 | PDCD6IP | Extracellular or secreted | -0.4751 | 0.988392437 | 30.0193 | 30.6257 | 30.4323 | 29.7131 | 29.8144 | 30.1245 |
| Phosducin-like protein 3 | Q9H2J4 | PDCL3 | Other | -0.0997 | 0.129530784 | 22.2229 | 22.7953 | 22.1902 | 22.2838 | 22.6643 | 21.9612 |
| 2,5-phosphodiesterase 12 | Q6L8Q7-2;Q6L8Q7 | PDE12 | Other | -1.2564 | 0.590164788 | 22.7151 | 22.3605 | 24.4256 | 20.5419 | 22.3124 | 22.8777 |
| Retinal rod rhodopsin-sensitive cGMP 3,5-cyclic phospho | O43924 | PDE6D | Other | -0.603 | 0.860094707 | 24.1988 | 24.796 | 24.3342 | 24.1139 | 24.0854 | 23.3207 |
| Pyruvate dehydrogenase E1 component subunit alpha, so | P08559-3;P08559;P08559-4; | PDHA1 | Extracellular or secreted | -1.223433333 | 0.266273587 | 21.3676 | 21.5927 | 26.4675 | 22.151 | 22.4291 | 21.1774 |
| Pyruvate dehydrogenase E1 component subunit beta, mi | P11177-2;P11177;P11177-3 | PDHB | Extracellular or secreted | -0.035166667 | 0.007095223 | 21.5989 | 22.2002 | 26.382 | 24.1506 | 22.8867 | 23.0383 |
| Pyruvate dehydrogenase protein X component, mitochon | O00330-3;O00330 | PDHX | Other | 0.873833333 | 0.202854023 | 23.3815 | 21.9381 | 25.6178 | 26.9557 | 22.7006 | 23.9026 |
| Protein disulfide-isomerase A3 | P30101 | PDIA3 | Extracellular or secreted | -0.391 | 0.343873134 | 34.3766 | 34.4004 | 34.6467 | 34.8012 | 34.1081 | 33.3414 |
| Protein disulfide-isomerase A4 | P13667 | PDIA4 | Extracellular or secreted | -0.5171 | 0.694761757 | 32.5089 | 32.2253 | 32.7306 | 32.4975 | 31.8835 | 31.5325 |
| Protein disulfide-isomerase A5 | Q14554 | PDIA5 | Extracellular or secreted | -1.296766667 | 1.155468333 | 26.2162 | 26.6432 | 27.5917 | 25.7924 | 25.5463 | 25.2221 |
| Protein disulfide-isomerase A6 | Q15084-3;Q15084;Q15084-4 | PDIA6 | Extracellular or secreted | -0.761866667 | 0.616589545 | 31.1202 | 30.9268 | 32.4196 | 30.6793 | 30.5488 | 30.9529 |
| PDZ and LIM domain protein 1 | O00151 | PDLIM1 | Other | -0.3917 | 0.375972959 | 28.2961 | 29.3485 | 29.4378 | 28.205 | 28.8127 | 28.8896 |
| PDZ and LIM domain protein 2 | Q96JY6;Q96JY6-5;Q96JY6-2; | PDLIM2 | Extracellular or secreted | -0.563866667 | 0.246809592 | 23.532 | 23.7085 | 23.4909 | 24.1551 | 21.4037 | 23.481 |
| PDZ and LIM domain protein 5 | Q96HC4;Q96HC4-7;Q96HC4- | PDLIM5 | Extracellular or secreted | -0.676866667 | 1.062243153 | 28.9144 | 29.1736 | 28.6598 | 27.7912 | 28.3176 | 28.6084 |
| PDZ and LIM domain protein 7 | Q9NR12-2;Q9NR12;Q9NR12 | PDLIM7 | Other | 1.410866667 | 0.301895155 | 20.3162 | 25.5002 | 25.4338 | 24.6022 | 25.6325 | 25.2481 |
| Pyruvate dehydrogenase phosphatase regulatory subuni | Q8NCN5 | PDPR | Other | -2.8708 | 0.743488669 | 22.2547 | 23.2367 | 27.0017 | 20.6841 | 21.3662 | 21.8304 |
| Sister chromatid cohesion protein PDS5 homolog A | Q29RF7;Q29RF7-3 | PDS5A | Extracellular or secreted | -3.006366667 | 1.251455383 | 28.1415 | 26.8782 | 29.8136 | 25.0852 | 24.797 | 25.932 |
| Sister chromatid cohesion protein PDS5 homolog B | Q9NTI5-2;Q9NTI5 | PDS5B | Other | -1.4906 | 0.989737039 | 27.3205 | 26.0567 | 26.8424 | 26.3747 | 24.6123 | 24.7608 |
| Pyridoxal kinase | O00764;O00764-3;O00764-2 | PDXK | Extracellular or secreted | -0.407566667 | 1.176451815 | 30.1948 | 29.8948 | 30.2642 | 29.7839 | 29.8658 | 29.4814 |
| Pyridoxal phosphate phosphatase | Q96GD0 | PDXP | Extracellular or secreted | -0.498933333 | 0.910116148 | 22.0045 | 22.4167 | 22.4881 | 21.6842 | 22.1955 | 21.5328 |
| PDZ domain-containing protein 11 | Q5EBL8;Q5EBL8-2 | PDZD11 | Extracellular or secreted | -0.369233333 | 0.625662997 | 23.9264 | 23.3044 | 23.9088 | 23.0981 | 23.6614 | 23.2724 |
| Astrocytic phosphoprotein PEA-15 | Q15121;Q15121-2 | PEA15 | Other | 0.3796 | 0.131724351 | 22.7629 | 22.3003 | 24.6707 | 25.1129 | 23.2384 | 22.5214 |
| Phosphatidylethanolamine-binding protein 1;Hippocamp | P30086 | PEBP1 | Extracellular or secreted | -0.099 | 0.070766264 | 32.8276 | 32.3272 | 31.6941 | 32.7434 | 32.3054 | 31.5031 |
| Proline-, glutamic acid- and leucine-rich protein 1 | Q8IZL8 | PELP1 | Other | -2.001633333 | 0.717214597 | 26.7375 | 24.5619 | 25.215 | 24.87 | 24.1831 | 21.4564 |
| Xaa-Pro dipeptidase | P12955;P12955-3;P12955-2 | PEPD | Extracellular or secreted | -0.281033333 | 0.180552998 | 29.274 | 29.1263 | 28.3502 | 29.5517 | 28.5463 | 27.8094 |
| Pescadillo homolog | O00541-2;O00541 | PES1 | Other | -0.371333333 | 0.126085679 | 21.0281 | 23.1675 | 23.93 | 23.5587 | 21.477 | 21.9759 |
| Peroxisomal membrane protein PEX14 | O75381-2;O75381 | PEX14 | Other | -1.0994 | 0.324096597 | 24.4681 | 20.6648 | 24.2575 | 22.111 | 21.2271 | 22.7541 |
| Platelet factor 4;Platelet factor 4, short form;Platelet fac | P02776;P10720 | PF4;PF4V1 | Extracellular or secreted | 2.6752 | 0.844286601 | 25.1973 | 27.8197 | 23.7368 | 27.4559 | 28.9381 | 28.3854 |
| Phosphoribosylformylglycinamidine synthase | O15067 | PFAS | Extracellular or secreted | -0.7985 | 0.978702976 | 30.3626 | 30.1519 | 29.6616 | 29.8251 | 29.173 | 28.7825 |
| Prefoldin subunit 1 | O60925 | PFDN1 | Other | 0.7313 | 0.577443873 | 27.4492 | 27.7994 | 27.0907 | 27.2499 | 28.8526 | 28.4307 |
| Prefoldin subunit 2 | Q9UHV9 | PFDN2 | Extracellular or secreted | 0.159633333 | 0.215606376 | 28.3061 | 28.1214 | 28.2049 | 27.8663 | 28.7547 | 28.4903 |
| Prefoldin subunit 4 | Q9NQP4 | PFDN4 | Other | 0.324566667 | 0.302242101 | 25.3363 | 26.4264 | 26.1006 | 26.8649 | 26.0079 | 25.9642 |
| Prefoldin subunit 5 | Q99471;Q99471-3 | PFDN5 | Other | 0.128066667 | 0.10493623 | 28.5586 | 28.8872 | 28.2497 | 28.0596 | 29.3944 | 28.6257 |
| Prefoldin subunit 6 | O15212 | PFDN6 | Other | 0.140833333 | 0.172760025 | 28.0394 | 28.8196 | 28.4573 | 28.159 | 28.7706 | 28.8092 |
| ATP-dependent 6-phosphofructokinase, liver type | P17858;P17858-2 | PFKL | Extracellular or secreted | -0.406933333 | 0.09930175 | 25.1003 | 24.7165 | 27.7633 | 23.3673 | 25.8282 | 27.1638 |
| ATP-dependent 6-phosphofructokinase, muscle type | P08237;P08237-3;P08237-2 | PFKM | Extracellular or secreted | -0.082333333 | 0.024266772 | 23.6579 | 23.2811 | 26.5673 | 23.7631 | 24.6289 | 24.8673 |
| ATP-dependent 6-phosphofructokinase, platelet type | Q01813;Q01813-2 | PFKP | Extracellular or secreted | 0.077533333 | 0.032790699 | 28.3364 | 27.9702 | 30.0798 | 28.1297 | 28.8007 | 29.6886 |
| Profilin-1 | P07737 | PFN1 | Extracellular or secreted | -0.0773 | 0.073605878 | 33.5665 | 34.3337 | 33.8345 | 33.3459 | 34.3499 | 33.807 |
| Profilin-2 | P35080-2 | PFN2 | Extracellular or secreted | -0.404633333 | 0.602150845 | 27.4392 | 27.7799 | 27.7256 | 26.7332 | 27.4977 | 27.4999 |
| Phosphoglycerate mutase 1 | P18669 | PGAM1 | Extracellular or secreted | -0.197166667 | 0.492336582 | 33.4108 | 33.1342 | 32.9349 | 33.1448 | 32.9587 | 32.7849 |
| 6-phosphogluconate dehydrogenase, decarboxylating | P52209-2;P52209 | PGD | Extracellular or secreted | 0.530233333 | 0.552040431 | 30.8492 | 31.9509 | 30.9275 | 31.4135 | 32.1133 | 31.7915 |
| Geranylgeranyl transferase type-1 subunit beta | P53609-2;P53609 | PGGT1B | Other | 0.889933333 | 0.254314356 | 20.0513 | 23.8555 | 23.8669 | 23.2386 | 23.7592 | 23.4457 |
| Phosphoglycerate kinase 1 | P00558;P00558-2 | PGK1 | Extracellular or secreted | 0.1469 | 0.206410596 | 35.1865 | 35.5293 | 34.9067 | 35.6832 | 35.4093 | 34.9707 |
| Phosphoglycerate kinase 2 | P07205 | PGK2 | Extracellular or secreted | 0.532766667 | 1.338092933 | 25.5357 | 25.9404 | 25.4763 | 26.3318 | 26.2114 | 26.0075 |
| 6-phosphogluconolactonase | O95336 | PGLS | Extracellular or secreted | 0.076066667 | 0.145481105 | 30.2543 | 30.408 | 30.1519 | 30.0647 | 30.6602 | 30.3175 |
| Phosphoglucomutase-1 | P36871;P36871-2;P36871-3 | PGM1 | Extracellular or secreted | -0.415333333 | 1.910227544 | 32.4217 | 32.3108 | 32.1752 | 31.9331 | 31.9287 | 31.7999 |
| Phosphoglucomutase-2 | Q96G03 | PGM2 | Extracellular or secreted | -0.290366667 | 0.597596813 | 30.0924 | 29.6003 | 29.7275 | 29.8294 | 29.4123 | 29.3074 |
| Glucose 1,6-bisphosphate synthase | Q6PCE3 | PGM2L1 | Other | -1.0041 | 0.470261757 | 24.8188 | 25.2798 | 25.3271 | 23.4211 | 23.2447 | 25.7476 |
| Phosphoacetylglucosamine mutase | O95394;O95394-3;O95394-4 | PGM3 | Other | -0.3899 | 1.231534951 | 28.9133 | 28.6732 | 28.6013 | 28.474 | 28.4287 | 28.1154 |
| Phosphoglycolate phosphatase | A6NDG6 | PGP | Other | 0.068033333 | 0.087823154 | 27.6259 | 27.8653 | 27.3522 | 27.2271 | 27.9408 | 27.8796 |
| Membrane-associated progesterone receptor componen | O00264;O00264-2 | PGRMC1 | Extracellular or secreted | 1.2508 | 1.005429548 | 25.8039 | 24.5606 | 25.8102 | 27.2405 | 26.8288 | 25.8578 |
| Membrane-associated progesterone receptor componen | O15173;O15173-2 | PGRMC2 | Other | -1.608366667 | 1.591275111 | 24.5422 | 23.7055 | 24.3504 | 22.1051 | 22.3795 | 23.2884 |
| Prohibitin | P35232;P35232-2 | PHB | Extracellular or secreted | 2.9781 | 1.334382572 | 21.7693 | 23.9946 | 21.871 | 25.8983 | 25.2334 | 25.4375 |
| Prohibitin-2 | Q99623;Q99623-2 | PHB2 | Extracellular or secreted | 5.233633333 | 1.813974022 | 20.2409 | 19.6149 | 20.7268 | 26.9974 | 25.171 | 24.1151 |
| PHD finger protein 10 | Q8WUB8-3;Q8WUB8-2;Q8W | PHF10 | Other | -0.484 | 1.073270713 | 21.5696 | 21.3643 | 21.5797 | 21.2704 | 20.705 | 21.0862 |
| PHD finger protein 3 | Q92576-2;Q92576 | PHF3 | Other | -2.0127 | 0.617164281 | 23.1949 | 22.086 | 26.2252 | 21.5838 | 21.4385 | 22.4457 |
| PHD finger-like domain-containing protein 5A | Q7RTV0 | PHF5A | Other | 0.0138 | 0.006976992 | 24.5528 | 25.1819 | 26.2349 | 24.7013 | 25.1686 | 26.1411 |
| PHD finger protein 6 | Q8IWS0-5;Q8IWS0-3;Q8IWS | PHF6 | Other | 0.7252 | 0.172878001 | 21.618 | 22.1518 | 22.1124 | 21.7494 | 20.7394 | 25.569 |
| D-3-phosphoglycerate dehydrogenase | O43175 | PHGDH | Extracellular or secreted | -0.373566667 | 0.193804325 | 27.4594 | 27.7808 | 29.5548 | 27.3265 | 28.0305 | 28.3173 |
| Phosphorylase b kinase regulatory subunit beta | Q93100-4;Q93100-2;Q9310 | PHKB | Extracellular or secreted | 2.335033333 | 0.524095105 | 20.6974 | 22.7309 | 23.3163 | 27.2633 | 21.5254 | 24.961 |
| 14 kDa phosphohistidine phosphatase | Q9NRX4;Q9NRX4-2 | PHPT1 | Extracellular or secreted | 0.1266 | 0.062027031 | 24.4529 | 26.3459 | 25.5562 | 24.8224 | 26.3639 | 25.5485 |
| 5-phosphohydroxy-L-lysine phospho-lyase | Q8IUZ5;Q8IUZ5-3 | PHYKPL | Extracellular or secreted | 0.329766667 | 0.283317876 | 22.7384 | 22.3217 | 22.3135 | 23.3434 | 23.0532 | 21.9663 |
| Elafin | P19957 | PI3 | Extracellular or secreted | 3.341533333 | 1.315135599 | 32.4783 | 34.0588 | 30.9163 | 36.6234 | 35.673 | 35.1816 |
| Phosphatidylinositol-binding clathrin assembly protein | Q13492-3;Q13492-2;Q13492 | PICALM | Extracellular or secreted | 0.172166667 | 0.127477586 | 26.5675 | 26.7153 | 27.8014 | 26.6947 | 27.163 | 27.743 |
| Peptidyl-prolyl cis-trans isomerase NIMA-interacting 4 | Q9Y237;Q9Y237-2 | PIN4 | Extracellular or secreted | -0.584666667 | 0.691727527 | 27.8199 | 27.8868 | 26.9748 | 26.5217 | 27.3642 | 27.0416 |
| Prolactin-inducible protein | P12273 | PIP | Extracellular or secreted | 2.229 | 1.112016783 | 23.5827 | 22.4488 | 24.9605 | 27.0314 | 25.1756 | 25.472 |
| Phosphatidylinositol 5-phosphate 4-kinase type-2 gamm | Q8TBX8-3;Q8TBX8 | PIP4K2C | Extracellular or secreted | 1.095466667 | 0.474451524 | 22.8898 | 23.2995 | 24.2415 | 23.225 | 24.3264 | 26.1658 |
| Pirin | O00625 | PIR | Other | -1.289066667 | 1.163329194 | 26.296 | 26.3059 | 26.4323 | 25.1236 | 25.6463 | 24.3971 |
| PITH domain-containing protein 1 | Q9GZP4-2;Q9GZP4 | PITHD1 | Other | -0.904866667 | 1.567454424 | 27.4975 | 26.888 | 26.9173 | 26.5006 | 25.9035 | 26.1841 |
| Phosphatidylinositol transfer protein alpha isoform | Q00169 | PITPNA | Extracellular or secreted | -0.393266667 | 0.458728102 | 26.7453 | 27.0851 | 27.0692 | 25.9342 | 26.9626 | 26.823 |
| Phosphatidylinositol transfer protein beta isoform | P48739;P48739-2;P48739-3 | PITPNB | Extracellular or secreted | -0.7211 | 0.706304702 | 28.2707 | 28.1483 | 28.6839 | 26.8762 | 27.9365 | 28.1269 |
| Presequence protease, mitochondrial | Q5JRX3;Q5JRX3-2;Q5JRX3-3 | PITRM1 | Other | -1.165366667 | 1.221267793 | 26.9786 | 27.7908 | 28.1527 | 26.4276 | 26.7771 | 26.2213 |
| Pituitary homeobox 1 | P78337 | PITX1 | Other | -0.2653 | 0.254472544 | 22.0783 | 22.2967 | 23.0419 | 22.7829 | 21.8172 | 22.0209 |
| Pyruvate kinase PKM | P14618;P14618-3 | PKM | Extracellular or secreted | 0.6474 | 0.962488105 | 34.3553 | 34.8436 | 34.5526 | 34.7298 | 35.4064 | 35.5575 |
| Pyruvate kinase PKM | P14618-2 | PKM | Extracellular or secreted | 0.191566667 | 0.10366857 | 28.11 | 28.1717 | 26.9777 | 28.9736 | 27.6843 | 27.1762 |

| | | | | | | | | | | | |
|---|---|---|---|---|---|---|---|---|---|---|---|
| Plakophilin-2 | Q99959-2;Q99959 | PKP2 | Extracellular or secreted | 1.895866667 | 1.40549854 | 22.198 | 21.763 | 21.1273 | 23.3055 | 24.5463 | 22.9241 |
| Plakophilin-3 | Q9Y446;Q9Y446-2 | PKP3 | Extracellular or secreted | 2.6328 | 0.797782708 | 20.8416 | 23.558 | 24.9487 | 27.4863 | 24.6276 | 25.1328 |
| Group XV phospholipase A2 | Q8NCC3;Q8NCC3-2 | PLA2G15 | Extracellular or secreted | 0.401766667 | 0.683658004 | 27.8931 | 27.8984 | 27.229 | 28.054 | 28.1845 | 27.9873 |
| Phospholipase A-2-activating protein | Q9Y263 | PLAA | Extracellular or secreted | -1.2955 | 0.350211561 | 20.7261 | 21.503 | 25.0595 | 22.2708 | 20.1718 | 20.9595 |
| Tissue-type plasminogen activator;Tissue-type plasminog | P00750;P00750-4 | PLAT | Extracellular or secreted | 2.224366667 | 0.800096788 | 28.7593 | 29.9584 | 26.4052 | 30.0345 | 30.6614 | 31.1001 |
| Urokinase-type plasminogen activator;Urokinase-type pl | P00749-2;P00749 | PLAU | Extracellular or secreted | -0.1327 | 0.068020351 | 28.3465 | 29.5887 | 27.3792 | 28.2115 | 28.3675 | 28.3373 |
| Urokinase plasminogen activator surface receptor | Q03405;Q03405-3;Q03405-2 | PLAUR | Extracellular or secreted | 0.683466667 | 0.421782441 | 29.6769 | 29.7233 | 28.1987 | 30.72 | 29.8577 | 29.0716 |
| Phospholipase B-like 1;Phospholipase B-like 1 chain A;Ph | Q6P4A8 | PLBD1 | Extracellular or secreted | -0.431633333 | 0.288839071 | 22.9088 | 22.5873 | 22.8213 | 21.4621 | 22.2159 | 23.3445 |
| Putative phospholipase B-like 2;Putative phospholipase E | Q8NHP8;Q8NHP8-2 | PLBD2 | Extracellular or secreted | 1.211933333 | 1.436926336 | 27.5547 | 27.6935 | 26.7225 | 28.8017 | 28.7172 | 28.0876 |
| 1-phosphatidylinositol 4,5-bisphosphate phosphodiester | Q01970;Q01970-2 | PLCB3 | Extracellular or secreted | -0.7857 | 2.044708125 | 29.4579 | 29.1859 | 29.1481 | 28.249 | 28.5054 | 28.6804 |
| 1-phosphatidylinositol 4,5-bisphosphate phosphodiester | P19174;P19174-2 | PLCG1 | Extracellular or secreted | -0.844033333 | 0.805009155 | 23.6943 | 23.6834 | 24.3395 | 22.2664 | 23.4362 | 23.4825 |
| 1-phosphatidylinositol 4,5-bisphosphate phosphodiester | Q4KWH8-3;Q4KWH8-4;Q4KV | PLCH1 | Extracellular or secreted | -0.9929 | 0.251230921 | 21.4416 | 24.6237 | 25.082 | 21.231 | 22.1421 | 24.7955 |
| Phospholipase D3 | Q8IV08 | PLD3 | Extracellular or secreted | 0.237533333 | 0.262127755 | 25.5188 | 25.4171 | 24.9556 | 24.9766 | 26.0369 | 25.5906 |
| Plectin | Q15149-2;Q15149-7;Q15149 | PLEC | Extracellular or secreted | 0.0364 | 0.074104115 | 22.7565 | 23.1945 | 23.1468 | 23.1112 | 23.2183 | 22.8775 |
| Plectin | Q15149-3 | PLEC | Extracellular or secreted | -0.6356 | 0.217466072 | 23.1249 | 20.8332 | 23.9802 | 22.3336 | 20.8218 | 22.8761 |
| Plectin | Q15149-4 | PLEC | Extracellular or secreted | -0.912933333 | 1.299584955 | 33.7403 | 33.8659 | 34.5658 | 32.9004 | 33.1406 | 33.3922 |
| Plectin | Q15149-9 | PLEC | Extracellular or secreted | 0.9442 | 0.539120903 | 21.2854 | 20.8763 | 22.788 | 22.2351 | 21.9534 | 23.5938 |
| Pleckstrin homology domain-containing family A membe | Q9HAU0-8;Q9HAU0;Q9HAU | PLEKHA5 | Other | -0.0663 | 0.059655857 | 25.1853 | 24.3177 | 24.0923 | 24.562 | 24.1206 | 24.7138 |
| Perilipin-3 | O60664-4;O60664;O60664-3 | PLIN3 | Other | 0.305566667 | 0.296890347 | 28.7196 | 28.4092 | 28.9937 | 28.6737 | 29.7403 | 28.6252 |
| Procollagen-lysine,2-oxoglutarate 5-dioxygenase 1 | Q02809;Q02809-2 | PLOD1 | Extracellular or secreted | 0.117366667 | 0.133845629 | 30.1801 | 30.843 | 29.9709 | 30.1169 | 30.7491 | 30.4801 |
| Procollagen-lysine,2-oxoglutarate 5-dioxygenase 2 | O00469-2;O00469;O00469-3 | PLOD2 | Extracellular or secreted | 0.437433333 | 0.804356698 | 29.9161 | 29.9604 | 29.3213 | 30.2387 | 30.2554 | 30.016 |
| Procollagen-lysine,2-oxoglutarate 5-dioxygenase 3 | O60568 | PLOD3 | Extracellular or secreted | 0.0072 | 0.014141183 | 30.2285 | 30.3433 | 30.1642 | 30.2836 | 30.5017 | 29.9723 |
| Proteolipid protein 2 | Q04941 | PLP2 | Extracellular or secreted | 4.5779 | 1.015265373 | 21.3245 | 20.284 | 21.8052 | 22.5035 | 27.6879 | 26.956 |
| Pleiotropic regulator 1 | O43660-2;O43660 | PLRG1 | Other | -1.026333333 | 1.234946803 | 30.2256 | 29.4245 | 29.2167 | 28.9073 | 28.6568 | 28.2237 |
| Plastin-1 | Q14651 | PLS1 | Extracellular or secreted | -0.766033333 | 0.407821942 | 27.0482 | 27.2154 | 27.6394 | 25.1279 | 27.2154 | 27.2616 |
| Plastin-3 | P13797;P13797-2;P13797-3 | PLS3 | Other | -0.452033333 | 1.387499005 | 30.2422 | 30.5909 | 30.2488 | 29.7168 | 30.0087 | 30.0003 |
| Phospholipid transfer protein | P55058;P55058-3;P55058-4; | PLTP | Extracellular or secreted | -0.872033333 | 1.01183819 | 26.3844 | 27.2507 | 26.4224 | 25.7271 | 25.3619 | 26.3524 |
| Plexin domain-containing protein 2 | Q6UX71;Q6UX71-2 | PLXDC2 | Extracellular or secreted | 1.169133333 | 0.917860882 | 26.4293 | 24.9637 | 25.0741 | 26.0858 | 27.247 | 26.6417 |
| Plexin-A1 | Q9UIW2 | PLXNA1 | Extracellular or secreted | -0.605466667 | 0.489725796 | 22.8648 | 22.8433 | 22.2816 | 21.7543 | 22.9688 | 21.4502 |
| Plexin-A3 | P51805 | PLXNA3 | Extracellular or secreted | 0.788033333 | 0.722313548 | 22.2155 | 22.5483 | 21.1776 | 22.4083 | 23.1815 | 22.7157 |
| Plexin-B2 | O15031 | PLXNB2 | Extracellular or secreted | 0.2355 | 1.179212186 | 27.6496 | 27.5037 | 27.3934 | 27.7575 | 27.8321 | 27.6636 |
| Peptidase M20 domain-containing protein 2 | Q8IYS1 | PM20D2 | Extracellular or secreted | -1.114866667 | 0.818645018 | 24.8256 | 24.5513 | 24.9279 | 22.6458 | 24.1414 | 24.173 |
| Protein PML | P29590-14;P29590-10;P2959 | PML | Other | 0.277833333 | 0.261462014 | 24.7073 | 25.8735 | 25.4893 | 25.1526 | 25.8764 | 25.8746 |
| Phosphomannomutase 2 | O15305;O15305-2 | PMM2 | Extracellular or secreted | -0.2075 | 0.113946929 | 24.0754 | 24.8672 | 25.1727 | 23.4049 | 24.8164 | 25.2715 |
| Mitochondrial-processing peptidase subunit beta | O75439 | PMPCB | Other | -0.7652 | 0.824836807 | 25.8412 | 25.6815 | 26.7073 | 24.9946 | 25.8842 | 25.0556 |
| Arginine/serine-rich protein PNISR | Q8TF01 | PNISR | Other | -0.389666667 | 0.258158238 | 22.0657 | 23.2365 | 22.5434 | 22.6632 | 22.7614 | 21.252 |
| Pinin | Q9H307 | PNN | Extracellular or secreted | -1.3521 | 0.631554255 | 27.881 | 25.7702 | 28.4413 | 25.6893 | 26.4659 | 25.881 |
| Purine nucleoside phosphorylase | P00491 | PNP | Extracellular or secreted | 0.010133333 | 0.007255346 | 32.3224 | 32.1228 | 31.141 | 32.193 | 32.0967 | 31.3269 |
| Pyridoxine-5-phosphate oxidase | Q9NVS9;Q9NVS9-3;Q9NVS9- | PNPO | Extracellular or secreted | -0.6868 | 0.822182651 | 28.1524 | 28.2236 | 27.2604 | 27.4909 | 27.302 | 26.7831 |
| Polyribonucleotide nucleotidyltransferase 1, mitochondr | Q8TCS8 | PNPT1 | Other | -1.6193 | 0.758571557 | 25.7615 | 26.1952 | 28.3066 | 24.0618 | 25.826 | 25.5176 |
| Podocalyxin | O00592-2;O00592 | PODXL | Extracellular or secreted | 1.375333333 | 1.044641031 | 25.5242 | 24.9123 | 24.9008 | 27.257 | 26.6242 | 25.5821 |
| GDP-fucose protein O-fucosyltransferase 1 | Q9H488 | POFUT1 | Extracellular or secreted | -1.590233333 | 1.90021686 | 27.5688 | 27.2278 | 28.1691 | 25.7916 | 26.5425 | 25.8609 |
| GDP-fucose protein O-fucosyltransferase 2 | Q9Y2G5-2;Q9Y2G5-1;Q9Y2G | POFUT2 | Extracellular or secreted | -2.816866667 | 1.912143974 | 23.6949 | 23.7581 | 25.0363 | 21.1558 | 21.147 | 21.7359 |
| Polymerase delta-interacting protein 2 | Q9Y2S7 | POLDIP2 | Other | -2.928833333 | 1.439597925 | 25.0571 | 24.6209 | 26.2674 | 21.4939 | 21.8194 | 23.8456 |
| Polymerase delta-interacting protein 3 | Q9BY77-2;Q9BY77 | POLDIP3 | Other | -0.232966667 | 0.110043213 | 21.6342 | 22.4531 | 23.6772 | 21.5422 | 23.2054 | 22.318 |
| DNA polymerase epsilon subunit 3 | Q9NRF9 | POLE3 | Other | -0.6401 | 0.691475321 | 25.2912 | 25.7556 | 25.6043 | 24.3627 | 25.5691 | 24.799 |
| DNA-directed RNA polymerases I and III subunit RPAC1 | O15160-2;O15160 | POLR1C | Other | -0.132033333 | 0.060834933 | 24.0659 | 24.4955 | 25.0736 | 23.2131 | 24.4939 | 25.5319 |
| DNA-directed RNA polymerase II subunit RPB1 | P24928 | POLR2A | Other | -0.389266667 | 0.126709506 | 24.2998 | 23.3637 | 26.2941 | 23.3483 | 23.7631 | 25.6784 |
| DNA-directed RNA polymerase II subunit RPB2 | P30876 | POLR2B | Other | -0.259566667 | 0.138827369 | 25.1914 | 25.0372 | 26.7205 | 25.6608 | 24.545 | 25.9646 |
| DNA-directed RNA polymerase II subunit RPB3 | P19387 | POLR2C | Other | -0.345133333 | 0.127505209 | 23.1426 | 23.3828 | 24.947 | 22.1605 | 23.335 | 24.9415 |
| DNA-directed RNA polymerase II subunit RPB4 | O15514 | POLR2D | Other | -1.4057 | 0.811885899 | 23.7944 | 23.8431 | 25.7548 | 23.6012 | 23.3545 | 22.2195 |
| DNA-directed RNA polymerases I, II, and III subunit RPAB | P19388 | POLR2E | Other | -0.775066667 | 1.125379358 | 26.3271 | 26.2281 | 26.7981 | 25.302 | 25.5611 | 26.165 |
| DNA-directed RNA polymerases I, II, and III subunit RPAB | P61218 | POLR2F | Other | -0.636066667 | 0.809738423 | 25.0088 | 24.0501 | 24.4191 | 24.3045 | 23.7006 | 23.5647 |
| DNA-directed RNA polymerase II subunit RPB7 | P62487 | POLR2G | Other | -0.729033333 | 1.363745371 | 25.2511 | 25.063 | 25.7133 | 24.4171 | 24.7796 | 24.6436 |
| DNA-directed RNA polymerases I, II, and III subunit RPAB | P52434;P52434-3;P52434-4 | POLR2H | Other | -0.723966667 | 1.441467329 | 28.5102 | 28.0272 | 27.9349 | 27.2968 | 27.7188 | 27.2848 |
| DNA-directed RNA polymerases I, II, and III subunit RPAB | P62875 | POLR2L | Other | -0.3721 | 0.082700818 | 20.5872 | 23.0898 | 24.1994 | 19.9132 | 23.1835 | 23.6634 |
| Protein O-linked-mannose beta-1,2-N-acetylglucosaminy | Q8WZA1;Q8WZA1-2 | POMGNT1 | Other | -0.024033333 | 0.016338023 | 21.0915 | 22.572 | 22.3062 | 22.0079 | 21.8048 | 22.0849 |
| Proteasome maturation protein | Q9Y244 | POMP | Other | 0.359833333 | 0.743646522 | 25.4794 | 25.7933 | 25.1568 | 25.9957 | 25.8477 | 25.6656 |
| Ribonucleases P/MRP protein subunit POP1 | Q99575 | POP1 | Extracellular or secreted | 0.0052 | 0.001098156 | 21.9668 | 22.1366 | 25.2162 | 21.2032 | 23.0297 | 25.1023 |
| NADPH--cytochrome P450 reductase | P16435 | POR | Other | -0.185966667 | 0.172090383 | 23.5888 | 23.3649 | 24.5892 | 23.6361 | 23.8051 | 23.5438 |
| Inorganic pyrophosphatase | Q15181 | PPA1 | Extracellular or secreted | -0.4092 | 0.539380487 | 29.8472 | 30.3159 | 30.4282 | 29.2402 | 30.0528 | 30.0707 |
| Inorganic pyrophosphatase 2, mitochondrial | Q9H2U2;Q9H2U2-2;Q9H2U2 | PPA2 | Extracellular or secreted | -0.662466667 | 0.380711438 | 26.9197 | 28.0018 | 28.9553 | 26.4984 | 27.9225 | 27.4685 |
| Amidophosphoribosyltransferase | Q06203 | PPAT | Other | 0.1022 | 0.075034617 | 23.993 | 24.5578 | 23.8158 | 23.8281 | 23.7952 | 25.0499 |
| Platelet basic protein;Connective tissue-activating peptic | P02775 | PPBP | Extracellular or secreted | 3.663233333 | 0.918501399 | 29.4373 | 30.435 | 25.4824 | 32.5713 | 33.1301 | 30.643 |
| Phosphopantothenoylcysteine decarboxylase | Q96CD2;Q96CD2-2 | PPCDC | Other | -0.262866667 | 0.236708471 | 25.2614 | 25.0463 | 24.008 | 24.5589 | 24.758 | 24.2102 |
| Phosphopantothenate--cysteine ligase | Q9HAB8-2;Q9HAB8 | PPCS | Extracellular or secreted | -0.601133333 | 0.119861591 | 24.6189 | 24.9743 | 24.5414 | 20.7545 | 25.2367 | 26.34 |
| Liprin-alpha-1 | Q13136-2;Q13136 | PPFIA1 | Extracellular or secreted | 1.685366667 | 0.226688585 | 24.9933 | 24.8949 | 19.8498 | 25.9792 | 28.3456 | 20.4693 |
| Liprin-beta-1 | Q86W92-3;Q86W92-4;Q86W | PPFIBP1 | Extracellular or secreted | -0.538466667 | 0.402641949 | 22.7446 | 22.8107 | 22.692 | 21.5795 | 21.85 | 23.2024 |
| Peptidyl-prolyl cis-trans isomerase A;Peptidyl-prolyl cis-t | P62937 | PPIA | Extracellular or secreted | 0.0479 | 0.040344484 | 33.0556 | 34.2286 | 33.7913 | 33.4363 | 34.1263 | 33.6566 |
| Peptidyl-prolyl cis-trans isomerase B | P23284 | PPIB | Extracellular or secreted | -0.256266667 | 0.212358896 | 32.8123 | 33.1331 | 32.6972 | 33.3724 | 32.5954 | 31.906 |
| Peptidyl-prolyl cis-trans isomerase C | P45877 | PPIC | Extracellular or secreted | 0.304166667 | 0.170205493 | 24.4194 | 25.9516 | 24.0533 | 24.5289 | 25.6011 | 25.2068 |
| Peptidyl-prolyl cis-trans isomerase D | Q08752 | PPID | Other | -0.420366667 | 0.278823053 | 25.4854 | 26.5392 | 26.7366 | 25.5567 | 25.2058 | 26.7376 |
| Peptidyl-prolyl cis-trans isomerase E | Q9UNP9;Q9UNP9-2;Q9UNP9 | PPIE | Other | -1.6183 | 1.620686507 | 29.1831 | 28.3447 | 28.0291 | 26.8555 | 27.2564 | 26.5901 |
| Peptidyl-prolyl cis-trans isomerase F, mitochondrial | P30405;P30405-2 | PPIF | Other | -1.810033333 | 1.340648202 | 29.2775 | 29.8874 | 29.3771 | 26.9804 | 28.5793 | 27.5522 |
| Peptidyl-prolyl cis-trans isomerase G | Q13427 | PPIG | Other | -2.009666667 | 0.562992957 | 26.267 | 25.2454 | 27.4665 | 21.6479 | 25.3931 | 25.9089 |
| Peptidyl-prolyl cis-trans isomerase H | O43447;O43447-2 | PPIH | Other | -1.094233333 | 0.882048444 | 25.7784 | 27.3468 | 26.7946 | 24.9003 | 25.8483 | 25.8885 |
| Peptidyl-prolyl cis-trans isomerase-like 1 | Q9Y3C6 | PPIL1 | Extracellular or secreted | -0.471566667 | 1.414290805 | 29.2291 | 29.1638 | 29.0601 | 28.8888 | 28.6555 | 28.494 |
| Peptidyl-prolyl cis-trans isomerase-like 4 | Q8WUA2 | PPIL4 | Other | -1.2459 | 0.606639937 | 24.6722 | 24.2464 | 24.412 | 21.6506 | 23.8874 | 24.0549 |
| Periplakin | O60437 | PPL | Extracellular or secreted | 0.7163 | 0.247283371 | 24.3776 | 25.5517 | 27.2735 | 25.011 | 27.6811 | 26.6596 |
| Protein phosphatase 1A | P35813;P35813-3;P35813-2 | PPM1A | Other | -0.368466667 | 0.558303833 | 26.4246 | 26.5558 | 26.8528 | 25.793 | 26.291 | 26.6438 |
| Protein phosphatase 1B | O75688;O75688-4;O75688-2 | PPM1B | Other | 0.018466667 | 0.021904706 | 23.9638 | 24.3299 | 23.7692 | 23.587 | 24.2548 | 24.2765 |
| Protein phosphatase 1F | P49593;P49593-2 | PPM1F | Other | -0.955233333 | 0.456457559 | 23.4229 | 24.0698 | 24.245 | 21.4878 | 24.1814 | 23.2028 |
| Protein phosphatase 1G | O15355 | PPM1G | Other | -0.168866667 | 0.253514338 | 29.0258 | 28.6708 | 29.4503 | 28.8706 | 28.6667 | 29.103 |
| Protein phosphatase methylesterase 1 | Q9Y570;Q9Y570-4;Q9Y570-2 | PPME1 | Other | -1.336433333 | 0.751095604 | 25.8998 | 26.6239 | 26.9059 | 23.8335 | 25.4794 | 26.1074 |
| Serine/threonine-protein phosphatase PP1-alpha catalyt | P62136;P62136-2;P62136-3 | PPP1CA | Extracellular or secreted | -0.902 | 0.984714757 | 30.7709 | 30.7042 | 30.6182 | 29.2135 | 29.8564 | 30.3174 |
| Serine/threonine-protein phosphatase PP1-beta catalytic | P62140 | PPP1CB | Extracellular or secreted | -0.743133333 | 1.209952519 | 27.188 | 27.2071 | 27.6791 | 26.1685 | 26.775 | 26.9013 |
| Serine/threonine-protein phosphatase PP1-gamma catal | P36873;P36873-2 | PPP1CC | Other | -0.837166667 | 1.460520592 | 27.1671 | 27.1187 | 27.3472 | 25.9994 | 26.5253 | 26.5968 |
| Serine/threonine-protein phosphatase 1 regulatory subu | Q96QC0 | PPP1R10 | Other | 0.247733333 | 0.100890681 | 21.0361 | 21.4037 | 21.6384 | 20.4752 | 21.1527 | 23.1935 |
| Protein phosphatase 1 regulatory subunit 11 | O60927 | PPP1R11 | Other | -0.691033333 | 0.293945028 | 22.0189 | 22.8283 | 24.251 | 22.6816 | 21.0077 | 23.3358 |
| Protein phosphatase 1 regulatory subunit 12A | O14974-5;O14974;O14974-4 | PPP1R12A | Other | -0.783133333 | 1.293484855 | 26.6917 | 26.0475 | 26.2257 | 25.5013 | 25.1983 | 25.9159 |
| RelA-associated inhibitor | Q8WUF5 | PPP1R13L | Extracellular or secreted | 0.9586 | 0.814628609 | 22.2431 | 21.1161 | 22.5582 | 22.8336 | 22.809 | 23.1506 |
| Protein phosphatase 1 regulatory subunit 14B | Q96C90 | PPP1R14B | Other | 0.1048 | 0.119915865 | 28.8464 | 28.568 | 28.2736 | 28.9313 | 28.9383 | 28.1328 |
| Protein phosphatase 1 regulatory subunit 14C | Q8TAE6 | PPP1R14C | Other | -0.486133333 | 0.647311261 | 24.7601 | 24.2132 | 24.3366 | 23.7662 | 24.5007 | 23.5846 |
| Protein phosphatase inhibitor 2;Protein phosphatase inh | P41236;Q6NXS1 | PPP1R2;PPP1R | Other | 0.97 | 0.747500978 | 25.4749 | 24.0035 | 25.4952 | 25.933 | 26.3102 | 25.6404 |
| Protein phosphatase 1 regulatory subunit 7 | Q15435;Q15435-3;Q15435-2 | PPP1R7 | Extracellular or secreted | 0.231566667 | 0.374271623 | 27.7288 | 27.8612 | 28.4491 | 28.0111 | 28.3668 | 28.3559 |
| Nuclear inhibitor of protein phosphatase 1;Activator of R | Q12972;Q12972-2 | PPP1R8 | Other | -0.600366667 | 0.208032672 | 23.4645 | 22.8093 | 25.5225 | 21.9206 | 23.5463 | 24.5283 |
| Neurabin-2 | Q96SB3 | PPP1R9B | Extracellular or secreted | -0.319533333 | 0.064644623 | 24.6598 | 20.9225 | 26.1572 | 22.7461 | 23.3392 | 24.6956 |
| Serine/threonine-protein phosphatase 2A catalytic subur | P67775;P67775-2 | PPP2CA | Extracellular or secreted | -0.026266667 | 0.025602066 | 25.9776 | 26.9731 | 26.0776 | 26.3392 | 26.1494 | 26.4609 |
| Serine/threonine-protein phosphatase 2A catalytic subur | P62714 | PPP2CB | Extracellular or secreted | -0.633366667 | 1.594882039 | 30.4265 | 30.437 | 30.1593 | 29.5316 | 29.9845 | 29.6066 |
| Serine/threonine-protein phosphatase 2A 65 kDa regulat | P30153 | PPP2R1A | Extracellular or secreted | -0.639766667 | 0.51963464 | 30.1563 | 31.0535 | 31.1573 | 29.371 | 30.2247 | 30.8521 |
| Serine/threonine-protein phosphatase 2A 65 kDa regulat | P30154-5;P30154;P30154-2; | PPP2R1B | Extracellular or secreted | -0.0989 | 0.033000705 | 22.4619 | 24.0772 | 25.2968 | 22.806 | 23.9255 | 24.8077 |
| Serine/threonine-protein phosphatase 2A 55 kDa regulat | P63151;P63151-2;Q66LE6 | PPP2R2A;PPP2 | Other | -0.535166667 | 0.557922674 | 27.6535 | 27.9536 | 28.3763 | 26.872 | 27.4143 | 28.0916 |
| Serine/threonine-protein phosphatase 2A activator | Q15257-3;Q15257-2;Q15257 | PPP2R4 | Extracellular or secreted | -0.622333333 | 0.807997268 | 28.1656 | 28.5121 | 28.4445 | 27.2611 | 28.2695 | 27.7246 |
| Serine/threonine-protein phosphatase 2A 56 kDa regulat | Q13362-2;Q13362-3;Q13362 | PPP2R5C | Other | -1.482966667 | 0.807815812 | 23.9202 | 23.2429 | 22.8108 | 23.2043 | 21.4299 | 20.8908 |
| Serine/threonine-protein phosphatase 2A 56 kDa regulat | Q14738;Q14738-2;Q14738-3 | PPP2R5D | Other | -0.538266667 | 0.361824719 | 26.1945 | 26.5382 | 27.3753 | 25.4008 | 25.9807 | 27.1117 |
| Serine/threonine-protein phosphatase 2B catalytic subur | Q08209-2;Q08209;Q08209-3 | PPP3CA | Extracellular or secreted | -0.0724 | 0.189477041 | 28.3835 | 27.9347 | 28.1578 | 28.0761 | 27.9955 | 28.1872 |
| Calcineurin subunit B type 1 | P63098 | PPP3R1 | Extracellular or secreted | 0.0004 | 0.000196884 | 25.5904 | 25.1747 | 24.6972 | 26.26 | 24.9586 | 24.2449 |
| Serine/threonine-protein phosphatase 4 catalytic subuni | P60510 | PPP4C | Extracellular or secreted | -0.166666667 | 0.251517625 | 28.4019 | 28.2036 | 27.6426 | 27.7062 | 28.1034 | 27.9385 |
| Serine/threonine-protein phosphatase 4 regulatory subu | Q8TF05-2;Q8TF05 | PPP4R1 | Other | 1.0715 | 0.496693958 | 28.1161 | 25.52 | 26.7951 | 28.6284 | 28.2091 | 26.8082 |
| Serine/threonine-protein phosphatase 4 regulatory subu | Q9NY27-3;Q9NY27;Q9NY27- | PPP4R2 | Other | 0.140433333 | 0.160594819 | 27.0839 | 26.2856 | 26.7975 | 27.2691 | 26.8494 | 26.4698 |
| Serine/threonine-protein phosphatase 5 | P53041 | PPP5C | Other | -0.752766667 | 0.845910467 | 28.2826 | 27.727 | 27.1196 | 27.1838 | 26.9347 | 26.7524 |
| Serine/threonine-protein phosphatase 6 catalytic subuni | O00743;O00743-3;O00743-2 | PPP6C | Other | -0.504666667 | 0.7353587 | 26.6922 | 26.8883 | 27.0544 | 25.9858 | 26.264 | 26.8711 |
| Serine/threonine-protein phosphatase 6 regulatory subu | Q5H9R7-3;Q5H9R7-4;Q5H9R | PPP6R3 | Extracellular or secreted | -0.316 | 0.246596617 | 27.1744 | 26.0446 | 27.2967 | 26.6831 | 25.926 | 26.9586 |
| Palmitoyl-protein thioesterase 1 | P50897;P50897-2 | PPT1 | Extracellular or secreted | -0.2888 | 0.823592604 | 29.3033 | 29.269 | 29.1262 | 28.6796 | 29.1022 | 29.0503 |
| Lysosomal thioesterase PPT2 | Q9UMR5;Q9UMR5-3;Q9UM | PPT2 | Extracellular or secreted | -0.998366667 | 1.214038959 | 26.4441 | 26.1108 | 25.4084 | 24.6613 | 24.9938 | 25.3131 |
| Peptidylprolyl isomerase domain and WD repeat-contain | Q96BP3;Q96BP3-2 | PPWD1 | Other | -0.296133333 | 0.221666864 | 24.5437 | 25.1554 | 26.1042 | 25.4148 | 24.7686 | 24.7315 |
| Polyglutamine-binding protein 1 | O60828-5;O60828-2;O60828 | PQBP1 | Other | 3.301966667 | 0.672202495 | 20.7295 | 21.0337 | 26.364 | 26.1263 | 25.8784 | 26.0284 |
| Melanoma antigen preferentially expressed in tumors | P78395 | PRAME | Extracellular or secreted | 1.281766667 | 0.34422509 | 23.7831 | 21.9558 | 20.5737 | 24.4549 | 24.7449 | 20.9581 |
| Protein regulator of cytokinesis 1 | O43663-3;O43663-2;O43663 | PRC1 | Extracellular or secreted | -1.498033333 | 1.114607064 | 23.7646 | 23.1486 | 24.1179 | 23.0603 | 22.1591 | 21.3176 |

| | | | | | | | | | | | |
|---|---|---|---|---|---|---|---|---|---|---|---|
| Proline-rich protein PRCC | Q92733 | PRCC | Other | 1.121333333 | 0.471122359 | 23.1572 | 20.8353 | 23.7938 | 23.9389 | 23.307 | 23.9044 |
| Lysosomal Pro-X carboxypeptidase | P42785;P42785-2 | PRCP | Extracellular or secreted | -0.030866667 | 0.035417702 | 29.1454 | 28.5675 | 28.5135 | 29.0689 | 28.7379 | 28.327 |
| Peroxiredoxin-1 | Q06830 | PRDX1 | Extracellular or secreted | -0.133366667 | 0.276955176 | 33.8396 | 34.0346 | 34.3868 | 33.7932 | 34.1461 | 33.9216 |
| Peroxiredoxin-2 | P32119 | PRDX2 | Extracellular or secreted | -0.121066667 | 0.146063657 | 32.0234 | 31.8843 | 32.3002 | 31.4782 | 32.422 | 31.9445 |
| Thioredoxin-dependent peroxide reductase, mitochondr | P30048-2;P30048 | PRDX3 | Extracellular or secreted | -0.8499 | 0.976187524 | 30.4682 | 30.5832 | 31.4081 | 29.5166 | 30.4821 | 29.9111 |
| Peroxiredoxin-4 | Q13162 | PRDX4 | Extracellular or secreted | -0.474366667 | 0.610363476 | 29.1597 | 28.6007 | 29.3551 | 28.9486 | 28.6844 | 28.0594 |
| Peroxiredoxin-5, mitochondrial | P30044-2;P30044;P30044-3; | PRDX5 | Extracellular or secreted | 1.130466667 | 0.908580589 | 28.6399 | 29.6381 | 29.868 | 29.6564 | 31.097 | 30.784 |
| Peroxiredoxin-6 | P30041 | PRDX6 | Extracellular or secreted | -0.855533333 | 1.470460399 | 31.2686 | 30.8558 | 31.5298 | 30.5079 | 30.2211 | 30.3586 |
| Prolyl endopeptidase | P48147 | PREP | Other | -0.310666667 | 0.964334961 | 29.2582 | 29.6532 | 29.3398 | 29.1193 | 29.1827 | 29.0172 |
| Prolyl endopeptidase-like | Q4J6C6-4;Q4J6C6-3;Q4J6C6; | PREPL | Other | 0.170233333 | 0.068447842 | 24.8975 | 25.1529 | 23.0437 | 25.5898 | 24.3162 | 23.6988 |
| Proteoglycan 4;Proteoglycan 4 C-terminal part | Q92954-5;Q92954-4;Q92954 | PRG4 | Extracellular or secreted | -1.816166667 | 1.812762014 | 23.6241 | 24.2423 | 23.1306 | 21.786 | 21.6012 | 22.1613 |
| Salivary acidic proline-rich phosphoprotein 1/2;Salivary a | P02810 | PRH1 | Extracellular or secreted | 1.8367 | 0.59791739 | 22.4136 | 21.6698 | 18.7765 | 23.6051 | 21.2704 | 23.4945 |
| cAMP-dependent protein kinase catalytic subunit alpha | P17612-2;P17612 | PRKACA | Extracellular or secreted | 0.872066667 | 0.394813385 | 24.0808 | 22.1043 | 24.8457 | 24.0323 | 24.5073 | 25.1074 |
| cAMP-dependent protein kinase type I-alpha regulatory | P10644;P10644-2 | PRKAR1A | Extracellular or secreted | -0.3616 | 0.302436575 | 25.9779 | 25.3272 | 25.3995 | 24.5882 | 25.0205 | 26.0111 |
| cAMP-dependent protein kinase type II-alpha regulatory | P13861-2;P13861 | PRKAR2A | Extracellular or secreted | -0.704766667 | 0.24654259 | 25.8353 | 26.0337 | 27.7023 | 24.0539 | 26.1319 | 27.2712 |
| Protein kinase C delta type;Protein kinase C delta type re | Q05655;Q05655-2 | PRKCD | Extracellular or secreted | -1.0121 | 1.348825997 | 21.7727 | 22.5852 | 21.804 | 21.3965 | 21.0956 | 20.6335 |
| Glucosidase 2 subunit beta | P14314-2;P14314 | PRKCSH | Extracellular or secreted | 0.1698 | 0.089665126 | 32.5406 | 31.6286 | 32.5565 | 33.5068 | 32.2175 | 31.5108 |
| DNA-dependent protein kinase catalytic subunit | P78527;P78527-2 | PRKDC | Other | -0.6662 | 0.18464249 | 28.5218 | 27.8942 | 31.3681 | 27.5961 | 27.8833 | 30.3061 |
| Interferon-inducible double-stranded RNA-dependent pr | O75569-3;O75569-2;O75569 | PRKRA | Other | 0.114866667 | 0.036579126 | 23.4911 | 21.1335 | 23.9881 | 21.9089 | 23.1558 | 23.8926 |
| Protein arginine N-methyltransferase 1 | Q99873-3;Q99873-2;Q99873 | PRMT1 | Other | 0.195933333 | 0.224504239 | 30.3476 | 30.2317 | 30.1164 | 29.903 | 30.4001 | 30.9804 |
| Protein arginine N-methyltransferase 3 | O60678-2;O60678 | PRMT3 | Other | -1.112833333 | 0.739092615 | 23.4701 | 22.7574 | 24.6298 | 21.7004 | 22.8951 | 22.9233 |
| Protein arginine N-methyltransferase 5;Protein arginine | O14744;O14744-2;O14744-5 | PRMT5 | Other | -0.4436 | 1.332781247 | 28.9084 | 29.2609 | 29.1814 | 28.5254 | 28.599 | 28.8955 |
| Protein arginine N-methyltransferase 6 | Q96LA8-2;Q96LA8 | PRMT6 | Other | -1.090033333 | 0.599955774 | 23.5328 | 21.3883 | 22.8096 | 21.0274 | 22.4939 | 20.9393 |
| Protein arginine N-methyltransferase 7 | Q9NVM4-4;Q9NVM4-3;Q9N | PRMT7 | Other | 0.2048 | 0.245390571 | 22.7827 | 22.9914 | 23.0143 | 22.7623 | 22.9139 | 23.7266 |
| Major prion protein | P04156-2;P04156 | PRNP | Extracellular or secreted | 2.1885 | 1.825298079 | 25.6854 | 25.79 | 25.127 | 28.0641 | 28.1544 | 26.9494 |
| Endothelial protein C receptor | Q9UNN8 | PROCR | Extracellular or secreted | 0.152933333 | 0.066406934 | 29.4652 | 29.0141 | 27.7933 | 29.7835 | 29.2499 | 27.698 |
| Prominin-2 | Q8N271-2;Q8N271;Q8N271- | PROM2 | Extracellular or secreted | -0.7953 | 0.215397182 | 24.0537 | 21.7008 | 24.1043 | 24.709 | 20.701 | 22.0629 |
| Proline synthase co-transcribed bacterial homolog prote | O94903 | PROSC | Extracellular or secreted | 0.658533333 | 0.139534358 | 25.9416 | 26.3836 | 21.3015 | 24.9388 | 25.5121 | 25.1514 |
| Pre-mRNA-processing factor 19 | Q9UMS4 | PRPF19 | Other | -1.595566667 | 1.575587129 | 30.8668 | 30.6823 | 31.4564 | 28.7161 | 29.5822 | 29.9205 |
| Pre-mRNA-splicing factor 38B | Q5VTL8-2;Q5VTL8 | PRPF38B | Other | 1.885633333 | 0.719657292 | 20.5334 | 20.458 | 23.3731 | 22.017 | 23.8232 | 24.1812 |
| U4/U6 small nuclear ribonucleoprotein Prp4 | O43172-2;O43172 | PRPF4 | Other | -1.292233333 | 0.394681988 | 24.9324 | 24.9922 | 25.7881 | 21.595 | 24.5212 | 25.7198 |
| Pre-mRNA-processing factor 40 homolog A | O75400-2;O75400-3;O75400 | PRPF40A | Other | -0.2693 | 0.122179127 | 26.4848 | 24.9615 | 27.4908 | 25.7212 | 25.9359 | 26.4721 |
| Serine/threonine-protein kinase PRP4 homolog | Q13523 | PRPF4B | Other | -0.7494 | 0.58017398 | 25.587 | 24.4635 | 26.1269 | 24.7893 | 24.4125 | 24.7274 |
| Pre-mRNA-processing-splicing factor 8 | Q6P2Q9 | PRPF8 | Other | -0.814466667 | 0.438381317 | 27.9048 | 26.84 | 29.1379 | 26.5671 | 26.959 | 27.9132 |
| Ribose-phosphate pyrophosphokinase 1 | P60891-2;P60891 | PRPS1 | Other | -1.011633333 | 0.522156153 | 24.8498 | 24.963 | 27.0541 | 24.2878 | 24.1631 | 25.3811 |
| Phosphoribosyl pyrophosphate synthase-associated prot | O60256-4;O60256-3;O60256 | PRPSAP2 | Other | 0.127633333 | 0.094128177 | 21.7386 | 21.1899 | 22.2441 | 21.317 | 21.6641 | 22.5744 |
| Protein PRRC1 | Q96M27;Q96M27-3;Q96M2 | PRRC1 | Other | -1.9415 | 0.798069254 | 24.8291 | 24.5853 | 24.1741 | 20.8295 | 23.1207 | 23.8138 |
| Protein PRRC2A | P48634;P48634-2;P48634-3; | PRRC2A | Extracellular or secreted | -1.7557 | 1.798853038 | 25.3951 | 25.2554 | 26.2574 | 24.4786 | 23.677 | 23.4852 |
| Protein PRRC2C | Q9Y520-3;Q9Y520-6;Q9Y520 | PRRC2C | Other | -0.106233333 | 0.138393767 | 27.1204 | 26.5679 | 27.3226 | 27.0525 | 27.0812 | 26.5585 |
| Brain-specific serine protease 4 | Q9GZN4 | PRSS22 | Extracellular or secreted | 1.694766667 | 1.625742316 | 26.0324 | 27.286 | 26.6264 | 27.9093 | 28.902 | 28.2178 |
| Trypsin-3 | P35030-5;P35030-3;P35030- | PRSS3 | Extracellular or secreted | 0.810166667 | 0.895736849 | 22.8382 | 21.9981 | 22.5973 | 23.0706 | 23.9365 | 22.857 |
| Prostasin;Prostasin light chain;Prostasin heavy chain | Q16651;Q16651-2 | PRSS8 | Extracellular or secreted | 1.0767 | 1.432049688 | 26.5598 | 26.6641 | 26.5013 | 27.6568 | 28.0429 | 27.2556 |
| Prosaposin;Saposin-A;Saposin-B-Val;Saposin-B;Saposin-C | P07602;P07602-2;P07602-3 | PSAP | Extracellular or secreted | 0.275066667 | 0.891682152 | 30.9028 | 30.9899 | 30.9647 | 31.2528 | 31.4088 | 31.021 |
| Phosphoserine aminotransferase | Q9Y617;Q9Y617-2 | PSAT1 | Extracellular or secreted | -0.612533333 | 0.495388912 | 32.8092 | 32.3668 | 31.6427 | 32.2182 | 31.9139 | 30.849 |
| PC4 and SFRS1-interacting protein | O75475 | PSIP1 | Other | -0.6119 | 0.810786987 | 28.9406 | 29.2359 | 29.9053 | 28.8561 | 28.5428 | 28.8472 |
| Proteasome subunit alpha type-1 | P25786;P25786-2 | PSMA1 | Extracellular or secreted | 0.253366667 | 0.162795483 | 33.4067 | 32.75 | 32.3074 | 34.005 | 32.8329 | 32.3863 |
| Proteasome subunit alpha type-2 | P25787 | PSMA2 | Extracellular or secreted | -0.064433333 | 0.040983153 | 32.6335 | 32.1264 | 31.3339 | 32.6544 | 31.8972 | 31.3489 |
| Proteasome subunit alpha type-3 | P25788-2;P25788 | PSMA3 | Extracellular or secreted | 0.199766667 | 0.109906177 | 32.558 | 31.612 | 31.4242 | 33.0956 | 31.8661 | 31.2318 |
| Proteasome subunit alpha type-4 | P25789;P25789-2 | PSMA4 | Extracellular or secreted | -0.061433333 | 0.040232406 | 32.7779 | 32.3102 | 31.801 | 32.9632 | 32.2667 | 31.4749 |
| Proteasome subunit alpha type-5 | P28066;P28066-2 | PSMA5 | Extracellular or secreted | 0.485866667 | 0.368667014 | 33.2784 | 32.8393 | 31.9164 | 33.8004 | 33.1975 | 32.4938 |
| Proteasome subunit alpha type-6 | P60900;P60900-2;P60900-3 | PSMA6 | Extracellular or secreted | 0.261033333 | 0.176096867 | 33.1956 | 32.5304 | 32.1968 | 33.7877 | 32.7326 | 32.1856 |
| Proteasome subunit alpha type-7 | O14818;O14818-4 | PSMA7 | Extracellular or secreted | -0.172533333 | 0.11451426 | 33.5715 | 32.8281 | 32.608 | 33.5771 | 32.9073 | 32.0056 |
| Proteasome subunit alpha type-7 | O14818-2 | PSMA7 | Extracellular or secreted | -1.212233333 | 0.312554218 | 24.9841 | 24.3822 | 22.7862 | 25.5762 | 21.2371 | 21.7025 |
| Proteasome subunit beta type-1 | P20618 | PSMB1 | Extracellular or secreted | -0.059466667 | 0.03050019 | 33.1419 | 32.4586 | 31.9474 | 33.4675 | 32.3227 | 31.5793 |
| Proteasome subunit beta type-10 | P40306 | PSMB10 | Other | -1.126833333 | 0.853502437 | 27.4106 | 26.7244 | 26.3835 | 26.328 | 26.083 | 24.727 |
| Proteasome subunit beta type-2 | P49721 | PSMB2 | Extracellular or secreted | -0.192166667 | 0.143155566 | 32.4249 | 31.9093 | 31.2814 | 32.1469 | 31.9468 | 30.9454 |
| Proteasome subunit beta type-3 | P49720 | PSMB3 | Extracellular or secreted | -0.9102 | 0.531869405 | 32.6185 | 31.7173 | 30.6062 | 31.178 | 31.2393 | 29.7941 |
| Proteasome subunit beta type-4 | P28070 | PSMB4 | Extracellular or secreted | 0.113833333 | 0.075502758 | 32.4971 | 31.8658 | 31.2584 | 32.6196 | 32.0696 | 31.2736 |
| Proteasome subunit beta type-5 | P28074;P28074-3;P28074-2 | PSMB5 | Extracellular or secreted | -0.1954 | 0.130794617 | 32.8755 | 32.2826 | 31.6314 | 32.7753 | 32.0876 | 31.3404 |
| Proteasome subunit beta type-6 | P28072 | PSMB6 | Extracellular or secreted | 0.259266667 | 0.154243116 | 32.2141 | 31.5157 | 30.7421 | 32.4405 | 31.9371 | 30.8721 |
| Proteasome subunit beta type-7 | Q99436 | PSMB7 | Extracellular or secreted | 0.087333333 | 0.057999534 | 32.3455 | 31.964 | 31.3231 | 32.7482 | 31.848 | 31.2984 |
| Proteasome subunit beta type-8 | P28062-2;P28062 | PSMB8 | Extracellular or secreted | -0.296733333 | 0.178665887 | 31.6261 | 30.5565 | 30.1145 | 31.193 | 30.5516 | 29.6623 |
| Proteasome subunit beta type-9 | P28065-2;P28065 | PSMB9 | Extracellular or secreted | -0.534566667 | 0.237126588 | 28.6191 | 28.4116 | 26.6975 | 28.3012 | 27.6885 | 26.1348 |
| 26S protease regulatory subunit 4 | P62191;P62191-2 | PSMC1 | Other | -0.1464 | 0.077268777 | 28.4099 | 27.9105 | 29.7023 | 27.9879 | 28.3017 | 29.2939 |
| 26S protease regulatory subunit 7 | P35998;P35998-2 | PSMC2 | Other | -0.2063 | 0.098634432 | 29.3629 | 29.0282 | 30.7968 | 28.9975 | 29.0136 | 30.5579 |
| 26S protease regulatory subunit 6A | P17980 | PSMC3 | Other | -0.359333333 | 0.176151267 | 28.575 | 28.749 | 30.1679 | 27.9294 | 28.5692 | 29.9153 |
| 26S protease regulatory subunit 6B | P43686;P43686-2 | PSMC4 | Other | -0.729833333 | 0.420204209 | 28.6234 | 28.295 | 29.6724 | 27.2427 | 27.8847 | 29.2739 |
| 26S protease regulatory subunit 8 | P62195;P62195-2 | PSMC5 | Extracellular or secreted | -0.315733333 | 0.159348085 | 28.9896 | 28.4477 | 30.2849 | 28.2689 | 28.588 | 29.9181 |
| 26S protease regulatory subunit 10B | P62333 | PSMC6 | Extracellular or secreted | -0.3382 | 0.178504267 | 29.4271 | 28.8531 | 30.5368 | 28.6151 | 28.885 | 30.3023 |
| 26S proteasome non-ATPase regulatory subunit 1 | Q99460;Q99460-2 | PSMD1 | Extracellular or secreted | -0.480533333 | 0.266629117 | 29.4797 | 29.1194 | 30.7254 | 28.6567 | 28.8786 | 30.3476 |
| 26S proteasome non-ATPase regulatory subunit 10 | O75832 | PSMD10 | Other | 1.004866667 | 0.815559707 | 24.7772 | 26.1151 | 26.2443 | 26.4361 | 26.5659 | 27.1492 |
| 26S proteasome non-ATPase regulatory subunit 11 | O00231;O00231-2 | PSMD11 | Extracellular or secreted | -0.2319 | 0.128036561 | 28.9984 | 28.4104 | 30.0291 | 28.3333 | 28.572 | 29.8369 |
| 26S proteasome non-ATPase regulatory subunit 12 | O00232;O00232-2 | PSMD12 | Extracellular or secreted | -0.6248 | 0.442719654 | 29.0306 | 28.6097 | 29.9238 | 27.8598 | 28.3936 | 29.4363 |
| 26S proteasome non-ATPase regulatory subunit 13 | Q9UNM6;Q9UNM6-2 | PSMD13 | Extracellular or secreted | -0.555366667 | 0.256180442 | 28.9161 | 29.0752 | 30.33 | 27.6259 | 28.9115 | 30.1178 |
| 26S proteasome non-ATPase regulatory subunit 14 | O00487 | PSMD14 | Extracellular or secreted | -0.6034 | 0.329692387 | 27.571 | 27.5158 | 28.8469 | 26.3781 | 27.276 | 28.4694 |
| 26S proteasome non-ATPase regulatory subunit 2 | Q13200;Q13200-3;Q13200-2 | PSMD2 | Extracellular or secreted | -0.116966667 | 0.067616479 | 30.0753 | 29.7121 | 31.1041 | 29.747 | 29.7402 | 31.0534 |
| 26S proteasome non-ATPase regulatory subunit 3 | O43242;O43242-2 | PSMD3 | Extracellular or secreted | -0.147966667 | 0.076404633 | 28.873 | 28.6463 | 30.1414 | 28.4612 | 28.698 | 30.0576 |
| 26S proteasome non-ATPase regulatory subunit 4 | P55036;P55036-2 | PSMD4 | Other | 1.021366667 | 0.700740063 | 26.285 | 26.7838 | 28.1415 | 27.5917 | 28.2237 | 28.459 |
| 26S proteasome non-ATPase regulatory subunit 5 | Q16401-2;Q16401 | PSMD5 | Other | -0.259666667 | 0.151410051 | 27.8392 | 28.0318 | 28.6085 | 26.7972 | 28.1901 | 27.7132 |
| 26S proteasome non-ATPase regulatory subunit 6 | Q15008;Q15008-4;Q15008-3 | PSMD6 | Extracellular or secreted | -0.5948 | 0.37188161 | 28.7714 | 28.588 | 29.8759 | 27.7694 | 28.1647 | 29.5168 |
| 26S proteasome non-ATPase regulatory subunit 7 | P51665 | PSMD7 | Extracellular or secreted | -1.390966667 | 0.532372308 | 27.7655 | 26.4823 | 28.8247 | 24.5787 | 26.6312 | 27.6897 |
| 26S proteasome non-ATPase regulatory subunit 8 | P48556 | PSMD8 | Extracellular or secreted | -1.2816 | 0.292502819 | 27.5974 | 26.6524 | 29.1526 | 23.624 | 27.039 | 28.8946 |
| 26S proteasome non-ATPase regulatory subunit 9 | O00233;O00233-2 | PSMD9 | Other | 0.036366667 | 0.073179563 | 27.288 | 27.3325 | 27.6088 | 27.169 | 27.6306 | 27.5388 |
| Proteasome activator complex subunit 1 | Q06323;Q06323-3;Q06323-2 | PSME1 | Extracellular or secreted | -0.682766667 | 0.665011054 | 29.9517 | 29.5059 | 29.6997 | 28.2574 | 29.3758 | 29.4758 |
| Proteasome activator complex subunit 2 | Q9UL46 | PSME2 | Extracellular or secreted | -1.010166667 | 1.257021457 | 29.7296 | 29.5681 | 29.7165 | 28.1439 | 28.8748 | 28.965 |
| Proteasome activator complex subunit 3 | P61289;P61289-3;P61289-2 | PSME3 | Other | -0.2753 | 0.354121536 | 28.4718 | 28.366 | 28.626 | 27.9234 | 27.9259 | 28.7886 |
| Proteasome activator complex subunit 4 | Q14997 | PSME4 | Other | -1.6198 | 0.396195903 | 24.7527 | 25.2031 | 24.8523 | 20.2619 | 24.6042 | 25.0826 |
| Proteasome inhibitor PI31 subunit | Q92530 | PSMF1 | Other | -0.1244 | 0.114449875 | 24.7599 | 24.5621 | 24.7184 | 23.8915 | 24.6177 | 25.158 |
| Proteasome assembly chaperone 1 | O95456-2;O95456 | PSMG1 | Other | -0.732566667 | 1.21223985 | 26.5581 | 26.9986 | 26.8534 | 25.6246 | 26.2569 | 26.3309 |
| Proteasome assembly chaperone 2 | Q969U7-2;Q969U7 | PSMG2 | Other | -0.9433 | 1.371599416 | 26.7153 | 27.0329 | 27.0221 | 25.7597 | 25.722 | 26.4587 |
| Proteasome assembly chaperone 3 | Q9BT73 | PSMG3 | Other | 0.301366667 | 0.292113591 | 24.5781 | 25.848 | 25.4546 | 25.7329 | 25.3948 | 25.6571 |
| Proteasome assembly chaperone 4 | Q5JS54;Q5JS54-2;Q5JS54-3 | PSMG4 | Other | 0.0763 | 0.119716562 | 25.5075 | 25.5991 | 24.9211 | 25.3077 | 25.5279 | 25.421 |
| Paraspeckle component 1 | Q8WXF1;Q8WXF1-2 | PSPC1 | Other | 0.192 | 0.070448736 | 27.5937 | 27.7095 | 29.2922 | 26.9395 | 28.6564 | 29.5755 |
| Phosphoserine phosphatase | P78330 | PSPH | Other | 0.107266667 | 0.14553225 | 28.0923 | 27.5754 | 27.4732 | 28.1543 | 27.8316 | 27.4768 |
| Polypyrimidine tract-binding protein 1 | P26599;P26599-2;P26599-3 | PTBP1 | Other | -1.2057 | 0.735895887 | 30.4698 | 30.6145 | 31.3662 | 28.5149 | 29.6415 | 30.677 |
| Phosphotriesterase-related protein | Q96BW5-2;Q96BW5 | PTER | Extracellular or secreted | -1.304933333 | 0.780396694 | 26.8747 | 26.5631 | 26.273 | 26.3453 | 25.279 | 24.1717 |
| Prostaglandin E synthase 3 | Q15185-4;Q15185;Q15185-3 | PTGES3 | Extracellular or secreted | 0.685566667 | 0.991907584 | 26.714 | 27.0995 | 27.5915 | 28.196 | 27.5956 | 27.6701 |
| Prostaglandin F2 receptor negative regulator | Q9P2B2 | PTGFRN | Other | -0.7172 | 1.058750637 | 31.0912 | 30.7881 | 30.282 | 30.3906 | 29.9584 | 29.6607 |
| Prostaglandin reductase 1 | Q14914;Q14914-2 | PTGR1 | Extracellular or secreted | -0.106066667 | 0.273015563 | 28.1098 | 28.222 | 27.9838 | 28.1634 | 28.1013 | 27.7327 |
| Prostaglandin G/H synthase 2 | P35354 | PTGS2 | Extracellular or secreted | 2.5755 | 1.955531373 | 20.7439 | 19.7499 | 20.7585 | 23.1768 | 22.9712 | 22.8308 |
| Focal adhesion kinase 1 | Q05397-7;Q05397;Q05397-5 | PTK2 | Extracellular or secreted | -0.375566667 | 0.661366255 | 25.6167 | 25.3417 | 24.9597 | 25.2015 | 24.611 | 24.9789 |
| Inactive tyrosine-protein kinase 7 | Q13308;Q13308-6;Q13308-4 | PTK7 | Extracellular or secreted | 0.733066667 | 1.53060744 | 30.5134 | 30.0948 | 30.0688 | 31.1053 | 31.1416 | 30.6293 |
| Prothymosin alpha;Prothymosin alpha, N-terminally proc | P06454-2;P06454 | PTMA | Extracellular or secreted | 0.5529 | 0.181895844 | 33.2539 | 31.9867 | 32.7735 | 35.307 | 32.2642 | 32.1016 |
| Parathymosin | P20962 | PTMS | Other | 4.928866667 | 0.974682478 | 25.3943 | 25.522 | 24.976 | 33.0235 | 26.9798 | 30.6756 |
| Protein tyrosine phosphatase type IVA 2;Protein tyrosine | Q12974-3;Q12974;Q93096;C | PTP4A2;PTP4A | Extracellular or secreted | -0.585566667 | 0.233082332 | 23.331 | 23.9333 | 23.7001 | 21.3258 | 23.5477 | 24.3342 |
| Tyrosine-protein phosphatase non-receptor type 1 | P18031 | PTPN1 | Extracellular or secreted | 0.4467 | 0.262409811 | 23.9239 | 24.157 | 25.8164 | 24.638 | 25.0423 | 25.5571 |
| Tyrosine-protein phosphatase non-receptor type 11 | Q06124-2;Q06124;Q06124-3 | PTPN11 | Other | -1.695133333 | 0.648142527 | 27.153 | 27.172 | 27.8386 | 23.7366 | 26.4181 | 26.9235 |
| Tyrosine-protein phosphatase non-receptor type 12 | Q05209-2;Q05209-3;Q05209 | PTPN12 | Extracellular or secreted | 0.744566667 | 0.912027614 | 25.8619 | 25.6136 | 26.1207 | 26.373 | 26.2335 | 27.2234 |
| Tyrosine-protein phosphatase non-receptor type 6 | P29350-2;P29350;P29350-3; | PTPN6 | Extracellular or secreted | -0.271266667 | 0.145966368 | 26.1886 | 25.9123 | 26.7257 | 24.9694 | 25.9299 | 27.1135 |
| Receptor-type tyrosine-protein phosphatase alpha | P18433-6;P18433 | PTPRA | Extracellular or secreted | -0.371966667 | 0.111498532 | 23.7011 | 22.8994 | 23.7442 | 24.4447 | 23.9147 | 20.8694 |
| Receptor-type tyrosine-protein phosphatase F | P10586-2;P10586 | PTPRF | Extracellular or secreted | -1.385366667 | 1.811993539 | 32.3271 | 31.711 | 31.7651 | 30.6441 | 30.4949 | 30.5081 |
| Receptor-type tyrosine-protein phosphatase gamma | P23470-2;P23470 | PTPRG | Extracellular or secreted | 1.354566667 | 1.153138651 | 26.5853 | 26.6756 | 25.4855 | 28.3122 | 27.563 | 26.9349 |
| Receptor-type tyrosine-protein phosphatase kappa | Q15262;Q15262-2;Q15262-3 | PTPRK | Extracellular or secreted | 1.170133333 | 1.180166999 | 27.7655 | 27.8748 | 26.7174 | 28.9614 | 28.7331 | 28.1736 |
| Receptor-type tyrosine-protein phosphatase S | Q13332-6;Q13332-2;Q13332 | PTPRS | Extracellular or secreted | -0.6547 | 0.993527279 | 28.8487 | 28.2898 | 29.041 | 27.7579 | 27.9846 | 28.4729 |
| Receptor-type tyrosine-protein phosphatase U | Q92729-3;Q92729-2;Q92729 | PTPRU | Extracellular or secreted | -0.283733333 | 0.156038605 | 25.8527 | 24.4901 | 23.7874 | 24.9271 | 24.1174 | 24.2345 |
| Polymerase I and transcript release factor | Q6NZI2;Q6NZI2-2 | PTRF | Extracellular or secreted | -0.306166667 | 0.121258829 | 25.9428 | 26.4963 | 28.6173 | 26.3077 | 26.3803 | 27.4499 |
| Putative peptidyl-tRNA hydrolase PTRHD1 | Q6GMV3 | PTRHD1 | Extracellular or secreted | 2.6539 | 1.559933299 | 22.5323 | 20.8076 | 21.5529 | 24.0768 | 24.2633 | 24.5144 |
| 6-pyruvoyl tetrahydrobiopterin synthase | Q03393 | PTS | Other | 0.482866667 | 0.291372332 | 22.5954 | 21.0585 | 21.5637 | 22.712 | 21.2335 | 22.7207 |
| Pituitary tumor-transforming gene 1 protein-interacting | P53801 | PTTG1IP | Extracellular or secreted | 2.714666667 | 0.925602073 | 19.4467 | 22.9896 | 22.157 | 24.6184 | 24.4067 | 23.7122 |

| | | | | | | | | | | | |
|---|---|---|---|---|---|---|---|---|---|---|---|
| Pentraxin-related protein PTX3 | P26022 | PTX3 | Extracellular or secreted | 3.246033333 | 1.086312598 | 28.318 | 27.8558 | 24.9037 | 30.9309 | 30.1042 | 29.7805 |
| Poly(U)-binding-splicing factor PUF60 | Q9UHX1-2;Q9UHX1-4;Q9UH | PUF60 | Other | -0.937133333 | 0.69645757 | 29.3618 | 29.1306 | 29.901 | 27.8454 | 28.2056 | 29.531 |
| Transcriptional activator protein Pur-alpha | Q00577 | PURA | Other | -0.113966667 | 0.460469177 | 26.3548 | 26.2642 | 26.4168 | 26.0909 | 26.2001 | 26.4029 |
| Transcriptional activator protein Pur-beta | Q96QR8 | PURB | Other | -0.749366667 | 2.023306098 | 27.3588 | 27.0811 | 27.4217 | 26.6882 | 26.6224 | 26.3029 |
| tRNA pseudouridine synthase A, mitochondrial | Q9Y606-2;Q9Y606 | PUS1 | Other | -0.4406 | 0.427721406 | 26.1781 | 26.2324 | 27.3058 | 25.5645 | 25.8423 | 25.9877 |
| Pseudouridylate synthase 7 homolog | Q96PZ0;Q96PZ0-2 | PUS7 | Other | -0.818766667 | 2.368916659 | 26.9947 | 27.2961 | 27.1841 | 26.3242 | 26.2767 | 26.4177 |
| Poliovirus receptor | P15151-3;P15151-2;P15151; | PVR | Extracellular or secreted | 0.936233333 | 1.240574055 | 28.4517 | 28.4605 | 27.6955 | 29.5727 | 29.1281 | 28.7156 |
| Nectin-1 | Q15223-3;Q15223-2;Q1522 | PVRL1 | Extracellular or secreted | 0.0397 | 0.05472949 | 25.8589 | 25.1548 | 25.2808 | 25.6851 | 25.2917 | 25.4368 |
| Nectin-2 | Q92692-2;Q92692 | PVRL2 | Extracellular or secreted | -0.3244 | 0.442157056 | 26.6642 | 25.7562 | 25.9757 | 25.7731 | 26.01 | 25.6398 |
| Nectin-4;Processed poliovirus receptor-related protein 4 | Q96NY8 | PVRL4 | Extracellular or secreted | 1.0579 | 1.649062393 | 24.86 | 24.3209 | 24.3408 | 25.5857 | 25.62 | 25.4897 |
| Periodic tryptophan protein 1 homolog | Q13610 | PWP1 | Other | -0.267433333 | 1.070759805 | 27.1243 | 27.0398 | 27.0247 | 26.9761 | 26.7001 | 26.7103 |
| Periodic tryptophan protein 2 homolog | Q15269 | PWP2 | Other | -0.104633333 | 0.022828591 | 21.7483 | 19.5739 | 24.5009 | 22.3013 | 21.9263 | 21.2816 |
| Peroxidasin homolog | Q92626 | PXDN | Extracellular or secreted | 1.792666667 | 0.706846392 | 29.8774 | 31.7594 | 28.4241 | 30.9562 | 32.0085 | 32.4742 |
| Paxillin | P49023-4;P49023-2;P49023; | PXN | Extracellular or secreted | -0.5771 | 0.26471323 | 20.8717 | 23.2138 | 23.2764 | 22.027 | 21.5813 | 22.0223 |
| Apoptosis-associated speck-like protein containing a CAR | Q9ULZ3;Q9ULZ3-3;Q9ULZ3-2 | PYCARD | Other | -1.0115 | 0.960378929 | 27.4543 | 27.6708 | 26.4291 | 25.6458 | 26.6752 | 26.1987 |
| Pyrroline-5-carboxylate reductase 1, mitochondrial | P32322;P32322-3;P32322-2 | PYCR1 | Other | -2.7031 | 0.844902851 | 24.4976 | 24.1178 | 27.5585 | 22.6879 | 20.9443 | 24.4324 |
| Pyrroline-5-carboxylate reductase 3 | Q53H96-2;Q53H96 | PYCRL | Other | 0.2211 | 0.167977093 | 24.2049 | 24.4854 | 25.7069 | 24.7042 | 25.132 | 25.3103 |
| Glycogen phosphorylase, brain form | P11216 | PYGB | Extracellular or secreted | 0.5499 | 0.559626438 | 30.4676 | 30.8775 | 31.021 | 30.6733 | 31.3876 | 31.9549 |
| Glycogen phosphorylase, liver form | P06737-2;P06737 | PYGL | Extracellular or secreted | -0.7325 | 3.790926146 | 32.4302 | 32.55 | 32.5051 | 31.7273 | 31.8403 | 31.7202 |
| Glutamine--tRNA ligase | P47897;P47897-2 | QARS | Other | -0.5189 | 0.316208725 | 28.2276 | 28.4191 | 29.6258 | 27.4583 | 28.0667 | 29.1908 |
| Dihydropteridine reductase | P09417;P09417-2 | QDPR | Extracellular or secreted | -0.176033333 | 0.248966905 | 28.5371 | 28.3359 | 28.1074 | 28.4241 | 28.3598 | 27.6684 |
| Glutamine-rich protein 1 | Q2TAL8 | QRICH1 | Other | -1.016633333 | 0.77397596 | 21.4758 | 21.6817 | 22.8656 | 21.1237 | 20.2018 | 21.6477 |
| Sulfhydryl oxidase 1 | O00391;O00391-2 | QSOX1 | Extracellular or secreted | 2.1229 | 0.979396288 | 31.3102 | 32.0252 | 29.5017 | 33.0282 | 33.1789 | 32.9987 |
| Sulfhydryl oxidase 2 | Q6ZRP7 | QSOX2 | Extracellular or secreted | -0.487866667 | 1.169844106 | 23.6205 | 24.0067 | 23.55 | 23.0882 | 23.5074 | 23.118 |
| Queuine tRNA-ribosyltransferase subunit QTRTD1 | Q9H974-2;Q9H974-3;Q9H97 | QTRTD1 | Other | 0.946333333 | 0.839920216 | 20.7593 | 21.5887 | 21.9734 | 22.9454 | 22.5545 | 21.6605 |
| Ras-related protein Rab-10 | P61026 | RAB10 | Extracellular or secreted | 1.191233333 | 0.631919882 | 25.66 | 24.1788 | 26.2302 | 26.7922 | 27.4237 | 25.4268 |
| Ras-related protein Rab-11B;Ras-related protein Rab-11A | Q15907;Q15907-2;P62491-2 | RAB11B;RAB11 | Extracellular or secreted | -0.346233333 | 0.532428625 | 29.3368 | 29.4408 | 29.7354 | 28.693 | 29.5133 | 29.268 |
| Rab11 family-interacting protein 1 | Q6WKZ4;Q6WKZ4-5;Q6WKZ | RAB11FIP1 | Other | 5.049466667 | 1.647151361 | 23.1016 | 21.5781 | 25.2136 | 28.0248 | 27.5029 | 29.514 |
| Ras-related protein Rab-13 | P51153 | RAB13 | Extracellular or secreted | 1.087233333 | 1.409669497 | 25.2553 | 24.8207 | 24.3292 | 25.4704 | 26.2882 | 25.9083 |
| Ras-related protein Rab-14 | P61106 | RAB14 | Extracellular or secreted | 0.471333333 | 0.663076328 | 27.3843 | 27.2197 | 28.0389 | 27.7205 | 28.3933 | 27.9431 |
| Ras-related protein Rab-18 | Q9NP72;Q9NP72-2;Q9NP72- | RAB18 | Extracellular or secreted | 0.659633333 | 0.359340681 | 23.0409 | 21.3992 | 22.9716 | 22.1697 | 24.0595 | 23.1614 |
| Ras-related protein Rab-1A | P62820;P62820-2;P62820-3 | RAB1A | Extracellular or secreted | 0.109933333 | 0.133398056 | 28.9261 | 28.6435 | 29.447 | 28.7606 | 29.4117 | 29.1741 |
| Ras-related protein Rab-1B;Putative Ras-related protein | Q9H0U4;Q92928 | RAB1B;RAB1C | Extracellular or secreted | 0.5197 | 0.503050788 | 26.4447 | 26.4383 | 27.6137 | 27.2022 | 27.5995 | 27.2541 |
| Ras-related protein Rab-21 | Q9UL25 | RAB21 | Extracellular or secreted | 0.464466667 | 0.503127408 | 24.3815 | 24.6782 | 24.9788 | 24.7332 | 24.8742 | 25.8245 |
| Ras-related protein Rab-22A | Q9UL26 | RAB22A | Extracellular or secreted | 2.158966667 | 0.942291346 | 22.9392 | 21.2144 | 24.1035 | 24.4271 | 25.1821 | 25.1248 |
| Ras-related protein Rab-27B | O00194 | RAB27B | Extracellular or secreted | 0.135033333 | 0.114602058 | 21.6581 | 21.3209 | 21.1283 | 20.8774 | 22.1995 | 21.4355 |
| Ras-related protein Rab-2A | P61019;P61019-2 | RAB2A | Extracellular or secreted | -0.2615 | 0.21617535 | 26.5839 | 26.6706 | 27.896 | 26.5082 | 26.9621 | 26.8957 |
| Ras-related protein Rab-34 | Q9BZG1-4;Q9BZG1-2;Q9BZG | RAB34 | Extracellular or secreted | 1.051133333 | 0.408089471 | 19.8431 | 22.2568 | 23.041 | 22.6152 | 22.3532 | 23.3259 |
| Ras-related protein Rab-35 | Q15286;Q15286-2 | RAB35 | Extracellular or secreted | 1.5936 | 2.893661615 | 24.69 | 24.2289 | 24.6325 | 25.8843 | 26.3339 | 26.114 |
| Ras-related protein Rab-3D | O95716 | RAB3D | Extracellular or secreted | 1.494266667 | 1.643795218 | 22.2286 | 21.3605 | 21.2032 | 22.6725 | 23.3864 | 23.2162 |
| Rab3 GTPase-activating protein catalytic subunit | Q15042;Q15042-3;Q15042-4 | RAB3GAP1 | Extracellular or secreted | -1.8807 | 0.630796378 | 23.0376 | 24.8943 | 26.0943 | 22.2086 | 21.428 | 24.7475 |
| Rab3 GTPase-activating protein non-catalytic subunit | Q9H2M9 | RAB3GAP2 | Extracellular or secreted | -1.156166667 | 0.352385965 | 22.5645 | 23.3275 | 25.705 | 20.7601 | 23.4772 | 23.8912 |
| Ras-related protein Rab-5C | P51148;P51148-2 | RAB5C | Extracellular or secreted | 0.682566667 | 1.152810844 | 26.268 | 26.6364 | 27.0335 | 27.0821 | 27.3639 | 27.5396 |
| Ras-related protein Rab-6A | P20340-2;P20340;P20340-4 | RAB6A | Extracellular or secreted | -0.290366667 | 0.251944449 | 27.9733 | 28.1484 | 29.2543 | 27.8486 | 28.476 | 28.1803 |
| Ras-related protein Rab-7a | P51149 | RAB7A | Extracellular or secreted | 1.0095 | 1.141530102 | 26.51 | 26.652 | 27.5054 | 27.9893 | 28 | 27.7066 |
| Ras-related protein Rab-8A | P61006;P61006-2 | RAB8A | Extracellular or secreted | 0.447133333 | 0.533204156 | 24.3444 | 24.556 | 24.73 | 24.7149 | 25.6242 | 24.6327 |
| Rab GTPase-binding effector protein 1 | Q15276;Q15276-2 | RABEP1 | Other | -0.353866667 | 0.418757334 | 22.217 | 22.9536 | 22.5492 | 21.7452 | 22.733 | 22.18 |
| Rab9 effector protein with kelch motifs | Q7Z6M1;Q7Z6M1-2 | RABEPK | Extracellular or secreted | -1.030033333 | 1.27805325 | 25.4463 | 24.7718 | 24.5685 | 23.892 | 24.0195 | 23.785 |
| Rab GTPase-activating protein 1 | Q9Y3P9-2;Q9Y3P9 | RABGAP1 | Other | 0.3141 | 0.432487365 | 22.7318 | 22.6025 | 22.7776 | 22.9049 | 22.6114 | 23.5379 |
| Geranylgeranyl transferase type-2 subunit alpha | Q92696 | RABGGTA | Other | -0.080433333 | 0.102277918 | 27.0793 | 27.3675 | 27.5392 | 26.7659 | 27.4387 | 27.5401 |
| Geranylgeranyl transferase type-2 subunit beta | P53611 | RABGGTB | Other | -0.054233333 | 0.026495054 | 26.3988 | 26.9124 | 24.8269 | 25.6119 | 26.317 | 26.0465 |
| Guanine nucleotide exchange factor MSS4 | P47224 | RABIF | Other | 0.9247 | 1.163980237 | 22.2975 | 22.8897 | 22.0011 | 23.2554 | 23.3218 | 23.3852 |
| Rab-like protein 6 | Q3YEC7;Q3YEC7-2;Q3YEC7-4 | RABL6 | Other | -0.918233333 | 0.528580946 | 24.6658 | 22.8431 | 24.8481 | 22.6395 | 23.0893 | 23.8735 |
| Ras-related C3 botulinum toxin substrate 1;Ras-related C | P63000;P63000-2;P60763 | RAC1;RAC3 | Extracellular or secreted | 1.2694 | 1.112468855 | 27.6075 | 28.7296 | 27.3964 | 28.9361 | 29.4745 | 29.1311 |
| Ras-related C3 botulinum toxin substrate 2 | P15153 | RAC2 | Extracellular or secreted | 2.318 | 2.231882856 | 22.3142 | 22.1202 | 21.9267 | 24.2185 | 24.9658 | 24.1308 |
| Rac GTPase-activating protein 1 | Q9H0H5 | RACGAP1 | Extracellular or secreted | 0.9434 | 0.346559841 | 22.5126 | 22.0849 | 21.0186 | 22.7275 | 24.5582 | 21.1606 |
| Double-strand-break repair protein rad21 homolog | O60216 | RAD21 | Other | -1.968266667 | 0.821299899 | 26.8489 | 27.2379 | 28.9958 | 24.3854 | 25.4371 | 27.3553 |
| UV excision repair protein RAD23 homolog A | P54725;P54725-2 | RAD23A | Other | 3.330066667 | 1.337657618 | 21.241 | 19.9106 | 22.4192 | 25.1418 | 22.7699 | 25.6493 |
| UV excision repair protein RAD23 homolog A | P54725-3 | RAD23A | Other | -0.651833333 | 1.417746958 | 29.4521 | 29.1453 | 28.8832 | 28.5582 | 28.6319 | 28.335 |
| UV excision repair protein RAD23 homolog B | P54727;P54727-2 | RAD23B | Other | -0.0062 | 0.00976895 | 31.4135 | 30.8098 | 31.1722 | 31.262 | 31.2048 | 30.9101 |
| DNA repair protein RAD50 | Q92878;Q92878-2;Q92878-3 | RAD50 | Other | -1.133566667 | 0.405004284 | 26.0648 | 24.8022 | 28.0855 | 24.2322 | 24.8272 | 26.4924 |
| mRNA export factor | P78406 | RAE1 | Other | -0.285633333 | 0.173845236 | 26.0823 | 25.1769 | 27.0204 | 25.215 | 26.1556 | 26.0521 |
| Ras-related protein Ral-A | P11233 | RALA | Extracellular or secreted | 0.713333333 | 0.795952582 | 25.3763 | 24.7889 | 24.5776 | 26.2509 | 25.4906 | 25.1413 |
| Ras-related protein Ral-B | P11234;P11234-3;P11234-2 | RALB | Extracellular or secreted | 1.2583 | 2.176311765 | 27.0399 | 26.4831 | 26.8081 | 28.1598 | 28.0568 | 27.8894 |
| RNA-binding protein Raly | Q9UKM9-2;Q9UKM9 | RALY | Other | -0.279966667 | 0.097123543 | 25.4837 | 25.1256 | 27.9637 | 25.8123 | 25.1404 | 26.7804 |
| GTP-binding nuclear protein Ran | P62826 | RAN | Extracellular or secreted | -0.234833333 | 0.517448605 | 30.7087 | 31.0206 | 31.1208 | 30.4425 | 30.727 | 30.9761 |
| Ran-specific GTPase-activating protein | P43487-2;P43487 | RANBP1 | Other | 0.451133333 | 0.275175916 | 28.2696 | 27.9066 | 29.8404 | 29.4391 | 28.9514 | 28.9795 |
| Ran-binding protein 10 | Q6VN20-3;Q6VN20;Q6VN20 | RANBP10 | Other | -0.899766667 | 0.306652107 | 23.7479 | 23.792 | 24.2181 | 20.8755 | 23.851 | 24.3322 |
| E3 SUMO-protein ligase RanBP2 | P49792 | RANBP2 | Other | -2.611966667 | 0.900598434 | 24.7021 | 24.3955 | 27.6949 | 22.6315 | 22.9374 | 23.3877 |
| Ran-binding protein 3 | Q9H6Z4-3;Q9H6Z4;Q9H6Z4- | RANBP3 | Other | -0.389166667 | 0.943639619 | 28.6158 | 28.1534 | 28.4424 | 27.8233 | 27.9384 | 28.2824 |
| Ran-binding protein 9 | Q96S59;Q96S59-2;Q96S59-3 | RANBP9 | Other | -0.1887 | 0.24457033 | 26.0236 | 25.875 | 26.3675 | 25.4892 | 25.8302 | 26.3806 |
| Ran GTPase-activating protein 1 | P46060 | RANGAP1 | Other | -0.5784 | 0.480773206 | 27.5826 | 27.6016 | 28.1748 | 26.5174 | 27.0542 | 28.0522 |
| Ran guanine nucleotide release factor | Q9HD47-4;Q9HD47-2;Q9HD | RANGRF | Extracellular or secreted | -3.061966667 | 1.129409572 | 24.0157 | 24.2909 | 22.5032 | 21.4674 | 18.5182 | 21.6383 |
| Ras-related protein Rap-1A | P62834 | RAP1A | Extracellular or secreted | 1.735066667 | 0.319242287 | 21.7449 | 20.5063 | 24.3768 | 20.2668 | 26.2657 | 25.3007 |
| Ras-related protein Rap-1b;Ras-related protein Rap-1b-li | P61224-3;P61224;P61224-4; | RAP1B | Extracellular or secreted | 0.511 | 1.116142393 | 26.9594 | 26.5089 | 26.9756 | 27.4245 | 27.5243 | 27.0281 |
| Rap1 GTPase-activating protein 2 | Q684P5-3;Q684P5-2;Q684P5 | RAP1GAP2 | Extracellular or secreted | -0.6263 | 1.196374985 | 22.8177 | 22.173 | 22.4115 | 21.9896 | 21.8031 | 21.7306 |
| Rap1 GTPase-GDP dissociation stimulator 1 | P52306-4;P52306;P52306-5; | RAP1GDS1 | Extracellular or secreted | -1.039 | 0.758079653 | 24.617 | 25.046 | 25.8385 | 23.2617 | 24.1234 | 24.9994 |
| Ras-related protein Rap-2b;Ras-related protein Rap-2a | P61225;P10114 | RAP2B;RAP2A | Extracellular or secreted | 0.7909 | 1.092871457 | 22.8579 | 21.9745 | 22.3762 | 23.6233 | 23.0736 | 22.8844 |
| Ras-related protein Rap-2c | Q9Y3L5 | RAP2C | Extracellular or secreted | 2.0293 | 1.382899028 | 22.8567 | 23.5768 | 22.2145 | 24.4006 | 25.9726 | 24.3627 |
| Retinoic acid receptor responder protein 1 | P49788;P49788-2 | RARRES1 | Extracellular or secreted | 2.733566667 | 1.510664593 | 32.5209 | 33.2664 | 31.5476 | 35.2119 | 35.0972 | 35.2265 |
| Arginine--tRNA ligase, cytoplasmic | P54136 | RARS | Extracellular or secreted | -0.269466667 | 0.2953741 | 29.0506 | 29.8769 | 30.0733 | 29.0693 | 29.4382 | 29.6849 |
| Ras GTPase-activating protein 1 | P20936-2;P20936-4;P20936; | RASA1 | Extracellular or secreted | 0.2146 | 0.158254024 | 21.4671 | 21.5425 | 22.5666 | 21.3849 | 22.6237 | 22.2114 |
| Ribonucleoprotein PTB-binding 1 | Q8IY67;Q8IY67-2 | RAVER1 | Other | -1.744733333 | 0.750999572 | 24.3852 | 23.7509 | 26.666 | 22.8599 | 22.763 | 23.945 |
| Histone-binding protein RBBP4 | Q09028-3;Q09028;Q09028-2 | RBBP4 | Other | -0.479133333 | 0.672505912 | 31.5611 | 30.7818 | 31.5892 | 30.6509 | 30.6827 | 31.1611 |
| Retinoblastoma-binding protein 5 | Q15291-2;Q15291 | RBBP5 | Other | -0.5223 | 1.513546532 | 26.2737 | 26.2686 | 25.905 | 25.4883 | 25.8113 | 25.5808 |
| E3 ubiquitin-protein ligase RBBP6 | Q7Z6E9-2;Q7Z6E9;Q7Z6E9-4 | RBBP6 | Other | -0.0572 | 0.026810208 | 23.5677 | 23.2148 | 24.8503 | 24.4743 | 24.1825 | 22.8044 |
| Histone-binding protein RBBP7 | Q16576;Q16576-2 | RBBP7 | Other | -0.637433333 | 0.978801478 | 29.5583 | 28.7346 | 29.3369 | 28.3595 | 28.5605 | 28.7975 |
| DNA endonuclease RBBP8 | Q99708-3;Q99708;Q99708-2 | RBBP8 | Other | -2.276966667 | 0.882143199 | 26.6619 | 25.1414 | 23.7455 | 24.4641 | 21.5043 | 22.7495 |
| Putative hydrolase RBBP9 | O75884;O75884-2 | RBBP9 | Extracellular or secreted | -0.0219 | 0.02039986 | 28.6744 | 28.8072 | 27.9968 | 28.1704 | 28.9777 | 28.2646 |
| RNA-binding protein 10 | P98175-2;P98175;P98175-5; | RBM10 | Other | -1.1866 | 0.440911993 | 25.3405 | 25.0425 | 26.1391 | 22.4096 | 24.7121 | 25.8406 |
| RNA-binding protein 12 | Q9NTZ6 | RBM12 | Other | -1.714733333 | 1.073521222 | 26.5132 | 26.3391 | 26.5148 | 23.6991 | 25.0251 | 25.4987 |
| RNA-binding protein 12B | Q8IXT5 | RBM12B | Other | 0.066933333 | 0.031750218 | 22.3723 | 23.0081 | 21.0624 | 21.5337 | 22.1484 | 22.9615 |
| Putative RNA-binding protein 15 | Q96T37-4;Q96T37-2;Q96T37 | RBM15 | Other | -1.1895 | 0.561571063 | 24.5724 | 23.0697 | 25.8514 | 23.7164 | 23.157 | 23.0516 |
| RNA-binding protein 25 | P49756 | RBM25 | Other | -1.595333333 | 1.38046054 | 28.5993 | 28.042 | 29.2548 | 26.3722 | 26.971 | 27.7669 |
| RNA-binding protein 26 | Q5T8P6-4;Q5T8P6;Q5T8P6-5 | RBM26 | Other | -1.195966667 | 0.619477001 | 23.8972 | 23.0986 | 25.5801 | 22.6873 | 23.0256 | 23.2751 |
| RNA-binding protein 3 | P98179 | RBM3 | Extracellular or secreted | 3.328933333 | 0.954310239 | 24.175 | 20.974 | 24.9663 | 26.7442 | 26.8635 | 26.4944 |
| RNA-binding protein 39 | Q14498-3;Q14498-2;Q14498 | RBM39 | Other | -1.412966667 | 0.569110031 | 25.8935 | 25.6082 | 26.5731 | 23.3415 | 24.0395 | 26.4549 |
| RNA-binding protein 4;RNA-binding protein 4B | Q9BWF3;Q9BWF3-4;Q9BQ04 | RBM4;RBM4B | Other | 1.3554 | 0.297818169 | 18.1131 | 22.0201 | 23.493 | 21.2355 | 22.574 | 23.8829 |
| RNA-binding protein 8A | Q9Y5S9;Q9Y5S9-2 | RBM8A | Other | -0.107033333 | 0.157632934 | 28.3937 | 28.4255 | 29.0943 | 28.4218 | 28.7027 | 28.4679 |
| RNA-binding motif, single-stranded-interacting protein 1 | P29558-2;P29558;Q6XE24-3 | RBMS1;RBMS3 | Other | -0.026433333 | 0.026147017 | 23.4646 | 23.7546 | 23.7275 | 23.4012 | 23.2288 | 24.2374 |
| RNA-binding motif protein, X chromosome;RNA-binding | P38159;P38159-2 | RBMX | Extracellular or secreted | -0.474933333 | 0.440306171 | 30.5809 | 30.1529 | 31.1659 | 29.5332 | 30.7629 | 30.1788 |
| RNA binding motif protein, X-linked-like-1 | Q96E39 | RBMXL1 | Other | -2.297966667 | 1.10804414 | 23.2099 | 23.0642 | 24.3556 | 20.1643 | 20.8234 | 22.7481 |
| Retinol-binding protein 1 | P09455;P09455-2;P09455-3 | RBP1 | Other | 0.3918 | 0.30370255 | 29.7166 | 30.2013 | 29.649 | 29.3608 | 30.4466 | 30.9349 |
| Retinol-binding protein 4;Plasma retinol-binding protein | P02753 | RBP4 | Extracellular or secreted | -1.0107 | 0.342428705 | 24.649 | 23.4234 | 22.5059 | 22.732 | 24.1654 | 20.6488 |
| E3 ubiquitin-protein ligase RBX1;E3 ubiquitin-protein liga | P62877 | RBX1 | Other | -2.045166667 | 0.655085543 | 26.7577 | 27.08 | 27.0512 | 22.5891 | 26.2376 | 25.9267 |
| Regulator of chromosome condensation | P18754;P18754-2 | RCC1 | Other | -1.2052 | 0.602735465 | 28.0748 | 28.2114 | 29.5024 | 26.019 | 27.6118 | 28.5422 |
| Protein RCC2 | Q9P258 | RCC2 | Extracellular or secreted | -1.0575 | 1.33373215 | 32.3968 | 32.143 | 31.4596 | 31.3105 | 31.0178 | 30.4986 |
| RING finger and CHY zinc finger domain-containing prote | Q96PM5-6;Q96PM5-2;Q96P | RCHY1 | Other | 0.223833333 | 0.089848866 | 19.9181 | 22.1237 | 19.502 | 20.4823 | 21.2094 | 20.5236 |
| Reticulocalbin-1 | Q15293;Q15293-2 | RCN1 | Extracellular or secreted | 1.348733333 | 0.609966831 | 29.3252 | 28.1233 | 30.7724 | 31.9019 | 30.5861 | 29.7791 |
| Reticulocalbin-2 | Q14257;Q14257-2 | RCN2 | Extracellular or secreted | 2.8171 | 0.662481824 | 24.8166 | 20.0558 | 24.9637 | 26.9589 | 25.8169 | 25.5116 |
| REST corepressor 1 | Q9UKL0 | RCOR1 | Extracellular or secreted | -2.221666667 | 1.121805612 | 25.2133 | 24.0316 | 26.5722 | 23.2432 | 22.0825 | 23.8264 |
| Retinol dehydrogenase 11 | Q8TC12-2;Q8TC12 | RDH11 | Extracellular or secreted | 2.244633333 | 0.885348201 | 20.2918 | 21.8559 | 23.463 | 23.9992 | 23.9306 | 24.4148 |
| Radixin | P35241;P35241-5;P35241-4 | RDX | Extracellular or secreted | -0.233933333 | 0.623733264 | 30.1852 | 30.0512 | 29.8807 | 30.0808 | 29.6659 | 29.6686 |
| ATP-dependent DNA helicase Q1 | P46063 | RECQL | Other | -0.442866667 | 0.172821276 | 27.2774 | 28.2373 | 29.4108 | 26.645 | 27.7347 | 29.2172 |
| Receptor expression-enhancing protein 5 | Q00765 | REEP5 | Extracellular or secreted | 2.288266667 | 0.914488341 | 22.2725 | 24.4669 | 25.4901 | 25.9412 | 25.8335 | 27.3196 |
| Transcription factor p65 | Q04206-2;Q04206-3;Q04206 | RELA | Other | -0.1866 | 0.200602347 | 22.712 | 22.9099 | 23.3153 | 22.3379 | 22.6712 | 23.3683 |
| Replication initiator 1 | Q9BWE0;Q9BWE0-4 | REPIN1 | Extracellular or secreted | 0.236366667 | 0.127769754 | 22.0162 | 21.2795 | 21.8338 | 23.1677 | 21.2468 | 21.4241 |
| RalBP1-associated Eps domain-containing protein 1 | Q96D71-2;Q96D71-4;Q96D7 | REPS1 | Extracellular or secreted | 0.1581 | 0.111254628 | 23.1279 | 23.4946 | 24.6623 | 23.6924 | 23.7917 | 24.275 |
| Oligoribonuclease, mitochondrial | Q9Y3B8-3 | REXO2 | Other | -0.337833333 | 1.014053976 | 28.2333 | 28.121 | 28.1923 | 27.6366 | 27.8455 | 28.051 |

| | | | | | | | | | | | |
|---|---|---|---|---|---|---|---|---|---|---|---|
| Raftlin | Q14699 | RFTN1 | Extracellular or secreted | 1.642766667 | 1.201592307 | 22.0316 | 21.0451 | 20.7808 | 23.3893 | 21.9335 | 23.463 |
| RanBP2-like and GRIP domain-containing protein 3 | A6NKT7 | RGPD3 | Other | -0.5407 | 0.220055551 | 22.3962 | 21.0826 | 24.0855 | 21.8471 | 21.6474 | 22.4477 |
| Regulator of G-protein signaling 10 | O43665-2;O43665;O43665-3 | RGS10 | Extracellular or secreted | 0.426866667 | 0.459425037 | 23.6955 | 23.9873 | 24.0264 | 24.7049 | 24.6543 | 23.6306 |
| Transforming protein RhoA | P61586 | RHOA | Extracellular or secreted | 0.1511 | 0.202443184 | 29.0531 | 29.2485 | 29.6644 | 29.0485 | 29.8046 | 29.5662 |
| Rho-related GTP-binding protein RhoG | P84095 | RHOG | Extracellular or secreted | 0.6828 | 1.302364611 | 24.3417 | 23.7255 | 24.207 | 24.5737 | 24.673 | 25.0759 |
| Telomere-associated protein RIF1 | Q5UIP0-2;Q5UIP0 | RIF1 | Extracellular or secreted | -3.073033333 | 1.172812035 | 26.0227 | 24.2127 | 26.4035 | 21.6242 | 21.3892 | 24.4064 |
| E3 ubiquitin-protein ligase RING1 | Q06587-2;Q06587 | RING1 | Other | -2.220866667 | 0.873932661 | 23.5001 | 23.1245 | 25.3317 | 21.3182 | 20.4176 | 23.5579 |
| Regulator of microtubule dynamics protein 1 | Q96DB5-2;Q96DB5;Q96DB5- | RMDN1 | Extracellular or secreted | -2.800066667 | 0.518823947 | 26.2115 | 26.6407 | 26.4076 | 19.5706 | 26.0416 | 25.2456 |
| Regulator of microtubule dynamics protein 3 | Q96TC7-2;Q96TC7 | RMDN3 | Other | -2.272233333 | 1.166776023 | 26.9897 | 25.8297 | 25.3634 | 24.1813 | 22.3867 | 24.7981 |
| Protein RMD5 homolog A | Q9H871;Q9H871-2 | RMND5A | Other | -0.656766667 | 0.525735445 | 23.1327 | 23.0263 | 23.7782 | 21.7404 | 22.966 | 23.2605 |
| Ribonuclease 4 | P34096 | RNASE4 | Extracellular or secreted | -1.058966667 | 1.751146515 | 26.1751 | 26.4037 | 26.6186 | 25.71 | 24.9975 | 25.313 |
| Ribonuclease H2 subunit C | Q8TDP1 | RNASEH2C | Other | -1.353366667 | 0.421221471 | 21.2128 | 23.1869 | 24.6182 | 21.5112 | 20.0834 | 23.3632 |
| Ribonuclease T2 | O00584 | RNASET2 | Extracellular or secreted | 0.831666667 | 0.521743693 | 29.5865 | 30.0345 | 28.1497 | 30.5694 | 30.3682 | 29.3281 |
| E3 ubiquitin-protein ligase RNF114 | Q9Y508;Q9Y508-2 | RNF114 | Other | 0.063866667 | 0.065571468 | 25.2173 | 26.1719 | 26.016 | 25.5643 | 25.5926 | 26.0399 |
| E3 ubiquitin-protein ligase RNF13 | O43567;O43567-2 | RNF13 | Other | 3.846333333 | 2.491429152 | 21.2985 | 20.5765 | 21.7058 | 25.5475 | 25.4338 | 24.1385 |
| E3 ubiquitin-protein ligase RING2 | Q99496;Q99496-2 | RNF2 | Other | -0.740266667 | 0.86829179 | 24.0964 | 24.0928 | 24.9747 | 23.1523 | 23.7295 | 24.0613 |
| E3 ubiquitin-protein ligase RNF213 | Q63HN8;Q63HN8-4 | RNF213 | Other | -1.3374 | 1.056289453 | 23.4396 | 22.2489 | 23.8233 | 22.2099 | 21.8765 | 21.4132 |
| RING-box protein 2 | Q9UBF6;Q9UBF6-4 | RNF7 | Other | -0.715833333 | 0.251043293 | 23.0033 | 23.9287 | 22.485 | 20.4283 | 23.3574 | 23.4838 |
| Ribonuclease inhibitor | P13489 | RNH1 | Extracellular or secreted | -0.432566667 | 0.843336663 | 30.49 | 30.1455 | 30.801 | 29.8368 | 30.0064 | 30.2956 |
| mRNA cap guanine-N7 methyltransferase | O43148;O43148-2 | RNMT | Other | -0.709166667 | 0.179542354 | 21.9501 | 23.7379 | 26.3125 | 22.3467 | 22.73 | 24.7963 |
| Aminopeptidase B | Q9H4A4 | RNPEP | Extracellular or secreted | -0.365066667 | 0.575159398 | 32.1181 | 31.9857 | 31.476 | 31.6262 | 31.7632 | 31.0952 |
| RNA-binding protein with serine-rich domain 1 | Q15287-3;Q15287-2;Q15287 | RNPS1 | Other | -0.6289 | 0.422627788 | 23.9874 | 23.8438 | 25.1214 | 22.7247 | 24.2817 | 24.0595 |
| Roundabout homolog 1 | Q9Y6N7-6;Q9Y6N7-5;Q9Y6N | ROBO1 | Extracellular or secreted | -0.459166667 | 1.043441064 | 28.0989 | 28.3357 | 27.7869 | 27.3599 | 27.7294 | 27.7547 |
| Rho-associated protein kinase 1 | Q13464 | ROCK1 | Extracellular or secreted | 0.408133333 | 0.28403786 | 26.5122 | 26.1824 | 26.2765 | 25.6832 | 27.2151 | 27.2972 |
| Rho-associated protein kinase 2 | O75116 | ROCK2 | Extracellular or secreted | -0.022333333 | 0.017702279 | 26.8307 | 26.9725 | 27.4589 | 27.7698 | 26.5529 | 26.8724 |
| Protein XRP2 | O75695 | RP2 | Extracellular or secreted | 0.578766667 | 1.29135549 | 23.7066 | 23.3554 | 23.9216 | 24.0394 | 24.296 | 24.3845 |
| Replication protein A 70 kDa DNA-binding subunit;Replic | P27694 | RPA1 | Other | -1.2315 | 0.881778398 | 28.1397 | 28.6202 | 29.1034 | 26.3974 | 27.5607 | 28.2107 |
| Replication protein A 32 kDa subunit | P15927;P15927-2;P15927-3 | RPA2 | Other | -1.2864 | 1.030477171 | 25.6296 | 25.4223 | 26.4683 | 23.7665 | 24.5194 | 25.3751 |
| Replication protein A 14 kDa subunit | P35244 | RPA3 | Other | -0.3802 | 0.270625207 | 26.2739 | 27.4726 | 26.9752 | 25.6556 | 27.0551 | 26.8704 |
| RNA polymerase II-associated protein 3 | Q9H6T3-3;Q9H6T3-2;Q9H6T | RPAP3 | Other | 0.273166667 | 0.226888736 | 23.0075 | 23.1635 | 23.5767 | 22.8095 | 23.4998 | 24.2579 |
| Ribulose-phosphate 3-epimerase;Ribulose-phosphate 3- | Q96AT9;Q96AT9-4;Q96AT9- | RPE;RPEL1 | Extracellular or secreted | 0.4714 | 0.565377972 | 27.0693 | 27.5226 | 26.6077 | 27.9495 | 27.6063 | 27.058 |
| Ribose-5-phosphate isomerase | P49247 | RPIA | Other | -0.952466667 | 0.905509951 | 27.848 | 27.5445 | 26.7853 | 27.0958 | 26.4144 | 25.8102 |
| 60S ribosomal protein L10 | P27635 | RPL10 | Other | 1.8919 | 2.80568363 | 25.748 | 25.7708 | 26.1469 | 27.7653 | 27.4769 | 28.0992 |
| 60S ribosomal protein L10a | P62906 | RPL10A | Extracellular or secreted | 1.743633333 | 2.004011103 | 29.281 | 28.7476 | 29.6636 | 31.188 | 31.2942 | 30.4409 |
| 60S ribosomal protein L11 | P62913-2;P62913 | RPL11 | Extracellular or secreted | 1.132066667 | 1.610581887 | 26.578 | 26.1237 | 26.869 | 27.6639 | 27.5191 | 27.7839 |
| 60S ribosomal protein L12 | P30050;P30050-2 | RPL12 | Extracellular or secreted | 0.819966667 | 1.170264471 | 29.6464 | 29.8465 | 30.4315 | 30.712 | 30.8871 | 30.7852 |
| 60S ribosomal protein L13 | P26373;P26373-2 | RPL13 | Other | 2.8132 | 2.550665405 | 25.5116 | 25.798 | 25.5288 | 28.8809 | 28.1032 | 28.2939 |
| 60S ribosomal protein L13a;Putative 60S ribosomal prote | P40429;Q6NVV1 | RPL13A;RPL13A | Other | 3.0065 | 3.219193061 | 24.8403 | 24.3673 | 24.5993 | 27.6226 | 27.5101 | 27.6937 |
| 60S ribosomal protein L14 | P50914 | RPL14 | Extracellular or secreted | 2.7758 | 2.778690461 | 24.9305 | 24.7169 | 25.0527 | 27.5647 | 27.3793 | 28.0835 |
| 60S ribosomal protein L15 | P61313 | RPL15 | Extracellular or secreted | 2.759666667 | 2.222015256 | 25.7831 | 24.4466 | 25.28 | 27.339 | 28.046 | 28.4037 |
| 60S ribosomal protein L17 | P18621;P18621-3;P18621-2 | RPL17 | Other | 1.730266667 | 2.369690798 | 26.4237 | 26.2357 | 26.1013 | 28.0892 | 27.6148 | 28.2475 |
| 60S ribosomal protein L18 | Q07020-2;Q07020 | RPL18 | Other | 2.6937 | 3.678725412 | 26.7141 | 26.2815 | 26.7394 | 29.2833 | 29.0727 | 29.4601 |
| 60S ribosomal protein L18a | Q02543 | RPL18A | Other | 2.267766667 | 3.432333035 | 24.7289 | 24.3473 | 24.8403 | 27.1465 | 26.7398 | 26.8335 |
| 60S ribosomal protein L19 | P84098 | RPL19 | Other | 1.418166667 | 2.245841755 | 24.2262 | 24.8198 | 24.8154 | 25.9423 | 25.8443 | 26.3293 |
| 60S ribosomal protein L21 | P46778 | RPL21 | Other | 2.6295 | 3.524637692 | 25.5628 | 25.5543 | 25.3084 | 28.2126 | 27.8298 | 28.2716 |
| 60S ribosomal protein L22 | P35268 | RPL22 | Extracellular or secreted | 1.640466667 | 2.32538891 | 25.5587 | 26.2858 | 25.7888 | 27.3924 | 27.8831 | 27.2792 |
| 60S ribosomal protein L23 | P62829 | RPL23 | Extracellular or secreted | 1.376766667 | 3.042725138 | 25.6639 | 25.3028 | 25.6268 | 26.736 | 26.9007 | 27.0871 |
| 60S ribosomal protein L23a | P62750 | RPL23A | Extracellular or secreted | 2.145866667 | 1.793356292 | 26.1217 | 27.4004 | 26.3002 | 29.1976 | 28.4002 | 28.6621 |
| 60S ribosomal protein L24 | P83731 | RPL24 | Extracellular or secreted | 2.612233333 | 2.845141378 | 24.2691 | 24.3113 | 24.0378 | 27.186 | 26.6777 | 26.5912 |
| 60S ribosomal protein L26;60S ribosomal protein L26-like | P61254;Q9UNX3 | RPL26;RPL26L1 | Extracellular or secreted | 2.400733333 | 1.263667081 | 24.3 | 26.4216 | 24.735 | 27.9498 | 27.2611 | 27.4479 |
| 60S ribosomal protein L27 | P61353 | RPL27 | Extracellular or secreted | 1.956533333 | 3.304351394 | 26.2033 | 26.3474 | 26.4897 | 28.2437 | 28.1191 | 28.5472 |
| 60S ribosomal protein L27a | P46776 | RPL27A | Other | 2.706366667 | 3.803047817 | 24.4288 | 24.7782 | 24.8467 | 27.5136 | 27.2212 | 27.438 |
| 60S ribosomal protein L28 | P46779-4;P46779-5;P46779; | RPL28 | Extracellular or secreted | 1.881833333 | 2.911535056 | 25.4954 | 25.6661 | 26.0462 | 27.9019 | 27.3365 | 27.6148 |
| 60S ribosomal protein L29 | P47914 | RPL29 | Other | 1.377766667 | 1.729972172 | 26.0081 | 25.8866 | 26.3947 | 27.8456 | 26.9525 | 27.6246 |
| 60S ribosomal protein L3 | P39023 | RPL3 | Extracellular or secreted | 1.918066667 | 3.818333269 | 27.2454 | 27.4622 | 27.5832 | 29.2291 | 29.2807 | 29.5352 |
| 60S ribosomal protein L30 | P62888 | RPL30 | Extracellular or secreted | 1.724066667 | 2.812576462 | 26.831 | 27.1558 | 27.3099 | 28.6878 | 29.1535 | 28.6276 |
| 60S ribosomal protein L31 | P62899;P62899-3;P62899-2 | RPL31 | Extracellular or secreted | 2.8615 | 1.624321765 | 25.466 | 26.7945 | 25.0109 | 28.7935 | 28.8061 | 28.2563 |
| 60S ribosomal protein L32 | P62910 | RPL32 | Other | 2.7446 | 3.837374904 | 25.1293 | 25.6059 | 25.3721 | 27.9269 | 28.0335 | 28.3807 |
| 60S ribosomal protein L34 | P49207 | RPL34 | Extracellular or secreted | 3.971266667 | 1.103573853 | 25.4595 | 25.5955 | 21.8536 | 27.9088 | 28.2192 | 28.6944 |
| 60S ribosomal protein L35 | P42766 | RPL35 | Other | 2.2892 | 2.027138354 | 24.8047 | 25.4214 | 24.4202 | 27.1053 | 27.0365 | 27.3721 |
| 60S ribosomal protein L35a | P18077 | RPL35A | Extracellular or secreted | 1.973733333 | 0.653001268 | 24.708 | 21.3668 | 24.8412 | 25.3783 | 25.4792 | 25.9797 |
| 60S ribosomal protein L36 | Q9Y3U8 | RPL36 | Other | 3.847433333 | 3.007065261 | 23.4294 | 22.6867 | 23.4307 | 27.2445 | 26.834 | 27.0106 |
| 60S ribosomal protein L36a;60S ribosomal protein L36a-l | P83881;Q969Q0 | RPL36A;RPL36A | Other | 1.400133333 | 2.259901979 | 26.167 | 26.5743 | 26.3868 | 27.7324 | 27.7846 | 27.8115 |
| 60S ribosomal protein L37a;Putative 60S ribosomal prote | P61513;A6NKH3 | RPL37A;RPL37A | Extracellular or secreted | 2.834966667 | 2.530825657 | 23.2784 | 24.2311 | 23.8264 | 26.0174 | 26.6823 | 27.1411 |
| 60S ribosomal protein L38 | P63173 | RPL38 | Other | 2.875733333 | 1.018196067 | 23.4609 | 20.4095 | 23.2472 | 24.9567 | 25.5282 | 25.2599 |
| Putative 60S ribosomal protein L39-like 5;60S ribosomal | Q59GN2;P62891 | RPL39P5;RPL39 | Other | 1.848366667 | 0.466150522 | 20.4596 | 23.3728 | 22.2123 | 21.0099 | 25.4922 | 25.0877 |
| 60S ribosomal protein L4 | P36578 | RPL4 | Extracellular or secreted | 2.757033333 | 2.944177385 | 26.4386 | 26.263 | 26.9986 | 29.4128 | 29.0569 | 29.5016 |
| 60S ribosomal protein L5 | P46777 | RPL5 | Extracellular or secreted | 2.346266667 | 2.314930771 | 27.015 | 27.3126 | 27.9838 | 29.5593 | 29.6971 | 30.0938 |
| 60S ribosomal protein L6 | Q02878 | RPL6 | Other | 1.0911 | 1.174658656 | 28.216 | 28.0915 | 29.1553 | 29.5474 | 29.3557 | 29.833 |
| 60S ribosomal protein L7 | P18124 | RPL7 | Extracellular or secreted | 2.296366667 | 3.046970525 | 27.2297 | 26.709 | 27.3081 | 29.2526 | 29.227 | 29.6563 |
| 60S ribosomal protein L7a | P62424 | RPL7A | Extracellular or secreted | 2.421733333 | 2.086936644 | 27.3278 | 27.0649 | 28.1308 | 29.981 | 29.6633 | 30.1444 |
| 60S ribosomal protein L8 | P62917 | RPL8 | Other | 1.6803 | 3.494856309 | 27.2909 | 27.035 | 27.3709 | 28.9834 | 28.7687 | 28.9856 |
| 60S ribosomal protein L9 | P32969 | RPL9 | Other | 3.587466667 | 0.952459327 | 21.1447 | 25.327 | 24.8038 | 27.4505 | 27.426 | 27.1614 |
| 60S acidic ribosomal protein P0;60S acidic ribosomal pro | P05388;P05388-2;Q8NHW5 | RPLP0;RPLP0P6 | Extracellular or secreted | 0.500366667 | 0.227221821 | 29.2807 | 29.4708 | 31.4465 | 30.0218 | 30.1097 | 31.5676 |
| 60S acidic ribosomal protein P1 | P05386 | RPLP1 | Extracellular or secreted | -0.157066667 | 0.096690733 | 29.7411 | 29.8243 | 31.3951 | 29.9828 | 30.1109 | 30.3956 |
| 60S acidic ribosomal protein P2 | P05387 | RPLP2 | Extracellular or secreted | -0.711266667 | 0.567570366 | 30.9382 | 31.7162 | 32.385 | 30.3579 | 30.9254 | 31.6223 |
| Dolichyl-diphosphooligosaccharide--protein glycosyltrans | P04843 | RPN1 | Other | 1.167533333 | 2.088243502 | 23.2973 | 23.7465 | 23.1962 | 24.2718 | 24.8549 | 24.6159 |
| Dolichyl-diphosphooligosaccharide--protein glycosyltrans | P04844-2;P04844 | RPN2 | Extracellular or secreted | -0.368466667 | 0.188603417 | 26.4421 | 24.753 | 26.7614 | 25.7479 | 26.2223 | 24.8809 |
| Ribonuclease P protein subunit p30 | P78346;P78346-2 | RPP30 | Other | -0.7737 | 0.153772751 | 24.2484 | 24.3943 | 25.3766 | 20.4516 | 25.9553 | 25.2913 |
| Ribonuclease P protein subunit p40 | O75818-2;O75818 | RPP40 | Other | -0.585666667 | 0.352269428 | 23.2397 | 23.7883 | 23.6541 | 21.7954 | 23.2611 | 23.8686 |
| Regulation of nuclear pre-mRNA domain-containing prot | Q96P16;Q96P16-3 | RPRD1A | Other | -0.145533333 | 0.062494837 | 22.9146 | 21.1662 | 23.0741 | 22.3864 | 21.2596 | 23.0723 |
| Regulation of nuclear pre-mRNA domain-containing prot | Q9NQG5 | RPRD1B | Other | -1.300433333 | 1.155333258 | 27.5674 | 27.4263 | 27.8236 | 25.5208 | 26.6962 | 26.699 |
| 40S ribosomal protein S10 | P46783 | RPS10 | Extracellular or secreted | 2.2171 | 2.934728568 | 24.0652 | 24.3602 | 23.7546 | 26.0821 | 26.6714 | 26.0778 |
| 40S ribosomal protein S11 | P62280 | RPS11 | Extracellular or secreted | 1.0716 | 1.137376276 | 27.5104 | 27.8179 | 28.327 | 28.4696 | 28.765 | 29.6355 |
| 40S ribosomal protein S12 | P25398 | RPS12 | Other | 0.3945 | 0.527191439 | 29.2643 | 29.4208 | 30.0688 | 29.552 | 30.1215 | 30.2639 |
| 40S ribosomal protein S13 | P62277 | RPS13 | Extracellular or secreted | 1.903033333 | 2.898928967 | 26.2083 | 25.7318 | 26.0244 | 27.6719 | 27.7674 | 28.2343 |
| 40S ribosomal protein S14 | P62263 | RPS14 | Extracellular or secreted | 1.5358 | 1.105032561 | 26.2962 | 25.8018 | 27.439 | 28.1851 | 27.7966 | 28.1627 |
| 40S ribosomal protein S15 | P62841 | RPS15 | Other | 1.546333333 | 0.965161869 | 24.8982 | 25.2532 | 25.0495 | 25.4931 | 27.0056 | 27.3412 |
| 40S ribosomal protein S15a | P62244 | RPS15A | Extracellular or secreted | 1.2605 | 1.18027455 | 25.8664 | 26.6082 | 26.9475 | 26.8506 | 27.6384 | 28.1146 |
| 40S ribosomal protein S16 | P62249 | RPS16 | Extracellular or secreted | 1.577633333 | 5.187936854 | 26.3358 | 26.4069 | 26.4533 | 27.9775 | 27.9308 | 28.0206 |
| 40S ribosomal protein S17 | P08708 | RPS17 | Extracellular or secreted | 1.516333333 | 1.321025602 | 26.3736 | 26.5737 | 26.4075 | 27.2552 | 28.3789 | 28.2697 |
| 40S ribosomal protein S18 | P62269 | RPS18 | Extracellular or secreted | 2.0025 | 2.707517753 | 27.1591 | 26.9015 | 27.5254 | 29.4061 | 28.788 | 29.3994 |
| 40S ribosomal protein S19 | P39019 | RPS19 | Extracellular or secreted | 3.264666667 | 3.408910357 | 25.5243 | 25.9215 | 25.172 | 29.1417 | 28.5865 | 28.6836 |
| 40S ribosomal protein S2 | P15880 | RPS2 | Extracellular or secreted | 1.547066667 | 1.928820344 | 27.1681 | 27.4547 | 27.8379 | 28.6303 | 28.9313 | 29.5403 |
| 40S ribosomal protein S20 | P60866;P60866-2 | RPS20 | Extracellular or secreted | 2.050566667 | 0.994345722 | 25.0338 | 26.7479 | 27.4436 | 28.288 | 28.523 | 28.566 |
| 40S ribosomal protein S21 | P63220 | RPS21 | Other | 0.8542 | 1.140927859 | 27.3839 | 27.3494 | 28.0467 | 27.9175 | 28.7396 | 28.6855 |
| 40S ribosomal protein S23 | P62266 | RPS23 | Other | 1.434133333 | 1.406427843 | 24.6138 | 24.881 | 24.5638 | 25.6376 | 25.977 | 26.7464 |
| 40S ribosomal protein S24 | P62847-2;P62847-3;P62847; | RPS24 | Other | 3.070366667 | 2.813230288 | 23.2959 | 23.5469 | 23.8976 | 26.2581 | 26.511 | 27.1824 |
| 40S ribosomal protein S25 | P62851 | RPS25 | Extracellular or secreted | 2.011566667 | 2.556195071 | 26.4534 | 26.3947 | 26.4545 | 28.6705 | 28.3972 | 28.2696 |
| 40S ribosomal protein S26;Putative 40S ribosomal protei | P62854;Q5JNZ5 | RPS26;RPS26P | Extracellular or secreted | 3.365233333 | 1.746560303 | 22.1257 | 22.8012 | 24.2741 | 26.2165 | 25.9949 | 27.0853 |
| 40S ribosomal protein S27;40S ribosomal protein S27-like | P42677;Q71UM5 | RPS27;RPS27L | Other | 2.267733333 | 1.918262124 | 26.1875 | 25.0909 | 25.8348 | 27.5054 | 27.6599 | 28.7511 |
| Ubiquitin-40S ribosomal protein S27a;Ubiquitin;40S ribo | P62979;P62987;P0CG47;P0C | RPS27A;UBA52 | Extracellular or secreted | 0.445033333 | 0.399623304 | 33.3089 | 33.1636 | 32.279 | 33.9167 | 33.4374 | 32.7325 |
| 40S ribosomal protein S28 | P62857 | RPS28 | Extracellular or secreted | 1.993233333 | 1.188781025 | 25.4925 | 24.8245 | 26.9264 | 27.2405 | 28.3471 | 27.6355 |
| 40S ribosomal protein S29 | P62273;P62273-2 | RPS29 | Extracellular or secreted | 1.962366667 | 1.06674282 | 24.8783 | 24.4531 | 24.5968 | 25.5835 | 26.46 | 27.7718 |
| 40S ribosomal protein S3 | P23396;P23396-2 | RPS3 | Extracellular or secreted | 1.110833333 | 1.465930707 | 29.1915 | 28.8895 | 29.7217 | 30.1689 | 30.0833 | 30.883 |
| 40S ribosomal protein S3a | P61247 | RPS3A | Extracellular or secreted | 1.828366667 | 1.776545668 | 27.8413 | 27.6737 | 28.052 | 29.1312 | 29.7573 | 30.1636 |
| 40S ribosomal protein S4, X isoform | P62701 | RPS4X | Extracellular or secreted | 1.591933333 | 2.043203908 | 27.2469 | 26.9895 | 27.6474 | 28.7363 | 28.5409 | 29.3824 |
| 40S ribosomal protein S5;40S ribosomal protein S5, N-te | P46782 | RPS5 | Extracellular or secreted | 2.5161 | 1.509908173 | 25.5944 | 26.4174 | 27.1654 | 28.8895 | 28.9351 | 28.9009 |
| 40S ribosomal protein S6 | P62753 | RPS6 | Other | 1.948733333 | 2.599173461 | 25.3782 | 25.9586 | 25.8403 | 27.301 | 27.6759 | 28.0464 |
| Ribosomal protein S6 kinase alpha-3;Ribosomal protein S | P51812;Q9UK32-2;Q9UK32 | RPS6KA3;RPS6 | Other | -0.231166667 | 0.07631585 | 22.9195 | 22.6532 | 25.7287 | 23.3574 | 23.2205 | 24.03 |
| 40S ribosomal protein S7 | P62081 | RPS7 | Extracellular or secreted | 2.067733333 | 2.415020797 | 25.7799 | 26.4006 | 26.5862 | 27.8412 | 28.6068 | 28.5219 |
| 40S ribosomal protein S8 | P62241 | RPS8 | Extracellular or secreted | 1.993833333 | 2.552899452 | 26.9792 | 26.8127 | 27.2982 | 28.7593 | 28.8561 | 29.4562 |
| 40S ribosomal protein S9 | P46781 | RPS9 | Extracellular or secreted | 2.028066667 | 2.875269365 | 26.8949 | 27.0245 | 27.4461 | 28.8746 | 29.0686 | 29.5065 |
| 40S ribosomal protein SA | P08865 | RPSA | Extracellular or secreted | 0.949733333 | 0.825032744 | 28.7388 | 28.8326 | 29.9369 | 29.3813 | 30.4615 | 30.5147 |
| RNA pseudouridylate synthase domain-containing prote | Q8IZ73;Q8IZ73-2 | RPUSD2 | Other | -0.5963 | 1.599025181 | 26.4192 | 26.0058 | 26.1864 | 25.8224 | 25.6004 | 25.3997 |
| Cell differentiation protein RCD1 homolog | Q92600;Q92600-2;Q92600-3 | RQCD1 | Extracellular or secreted | -0.785733333 | 0.520205339 | 22.7459 | 23.3578 | 24.0851 | 21.7872 | 22.4493 | 23.5951 |
| Ras-related GTP-binding protein C;Ras-related GTP-bindi | Q9HB90;Q9NQL2 | RRAGC;RRAGD | Other | 1.263933333 | 0.543933165 | 20.2707 | 22.5467 | 23.113 | 22.7931 | 22.8152 | 24.1139 |
| Ras-related protein R-Ras2 | P62070;P62070-4;P62070-3; | RRAS2 | Extracellular or secreted | 0.2082 | 0.219399701 | 25.3569 | 24.8575 | 25.7083 | 25.0823 | 25.44 | 26.025 |
| Ribosome-binding protein 1 | Q9P2E9;Q9P2E9-3 | RRBP1 | Other | -0.7487 | 0.681216223 | 31.2098 | 30.3225 | 31.7529 | 30.5564 | 30.1314 | 30.3513 |

| | | | | | | | | | | | |
|---|---|---|---|---|---|---|---|---|---|---|---|
| Ribonucleoside-diphosphate reductase large subunit | P23921 | RRM1 | Extracellular or secreted | -1.291733333 | 0.556416077 | 27.1666 | 26.8044 | 27.2888 | 24.1873 | 25.9859 | 27.2114 |
| Ribonucleoside-diphosphate reductase subunit M2 | P31350;P31350-2 | RRM2 | Other | -1.2965 | 1.245662878 | 29.675 | 29.5654 | 28.5279 | 27.793 | 27.8528 | 28.233 |
| RRP15-like protein | Q9Y3B9 | RRP15 | Other | -0.3603 | 0.332928356 | 22.192 | 21.9787 | 23.1951 | 22.4771 | 22.0983 | 21.7095 |
| Ribosomal RNA-processing protein 8 | O43159 | RRP8 | Extracellular or secreted | -2.210333333 | 1.547476858 | 24.504 | 23.2211 | 23.4564 | 21.5539 | 21.407 | 21.5896 |
| U3 small nucleolar RNA-interacting protein 2 | O43818 | RRP9 | Other | -0.9303 | 1.029606916 | 26.1323 | 25.301 | 26.4233 | 25.4829 | 24.8335 | 24.7493 |
| Remodeling and spacing factor 1 | Q96T23-2;Q96T23;Q96T23-3 | RSF1 | Extracellular or secreted | -0.2067 | 0.254278362 | 27.7326 | 27.1033 | 27.7595 | 27.1858 | 26.9954 | 27.7941 |
| Ribosomal L1 domain-containing protein 1 | O76021 | RSL1D1 | Other | 0.2355 | 0.523337984 | 25.0498 | 25.3194 | 25.6069 | 25.6772 | 25.3453 | 25.6601 |
| Ras suppressor protein 1 | Q15404;Q15404-2 | RSU1 | Extracellular or secreted | -0.447366667 | 0.593027943 | 26.9134 | 27.4753 | 27.6479 | 26.3983 | 27.0827 | 27.2135 |
| RNA 3-terminal phosphate cyclase | O00442;O00442-2 | RTCA | Other | 0.4403 | 0.233420504 | 22.7214 | 22.8292 | 23.3948 | 22.3476 | 23.2718 | 24.6469 |
| tRNA-splicing ligase RtcB homolog | Q9Y3I0 | RTCB | Other | -0.626566667 | 0.30480401 | 27.927 | 28.6179 | 29.2873 | 26.7914 | 27.9037 | 29.2574 |
| RNA polymerase-associated protein RTF1 homolog | Q92541 | RTF1 | Extracellular or secreted | -0.9785 | 0.625955014 | 26.6576 | 24.6776 | 26.1095 | 24.9991 | 24.6227 | 24.8874 |
| Protein RTF2 homolog | Q9BY42 | RTFDC1 | Other | 0.230266667 | 0.243734128 | 24.5316 | 24.4612 | 25.3352 | 24.5584 | 25.0566 | 25.4038 |
| Reticulon-3 | O95197-3;O95197-4;O95197 | RTN3 | Extracellular or secreted | -0.288933333 | 0.225505909 | 26.7981 | 26.056 | 27.0452 | 25.8491 | 26.0497 | 27.1337 |
| Reticulon-4 | Q9NQC3-2;Q9NQC3-5;Q9NQ | RTN4 | Extracellular or secreted | 0.573066667 | 0.346340919 | 28.1014 | 27.5879 | 28.783 | 28.6647 | 27.7707 | 29.7561 |
| Reticulon-4 receptor | Q9BZR6 | RTN4R | Extracellular or secreted | 0.202133333 | 0.177591117 | 23.194 | 24.1083 | 23.0627 | 23.8787 | 23.9902 | 23.1025 |
| Reticulon-4 receptor-like 2 | Q86UN3;Q86UN3-2 | RTN4RL2 | Extracellular or secreted | 1.1285 | 0.602438631 | 24.6154 | 25.0442 | 22.796 | 25.6477 | 25.7689 | 24.4245 |
| RUN and FYVE domain-containing protein 1 | Q96T51;Q96T51-2;Q96T51-3 | RUFY1 | Other | 0.523233333 | 0.152106734 | 22.9122 | 23.1537 | 26.2464 | 23.5177 | 24.5238 | 25.8405 |
| RuvB-like 1 | Q9Y265;Q9Y265-2 | RUVBL1 | Extracellular or secreted | -0.516666667 | 0.33851329 | 29.0748 | 28.8915 | 30.1685 | 28.0094 | 28.8846 | 29.6908 |
| RuvB-like 2 | Q9Y230;Q9Y230-2 | RUVBL2 | Extracellular or secreted | -0.390866667 | 0.255331352 | 29.1682 | 28.6172 | 30.164 | 28.3355 | 28.7391 | 29.7022 |
| RWD domain-containing protein 1 | Q9H446;Q9H446-2 | RWDD1 | Other | -0.961566667 | 1.073962677 | 24.386 | 23.4747 | 24.0306 | 23.499 | 22.4022 | 23.1054 |
| RWD domain-containing protein 4 | Q6NW29;Q6NW29-2 | RWDD4 | Other | 0.324 | 0.251783089 | 23.1686 | 22.4471 | 23.4368 | 22.6347 | 23.3327 | 24.0571 |
| Protein S100-A10 | P60903 | S100A10 | Extracellular or secreted | 0.595533333 | 0.767416153 | 25.8918 | 25.0301 | 24.9935 | 26.0288 | 25.9618 | 25.7114 |
| Protein S100-A11;Protein S100-A11, N-terminally proces | P31949 | S100A11 | Extracellular or secreted | 0.438166667 | 0.300735748 | 31.6621 | 32.4005 | 30.7178 | 32.2664 | 32.4319 | 31.3966 |
| Protein S100-A13 | Q99584 | S100A13 | Extracellular or secreted | 1.484033333 | 1.310628053 | 29.5295 | 29.2666 | 28.2058 | 31.053 | 30.489 | 29.912 |
| Protein S100-A14 | Q9HCY8 | S100A14 | Extracellular or secreted | 2.801333333 | 1.591758783 | 27.7959 | 27.6998 | 27.439 | 31.2787 | 30.4967 | 29.5633 |
| Protein S100-A16 | Q96FQ6 | S100A16 | Extracellular or secreted | 1.580933333 | 1.221595757 | 29.8074 | 30.2271 | 28.6376 | 30.7188 | 31.564 | 31.1321 |
| Protein S100-A3 | P33764 | S100A3 | Other | 1.449433333 | 0.704515054 | 21.0564 | 22.3059 | 19.6341 | 23.3598 | 21.9879 | 21.997 |
| Protein S100-A4 | P26447 | S100A4 | Extracellular or secreted | 0.266566667 | 0.196796823 | 22.8121 | 24.1032 | 22.8304 | 23.1871 | 24.0875 | 23.2708 |
| Protein S100-A6 | P06703 | S100A6 | Extracellular or secreted | -0.0835 | 0.027805229 | 33.7696 | 32.262 | 30.8935 | 33.3276 | 31.7966 | 31.5504 |
| Protein S100-A7 | P31151 | S100A7 | Extracellular or secreted | 1.746266667 | 0.658805181 | 30.5687 | 30.9167 | 28.4439 | 32.8335 | 32.4175 | 29.9171 |
| Protein S100-A7A | Q86SG5 | S100A7A | Other | 0.8945 | 0.337648025 | 21.7311 | 21.9382 | 20.235 | 23.4609 | 22.7258 | 20.4011 |
| Protein S100-A8;Protein S100-A8, N-terminally processe | P05109 | S100A8 | Extracellular or secreted | 0.671666667 | 0.410982916 | 32.1558 | 32.2788 | 30.7693 | 33.1329 | 32.6325 | 31.4535 |
| Protein S100-A9 | P06702 | S100A9 | Extracellular or secreted | 0.695633333 | 0.366342399 | 32.3164 | 32.6805 | 31.5938 | 34.0958 | 32.8667 | 31.7151 |
| Protein S100-P | P25815 | S100P | Extracellular or secreted | -0.451466667 | 0.34143045 | 30.3498 | 30.522 | 29.5065 | 30.285 | 29.9201 | 28.8188 |
| Protein SAAL1 | Q96ER3 | SAAL1 | Extracellular or secreted | -0.892833333 | 0.656689021 | 27.6964 | 26.5476 | 26.852 | 26.9406 | 26.2394 | 25.2375 |
| SUMO-activating enzyme subunit 1;SUMO-activating enz | Q9UBE0;Q9UBE0-2;Q9UBE0- | SAE1 | Other | -0.271033333 | 0.379832111 | 27.3922 | 27.2035 | 27.8802 | 26.8086 | 27.2872 | 27.567 |
| Scaffold attachment factor B1 | Q15424;Q15424-4;Q15424-3 | SAFB | Other | -0.028366667 | 0.014642166 | 28.5906 | 27.4226 | 29.428 | 28.8358 | 28.4231 | 28.0972 |
| Scaffold attachment factor B2 | Q14151 | SAFB2 | Extracellular or secreted | -2.895533333 | 1.052560821 | 26.4324 | 25.5429 | 26.9841 | 22.5154 | 25.4547 | 22.3027 |
| SAM domain-containing protein SAMSN-1;SAM and SH3 | Q9NSI8;Q9NSI8-3;O94885 | SAMSN1;SASH | Other | -1.239266667 | 0.77066235 | 24.0229 | 23.5556 | 22.8667 | 21.9461 | 23.4143 | 21.367 |
| Histone deacetylase complex subunit SAP18 | O00422 | SAP18 | Other | 0.3248 | 0.102133701 | 24.6873 | 24.4184 | 27.3348 | 24.6703 | 25.828 | 26.9166 |
| SAP30-binding protein | Q9UHR5-2;Q9UHR5 | SAP30BP | Other | 0.166166667 | 0.079332903 | 21.3589 | 22.3576 | 23.2259 | 21.519 | 22.6962 | 23.2257 |
| Specifically androgen-regulated gene protein | Q9BW04-2;Q9BW04 | SARG | Extracellular or secreted | 0.781633333 | 0.629153171 | 23.0478 | 24.5912 | 24.2859 | 24.4876 | 24.8891 | 24.8931 |
| SAP domain-containing ribonucleoprotein | P82979 | SARNP | Other | 0.934333333 | 0.497426196 | 26.5377 | 26.3295 | 28.5586 | 27.792 | 28.0627 | 28.3741 |
| Serine--tRNA ligase, cytoplasmic | P49591 | SARS | Extracellular or secreted | -0.086233333 | 0.053684637 | 27.4904 | 28.1475 | 28.7621 | 27.3289 | 28.0514 | 28.761 |
| U4/U6.U5 tri-snRNP-associated protein 1 | O43290 | SART1 | Other | -0.634733333 | 0.273822492 | 23.9901 | 21.8533 | 24.1372 | 21.6082 | 23.0278 | 23.4404 |
| Squamous cell carcinoma antigen recognized by T-cells 3 | Q15020;Q15020-4 | SART3 | Other | -1.7758 | 0.532156557 | 26.3606 | 25.4363 | 28.1923 | 23.7293 | 23.662 | 27.2705 |
| Ribosome maturation protein SBDS | Q9Y3A5 | SBDS | Other | -0.440166667 | 0.49083751 | 28.3869 | 29.0504 | 28.485 | 27.5759 | 28.3867 | 28.6392 |
| Suprabasin | Q6UWP8 | SBSN | Extracellular or secreted | 2.920066667 | 0.965833248 | 23.513 | 24.1231 | 25.9836 | 29.6818 | 25.9677 | 26.7304 |
| Protein SCAF11 | Q99590;Q99590-2 | SCAF11 | Other | -0.927766667 | 1.376256788 | 26.3989 | 25.854 | 26.6905 | 25.6801 | 25.2623 | 25.2177 |
| Splicing factor, arginine/serine-rich 15 | O95104-3;O95104;O95104-2 | SCAF4 | Other | -0.494066667 | 0.154642861 | 24.1234 | 24.0683 | 26.7831 | 26.0576 | 23.5205 | 23.9145 |
| Secretory carrier-associated membrane protein 3 | O14828-2;O14828 | SCAMP3 | Extracellular or secreted | 0.866966667 | 0.319868801 | 20.3963 | 22.5104 | 22.8033 | 23.6569 | 21.1501 | 23.5039 |
| Lysosome membrane protein 2 | Q14108;Q14108-2 | SCARB2 | Extracellular or secreted | 1.046933333 | 1.064289888 | 24.6591 | 24.2213 | 25.3656 | 26.2121 | 26.0081 | 25.1666 |
| Acyl-CoA desaturase | O00767 | SCD | Other | 3.238833333 | 1.927597399 | 21.8887 | 20.1991 | 20.9585 | 23.8592 | 24.3302 | 24.5734 |
| Sec1 family domain-containing protein 1 | Q8WVM8;Q8WVM8-2;Q8W\ | SCFD1 | Extracellular or secreted | -0.540266667 | 0.287994039 | 26.6875 | 27.3298 | 28.4077 | 26.0679 | 26.7367 | 27.9996 |
| Protein SCO1 homolog, mitochondrial | O75880 | SCO1 | Other | -0.630166667 | 0.540898727 | 25.7497 | 24.9123 | 26.4248 | 25.0729 | 25.4168 | 24.7066 |
| Non-specific lipid-transfer protein | P22307-6;P22307-2;P22307- | SCP2 | Extracellular or secreted | -0.6441 | 0.316809981 | 24.3208 | 24.5306 | 26.3548 | 23.4248 | 25.1643 | 24.6848 |
| Retinoid-inducible serine carboxypeptidase | Q9HB40;Q9HB40-2 | SCPEP1 | Extracellular or secreted | -1.2145 | 0.213545474 | 25.0669 | 25.2343 | 26.8642 | 20.5631 | 25.8692 | 27.0896 |
| Protein scribble homolog | Q14160;Q14160-3;Q14160-2 | SCRIB | Extracellular or secreted | -0.4905 | 0.223199309 | 25.6621 | 25.8304 | 26.199 | 23.841 | 26.2728 | 26.1062 |
| Secernin-1 | Q12765;Q12765-2;Q12765-3 | SCRN1 | Other | 0.659 | 0.52842725 | 29.0595 | 28.5597 | 28.1774 | 30.0528 | 29.2628 | 28.458 |
| Secernin-2 | Q96FV2-2;Q96FV2 | SCRN2 | Extracellular or secreted | -0.574433333 | 1.149105588 | 25.2476 | 25.1516 | 25.1287 | 24.9209 | 24.3447 | 24.539 |
| Syndecan-1 | P18827 | SDC1 | Extracellular or secreted | 1.6426 | 0.824486521 | 27.1468 | 26.9127 | 25.3145 | 29.3984 | 27.9507 | 26.9527 |
| Syndecan-2 | P34741 | SDC2 | Extracellular or secreted | 1.790666667 | 0.319245003 | 26.5745 | 26.5473 | 20.3428 | 26.7528 | 26.3516 | 25.7322 |
| Syndecan-4 | P31431;P31431-2 | SDC4 | Extracellular or secreted | 2.3515 | 1.260930996 | 30.3211 | 29.9867 | 28.4839 | 33.1614 | 31.7867 | 30.8981 |
| Syntenin-1 | O00560;O00560-3 | SDCBP | Extracellular or secreted | 0.7711 | 0.534863023 | 28.3336 | 28.9641 | 27.1266 | 28.7484 | 29.4312 | 28.558 |
| Syntenin-1 | O00560-2 | SDCBP | Extracellular or secreted | 0.017066667 | 0.015327833 | 24.0031 | 23.722 | 22.9081 | 23.3635 | 23.8447 | 23.4762 |
| Protein SDE2 homolog | Q6IQ49-3;Q6IQ49-2;Q6IQ49 | SDE2 | Other | -1.4098 | 0.580594628 | 24.7979 | 24.3933 | 25.3045 | 21.5957 | 24.1091 | 24.5615 |
| Stromal cell-derived factor 2 | Q99470 | SDF2 | Extracellular or secreted | 0.504366667 | 0.311075283 | 20.7555 | 21.7768 | 22.7878 | 22.5825 | 22.3896 | 21.8611 |
| Stromal cell-derived factor 2-like protein 1 | Q9HCN8 | SDF2L1 | Extracellular or secreted | 1.605666667 | 0.936062363 | 22.3471 | 23.9587 | 21.8026 | 24.7788 | 24.3647 | 23.7819 |
| 45 kDa calcium-binding protein | Q9BRK5;Q9BRK5-6;Q9BRK5- | SDF4 | Extracellular or secreted | 1.726066667 | 1.794818211 | 28.9534 | 27.8274 | 28.3773 | 30.4907 | 30.0515 | 29.7941 |
| Succinate dehydrogenase [ubiquinone] flavoprotein subu | P31040;P31040-2;P31040-3 | SDHA | Other | -2.6358 | 1.758933569 | 28.7523 | 28.8352 | 30.3008 | 27.2469 | 26.9315 | 25.8025 |
| Succinate dehydrogenase assembly factor 2, mitochondr | Q9NX18 | SDHAF2 | Other | -0.876133333 | 0.395227358 | 21.2283 | 22.8029 | 21.8418 | 22.5424 | 20.8572 | 19.845 |
| Succinate dehydrogenase [ubiquinone] iron-sulfur subun | P21912 | SDHB | Extracellular or secreted | 0.1789 | 0.027059536 | 22.2913 | 21.0565 | 23.4512 | 26.325 | 19.5205 | 21.4902 |
| Protein SEC13 homolog | P55735;P55735-2;P55735-4; | SEC13 | Extracellular or secreted | -0.3914 | 0.835319019 | 28.1672 | 27.7272 | 28.0995 | 27.3711 | 27.5205 | 27.9281 |
| SEC14-like protein 2 | O76054;O76054-5;O76054-4 | SEC14L2 | Extracellular or secreted | 0.1503 | 0.156649606 | 24.4403 | 25.469 | 25.272 | 24.9111 | 25.3448 | 25.3763 |
| Protein transport protein Sec16A | O15027-2;O15027-4;O15027 | SEC16A | Extracellular or secreted | 0.235033333 | 0.117323956 | 21.0136 | 20.9671 | 22.9643 | 22.2591 | 21.9587 | 21.4323 |
| Vesicle-trafficking protein SEC22b | O75396 | SEC22B | Other | -1.117866667 | 0.381473135 | 25.63 | 26.0732 | 26.1375 | 22.6402 | 26.0333 | 25.8136 |
| Protein transport protein Sec23A | Q15436;Q15436-2 | SEC23A | Other | -0.383 | 0.489284109 | 27.7708 | 28.153 | 28.4427 | 27.2548 | 27.7602 | 28.2025 |
| Protein transport protein Sec23B | Q15437 | SEC23B | Other | 0.445366667 | 0.801895391 | 26.5997 | 27.0072 | 27.3023 | 27.1892 | 27.3662 | 27.6899 |
| SEC23-interacting protein | Q9Y6Y8;Q9Y6Y8-2 | SEC23IP | Other | -0.953666667 | 0.677031365 | 24.9177 | 25.1226 | 25.283 | 23.1428 | 24.935 | 24.3845 |
| Protein transport protein Sec24A | O95486 | SEC24A | Other | -1.08 | 0.816853566 | 23.1074 | 22.9171 | 23.7526 | 21.3335 | 22.1296 | 23.074 |
| Protein transport protein Sec24B | O95487-2;O95487;O95487-3 | SEC24B | Other | -0.913566667 | 0.541329081 | 23.2053 | 23.1813 | 23.9041 | 22.0581 | 21.7124 | 23.7795 |
| Protein transport protein Sec24C | P53992 | SEC24C | Other | -0.9705 | 0.529736051 | 27.0169 | 26.8942 | 27.6662 | 25.1047 | 26.0673 | 27.4938 |
| Protein transport protein Sec31A | O94979-6;O94979-3;O94979 | SEC31A | Extracellular or secreted | -0.0669 | 0.032118203 | 26.8679 | 27.2195 | 27.868 | 26.0429 | 27.6174 | 28.0944 |
| Protein transport protein Sec61 subunit beta | P60468 | SEC61B | Other | -0.336466667 | 0.359563638 | 24.9904 | 24.3934 | 25.5213 | 24.727 | 24.9169 | 24.2518 |
| Translocation protein SEC63 homolog | Q9UGP8 | SEC63 | Other | -1.736166667 | 0.580826924 | 23.9594 | 20.9384 | 24.0884 | 22.6761 | 21.3043 | 19.7973 |
| Secreted and transmembrane protein 1 | Q8WVN6 | SECTM1 | Extracellular or secreted | -0.775166667 | 0.282538363 | 27.6317 | 28.0109 | 25.2314 | 27.1265 | 26.5365 | 24.8855 |
| Nucleoporin SEH1 | Q96EE3;Q96EE3-1 | SEH1L | Other | -1.312966667 | 0.577828007 | 23.5374 | 21.9882 | 24.2645 | 21.8711 | 20.6847 | 23.2954 |
| Selenoprotein M | Q8WWX9 | SELM | Extracellular or secreted | 0.506766667 | 0.213329357 | 21.6145 | 21.4376 | 22.6669 | 24.0031 | 21.8531 | 21.3831 |
| Semaphorin-3A | Q14563 | SEMA3A | Extracellular or secreted | 0.404633333 | 0.149744314 | 22.3794 | 24.9096 | 21.9945 | 23.7628 | 23.8282 | 22.9064 |
| Semaphorin-3B | Q13214-2;Q13214 | SEMA3B | Extracellular or secreted | 0.721733333 | 0.500031785 | 26.9618 | 28.2599 | 26.6082 | 27.4029 | 27.9213 | 28.6709 |
| Semaphorin-3C | Q99985 | SEMA3C | Extracellular or secreted | 0.698766667 | 0.234057093 | 24.5274 | 26.0394 | 23.3092 | 23.6303 | 25.8581 | 26.4839 |
| Semaphorin-3F | Q13275-2;Q13275 | SEMA3F | Extracellular or secreted | -2.706133333 | 2.358923453 | 25.9003 | 26.8615 | 26.72 | 23.7209 | 23.5732 | 24.0693 |
| Semaphorin-4B | Q9NPR2-2;Q9NPR2 | SEMA4B | Extracellular or secreted | -0.3174 | 0.616200235 | 27.2634 | 27.007 | 27.373 | 26.5468 | 26.9342 | 27.2102 |
| Semaphorin-4C | Q9C0C4 | SEMA4C | Extracellular or secreted | 0.797366667 | 1.231447044 | 24.7355 | 24.0547 | 24.0707 | 25.0848 | 25.4372 | 24.731 |
| Semaphorin-7A | O75326;O75326-2 | SEMA7A | Extracellular or secreted | 0.7875 | 0.272660736 | 30.2929 | 31.1545 | 27.8669 | 31.4612 | 30.6688 | 29.5468 |
| Sentrin-specific protease 8 | Q96LD8 | SENP8 | Other | 0.811133333 | 0.506087044 | 20.4691 | 22.2768 | 22.2347 | 21.8764 | 22.7805 | 22.7571 |
| 15 kDa selenoprotein | O60613 | SEP15 | Extracellular or secreted | 0.824766667 | 0.983518909 | 24.8791 | 25.1033 | 25.4469 | 25.3923 | 26.4854 | 26.0259 |
| Selenide, water dikinase 1 | P49903;P49903-2;P49903-4; | SEPHS1 | Extracellular or secreted | -0.938433333 | 2.327140657 | 29.0342 | 28.6339 | 28.7022 | 27.6832 | 28.0103 | 27.8615 |
| Septin-10 | Q9P0V9;Q9P0V9-2;Q9P0V9- | SEPT10 | Other | -0.6673 | 0.261095323 | 25.6672 | 25.7776 | 27.0897 | 23.8975 | 25.7195 | 26.9156 |
| Septin-11 | Q9NVA2;Q9NVA2-2 | SEPT11 | Extracellular or secreted | 0.0478 | 0.035016933 | 28.0014 | 28.1867 | 29.0147 | 28.0585 | 28.1604 | 29.1273 |
| Septin-2 | Q15019;Q15019-3;Q15019-2 | SEPT2 | Extracellular or secreted | 0.161 | 0.108492233 | 28.0106 | 28.3657 | 29.2287 | 28.1251 | 28.5073 | 29.4556 |
| Neuronal-specific septin-3 | Q9UH03-2;Q9UH03 | SEPT3 | Other | 0.1269 | 0.048780445 | 24.055 | 23.7816 | 22.1223 | 22.1748 | 23.8023 | 24.3625 |
| Septin-7 | Q16181-2;Q16181 | SEPT7 | Extracellular or secreted | -0.133033333 | 0.083234859 | 27.9581 | 28.2445 | 28.8015 | 27.399 | 28.1035 | 29.1025 |
| Septin-8 | Q92599-2;Q92599-4;Q92599 | SEPT8 | Other | -0.218066667 | 0.04164514 | 21.7921 | 24.5506 | 24.375 | 20.3915 | 24.2911 | 25.3809 |
| Septin-9 | Q9UHD8;Q9UHD8-7;Q9UHD | SEPT9 | Extracellular or secreted | 0.542433333 | 0.320239313 | 27.3157 | 27.0993 | 28.5018 | 27.1624 | 28.3981 | 28.9836 |
| Plasminogen activator inhibitor 1 RNA-binding protein | Q8NC51-3;Q8NC51-4 | SERBP1 | Extracellular or secreted | 1.5492 | 1.065621689 | 28.8617 | 28.6564 | 30.2524 | 30.9698 | 30.8032 | 30.6451 |
| Small EDRK-rich factor 2 | P84101-4;P84101;P84101-3; | SERF2 | Other | 2.5712 | 0.504711361 | 27.5619 | 21.6747 | 26.9005 | 30.0911 | 27.0642 | 26.6954 |
| Alpha-1-antitrypsin;Short peptide from AAT | P01009;P01009-2;P01009-3 | SERPINA1 | Extracellular or secreted | 3.242233333 | 0.898941078 | 28.236 | 28.2665 | 24.2115 | 30.6719 | 30.5465 | 29.2223 |
| Alpha-1-antichymotrypsin;Alpha-1-antichymotrypsin His | P01011 | SERPINA3 | Extracellular or secreted | 1.083733333 | 0.704890931 | 29.6309 | 30.3816 | 28.5572 | 29.833 | 31.4108 | 30.5771 |
| Plasma serine protease inhibitor | P05154 | SERPINA5 | Extracellular or secreted | 1.555933333 | 0.801344442 | 22.1889 | 20.951 | 19.6999 | 23.2355 | 21.5632 | 22.7089 |
| Leukocyte elastase inhibitor | P30740;P30740-2 | SERPINB1 | Extracellular or secreted | 0.4791 | 0.696699176 | 30.9493 | 31.1427 | 30.3999 | 31.7293 | 31.2187 | 30.9812 |
| Serpin B12 | Q96P63;Q96P63-2 | SERPINB12 | Extracellular or secreted | 2.224133333 | 0.695349621 | 24.2214 | 21.3507 | 25.441 | 26.9406 | 24.781 | 25.9639 |
| Plasminogen activator inhibitor 2 | P05120 | SERPINB2 | Extracellular or secreted | -1.599033333 | 0.895643466 | 31.2426 | 33.1766 | 31.1224 | 30.3035 | 30.6023 | 29.8387 |
| Serpin B3 | P29508;P29508-2 | SERPINB3 | Extracellular or secreted | 0.684566667 | 0.837274255 | 31.1333 | 31.803 | 31.1697 | 31.4542 | 32.389 | 32.3165 |
| Serpin B4 | P48594 | SERPINB4 | Extracellular or secreted | 0.792166667 | 0.98672989 | 29.6604 | 30.1026 | 29.4198 | 29.965 | 30.9959 | 30.5984 |
| Serpin B5 | P36952 | SERPINB5 | Extracellular or secreted | -0.840066667 | 0.823498447 | 31.0232 | 31.5415 | 31.6344 | 29.8667 | 30.6028 | 31.2094 |
| Serpin B6 | P35237 | SERPINB6 | Extracellular or secreted | 0.0063 | 0.005512521 | 31.5027 | 31.4711 | 30.758 | 31.7119 | 31.3067 | 30.7321 |
| Serpin B7 | O75635-2;O75635 | SERPINB7 | Extracellular or secreted | 2.167166667 | 0.6836633 | 29.9364 | 31.2573 | 27.166 | 32.0135 | 32.0367 | 30.811 |
| Serpin B8 | P50452;P50452-2 | SERPINB8 | Extracellular or secreted | 0.645166667 | 0.730093722 | 29.6487 | 29.6949 | 28.7153 | 30.4242 | 29.9691 | 29.6011 |

| | | | | | | | | | | | |
|---|---|---|---|---|---|---|---|---|---|---|---|
| Serpin B9 | P50453 | SERPINB9 | Extracellular or secreted | -0.6277 | 0.800047686 | 24.2603 | 24.5449 | 23.5431 | 23.1717 | 23.6516 | 23.6419 |
| Antithrombin-III | P01008 | SERPINC1 | Extracellular or secreted | -2.0659 | 0.985751736 | 21.6074 | 23.7886 | 21.3517 | 20.9264 | 20.5442 | 19.0794 |
| Plasminogen activator inhibitor 1 | P05121-2;P05121 | SERPINE1 | Extracellular or secreted | -0.864066667 | 0.410728579 | 23.5284 | 22.8523 | 21.8577 | 22.8695 | 20.4536 | 22.3231 |
| Glia-derived nexin | P07093 | SERPINE2 | Extracellular or secreted | 3.1501 | 0.683811965 | 25.1521 | 26.9221 | 21.0295 | 25.8153 | 27.9086 | 28.8301 |
| Glia-derived nexin | P07093-2;P07093-3 | SERPINE2 | Extracellular or secreted | 1.858233333 | 0.770456336 | 29.2684 | 30.7768 | 27.8071 | 30.0467 | 30.9672 | 32.4131 |
| Serpin H1 | P50454 | SERPINH1 | Extracellular or secreted | -0.8761 | 0.439520885 | 29.8976 | 30.1148 | 32.2103 | 29.3238 | 29.8054 | 30.4652 |
| Neuroserpin | Q99574 | SERPINI1 | Extracellular or secreted | 2.6658 | 1.595799852 | 25.6343 | 25.8755 | 24.1694 | 28.2155 | 27.9634 | 27.4977 |
| Protein SET | Q01105 | SET | Other | 0.014133333 | 0.008322255 | 33.4993 | 32.9543 | 32.305 | 33.7684 | 32.7384 | 32.2942 |
| Protein SET | Q01105-2;Q01105-3;Q01105 | SET | Other | 2.2802 | 0.972100299 | 27.5289 | 28.7326 | 25.9134 | 31.1308 | 28.9382 | 28.9465 |
| Histone-lysine N-methyltransferase setd3 | Q86TU7;Q86TU7-3;Q86TU7- | SETD3 | Other | -1.142633333 | 0.303502389 | 25.0115 | 24.1742 | 25.1335 | 20.9213 | 24.5237 | 25.4463 |
| Histone-lysine N-methyltransferase SETD7 | Q8WTS6 | SETD7 | Other | -2.045733333 | 1.721782091 | 23.5181 | 23.2706 | 22.8055 | 20.49 | 21.1118 | 21.8552 |
| Seizure 6-like protein 2 | Q6UXD5-6;Q6UXD5-2;Q6UX | SEZ6L2 | Extracellular or secreted | -0.142933333 | 0.077098711 | 24.8905 | 23.4727 | 23.8682 | 24.9261 | 23.4288 | 23.4477 |
| Splicing factor 1 | Q15637-4;Q15637-6;Q15637 | SF1 | Other | -0.268333333 | 0.272692059 | 28.0787 | 28.4155 | 28.7694 | 27.5445 | 28.2295 | 28.6846 |
| Splicing factor 3A subunit 1 | Q15459;Q15459-2 | SF3A1 | Other | -0.7281 | 0.437540968 | 29.5133 | 29.465 | 30.8516 | 28.2418 | 29.2727 | 30.1311 |
| Splicing factor 3A subunit 2 | Q15428 | SF3A2 | Other | -0.446066667 | 0.249094068 | 26.6849 | 27.0091 | 28.5858 | 26.3778 | 26.8832 | 27.6806 |
| Splicing factor 3A subunit 3 | Q12874 | SF3A3 | Other | -0.216733333 | 0.106873687 | 27.7262 | 27.7752 | 29.3465 | 27.1818 | 28.0952 | 28.9207 |
| Splicing factor 3B subunit 1 | O75533 | SF3B1 | Other | 0.384433333 | 0.154113512 | 27.7762 | 27.2106 | 29.7236 | 27.8428 | 28.4033 | 29.6176 |
| Splicing factor 3B subunit 2 | Q13435 | SF3B2 | Other | -0.3286 | 0.379653612 | 30.0758 | 30.0593 | 30.4176 | 29.3102 | 29.8401 | 30.4166 |
| Splicing factor 3B subunit 3 | Q15393 | SF3B3 | Other | -0.8514 | 1.116012591 | 31.0327 | 30.5053 | 31.2403 | 29.5999 | 30.0666 | 30.5576 |
| Splicing factor 3B subunit 4 | Q15427 | SF3B4 | Other | -0.1391 | 0.433971841 | 26.6538 | 26.4603 | 26.5451 | 26.388 | 26.224 | 26.6299 |
| Splicing factor 3B subunit 5 | Q9BWJ5 | SF3B5 | Other | -0.207266667 | 0.138058147 | 24.6787 | 24.4687 | 25.5943 | 25.1958 | 25.0782 | 23.8459 |
| Splicing factor 3B subunit 6 | Q9Y3B4 | SF3B6 | Other | 0.022333333 | 0.008706075 | 24.4627 | 23.8811 | 26.0932 | 24.8738 | 23.9178 | 25.7124 |
| 14-3-3 protein sigma | P31947;P31947-2 | SFN | Extracellular or secreted | 1.0848 | 1.253675877 | 32.6015 | 32.4939 | 32.5786 | 33.8043 | 34.0127 | 33.1114 |
| Splicing factor, proline- and glutamine-rich | P23246;P23246-2 | SFPQ | Other | -0.249366667 | 0.144046752 | 29.7922 | 30.2537 | 31.4556 | 29.5672 | 30.2142 | 30.972 |
| Secreted frizzled-related protein 1 | Q8N474 | SFRP1 | Extracellular or secreted | -0.068033333 | 0.022839142 | 29.8184 | 30.501 | 27.4498 | 28.8306 | 29.5539 | 29.1806 |
| Vesicle transport protein SFT2B | O95562 | SFT2D2 | Extracellular or secreted | 2.1573 | 0.757211772 | 21.5781 | 21.0232 | 20.379 | 24.4595 | 23.9794 | 21.0133 |
| Epsilon-sarcoglycan | O43556;O43556-3;O43556-4 | SGCE | Extracellular or secreted | -0.911933333 | 0.286111661 | 24.7 | 23.523 | 21.3079 | 23.6618 | 20.8629 | 22.2704 |
| N-sulphoglucosamine sulphohydrolase | P51688 | SGSH | Extracellular or secreted | -0.029566667 | 0.019944484 | 23.3572 | 23.2727 | 23.9977 | 22.6591 | 23.9172 | 23.9626 |
| Small glutamine-rich tetratricopeptide repeat-containing | O43765 | SGTA | Other | -0.358433333 | 0.522987685 | 28.4597 | 28.1554 | 27.5856 | 27.5131 | 27.9369 | 27.6754 |
| SH3 domain-binding glutamic acid-rich-like protein | O75368 | SH3BGRL | Extracellular or secreted | 0.852633333 | 0.649390401 | 29.0476 | 28.9657 | 28.137 | 30.3552 | 29.682 | 28.671 |
| SH3 domain-binding glutamic acid-rich-like protein 3 | Q9H299 | SH3BGRL3 | Extracellular or secreted | 0.629066667 | 0.296368828 | 30.1097 | 30.3682 | 28.7753 | 31.4391 | 30.6289 | 29.0724 |
| Endophilin-A2 | Q99961-2;Q99961;Q99961-3 | SH3GL1 | Other | -0.326866667 | 0.231487705 | 22.4021 | 23.4371 | 23.2529 | 22.1682 | 22.3518 | 23.5915 |
| Endophilin-B1 | Q9Y371;Q9Y371-2;Q9Y371-3 | SH3GLB1 | Extracellular or secreted | 0.602466667 | 0.338407133 | 20.8405 | 21.2005 | 21.3982 | 20.4522 | 22.2064 | 22.588 |
| SH3 domain-containing kinase-binding protein 1 | Q96B97-3;Q96B97-2;Q96B97 | SH3KBP1 | Extracellular or secreted | 1.552933333 | 2.43800203 | 26.3545 | 25.8389 | 26.3696 | 27.4951 | 27.6252 | 28.1015 |
| SHC-transforming protein 1 | P29353-2;P29353-7;P29353; | SHC1 | Extracellular or secreted | 0.239666667 | 0.360996846 | 22.1811 | 22.4065 | 22.8687 | 23.1022 | 22.5505 | 22.5226 |
| Serine hydroxymethyltransferase, cytosolic | P34896-2;P34896;P34896-3; | SHMT1 | Extracellular or secreted | -0.049633333 | 0.067694937 | 27.8611 | 27.297 | 28.0828 | 27.548 | 27.767 | 27.777 |
| Serine hydroxymethyltransferase, mitochondrial | P34897-3;P34897;P34897-2 | SHMT2 | Extracellular or secreted | -1.032766667 | 1.183746037 | 30.2323 | 30.8383 | 31.3342 | 29.8727 | 30.1248 | 29.309 |
| Leucine-rich repeat protein SHOC-2 | Q9UQ13-2;Q9UQ13 | SHOC2 | Other | -0.296433333 | 0.814366889 | 22.5288 | 22.6986 | 22.2352 | 22.1581 | 22.1568 | 22.2584 |
| Sedoheptulokinase | Q9UHJ6 | SHPK | Other | -0.1203 | 0.141791952 | 26.3677 | 26.8339 | 26.5291 | 26.8827 | 26.5429 | 25.9442 |
| Sialate O-acetylesterase | Q9HAT2-2;Q9HAT2 | SIAE | Extracellular or secreted | 0.092266667 | 0.142229765 | 27.0378 | 27.2709 | 27.0116 | 27.5153 | 27.296 | 26.7858 |
| Paired amphipathic helix protein Sin3a | Q96ST3 | SIN3A | Extracellular or secreted | -2.0631 | 0.762135193 | 22.9271 | 23.6044 | 26.2243 | 22.4852 | 21.85 | 22.2313 |
| Tyrosine-protein phosphatase non-receptor type substra | P78324;P78324-2;P78324-4 | SIRPA | Extracellular or secreted | 0.983333333 | 1.534807537 | 27.4887 | 27.5825 | 27.2425 | 28.7697 | 28.4712 | 28.0228 |
| Signal-regulatory protein beta-1 | O00241 | SIRPB1 | Extracellular or secreted | 6.2331 | 1.637579713 | 21.4505 | 22.8139 | 19.0262 | 26.7589 | 27.3644 | 27.8666 |
| NAD-dependent protein deacylase sirtuin-5, mitochondr | Q9NXA8-2;Q9NXA8;Q9NXA8 | SIRT5 | Other | -2.672633333 | 0.848777597 | 25.584 | 25.6511 | 26.8754 | 20.9947 | 24.4984 | 24.5995 |
| Helicase SKI2W | Q15477 | SKIV2L | Other | 0.720066667 | 0.235634393 | 20.4627 | 23.1225 | 23.9495 | 22.6834 | 22.8033 | 24.2082 |
| Superkiller viralicidic activity 2-like 2 | P42285 | SKIV2L2 | Other | -1.499 | 0.88668495 | 27.8332 | 27.9048 | 28.5915 | 25.7017 | 26.2952 | 27.8356 |
| S-phase kinase-associated protein 1 | P63208;P63208-2 | SKP1 | Extracellular or secreted | 0.268033333 | 0.375258224 | 30.0711 | 29.6485 | 29.6918 | 30.4939 | 30.1112 | 29.6104 |
| Solute carrier family 12 member 4;Solute carrier family 1 | Q9UP95-5;Q9UP95-2;Q9UP9 | SLC12A4;SLC12 | Extracellular or secreted | 0.1465 | 0.170202544 | 21.5429 | 22.0067 | 21.1376 | 21.6151 | 22.102 | 21.4096 |
| Monocarboxylate transporter 1 | P53985;P53985-2 | SLC16A1 | Extracellular or secreted | 1.3732 | 1.286173268 | 27.4737 | 26.366 | 26.2343 | 28.4446 | 28.1183 | 27.6307 |
| Monocarboxylate transporter 4 | O15427 | SLC16A3 | Extracellular or secreted | 1.267466667 | 1.270182228 | 29.8945 | 28.6321 | 29.1281 | 30.9462 | 30.4277 | 30.0832 |
| Monocarboxylate transporter 2 | O60669 | SLC16A7 | Extracellular or secreted | 2.653366667 | 1.880336196 | 21.955 | 20.8933 | 21.3253 | 24.0287 | 24.0597 | 24.0453 |
| Neutral amino acid transporter A | P43007;P43007-2 | SLC1A4 | Extracellular or secreted | 1.447866667 | 0.744139454 | 24.7065 | 23.7816 | 22.6684 | 26.24 | 25.3168 | 23.9433 |
| Neutral amino acid transporter B(0) | Q15758;Q15758-3;Q15758-2 | SLC1A5 | Extracellular or secreted | 2.278133333 | 1.607891871 | 27.0529 | 25.6814 | 25.6565 | 29.2115 | 28.3825 | 27.6312 |
| Calcium-binding mitochondrial carrier protein SCaMC-1 | Q6NUK1;Q6NUK1-2 | SLC25A24 | Other | -1.447866667 | 0.663034185 | 23.0789 | 22.8826 | 24.5373 | 23.6065 | 21.6371 | 20.9116 |
| Phosphate carrier protein, mitochondrial | Q00325-2;Q00325 | SLC25A3 | Extracellular or secreted | 2.296133333 | 2.038111398 | 22.9933 | 23.2243 | 22.8338 | 25.8045 | 24.7738 | 25.3615 |
| ADP/ATP translocase 2;ADP/ATP translocase 2, N-termin | P05141;P12236;P12235 | SLC25A5;SLC25 | Extracellular or secreted | 3.296733333 | 0.603822484 | 26.1479 | 22.6969 | 22.1248 | 30.8527 | 24.1749 | 25.8322 |
| Equilibrative nucleoside transporter 1 | Q99808;Q99808-2 | SLC29A1 | Extracellular or secreted | 0.9802 | 0.921118386 | 25.1042 | 24.0325 | 23.8484 | 25.8028 | 25.3134 | 24.8095 |
| Equilibrative nucleoside transporter 2 | Q14542-3;Q14542 | SLC29A2 | Extracellular or secreted | 1.8369 | 0.966889876 | 21.8768 | 22.2596 | 21.6334 | 24.6425 | 24.2335 | 22.4045 |
| Solute carrier family 2, facilitated glucose transporter me | P11166 | SLC2A1 | Extracellular or secreted | 3.128666667 | 3.3340098 | 25.1325 | 25.0595 | 25.7484 | 28.7205 | 28.4961 | 28.1098 |
| Zinc transporter 1 | Q9Y6M5 | SLC30A1 | Extracellular or secreted | 2.014866667 | 0.790340083 | 24.4741 | 21.4796 | 21.6759 | 25.2767 | 24.3054 | 24.0921 |
| Solute carrier family 35 member F6 | Q8N357 | SLC35F6 | Extracellular or secreted | 1.7517 | 1.525082502 | 21.4657 | 22.3667 | 21.9082 | 23.6096 | 24.4008 | 22.9853 |
| Sodium-coupled neutral amino acid transporter 2 | Q96QD8 | SLC38A2 | Extracellular or secreted | 2.527066667 | 1.43336477 | 24.2513 | 24.3365 | 22.5477 | 26.6063 | 25.8727 | 26.2377 |
| Zinc transporter ZIP10 | Q9ULF5 | SLC39A10 | Extracellular or secreted | 1.0875 | 1.040697947 | 24.6534 | 23.4958 | 24.285 | 25.419 | 24.5503 | 25.7274 |
| Zinc transporter ZIP6 | Q13433 | SLC39A6 | Extracellular or secreted | 1.4615 | 0.845169494 | 23.6026 | 21.7592 | 23.0576 | 25.4412 | 23.6971 | 23.6656 |
| Zinc transporter ZIP8 | Q9C0K1-2;Q9C0K1-3;Q9C0K1 | SLC39A8 | Extracellular or secreted | 3.465333333 | 1.065062902 | 21.0377 | 21.4831 | 24.8172 | 26.0919 | 26.6024 | 25.0397 |
| 4F2 cell-surface antigen heavy chain | P08195-2;P08195-3;P08195; | SLC3A2 | Extracellular or secreted | -0.1352 | 0.128574268 | 31.7276 | 31.3324 | 30.872 | 31.2982 | 31.6143 | 30.6139 |
| Solute carrier family 43 member 3 | Q8NBI5;Q8NBI5-2 | SLC43A3 | Other | 2.3452 | 1.628451069 | 25.001 | 24.1768 | 23.3026 | 26.8707 | 26.6455 | 25.9998 |
| Choline transporter-like protein 1 | Q8WWI5-3;Q8WWI5-2;Q8W | SLC44A1 | Extracellular or secreted | 0.715566667 | 0.710403268 | 25.2388 | 25.8851 | 24.5567 | 26.0799 | 26.0939 | 25.6535 |
| Choline transporter-like protein 2 | Q8IWA5;Q8IWA5-3;Q8IWA5 | SLC44A2 | Extracellular or secreted | 1.0852 | 1.318886642 | 27.2344 | 26.9038 | 26.4063 | 28.4603 | 27.8931 | 27.4467 |
| Kanadaptin | Q9BWU0 | SLC4A1AP | Extracellular or secreted | 0.4749 | 0.136010566 | 23.6664 | 20.6539 | 23.2535 | 23.9398 | 21.2462 | 23.8125 |
| High affinity cationic amino acid transporter 1 | P30825 | SLC7A1 | Extracellular or secreted | 1.599933333 | 1.249451091 | 25.465 | 24.5505 | 23.8446 | 26.0134 | 26.8856 | 25.7609 |
| Large neutral amino acids transporter small subunit 1 | Q01650 | SLC7A5 | Extracellular or secreted | 0.555566667 | 0.59695518 | 27.9549 | 26.806 | 27.1903 | 28.0777 | 28.1193 | 27.4209 |
| Na(+)/H(+) exchange regulatory cofactor NHE-RF1 | O14745;O14745-2 | SLC9A3R1 | Extracellular or secreted | -0.321566667 | 0.286755774 | 26.527 | 26.2086 | 26.7292 | 25.6472 | 26.9572 | 25.8957 |
| Na(+)/H(+) exchange regulatory cofactor NHE-RF2 | Q15599;Q15599-2;Q15599-3 | SLC9A3R2 | Extracellular or secreted | 0.6023 | 0.871309599 | 24.1618 | 24.839 | 24.3521 | 24.7045 | 24.9255 | 25.5298 |
| SRA stem-loop-interacting RNA-binding protein, mitocho | Q9GZT3-2;Q9GZT3 | SLIRP | Other | -1.346466667 | 0.308655689 | 22.2078 | 22.5115 | 27.1323 | 22.9913 | 21.6968 | 23.1241 |
| Slit homolog 1 protein | O75093-2;O75093 | SLIT1 | Extracellular or secreted | 1.630966667 | 0.209933767 | 21.4612 | 26.9139 | 22.5957 | 27.9729 | 27.5349 | 20.3559 |
| Slit homolog 2 protein;Slit homolog 2 protein N-product; | O94813-3;O94813-2;O94813 | SLIT2 | Extracellular or secreted | 1.077766667 | 0.853294506 | 24.9515 | 24.2819 | 23.3019 | 25.0953 | 24.8956 | 25.7777 |
| STE20-like serine/threonine-protein kinase | Q9H2G2-2;Q9H2G2 | SLK | Extracellular or secreted | -0.786966667 | 0.513771125 | 25.4327 | 25.1109 | 26.6258 | 24.2786 | 24.6398 | 25.8901 |
| Antileukoproteinase | P03973 | SLPI | Extracellular or secreted | 1.2299 | 0.751486805 | 31.8061 | 33.0867 | 30.9797 | 33.4752 | 33.0423 | 33.0447 |
| SAFB-like transcription modulator | Q9NWH9 | SLTM | Extracellular or secreted | -1.185433333 | 0.52084751 | 24.3028 | 23.8149 | 24.4625 | 23.9461 | 23.7921 | 21.2857 |
| Small acidic protein | O00193 | SMAP | Extracellular or secreted | 1.290933333 | 1.375816996 | 25.191 | 24.6635 | 25.8412 | 26.9208 | 26.3105 | 26.3372 |
| Transcription activator BRG1 | P51532-5;P51532-2;P51532- | SMARCA4 | Extracellular or secreted | -2.1906 | 0.874524268 | 27.4937 | 26.5633 | 27.8913 | 23.758 | 24.6732 | 26.9453 |
| SWI/SNF-related matrix-associated actin-dependent regu | O60264 | SMARCA5 | Other | -0.390533333 | 0.287921265 | 26.9612 | 26.2238 | 27.2883 | 26.4851 | 25.6445 | 27.1721 |
| SWI/SNF-related matrix-associated actin-dependent regu | Q12824-2;Q12824 | SMARCB1 | Other | -2.543666667 | 0.863432444 | 24.3004 | 25.3069 | 27.0318 | 22.3243 | 21.5681 | 25.1157 |
| SWI/SNF complex subunit SMARCC1 | Q92922 | SMARCC1 | Other | -2.379866667 | 0.797468466 | 28.3283 | 27.9346 | 29.3434 | 23.9135 | 27.0183 | 27.5349 |
| SWI/SNF complex subunit SMARCC2 | Q8TAQ2;Q8TAQ2-2;Q8TAQ2 | SMARCC2 | Other | -1.090366667 | 1.045842414 | 27.1451 | 26.9308 | 27.2525 | 25.4232 | 25.9391 | 26.695 |
| SWI/SNF-related matrix-associated actin-dependent regu | Q96GM5-2;Q96GM5 | SMARCD1 | Other | -1.5814 | 0.884821701 | 23.6841 | 23.2101 | 24.7304 | 23.5533 | 22.04 | 21.2871 |
| SWI/SNF-related matrix-associated actin-dependent regu | Q92925-3;Q92925-2;Q92925 | SMARCD2 | Other | -1.7389 | 0.748982096 | 25.8719 | 25.8753 | 28.0097 | 23.4846 | 24.8331 | 26.2225 |
| SWI/SNF-related matrix-associated actin-dependent regu | Q969G3-2;Q969G3;Q969G3- | SMARCE1 | Other | -0.850633333 | 0.647391939 | 26.0353 | 25.59 | 27.2537 | 25.4285 | 25.6549 | 25.2437 |
| Structural maintenance of chromosomes protein 1A | Q14683 | SMC1A | Other | -2.014866667 | 1.166514913 | 29.0923 | 28.8178 | 30.6467 | 26.5432 | 27.4247 | 28.5443 |
| Structural maintenance of chromosomes protein 1B | Q8NDV3-2;Q8NDV3-3;Q8ND | SMC1B | Other | -2.282533333 | 1.322270733 | 23.5606 | 23.0769 | 25.0044 | 22.7129 | 21.1835 | 20.8979 |
| Structural maintenance of chromosomes protein 2 | O95347;O95347-2 | SMC2 | Extracellular or secreted | -0.911566667 | 0.489344756 | 25.8057 | 25.2845 | 27.3917 | 24.5519 | 24.9854 | 26.2099 |
| Structural maintenance of chromosomes protein 3 | Q9UQE7 | SMC3 | Extracellular or secreted | -3.046633333 | 1.174158734 | 28.9827 | 28.5038 | 30.5235 | 25.1812 | 25.474 | 28.2149 |
| Structural maintenance of chromosomes protein 4 | Q9NTJ3;Q9NTJ3-2 | SMC4 | Other | -1.363033333 | 0.653469329 | 29.0572 | 26.8406 | 28.7453 | 26.8113 | 25.7651 | 27.9776 |
| Serine/threonine-protein phosphatase 4 regulatory subu | Q6IN85-2;Q6IN85;Q6IN85-5; | SMEK1 | Other | -0.7195 | 0.294225435 | 24.0438 | 25.5035 | 26.2522 | 23.263 | 24.529 | 25.849 |
| Serine/threonine-protein phosphatase 4 regulatory subu | Q5MIZ7-3;Q5MIZ7-2;Q5MIZ | SMEK2 | Other | -0.230266667 | 0.125755132 | 22.9523 | 22.5998 | 22.7507 | 23.7767 | 21.8103 | 22.025 |
| SPARC-related modular calcium-binding protein 1 | Q9H4F8;Q9H4F8-2 | SMOC1 | Extracellular or secreted | 0.796066667 | 0.367650117 | 31.5367 | 31.4827 | 29.0873 | 31.4802 | 31.7514 | 31.2633 |
| Spermine oxidase | Q9NWM0-5;Q9NWM0;Q9NV | SMOX | Other | 0.586766667 | 0.37567385 | 20.7406 | 21.8048 | 21.9328 | 22.1513 | 21.1346 | 22.9526 |
| Sphingomyelin phosphodiesterase | P17405-3;P17405-2;P17405- | SMPD1 | Extracellular or secreted | 0.793066667 | 0.72338415 | 21.6934 | 22.9712 | 22.1861 | 22.4102 | 23.467 | 23.3527 |
| Acid sphingomyelinase-like phosphodiesterase 3b | Q92485;Q92485-2 | SMPDL3B | Extracellular or secreted | 0.0525 | 0.051575617 | 26.8859 | 27.1677 | 27.1521 | 27.0416 | 27.7145 | 26.6071 |
| Spermine synthase | P52788;P52788-2 | SMS | Extracellular or secreted | 0.3503 | 0.491206034 | 28.7748 | 28.5502 | 28.0935 | 29.1007 | 29.015 | 28.3537 |
| WD40 repeat-containing protein SMU1;WD40 repeat-co | Q2TAY7;Q2TAY7-2 | SMU1 | Other | -3.834466667 | 0.860981262 | 28.3925 | 28.1735 | 28.5791 | 21.5332 | 25.1347 | 26.9738 |
| SET and MYND domain-containing protein 5 | Q6GMV2 | SMYD5 | Other | -2.303766667 | 0.868601173 | 23.9483 | 24.3072 | 24.3391 | 20.7859 | 23.7984 | 21.099 |
| Synaptosomal-associated protein 23 | O00161;O00161-2 | SNAP23 | Extracellular or secreted | 3.483 | 1.698606689 | 23.3794 | 21.3626 | 23.4394 | 26.9028 | 25.7248 | 26.0028 |
| SNARE-associated protein Snapin | O95295 | SNAPIN | Other | 0.4347 | 0.266010371 | 21.5506 | 22.4863 | 22.7083 | 21.8367 | 22.5325 | 23.6801 |
| Staphylococcal nuclease domain-containing protein 1 | Q7KZF4 | SND1 | Extracellular or secreted | -0.173833333 | 0.108763104 | 29.8614 | 30.1486 | 30.812 | 29.418 | 29.8244 | 31.0581 |
| U5 small nuclear ribonucleoprotein 200 kDa helicase | O75643 | SNRNP200 | Other | -1.076966667 | 0.575265716 | 29.3279 | 28.7487 | 30.5377 | 27.3498 | 28.4687 | 29.5649 |
| U5 small nuclear ribonucleoprotein 40 kDa protein | Q96DI7;Q96DI7-2 | SNRNP40 | Other | -1.810033333 | 0.420436218 | 25.3628 | 25.2611 | 27.2817 | 21.069 | 25.0184 | 26.3881 |
| U1 small nuclear ribonucleoprotein 70 kDa | P08621;P08621-2;P08621-3 | SNRNP70 | Other | 1.8437 | 1.237576244 | 26.6196 | 26.2779 | 28.077 | 28.1862 | 29.0052 | 29.3142 |
| U1 small nuclear ribonucleoprotein A | P09012 | SNRPA | Other | -0.399666667 | 0.553789884 | 28.8296 | 28.7278 | 28.9426 | 27.9178 | 28.5334 | 28.8498 |
| U2 small nuclear ribonucleoprotein A | P09661 | SNRPA1 | Other | 1.127933333 | 0.436024611 | 24.885 | 25.4568 | 27.7255 | 25.8361 | 27.5256 | 28.0894 |
| Small nuclear ribonucleoprotein-associated proteins B ar | P14678-2;P14678;P14678-3; | SNRPB;SNRPN | Extracellular or secreted | -0.6947 | 0.656056815 | 29.4158 | 28.6478 | 29.1057 | 27.5667 | 28.7646 | 28.7539 |
| U2 small nuclear ribonucleoprotein B | P08579 | SNRPB2 | Other | 0.803833333 | 0.526075177 | 26.8169 | 25.6188 | 25.8373 | 27.3938 | 25.8046 | 27.4861 |
| U1 small nuclear ribonucleoprotein C | P09234 | SNRPC | Other | 0.459033333 | 0.326132199 | 24.5628 | 24.7227 | 24.9954 | 24.2056 | 25.5374 | 25.915 |
| Small nuclear ribonucleoprotein Sm D1 | P62314 | SNRPD1 | Other | -1.2067 | 0.608594788 | 30.4552 | 28.5307 | 29.3935 | 26.9696 | 29.3115 | 28.4782 |
| Small nuclear ribonucleoprotein Sm D2 | P62316;P62316-2 | SNRPD2 | Extracellular or secreted | -0.22 | 0.230634709 | 30.3823 | 29.9905 | 30.5767 | 29.5908 | 30.6926 | 30.0061 |
| Small nuclear ribonucleoprotein Sm D3 | P62318-2;P62318 | SNRPD3 | Extracellular or secreted | -0.632333333 | 0.440652117 | 30.0354 | 29.5703 | 30.045 | 28.3367 | 30.2038 | 29.2132 |
| Small nuclear ribonucleoprotein E | P62304 | SNRPE | Extracellular or secreted | -0.397366667 | 0.28498173 | 28.0407 | 27.5726 | 27.7564 | 26.3809 | 27.9358 | 27.8609 |
| Small nuclear ribonucleoprotein F | P62306 | SNRPF | Other | -0.708966667 | 0.458927518 | 28.3554 | 27.9607 | 28.0987 | 26.2993 | 28.2432 | 27.7454 |

| | | | | | | | | | | | |
|---|---|---|---|---|---|---|---|---|---|---|---|
| Small nuclear ribonucleoprotein G;Putative small nuclear | P62308;A8MWD9 | SNRPG;SNRPG | Other | -1.2598 | 0.584412152 | 27.0279 | 26.2934 | 26.9492 | 23.8686 | 26.4638 | 26.1587 |
| Beta-2-syntrophin | Q13425;Q13425-2 | SNTB2 | Extracellular or secreted | 0.485166667 | 0.24263687 | 21.7306 | 20.9091 | 22.4124 | 20.8757 | 22.7731 | 22.8588 |
| SNW domain-containing protein 1 | Q13573 | SNW1 | Other | -0.192866667 | 0.292787722 | 23.6039 | 24.1971 | 24.3969 | 23.7288 | 23.8907 | 23.9998 |
| Sorting nexin-1 | Q13596;Q13596-2;Q13596-3 | SNX1 | Other | 0.073166667 | 0.030506779 | 24.793 | 24.845 | 25.7591 | 24.0936 | 24.9695 | 26.5535 |
| Sorting nexin-12 | Q9UMY4-2;Q9UMY4;Q9UMY | SNX12 | Extracellular or secreted | -0.2432 | 0.331248399 | 25.6956 | 26.137 | 26.5802 | 25.7127 | 25.7846 | 26.1859 |
| Sorting nexin-2 | O60749-2;O60749 | SNX2 | Extracellular or secreted | -0.623566667 | 0.360462435 | 25.7183 | 26.6619 | 27.3175 | 24.9025 | 26.1555 | 26.769 |
| Sorting nexin-25 | Q9H3E2;Q9H3E2-2 | SNX25 | Other | 1.646766667 | 1.464923372 | 26.8873 | 26.5777 | 25.8899 | 28.3697 | 28.6211 | 27.3044 |
| Sorting nexin-5 | Q9Y5X3 | SNX5 | Extracellular or secreted | -0.618566667 | 0.367593608 | 23.4665 | 23.5204 | 23.71 | 21.7027 | 23.4171 | 23.7214 |
| Sorting nexin-6;Sorting nexin-6, N-terminally processed | Q9UNH7-2;Q9UNH7 | SNX6 | Other | -1.040533333 | 0.585367268 | 24.9596 | 25.4966 | 26.0025 | 23.3547 | 24.3099 | 25.6725 |
| Sorting nexin-9 | Q9Y5X1 | SNX9 | Extracellular or secreted | -0.469333333 | 0.599421421 | 28.2417 | 27.9583 | 28.7558 | 27.4122 | 27.8223 | 28.3133 |
| Superoxide dismutase [Cu-Zn] | P00441 | SOD1 | Extracellular or secreted | -0.513366667 | 0.361562192 | 33.6749 | 33.2699 | 33.0331 | 33.7071 | 32.832 | 31.8987 |
| Superoxide dismutase [Mn], mitochondrial | P04179;P04179-4;P04179-2; | SOD2 | Extracellular or secreted | 0.495033333 | 0.368379002 | 34.1554 | 33.7775 | 33.3907 | 34.7505 | 34.7644 | 33.2938 |
| Protein SON | P18583-6;P18583-2;P18583- | SON | Other | -0.799733333 | 0.272908037 | 23.2671 | 20.3067 | 23.6649 | 22.1858 | 21.3713 | 21.2824 |
| Vinexin | O60504;O60504-2 | SORBS3 | Other | -2.549233333 | 0.587191396 | 26.2142 | 26.2427 | 26.5805 | 25.8234 | 25.0031 | 20.5632 |
| Sorbitol dehydrogenase | Q00796 | SORD | Extracellular or secreted | -0.354433333 | 0.4153399 | 30.6288 | 30.7822 | 29.7549 | 29.9453 | 30.1998 | 29.9575 |
| Transcription factor Sp1 | P08047-3;P08047;P08047-2 | SP1 | Other | 0.133266667 | 0.035352847 | 23.5275 | 20.7094 | 23.955 | 23.5331 | 21.3517 | 23.7069 |
| Nuclear autoantigen Sp-100 | P23497;P23497-7;P23497-6; | SP100 | Other | 0.3895 | 0.153500547 | 22.8179 | 20.1803 | 22.4558 | 22.4353 | 21.4007 | 22.7865 |
| Transcription factor Sp3 | Q02447-4;Q02447-3;Q02447 | SP3 | Extracellular or secreted | -1.5151 | 0.612232867 | 22.4966 | 22.2733 | 24.5722 | 21.4515 | 20.2385 | 23.1068 |
| Sperm-associated antigen 7 | O75391 | SPAG7 | Other | 2.984766667 | 1.607012362 | 20.1904 | 21.6135 | 20.8168 | 22.5772 | 24.7155 | 24.2823 |
| C-Jun-amino-terminal kinase-interacting protein 4 | O60271-4;O60271-2;O60271 | SPAG9 | Extracellular or secreted | -0.4606 | 0.525313955 | 26.6396 | 26.0284 | 26.9559 | 25.6919 | 25.9427 | 26.6075 |
| SPARC | P09486 | SPARC | Extracellular or secreted | 0.243 | 0.200344733 | 24.4346 | 25.1853 | 24.7868 | 24.2957 | 25.1704 | 25.6696 |
| Signal peptidase complex subunit 2 | Q15005 | SPCS2 | Other | -0.357 | 0.103755197 | 19.9409 | 23.4814 | 23.2132 | 21.8023 | 22.4204 | 21.3418 |
| Protein Spindly | Q96EA4-2;Q96EA4-3;Q96EA | SPDL1 | Other | 0.9766 | 0.288882122 | 22.0915 | 20.2005 | 23.1665 | 24.8105 | 22.3018 | 21.276 |
| Cytospin-A | Q69YQ0-2;Q69YQ0 | SPECC1L | Other | 0.328866667 | 0.469683114 | 21.153 | 21.0468 | 21.8053 | 21.7659 | 21.9212 | 21.3046 |
| Msx2-interacting protein | Q96T58 | SPEN | Extracellular or secreted | 2.402166667 | 0.835845217 | 22.2838 | 22.6331 | 23.2828 | 27.2022 | 24.5797 | 23.6243 |
| Spartin | Q8N0X7 | SPG20 | Extracellular or secreted | -0.239666667 | 0.832452133 | 24.9611 | 24.6474 | 24.6271 | 24.5206 | 24.5575 | 24.4385 |
| Spindlin-1;Spindlin-3 | Q9Y657;Q5JUX0 | SPIN1;SPIN3 | Other | -1.674566667 | 0.71969039 | 24.3939 | 24.975 | 25.7945 | 21.744 | 23.6069 | 24.7888 |
| Serine protease inhibitor Kazal-type 6 | Q6UWN8 | SPINK6 | Extracellular or secreted | 6.9582 | 1.023614869 | 21.4735 | 26.8756 | 18.4115 | 30.3119 | 27.6188 | 29.7045 |
| Kunitz-type protease inhibitor 1 | O43278 | SPINT1 | Extracellular or secreted | 1.164833333 | 1.596490916 | 31.4905 | 31.1219 | 30.6909 | 32.5824 | 32.4238 | 31.7916 |
| Kunitz-type protease inhibitor 1 | O43278-2 | SPINT1 | Extracellular or secreted | 1.399066667 | 1.254354391 | 26.1377 | 25.7128 | 25.1703 | 27.5177 | 27.4488 | 26.2515 |
| Testican-2 | Q92563 | SPOCK2 | Extracellular or secreted | 1.323366667 | 0.748149493 | 26.5779 | 24.3228 | 24.987 | 26.292 | 27.3634 | 26.2024 |
| Signal peptide peptidase-like 2A | Q8TCT8 | SPPL2A | Extracellular or secreted | 1.342033333 | 0.36098801 | 24.2879 | 24.3226 | 23.1532 | 26.32 | 26.9387 | 22.5311 |
| Sepiapterin reductase | P35270 | SPR | Extracellular or secreted | -1.4072 | 1.223082493 | 25.1543 | 25.6089 | 26.1116 | 23.3669 | 24.6895 | 24.5968 |
| Small proline-rich protein 2F;Small proline-rich protein 2 | Q96RM1;P35326;P35325;P2 | SPRR2F;SPRR2 | Extracellular or secreted | 2.405133333 | 1.91614222 | 22.1244 | 20.7316 | 21.577 | 23.327 | 23.7326 | 24.5888 |
| SPRY domain-containing protein 4 | Q8WW59 | SPRYD4 | Other | -1.078533333 | 1.584481537 | 26.6623 | 26.5271 | 27.3082 | 25.664 | 26.0857 | 25.5123 |
| Spectrin alpha chain, non-erythrocytic 1 | Q13813;Q13813-3 | SPTAN1 | Extracellular or secreted | -0.877 | 1.888274516 | 33.8882 | 33.3923 | 33.7725 | 32.6179 | 32.7248 | 33.0793 |
| Spectrin alpha chain, non-erythrocytic 1 | Q13813-2 | SPTAN1 | Extracellular or secreted | -0.355833333 | 0.166332375 | 23.2708 | 23.3664 | 21.2772 | 23.0922 | 21.815 | 21.9397 |
| Spectrin beta chain, non-erythrocytic 1 | Q01082;Q01082-2 | SPTBN1 | Extracellular or secreted | -0.816066667 | 1.030335244 | 32.8066 | 32.635 | 33.4197 | 31.7532 | 31.9734 | 32.6865 |
| Spectrin beta chain, non-erythrocytic 2 | O15020;O15020-2 | SPTBN2 | Extracellular or secreted | -1.724333333 | 0.695026478 | 24.8385 | 25.6729 | 27.6992 | 23.1643 | 24.1661 | 25.7072 |
| Sulfide:quinone oxidoreductase, mitochondrial | Q9Y6N5 | SQRDL | Other | 1.4835 | 0.733699476 | 21.3754 | 22.9678 | 23.8192 | 24.3537 | 25.1148 | 23.1444 |
| Sequestosome-1 | Q13501;Q13501-2 | SQSTM1 | Extracellular or secreted | 2.9355 | 1.429047187 | 22.4268 | 21.4 | 23.9306 | 24.7314 | 25.2074 | 26.6251 |
| Steroid receptor RNA activator 1 | Q9HD15 | SRA1 | Extracellular or secreted | 0.552166667 | 0.349055853 | 22.7854 | 23.4541 | 21.8121 | 22.3533 | 23.8387 | 23.5161 |
| Proto-oncogene tyrosine-protein kinase Src | P12931;P12931-2 | SRC | Extracellular or secreted | 2.000766667 | 0.45142607 | 26.1551 | 21.3043 | 26.4609 | 26.628 | 26.573 | 26.7216 |
| Splicing regulatory glutamine/lysine-rich protein 1 | Q8WXA9;Q8WXA9-2 | SREK1 | Other | -0.795333333 | 0.799188036 | 22.1608 | 22.2492 | 22.8835 | 21.9128 | 22.1001 | 20.8946 |
| SLIT-ROBO Rho GTPase-activating protein 1 | Q7Z6B7-2;Q7Z6B7 | SRGAP1 | Other | 0.2682 | 0.136680482 | 21.2306 | 22.0168 | 20.7901 | 22.8211 | 20.8025 | 21.2185 |
| Serglycin | P10124 | SRGN | Extracellular or secreted | 2.462666667 | 1.312413831 | 21.2303 | 21.633 | 20.7146 | 24.9458 | 22.8114 | 23.2087 |
| Sorcin | P30626-2;P30626;P30626-3 | SRI | Extracellular or secreted | -0.0256 | 0.026281397 | 29.3389 | 29.9066 | 29.1224 | 29.0098 | 29.793 | 29.4883 |
| Spermidine synthase | P19623 | SRM | Other | -0.198 | 0.330918745 | 28.0931 | 28.2623 | 28.5125 | 27.6765 | 28.2978 | 28.2996 |
| Signal recognition particle 14 kDa protein | P37108 | SRP14 | Extracellular or secreted | 0.075466667 | 0.051380501 | 27.6407 | 27.2388 | 28.0419 | 26.9033 | 27.841 | 28.4035 |
| Signal recognition particle 19 kDa protein | P09132 | SRP19 | Other | 1.8656 | 1.172913645 | 23.465 | 22.4699 | 24.4292 | 25.2631 | 25.7007 | 24.9971 |
| Signal recognition particle subunit SRP68 | Q9UHB9;Q9UHB9-4;Q9UHB9 | SRP68 | Other | -0.740033333 | 0.172680923 | 23.3785 | 21.6046 | 24.9574 | 20.5488 | 22.2017 | 24.9699 |
| Signal recognition particle subunit SRP72 | O76094;O76094-2 | SRP72 | Extracellular or secreted | -0.309633333 | 0.132290774 | 24.2985 | 22.9383 | 25.2642 | 22.8629 | 24.0362 | 24.673 |
| Signal recognition particle 9 kDa protein | P49458;P49458-2 | SRP9 | Extracellular or secreted | 0.136533333 | 0.203138698 | 27.495 | 27.288 | 27.5439 | 27.2046 | 28.0089 | 27.523 |
| SRSF protein kinase 1 | Q96SB4-4;Q96SB4;Q96SB4-3 | SRPK1 | Extracellular or secreted | -0.751833333 | 0.478133102 | 25.1413 | 24.2274 | 26.1268 | 23.9215 | 24.136 | 25.1825 |
| Signal recognition particle receptor subunit alpha | P08240;P08240-2 | SRPR | Extracellular or secreted | -0.819366667 | 0.212639375 | 25.8499 | 25.4356 | 28.1048 | 27.7741 | 23.5328 | 25.6253 |
| Sushi repeat-containing protein SRPX | P78539-4;P78539-3;P78539- | SRPX | Extracellular or secreted | 1.6422 | 1.200158381 | 23.6249 | 22.8483 | 22.9197 | 23.9018 | 24.7749 | 25.6428 |
| Sushi repeat-containing protein SRPX2 | O60687 | SRPX2 | Extracellular or secreted | 4.023 | 0.736735821 | 26.2959 | 26.9898 | 20.8014 | 28.5355 | 29.0009 | 29.0197 |
| Serine/arginine repetitive matrix protein 1 | Q8IYB3-2;Q8IYB3 | SRRM1 | Other | -1.1519 | 1.170275243 | 28.6737 | 27.5539 | 28.2356 | 26.995 | 26.9093 | 27.1032 |
| Serine/arginine repetitive matrix protein 2 | Q9UQ35;Q9UQ35-2 | SRRM2 | Other | -0.153366667 | 0.106451196 | 27.8827 | 26.2322 | 27.3455 | 27.0064 | 26.9241 | 27.0698 |
| Serrate RNA effector molecule homolog | Q9BXP5-5;Q9BXP5-4;Q9BXP | SRRT | Other | -0.137 | 0.059095547 | 26.5307 | 25.4705 | 27.9612 | 26.6352 | 25.9692 | 26.947 |
| Serine/arginine-rich splicing factor 1 | Q07955;Q07955-3;Q07955-2 | SRSF1 | Extracellular or secreted | 1.3426 | 1.124110197 | 29.0547 | 28.6967 | 30.1803 | 31.0122 | 30.812 | 30.1353 |
| Serine/arginine-rich splicing factor 10 | O75494-5;O75494-4;O75494 | SRSF10 | Other | -1.391433333 | 0.783250105 | 22.4354 | 22.128 | 24.2869 | 21.8603 | 21.763 | 21.0527 |
| Serine/arginine-rich splicing factor 11 | Q05519-2;Q05519 | SRSF11 | Other | -0.1067 | 0.072055659 | 25.5657 | 25.9378 | 26.6238 | 25.2798 | 25.829 | 26.6984 |
| Serine/arginine-rich splicing factor 2 | Q01130-2;Q01130 | SRSF2 | Extracellular or secreted | 1.479 | 1.115809791 | 27.8527 | 27.1784 | 28.885 | 29.9184 | 29.544 | 28.8907 |
| Serine/arginine-rich splicing factor 3 | P84103-2;P84103 | SRSF3 | Other | 0.6401 | 0.554021384 | 25.5137 | 26.1017 | 27.005 | 26.463 | 27.1865 | 26.8912 |
| Serine/arginine-rich splicing factor 4 | Q08170 | SRSF4 | Other | 1.801866667 | 1.017353882 | 25.9001 | 27.3996 | 28.1702 | 29.4023 | 29.1639 | 28.3093 |
| Serine/arginine-rich splicing factor 5 | Q13243-3;Q13243;Q13243-2 | SRSF5 | Other | 0.5921 | 0.165487659 | 23.5957 | 21.2685 | 24.435 | 21.7745 | 24.6939 | 24.6071 |
| Serine/arginine-rich splicing factor 6 | Q13247-3;Q13247;Q13247-2 | SRSF6 | Other | 1.376533333 | 0.503814768 | 26.2296 | 23.281 | 26.5148 | 26.7242 | 27.0403 | 26.3905 |
| Serine/arginine-rich splicing factor 7 | Q16629-3;Q16629-2;Q16629 | SRSF7 | Extracellular or secreted | 0.6111 | 0.487140057 | 26.8923 | 26.7893 | 28.2263 | 27.559 | 28.3724 | 27.8098 |
| Serine/arginine-rich splicing factor 9 | Q13242 | SRSF9 | Other | 0.735366667 | 0.36347272 | 24.8283 | 24.4763 | 26.7396 | 25.2418 | 26.3218 | 26.6867 |
| Lupus La protein | P05455 | SSB | Other | 0.499566667 | 0.853356051 | 30.625 | 30.3102 | 31.0664 | 31.058 | 31.3024 | 31.1399 |
| Single-stranded DNA-binding protein, mitochondrial | Q04837 | SSBP1 | Extracellular or secreted | -1.109066667 | 0.770151689 | 29.4191 | 28.7672 | 29.6834 | 29.0624 | 28.3041 | 27.176 |
| Protein phosphatase Slingshot homolog 3 | Q8TE77-3;Q8TE77;Q8TE77-2 | SSH3 | Other | 0.1947 | 0.046568736 | 21.1402 | 22.2296 | 24.6931 | 21.4251 | 22.5022 | 24.7197 |
| Translocon-associated protein subunit delta | P51571 | SSR4 | Extracellular or secreted | 1.9483 | 1.201877481 | 23.541 | 23.0012 | 21.5505 | 25.5195 | 24.2489 | 24.1692 |
| FACT complex subunit SSRP1 | Q08945 | SSRP1 | Other | -0.864266667 | 0.907390962 | 29.1144 | 29.0625 | 29.6638 | 27.8185 | 28.3625 | 29.0669 |
| Sjoegren syndrome/scleroderma autoantigen 1 | O60232 | SSSCA1 | Other | 1.3514 | 1.078352613 | 24.7382 | 25.9865 | 26.1779 | 26.8633 | 26.8076 | 27.2859 |
| Hsc70-interacting protein;Putative protein FAM10A5 | P50502;Q8NFI4 | ST13;ST13P5 | Extracellular or secreted | -0.532733333 | 1.07796493 | 30.4029 | 30.1234 | 30.126 | 29.3164 | 29.8968 | 29.8409 |
| Suppressor of tumorigenicity 14 protein | Q9Y5Y6 | ST14 | Extracellular or secreted | -0.469333333 | 0.569975226 | 29.1821 | 29.0588 | 28.1928 | 28.3368 | 28.429 | 28.2599 |
| CMP-N-acetylneuraminate-beta-galactosamide-alpha-2,3 | Q11206-3;Q11206-2;Q11206 | ST3GAL4 | Extracellular or secreted | 0.2409 | 0.215532677 | 24.4256 | 24.5206 | 23.9708 | 23.9705 | 24.3974 | 25.2718 |
| Beta-galactoside alpha-2,6-sialyltransferase 1 | P15907 | ST6GAL1 | Extracellular or secreted | -0.785433333 | 0.377284743 | 22.3591 | 22.9919 | 21.0336 | 21.7747 | 20.0575 | 22.1961 |
| CMP-N-acetylneuraminate-poly-alpha-2,8-sialyltransfera | Q92187 | ST8SIA4 | Extracellular or secreted | 1.282833333 | 0.929948119 | 24.8302 | 25.4297 | 23.6787 | 25.2335 | 26.1874 | 26.3662 |
| Cohesin subunit SA-1 | Q8WVM7-2;Q8WVM7 | STAG1 | Other | -2.820066667 | 0.873618363 | 24.7525 | 24.8891 | 28.4695 | 22.4026 | 22.5445 | 24.7038 |
| Cohesin subunit SA-2 | Q8N3U4;Q8N3U4-2 | STAG2 | Extracellular or secreted | -1.3197 | 0.545160791 | 25.4413 | 24.8251 | 27.0133 | 23.0374 | 24.3516 | 25.9316 |
| Signal transducing adapter molecule 1 | Q92783-2;Q92783 | STAM | Other | 0.0189 | 0.016077249 | 25.7919 | 25.9225 | 26.8747 | 25.9293 | 26.1975 | 26.519 |
| STAM-binding protein | O95630;O95630-2 | STAMBP | Extracellular or secreted | -0.226666667 | 1.462355412 | 25.5907 | 25.4154 | 25.5493 | 25.3132 | 25.2451 | 25.3171 |
| PCTP-like protein | Q9Y365 | STARD10 | Other | -1.077366667 | 0.75551091 | 23.8786 | 24.9137 | 25.1692 | 22.57 | 23.9148 | 24.2446 |
| Signal transducer and activator of transcription 1-alpha/b | P42224;P42224-2 | STAT1 | Other | -0.169933333 | 0.063124369 | 25.6451 | 25.5886 | 26.7012 | 24.4408 | 25.6557 | 27.3186 |
| Signal transducer and activator of transcription 3 | P40763-3;P40763-2;P40763 | STAT3 | Extracellular or secreted | 0.9276 | 0.442179127 | 22.5787 | 23.1782 | 24.7875 | 24.8777 | 23.233 | 25.2165 |
| Signal transducer and activator of transcription 6 | P42226-2;P42226-3;P42226 | STAT6 | Other | -0.3409 | 0.197995103 | 21.4962 | 21.8996 | 22.8707 | 22.2291 | 20.7125 | 22.3022 |
| Double-stranded RNA-binding protein Staufen homolog 1 | O95793-2;O95793-3;O95793 | STAU1 | Extracellular or secreted | 1.187833333 | 1.809288737 | 24.2925 | 23.804 | 23.6057 | 24.7861 | 25.4939 | 24.9857 |
| Stanniocalcin-1 | P52823;P52823-2 | STC1 | Extracellular or secreted | 1.605066667 | 0.916837248 | 28.2625 | 29.0951 | 28.0995 | 28.7667 | 30.8079 | 30.6977 |
| Stanniocalcin-2 | O76061 | STC2 | Extracellular or secreted | 0.049566667 | 0.0162806 | 30.1598 | 27.8974 | 27.3404 | 29.2331 | 28.7494 | 27.5638 |
| Metalloreductase STEAP4 | Q687X5;Q687X5-2 | STEAP4 | Extracellular or secreted | 1.664333333 | 2.113647393 | 27.7649 | 26.9922 | 27.5848 | 28.6561 | 29.4737 | 29.2051 |
| Stress-induced-phosphoprotein 1 | P31948;P31948-2;P31948-3 | STIP1 | Other | -0.233966667 | 0.381040063 | 31.9598 | 32.3059 | 31.981 | 31.4505 | 32.2151 | 31.8792 |
| Serine/threonine-protein kinase 24;Serine/threonine-pro | Q9Y6E0-2;Q9Y6E0 | STK24 | Extracellular or secreted | -0.203733333 | 0.296817486 | 25.5189 | 25.4305 | 26.0132 | 25.2382 | 25.2417 | 25.8715 |
| Serine/threonine-protein kinase 26 | Q9P289;Q9P289-2;Q9P289-3 | STK26 | Extracellular or secreted | 1.108466667 | 0.508754719 | 23.8921 | 21.6496 | 24.306 | 24.6974 | 24.0066 | 24.4691 |
| Serine/threonine-protein kinase 3;Serine/threonine-prot | Q13188;Q13188-2 | STK3 | Other | -1.123433333 | 0.316935741 | 23.7573 | 23.8977 | 24.1978 | 20.9973 | 22.1175 | 25.3677 |
| Stathmin | P16949;P16949-2 | STMN1 | Extracellular or secreted | 1.914966667 | 1.30850548 | 27.9834 | 28.1196 | 27.8373 | 29.771 | 30.7456 | 29.1686 |
| Stathmin-2 | Q93045;Q93045-2 | STMN2 | Other | 4.104333333 | 1.09485218 | 21.135 | 25.0985 | 21.0085 | 26.133 | 27.3997 | 26.0223 |
| Stonin-2 | Q8WXE9;Q8WXE9-3 | STON2 | Other | -1.680966667 | 0.837538095 | 24.8574 | 22.9703 | 22.8332 | 23.1762 | 21.4381 | 21.0037 |
| Stimulated by retinoic acid gene 6 protein homolog | Q9BX79-3;Q9BX79;Q9BX79-6 | STRA6 | Extracellular or secreted | 3.0051 | 3.243036617 | 23.8452 | 23.472 | 23.4502 | 26.6468 | 26.2227 | 26.9132 |
| STE20-related kinase adapter protein beta | Q9C0K7-3;Q9C0K7-2;Q9C0K7 | STRADB | Other | 1.029866667 | 0.347912484 | 23.2974 | 23.5613 | 21.5056 | 21.8047 | 25.0752 | 24.574 |
| Serine-threonine kinase receptor-associated protein | Q9Y3F4;Q9Y3F4-2 | STRAP | Other | -0.072333333 | 0.126646759 | 30.1999 | 30.0157 | 30.5484 | 30.0256 | 30.0618 | 30.4596 |
| Spermatid perinuclear RNA-binding protein | Q96SI9-2;Q96SI9 | STRBP | Other | -1.9577 | 1.163950528 | 28.2154 | 27.898 | 28.5508 | 25.1099 | 26.6269 | 27.0543 |
| Striatin-interacting protein 1;Striatin-interacting protein | Q5VSL9-3;Q5VSL9-2;Q5VSL9 | STRIP1;STRIP2 | Extracellular or secreted | -0.200833333 | 0.144631016 | 21.1752 | 22.0983 | 22.8214 | 22.0073 | 21.8019 | 21.6832 |
| Striatin | O43815-2;O43815 | STRN | Extracellular or secreted | -1.2195 | 0.946620536 | 25.3851 | 25.8836 | 26.518 | 23.76 | 25.0599 | 25.3083 |
| Striatin-3 | Q13033-2;Q13033 | STRN3 | Extracellular or secreted | -0.376566667 | 0.396805652 | 23.9635 | 24.7761 | 24.2232 | 23.7473 | 23.5203 | 24.5655 |
| Striatin-4 | Q9NRL3;Q9NRL3-3 | STRN4 | Other | 0.285966667 | 0.19298523 | 22.3991 | 21.2807 | 22.7521 | 22.5113 | 21.7844 | 22.9941 |
| E3 ubiquitin-protein ligase CHIP | Q9UNE7-2;Q9UNE7 | STUB1 | Extracellular or secreted | 0.394966667 | 0.27493602 | 22.5409 | 23.6513 | 24.2944 | 23.5665 | 23.8672 | 24.2378 |
| Syntaxin-10 | O60499;O60499-2 | STX10 | Other | 1.581733333 | 0.69696437 | 23.6648 | 20.8793 | 23.2242 | 24.1845 | 24.7372 | 23.5918 |
| Syntaxin-12 | Q86Y82 | STX12 | Extracellular or secreted | -0.4591 | 0.255451468 | 26.8076 | 27.026 | 27.5729 | 25.7536 | 27.9013 | 26.3743 |
| Syntaxin-3 | Q13277-2;Q13277-3;Q13277 | STX3 | Extracellular or secreted | 0.664733333 | 0.881970106 | 22.6052 | 21.6585 | 22.1373 | 22.8708 | 22.7021 | 22.8223 |
| Syntaxin-4 | Q12846;Q12846-2 | STX4 | Extracellular or secreted | 3.3521 | 1.374738022 | 21.2755 | 20.6192 | 23.5533 | 25.9377 | 25.5378 | 24.0288 |
| Syntaxin-6 | O43752 | STX6 | Extracellular or secreted | 0.586166667 | 0.313439607 | 25.6653 | 23.6253 | 25.6564 | 25.5907 | 25.9732 | 25.1416 |
| Syntaxin-7 | O15400-2;O15400 | STX7 | Extracellular or secreted | 0.641666667 | 0.940373028 | 26.9574 | 26.7552 | 26.1903 | 27.2736 | 27.6609 | 26.8934 |
| Syntaxin-8 | Q9UNK0 | STX8 | Extracellular or secreted | 0.177133333 | 0.055709201 | 23.353 | 20.0463 | 22.8233 | 22.3959 | 21.9168 | 22.4413 |
| Syntaxin-binding protein 2 | Q15833-2;Q15833;Q15833-3 | STXBP2 | Extracellular or secreted | -1.305433333 | 0.694329575 | 25.2848 | 26.314 | 27.3731 | 23.8018 | 25.6322 | 25.6216 |
| Syntaxin-binding protein 3 | O00186 | STXBP3 | Extracellular or secreted | 1.692366667 | 1.960510362 | 24.1971 | 24.6206 | 25.1437 | 25.8628 | 26.4287 | 26.747 |
| Syntaxin-binding protein 5 | Q5T5C0-2;Q5T5C0 | STXBP5 | Extracellular or secreted | 1.019866667 | 0.08870225 | 30.4461 | 20.8004 | 28.614 | 31.1619 | 29.6959 | 22.0623 |

| | | | | | | | | | | | |
|---|---|---|---|---|---|---|---|---|---|---|---|
| Activated RNA polymerase II transcriptional coactivator p | P53999 | SUB1 | Extracellular or secreted | 1.3395 | 1.145553145 | 28.6996 | 28.3232 | 28.3886 | 30.5447 | 29.7457 | 29.1395 |
| Succinyl-CoA ligase [ADP-forming] subunit beta, mitocho | Q9P2R7-2;Q9P2R7 | SUCLA2 | Extracellular or secreted | -1.727266667 | 0.710570629 | 25.1129 | 25.1539 | 26.9353 | 22.2494 | 24.5812 | 25.1897 |
| Succinyl-CoA ligase [ADP/GDP-forming] subunit alpha, m | P53597 | SUCLG1 | Extracellular or secreted | -2.4544 | 1.434094469 | 28.4543 | 28.2606 | 29.968 | 26.7321 | 26.3747 | 26.2129 |
| Succinyl-CoA ligase [GDP-forming] subunit beta, mitocho | Q96I99;Q96I99-2 | SUCLG2 | Extracellular or secreted | -2.4315 | 1.487924132 | 29.7979 | 29.445 | 31.0336 | 27.6084 | 27.5185 | 27.8551 |
| Sin3 histone deacetylase corepressor complex componer | Q9H7L9 | SUDS3 | Extracellular or secreted | -1.449133333 | 0.714257852 | 23.4799 | 21.4988 | 23.9867 | 21.6932 | 21.2581 | 21.6667 |
| Suppressor of G2 allele of SKP1 homolog | Q9Y2Z0-2;Q9Y2Z0 | SUGT1 | Other | -0.661 | 0.604991374 | 26.8681 | 26.3394 | 27.0186 | 25.385 | 26.0478 | 26.8103 |
| Sulfatase-modifying factor 2 | Q8NBJ7;Q8NBJ7-2;Q8NBJ7-3 | SUMF2 | Extracellular or secreted | 0.043866667 | 0.041293181 | 29.0486 | 28.3145 | 29.1155 | 29.2113 | 29.0273 | 28.3716 |
| Small ubiquitin-related modifier 1 | P63165 | SUMO1 | Extracellular or secreted | -0.350833333 | 0.399594529 | 28.4224 | 27.7524 | 28.4956 | 28.4105 | 27.7653 | 27.4421 |
| Small ubiquitin-related modifier 2;Small ubiquitin-related | P61956-2;P61956;P55854;P5 | SUMO2;SUMO | Other | 0.361333333 | 0.146565118 | 29.8199 | 28.7823 | 28.8081 | 31.0996 | 28.976 | 28.4187 |
| Small ubiquitin-related modifier 4 | Q6EEV6 | SUMO4 | Other | 1.8899 | 0.258059408 | 20.9656 | 21.924 | 27.3374 | 27.8841 | 26.9453 | 21.0673 |
| FACT complex subunit SPT16 | Q9Y5B9 | SUPT16H | Other | -1.162133333 | 1.297525576 | 30.8604 | 30.0417 | 30.4084 | 28.7633 | 29.1771 | 29.8837 |
| Transcription elongation factor SPT5 | O00267-2;O00267 | SUPT5H | Other | -0.222266667 | 0.360914441 | 28.1042 | 27.9896 | 28.1776 | 27.6562 | 27.623 | 28.3254 |
| Transcription elongation factor SPT6 | Q7KZ85 | SUPT6H | Other | -1.7348 | 0.397125659 | 25.2537 | 25.9752 | 27.1456 | 21.9058 | 23.8215 | 27.4428 |
| Sushi, von Willebrand factor type A, EGF and pentraxin d | Q4LDE5;Q4LDE5-4 | SVEP1 | Extracellular or secreted | 1.636833333 | 1.07795378 | 27.4904 | 26.9067 | 25.5643 | 27.7059 | 28.2881 | 28.8779 |
| Supervillin | O95425-2;O95425-4;O95425 | SVIL | Extracellular or secreted | -1.566366667 | 0.562727457 | 25.5379 | 22.6642 | 23.6556 | 23.3773 | 20.5843 | 23.197 |
| Switch-associated protein 70 | Q9UH65 | SWAP70 | Extracellular or secreted | -0.947033333 | 0.489793108 | 25.2277 | 26.3851 | 26.9183 | 23.9372 | 25.5912 | 26.1616 |
| Symplekin | Q92797;Q92797-2 | SYMPK | Extracellular or secreted | -1.075366667 | 0.374832844 | 23.7597 | 22.9168 | 26.4018 | 22.4978 | 23.3372 | 24.0172 |
| Heterogeneous nuclear ribonucleoprotein Q | O60506;O60506-2 | SYNCRIP | Other | -0.895366667 | 0.864038475 | 30.4626 | 30.7509 | 31.117 | 29.1225 | 30.0785 | 30.4434 |
| Heterogeneous nuclear ribonucleoprotein Q | O60506-3;O60506-4;O60506 | SYNCRIP | Other | 0.0143 | 0.005568091 | 24.5838 | 24.3259 | 26.4881 | 24.4675 | 24.8887 | 26.0845 |
| Nesprin-2 | Q8WXH0;Q8WXH0-2;Q8WXI | SYNE2 | Extracellular or secreted | -4.021933333 | 0.917711694 | 27.2202 | 28.5409 | 26.3373 | 25.6616 | 20.1638 | 24.2072 |
| Nesprin-3 | Q6ZMZ3-3;Q6ZMZ3-2;Q6ZM | SYNE3 | Other | -4.408066667 | 2.088263513 | 28.3548 | 27.9025 | 26.6733 | 23.0695 | 22.146 | 24.4909 |
| Synaptophysin-like protein 1 | Q16563-2;Q16563 | SYPL1 | Extracellular or secreted | 0.670166667 | 1.384188226 | 23.9809 | 23.8147 | 24.2773 | 24.527 | 25.0418 | 24.5146 |
| SUZ domain-containing protein 1 | Q7Z422-2;Q7Z422-4;Q7Z422 | SZRD1 | Other | 0.7735 | 1.467481915 | 25.7728 | 25.607 | 25.379 | 25.9842 | 26.51 | 26.5851 |
| Transforming acidic coiled-coil-containing protein 1 | O75410-5;O75410-4;O75410 | TACC1 | Other | -0.361966667 | 1.335507229 | 26.327 | 26.5628 | 26.6374 | 26.0128 | 26.3042 | 26.1243 |
| Transforming acidic coiled-coil-containing protein 3 | Q9Y6A5 | TACC3 | Other | -0.0758 | 0.084817638 | 24.3147 | 23.4262 | 23.4269 | 23.6697 | 23.5723 | 23.6984 |
| Tumor-associated calcium signal transducer 2 | P09758 | TACSTD2 | Extracellular or secreted | 1.198733333 | 1.140076408 | 29.2321 | 29.3325 | 28.7234 | 31.0263 | 30.1589 | 29.699 |
| Transcription initiation factor TFIID subunit 10 | Q12962 | TAF10 | Other | 0.865466667 | 0.484277348 | 20.5806 | 20.8149 | 22.1865 | 21.7366 | 23.205 | 21.2368 |
| TATA-binding protein-associated factor 2N | Q92804-2;Q92804 | TAF15 | Other | 0.234033333 | 0.193090005 | 26.6527 | 26.8992 | 28.0447 | 27.3668 | 27.3869 | 27.545 |
| Transgelin | Q01995 | TAGLN | Other | -4.195833333 | 1.584572972 | 26.6225 | 28.9198 | 25.8108 | 23.1339 | 24.0516 | 21.5801 |
| Transgelin-2 | P37802;P37802-2 | TAGLN2 | Extracellular or secreted | -1.4021 | 1.668898685 | 31.9317 | 32.4512 | 32.0703 | 30.2141 | 31.1858 | 30.847 |
| Transaldolase | P37837 | TALDO1 | Extracellular or secreted | -0.0284 | 0.0231162 | 33.1872 | 33.0234 | 32.4618 | 33.4006 | 32.9581 | 32.2285 |
| TAR DNA-binding protein 43 | Q13148 | TARDBP | Other | -1.509333333 | 0.525261404 | 26.8891 | 26.6859 | 28.6028 | 25.8653 | 24.065 | 27.7195 |
| Threonine--tRNA ligase, cytoplasmic | P26639;P26639-2 | TARS | Extracellular or secreted | 0.1275 | 0.264389886 | 29.7867 | 30.0652 | 29.4868 | 29.7486 | 30.026 | 29.9466 |
| Putative deoxyribonuclease TATDN1 | Q6P1N9;Q6P1N9-2 | TATDN1 | Other | -0.4503 | 0.436210132 | 28.4689 | 28.7776 | 27.6117 | 27.6159 | 28.3648 | 27.5266 |
| Putative deoxyribonuclease TATDN3 | Q17R31-5;Q17R31-4;Q17R31 | TATDN3 | Other | -0.0254 | 0.016675131 | 23.3191 | 22.8365 | 22.3748 | 23.5068 | 22.8891 | 22.0583 |
| Tax1-binding protein 3 | O14907 | TAX1BP3 | Extracellular or secreted | 0.800633333 | 1.220415773 | 26.1982 | 25.4317 | 26.1115 | 26.5894 | 26.9791 | 26.5748 |
| TBC1 domain family member 10B | Q4KMP7 | TBC1D10B | Extracellular or secreted | -0.156 | 0.10539588 | 26.19 | 25.5132 | 24.4986 | 25.4735 | 25.2201 | 25.0402 |
| TBC1 domain family member 22A | Q8WUA7-4;Q8WUA7;Q8WU | TBC1D22A | Other | -0.529766667 | 0.477929872 | 23.147 | 22.2187 | 22.7192 | 22.5732 | 22.536 | 21.3864 |
| TBC1 domain family member 23 | Q9NUY8-2;Q9NUY8 | TBC1D23 | Other | -0.408533333 | 0.374272556 | 24.6988 | 24.9471 | 25.4232 | 24.5226 | 23.9921 | 25.3288 |
| TBC1 domain family member 4 | O60343-2;O60343-3;O60343 | TBC1D4 | Extracellular or secreted | -2.406933333 | 1.139685702 | 24.6882 | 22.667 | 23.4918 | 22.3155 | 21.5871 | 19.7236 |
| TBC1 domain family member 5 | Q92609;Q92609-2 | TBC1D5 | Extracellular or secreted | -0.095433333 | 0.052708405 | 21.8399 | 23.1575 | 22.5296 | 21.4611 | 23.0936 | 22.686 |
| TBC1 domain family member 9B | Q66K14-2;Q66K14 | TBC1D9B | Other | -1.7336 | 0.734578303 | 24.2896 | 21.2455 | 23.3307 | 22.0648 | 20.9137 | 20.6865 |
| Tubulin-specific chaperone A | O75347;O75347-2 | TBCA | Extracellular or secreted | 0.168266667 | 0.087652735 | 30.0237 | 29.9964 | 30.0737 | 31.2914 | 30.2316 | 29.0756 |
| Tubulin-folding cofactor B | Q99426;Q99426-2 | TBCB | Other | -0.257533333 | 0.453956024 | 25.2464 | 25.878 | 25.6332 | 25.0774 | 25.2811 | 25.6265 |
| Tubulin-specific chaperone D | Q9BTW9;Q9BTW9-4;Q9BTW | TBCD | Extracellular or secreted | -0.9403 | 0.537666915 | 27.0692 | 27.6419 | 28.6748 | 25.8343 | 26.8078 | 27.9229 |
| Tubulin-specific chaperone E | Q15813;Q15813-2 | TBCE | Other | -1.6722 | 0.535415958 | 24.794 | 24.9479 | 26.2511 | 21.3558 | 24.3566 | 25.264 |
| F-box-like/WD repeat-containing protein TBL1XR1 | Q9BZK7 | TBL1XR1 | Other | -0.180433333 | 0.324079956 | 29.3432 | 29.0389 | 28.7039 | 29.1034 | 28.7682 | 28.6731 |
| Transducin beta-like protein 2 | Q9Y4P3 | TBL2 | Extracellular or secreted | 1.022933333 | 1.098354955 | 21.7182 | 21.2755 | 22.4459 | 22.9593 | 22.5586 | 22.9905 |
| Transducin beta-like protein 3 | Q12788 | TBL3 | Other | -0.263233333 | 0.08066644 | 22.2416 | 21.0386 | 24.4758 | 22.294 | 23.2088 | 21.4635 |
| Transcription elongation factor A protein 1 | P23193;P23193-2 | TCEA1 | Other | -0.740866667 | 0.295080474 | 26.1656 | 26.218 | 26.3988 | 23.7051 | 26.1288 | 26.7259 |
| Transcription elongation factor A protein-like 3 | Q969E4 | TCEAL3 | Other | 0.718633333 | 0.330097915 | 25.5454 | 23.9019 | 24.7806 | 26.9429 | 24.6658 | 24.7751 |
| Transcription elongation factor A protein-like 4 | Q96EI5;Q96EI5-2 | TCEAL4 | Other | 0.530233333 | 0.326582957 | 20.5962 | 20.2811 | 21.6461 | 22.41 | 20.9043 | 20.7998 |
| Transcription elongation factor B polypeptide 1 | Q15369;Q15369-2 | TCEB1 | Other | -0.659833333 | 0.605225819 | 28.0439 | 28.2008 | 28.0396 | 27.9593 | 27.7204 | 26.6251 |
| Transcription elongation factor B polypeptide 2 | Q15370;Q15370-2 | TCEB2 | Extracellular or secreted | -0.225033333 | 0.191386895 | 28.331 | 28.5949 | 27.7301 | 28.4313 | 28.2824 | 27.2672 |
| Transcription elongation regulator 1 | O14776-2;O14776 | TCERG1 | Other | -2.6083 | 0.952943953 | 29.1531 | 28.3987 | 28.749 | 24.2204 | 26.8351 | 27.4204 |
| Treacle protein | Q13428-3;Q13428;Q13428-4 | TCOF1 | Other | -1.635233333 | 2.192503892 | 30.0942 | 29.4942 | 29.7741 | 28.4119 | 28.3626 | 27.6823 |
| T-complex protein 1 subunit alpha | P17987 | TCP1 | Extracellular or secreted | -0.869433333 | 0.871768206 | 31.2811 | 31.4365 | 32.3005 | 30.2385 | 30.7629 | 31.4084 |
| Teneurin-3 | Q9P273 | TENM3 | Other | -1.978133333 | 0.973150873 | 26.8653 | 26.7823 | 24.9223 | 24.9943 | 24.8392 | 22.802 |
| Telomeric repeat-binding factor 2-interacting protein 1 | Q9NYB0 | TERF2IP | Other | 0.303466667 | 0.070408817 | 24.1909 | 21.0264 | 25.025 | 22.1457 | 23.8469 | 25.1601 |
| Testin | Q9UGI8-2;Q9UGI8 | TES | Extracellular or secreted | -0.326 | 0.32541992 | 27.3598 | 27.8579 | 28.0222 | 26.7505 | 27.5921 | 27.9193 |
| Testis-expressed sequence 264 protein | Q9Y6I9 | TEX264 | Extracellular or secreted | -1.798633333 | 1.259028167 | 24.0715 | 22.8758 | 24.671 | 21.8997 | 21.5934 | 22.7293 |
| Serotransferrin | P02787 | TF | Extracellular or secreted | 12.47253333 | 3.910335319 | 21.8883 | 21.459 | 21.5937 | 34.7381 | 33.9187 | 33.7018 |
| Transcription factor A, mitochondrial | Q00059;Q00059-2 | TFAM | Extracellular or secreted | 2.982433333 | 1.686683045 | 21.5447 | 21.0464 | 19.4989 | 24.0977 | 24.1618 | 22.7778 |
| Protein TFG | Q92734-2;Q92734;Q92734-4 | TFG | Extracellular or secreted | -0.084733333 | 0.156821706 | 29.794 | 29.3331 | 29.6728 | 29.317 | 29.4241 | 29.8046 |
| Tissue factor pathway inhibitor 2 | P48307-2;P48307 | TFPI2 | Extracellular or secreted | -0.172033333 | 0.063397308 | 23.4599 | 23.5695 | 21.3198 | 22.2181 | 23.7721 | 21.8429 |
| Transferrin receptor protein 1;Transferrin receptor prote | P02786 | TFRC | Extracellular or secreted | -0.9023 | 0.538949258 | 31.9116 | 31.4284 | 29.8963 | 30.9242 | 29.9542 | 29.651 |
| Transforming growth factor beta-1;Latency-associated pe | P01137 | TGFB1 | Extracellular or secreted | 0.9123 | 0.807724009 | 26.9898 | 26.971 | 25.7183 | 27.4067 | 27.3291 | 27.6802 |
| Transforming growth factor beta-2;Latency-associated pe | P61812 | TGFB2 | Extracellular or secreted | 1.967766667 | 0.820088069 | 30.9803 | 32.1026 | 29.0214 | 32.1393 | 33.0868 | 32.7815 |
| Transforming growth factor-beta-induced protein ig-h3 | Q15582 | TGFBI | Extracellular or secreted | -1.777466667 | 0.70203848 | 31.8149 | 33.0449 | 29.818 | 28.7063 | 29.8611 | 30.778 |
| Protein-glutamine gamma-glutamyltransferase 2 | P21980;P21980-2 | TGM2 | Extracellular or secreted | 0.836966667 | 0.502237604 | 28.4409 | 28.8264 | 28.9132 | 28.3358 | 29.9269 | 30.4287 |
| Protein-glutamine gamma-glutamyltransferase E;Protein | Q08188 | TGM3 | Extracellular or secreted | 1.643933333 | 0.393349537 | 23.8901 | 21.7532 | 25.6872 | 27.9336 | 23.3469 | 24.9818 |
| Trans-Golgi network integral membrane protein 2 | O43493-4;O43493-2;O43493 | TGOLN2 | Extracellular or secreted | 0.466 | 0.2968681 | 24.2792 | 23.9591 | 24.6874 | 25.9039 | 24.2254 | 24.1944 |
| Thrombospondin-1 | P07996;P07996-2 | THBS1 | Extracellular or secreted | -0.356133333 | 0.525458669 | 34.6989 | 34.9403 | 34.0777 | 34.3607 | 34.187 | 34.1008 |
| Probable tRNA(His) guanylyltransferase | Q9NWX6 | THG1L | Other | -0.8439 | 0.506028286 | 22.7352 | 23.5181 | 22.9894 | 20.9905 | 23.014 | 22.7065 |
| THO complex subunit 2 | Q8NI27 | THOC2 | Other | -2.8138 | 1.545876538 | 25.0376 | 25.1092 | 25.4584 | 21.7273 | 21.9917 | 23.4448 |
| THO complex subunit 3 | Q96J01;Q96J01-2 | THOC3 | Extracellular or secreted | -1.164766667 | 0.563071242 | 25.6516 | 24.9928 | 27.5889 | 24.1924 | 25.3628 | 25.1838 |
| THO complex subunit 5 homolog | Q13769 | THOC5 | Other | -0.0485 | 0.010120696 | 24.2172 | 20.6771 | 24.3134 | 23.8563 | 20.9782 | 24.2277 |
| THO complex subunit 6 homolog | Q86W42-3;Q86W42 | THOC6 | Other | -1.393333333 | 0.598393033 | 23.0873 | 20.8407 | 22.8772 | 21.3352 | 19.3979 | 21.8921 |
| Thimet oligopeptidase | P52888 | THOP1 | Other | -0.251533333 | 0.183995968 | 30.194 | 29.8746 | 29.2066 | 30.2921 | 29.4016 | 28.8269 |
| Thyroid hormone receptor-associated protein 3 | Q9Y2W1 | THRAP3 | Extracellular or secreted | -0.8374 | 0.93264741 | 30.0718 | 29.1182 | 30.154 | 29.4149 | 28.6792 | 28.7377 |
| THUMP domain-containing protein 1 | Q9NXG2 | THUMPD1 | Other | -0.067266667 | 0.081011625 | 26.2791 | 26.3987 | 26.934 | 26.3002 | 26.2163 | 26.8935 |
| Thymocyte nuclear protein 1 | Q9P016;Q9P016-2 | THYN1 | Other | -1.061 | 1.225732306 | 25.5005 | 25.4213 | 25.6797 | 23.9337 | 24.5522 | 24.9326 |
| Nucleolysin TIA-1 isoform p40 | P31483;P31483-3 | TIA1 | Other | -2.412233333 | 0.468883082 | 26.8823 | 26.8673 | 27.6115 | 20.8614 | 26.1675 | 27.0955 |
| Nucleolysin TIAR | Q01085 | TIAL1 | Extracellular or secreted | -1.168466667 | 0.551390522 | 24.0921 | 24.3845 | 24.6709 | 21.7106 | 23.4769 | 24.4546 |
| Nucleolysin TIAR | Q01085-2 | TIAL1 | Extracellular or secreted | -0.796266667 | 0.505411344 | 25.8931 | 26.0724 | 26.3923 | 24.2694 | 25.3649 | 26.3347 |
| Fructose-2,6-bisphosphatase TIGAR | Q9NQ88 | TIGAR | Other | -0.443833333 | 0.3407986 | 25.1187 | 25.1393 | 24.9681 | 23.6758 | 25.2418 | 24.977 |
| Mitochondrial import inner membrane translocase subu | P62072 | TIMM10 | Other | -0.7886 | 0.312813026 | 25.9943 | 26.1161 | 25.4144 | 26.1793 | 25.7628 | 23.2169 |
| Mitochondrial import inner membrane translocase subu | Q9Y5L4 | TIMM13 | Other | 0.2384 | 0.177410613 | 24.6498 | 25.2366 | 25.6112 | 24.7896 | 26.2072 | 25.216 |
| Mitochondrial import inner membrane translocase subu | O43615 | TIMM44 | Other | -1.693633333 | 0.588052019 | 23.2392 | 23.3131 | 26.5601 | 22.4918 | 23.3362 | 22.2035 |
| Mitochondrial import inner membrane translocase subu | Q9Y5J7 | TIMM9 | Other | -0.8632 | 0.692811018 | 27.9311 | 28.3313 | 28.1034 | 27.714 | 27.7438 | 26.3184 |
| Metalloproteinase inhibitor 1 | P01033 | TIMP1 | Extracellular or secreted | 2.508466667 | 0.991607154 | 33.4265 | 34.46 | 31.4771 | 35.6976 | 35.7164 | 35.475 |
| Metalloproteinase inhibitor 2 | P16035 | TIMP2 | Extracellular or secreted | 0.6611 | 0.611941257 | 29.1641 | 29.5523 | 28.1769 | 29.5853 | 29.9671 | 29.3242 |
| Metalloproteinase inhibitor 4 | Q99727 | TIMP4 | Extracellular or secreted | 1.0139 | 0.700969137 | 24.3337 | 24.2283 | 22.6613 | 24.6101 | 24.9113 | 24.7436 |
| Tubulointerstitial nephritis antigen-like | Q9GZM7;Q9GZM7-3 | TINAGL1 | Extracellular or secreted | 2.655666667 | 0.997223908 | 31.0775 | 31.6284 | 28.6185 | 33.2184 | 33.2139 | 32.8591 |
| TIP41-like protein | O75663;O75663-2 | TIPRL | Other | -0.598066667 | 1.059102938 | 28.1724 | 27.8843 | 28.5756 | 27.2714 | 27.7864 | 27.7803 |
| Tight junction protein ZO-1 | Q07157-2;Q07157 | TJP1 | Extracellular or secreted | -0.540333333 | 1.095411978 | 28.2973 | 27.9493 | 27.9922 | 27.2182 | 27.5405 | 27.8591 |
| Tight junction protein ZO-2 | Q9UDY2-3;Q9UDY2;Q9UDY2 | TJP2 | Extracellular or secreted | -0.015933333 | 0.026661605 | 26.9981 | 26.504 | 27.0221 | 26.8002 | 26.6606 | 27.0156 |
| Thymidine kinase, cytosolic | P04183 | TK1 | Other | -0.4072 | 0.540102358 | 24.4906 | 25.3863 | 25.3005 | 24.7178 | 24.6836 | 24.5544 |
| Transketolase | P29401;P29401-2 | TKT | Extracellular or secreted | -0.343333333 | 0.370057687 | 35.1374 | 34.9709 | 34.5509 | 35.0529 | 34.6555 | 33.9208 |
| TLD domain-containing protein 1 | Q6P9B6 | TLDC1 | Other | 0.683566667 | 0.567399434 | 24.4499 | 25.6203 | 25.9522 | 25.6329 | 26.1867 | 26.2535 |
| Transducin-like enhancer protein 3;Transducin-like enha | Q04726-2;Q04726-7;Q04726 | TLE3;TLE2;TLE4 | Other | -2.360233333 | 1.441944692 | 24.3604 | 24.6704 | 25.011 | 22.8821 | 22.8292 | 21.2498 |
| Tolloid-like protein 1 | O43897 | TLL1 | Extracellular or secreted | 1.7305 | 1.268401199 | 24.4596 | 23.9886 | 22.9696 | 25.4554 | 25.7129 | 25.441 |
| Talin-1 | Q9Y490 | TLN1 | Extracellular or secreted | -0.919566667 | 0.458597432 | 29.2619 | 29.703 | 30.02 | 27.3372 | 28.9631 | 29.9259 |
| Transmembrane 9 superfamily member 2 | Q99805 | TM9SF2 | Extracellular or secreted | 0.1959 | 0.408240181 | 24.7013 | 24.1278 | 24.5459 | 24.6737 | 24.8147 | 24.4743 |
| Transmembrane 9 superfamily member 4 | Q92544 | TM9SF4 | Other | -0.748366667 | 1.045411168 | 23.1654 | 22.5785 | 23.3609 | 22.5547 | 22.4981 | 21.8069 |
| Translation machinery-associated protein 16 | Q96EY4 | TMA16 | Other | -1.945333333 | 0.732480427 | 23.4446 | 23.4081 | 25.4439 | 20.3795 | 23.7473 | 22.3338 |
| Translation machinery-associated protein 7 | Q9Y2S6 | TMA7 | Other | 1.610166667 | 0.680386626 | 25.0162 | 27.3641 | 27.2376 | 29.5658 | 27.9129 | 26.9697 |
| Protein lifeguard 3 | Q969X1 | TMBIM1 | Extracellular or secreted | 2.863 | 3.839040118 | 22.8176 | 22.9428 | 22.4893 | 25.7666 | 25.6268 | 25.4453 |
| Transmembrane emp24 domain-containing protein 10 | P49755 | TMED10 | Extracellular or secreted | 0.7015 | 0.665206082 | 21.9648 | 21.8076 | 21.8543 | 22.0479 | 23.3489 | 22.3344 |
| Transmembrane protein 106B | Q9NUM4 | TMEM106B | Extracellular or secreted | 2.0095 | 0.565526117 | 20.2578 | 24.7793 | 22.6906 | 23.0308 | 25.5534 | 25.172 |
| Transmembrane protein 132A | Q24JP5;Q24JP5-2;Q24JP5-4; | TMEM132A | Extracellular or secreted | 1.438566667 | 0.802593426 | 29.6505 | 30.2877 | 28.0831 | 30.9307 | 30.7125 | 30.6938 |
| Transmembrane protein 154 | Q6P9G4 | TMEM154 | Other | 3.465066667 | 1.543176333 | 20.2922 | 22.5298 | 22.6753 | 25.9046 | 25.4119 | 24.576 |
| Transmembrane protein 205 | Q6UW68 | TMEM205 | Extracellular or secreted | 3.415766667 | 2.568193444 | 21.401 | 21.3721 | 21.2648 | 24.5621 | 24.5478 | 25.1753 |
| CSC1-like protein 1 | O94886 | TMEM63A | Extracellular or secreted | 1.093466667 | 1.262106248 | 22.8852 | 22.6207 | 22.8196 | 24.3813 | 23.8449 | 23.3797 |
| Transmembrane protein 87A | Q8NBN3-3;Q8NBN3 | TMEM87A | Extracellular or secreted | 0.101833333 | 0.173568357 | 24.5667 | 25.0893 | 24.6199 | 24.6388 | 25.1394 | 24.8032 |
| TATA element modulatory factor | P82094;P82094-2 | TMF1 | Other | -0.517666667 | 0.647367887 | 24.8256 | 24.7017 | 24.4448 | 24.6674 | 23.6125 | 24.1392 |
| Tropomodulin-3 | Q9NYL9 | TMOD3 | Other | -0.536833333 | 0.953526973 | 27.8997 | 28.1067 | 28.5158 | 27.2606 | 27.7682 | 27.8829 |
| Lamina-associated polypeptide 2, isoforms beta/gamma; | P42167;P42167-3;P42167-2 | TMPO | Other | 0.0948 | 0.042964483 | 27.7673 | 27.1444 | 29.5096 | 28.1498 | 28.2784 | 28.2775 |
| Thymosin beta-10 | P63313 | TMSB10 | Other | 1.778433333 | 0.607468271 | 27.3687 | 27.4499 | 26.5275 | 31.1137 | 27.882 | 27.6857 |

| | | | | | | | | | | |
|---|---|---|---|---|---|---|---|---|---|---|
| Thymosin beta-4;Hematopoietic system regulatory pepti | P62328 | TMSB4X | Extracellular or secreted | 2.5815 | 1.028368257 | 26.0891 | 26.0538 | 25.4827 | 30.2255 | 27.5873 | 27.5573 |
| Thioredoxin-related transmembrane protein 1 | Q9H3N1 | TMX1 | Other | -0.125766667 | 0.044065269 | 24.3464 | 24.4497 | 27.1091 | 25.5542 | 25.1341 | 24.8396 |
| Tenascin | P24821-4;P24821;P24821-2; | TNC | Extracellular or secreted | 1.547833333 | 0.380241973 | 31.3329 | 31.2961 | 26.7433 | 31.3429 | 31.2268 | 31.4461 |
| Tumor necrosis factor;Tumor necrosis factor, membrane | P01375 | TNF | Extracellular or secreted | 2.463466667 | 1.089816959 | 22.6168 | 20.0168 | 20.8455 | 23.7613 | 23.3818 | 23.7264 |
| Tumor necrosis factor alpha-induced protein 2 | Q03169 | TNFAIP2 | Extracellular or secreted | 1.628 | 0.776107064 | 20.9389 | 22.8844 | 23.5345 | 24.3379 | 23.7835 | 24.1204 |
| Tumor necrosis factor-inducible gene 6 protein | P98066 | TNFAIP6 | Extracellular or secreted | 1.9587 | 1.11415081 | 29.3959 | 29.9004 | 28.7146 | 30.1056 | 31.4107 | 32.3707 |
| Tumor necrosis factor receptor superfamily member 10B | O14763-2;O14763 | TNFRSF10B | Extracellular or secreted | 2.798066667 | 1.304842857 | 24.2349 | 23.9713 | 21.7568 | 25.8199 | 27.1773 | 25.36 |
| Tumor necrosis factor receptor superfamily member 11B | O00300 | TNFRSF11B | Extracellular or secreted | 8.058466667 | 2.653115826 | 20.5189 | 23.3251 | 21.1885 | 28.2712 | 30.0389 | 30.8978 |
| Tumor necrosis factor receptor superfamily member 12A | Q9NP84 | TNFRSF12A | Extracellular or secreted | 2.004966667 | 0.982709439 | 22.9495 | 24.6854 | 22.0752 | 26.0205 | 25.3478 | 24.3567 |
| Tumor necrosis factor receptor superfamily member 1A; | P19438;P19438-5;P19438-4; | TNFRSF1A | Extracellular or secreted | 2.7337 | 1.124122107 | 21.0656 | 22.7684 | 20.5608 | 25.9241 | 22.9463 | 23.7255 |
| Tumor necrosis factor receptor superfamily member 21 | O75509 | TNFRSF21 | Extracellular or secreted | 0.111666667 | 0.039985684 | 26.2621 | 25.2315 | 25.6613 | 27.5194 | 25.3018 | 24.6687 |
| Tumor necrosis factor ligand superfamily member 13 | O43508-2;O75888-5;O75888 | TNFSF13 | Extracellular or secreted | -0.4104 | 0.096200439 | 23.651 | 26.232 | 21.3805 | 22.8981 | 24.1338 | 23.0004 |
| 182 kDa tankyrase-1-binding protein | Q9C0C2 | TNKS1BP1 | Extracellular or secreted | 0.4333 | 0.845441642 | 28.1678 | 28.0685 | 28.5804 | 28.5116 | 28.5442 | 29.0608 |
| Transportin-1 | Q92973-2;Q92973;Q92973-3 | TNPO1 | Extracellular or secreted | -1.235166667 | 0.799562268 | 27.9778 | 28.7236 | 28.9246 | 26.2911 | 27.3012 | 28.3282 |
| Transportin-2 | O14787-2;O14787 | TNPO2 | Other | -0.2916 | 0.2694868 | 25.2036 | 25.4327 | 26.2779 | 25.487 | 24.8011 | 25.7513 |
| Transportin-3 | Q9Y5L0;Q9Y5L0-3;Q9Y5L0-1 | TNPO3 | Other | -0.8774 | 0.553466139 | 25.5629 | 26.107 | 27.2139 | 24.8382 | 24.9862 | 26.4272 |
| Tensin-3 | Q68CZ2;Q68CZ2-3;Q68CZ2-2 | TNS3 | Other | 0.733233333 | 0.256097055 | 23.9705 | 21.0406 | 24.1708 | 23.0339 | 24.0159 | 24.3318 |
| Toll-interacting protein | Q9H0E2 | TOLLIP | Extracellular or secreted | 1.1375 | 1.312850901 | 23.4545 | 23.6531 | 24.4749 | 24.7013 | 25.0678 | 25.2259 |
| Target of Myb protein 1 | O60784;O60784-2;O60784-3 | TOM1 | Extracellular or secreted | -0.248733333 | 0.168588533 | 22.5845 | 22.5977 | 23.1764 | 21.5493 | 23.1458 | 22.9173 |
| Mitochondrial import receptor subunit TOM22 homolog | Q9NS69 | TOMM22 | Other | -1.5034 | 0.418447982 | 23.2966 | 22.8187 | 27.0734 | 23.1708 | 22.1221 | 23.3856 |
| Mitochondrial import receptor subunit TOM34 | Q15785 | TOMM34 | Other | 0.517366667 | 0.493585948 | 23.693 | 22.6518 | 23.9482 | 24.1074 | 23.8807 | 23.857 |
| Mitochondrial import receptor subunit TOM70 | O94826 | TOMM70A | Extracellular or secreted | -1.746733333 | 1.078795202 | 25.3428 | 26.5197 | 27.1884 | 23.6248 | 24.7103 | 25.4756 |
| DNA topoisomerase 1 | P11387 | TOP1 | Other | 1.5044 | 1.00735753 | 28.039 | 28.125 | 29.7578 | 29.9205 | 29.8716 | 30.6429 |
| DNA topoisomerase 2-alpha | P11388;P11388-2;P11388-3; | TOP2A | Other | -1.3573 | 0.733996871 | 28.711 | 27.1099 | 29.4805 | 27.7887 | 27.013 | 26.4278 |
| DNA topoisomerase 2-beta | Q02880-2;Q02880 | TOP2B | Other | -2.453433333 | 0.75789812 | 24.996 | 21.9666 | 26.0885 | 23.0371 | 21.5543 | 21.0994 |
| Torsin-1A-interacting protein 1 | Q5JTV8-3;Q5JTV8-2 | TOR1AIP1 | Other | 0.145233333 | 0.154158436 | 25.275 | 24.6291 | 25.6145 | 25.0767 | 25.7041 | 25.1735 |
| Torsin-1A-interacting protein 2 | Q8NFQ8 | TOR1AIP2 | Other | -0.234733333 | 0.068781318 | 23.8758 | 22.2176 | 24.4781 | 24.7441 | 23.6762 | 21.447 |
| Torsin-1B | O14657 | TOR1B | Extracellular or secreted | 0.670166667 | 0.423235617 | 21.4496 | 22.9406 | 23.391 | 22.7913 | 23.4011 | 23.5993 |
| Tumor suppressor p53-binding protein 1 | Q12888;Q12888-3;Q12888-2 | TP53BP1 | Other | 0.049766667 | 0.020137937 | 26.1052 | 25.3247 | 27.244 | 25.2881 | 26.1595 | 27.3756 |
| Quinone oxidoreductase PIG3 | Q53FA7;Q53FA7-2 | TP53I3 | Extracellular or secreted | 0.6441 | 0.438067561 | 28.807 | 28.7123 | 27.9096 | 30.0974 | 29.0097 | 28.2541 |
| Trophoblast glycoprotein | Q13641 | TPBG | Extracellular or secreted | 1.454666667 | 1.230548882 | 25.8675 | 25.5254 | 24.4508 | 26.3482 | 27.4101 | 26.4494 |
| Tumor protein D52 | P55327-2;P55327-4;P55327; | TPD52 | Extracellular or secreted | 0.8684 | 0.694324439 | 27.1399 | 28.0636 | 27.8651 | 28.9174 | 29.1265 | 27.6299 |
| Tumor protein D54 | O43399;O43399-5;O43399-7 | TPD52L2 | Other | 0.7617 | 0.781209838 | 28.1747 | 28.8634 | 28.972 | 29.5219 | 30.0045 | 28.7688 |
| Triosephosphate isomerase | P60174-1;P60174;P60174-4 | TPI1 | Extracellular or secreted | 0.259 | 0.180431238 | 36.1439 | 35.891 | 35.4279 | 36.9498 | 36.0117 | 35.2783 |
| Tropomyosin alpha-1 chain | P09493-3;P09493-9;P09493- | TPM1 | Extracellular or secreted | -0.356033333 | 0.589148943 | 26.9484 | 26.277 | 26.696 | 26.022 | 26.6429 | 26.1884 |
| Tropomyosin alpha-1 chain | P09493-5 | TPM1 | Extracellular or secreted | -0.256166667 | 0.207448233 | 23.0059 | 23.6182 | 23.2025 | 22.4562 | 22.7543 | 23.8476 |
| Tropomyosin alpha-3 chain | P06753 | TPM3 | Extracellular or secreted | -0.424633333 | 0.050150733 | 22.385 | 28.203 | 27.5572 | 28.4403 | 21.1956 | 27.2354 |
| Tropomyosin alpha-3 chain | P06753-2;P06753-3;P06753- | TPM3 | Extracellular or secreted | -0.186566667 | 0.2314605 | 33.1878 | 32.6264 | 32.536 | 32.9494 | 32.7045 | 32.1366 |
| Tropomyosin alpha-4 chain | P67936 | TPM4 | Extracellular or secreted | -0.517166667 | 0.488753422 | 32.5001 | 32.2526 | 32.171 | 32.5291 | 31.6892 | 31.1539 |
| Tropomyosin alpha-4 chain | P67936-2 | TPM4 | Extracellular or secreted | -3.433066667 | 1.65704345 | 25.2954 | 25.5969 | 24.3982 | 22.7836 | 21.8197 | 20.388 |
| Thiopurine S-methyltransferase | P51580 | TPMT | Extracellular or secreted | 0.402766667 | 0.217713735 | 20.8981 | 21.9115 | 21.8681 | 22.3788 | 20.7434 | 22.7638 |
| Tripeptidyl-peptidase 1 | O14773 | TPP1 | Extracellular or secreted | 0.268433333 | 0.188287681 | 29.0688 | 29.3623 | 27.7791 | 28.9708 | 29.3575 | 28.6872 |
| Tripeptidyl-peptidase 2 | P29144 | TPP2 | Other | 0.024433333 | 0.019391188 | 28.7975 | 29.3474 | 29.5475 | 28.5627 | 29.5236 | 29.6794 |
| Nucleoprotein TPR | P12270 | TPR | Other | -1.0246 | 1.463772781 | 31.7299 | 30.9974 | 31.7477 | 30.2491 | 30.4497 | 30.7024 |
| EKC/KEOPS complex subunit TPRKB | Q9Y3C4;Q9Y3C4-3;Q9Y3C4-2 | TPRKB | Other | -0.824933333 | 0.668476254 | 21.6621 | 23.0081 | 22.7998 | 22.3413 | 21.0685 | 21.5854 |
| Translationally-controlled tumor protein | P13693;P13693-2 | TPT1 | Extracellular or secreted | -0.150933333 | 0.113156015 | 32.7183 | 32.9331 | 31.551 | 32.4575 | 32.4204 | 31.8717 |
| Transformer-2 protein homolog alpha | Q13595-2;Q13595-4;Q13595 | TRA2A | Other | -0.105466667 | 0.025878113 | 23.8767 | 20.9314 | 24.3444 | 21.2591 | 23.8409 | 23.7361 |
| Transformer-2 protein homolog beta | P62995;P62995-3 | TRA2B | Other | -0.195833333 | 0.166735844 | 26.2255 | 26.18 | 26.5831 | 25.3359 | 26.6185 | 26.4467 |
| Heat shock protein 75 kDa, mitochondrial | Q12931-2;Q12931 | TRAP1 | Extracellular or secreted | -1.374533333 | 0.844295612 | 28.0429 | 28.1218 | 29.8872 | 27.0541 | 27.5673 | 27.3069 |
| Trafficking protein particle complex subunit 1 | Q9Y5R8 | TRAPPC1 | Other | 0.488866667 | 0.589282503 | 21.4011 | 21.8682 | 22.4901 | 22.5983 | 22.4148 | 22.2129 |
| Trafficking protein particle complex subunit 3 | O43617;O43617-2 | TRAPPC3 | Other | -0.0366 | 0.055258171 | 27.7202 | 27.5928 | 27.7257 | 27.3544 | 28.0578 | 27.5167 |
| Trafficking protein particle complex subunit 6B | Q86SZ2;Q86SZ2-2 | TRAPPC6B | Other | 0.1432 | 0.072372173 | 21.2468 | 23.4102 | 22.8118 | 22.6363 | 22.854 | 22.4081 |
| Tripartite motif-containing protein 16 | O95361 | TRIM16 | Other | -1.671 | 1.433206841 | 26.4253 | 27.4611 | 27.8358 | 25.007 | 25.7503 | 25.9519 |
| E3 ubiquitin/ISG15 ligase TRIM25 | Q14258 | TRIM25 | Other | -0.105633333 | 0.082598142 | 25.5145 | 25.5886 | 26.6645 | 25.5728 | 25.5597 | 26.3182 |
| Transcription intermediary factor 1-beta | Q13263;Q13263-2 | TRIM28 | Extracellular or secreted | -1.061266667 | 0.564154884 | 28.3654 | 28.0517 | 29.8196 | 26.7585 | 27.4074 | 28.887 |
| Tripartite motif-containing protein 29 | Q14134-2;Q14134 | TRIM29 | Other | -2.920266667 | 1.233342696 | 25.8346 | 27.1094 | 28.4551 | 22.8578 | 24.1142 | 25.6663 |
| Tripartite motif-containing protein 47 | Q96LD4;Q96LD4-2 | TRIM47 | Other | -1.770233333 | 0.589013091 | 22.9802 | 21.6974 | 23.7882 | 22.3255 | 22.0171 | 18.8125 |
| Cdc42-interacting protein 4 | Q15642-4;Q15642-2;Q15642 | TRIP10 | Extracellular or secreted | -0.815933333 | 0.945396372 | 29.1715 | 29.0221 | 29.6199 | 27.8068 | 28.7079 | 28.851 |
| Thyroid receptor-interacting protein 11 | Q15643 | TRIP11 | Other | -0.821766667 | 0.227475534 | 24.2814 | 24.6632 | 23.1182 | 20.6709 | 24.5323 | 24.3943 |
| Thyroid receptor-interacting protein 6 | Q15654 | TRIP6 | Extracellular or secreted | 0.150033333 | 0.108713968 | 24.7398 | 25.7209 | 25.9001 | 25.3157 | 25.2056 | 26.2896 |
| tRNA (guanine(26)-N(2))-dimethyltransferase | Q9NXH9-2;Q9NXH9 | TRMT1 | Other | -1.600666667 | 1.113106851 | 24.4601 | 25.3381 | 25.9792 | 24.0235 | 22.6558 | 24.2961 |
| Multifunctional methyltransferase subunit TRM112-like p | Q9UI30-2;Q9UI30 | TRMT112 | Extracellular or secreted | 0.657433333 | 1.044697368 | 23.0169 | 22.5046 | 22.7349 | 23.4916 | 22.9656 | 23.7715 |
| tRNA (adenine(58)-N(1))-methyltransferase non-catalytic | Q9UJA5;Q9UJA5-2;Q9UJA5-3 | TRMT6 | Other | -0.599633333 | 0.372790162 | 25.9884 | 25.9293 | 27.0802 | 24.7426 | 25.8195 | 26.6369 |
| tRNA (adenine(58)-N(1))-methyltransferase catalytic sub | Q96FX7 | TRMT61A | Other | -0.591233333 | 0.653669454 | 25.954 | 25.8541 | 26.6862 | 25.1169 | 25.4354 | 26.1683 |
| CCA tRNA nucleotidyltransferase 1, mitochondrial | Q96Q11-2;Q96Q11 | TRNT1 | Other | -0.617833333 | 2.075616609 | 26.3183 | 26.4002 | 26.6378 | 25.7124 | 25.9844 | 25.806 |
| 60 kDa SS-A/Ro ribonucleoprotein | P10155;P10155-3;P10155-5; | TROVE2 | Other | 0.523966667 | 0.685640896 | 29.8132 | 29.4568 | 29.0906 | 30.4773 | 29.9157 | 29.5395 |
| tRNA-splicing endonuclease subunit Sen15 | Q8WW01-2;Q8WW01 | TSEN15 | Other | -0.5813 | 0.575386443 | 22.7351 | 22.3638 | 22.7144 | 21.3007 | 22.159 | 22.6097 |
| Elongation factor Ts, mitochondrial | P43897;P43897-2 | TSFM | Other | 0.2372 | 0.056146138 | 22.1945 | 20.5583 | 24.9287 | 21.5243 | 23.2975 | 23.5713 |
| Tumor susceptibility gene 101 protein | Q99816;Q99816-2 | TSG101 | Extracellular or secreted | -0.483266667 | 0.910966984 | 24.8072 | 24.9656 | 24.7933 | 24.6586 | 24.4586 | 23.9991 |
| Tsukushin | Q8WUA8 | TSKU | Extracellular or secreted | -0.161933333 | 0.054794948 | 26.3124 | 26.3829 | 25.5278 | 24.1954 | 26.1224 | 27.4195 |
| Translin | Q15631 | TSN | Other | -0.838333333 | 1.550962529 | 29.8282 | 29.4612 | 29.7196 | 28.4548 | 29.0609 | 28.9783 |
| Translin-associated protein X | Q99598 | TSNAX | Other | 0.1093 | 0.411328075 | 29.2574 | 28.9963 | 29.044 | 29.0881 | 29.3584 | 29.1791 |
| Tetraspanin-14 | Q8NG11;Q8NG11-2;Q8NG11 | TSPAN14 | Extracellular or secreted | -0.346833333 | 0.339005241 | 24.9932 | 25.5301 | 24.2421 | 24.5848 | 24.8067 | 24.3334 |
| Tetraspanin-4 | O14817 | TSPAN4 | Extracellular or secreted | -1.2428 | 0.402028994 | 25.0429 | 25.9608 | 23.8787 | 25.2985 | 21.5746 | 24.2809 |
| Tetraspanin-9 | O75954 | TSPAN9 | Extracellular or secreted | 2.938166667 | 1.641831716 | 21.9413 | 22.4638 | 20.4219 | 24.8461 | 24.9252 | 23.8702 |
| Translocator protein | P30536 | TSPO | Extracellular or secreted | 2.006666667 | 0.995376546 | 21.2983 | 20.9968 | 20.6866 | 24.1514 | 23.1783 | 21.672 |
| Pre-rRNA-processing protein TSR1 homolog | Q2NL82 | TSR1 | Other | -1.589733333 | 0.928766851 | 23.714 | 21.8488 | 23.8536 | 22.2946 | 20.9899 | 21.3627 |
| Pre-rRNA-processing protein TSR2 homolog | Q969E8 | TSR2 | Other | 0.710866667 | 1.447335038 | 23.5891 | 23.1336 | 23.4651 | 23.9546 | 24.4565 | 23.9093 |
| Protein TSSC1 | Q53HC9 | TSSC1 | Other | 0.664833333 | 0.355324672 | 22.7582 | 24.3255 | 24.7101 | 24.0525 | 24.1507 | 25.5851 |
| Thiosulfate sulfurtransferase | Q16762 | TST | Extracellular or secreted | 0.532333333 | 0.26023219 | 22.6359 | 21.246 | 23.8239 | 23.0087 | 23.1848 | 23.1093 |
| GDP-L-fucose synthase | Q13630 | TSTA3 | Extracellular or secreted | 0.324266667 | 0.268655179 | 28.988 | 28.2896 | 28.5757 | 29.7403 | 28.7692 | 28.3166 |
| Thiosulfate sulfurtransferase/rhodanese-like domain-cor | Q8NFU3-2;Q8NFU3;Q8NFU3 | TSTD1 | Other | 2.622733333 | 1.068646705 | 21.9617 | 22.2089 | 20.9052 | 22.6351 | 25.7622 | 24.5467 |
| Tetratricopeptide repeat protein 1 | Q99614 | TTC1 | Other | 1.8129 | 0.529077485 | 24.9736 | 21.1212 | 25.0297 | 25.7239 | 25.3486 | 25.4907 |
| Tetratricopeptide repeat protein 13 | Q8NBP0;Q8NBP0-2 | TTC13 | Extracellular or secreted | 2.5539 | 0.839553481 | 24.2185 | 21.062 | 20.6873 | 24.0786 | 24.6567 | 24.8942 |
| Tetratricopeptide repeat protein 38 | Q5R3I4 | TTC38 | Extracellular or secreted | 0.1939 | 0.089254624 | 22.9464 | 22.976 | 23.1876 | 22.0567 | 24.5492 | 23.0858 |
| Tubulin--tyrosine ligase-like protein 12 | Q14166 | TTLL12 | Other | -0.8559 | 0.447812409 | 27.0899 | 27.3244 | 29.1494 | 26.183 | 26.9403 | 27.8727 |
| Titin | Q8WZ42-5;Q8WZ42-11;Q8W | TTN | Extracellular or secreted | 1.642266667 | 1.205842043 | 24.6677 | 25.4221 | 24.7679 | 27.2867 | 26.8689 | 25.6289 |
| Tubulin alpha-1B chain | P68363;P68363-2 | TUBA1B | Extracellular or secreted | -0.040433333 | 0.01985471 | 32.2197 | 31.9293 | 33.4386 | 31.8305 | 32.1758 | 33.46 |
| Tubulin alpha-1C chain | Q9BQE3 | TUBA1C | Other | 0.198833333 | 0.096426528 | 27.7535 | 27.4906 | 29.0941 | 27.5687 | 27.9917 | 29.3743 |
| Tubulin alpha-4A chain | P68366-2;P68366 | TUBA4A | Extracellular or secreted | -0.125966667 | 0.057808657 | 26.1741 | 26.4992 | 27.8827 | 26.1827 | 26.1858 | 27.8096 |
| Tubulin alpha chain-like 3 | A6NHL2-2;A6NHL2 | TUBAL3 | Other | -0.543866667 | 0.359362814 | 23.0117 | 23.1423 | 24.0937 | 21.9498 | 22.9244 | 23.7419 |
| Tubulin beta chain | P07437 | TUBB | Extracellular or secreted | 0.1807 | 0.074538311 | 29.7361 | 29.4179 | 31.2921 | 29.4626 | 29.9859 | 31.5397 |
| Tubulin beta-2A chain | Q13885 | TUBB2A | Extracellular or secreted | -0.289766667 | 0.149662376 | 26.4759 | 26.2285 | 27.5605 | 26.3125 | 25.5337 | 27.5494 |
| Tubulin beta-3 chain | Q13509;Q13509-2 | TUBB3 | Extracellular or secreted | 0.290466667 | 0.146010464 | 26.1674 | 26.5779 | 28.0953 | 26.7153 | 26.8758 | 28.1209 |
| Tubulin beta-4A chain | P04350 | TUBB4A | Extracellular or secreted | -0.307233333 | 0.132661903 | 21.7284 | 21.5375 | 23.0834 | 20.5844 | 21.8643 | 22.9789 |
| Tubulin beta-4B chain | P68371 | TUBB4B | Extracellular or secreted | -0.048466667 | 0.023975327 | 32.0625 | 31.8144 | 33.4644 | 31.7847 | 32.1645 | 33.2467 |
| Tubulin beta-6 chain | Q9BUF5 | TUBB6 | Extracellular or secreted | 0.265966667 | 0.131204977 | 24.8191 | 25.0421 | 26.8165 | 25.1039 | 26.0263 | 26.3454 |
| Tubulin beta-8 chain | Q3ZCM7 | TUBB8 | Extracellular or secreted | 0.157633333 | 0.056006952 | 28.1749 | 28.2483 | 30.3632 | 27.9531 | 29.0852 | 30.221 |
| Tubulin gamma-1 chain;Tubulin gamma-2 chain | P23258;Q9NRH3 | TUBG1;TUBG2 | Other | -0.8013 | 0.224308673 | 22.4048 | 22.8868 | 26.3279 | 22.3121 | 22.8326 | 24.0709 |
| Gamma-tubulin complex component 2 | Q9BSJ2;Q9BSJ2-4;Q9BSJ2-3 | TUBGCP2 | Other | -0.483833333 | 0.178092357 | 20.9089 | 23.0962 | 23.9941 | 22.5923 | 21.405 | 22.5504 |
| Gamma-tubulin complex component 4 | Q9UGJ1-2;Q9UGJ1 | TUBGCP4 | Other | -0.014266667 | 0.00551429 | 22.2783 | 21.7978 | 20.1407 | 21.128 | 22.4396 | 20.6064 |
| Elongation factor Tu, mitochondrial | P49411 | TUFM | Extracellular or secreted | 0.757633333 | 0.24088499 | 25.4296 | 24.6605 | 28.3565 | 27.3932 | 26.9036 | 26.4227 |
| Twinfilin-1 | Q12792;Q12792-3;Q12792-4 | TWF1 | Extracellular or secreted | 0.3761 | 0.282806339 | 27.413 | 28.1071 | 28.1872 | 27.5215 | 28.2071 | 29.107 |
| Twinfilin-2 | Q6IBS0 | TWF2 | Extracellular or secreted | -0.344566667 | 0.587594359 | 29.8075 | 29.903 | 29.81 | 29.053 | 29.7108 | 29.723 |
| Twisted gastrulation protein homolog 1 | Q9GZX9 | TWSG1 | Extracellular or secreted | 1.6692 | 1.048420939 | 25.6959 | 26.6037 | 24.5328 | 27.8405 | 27.3279 | 26.6716 |
| Alpha-taxilin | P40222 | TXLNA | Extracellular or secreted | 0.588566667 | 0.915589421 | 27.1045 | 26.4021 | 27.1512 | 27.7371 | 27.2209 | 27.4655 |
| Gamma-taxilin | Q9NUQ3;Q9NUQ3-2 | TXLNG | Other | 1.4311 | 1.200215538 | 30.2548 | 29.285 | 28.7914 | 31.5617 | 30.5841 | 30.4787 |
| Thioredoxin | P10599;P10599-2 | TXN | Extracellular or secreted | -0.095766667 | 0.074994703 | 33.2049 | 33.4886 | 32.601 | 33.4757 | 33.2379 | 32.2936 |
| Thioredoxin, mitochondrial | Q99757 | TXN2 | Other | -2.528466667 | 1.331904746 | 28.1819 | 27.3333 | 26.9469 | 25.4182 | 25.8516 | 23.6069 |
| Thioredoxin domain-containing protein 12 | O95881 | TXNDC12 | Extracellular or secreted | -1.303133333 | 1.218286031 | 29.6479 | 29.7926 | 29.8745 | 28.5998 | 28.9981 | 27.8077 |
| Thioredoxin domain-containing protein 17 | Q9BRA2 | TXNDC17 | Extracellular or secreted | -0.018233333 | 0.005665169 | 31.5316 | 30.9087 | 29.4838 | 31.8037 | 31.1045 | 28.9612 |
| Thioredoxin domain-containing protein 5 | Q8NBS9;Q8NBS9-2 | TXNDC5 | Extracellular or secreted | -1.250466667 | 1.359395715 | 31.6781 | 31.0608 | 31.8496 | 30.8595 | 30.2769 | 29.7007 |
| Thioredoxin-like protein 1 | O43396 | TXNL1 | Extracellular or secreted | -0.035733333 | 0.054783408 | 28.0174 | 28.3896 | 28.115 | 27.7932 | 28.1668 | 28.4548 |
| Thioredoxin-like protein 4A | P83876 | TXNL4A | Other | -1.005866667 | 0.765933259 | 24.2191 | 23.6123 | 24.6604 | 22.1801 | 23.7491 | 23.545 |
| Thioredoxin reductase 1, cytoplasmic | Q16881-4;Q16881-3;Q16881 | TXNRD1 | Extracellular or secreted | -0.6337 | 0.423274206 | 33.3108 | 32.6917 | 32.1479 | 32.8803 | 32.2817 | 31.0873 |
| Thioredoxin reductase 2, mitochondrial | Q9NNW7-2;Q9NNW7;Q9NN | TXNRD2 | Extracellular or secreted | -0.422966667 | 0.318234609 | 28.8763 | 28.6456 | 28.6891 | 29.0666 | 28.4841 | 27.3914 |
| Thymidine phosphorylase | P19971;P19971-2 | TYMP | Other | 0.466033333 | 0.702523373 | 27.6564 | 28.3831 | 28.4334 | 28.5181 | 28.5944 | 28.7585 |
| Tyrosine-protein kinase receptor TYRO3 | Q06418 | TYRO3 | Extracellular or secreted | 0.0069 | 0.003035859 | 24.5317 | 23.6282 | 22.209 | 23.9242 | 23.328 | 23.1374 |
| tRNA wybutosine-synthesizing protein 5 | A2RUC4 | TYW5 | Other | 0.0474 | 0.019862791 | 24.082 | 23.8648 | 22.7706 | 24.8621 | 23.4391 | 22.5584 |

| | | | | | | | | | | | |
|---|---|---|---|---|---|---|---|---|---|---|---|
| Splicing factor U2AF 35 kDa subunit | P0DN76;Q01081;Q01081-2;( | U2AF1 | Other | 0.0846 | 0.056332084 | 25.5855 | 25.8507 | 27.0846 | 26.1256 | 25.9939 | 26.6551 |
| Splicing factor U2AF 65 kDa subunit | P26368-2;P26368 | U2AF2 | Other | -1.679 | 1.624845278 | 29.7474 | 29.4055 | 29.5505 | 27.3059 | 27.9999 | 28.3606 |
| U2 snRNP-associated SURP motif-containing protein | O15042-2;O15042;O15042-3 | U2SURP | Other | -1.617166667 | 0.908740206 | 27.7164 | 25.7436 | 27.2286 | 26.2463 | 24.2371 | 25.3537 |
| UDP-N-acetylhexosamine pyrophosphorylase;UDP-N-ace | Q16222-3;Q16222;Q16222-2 | UAP1 | Extracellular or secreted | -0.577866667 | 0.641408128 | 28.412 | 28.8483 | 28.8576 | 27.5518 | 28.095 | 28.7375 |
| Ubiquitin-like modifier-activating enzyme 1 | P22314-2;P22314 | UBA1 | Extracellular or secreted | -0.685533333 | 0.736732062 | 31.725 | 32.2683 | 32.3981 | 30.8502 | 31.4224 | 32.0622 |
| SUMO-activating enzyme subunit 2 | Q9UBT2;Q9UBT2-2 | UBA2 | Other | -0.521466667 | 0.530166112 | 28.4743 | 28.597 | 29.3262 | 27.664 | 28.3411 | 28.828 |
| NEDD8-activating enzyme E1 catalytic subunit | Q8TBC4;Q8TBC4-2 | UBA3 | Other | -1.070866667 | 0.945846209 | 28.05 | 28.3129 | 28.536 | 26.3961 | 27.5658 | 27.7244 |
| Ubiquitin-like modifier-activating enzyme 5 | Q9GZZ9-2;Q9GZZ9 | UBA5 | Other | 2.0543 | 2.806394246 | 21.2321 | 21.2975 | 21.0797 | 22.9652 | 23.4223 | 23.3847 |
| Ubiquitin-like modifier-activating enzyme 6 | A0AVT1;A0AVT1-2 | UBA6 | Other | -1.2756 | 0.720583666 | 27.6048 | 27.4591 | 28.5085 | 25.3992 | 26.6154 | 27.731 |
| Ubiquitin-associated protein 2 | Q5T6F2 | UBAP2 | Extracellular or secreted | -1.015333333 | 0.420621271 | 23.2831 | 23.0777 | 23.8619 | 22.0652 | 24.0988 | 21.0127 |
| Ubiquitin-associated protein 2-like | Q14157-4;Q14157-1;Q14157 | UBAP2L | Other | 2.066366667 | 0.537425201 | 20.8684 | 20.0665 | 24.6619 | 22.3789 | 24.3292 | 25.0878 |
| Ubiquitin-conjugating enzyme E2 A | P49459;P49459-3;P49459-2 | UBE2A | Other | -0.3555 | 0.186858989 | 25.7846 | 26.301 | 24.2372 | 25.7228 | 25.0182 | 24.5153 |
| Ubiquitin-conjugating enzyme E2 D2;Ubiquitin-conjugati | P62837-2;P62837;P61077;P6 | UBE2D2;UBE2D | Extracellular or secreted | -2.646633333 | 2.300209438 | 27.5174 | 27.8519 | 27.0314 | 24.9903 | 25.3225 | 24.148 |
| Ubiquitin-conjugating enzyme E2 E2;Ubiquitin-conjugati | Q96LR5;Q969T4 | UBE2E2;UBE2E | Other | -1.523066667 | 0.613770221 | 25.1468 | 24.6407 | 24.8178 | 23.5813 | 24.8355 | 21.6193 |
| Ubiquitin-conjugating enzyme E2 G1;Ubiquitin-conjugati | P62253 | UBE2G1 | Extracellular or secreted | -0.841833333 | 1.82630314 | 25.5612 | 25.2411 | 25.105 | 24.2165 | 24.741 | 24.4243 |
| Ubiquitin-conjugating enzyme E2 G2 | P60604-2;P60604 | UBE2G2 | Other | -1.257766667 | 0.686395385 | 25.4038 | 25.9117 | 26.343 | 23.3151 | 24.8744 | 25.6957 |
| Ubiquitin-conjugating enzyme E2 H | P62256-2;P62256 | UBE2H | Other | 0.0475 | 0.045114194 | 25.9656 | 26.2248 | 25.1214 | 25.9482 | 25.8397 | 25.6664 |
| SUMO-conjugating enzyme UBC9 | P63279 | UBE2I | Other | -0.3912 | 0.810074856 | 29.4361 | 29.5403 | 28.9698 | 28.9811 | 29.1192 | 28.6723 |
| Ubiquitin-conjugating enzyme E2 K | P61086;P61086-2;P61086-3 | UBE2K | Extracellular or secreted | -0.911233333 | 2.213781782 | 27.9306 | 28.2568 | 27.8272 | 26.8877 | 27.2028 | 27.1904 |
| Ubiquitin-conjugating enzyme E2 L3 | P68036-2;P68036;P68036-3; | UBE2L3 | Extracellular or secreted | -0.6404 | 0.82045966 | 29.1798 | 29.5912 | 29.134 | 28.0937 | 29.0971 | 28.793 |
| NEDD8-conjugating enzyme Ubc12 | P61081 | UBE2M | Extracellular or secreted | -0.793866667 | 0.806793829 | 27.6591 | 28.3343 | 27.6031 | 26.3387 | 27.3726 | 27.5036 |
| Ubiquitin-conjugating enzyme E2 N | P61088 | UBE2N | Extracellular or secreted | -0.6414 | 0.666813016 | 29.2929 | 29.3832 | 29.2932 | 27.9789 | 29.1618 | 28.9044 |
| E2/E3 hybrid ubiquitin-protein ligase UBE2O | Q9C0C9 | UBE2O | Other | -1.134866667 | 0.494892543 | 24.5118 | 24.5762 | 25.2432 | 21.9126 | 24.3861 | 24.6279 |
| Ubiquitin-conjugating enzyme E2 R2 | Q712K3 | UBE2R2 | Other | 0.060033333 | 0.078794731 | 26.4775 | 25.779 | 25.8167 | 26.3465 | 25.8854 | 26.0214 |
| Ubiquitin-conjugating enzyme E2 S | Q16763 | UBE2S | Other | -1.426766667 | 0.545656335 | 25.8851 | 25.0854 | 24.0449 | 21.6434 | 24.4439 | 24.6478 |
| Ubiquitin-conjugating enzyme E2 T | Q9NPD8 | UBE2T | Other | -1.1566 | 0.748517397 | 25.6218 | 25.1338 | 24.8496 | 22.8721 | 24.5644 | 24.6989 |
| Ubiquitin-conjugating enzyme E2 variant 1 | Q13404;Q13404-7;Q13404-2 | UBE2V1 | Extracellular or secreted | -0.8652 | 1.495735325 | 29.4572 | 29.2888 | 29.3936 | 28.1861 | 28.5638 | 28.7941 |
| Ubiquitin-conjugating enzyme E2 variant 2 | Q15819 | UBE2V2 | Extracellular or secreted | -0.352366667 | 0.420471503 | 27.321 | 27.1445 | 27.119 | 26.5727 | 27.4703 | 26.4844 |
| Ubiquitin-conjugating enzyme E2 Z | Q9H832-2;Q9H832 | UBE2Z | Other | 0.837766667 | 0.268814017 | 21.1847 | 23.0671 | 24.7001 | 22.4534 | 24.2613 | 24.7505 |
| Ubiquitin-protein ligase E3A | Q05086-2;Q05086-3;Q05086 | UBE3A | Other | -1.927466667 | 0.83175165 | 26.5941 | 26.7498 | 27.8477 | 23.6404 | 25.0779 | 26.6909 |
| Ubiquitin conjugation factor E4 A | Q14139;Q14139-2 | UBE4A | Other | -0.418066667 | 0.162193558 | 23.3257 | 23.6721 | 23.9277 | 21.6168 | 23.3499 | 24.7046 |
| Ubiquitin conjugation factor E4 B | O95155-2;O95155;O95155-4 | UBE4B | Other | 0.245533333 | 0.154823438 | 21.231 | 22.8312 | 22.2053 | 23.0595 | 22.0185 | 21.9261 |
| Ubiquitin domain-containing protein UBFD1 | O14562 | UBFD1 | Other | -1.066 | 0.618414805 | 23.7108 | 23.2794 | 23.2653 | 21.1007 | 23.2988 | 22.658 |
| Ubiquitin-like protein 4A | P11441 | UBL4A | Other | 0.170266667 | 0.075265232 | 23.9115 | 24.6674 | 23.8061 | 22.9424 | 24.5692 | 25.3842 |
| Ubiquitin-like domain-containing CTD phosphatase 1 | Q8WVY7 | UBLCP1 | Other | -0.431933333 | 1.056720191 | 25.4648 | 25.5465 | 25.0916 | 24.8032 | 24.806 | 25.1979 |
| Ubiquilin-1 | Q9UMX0;Q9UMX0-2;Q9UM) | UBQLN1 | Extracellular or secreted | -0.4692 | 0.726883954 | 29.2916 | 29.3514 | 29.3579 | 28.4303 | 28.9089 | 29.2541 |
| Ubiquilin-2 | Q9UHD9 | UBQLN2 | Extracellular or secreted | -0.2871 | 0.346499085 | 26.0914 | 26.1277 | 25.4884 | 25.0985 | 25.7337 | 26.014 |
| Ubiquilin-4 | Q9NRR5 | UBQLN4 | Other | -0.757233333 | 1.367414071 | 25.8158 | 25.1543 | 25.2932 | 24.379 | 24.8267 | 24.7859 |
| E3 ubiquitin-protein ligase UBR1 | Q8IWV7;Q8IWV7-2 | UBR1 | Other | 0.240766667 | 0.270605689 | 24.2615 | 24.7737 | 24.698 | 24.7878 | 24.3031 | 25.3646 |
| E3 ubiquitin-protein ligase UBR4 | Q5T4S7-3;Q5T4S7-4;Q5T4S7 | UBR4 | Other | -0.449833333 | 0.511079312 | 29.6479 | 29.4898 | 30.349 | 29.0578 | 29.1407 | 29.9387 |
| Putative E3 ubiquitin-protein ligase UBR7 | Q8N806 | UBR7 | Other | 0.2202 | 0.368043147 | 23.7294 | 23.5728 | 23.4435 | 23.6737 | 24.2306 | 23.502 |
| Nucleolar transcription factor 1 | P17480-2;P17480 | UBTF | Other | -0.266233333 | 0.858122623 | 25.0578 | 24.7468 | 24.7033 | 24.4011 | 24.7002 | 24.6079 |
| UBX domain-containing protein 1 | Q04323;Q04323-2 | UBXN1 | Other | 0.050666667 | 0.020542955 | 24.568 | 25.3575 | 26.812 | 24.66 | 26.0501 | 26.1794 |
| UBX domain-containing protein 4 | Q92575 | UBXN4 | Other | -1.087566667 | 0.97252627 | 26.1321 | 25.9818 | 26.7179 | 24.3563 | 25.4807 | 25.7321 |
| UBX domain-containing protein 7 | O94888 | UBXN7 | Other | -0.032666667 | 0.016526729 | 23.4041 | 24.7053 | 24.557 | 23.1734 | 24.7389 | 24.6561 |
| Ubiquitin carboxyl-terminal hydrolase isozyme L3 | P15374 | UCHL3 | Extracellular or secreted | 1.3233 | 2.316115674 | 29.4473 | 28.9937 | 29.5308 | 30.8765 | 30.7362 | 30.329 |
| Ubiquitin carboxyl-terminal hydrolase isozyme L5 | Q9Y5K5-2;Q9Y5K5-4;Q9Y5K5 | UCHL5 | Other | -0.6926 | 0.403107102 | 27.2928 | 26.9184 | 28.298 | 26.0063 | 26.4691 | 27.956 |
| Ubiquitin-fold modifier-conjugating enzyme 1 | Q9Y3C8 | UFC1 | Extracellular or secreted | 0.239066667 | 0.467723081 | 24.5986 | 25.1319 | 24.5464 | 24.9242 | 25.1969 | 24.873 |
| Ubiquitin fusion degradation protein 1 homolog | Q92890;Q92890-1;Q92890-3 | UFD1L | Other | -0.4183 | 0.976178427 | 27.9359 | 27.5582 | 27.4369 | 27.0013 | 27.4575 | 27.2173 |
| E3 UFM1-protein ligase 1 | O94874;O94874-2;O94874-3 | UFL1 | Other | -0.921133333 | 0.209488197 | 22.1388 | 23.7455 | 27.3658 | 22.7656 | 23.425 | 24.2961 |
| Ubiquitin-fold modifier 1 | P61960;P61960-2 | UFM1 | Extracellular or secreted | 0.2723 | 0.305467016 | 27.317 | 27.8734 | 27.026 | 27.226 | 28.1416 | 27.6657 |
| UDP-glucose 6-dehydrogenase | O60701-2;O60701;O60701-3 | UGDH | Extracellular or secreted | 0.1876 | 0.054404528 | 25.4363 | 25.7529 | 26.9459 | 24.2039 | 26.7159 | 27.7781 |
| UDP-glucose:glycoprotein glucosyltransferase 1 | Q9NYU2-2;Q9NYU2 | UGGT1 | Extracellular or secreted | 0.991633333 | 0.373214048 | 24.0288 | 24.6723 | 27.1287 | 25.4778 | 26.0762 | 27.2507 |
| UTP--glucose-1-phosphate uridylyltransferase | Q16851 | UGP2 | Extracellular or secreted | -0.6636 | 1.54596076 | 31.4149 | 31.6653 | 31.7259 | 30.7122 | 30.8748 | 31.2283 |
| UTP--glucose-1-phosphate uridylyltransferase | Q16851-2 | UGP2 | Extracellular or secreted | 0.1719 | 0.225770116 | 24.3939 | 24.4661 | 24.7464 | 24.9626 | 24.1788 | 24.9807 |
| E3 ubiquitin-protein ligase UHRF1 | Q96T88;Q96T88-2 | UHRF1 | Extracellular or secreted | -1.1479 | 1.719624311 | 24.2291 | 24.4712 | 23.7071 | 22.8436 | 23.3753 | 22.7448 |
| NKG2D ligand 2;Retinoic acid early transcript 1G protein; | Q9BZM5;Q6H3X3-2;Q5VY80 | ULBP2;RAET1G | Extracellular or secreted | 1.849966667 | 1.352915226 | 27.7322 | 26.6695 | 26.0465 | 28.9891 | 28.8483 | 28.1607 |
| Uridine 5-monophosphate synthase;Orotate phosphorib | P11172-2;P11172-3;P11172 | UMPS | Other | -2.250933333 | 0.995289961 | 25.3089 | 24.4168 | 23.9012 | 21.7173 | 21.1895 | 23.9673 |
| Protein unc-45 homolog A | Q9H3U1-2;Q9H3U1 | UNC45A | Other | -0.275433333 | 0.11298752 | 25.4575 | 26.2342 | 27.7619 | 25.5613 | 25.7298 | 27.3362 |
| Regulator of nonsense transcripts 1 | Q92900-2;Q92900 | UPF1 | Other | -0.697866667 | 0.233918276 | 25.591 | 25.7432 | 28.0809 | 24.4934 | 25.4444 | 27.3837 |
| Regulator of nonsense transcripts 2 | Q9HAU5;Q9HAU5-2 | UPF2 | Other | -0.2459 | 0.151721082 | 22.4218 | 23.0678 | 24.1548 | 23.3167 | 22.3365 | 23.2535 |
| Uridine phosphorylase 1 | Q16831 | UPP1 | Other | 1.2727 | 0.389417706 | 21.6601 | 25.3988 | 25.2496 | 24.8624 | 25.7428 | 25.5214 |
| Cytochrome b-c1 complex subunit 1, mitochondrial | P31930 | UQCRC1 | Other | -1.046 | 0.39498016 | 25.8451 | 25.2691 | 28.4908 | 25.4586 | 25.829 | 25.1794 |
| Cytochrome b-c1 complex subunit Rieske, mitochondrial | P47985 | UQCRFS1 | Other | -2.424633333 | 1.032656031 | 24.0664 | 24.4237 | 26.7221 | 23.1904 | 23.5121 | 21.2358 |
| Cytochrome b-c1 complex subunit 6, mitochondrial | P07919 | UQCRH | Other | -1.069533333 | 0.564985711 | 27.7512 | 26.5138 | 28.5544 | 27.6869 | 26.2308 | 25.6931 |
| Uroporphyrinogen decarboxylase | P06132 | UROD | Other | -0.2879 | 0.313485122 | 27.6602 | 28.1162 | 27.8797 | 26.9394 | 27.9884 | 27.8646 |
| General vesicular transport factor p115 | O60763;O60763-2 | USO1 | Other | -0.062433333 | 0.030103856 | 28.5751 | 28.5024 | 29.5786 | 27.8542 | 28.7123 | 29.9023 |
| Ubiquitin carboxyl-terminal hydrolase 10 | Q14694;Q14694-3;Q14694-2 | USP10 | Other | -1.1526 | 0.717525109 | 23.7653 | 23.7775 | 25.3886 | 22.7176 | 22.6055 | 24.1505 |
| Ubiquitin carboxyl-terminal hydrolase 11 | P51784 | USP11 | Extracellular or secreted | 1.282433333 | 1.035221994 | 25.637 | 27.0331 | 25.6301 | 26.9615 | 27.8257 | 27.3603 |
| Ubiquitin carboxyl-terminal hydrolase 14 | P54578;P54578-2;P54578-3 | USP14 | Extracellular or secreted | -0.085333333 | 0.171248403 | 28.5915 | 28.4846 | 28.6241 | 28.175 | 28.4964 | 28.7728 |
| Ubiquitin carboxyl-terminal hydrolase 15 | Q9Y4E8-2;Q9Y4E8-3;Q9Y4E8 | USP15 | Other | -0.381 | 0.971533474 | 26.5322 | 26.2906 | 26.6421 | 25.825 | 26.1901 | 26.3068 |
| Ubiquitin carboxyl-terminal hydrolase 24 | Q9UPU5 | USP24 | Other | -1.832133333 | 0.748455524 | 24.9207 | 26.2554 | 27.9326 | 23.426 | 24.3823 | 25.804 |
| U4/U6.U5 tri-snRNP-associated protein 2 | Q53GS9;Q53GS9-3;Q53GS9- | USP39 | Other | -1.0102 | 0.508894955 | 26.5449 | 27.0364 | 27.6894 | 24.6117 | 26.6041 | 27.0243 |
| Ubiquitin carboxyl-terminal hydrolase 47 | Q96K76-2;Q96K76-4;Q96K76 | USP47 | Other | -0.167166667 | 0.07858304 | 21.6349 | 21.6872 | 23.7459 | 22.4756 | 21.9463 | 22.1446 |
| Ubiquitin carboxyl-terminal hydrolase 5 | P45974-2;P45974 | USP5 | Other | 0.466066667 | 0.348058741 | 28.604 | 27.8365 | 29.4704 | 28.7605 | 28.9347 | 29.6139 |
| Ubiquitin carboxyl-terminal hydrolase 7 | Q93009-3;Q93009 | USP7 | Other | -2.145733333 | 0.4161395 | 28.4745 | 28.8301 | 29.0333 | 22.757 | 28.5728 | 28.5709 |
| Ubiquitin carboxyl-terminal hydrolase 8 | P40818-2;P40818 | USP8 | Extracellular or secreted | -1.192633333 | 0.437523967 | 23.2791 | 22.2196 | 23.8372 | 22.7527 | 19.9208 | 23.0845 |
| Probable ubiquitin carboxyl-terminal hydrolase FAF-X;Pr | Q93008-1;Q93008;O00507 | USP9X;USP9Y | Other | -0.328533333 | 0.073333235 | 24.77 | 24.1853 | 26.9225 | 25.7839 | 22.3956 | 26.7127 |
| U3 small nucleolar RNA-associated protein 15 homolog | Q8TED0-3;Q8TED0;Q8TED0- | UTP15 | Other | -1.923233333 | 1.104734744 | 24.4583 | 23.7548 | 25.9301 | 22.7088 | 23.3065 | 22.3582 |
| Something about silencing protein 10 | Q9NQZ2 | UTP3 | Other | -0.0425 | 0.005477201 | 19.3765 | 24.743 | 25.9174 | 22.5966 | 21.0668 | 26.246 |
| Utrophin | P46939;P46939-2 | UTRN | Extracellular or secreted | -0.974833333 | 0.902226088 | 23.8496 | 24.4072 | 24.459 | 22.4561 | 23.5475 | 23.7877 |
| UDP-glucuronic acid decarboxylase 1 | Q8NBZ7;Q8NBZ7-2;Q8NBZ7- | UXS1 | Extracellular or secreted | -0.083166667 | 0.020734531 | 22.6413 | 23.1055 | 22.7184 | 20.4249 | 23.0503 | 24.7405 |
| Vesicle-associated membrane protein 3 | Q15836 | VAMP3 | Extracellular or secreted | 1.566533333 | 1.671922143 | 24.4558 | 23.8945 | 24.6588 | 26.4894 | 25.8617 | 25.3576 |
| Vesicle-associated membrane protein 7 | P51809;P51809-3;P51809-2 | VAMP7 | Extracellular or secreted | -0.041366667 | 0.074591655 | 24.8854 | 24.6478 | 25.2013 | 24.7479 | 25.0843 | 24.7782 |
| Vesicle-associated membrane protein 8 | Q9BV40 | VAMP8 | Extracellular or secreted | 1.832 | 0.825754076 | 24.8207 | 22.1019 | 24.0648 | 25.7066 | 25.4912 | 25.2856 |
| Vesicle-associated membrane protein-associated protein | Q9P0L0;Q9P0L0-2 | VAPA | Extracellular or secreted | -0.782266667 | 1.033626312 | 29.1432 | 28.9014 | 29.1025 | 27.743 | 28.6467 | 28.4106 |
| Vesicle-associated membrane protein-associated protein | O95292 | VAPB | Other | -1.083466667 | 0.624585242 | 27.8341 | 27.3688 | 29.54 | 26.793 | 27.0244 | 27.6751 |
| Valine--tRNA ligase | P26640 | VARS | Other | -0.475933333 | 0.367658256 | 29.2462 | 29.2246 | 30.553 | 28.7595 | 29.0555 | 29.781 |
| Vasorin | Q6EMK4 | VASN | Extracellular or secreted | 0.9846 | 0.564029186 | 23.9402 | 22.1358 | 21.9854 | 23.3924 | 24.5074 | 23.1154 |
| Vasodilator-stimulated phosphoprotein | P50552 | VASP | Extracellular or secreted | -0.203833333 | 0.161803974 | 28.3076 | 28.9748 | 28.7054 | 27.8093 | 28.3319 | 29.2351 |
| Synaptic vesicle membrane protein VAT-1 homolog | Q99536;Q99536-3;Q99536-2 | VAT1 | Extracellular or secreted | -0.014833333 | 0.013553014 | 29.0475 | 29.0827 | 29.5962 | 28.7423 | 29.1376 | 29.802 |
| Guanine nucleotide exchange factor VAV2 | P52735-3;P52735-2;P52735 | VAV2 | Extracellular or secreted | 0.515966667 | 0.388818509 | 25.9239 | 27.5938 | 27.1233 | 27.2982 | 27.3867 | 27.504 |
| Prefoldin subunit 3 | P61758 | VBP1 | Other | -0.370833333 | 1.027666076 | 29.1389 | 28.9328 | 28.8681 | 28.3648 | 28.8342 | 28.6283 |
| Versican core protein | P13611-2;P13611-5;P13611; | VCAN | Extracellular or secreted | -0.193566667 | 0.033679285 | 28.9104 | 29.3247 | 23.6517 | 26.804 | 27.2654 | 27.2367 |
| Vinculin | P18206-2;P18206 | VCL | Extracellular or secreted | -0.034533333 | 0.073904855 | 31.6694 | 31.4593 | 31.1595 | 31.5069 | 31.3516 | 31.3261 |
| Transitional endoplasmic reticulum ATPase | P55072 | VCP | Extracellular or secreted | 0.1234 | 0.529328377 | 33.4615 | 33.3556 | 33.6537 | 33.692 | 33.5487 | 33.6003 |
| Voltage-dependent anion-selective channel protein 1 | P21796 | VDAC1 | Extracellular or secreted | 3.342966667 | 3.472842605 | 23.1853 | 22.9015 | 22.6304 | 26.6661 | 26.0894 | 25.9906 |
| Voltage-dependent anion-selective channel protein 2 | P45880-2;P45880;P45880-1 | VDAC2 | Extracellular or secreted | 3.129133333 | 2.372939571 | 20.8473 | 21.1878 | 22.1209 | 25.1816 | 24.4757 | 23.8861 |
| Voltage-dependent anion-selective channel protein 3 | Q9Y277;Q9Y277-2 | VDAC3 | Extracellular or secreted | 2.2794 | 1.499022205 | 22.1717 | 21.7328 | 21.3606 | 25.0632 | 23.4343 | 23.6058 |
| Vascular endothelial growth factor A | P15692-5;P15692-8;P15692- | VEGFA | Extracellular or secreted | 2.050233333 | 1.226937822 | 26.4097 | 26.1626 | 25.2123 | 28.914 | 28.213 | 26.8083 |
| Vascular endothelial growth factor C | P49767 | VEGFC | Extracellular or secreted | 1.751233333 | 0.641255435 | 26.8747 | 28.6346 | 25.0588 | 28.1847 | 28.6834 | 28.9537 |
| Neurosecretory protein VGF;Neuroendocrine regulatory | O15240 | VGF | Extracellular or secreted | 2.0361 | 1.743041527 | 25.2135 | 25.4892 | 24.1726 | 27.1654 | 27.4366 | 26.3816 |
| Vimentin | P08670 | VIM | Extracellular or secreted | 3.453966667 | 1.192030523 | 28.8655 | 25.6015 | 27.6063 | 30.5427 | 30.7925 | 31.1 |
| Protein VPRBP | Q9Y4B6-3;Q9Y4B6-2;Q9Y4B6 | VPRBP | Other | 0.836666667 | 0.459082962 | 24.7164 | 24.4709 | 24.6268 | 26.7627 | 25.1096 | 24.4518 |
| Vacuolar protein sorting-associated protein 26A | O75436;O75436-2 | VPS26A | Extracellular or secreted | -0.621133333 | 1.078417547 | 28.6798 | 28.881 | 28.8028 | 27.7583 | 28.3628 | 28.3791 |
| Vacuolar protein sorting-associated protein 26B | Q4G0F5 | VPS26B | Other | -1.540333333 | 0.831054618 | 23.2926 | 22.9505 | 23.3305 | 22.292 | 22.3671 | 20.2935 |
| Vacuolar protein sorting-associated protein 28 homolog | Q9UK41;Q9UK41-2 | VPS28 | Extracellular or secreted | 0.3735 | 0.264084127 | 23.8811 | 25.2085 | 23.7696 | 24.1324 | 25.2118 | 24.6355 |
| Vacuolar protein sorting-associated protein 29 | Q9UBQ0;Q9UBQ0-2 | VPS29 | Extracellular or secreted | -1.0347 | 0.730283751 | 27.668 | 27.5034 | 26.8309 | 25.3349 | 26.363 | 27.2003 |
| Vacuolar protein sorting-associated protein 35 | Q96QK1 | VPS35 | Extracellular or secreted | -1.118 | 0.616111514 | 28.1791 | 29.1286 | 29.577 | 26.6417 | 27.93 | 28.959 |
| Vacuolar protein sorting-associated protein 45 | Q9NRW7 | VPS45 | Other | -1.851766667 | 0.457309095 | 26.6337 | 25.6239 | 26.3027 | 21.3334 | 25.3395 | 26.3321 |
| Vacuolar protein sorting-associated protein 4A | Q9UN37 | VPS4A | Extracellular or secreted | -0.350966667 | 0.135609581 | 20.1893 | 23.1508 | 21.4126 | 21.7095 | 20.615 | 21.3753 |
| Vacuolar protein sorting-associated protein 4B | O75351 | VPS4B | Extracellular or secreted | 0.364733333 | 0.292995678 | 23.707 | 23.9499 | 25.0398 | 24.534 | 24.1412 | 25.1157 |
| Serine/threonine-protein kinase VRK1 | Q99986 | VRK1 | Other | -1.086766667 | 0.472608381 | 22.3076 | 20.8306 | 23.8357 | 20.8296 | 21.7173 | 21.1667 |
| Vacuolar protein sorting-associated protein VTA1 homol | Q9NP79;Q9NP79-2 | VTA1 | Extracellular or secreted | 0.253633333 | 0.171504686 | 25.001 | 26.084 | 26.7184 | 25.8064 | 26.4164 | 26.3415 |
| von Willebrand factor A domain-containing protein 9 | Q96SY0-4;Q96SY0-3;Q96SY0 | VWA9 | Other | -3.658 | 3.643345524 | 25.1328 | 24.8962 | 25.0407 | 21.2927 | 21.1632 | 21.6398 |
| Tryptophan--tRNA ligase, cytoplasmic;T1-TrpRS;T2-TrpRS | P23381;P23381-2 | WARS | Extracellular or secreted | 0.2807 | 0.706152067 | 29.5196 | 29.3204 | 29.5039 | 29.4515 | 29.7613 | 29.9732 |
| Wiskott-Aldrich syndrome protein family member 2 | Q9Y6W5;Q9Y6W5-2 | WASF2 | Extracellular or secreted | 0.286366667 | 0.363416621 | 25.2973 | 26.146 | 26.1891 | 26.3384 | 26.1628 | 25.9903 |
| Neural Wiskott-Aldrich syndrome protein | O00401 | WASL | Extracellular or secreted | -0.608533333 | 0.233641352 | 23.6283 | 24.0692 | 24.732 | 21.7339 | 24.1319 | 24.7381 |
| WW domain-binding protein 11 | Q9Y2W2 | WBP11 | Other | -0.195433333 | 0.18322608 | 27.0087 | 26.4103 | 26.843 | 25.8703 | 26.7641 | 27.0413 |

| | | | | | | | | | | | |
|---|---|---|---|---|---|---|---|---|---|---|---|
| WW domain-binding protein 4 | O75554-2;O75554 | WBP4 | Extracellular or secreted | -1.132033333 | 0.38430012 | 24.0331 | 24.124 | 24.1638 | 25.1812 | 21.8397 | 21.9039 |
| Williams-Beuren syndrome chromosomal region 16 prot | Q96I51-3;Q96I51;Q96I51-2 | WBSCR16 | Other | 0.351266667 | 0.405416337 | 22.5683 | 22.3787 | 23.1757 | 22.7538 | 22.8114 | 23.6113 |
| WD repeat and HMG-box DNA-binding protein 1 | O75717;O75717-2 | WDHD1 | Other | -0.807366667 | 1.420883456 | 27.2925 | 26.8721 | 27.398 | 26.3314 | 26.0563 | 26.7528 |
| WD repeat-containing protein 1 | O75083;O75083-3 | WDR1 | Extracellular or secreted | -0.2431 | 0.225913021 | 31.2619 | 32.2212 | 31.9391 | 30.9673 | 31.7207 | 32.0049 |
| Ribosome biogenesis protein WDR12 | Q9GZL7 | WDR12 | Other | -0.992933333 | 1.163169898 | 30.7788 | 29.8469 | 30.0632 | 29.6763 | 29.3339 | 28.6999 |
| WD repeat-containing protein 26 | Q9H7D7-2;Q9H7D7 | WDR26 | Other | -0.302166667 | 0.408075196 | 27.1847 | 26.9449 | 27.7361 | 26.5745 | 27.1326 | 27.2521 |
| WD repeat-containing protein 3 | Q9UNX4 | WDR3 | Other | -3.8013 | 2.177059346 | 24.6199 | 24.5561 | 26.2404 | 22.1196 | 20.9774 | 20.9155 |
| WD repeat-containing protein 36 | Q8NI36 | WDR36 | Other | -0.099833333 | 0.039568417 | 21.468 | 21.3228 | 23.5842 | 22.8452 | 21.4338 | 21.7965 |
| WD repeat-containing protein 43 | Q15061 | WDR43 | Other | -2.9532 | 1.473748198 | 24.0723 | 24.5025 | 25.6857 | 21.1285 | 21.0306 | 23.2418 |
| WD repeat-containing protein 44 | Q5JSH3-2;Q5JSH3;Q5JSH3-4 | WDR44 | Other | 0.5652 | 0.852879767 | 23.6618 | 23.2683 | 24.1388 | 24.0527 | 24.2705 | 24.4413 |
| WD repeat domain phosphoinositide-interacting protein | Q5MNZ6 | WDR45B | Other | -0.239166667 | 0.201488321 | 24.6405 | 25.4653 | 24.1208 | 24.069 | 24.7223 | 24.7178 |
| WD repeat-containing protein 48 | Q8TAF3-4;Q8TAF3-3;Q8TAF3 | WDR48 | Other | -0.2705 | 0.266015091 | 24.8668 | 24.6936 | 23.8434 | 24.6951 | 24.0198 | 23.8774 |
| WD repeat-containing protein 5 | P61964 | WDR5 | Other | -1.064033333 | 1.624419189 | 29.4934 | 28.9534 | 29.0016 | 28.259 | 28.3674 | 27.6299 |
| WD repeat-containing protein 61;WD repeat-containing | Q9GZS3 | WDR61 | Other | -0.010833333 | 0.009934088 | 25.5011 | 25.7183 | 26.2409 | 25.2519 | 26.0064 | 26.1695 |
| WD repeat-containing protein 75 | Q8IWA0 | WDR75 | Other | -0.9163 | 0.415805515 | 24.8739 | 23.1913 | 25.5562 | 24.1285 | 24.3498 | 22.3942 |
| Methylosome protein 50 | Q9BQA1;Q9BQA1-2 | WDR77 | Other | -0.425766667 | 3.101093589 | 28.3171 | 28.3616 | 28.2496 | 27.9126 | 27.8174 | 27.921 |
| WD repeat-containing protein 82 | Q6UXN9 | WDR82 | Other | -0.947333333 | 1.056259818 | 28.2618 | 28.7545 | 28.4211 | 26.9096 | 28.0486 | 27.6372 |
| WD repeat-containing protein 89 | Q96FK6 | WDR89 | Other | -1.473733333 | 1.003050967 | 23.5025 | 22.8107 | 24.696 | 22.056 | 22.6765 | 21.8555 |
| WD repeat-containing protein 92 | Q96MX6-2;Q96MX6 | WDR92 | Other | -1.753166667 | 0.993097915 | 23.9993 | 23.5039 | 23.9603 | 22.1251 | 23.1348 | 20.9441 |
| WAP four-disulfide core domain protein 2 | Q14508;Q14508-2;Q14508-3 | WFDC2 | Extracellular or secreted | 1.403133333 | 1.592518965 | 30.8189 | 31.5193 | 30.4683 | 32.7565 | 32.3422 | 31.9172 |
| Wolframin | O76024 | WFS1 | Other | -1.069166667 | 1.095849047 | 25.6302 | 24.4689 | 24.6812 | 23.7572 | 24.1403 | 23.6753 |
| Partner of Y14 and mago | Q9BRP8-2;Q9BRP8 | WIBG | Other | 1.483266667 | 0.586885731 | 24.9192 | 21.7379 | 24.1431 | 24.9024 | 25.5001 | 24.8475 |
| WAS/WASL-interacting protein family member 1 | O43516-2;O43516;O43516-3 | WIPF1 | Other | -0.115866667 | 0.063468683 | 23.113 | 23.4586 | 21.7521 | 22.7325 | 22.0135 | 23.2301 |
| Serine/threonine-protein kinase WNK1 | Q9H4A3-2;Q9H4A3;Q9H4A3 | WNK1 | Extracellular or secreted | -1.430466667 | 1.011450718 | 26.2594 | 25.377 | 27.2229 | 24.2334 | 25.0669 | 25.2676 |
| Protein Wnt-5a | P41221-2;P41221 | WNT5A | Extracellular or secreted | -0.948333333 | 0.33884101 | 23.8813 | 25.2766 | 24.2484 | 21.4992 | 24.6865 | 24.3756 |
| Telomerase Cajal body protein 1 | Q9BUR4 | WRAP53 | Other | -0.136733333 | 0.219370446 | 23.3236 | 23.4784 | 23.2894 | 22.9443 | 23.6621 | 23.0748 |
| ATPase WRNIP1 | Q96S55-2;Q96S55;Q96S55-3 | WRNIP1 | Other | -0.957166667 | 0.258188686 | 26.5542 | 23.0903 | 27.3608 | 24.8766 | 23.7529 | 25.5043 |
| Pre-mRNA-splicing regulator WTAP | Q15007;Q15007-2 | WTAP | Other | -1.948733333 | 1.494114852 | 25.7948 | 25.4338 | 26.9827 | 23.5495 | 24.6807 | 24.1349 |
| Pre-mRNA-splicing factor SYF1 | Q9HCS7 | XAB2 | Other | -0.128566667 | 0.030905845 | 23.9101 | 26.4107 | 22.5812 | 24.0662 | 22.7973 | 25.6528 |
| Skin-specific protein 32 | Q5T750 | XP32 | Extracellular or secreted | 3.459233333 | 1.088113502 | 21.3837 | 22.1391 | 25.1192 | 27.0363 | 25.7526 | 26.2308 |
| Xaa-Pro aminopeptidase 1 | Q9NQW7-3;Q9NQW7;Q9NQ | XPNPEP1 | Extracellular or secreted | 0.029966667 | 0.037244633 | 29.1944 | 29.272 | 28.8219 | 29.5732 | 29 | 28.805 |
| Exportin-1 | O14980 | XPO1 | Other | -1.516133333 | 0.837450177 | 29.6922 | 30.2279 | 30.6868 | 27.4279 | 28.8376 | 29.793 |
| Exportin-4 | Q9C0E2 | XPO4 | Other | -1.2104 | 0.628187859 | 23.5255 | 24.5179 | 26.0147 | 23.0044 | 23.1674 | 24.2551 |
| Exportin-5 | Q9HAV4 | XPO5 | Other | -2.644833333 | 0.862098546 | 26.3784 | 26.4919 | 27.8241 | 22.0457 | 24.7402 | 25.974 |
| Exportin-7 | Q9UIA9 | XPO7 | Other | -0.864766667 | 0.507375014 | 26.4229 | 27.7155 | 28.2762 | 25.7035 | 26.6694 | 27.4474 |
| Exportin-T | O43592 | XPOT | Other | -1.271666667 | 0.98572912 | 26.0081 | 26.8469 | 27.3408 | 24.6553 | 25.4866 | 26.2389 |
| DNA repair protein XRCC4 | Q13426-3;Q13426-2;Q13426 | XRCC4 | Other | -0.9458 | 0.89961109 | 24.1446 | 23.458 | 23.8026 | 22.4954 | 22.4196 | 23.6528 |
| X-ray repair cross-complementing protein 5 | P13010 | XRCC5 | Extracellular or secreted | -0.9175 | 0.720109568 | 31.138 | 31.2088 | 32.181 | 30.0194 | 30.2491 | 31.5068 |
| X-ray repair cross-complementing protein 6 | P12956;P12956-2 | XRCC6 | Other | -0.954166667 | 0.700161812 | 31.1266 | 31.0352 | 32.0386 | 29.66 | 30.2852 | 31.3927 |
| 5-3 exoribonuclease 2 | Q9H0D6;Q9H0D6-2 | XRN2 | Other | -1.802966667 | 0.988924443 | 28.7579 | 28.7061 | 29.4722 | 25.9311 | 27.3489 | 28.2473 |
| Xylosyltransferase 2 | Q9H1B5;Q9H1B5-2 | XYLT2 | Extracellular or secreted | -0.0029 | 0.002461947 | 24.4574 | 24.8476 | 23.6649 | 24.0606 | 24.4893 | 24.4113 |
| Transcriptional coactivator YAP1 | P46937-5;P46937-3;P46937- | YAP1 | Other | -0.396166667 | 0.262889843 | 24.284 | 25.8842 | 24.2476 | 24.0879 | 24.4585 | 24.6809 |
| Tyrosine--tRNA ligase, cytoplasmic;Tyrosine--tRNA ligase | P54577 | YARS | Extracellular or secreted | -0.412666667 | 0.259264369 | 28.4784 | 28.6083 | 29.5752 | 27.6271 | 28.3735 | 29.4233 |
| Tyrosine--tRNA ligase, mitochondrial | Q9Y2Z4 | YARS2 | Other | -1.3542 | 0.874410117 | 21.7678 | 23.2181 | 23.7115 | 21.6344 | 21.1242 | 21.8762 |
| Nuclease-sensitive element-binding protein 1 | P67809 | YBX1 | Extracellular or secreted | 0.1557 | 0.257366051 | 31.4249 | 31.14 | 31.4849 | 31.9211 | 31.3104 | 31.2854 |
| Y-box-binding protein 3 | P16989;P16989-2;P16989-3 | YBX3 | Other | 1.0255 | 1.789273932 | 27.6982 | 27.1033 | 27.3295 | 28.7686 | 28.1435 | 28.2954 |
| Tyrosine-protein kinase Yes | P07947 | YES1 | Extracellular or secreted | 0.5276 | 1.00483071 | 24.162 | 24.0222 | 24.3588 | 25.084 | 24.6287 | 24.4131 |
| Synaptobrevin homolog YKT6 | O15498;O15498-2 | YKT6 | Extracellular or secreted | 0.5137 | 0.224807202 | 24.8605 | 24.3437 | 24.954 | 26.8442 | 24.2269 | 24.6282 |
| YLP motif-containing protein 1 | P49750;P49750-3;P49750-1 | YLPM1 | Other | -0.3486 | 0.194651161 | 23.8479 | 23.8021 | 25.3677 | 23.3802 | 23.7117 | 24.88 |
| ATP-dependent zinc metalloprotease YME1L1 | Q96TA2-3;Q96TA2-2;Q96TA | YME1L1 | Other | -0.570366667 | 1.505271034 | 25.6472 | 26.0099 | 25.6148 | 25.0087 | 25.4175 | 25.1346 |
| YTH domain-containing family protein 2 | Q9Y5A9-2;Q9Y5A9 | YTHDF2 | Other | -0.223033333 | 0.100400971 | 26.1078 | 25.7981 | 25.4983 | 26.951 | 24.4286 | 25.3555 |
| YTH domain-containing family protein 3 | Q7Z739 | YTHDF3 | Other | -2.504533333 | 0.726779371 | 25.2943 | 23.9376 | 25.6783 | 20.5219 | 21.9087 | 24.966 |
| 14-3-3 protein beta/alpha;14-3-3 protein beta/alpha, N-t | P31946-2;P31946 | YWHAB | Extracellular or secreted | 2.1183 | 2.597632059 | 30.5504 | 30.0513 | 29.8937 | 32.5467 | 32.4861 | 31.8175 |
| 14-3-3 protein epsilon | P62258;P62258-2 | YWHAE | Extracellular or secreted | 0.640033333 | 0.724352745 | 34.9541 | 34.5083 | 34.5575 | 35.9199 | 35.2634 | 34.7567 |
| 14-3-3 protein gamma;14-3-3 protein gamma, N-termina | P61981 | YWHAG | Extracellular or secreted | 0.787233333 | 0.725787069 | 32.7562 | 31.9007 | 32.1741 | 33.8742 | 32.6729 | 32.6456 |
| 14-3-3 protein eta | Q04917 | YWHAH | Extracellular or secreted | 0.7221 | 1.363712356 | 30.4864 | 30.2099 | 29.9816 | 31.2788 | 30.9479 | 30.6175 |
| 14-3-3 protein theta | P27348 | YWHAQ | Extracellular or secreted | 0.981833333 | 2.458912258 | 30.563 | 30.2249 | 30.5079 | 31.5309 | 31.5307 | 31.1797 |
| 14-3-3 protein zeta/delta | P63104;P63104-2 | YWHAZ | Extracellular or secreted | 1.412133333 | 1.496608941 | 32.9678 | 32.6925 | 32.8142 | 34.7061 | 34.2941 | 33.7107 |
| Mitogen-activated protein kinase kinase kinase MLT | Q9NYL2;Q9NYL2-3 | ZAK | Other | -1.4636 | 0.42331145 | 24.3564 | 24.6492 | 25.4209 | 20.7482 | 24.6271 | 24.6604 |
| Zinc finger and BTB domain-containing protein 7A | O95365 | ZBTB7A | Other | -0.4652 | 0.287957059 | 23.0344 | 21.4671 | 21.6755 | 22.3442 | 20.8635 | 21.5737 |
| Protein archease | Q8IWT0;Q8IWT0-2 | ZBTB8OS | Extracellular or secreted | 0.698433333 | 0.420796543 | 26.114 | 25.8904 | 25.4774 | 27.333 | 26.9407 | 25.3034 |
| Zinc finger CCCH domain-containing protein 13 | Q5T200-2;Q5T200 | ZC3H13 | Other | -2.188233333 | 1.00880521 | 27.0062 | 24.3975 | 25.6641 | 23.6131 | 23.6047 | 23.2853 |
| Zinc finger CCCH domain-containing protein 14 | Q6PJT7-5;Q6PJT7-9;Q6PJT7- | ZC3H14 | Other | -1.4959 | 0.890422634 | 24.2155 | 24.4208 | 26.0526 | 22.3711 | 23.7706 | 24.0595 |
| Zinc finger CCCH domain-containing protein 15 | Q8WU90;Q8WU90-2 | ZC3H15 | Other | -2.182933333 | 1.265588483 | 27.4385 | 25.859 | 27.9826 | 24.4581 | 24.5716 | 25.7016 |
| Zinc finger CCCH domain-containing protein 18 | Q86VM9;Q86VM9-2 | ZC3H18 | Other | -0.596133333 | 0.242838041 | 27.6629 | 26.1016 | 27.1347 | 27.9375 | 25.1264 | 26.0469 |
| Zinc finger CCCH domain-containing protein 4 | Q9UPT8 | ZC3H4 | Other | 0.8766 | 0.610008041 | 23.2932 | 24.0093 | 24.8922 | 24.0517 | 25.4907 | 25.2821 |
| Nuclear-interacting partner of ALK | Q86WB0;Q86WB0-3;Q86WB | ZC3HC1 | Other | -0.721666667 | 0.159179591 | 24.4538 | 21.09 | 24.4334 | 20.0472 | 23.9265 | 23.8385 |
| Zinc finger CCHC domain-containing protein 8 | Q6NZY4 | ZCCHC8 | Other | 0.019766667 | 0.00594882 | 21.1139 | 22.8072 | 24.5717 | 22.7113 | 23.4544 | 22.3864 |
| Palmitoyltransferase ZDHHC5 | Q9C0B5-2;Q9C0B5 | ZDHHC5 | Extracellular or secreted | 1.364433333 | 1.20125585 | 24.6577 | 23.4526 | 24.5057 | 25.6452 | 25.6635 | 25.4006 |
| AN1-type zinc finger protein 6 | Q6FIF0-2;Q6FIF0 | ZFAND6 | Other | 0.6681 | 0.565288129 | 25.4379 | 25.4994 | 24.642 | 26.6882 | 25.2448 | 25.6506 |
| E3 ubiquitin-protein ligase ZFP91 | Q96JP5-2;Q96JP5 | ZFP91 | Other | -0.732866667 | 0.647608497 | 24.578 | 23.6875 | 25.1685 | 24.0375 | 23.396 | 23.8019 |
| Zinc finger protein-like 1 | O95159 | ZFPL1 | Other | -0.213766667 | 0.065571326 | 21.4923 | 24.4923 | 23.787 | 21.7166 | 23.4853 | 23.9284 |
| Zinc finger RNA-binding protein | Q96KR1 | ZFR | Other | -2.3412 | 1.293175842 | 24.8038 | 25.4177 | 25.3755 | 22.4463 | 22.0643 | 24.0628 |
| Zinc finger protein 185 | O15231-3;O15231-2 | ZNF185 | Extracellular or secreted | -2.0609 | 0.803334664 | 23.7291 | 23.4486 | 24.559 | 21.7032 | 23.5986 | 20.2522 |
| Zinc finger protein 185 | O15231-7;O15231-4;O15231 | ZNF185 | Extracellular or secreted | -2.4071 | 0.4809648 | 25.9142 | 26.3553 | 26.71 | 20.1441 | 25.9705 | 25.6436 |
| BUB3-interacting and GLEBS motif-containing protein ZN | O43670-2;O43670;O43670-4 | ZNF207 | Extracellular or secreted | 0.0192 | 0.017963336 | 27.3075 | 28.0945 | 27.4665 | 27.1868 | 28.0922 | 27.6471 |
| DBIRD complex subunit ZNF326 | Q5BKZ1-3;Q5BKZ1 | ZNF326 | Other | -2.064266667 | 0.572910183 | 22.6842 | 22.6508 | 26.7398 | 21.7619 | 22.0339 | 22.0862 |
| Zinc finger protein 483 | Q8TF39-2;Q8TF39 | ZNF483 | Other | 2.830733333 | 1.559313101 | 21.5295 | 23.2135 | 23.5465 | 26.5135 | 24.632 | 25.6362 |
| Zinc finger protein 579 | Q8NAF0 | ZNF579 | Other | 0.057 | 0.074408973 | 21.05 | 20.753 | 20.8964 | 21.0278 | 20.5027 | 21.3399 |
| Zinc finger protein 622 | Q969S3 | ZNF622 | Other | -0.919133333 | 0.782851393 | 22.7599 | 22.5111 | 22.463 | 21.3748 | 21.0865 | 22.5153 |
| Zinc finger protein 706 | Q9Y5V0 | ZNF706 | Other | -1.1303 | 0.356832487 | 24.1405 | 25.1539 | 22.6159 | 23.9586 | 23.8633 | 20.6975 |
| Zinc finger protein ZPR1 | O75312 | ZPR1 | Other | 0.090266667 | 0.103389082 | 25.6837 | 26.0266 | 25.8852 | 25.4412 | 25.9997 | 26.4254 |
| Zinc finger Ran-binding domain-containing protein 2 | O95218-2;O95218 | ZRANB2 | Other | 0.218866667 | 0.127532703 | 28.4231 | 26.9292 | 27.0474 | 28.494 | 27.2209 | 27.3414 |
| Centromere/kinetochore protein zw10 homolog | O43264;O43264-2 | ZW10 | Other | -0.7649 | 0.184892394 | 21.6974 | 21.6928 | 24.603 | 22.0169 | 19.7007 | 23.9809 |
| Protein zwilch homolog | Q9H900-2;Q9H900 | ZWILCH | Other | -0.5883 | 0.777003941 | 22.005 | 21.8846 | 21.8343 | 21.6324 | 20.7602 | 21.5664 |
| Zyxin | Q15942;Q15942-2 | ZYX | Extracellular or secreted | -1.4461 | 0.507791145 | 26.1411 | 27.1002 | 26.5207 | 22.9929 | 26.0474 | 26.3834 |
| | P0DPI2;A0A0B4J2D5;P0DPI2-2;A0A0B4J2D5 | | NA | -1.243166667 | 1.352330668 | 30.7248 | 30.7634 | 30.8922 | 29.5263 | 30.0527 | 29.0719 |
| | P0DP25;P0DP24;P0DP23 | | NA | 0.3235 | 0.103766784 | 30.1077 | 29.6417 | 31.5085 | 32.6302 | 30.0166 | 29.5816 |
| | P0DPB6 | | NA | -0.234566667 | 0.149408918 | 22.763 | 22.4561 | 22.6979 | 22.9015 | 21.3228 | 22.989 |
| | P55196-3 | | NA | -0.309466667 | 0.22899995 | 27.0273 | 27.5225 | 28.0149 | 26.4951 | 27.1361 | 28.0051 |
| | Q16787-1;Q16787-4 | | NA | 0.048266667 | 0.018185951 | 32.8464 | 34.2478 | 31.5013 | 32.275 | 33.1861 | 33.2792 |
| | Q9UPN3-3;Q9UPN3-2 | | NA | -0.8065 | 0.514335563 | 28.0844 | 28.0104 | 28.1587 | 26.1713 | 27.4672 | 28.1955 |

Table S6. Proteins acquired by Mass spectrometer that were differentially expressed in PDX401 2D and 3D conditioned media.

| Protein.names | Majority.protein.IDs | Gene.names | Extracellular or secreted | log2_fold-change | -log10_p-value | 2D_CM1 | 2D_CM2 | 2D_CM3 | 2D_CM4 | 2D_CM5 | 2D_CM6 | 3D_CM2 | 3D_CM3 | 3D_CM6 |
|---|---|---|---|---|---|---|---|---|---|---|---|---|---|---|
| Alpha-2-macroglobulin | P01023 | A2M | Extracellular or secreted | -0.76579909 | 0.408129346 | 22.99447967 | 22.04775914 | 21.55232989 | 22.41502457 | 21.04019338 | 21.60414425 | 20.49726622 | 20.46493257 | 22.56736939 |
| Alpha-2-macroglobulin-like protein 1 | A8K2U0;A8K2U0-2 | A2ML1 | Extracellular or secreted | 3.460055327 | 0.653002011 | 21.15171276 | 23.02325959 | 20.8917837 | 21.49706826 | 25.21626777 | 20.82919624 | 23.71565371 | 23.36719716 | 29.60195927 |
| Acetoacetyl-CoA synthetase | Q86V21;Q86V21-2;Q86V21-3 | AACS | Other | 2.069280537 | 1.1986237 | 24.36365943 | 24.64921758 | 24.32051577 | 22.57604787 | 23.54146182 | 23.54674942 | 26.24083572 | 26.80906744 | 24.65676439 |
| Alpha- and gamma-adaptin-binding pr | Q6PD74 | AAGAB | Other | 0.507913905 | 0.602789988 | 23.54792627 | 23.62399577 | 23.64801081 | 21.85726971 | 24.05763285 | 23.8322905 | 24.23396074 | 24.11695942 | 23.4563845 |
| Mth938 domain-containing protein | Q9H7C9 | AAMDC | Other | -0.275257807 | 0.117304483 | 27.85869465 | 27.69272084 | 27.62623986 | 27.17245528 | 26.76900728 | 25.45268214 | 27.55603506 | 27.58412677 | 25.31996478 |
| Angio-associated migratory cell protei | Q13685 | AAMP | Extracellular or secreted | 0.565579152 | 0.992049333 | 27.27994264 | 27.39525641 | 27.31881047 | 28.19394535 | 28.46427514 | 27.01797022 | 28.44699246 | 28.2587619 | 27.82608321 |
| Protein AAR2 homolog | Q9Y312 | AAR2 | Other | 0.194127999 | 0.357798686 | 25.24230858 | 25.43523532 | 25.27198841 | 24.06931799 | 24.863677 | 24.5712638 | 25.27653906 | 24.91479022 | 25.11795027 |
| Alanine--tRNA ligase, cytoplasmic | P49588;P49588-2 | AARS | Extracellular or secreted | 1.523926608 | 3.141460855 | 29.94851611 | 30.06573816 | 29.92818328 | 30.65753121 | 30.6766546 | 29.53530617 | 31.79292692 | 31.85322358 | 31.33159409 |
| Alanyl-tRNA editing protein Aarsd1 | Q9BTE6;Q9BTE6-2;Q9BTE6-3 | AARSD1 | Other | 3.430807973 | 0.895431113 | 20.63300984 | 21.47961377 | 20.18640445 | 21.74414894 | 22.70261849 | 22.14311038 | 26.45710504 | 26.18876781 | 22.09100401 |
| L-aminoadipate-semialdehyde dehydr | Q9NRN7 | AASDHPPT | Extracellular or secreted | 0.459071802 | 0.575604002 | 25.94491916 | 26.21756254 | 26.36496206 | 26.12664052 | 25.90364082 | 26.27419442 | 26.81763091 | 26.97650906 | 25.9990352 |
| Protein AATF | Q9NY61 | AATF | Other | -3.043184743 | 1.523498177 | 24.03246214 | 24.36912924 | 24.03540188 | 23.77936509 | 24.39956531 | 24.45287062 | 20.09786546 | 22.07818444 | 21.22879301 |
| Multidrug resistance-associated prote | P33527;P33527-9;P33527-4;P | ABCC1 | Extracellular or secreted | 0.267869551 | 0.386347753 | 25.57007689 | 25.68592164 | 25.79572995 | 25.91195953 | 26.12382192 | 26.13963706 | 25.72343472 | 26.08565755 | 26.60808987 |
| ATP-binding cassette sub-family E me | P61221 | ABCE1 | Other | 0.947768041 | 1.701104653 | 28.00731511 | 27.82450597 | 28.06141825 | 28.63318693 | 28.69321952 | 28.78684197 | 29.4680295 | 29.51964603 | 28.85887247 |
| ATP-binding cassette sub-family F me | Q8NE71;Q8NE71-2 | ABCF1 | Other | -0.481621179 | 2.591411075 | 29.70361916 | 29.54329425 | 29.66910324 | 29.63649239 | 29.71654687 | 29.99050734 | 29.2099251 | 29.12373807 | 29.35125493 |
| ATP-binding cassette sub-family F me | Q9UG63;Q9UG63-2 | ABCF2 | Extracellular or secreted | -0.722632454 | 0.88535697 | 26.77492436 | 27.04603971 | 26.91113667 | 26.02254899 | 25.72917822 | 26.93145585 | 25.56421486 | 25.50682038 | 26.4687093 |
| ATP-binding cassette sub-family F me | Q9NUQ8-2;Q9NUQ8 | ABCF3 | Other | 1.081278748 | 1.163178837 | 20.75384599 | 22.58648523 | 22.69971054 | 22.87938432 | 22.97750074 | 24.2454674 | 23.89272883 | 24.01111735 | 23.41118718 |
| Mycophenolic acid acyl-glucuronide e | Q9NUJ1;Q9NUJ1-2 | ABHD10 | Extracellular or secreted | -0.166066283 | 0.263338583 | 28.73966397 | 28.75225813 | 28.6477707 | 28.39558329 | 28.51874601 | 28.79411295 | 28.69265434 | 28.71200991 | 28.02120442 |
| Alpha/beta hydrolase domain-contain | Q8NFV4-4;Q8NFV4;Q8NFV4- | ABHD11 | Other | -0.137377604 | 0.077424352 | 25.95553356 | 26.14765476 | 26.12145242 | 24.42431906 | 24.55818292 | 24.89518141 | 25.68471845 | 25.79061875 | 24.16369205 |
| Alpha/beta hydrolase domain-contain | Q96IU4 | ABHD14B | Extracellular or secreted | -0.433964165 | 1.356352803 | 28.54285163 | 28.63304832 | 28.50782615 | 28.07500281 | 28.15716164 | 27.74643939 | 27.93393355 | 27.96444082 | 27.63089809 |
| Abl interactor 1 | Q8IZP0-10;Q8IZP0-8;Q8IZP0- | ABI1 | Extracellular or secreted | -0.118518987 | 0.18769674 | 28.34511188 | 28.28091615 | 28.42522391 | 27.6063964 | 27.76540741 | 27.75078967 | 28.08901657 | 28.11877216 | 27.52357703 |
| Abl interactor 2 | Q9NYB9-2;Q9NYB9-4;Q9NYB | ABI2 | Other | 0.207943576 | 0.446390716 | 24.5383661 | 24.35258984 | 24.04576899 | 24.07651226 | 23.96163975 | 24.01852213 | 24.59383105 | 24.48928498 | 24.03741424 |
| Active breakpoint cluster region-relat | Q12979-4;Q12979-2;Q12979 | ABR | Extracellular or secreted | -0.076906804 | 0.092870758 | 25.93573284 | 26.17346594 | 26.09204511 | 25.39434075 | 24.42098305 | 25.57551132 | 25.69490046 | 25.64165864 | 25.22875999 |
| Costars family protein ABRACL | Q9P1F3 | ABRACL | Other | 0.498589205 | 0.441026564 | 27.89562058 | 27.78131769 | 27.97114567 | 28.19980644 | 28.12356542 | 26.5537522 | 28.90423251 | 28.35317059 | 27.50096852 |
| 3-ketoacyl-CoA thiolase, peroxisomal | P09110;P09110-2 | ACAA1 | Other | -0.362384388 | 0.458831516 | 25.70382375 | 25.83023646 | 26.2556951 | 24.06825215 | 25.53563989 | 26.41451363 | 25.44211914 | 25.25555104 | 25.11925715 |
| 3-ketoacyl-CoA thiolase, mitochondri | P42765 | ACAA2 | Extracellular or secreted | -0.243987155 | 0.479872482 | 27.07490076 | 26.97847429 | 26.95182629 | 26.58677727 | 26.77026827 | 27.930159 | 26.91250784 | 26.96664257 | 26.53509107 |
| Acetyl-CoA carboxylase 1;Biotin carbo | Q13085-3;Q13085-2;Q13085 | ACACA | Extracellular or secreted | 1.523553597 | 1.925426724 | 26.23392418 | 26.34126499 | 26.23377793 | 25.0542827 | 26.01830988 | 27.2131741 | 27.58046538 | 27.26400799 | 28.27355432 |
| Medium-chain specific acyl-CoA dehy | P11310;P11310-2 | ACADM | Extracellular or secreted | -0.179626327 | 0.121990729 | 24.0988462 | 23.93237986 | 24.22302479 | 24.73531224 | 24.71800811 | 25.9111824 | 23.93346087 | 24.01869191 | 25.31834503 |
| Short-chain specific acyl-CoA dehydro | P16219 | ACADS | Other | -0.513145321 | 0.618962399 | 22.22101944 | 22.09306933 | 22.59267967 | 22.84345186 | 21.82638292 | 20.71497328 | 21.7730587 | 21.80687683 | 21.02641676 |
| Short/branched chain specific acyl-Co | P45954-2;P45954 | ACADSB | Extracellular or secreted | -0.571339319 | 0.242389316 | 22.27590965 | 22.17968728 | 21.86341479 | 23.84957309 | 23.91634018 | 24.7856141 | 21.01285517 | 23.46475811 | 23.24363831 |
| Very long-chain specific acyl-CoA deh | P49748;P49748-3;P49748-2 | ACADVL | Other | -0.509005622 | 0.933956613 | 28.57192197 | 28.77445315 | 28.814395 | 28.42614419 | 28.54521072 | 30.18352431 | 28.37106704 | 28.26936885 | 28.49037191 |
| Arf-GAP with coiled-coil, ANK repeat a | Q15057 | ACAP2 | Other | -0.351487763 | 0.529450251 | 24.8336904 | 25.04564403 | 24.86173818 | 24.65036855 | 24.72411162 | 24.50615453 | 24.66746978 | 24.67422158 | 23.91469899 |
| Acetyl-CoA acetyltransferase, mitoch | P24752 | ACAT1 | Extracellular or secreted | -0.184262333 | 0.214952752 | 26.99453213 | 26.63547214 | 27.0584795 | 27.52805653 | 27.66152913 | 28.87575352 | 27.22278532 | 27.10219471 | 27.49914444 |
| Acetyl-CoA acetyltransferase, cytosoli | Q9BWD1;Q9BWD1-2 | ACAT2 | Extracellular or secreted | 2.780863247 | 3.430313693 | 30.26751713 | 30.35604982 | 30.41588799 | 29.8134014 | 29.76363576 | 29.82069223 | 32.61056476 | 32.71812409 | 33.23249306 |
| Golgi resident protein GCP60 | Q9H3P7 | ACBD3 | Other | -0.118028889 | 0.662584038 | 28.71881818 | 28.67340629 | 28.53815885 | 28.22715386 | 28.35904904 | 28.46532657 | 28.39186063 | 28.44375594 | 28.30125315 |
| Acyl-CoA-binding domain-containing | Q5T8D3-4;Q5T8D3-2;Q5T8D3 | ACBD5 | Other | -1.023566412 | 0.912346812 | 23.57600172 | 23.98224439 | 23.62176215 | 24.07341004 | 23.76172246 | 20.58912088 | 22.56787997 | 22.17392718 | 21.98962444 |
| Angiotensin-converting enzyme;Angic | P12821-2;P12821 | ACE | Extracellular or secreted | 0.174433244 | 0.142655052 | 23.54274095 | 24.1808622 | 24.09916735 | 21.76717634 | 21.80451712 | 23.72754326 | 23.48037507 | 23.19248246 | 23.41144582 |
| Apoptotic chromatin condensation in | Q9UKV3-5;Q9UKV3 | ACIN1 | Extracellular or secreted | -0.533926854 | 0.718474815 | 29.65996531 | 29.57931479 | 29.63008225 | 29.46777692 | 29.59759724 | 29.58464651 | 28.79955767 | 28.75519057 | 29.60316271 |
| ATP-citrate synthase | P53396;P53396-2;P53396-3 | ACLY | Extracellular or secreted | 1.495063003 | 5.74151527 | 31.835601 | 31.84879145 | 31.80489732 | 31.55362039 | 31.6279276 | 31.33942052 | 33.06384336 | 33.15742369 | 33.2690511 |
| Cytoplasmic aconitate hydratase | P21399 | ACO1 | Extracellular or secreted | -0.390057131 | 0.267410449 | 30.14879929 | 30.10898811 | 30.09760609 | 29.4344998 | 29.48870674 | 29.51656864 | 29.9654045 | 29.90526594 | 28.3567425 |
| Aconitate hydratase, mitochondrial | Q99798 | ACO2 | Other | -0.234401272 | 0.475121049 | 29.41088402 | 29.42622418 | 29.5164935 | 29.35314955 | 29.29896148 | 29.56846503 | 29.38806673 | 29.3740767 | 28.82174163 |
| Acyl-coenzyme A thioesterase 1;Acyl- | Q86TX2;P49753 | ACOT1;ACOT2 | Other | -0.023444998 | 0.032366569 | 25.06023316 | 25.90039661 | 25.78171789 | 25.13210431 | 25.79439187 | 25.39145915 | 25.75519057 | 25.59821156 | 25.10641438 |
| Acyl-coenzyme A thioesterase 13;Acyl | Q9NPJ3;Q9NPJ3-2 | ACOT13 | Extracellular or secreted | 0.09239321 | 0.042871509 | 26.83593224 | 27.1559562 | 27.11684047 | 26.9502695 | 26.79934151 | 25.48369968 | 27.4689423 | 27.4912898 | 25.48796733 |
| Cytosolic acyl coenzyme A thioester h | O00154-4;O00154-6;O00154 | ACOT7 | Extracellular or secreted | 0.491477132 | 1.408992894 | 28.97103601 | 28.82195446 | 28.80654992 | 28.93474913 | 28.92928437 | 28.69511293 | 29.40843626 | 29.51930859 | 29.12602996 |
| Acyl-coenzyme A thioesterase 9, mito | Q9Y305-3;Q9Y305-2;Q9Y305 | ACOT9 | Other | 0.088271738 | 0.043683891 | 23.19723049 | 24.39368634 | 24.78191796 | 25.23172891 | 24.88557871 | 26.31061764 | 24.34014458 | 24.32943997 | 25.9956107 |
| Peroxisomal acyl-coenzyme A oxidase | Q15067-2;Q15067;Q15067-3 | ACOX1 | Extracellular or secreted | -0.396052627 | 0.596858967 | 25.81098028 | 26.27230885 | 26.41564224 | 25.25208935 | 25.58806494 | 27.24199958 | 25.74816828 | 25.45785652 | 25.89635994 |
| Low molecular weight phosphotyrosin | P24666;P24666-4;P24666-3 | ACP1 | Extracellular or secreted | 0.67266188 | 1.936977413 | 27.29649113 | 27.50294201 | 27.48100672 | 27.88669535 | 27.90190458 | 27.68244301 | 28.25300065 | 28.53993429 | 28.10079209 |
| Lysosomal acid phosphatase | P11117 | ACP2 | Extracellular or secreted | -1.470337549 | 2.938961461 | 25.00876883 | 25.15638063 | 25.18555784 | 25.536648 | 25.85367125 | 24.18268168 | 23.77313921 | 23.60229562 | 23.67540664 |
| Lysophosphatidic acid phosphatase ty | Q9NPH0;Q9NPH0-2 | ACP6 | Extracellular or secreted | 0.476318871 | 0.338852172 | 25.84735236 | 25.10569531 | 25.24459669 | 24.65047812 | 25.53243298 | 22.17337473 | 24.93629466 | 24.99817445 | 25.77145259 |
| Acyl-CoA synthetase family member 2 | Q96CM8-3;Q96CM8;Q96CM8 | ACSF2 | Extracellular or secreted | -1.913839708 | 2.091061789 | 23.40028221 | 23.48099132 | 23.53005288 | 24.05308137 | 24.29747103 | 25.71978444 | 22.17861501 | 22.7449496 | 21.57574789 |
| Acyl-CoA synthetase family member 3 | Q4G176 | ACSF3 | Other | -0.268097108 | 0.299706908 | 22.75184967 | 22.54710258 | 23.32099771 | 23.68795142 | 23.40067309 | 24.39290067 | 22.87582733 | 22.77456751 | 23.59605141 |
| Long-chain-fatty-acid--CoA ligase 1 | P33121-3;P33121;P33121-2 | ACSL1 | Extracellular or secreted | 0.940871325 | 0.70952481 | 23.49020357 | 23.83759329 | 23.5920637 | 24.51659118 | 22.7665621 | 22.34642263 | 23.75776111 | 23.97447551 | 25.36509559 |
| Long-chain-fatty-acid--CoA ligase 3 | O95573 | ACSL3 | Extracellular or secreted | -1.526390336 | 0.441295961 | 25.18680491 | 24.81440723 | 25.18419616 | 23.86910129 | 23.90085723 | 28.08957227 | 21.73863492 | 23.13221416 | 26.07244945 |
| Long-chain-fatty-acid--CoA ligase 4 | O60488-2;O60488 | ACSL4 | Extracellular or secreted | 0.681166668 | 0.624104068 | 26.23134439 | 26.05967634 | 26.459671 | 26.15160825 | 25.25241424 | 27.78624998 | 26.74239723 | 26.51505757 | 27.75652731 |
| Acetyl-coenzyme A synthetase, cytopl | Q9NR19;Q9NR19-2 | ACSS2 | Other | 2.724187765 | 1.942801191 | 27.37260379 | 27.48431451 | 27.07040351 | 27.26508175 | 27.44387448 | 27.56310944 | 29.89078811 | 29.64560797 | 30.73586096 |
| Actin, alpha skeletal muscle;Actin, alp | P68133;P68032;P63267;P627 | ACTA1;ACTC1;AC | Extracellular or secreted | -0.139765149 | 0.11100536 | 31.2923608 | 31.30399836 | 31.33015195 | 31.35982404 | 31.43285842 | 33.9149185 | 31.33527301 | 31.5861151 | 31.97637249 |
| Actin, cytoplasmic 1;Actin, cytoplasmi | P60709 | ACTB | Extracellular or secreted | 1.397474527 | 2.87408732 | 27.97114567 | 27.47908009 | 27.80230316 | 27.77033129 | 27.89411759 | 29.05915026 | 29.38067594 | 29.3346019 | 29.46520978 |
| Beta-actin-like protein 2 | Q562R1 | ACTBL2 | Extracellular or secreted | 3.311473995 | 1.043834036 | 30.52561319 | 30.12275119 | 29.23448161 | 25.25031924 | 21.82459138 | 29.31790532 | 29.55853694 | 30.76390481 | 32.7498112 |
| Actin, cytoplasmic 2;Actin, cytoplasmi | P63261 | ACTG1 | Extracellular or secreted | 0.620994886 | 2.454752139 | 35.63325407 | 35.58827274 | 35.58332103 | 35.74842723 | 35.77377157 | 36.04209266 | 36.25462333 | 36.26690658 | 36.5260244 |
| Actin-like protein 6A | O96019;O96019-2 | ACTL6A | Extracellular or secreted | 0.24489396 | 0.406606292 | 30.55334574 | 30.67101167 | 30.59396992 | 30.01611124 | 30.04100749 | 30.2015134 | 30.80439733 | 30.81367662 | 30.15508766 |
| Alpha-actinin-1 | P12814;P12814-3;P12814-2;P | ACTN1 | Extracellular or secreted | 0.744005522 | 1.915393224 | 32.55962673 | 32.55304814 | 32.46246255 | 32.4588012 | 32.62093326 | 32.00808357 | 33.05609381 | 33.04286728 | 33.4645332 |
| Alpha-actinin-2 | P35609-2;P35609 | ACTN2 | Extracellular or secreted | 0.363310596 | 0.204660119 | 26.35248882 | 26.69988525 | 26.29792576 | 24.42130416 | 24.95841288 | 24.08925908 | 26.65608242 | 25.99033962 | 24.85314772 |
| Alpha-actinin-4 | O43707;O43707-2;O43707-3 | ACTN4 | Extracellular or secreted | 0.476205763 | 1.768003197 | 35.70336751 | 35.73809302 | 35.64065238 | 35.28171355 | 35.3735725 | 34.9905327 | 35.84377961 | 35.83255327 | 36.11625024 |
| Actin-related protein 10 | Q9NZ32 | ACTR10 | Other | 1.640521831 | 1.068514415 | 24.56139144 | 25.11199495 | 24.98901884 | 20.61799967 | 22.72419992 | 25.38919569 | 25.40223558 | 25.30539076 | 25.91083941 |
| Alpha-centractin | P61163 | ACTR1A | Extracellular or secreted | 0.225735875 | 0.550713293 | 29.15274481 | 29.29010989 | 29.18690406 | 28.55530377 | 28.69052461 | 28.74474049 | 29.27810446 | 29.32422945 | 28.88503753 |
| Beta-centractin | P42025 | ACTR1B | Extracellular or secreted | 0.137052574 | 0.135948087 | 26.81518939 | 26.95282619 | 26.75251336 | 26.07783684 | 26.37316824 | 25.83067168 | 26.9699389 | 26.87290967 | 25.96941199 |
| Actin-related protein 2 | P61160;P61160-2 | ACTR2 | Extracellular or secreted | 0.12398105 | 0.227075445 | 30.87598782 | 31.00617836 | 30.90934934 | 30.19342843 | 30.31559678 | 30.75620915 | 30.97589438 | 30.95767699 | 30.46674673 |
| Actin-related protein 3 | P61158 | ACTR3 | Extracellular or secreted | -0.045609913 | 0.091149252 | 32.02696559 | 32.04540839 | 31.98162126 | 31.13723965 | 31.19876491 | 31.75306355 | 31.6539919 | 31.75094936 | 31.52976068 |
| Actin-related protein 3B | Q9P1U1;Q9P1U1-3;Q9P1U1- | ACTR3B | Extracellular or secreted | -0.818041733 | 0.505741669 | 26.52197597 | 26.38442753 | 26.29673617 | 25.60986588 | 25.51171405 | 25.54798508 | 25.58058038 | 25.94824316 | 23.9534036 |
| Actin-related protein 5 | Q9H9F9 | ACTR5 | Other | 0.514013143 | 0.194581404 | 21.25046096 | 20.34105242 | 22.23435694 | 20.96200028 | 21.51387559 | 22.16189457 | 23.02250666 | 22.64976578 | 20.10158737 |
| Aminoacylase-1 | Q03154;Q03154-4;Q03154-3 | ACY1 | Extracellular or secreted | 0.144529584 | 0.377856373 | 28.62470701 | 28.76932263 | 28.58974434 | 28.65649164 | 28.84272084 | 27.91575914 | 28.8520703 | 28.76113276 | 28.51975849 |
| Acylphosphatase-1 | P07311 | ACYP1 | Extracellular or secreted | -0.315515652 | 0.574090655 | 28.90213467 | 29.04478981 | 28.98236953 | 29.44730782 | 29.41904688 | 27.95654198 | 28.85102784 | 28.7004476 | 28.37807295 |
| Acylphosphatase-2 | P14621 | ACYP2 | Extracellular or secreted | 0.702916925 | 0.819181643 | 26.30016233 | 26.19289252 | 26.19649902 | 26.14035892 | 26.37280304 | 24.08237192 | 26.85638107 | 26.8003294 | 26.09458418 |
| Adenosine deaminase | P00813 | ADA | Extracellular or secreted | -0.67149854 | 2.401991797 | 28.68113636 | 28.61841922 | 28.64271928 | 28.44604594 | 28.38574446 | 27.80937416 | 27.85833896 | 27.83418487 | 27.58470026 |
| Disintegrin and metalloproteinase do | O14672;O14672-2 | ADAM10 | Extracellular or secreted | -0.202900014 | 0.206858266 | 28.31125017 | 28.34307956 | 28.35237095 | 28.39312987 | 28.45206939 | 29.06183023 | 27.84122168 | 28.05128786 | 28.95565549 |
| Disintegrin and metalloproteinase do | Q13444-10;Q13444-9;Q1344 | ADAM15 | Extracellular or secreted | 1.03983763 | 0.669187409 | 21.70702694 | 24.3595099 | 24.14802343 | 24.75047025 | 24.51869349 | 24.35460882 | 24.39021305 | 24.5168917 | 26.13157456 |
| Disintegrin and metalloproteinase do | P78536-2;P78536 | ADAM17 | Extracellular or secreted | 1.650544506 | 2.58602874 | 23.61649949 | 21.91581914 | 23.37663309 | 22.78302779 | 23.52909974 | 22.2588992 | 24.18517972 | 24.59297615 | 24.91346688 |
| Disintegrin and metalloproteinase do | Q13443-2;Q13443 | ADAM9 | Extracellular or secreted | 0.638426527 | 0.59569596 | 23.91679574 | 23.93624973 | 23.98615661 | 21.61471691 | 23.97814694 | 24.67766631 | 24.00212964 | 24.08812699 | 24.87988907 |
| ADAMTS-like protein 3 | P82987-2;P82987 | ADAMTSL3 | Extracellular or secreted | -0.708059689 | 0.734925861 | 20.83415101 | 21.45016315 | 21.5712002 | 21.72504072 | 21.15789213 | 22.55159187 | 20.52046906 | 20.43063631 | 21.56973511 |
| Double-stranded RNA-specific adenos | P55265-5;P55265;P55265-4 | ADAR | Other | -0.785454381 | 1.779958593 | 29.25770589 | 29.44925764 | 29.37083847 | 28.57639825 | 28.56949755 | 30.00490682 | 28.52690065 | 28.32268325 | 28.40835527 |
| tRNA-specific adenosine deaminase 2 | Q7Z6V5-2;Q7Z6V5 | ADAT2 | Other | 0.223640366 | 0.106846124 | 20.97214434 | 22.87233671 | 21.28032332 | 22.81246955 | 22.06352609 | 22.66422851 | 22.84867566 | 23.13105247 | 21.02370724 |
| Alpha-adducin | P35611-2;P35611-6;P35611;P | ADD1 | Extracellular or secreted | -0.459051616 | 0.718075479 | 27.72747835 | 27.94781972 | 27.78462856 | 27.25448818 | 27.28665533 | 27.5458367 | 27.07030114 | 26.74136885 | 27.58462859 |
| Beta-adducin | P35612;P35612-2;P35612-8;P | ADD2 | Extracellular or secreted | -0.130311926 | 0.479054756 | 25.73402031 | 25.53744806 | 25.58112655 | 25.5693237 | 25.16238641 | 25.263023 | 25.25407363 | 25.26266465 | 25.51598996 |
| Gamma-adducin | Q9UEY8-2;Q9UEY8 | ADD3 | Extracellular or secreted | -0.728585835 | 0.744975144 | 25.46210756 | 26.1104027 | 26.09018836 | 24.87853331 | 24.90811551 | 24.42534398 | 24.7265562 | 25.12730986 | 23.89772214 |
| Alcohol dehydrogenase class-3 | P11766 | ADH5 | Extracellular or secreted | -0.04264815 | 0.062372598 | 29.24465567 | 29.33131645 | 29.17169205 | 29.16079597 | 29.1088386 | 28.63688992 | 29.23235582 | 29.31717217 | 28.64962189 |
| 1,2-dihydroxy-3-keto-5-methylthiope | Q9BV57;Q9BV57-2 | ADI1 | Extracellular or secreted | 2.137905542 | 2.618459057 | 24.19970349 | 23.76597641 | 23.75236023 | 25.87232262 | 25.78344251 | 24.0229299 | 26.81188676 | 26.87829943 | 26.42189802 |
| Adenosine kinase | P55263;P55263-2;P55263-3;P | ADK | Other | 0.092808455 | 0.63866194 | 27.81164182 | 27.7933999 | 27.90477802 | 28.01291885 | 28.12888357 | 28.0333867 | 28.10004013 | 28.06717526 | 27.95371441 |
| Activity-dependent neuroprotector ho | Q9H2P0 | ADNP | Extracellular or secreted | -0.786966429 | 0.944929957 | 28.05050964 | 27.99674233 | 27.8847756 | 27.31250576 | 26.19308058 | 28.2194656 | 26.97213228 | 27.23461865 | 26.26088954 |
| 2-aminoethanethiol dioxygenase | Q96SZ5 | ADO | Other | 1.260481924 | 1.623122787 | 23.82938983 | 22.96573145 | 23.26670878 | 24.01239675 | 24.02538224 | 23.09414922 | 25.0123115 | 25.1492646 | 24.21674882 |
| Poly(ADP-ribose) glycohydrolase ARH | Q9NX46 | ADPRHL2 | Extracellular or secreted | -0.110802284 | 0.607079179 | 27.83900007 | 27.73969616 | 27.64725606 | 27.73498936 | 27.68612208 | 27.25862713 | 27.54406852 | 27.49777486 | 27.5785952 |
| Beta-adrenergic receptor kinase 1;Bet | P25098;P35626 | ADRBK1;ADRBK2 | Extracellular or secreted | -0.876135042 | 0.27185486 | 23.7459139 | 24.32560246 | 24.34374038 | 24.19708048 | 23.83151756 | 24.45738689 | 23.95446897 | 24.88553217 | 20.98221457 |
| Proteasomal ubiquitin receptor ADRM | Q16186 | ADRM1 | Extracellular or secreted | -0.355760236 | 1.777218787 | 27.11505508 | 27.0349613 | 26.86088616 | 27.47220035 | 27.31259232 | 26.92546932 | 26.64787361 | 26.78643695 | 26.858991 |
| Adenylosuccinate lyase | P30566;P30566-2 | ADSL | Other | 0.17101195 | 0.264395615 | 29.53365497 | 29.67551096 | 29.64423313 | 29.27068368 | 29.36231381 | 29.4278032 | 29.96705502 | 29.80259591 | 29.20048479 |
| Adenylosuccinate synthetase isozyme | P30520 | ADSS | Extracellular or secreted | 0.682264722 | 3.964405884 | 30.08271199 | 30.16146888 | 29.96691755 | 29.7322582 | 29.78230237 | 29.76339832 | 30.65070066 | 30.63616392 | 30.50445824 |
| Adenylosuccinate synthetase isozyme | Q8N142;Q8N142-2 | ADSSL1 | Other | -0.73220905 | 0.376378305 | 23.83720829 | 24.04435209 | 23.57842245 | 22.66248756 | 22.87897242 | 23.44202419 | 21.22963146 | 23.39976086 | 23.39571403 |
| AFG3-like protein 2 | Q9Y4W6 | AFG3L2 | Other | -0.080330311 | 0.111106208 | 24.85600105 | 25.08480644 | 24.82905104 | 25.10733267 | 24.97281155 | 25.19775542 | 25.15332967 | 25.18430968 | 24.4452488 |
| Aftiphilin | Q6ULP2-8;Q6ULP2-6;Q6ULP2 | AFTPH | Other | 1.153479457 | 0.993113121 | 23.47704278 | 23.57968883 | 23.4302024 | 21.62764779 | 21.07856652 | 24.48646434 | 24.25219765 | 23.52157169 | 24.52647536 |
| N(4)-(beta-N-acetylglucosaminyl)-L-as | P20933 | AGA | Extracellular or secreted | -1.828952756 | 0.493757599 | 26.85638107 | 26.31485814 | 26.06198469 | 27.08222978 | 27.18886212 | 21.08881067 | 24.74811708 | 25.7004939 | 21.36109398 |
| Arf-GAP with GTPase, ANK repeat and | Q96P47-3;Q96P47-6;Q9UPQ3 | AGAP3;AGAP1 | Other | 0.066754678 | 0.075530044 | 22.95375881 | 23.03289928 | 22.86310035 | 21.07402627 | 22.78271905 | 22.85321436 | 22.71693079 | 22.49950336 | 22.76368894 |
| Arf-GAP domain and FG repeat-contai | P52594-2;P52594-3;P52594;P | AGFG1 | Other | 0.57819117 | 1.004231081 | 25.26309466 | 24.90334202 | 25.17047005 | 25.50497339 | 25.3688296 | 23.71732834 | 25.67839189 | 25.73838238 | 25.28181827 |
| Glycogen debranching enzyme;4-alph | P35573;P35573-2;P35573-3 | AGL | Other | -0.191666316 | 0.532574044 | 28.40989328 | 28.53897287 | 28.58276381 | 28.30168925 | 28.49716575 | 28.13330044 | 28.28854866 | 28.41041907 | 27.95792602 |
| Protein argonaute-1 | Q9UL18 | AGO1 | Other | -0.985239636 | 0.559853125 | 23.30229084 | 23.66925533 | 23.46699948 | 22.2559085 | 22.28534351 | 23.47444567 | 22.84255061 | 22.67356958 | 20.75528257 |
| Protein argonaute-2 | Q9UKV8;Q9UKV8-2 | AGO2 | Other | -0.292654396 | 0.522280267 | 27.8003294 | 27.91245073 | 27.85946502 | 26.96135272 | 26.76293916 | 27.09032971 | 27.1330064 | 27.35850419 | 26.8239596 |
| Anterior gradient protein 2 homolog | O95994 | AGR2 | Extracellular or secreted | -1.518930359 | 1.837908321 | 28.41671359 | 28.44978895 | 28.34604241 | 27.93769825 | 27.86979554 | 28.63985928 | 26.99991693 | 27.04239114 | 26.23084987 |
| Agrin;Agrin N-terminal 110 kDa subur | O00468-6;O00468-3;O00468 | AGRN | Extracellular or secreted | 1.943656372 | 2.146778871 | 30.43191254 | 30.29427953 | 30.29208648 | 30.69693756 | 30.739004 | 33.22029048 | 32.57601073 | 32.86315561 | 33.22905807 |
| Cytosolic carboxypeptidase 1 | Q9UPW5-2;Q9UPW5;Q9UPW | AGTPBP1 | Extracellular or secreted | -0.081188958 | 0.091295391 | 22.12801096 | 20.82912542 | 21.83824904 | 21.75061454 | 22.64980128 | 22.22789014 | 21.9457453 | 22.10216667 | 21.42036685 |
| Protein ELYS | Q8WYP5;Q8WYP5-3;Q8WYP5 | AHCTF1 | Extracellular or secreted | -0.077879769 | 0.059738817 | 22.71577938 | 23.96446285 | 23.60501193 | 21.76660059 | 24.111 | 23.95224855 | 22.82032156 | 23.25378517 | 23.74980561 |
| Adenosylhomocysteinase | P23526;P23526-2 | AHCY | Extracellular or secreted | 0.401996548 | 1.307186584 | 33.01376752 | 33.0904149 | 33.09881421 | 32.89164292 | 32.92597505 | 32.79934923 | 33.48711634 | 33.47761896 | 33.15123626 |
| Putative adenosylhomocysteinase 2 | O43865;O43865-2 | AHCYL1 | Extracellular or secreted | 1.269765891 | 5.090132728 | 26.47512632 | 26.53442329 | 26.42476755 | 26.62393997 | 26.67774695 | 27.06363125 | 27.81659377 | 27.91661353 | 27.97590804 |
| Putative adenosylhomocysteinase 3 | Q96HN2-2;Q96HN2;Q96HN2 | AHCYL2 | Extracellular or secreted | 0.797925408 | 0.763725638 | 22.04009303 | 22.83951675 | 23.61717239 | 24.11283017 | 24.55929213 | 24.67120068 | 24.16783156 | 24.08861228 | 25.05738496 |
| Neuroblast differentiation-associated | Q09666 | AHNAK | Extracellular or secreted | 0.911289482 | 1.746924225 | 31.1825183 | 31.07495178 | 31.21627702 | 31.2792453 | 31.28504625 | 31.78713788 | 31.96302856 | 32.11523496 | 32.56819319 |
| Protein AHNAK2 | Q8IVF2-3;Q8IVF2 | AHNAK2 | Extracellular or secreted | 1.050081144 | 1.17456275 | 27.6810358 | 27.9816893 | 28.01632382 | 28.30499928 | 28.23068499 | 28.56598033 | 28.6845847 | 29.06925139 | 29.78676409 |
| Activator of 90 kDa heat shock protei | O95433;O95433-2 | AHSA1 | Extracellular or secreted | 0.268375987 | 0.342007568 | 31.23083384 | 31.0732287 | 31.04446414 | 30.94062736 | 30.85830931 | 30.47981251 | 31.38082041 | 31.5929904 | 30.64495508 |
| Axin interactor, dorsalization-associat | Q96BJ3;Q96BJ3-3 | AIDA | Other | 0.962261006 | 1.522388495 | 25.36064886 | 25.12368381 | 24.59502705 | 23.61952506 | 23.87126531 | 25.19053968 | 25.79577948 | 25.76164639 | 25.20970204 |
| Allograft inflammatory factor 1-like | Q9BQI0-3;Q9BQI0;Q9BQI0-2; | AIF1L | Extracellular or secreted | 0.469839659 | 1.52241452 | 26.70767256 | 26.90517984 | 26.86926606 | 27.14530467 | 27.03873333 | 26.27884445 | 27.12238094 | 27.50460977 | 27.25502871 |
| Apoptosis-inducing factor 1, mitochor | O95831-3;O95831;O95831-2 | AIFM1 | Other | -0.737877672 | 0.888873013 | 31.61894425 | 31.56656108 | 31.61311447 | 31.27147423 | 31.29739887 | 30.34868267 | 30.81512829 | 30.91876151 | 29.91056497 |
| Absent in melanoma 1 protein | Q9Y4K1 | AIM1 | Extracellular or secreted | -0.329156475 | 0.468339832 | 24.4880593 | 23.90994727 | 24.45776261 | 23.20959794 | 23.92143419 | 23.43033003 | 23.72702384 | 23.1190513 | 23.87502111 |
| Aminoacyl tRNA synthase complex-int | Q12904;Q12904-2 | AIMP1 | Extracellular or secreted | -0.724022549 | 3.318453144 | 31.04478981 | 30.83748201 | 30.80585836 | 30.52215562 | 30.6357316 | 30.34297363 | 29.91883258 | 29.95282619 | 30.0507691 |
| Aminoacyl tRNA synthase complex-int | Q13155 | AIMP2 | Other | -0.41979322 | 1.548740426 | 27.87783155 | 28.14457555 | 28.15185015 | 27.96537697 | 28.08085842 | 28.30007503 | 27.56114376 | 27.57231963 | 27.86744079 |
| AH receptor-interacting protein | O00170 | AIP | Extracellular or secreted | -0.156439579 | 1.167202508 | 28.1625977 | 28.14559621 | 28.12376273 | 28.06229356 | 28.17288442 | 28.29959478 | 28.07658362 | 28.03291399 | 27.90454836 |
| Adenylate kinase isoenzyme 1 | P00568 | AK1 | Extracellular or secreted | -0.103161768 | 0.2531847 | 29.69009162 | 29.70307455 | 29.73649011 | 29.64048001 | 29.57774923 | 29.33515585 | 29.59429034 | 29.70607563 | 29.23166941 |
| Adenylate kinase 2, mitochondrial;Ad | P54819-2;P54819;P54819-5;P | AK2 | Extracellular or secreted | -0.544147371 | 1.642171132 | 31.4045132 | 31.31753879 | 31.28967028 | 31.49070662 | 31.52056422 | 30.47121208 | 30.81222349 | 30.79164679 | 30.51079021 |
| GTP:AMP phosphotransferase AK3, m | Q9UIJ7;Q9UIJ7-3;Q9UIJ7-2 | AK3 | Other | -0.318825972 | 0.225413337 | 29.73816652 | 29.57163271 | 29.53693696 | 29.7503744 | 29.65587777 | 28.96914298 | 29.71860591 | 29.72554571 | 28.21043614 |
| Adenylate kinase 4, mitochondrial | P27144 | AK4 | Extracellular or secreted | 0.122592034 | 0.092408114 | 26.1360419 | 25.43389917 | 25.9624343 | 26.46319956 | 26.30855346 | 28.08273736 | 26.46419723 | 26.00174319 | 27.09526856 |
| Adenylate kinase isoenzyme 6 | Q9Y3D8-2;Q9Y3D8 | AK6 | Other | 0.452211767 | 0.446222422 | 23.52611712 | 23.30368489 | 23.28263147 | 23.79610141 | 23.38640248 | 23.10022665 | 24.52695288 | 23.82220981 | 23.20505462 |
| A-kinase anchor protein 1, mitochond | Q92667;Q92667-2 | AKAP1 | Extracellular or secreted | -0.065275635 | 0.058516269 | 22.15742959 | 22.22664503 | 22.22667442 | 22.22623343 | 22.0380509 | 22.9271978 | 21.63820963 | 22.21711875 | 22.8499603 |
| A-kinase anchor protein 12 | Q02952-3;Q02952-2;Q02952 | AKAP12 | Extracellular or secreted | -0.079892777 | 0.128962068 | 28.48123774 | 28.43757855 | 28.51080907 | 28.42173754 | 28.35051745 | 27.36269004 | 28.1844232 | 27.9231471 | 28.43503657 |
| A-kinase anchor protein 13 | Q12802-4;Q12802;Q12802-2 | AKAP13 | Other | -0.21529329 | 0.209257523 | 26.68872519 | 26.21808013 | 26.65758232 | 25.05448972 | 25.25619918 | 25.54377361 | 26.12878526 | 25.78266794 | 25.15209201 |
| A-kinase anchor protein 8 | O43823 | AKAP8 | Other | -0.11123285 | 0.073004182 | 25.57242083 | 25.74957547 | 26.27767448 | 26.33422254 | 25.3757718 | 26.27547379 | 26.18343912 | 26.41781634 | 24.85761544 |
| A-kinase anchor protein 8-like | Q9ULX6;Q9ULX6-2 | AKAP8L | Other | -0.817788485 | 1.596921702 | 27.95321485 | 27.77316437 | 27.73046153 | 27.33065419 | 27.32620275 | 27.58899421 | 26.87396578 | 26.4499306 | 27.07408412 |
| A-kinase anchor protein 9 | Q99996-5;Q99996-3;Q99996 | AKAP9 | Extracellular or secreted | 1.652362659 | 0.964630756 | 21.31641744 | 22.36869888 | 22.31903605 | 22.35784861 | 20.79658225 | 22.07799306 | 22.29605191 | 23.78875338 | 24.49057084 |
| Alcohol dehydrogenase [NADP(+)] | P14550 | AKR1A1 | Extracellular or secreted | 0.234230144 | 0.302011012 | 30.41387235 | 30.49581335 | 30.43051747 | 29.92846569 | 29.90841906 | 30.16410942 | 30.71726623 | 30.75708392 | 29.89893895 |
| Aldose reductase | P15121 | AKR1B1 | Extracellular or secreted | -0.224592841 | 1.222687569 | 31.72740531 | 31.85734552 | 31.728541 | 31.41008542 | 31.34345025 | 31.61894425 | 31.29027471 | 31.38879553 | 31.49003712 |
| Aldo-keto reductase family 1 member | Q04828 | AKR1C1 | Extracellular or secreted | -3.347251317 | 1.22304785 | 27.09978938 | 27.11584886 | 26.93966099 | 24.6748681 | 24.92420106 | 25.88908855 | 23.54309509 | 20.76529519 | 23.97158425 |
| Aflatoxin B1 aldehyde reductase mem | O43488 | AKR7A2 | Extracellular or secreted | -0.272756595 | 1.694329437 | 26.36217166 | 26.50332121 | 26.42908514 | 26.44248303 | 26.60031176 | 26.88890282 | 26.16886459 | 26.30884852 | 26.31715491 |
| RAC-alpha serine/threonine-protein k | P31749;P31749-2 | AKT1 | Extracellular or secreted | 0.204472713 | 0.205236902 | 27.58985147 | 27.67350739 | 27.632147 | 26.90942088 | 27.17540899 | 26.86738188 | 27.78362985 | 27.91735359 | 26.8362935 |
| Proline-rich AKT1 substrate 1 | Q96B36;Q96B36-2;Q96B36-3 | AKT1S1 | Other | 0.713770043 | 0.954035846 | 23.7293079 | 23.87041891 | 23.71261348 | 24.83918033 | 24.88082334 | 23.30910882 | 25.17828067 | 24.8723461 | 24.26140975 |
| RAC-beta serine/threonine-protein ki | P31751;P31751-2 | AKT2 | Extracellular or secreted | 0.622171756 | 1.237992114 | 24.31941358 | 23.71334791 | 24.33435895 | 23.53753693 | 23.24139958 | 23.68281803 | 24.58806494 | 24.64163108 | 24.05125674 |
| RAC-gamma serine/threonine-protein | Q9Y243-2;Q9Y243 | AKT3 | Extracellular or secreted | 0.01587808 | 0.019608118 | 22.97791993 | 23.14847348 | 23.63200829 | 23.21397872 | 23.41299669 | 22.26023273 | 23.30215137 | 23.33483629 | 22.7334515 |
| AKT-interacting protein | Q9H8T0;Q9H8T0-2 | AKTIP | Extracellular or secreted | 0.378495892 | 0.472326711 | 24.508393 | 24.92061672 | 24.48842711 | 24.41041098 | 22.50970145 | 24.55408901 | 24.54492341 | 24.75460456 | 24.53177884 |
| Delta-aminolevulinic acid dehydratase | P13716;P13716-2 | ALAD | Extracellular or secreted | -0.47792652 | 2.801541208 | 27.16230957 | 27.17835666 | 27.24145414 | 27.08567781 | 27.23580579 | 26.76103762 | 26.51749255 | 26.6896585 | 26.69139019 |

| | | | | | | | | | | | | | | |
|---|---|---|---|---|---|---|---|---|---|---|---|---|---|---|
| CD166 antigen | Q13740-2;Q13740 | ALCAM | Extracellular or secreted | -0.213696403 | 0.045017805 | 25.40444621 | 25.05303992 | 24.85695091 | 24.77172964 | 25.46185785 | 28.58089661 | 23.46886464 | 24.64174132 | 28.31276541 |
| Aldehyde dehydrogenase family 16 m | Q8IZ83;Q8IZ83-3 | ALDH16A1 | Extracellular or secreted | 0.096926944 | 0.172220828 | 30.38360384 | 30.54120904 | 30.56819319 | 29.85967236 | 30.03433167 | 29.62114729 | 30.37243774 | 30.45316901 | 29.96925279 |
| Delta-1-pyrroline-5-carboxylate synth | P54886-2;P54886 | ALDH18A1 | Other | -0.190009243 | 0.206161801 | 26.49304746 | 25.87584139 | 26.37119171 | 26.72142821 | 26.92953835 | 28.21632328 | 26.42114362 | 26.38613931 | 26.92637455 |
| Aldehyde dehydrogenase family 1 me | P47895 | ALDH1A3 | Extracellular or secreted | -0.485085655 | 0.662970429 | 34.79047233 | 34.9024384 | 34.83742561 | 34.64753056 | 34.63763284 | 33.34281472 | 34.29453973 | 34.29138675 | 33.53797378 |
| Aldehyde dehydrogenase X, mitochon | P30837 | ALDH1B1 | Other | -0.178272713 | 0.225315103 | 25.19648026 | 25.16392234 | 25.77826245 | 26.54546851 | 26.28300256 | 26.86218765 | 25.73311527 | 25.59391651 | 26.05281196 |
| Fatty aldehyde dehydrogenase | P51648;P51648-2 | ALDH3A2 | Extracellular or secreted | -0.416130575 | 0.147532425 | 21.51325725 | 21.73945925 | 22.4914519 | 21.49469415 | 20.66653251 | 27.05196198 | 23.09596152 | 21.3093945 | 21.82493078 |
| Succinate-semialdehyde dehydrogena | P51649;P51649-2 | ALDH5A1 | Other | -0.7927913 | 1.008870013 | 24.50239577 | 24.71486806 | 25.09792249 | 26.08662964 | 25.75381434 | 26.65785486 | 24.55835812 | 25.1283723 | 24.34163826 |
| Alpha-aminoadipic semialdehyde deh | P49419-2;P49419;P49419-4 | ALDH7A1 | Extracellular or secreted | -0.089319302 | 0.262042271 | 28.3948477 | 28.46252106 | 28.42157704 | 28.31064364 | 28.40393449 | 29.21604569 | 28.44565137 | 28.47196787 | 28.41920766 |
| 4-trimethylaminobutyraldehyde dehy | P49189;P49189-2 | ALDH9A1 | Extracellular or secreted | -0.381369876 | 2.33453703 | 30.97131015 | 31.0766091 | 31.01471544 | 30.84086166 | 30.91006454 | 30.76690845 | 30.48202679 | 30.4994296 | 30.66466865 |
| Fructose-bisphosphate aldolase A | P04075;P04075-2 | ALDOA | Extracellular or secreted | -0.068264681 | 0.195121143 | 35.1354136 | 35.15206026 | 35.05366856 | 35.18441876 | 35.23167084 | 34.48147354 | 35.08230752 | 35.01350964 | 34.81874158 |
| Fructose-bisphosphate aldolase C | P09972 | ALDOC | Extracellular or secreted | -0.154428432 | 0.21570494 | 31.37150332 | 31.44815504 | 31.24201094 | 31.98681711 | 32.00259511 | 30.60779013 | 31.11421121 | 31.07940946 | 31.67252985 |
| Alpha-ketoglutarate-dependent dioxy | Q9NXW9 | ALKBH4 | Other | 0.258551392 | 0.295484238 | 21.98332348 | 20.78839115 | 21.46454264 | 22.19224965 | 22.37488558 | 21.34482063 | 22.35016333 | 22.05900375 | 21.44059366 |
| RNA demethylase ALKBH5 | Q6P6C2;Q6P6C2-3;Q6P6C2-1 | ALKBH5 | Extracellular or secreted | -1.106448499 | 1.770333454 | 26.83918033 | 26.80180998 | 26.91376359 | 26.28206582 | 26.45624348 | 26.36320824 | 25.37345038 | 25.21804316 | 25.91729668 |
| Alkaline phosphatase, placental type; | P05187;P10696 | ALPP;ALPPL2 | Extracellular or secreted | 0.637070971 | 0.48613192 | 27.56985966 | 27.4867712 | 27.63644053 | 28.42834247 | 28.45281565 | 26.0182462 | 28.52454877 | 28.83677504 | 27.34612697 |
| THO complex subunit 4 | Q86V81 | ALYREF | Other | -0.541002198 | 0.982002671 | 29.98901884 | 29.89414651 | 29.84335002 | 30.28515652 | 30.32420799 | 28.56150798 | 29.30190725 | 29.4511262 | 29.07265389 |
| Putative N-acetylglucosamine-6-phos | Q9Y303;Q9Y303-2;Q9Y303-3 | AMDHD2 | Other | 0.372151239 | 0.243637244 | 26.14971038 | 26.18118457 | 26.13786014 | 25.01026379 | 24.41273832 | 23.9303959 | 26.30497318 | 25.9446958 | 24.77786129 |
| AMP deaminase 2 | Q01433-5;Q01433-2;Q01433 | AMPD2 | Other | -0.638846219 | 1.628440011 | 26.67263101 | 26.78006632 | 27.01786405 | 27.12129431 | 27.39950012 | 27.53489819 | 26.73969616 | 26.36242251 | 26.24446967 |
| Anaphase-promoting complex subuni | Q9H1A4 | ANAPC1 | Other | -0.262612335 | 0.35001364 | 22.82104211 | 23.60591622 | 22.57941262 | 22.26235641 | 21.4348655 | 21.84563553 | 22.07714821 | 22.09780196 | 22.31182702 |
| Anaphase-promoting complex subuni | Q9UM13 | ANAPC10 | Other | 0.149182166 | 0.158961708 | 23.53860292 | 22.87803744 | 22.69523509 | 21.91505147 | 22.31326725 | 21.58306883 | 22.45653618 | 23.07628408 | 22.37635773 |
| Angiogenin | P03950 | ANG | Extracellular or secreted | -1.379651653 | 0.567936862 | 24.09796267 | 24.08310271 | 24.03716285 | 24.06010944 | 22.07209761 | 25.74511891 | 23.17247817 | 20.84955559 | 23.88678836 |
| Protein angel homolog 2 | Q5VTE6-2;Q5VTE6 | ANGEL2 | Other | 0.659012739 | 1.00127195 | 21.12857315 | 22.362175 | 21.88632323 | 20.46564752 | 22.31772067 | 22.41833442 | 22.45430231 | 22.45600842 | 22.35611449 |
| Ankyrin-2 | Q01484;Q01484-5;Q01484-2 | ANK2 | Extracellular or secreted | 1.080465737 | 0.503463686 | 21.35973696 | 21.07223409 | 25.08334622 | 26.48260771 | 21.73918969 | 24.43128692 | 24.290668 | 23.95349241 | 25.0814376 |
| Ankyrin-3 | Q12955;Q12955-7;Q12955-5 | ANK3 | Extracellular or secreted | 0.314204836 | 0.190695938 | 28.23721996 | 28.40356886 | 28.01147997 | 25.7969678 | 27.08800564 | 27.29272281 | 28.16652966 | 28.39357998 | 26.79748738 |
| Rabankyrin-5 | Q9P2R3;Q9P2R3-2;Q9P2R3-4 | ANKFY1 | Extracellular or secreted | 0.109455968 | 0.151296292 | 27.65259938 | 27.81836256 | 27.74842423 | 27.69032478 | 27.65655983 | 27.45222653 | 28.00179687 | 28.05393069 | 27.281889 |
| Ankyrin repeat domain-containing pro | O75179-7;O75179-2;O75179 | ANKRD17 | Other | 0.274855165 | 0.985620094 | 26.11127865 | 25.99172406 | 26.40593971 | 26.22237999 | 26.03149492 | 26.40579368 | 26.28926132 | 26.45821647 | 26.66139321 |
| Ankyrin repeat domain-containing pro | Q8IVF6;A2A2Z9 | ANKRD18A;ANKR | Other | 0.320603425 | 0.566148419 | 30.27475428 | 29.74797628 | 30.04048501 | 29.11857416 | 29.20226102 | 29.73469872 | 30.18139312 | 30.20594672 | 29.63384517 |
| Serine/threonine-protein phosphatas | O15084;O15084-4;O15084-1 | ANKRD28 | Other | 0.035470749 | 0.140964319 | 25.46799453 | 25.48006685 | 25.34495956 | 25.37507577 | 25.50020876 | 25.4452488 | 25.55870844 | 25.55362039 | 25.30086055 |
| Ankyrin repeat and SAM domain-cont | Q92625 | ANKS1A | Other | 0.705413579 | 0.631940319 | 22.60517022 | 22.91683217 | 23.32347373 | 23.0575998 | 23.02301453 | 21.22565713 | 24.09071333 | 23.41982648 | 22.68157472 |
| Actin-binding protein anillin | Q9NQW6;Q9NQW6-2 | ANLN | Other | 0.03689714 | 0.05112531 | 25.99642987 | 25.90253723 | 25.86660393 | 26.19917927 | 26.24738836 | 25.96161767 | 26.0082773 | 25.70210693 | 26.48718536 |
| Acidic leucine-rich nuclear phosphopr | P39687 | ANP32A | Other | -0.023940694 | 0.084769427 | 32.13154758 | 32.34857716 | 32.26899535 | 32.43804972 | 32.5151215 | 32.05122303 | 32.2664005 | 32.1451285 | 32.39340609 |
| Acidic leucine-rich nuclear phosphopr | Q92688-2;Q92688 | ANP32B | Extracellular or secreted | 0.1938436 | 0.962319149 | 30.99543548 | 31.20518936 | 31.16812098 | 31.21673957 | 31.43260956 | 30.73012137 | 31.388231 | 31.29098871 | 31.27641925 |
| Acidic leucine-rich nuclear phosphopr | Q9BTT0;Q9BTT0-2 | ANP32E | Other | 0.895507697 | 0.778091348 | 29.07367563 | 29.27464322 | 29.08575378 | 28.71102529 | 28.8342753 | 28.9181786 | 30.49037191 | 30.08841009 | 29.056517 |
| Annexin A1 | P04083 | ANXA1 | Extracellular or secreted | 0.501064381 | 0.570940071 | 32.70126181 | 32.84312909 | 33.0492766 | 32.50828909 | 32.71319467 | 31.72488738 | 33.46672242 | 33.37126962 | 32.43522042 |
| Annexin A11 | P50995-2;P50995 | ANXA11 | Extracellular or secreted | -0.024957169 | 0.02338719 | 24.08318389 | 24.22589524 | 24.43160574 | 24.24590256 | 24.84191748 | 25.65417121 | 24.16254008 | 24.43039385 | 25.07353263 |
| Annexin A2 | P07355;P07355-2 | ANXA2 | Extracellular or secreted | 0.344882234 | 0.55562408 | 33.13028985 | 33.20766389 | 33.46258444 | 32.76487377 | 32.88046503 | 33.86230591 | 33.31108232 | 33.36801986 | 34.00963596 |
| Annexin A3 | P12429 | ANXA3 | Extracellular or secreted | 0.780967556 | 1.158898746 | 32.98571498 | 33.01416672 | 33.0407626 | 32.39690539 | 32.48202679 | 31.15182596 | 33.58508545 | 33.41223257 | 32.88128587 |
| Annexin A4 | P09525;P09525-2 | ANXA4 | Extracellular or secreted | 0.226375783 | 0.500387404 | 27.39550158 | 27.37476078 | 27.74175458 | 27.45159787 | 27.47429093 | 28.52050802 | 27.70589442 | 27.87753905 | 28.07490076 |
| Annexin A5 | P08758 | ANXA5 | Extracellular or secreted | 0.150419606 | 0.221856996 | 29.68324652 | 29.90339948 | 30.01757207 | 29.04411233 | 29.12178833 | 29.57699284 | 29.95504572 | 29.88122022 | 29.28854866 |
| Annexin A6 | P08133;P08133-2 | ANXA6 | Extracellular or secreted | -1.935587101 | 0.303104989 | 28.59519783 | 28.5911364 | 28.69564397 | 28.25295555 | 28.32680278 | 28.94405344 | 29.18725812 | 28.78125514 | 21.92762042 |
| Annexin A7 | P20073-2;P20073 | ANXA7 | Extracellular or secreted | 0.279947447 | 0.407664103 | 26.54223911 | 26.5803216 | 27.01563283 | 26.34536572 | 26.23768496 | 27.83116718 | 26.73337391 | 27.00088406 | 27.38179007 |
| Annexin A8 | P13928;P13928-3 | ANXA8 | Extracellular or secreted | 0.038914819 | 0.042845915 | 25.12163026 | 25.37325123 | 25.15007853 | 24.81601997 | 24.54827913 | 24.0997292 | 25.31063497 | 24.90334202 | 24.45726163 |
| Annexin A8-like protein 2 | Q5VT79;Q5VT79-2 | ANXA8L2 | Extracellular or secreted | 0.135116594 | 0.195626722 | 29.93142767 | 29.95712344 | 29.9005233 | 29.61180501 | 29.59478864 | 28.99337176 | 29.98996625 | 30.04648212 | 29.36342132 |
| AP-1 complex subunit beta-1 | Q10567-3;Q10567-2;Q10567 | AP1B1 | Other | -0.18630404 | 0.419837461 | 30.31592063 | 30.41064146 | 30.38061401 | 29.62578714 | 29.66991421 | 30.0383932 | 30.01836825 | 30.02287171 | 29.62048324 |
| AP-1 complex subunit gamma-1 | O43747;O43747-2 | AP1G1 | Other | 0.018724548 | 0.033691375 | 29.62907483 | 29.93170945 | 30.0235328 | 29.20513108 | 29.04944539 | 29.56427302 | 29.74920781 | 29.63476273 | 29.37378639 |
| AP-1 complex subunit gamma-like 2 | O75843 | AP1G2 | Other | -0.378177787 | 0.076473319 | 25.14755772 | 25.56191579 | 25.24307168 | 20.58386694 | 20.96758112 | 25.04046933 | 24.93984031 | 20.47796876 | 24.71988886 |
| AP-1 complex subunit mu-1 | Q9BXS5;Q9BXS5-2 | AP1M1 | Extracellular or secreted | -0.021387454 | 0.059588918 | 27.10559541 | 27.13565058 | 27.18072944 | 26.63878964 | 26.64182399 | 26.90437608 | 27.06537866 | 26.80538177 | 26.86855978 |
| AP-1 complex subunit mu-2 | Q9Y6Q5 | AP1M2 | Other | -0.262296103 | 0.512989086 | 29.68291178 | 30.00704772 | 29.84680561 | 28.88930909 | 29.05336108 | 29.37301885 | 29.45695235 | 29.2456986 | 28.93668781 |
| AP-1 complex subunit mu-2 | Q9Y6Q5-2 | AP1M2 | Other | 0.172431004 | 0.171660866 | 22.94343986 | 23.31063497 | 22.87483355 | 20.98038845 | 22.21534221 | 23.21520888 | 23.02375909 | 22.78236799 | 22.48108989 |
| AP-1 complex subunit sigma-1A | P61966;P61966-2 | AP1S1 | Extracellular or secreted | -0.244645294 | 0.682145541 | 27.33406906 | 27.44158103 | 27.49106038 | 27.23863273 | 27.42205848 | 27.1574026 | 27.29017582 | 27.18271956 | 26.83557089 |
| AP-1 complex subunit sigma-2 | P56377;P56377-2 | AP1S2 | Other | 0.634629149 | 0.540227624 | 23.44202419 | 20.212943 | 23.54321311 | 22.66472764 | 23.06261472 | 21.5450724 | 23.01806363 | 22.89943806 | 23.22168329 |
| AP-1 complex subunit sigma-3 | Q96PC3-4;Q96PC3-3;Q96PC3 | AP1S3 | Other | 0.546565961 | 2.203809131 | 24.22942028 | 24.45807563 | 24.64019727 | 23.97648721 | 23.98346266 | 24.66139321 | 24.86438567 | 24.78096741 | 24.96886293 |
| AP-2 complex subunit alpha-1 | O95782-2;O95782 | AP2A1 | Extracellular or secreted | -0.252399738 | 0.571099343 | 30.37844509 | 30.65616769 | 30.6169306 | 29.67813327 | 29.63865156 | 30.4016173 | 30.16470887 | 29.91712592 | 29.84593875 |
| AP-2 complex subunit alpha-2 | O94973;O94973-2;O94973-3 | AP2A2 | Extracellular or secreted | -0.060856329 | 0.067598943 | 28.44494088 | 28.61185776 | 28.56804819 | 27.71115662 | 27.76673473 | 27.92987692 | 28.35741387 | 28.39083624 | 27.58548844 |
| AP-2 complex subunit beta | P63010;P63010-2;P63010-3 | AP2B1 | Extracellular or secreted | -0.106847742 | 0.273641184 | 31.68645607 | 31.86045701 | 31.78171633 | 31.00463899 | 31.03170524 | 31.57569524 | 31.45395394 | 31.40024352 | 31.29559376 |
| AP-2 complex subunit mu | Q96CW1-2;Q96CW1 | AP2M1 | Extracellular or secreted | -0.280430986 | 0.613481746 | 29.6628876 | 29.84495169 | 29.67317038 | 28.79498053 | 28.79333788 | 29.51966477 | 29.26342603 | 29.14930821 | 28.89046923 |
| AP-2 complex subunit sigma | P53680;P53680-2 | AP2S1 | Extracellular or secreted | -0.694309181 | 2.354104642 | 26.6477364 | 27.31345755 | 27.26945803 | 26.55332743 | 26.68391576 | 26.87044242 | 26.06884665 | 26.12644359 | 26.390951 |
| AP-3 complex subunit beta-1 | O00203-3;O00203 | AP3B1 | Other | -0.396853852 | 1.804457671 | 26.24952379 | 26.41800943 | 26.48491371 | 25.72142821 | 26.19507253 | 26.27453214 | 25.82048225 | 25.93086959 | 25.7298265 |
| AP-3 complex subunit delta-1 | O14617-5;O14617;O14617-4 | AP3D1 | Other | -0.573477252 | 2.306304107 | 27.36126821 | 27.34384201 | 27.42382236 | 26.75225813 | 26.94871103 | 27.50725907 | 26.61203359 | 26.64113492 | 26.69498014 |
| AP-3 complex subunit mu-1 | Q9Y2T2 | AP3M1 | Other | -0.487774709 | 1.621899864 | 24.81821626 | 24.74314235 | 24.75776111 | 24.73039675 | 25.07157005 | 25.57712614 | 24.27056339 | 24.60924593 | 24.50597288 |
| AP-3 complex subunit sigma-1 | Q92572 | AP3S1 | Other | -0.526670084 | 2.523721259 | 23.76364842 | 24.00787112 | 23.55227227 | 24.14623739 | 24.11394306 | 24.16184845 | 23.42815874 | 23.39649817 | 23.46824319 |
| Acylamino-acid-releasing enzyme | P13798 | APEH | Extracellular or secreted | 0.014999596 | 0.028878731 | 31.92252343 | 31.96650506 | 31.90102704 | 32.32184502 | 32.44684953 | 31.32136121 | 31.92348009 | 31.91056497 | 32.15100938 |
| DNA-(apurinic or apyrimidinic site) lya | P27695 | APEX1 | Other | 0.067981285 | 0.083239291 | 32.74236109 | 32.69548634 | 32.62881852 | 32.71738882 | 32.7681121 | 32.329564 | 32.93494593 | 33.02633906 | 32.18352431 |
| Apoptosis inhibitor 5 | Q9BZZ5-2;Q9BZZ5;Q9BZZ5-3; | API5 | Other | 1.183058683 | 1.94689923 | 29.22319975 | 29.36202112 | 29.495699 | 28.74332857 | 28.77291276 | 29.25376714 | 30.64151395 | 30.4039548 | 29.92917148 |
| Methylthioribulose-1-phosphate dehy | Q96GX9;Q96GX9-3 | APIP | Other | 0.495696239 | 2.263323139 | 28.46835973 | 28.53812184 | 28.44573029 | 28.44537511 | 28.3933754 | 27.87806551 | 28.92042364 | 28.94268398 | 28.70849503 |
| Amyloid-like protein 2 | Q06481-3;Q06481-6;Q06481 | APLP2 | Extracellular or secreted | 0.862225927 | 0.972280908 | 26.7679977 | 26.96091102 | 27.13662868 | 28.45203011 | 28.53819586 | 28.46466465 | 28.26561834 | 28.296141 | 29.18513245 |
| Adipocyte plasma membrane-associat | Q9HDC9;Q9HDC9-2 | APMAP | Extracellular or secreted | -0.685640121 | 0.227223152 | 24.82444527 | 24.95783748 | 25.42547204 | 23.0038459 | 24.90999303 | 28.37422184 | 23.09579169 | 24.24981309 | 26.34538264 |
| NAD(P)H-hydrate epimerase | Q8NCW5;Q8NCW5-2 | APOA1BP | Extracellular or secreted | -0.123168409 | 0.212053147 | 31.55768825 | 31.71971591 | 31.65727565 | 31.62078035 | 31.69722762 | 30.63918653 | 31.56687858 | 31.45865455 | 31.05089881 |
| Apolipoprotein B-100;Apolipoprotein | P04114 | APOB | Extracellular or secreted | -3.458773144 | 2.768520887 | 25.10972546 | 25.17843265 | 25.25598317 | 25.56749705 | 25.33073966 | 21.62016346 | 21.36491545 | 20.6060018 | 21.68403405 |
| Apolipoprotein C-III | P02656 | APOC3 | Extracellular or secreted | 3.943719984 | 3.66954299 | 21.24404976 | 21.14452973 | 19.70401785 | 23.08728405 | 22.21361118 | 20.58811309 | 25.25551502 | 24.92718876 | 25.639259 |
| Amyloid beta A4 protein;N-APP;Solub | P05067-7;P05067-11;P05067 | APP | Extracellular or secreted | 0.209825002 | 0.319969644 | 30.02683371 | 29.87595853 | 29.76903882 | 31.10718049 | 31.08112518 | 30.6786706 | 30.43489743 | 30.58799343 | 30.87598782 |
| DCC-interacting protein 13-alpha | Q9UKG1 | APPL1 | Extracellular or secreted | -0.097750331 | 0.284653114 | 28.17517101 | 28.04192137 | 27.9837562 | 27.89746826 | 27.92048043 | 27.62387022 | 27.8929025 | 28.01270577 | 27.62247449 |
| Adenine phosphoribosyltransferase | P07741;P07741-2 | APRT | Extracellular or secreted | 1.329679077 | 1.928740142 | 27.79680694 | 27.87853331 | 28.03852403 | 27.80912879 | 27.61925918 | 29.58392958 | 29.10204449 | 29.32883675 | 29.9212469 |
| Intron-binding protein aquarius | O60306 | AQR | Other | -0.288924735 | 0.967529834 | 28.71499248 | 28.9671375 | 29.12257842 | 28.84526583 | 28.92855041 | 29.1458634 | 28.8766612 | 28.63910026 | 28.47965835 |
| Serine/threonine-protein kinase A-Ra | P10398 | ARAF | Other | 0.744979833 | 0.59536852 | 22.53488038 | 23.1400234 | 22.57554017 | 22.45716141 | 22.14208212 | 22.66079502 | 23.77424576 | 23.83691947 | 22.37901552 |
| Coatomer subunit delta | P48444;P48444-2 | ARCN1 | Extracellular or secreted | -0.561849077 | 1.845205425 | 30.83575157 | 30.76469585 | 30.72476543 | 30.38535362 | 30.39407084 | 29.97514274 | 30.09948843 | 30.01038652 | 29.74446784 |
| ADP-ribosylation factor 1 | P84077 | ARF1 | Extracellular or secreted | 0.365250088 | 0.813513341 | 29.57109017 | 30.05736946 | 30.14321356 | 30.3936618 | 30.19765233 | 30.60151657 | 30.27197506 | 30.47276201 | 30.83326514 |
| ADP-ribosylation factor 4 | P18085 | ARF4 | Extracellular or secreted | 0.380985661 | 0.310015533 | 24.73024125 | 25.314789 | 25.26391847 | 24.70567035 | 24.92011693 | 26.71973223 | 25.10381604 | 25.42748754 | 26.43888752 |
| ADP-ribosylation factor 5 | P84085 | ARF5 | Extracellular or secreted | -0.045053211 | 0.024024818 | 27.19432117 | 27.14297021 | 27.14326222 | 26.89585167 | 27.03883796 | 28.59224196 | 26.57529491 | 26.90655668 | 28.38673137 |
| ADP-ribosylation factor 6 | P62330 | ARF6 | Extracellular or secreted | 1.098251368 | 3.815679044 | 25.0600682 | 24.9624343 | 25.42274825 | 25.13692198 | 25.5661328 | 25.76293916 | 26.45985857 | 26.30394612 | 26.48657175 |
| ADP-ribosylation factor GTPase-activa | Q8N6T3-5;Q8N6T3-3;Q8N6T3 | ARFGAP1 | Other | -0.683379077 | 0.339638835 | 26.17399959 | 23.49801844 | 23.79153812 | 25.78626245 | 24.98793533 | 26.22208513 | 25.09555027 | 23.03883796 | 25.04539407 |
| ADP-ribosylation factor GTPase-activa | Q8N6H7;Q8N6H7-2 | ARFGAP2 | Extracellular or secreted | 0.220734052 | 0.470805529 | 25.13076975 | 25.3390233 | 25.17603706 | 25.14608198 | 25.16096422 | 25.70909349 | 25.7161249 | 25.62176215 | 25.15530001 |
| ADP-ribosylation factor GTPase-activa | Q9NP61-2;Q9NP61 | ARFGAP3 | Other | -0.143035665 | 0.219356014 | 24.67949299 | 24.99403571 | 25.08917825 | 23.90481247 | 24.44644828 | 24.26148148 | 24.64597896 | 24.55806611 | 24.05457252 |
| Brefeldin A-inhibited guanine nucleot | Q9Y6D6 | ARFGEF1 | Other | 0.478378471 | 0.743497437 | 27.08304183 | 27.17178748 | 27.14899318 | 26.95914287 | 27.01563283 | 27.04510239 | 27.75072579 | 27.82238002 | 27.07387989 |
| Brefeldin A-inhibited guanine nucleot | Q9Y6D5 | ARFGEF2 | Other | 0.410447061 | 0.85932919 | 26.6477364 | 26.64182399 | 26.59690449 | 26.56152255 | 26.66451608 | 26.43076071 | 27.28885664 | 27.02886332 | 26.68525332 |
| Brefeldin A-inhibited guanine nucleot | Q5TH69 | ARFGEF3 | Other | 0.900411872 | 1.357826722 | 21.00182049 | 20.97309193 | 21.50726815 | 21.35589949 | 21.84773944 | 22.21138675 | 22.38650773 | 21.93840515 | 22.82492586 |
| Arfaptin-1 | P53367;P53367-2 | ARFIP1 | Other | -0.158770114 | 0.390594868 | 25.5210025 | 25.33912527 | 25.40256088 | 25.98263609 | 25.57071389 | 25.86384239 | 25.18404478 | 25.54300656 | 25.63657882 |
| Arfaptin-2 | P53365-3;P53365;P53365-2 | ARFIP2 | Extracellular or secreted | -0.511838989 | 0.154568768 | 21.53770144 | 21.83358159 | 20.56994147 | 22.18586026 | 22.2875433 | 23.38112994 | 20.43959149 | 20.22747173 | 23.69529882 |
| Arginase-1 | P05089;P05089-2;P05089-3 | ARG1 | Extracellular or secreted | -1.159365207 | 1.470617716 | 21.96349303 | 20.80586655 | 22.08310271 | 23.18948436 | 23.57819208 | 22.17952022 | 21.25855297 | 21.08899035 | 21.07419054 |
| Arginine and glutamate-rich protein 1 | Q9NWB6;Q9NWB6-3;Q9NWB | ARGLU1 | Other | -0.828672545 | 0.272370119 | 25.40415382 | 22.16032578 | 21.15569028 | 25.4203085 | 25.73840815 | 25.9794777 | 21.65939613 | 24.34624535 | 24.437523 |
| Rho GTPase-activating protein 1 | Q07960 | ARHGAP1 | Extracellular or secreted | 0.200054672 | 0.528383062 | 28.44908049 | 28.71214114 | 28.45900648 | 27.9012141 | 28.11728648 | 28.4207743 | 28.72601693 | 28.60434747 | 28.29955111 |
| Rho GTPase-activating protein 11A | Q6P4F7-3;Q6P4F7;Q6P4F7-2 | ARHGAP11A | Other | 0.247960833 | 0.120448437 | 24.77374289 | 24.66838989 | 25.93773192 | 28.01163991 | 28.37380713 | 24.40684805 | 25.84983553 | 26.18995558 | 26.79017128 |
| Rho GTPase-activating protein 12 | Q8IWW6-3;Q8IWW6-2;Q8IW | ARHGAP12 | Other | 0.316090314 | 0.971152769 | 25.05506923 | 25.80717664 | 25.84083765 | 25.31402832 | 25.15830828 | 25.71588932 | 25.92680433 | 25.85469397 | 25.61242737 |
| Rho GTPase-activating protein 17 | Q68EM7-5;Q68EM7-2;Q68EM | ARHGAP17 | Extracellular or secreted | -0.052128698 | 0.044207102 | 26.11886125 | 25.83566725 | 26.13516128 | 26.08474563 | 26.38885094 | 25.56179929 | 26.28750103 | 26.38531659 | 25.2333391 |
| Rho GTPase-activating protein 18 | Q8N392;Q8N392-2 | ARHGAP18 | Other | 0.598973381 | 0.36479915 | 26.63491848 | 26.1733325 | 26.15110498 | 25.16419096 | 24.3191379 | 25.32006811 | 26.45538141 | 27.09536917 | 25.12754603 |
| Rho GTPase-activating protein 28 | Q9P2N2-3;Q9P2N2-5;Q9P2N | ARHGAP28 | Other | -3.85198335 | 1.155361579 | 28.39431621 | 27.94815403 | 28.24213591 | 29.10527069 | 28.1519469 | 28.39766543 | 25.7976358 | 22.32879448 | 25.43736425 |
| Rho GTPase-activating protein 35 | Q9NRY4 | ARHGAP35 | Other | 0.198639008 | 0.183623732 | 21.92317658 | 21.45180526 | 21.8711934 | 21.82064244 | 21.39494623 | 22.35927539 | 21.26347847 | 22.38584976 | 22.35710844 |
| Rho GDP-dissociation inhibitor 1 | P52565;P52565-2 | ARHGDIA | Extracellular or secreted | 0.910284199 | 2.585770372 | 29.53032828 | 29.92294869 | 29.9141344 | 30.617894 | 30.60293272 | 30.20443159 | 30.88314471 | 31.04498518 | 31.19905755 |
| Rho GDP-dissociation inhibitor 2 | P52566 | ARHGDIB | Extracellular or secreted | 1.142344528 | 2.86382993 | 29.37324708 | 29.60591975 | 29.62547364 | 28.91624336 | 28.95867285 | 28.52267946 | 30.2807392 | 30.31635232 | 30.33106013 |
| Rho guanine nucleotide exchange fact | Q92888;Q92888-3;Q92888-2 | ARHGEF1 | Extracellular or secreted | 0.538623276 | 2.059583032 | 26.99593411 | 27.06773944 | 27.13349643 | 27.22894343 | 27.06845718 | 26.64910791 | 27.51713207 | 27.42157704 | 27.74899997 |
| Rho guanine nucleotide exchange fact | Q5VV41;Q5VV41-2 | ARHGEF16 | Extracellular or secreted | 0.056721347 | 0.081021178 | 26.16334656 | 26.5140192 | 26.40353636 | 25.65119012 | 25.68094193 | 25.46920632 | 26.26282592 | 26.1626169 | 25.68584146 |
| Rho guanine nucleotide exchange fact | Q92974-3;Q92974-2;Q92974 | ARHGEF2 | Extracellular or secreted | -0.268741912 | 0.164749839 | 24.38811191 | 24.96706052 | 25.17538995 | 24.26913695 | 24.16138718 | 24.77746001 | 24.52766885 | 23.31021891 | 25.22515977 |
| Rho guanine nucleotide exchange fact | Q14155-6;Q14155-1;Q14155 | ARHGEF7 | Extracellular or secreted | -0.826990219 | 1.135469279 | 26.83436573 | 26.42279636 | 25.54392107 | 24.54857312 | 26.41149431 | 26.71999328 | 25.19385138 | 25.03942379 | 25.5263261 |
| AT-rich interactive domain-containing | O14497;O14497-3;O14497-2 | ARID1A | Other | -0.925816543 | 1.469890729 | 27.15788442 | 27.01733312 | 26.93021541 | 26.37218862 | 26.42112756 | 27.0849686 | 25.88529942 | 26.32104934 | 25.50806047 |
| AT-rich interactive domain-containing | Q68CP9-3;Q68CP9 | ARID2 | Extracellular or secreted | -0.94250577 | 2.643079568 | 24.84933446 | 24.22633634 | 24.36606336 | 24.86716975 | 24.56401129 | 24.60246554 | 23.8345104 | 23.66589884 | 23.40976383 |
| AT-rich interactive domain-containing | Q4LE39-3;Q4LE39-2;Q4LE39; | ARID4B | Other | -0.766277178 | 1.744561206 | 24.42981944 | 24.14561564 | 24.0759413 | 23.57761599 | 24.34678639 | 24.50960154 | 23.56720689 | 23.59593763 | 23.0807141 |
| E3 ubiquitin-protein ligase ARIH1 | Q9Y4X5 | ARIH1 | Other | 0.168252232 | 0.483463359 | 25.80855201 | 26.05618621 | 25.95062549 | 26.12083966 | 25.95522314 | 25.96831365 | 26.29189773 | 26.26022555 | 25.8825035 |
| E3 ubiquitin-protein ligase ARIH2 | O95376 | ARIH2 | Other | 0.146333316 | 0.276573603 | 26.27501193 | 25.88943675 | 25.83607675 | 26.0651321 | 26.15236284 | 25.97127726 | 26.21673032 | 26.4853284 | 25.83159004 |
| ADP-ribosylation factor-like protein 1 | P40616-2;P40616 | ARL1 | Extracellular or secreted | -0.103592701 | 0.072835766 | 24.1754661 | 24.72816632 | 24.88683487 | 25.23319279 | 25.48237691 | 26.07932299 | 24.3803374 | 24.8098525 | 25.79171199 |
| ADP-ribosylation factor-like protein 2 | P36404-2;P36404 | ARL2 | Extracellular or secreted | 0.076579508 | 0.071608705 | 22.64480247 | 22.79205469 | 22.46587923 | 23.50560279 | 22.50147062 | 21.14784281 | 22.61732935 | 22.95649102 | 22.18474446 |
| ADP-ribosylation factor-like protein 2- | Q9Y2Y0-2;Q9Y2Y0 | ARL2BP | Other | 0.643039658 | 2.077833913 | 25.46179541 | 25.93170382 | 26.20821411 | 26.07148822 | 25.97070696 | 25.24030804 | 26.47735165 | 26.59334667 | 26.30052894 |
| ADP-ribosylation factor-like protein 3 | P36405 | ARL3 | Extracellular or secreted | -0.230682841 | 0.538831361 | 25.69540503 | 25.82303636 | 25.77605466 | 25.29031646 | 25.5431541 | 26.05740562 | 25.46273166 | 25.73861431 | 25.19929162 |
| ADP-ribosylation factor-like protein 6- | Q15041;Q15041-3;Q15041-2 | ARL6IP1 | Other | 0.482435702 | 0.472768695 | 25.07964851 | 25.13555274 | 25.10085223 | 25.24887264 | 25.39414446 | 22.49270444 | 25.32285513 | 25.14592655 | 25.20441293 |
| ADP-ribosylation factor-like protein 6- | Q66PJ3-7;Q66PJ3-4;Q66PJ3-3 | ARL6IP4 | Other | -0.714441254 | 1.386053267 | 24.08594115 | 23.98797869 | 24.27994264 | 24.24995772 | 24.19940396 | 24.94167702 | 23.81039198 | 23.73593195 | 23.1828029 |
| PRA1 family protein 3 | O75915 | ARL6IP5 | Extracellular or secreted | 0.01353432 | 0.01053045 | 25.1455379 | 25.19629264 | 25.0284629 | 24.11426087 | 24.3258083 | 26.48250001 | 24.70598667 | 24.89115328 | 25.58989432 |
| ADP-ribosylation factor-like protein 8 | Q96BM9 | ARL8A | Extracellular or secreted | -0.271187798 | 0.247920059 | 25.58932289 | 25.36909595 | 25.46444654 | 25.03485638 | 25.1655333 | 26.27108012 | 25.30382422 | 24.51207589 | 25.81770409 |
| ADP-ribosylation factor-like protein 8 | Q9NVJ2 | ARL8B | Extracellular or secreted | -0.883196887 | 0.537472465 | 21.65167189 | 22.00023938 | 21.95386536 | 22.50705033 | 23.26585079 | 23.04280024 | 20.65269176 | 21.18089254 | 22.72756404 |
| Armadillo repeat-containing protein 1 | Q9NVT9;Q9NVT9-2 | ARMC1 | Other | -0.451142746 | 1.110206397 | 23.80459427 | 24.1673722 | 24.31326725 | 24.94793117 | 24.84028542 | 24.85095633 | 24.25414573 | 23.91031334 | 23.944316 |
| Armadillo repeat-containing protein 1 | Q8N2F6-4;Q8N2F6-3;Q8N2F6 | ARMC10 | Other | -0.506555261 | 0.424251928 | 24.39316261 | 24.95455772 | 24.62906441 | 22.97051389 | 23.52205083 | 25.24441523 | 23.28855746 | 23.50869523 | 24.53996386 |
| Armadillo repeat-containing protein 6 | Q6NXE6-2;Q6NXE6 | ARMC6 | Other | 2.409039614 | 2.182336132 | 21.18052838 | 22.01171455 | 22.26028516 | 22.86368645 | 23.13078546 | 23.29740106 | 25.31520375 | 25.22062837 | 24.06348725 |
| Armadillo repeat-containing protein 8 | Q8IUR7-2;Q8IUR7;Q8IUR7-8; | ARMC8 | Other | 0.004970809 | 0.010132992 | 24.9983036 | 25.49606868 | 25.20944176 | 24.78396699 | 25.1792303 | 24.86853623 | 25.34282532 | 24.96499157 | 24.97486932 |
| Protein-glutamate O-methyltransfera | Q9H993 | ARMT1 | Other | -0.279261011 | 0.655760577 | 29.24757851 | 29.30380245 | 29.31984422 | 28.73956741 | 28.89168638 | 28.54078425 | 28.98340286 | 28.73282425 | 28.46762147 |
| Aryl hydrocarbon receptor nuclear tra | P27540-2;P27540-3;P27540-4 | ARNT | Other | -0.105512676 | 0.357616615 | 22.66109959 | 22.2862464 | 22.8814583 | 22.61380475 | 22.84758654 | 22.71822766 | 22.70117107 | 22.38995057 | 22.59655194 |
| Actin-related protein 2/3 complex sub | Q92747;Q92747-2 | ARPC1A | Extracellular or secreted | -0.008188671 | 0.033959826 | 28.89547613 | 29.19232348 | 28.84926884 | 28.83297855 | 28.88518303 | 28.60819565 | 28.92093468 | 28.88046138 | 28.80575076 |
| Actin-related protein 2/3 complex sub | O15143 | ARPC1B | Extracellular or secreted | 0.206401283 | 0.71329196 | 31.06027182 | 31.12355308 | 31.10817808 | 30.40598433 | 30.42512384 | 30.87166189 | 31.02913986 | 31.0768639 | 31.0105866 |
| Actin-related protein 2/3 complex sub | O15144 | ARPC2 | Extracellular or secreted | 0.091874805 | 0.176576644 | 31.49847883 | 31.61535273 | 31.57375568 | 30.6769908 | 30.72736473 | 31.06451807 | 31.39350838 | 31.38571361 | 31.07463285 |
| Actin-related protein 2/3 complex sub | O15145 | ARPC3 | Extracellular or secreted | 0.001063183 | 0.002376912 | 30.16925656 | 30.27375438 | 30.15110014 | 29.68378194 | 29.61721093 | 29.5327265 | 29.86050141 | 30.0383932 | 29.81821016 |
| Actin-related protein 2/3 complex sub | P59998;P59998-3;P59998-4;P | ARPC4 | Extracellular or secreted | 0.115110881 | 0.255411572 | 29.97185829 | 30.28283911 | 30.23397902 | 29.64939919 | 29.66895113 | 29.6253517 | 30.12028103 | 30.18824904 | 29.7529918 |
| Actin-related protein 2/3 complex sub | O15511;O15511-2 | ARPC5 | Extracellular or secreted | 0.155832186 | 0.528903512 | 29.12030576 | 29.25739119 | 29.17605134 | 28.74104733 | 28.66071343 | 28.8440388 | 29.25223375 | 29.1679416 | 28.94709513 |
| Actin-related protein 2/3 complex sub | Q9BPX5 | ARPC5L | Extracellular or secreted | 0.401656098 | 1.045367887 | 28.90867673 | 29.02699197 | 28.92987692 | 28.37575523 | 28.3583365 | 28.48258463 | 29.223522 | 29.23018107 | 28.79237622 |
| Arpin | Q7Z6K5;Q7Z6K5-2 | ARPIN | Other | -0.140777637 | 0.113678079 | 27.09727956 | 27.10069185 | 27.2850683 | 26.77756034 | 26.95371441 | 26.02438912 | 26.81421162 | 27.37658342 | 26.00622385 |
| cAMP-regulated phosphoprotein 19 | P56211-2;P56211 | ARPP19 | Other | 1.627960002 | 0.912837444 | 20.59974787 | 25.995546 | 25.38155906 | 25.86036524 | 25.80727492 | 24.77308889 | 27.00367437 | 25.84117369 | 26.24782294 |
| Arrestin-C | P36575-2;P36575 | ARR3 | Other | 0.282321611 | 0.359465397 | 22.94293897 | 23.1187979 | 23.69890556 | 22.46145852 | 21.42977513 | 22.29825815 | 23.07892412 | 23.03473046 | 22.70837737 |
| Beta-arrestin-1 | P49407-2;P49407 | ARRB1 | Extracellular or secreted | 0.204827355 | 0.26645992 | 27.85001444 | 27.90821861 | 27.73195082 | 27.26874444 | 27.12819528 | 27.00731511 | 27.85780525 | 28.02458992 | 27.17930624 |
| Beta-arrestin-2 | P32121;P32121-3;P32121-5;P | ARRB2 | Extracellular or secreted | 0.755711736 | 1.031466425 | 23.02744237 | 23.47555929 | 23.08258308 | 24.09940817 | 24.39368634 | 23.05084174 | 24.68078101 | 24.39865237 | 23.75246232 |
| Arrestin domain-containing protein 1 | Q8N5I2 | ARRDC1 | Extracellular or secreted | 1.53431425 | 0.77997844 | 21.00082378 | 23.73220967 | 23.43288032 | 25.0814376 | 24.13453474 | 25.92746006 | 24.61031658 | 24.8757711 | 26.77152816 |
| Arylsulfatase A;Arylsulfatase A compo | P15289-2;P15289 | ARSA | Extracellular or secreted | 0.012797205 | 0.030162673 | 25.77152816 | 26.12949292 | 25.56235261 | 25.93943681 | 25.87375462 | 25.16618485 | 25.77725933 | 25.6623172 | 25.82019007 |
| Ecto-ADP-ribosyltransferase 3 | Q13508-2;Q13508-3;Q13508 | ART3 | Extracellular or secreted | 0.824254835 | 0.29940192 | 22.20051657 | 20.9793315 | 21.66814858 | 21.09925 | 22.3621346 | 25.87471631 | 23.2831969 | 21.64279914 | 24.63881725 |
| Acid ceramidase;Acid ceramidase sub | Q13510;Q13510-2;Q13510-3 | ASAH1 | Extracellular or secreted | -0.167656358 | 0.438146624 | 29.50208842 | 29.60830142 | 29.46557958 | 29.63147064 | 29.60429444 | 29.30562994 | 29.22533907 | 29.19220589 | 29.63816819 |
| Ankyrin repeat and SOCS box protein | Q96DX5-3;Q96DX5-2;Q96DX5 | ASB9 | Other | 0.929335205 | 3.395616374 | 25.08667013 | 25.03691141 | 25.31575656 | 25.21070553 | 25.03376476 | 24.43923637 | 26.08824833 | 25.94545509 | 25.81582458 |
| Activating signal cointegrator 1 compl | Q8N9N2-2;Q8N9N2 | ASCC1 | Other | 0.210392133 | 0.596253707 | 24.80429885 | 24.61475971 | 24.79897088 | 24.47023088 | 24.48732338 | 24.42091882 | 24.93732784 | 24.9497131 | 24.54238673 |
| Histone chaperone ASF1A | Q9Y294 | ASF1A | Other | 0.503117099 | 0.514351996 | 24.74051988 | 24.70767256 | 24.47302146 | 21.97047669 | 24.20888387 | 24.48578903 | 24.7161249 | 24.37636813 | 24.71004 |
| Histone chaperone ASF1B | Q9NVP2 | ASF1B | Other | 0.639389302 | 1.321641531 | 24.40704262 | 24.10389606 | 23.74232013 | 24.64663825 | 24.57068494 | 24.91788848 | 25.2190778 | 25.22147679 | 24.67184855 |
| Set1/Ash2 histone methyltransferase | Q9UBL3-3;Q9UBL3;Q9UBL3-2 | ASH2L | Other | 0.067261823 | 0.138678393 | 25.48203833 | 25.80208125 | 25.66562781 | 25.51298008 | 25.90106446 | 25.96170598 | 25.86735831 | 26.00973005 | 25.48744606 |

| | | | | | | | | | | | | | | |
|---|---|---|---|---|---|---|---|---|---|---|---|---|---|---|
| Argininosuccinate lyase | P04424-2;P04424;P04424-3 | ASL | Extracellular or secreted | 0.088447136 | 0.213032799 | 26.53826248 | 26.4594834 | 26.3173102 | 25.95814734 | 26.17163479 | 25.50560951 | 26.13286916 | 26.30127932 | 26.30641679 |
| N-acetylserotonin O-methyltransferas | O95671-2;O95671;O95671-3 | ASMTL | Other | 0.362341961 | 0.189267192 | 25.80474196 | 25.42681602 | 25.66038703 | 23.96622449 | 25.54223911 | 25.72956722 | 26.36284051 | 26.370826 | 24.41834729 |
| ATPase ASNA1 | O43681 | ASNA1 | Extracellular or secreted | 0.058300247 | 0.220593786 | 28.35136025 | 28.39472507 | 28.32388599 | 28.30186365 | 28.26816439 | 28.23169229 | 28.47533509 | 28.45226582 | 28.18314566 |
| Asparagine synthetase [glutamine-hy | P08243-2;P08243;P08243-3 | ASNS | Other | 0.737009101 | 1.129090474 | 27.72084136 | 27.80415111 | 27.78898979 | 28.74567032 | 28.81833208 | 28.18735253 | 28.69265434 | 28.63727007 | 29.4137715 |
| Aspartyl/asparaginyl beta-hydroxylase | Q12797;Q12797-10 | ASPH | Extracellular or secreted | 0.04721792 | 0.016431198 | 22.99291276 | 22.4064199 | 22.65510529 | 22.72336181 | 23.07667535 | 26.16252087 | 22.52886136 | 22.70777786 | 24.91351253 |
| Tether containing UBX domain for GLU | Q9BZE9;Q9BZE9-2;Q9BZE9-3 | ASPSCR1 | Extracellular or secreted | -0.176239894 | 0.380226924 | 26.10183417 | 25.59653488 | 25.61231487 | 25.07267433 | 25.69423629 | 26.05320569 | 25.41760712 | 25.77618019 | 25.34289312 |
| Isoaspartyl peptidase/L-asparaginase; | Q7L266;Q7L266-2 | ASRGL1 | Other | 0.278576598 | 0.400686239 | 25.30594734 | 25.26799479 | 25.23870561 | 25.47735165 | 25.19155658 | 24.06414541 | 25.46774583 | 25.69178952 | 24.94904513 |
| Argininosuccinate synthase | P00966 | ASS1 | Extracellular or secreted | -1.630687144 | 1.464046541 | 30.33734825 | 30.60867154 | 30.55937601 | 30.12188712 | 30.24104492 | 29.71979749 | 28.96046919 | 29.0401453 | 27.90138676 |
| Protein asunder homolog | Q9NVM9;Q9NVM9-2 | ASUN | Other | -0.728009867 | 0.81920732 | 27.02917937 | 27.264187 | 27.08648792 | 25.94395846 | 26.47275813 | 26.77127627 | 26.31219413 | 26.43507632 | 25.35262352 |
| ATPase family AAA domain-containing | Q9NVI7-2;Q9NVI7;Q9NVI7-3 | ATAD3A | Other | -0.402958552 | 0.185492318 | 23.297541 | 20.80456646 | 25.60721516 | 23.40678319 | 23.84794967 | 26.65621884 | 23.32841307 | 23.43122315 | 23.84162529 |
| Arginyl-tRNA--protein transferase 1 | O95260-2;O95260 | ATE1 | Other | -2.132435522 | 1.25122983 | 27.19666786 | 26.95980618 | 26.69431601 | 27.041347 | 27.66064544 | 28.04884803 | 26.11531311 | 25.22051767 | 24.06767791 |
| Activating transcription factor 7-intera | Q6VMQ6-2;Q6VMQ6-5;Q6VM | ATF7IP | Other | 0.205951471 | 0.292289368 | 23.69922337 | 23.62176215 | 23.7370674 | 22.59208652 | 23.56278929 | 24.19422722 | 23.63798861 | 23.51484694 | 24.16859684 |
| Autophagy-related protein 101 | Q9BSB4 | ATG101 | Other | 0.231305829 | 0.27200592 | 21.24202503 | 21.16398374 | 20.99520276 | 21.00058325 | 21.38336861 | 20.47149661 | 21.46123962 | 20.68551534 | 21.67549253 |
| Ubiquitin-like protein ATG12 | O94817 | ATG12 | Other | 0.819376097 | 1.062019243 | 20.29015607 | 22.0263789 | 21.86566985 | 19.86886776 | 21.94369608 | 20.77240652 | 21.98092053 | 21.80884166 | 22.05195369 |
| Autophagy-related protein 16-1 | Q676U5-2;Q676U5;Q676U5- | ATG16L1 | Other | -0.098431814 | 0.146782834 | 26.52993377 | 25.92193353 | 25.88978486 | 26.12573444 | 24.80272224 | 25.30138399 | 25.69189598 | 25.69824323 | 25.60031176 |
| Ubiquitin-like-conjugating enzyme AT | Q9NT62;Q9NT62-2 | ATG3 | Other | -0.224288404 | 0.08015023 | 24.30396353 | 24.47227783 | 24.59160726 | 24.79684406 | 22.98235561 | 24.10485594 | 24.80857656 | 24.97175964 | 22.1727507 |
| Cysteine protease ATG4B | Q9Y4P1-6;Q9Y4P1;Q9Y4P1-2 | ATG4B | Other | 0.01195797 | 0.014908032 | 25.54489394 | 25.5167114 | 25.54695544 | 25.49448257 | 25.91873024 | 25.59115068 | 25.91520067 | 25.79978615 | 25.12734923 |
| Cysteine protease ATG4C | Q96DT6 | ATG4C | Extracellular or secreted | 0.031213719 | 0.04048964 | 21.62693886 | 21.73983004 | 20.76606309 | 21.52899663 | 22.5171455 | 21.02695293 | 21.61072209 | 21.70998744 | 21.37589516 |
| Autophagy protein 5 | Q9H1Y0;Q9H1Y0-2 | ATG5 | Extracellular or secreted | -0.212518248 | 0.445124598 | 25.74085438 | 25.89585167 | 25.81144584 | 25.66752392 | 25.53005288 | 25.1669127 | 25.61102052 | 25.57842245 | 25.07932299 |
| Ubiquitin-like modifier-activating enz | O95352;O95352-2;O95352-3 | ATG7 | Other | 0.391300879 | 0.740500354 | 26.11002428 | 26.263023 | 26.32290669 | 25.8687717 | 25.63959022 | 25.92675909 | 26.64099707 | 26.59020851 | 26.00823455 |
| Acid trehalase-like protein 1 | Q32M88 | ATHL1 | Extracellular or secreted | 0.093746005 | 0.215760278 | 25.93154603 | 25.83593224 | 25.48784469 | 26.32042969 | 26.41746226 | 25.53287881 | 25.99653762 | 26.14963286 | 25.8981144 |
| Bifunctional purine biosynthesis prote | P31939;P31939-2 | ATIC | Extracellular or secreted | 0.373730104 | 1.045466122 | 32.33202111 | 32.22412028 | 32.24167007 | 32.30083874 | 32.28466024 | 31.60071938 | 32.62849264 | 32.713461 | 32.27125158 |
| Atlastin-2 | Q8NHH9-5;Q8NHH9;Q8NHH9 | ATL2 | Other | 0.075394232 | 0.097370434 | 25.76192532 | 25.89776829 | 25.78341753 | 25.88203698 | 26.28162375 | 26.79017128 | 25.9201851 | 25.91326142 | 26.59120776 |
| Atlastin-3 | Q6DD88 | ATL3 | Other | 0.064431587 | 0.061503648 | 27.32740257 | 27.67808289 | 27.7134528 | 27.22278532 | 27.54377361 | 28.47363311 | 27.53341363 | 27.3069556 | 28.33249069 |
| Copper transport protein ATOX1 | O00244 | ATOX1 | Other | 1.053501651 | 0.397201435 | 25.88550889 | 25.12809693 | 25.35608761 | 25.19174481 | 25.48376117 | 23.63223022 | 27.26847675 | 27.0372676 | 24.19347543 |
| Probable cation-transporting ATPase 1 | Q9H7F0;Q9H7F0-2 | ATP13A3 | Extracellular or secreted | 1.769191615 | 1.712795117 | 21.38697223 | 22.44151772 | 23.99394936 | 22.10729003 | 20.98407878 | 24.02276062 | 24.16246325 | 24.21415664 | 24.39923932 |
| Sodium/potassium-transporting ATPa | P05023-4;P05023;P05023-3;P | ATP1A1 | Extracellular or secreted | 1.415491257 | 4.752649673 | 30.88401863 | 30.74358539 | 31.0313766 | 30.76430038 | 30.92691175 | 31.45513053 | 32.22291197 | 32.47290723 | 32.45331622 |
| Sodium/potassium-transporting ATPa | P05026-2;P05026 | ATP1B1 | Extracellular or secreted | 0.968425838 | 1.917319987 | 27.50460977 | 27.59733084 | 27.74278269 | 27.29998772 | 27.41175289 | 27.38277968 | 28.21729443 | 28.42029244 | 28.73731244 |
| Sodium/potassium-transporting ATPa | P54709;P54709-2 | ATP1B3 | Extracellular or secreted | 1.5495792 | 3.676861027 | 27.62964811 | 27.58348131 | 27.90902023 | 27.48393027 | 27.59562468 | 28.56067015 | 29.07839178 | 29.49958167 | 29.45195153 |
| Sarcoplasmic/endoplasmic reticulum | P16615-5;P16615-2;P16615-3 | ATP2A2 | Extracellular or secreted | -0.125282744 | 0.032979664 | 22.07129999 | 24.47846303 | 24.52067287 | 24.50960154 | 24.33987283 | 28.45473844 | 23.44493298 | 23.79887202 | 26.56767112 |
| Plasma membrane calcium-transporti | P20020-6;P20020;P20020-4;P | ATP2B1 | Extracellular or secreted | 0.60899438 | 3.432858998 | 27.37823836 | 27.26508175 | 27.61168191 | 27.41183369 | 27.16470887 | 27.3948477 | 27.96504663 | 28.08744933 | 27.88768332 |
| Plasma membrane calcium-transporti | P23634-5;P23634-4;P23634-3 | ATP2B4 | Extracellular or secreted | 0.741231536 | 1.815083455 | 25.20377843 | 25.35790042 | 24.89901388 | 25.42306896 | 25.30016233 | 24.69189598 | 25.58078163 | 26.02749299 | 26.05333 |
| ATP synthase subunit alpha, mitochon | P25705;P25705-2;P25705-3 | ATP5A1 | Extracellular or secreted | -0.873146683 | 0.867211602 | 29.21539776 | 29.31997339 | 29.7206457 | 30.18813111 | 30.17949608 | 31.52547318 | 28.69454849 | 28.89417543 | 29.86639464 |
| ATP synthase subunit beta, mitochon | P06576 | ATP5B | Extracellular or secreted | -0.919929771 | 1.750794262 | 30.99118343 | 30.99462653 | 31.1742068 | 31.99361471 | 31.94962962 | 32.32189877 | 30.51116736 | 30.44134357 | 31.00027968 |
| ATP synthase subunit gamma, mitoch | P36542-2;P36542 | ATP5C1 | Extracellular or secreted | -1.844106857 | 0.419905776 | 25.25601917 | 25.09924763 | 25.59303316 | 24.88548562 | 24.80247574 | 28.89469585 | 21.08865638 | 23.8909678 | 26.75353384 |
| ATP synthase subunit delta, mitochon | P30049 | ATP5D | Other | 1.459917863 | 1.093395563 | 25.18082427 | 24.72935977 | 24.84522395 | 26.15911713 | 22.48728397 | 27.39059027 | 26.6007371 | 26.65020419 | 26.52501198 |
| ATP synthase F(0) complex subunit B1 | P24539 | ATP5F1 | Extracellular or secreted | -1.827450238 | 0.402443115 | 24.46157687 | 25.78701018 | 25.84505641 | 22.36807595 | 22.63725901 | 29.00576356 | 20.30829999 | 23.37065974 | 25.89106054 |
| ATP synthase subunit d, mitochondria | O75947;O75947-2 | ATP5H | Extracellular or secreted | -0.316893469 | 0.085101225 | 23.70767256 | 24.20500986 | 23.87942171 | 23.93292046 | 23.29108974 | 27.52544518 | 23.35984498 | 22.61822596 | 26.3420284 |
| ATP synthase subunit f, mitochondrial | P56134-3;P56134;P56134-4;P | ATP5J2 | Extracellular or secreted | -0.807895522 | 0.389622989 | 22.4586389 | 21.46741144 | 23.90048875 | 23.31492726 | 21.70053769 | 26.4953521 | 22.0399592 | 21.76490831 | 23.44012399 |
| ATP synthase subunit O, mitochondria | P48047 | ATP5O | Extracellular or secreted | -1.375825129 | 0.420750917 | 23.24042451 | 20.32725877 | 25.3889987 | 24.60048191 | 24.43045766 | 28.93562045 | 21.66151922 | 22.8927103 | 24.77991609 |
| V-type proton ATPase subunit S1 | Q15904 | ATP6AP1 | Extracellular or secreted | 3.240002647 | 1.212116002 | 22.82440642 | 22.81778702 | 23.6655736 | 21.4378317 | 21.36671689 | 23.88511319 | 25.02195627 | 24.83156588 | 27.86520022 |
| Renin receptor | O75787-2;O75787 | ATP6AP2 | Extracellular or secreted | 1.179858914 | 1.96547002 | 24.14763535 | 25.78753336 | 24.20500986 | 24.07773499 | 24.35070291 | 24.51123147 | 25.437396 | 25.5310649 | 26.11103981 |
| V-type proton ATPase subunit d 1 | P61421 | ATP6V0D1 | Extracellular or secreted | -0.356606543 | 1.084527514 | 26.20394641 | 25.88978486 | 26.33944813 | 26.79215898 | 26.54328687 | 26.89133872 | 25.97696785 | 26.02852613 | 26.25466838 |
| V-type proton ATPase catalytic subun | P38606;P38606-2 | ATP6V1A | Extracellular or secreted | -0.08806018 | 0.425698077 | 29.90684336 | 29.85939097 | 29.81318731 | 29.45724597 | 29.43646699 | 29.5413937 | 29.57900901 | 29.63162676 | 29.53244784 |
| V-type proton ATPase subunit B, brain | P21281 | ATP6V1B2 | Extracellular or secreted | 0.068346528 | 0.204602368 | 29.30530377 | 29.33904879 | 29.22662574 | 28.68985842 | 28.62058811 | 29.02633246 | 29.05450007 | 29.09393472 | 29.16048344 |
| V-type proton ATPase subunit C 1 | P21283 | ATP6V1C1 | Extracellular or secreted | 0.077334381 | 0.12551055 | 27.39664515 | 27.43714197 | 27.55800771 | 27.36076606 | 26.95958511 | 27.47514178 | 27.59917689 | 27.68418337 | 27.04228676 |
| V-type proton ATPase subunit D | Q9Y5K8 | ATP6V1D | Extracellular or secreted | 0.307423721 | 0.634091437 | 26.12419674 | 26.01913749 | 25.93739519 | 25.32563677 | 25.38686291 | 25.22365088 | 26.12072103 | 26.15726767 | 25.65272246 |
| V-type proton ATPase subunit E 1 | P36543;P36543-3;P36543-2 | ATP6V1E1 | Extracellular or secreted | -0.216235291 | 0.376980391 | 24.92220582 | 25.34624535 | 24.95149283 | 26.2464282 | 26.23222314 | 25.39936973 | 25.26538584 | 25.23128945 | 25.40360137 |
| V-type proton ATPase subunit F | Q16864;Q16864-2 | ATP6V1F | Extracellular or secreted | 1.037734794 | 1.036082792 | 23.57646313 | 23.33919324 | 24.22354042 | 24.58233313 | 24.46188906 | 24.44688995 | 24.72910041 | 24.7869105 | 25.91234793 |
| V-type proton ATPase subunit G 1 | O75348 | ATP6V1G1 | Extracellular or secreted | 0.605536447 | 2.04176443 | 26.22600552 | 26.07938403 | 25.88199032 | 25.67303556 | 25.76577413 | 26.21706327 | 26.54237196 | 26.4086792 | 26.78718459 |
| V-type proton ATPase subunit H | Q9UI12-2;Q9UI12 | ATP6V1H | Extracellular or secreted | 0.404687897 | 2.058676449 | 26.66830872 | 26.52008833 | 26.70041453 | 26.22685078 | 26.32681992 | 26.81311084 | 26.93853975 | 27.05982072 | 26.84349978 |
| ATP synthase mitochondrial F1 compl | Q5TC12-3;Q5TC12;Q5TC12-2 | ATPAF1 | Extracellular or secreted | 0.249843988 | 0.667685667 | 29.61932915 | 29.78698214 | 29.75080564 | 30.37885847 | 30.12003379 | 30.72492802 | 30.29985675 | 30.26035568 | 30.37978814 |
| ATP synthase mitochondrial F1 compl | Q8N5M1 | ATPAF2 | Other | -0.260800889 | 0.320346999 | 26.04936749 | 25.95187074 | 26.03695332 | 26.20170502 | 25.47812353 | 26.11107962 | 25.60481403 | 26.26908344 | 25.25824972 |
| Serine/threonine-protein kinase ATR | Q13535-2;Q13535-3;Q13535 | ATR | Other | -3.571924861 | 3.362945629 | 24.73412371 | 24.74693906 | 24.93732784 | 25.31080829 | 24.70445714 | 22.19180276 | 21.05247745 | 20.7730961 | 20.77138125 |
| Attractin | O75882-2;O75882;O75882-3 | ATRN | Extracellular or secreted | 0.452008412 | 0.273334776 | 26.27277158 | 26.4093754 | 26.29416119 | 27.52999333 | 27.62247449 | 28.39966309 | 27.03957022 | 26.93898835 | 28.6416862 |
| Transcriptional regulator ATRX | P46100-4;P46100;P46100-6;P | ATRX | Other | -0.526385475 | 1.644219339 | 25.32865274 | 24.8827367 | 25.0013996 | 24.5020315 | 24.7649647 | 24.99178893 | 24.47450756 | 24.53759617 | 24.14452693 |
| Ataxin-10 | Q9UBB4;Q9UBB4-2 | ATXN10 | Extracellular or secreted | -0.088671128 | 0.09645133 | 27.77442173 | 27.96113188 | 27.97092633 | 28.00077664 | 28.0973298 | 27.91410588 | 28.09958876 | 28.23799487 | 27.25574912 |
| Ataxin-2 | Q99700-2;Q99700-4;Q99700 | ATXN2 | Other | -0.552645464 | 1.920839026 | 24.23534932 | 24.67012026 | 24.78551433 | 24.28432709 | 24.05133973 | 23.84345186 | 23.77936509 | 23.64008691 | 23.8576629 |
| Ataxin-2-like protein | Q8WWM7;Q8WWM7-6;Q8W | ATXN2L | Other | 0.167247818 | 0.853225388 | 28.330013 | 28.35228675 | 28.28709586 | 28.64680989 | 28.63606017 | 28.09239802 | 28.52805653 | 28.49235994 | 28.65365884 |
| Aurora kinase A | O14965 | AURKA | Other | -0.569022694 | 1.130032789 | 25.50148492 | 25.63826488 | 26.15558954 | 26.08691304 | 26.04891037 | 25.53910602 | 25.63034268 | 25.17478064 | 24.97294299 |
| Aurora kinase B | Q96GD4-4;Q96GD4-2;Q96GD | AURKB | Extracellular or secreted | -0.809134949 | 1.13180012 | 22.61950268 | 22.77589396 | 21.76036898 | 22.32311635 | 22.65252553 | 23.3321065 | 21.69867245 | 21.32965187 | 22.27602783 |
| Late secretory pathway protein AVL9 | Q8NBF6-2;Q8NBF6 | AVL9 | Other | -0.4437768 | 0.12951679 | 24.52737057 | 24.90412338 | 24.89555125 | 24.46531878 | 24.37032716 | 24.33053453 | 25.02081246 | 25.58081037 | 21.8136596 |
| Zinc-alpha-2-glycoprotein | P25311 | AZGP1 | Extracellular or secreted | -1.692040423 | 0.681895727 | 23.35729641 | 22.67641855 | 24.46076484 | 25.23165568 | 24.39335903 | 23.80587374 | 23.20602407 | 20.39734188 | 23.2831969 |
| Beta-1,3-glucosyltransferase | Q6Y288 | B3GALTL | Extracellular or secreted | 0.175288455 | 0.38301975 | 23.70904089 | 23.98849886 | 24.01699323 | 23.1280025 | 23.01162925 | 23.89513517 | 23.80390486 | 23.64801081 | 23.94859965 |
| Beta-1,4-galactosyltransferase 1;Lacto | P15291-2;P15291 | B4GALT1 | Extracellular or secreted | -1.254459412 | 1.714713023 | 25.741806 | 25.70606574 | 25.67111968 | 25.90455984 | 25.60902043 | 25.90653374 | 24.33749288 | 24.26842321 | 24.9002584 |
| Beta-1,4-galactosyltransferase 5 | O43286 | B4GALT5 | Extracellular or secreted | -0.415292825 | 0.822268062 | 24.24372546 | 24.09844466 | 24.03405874 | 23.32745396 | 23.09588824 | 23.47864818 | 23.62343769 | 23.18756774 | 23.08222572 |
| BRISC and BRCA1-A complex member | Q9NWV8;Q9NWV8-3 | BABAM1 | Other | -0.10058323 | 0.155473865 | 26.90999303 | 27.04353883 | 26.8811035 | 27.08729757 | 26.89573613 | 26.14612083 | 26.8766612 | 26.96851141 | 26.33497265 |
| BAG family molecular chaperone regu | Q99933-4;Q99933-3;Q99933 | BAG1 | Other | -1.740412102 | 0.382579324 | 25.58437053 | 25.48419156 | 25.7206457 | 25.49963108 | 25.50691116 | 25.10952621 | 25.54904338 | 25.34329986 | 20.33905857 |
| BAG family molecular chaperone regu | O95816;O95816-2 | BAG2 | Other | 0.444610222 | 0.811185054 | 27.48631089 | 27.5803216 | 27.60434747 | 26.9446958 | 27.05320569 | 27.2499758 | 28.03039023 | 27.91547423 | 27.34739483 |
| BAG family molecular chaperone regu | O95817 | BAG3 | Extracellular or secreted | 0.69578972 | 2.735986311 | 27.76395229 | 27.84218132 | 27.85715268 | 27.95632041 | 28.0877528 | 27.41183369 | 28.50979028 | 28.36252284 | 28.67465262 |
| BAG family molecular chaperone regu | Q9UL15;Q9UL15-2 | BAG5 | Other | 0.283920817 | 1.082137879 | 27.10169393 | 27.3569943 | 27.39116412 | 27.71790355 | 27.88623019 | 27.12966978 | 27.65129963 | 27.68224206 | 27.8100487 |
| Large proline-rich protein BAG6 | P46379-2;P46379;P46379-3;P | BAG6 | Other | -0.277257128 | 0.473044538 | 26.2557131 | 26.08450235 | 26.2641512 | 26.65198385 | 26.67653692 | 27.23790373 | 26.05384785 | 26.05492438 | 26.64485197 |
| Brain-specific angiogenesis inhibitor 1 | Q9UQB8-3;Q9UQB8-5;Q9UQ | BAIAP2 | Extracellular or secreted | 1.092667313 | 2.937517617 | 25.23976196 | 25.00987952 | 24.88804347 | 24.881757 | 24.13375118 | 25.4172208 | 25.8276224 | 26.0993881 | 26.1361984 |
| Brain-specific angiogenesis inhibitor 1 | Q9UHR4 | BAIAP2L1 | Extracellular or secreted | -0.219884569 | 0.42588399 | 26.44917497 | 26.18544441 | 26.31934467 | 26.95515661 | 27.03779124 | 26.76141813 | 26.23770319 | 26.212228 | 26.74458012 |
| Barrier-to-autointegration factor;Barr | O75531 | BANF1 | Extracellular or secreted | 0.759945635 | 1.335085281 | 29.68266067 | 29.55438913 | 29.72245462 | 29.63289236 | 29.84696996 | 30.02088663 | 30.87136814 | 30.42602418 | 30.21257127 |
| Chromatin complexes subunit BAP18 | Q8IXM2;Q8IXM2-3 | BAP18 | Other | 0.534927972 | 1.619636028 | 26.9139918 | 26.72234061 | 27.06855968 | 26.1251235 | 26.06445382 | 26.70898828 | 27.15392879 | 27.05216934 | 27.20041463 |
| Brain acid soluble protein 1 | P80723;P80723-2 | BASP1 | Extracellular or secreted | -1.813051374 | 0.804233327 | 28.68137097 | 28.83315956 | 28.84908983 | 28.37550668 | 28.29131813 | 29.80717971 | 27.92710962 | 25.31150138 | 27.74104733 |
| Apoptosis regulator BAX | Q07812-8;Q07812;Q07812-2 | BAX | Extracellular or secreted | 0.6892922 | 0.786924179 | 27.24644633 | 27.22765883 | 27.25592916 | 26.68525332 | 26.63920379 | 27.63207765 | 28.18267221 | 28.09597273 | 27.1325162 |
| Tyrosine-protein kinase BAZ1B | Q9UIG0-2;Q9UIG0 | BAZ1B | Other | -4.010809 | 4.383937934 | 28.06578951 | 26.7278029 | 28.13011183 | 28.83653429 | 26.75646368 | 28.00678028 | 23.61548955 | 23.92469944 | 23.68912526 |
| Basal cell adhesion molecule | P50895 | BCAM | Extracellular or secreted | -1.055534346 | 3.25051264 | 28.81726494 | 28.7521305 | 28.67838517 | 29.37799024 | 29.40019261 | 29.29085692 | 28.02189275 | 27.80415111 | 28.16576329 |
| B-cell receptor-associated protein 31 | P51572;P51572-2 | BCAP31 | Extracellular or secreted | -0.19259 | 0.174364162 | 26.77881388 | 26.29958605 | 26.57127827 | 25.93116275 | 25.67087664 | 27.59263405 | 25.82514921 | 26.11844544 | 26.90081117 |
| Breast cancer anti-estrogen resistance | P56945-4;P56945-5;P56945-8 | BCAR1 | Extracellular or secreted | 1.302505786 | 1.138560945 | 23.26728048 | 20.49308777 | 22.55325711 | 21.58364216 | 20.69863962 | 21.00200023 | 23.3325163 | 23.27582896 | 22.09812578 |
| Pre-mRNA-splicing factor SPF27 | O75934 | BCAS2 | Other | 0.028748922 | 0.112860279 | 26.37532439 | 26.34866157 | 26.09780196 | 26.32935442 | 26.35743066 | 26.50685064 | 26.2168968 | 26.41511029 | 26.4619515 |
| Branched-chain-amino-acid aminotra | O15382;O15382-2 | BCAT2 | Other | -1.366552075 | 3.714497645 | 30.54000822 | 30.59646024 | 30.54637042 | 30.95843777 | 30.96285623 | 30.04492006 | 29.44413128 | 29.23986209 | 29.04087689 |
| BRCA2 and CDKN1A-interacting prote | Q9P287;Q9P287-3;Q9P287-4 | BCCIP | Other | 0.097161621 | 0.143023529 | 29.50339704 | 29.60865392 | 29.57116252 | 29.28528883 | 29.39770623 | 28.56318219 | 29.66636285 | 29.56356407 | 29.0262533 |
| Bcl-2-like protein 1 | Q07817-2;Q07817-3;Q07817 | BCL2L1 | Extracellular or secreted | -0.435199107 | 0.16449018 | 22.4066794 | 21.48880709 | 22.08440097 | 20.25185063 | 22.20514028 | 22.85617443 | 22.47874691 | 22.18225734 | 19.67992483 |
| Bcl-2-like protein 13 | Q9BXK5;Q9BXK5-2 | BCL2L13 | Other | -0.297474533 | 0.240880719 | 28.64756487 | 28.77149667 | 28.63640596 | 27.54701429 | 27.60533697 | 27.7978584 | 28.36331687 | 28.20648246 | 27.04061565 |
| B-cell CLL/lymphoma 7 protein family | Q8WUZ0;Q8WUZ0-2 | BCL7C | Other | 1.590876261 | 1.546937653 | 21.81108267 | 20.98549365 | 22.52615989 | 20.52670749 | 22.24453385 | 24.13375118 | 23.40262593 | 23.40898686 | 24.07488035 |
| Bcl-2-associated transcription factor 1 | Q9NYF8-2;Q9NYF8;Q9NYF8-4 | BCLAF1 | Other | -0.535708321 | 0.770501442 | 27.78494051 | 27.92722267 | 28.23904263 | 27.94055735 | 27.84774661 | 28.37608656 | 27.00838417 | 27.51885854 | 27.9234305 |
| Breakpoint cluster region protein | P11274-2;P11274 | BCR | Extracellular or secreted | 0.329771346 | 0.44168701 | 24.33558608 | 22.9274691 | 23.56627799 | 24.26018965 | 23.7716289 | 23.8305508 | 23.61189293 | 24.41222146 | 24.31105091 |
| Protein bicaudal D homolog 2 | Q8TD16;Q8TD16-2 | BICD2 | Extracellular or secreted | 1.615498642 | 2.113098592 | 24.94073656 | 24.93665411 | 24.52246995 | 24.51352232 | 24.80503729 | 25.09018836 | 26.17337063 | 26.22890674 | 26.84852285 |
| BH3-interacting domain death agonist | P55957;P55957-2;P55957-4 | BID | Extracellular or secreted | 0.706025069 | 0.534624408 | 27.04978292 | 26.99744242 | 27.51389876 | 27.39811412 | 27.49952464 | 27.33142324 | 28.48181513 | 28.52671414 | 27.00463899 |
| Myc box-dependent-interacting prote | O00499-9;O00499-10;O0049 | BIN1 | Extracellular or secreted | -0.317557132 | 0.797322691 | 25.11878206 | 25.24212682 | 25.2643839 | 24.70851476 | 25.20224701 | 24.11330723 | 24.4865871 | 24.66068624 | 24.72473616 |
| Baculoviral IAP repeat-containing prot | Q9NR09 | BIRC6 | Other | 0.141681573 | 0.166713096 | 26.53203161 | 26.47767589 | 26.56361862 | 26.27289613 | 26.3669138 | 26.39791835 | 27.11038279 | 26.5339037 | 26.06828543 |
| Bleomycin hydrolase | Q13867 | BLMH | Extracellular or secreted | -0.317321916 | 1.06087234 | 31.48346905 | 31.62566523 | 31.47416519 | 31.66661681 | 31.86599661 | 30.88743639 | 31.34884094 | 31.08182359 | 31.11904437 |
| Biogenesis of lysosome-related organ | P78537-2;P78537 | BLOC1S1 | Extracellular or secreted | 0.579230425 | 0.946123417 | 23.4693616 | 23.86834783 | 23.9400196 | 21.97591168 | 23.26470603 | 23.6292312 | 23.7267121 | 23.99963742 | 24.08513073 |
| Biogenesis of lysosome-related organ | Q6QNY1 | BLOC1S2 | Other | 1.089973111 | 0.76813017 | 24.98910549 | 24.93719312 | 25.02474842 | 22.30113182 | 21.277775 | 24.85005022 | 25.01818252 | 25.10337586 | 24.83836298 |
| Biogenesis of lysosome-related organ | Q6QNY0 | BLOC1S3 | Other | -0.292406934 | 0.080970403 | 23.4537499 | 23.46063987 | 23.81753333 | 22.46805553 | 22.02771846 | 21.94849105 | 23.60761026 | 20.2299308 | 23.87333221 |
| Biogenesis of lysosome-related organ | Q9UL45 | BLOC1S6 | Other | 0.276716071 | 0.419931279 | 23.43288032 | 24.08513073 | 23.98102509 | 23.93571037 | 23.68731075 | 23.42764738 | 24.22331945 | 24.3280021 | 23.55367897 |
| Biliverdin reductase A | P53004 | BLVRA | Extracellular or secreted | 0.126710409 | 0.207126597 | 28.69803122 | 28.86732296 | 28.78159909 | 28.2124785 | 28.2414996 | 28.49944861 | 28.91430552 | 28.84343988 | 28.27257583 |
| Flavin reductase (NADPH) | P30043 | BLVRB | Extracellular or secreted | -0.405229094 | 0.521165956 | 30.83778274 | 30.71587624 | 30.69544485 | 30.41689476 | 30.22745227 | 29.91698361 | 30.2783039 | 30.42612419 | 29.48510187 |
| Bone morphogenetic protein 1 | P13497;P13497-5;P13497-2;P | BMP1 | Extracellular or secreted | 0.061762047 | 0.021378103 | 22.56796809 | 23.16911699 | 20.85968503 | 23.74990788 | 23.64768151 | 25.03615684 | 21.71151581 | 23.69274745 | 24.29628106 |
| BMP-2-inducible protein kinase | Q9NSY1-2;Q9NSY1-3;Q9NSY1 | BMP2K | Other | 0.01223831 | 0.011772189 | 22.20993997 | 22.22702712 | 22.93016126 | 22.82783424 | 21.24175177 | 21.52725843 | 22.47395045 | 22.29471161 | 21.75003927 |
| Bone morphogenetic protein 7 | P18075 | BMP7 | Extracellular or secreted | 0.738808395 | 0.807101723 | 22.30069999 | 22.86870577 | 22.87837428 | 23.22714467 | 23.56801919 | 23.7163866 | 23.71910551 | 23.28714872 | 24.4898362 |
| Ribosome biogenesis protein BMS1 ho | Q14692 | BMS1 | Other | -2.744812433 | 1.741426155 | 26.49993516 | 26.19889836 | 26.33868334 | 25.09454391 | 25.25806996 | 26.63325623 | 22.37243313 | 23.19586478 | 24.20895827 |
| Vesicle transport protein SEC20 | Q12981;Q12981-1 | BNIP1 | Other | -1.035366375 | 0.74275179 | 23.02607304 | 23.30215137 | 23.55964222 | 24.26656581 | 22.67083343 | 23.49753125 | 22.95624285 | 22.81213018 | 21.2869264 |
| BCL2/adenovirus E1B 19 kDa protein- | Q12982;Q12982-2 | BNIP2 | Other | 1.133876994 | 1.226937502 | 21.80323234 | 19.85117172 | 20.89572514 | 22.77593385 | 21.67274649 | 21.19124243 | 22.15013277 | 23.09921552 | 22.24730868 |
| Biorientation of chromosomes in cell | Q8NFC6 | BOD1L1 | Other | -0.338545076 | 1.035764749 | 27.17645563 | 26.52607233 | 26.71502522 | 26.84098167 | 26.12145242 | 26.39790203 | 26.3757718 | 26.38101439 | 26.11652323 |
| BolA-like protein 2 | Q9H3K6;Q9H3K6-2 | BOLA2 | Extracellular or secreted | 1.147038216 | 1.292612926 | 28.04306943 | 27.56514509 | 27.63844442 | 28.08967329 | 28.2337048 | 27.02686009 | 29.30510804 | 29.18416305 | 28.25029213 |
| BolA-like protein 3 | Q53S33 | BOLA3 | Extracellular or secreted | -1.932708032 | 0.759146329 | 24.83282165 | 20.27989144 | 22.10523915 | 24.43726899 | 24.74447747 | 25.31806915 | 19.9274246 | 22.11714108 | 23.01619415 |
| Ribosome biogenesis protein BOP1 | Q14137;Q14137-2 | BOP1 | Other | 0.137141172 | 0.088505336 | 23.60410706 | 23.31105091 | 22.69557495 | 24.03741424 | 24.03699523 | 26.26018965 | 23.9791942 | 23.75491033 | 24.649985 |
| Valacyclovir hydrolase | Q86WA6-2;Q86WA6 | BPHL | Extracellular or secreted | -0.727969597 | 1.20107439 | 27.48608068 | 27.63816819 | 27.65928485 | 27.70885677 | 27.6338798 | 27.18839054 | 27.01977378 | 27.06002695 | 26.39362089 |
| 3(2),5-bisphosphate nucleotidase 1 | O95861;O95861-2;O95861-4 | BPNT1 | Extracellular or secreted | -0.056315715 | 0.110879765 | 29.58375029 | 29.87532877 | 29.71082829 | 29.19448556 | 29.28219841 | 28.75315124 | 29.46828204 | 29.39345723 | 29.16918487 |
| Nucleosome-remodeling factor subun | Q12830-4;Q12830-2;Q12830 | BPTF | Extracellular or secreted | -1.384285785 | 1.713147432 | 24.67702105 | 24.41073445 | 25.32453849 | 23.55332743 | 24.00032535 | 24.71539188 | 23.38086581 | 23.35997899 | 22.44696716 |
| BRCA1-associated ATM activator 1 | Q6PJG6;Q6PJG6-2 | BRAT1 | Other | 0.20294357 | 0.207368377 | 22.12335228 | 22.23069598 | 22.19592484 | 22.39604515 | 23.67899502 | 22.34163826 | 23.22010432 | 22.68416732 | 22.18788484 |
| Lys-63-specific deubiquitinase BRCC36 | P46736-2;P46736;P46736-3;P | BRCC3 | Other | -0.15075488 | 0.250461363 | 28.14253207 | 28.29701616 | 28.14326222 | 27.52857823 | 27.79358595 | 27.36536263 | 27.82559809 | 27.99085894 | 27.36644695 |
| Bromodomain-containing protein 2 | P25440-4;P25440-3;P25440;P | BRD2 | Other | -1.119718884 | 2.419774318 | 25.52784779 | 25.47978939 | 25.44468027 | 26.13588539 | 25.4945436 | 25.18343912 | 24.29010549 | 24.23622563 | 24.74760501 |
| Bromodomain-containing protein 3 | Q15059;Q15059-2 | BRD3 | Other | -0.886305321 | 1.37942048 | 27.33858134 | 27.0658922 | 27.15865499 | 27.12514321 | 26.85281446 | 27.92925615 | 26.70688254 | 26.47991271 | 25.88945996 |
| Bromodomain-containing protein 4 | O60885;O60885-2;O60885-3 | BRD4 | Other | -0.451888584 | 1.146800981 | 24.00023938 | 24.42566411 | 23.97412536 | 23.57427014 | 23.68238943 | 24.11068147 | 23.69370474 | 23.62366094 | 23.21065351 |
| Bromodomain-containing protein 7 | Q9NPI1;Q9NPI1-2 | BRD7 | Other | -0.304950208 | 0.180427789 | 24.18850373 | 24.0857791 | 23.97465055 | 21.60020245 | 23.22690957 | 24.3043814 | 23.03420991 | 22.5321946 | 24.20895827 |
| BRCA1-A complex subunit BRE | Q9NXR7;Q9NXR7-4;Q9NXR7- | BRE | Other | -0.429595095 | 1.321171373 | 26.36781371 | 26.07856993 | 26.09665645 | 25.357632 | 25.63231344 | 25.47218485 | 25.47357892 | 25.43323063 | 25.30699035 |
| BRI3-binding protein | Q8WY22 | BRI3BP | Other | -0.527310544 | 0.187217118 | 21.48291335 | 20.61180776 | 22.15437313 | 20.86340288 | 21.05337972 | 24.6258359 | 19.82826935 | 21.09736103 | 22.88829436 |
| Ribosome biogenesis protein BRX1 ho | Q8TDN6 | BRIX1 | Other | 0.079681493 | 0.073687124 | 24.69922337 | 24.72296592 | 25.11012388 | 25.24212682 | 25.32793359 | 27.18744692 | 25.44950561 | 25.61369236 | 25.32075676 |
| Protein BRICK1 | Q8WUW1;Q8WUW1-2 | BRK1 | Extracellular or secreted | 0.629637106 | 1.211437517 | 26.13230046 | 25.93874164 | 25.74752819 | 25.6916298 | 25.9730306 | 26.05540028 | 26.84553503 | 26.20869786 | 26.60399391 |
| BRO1 domain-containing protein BRO | Q5VW32;Q5VW32-2 | BROX | Extracellular or secreted | -0.225167944 | 0.274443082 | 29.05406011 | 28.95479619 | 28.91527476 | 27.72416367 | 27.55756957 | 28.61687803 | 28.45147997 | 28.45763738 | 27.82674999 |
| Basigin | P35613-4;P35613-2;P35613 | BSG | Extracellular or secreted | 1.351498593 | 3.213098683 | 28.34218951 | 28.29021977 | 28.48945343 | 28.51987095 | 28.4740201 | 28.80654992 | 29.62524718 | 29.9938846 | 29.89651585 |
| Basigin | P35613-3 | BSG | Extracellular or secreted | 0.987746921 | 0.471745641 | 22.84028542 | 22.39926541 | 22.93876407 | 21.73653076 | 22.96533693 | 23.30549513 | 22.34794895 | 23.6224326 | 25.08569807 |
| TATA-binding protein-associated facto | O14981 | BTAF1 | Other | 1.002325148 | 0.535794848 | 25.92895131 | 26.03896352 | 26.30323197 | 21.09391371 | 26.09729966 | 26.46881804 | 26.39635118 | 26.28577387 | 26.29043951 |
| Transcription factor BTF3 | P20290;P20290-2 | BTF3 | Other | 0.974084247 | 1.181718559 | 27.34840831 | 27.42678403 | 27.75276855 | 28.16466092 | 28.27927851 | 27.84212136 | 29.04762656 | 29.12235625 | 28.15928077 |
| Transcription factor BTF3 homolog 4 | Q96K17;Q96K17-2 | BTF3L4 | Other | 0.426397947 | 0.685290351 | 30.01051991 | 29.93142767 | 30.07444146 | 29.58378615 | 29.48569691 | 29.71220675 | 30.38740948 | 30.53380347 | 29.75702032 |
| Mitotic checkpoint serine/threonine-p | O60566;O60566-3;O60566-2 | BUB1B | Other | 0.165596275 | 0.420919776 | 23.07517423 | 22.69795171 | 22.63617498 | 23.41415874 | 23.6994352 | 23.23211698 | 23.36372626 | 23.29530035 | 23.21526814 |
| Mitotic checkpoint protein BUB3 | O43684-2;O43684 | BUB3 | Other | 0.090034421 | 0.272387153 | 29.37418037 | 29.47154158 | 29.36603 | 29.4915383 | 29.37695594 | 30.03930874 | 29.61966148 | 29.45475804 | 29.7554612 |
| BUD13 homolog | Q9BRD0 | BUD13 | Other | -1.50185457 | 1.881111397 | 22.68764414 | 24.26792338 | 24.08350854 | 23.72296592 | 24.11195516 | 21.79723414 | 21.69000842 | 21.8114208 | 22.32862271 |
| Protein BUD31 homolog | P41223;P41223-2 | BUD31 | Other | -0.261584716 | 0.328739491 | 24.10181413 | 23.83663059 | 24.33114985 | 24.86499957 | 24.72338264 | 25.34252017 | 24.18177223 | 23.8930993 | 24.74062281 |
| Bystin | Q13895 | BYSL | Other | 0.424065731 | 0.73370176 | 24.49801844 | 24.81347786 | 24.88748578 | 25.31706863 | 25.60379588 | 25.66584464 | 25.43510812 | 25.27937594 | 25.95055874 |
| Basic leucine zipper and W2 domain-c | Q7L1Q6;Q7L1Q6-4;Q7L1Q6-3 | BZW1 | Other | 0.203347219 | 0.375357395 | 28.75197094 | 28.83617309 | 28.87841637 | 29.50354869 | 29.48287309 | 28.99080486 | 29.36467408 | 29.54298074 | 28.92428036 |
| Basic leucine zipper and W2 domain-c | Q9Y6E2 | BZW2 | Other | 0.051884956 | 0.109721448 | 28.45238366 | 28.4326892 | 28.47080494 | 28.64948485 | 28.60925298 | 28.22536206 | 28.71158333 | 28.64357962 | 28.22048077 |
| UPF0693 protein C10orf32 | Q96B45 | C10orf32 | Other | -0.187346369 | 0.224538541 | 23.50272681 | 22.13611624 | 22.21285138 | 22.70545943 | 22.44954969 | 22.78863391 | 22.72640029 | 22.68090975 | 21.92831959 |
| Ester hydrolase C11orf54 | Q9H0W9;Q9H0W9-2;Q9H0W | C11orf54 | Extracellular or secreted | -0.439878133 | 1.281637448 | 28.64330436 | 28.81335553 | 28.88401863 | 28.90833315 | 29.0150213 | 27.85887247 | 28.35447431 | 28.20419835 | 28.18314566 |
| UPF0696 protein C11orf68 | Q9H3H3;Q9H3H3-2;Q9H3H3- | C11orf68 | Other | 0.201558707 | 0.476955555 | 26.66329489 | 26.83737674 | 26.63478004 | 26.4964497 | 26.76243233 | 25.64375162 | 26.65430781 | 26.84840345 | 26.62100752 |
| Protein Hikeshi | Q53FT3 | C11orf73 | Extracellular or secreted | 0.232420047 | 0.610523614 | 27.21715574 | 27.40997418 | 27.47568297 | 26.78019151 | 26.43993383 | 27.47583756 | 27.38673137 | 27.39966309 | 27.31025358 |
| Uncharacterized protein C11orf84 | Q9BUA3 | C11orf84 | Other | 0.589643988 | 0.578613746 | 22.35361315 | 22.25649435 | 22.44510985 | 22.21231707 | 19.8465457 | 21.00720247 | 22.01474789 | 22.08128318 | 22.73354219 |
| UPF0160 protein MYG1, mitochondria | Q9HB07 | C12orf10 | Extracellular or secreted | 0.944047213 | 1.881909025 | 25.81137234 | 26.31670621 | 26.25428993 | 26.15013665 | 26.34465486 | 24.71816493 | 27.07520688 | 26.87537271 | 26.6792245 |

| | | | | | | | | | | | | | | |
|---|---|---|---|---|---|---|---|---|---|---|---|---|---|---|
| Uncharacterized protein C12orf29 | Q8N999-3;Q8N999 | C12orf29 | Other | 1.694469005 | 0.959214955 | 26.35952666 | 25.87897758 | 26.03275638 | 25.15572464 | 23.8930993 | 21.08400191 | 25.91597563 | 26.07263345 | 27.29684117 |
| Protein C10 | Q99622 | C12orf57 | Other | 2.638093132 | 2.036761146 | 21.19430694 | 20.34336718 | 21.75945044 | 24.26977902 | 24.1718829 | 22.27278271 | 25.11739548 | 25.07512525 | 24.72754326 |
| Uncharacterized protein C14orf142 | Q9BXV9 | C14orf142 | Other | 0.597501872 | 0.561817311 | 25.48937687 | 25.36942881 | 25.41969792 | 25.5463373 | 25.4241909 | 23.58083912 | 25.8250764 | 26.32920042 | 25.05316425 |
| UPF0568 protein C14orf166 | Q9Y224 | C14orf166 | Other | -0.344586136 | 0.780545036 | 29.0427825 | 29.17214527 | 29.20762317 | 29.25959268 | 29.14214251 | 29.10309568 | 28.78674852 | 29.10439608 | 28.5387879 |
| Uncharacterized protein C15orf52 | Q6ZUT6-4;Q6ZUT6;Q6ZUT6-2 | C15orf52 | Other | 0.450586503 | 0.918568741 | 24.72093267 | 24.11028321 | 24.16859684 | 24.55238955 | 24.81088225 | 23.46824319 | 24.92179736 | 24.89300669 | 24.45261931 |
| UPF0505 protein C16orf62 | Q7Z3J2;Q7Z3J2-2 | C16orf62 | Other | 0.94147697 | 1.309330081 | 23.96296377 | 24.22633634 | 24.4368879 | 23.16429072 | 22.26810124 | 23.46363612 | 24.83006717 | 24.66177375 | 24.09369802 |
| Uncharacterized protein C17orf59 | Q96GS4 | C17orf59 | Other | 1.170015235 | 1.346937659 | 23.43084045 | 23.70535396 | 23.60354123 | 21.47925099 | 24.25154771 | 22.19203908 | 24.18895641 | 24.674491 | 23.977885 |
| Protein Njmu-R1 | Q9HAS0 | C17orf75 | Extracellular or secreted | -0.040149076 | 0.216903948 | 24.83306302 | 24.92537877 | 24.98628683 | 24.81518939 | 25.03565357 | 24.89998193 | 24.92261416 | 24.75353384 | 24.95118154 |
| Uncharacterized protein C17orf85 | Q53F19 | C17orf85 | Other | 0.007973807 | 0.005252299 | 21.9820514 | 22.64513242 | 22.47372755 | 24.16246325 | 21.70528195 | 23.52061293 | 22.17046419 | 22.76066716 | 23.33742482 |
| Uncharacterized protein C18orf25 | Q96B23-2;Q96B23 | C18orf25 | Other | 0.680483871 | 0.528347492 | 21.73978885 | 24.89869105 | 24.85238587 | 24.51929359 | 24.5234873 | 23.81909384 | 25.30709461 | 24.8110783 | 24.09964895 |
| Uncharacterized protein C18orf8 | Q96DM3 | C18orf8 | Other | 1.886234135 | 1.221535506 | 25.14903196 | 24.70846214 | 24.78216799 | 22.70965096 | 22.94802923 | 24.64041795 | 26.61540536 | 26.66370207 | 24.84847509 |
| Uncharacterized protein C19orf43 | Q9BQ61 | C19orf43 | Other | 0.620782362 | 2.182924518 | 27.45646283 | 27.69750106 | 27.70001758 | 28.3254738 | 28.2478501 | 28.12114607 | 28.62787543 | 28.46754374 | 28.54115364 |
| UPF0692 protein C19orf54 | Q5BKX5;Q5BKX5-3 | C19orf54 | Other | -0.704232412 | 0.508605469 | 22.50076798 | 23.15322915 | 22.88745789 | 22.49848112 | 22.79042487 | 22.7696127 | 22.23192662 | 22.8736326 | 21.08173039 |
| UPF0587 protein C1orf123 | Q9NWV4 | C1orf123 | Extracellular or secreted | 0.388746034 | 1.855451697 | 28.24567593 | 28.11083079 | 28.07877349 | 28.41828294 | 28.49537497 | 27.97563477 | 28.68214158 | 28.69172297 | 28.45466 |
| UPF0688 protein C1orf174 | Q8IYL3 | C1orf174 | Other | 0.186047662 | 0.659879778 | 23.72546445 | 23.96305199 | 23.73510561 | 23.80597211 | 24.03439464 | 23.77735967 | 24.0736552 | 23.82036052 | 24.18480151 |
| Uncharacterized protein C1orf50 | Q9BV19 | C1orf50 | Other | 0.236763869 | 0.094319172 | 21.14444013 | 22.93635356 | 23.49570281 | 23.22352569 | 23.73665462 | 23.22881869 | 23.29263506 | 21.78423395 | 24.51617035 |
| Complement component 1 Q subcom | Q07021 | C1QBP | Extracellular or secreted | -0.219277645 | 0.61402302 | 31.06072277 | 31.28488083 | 31.19054674 | 31.90192615 | 31.84860493 | 31.0119864 | 31.27086187 | 31.12083719 | 31.09980192 |
| UPF0687 protein C20orf27 | Q9GZN8;Q9GZN8-2 | C20orf27 | Other | -0.029033523 | 0.040477475 | 27.84272084 | 27.99841121 | 27.84379926 | 28.50134824 | 27.86183281 | 27.34933671 | 28.0207277 | 28.10504584 | 27.48585043 |
| UPF0769 protein C21orf59 | P57076 | C21orf59 | Other | 0.348704478 | 0.68565753 | 25.64589652 | 25.82553744 | 25.61051372 | 26.18306045 | 26.08573859 | 25.54548324 | 26.3383093 | 26.37868487 | 25.77723424 |
| C2 domain-containing protein 5 | Q86YS7;Q86YS7-2;Q86YS7-3; | C2CD5 | Extracellular or secreted | -0.132191774 | 0.29803049 | 25.13182965 | 25.03124249 | 25.21178257 | 24.71082829 | 24.69242821 | 25.50563979 | 24.82318218 | 24.7430396 | 25.1790784 |
| Ashwin | Q9BVC5-2;Q9BVC5 | C2orf49 | Other | 0.877401167 | 1.860584857 | 21.55638732 | 23.23429428 | 22.2049651 | 22.71870559 | 22.13771164 | 22.18068121 | 23.38297752 | 23.30646894 | 22.9591296 |
| UPF0565 protein C2orf69 | Q8N8R5 | C2orf69 | Extracellular or secreted | -0.30364159 | 0.176407751 | 24.17249343 | 21.45568531 | 24.27731976 | 22.7408431 | 24.46780801 | 23.90334202 | 23.15190269 | 22.41716039 | 24.02875796 |
| Complement C3;Complement C3 beta | P01024 | C3 | Extracellular or secreted | 0.702899662 | 1.689312286 | 31.00998627 | 31.02240877 | 31.00390219 | 30.98114489 | 31.02663587 | 31.9030762 | 31.6042149 | 31.82971645 | 32.14834474 |
| UPF0609 protein C4orf27 | Q9NWY4 | C4orf27 | Other | -0.327355319 | 0.495656194 | 25.81682566 | 25.90081117 | 25.58883699 | 25.59166432 | 25.67430241 | 25.36603 | 25.82730742 | 25.11676117 | 25.04310073 |
| UPF0489 protein C5orf22 | Q49AR2;Q49AR2-2 | C5orf22 | Other | 0.236675745 | 0.353794354 | 27.41482005 | 27.2178953 | 27.22158742 | 27.61308811 | 27.72708878 | 27.1710239 | 27.93837149 | 27.80783993 | 27.14656759 |
| Putative uncharacterized protein C5o | C9J3I9 | C5orf58 | Other | -5.155699579 | 2.065548574 | 27.41683437 | 27.47560567 | 27.44758372 | 27.38730676 | 27.44529617 | 27.84134167 | 22.76127608 | 22.94166807 | 21.33694129 |
| Uncharacterized protein C6orf132 | Q5T0Z8 | C6orf132 | Other | -0.882268867 | 0.382417097 | 23.35110746 | 23.17840226 | 23.17273758 | 21.76555658 | 22.12244577 | 23.27042081 | 20.33293902 | 22.18216639 | 23.26842321 |
| Uncharacterized protein C6orf203 | Q9P0P8 | C6orf203 | Other | -1.122778885 | 0.741341506 | 24.52623654 | 24.69264104 | 25.45525597 | 24.59337516 | 25.27009995 | 24.63644053 | 22.64965615 | 24.58600411 | 23.98302768 |
| Uncharacterized protein C7orf50 | Q9BRJ6 | C7orf50 | Other | 1.390401488 | 1.950849093 | 23.26083571 | 22.00326594 | 22.94966858 | 23.56732296 | 23.74929414 | 24.68244301 | 24.572363 | 25.07769425 | 24.62756238 |
| UPF0562 protein C7orf55 | Q96HJ9 | C7orf55 | Other | -1.848567731 | 0.741536876 | 23.06126373 | 23.04181487 | 22.96119814 | 22.84789234 | 22.68483079 | 23.11383181 | 22.86540555 | 19.77011409 | 20.674193 |
| UPF0562 protein C7orf55;Putative RN | Q96HJ9-2;Q9Y383-3;Q9Y383- | C7orf55;LUC7L2 | Other | -1.515174908 | 0.771758719 | 24.47407427 | 24.34082372 | 24.04751736 | 23.69710332 | 23.28192437 | 25.60263544 | 21.23841407 | 23.67325127 | 23.26484917 |
| Putative methyltransferase C9orf114 | Q5T280 | C9orf114 | Other | -1.022651716 | 0.888991229 | 23.70155183 | 22.41720148 | 22.54738504 | 23.66893085 | 23.01055406 | 23.93571037 | 21.6076554 | 21.90260623 | 23.06245003 |
| Protein PAXX | Q9BUH6 | C9orf142 | Extracellular or secreted | -0.340213457 | 0.437020659 | 25.41499743 | 25.14592655 | 24.934766 | 24.84210937 | 24.80020595 | 25.0736552 | 24.35991199 | 25.27756807 | 24.44770982 |
| UPF0553 protein C9orf64 | Q5T6V5 | C9orf64 | Other | -0.489686645 | 0.751235417 | 27.32474448 | 27.46552119 | 27.36744715 | 27.1178314 | 27.31622283 | 25.83282165 | 26.81921568 | 26.7494476 | 26.17457113 |
| Uncharacterized protein C9orf78 | Q9NZ63 | C9orf78 | Other | 3.136811633 | 1.088671681 | 22.00536483 | 20.74751947 | 21.25291947 | 22.0139251 | 21.64754831 | 22.736369 | 25.62460941 | 26.13118203 | 22.85646656 |
| Carbonic anhydrase 13 | Q8N1Q1 | CA13 | Other | -0.721035585 | 1.512412359 | 26.91456214 | 26.94134121 | 27.01414344 | 27.08719639 | 27.29789078 | 26.38980291 | 26.51055255 | 26.26123038 | 25.88757874 |
| Carbonic anhydrase 2 | P00918 | CA2 | Extracellular or secreted | -0.542803256 | 0.575849952 | 32.72026651 | 32.77150061 | 32.74793228 | 32.33657703 | 32.52990027 | 31.34815501 | 32.24513188 | 32.18562281 | 31.16800139 |
| Calcium-binding protein 39 | Q9Y376 | CAB39 | Extracellular or secreted | 0.339533419 | 1.931865431 | 28.69494694 | 28.78612532 | 28.67841875 | 28.69478094 | 28.75146024 | 28.27155215 | 29.04273033 | 29.05211751 | 28.86239459 |
| Voltage-dependent R-type calcium ch | Q15878-2;Q15878-3;Q15878 | CACNA1E | Extracellular or secreted | 3.522434583 | 1.283508996 | 21.84790164 | 22.11824655 | 20.85661886 | 28.1283428 | 27.95748327 | 21.20516803 | 26.83677504 | 27.1915001 | 27.59590918 |
| Cactin | Q8WUQ7;Q8WUQ7-2 | CACTIN | Extracellular or secreted | -0.731053715 | 0.949421504 | 22.63180852 | 23.76243233 | 23.99748549 | 22.58301089 | 22.19132313 | 22.00131585 | 21.95822701 | 21.86661262 | 22.56568733 |
| Calcyclin-binding protein | Q9HB71;Q9HB71-3 | CACYBP | Extracellular or secreted | 0.238454327 | 0.420786525 | 31.68022267 | 31.59543795 | 31.61530888 | 31.74470842 | 31.75998277 | 31.07839178 | 31.95930182 | 32.08527892 | 31.40780848 |
| CAD protein;Glutamine-dependent ca | P27708 | CAD | Extracellular or secreted | 0.893494299 | 0.985904245 | 31.25496116 | 31.41588799 | 31.3677075 | 31.10724286 | 31.18612482 | 31.21361447 | 32.537349 | 32.39412196 | 31.52178133 |
| Cell adhesion molecule 1 | Q9BY67-2;Q9BY67-5;Q9BY67 | CADM1 | Extracellular or secreted | -0.255432401 | 0.481256222 | 25.85031257 | 25.97289918 | 26.01917992 | 25.81570245 | 26.20500986 | 26.18465019 | 25.53851411 | 25.56587141 | 26.15319436 |
| Cell adhesion molecule 4 | Q8NFZ8 | CADM4 | Extracellular or secreted | -0.201491319 | 0.44179211 | 29.45538533 | 29.4544443 | 29.33600766 | 29.61642236 | 29.84571448 | 29.03873333 | 29.05842788 | 29.13437806 | 29.57607383 |
| Calretinin | P22676 | CALB2 | Other | 2.255205636 | 0.452351947 | 21.49765306 | 21.37840705 | 22.55915206 | 21.56699794 | 21.66886595 | 22.5625797 | 22.22016338 | 22.27500482 | 27.88727659 |
| Caldesmon | Q05682-5;Q05682-4;Q05682 | CALD1 | Extracellular or secreted | 1.031789191 | 4.490317215 | 27.90546679 | 27.67148416 | 27.86891296 | 28.24889073 | 28.18952206 | 27.75378884 | 28.89769906 | 28.94458411 | 29.07211719 |
| Calreticulin | P27797 | CALR | Extracellular or secreted | -0.623090281 | 1.989443276 | 34.49595627 | 34.45120482 | 34.38753787 | 34.47586896 | 34.52729226 | 34.63444468 | 33.76228183 | 33.81067314 | 34.04392662 |
| Calumenin | O43852;O43852-3;O43852-6 | CALU | Extracellular or secreted | 0.667639574 | 2.01845605 | 29.61754375 | 29.82776472 | 29.90167446 | 30.42342167 | 30.581633 | 30.27386552 | 30.62479415 | 30.70775483 | 30.98332131 |
| Calcium/calmodulin-dependent prote | Q13557-8;Q13557-12;Q1355 | CAMK2D | Extracellular or secreted | 0.181690909 | 0.29071153 | 26.36327509 | 26.53244784 | 26.47455398 | 26.02000702 | 26.02261248 | 27.3414686 | 26.68190709 | 26.32601411 | 26.91433403 |
| Calcium/calmodulin-dependent prote | Q13555-10;Q13555-3;Q1355 | CAMK2G | Extracellular or secreted | 0.715254201 | 0.40429258 | 24.10980515 | 23.40028221 | 19.72519938 | 22.88043102 | 22.99218669 | 24.80557857 | 23.56651027 | 23.22646864 | 24.3095252 |
| Calmodulin-regulated spectrin-associ | Q9P1Y5;Q9P1Y5-2 | CAMSAP3 | Other | -0.7612088 | 1.805815816 | 24.99494208 | 24.66876858 | 24.63084256 | 24.3415704 | 24.4898362 | 23.79748738 | 23.85443242 | 23.9097642 | 23.41390059 |
| Cullin-associated NEDD8-dissociated | Q86VP6;Q86VP6-2 | CAND1 | Extracellular or secreted | -0.012338823 | 0.016537806 | 32.66966083 | 32.8696779 | 32.81764615 | 32.15906418 | 32.1897518 | 32.42442319 | 32.7714219 | 32.66331186 | 32.0933618 |
| Calnexin | P27824;P27824-2;P27824-3 | CANX | Extracellular or secreted | -0.988249876 | 0.932134071 | 31.68311263 | 31.49961968 | 31.65782931 | 31.76026044 | 31.86215808 | 32.81867492 | 30.50691872 | 30.46548227 | 31.70367691 |
| Adenylyl cyclase-associated protein 1 | Q01518 | CAP1 | Extracellular or secreted | 0.418310746 | 1.381341328 | 32.55362039 | 32.53725642 | 32.58174523 | 32.15569087 | 32.12786331 | 32.48483786 | 32.69842871 | 32.71881818 | 33.05819239 |
| Adenylyl cyclase-associated protein 1 | Q01518-2 | CAP1 | Extracellular or secreted | -0.179341898 | 0.303079056 | 24.27689397 | 23.89476523 | 24.66665744 | 24.41654448 | 25.24081753 | 24.34786787 | 24.02419887 | 24.25889665 | 24.60065204 |
| Macrophage-capping protein | P40121;P40121-2 | CAPG | Extracellular or secreted | -0.286952334 | 0.832080216 | 31.62418407 | 31.55640971 | 31.43166352 | 30.74703192 | 30.80408955 | 30.98712216 | 30.93030002 | 31.01025312 | 30.77384033 |
| Calpain-1 catalytic subunit | P07384 | CAPN1 | Extracellular or secreted | -0.645095383 | 1.128988895 | 30.96113188 | 30.99530069 | 30.95039466 | 30.52421247 | 30.5171884 | 30.98935727 | 29.986851 | 29.94363436 | 30.60302118 |
| Calpain-2 catalytic subunit | P17655;P17655-2 | CAPN2 | Extracellular or secreted | -0.461022825 | 2.953162066 | 30.66898494 | 30.75374103 | 30.71644875 | 30.36348398 | 30.46324244 | 30.3292432 | 30.02300396 | 30.14879929 | 30.09270045 |
| Calpain-5 | O15484 | CAPN5 | Extracellular or secreted | -1.076536408 | 1.585114068 | 22.75225813 | 22.51537645 | 22.60374495 | 21.68177162 | 22.87800001 | 21.22079636 | 20.9041729 | 21.6936806 | 20.99851105 |
| Calpain-7 | Q9Y6W3 | CAPN7 | Extracellular or secreted | 1.636772244 | 0.57417255 | 20.44891533 | 18.70596725 | 20.12929227 | 23.12245203 | 22.23053017 | 23.04705134 | 20.84016021 | 24.07005541 | 23.8422053 |
| Calpain small subunit 1 | P04632 | CAPNS1 | Extracellular or secreted | -0.579009251 | 2.078173536 | 30.35026452 | 30.23626213 | 30.26840982 | 29.70915267 | 29.72622811 | 29.95393638 | 29.30088236 | 29.64727322 | 29.43694348 |
| Caprin-1 | Q14444-2;Q14444 | CAPRIN1 | Extracellular or secreted | 0.052018182 | 0.299749588 | 30.48644516 | 30.27308739 | 30.31246249 | 30.59076175 | 30.67463579 | 30.35930044 | 30.45336528 | 30.56229073 | 30.48874504 |
| Calcyphosin | Q13938-4;Q13938;Q13938-3 | CAPS | Extracellular or secreted | 0.763338012 | 2.223660464 | 30.39140999 | 30.57551492 | 30.48654107 | 29.54589193 | 29.73713512 | 30.17830917 | 30.93740361 | 30.83288803 | 30.97712348 |
| F-actin-capping protein subunit alpha | P52907 | CAPZA1 | Extracellular or secreted | -0.159965477 | 0.541729248 | 30.18695128 | 30.13613971 | 30.14029065 | 29.94586806 | 29.92479004 | 29.90784627 | 29.96884096 | 30.01358451 | 29.6586211 |
| F-actin-capping protein subunit alpha | P47755;P47755-2 | CAPZA2 | Extracellular or secreted | -0.249554201 | 1.101500325 | 30.63200829 | 30.74759221 | 30.65880835 | 30.29142792 | 30.26450023 | 30.22020398 | 30.26214938 | 30.32023169 | 30.07622681 |
| F-actin-capping protein subunit beta | P47756-2;P47756 | CAPZB | Extracellular or secreted | -0.225368307 | 1.415807048 | 31.42297075 | 31.49667065 | 31.39033402 | 31.16901756 | 31.15061605 | 31.06464655 | 30.95525363 | 31.11916808 | 31.09660116 |
| Calcium-regulated heat stable protein | Q9Y2V2 | CARHSP1 | Extracellular or secreted | 0.1358573 | 0.755112257 | 29.40186139 | 29.31471122 | 29.32594551 | 29.13239363 | 29.19044078 | 28.82941403 | 29.38687524 | 29.325774 | 29.29230594 |
| ATP-dependent (S)-NAD(P)H-hydrate | Q8IW45;Q8IW45-4;Q8IW45-1 | CARKD | Other | -1.056809715 | 0.346298088 | 24.71790355 | 25.16951463 | 24.83272509 | 22.29170948 | 21.48367988 | 24.69508638 | 24.12998414 | 23.60602922 | 20.688867 |
| Histone-arginine methyltransferase C | Q86X55-1;Q86X55;Q86X55-2 | CARM1 | Extracellular or secreted | -0.290328276 | 0.961928675 | 27.11137816 | 26.99258867 | 26.86017577 | 27.5515684 | 27.35682644 | 27.29998772 | 26.83833894 | 27.12060239 | 26.75633642 |
| Cysteine--tRNA ligase, cytoplasmic | P49589-3;P49589;P49589-2 | CARS | Other | -0.02449172 | 0.059659678 | 28.97708253 | 29.03603628 | 29.04411233 | 28.85135555 | 29.04567527 | 28.5574235 | 29.02199861 | 29.00254817 | 28.65782079 |
| Protein CASC3 | O15234 | CASC3 | Other | 0.472349854 | 0.229383626 | 23.48480618 | 21.93329877 | 20.96958326 | 23.11462626 | 21.11413375 | 23.22275953 | 22.27491448 | 21.96466037 | 24.09707859 |
| Peripheral plasma membrane protein | O14936-3;O14936-4;O14936 | CASK | Extracellular or secreted | -0.125777141 | 0.508266887 | 28.58305085 | 28.53096076 | 28.54130137 | 28.10394607 | 28.22949362 | 28.82429351 | 28.36114269 | 28.26472392 | 28.40332506 |
| Caspase-14;Caspase-14 subunit p17, r | P31944 | CASP14 | Extracellular or secreted | -4.221620684 | 1.384865652 | 26.97311821 | 26.81213166 | 27.03170524 | 26.38368623 | 26.26760196 | 26.04570651 | 23.5995742 | 22.94440538 | 20.54813328 |
| Caspase-2;Caspase-2 subunit p18;Cas | P42575;P42575-2 | CASP2 | Other | 0.595989757 | 0.442730714 | 23.71732834 | 23.18152961 | 23.53599578 | 24.11886125 | 24.07887527 | 23.63234118 | 24.90531759 | 24.71575843 | 23.29935896 |
| Caspase-3;Caspase-3 subunit p17;Cas | P42574 | CASP3 | Extracellular or secreted | 0.762495199 | 1.487501956 | 26.0021511 | 26.35894008 | 26.13789922 | 25.99552444 | 26.03649225 | 25.78309276 | 26.81616649 | 27.11961337 | 26.50875567 |
| Caspase-6;Caspase-6 subunit p18;Cas | P55212 | CASP6 | Other | 0.11770901 | 0.174020297 | 25.59057974 | 25.60339974 | 25.86396051 | 24.7915878 | 25.26223452 | 24.65337867 | 25.51126163 | 25.65102584 | 25.07341004 |
| Caspase-7;Caspase-7 subunit p20;Cas | P55210-4;P55210;P55210-3 | CASP7 | Other | -1.926137988 | 1.063860557 | 24.13226124 | 24.79565564 | 24.53931314 | 24.12801824 | 24.23527627 | 24.4633243 | 21.46451133 | 22.30000521 | 23.60399391 |
| Caspase-8;Caspase-8 subunit p18;Cas | Q14790-2;Q14790;Q14790-4 | CASP8 | Extracellular or secreted | 1.224550898 | 0.319015084 | 24.58520188 | 23.59354614 | 22.56362735 | 25.07687919 | 22.34429591 | 22.03066175 | 26.03795877 | 25.93132058 | 21.80147945 |
| Calpastatin | P20810-5;P20810-2;P20810-3 | CAST | Other | -0.225965743 | 0.319343203 | 30.82168082 | 30.79436088 | 30.82941403 | 31.01896511 | 31.09710371 | 29.4034876 | 30.58190232 | 30.6084953 | 30.11421121 |
| Catalase | P04040 | CAT | Extracellular or secreted | -1.298324541 | 1.674263674 | 31.04739255 | 31.13356992 | 31.08106167 | 30.9549071 | 31.02966647 | 31.12466266 | 29.6367171 | 29.49933455 | 30.15460491 |
| Caveolin-1 | Q03135;Q03135-2 | CAV1 | Extracellular or secreted | 0.53314674 | 0.320150794 | 24.94730697 | 24.41956934 | 24.61386094 | 25.14670353 | 25.36739715 | 27.02178687 | 25.29210843 | 25.15313636 | 26.91250784 |
| Core-binding factor subunit beta | Q13951-2;Q13951 | CBFB | Other | 0.584631556 | 1.039547328 | 27.04614382 | 26.78979828 | 26.84337997 | 26.66356636 | 26.74688782 | 26.36799696 | 27.44624318 | 27.61946909 | 26.917069 |
| E3 ubiquitin-protein ligase CBL | P22681 | CBL | Extracellular or secreted | 0.037382045 | 0.021449095 | 23.54108716 | 23.43122315 | 22.94981104 | 21.80315324 | 20.38423622 | 21.35687352 | 22.56420905 | 22.59375128 | 21.68737798 |
| Carbonyl reductase [NADPH] 1 | P16152;P16152-2 | CBR1 | Extracellular or secreted | -0.049258808 | 0.085377615 | 31.06213913 | 30.96037252 | 30.92994744 | 30.47121208 | 30.45689361 | 30.81145809 | 30.87884022 | 30.93177989 | 30.3876149 |
| Carbonyl reductase [NADPH] 3 | O75828 | CBR3 | Extracellular or secreted | 0.274262859 | 1.012314335 | 28.35009587 | 28.41453782 | 28.39582841 | 27.86319254 | 27.7796907 | 28.02780934 | 28.40904353 | 28.55928483 | 28.27003756 |
| Carbonyl reductase family member 4 | Q8N4T8;Q8N4T8-2 | CBR4 | Other | -1.011867173 | 3.506695701 | 23.31174388 | 23.88576488 | 23.66903902 | 23.41828294 | 23.60794882 | 23.03489835 | 22.29935896 | 22.53535512 | 22.59352334 |
| COBW domain-containing protein 1;C | Q9BRT8-4;Q5RIA9-3;Q5JTY5- | CBWD1;CBWD5;C | Other | 0.923936556 | 0.556149039 | 22.51881353 | 22.50867105 | 22.16210977 | 20.77138056 | 21.54904528 | 20.79352393 | 23.60320163 | 22.85214771 | 21.46823239 |
| Chromobox protein homolog 1 | P83916 | CBX1 | Other | -0.121961007 | 0.321328212 | 28.96636753 | 28.78870982 | 28.99781925 | 28.98261433 | 29.3174741 | 28.54638882 | 29.04406021 | 28.64519566 | 28.74454804 |
| Chromobox protein homolog 3 | Q13185 | CBX3 | Other | 0.088673331 | 0.194218999 | 31.12786331 | 31.28890063 | 31.22377515 | 31.70318183 | 31.58279969 | 30.81145809 | 31.61429986 | 31.32071588 | 31.19999361 |
| Chromobox protein homolog 5 | P45973 | CBX5 | Other | 0.109066791 | 0.872287193 | 29.20955332 | 29.22779652 | 29.29673179 | 29.01494152 | 29.15139051 | 29.28197741 | 29.32257581 | 29.22087281 | 29.3749473 |
| Coiled-coil and C2 domain-containing | Q6P1N0;Q6P1N0-2 | CC2D1A | Extracellular or secreted | -1.51610576 | 2.546119866 | 26.1299645 | 26.55057066 | 26.31943081 | 26.21207954 | 25.96527788 | 24.6837284 | 24.71570607 | 24.5944007 | 24.07210185 |
| Coiled-coil and C2 domain-containing | Q5T0F9-3;Q5T0F9-2;Q5T0F9 | CC2D1B | Other | 0.700370271 | 0.385416006 | 21.61099237 | 22.91457127 | 22.83126626 | 21.37937531 | 21.33225241 | 19.8459494 | 22.43941394 | 21.24266474 | 23.37623564 |
| Cell division cycle and apoptosis regul | Q8IX12-2;Q8IX12 | CCAR1 | Other | -0.253109086 | 0.799012188 | 28.87173532 | 29.02366498 | 29.04111196 | 28.5831226 | 28.61782396 | 28.88878672 | 28.74970333 | 28.655093 | 28.34899918 |
| Cell cycle and apoptosis regulator pro | Q8N163;Q8N163-2 | CCAR2 | Other | -0.773287398 | 2.148407385 | 31.14795068 | 31.13032053 | 31.13907103 | 30.31904744 | 30.34159585 | 31.32017788 | 30.14588769 | 30.02656992 | 30.2067619 |
| Kynurenine--oxoglutarate transamina | Q16773;Q16773-2 | CCBL1 | Other | 0.079579982 | 0.061779587 | 24.46842965 | 24.59951745 | 23.77936509 | 24.76819967 | 24.80094649 | 24.49996556 | 24.29656114 | 24.03917275 | 25.36121801 |
| Kynurenine--oxoglutarate transamina | Q6YP21-3;Q6YP21 | CCBL2 | Other | 0.035186323 | 0.105078549 | 27.58663412 | 27.49297109 | 27.39941862 | 27.54074731 | 27.63360269 | 27.10689356 | 27.34291007 | 27.47630123 | 27.66648137 |
| Coiled-coil domain-containing protein | Q8WUD4 | CCDC12 | Other | 0.619342246 | 1.203608365 | 25.68139778 | 25.8089202 | 26.06264354 | 26.26774482 | 25.35292654 | 24.57091651 | 26.21258054 | 26.48166888 | 26.03605201 |
| Coiled-coil domain-containing protein | Q96CT7 | CCDC124 | Extracellular or secreted | 2.338844343 | 3.483357644 | 25.98520127 | 25.5842845 | 25.90712998 | 26.98174373 | 27.00206524 | 27.25439807 | 28.91669894 | 28.77159112 | 28.68565435 |
| Coiled-coil domain-containing protein | Q96JG6-3;Q96JG6 | CCDC132 | Extracellular or secreted | -0.921645075 | 1.004224132 | 24.15645778 | 24.53889888 | 24.65567309 | 24.29530035 | 24.03473046 | 23.95633149 | 23.86759397 | 23.4693616 | 22.71680523 |
| Coiled-coil domain-containing protein | P0C221 | CCDC175 | Other | -0.134000024 | 0.387610523 | 25.25814187 | 25.16855858 | 25.47016881 | 24.85071793 | 24.71324302 | 25.46444654 | 25.16798465 | 25.01197041 | 24.88068324 |
| Coiled-coil domain-containing protein | O60826 | CCDC22 | Other | 0.084195548 | 0.227368773 | 25.83771358 | 25.25068066 | 25.42191406 | 25.77740984 | 25.10545554 | 25.85145683 | 25.71127479 | 25.67688659 | 25.48674052 |
| Coiled-coil domain-containing protein | Q86WR0;Q86WR0-2 | CCDC25 | Extracellular or secreted | 0.005663437 | 0.024572713 | 26.6292312 | 26.73414956 | 26.64924499 | 26.48408398 | 26.62352141 | 26.45586737 | 26.70332212 | 26.48406861 | 26.61764883 |
| Coiled-coil domain-containing protein | Q8N5R6;Q8N5R6-2;Q8N5R6- | CCDC33 | Other | -3.730315071 | 0.639910769 | 27.07541092 | 27.06208765 | 26.87946846 | 27.30129677 | 27.74624716 | 22.18881287 | 19.65852182 | 21.27523698 | 27.0019579 |
| Coiled-coil domain-containing protein | Q96MW1;Q96MW1-2 | CCDC43 | Other | 0.546137317 | 0.639213182 | 25.56569713 | 25.21741463 | 25.5134922 | 24.99416523 | 24.39865237 | 23.07841927 | 25.62151065 | 25.22809939 | 25.17272232 |
| Coiled-coil domain-containing protein | Q8IVM0;Q8IVM0-2 | CCDC50 | Other | 0.726123287 | 0.325853473 | 20.92407471 | 22.65138181 | 22.66978903 | 20.62076229 | 22.32479931 | 22.32938573 | 23.51364279 | 23.37570553 | 21.04911798 |
| WASH complex subunit CCDC53 | Q9Y3C0 | CCDC53 | Other | 0.279134371 | 0.692255619 | 23.70767256 | 24.26492074 | 24.6140295 | 24.69349206 | 24.69980584 | 24.16951463 | 24.82313357 | 24.67076861 | 24.41821859 |
| Coiled-coil domain-containing protein | Q4VC31 | CCDC58 | Other | -0.425518588 | 0.782693016 | 28.94332695 | 28.88567762 | 28.89217874 | 29.07405859 | 28.80888338 | 27.62387022 | 28.49621348 | 28.38319182 | 27.95803668 |
| Coiled-coil domain-containing protein | Q16204 | CCDC6 | Other | -0.086128594 | 0.15921015 | 26.4194086 | 26.41660891 | 26.48055997 | 27.13643311 | 27.01987981 | 25.97517555 | 26.68297873 | 26.49147331 | 26.29119515 |
| Coiled-coil domain-containing protein | Q8NA47-2;Q8NA47 | CCDC63 | Other | -4.31106421 | 2.883208734 | 25.9464148 | 26.31414937 | 26.34068791 | 25.79919327 | 25.47060327 | 26.41100934 | 21.1321357 | 21.85327148 | 22.22242917 |
| Coiled-coil domain-containing protein | Q9H6F5 | CCDC86 | Other | -0.958146099 | 0.674260205 | 27.52238015 | 27.33994077 | 27.5140493 | 27.94503083 | 28.1711671 | 27.2201117 | 27.21187538 | 27.18763569 | 25.58239056 |
| Girdin | Q3V6T2-2;Q3V6T2;Q3V6T2-5 | CCDC88A | Extracellular or secreted | -0.121934032 | 0.323780854 | 25.82398388 | 26.12078035 | 26.19146245 | 25.61669578 | 25.82257453 | 26.41207606 | 26.08754037 | 25.79168715 | 25.74875691 |
| Coiled-coil domain-containing protein | Q9Y3X0 | CCDC9 | Other | 0.16843869 | 0.170284723 | 25.72234061 | 26.0766142 | 26.06249944 | 25.0898652 | 25.1492646 | 23.63854109 | 25.39972827 | 25.36926239 | 25.60588797 |
| Coiled-coil domain-containing protein | Q567U6 | CCDC93 | Other | 1.371631017 | 0.971459515 | 25.41163977 | 25.06858018 | 21.09217898 | 24.01137332 | 24.196105 | 22.30513995 | 25.22464472 | 25.38972087 | 24.54303607 |
| Coiled-coil domain-containing protein | Q96F63 | CCDC97 | Other | 0.248269626 | 0.266816219 | 25.3405521 | 25.69858768 | 25.27976557 | 25.73820196 | 25.41580339 | 25.01937083 | 26.16273214 | 25.79743791 | 25.0307796 |
| G2/mitotic-specific cyclin-B1 | P14635;P14635-2 | CCNB1 | Other | 0.343813541 | 0.092166593 | 23.33612443 | 21.43108484 | 22.40794985 | 23.71570138 | 21.77836009 | 22.48795042 | 24.07103806 | 24.1091675 | 20.42982057 |
| Cyclin-C | P24863;P24863-2 | CCNC | Other | 0.819093587 | 0.542960048 | 23.29656114 | 23.59274809 | 20.66562197 | 22.08318949 | 22.59951951 | 23.4178968 | 24.19069038 | 22.44119892 | 23.65315997 |
| G1/S-specific cyclin-E1 | P24864-3;P24864;P24864-2 | CCNE1 | Other | 0.119818053 | 0.235419767 | 24.40872778 | 24.03825189 | 24.35029825 | 23.54557161 | 23.34989347 | 24.15189852 | 23.9035259 | 24.27795819 | 24.10029082 |
| Cyclin-K | O75909;O75909-4;O75909-1 | CCNK | Other | 0.022043505 | 0.031673575 | 25.8317833 | 26.26361407 | 26.34844208 | 25.69088422 | 25.40460862 | 26.41989076 | 26.25616318 | 26.10253514 | 25.68704372 |
| Cyclin-L1 | Q9UK58-5;Q9UK58-4;Q9UK5 | CCNL1 | Extracellular or secreted | -1.573288505 | 1.40785816 | 20.12804972 | 21.93998374 | 22.01719717 | 21.06415375 | 20.36570303 | 23.70914915 | 19.93963127 | 19.4429354 | 20.5096861 |
| Copper chaperone for superoxide dis | O14618 | CCS | Other | 0.40685534 | 0.750631777 | 27.60448887 | 27.69842871 | 27.52701255 | 27.3178622 | 27.29920173 | 27.129375 | 28.03212579 | 28.05568988 | 27.42093488 |
| T-complex protein 1 subunit beta | P78371;P78371-2 | CCT2 | Extracellular or secreted | 0.088505102 | 0.087920461 | 32.46158467 | 32.64626486 | 32.55741437 | 32.34517006 | 32.35160772 | 31.67118043 | 32.73862156 | 32.75224616 | 31.79125864 |
| T-complex protein 1 subunit gamma | P49368;P49368-2 | CCT3 | Extracellular or secreted | 0.039108992 | 0.044042926 | 32.56547198 | 32.49945336 | 32.46822862 | 32.38378407 | 32.42829754 | 31.85504463 | 32.63951001 | 32.71798115 | 31.85997591 |
| T-complex protein 1 subunit delta | P50991;P50991-2 | CCT4 | Extracellular or secreted | 0.30681002 | 0.359686885 | 31.99755682 | 32.17819042 | 32.15668561 | 31.60730512 | 31.67753695 | 31.59930106 | 32.49731329 | 32.4835171 | 31.54788767 |
| T-complex protein 1 subunit epsilon | P48643;P48643-2 | CCT5 | Extracellular or secreted | 0.275599759 | 0.45698455 | 32.54568023 | 32.61267518 | 32.58633884 | 32.44808117 | 32.43081653 | 32.04934152 | 32.96292517 | 32.93190314 | 32.26843771 |
| T-complex protein 1 subunit zeta | P40227;P40227-2 | CCT6A | Extracellular or secreted | 0.0449508 | 0.038830738 | 31.85180976 | 31.98328733 | 31.91056497 | 31.40659264 | 31.51704757 | 31.0123861 | 32.08964172 | 31.92326755 | 30.9627873 |
| T-complex protein 1 subunit zeta-2 | Q92526;Q92526-2;Q92526-3 | CCT6B | Other | 0.837552716 | 0.117784355 | 21.94734611 | 27.78000373 | 27.71082829 | 26.80267294 | 20.56026049 | 22.08779599 | 20.96757851 | 27.90213467 | 27.08739874 |
| T-complex protein 1 subunit eta | Q99832;Q99832-3;Q99832-4 | CCT7 | Extracellular or secreted | 0.15461774 | 0.188974207 | 31.92606344 | 31.88369097 | 31.82982984 | 31.54231661 | 31.54462131 | 31.13613971 | 31.99546918 | 32.14858718 | 31.2511278 |
| T-complex protein 1 subunit theta | P50990;P50990-2;P50990-3 | CCT8 | Extracellular or secreted | 0.194655019 | 0.329566124 | 32.66991421 | 32.74979123 | 32.73204385 | 32.75294396 | 32.79985098 | 32.3134089 | 33.1172648 | 33.0413013 | 32.43437552 |
| Vacuolar fusion protein CCZ1 homolo | P86791;P86790 | CCZ1;CCZ1B | Other | -0.172618029 | 0.22961122 | 24.8891582 | 25.31094694 | 25.12746731 | 25.33234556 | 25.37785789 | 26.40101503 | 25.57380804 | 24.83667874 | 25.2910546 |
| CD109 antigen | Q6YHK3;Q6YHK3-4;Q6YHK3-2 | CD109 | Extracellular or secreted | 0.774248485 | 0.937071614 | 29.09325474 | 29.18553421 | 29.24619713 | 29.10814068 | 29.07706771 | 30.60178221 | 29.86565746 | 29.82624994 | 30.7868264 |
| CD151 antigen | P48509 | CD151 | Extracellular or secreted | 2.668906012 | 1.893413827 | 25.66484155 | 26.35863832 | 26.28875106 | 24.29663115 | 21.91109725 | 26.22166116 | 27.82438457 | 28.04437294 | 27.50877078 |
| CD276 antigen | Q5ZPR3-4;Q5ZPR3;Q5ZPR3-3 | CD276 | Extracellular or secreted | 0.442822305 | 1.610687167 | 26.53237352 | 26.75021465 | 26.77026827 | 26.00705842 | 26.17554224 | 26.07928229 | 26.77329015 | 26.76483819 | 26.94770827 |
| CD2-associated protein | Q9Y5K6 | CD2AP | Extracellular or secreted | 0.135353512 | 1.650089223 | 28.14627624 | 28.11674134 | 28.07694034 | 28.22719977 | 28.03857635 | 28.01860703 | 28.17902143 | 28.27914565 | 28.26006399 |
| CD2 antigen cytoplasmic tail-binding | O95400 | CD2BP2 | Other | 0.680388804 | 2.034030642 | 24.85856424 | 24.92944805 | 25.19745549 | 24.90260623 | 25.54554216 | 24.32765954 | 25.72192358 | 25.44805655 | 25.75182413 |
| DNA-directed RNA polymerase I subu | O15446;O15446-2 | CD3EAP | Other | -0.552429545 | 1.699616299 | 27.30990678 | 27.17216912 | 27.21752556 | 27.99318278 | 27.86224678 | 26.98228792 | 26.84361958 | 26.98793533 | 26.77981592 |
| CD44 antigen | P16070-8;P16070-4;P16070-3 | CD44 | Extracellular or secreted | 0.895174303 | 1.566861398 | 30.27763901 | 30.11408707 | 30.2570989 | 30.59405893 | 30.84146165 | 30.62592646 | 31.0235328 | 31.28862566 | 31.72850046 |
| Membrane cofactor protein | P15529-7;P15529-4;P15529-6 | CD46 | Extracellular or secreted | 1.190073801 | 2.496126051 | 26.17491395 | 26.33604598 | 26.55940883 | 26.01448401 | 25.88939033 | 26.70938277 | 27.30190725 | 27.3106003 | 27.79952679 |
| Leukocyte surface antigen CD47 | Q08722-2;Q08722-3;Q08722 | CD47 | Extracellular or secreted | -0.550704284 | 0.481995156 | 27.56456377 | 27.78400444 | 27.83213353 | 27.44181846 | 27.11376426 | 27.16585911 | 26.92569568 | 27.68498591 | 26.18827734 |
| Complement decay-accelerating facto | P08174-4;P08174;P08174-3;P | CD55 | Extracellular or secreted | 1.182189623 | 2.269682514 | 24.66876858 | 24.34577177 | 24.21356349 | 25.63361655 | 24.7573542 | 25.37693111 | 25.79847655 | 25.87330874 | 26.37278643 |
| CD59 glycoprotein | P13987 | CD59 | Extracellular or secreted | 0.827215159 | 2.946504543 | 28.57036647 | 27.80943549 | 28.13232008 | 28.25399251 | 28.70984286 | 28.52133206 | 29.099664 | 29.07262834 | 29.30799788 |
| CD63 antigen | P08962-3;P08962-2;P08962 | CD63 | Extracellular or secreted | 2.18180809 | 1.648285091 | 25.32930309 | 22.9982928 | 22.98070261 | 24.54439285 | 24.85086097 | 27.30903941 | 26.38595835 | 26.65676439 | 27.5089974 |
| CD81 antigen | P60033 | CD81 | Extracellular or secreted | 0.544417782 | 0.735714496 | 27.35808493 | 27.77045732 | 27.69178952 | 28.06110919 | 27.9891813 | 28.7138133 | 28.1654758 | 28.22527011 | 29.03472522 |
| CD82 antigen | P27701-2;P27701 | CD82 | Extracellular or secreted | 1.294018218 | 0.573503706 | 26.31629189 | 26.18449887 | 26.16614653 | 22.07454032 | 25.17077564 | 28.19821458 | 26.22982364 | 26.38147655 | 28.32598839 |

| | | | | | | | | | | | | | |
|---|---|---|---|---|---|---|---|---|---|---|---|---|---|
| CD9 antigen | P21926 | CD9 | Extracellular or secreted | 0.685480877 | 0.943382616 | 29.99577241 | 30.03590523 | 30.20174707 | 30.16506842 | 30.16877853 | 30.87195558 | 30.45826341 | 30.89843447 | 31.41935837 |
| CD97 antigen;CD97 antigen subunit al | P48960-2;P48960-3;P48960 | CD97 | Extracellular or secreted | 0.287325518 | 1.198258422 | 26.47438378 | 26.30858817 | 26.53724068 | 26.72663415 | 27.01169322 | 26.84397892 | 26.83749705 | 26.88925106 | 27.08648792 |
| Cell division cycle protein 123 homolo | O75794 | CDC123 | Other | 0.900692352 | 0.882719252 | 23.08632998 | 24.94547741 | 24.32759102 | 26.02466388 | 25.61540536 | 24.66968786 | 26.29570322 | 25.44915923 | 25.29179236 |
| Cell division cycle protein 16 homolog | Q13042-3;Q13042-2;Q13042 | CDC16 | Other | -0.603422924 | 0.689617842 | 24.59445765 | 23.59548241 | 23.82318218 | 22.36922245 | 21.92036791 | 23.75144109 | 22.63809912 | 22.64717645 | 22.9315325 |
| Cell division cycle protein 23 homolog | Q9UJX2;Q9UJX2-3 | CDC23 | Extracellular or secreted | -1.229300271 | 1.036536853 | 24.44354255 | 24.82211254 | 24.77183037 | 24.35555005 | 24.23169229 | 26.34783409 | 23.32003367 | 22.98337568 | 24.49497079 |
| Cell division cycle protein 27 homolog | P30260;P30260-2 | CDC27 | Other | -0.798900577 | 1.601410223 | 23.59343216 | 24.42668807 | 24.14546016 | 23.37967661 | 23.48566621 | 23.49044842 | 22.58927716 | 23.04640198 | 23.22830494 |
| Ubiquitin-conjugating enzyme E2 R1 | P49427 | CDC34 | Other | 0.474680366 | 0.698135354 | 23.68241279 | 21.9169597 | 22.21377112 | 22.54519451 | 23.22178652 | 22.25290137 | 23.2403226 | 23.36145229 | 22.73877921 |
| Hsp90 co-chaperone Cdc37;Hsp90 co- | Q16543 | CDC37 | Extracellular or secreted | 0.085755965 | 0.177417886 | 30.52477292 | 30.41538435 | 30.39907223 | 30.78729359 | 30.9616839 | 30.3372419 | 30.79443835 | 30.82297257 | 30.35258143 |
| Pre-mRNA-processing factor 17 | O60508 | CDC40 | Extracellular or secreted | -0.919550003 | 0.934553302 | 26.27861407 | 26.41385218 | 26.46877144 | 25.38557991 | 24.69290704 | 25.77399435 | 25.47029296 | 25.07626759 | 24.20164894 |
| Cell division control protein 42 homol | P60953;P60953-1 | CDC42 | Extracellular or secreted | 0.533300509 | 1.937667791 | 29.12691617 | 29.0807822 | 29.0762523 | 28.96776966 | 29.05403423 | 29.66061144 | 29.59981537 | 29.58274586 | 29.9005233 |
| Serine/threonine-protein kinase MRC | Q5VT25-6;Q5VT25-3;Q5VT25 | CDC42BPA | Extracellular or secreted | -2.239804007 | 0.434095182 | 27.50096852 | 27.58204594 | 27.47049467 | 26.17245528 | 26.02544561 | 26.3146853 | 26.56270196 | 20.71838179 | 26.53255188 |
| Serine/threonine-protein kinase MRC | Q9Y5S2 | CDC42BPB | Extracellular or secreted | 0.016930133 | 0.038085674 | 27.52783288 | 27.59526898 | 27.64519566 | 27.08172202 | 27.01754552 | 27.5202832 | 27.20955332 | 27.47460039 | 27.56056083 |
| Cdc42 effector protein 4 | Q9H3Q1;Q9H3Q1-2 | CDC42EP4 | Extracellular or secreted | 0.071498188 | 0.123801799 | 25.15332967 | 25.02757735 | 24.69779267 | 25.18737141 | 25.2180062 | 24.19970349 | 24.86702832 | 25.26992167 | 24.81943498 |
| CDC42 small effector protein 1 | Q9NRR8 | CDC42SE1 | Extracellular or secreted | 1.322598274 | 0.653381886 | 21.1441595 | 21.4822717 | 20.61995989 | 22.04922721 | 21.30114068 | 21.62398214 | 23.87088919 | 22.9556045 | 21.25167169 |
| Cell division cycle 5-like protein | Q99459 | CDC5L | Other | 0.369957665 | 0.957156592 | 25.27657455 | 25.26660156 | 25.68171947 | 25.64617128 | 25.64270551 | 26.54284425 | 26.05125674 | 25.97145269 | 26.11547187 |
| Parafibromin | Q6P1J9 | CDC73 | Other | -0.968536411 | 1.68354929 | 26.90403148 | 26.9359576 | 26.93629466 | 26.39858714 | 26.09526856 | 26.63436461 | 25.90589137 | 25.88462423 | 25.25612718 |
| Cell division cycle-associated protein 3 | Q99618 | CDCA3 | Other | 1.343881423 | 2.129133388 | 21.64223104 | 20.64335814 | 21.4229889 | 21.56860851 | 22.48226512 | 20.53815362 | 22.31329493 | 22.89026279 | 22.97688921 |
| CUB domain-containing protein 1 | Q9H5V8;Q9H5V8-2 | CDCP1 | Extracellular or secreted | 1.066624604 | 1.601542082 | 27.68832502 | 27.75347008 | 28.00034685 | 28.13089737 | 28.27848115 | 28.2998131 | 28.91812172 | 28.81613597 | 29.54128291 |
| Cadherin-1;E-Cad/CTF1;E-Cad/CTF2;E | P12830;P12830-2 | CDH1 | Extracellular or secreted | 0.603519712 | 1.004817159 | 31.59312401 | 31.5719039 | 31.58866367 | 31.56074302 | 31.65121407 | 32.15698691 | 31.99701836 | 32.14004681 | 32.73481175 |
| Cadherin-3 | P22223;P22223-2 | CDH3 | Extracellular or secreted | 1.093761685 | 0.878442603 | 27.71809959 | 27.65259938 | 27.62909221 | 28.32070512 | 28.43714197 | 29.10479596 | 28.74647142 | 28.77048883 | 30.19554192 |
| Cyclin-dependent kinase 1 | P06493;P06493-2 | CDK1 | Extracellular or secreted | 0.474017877 | 0.817615209 | 27.45199082 | 28.00672679 | 28.26593125 | 28.09491634 | 28.28471539 | 28.1921118 | 28.71917734 | 28.77316437 | 28.07790813 |
| Cyclin-dependent kinase 11B;Cyclin-d | P21127-6;P21127-9;P21127-8 | CDK11B;CDK11A | Other | -0.193210579 | 0.927060966 | 25.58244799 | 25.64746194 | 25.82048225 | 25.82757395 | 26.12577384 | 26.15365825 | 25.7170145 | 25.71064439 | 25.57140848 |
| Cyclin-dependent kinase 16 | Q00536;Q00536-3;Q00536-2 | CDK16 | Extracellular or secreted | -6.677168444 | 3.158446151 | 27.85061064 | 27.79507345 | 27.82747703 | 27.6528045 | 27.31639548 | 27.48807462 | 20.5408709 | 21.49015613 | 20.90268551 |
| Cyclin-dependent kinase 2 | P24941;P24941-2 | CDK2 | Other | -0.130232724 | 0.259986068 | 26.72624436 | 26.72051524 | 26.71423923 | 26.22289585 | 26.15452765 | 26.53161526 | 26.64650092 | 26.42487965 | 26.07294005 |
| Cyclin-dependent kinase 2-associated | O14519-2;O14519 | CDK2AP1 | Other | 0.647286672 | 0.444083391 | 21.00963183 | 21.3862709 | 21.55889525 | 21.77140726 | 21.82314329 | 21.89559747 | 21.6658338 | 21.67874111 | 23.31975811 |
| Cyclin-dependent kinase 4 | P11802;P11802-2 | CDK4 | Other | 0.644186413 | 0.771835442 | 26.54161894 | 26.43409013 | 26.34138377 | 26.90437608 | 26.74278269 | 26.93438358 | 27.49350563 | 27.7135839 | 26.6747873 |
| Cyclin-dependent-like kinase 5 | Q00535;Q00535-2 | CDK5 | Extracellular or secreted | 0.238553833 | 0.50950641 | 23.83518534 | 24.07447208 | 24.15406404 | 23.7020805 | 23.80931282 | 24.90187007 | 24.41628676 | 24.4665017 | 24.07136547 |
| CDK5 regulatory subunit-associated p | Q96JB5;Q96JB5-4;Q96JB5-2 | CDK5RAP3 | Other | 0.204800404 | 0.984051951 | 27.01211965 | 26.8606494 | 27.09285164 | 26.66166503 | 26.89307614 | 26.76230559 | 27.24272652 | 27.01147997 | 27.00152846 |
| Cyclin-dependent kinase 6 | Q00534 | CDK6 | Other | 0.819471219 | 0.875516437 | 26.81921568 | 27.07938403 | 26.72793271 | 27.55288787 | 27.76065701 | 26.46475811 | 28.26373942 | 28.23228719 | 27.16480475 |
| Cyclin-dependent kinase 9 | P50750;P50750-2 | CDK9 | Other | 0.05695192 | 0.063927407 | 21.43675215 | 22.636017 | 21.64353014 | 21.40209927 | 22.62843041 | 23.19356567 | 22.25557265 | 22.28280112 | 22.1026793 |
| Cyclin-dependent kinase inhibitor 2A | P42771;P42771-4;P42771-2 | CDKN2A | Extracellular or secreted | 1.222276031 | 1.107487812 | 26.99258867 | 27.30556471 | 27.37260379 | 28.05185829 | 28.20671533 | 27.2216796 | 29.3021906 | 29.00694075 | 27.93320195 |
| CDKN2A-interacting protein | Q9NXV6 | CDKN2AIP | Other | -0.978302721 | 2.843560339 | 24.5234873 | 24.60727161 | 25.29642111 | 24.39884805 | 24.06043933 | 24.79109093 | 23.68955187 | 23.47035502 | 23.74396411 |
| CDKN2AIP N-terminal-like protein | Q96HQ2;Q96HQ2-2 | CDKN2AIPNL | Other | -0.29409217 | 0.847115462 | 23.70935647 | 23.16117187 | 23.58267768 | 24.15784588 | 24.10773174 | 23.65981563 | 23.40820948 | 23.28602073 | 23.61279292 |
| Protein CDV3 homolog | Q9UKY7;Q9UKY7-2 | CDV3 | Extracellular or secreted | 0.358729337 | 0.455933884 | 29.34460407 | 29.67741093 | 29.6623953 | 30.58629409 | 30.4666495 | 28.72133042 | 30.37191869 | 30.25653664 | 29.67707484 |
| Carcinoembryonic antigen-related cel | P13688-2;P13688-8;P13688-1 | CEACAM1 | Extracellular or secreted | 1.115792639 | 1.170259037 | 26.82432387 | 27.32688848 | 27.13418219 | 25.25771039 | 25.79520973 | 24.44897026 | 27.57701085 | 27.24762378 | 26.91638574 |
| Carcinoembryonic antigen-related cel | P40199 | CEACAM6 | Extracellular or secreted | 0.486968393 | 1.277421628 | 21.35105353 | 21.78244861 | 21.88311907 | 20.82970446 | 21.57034728 | 21.01033908 | 21.69892675 | 21.86791232 | 22.10757213 |
| CCAAT/enhancer-binding protein beta | P17676-3;P17676-2;P17676 | CEBPB | Other | -0.490437721 | 0.808587603 | 25.01120268 | 25.09522831 | 25.08046201 | 24.17417108 | 24.35635632 | 25.01303603 | 24.4862802 | 24.56581332 | 23.84182153 |
| CCAAT/enhancer-binding protein zeta | Q03701 | CEBPZ | Other | -3.561982805 | 1.026625255 | 26.9280703 | 26.10970554 | 27.38046952 | 24.86145423 | 28.45316901 | 26.29803067 | 25.55888357 | 21.18567076 | 22.58494689 |
| Cat eye syndrome critical region prote | Q9BXW7-2;Q9BXW7 | CECR5 | Extracellular or secreted | -0.324421813 | 0.247327013 | 26.5734758 | 26.96388986 | 26.88739281 | 27.70186906 | 27.62219518 | 28.83605267 | 26.63256306 | 26.77039431 | 27.91621488 |
| CUGBP Elav-like family member 1 | Q92879-5;Q92879-2;Q92879 | CELF1 | Other | -0.487558439 | 0.617599044 | 25.03833563 | 25.50621507 | 25.31399374 | 25.90563894 | 25.98043684 | 26.54538013 | 25.60167242 | 25.41432007 | 24.66633237 |
| Cadherin EGF LAG seven-pass G-type | Q9NYQ6 | CELSR1 | Extracellular or secreted | -0.088224428 | 0.200024014 | 22.20072144 | 22.89971463 | 22.23561227 | 21.85490794 | 21.83753554 | 22.38763873 | 22.14448025 | 22.27233021 | 22.02658153 |
| Cadherin EGF LAG seven-pass G-type | Q9HCU4 | CELSR2 | Extracellular or secreted | -0.053253263 | 0.039158833 | 22.6418295 | 21.53488038 | 22.33606302 | 21.87019311 | 22.2893739 | 22.81182562 | 21.61103741 | 21.99106229 | 22.98022327 |
| Centromere protein F | P49454 | CENPF | Other | -3.167200895 | 1.879689537 | 25.6156018 | 25.13057339 | 24.74924299 | 25.36068235 | 25.08318389 | 27.85382589 | 23.12793955 | 21.21839767 | 23.04861525 |
| Centromere protein V | Q7Z7K6;Q7Z7K6-3;Q7Z7K6-2 | CENPV | Other | -0.460981219 | 0.56798842 | 22.30530861 | 22.70453101 | 23.03850309 | 23.03905559 | 22.97235582 | 24.89813747 | 22.85327148 | 22.54874949 | 22.69398117 |
| Centrosomal protein of 164 kDa | Q9UPV0-2;Q9UPV0 | CEP164 | Extracellular or secreted | 0.237954321 | 0.160183573 | 25.68281803 | 26.35107375 | 26.43213164 | 27.48323838 | 27.09526856 | 27.95460209 | 26.56034216 | 26.69178952 | 27.96129751 |
| Centrosomal protein of 170 kDa | Q5SW79;Q5SW79-2;Q5SW79 | CEP170 | Other | 0.197999052 | 0.210889797 | 24.80439733 | 24.06858018 | 24.52963596 | 24.23768496 | 23.86787671 | 25.58072413 | 24.44834018 | 24.43300771 | 25.2570989 |
| Centrosomal protein of 170 kDa prote | Q9Y4F5-2;Q9Y4F5;Q9Y4F5-3 | CEP170B | Other | 0.488234765 | 0.427921046 | 23.04626181 | 22.80433824 | 23.44392189 | 23.38018798 | 20.66915452 | 22.66179683 | 23.6846382 | 23.12237304 | 22.66052369 |
| Centrosome-associated protein 350 | Q5VT06 | CEP350 | Other | -0.629305898 | 0.200430773 | 23.99455371 | 22.14387261 | 21.73204376 | 22.66523254 | 24.85414703 | 25.82619234 | 20.91430616 | 23.64910791 | 24.15668923 |
| Centrosomal protein of 41 kDa | Q9BYV8;Q9BYV8-5;Q9BYV8-2 | CEP41 | Extracellular or secreted | 0.389362625 | 0.260080787 | 23.97963034 | 24.35957692 | 24.43364453 | 22.34346907 | 21.48783464 | 24.53082684 | 23.65425317 | 23.51929359 | 24.56203229 |
| Centrin-2 | P41208 | CETN2 | Other | 2.625277791 | 2.85982195 | 25.03061124 | 25.88716036 | 25.74224302 | 25.26338125 | 24.97971755 | 25.94777514 | 27.71613799 | 28.61235003 | 27.97278964 |
| Cilia- and flagella-associated protein 2 | Q9Y6A4 | CFAP20 | Extracellular or secreted | 0.04506678 | 0.029309826 | 25.71101216 | 25.7370674 | 26.44156521 | 25.76602698 | 24.18359056 | 26.07471706 | 25.3281391 | 25.24107221 | 26.52297871 |
| Cilia- and flagella-associated protein 3 | Q96G28;Q96G28-2 | CFAP36 | Other | 1.633302953 | 1.360225833 | 22.63169752 | 21.04272841 | 19.38794402 | 23.66155631 | 22.21740548 | 22.53002906 | 23.81567802 | 23.36666372 | 23.45324752 |
| Craniofacial development protein 1 | Q9UEE9;Q9UEE9-2 | CFDP1 | Extracellular or secreted | 0.037481303 | 0.086053718 | 27.86319254 | 27.7222103 | 27.75995897 | 27.98272312 | 27.74836025 | 27.0372676 | 27.71881818 | 27.58168687 | 27.86879524 |
| Cofilin-1 | P23528 | CFL1 | Extracellular or secreted | -0.046957404 | 0.104732085 | 34.02515119 | 34.1669845 | 34.11590156 | 34.17719555 | 34.17086874 | 33.41462855 | 34.10266716 | 34.04848105 | 33.74334462 |
| Cofilin-2 | Q9Y281;Q9Y281-3 | CFL2 | Extracellular or secreted | 0.411026165 | 1.427975573 | 26.858991 | 26.84397892 | 26.63173082 | 26.69325276 | 26.51422994 | 26.32966237 | 27.14559621 | 27.18735253 | 26.83605267 |
| Cingulin | Q9P2M7;Q9P2M7-2 | CGN | Other | 0.026576468 | 0.054545759 | 28.07270499 | 28.03170524 | 27.99507151 | 27.81127434 | 27.88547398 | 27.70701424 | 27.74586263 | 27.8430804 | 28.24240853 |
| Putative glutathione-specific gamma-g | Q8WUX2 | CHAC2 | Other | 1.810829884 | 1.161403354 | 24.49130509 | 21.22369847 | 24.45951467 | 25.70104941 | 25.35598684 | 21.62756532 | 25.86606148 | 25.78922617 | 25.2067619 |
| Chromatin assembly factor 1 subunit | Q13111-2;Q13111-3;Q13111 | CHAF1A | Other | -0.495622181 | 0.253368273 | 20.78258952 | 20.88375301 | 20.98558348 | 22.22696835 | 22.04510224 | 22.3403076 | 21.96757882 | 19.70824576 | 21.46946098 |
| Chromatin assembly factor 1 subunit | Q13112 | CHAF1B | Other | 0.146984267 | 0.726774504 | 26.07488035 | 26.04797492 | 26.05057192 | 26.29012307 | 26.45849811 | 26.37394814 | 26.42921287 | 26.4290532 | 26.23068499 |
| Chromosome alignment-maintaining | Q96JM3 | CHAMP1 | Other | 0.465529978 | 1.818046442 | 24.50597288 | 25.0129082 | 24.8632989 | 24.55666368 | 24.50172787 | 25.15360027 | 25.222251 | 25.0857791 | 25.38564573 |
| Coiled-coil-helix-coiled-coil-helix dom | Q9Y6H1;Q5T1J5 | CHCHD2;CHCHD2 | Other | 1.021296907 | 1.092101714 | 24.77489924 | 24.33408611 | 24.28841665 | 25.71174741 | 25.57542476 | 25.33776507 | 26.62338187 | 25.3741472 | 26.07753127 |
| Mitochondrial intermembrane space | Q8N4Q1;Q8N4Q1-2 | CHCHD4 | Other | -0.192219592 | 0.283311984 | 28.48316149 | 28.77077236 | 28.80568927 | 28.82935353 | 28.88107432 | 27.39835881 | 28.32268325 | 28.62411433 | 28.06074853 |
| Coiled-coil-helix-coiled-coil-helix dom | Q9BSY4;Q9BSY4-2 | CHCHD5 | Other | 0.749745606 | 0.211989959 | 23.56337132 | 23.26212697 | 23.39218009 | 23.36599664 | 24.11187559 | 22.43257442 | 25.59448612 | 25.11672152 | 21.60209169 |
| Chromodomain-helicase-DNA-binding | O14647-2;O14647;O14647-3 | CHD2 | Extracellular or secreted | -1.766490897 | 1.003682187 | 23.53409671 | 23.10376803 | 23.40509575 | 22.87958998 | 23.28912057 | 22.61146383 | 22.17963262 | 21.79264674 | 20.13981538 |
| Chromodomain-helicase-DNA-binding | Q12873-2;Q12873;Q12873-3 | CHD3 | Other | -2.642743617 | 1.038919812 | 25.86910129 | 25.28047373 | 25.629259 | 23.26670878 | 23.27426553 | 25.84639616 | 23.29361758 | 23.08773863 | 20.27351517 |
| Chromodomain-helicase-DNA-binding | Q14839;Q14839-2 | CHD4 | Other | -0.931872806 | 2.449739378 | 30.25822276 | 30.29493679 | 30.36650949 | 29.73926158 | 29.69365157 | 30.90906316 | 29.33517715 | 29.25932329 | 29.24070382 |
| Chromodomain-helicase-DNA-binding | Q9HCK8-2;Q9HCK8 | CHD8 | Other | -0.275269676 | 0.254243285 | 23.08093616 | 22.56101845 | 21.25281263 | 22.71104155 | 22.36251497 | 22.07633284 | 21.98672951 | 21.50688492 | 22.70290485 |
| Serine/threonine-protein kinase Chk1 | O14757-2;O14757-3;O14757 | CHEK1 | Extracellular or secreted | 0.06950096 | 0.09066998 | 21.52806801 | 20.3901832 | 22.08315579 | 21.31444764 | 22.20121255 | 21.1169277 | 21.53886337 | 21.40288021 | 21.58375674 |
| Serine/threonine-protein kinase Chk2 | O96017;O96017-13;O96017- | CHEK2 | Extracellular or secreted | -0.702347069 | 0.652370291 | 25.20735786 | 24.54173709 | 24.4865871 | 24.41409421 | 24.37125819 | 25.1667595 | 23.20874994 | 24.59838196 | 24.17972386 |
| Calcium homeostasis endoplasmic ret | Q8IWX8 | CHERP | Other | -0.128259562 | 0.161306856 | 26.74201167 | 26.56733747 | 26.25389335 | 26.00485326 | 26.15156955 | 27.29876488 | 26.31617103 | 26.00534597 | 26.80291941 |
| Chitinase domain-containing protein 1 | Q9BWS9-3;Q9BWS9;Q9BWS9 | CHID1 | Extracellular or secreted | 0.713807469 | 1.873635781 | 23.87988907 | 23.77374289 | 23.9097642 | 23.63500155 | 23.05097455 | 24.28347953 | 24.48185362 | 24.23154582 | 24.69444886 |
| Rab proteins geranylgeranyltransferas | P24386 | CHM | Other | 0.330701209 | 0.737206998 | 27.03653417 | 26.67855308 | 26.65621884 | 26.08705472 | 26.36370955 | 26.14485752 | 26.67505661 | 27.14102193 | 26.65948902 |
| Rab proteins geranylgeranyltransferas | P26374 | CHML | Other | -0.49766245 | 0.13243703 | 22.67243138 | 20.5289731 | 22.02049034 | 22.26732939 | 23.06785668 | 21.48520421 | 22.50690511 | 23.02430034 | 18.99694976 |
| Charged multivesicular body protein 1 | Q9HD42 | CHMP1A | Extracellular or secreted | -0.022839573 | 0.060205004 | 27.88180367 | 27.98945202 | 27.94547741 | 28.26293342 | 28.34823945 | 28.23338482 | 28.18640823 | 28.19952565 | 27.87619279 |
| Charged multivesicular body protein 1 | Q7LBR1 | CHMP1B | Extracellular or secreted | -0.165731553 | 0.135476298 | 27.7498951 | 25.68056642 | 25.90117957 | 26.58132772 | 26.22905349 | 26.67343999 | 26.24239944 | 26.93055382 | 25.73758322 |
| Charged multivesicular body protein 3 | Q9Y3E7;Q9Y3E7-3;Q9Y3E7-2; | CHMP3 | Extracellular or secreted | 0.596711995 | 2.466806518 | 27.05889231 | 26.6065376 | 26.71881818 | 26.68565435 | 26.83942064 | 27.01467554 | 27.39762463 | 27.57607383 | 27.27843684 |
| Charged multivesicular body protein 4 | Q9BY43;Q9BY43-2 | CHMP4A | Extracellular or secreted | -0.538150583 | 0.417287328 | 26.95293725 | 26.68485219 | 26.64127276 | 26.91421996 | 26.5579493 | 27.53311653 | 26.74919183 | 26.89839122 | 25.3801392 |
| Charged multivesicular body protein 4 | Q9H444 | CHMP4B | Extracellular or secreted | 1.94477243 | 4.321046633 | 27.24744269 | 26.93741765 | 27.23086818 | 27.19901073 | 26.68605527 | 28.00629877 | 29.29366581 | 29.10414609 | 29.09005203 |
| Charged multivesicular body protein 5 | Q9NZZ3;Q9NZZ3-2 | CHMP5 | Extracellular or secreted | 1.677047985 | 3.617062767 | 24.16790811 | 24.54398005 | 25.17165388 | 24.08082793 | 25.00812766 | 25.18146895 | 26.47773764 | 26.25445213 | 26.37593747 |
| Cysteine and histidine-rich domain-co | Q9UHD1;Q9UHD1-2 | CHORDC1 | Other | 0.005934176 | 0.0048586 | 30.05581915 | 29.98196143 | 29.94265602 | 30.69170633 | 30.69900819 | 29.40403603 | 30.33212785 | 30.60195927 | 29.471309 |
| Chromatin accessibility complex prote | Q9NRG0 | CHRAC1 | Other | 0.544878072 | 0.581395132 | 24.6865095 | 24.37815565 | 24.52073281 | 24.43974365 | 24.12004862 | 23.376898 | 24.98932208 | 25.29281056 | 24.1135457 |
| Chromatin target of PRMT1 protein | Q9Y3Y2-4;Q9Y3Y2;Q9Y3Y2-3 | CHTOP | Other | 0.442741318 | 0.248575401 | 23.33429075 | 23.33619925 | 23.5584749 | 23.54403903 | 21.813066 | 23.65774585 | 24.02867366 | 24.48308459 | 22.43837359 |
| Inhibitor of nuclear factor kappa-B kin | O15111 | CHUK | Extracellular or secreted | -0.658346949 | 0.463755151 | 19.64656574 | 22.57957514 | 22.01184106 | 20.89296964 | 22.26088852 | 19.66421115 | 21.25193771 | 20.12313122 | 20.17791585 |
| Protein Churchill | Q8WUH1;Q8WUH1-2 | CHURC1 | Other | 0.67809688 | 0.586556386 | 21.19324121 | 20.64654429 | 21.01951749 | 22.09284955 | 21.66024952 | 20.97267685 | 22.03607297 | 22.6510368 | 21.13972032 |
| Probable cytosolic iron-sulfur protein | O76071 | CIAO1 | Other | 0.280237105 | 1.913212011 | 27.25979469 | 27.49060144 | 27.37376565 | 27.3778248 | 27.306608 | 27.15826975 | 27.6082309 | 27.51239242 | 27.70352016 |
| Anamorsin | Q6FI81-3;Q6FI81 | CIAPIN1 | Other | 0.231046649 | 0.348460047 | 27.86926606 | 27.89752597 | 27.70021607 | 28.03758181 | 28.03768653 | 27.34375732 | 28.13530809 | 28.40956963 | 27.5912791 |
| Calcium and integrin-binding protein 1 | Q99828;Q99828-2 | CIB1 | Extracellular or secreted | 0.431316486 | 0.102559251 | 22.65667475 | 22.72338264 | 20.47937561 | 22.24629409 | 21.86908736 | 22.27745071 | 19.83828971 | 22.98836016 | 24.59343216 |
| Protein capicua homolog | Q96RK0 | CIC | Other | -0.948933042 | 0.471040118 | 22.34697571 | 22.3794651 | 22.4542772 | 22.68296802 | 21.47960371 | 22.08141322 | 21.43023755 | 19.90483322 | 22.53048159 |
| Cold-inducible RNA-binding protein | Q14011;Q14011-2 | CIRBP | Other | 1.012196138 | 2.239646376 | 29.34252864 | 29.32648137 | 29.23327052 | 29.57564115 | 29.60036493 | 28.10794121 | 30.16710418 | 30.024722 | 30.43787615 |
| Cirhin | Q969X6-2;Q969X6-3;Q969X6 | CIRH1A | Extracellular or secreted | -1.391343343 | 1.184634461 | 23.63928 | 22.02924678 | 20.97746795 | 23.16034071 | 22.47212848 | 23.93669904 | 20.49363352 | 21.52836063 | 21.9115573 |
| Citron Rho-interacting kinase | O14578-3;O14578;O14578-4 | CIT | Other | 0.377306742 | 0.527707415 | 22.23521765 | 22.10424809 | 22.13777417 | 22.41488133 | 21.07676636 | 22.66075151 | 22.12119237 | 22.39115425 | 22.93439316 |
| Cip1-interacting zinc finger protein | Q9ULV3-5;Q9ULV3-3;Q9ULV3 | CIZ1 | Other | 0.320914799 | 0.26410709 | 19.91591671 | 22.09451975 | 22.12483568 | 22.10722091 | 21.9556045 | 23.45412656 | 22.53969178 | 21.90440824 | 22.34475643 |
| Cytoskeleton-associated protein 4 | Q07065 | CKAP4 | Extracellular or secreted | -0.79989909 | 0.42929444 | 26.75531793 | 26.75251336 | 27.28453891 | 26.67949299 | 26.27804684 | 29.69149003 | 25.50357902 | 26.17470446 | 27.64271928 |
| Cytoskeleton-associated protein 5 | Q14008-2;Q14008;Q14008-3 | CKAP5 | Other | -0.631486973 | 2.301708548 | 29.83142393 | 30.00517461 | 29.94600756 | 29.89071564 | 29.80532026 | 29.8528591 | 29.18075315 | 29.19199419 | 29.39854229 |
| Creatine kinase B-type | P12277 | CKB | Extracellular or secreted | -0.344697753 | 0.301361879 | 26.45444038 | 26.34267275 | 26.35319583 | 28.25763846 | 28.34129892 | 28.79912532 | 27.12346675 | 26.97901971 | 27.13760612 |
| Creatine kinase U-type, mitochondrial | P12532;P12532-2 | CKMT1A | Extracellular or secreted | -1.079855808 | 3.065715157 | 27.39795098 | 27.21872685 | 27.19863613 | 27.24544927 | 27.34316429 | 27.91530326 | 26.22376134 | 26.53996386 | 26.15632276 |
| Cyclin-dependent kinases regulatory s | P61024 | CKS1B | Other | 0.435351299 | 0.449223318 | 26.08794496 | 25.96388986 | 25.88822932 | 26.74663159 | 26.81274372 | 25.29900953 | 26.66343063 | 27.11217396 | 25.92967379 |
| Cyclin-dependent kinases regulatory s | P33552 | CKS2 | Other | 1.066267636 | 0.788438154 | 22.44211258 | 21.61485998 | 22.54041539 | 22.56153129 | 22.6943851 | 22.65044525 | 24.46768365 | 23.23268792 | 22.75030613 |
| CLIP-associating protein 1 | Q7Z460-2;Q7Z460-4;Q7Z460- | CLASP1 | Extracellular or secreted | 0.45881128 | 0.629075179 | 25.97862703 | 25.70131387 | 26.36974496 | 25.15325235 | 25.16641473 | 25.89173272 | 25.64979318 | 26.54142693 | 26.31575656 |
| CLIP-associating protein 2 | O75122-3 | CLASP2 | Extracellular or secreted | 0.0682471 | 0.073104309 | 23.98884553 | 24.08634618 | 23.98433223 | 22.0165441 | 23.8089202 | 23.97762302 | 23.66871449 | 23.63997655 | 23.82735589 |
| Claudin-1 | O95832 | CLDN1 | Extracellular or secreted | 4.234541457 | 3.297419692 | 22.50823582 | 23.89559747 | 24.22493905 | 22.83705426 | 21.93589616 | 25.86452737 | 27.52342747 | 27.27949992 | 28.53382203 |
| Claudin-3 | O15551 | CLDN3 | Extracellular or secreted | 2.71814325 | 1.731607048 | 22.21922277 | 20.83549093 | 20.58532086 | 21.97813319 | 22.65562942 | 21.8959672 | 23.84114969 | 23.86665109 | 25.53151116 |
| Claudin-4 | O14493 | CLDN4 | Extracellular or secreted | 2.146823223 | 3.237530132 | 24.48861098 | 25.05233521 | 24.78611286 | 24.76531888 | 24.30632986 | 26.02299338 | 27.19178246 | 26.61357996 | 27.34595784 |
| Claudin-7 | O95471-2;O95471 | CLDN7 | Extracellular or secreted | 2.859620095 | 1.710505969 | 24.47326925 | 24.52748989 | 24.4388558 | 21.54561602 | 21.80257123 | 27.22775063 | 26.85495548 | 26.48270002 | 27.24898118 |
| Claudin domain-containing protein 1 | Q9NY35-2;Q9NY35 | CLDND1 | Extracellular or secreted | 1.393584033 | 3.334581495 | 22.16257081 | 22.09113317 | 22.39495561 | 23.03434426 | 22.73440284 | 23.23047611 | 23.82162608 | 23.95144836 | 24.23161906 |
| Protein CLEC16A | Q2KHT3-2;Q2KHT3 | CLEC16A | Extracellular or secreted | 0.229847869 | 0.427219202 | 28.4392443 | 28.44517774 | 28.50880855 | 28.65730973 | 28.82040313 | 28.6920224 | 28.59448612 | 28.65058084 | 29.22595957 |
| Chloride intracellular channel protein | O00299 | CLIC1 | Extracellular or secreted | 0.420451799 | 1.188234045 | 32.26631671 | 32.22750965 | 32.2456986 | 32.00145464 | 31.94715785 | 32.22837009 | 32.52309092 | 32.81941747 | 32.37710079 |
| Chloride intracellular channel protein | Q9Y696 | CLIC4 | Extracellular or secreted | 0.39827104 | 1.30979571 | 28.52525847 | 28.58742125 | 28.48880249 | 28.13951021 | 27.82844585 | 28.46060863 | 28.54373674 | 28.92331715 | 28.74278269 |
| Clathrin interactor 1 | Q14677-3;Q14677-2;Q14677 | CLINT1 | Other | -0.743623603 | 0.885977494 | 28.0560518 | 27.72533443 | 27.71679215 | 28.05775678 | 28.12740827 | 28.03307158 | 26.77618019 | 27.04614382 | 27.80501268 |
| CAP-Gly domain-containing linker pro | P30622-2;P30622-1;P30622 | CLIP1 | Extracellular or secreted | 1.298708078 | 3.173459451 | 27.50248682 | 27.54642562 | 27.72005853 | 27.57888308 | 27.78069214 | 27.34857716 | 28.89593832 | 29.04476376 | 28.69398383 |
| Calmin | Q96JQ2 | CLMN | Other | 0.031741978 | 0.041257958 | 26.73389105 | 26.48448354 | 26.2747454 | 25.98082903 | 25.91369513 | 26.01241807 | 26.14530467 | 25.95298167 | 26.69697071 |
| Methylosome subunit pICln | P54105 | CLNS1A | Extracellular or secreted | 0.802619193 | 3.547864048 | 27.34163826 | 27.46878697 | 27.33636953 | 27.6345031 | 27.59327542 | 27.9644959 | 28.36026373 | 28.32036083 | 28.39676762 |
| Caseinolytic peptidase B protein hom | Q9H078-3;Q9H078-2;Q9H078 | CLPB | Other | -0.315730411 | 0.934629582 | 26.54022999 | 26.42779122 | 26.21361911 | 25.8876717 | 25.91520067 | 26.10940665 | 26.00889703 | 25.97543798 | 25.61543342 |
| ATP-dependent Clp protease proteoly | Q16740 | CLPP | Other | -0.317474678 | 0.421975389 | 26.25625318 | 26.24025344 | 26.14493529 | 26.63089809 | 26.63906575 | 27.74554211 | 26.18876781 | 25.95688534 | 26.73039675 |
| Colipase | P04118 | CLPS | Extracellular or secreted | 0.664530433 | 0.303723353 | 21.42582377 | 23.07579445 | 21.09084229 | 24.71177366 | 24.2617684 | 22.97650581 | 24.23300989 | 24.31395915 | 22.21787645 |
| Cleft lip and palate transmembrane p | O96005;O96005-4 | CLPTM1 | Extracellular or secreted | -0.400699014 | 0.176879382 | 20.80697566 | 21.26124209 | 22.44346682 | 21.99168515 | 21.28048081 | 25.24633759 | 20.96444523 | 21.43354265 | 22.91500914 |
| Calsyntenin-1;Soluble Alc-alpha;CTF1- | O94985-2;O94985 | CLSTN1 | Extracellular or secreted | 0.473888992 | 0.336215967 | 29.87978977 | 29.84456266 | 29.8651117 | 30.87920551 | 30.93318787 | 32.38805133 | 30.60434747 | 30.73860143 | 31.97367251 |
| Calsyntenin-2 | Q9H4D0 | CLSTN2 | Extracellular or secreted | -0.87219202 | 0.301473681 | 23.5380108 | 21.03412289 | 23.91570216 | 24.80055159 | 24.98324519 | 27.08365056 | 21.83017365 | 23.3267685 | 24.90412338 |
| Calsyntenin-3 | Q9BQT9;Q9BQT9-2 | CLSTN3 | Extracellular or secreted | 0.09021616 | 0.053956093 | 26.69085759 | 26.52710207 | 26.81225409 | 27.61911922 | 27.6719565 | 29.13033281 | 27.01329166 | 27.12671928 | 28.35644868 |
| Clathrin light chain A | P09496-2;P09496-4;P09496-3 | CLTA | Extracellular or secreted | 0.769066509 | 4.090030005 | 27.51389876 | 27.56797568 | 27.61701821 | 27.37044357 | 27.71469778 | 27.30225598 | 28.18181961 | 28.29136205 | 28.37716286 |
| Clathrin light chain B | P09497-2;P09497 | CLTB | Extracellular or secreted | 0.795410175 | 2.69370419 | 27.20004039 | 27.31371702 | 27.58792192 | 27.0964755 | 26.98076367 | 26.77530123 | 27.78419171 | 28.14515887 | 27.93398981 |
| Clathrin heavy chain 1 | Q00610-2;Q00610 | CLTC | Extracellular or secreted | -0.314312693 | 0.845048057 | 34.20514565 | 34.30409359 | 34.26932983 | 33.83026442 | 33.88876132 | 33.7753915 | 33.90875904 | 33.83469725 | 33.45009879 |
| Clathrin heavy chain 2 | P53675-2;P53675 | CLTCL1 | Extracellular or secreted | 2.298636218 | 1.291297965 | 20.11329666 | 21.0860823 | 20.36952494 | 22.87310492 | 22.71143346 | 22.75046002 | 25.31523831 | 23.12661689 | 23.40600461 |
| Clusterin;Clusterin beta chain;Clusteri | P10909-4;P10909;P10909-5;P | CLU | Extracellular or secreted | -0.729441089 | 0.311742943 | 25.15526141 | 25.36992796 | 25.01418601 | 26.89342338 | 27.02464275 | 29.19783977 | 24.91920778 | 25.06212884 | 27.15798076 |
| Clustered mitochondria protein homo | O75153 | CLUH | Other | 0.515052045 | 1.080591723 | 29.01690824 | 29.19643336 | 29.13325143 | 29.58188437 | 29.67569602 | 29.30515153 | 30.03708428 | 29.99563765 | 29.46709669 |
| N-acylneuraminate cytidylyltransferas | Q8NFW8 | CMAS | Other | -0.658977615 | 3.385740048 | 26.82323078 | 26.92445027 | 27.10069185 | 26.94246028 | 26.91479022 | 27.41635119 | 26.34245235 | 26.27177475 | 26.46982735 |
| COX assembly mitochondrial protein h | Q7Z7K0 | CMC1 | Other | 0.183755621 | 0.225999706 | 25.42066188 | 25.16491981 | 25.28026132 | 25.31492726 | 25.63284037 | 24.64014209 | 25.65624612 | 25.74870573 | 24.87319138 |
| UMP-CMP kinase | P30085;P30085-2 | CMPK1 | Extracellular or secreted | 0.472620087 | 2.089613099 | 29.89922714 | 29.87764144 | 29.87570959 | 29.87063349 | 29.81792056 | 29.38337725 | 30.19999361 | 30.42672406 | 30.15339733 |
| CKLF-like MARVEL transmembrane do | Q9NX76 | CMTM6 | Extracellular or secreted | 2.232050112 | 1.606494557 | 20.88073051 | 20.77607384 | 20.98447131 | 22.26562255 | 23.52706732 | 20.13683595 | 22.62046506 | 24.04643529 | 24.31465073 |
| Cap-specific mRNA (nucleoside-2-O-)- | Q8N1G2 | CMTR1 | Other | -0.729764097 | 0.333372971 | 27.38853895 | 27.90517984 | 27.24698989 | 26.22749359 | 27.31371702 | 27.23726555 | 27.29911437 | 27.29377542 | 24.87741034 |
| Cellular nucleic acid-binding protein | P62633-2;P62633;P62633-7;P | CNBP | Other | 0.234102202 | 0.276956549 | 29.27342093 | 28.8758121 | 28.90635597 | 30.46732995 | 30.31753879 | 30.34148981 | 30.03891644 | 30.15532897 | 29.59903497 |
| Cellular nucleic acid-binding protein | P62633-8;P62633-4 | CNBP | Other | 0.140802395 | 0.108962896 | 23.30033692 | 22.9433862 | 23.1281284 | 24.79277957 | 24.80572616 | 24.30709461 | 24.42534398 | 24.26140975 | 23.3743794 |
| Cytosolic non-specific dipeptidase | Q96KP4;Q96KP4-2 | CNDP2 | Extracellular or secreted | 0.358031292 | 0.445193587 | 31.54655441 | 31.62370455 | 31.57974637 | 31.503416 | 31.54157833 | 30.3661968 | 32.0644217 | 31.93343413 | 31.15683627 |
| Protein cornichon homolog 4 | Q9P003 | CNIH4 | Other | 0.410705386 | 0.138772524 | 22.87141573 | 20.28405036 | 21.78711727 | 22.3074204 | 21.88568644 | 25.45513053 | 21.14767588 | 23.47072738 | 23.90912327 |
| Calponin-2 | Q99439;Q99439-2 | CNN2 | Extracellular or secreted | 0.577987994 | 2.397680135 | 26.69630749 | 26.62839703 | 26.82092042 | 27.15093073 | 26.95681889 | 27.41514253 | 27.50536721 | 27.46084294 | 27.60201238 |
| Calponin-3 | Q15417;Q15417-3;Q15417-2 | CNN3 | Other | 1.096442845 | 2.444888263 | 26.43383551 | 26.31584292 | 26.49095331 | 26.87970215 | 26.76963792 | 26.0123115 | 27.48730805 | 27.36536263 | 27.88779951 |
| Metal transporter CNNM4 | Q6P4Q7;Q6P4Q7-2 | CNNM4 | Extracellular or secreted | -0.184312556 | 0.126442871 | 23.50603343 | 23.38495445 | 23.45525597 | 23.01901444 | 23.00725522 | 23.95287061 | 22.30537166 | 23.98641705 | 23.31796568 |
| CCR4-NOT transcription complex subu | A5YKK6;A5YKK6-2;A5YKK6-3; | CNOT1 | Extracellular or secreted | -0.416304405 | 1.425922943 | 28.69125706 | 28.80168665 | 28.81674633 | 27.95487937 | 28.21516629 | 28.54723499 | 28.02173393 | 28.00088406 | 28.24195414 |
| CCR4-NOT transcription complex subu | Q9H9A5-2;Q9H9A5;Q9H9A5- | CNOT10 | Other | -0.345343652 | 0.951567353 | 25.76336982 | 25.63173082 | 25.45305909 | 24.7741452 | 25.26710185 | 26.01563283 | 25.10433608 | 25.06693926 | 25.2452135 |
| CCR4-NOT transcription complex subu | Q9UKZ1 | CNOT11 | Other | -0.332718214 | 0.452606857 | 24.60444645 | 24.56302213 | 24.44619584 | 23.93346087 | 24.04101271 | 24.70097007 | 24.04360141 | 23.57865278 | 24.5241452 |
| CCR4-NOT transcription complex subu | Q9NZN8-2;Q9NZN8-5;Q9NZN | CNOT2 | Other | -0.12249848 | 0.17908008 | 25.18393124 | 24.76501531 | 24.90145581 | 23.95491265 | 24.65703708 | 24.87267488 | 24.21297009 | 24.89180224 | 24.69524571 |
| CCR4-NOT transcription complex subu | O75175 | CNOT3 | Other | -0.697492609 | 1.409066488 | 26.23284526 | 26.01239675 | 26.22330104 | 25.6037393 | 25.96657656 | 25.11747475 | 25.51385359 | 24.87262792 | 25.09920749 |
| CCR4-NOT transcription complex subu | Q96LI5;Q9ULM6 | CNOT6L;CNOT6 | Other | 0.229388994 | 0.551306403 | 21.06532935 | 21.22340784 | 21.18646491 | 20.92465153 | 22.17928132 | 21.41998075 | 21.51710803 | 21.69481015 | 21.47580664 |

| | | | | | | | | | | | | | | |
|---|---|---|---|---|---|---|---|---|---|---|---|---|---|---|
| CCR4-NOT transcription complex subu | Q9UIV1;Q9UIV1-2 | CNOT7 | Other | -0.008125348 | 0.026784857 | 26.46259126 | 26.44589601 | 26.40031478 | 26.45693277 | 26.35492824 | 25.90499617 | 26.30465992 | 26.43671319 | 26.24708046 |
| 2,3-cyclic-nucleotide 3-phosphodieste | P09543-2;P09543 | CNP | Extracellular or secreted | 0.538118953 | 2.090273663 | 27.32130745 | 27.29386311 | 27.23214993 | 26.89919832 | 26.88936712 | 27.66587174 | 27.66010136 | 27.68023106 | 27.92490327 |
| Protein canopy homolog 2 | Q9Y2B0 | CNPY2 | Extracellular or secreted | -0.628368034 | 1.831927434 | 29.21558291 | 29.19882344 | 29.14068072 | 29.23895155 | 29.38935983 | 28.55651021 | 28.63803005 | 28.61147086 | 28.23534932 |
| Protein canopy homolog 3 | Q9BT09 | CNPY3 | Extracellular or secreted | -0.715661751 | 3.041389792 | 27.06825215 | 27.13965658 | 27.07581893 | 26.72987835 | 26.90161692 | 26.46661061 | 26.21491626 | 26.03818909 | 26.29082617 |
| Cytochrome c oxidase assembly facto | Q5JTJ3-3;Q5JTJ3;Q5JTJ3-2 | COA6 | Other | 0.277154508 | 0.523670424 | 27.35800107 | 27.61701821 | 27.71450127 | 27.58348131 | 27.73065588 | 26.59192108 | 27.9647713 | 27.77184296 | 27.39263868 |
| Cytochrome c oxidase assembly facto | Q96BR5 | COA7 | Other | 0.510515269 | 1.168669406 | 27.59676234 | 27.79538316 | 27.62226501 | 27.8363537 | 28.04754856 | 27.25475847 | 28.36628019 | 28.37654202 | 27.86525922 |
| Bifunctional coenzyme A synthase;Ph | Q13057;Q13057-2 | COASY | Extracellular or secreted | 0.072284504 | 0.037012504 | 24.71675291 | 25.2704921 | 25.04938826 | 24.99851881 | 24.77213251 | 25.6614204 | 25.52294879 | 25.97832153 | 23.94993568 |
| Cochlin | O43405;O43405-2 | COCH | Extracellular or secreted | 1.38399892 | 0.737326945 | 20.99439202 | 20.3440462 | 21.02469433 | 23.51400414 | 23.48222302 | 25.71311189 | 23.55859167 | 23.45412656 | 24.67551432 |
| Conserved oligomeric Golgi complex s | Q8WTW3 | COG1 | Other | 0.028134414 | 0.039180812 | 23.64273305 | 23.89818361 | 23.34840831 | 23.69646669 | 22.89565294 | 23.92678171 | 23.39322809 | 23.97263629 | 23.42265202 |
| Conserved oligomeric Golgi complex s | Q14746-2;Q14746 | COG2 | Other | -0.676758674 | 0.693095573 | 25.10245505 | 25.0209396 | 25.08277796 | 24.13453474 | 24.41079913 | 24.52043309 | 24.68779128 | 23.39100019 | 24.02690229 |
| Conserved oligomeric Golgi complex s | Q96JB2;Q96JB2-2 | COG3 | Extracellular or secreted | -0.870073289 | 1.21316443 | 25.18725812 | 24.86839493 | 24.80237713 | 25.31868981 | 26.46923738 | 25.11159705 | 24.84191748 | 24.50966194 | 23.91697792 |
| Conserved oligomeric Golgi complex s | Q9H9E3;Q9H9E3-3;Q9H9E3-2 | COG4 | Other | -0.359408421 | 0.5259526 | 24.98971186 | 25.33019257 | 25.23910639 | 25.25511878 | 25.32573969 | 25.36479512 | 25.0752885 | 25.21874532 | 24.38007312 |
| Conserved oligomeric Golgi complex s | Q9UP83;Q9UP83-2;Q9UP83- | COG5 | Other | -0.494766053 | 0.767509369 | 25.01354725 | 25.25868104 | 25.28662008 | 24.437396 | 25.04255814 | 25.12056285 | 24.94418194 | 24.56412762 | 24.08707496 |
| Conserved oligomeric Golgi complex s | Q9Y2V7;Q9Y2V7-2 | COG6 | Other | -0.393546857 | 0.632889401 | 25.45848246 | 25.02216798 | 25.20336772 | 24.57016377 | 24.70998744 | 24.38962241 | 24.87722309 | 24.53195727 | 24.08707496 |
| Conserved oligomeric Golgi complex s | P83436 | COG7 | Other | -0.5409143 | 0.756533646 | 25.55294649 | 25.44493298 | 25.08841009 | 25.18680491 | 25.33875134 | 24.74816828 | 24.75699806 | 25.12099782 | 24.17926827 |
| Conserved oligomeric Golgi complex s | Q96MW5 | COG8 | Other | 1.127808668 | 0.762191016 | 25.5167715 | 25.98585271 | 26.085151 | 25.13758657 | 21.57269013 | 25.66196398 | 26.433358 | 26.16079116 | 25.7692848 |
| Collagen alpha-1(XII) chain | Q99715;Q99715-4 | COL12A1 | Extracellular or secreted | 1.786997839 | 0.539110398 | 22.1880962 | 22.0114757 | 21.87368892 | 21.5634351 | 22.25554349 | 23.6053511 | 22.46819346 | 23.11876622 | 26.52282909 |
| Collagen alpha-1(XVIII) chain;Endostat | P39060-2;P39060-1;P39060 | COL18A1 | Extracellular or secreted | -1.138619379 | 1.287801837 | 25.81516495 | 25.791861 | 25.52880176 | 26.57873915 | 26.46538107 | 28.5283174 | 25.22298795 | 25.22129239 | 25.49399418 |
| Collagen alpha-1(IV) chain;Arresten;C | P02462;P29400;P29400-2 | COL4A1;COL4A5 | Extracellular or secreted | 0.014332996 | 0.009227661 | 23.08035628 | 23.2454529 | 22.93515282 | 23.38850611 | 22.93743561 | 25.47072738 | 23.44909624 | 22.97826916 | 24.14444914 |
| Collagen alpha-2(IV) chain;Canstatin | P08572 | COL4A2 | Extracellular or secreted | 0.831922771 | 0.485782069 | 24.00204377 | 24.68238943 | 24.87890744 | 26.14932275 | 26.25066259 | 28.1654758 | 25.62405156 | 26.60554892 | 27.33056872 |
| Collagen type IV alpha-3-binding prot | Q9Y5P4-2;Q9Y5P4;Q9Y5P4-3 | COL4A3BP | Other | 0.720509659 | 1.504259309 | 24.4794502 | 24.16890283 | 24.25645116 | 23.5809541 | 23.39728189 | 24.70445714 | 25.07500281 | 24.51225677 | 24.86801806 |
| Collagen alpha-1(V) chain | P20908-2;P20908 | COL5A1 | Extracellular or secreted | -1.790459339 | 1.662288415 | 23.42572813 | 22.73694358 | 21.06847082 | 23.93571037 | 24.53966812 | 24.46263806 | 21.48535775 | 21.12643021 | 22.10141356 |
| Collagen alpha-1(VI) chain | P12109 | COL6A1 | Extracellular or secreted | 0.93069982 | 0.935957113 | 28.63626765 | 28.67269844 | 28.59644247 | 29.61175225 | 29.70416357 | 30.35667955 | 29.72987835 | 29.91213661 | 30.93908647 |
| Collagen alpha-2(VI) chain | P12110;P12110-3;P12110-2 | COL6A2 | Extracellular or secreted | 0.173155933 | 0.069460248 | 23.70598667 | 23.98806539 | 24.19422722 | 24.79119032 | 25.13492636 | 27.41385218 | 24.45907685 | 24.26535007 | 26.40916495 |
| Collagen alpha-3(IX) chain | Q14050 | COL9A3 | Extracellular or secreted | 1.984440902 | 3.022596555 | 22.92556893 | 23.10717301 | 23.08136446 | 23.5168917 | 23.50142418 | 23.84555896 | 25.01171455 | 25.25223375 | 25.67836502 |
| Procollagen galactosyltransferase 1 | Q8NBJ5 | COLGALT1 | Extracellular or secreted | 0.030670465 | 0.03604238 | 27.85037219 | 27.8046435 | 27.81029391 | 27.61210391 | 27.81102929 | 29.03081117 | 27.83557089 | 27.7917244 | 28.42434309 |
| COMM domain-containing protein 1 | Q8N668-2;Q8N668 | COMMD1 | Extracellular or secreted | 0.782995136 | 1.199545751 | 23.89068955 | 23.85224298 | 23.41751055 | 23.29250626 | 23.24542702 | 24.306608 | 24.91369513 | 24.39218009 | 24.04560237 |
| COMM domain-containing protein 10 | Q9Y6G5 | COMMD10 | Other | 0.434294901 | 0.871177966 | 23.35218571 | 23.10102863 | 22.59354614 | 22.78036675 | 22.86404555 | 24.11806913 | 23.42265202 | 23.81636183 | 23.4684918 |
| COMM domain-containing protein 2 | Q86X83-2;Q86X83 | COMMD2 | Other | 0.279076575 | 0.494883866 | 25.20486065 | 25.27742618 | 25.05635161 | 24.97548171 | 25.05854143 | 24.88087004 | 25.5438031 | 25.59169285 | 24.92849958 |
| COMM domain-containing protein 3 | Q9UBI1 | COMMD3 | Other | 0.131975792 | 0.235692507 | 24.79104124 | 24.70619752 | 24.48111453 | 24.44764677 | 24.57369249 | 24.80375708 | 24.73365836 | 25.12463062 | 24.43936321 |
| COMM domain-containing protein 4 | Q9H0A8;Q9H0A8-3;Q9H0A8- | COMMD4 | Other | 0.065774526 | 0.308854852 | 24.93966099 | 25.00255889 | 25.38452635 | 24.88054312 | 25.00311674 | 25.05225228 | 25.04464392 | 25.21719272 | 25.06681612 |
| COMM domain-containing protein 5 | Q9GZQ3 | COMMD5 | Other | -0.230557902 | 0.253333294 | 24.23513016 | 24.63278491 | 24.55062937 | 23.82667727 | 23.57484757 | 22.91265402 | 23.8374008 | 24.1049359 | 23.23235124 |
| COMM domain-containing protein 7 | Q86VX2-2;Q86VX2 | COMMD7 | Extracellular or secreted | 1.380320808 | 2.780899837 | 21.32598778 | 21.6380213 | 21.404465 | 21.79817358 | 21.72370736 | 22.95014934 | 23.11818005 | 23.45550683 | 22.98752772 |
| COMM domain-containing protein 8 | Q9NX08 | COMMD8 | Other | 0.199672596 | 0.244531488 | 25.0867511 | 24.7321579 | 25.24099945 | 24.68099557 | 24.58514456 | 24.5903513 | 25.11791065 | 25.45973353 | 24.47957355 |
| COMM domain-containing protein 9 | Q9P000;Q9P000-2 | COMMD9 | Other | -0.083302024 | 0.181315623 | 25.42911707 | 25.75837125 | 25.75984471 | 25.44770982 | 25.37391496 | 25.52909974 | 25.4928336 | 25.72020208 | 25.18608703 |
| Catechol O-methyltransferase | P21964-2;P21964 | COMT | Extracellular or secreted | 1.025216927 | 4.020056962 | 26.55065873 | 26.67020132 | 26.77341592 | 26.50967704 | 26.41191449 | 27.16748706 | 27.79761107 | 27.65615063 | 27.66356636 |
| Coatomer subunit alpha;Xenin;Proxer | P53621;P53621-2 | COPA | Extracellular or secreted | -0.482755346 | 1.343203301 | 31.43688393 | 31.63330821 | 31.55201752 | 30.88823512 | 30.85102784 | 30.81084548 | 30.86960436 | 30.79040436 | 30.47788429 |
| Coatomer subunit beta | P53618 | COPB1 | Extracellular or secreted | -0.364881747 | 0.782751305 | 30.6563382 | 30.78034798 | 30.74342488 | 30.09622413 | 30.11743511 | 30.07839178 | 30.31083862 | 30.10562038 | 29.7249768 |
| Coatomer subunit beta | P35606;P35606-2 | COPB2 | Other | -0.485452744 | 2.201968911 | 30.84505641 | 30.88743639 | 30.82380779 | 30.57235577 | 30.66966083 | 30.32066209 | 30.34667651 | 30.18789522 | 30.06855968 |
| Coatomer subunit epsilon | O14579;O14579-3;O14579-2 | COPE | Other | -0.515264455 | 1.91692735 | 30.17414726 | 30.04544094 | 30.05413776 | 29.66173298 | 29.88547398 | 29.29721299 | 29.42638416 | 29.40173935 | 29.18515609 |
| Coatomer subunit gamma-1 | Q9Y678 | COPG1 | Other | -0.495569072 | 0.916637581 | 30.59903497 | 30.64013519 | 30.63183488 | 30.02867893 | 30.10898811 | 29.50517789 | 30.01836825 | 29.87802165 | 29.37382787 |
| Coatomer subunit gamma-2 | Q9UBF2;Q9UBF2-2 | COPG2 | Other | -0.411121841 | 0.982420397 | 28.49121333 | 28.53363641 | 28.4732073 | 27.89994737 | 27.96977426 | 27.80248806 | 28.06999398 | 27.74836025 | 27.53341363 |
| COP9 signalosome complex subunit 2 | P61201;P61201-2 | COPS2 | Other | -0.157067712 | 0.261240507 | 29.53421177 | 29.51470776 | 29.44436828 | 29.08154427 | 29.03742471 | 29.07681295 | 29.3290293 | 29.34008938 | 28.70421305 |
| COP9 signalosome complex subunit 3 | Q9UNS2;Q9UNS2-2 | COPS3 | Other | -0.042043877 | 0.059308803 | 29.30602124 | 29.31663285 | 29.18108502 | 28.65451268 | 28.63595642 | 28.86183281 | 29.14882352 | 29.16672119 | 28.53634416 |
| COP9 signalosome complex subunit 4 | Q9BT78;Q9BT78-2 | COPS4 | Extracellular or secreted | -0.248839891 | 0.513372081 | 29.79982011 | 29.7459107 | 29.68254347 | 29.58457483 | 29.58792192 | 29.13960779 | 29.5516784 | 29.49060144 | 28.9813899 |
| COP9 signalosome complex subunit 5 | Q92905 | COPS5 | Other | -0.332078023 | 1.116485237 | 29.05654284 | 29.22846184 | 29.06352839 | 28.989858 | 28.97281703 | 28.38841578 | 28.76695583 | 28.66729381 | 28.41932823 |
| COP9 signalosome complex subunit 6 | Q7L5N1 | COPS6 | Extracellular or secreted | -0.095663263 | 0.159990914 | 29.31324129 | 29.32568823 | 29.2783039 | 28.75429872 | 28.82841559 | 28.77687043 | 29.0741607 | 29.19898732 | 28.57827127 |
| COP9 signalosome complex subunit 7 | Q9UBW8 | COPS7A | Other | -0.047012336 | 0.134511087 | 26.54178139 | 26.53219515 | 26.29140596 | 26.69471453 | 26.47365633 | 26.4575278 | 26.55707286 | 26.22458952 | 26.57294119 |
| COP9 signalosome complex subunit 7 | Q9H9Q2;Q9H9Q2-2;Q9H9Q2 | COPS7B | Other | -0.48895762 | 1.727859838 | 27.8496566 | 27.64567668 | 27.56768562 | 27.14870233 | 27.07581893 | 27.27266481 | 27.12809693 | 26.87876715 | 26.80636554 |
| COP9 signalosome complex subunit 8 | Q99627-2;Q99627 | COPS8 | Extracellular or secreted | -0.015578243 | 0.020803399 | 28.10609483 | 27.72955426 | 27.93651933 | 27.42654412 | 27.63830631 | 27.00077664 | 27.74265421 | 27.88605572 | 27.24345309 |
| Coatomer subunit zeta-1 | P61923;P61923-4;P61923-5 | COPZ1 | Other | -0.670507619 | 1.449490645 | 27.69962053 | 27.48054456 | 27.48254617 | 27.30373714 | 27.53697401 | 27.05206567 | 26.95803668 | 26.88843837 | 26.41974613 |
| Ubiquinone biosynthesis protein COQ | O75208 | COQ9 | Other | -0.497136669 | 0.613102295 | 24.12226244 | 24.10229485 | 24.11195516 | 24.466066 | 24.56859912 | 24.29298604 | 24.26298717 | 23.87051298 | 23.20717165 |
| Coronin-1A | P31146 | CORO1A | Extracellular or secreted | -1.229938079 | 4.989358602 | 27.2051777 | 27.33005576 | 27.34375732 | 27.30773738 | 27.16605073 | 26.79563087 | 26.04214062 | 26.00795664 | 25.83429339 |
| Coronin-1B | Q9BR76 | CORO1B | Extracellular or secreted | 0.016710077 | 0.043274858 | 30.32656708 | 30.33511325 | 30.23375051 | 29.96705502 | 29.86618827 | 29.50248682 | 30.07022435 | 30.05840208 | 30.03708428 |
| Coronin-1C | Q9ULV4;Q9ULV4-2;Q9ULV4-3 | CORO1C | Extracellular or secreted | 0.13131206 | 0.770500038 | 29.91968515 | 29.78198985 | 29.76828172 | 29.65066643 | 29.34496379 | 29.80544327 | 29.81347786 | 29.86369472 | 29.8522787 |
| Coronin-7 | P57737-4;P57737;P57737-3;P | CORO7 | Other | -0.269120575 | 0.258710273 | 25.85102784 | 25.70387654 | 25.87790174 | 25.53433423 | 25.58614732 | 26.40525809 | 25.93759724 | 25.91868475 | 24.81562916 |
| Coactosin-like protein | Q14019 | COTL1 | Extracellular or secreted | -0.238312782 | 0.404854752 | 30.82244081 | 31.02617413 | 30.96739864 | 30.79474819 | 30.87927855 | 29.45124413 | 30.58844029 | 30.38740948 | 30.27985411 |
| Cytochrome c oxidase copper chaper | Q14061 | COX17 | Other | 0.415442091 | 0.294308613 | 22.72490266 | 22.96013772 | 23.0304597 | 21.48354259 | 20.68634803 | 21.86767223 | 22.8650279 | 23.08542253 | 21.6724073 |
| Cytochrome c oxidase subunit 4 isofo | P13073 | COX4I1 | Extracellular or secreted | -0.018824764 | 0.00432391 | 22.85369028 | 22.82320162 | 23.12420463 | 22.20457711 | 22.08323259 | 28.80621187 | 22.31837952 | 22.95519653 | 25.61750872 |
| Cytochrome c oxidase subunit 5A, mit | P20674 | COX5A | Extracellular or secreted | 0.0621572 | 0.05387098 | 23.18793013 | 23.48763006 | 22.68048057 | 22.04940471 | 23.3540707 | 23.27696494 | 23.68131735 | 22.55442645 | 22.96896836 |
| Cytochrome c oxidase subunit 6B1 | P14854 | COX6B1 | Other | 1.685626845 | 1.209268097 | 25.94552207 | 23.76880542 | 25.74224302 | 26.87829943 | 27.0753089 | 25.13864147 | 28.04702845 | 27.87882561 | 26.40543664 |
| Cytochrome c oxidase subunit 7A2, m | P14406 | COX7A2 | Other | 0.917429101 | 0.595056453 | 20.63404159 | 22.02653718 | 21.88644685 | 22.61235242 | 23.15120498 | 25.75582727 | 23.36852991 | 23.5818736 | 23.83508894 |
| Ceruloplasmin | P00450 | CP | Extracellular or secreted | -1.545591549 | 3.464434021 | 28.07081293 | 28.03354424 | 28.0296533 | 28.33896381 | 28.49331474 | 29.30114411 | 26.90173199 | 27.01488833 | 26.5803216 |
| Carboxypeptidase A4 | Q9UI42-2;Q9UI42 | CPA4 | Extracellular or secreted | 0.340552053 | 0.127487828 | 22.21717793 | 22.90686858 | 21.72863341 | 24.83393162 | 24.36866311 | 23.80823285 | 22.74861873 | 22.89363631 | 25.31115488 |
| Carboxypeptidase D | O75976;O75976-2 | CPD | Extracellular or secreted | -1.163689443 | 3.025996276 | 27.225454 | 27.27941136 | 27.22195611 | 27.43451968 | 27.33756093 | 27.73182138 | 26.11689995 | 26.07728678 | 26.43010667 |
| Carboxypeptidase E | P16870-2;P16870 | CPE | Extracellular or secreted | -2.819236855 | 1.387511553 | 24.88655582 | 24.44682686 | 24.9389435 | 24.87867362 | 25.07643071 | 27.8192766 | 21.66678745 | 21.83035737 | 24.06849818 |
| Copine-1 | Q99829 | CPNE1 | Extracellular or secreted | 0.008017296 | 0.041847846 | 28.45077234 | 28.38364504 | 28.42001127 | 28.48200755 | 28.42422294 | 28.24295361 | 28.41409421 | 28.50112042 | 28.31064364 |
| Copine-2 | Q96FN4;Q96FN4-2 | CPNE2 | Extracellular or secreted | -2.130806786 | 0.958476824 | 24.66638656 | 25.00581709 | 24.95211522 | 24.8721582 | 24.89933664 | 24.74709277 | 22.22055356 | 21.710765 | 24.24771431 |
| Copine-3 | O75131 | CPNE3 | Extracellular or secreted | 0.043798502 | 0.048977543 | 27.73679653 | 27.8607086 | 27.94050135 | 28.07127339 | 28.11152741 | 29.01643009 | 27.85887247 | 27.99004742 | 28.65109429 |
| Oxygen-dependent coproporphyrinog | P36551 | CPOX | Other | -0.106013956 | 0.469074324 | 29.25658163 | 29.17726387 | 29.17386143 | 29.18205649 | 29.38203753 | 29.03033761 | 29.05827304 | 29.2525947 | 28.97215968 |
| Serine/threonine-protein phosphatas | Q9BRF8;Q9BRF8-2 | CPPED1 | Extracellular or secreted | 0.079778509 | 0.225669397 | 28.30769396 | 28.32727406 | 28.37442915 | 28.75168369 | 28.66658295 | 27.84637225 | 28.55054131 | 28.3543482 | 28.47146405 |
| Cleavage and polyadenylation specific | Q10570 | CPSF1 | Other | -0.550357855 | 2.01703923 | 28.44059136 | 28.47622396 | 28.42818271 | 28.05133973 | 28.14301888 | 28.14019312 | 27.65082047 | 27.60935867 | 27.92852217 |
| Cleavage and polyadenylation specific | Q9P2I0 | CPSF2 | Other | 0.003996092 | 0.007762667 | 27.04082464 | 27.14676179 | 26.78494051 | 26.26099718 | 26.64196176 | 26.79983554 | 26.92184275 | 26.87537271 | 26.55243352 |
| Cleavage and polyadenylation specific | Q9UKF6 | CPSF3 | Other | -0.126868486 | 0.364572693 | 27.7612913 | 27.90351441 | 27.76357245 | 27.58125587 | 27.58462859 | 27.54598396 | 27.79457779 | 27.53660354 | 27.3583365 |
| Cleavage and polyadenylation specific | O95639-3;O95639-2;O95639 | CPSF4 | Other | -0.606875015 | 1.085942279 | 25.11735584 | 24.97031201 | 25.25511878 | 24.47673385 | 25.1178314 | 25.18835281 | 24.24001682 | 24.83951675 | 24.16269373 |
| Cleavage and polyadenylation specific | Q16630-3;Q16630;Q16630-2 | CPSF6 | Other | -0.520201138 | 4.068617174 | 29.36648864 | 29.15846238 | 29.23061628 | 29.02559346 | 29.22871412 | 29.01201305 | 28.58649096 | 28.69385094 | 28.66999866 |
| Cleavage and polyadenylation specific | Q8N684-2;Q8N684;Q8N684- | CPSF7 | Other | -0.687592532 | 1.453210801 | 26.5903513 | 26.49859676 | 26.52040311 | 26.6646517 | 26.59291914 | 26.50084698 | 25.8073732 | 25.67047148 | 26.14326222 |
| Carnitine O-palmitoyltransferase 1, liv | P50416;P50416-2 | CPT1A | Other | -1.90960998 | 0.750140327 | 24.91929872 | 24.93026053 | 25.89781444 | 25.6443019 | 25.06570735 | 28.56219974 | 22.07175825 | 24.3321065 | 25.37709665 |
| Carnitine O-palmitoyltransferase 2, m | P23786 | CPT2 | Other | 0.164058131 | 0.091757274 | 23.06731684 | 22.88671395 | 23.5731146 | 24.24655505 | 23.90875689 | 26.97125533 | 24.43466284 | 24.57427014 | 23.81009775 |
| Cellular retinoic acid-binding protein 2 | P29373 | CRABP2 | Extracellular or secreted | -2.50790477 | 2.328070365 | 24.75826958 | 24.55350321 | 24.90517984 | 24.51490712 | 24.44575397 | 23.96031451 | 21.65062055 | 22.57890611 | 21.81572315 |
| Carnitine O-acetyltransferase | P43155-2;P43155;P43155-3 | CRAT | Other | -0.805582653 | 1.235925607 | 27.32997025 | 27.29403846 | 27.29131813 | 27.16432525 | 27.36452798 | 28.5590295 | 26.56711983 | 26.34175701 | 27.17597998 |
| Protein cereblon | Q96SW2-2;Q96SW2 | CRBN | Other | -1.07351018 | 0.805972726 | 28.46060863 | 28.30630379 | 28.46964107 | 28.11728648 | 29.40541635 | 28.56985966 | 27.62561298 | 26.54215053 | 28.27626394 |
| Cyclic AMP-responsive element-bindi | P16220-3;P16220-2;P16220;( | CREB1 | Other | 0.660131754 | 1.048379011 | 25.87497422 | 25.73409786 | 25.79679457 | 25.14899318 | 24.5944007 | 25.56566808 | 26.40143827 | 26.29685867 | 25.63956262 |
| Protein CREG1 | O75629 | CREG1 | Extracellular or secreted | 0.246284221 | 0.291570368 | 26.1086492 | 26.1964615 | 26.59334667 | 26.87513832 | 27.07653265 | 25.39034427 | 27.0820267 | 26.54436336 | 26.2326989 |
| Cysteine-rich with EGF-like domain pr | Q6UXH1-4;Q6UXH1;Q6UXH1 | CRELD2 | Extracellular or secreted | -0.925668075 | 2.208311694 | 26.71437026 | 27.12119549 | 27.08618418 | 27.18820186 | 27.27408776 | 27.18366627 | 26.29944631 | 25.89968237 | 26.30772001 |
| Cysteine-rich motor neuron 1 protein | Q9NZV1 | CRIM1 | Extracellular or secreted | 0.73492572 | 0.305492486 | 24.3652625 | 23.4230369 | 24.92881581 | 26.63242439 | 26.59733084 | 27.96013772 | 25.56007973 | 25.75709982 | 27.84110168 |
| Cysteine-rich protein 2 | P52943;P52943-2 | CRIP2 | Other | -0.788719455 | 1.3665807 | 29.20531755 | 28.68348079 | 28.60798409 | 28.25696398 | 28.2994201 | 28.88366912 | 27.46256006 | 27.91649964 | 28.22319975 |
| Cysteine-rich secretory protein LCCL d | Q9H0B8-3;Q9H0B8-2;Q9H0B | CRISPLD2 | Extracellular or secreted | -1.405432217 | 0.758664133 | 22.49609307 | 21.86520709 | 21.93992402 | 23.09818762 | 22.87150973 | 25.12652237 | 20.25003601 | 21.56006582 | 22.67232347 |
| Adapter molecule crk | P46108 | CRK | Extracellular or secreted | -0.407263025 | 1.057046005 | 29.76121203 | 29.67969432 | 29.64650092 | 29.90339948 | 29.94391376 | 28.89238143 | 29.37705941 | 29.34144739 | 28.9732551 |
| Adapter molecule crk | P46108-2 | CRK | Extracellular or secreted | 0.496298928 | 0.142068828 | 22.79115966 | 21.33768031 | 22.16174425 | 22.07071784 | 23.26931676 | 22.20786715 | 24.07202005 | 23.91852553 | 20.41759419 |
| Crk-like protein | P46109 | CRKL | Extracellular or secreted | -0.084296119 | 0.234296945 | 29.19305237 | 29.05602595 | 28.9509091 | 29.10988484 | 29.10394607 | 28.38993417 | 28.88904793 | 29.02543505 | 28.73450492 |
| Cytokine receptor-like factor 1 | O75462 | CRLF1 | Extracellular or secreted | 0.145863952 | 0.054912636 | 22.81174078 | 19.97197008 | 19.04356136 | 22.16456694 | 21.19508379 | 21.74529846 | 20.03199766 | 22.53196226 | 21.33974265 |
| Cytokine receptor-like factor 3 | Q8IUI8-2;Q8IUI8 | CRLF3 | Extracellular or secreted | 0.69103927 | 1.51961119 | 26.89988977 | 26.90701533 | 26.85709334 | 26.96960961 | 27.02030382 | 26.83038155 | 27.5597589 | 27.85453942 | 27.40096619 |
| Dihydropyrimidinase-related protein 1 | Q14194;Q14194-2 | CRMP1 | Other | -0.073617924 | 0.20870915 | 27.03380676 | 27.03695332 | 26.83195239 | 26.77039431 | 26.5869204 | 26.61090791 | 26.85887247 | 26.83942064 | 26.51632066 |
| Crooked neck-like protein 1 | Q9BZJ0-2;Q9BZJ0-3;Q9BZJ0 | CRNKL1 | Other | -1.249122667 | 0.570150976 | 23.16175621 | 23.11513448 | 23.05872307 | 22.47674495 | 23.62555725 | 23.70872524 | 22.92909584 | 20.29096314 | 22.60589362 |
| Rootletin | Q5TZA2-2;Q5TZA2 | CROCC | Extracellular or secreted | 0.54065131 | 1.841440877 | 28.40202409 | 29.00415676 | 28.80384329 | 28.18517972 | 28.54277784 | 28.05325749 | 29.0585311 | 29.05260988 | 29.00643254 |
| Lambda-crystallin homolog | Q9Y2S2-2;Q9Y2S2 | CRYL1 | Extracellular or secreted | -0.880477864 | 1.099950202 | 27.19516642 | 27.18167746 | 27.32251134 | 26.96157351 | 26.83749705 | 26.11501538 | 26.413255 | 26.30997615 | 25.44205584 |
| Quinone oxidoreductase | Q08257;Q08257-3;Q08257-2 | CRYZ | Extracellular or secreted | -0.336750533 | 0.800954292 | 30.39815491 | 30.39815491 | 30.3075854 | 30.25282024 | 30.14734422 | 29.94391376 | 29.98820628 | 30.13246717 | 29.59306166 |
| Quinone oxidoreductase-like protein 1 | O95825;O95825-2 | CRYZL1 | Other | -0.072873444 | 0.219308584 | 27.4221387 | 27.17331343 | 26.9938846 | 27.03044286 | 26.79711627 | 26.65744603 | 26.98587441 | 27.04113808 | 26.79153812 |
| Citrate synthase, mitochondrial | O75390 | CS | Extracellular or secreted | -0.494257418 | 0.565019573 | 30.47121208 | 30.5549014 | 30.44124462 | 30.93220243 | 31.04329111 | 30.12780183 | 30.42981944 | 30.43708243 | 29.43565262 |
| Cold shock domain-containing protein | O75534-2;O75534;O75534-3 | CSDE1 | Extracellular or secreted | 0.775406098 | 1.905758088 | 27.37011094 | 27.16672119 | 27.38112994 | 28.34278294 | 28.30155843 | 27.88471738 | 28.57776724 | 28.63138391 | 28.34057757 |
| Exportin-2 | P55060;P55060-3;P55060-4 | CSE1L | Extracellular or secreted | 0.035658782 | 0.04317256 | 32.57389108 | 32.92291326 | 32.90433301 | 32.24343039 | 32.19095876 | 32.61883489 | 32.8433912 | 32.87067023 | 32.1200956 |
| Tyrosine-protein kinase CSK | P41240 | CSK | Extracellular or secreted | 0.254081393 | 0.521412725 | 24.35507951 | 24.50312404 | 24.59297615 | 23.62232088 | 24.16882634 | 23.73189904 | 24.60478576 | 24.5307078 | 24.11386359 |
| Casein kinase I isoform alpha | P48729;P48729-3;P48729-2 | CSNK1A1 | Other | -0.141824958 | 0.225651567 | 27.10918743 | 27.31172656 | 27.17245528 | 26.31103358 | 26.74239723 | 28.02289816 | 26.86266062 | 26.88623019 | 27.16048344 |
| Casein kinase II subunit alpha;Casein k | P68400;Q8NEV1;P68400-2 | CSNK2A1;CSNK2A | Extracellular or secreted | 0.583275411 | 1.354103016 | 27.58075288 | 27.62846657 | 27.69052461 | 27.8919181 | 28.09753074 | 28.72997557 | 28.72152599 | 28.61652753 | 28.22135694 |
| Casein kinase II subunit alpha | P19784 | CSNK2A2 | Other | 0.434518025 | 0.738194098 | 26.10880869 | 25.9700706 | 25.85993889 | 26.89133872 | 26.82323078 | 27.58089661 | 26.99690392 | 26.81311084 | 27.11068147 |
| Casein kinase II subunit beta | P67870 | CSNK2B | Extracellular or secreted | 0.379759754 | 0.748089049 | 27.03873333 | 27.16106036 | 27.0431216 | 27.26990384 | 27.57499189 | 28.2714631 | 28.04369526 | 27.78998479 | 27.48523626 |
| Cysteine and glycine-rich protein 1 | P21291 | CSRP1 | Extracellular or secreted | 0.327841149 | 0.668415032 | 27.40365012 | 27.14287286 | 27.19601117 | 27.25664911 | 27.09687758 | 26.93438358 | 27.67640241 | 27.6969044 | 27.12543885 |
| Cystatin-C | P01034 | CST3 | Extracellular or secreted | 0.503590588 | 0.535104291 | 29.49304746 | 29.39353906 | 29.27861407 | 30.68912526 | 30.63728734 | 31.05517269 | 30.18245911 | 30.43370421 | 31.16800139 |
| Cystatin-A;Cystatin-A, N-terminally pr | P01040 | CSTA | Extracellular or secreted | -1.640892093 | 1.25288278 | 24.58594682 | 23.4683675 | 24.17812867 | 23.14010703 | 24.02504424 | 21.26590161 | 21.38501046 | 21.25341936 | 22.77064183 |
| Cystatin-B | P04080 | CSTB | Extracellular or secreted | 0.219983426 | 0.60348017 | 31.23574874 | 31.23180672 | 31.20250625 | 31.23437882 | 31.20396508 | 30.22031932 | 31.32715626 | 31.28499111 | 31.21216537 |
| Cleavage stimulation factor subunit 1 | Q05048 | CSTF1 | Other | -0.008974797 | 0.01053349 | 29.14688315 | 29.28341329 | 29.27949992 | 28.73595777 | 28.81735644 | 29.10831519 | 29.34545032 | 29.26742784 | 28.54591033 |
| Cleavage stimulation factor subunit 2 | P33240-2;P33240 | CSTF2 | Other | 0.080807784 | 0.246931721 | 28.5911364 | 28.59167146 | 28.6284318 | 28.75865717 | 28.91139386 | 28.19905755 | 28.73560276 | 28.82738617 | 28.51960854 |
| Cleavage stimulation factor subunit 2 | Q9H0L4 | CSTF2T | Other | -0.305140439 | 0.568825453 | 24.46494503 | 24.62488825 | 24.58250541 | 24.33633548 | 24.46942371 | 24.41338414 | 24.27845456 | 24.46699948 | 23.78486565 |
| Cleavage stimulation factor subunit 3 | Q12996 | CSTF3 | Other | -0.472383095 | 0.810699976 | 28.21220017 | 28.20167698 | 28.28065072 | 27.88849643 | 27.81671582 | 27.97629054 | 27.79457779 | 27.82777986 | 27.14850839 |
| C-terminal-binding protein 1 | Q13363-2;Q13363 | CTBP1 | Other | -0.141397473 | 0.264290936 | 26.20929301 | 26.44317892 | 26.44892301 | 26.92286365 | 26.80550478 | 27.2266487 | 26.4592645 | 26.30917823 | 26.83557089 |
| C-terminal-binding protein 2 | P56545;P56545-3;P56545-2 | CTBP2 | Other | 0.983775541 | 0.644948997 | 24.77560265 | 25.30740735 | 25.30445104 | 24.69736849 | 21.21490294 | 26.25562307 | 25.50157603 | 25.68062007 | 25.54680829 |
| Di-N-acetylchitobiase | Q01459 | CTBS | Extracellular or secreted | -0.78761759 | 0.772258856 | 22.98877621 | 23.05599597 | 23.23950705 | 23.48763006 | 23.78676098 | 22.22652744 | 22.0814074 | 21.83402314 | 23.11431554 |
| Transcriptional repressor CTCF | P49711;P49711-2 | CTCF | Other | -0.614848964 | 0.222829357 | 25.14433243 | 24.85590603 | 25.01448401 | 24.85998627 | 24.59974444 | 25.60016995 | 25.40197528 | 25.37049346 | 22.42029593 |
| RNA polymerase II subunit A C-termin | Q9Y5B0;Q9Y5B0-4 | CTDP1 | Other | 0.490080763 | 1.387127108 | 22.72862306 | 22.27127774 | 21.67277429 | 22.44417979 | 22.63291101 | 22.91767 | 23.04341784 | 22.85585852 | 22.90468386 |
| CTD small phosphatase-like protein | O15194-2;O15194 | CTDSPL | Extracellular or secreted | 0.991856764 | 1.652039951 | 23.31962031 | 22.18848864 | 21.9442384 | 22.74507864 | 22.48079414 | 21.39030214 | 23.4693616 | 22.94419087 | 23.59627895 |
| Catenin alpha-1 | P35221;P35221-2;P35221-3 | CTNNA1 | Extracellular or secreted | -0.283462102 | 0.710118742 | 32.30081147 | 32.22317098 | 32.31224608 | 31.94994264 | 31.9838649 | 32.11619601 | 31.9395419 | 32.08340963 | 31.5697782 |
| Catenin alpha-2 | P26232-2;P26232-5;P26232;F | CTNNA2 | Extracellular or secreted | -1.514461674 | 2.275750545 | 28.8969777 | 28.61097829 | 28.59249148 | 29.14299454 | 29.27050547 | 28.61582627 | 27.54333113 | 27.5528146 | 26.92535613 |
| Catenin beta-1 | P35222 | CTNNB1 | Extracellular or secreted | 0.539504589 | 1.393115759 | 31.82612111 | 31.83269944 | 31.77214982 | 30.84961186 | 31.03183668 | 31.95684658 | 32.10487093 | 32.21656613 | 31.93170945 |
| Beta-catenin-like protein 1 | Q8WYA6;Q8WYA6-4;Q8WYA | CTNNBL1 | Other | 0.306210607 | 0.379942716 | 25.93244744 | 25.96892883 | 25.84878548 | 26.22751195 | 26.27241565 | 25.82021442 | 26.75595457 | 26.45502075 | 25.74280838 |
| Catenin delta-1 | O60716-21;O60716-19;O607 | CTNND1 | Extracellular or secreted | 1.631166535 | 4.456431319 | 29.1969492 | 29.06363125 | 29.37318484 | 29.05333518 | 29.0976312 | 29.88951218 | 30.89929918 | 31.09578414 | 30.73553821 |
| CTP synthase 1 | P17812;P17812-2 | CTPS1 | Other | 0.511034561 | 0.943881096 | 29.6747873 | 29.70162123 | 29.63718368 | 29.64783931 | 29.71199351 | 29.34379967 | 30.22054996 | 30.42031252 | 29.75085355 |
| CTP synthase 2 | Q9NRF8 | CTPS2 | Other | 0.056692269 | 0.08874392 | 27.17207372 | 27.30059876 | 27.24445152 | 26.71161615 | 26.79711627 | 26.91752432 | 27.33807123 | 27.21363764 | 26.69005831 |
| RNA polymerase-associated protein C | Q6PD62 | CTR9 | Other | 0.07343762 | 0.114897514 | 25.6098377 | 25.58789332 | 25.34431623 | 25.2545062 | 25.18517972 | 24.9449415 | 25.7525644 | 25.3807007 | 25.05038509 |
| Lysosomal protective protein;Lysoson | P10619-2;P10619 | CTSA | Extracellular or secreted | -0.109891705 | 0.401613356 | 28.37832105 | 28.62097257 | 28.27017126 | 28.57535262 | 28.62170626 | 27.97180349 | 28.25552403 | 28.21572176 | 28.41824272 |
| Cathepsin B;Cathepsin B light chain;C | P07858 | CTSB | Extracellular or secreted | 0.106470908 | 0.197353626 | 26.93415859 | 26.47299048 | 26.44423793 | 26.85697465 | 27.01211965 | 27.82638633 | 26.94134121 | 27.01977378 | 27.13173154 |
| Dipeptidyl peptidase 1;Dipeptidyl pep | P53634 | CTSC | Extracellular or secreted | -0.050671656 | 0.088183254 | 30.19964266 | 30.25146645 | 30.0768639 | 30.36306617 | 30.50332121 | 29.11010894 | 30.06753431 | 30.0805535 | 29.95213189 |
| Cathepsin D;Cathepsin D light chain;C | P07339 | CTSD | Extracellular or secreted | 1.045679527 | 3.794423446 | 30.04035436 | 30.13858289 | 30.04791253 | 30.51455729 | 30.4027357 | 30.64400959 | 31.33122034 | 31.33474041 | 31.36515401 |
| Pro-cathepsin H;Cathepsin H mini cha | P09668 | CTSH | Extracellular or secreted | -0.660833412 | 0.505985946 | 27.38541534 | 27.51780789 | 27.42654412 | 27.6314533 | 27.43340575 | 27.70404811 | 26.40631285 | 26.31948249 | 27.84104168 |
| Cathepsin L1;Cathepsin L1 heavy chai | P07711 | CTSL | Extracellular or secreted | 0.133501912 | 0.046778435 | 24.99640832 | 25.91024471 | 25.41328729 | 25.49619062 | 25.6899517 | 24.96468318 | 26.55127502 | 23.72327847 | 26.36133516 |
| Cathepsin L2 | O60911 | CTSV | Extracellular or secreted | 0.992506559 | 1.715531642 | 28.58290734 | 28.5376406 | 28.44671645 | 29.11897013 | 28.97465055 | 29.7945468 | 29.5762 | 29.80977279 | 30.31926283 |
| Cathepsin Z | Q9UBR2 | CTSZ | Extracellular or secreted | -0.099046898 | 0.082562138 | 28.09652576 | 28.2640975 | 28.1257837 | 28.35812687 | 28.47961981 | 28.65403459 | 27.93494593 | 27.74720804 | 29.00979945 |
| Src substrate cortactin | Q14247;Q14247-3;Q14247-2 | CTTN | Extracellular or secreted | 2.15703077 | 5.040604675 | 29.24177234 | 29.11562566 | 29.25342904 | 29.65222326 | 29.51453847 | 29.26463445 | 31.57947664 | 31.59934541 | 31.31338187 |
| Cytoplasmic tRNA 2-thiolation protein | Q7Z7A3 | CTU1 | Other | 0.133096047 | 0.241024575 | 22.75125719 | 22.88663954 | 23.2369583 | 22.33921108 | 22.78504531 | 22.52207478 | 23.07177461 | 23.05743454 | 22.53067209 |
| Cullin-1 | Q13616 | CUL1 | Other | 0.379212421 | 0.775574769 | 29.28332497 | 29.34625381 | 29.22250897 | 28.60052445 | 28.48998928 | 29.23095978 | 29.54124598 | 29.62639654 | 29.05677537 |
| Cullin-2 | Q13617;Q13617-2 | CUL2 | Other | -0.119058675 | 0.157831262 | 29.37175255 | 29.49184408 | 29.4078895 | 29.0115866 | 28.98277751 | 29.22832421 | 29.39224971 | 29.37976749 | 28.617894 |
| Cullin-3 | Q13618-2;Q13618;Q13618-3 | CUL3 | Extracellular or secreted | -0.050412989 | 0.057629501 | 28.71155051 | 28.76594481 | 28.75865717 | 28.32324179 | 28.2084374 | 28.49609917 | 28.72728358 | 28.80768642 | 27.94575646 |
| Cullin-4A | Q13619;Q13619-2 | CUL4A | Other | 0.007005237 | 0.013853105 | 27.4270239 | 27.67855308 | 27.48577367 | 27.01595179 | 27.24454226 | 27.62135693 | 27.47049467 | 27.62212535 | 27.16499651 |

| | | | | | | | | | | | | | |
|---|---|---|---|---|---|---|---|---|---|---|---|---|---|
| Cullin-4B | Q13620-1;Q13620;Q13620-3 | CUL4B | Extracellular or secreted | 0.075737447 | 0.161687013 | 28.63030796 | 28.8738778 | 28.83318973 | 28.33133781 | 28.38681358 | 28.92249507 | 28.83448629 | 28.92331715 | 28.45841988 |
| Cullin-5 | Q93034 | CUL5 | Other | -0.796753744 | 0.491663115 | 27.09113721 | 27.0012063 | 27.57159654 | 26.80783993 | 26.56459284 | 26.81677684 | 26.83050244 | 26.75289612 | 24.95291504 |
| Protein CutA | O60888-3;O60888;O60888-2 | CUTA | Extracellular or secreted | -0.180299355 | 0.480878368 | 28.61617695 | 28.61037995 | 28.62446299 | 29.11286993 | 29.03527601 | 28.01222623 | 28.59416574 | 28.47877159 | 28.39186063 |
| Copper homeostasis protein cutC hom | Q9NTM9 | CUTC | Other | -0.153980609 | 0.271142142 | 23.9295835 | 24.07447208 | 24.22309846 | 24.0661181 | 24.74201167 | 24.02555122 | 24.30361522 | 23.6575278 | 24.10733267 |
| Homeobox protein cut-like 1 | P39880-3;P39880-5;P39880-2 | CUX1 | Other | -2.486851979 | 3.651149823 | 26.59946069 | 27.3967268 | 27.10499587 | 26.01305733 | 26.04264163 | 26.76116447 | 24.29796073 | 23.7432451 | 24.45726163 |
| CUB and zona pellucida-like domain-c | Q86UP6-2;Q86UP6;Q86UP6- | CUZD1 | Other | -0.329938342 | 0.05990489 | 25.39659616 | 20.66282095 | 25.78917641 | 26.15992552 | 25.98791365 | 26.5329531 | 26.23471001 | 26.58118403 | 21.45898383 |
| Spliceosome-associated protein CWC | Q9P013 | CWC15 | Other | 1.188757512 | 3.764432173 | 25.33118403 | 24.63782282 | 24.97517555 | 25.22828292 | 25.21874532 | 24.50275995 | 26.24984925 | 26.2642228 | 25.99918578 |
| Pre-mRNA-splicing factor CWC22 hom | Q9HCG8 | CWC22 | Other | -1.772548148 | 2.561238338 | 23.1911725 | 23.50385194 | 23.56883102 | 23.50651776 | 23.10666197 | 21.5978707 | 21.77334046 | 21.02237692 | 21.12409112 |
| Peptidyl-prolyl cis-trans isomerase CW | Q6UX04;Q6UX04-2 | CWC27 | Other | -0.519681194 | 0.840043432 | 25.21152266 | 25.52629625 | 25.71203616 | 24.94637017 | 25.17096661 | 25.45170791 | 24.87680169 | 25.19531663 | 24.37828798 |
| CWF19-like protein 1 | Q69YN2;Q69YN2-3 | CWF19L1 | Other | 0.056533683 | 0.062062567 | 26.53336907 | 27.0581698 | 26.67774695 | 25.4822538 | 26.1959361 | 26.06463884 | 26.84050154 | 26.26760196 | 26.06755483 |
| Fractalkine;Processed fractalkine | P78423 | CX3CL1 | Extracellular or secreted | 1.320964698 | 0.585324249 | 23.10352794 | 23.36706382 | 23.34340153 | 23.54403903 | 23.45458801 | 25.455789 | 24.11195516 | 24.23454557 | 26.75059802 |
| Coxsackievirus and adenovirus recept | P78310-2;P78310-6;P78310;P | CXADR | Extracellular or secreted | 0.450165504 | 0.842641783 | 26.67384432 | 26.42438314 | 26.56012347 | 26.4363955 | 26.80488963 | 26.58491526 | 27.27737297 | 27.18006546 | 26.63533375 |
| Growth-regulated alpha protein;GRO- | P09341 | CXCL1 | Extracellular or secreted | 2.226677268 | 1.423777922 | 24.72754326 | 24.20858624 | 24.25010234 | 24.44588023 | 24.40665345 | 26.70490562 | 26.82383815 | 26.09683738 | 28.13119184 |
| C-X-C motif chemokine 16 | Q9H2A7 | CXCL16 | Extracellular or secreted | 0.319275231 | 0.086253012 | 20.70062713 | 20.45480537 | 23.56117832 | 24.79872373 | 25.00328834 | 25.0520864 | 21.66533915 | 24.03716285 | 25.04067835 |
| Interleukin-8;MDNCF-a;Interleukin-8; | P10145 | CXCL8 | Extracellular or secreted | 0.985126199 | 0.714133524 | 24.71889655 | 24.60715871 | 24.57444339 | 23.84632442 | 23.8781123 | 23.52336765 | 24.49953985 | 24.81372249 | 26.21626777 |
| Cytochrome b5 | P00167-2;P00167;P00167-3 | CYB5A | Extracellular or secreted | 0.787070603 | 1.116301647 | 25.98873721 | 25.83004299 | 26.50453401 | 26.19475325 | 26.50972234 | 24.67066057 | 27.00849103 | 26.9560434 | 26.24590256 |
| Cytochrome b5 type B | O43169 | CYB5B | Other | -0.592981491 | 0.401592991 | 26.98054579 | 27.34078977 | 27.52678875 | 26.44145439 | 26.7948876 | 28.55266804 | 25.83388338 | 26.54414223 | 27.66159708 |
| NADH-cytochrome b5 reductase 3;NA | P00387-2;P00387;P00387-3 | CYB5R3 | Extracellular or secreted | 0.409867752 | 0.238198441 | 25.22523333 | 24.86273156 | 25.56764211 | 25.58003401 | 25.02947428 | 28.87352583 | 25.90612083 | 25.94888923 | 26.94391376 |
| Cytochrome c1, heme protein, mitoch | P08574 | CYC1 | Other | -0.887743729 | 0.292221862 | 21.30186725 | 23.3937518 | 24.1828332 | 24.17911638 | 23.89864493 | 26.71463228 | 21.09459431 | 23.64669318 | 24.43090424 |
| Cytochrome c | P99999 | CYCS | Other | 0.018700431 | 0.01941813 | 31.9584032 | 32.24903771 | 32.12869309 | 32.08248994 | 32.154122 | 31.02966647 | 32.24872677 | 32.17911044 | 31.42947029 |
| Cytoplasmic FMR1-interacting protein | Q7L576;Q7L576-2 | CYFIP1 | Extracellular or secreted | 0.122160362 | 0.374543004 | 29.78128642 | 29.80241103 | 29.84876161 | 29.14165541 | 29.20515439 | 29.60068394 | 29.70272786 | 29.78506527 | 29.56866434 |
| Cytoplasmic FMR1-interacting protein | Q96F07-2;Q96F07 | CYFIP2 | Extracellular or secreted | 0.689821868 | 0.397244536 | 24.67583732 | 25.27277158 | 24.95118154 | 23.76971357 | 20.45132294 | 24.99438106 | 24.95628717 | 24.63793335 | 24.53284909 |
| Cysteine and histidine-rich protein 1 | Q6ZMK1-2;Q6ZMK1;Q6ZMK1 | CYHR1 | Extracellular or secreted | 2.303951759 | 1.4661649 | 25.30997615 | 25.42895739 | 25.20825133 | 21.77172632 | 21.57963704 | 21.8599776 | 25.53180858 | 26.11967273 | 25.83963689 |
| Lanosterol 14-alpha demethylase | Q16850;Q16850-2 | CYP51A1 | Extracellular or secreted | 2.501546563 | 0.775099199 | 21.47333491 | 20.62144366 | 22.88353106 | 20.27291524 | 21.47056898 | 25.0603981 | 22.79378189 | 23.71607255 | 26.88588122 |
| Protein CYR61 | O00622 | CYR61 | Extracellular or secreted | -0.918367167 | 0.457778473 | 22.07642503 | 23.33497265 | 21.7365763 | 25.06447438 | 25.58477193 | 24.23972554 | 23.29010549 | 23.47877159 | 21.49449433 |
| Dystroglycan;Alpha-dystroglycan;Beta | Q14118 | DAG1 | Extracellular or secreted | 0.905455906 | 1.325265648 | 29.81491446 | 29.93536754 | 29.96526687 | 30.5171884 | 30.68970849 | 30.89345231 | 30.86297093 | 31.03597076 | 31.72537506 |
| Bifunctional ATP-dependent dihydrox | Q3LXA3;Q3LXA3-2 | DAK | Extracellular or secreted | -0.065875317 | 0.047797003 | 29.72177043 | 29.82282066 | 29.80532026 | 28.75318313 | 28.85768663 | 28.52118227 | 29.58790404 | 29.57419795 | 28.38125374 |
| Death-associated protein 1 | P51397 | DAP | Other | -0.417359349 | 0.26849006 | 26.48123774 | 26.23328424 | 26.48434524 | 25.96380168 | 25.730319 | 23.35097262 | 25.71287582 | 25.70780418 | 24.44922221 |
| Aspartate--tRNA ligase, cytoplasmic | P14868;P14868-2 | DARS | Extracellular or secreted | -0.457292566 | 1.913468529 | 29.11916808 | 29.2583351 | 29.24029439 | 28.84625267 | 28.81589175 | 28.80565852 | 28.58534517 | 28.69634066 | 28.38923673 |
| Aspartate--tRNA ligase, mitochondrial | Q6PI48 | DARS2 | Other | -0.062938482 | 0.043802954 | 25.17698817 | 24.97355619 | 25.41041098 | 26.05136048 | 26.15516488 | 27.52447404 | 25.63023157 | 25.37085925 | 26.45607111 |
| Death domain-associated protein 6 | Q9UER7;Q9UER7-3;Q9UER7- | DAXX | Other | -0.01575817 | 0.016923122 | 26.26162495 | 25.01818252 | 25.32034361 | 25.09936804 | 24.96494752 | 24.40399135 | 24.93400107 | 25.55039452 | 25.00255889 |
| DAZ-associated protein 1 | Q96EP5-2;Q96EP5 | DAZAP1 | Other | 0.12216086 | 0.461607401 | 29.29533101 | 29.18860277 | 29.1068187 | 29.25891911 | 29.47221972 | 29.41667333 | 29.52785152 | 29.48805546 | 29.21985791 |
| Acyl-CoA-binding protein | P07108;P07108-3;P07108-2;P | DBI | Extracellular or secreted | 0.348233448 | 0.233993204 | 26.49868806 | 25.68477194 | 26.06597435 | 27.3800566 | 27.40827428 | 24.16338495 | 26.98945202 | 26.79389597 | 25.86192745 |
| Drebrin | Q16643;Q16643-3;Q16643-2 | DBN1 | Extracellular or secreted | 1.540851584 | 3.987240016 | 29.06650821 | 28.91316439 | 29.00803681 | 29.12551276 | 29.2950025 | 28.7206457 | 30.73141679 | 30.56529038 | 30.39028276 |
| Drebrin-like protein | Q9UJU6;Q9UJU6-5 | DBNL | Extracellular or secreted | 0.75378299 | 4.361645623 | 25.91248499 | 26.10952621 | 26.07332832 | 26.08405624 | 26.15383217 | 26.39974456 | 26.84278077 | 26.90150184 | 26.8835526 |
| Drebrin-like protein | Q9UJU6-2;Q9UJU6-3;Q9UJU6 | DBNL | Extracellular or secreted | -0.121958926 | 0.088675021 | 22.09287583 | 22.19593244 | 21.36667679 | 21.93905115 | 21.86964462 | 22.15196044 | 20.92501804 | 22.14030431 | 22.37687151 |
| Lariat debranching enzyme | Q9UK59 | DBR1 | Other | -0.071077046 | 0.064110108 | 27.99377665 | 27.89405975 | 27.87595853 | 27.42989923 | 27.41207606 | 26.84326015 | 27.78487812 | 27.89035326 | 26.83605267 |
| DDB1- and CUL4-associated factor 6 | Q58WW2-4;Q58WW2;Q58W | DCAF6 | Other | 0.957861845 | 0.46099817 | 23.6292312 | 23.25767443 | 23.10631053 | 20.77258213 | 21.81035166 | 23.04591893 | 24.35009587 | 24.28897982 | 22.04554429 |
| DDB1- and CUL4-associated factor 7 | P61962 | DCAF7 | Other | -0.112399468 | 0.231131137 | 26.1253206 | 26.02529775 | 25.91326142 | 25.13473056 | 25.42294069 | 26.4171403 | 25.71347902 | 25.74893601 | 25.71973223 |
| DDB1- and CUL4-associated factor 8 | Q5TAQ9;Q5TAQ9-2 | DCAF8 | Other | 1.233538987 | 3.303325756 | 24.85167128 | 25.05287414 | 25.02757735 | 25.11008404 | 24.43999722 | 25.77459767 | 26.40932684 | 26.29342464 | 26.12626634 |
| Discoidin, CUB and LCCL domain-cont | Q8N8Z6-2;Q8N8Z6 | DCBLD1 | Other | 0.863895194 | 0.499624413 | 22.16481243 | 22.01440738 | 21.64306348 | 21.88874028 | 21.94867094 | 23.50045193 | 22.08670252 | 22.78835512 | 24.29670116 |
| Discoidin, CUB and LCCL domain-cont | Q96PD2;Q96PD2-2 | DCBLD2 | Extracellular or secreted | 1.042606845 | 0.773658784 | 25.8515998 | 26.0452274 | 26.18143103 | 26.91854828 | 26.75085355 | 27.44561191 | 27.21771044 | 26.91090802 | 28.59583806 |
| Dermcidin;Survival-promoting peptide | P81605;P81605-2 | DCD | Extracellular or secreted | -3.069204195 | 1.51308808 | 26.56874406 | 26.3107563 | 26.98174373 | 27.35758167 | 27.22756703 | 26.83532993 | 24.72832205 | 24.05763285 | 22.64729388 |
| Deoxycytidine kinase | P27707 | DCK | Other | -0.052616036 | 0.100757605 | 27.80421267 | 27.66912014 | 27.58742125 | 27.10739254 | 27.62261412 | 26.69205567 | 27.49373466 | 27.381295 | 27.20853043 |
| Serine/threonine-protein kinase DCLK | O15075-2;O15075 | DCLK1 | Extracellular or secreted | -0.932278767 | 0.379498202 | 23.27099104 | 22.92368438 | 23.35393614 | 21.53365324 | 22.60713613 | 23.07448841 | 23.07625128 | 22.43644316 | 20.07241393 |
| mRNA-decapping enzyme 1A | Q9NPI6-2;Q9NPI6 | DCP1A | Other | 0.687072331 | 1.215358429 | 22.65661165 | 23.39047548 | 22.60025504 | 21.66172079 | 22.43229243 | 22.32248461 | 23.50796977 | 23.25537095 | 22.82979627 |
| m7GpppX diphosphatase | Q96C86 | DCPS | Other | 0.055014503 | 0.066222946 | 29.39140999 | 29.47032011 | 29.26136491 | 28.58398336 | 28.46517085 | 28.054086 | 29.08428438 | 29.12388605 | 28.57004069 |
| Deoxycytidylate deaminase | P32321;P32321-2 | DCTD | Extracellular or secreted | 0.387548125 | 2.117909942 | 26.50413996 | 26.84122168 | 26.87114778 | 26.49307801 | 26.42122389 | 26.31544563 | 26.93988513 | 27.02654354 | 26.91934419 |
| Dynactin subunit 1 | Q14203-3;Q14203-4;Q14203 | DCTN1 | Other | 0.07020603 | 0.12597208 | 30.0353809 | 30.17367085 | 30.14187463 | 29.55217331 | 29.65220616 | 29.57383692 | 30.11916808 | 30.06162425 | 29.59439714 |
| Dynactin subunit 2 | Q13561;Q13561-3;Q13561-2 | DCTN2 | Extracellular or secreted | 0.539190338 | 1.056350624 | 28.9517985 | 28.92368551 | 28.87032483 | 28.90738788 | 28.97801052 | 28.18791881 | 29.55420614 | 29.52494102 | 28.94798689 |
| Dynactin subunit 3 | O75935-3;O75935 | DCTN3 | Other | 0.158371279 | 0.750981391 | 25.31951695 | 25.33114985 | 25.08517127 | 25.6131865 | 25.60631168 | 25.41212453 | 25.683889 | 25.46995153 | 25.50500369 |
| Dynactin subunit 4 | Q9UJW0;Q9UJW0-3;Q9UJW0 | DCTN4 | Other | 0.108000437 | 0.468496479 | 26.32753962 | 26.26874444 | 26.22009325 | 25.78291785 | 25.91902588 | 25.91567937 | 26.17131029 | 26.27150762 | 26.0981836 |
| Dynactin subunit 5 | Q9BTE1;Q9BTE1-3;Q9BTE1-2 | DCTN5 | Other | 0.139992677 | 0.244589096 | 23.16498117 | 23.21319264 | 23.05885516 | 22.41020392 | 22.80274196 | 22.41175613 | 23.19684042 | 23.11778386 | 22.63621924 |
| Dynactin subunit 6 | O00399 | DCTN6 | Other | 1.627639479 | 1.119919298 | 24.30382422 | 23.21083239 | 22.84812165 | 19.39394429 | 22.6879855 | 23.23341956 | 24.55162707 | 24.67648311 | 23.49387206 |
| dCTP pyrophosphatase 1 | Q9H773 | DCTPP1 | Other | 0.550417139 | 2.745432645 | 30.04322592 | 30.11458357 | 29.98141712 | 30.27275378 | 30.40517286 | 29.70834702 | 30.70972785 | 30.5862046 | 30.6180691 |
| DCN1-like protein 1 | Q96GG9 | DCUN1D1 | Other | -0.115126543 | 2.025588521 | 28.5280938 | 28.57054743 | 28.52682605 | 28.57354804 | 28.4646257 | 28.546168 | 28.38471571 | 28.45708938 | 28.41771978 |
| DCN1-like protein 4 | Q92564-2;Q92564;Q92564-3 | DCUN1D4 | Other | 0.01975554 | 0.016195615 | 22.51463027 | 22.12571867 | 22.70895672 | 21.71644227 | 20.54156247 | 23.14871578 | 22.10126913 | 22.47251335 | 21.86349722 |
| DCN1-like protein 5 | Q9BTE7 | DCUN1D5 | Other | 0.415972859 | 0.942335533 | 21.40556323 | 21.96095078 | 22.08258308 | 21.49045174 | 21.64632957 | 22.49572721 | 22.19883468 | 22.55109896 | 22.03878774 |
| L-xylulose reductase | Q7Z4W1 | DCXR | Extracellular or secreted | -0.586047041 | 1.386125492 | 26.62867514 | 26.56794668 | 26.63131452 | 26.8811035 | 26.71960168 | 27.17997058 | 25.89610583 | 26.20504716 | 26.44501194 |
| DET1- and DDB1-associated protein 1 | Q9BW61 | DDA1 | Other | 1.809121973 | 3.177881305 | 21.44052645 | 21.53684464 | 22.61024898 | 22.7215076 | 22.9412203 | 22.01672292 | 24.13617884 | 24.03691141 | 23.88781112 |
| N(G),N(G)-dimethylarginine dimethyla | O94760;O94760-2 | DDAH1 | Extracellular or secreted | -0.395332416 | 1.386705684 | 29.51265614 | 29.35070712 | 29.34049268 | 29.41324693 | 29.65517832 | 28.66729381 | 28.8003294 | 28.93331453 | 29.05014633 |
| N(G),N(G)-dimethylarginine dimethyla | O95865 | DDAH2 | Extracellular or secreted | -0.568352385 | 1.621080989 | 30.1106068 | 30.12275119 | 30.07048029 | 29.55101826 | 29.53493528 | 29.04265206 | 29.11040768 | 29.31239757 | 29.08835954 |
| DNA damage-binding protein 1 | Q16531 | DDB1 | Extracellular or secreted | -0.169400649 | 0.429480495 | 32.82466149 | 33.08226786 | 32.94342477 | 33.21146929 | 33.33218121 | 32.46677103 | 33.00110225 | 32.83855309 | 32.58253054 |
| DNA damage-binding protein 2 | Q92466;Q92466-4;Q92466-5 | DDB2 | Other | -0.638043785 | 1.999291228 | 28.53449008 | 28.64646659 | 28.42414283 | 27.8275376 | 27.95798135 | 27.7222103 | 27.62686647 | 27.62261412 | 27.39280243 |
| Phospholipase DDHD2 | O94830;O94830-2 | DDHD2 | Other | 0.170183161 | 0.14865815 | 24.18184804 | 23.86542443 | 23.85052719 | 21.82898193 | 22.53146539 | 23.99765776 | 23.69720939 | 23.78725933 | 23.15403312 |
| Protein DDI1 homolog 2 | Q5TDH0;Q5TDH0-3;Q5TDH0- | DDI2 | Other | 0.136535066 | 0.190836028 | 26.74033973 | 26.92161579 | 26.92694002 | 26.89168638 | 27.07999431 | 25.35840358 | 26.9563758 | 26.8075943 | 26.60512499 |
| Dolichyl-diphosphooligosaccharide--p | P39656;P39656-3;P39656-2 | DDOST | Extracellular or secreted | -0.364821482 | 0.105740272 | 25.74170315 | 25.79691831 | 26.36401025 | 21.64882685 | 23.52145188 | 29.04629997 | 23.9680719 | 24.53943147 | 26.45763738 |
| Epithelial discoidin domain-containing | Q08345;Q08345-5;Q08345-2 | DDR1 | Extracellular or secreted | 0.566906795 | 0.785526806 | 28.58598979 | 28.50058869 | 28.30334521 | 28.36277364 | 28.55354715 | 29.00209207 | 28.80322743 | 28.88602664 | 29.66563459 |
| DDRGK domain-containing protein 1 | Q96HY6 | DDRGK1 | Other | 2.135345176 | 3.749545318 | 21.86733003 | 21.72795867 | 22.51842937 | 22.03015656 | 23.06092235 | 23.14499362 | 24.35104004 | 24.38442753 | 24.84546325 |
| D-dopachrome decarboxylase;D-dopa | P30046;P30046-2;A6NHG4 | DDT;DDTL | Extracellular or secreted | 0.385397157 | 0.967649722 | 29.56309125 | 29.89792982 | 29.83413965 | 29.49882118 | 29.55528548 | 29.14425949 | 29.90081117 | 30.25979469 | 29.74234904 |
| ATP-dependent RNA helicase DDX1 | Q92499;Q92499-3;Q92499-2 | DDX1 | Other | -0.073392551 | 0.324614194 | 30.13894901 | 30.07265389 | 30.01850091 | 29.72185189 | 29.67258043 | 29.99253464 | 29.9294537 | 29.90870536 | 29.75019868 |
| Probable ATP-dependent RNA helicas | Q13206 | DDX10 | Other | -1.418449189 | 0.971307548 | 23.04193181 | 22.0935852 | 22.20669484 | 24.29600093 | 23.28065072 | 22.88487106 | 21.73549818 | 20.49206952 | 22.41895201 |
| Probable ATP-dependent RNA helicas | Q92841;Q92841-1;Q92841-3 | DDX17 | Extracellular or secreted | -0.764646233 | 3.483942045 | 31.02525022 | 31.08777808 | 31.10618222 | 30.89004879 | 30.835601 | 31.50564169 | 30.25945799 | 30.44084874 | 30.23100557 |
| ATP-dependent RNA helicase DDX18 | Q9NVP1 | DDX18 | Other | -1.640984752 | 1.232388571 | 25.45044986 | 25.39443888 | 25.44594336 | 25.34999467 | 25.63064818 | 26.72377321 | 24.90122562 | 23.22146234 | 23.95198187 |
| ATP-dependent RNA helicase DDX19A | Q9NUU7;Q9NUU7-2;Q9UMR | DDX19A;DDX19B | Other | 0.641107661 | 1.876298363 | 26.33325023 | 26.29426638 | 26.12119549 | 26.97202269 | 26.91479022 | 26.76989009 | 27.32757389 | 27.33159409 | 26.96686256 |
| Probable ATP-dependent RNA helicas | Q9UHI6 | DDX20 | Extracellular or secreted | 0.578492693 | 0.66558927 | 21.71389325 | 23.34394364 | 24.28736012 | 22.40776877 | 22.19376116 | 23.36653033 | 23.73148477 | 23.51002429 | 23.15059765 |
| Nucleolar RNA helicase 2 | Q9NR30;Q9NR30-2 | DDX21 | Other | -0.775250198 | 2.048938659 | 29.28861466 | 29.1237874 | 29.21222336 | 29.29142792 | 29.06814962 | 30.30595603 | 28.56681508 | 28.60618458 | 28.64632924 |
| Probable ATP-dependent RNA helicas | Q9BUQ8 | DDX23 | Extracellular or secreted | -2.063430233 | 2.319017405 | 27.56732296 | 27.30469473 | 27.59056546 | 26.67249613 | 26.65717341 | 27.59988629 | 24.7297228 | 25.06336381 | 25.71269219 |
| ATP-dependent RNA helicase DDX24 | Q9GZR7;Q9GZR7-2 | DDX24 | Other | -0.86228573 | 2.155338421 | 25.17004974 | 24.68308585 | 25.1980553 | 24.95464646 | 24.73371008 | 25.4349173 | 23.972461 | 24.45368711 | 24.07422706 |
| Probable ATP-dependent RNA helicas | Q96GQ7 | DDX27 | Other | -1.604719271 | 1.536436148 | 26.45133061 | 26.48348443 | 26.58305085 | 26.44072599 | 26.46961002 | 26.16702759 | 24.98589612 | 25.24651881 | 24.251042 |
| ATP-dependent RNA helicase DDX39A | O00148;O00148-2;O00148-3 | DDX39A | Other | 0.969453234 | 1.920888882 | 26.73918109 | 26.5767226 | 26.65389797 | 26.09735994 | 26.16993509 | 27.73440801 | 27.8760171 | 27.68598846 | 27.3321065 |
| Spliceosome RNA helicase DDX39B | Q13838;Q13838-2 | DDX39B | Other | 0.094323117 | 0.105166123 | 32.80458965 | 32.85184698 | 32.78554083 | 32.22014631 | 32.26681934 | 32.33073966 | 32.94230646 | 32.9102969 | 32.06020738 |
| ATP-dependent RNA helicase DDX3X; | O00571-2;O00571;O15523-2 | DDX3X;DDX3Y | Extracellular or secreted | -0.400417457 | 1.248359929 | 28.73498936 | 28.76113276 | 28.81561695 | 28.3294143 | 28.47961981 | 29.12721145 | 28.06896963 | 28.3965635 | 28.45720683 |
| Probable ATP-dependent RNA helicas | Q9UJV9 | DDX41 | Other | -0.828397056 | 1.22721223 | 22.44387132 | 23.38271373 | 22.43022793 | 22.04606886 | 22.43055975 | 22.31442946 | 21.79835788 | 22.06045582 | 21.17993065 |
| ATP-dependent RNA helicase DDX42 | Q86XP3;Q86XP3-2 | DDX42 | Other | -0.629823549 | 1.980613901 | 29.06278247 | 29.1231707 | 29.00450505 | 28.59441493 | 28.56942512 | 29.12910473 | 28.11644391 | 28.22015784 | 28.5156291 |
| Probable ATP-dependent RNA helicas | Q7L014 | DDX46 | Other | -0.856006423 | 2.120133264 | 28.05837627 | 27.88593939 | 27.91593006 | 27.70206729 | 27.66871449 | 28.20978569 | 27.12139313 | 26.78917641 | 27.24181779 |
| Probable ATP-dependent RNA helicas | Q9H0S4;Q9H0S4-2 | DDX47 | Other | -0.768547407 | 0.789103014 | 23.29726111 | 23.51773282 | 23.80321511 | 23.6462537 | 23.63920379 | 25.92589938 | 22.93054028 | 22.89663248 | 23.78196797 |
| Probable ATP-dependent RNA helicas | P17844;P17844-2 | DDX5 | Extracellular or secreted | -0.605774724 | 2.135002966 | 28.22384418 | 28.32216747 | 28.28104885 | 28.49769874 | 28.57398134 | 28.9912375 | 27.67337259 | 27.94201275 | 28.01227952 |
| ATP-dependent RNA helicase DDX51 | Q8N8A6 | DDX51 | Other | -1.2749352 | 0.48334742 | 23.69954111 | 23.50675986 | 23.3295084 | 22.07668233 | 22.95611875 | 23.80163732 | 22.26999298 | 20.08877964 | 23.50154566 |
| ATP-dependent RNA helicase DDX54 | Q8TDD1;Q8TDD1-2 | DDX54 | Other | -1.422546821 | 0.759316168 | 23.21911474 | 22.87105846 | 22.93630365 | 22.85770086 | 22.88816427 | 22.50941684 | 22.1644971 | 20.07382777 | 22.13491408 |
| Probable ATP-dependent RNA helicas | P26196 | DDX6 | Other | 0.066837191 | 0.078575956 | 30.71227236 | 30.93220243 | 30.86053101 | 30.55873763 | 30.46324244 | 30.42602418 | 30.89684782 | 31.09156601 | 30.18860277 |
| 2,4-dienoyl-CoA reductase, mitochon | Q16698-2;Q16698 | DECR1 | Extracellular or secreted | -1.391152267 | 0.949962609 | 25.06315805 | 24.52599769 | 24.80656221 | 25.72606242 | 25.32264887 | 27.41425554 | 23.2011703 | 23.93427109 | 25.12044419 |
| Protein DEK | P35659;P35659-2 | DEK | Other | -0.49502954 | 1.057162763 | 29.61871676 | 29.52193106 | 29.62837965 | 29.62146174 | 29.3530654 | 30.92634627 | 29.24911685 | 29.2828612 | 29.31788376 |
| Density-regulated protein | O43583 | DENR | Other | 0.381112355 | 1.307149336 | 26.89377053 | 27.09557039 | 26.8460135 | 27.59184976 | 27.71063126 | 27.35194991 | 27.56347316 | 27.61701821 | 27.70773837 |
| Deoxyribose-phosphate aldolase | Q9Y315 | DERA | Extracellular or secreted | -0.06053354 | 0.070882527 | 29.21064515 | 29.32755247 | 29.27828174 | 28.41033819 | 28.30825833 | 28.16920876 | 28.89853538 | 28.96765974 | 28.30434659 |
| DNA fragmentation factor subunit alp | O00273;O00273-2 | DFFA | Other | 0.704731847 | 0.53844978 | 27.07214275 | 26.81115182 | 27.11416156 | 26.60907681 | 26.80501268 | 24.96692855 | 27.87367249 | 27.59910594 | 26.33065419 |
| DNA fragmentation factor subunit bet | O76075;O76075-2 | DFFB | Other | 1.13340022 | 1.254605744 | 22.67381198 | 20.0635601 | 22.44710442 | 21.22435327 | 22.0071012 | 20.82229548 | 22.95829782 | 22.93064856 | 22.13036751 |
| Dihydrofolate reductase;Dihydrofolat | P00374-2;P00374;Q86XF0 | DHFR;DHFRL1 | Other | 1.079842554 | 0.471422025 | 21.70872854 | 21.61902278 | 22.47436773 | 23.0500612 | 20.81758283 | 23.04947967 | 23.77986601 | 24.31008019 | 21.50920283 |
| Deoxyhypusine synthase | P49366;P49366-2;P49366-3 | DHPS | Other | 0.08836914 | 0.162511818 | 30.139559 | 30.15388048 | 30.11421121 | 29.98440829 | 29.998734 | 29.53027246 | 30.23386477 | 30.26919047 | 29.7225849 |
| Dehydrogenase/reductase SDR family | Q6UWP2-2;Q6UWP2;Q6UWP | DHRS11 | Extracellular or secreted | -0.272132584 | 0.272422654 | 24.43676084 | 24.76850258 | 24.19715549 | 24.50197078 | 24.69418315 | 23.38548117 | 23.91433403 | 24.70282032 | 23.5584749 |
| Probable 2-oxoglutarate dehydrogena | Q96HY7 | DHTKD1 | Other | -0.352463991 | 0.266480734 | 25.65739151 | 25.8947421 | 26.06607703 | 26.34883039 | 26.55591808 | 28.45340453 | 25.88899569 | 25.69112391 | 26.85067025 |
| Pre-mRNA-splicing factor ATP-depend | O43143 | DHX15 | Other | -0.462686995 | 2.344298618 | 30.48212298 | 30.63495309 | 30.59156446 | 30.03590523 | 30.15351813 | 30.28769037 | 29.90827588 | 29.95573862 | 29.84080164 |
| Putative pre-mRNA-splicing factor ATI | O60231 | DHX16 | Other | -0.334891832 | 1.012752031 | 26.23152751 | 26.17092842 | 25.90726753 | 26.22221414 | 25.98680761 | 26.77618019 | 25.66847104 | 25.90331903 | 26.07099713 |
| ATP-dependent RNA helicase DHX29 | Q7Z478 | DHX29 | Extracellular or secreted | -1.032163951 | 0.588906581 | 25.9410949 | 26.0383147 | 25.86235321 | 26.74816828 | 25.72088049 | 27.14043694 | 25.89952104 | 23.87389539 | 25.85571597 |
| Putative ATP-dependent RNA helicase | Q7L2E3-3;Q7L2E3;Q7L2E3-2 | DHX30 | Extracellular or secreted | 0.235973535 | 0.262794368 | 25.14631509 | 25.13422137 | 25.42755147 | 26.15934815 | 25.82335227 | 27.37252077 | 25.98089439 | 25.92372517 | 26.3349556 |
| ATP-dependent RNA helicase DHX36 | Q9H2U1-3;Q9H2U1-2;Q9H2U | DHX36 | Extracellular or secreted | 0.286991435 | 0.890376222 | 26.09121794 | 26.13284955 | 26.15946364 | 25.40947251 | 26.10171397 | 26.03948655 | 26.34911733 | 26.42415885 | 26.0548002 |
| Pre-mRNA-splicing factor ATP-depend | Q92620 | DHX38 | Other | -0.04423544 | 0.053062692 | 23.95269291 | 24.45104757 | 24.69535192 | 24.81484724 | 24.90260623 | 23.95011373 | 23.96349303 | 24.72009768 | 24.56703277 |
| ATP-dependent RNA helicase DHX8 | Q14562 | DHX8 | Extracellular or secreted | -1.25476903 | 3.065009799 | 26.44570661 | 26.06177874 | 26.35460882 | 25.90848205 | 25.47932684 | 25.58877982 | 24.43980705 | 24.84090966 | 24.87431764 |
| ATP-dependent RNA helicase A | Q08211 | DHX9 | Other | 0.386711988 | 0.804680686 | 31.61135653 | 31.73109305 | 31.66801278 | 31.04771756 | 31.04166032 | 32.14570552 | 31.96344204 | 32.19101762 | 31.62844918 |
| Diablo homolog, mitochondrial | Q9NR28-2;Q9NR28;Q9NR28- | DIABLO | Extracellular or secreted | 0.390534355 | 0.347238667 | 27.65218906 | 27.67418117 | 27.60625519 | 26.91570216 | 27.14053446 | 26.88832223 | 28.08192515 | 28.14821744 | 26.88005261 |
| Protein diaphanous homolog 1 | O60610-2;O60610-3;O60610 | DIAPH1 | Extracellular or secreted | -0.235240479 | 2.314922905 | 28.91667047 | 29.11512952 | 29.06252517 | 29.00744878 | 29.10534563 | 28.98359313 | 28.79160022 | 28.73534452 | 28.86269018 |
| Protein diaphanous homolog 2 | O60879-2;O60879-3;O60879 | DIAPH2 | Other | -1.248586111 | 1.356219379 | 25.50567008 | 24.87201725 | 24.77715898 | 24.35359969 | 25.13989073 | 23.87998253 | 23.74427215 | 22.8117831 | 23.96234604 |
| Endoribonuclease Dicer | Q9UPY3-2;Q9UPY3;Q9UPY3-3 | DICER1 | Other | 0.585369293 | 0.594345361 | 23.08409274 | 23.39362089 | 23.95677459 | 24.53949064 | 23.89133872 | 25.94808717 | 24.305843 | 24.80326439 | 25.05370287 |
| Death-inducer obliterator 1 | Q9BTC0 | DIDO1 | Other | -0.827372366 | 0.621529694 | 24.17317998 | 24.408663 | 24.78735899 | 23.79491238 | 24.37145762 | 25.4474576 | 23.72660817 | 22.76271617 | 24.52007334 |
| Digestive organ expansion factor hom | Q68CQ4 | DIEXF | Other | 0.068512591 | 0.144585567 | 26.20594206 | 26.4234377 | 26.38898228 | 25.28985933 | 25.86268427 | 25.84385915 | 26.04489401 | 26.14645106 | 26.0215751 |
| Disco-interacting protein 2 homolog B | Q9P265 | DIP2B | Extracellular or secreted | 1.938628581 | 3.518112889 | 24.22582171 | 24.91328425 | 24.56214878 | 24.87347303 | 23.65774585 | 25.4737647 | 26.44332122 | 26.79872373 | 26.42695994 |
| Exosome complex exonuclease RRP44 | Q9Y2L1;Q9Y2L1-2 | DIS3 | Other | -0.316134149 | 1.911885416 | 28.47803864 | 28.51713207 | 28.50419301 | 28.66336276 | 28.90083996 | 28.82012919 | 28.41437653 | 28.22094198 | 28.35812687 |
| DIS3-like exonuclease 2 | Q8IYB7;Q8IYB7-2;Q8IYB7-3 | DIS3L2 | Other | -0.006780707 | 0.005398033 | 26.18124145 | 26.18152582 | 26.13377077 | 25.51108062 | 25.61251173 | 25.42598418 | 26.16447871 | 26.24494141 | 25.09329505 |
| H/ACA ribonucleoprotein complex sub | O60832;O60832-2 | DKC1 | Other | -0.186347768 | 0.737683454 | 28.59327542 | 28.71826294 | 28.50104447 | 28.28202162 | 28.56201773 | 29.0875252 | 28.52544518 | 28.31790532 | 28.46967988 |
| Dickkopf-related protein 1 | O94907 | DKK1 | Extracellular or secreted | -0.864193727 | 0.350212394 | 24.84402683 | 24.13108388 | 24.74170315 | 26.2816768 | 26.02974807 | 28.0550175 | 23.79461497 | 24.16392234 | 26.49050963 |
| Dihydrolipoyllysine-residue acetyltran | P10515 | DLAT | Other | 0.200818614 | 0.12910997 | 24.31768973 | 24.89582856 | 24.92858994 | 24.37510892 | 22.52592602 | 26.85210008 | 24.77720915 | 24.6798151 | 25.09305321 |
| Dihydrolipoyl dehydrogenase, mitoch | P09622;P09622-2;P09622-3 | DLD | Other | -0.838266037 | 3.140167488 | 31.8977856 | 31.76010178 | 31.73190228 | 32.22196187 | 32.2787193 | 31.45708938 | 31.15297444 | 31.08277542 | 30.92323213 |
| Disks large homolog 1 | Q12959-5;Q12959-3;Q12959 | DLG1 | Extracellular or secreted | -0.328571047 | 1.42631289 | 29.24812164 | 29.19206476 | 29.20613309 | 28.80415111 | 28.77137073 | 28.95140945 | 28.60473628 | 28.62488128 | 28.87129469 |
| Disks large homolog 3 | Q92796-3;Q92796-2;Q92796 | DLG3 | Extracellular or secreted | 0.46094554 | 0.476499414 | 24.13029844 | 24.19917927 | 24.14172362 | 22.10748554 | 22.92488063 | 23.72619238 | 23.61504046 | 23.85119468 | 24.53148141 |
| Disks large-associated protein 4 | Q9Y2H0-3;Q9Y2H0;Q9Y2H0-1 | DLGAP4 | Other | 1.915576607 | 1.676392497 | 26.03286146 | 24.84928673 | 25.28139382 | 22.25790457 | 24.13641356 | 25.97368756 | 26.6087949 | 26.71711912 | 26.68658965 |
| Disks large-associated protein 5 | Q15398-3;Q15398-1;Q15398 | DLGAP5 | Other | -0.256386347 | 0.133690599 | 23.54980722 | 23.39296616 | 21.41547162 | 23.52312832 | 23.50700193 | 22.61009125 | 23.53469044 | 23.13930528 | 21.55607849 |
| Dihydrolipoyllysine-residue succinyltr | P36957;P36957-2 | DLST | Extracellular or secreted | -0.191198074 | 0.197695097 | 25.86154888 | 26.08903678 | 26.35502909 | 26.71371499 | 26.71253477 | 28.30630379 | 26.34502726 | 26.32981632 | 26.77064635 |
| Deleted in malignant brain tumors 1 p | Q9UGM3-4;Q9UGM3-8;Q9U | DMBT1 | Extracellular or secreted | 1.741031398 | 0.742151228 | 24.90288219 | 25.54200288 | 25.41515866 | 21.67400965 | 21.05273076 | 28.06203617 | 25.53171936 | 26.00793526 | 27.00784974 |
| Dynein heavy chain 8, axonemal | Q96JB1;Q96JB1-2 | DNAH8 | Other | -0.640627252 | 0.170159196 | 28.39018024 | 26.21152266 | 26.3191379 | 22.56340505 | 26.61610682 | 21.65083297 | 23.19043418 | 24.20418902 | 26.55908786 |
| DnaJ homolog subfamily A member 1 | P31689;P31689-2 | DNAJA1 | Extracellular or secreted | -0.21249504 | 0.698312441 | 28.59067253 | 28.57582145 | 28.59281224 | 29.06884153 | 29.0787226 | 29.23986209 | 28.73133586 | 28.72913283 | 28.4754124 |
| DnaJ homolog subfamily A member 2 | O60884 | DNAJA2 | Extracellular or secreted | 0.02648444 | 0.044127653 | 28.67485463 | 28.27061685 | 28.65972719 | 28.60151657 | 28.5965491 | 29.09065781 | 28.90377297 | 28.78665505 | 28.33598638 |
| DnaJ homolog subfamily A member 3, | Q96EY1-2;Q96EY1;Q96EY1-3 | DNAJA3 | Extracellular or secreted | 0.138182924 | 0.134509635 | 23.78246798 | 24.38073373 | 23.98754507 | 25.1808622 | 25.06225238 | 24.06972771 | 24.74237153 | 23.9627873 | 24.94118447 |
| DnaJ homolog subfamily B member 1 | P25685;P25685-2 | DNAJB1 | Extracellular or secreted | -0.128160433 | 0.345917695 | 27.15267229 | 27.32705986 | 27.35732997 | 27.56985966 | 27.74592672 | 27.34823945 | 27.53103515 | 27.23443593 | 27.1005916 |
| DnaJ homolog subfamily B member 1 | Q9UBS4 | DNAJB11 | Extracellular or secreted | -0.026785152 | 0.03990472 | 28.21001803 | 28.29123029 | 28.26239584 | 28.18989904 | 28.2525947 | 29.1574267 | 28.19891709 | 28.1892864 | 28.71322335 |
| DnaJ homolog subfamily C member 1 | Q8IXB1-2;Q8IXB1 | DNAJC10 | Other | -0.224471602 | 0.556144316 | 26.60314502 | 26.34661734 | 26.58820794 | 25.86027051 | 25.84876161 | 26.98966856 | 26.14033942 | 26.17306558 | 26.13151569 |
| DnaJ homolog subfamily C member 1 | Q9NVH1-3;Q9NVH1;Q9NVH1 | DNAJC11 | Extracellular or secreted | 0.063626795 | 0.023896961 | 21.82690997 | 21.89182078 | 21.94563815 | 21.34774705 | 20.12238883 | 25.30389388 | 20.97991066 | 22.77676851 | 22.65340054 |
| DnaJ homolog subfamily C member 1 | O75165 | DNAJC13 | Extracellular or secreted | 0.741123748 | 0.564179311 | 24.65337867 | 24.41331957 | 24.33633548 | 23.60512499 | 21.2056085 | 23.89762982 | 25.00688726 | 24.45406379 | 23.81811872 |
| DnaJ homolog subfamily C member 1 | Q9NVM6 | DNAJC17 | Other | 0.153281487 | 0.166863616 | 22.15545444 | 22.80780063 | 22.72406997 | 23.95171514 | 24.19670537 | 22.36153261 | 23.26069217 | 23.26670878 | 23.0310826 |
| DnaJ homolog subfamily C member 2 | Q99543;Q99543-2 | DNAJC2 | Other | -0.553840796 | 1.604174824 | 27.64079027 | 27.6150545 | 27.62693608 | 27.64388921 | 27.46901996 | 27.17092842 | 27.12070126 | 27.07887527 | 26.7222103 |
| DnaJ homolog subfamily C member 2 | Q5F1R6;Q5F1R6-3;Q5F1R6-2 | DNAJC21 | Other | -0.14821222 | 0.441814319 | 24.35897361 | 24.6984817 | 24.29991788 | 23.76233094 | 24.36919582 | 23.87633333 | 24.00307384 | 24.17150118 | 24.06340496 |
| DnaJ homolog subfamily C member 3 | Q13217 | DNAJC3 | Extracellular or secreted | -0.985098926 | 0.662435612 | 22.83676541 | 25.52954661 | 24.6957767 | 25.48440671 | 25.53299768 | 27.61224455 | 24.1891827 | 24.96969743 | 23.73169192 |
| DnaJ homolog subfamily C member 5 | Q9H3Z4-2;Q9H3Z4 | DNAJC5 | Extracellular or secreted | 1.357963245 | 0.745179534 | 20.9316877 | 25.39326082 | 24.89101417 | 22.33642881 | 22.19353572 | 26.24852886 | 24.77520074 | 25.22725486 | 25.06866218 |
| DnaJ homolog subfamily C member 7 | Q99615-2;Q99615 | DNAJC7 | Extracellular or secreted | 0.124271807 | 0.353458786 | 26.01335556 | 26.25833959 | 26.23770319 | 26.70833057 | 26.70147251 | 26.09172236 | 26.62198566 | 26.30766791 | 26.44862375 |

| | | | | | | | | | | | | | | |
|---|---|---|---|---|---|---|---|---|---|---|---|---|---|---|
| DnaJ homolog subfamily C member 8 | O75937 | DNAJC8 | Extracellular or secreted | 0.07296531 | 0.285890374 | 29.68418337 | 29.56274563 | 29.52770241 | 29.43841167 | 29.24204503 | 29.05198791 | 29.42484362 | 29.58836881 | 29.4592215 |
| DnaJ homolog subfamily C member 9 | Q8WXX5 | DNAJC9 | Extracellular or secreted | -0.242806364 | 0.581893862 | 28.54011911 | 28.27475428 | 28.32410066 | 28.49857394 | 28.75837125 | 27.78250548 | 27.93550806 | 28.03170524 | 28.39357998 |
| Dynein light chain 1, axonemal | Q4LDG9-2;Q4LDG9-3;Q4LDG | DNAL1 | Other | 0.87231646 | 1.239311204 | 19.99808764 | 22.45146086 | 21.12540152 | 21.71950098 | 21.03886638 | 22.04376432 | 22.27130458 | 22.31997856 | 22.21420709 |
| Deoxyribonuclease-1-like 1 | P49184 | DNASE1L1 | Extracellular or secreted | 1.662447395 | 1.670838594 | 21.3054673 | 21.01243086 | 21.29255081 | 21.67163263 | 21.04126343 | 20.94870658 | 22.32657651 | 22.91632196 | 23.38046952 |
| Deoxyribonuclease-2-alpha | O00115-2;O00115 | DNASE2 | Extracellular or secreted | 5.91E-05 | 6.70E-05 | 28.29867749 | 28.0759719 | 28.08035018 | 28.47223909 | 28.5067296 | 27.86873638 | 27.99792689 | 27.92019646 | 28.73340624 |
| Dynamin-1 | Q05193-3;Q05193;Q05193-5 | DNM1 | Extracellular or secreted | -0.235650085 | 0.94702608 | 25.99483421 | 25.90152486 | 26.13484805 | 26.30511239 | 26.09729966 | 26.67909024 | 25.93777681 | 26.06525538 | 25.84637225 |
| Dynamin-1-like protein | O00429-3;O00429;O00429-4 | DNM1L | Other | 0.119403605 | 0.970712345 | 28.28475951 | 28.56245451 | 28.544695 | 28.61747369 | 28.63930731 | 28.66166503 | 28.61006308 | 28.6907577 | 28.71256756 |
| Dynamin-2 | P50570-2;P50570-5;P50570;I | DNM2 | Extracellular or secreted | -0.282609967 | 1.811869162 | 27.36761378 | 27.39345723 | 27.44632207 | 27.40186139 | 27.32337065 | 27.67962721 | 27.26597595 | 27.06599489 | 27.12632542 |
| DNA (cytosine-5)-methyltransferase 1 | P26358;P26358-2;P26358-3 | DNMT1 | Other | -0.446454896 | 0.770376641 | 28.10379604 | 28.00007816 | 28.05791168 | 28.37807295 | 28.12814611 | 28.94382994 | 27.99851881 | 28.08451249 | 27.38352145 |
| Aspartyl aminopeptidase | Q9ULA0 | DNPEP | Extracellular or secreted | -0.096785697 | 0.14788769 | 30.14503737 | 30.20373176 | 30.18091909 | 30.02973228 | 30.09093538 | 29.20261133 | 30.09722932 | 30.03891644 | 29.49998076 |
| 2-deoxynucleoside 5-phosphate N-hy | O43598;O43598-2 | DNPH1 | Extracellular or secreted | -0.144073598 | 0.145062478 | 28.0378436 | 28.02205154 | 27.97306346 | 27.3711086 | 27.55479165 | 26.44496456 | 27.63415686 | 27.76426874 | 26.87126531 |
| Deoxynucleotidyltransferase terminal | Q5QJE6 | DNTTIP2 | Other | -2.968095842 | 1.981981729 | 23.43211571 | 23.49252803 | 23.10309568 | 23.68078101 | 23.78646188 | 24.39008182 | 21.37343142 | 19.89752852 | 20.76728459 |
| Dedicator of cytokinesis protein 7 | Q96N67-4;Q96N67-3;Q96N6 | DOCK7 | Other | 0.19146984 | 0.719413571 | 25.05461392 | 24.6991704 | 24.96327253 | 24.75674362 | 24.6549633 | 24.34163826 | 24.93589018 | 24.79476368 | 25.07895668 |
| Dedicator of cytokinesis protein 9 | Q9BZ29-3;Q9BZ29-5;Q9BZ29 | DOCK9 | Other | 0.975087402 | 1.014789795 | 22.17252395 | 23.08393049 | 22.69406622 | 19.9327922 | 21.39909558 | 21.6324965 | 22.39498177 | 22.86387546 | 23.12385744 |
| Deoxyhypusine hydroxylase | Q9BU89 | DOHH | Other | 2.496075465 | 2.060609801 | 23.4374595 | 20.75250667 | 21.55679229 | 23.97009255 | 23.85224298 | 21.93131665 | 25.63657882 | 25.32247696 | 24.27937594 |
| Protein DPCD | Q9BVM2 | DPCD | Other | -0.246871586 | 0.390755184 | 23.70925129 | 22.93797432 | 22.70864105 | 22.97233829 | 23.38192205 | 22.23575833 | 22.77116542 | 23.04506905 | 22.41609343 |
| Zinc finger protein ubi-d4 | Q92785;Q92785-2 | DPF2 | Other | -0.554341377 | 0.921158653 | 25.57649196 | 25.33391556 | 24.96247843 | 25.15034974 | 25.39100019 | 25.0987659 | 24.26856599 | 25.04364312 | 24.78126765 |
| Diphthamide biosynthesis protein 1 | Q9BZG8-3;Q9BZG8-2;Q9BZG | DPH1 | Other | -0.136439952 | 0.056848465 | 23.28354828 | 21.63989115 | 22.27532554 | 21.79080293 | 21.69848169 | 21.08855031 | 22.75391123 | 22.37161714 | 20.35345173 |
| Diphthamide biosynthesis protein 2 | Q9BQC3;Q9BQC3-3;Q9BQC3- | DPH2 | Other | 0.897850772 | 0.582983586 | 23.37491 | 23.67206444 | 23.60252218 | 23.39584475 | 23.51364279 | 22.86122703 | 25.14534354 | 24.57669377 | 23.18162059 |
| Diphthine synthase | Q9H2P9-3;Q9H2P9-2;Q9H2P | DPH5 | Other | 0.330953894 | 0.41372447 | 22.22352715 | 21.67686508 | 22.6526787 | 22.74028311 | 22.45816327 | 21.62644876 | 23.05783113 | 22.52971936 | 22.09429422 |
| Dolichol-phosphate mannosyltransfer | O60762 | DPM1 | Other | -0.764587809 | 0.251690024 | 21.04353466 | 20.43822274 | 21.62946086 | 22.19244803 | 20.16533395 | 26.7612913 | 20.40197528 | 20.63803934 | 22.78136772 |
| Dolichol-phosphate mannosyltransfer | Q9P2X0;Q9P2X0-2 | DPM3 | Other | -0.078226801 | 0.018390575 | 23.42266451 | 22.43620159 | 21.38041667 | 24.01934809 | 20.81133449 | 26.2242215 | 22.06824018 | 21.56355753 | 25.28061532 |
| Dipeptidyl peptidase 3 | Q9NY33;Q9NY33-4 | DPP3 | Extracellular or secreted | 0.081860229 | 0.153957555 | 31.84412114 | 31.82304852 | 31.7677373 | 32.14534111 | 32.24640668 | 30.87180875 | 31.90852643 | 31.78515104 | 31.90113497 |
| Dipeptidyl peptidase 2 | Q9UHL4 | DPP7 | Extracellular or secreted | -1.19555727 | 3.443706401 | 29.48979793 | 29.59510888 | 29.46791293 | 29.38339785 | 29.43604993 | 29.05100257 | 28.24658224 | 28.02289816 | 28.35548284 |
| Dipeptidyl peptidase 8 | Q6V1X1-4;Q6V1X1-2;Q6V1X1 | DPP8 | Other | -0.473562544 | 1.376726999 | 24.49570281 | 24.72681602 | 24.37550668 | 24.66573623 | 24.77318952 | 23.64207197 | 23.88529942 | 24.12028598 | 23.91323859 |
| Dipeptidyl peptidase 9 | Q86TI2;Q86TI2-4;Q86TI2-2 | DPP9 | Other | 0.065001075 | 0.219419386 | 29.0847659 | 29.22504022 | 29.07854448 | 28.76195701 | 28.80648846 | 28.72448898 | 29.03149492 | 28.8603534 | 29.14379743 |
| Protein dpy-30 homolog | Q9C005 | DPY30 | Other | 2.034022663 | 3.257349034 | 25.1828332 | 25.0606867 | 25.60540763 | 24.05606214 | 24.50911825 | 25.55007154 | 27.36210475 | 26.64705015 | 27.07500281 |
| Dihydropyrimidinase-related protein | Q16555;Q16555-2 | DPYSL2 | Extracellular or secreted | 0.718990682 | 2.569846064 | 31.35835746 | 31.37285284 | 31.37337155 | 30.90117094 | 30.92450691 | 30.73020237 | 31.67644444 | 31.80000533 | 32.01075332 |
| Dihydropyrimidinase-related protein | Q14195-2;Q14195 | DPYSL3 | Extracellular or secreted | -0.618843154 | 0.693318067 | 25.97105794 | 25.74658034 | 25.50582149 | 25.75437518 | 25.99610657 | 25.67780071 | 25.41950505 | 25.56104177 | 24.48879483 |
| Dihydropyrimidinase-related protein | Q9BPU6 | DPYSL5 | Other | -0.696439526 | 3.366800148 | 28.97459585 | 28.95396413 | 28.93811907 | 28.56263646 | 28.4676992 | 28.55811722 | 28.1212449 | 28.01966775 | 27.99733473 |
| Protein Dr1 | Q01658 | DR1 | Other | -0.24491945 | 0.441113774 | 26.34773273 | 26.39782045 | 26.51523808 | 26.47792288 | 26.84804521 | 26.02462162 | 26.51005448 | 26.24874599 | 25.81213166 |
| Dr1-associated corepressor | Q14919;Q14919-2 | DRAP1 | Other | -0.152772744 | 0.675936745 | 25.01111735 | 25.05295703 | 24.95681889 | 25.2445604 | 25.08917825 | 25.12008818 | 24.78066711 | 24.91811603 | 25.08025868 |
| Developmentally-regulated GTP-bindi | Q9Y295 | DRG1 | Other | -0.944702086 | 2.406102258 | 27.24372546 | 27.28091615 | 27.23808601 | 26.53109466 | 26.46273166 | 26.73156246 | 25.7332446 | 26.22350359 | 25.95320375 |
| Developmentally-regulated GTP-bindi | P55039 | DRG2 | Other | 0.583687312 | 1.018824843 | 23.30075584 | 23.50833255 | 23.32567108 | 21.96327075 | 23.45034034 | 24.01987981 | 23.70112875 | 23.81919132 | 24.01486705 |
| Desmocollin-1 | Q08554-2;Q08554 | DSC1 | Extracellular or secreted | -2.69785559 | 2.233640752 | 26.0654403 | 24.98006634 | 25.35181515 | 26.64209952 | 26.67492196 | 24.59832516 | 23.55964222 | 23.34989347 | 22.15323175 |
| Desmocollin-2 | Q02487;Q02487-2 | DSC2 | Extracellular or secreted | 0.845121476 | 3.667834667 | 27.79029559 | 27.64265043 | 27.25808794 | 27.63186956 | 27.7991562 | 28.02707109 | 28.60960525 | 28.54546851 | 28.45485607 |
| Desmocollin-3 | Q14574-2;Q14574 | DSC3 | Extracellular or secreted | 0.797628324 | 0.974724092 | 27.12020686 | 27.05372358 | 27.04791253 | 27.74804028 | 27.84050154 | 29.23708316 | 28.21549034 | 28.21900392 | 28.98212468 |
| Sister chromatid cohesion protein DC | Q9BVC3 | DSCC1 | Other | 0.683056902 | 0.777713556 | 24.43764998 | 24.71161615 | 23.85357608 | 23.92052587 | 24.31416666 | 24.71182617 | 25.31375161 | 25.37567239 | 24.33442715 |
| Down syndrome critical region protei | O14972-2;O14972 | DSCR3 | Other | 0.057208682 | 0.039883506 | 23.86599071 | 23.71586315 | 23.77334046 | 22.91676423 | 23.25811161 | 22.70416221 | 23.81899636 | 23.90398552 | 22.56576035 |
| Desmoglein-1 | Q02413 | DSG1 | Extracellular or secreted | -3.036236113 | 1.267117668 | 25.55429398 | 25.21007378 | 25.95883323 | 27.10888844 | 26.17658878 | 24.32251134 | 23.96560816 | 21.02709996 | 23.06417831 |
| Desmoglein-2 | Q14126 | DSG2 | Extracellular or secreted | 0.701543869 | 1.242563198 | 29.85784974 | 29.65517832 | 29.71878552 | 29.61076714 | 29.81555588 | 30.00463899 | 30.23306476 | 30.34286769 | 30.86008694 |
| Desmoglein-3 | P32926 | DSG3 | Extracellular or secreted | 1.65926839 | 1.552610715 | 24.70038807 | 24.88044971 | 24.51201559 | 24.01877679 | 24.22346677 | 25.18499063 | 25.73140708 | 26.04420616 | 26.96223571 |
| Kinetochore-associated protein DSN1 | Q9H410-4;Q9H410;Q9H410-2 | DSN1 | Other | -0.425131152 | 0.671834758 | 25.47199887 | 25.85409946 | 25.48213068 | 24.40626418 | 25.22615256 | 24.16484311 | 24.64685795 | 24.94449474 | 24.43599828 |
| Desmoplakin | P15924;P15924-3;P15924-2 | DSP | Extracellular or secreted | -0.346562153 | 1.227565435 | 30.59040485 | 30.44480268 | 30.61219182 | 30.54756596 | 30.49161475 | 31.3719706 | 30.41134883 | 30.33276809 | 30.24547194 |
| Dystonin | Q03001-14;Q03001;Q03001- | DST | Extracellular or secreted | 0.562990824 | 0.543104757 | 22.364047 | 23.15758379 | 22.77424576 | 22.24457915 | 23.1972905 | 24.90508801 | 23.91314727 | 23.09863742 | 23.99860489 |
| Destrin | P60981;P60981-2 | DSTN | Extracellular or secreted | 0.535452234 | 1.970337958 | 30.30595603 | 30.37025648 | 30.31365216 | 30.22227864 | 30.46985452 | 30.41508207 | 30.72362676 | 30.96016535 | 30.97110455 |
| D-tyrosyl-tRNA(Tyr) deacylase 1 | Q8TEA8 | DTD1 | Other | 0.018213007 | 0.042432976 | 29.08603229 | 29.22802598 | 28.97686409 | 28.84323019 | 28.79029559 | 28.86679256 | 28.78724688 | 28.94181692 | 29.22119558 |
| Probable D-tyrosyl-tRNA(Tyr) deacyla | Q96FN9 | DTD2 | Other | 0.308889653 | 0.284827293 | 22.9271978 | 23.27013561 | 23.73665462 | 22.75599022 | 22.72357013 | 21.61818115 | 23.43529892 | 22.47325686 | 23.53397794 |
| Dysbindin | Q96EV8-3;Q96EV8-2;Q96EV8 | DTNBP1 | Extracellular or secreted | 1.185059736 | 1.193800641 | 21.64062941 | 22.57591924 | 21.64566375 | 22.02105935 | 21.57589061 | 20.72520282 | 22.43285484 | 22.57019852 | 23.64430844 |
| Thymidylate kinase | P23919;P23919-2 | DTYMK | Other | 1.123184029 | 1.625454931 | 26.58190232 | 26.99927181 | 26.81396707 | 27.05693038 | 26.3274197 | 27.14530467 | 28.16283776 | 28.20983216 | 27.45928014 |
| tRNA-dihydrouridine(47) synthase [N | Q96G46-2;Q96G46 | DUS3L | Other | 1.128578633 | 0.907457566 | 20.41741586 | 22.98525979 | 22.14362597 | 23.14258993 | 20.55069538 | 23.33087653 | 23.87502111 | 23.15855484 | 22.63739169 |
| Dual specificity protein phosphatase 1 | Q9UNI6 | DUSP12 | Other | 0.65477083 | 0.853734351 | 22.09258554 | 22.7267121 | 22.60794882 | 22.57119435 | 22.79195537 | 22.63185291 | 23.59217779 | 23.42918093 | 22.65407831 |
| Dual specificity protein phosphatase 3 | P51452;P51452-2 | DUSP3 | Extracellular or secreted | 0.015189623 | 0.02559634 | 26.11822759 | 26.22892509 | 26.1013533 | 26.98402794 | 27.00185055 | 26.00955922 | 26.42529595 | 26.55860627 | 26.28363849 |
| Deoxyuridine 5-triphosphate nucleoti | P33316-2 | DUT | Extracellular or secreted | 0.307335837 | 0.32527956 | 30.03852403 | 29.86471332 | 29.83415472 | 30.29175724 | 30.24717103 | 28.85599512 | 30.39652267 | 30.56556277 | 29.5260798 |
| Segment polarity protein dishevelled | O14641 | DVL2 | Extracellular or secreted | 0.735992961 | 1.906954093 | 24.15784588 | 23.57507847 | 23.44417473 | 23.82152877 | 23.67572966 | 23.80950909 | 24.66757806 | 24.57657845 | 24.20575567 |
| Cytoplasmic dynein 1 heavy chain 1 | Q14204 | DYNC1H1 | Extracellular or secreted | -0.182465557 | 1.309582733 | 32.95561739 | 33.02529973 | 32.99967505 | 32.8053587 | 32.79604493 | 32.8086993 | 32.81829398 | 32.69297436 | 32.63668253 |
| Cytoplasmic dynein 1 intermediate ch | O14576-4;O14576-3;O14576 | DYNC1I1 | Other | -0.629096395 | 0.134116558 | 25.65899897 | 25.98102509 | 25.7942927 | 25.46864716 | 25.64474197 | 21.70062207 | 26.02405089 | 25.87424727 | 21.33857664 |
| Cytoplasmic dynein 1 intermediate ch | Q13409-3;Q13409-2;Q13409 | DYNC1I2 | Other | 0.333351213 | 0.956701312 | 29.97404876 | 30.04152978 | 30.14479433 | 29.78536154 | 29.78729359 | 29.04835437 | 30.2266487 | 30.20489796 | 29.95919816 |
| Cytoplasmic dynein 1 light intermedia | Q9Y6G9 | DYNC1LI1 | Extracellular or secreted | 0.082827616 | 0.314780024 | 28.09421166 | 28.13775267 | 28.15990628 | 27.71004 | 27.77918973 | 27.65996531 | 28.12794939 | 27.94503083 | 27.94603545 |
| Cytoplasmic dynein 1 light intermedia | O43237;O43237-2 | DYNC1LI2 | Other | -0.405443542 | 1.400376484 | 28.62933544 | 28.82392924 | 28.69282059 | 28.48972138 | 28.58028566 | 28.25200812 | 28.31738784 | 28.24667283 | 27.95365891 |
| Cytoplasmic dynein 2 heavy chain 1 | Q8NCM8;Q8NCM8-2 | DYNC2H1 | Extracellular or secreted | 1.336548808 | 1.086405523 | 26.4745385 | 26.16587827 | 26.421256 | 22.78041683 | 25.76321784 | 25.7830178 | 26.81127434 | 27.50809071 | 26.384444 |
| Dynein light chain 1, cytoplasmic | P63167 | DYNLL1 | Extracellular or secreted | -0.105883737 | 0.210890325 | 29.83340088 | 30.02075419 | 30.01995932 | 29.29743167 | 29.41035841 | 30.08575378 | 29.8144714 | 29.81444084 | 29.38726566 |
| Dynein light chain 2, cytoplasmic | Q96FJ2 | DYNLL2 | Extracellular or secreted | -0.717929068 | 1.252236452 | 27.37351676 | 27.21891157 | 26.97388459 | 26.71371499 | 26.70319009 | 27.24136321 | 26.34005966 | 26.68271089 | 25.93573284 |
| Dynein light chain roadblock-type 1 | Q9NP97 | DYNLRB1 | Other | 0.756664549 | 1.769429937 | 28.1777391 | 28.05180645 | 27.95149283 | 27.12602996 | 27.50642696 | 27.95941929 | 28.8657017 | 28.45242293 | 28.33832631 |
| Dynein light chain Tctex-type 1 | P63172 | DYNLT1 | Extracellular or secreted | -0.077436579 | 0.078137271 | 25.62611451 | 25.73055223 | 25.78079225 | 25.64781873 | 25.87117129 | 25.28528001 | 25.90932931 | 25.89427953 | 24.93494593 |
| Dynein light chain Tctex-type 3 | P51808 | DYNLT3 | Extracellular or secreted | 0.431623075 | 0.87558672 | 27.04353883 | 26.80525875 | 26.82128546 | 26.6477364 | 27.03443662 | 27.00613823 | 27.46021802 | 26.95759397 | 27.55625438 |
| Dual specificity tyrosine-phosphorylat | Q13627-4;Q13627-3;Q13627 | DYRK1A | Other | -0.191469319 | 0.227107224 | 22.35135208 | 24.39512564 | 23.8944877 | 23.22166854 | 23.72171503 | 23.33987283 | 23.59115068 | 23.35366698 | 22.94288529 |
| ELL-associated factor 1 | Q96JC9 | EAF1 | Extracellular or secreted | -0.350002227 | 0.262650819 | 23.43898267 | 23.24231584 | 23.45938959 | 23.90939799 | 22.24642632 | 23.21206469 | 23.4612646 | 23.26227037 | 21.9807469 |
| E2F-associated phosphoprotein | Q56P03 | EAPP | Other | 0.295731907 | 0.178348988 | 23.03091428 | 22.90688693 | 23.53385915 | 20.99279177 | 21.31419297 | 21.51620541 | 23.06887535 | 22.89241385 | 21.57333177 |
| Probable rRNA-processing protein EB | Q99848 | EBNA1BP2 | Other | -1.720392066 | 2.832064376 | 26.26454497 | 26.47995895 | 26.60851293 | 25.29042193 | 25.32831034 | 26.92229657 | 24.62617022 | 24.61565792 | 24.0440185 |
| Endothelin-converting enzyme 1 | P42892-3;P42892-2;P42892-4 | ECE1 | Extracellular or secreted | 0.439187862 | 1.152313334 | 22.39312332 | 22.13304561 | 22.105388 | 21.63791806 | 22.71353671 | 21.58622153 | 22.74310125 | 22.41356492 | 22.44551403 |
| Delta(3,5)-Delta(2,4)-dienoyl-CoA isor | Q13011 | ECH1 | Extracellular or secreted | -0.192070376 | 1.067187503 | 32.06345768 | 32.1208063 | 32.17949608 | 32.13940653 | 32.12863165 | 31.93898134 | 31.93635083 | 32.00162241 | 31.77120542 |
| Enoyl-CoA hydratase, mitochondrial | P30084 | ECHS1 | Extracellular or secreted | -0.153669383 | 0.490307966 | 29.16307778 | 29.25653664 | 29.23354482 | 29.17509961 | 29.11107962 | 29.99523329 | 29.14969584 | 29.10744242 | 29.24913946 |
| Enoyl-CoA delta isomerase 1, mitocho | P42126;P42126-2 | ECI1 | Extracellular or secreted | -0.60402838 | 1.180113558 | 28.27502081 | 28.40831477 | 28.28731608 | 28.47661025 | 28.7278029 | 27.82304852 | 27.91536025 | 27.93651933 | 27.33509195 |
| Extracellular matrix protein 1 | Q16610;Q16610-4;Q16610-2 | ECM1 | Extracellular or secreted | 2.062976143 | 0.568924355 | 23.36839669 | 22.10248709 | 22.27523222 | 22.4196722 | 22.8439885 | 27.00227989 | 23.63610859 | 24.47982022 | 28.07902792 |
| Proteasome-associated protein ECM2 | Q5VYK3 | ECM29 | Other | 0.296286269 | 1.968626871 | 29.61208633 | 29.77262965 | 29.7289545 | 29.61046796 | 29.69225526 | 29.8942188 | 29.94461203 | 29.9744591 | 30.12509393 |
| Enhancer of mRNA-decapping protein | Q96F86 | EDC3 | Other | -0.663633926 | 0.606888493 | 23.01689124 | 22.89642923 | 23.76981444 | 23.62733973 | 23.7004939 | 23.81577573 | 22.27053487 | 22.52248192 | 23.62945357 |
| Enhancer of mRNA-decapping protein | Q6P2E9;Q6P2E9-2 | EDC4 | Other | -0.243161065 | 0.355474658 | 25.83340088 | 26.41858857 | 25.81633742 | 25.95215966 | 25.90090328 | 26.22309846 | 25.5472497 | 25.51406436 | 26.28144688 |
| Endothelial differentiation-related fac | O60869-2;O60869-3;O60869 | EDF1 | Extracellular or secreted | 0.463460749 | 0.658921973 | 27.5767226 | 25.80695547 | 25.5431541 | 27.10868907 | 27.03821002 | 26.85982044 | 27.0981836 | 27.01062661 | 27.24834789 |
| Early endosome antigen 1 | Q15075 | EEA1 | Extracellular or secreted | 0.025248053 | 0.089706501 | 29.89178776 | 29.66475341 | 29.67512393 | 29.82383815 | 29.97555277 | 29.36659286 | 29.71046705 | 29.70048068 | 29.86362088 |
| Polycomb protein EED | O75530-3;O75530;O75530-2 | EED | Other | 0.354813509 | 0.357554865 | 23.88091673 | 24.49911402 | 24.23403386 | 24.02698669 | 23.74334784 | 24.92944805 | 24.62103548 | 23.93265019 | 25.16767846 |
| Elongation factor 1-alpha 1;Putative e | P68104;Q5VTE0;P68104-2 | EEF1A1;EEF1A1P | Extracellular or secreted | 0.436805345 | 1.497229451 | 34.09796706 | 34.14102193 | 34.17860599 | 34.58110544 | 34.5372217 | 34.65421603 | 34.9240644 | 34.90126986 | 34.58015085 |
| Elongation factor 1-alpha 2 | Q05639 | EEF1A2 | Other | -0.319214551 | 0.883939078 | 32.19210592 | 32.18553421 | 32.16449909 | 31.73416571 | 31.68624733 | 32.37952996 | 31.87635382 | 31.86728613 | 31.4697575 |
| Elongation factor 1-beta | P24534 | EEF1B2 | Other | 1.239366744 | 4.228590125 | 31.20850717 | 31.25411644 | 31.22101115 | 30.59209935 | 30.71612163 | 31.22031932 | 32.23320765 | 32.37077612 | 32.22020398 |
| Elongation factor 1-delta | P29692;P29692-2 | EEF1D | Other | 0.44760492 | 2.461873121 | 32.23240729 | 32.14543222 | 32.11975558 | 32.03081774 | 32.18845539 | 32.02528323 | 32.63547214 | 32.61756564 | 32.4608527 |
| Elongation factor 1-delta | P29692-3 | EEF1D | Other | 0.488754802 | 1.237783333 | 26.36401025 | 26.66139321 | 26.85317152 | 26.38625445 | 26.49551984 | 27.22912685 | 27.28982416 | 27.31622283 | 26.85495548 |
| Eukaryotic translation elongation fact | O43324;O43324-2 | EEF1E1 | Extracellular or secreted | -0.510171 | 0.948768273 | 27.44181846 | 27.52065788 | 27.49914444 | 27.16748706 | 27.29824049 | 26.86926606 | 27.02347993 | 26.96609244 | 26.37822182 |
| Elongation factor 1-gamma | P26641;P26641-2 | EEF1G | Extracellular or secreted | -0.133840689 | 0.496529137 | 33.35534106 | 33.35507846 | 33.28071155 | 33.19325216 | 33.20182009 | 33.15963566 | 33.20780932 | 33.24274923 | 32.92083888 |
| Elongation factor 2 | P13639 | EEF2 | Extracellular or secreted | 0.306918453 | 0.884673452 | 33.84698116 | 33.87637212 | 33.86239829 | 34.06904007 | 34.04498518 | 33.90463807 | 34.33803934 | 34.39993806 | 33.98498541 |
| Eukaryotic elongation factor 2 kinase | O00418 | EEF2K | Other | 2.074107516 | 2.904758552 | 21.53751207 | 22.34362933 | 21.62256119 | 21.15064061 | 20.39637301 | 22.59255177 | 23.81333106 | 23.98962525 | 23.24100023 |
| Selenocysteine-specific elongation fac | P57772-2;P57772 | EEFSEC | Other | 0.488642348 | 0.492690284 | 22.98341047 | 23.08191703 | 23.14900094 | 23.22143254 | 22.47557021 | 23.71397713 | 23.92079842 | 24.00889703 | 22.84888574 |
| EGF-containing fibulin-like extracellul | Q12805-2;Q12805-4;Q12805 | EFEMP1 | Extracellular or secreted | 0.382654655 | 0.20646791 | 28.63131452 | 28.58724239 | 28.50157603 | 30.37637641 | 30.31083862 | 30.37730769 | 29.14299454 | 29.37085925 | 31.026438 |
| EF-hand domain-containing protein D | Q9BUP0;Q9BUP0-2 | EFHD1 | Extracellular or secreted | -0.926170318 | 0.265366919 | 25.06031563 | 25.47586848 | 25.11012388 | 25.11957379 | 25.13402547 | 24.53569921 | 25.6572552 | 25.16407584 | 21.61796123 |
| EF-hand domain-containing protein D | Q96C19 | EFHD2 | Other | -0.057316118 | 0.349231945 | 28.3364121 | 28.51653107 | 28.35485259 | 28.53567697 | 28.44797776 | 28.1641334 | 28.41880568 | 28.27453214 | 28.31250576 |
| Ephrin-A1;Ephrin-A1, secreted form | P20827;P20827-2 | EFNA1 | Extracellular or secreted | -0.151819242 | 0.125531676 | 27.12415729 | 26.9312304 | 26.83328022 | 26.78031668 | 26.84290063 | 27.42838241 | 26.28259612 | 26.59889304 | 27.63318693 |
| Ephrin-A5 | P52803 | EFNA5 | Extracellular or secreted | -0.406080787 | 0.103218329 | 20.51815832 | 24.08042135 | 23.40028221 | 24.27994264 | 24.30180261 | 24.17881254 | 21.77591405 | 21.84083632 | 25.5447171 |
| Ephrin-B1 | P98172 | EFNB1 | Extracellular or secreted | 0.0988496 | 0.060280811 | 25.31672347 | 25.18710707 | 25.23721084 | 25.80720121 | 25.62736756 | 26.47619305 | 25.05150569 | 25.35238779 | 26.71855691 |
| Ephrin-B3 | Q15768 | EFNB3 | Extracellular or secreted | -1.59412087 | 0.617303792 | 25.32378294 | 24.73531224 | 24.34197752 | 25.18044491 | 25.16721905 | 26.0573643 | 21.62460023 | 24.85319532 | 24.14289233 |
| Elongation factor Tu GTP-binding dom | Q7Z2Z2;Q7Z2Z2-2 | EFTUD1 | Other | 0.350778944 | 0.638498994 | 27.34367261 | 27.59626473 | 27.56920779 | 27.54546851 | 27.65969316 | 27.58491526 | 28.06563545 | 28.14831443 | 27.48799798 |
| 116 kDa U5 small nuclear ribonucleop | Q15029;Q15029-3;Q15029-2 | EFTUD2 | Other | -0.245161734 | 0.949427221 | 30.90361497 | 30.90720162 | 30.78144276 | 30.31753879 | 30.48855353 | 30.60284425 | 30.4752771 | 30.54277784 | 30.24705782 |
| Epidermal growth factor-like protein | Q9UHF1 | EGFL7 | Extracellular or secreted | -0.821985341 | 0.758184109 | 22.99470907 | 22.5529289 | 22.41588719 | 23.78506527 | 23.72640029 | 25.0631169 | 22.24603309 | 22.24559797 | 23.31146673 |
| Epidermal growth factor receptor | P00533;P00533-4;P00533-3; | EGFR | Extracellular or secreted | -3.612121667 | 1.127985007 | 27.04635202 | 27.00260181 | 26.92309042 | 26.77668222 | 25.62039235 | 29.24209047 | 22.3672505 | 22.25853728 | 25.84345186 |
| Egl nine homolog 1 | Q9GZT9-3;Q9GZT9;Q9GZT9-2 | EGLN1 | Other | 0.075481218 | 0.111843217 | 21.14904747 | 20.89622596 | 21.03079644 | 21.04346791 | 21.68174627 | 20.28104708 | 20.75041913 | 21.24574302 | 21.27144707 |
| EH domain-binding protein 1 | Q8NDI1-3;Q8NDI1-2;Q8NDI1 | EHBP1 | Extracellular or secreted | 0.536399028 | 0.563605078 | 22.98259257 | 23.77172964 | 24.16966754 | 23.7370674 | 23.53064827 | 23.59821156 | 24.38185606 | 23.46313718 | 24.65916234 |
| EH domain-binding protein 1-like prot | Q8N3D4 | EHBP1L1 | Other | 0.393786797 | 0.294257422 | 24.26198355 | 24.09860529 | 24.56354589 | 23.97797232 | 22.0078825 | 25.14499362 | 23.83807439 | 24.30848402 | 25.06229356 |
| EH domain-containing protein 1 | Q9H4M9 | EHD1 | Extracellular or secreted | 0.524046115 | 1.003350828 | 28.62226501 | 28.58323023 | 28.70091056 | 28.36365107 | 28.39050828 | 28.62749281 | 28.80328903 | 28.97448645 | 29.43839184 |
| EH domain-containing protein 2 | Q9NZN4;Q9NZN4-2 | EHD2 | Extracellular or secreted | 0.857071856 | 1.151968061 | 28.40259341 | 28.29924541 | 28.44193715 | 29.59676234 | 28.7959095 | 28.21117915 | 29.16703237 | 29.22204827 | 30.05594841 |
| EH domain-containing protein 4 | Q9H223 | EHD4 | Extracellular or secreted | 0.720996751 | 1.536941311 | 25.9174788 | 26.16752534 | 26.1393248 | 25.88771818 | 25.57813448 | 26.60484231 | 26.42852618 | 26.84230123 | 27.03967479 |
| Histone-lysine N-methyltransferase E | Q96KQ7-2;Q96KQ7;Q96KQ7- | EHMT2 | Extracellular or secreted | -0.339685727 | 0.730177796 | 23.21886355 | 23.62912001 | 23.51472657 | 23.57137955 | 23.46749709 | 24.32113538 | 23.51135213 | 23.36892949 | 22.96202227 |
| Eukaryotic translation initiation factor | P41567 | EIF1 | Other | 1.191802251 | 3.049074601 | 26.86407862 | 26.88553217 | 26.76989009 | 26.42407874 | 26.58218954 | 27.44063096 | 27.97059727 | 27.81543372 | 28.27257583 |
| Probable RNA-binding protein EIF1AD | Q8N9N8 | EIF1AD | Other | 1.574519557 | 5.075990648 | 23.62410736 | 23.97928144 | 24.02055816 | 24.29193285 | 24.30340619 | 23.82259884 | 25.63843061 | 25.68056642 | 25.42550405 |
| Eukaryotic translation initiation factor | P47813;O14602 | EIF1AX;EIF1AY | Other | -0.369272841 | 2.192723854 | 29.14756258 | 29.17555176 | 29.08159506 | 29.03396425 | 28.99682312 | 28.63944532 | 28.74323225 | 28.55760609 | 28.62881418 |
| Eukaryotic translation initiation factor | Q9BY44;Q9BY44-3;Q9BY44-4 | EIF2A | Extracellular or secreted | 0.263280369 | 1.066494433 | 28.39415264 | 28.32070512 | 28.25619918 | 28.23466433 | 28.22513218 | 28.68849177 | 28.57423404 | 28.79801296 | 28.47726672 |
| Interferon-induced, double-stranded | P19525;P19525-2 | EIF2AK2 | Other | 0.177380313 | 0.252989418 | 25.08836965 | 25.55036516 | 25.63483542 | 25.85281446 | 25.95060324 | 26.68912526 | 25.69003166 | 25.89582856 | 26.32933731 |
| Translation initiation factor eIF-2B sul | Q14232;Q14232-2 | EIF2B1 | Extracellular or secreted | -0.087013389 | 0.564674326 | 27.14501306 | 27.22857652 | 27.18659714 | 27.2216796 | 27.22582171 | 27.33398379 | 27.01998582 | 27.2036851 | 27.18612482 |
| Translation initiation factor eIF-2B sul | P49770 | EIF2B2 | Other | 0.117905963 | 1.103842754 | 27.64732469 | 27.34274056 | 27.34933671 | 27.37674899 | 27.38269724 | 27.27159667 | 27.54495288 | 27.50453401 | 27.48945343 |
| Translation initiation factor eIF-2B sul | Q9NR50-3;Q9NR50-2;Q9NR5 | EIF2B3 | Other | -0.045293474 | 0.100128035 | 27.32628848 | 27.22894343 | 27.40568008 | 27.19507253 | 27.37600374 | 26.87911785 | 27.33253337 | 27.32645994 | 26.91067932 |
| Translation initiation factor eIF-2B sul | Q9UI10;Q9UI10-3;Q9UI10-2 | EIF2B4 | Other | 0.240420666 | 0.81534727 | 26.88913499 | 26.93809101 | 27.05330928 | 26.59192108 | 26.80796273 | 26.61414187 | 27.22921855 | 27.09486602 | 26.84445791 |
| Translation initiation factor eIF-2B sul | Q13144 | EIF2B5 | Other | 0.339319076 | 2.458710301 | 26.71148487 | 27.11495583 | 27.01510108 | 27.1375084 | 27.10699337 | 27.16758276 | 27.44576975 | 27.3378161 | 27.36118453 |
| Eukaryotic translation initiation factor | P41214-2;P41214 | EIF2D | Other | -0.845078741 | 1.318404739 | 22.34470226 | 22.09145605 | 21.83437537 | 22.12019104 | 22.2250862 | 21.21673402 | 20.84176396 | 21.61791074 | 20.92136155 |
| Eukaryotic translation initiation factor | P05198 | EIF2S1 | Extracellular or secreted | -0.022242108 | 0.044777982 | 30.90203401 | 30.93592951 | 30.92662904 | 30.63495309 | 30.67505661 | 30.27774985 | 30.83703079 | 30.83951075 | 30.43290819 |
| Eukaryotic translation initiation factor | P20042 | EIF2S2 | Other | -0.649243258 | 2.200186558 | 30.50322642 | 30.51681282 | 30.58718873 | 30.40059981 | 30.47701575 | 29.63166144 | 29.56797568 | 29.85372181 | 29.68882522 |
| Eukaryotic translation initiation factor | P41091;Q2VIR3;Q2VIR3-2 | EIF2S3;EIF2S3L | Extracellular or secreted | -0.078342536 | 0.298807957 | 31.44815504 | 31.47281042 | 31.42196822 | 31.2440545 | 31.34936835 | 31.1832285 | 31.31403049 | 31.41099519 | 31.09973923 |
| Eukaryotic translation initiation factor | Q14152;Q14152-2 | EIF3A | Other | 0.280988775 | 1.125516185 | 31.60774605 | 31.6752249 | 31.65450414 | 31.4889844 | 31.50270495 | 31.36848828 | 31.93262484 | 31.91748164 | 31.6416862 |
| Eukaryotic translation initiation factor | P55884;P55884-2 | EIF3B | Extracellular or secreted | 0.074813285 | 0.28508417 | 31.33282144 | 31.40324378 | 31.32254358 | 31.3702045 | 31.40578151 | 31.00202499 | 31.48951086 | 31.44223385 | 31.21100504 |
| Eukaryotic translation initiation factor | Q99613;Q99613-2;B5ME19 | EIF3C;EIF3CL | Other | 0.147043326 | 0.603717115 | 30.89460913 | 30.94020728 | 30.80254969 | 30.67749494 | 30.791181 | 30.67337259 | 31.05756313 | 31.01325172 | 30.76002244 |
| Eukaryotic translation initiation factor | O15371-2;O15371-3;O15371 | EIF3D | Other | 0.302323571 | 1.763099355 | 29.79071506 | 29.82439974 | 29.69494694 | 30.13528362 | 30.17271755 | 29.70259577 | 30.21697079 | 30.23922477 | 30.1111045 |
| Eukaryotic translation initiation factor | P60228 | EIF3E | Extracellular or secreted | 0.408423519 | 2.943221495 | 29.96264942 | 30.11594806 | 30.03315037 | 29.76692424 | 29.83155984 | 29.92450691 | 30.4139732 | 30.37471933 | 30.25394744 |
| Eukaryotic translation initiation factor | O00303 | EIF3F | Other | 0.303490519 | 0.885975985 | 29.3893188 | 29.47465841 | 29.37894113 | 29.27135178 | 29.40391418 | 29.24047637 | 29.7764626 | 29.80330443 | 29.41003486 |
| Eukaryotic translation initiation factor | O75821 | EIF3G | Other | 0.702129441 | 2.647612861 | 29.42566411 | 29.37449134 | 29.45730469 | 29.41711615 | 29.30101324 | 29.31088195 | 30.17080907 | 30.10873892 | 29.97007609 |
| Eukaryotic translation initiation factor | O15372 | EIF3H | Extracellular or secreted | 0.058648057 | 0.17507671 | 29.19450905 | 29.49407813 | 29.40413757 | 29.32433676 | 29.27974344 | 29.01786405 | 29.51703818 | 29.36584233 | 29.15039816 |
| Eukaryotic translation initiation factor | Q13347 | EIF3I | Extracellular or secreted | 0.493865049 | 3.71504691 | 30.06971235 | 29.87550454 | 29.70663558 | 29.9005233 | 30.09244843 | 29.83143903 | 30.43489743 | 30.44726841 | 30.33756093 |
| Eukaryotic translation initiation factor | O75822;O75822-2;O75822-3 | EIF3J | Other | 0.441339081 | 1.519901084 | 29.84354471 | 29.7003153 | 29.68403285 | 30.2414996 | 30.15448419 | 29.29469583 | 30.22974572 | 30.41326711 | 30.14029065 |
| Eukaryotic translation initiation factor | Q9UBQ5;Q9UBQ5-2 | EIF3K | Extracellular or secreted | 0.543171682 | 1.629550631 | 28.88657907 | 28.9517985 | 28.80636554 | 28.45151927 | 28.40527432 | 28.87783155 | 29.43256974 | 29.36473668 | 29.02189275 |
| Eukaryotic translation initiation factor | Q9Y262;Q9Y262-2 | EIF3L | Other | 0.352425039 | 1.395125295 | 30.4201117 | 30.4882662 | 30.35930044 | 30.21164334 | 30.22250897 | 30.26270945 | 30.75724291 | 30.78534595 | 30.4969563 |
| Eukaryotic translation initiation factor | Q7L2H7;Q7L2H7-2 | EIF3M | Other | 0.357236459 | 0.573230749 | 29.35392773 | 29.82122463 | 29.70801805 | 29.09748051 | 29.13535702 | 29.15518419 | 30.11830186 | 29.77976897 | 29.30923462 |
| Eukaryotic initiation factor 4A-I | P60842;P60842-2 | EIF4A1 | Extracellular or secreted | 0.793765306 | 0.928749903 | 31.68649782 | 31.84340992 | 31.84430824 | 31.73072875 | 31.67762095 | 32.16311978 | 32.90422533 | 32.95232285 | 31.99759047 |
| Eukaryotic initiation factor 4A-II;Euka | Q14240;Q14240-2 | EIF4A2 | Other | 0.65990919 | 1.649995185 | 28.75340632 | 28.95742791 | 28.8594354 | 28.41381184 | 28.40112899 | 29.14316489 | 29.60362258 | 29.52801925 | 29.11227341 |
| Eukaryotic initiation factor 4A-III;Euka | P38919 | EIF4A3 | Other | 0.242614821 | 0.699724834 | 28.55610817 | 28.5276092 | 28.51747002 | 28.22531609 | 28.28753626 | 29.33951609 | 28.79079273 | 28.78144276 | 28.88238688 |
| Eukaryotic translation initiation factor | P23588;P23588-2 | EIF4B | Other | 0.82191119 | 2.300623909 | 29.76883381 | 29.63514343 | 29.65727565 | 29.71938953 | 29.77461023 | 28.68825831 | 30.50294201 | 30.43817368 | 30.14637336 |
| Eukaryotic translation initiation factor | P06730;P06730-2;P06730-3 | EIF4E | Extracellular or secreted | 0.297085081 | 1.820912517 | 28.30330166 | 28.37077612 | 28.32637421 | 28.12932586 | 28.22554593 | 28.32452991 | 28.61729852 | 28.65112852 | 28.46275506 |
| Eukaryotic translation initiation factor | O60573;O60573-2 | EIF4E2 | Other | -0.427629516 | 1.125208591 | 25.69346548 | 25.78476582 | 25.58181615 | 26.37167363 | 26.1788885 | 25.88033293 | 25.69232178 | 25.5617119 | 25.20854903 |
| Eukaryotic translation initiation factor | Q04637-4;Q04637-5;Q04637 | EIF4G1 | Other | -0.119550348 | 0.250120987 | 30.57992615 | 30.46684395 | 30.49676587 | 30.63365466 | 30.62270139 | 30.79969661 | 30.23591989 | 30.39499077 | 30.81023261 |

| | | | | | | | | | | | | | | |
|---|---|---|---|---|---|---|---|---|---|---|---|---|---|---|
| Eukaryotic translation initiation factor | P78344-2;P78344 | EIF4G2 | Other | 0.991664934 | 1.205568616 | 25.55628362 | 26.35050059 | 26.0238183 | 24.65381599 | 23.98302768 | 26.41813815 | 26.65635525 | 26.68845842 | 26.1229733 |
| Eukaryotic translation initiation factor | Q15056-2;Q15056 | EIF4H | Other | 1.339342271 | 1.981703083 | 28.01903141 | 27.65874026 | 27.7182956 | 27.88959921 | 28.09592244 | 26.7933999 | 29.16065173 | 29.39110265 | 28.55376685 |
| Eukaryotic translation initiation factor | P55010 | EIF5 | Extracellular or secreted | 0.309625144 | 1.024446206 | 29.42576414 | 29.46359325 | 29.29961661 | 29.62712748 | 29.72812739 | 29.34117165 | 29.9119938 | 29.9022497 | 29.5573322 |
| Eukaryotic translation initiation factor | P63241;P63241-2;Q6IS14 | EIF5A;EIF5AL1 | Extracellular or secreted | 0.258264449 | 1.538223608 | 32.55912525 | 32.5443219 | 32.6627603 | 32.69426204 | 32.66065819 | 32.35791196 | 32.82809019 | 32.95442182 | 32.73180115 |
| Eukaryotic translation initiation factor | O60841 | EIF5B | Other | -0.39490053 | 1.951313615 | 30.28239728 | 30.33596509 | 30.26863291 | 30.53073756 | 30.49065881 | 29.82818858 | 29.92620486 | 29.90037934 | 29.85700433 |
| Eukaryotic translation initiation factor | P56537;P56537-2 | EIF6 | Extracellular or secreted | 0.194080632 | 0.664752209 | 30.81367662 | 30.77407606 | 30.79668319 | 30.83906016 | 30.67244555 | 30.06753431 | 30.86363565 | 30.9816893 | 30.71865489 |
| Zinc phosphodiesterase ELAC protein | Q9BQ52;Q9BQ52-4;Q9BQ52- | ELAC2 | Other | 0.14690977 | 0.184254891 | 29.0120397 | 28.8860848 | 28.930159 | 28.91202236 | 29.10259521 | 28.83815857 | 29.34625381 | 29.40196308 | 28.53304225 |
| ELAV-like protein 1 | Q15717;Q15717-2 | ELAVL1 | Other | -0.606084795 | 3.52903684 | 30.20501452 | 30.30508628 | 30.23956623 | 30.36317064 | 30.38689579 | 30.44509879 | 29.74019495 | 29.62665763 | 29.78730916 |
| Engulfment and cell motility protein 3 | Q96BJ8-2;Q96BJ8;Q96BJ8-3 | ELMO3 | Other | 0.360428309 | 0.754560611 | 21.35272453 | 20.77354169 | 21.5625904 | 20.74032429 | 20.78274102 | 21.25315038 | 21.13733641 | 21.74238181 | 21.43410286 |
| Elongator complex protein 2 | Q6IA86-3;Q6IA86;Q6IA86-6;( | ELP2 | Extracellular or secreted | 0.023187602 | 0.043375443 | 27.12858863 | 27.04915973 | 27.16901756 | 26.8811035 | 27.28806456 | 27.14063196 | 27.18290895 | 27.3925568 | 26.82238002 |
| Elongator complex protein 3 | Q9H9T3;Q9H9T3-2;Q9H9T3-4 | ELP3 | Other | 0.165756654 | 0.580284044 | 24.8534333 | 24.89905999 | 25.20791634 | 24.84675481 | 25.32889238 | 24.96547606 | 25.24804018 | 25.31527286 | 24.98472336 |
| Elongator complex protein 4 | Q96EB1;Q96EB1-3;Q96EB1-2 | ELP4 | Other | -0.303099802 | 0.715992106 | 23.86504679 | 23.55180306 | 24.36746381 | 24.5307078 | 24.56424394 | 24.05961446 | 24.10045125 | 23.66438044 | 23.79530883 |
| Elongator complex protein 6 | Q0PNE2-2;Q0PNE2 | ELP6 | Other | 0.223861995 | 0.291993236 | 24.62254431 | 25.07557414 | 24.97526303 | 23.98311469 | 24.45882661 | 24.7571507 | 24.97827789 | 25.25742266 | 24.37212218 |
| ER membrane protein complex subun | Q9NPA0 | EMC7 | Other | 0.883101825 | 0.663024375 | 23.20639676 | 23.30688608 | 21.47001294 | 20.62171647 | 21.5926072 | 24.75322777 | 23.1973355 | 23.47728988 | 23.45010371 |
| Emerin | P50402 | EMD | Other | -0.577117104 | 1.77480502 | 25.14172362 | 24.60941504 | 24.891246 | 24.60342804 | 24.51502748 | 25.41751055 | 24.07496199 | 24.33319904 | 24.44966303 |
| Ribosomal RNA small subunit methylt | Q92979 | EMG1 | Other | 0.491940752 | 1.233067761 | 25.02233733 | 25.11612658 | 25.44515406 | 26.07287873 | 26.04543573 | 26.04666425 | 26.23145427 | 26.07502321 | 26.04364312 |
| Echinoderm microtubule-associated p | O00423;O00423-3 | EML1 | Other | -0.666570089 | 2.244714523 | 27.02506536 | 26.61526502 | 26.4754356 | 26.17820467 | 26.26277217 | 26.21824645 | 25.84570252 | 25.94344433 | 25.59863752 |
| Echinoderm microtubule-associated p | O95834;O95834-2;O95834-3 | EML2 | Other | -0.245858786 | 0.856882972 | 30.76690845 | 30.73658688 | 30.67505661 | 30.46557958 | 30.48884079 | 30.10324578 | 30.25135357 | 30.14321356 | 30.48596556 |
| Echinoderm microtubule-associated p | Q32P44;Q32P44-2 | EML3 | Other | -0.177437767 | 0.705226873 | 27.14870233 | 27.19648026 | 27.11763327 | 26.94167702 | 27.16681695 | 26.48019014 | 26.86455098 | 26.90173199 | 26.72715371 |
| Echinoderm microtubule-associated p | Q9HC35;Q9HC35-2 | EML4 | Other | -0.612411336 | 2.621968381 | 31.12034284 | 31.05549596 | 31.00698086 | 30.70314057 | 30.72175413 | 30.84617794 | 30.42382236 | 30.32291958 | 30.14297021 |
| Protein enabled homolog | Q8N8S7-3;Q8N8S7-2;Q8N8S7 | ENAH | Extracellular or secreted | -0.186795804 | 0.520597524 | 28.41300476 | 28.25696398 | 28.2295853 | 27.92218313 | 27.97355619 | 28.38948292 | 28.15711344 | 28.12607921 | 27.74880808 |
| Endonuclease domain-containing 1 pr | O94919 | ENDOD1 | Extracellular or secreted | -1.561859011 | 1.788193458 | 24.88655582 | 25.06361068 | 24.77996617 | 23.9107708 | 24.22435032 | 23.95491265 | 22.51072458 | 22.67992245 | 23.53385915 |
| Alpha-enolase | P06733;P06733-2 | ENO1 | Extracellular or secreted | 0.146852286 | 0.322478437 | 35.18685537 | 35.34397302 | 35.28636891 | 35.4562021 | 35.51359291 | 34.38785881 | 35.43657372 | 35.44464844 | 35.14676027 |
| Gamma-enolase | P09104;P09104-2 | ENO2 | Extracellular or secreted | -0.294859911 | 0.58270911 | 31.01458244 | 31.15689653 | 31.17003302 | 31.45733405 | 31.60827497 | 30.00611147 | 30.79567731 | 30.68369829 | 30.8426609 |
| Enolase-phosphatase E1 | Q9UHY7 | ENOPH1 | Extracellular or secreted | -0.182112677 | 0.346037488 | 29.0995386 | 29.02086014 | 28.94371818 | 29.37635571 | 29.45140136 | 27.99048029 | 28.91965674 | 28.80359698 | 28.67158539 |
| Mitochondrial enolase superfamily m | Q7L5Y1;Q7L5Y1-4;Q7L5Y1-5; | ENOSF1 | Other | 0.59997467 | 1.068773791 | 26.23803133 | 26.90426122 | 26.5024413 | 26.55184705 | 26.17924928 | 26.11576951 | 27.2333391 | 27.22875999 | 26.58362477 |
| Alpha-endosulfine | O43768-2;O43768;O43768-9 | ENSA | Other | 0.934291027 | 1.170534037 | 27.56383678 | 27.46154564 | 27.59910594 | 27.91285042 | 27.89694884 | 25.89899083 | 28.61178742 | 28.60770196 | 27.75002293 |
| Ectonucleoside triphosphate diphospl | Q9Y5L3;Q9Y5L3-3;Q9Y5L3-2 | ENTPD2 | Extracellular or secreted | -0.645171219 | 0.495847788 | 23.90490432 | 23.52921892 | 23.4867712 | 24.61144272 | 23.88771818 | 23.0660031 | 22.13162592 | 23.69625442 | 23.47963522 |
| Transcription and mRNA export factor | Q9NPA8-2;Q9NPA8 | ENY2 | Other | 1.782415158 | 1.225660291 | 24.8247366 | 20.7790495 | 22.10795826 | 24.2128217 | 24.39819569 | 25.46082732 | 25.57398134 | 25.57940111 | 25.08565755 |
| Histone acetyltransferase p300 | Q09472 | EP300 | Other | -0.565164417 | 0.267760534 | 21.21666709 | 23.35083777 | 23.43084045 | 22.35291307 | 22.82613414 | 24.43612541 | 22.92667314 | 20.96369118 | 23.2209014 |
| Protein 4.1 | P11171-2;P11171;P11171-5;F | EPB41 | Extracellular or secreted | 1.88345403 | 1.175560787 | 24.6690931 | 25.06812911 | 25.55321022 | 25.18861691 | 20.32873524 | 25.11310848 | 26.33971995 | 26.29082617 | 25.9802625 |
| Band 4.1-like protein 1 | Q9H4G0-2;Q9H4G0;Q9H4G0- | EPB41L1 | Extracellular or secreted | 0.042347477 | 0.095479704 | 26.89931359 | 26.78519003 | 26.8059967 | 26.3451288 | 26.8718528 | 27.00987952 | 26.65853599 | 26.74688782 | 27.08029935 |
| Band 4.1-like protein 2 | O43491;O43491-3;O43491-2 | EPB41L2 | Extracellular or secreted | 0.538556964 | 2.916416646 | 28.50039874 | 28.54881562 | 28.46513192 | 28.52723633 | 28.46657171 | 28.49002755 | 29.02149568 | 28.99002037 | 29.10324578 |
| Band 4.1-like protein 2 | O43491-4 | EPB41L2 | Extracellular or secreted | 0.751949893 | 0.954535514 | 23.44012399 | 22.37087255 | 22.51515986 | 21.41698041 | 23.51455067 | 21.95459171 | 23.53978642 | 23.50978273 | 22.81242012 |
| Band 4.1-like protein 3;Band 4.1-like p | Q9Y2J2-2;Q9Y2J2-4;Q9Y2J2;( | EPB41L3 | Extracellular or secreted | 0.341639567 | 0.259655377 | 25.11644391 | 25.12636481 | 25.00680168 | 25.70619752 | 25.60025504 | 23.64987539 | 25.89451083 | 25.71090709 | 24.52246995 |
| Band 4.1-like protein 5 | Q9HCM4-2;Q9HCM4-4;Q9HC | EPB41L5 | Extracellular or secreted | 1.492015729 | 3.584372908 | 22.32292387 | 22.03126774 | 22.75493071 | 21.92875257 | 21.97627742 | 22.94102324 | 23.73748007 | 23.84047753 | 23.87567737 |
| Epithelial cell adhesion molecule | P16422 | EPCAM | Extracellular or secreted | 0.743278697 | 1.585964719 | 30.24036264 | 30.14952627 | 30.27330976 | 30.76002244 | 30.75182732 | 30.8843826 | 31.02128386 | 31.13387609 | 31.60439166 |
| Mammalian ependymin-related prote | Q9UM22;Q9UM22-3 | EPDR1 | Extracellular or secreted | -1.180080656 | 1.944394302 | 25.38442753 | 25.94105011 | 25.96549808 | 25.58454257 | 26.17738745 | 24.26112276 | 24.48173046 | 24.64680302 | 23.9882388 |
| Ephrin type-A receptor 1 | P21709;P21709-3;P21709-2 | EPHA1 | Extracellular or secreted | 0.411821767 | 0.370237661 | 23.71156364 | 23.65610971 | 23.65184703 | 24.46457118 | 24.64614381 | 25.05279124 | 23.98910549 | 24.51014505 | 25.32772806 |
| Ephrin type-A receptor 2 | P29317 | EPHA2 | Extracellular or secreted | 2.369734868 | 0.789948322 | 22.98539239 | 23.18812639 | 22.34894855 | 23.25839351 | 23.83720829 | 25.66383777 | 24.52067287 | 25.03812628 | 28.1913589 |
| Ephrin type-A receptor 4 | P54764;P54764-2 | EPHA4 | Extracellular or secreted | 0.500155129 | 1.199180719 | 26.13594408 | 25.94138599 | 25.84373937 | 26.87759756 | 26.97541611 | 26.52278421 | 26.72494428 | 26.82456666 | 27.0993881 |
| Ephrin type-B receptor 2 | P29323-2;P29323-3;P29323 | EPHB2 | Extracellular or secreted | -0.55123107 | 0.674947142 | 26.38314237 | 26.25735072 | 26.38566219 | 26.19304297 | 26.21445312 | 26.84326015 | 25.48888675 | 25.54654337 | 26.44933243 |
| Ephrin type-B receptor 3 | P54753 | EPHB3 | Extracellular or secreted | 0.748726254 | 2.226179791 | 25.62050421 | 25.47602304 | 25.46808778 | 25.81782605 | 25.12234144 | 26.16802292 | 26.12736891 | 26.44044086 | 26.51477171 |
| Ephrin type-B receptor 4 | P54760;P54760-2 | EPHB4 | Extracellular or secreted | 1.131132112 | 2.542539217 | 27.24218135 | 27.17512341 | 27.30321455 | 27.88349434 | 27.83080463 | 28.25493864 | 28.57040266 | 28.60508966 | 29.06278247 |
| Epoxide hydrolase 1 | P07099 | EPHX1 | Other | -2.107351855 | 1.934057034 | 25.86853623 | 25.63142554 | 26.12470949 | 25.99978793 | 26.05845885 | 27.95598799 | 23.59662018 | 23.94010924 | 24.96066803 |
| Epsin-1 | Q9Y6I3-3;Q9Y6I3;Q9Y6I3-1 | EPN1 | Extracellular or secreted | 0.337353 | 1.41284787 | 25.56048794 | 25.34702304 | 25.63433691 | 25.07749053 | 24.84603742 | 25.36743048 | 25.67392517 | 25.68939191 | 25.56514509 |
| Epsin-2 | O95208-2;O95208 | EPN2 | Other | -0.042418886 | 0.070722244 | 24.63089809 | 25.08630568 | 25.1567278 | 25.44742607 | 25.39813044 | 24.71198366 | 25.35039942 | 24.84785412 | 24.89022568 |
| Epsin-3 | Q9H201-2;Q9H201 | EPN3 | Extracellular or secreted | 0.301323307 | 0.813752888 | 22.90459199 | 23.47654845 | 23.46724831 | 23.04878153 | 23.06388218 | 22.80858638 | 23.66979597 | 23.42098305 | 23.19801032 |
| Epiplakin | P58107 | EPPK1 | Other | -0.781292363 | 1.453366733 | 28.83872964 | 28.68932525 | 28.8483736 | 28.52118227 | 28.56048794 | 28.91439106 | 27.81775288 | 27.72318732 | 28.30142761 |
| Bifunctional glutamate/proline--tRNA | P07814 | EPRS | Other | -0.557472276 | 1.298642755 | 31.910422 | 31.94199177 | 31.8982903 | 31.66902719 | 31.64086782 | 31.7527845 | 31.06445382 | 31.13013639 | 31.53968475 |
| Epidermal growth factor receptor sub | P42566 | EPS15 | Extracellular or secreted | -0.053780519 | 0.136347432 | 26.55623976 | 26.51595989 | 26.44782015 | 26.85174275 | 26.75340632 | 26.23304647 | 26.42881367 | 26.35544923 | 26.73350322 |
| Epidermal growth factor receptor sub | Q9UBC2-3;Q9UBC2;Q9UBC2- | EPS15L1 | Extracellular or secreted | -0.165496968 | 2.115322161 | 27.8501337 | 27.95476847 | 27.93887622 | 27.95254851 | 27.8512066 | 27.83334055 | 27.76780832 | 27.67451794 | 27.75161985 |
| Epidermal growth factor receptor kina | Q12929 | EPS8 | Extracellular or secreted | -0.245232466 | 0.122787656 | 23.40080336 | 23.41299669 | 23.15121726 | 22.22484161 | 22.54486186 | 23.52501021 | 23.66687411 | 23.23375599 | 21.493538 |
| Epidermal growth factor receptor kina | Q8TE68;Q8TE68-4;Q8TE68-3; | EPS8L1 | Extracellular or secreted | -0.693664387 | 0.835086433 | 23.24052641 | 22.50978273 | 22.46617805 | 22.77750527 | 21.79137913 | 21.35746208 | 22.23470635 | 21.2235571 | 21.53216023 |
| Epidermal growth factor receptor kina | Q9H6S3;Q9H6S3-3 | EPS8L2 | Extracellular or secreted | 0.861255885 | 0.546421731 | 25.59052263 | 25.67020132 | 25.71465848 | 25.29757598 | 24.54657281 | 25.57375026 | 27.33807123 | 26.17405675 | 25.26828042 |
| Endoplasmic reticulum aminopeptida | Q9NZ08;Q9NZ08-2 | ERAP1 | Extracellular or secreted | -0.325324043 | 0.219378348 | 25.14297021 | 24.77961557 | 24.56302213 | 23.31589473 | 21.69056439 | 25.58106907 | 23.8692896 | 23.82890582 | 23.86240051 |
| Protein LAP2 | Q96RT1-6;Q96RT1-4;Q96RT1 | ERBB2IP | Extracellular or secreted | 0.585427274 | 0.958022046 | 22.81354636 | 22.75340121 | 22.97023301 | 22.64671515 | 23.04355135 | 22.61083471 | 23.41776806 | 23.76809869 | 22.98955596 |
| Receptor tyrosine-protein kinase erbB | P21860-4;P21860;P21860-3 | ERBB3 | Extracellular or secreted | 1.073873717 | 0.610449605 | 20.40753275 | 22.17952645 | 22.4853714 | 22.02755349 | 21.41065577 | 22.89683675 | 23.38890019 | 23.86551883 | 21.67094044 |
| ELKS/Rab6-interacting/CAST family m | Q8IUD2;Q8IUD2-3;Q8IUD2-2 | ERC1 | Extracellular or secreted | 0.213154353 | 0.924842777 | 27.62540396 | 27.90270972 | 27.78419171 | 27.81860636 | 27.82256237 | 27.597615 | 28.00136739 | 27.80495116 | 28.10868907 |
| DNA excision repair protein ERCC-1 | P07992-4;P07992;P07992-2;F | ERCC1 | Other | -0.079677985 | 0.271680654 | 24.38112994 | 24.34820567 | 24.14226913 | 24.43961685 | 24.52306848 | 24.50288132 | 24.276823 | 24.15452765 | 24.49820109 |
| DNA repair endonuclease XPF | Q92889 | ERCC4 | Other | -0.66966206 | 1.161951532 | 25.90037358 | 25.82065267 | 25.71001372 | 25.58772167 | 25.93269523 | 25.25122263 | 25.36996123 | 24.63006489 | 25.09232745 |
| DNA excision repair protein ERCC-6-lik | Q2NKX8 | ERCC6L | Extracellular or secreted | -0.699608233 | 0.164122274 | 24.10453605 | 24.10333584 | 24.75689629 | 24.24583005 | 24.23724731 | 23.06565805 | 25.52746006 | 24.09619397 | 20.53427306 |
| Enhancer of rudimentary homolog | P84090 | ERH | Other | -0.148902065 | 0.282262801 | 30.49371557 | 30.51972101 | 30.47585688 | 30.65010145 | 30.77729382 | 30.64185843 | 30.20932091 | 30.29898332 | 30.82426316 |
| ERI1 exoribonuclease 3 | O43414-3;O43414-2;O43414 | ERI3 | Other | 0.184775 | 0.152162878 | 24.82677423 | 25.19166952 | 25.47892585 | 24.46295003 | 24.65910788 | 24.68709713 | 25.44256213 | 25.51044692 | 24.25457828 |
| Glutamate-rich protein 5 | Q6P6B1 | ERICH5 | Other | -0.738778065 | 0.409930351 | 25.53629228 | 25.26770911 | 25.75959078 | 25.36202112 | 25.43103181 | 21.43241312 | 24.34624535 | 24.62811887 | 23.20383069 |
| ERO1-like protein alpha | Q96HE7 | ERO1L | Extracellular or secreted | -0.075059032 | 0.673171646 | 30.36775956 | 30.57596567 | 30.51079021 | 30.24932033 | 30.3504753 | 30.30236494 | 30.34466755 | 30.28923053 | 30.31926283 |
| ERO1-like protein beta | Q86YB8;Q86YB8-2 | ERO1LB | Extracellular or secreted | -0.285990881 | 0.485873078 | 23.40483597 | 23.14619077 | 23.3607828 | 22.91739685 | 22.79489255 | 21.74801385 | 22.62395113 | 22.77681773 | 22.42731489 |
| Endoplasmic reticulum resident prote | P30040 | ERP29 | Extracellular or secreted | 0.103600521 | 0.254290651 | 28.58348131 | 28.52196849 | 28.39018024 | 28.56641591 | 28.71953641 | 28.59145746 | 28.48876419 | 28.55340067 | 28.95515661 |
| Endoplasmic reticulum resident prote | Q9BS26 | ERP44 | Extracellular or secreted | -0.19976892 | 0.509230424 | 29.64838803 | 29.68752434 | 29.63032532 | 30.04152978 | 30.12102253 | 30.68570447 | 29.75222622 | 29.6586211 | 29.89709315 |
| N-acetyltransferase ESCO1 | Q5FWF5-2;Q5FWF5 | ESCO1 | Other | 1.726215579 | 1.037422646 | 23.23128213 | 23.00224984 | 19.35565663 | 23.73964466 | 19.78069551 | 21.42028662 | 24.22530689 | 22.68415083 | 23.53409671 |
| S-formylglutathione hydrolase | P10768 | ESD | Extracellular or secreted | 0.542136969 | 1.376512631 | 29.06499338 | 29.26776268 | 29.29338078 | 29.01219959 | 29.01884576 | 29.62691868 | 29.54373674 | 29.67850271 | 30.0462219 |
| ESF1 homolog | Q9H501 | ESF1 | Extracellular or secreted | 1.933381619 | 1.432201341 | 22.22643924 | 22.37422018 | 22.52217059 | 20.58892974 | 20.56147839 | 23.19172975 | 23.39506025 | 23.2598306 | 24.87773796 |
| Epithelial splicing regulatory protein 1 | Q6NXG1-2;Q6NXG1-5;Q6NX( | ESRP1 | Extracellular or secreted | -0.086996272 | 0.217974301 | 26.61231487 | 26.57773123 | 26.73169192 | 26.89041124 | 26.84947764 | 27.44671645 | 26.87302706 | 26.84565467 | 26.57450114 |
| Steroid hormone receptor ERR1 | P11474-2;P11474 | ESRRA | Extracellular or secreted | -0.101751059 | 0.101681305 | 21.5274201 | 19.84633207 | 20.37411403 | 22.01674807 | 19.95162622 | 20.29445921 | 20.58242502 | 20.52229034 | 20.59538132 |
| Extended synaptotagmin-1 | Q9BSJ8;Q9BSJ8-2 | ESYT1 | Extracellular or secreted | -1.405820958 | 0.882727669 | 26.47835502 | 26.37351676 | 26.43348535 | 26.38327423 | 26.16616569 | 26.66139321 | 24.41828294 | 24.46070235 | 26.15164696 |
| Eukaryotic peptide chain release facto | P62495;P62495-2 | ETF1 | Other | 0.051030252 | 0.055822079 | 29.78236487 | 29.85992409 | 29.88980226 | 29.56978725 | 29.77723111 | 28.93547995 | 29.94125725 | 29.97937412 | 29.13975415 |
| Electron transfer flavoprotein subunit | P13804-2;P13804 | ETFA | Extracellular or secreted | -0.467088811 | 0.719305192 | 27.7866239 | 27.65259938 | 27.90368679 | 27.89752597 | 27.62002871 | 29.2461065 | 27.26508175 | 27.45002503 | 27.93691241 |
| Electron transfer flavoprotein subunit | P38117;P38117-2 | ETFB | Extracellular or secreted | -0.27569491 | 0.435452061 | 27.73634495 | 27.65403459 | 27.80925148 | 28.06018161 | 28.00442468 | 29.27315411 | 27.57693879 | 27.78163036 | 28.08304183 |
| Persulfide dioxygenase ETHE1, mitoch | O95571 | ETHE1 | Other | -0.074751818 | 0.155930311 | 26.85317152 | 26.81738694 | 26.90276721 | 27.0155265 | 27.15460491 | 27.7991562 | 26.86549523 | 26.95692964 | 27.22462632 |
| Ethanolamine kinase 1 | Q9HBU6;Q9HBU6-2 | ETNK1 | Extracellular or secreted | 1.465474465 | 1.244152955 | 20.06797942 | 20.93921042 | 22.7530543 | 23.14570892 | 22.69536255 | 23.22515977 | 23.80143998 | 24.09796267 | 22.91025844 |
| RNA-binding protein EWS | Q01844-6;Q01844-3;Q01844 | EWSR1 | Extracellular or secreted | -0.810982176 | 1.450561994 | 29.4508903 | 29.66879901 | 29.57547886 | 29.82070744 | 29.78757383 | 29.37212633 | 28.6873241 | 28.5476395 | 29.16987776 |
| Exocyst complex component 1 | Q9NV70-2;Q9NV70 | EXOC1 | Extracellular or secreted | 0.270911997 | 0.411635876 | 23.91761537 | 24.95828012 | 25.1208792 | 23.99213481 | 24.73608684 | 25.15630347 | 24.84914354 | 24.63771229 | 25.26653007 |
| Exocyst complex component 2 | Q96KP1 | EXOC2 | Extracellular or secreted | 1.427228713 | 0.612530038 | 25.67117368 | 25.83373865 | 25.35793397 | 19.58672045 | 21.69606599 | 25.45951467 | 25.21623075 | 25.58359608 | 25.284433 |
| Exocyst complex component 3 | O60645-2;O60645-3;O60645 | EXOC3 | Other | 0.213791869 | 0.421611174 | 22.79453565 | 23.36959522 | 22.91522803 | 22.7236951 | 22.82246269 | 22.64577012 | 23.33742482 | 23.20641166 | 22.73318252 |
| Exocyst complex component 4 | Q96A65;Q96A65-2 | EXOC4 | Extracellular or secreted | -0.350017071 | 0.236536965 | 26.42138442 | 26.27992494 | 26.31148406 | 25.52710207 | 25.92994462 | 26.32659708 | 25.88855449 | 26.64003173 | 24.81958115 |
| Exocyst complex component 5 | O00471 | EXOC5 | Other | -0.031638523 | 0.033340835 | 25.89307614 | 26.16823338 | 25.97316202 | 25.97425668 | 25.63588725 | 25.20489796 | 25.21674882 | 26.10868907 | 26.00440325 |
| Exocyst complex component 7 | Q9UPT5-2;Q9UPT5-5;Q9UPT | EXOC7 | Extracellular or secreted | -0.146889274 | 0.425078548 | 26.30358038 | 25.65549228 | 25.58534517 | 25.73544137 | 25.69659935 | 26.2761308 | 25.72218424 | 25.87546646 | 25.59297615 |
| Exocyst complex component 8 | Q8IYI6 | EXOC8 | Extracellular or secreted | 0.743854984 | 1.198931181 | 24.55561096 | 23.33742482 | 23.78416674 | 23.12659295 | 22.28384492 | 23.6070458 | 24.06480328 | 24.08001464 | 24.43409013 |
| Exosome complex component CSL4 | Q9Y3B2 | EXOSC1 | Other | -0.576649175 | 1.137367694 | 25.11410197 | 25.65201122 | 25.59457155 | 24.86188013 | 24.30542555 | 24.38890019 | 24.53647015 | 24.13320243 | 24.5588252 |
| Exosome component 10 | Q01780;Q01780-2 | EXOSC10 | Other | -1.193208578 | 2.189163793 | 25.13104462 | 25.11688013 | 25.35228675 | 26.87208773 | 25.58757862 | 25.18275744 | 24.29024614 | 24.15329101 | 24.59815475 |
| Exosome complex component RRP4 | Q13868;Q13868-2;Q13868-3 | EXOSC2 | Other | -0.206552164 | 0.477454902 | 27.74496499 | 27.75442616 | 27.66566847 | 27.36327509 | 27.60272038 | 27.20685504 | 27.41956934 | 27.58348131 | 27.04624792 |
| Exosome complex component RRP40 | Q9NQT5;Q9NQT5-2 | EXOSC3 | Other | -0.367459285 | 0.440361465 | 25.87424727 | 25.88732308 | 25.381295 | 24.64636358 | 24.13688288 | 25.02512875 | 24.52743023 | 25.28057992 | 24.56523227 |
| Exosome complex component RRP41 | Q9NPD3 | EXOSC4 | Other | -0.213630527 | 0.773344329 | 27.14228861 | 27.09828401 | 27.025171 | 26.5710757 | 26.48575832 | 26.48303844 | 26.64196176 | 26.60808987 | 26.51186483 |
| Exosome complex component RRP46 | Q9NQT4 | EXOSC5 | Other | -0.189292748 | 0.288019594 | 26.33012416 | 26.21154123 | 26.07938403 | 25.39751039 | 25.43793566 | 26.02854721 | 26.10329581 | 25.70385015 | 25.36749714 |
| Exosome complex component MTR3 | Q5RKV6 | EXOSC6 | Other | -0.533339487 | 1.339185938 | 26.75697262 | 27.02062174 | 26.96377964 | 26.40395886 | 26.40381262 | 26.57801928 | 26.39613882 | 25.86940727 | 26.19801782 |
| Exosome complex component RRP42 | Q15024 | EXOSC7 | Other | -0.553222981 | 1.489680387 | 27.93427109 | 28.19708986 | 28.11812855 | 27.33005576 | 27.30399836 | 27.16307778 | 27.25412771 | 27.12809693 | 26.98141712 |
| Exosome complex component RRP43 | Q96B26 | EXOSC8 | Other | -0.161017596 | 0.973130691 | 26.85078946 | 27.06937945 | 26.99410047 | 26.86584916 | 26.83677504 | 26.82274469 | 26.79934151 | 26.61820916 | 26.81921568 |
| Exosome complex component RRP45 | Q06265;Q06265-2;Q06265-3 | EXOSC9 | Extracellular or secreted | -0.378452595 | 1.505422634 | 28.05336108 | 28.12346675 | 28.12455174 | 27.57072836 | 27.5236518 | 27.69365157 | 27.44624318 | 27.61883926 | 27.34426543 |
| Exostosin-1 | Q16394 | EXT1 | Other | 1.032404713 | 0.286709067 | 21.65899883 | 20.98342787 | 19.82733455 | 23.08275596 | 22.22729159 | 21.19868483 | 21.68897591 | 20.82329882 | 25.07418622 |
| Exostosin-2 | Q93063;Q93063-2;Q93063-3 | EXT2 | Extracellular or secreted | 1.08206398 | 0.477708466 | 21.23186804 | 21.30958071 | 21.6095503 | 21.35557693 | 21.54108716 | 27.17845165 | 23.06749738 | 23.10445607 | 24.18729589 |
| Exostosin-like 2;Processed exostosin-l | Q9UBQ6 | EXTL2 | Extracellular or secreted | -0.070989571 | 0.047458181 | 21.1305363 | 22.28124064 | 21.00008459 | 21.47055102 | 22.36652045 | 24.15638063 | 21.69871484 | 21.8754149 | 22.41555836 |
| Histone-lysine N-methyltransferase EZ | Q15910-5;Q15910-3;Q15910 | EZH2 | Other | 0.362676831 | 0.125428945 | 19.16233723 | 20.30930315 | 20.87004257 | 23.09638179 | 22.73501064 | 19.35638319 | 22.68710147 | 21.3361988 | 19.82945951 |
| Ezrin | P15311 | EZR | Extracellular or secreted | 0.273148194 | 1.453826696 | 32.74207996 | 32.79017128 | 32.76556549 | 33.00132041 | 33.08461386 | 32.41339322 | 33.08017226 | 33.1104668 | 33.02737764 |
| Junctional adhesion molecule A | Q9Y624;Q9Y624-2 | F11R | Extracellular or secreted | 0.251901884 | 0.495168201 | 30.05284305 | 29.82500663 | 29.72203111 | 29.5156291 | 29.48949172 | 30.21604569 | 29.80167124 | 29.94907297 | 30.41548509 |
| Fatty acid-binding protein, epidermal | Q01469 | FABP5 | Extracellular or secreted | 0.375875811 | 0.343658703 | 30.84131168 | 30.69278735 | 30.61227971 | 30.92145973 | 30.9267704 | 32.01594515 | 30.8086993 | 31.20308996 | 32.12111519 |
| FAS-associated death domain protein | Q13158 | FADD | Extracellular or secreted | 0.40298989 | 0.482217884 | 25.60453129 | 25.77961557 | 25.65649164 | 26.67935875 | 26.70911979 | 24.97000475 | 26.69922337 | 26.39773886 | 25.81156833 |
| FAS-associated factor 1 | Q9UNN5;Q9UNN5-2 | FAF1 | Extracellular or secreted | -0.175439761 | 0.359568153 | 28.5021074 | 28.65836573 | 28.57805528 | 28.34396905 | 28.3332161 | 27.96614747 | 28.46291104 | 28.31436549 | 27.8873347 |
| FAS-associated factor 2 | Q96CS3 | FAF2 | Other | -0.30475043 | 0.210328366 | 24.66828167 | 24.46238844 | 24.90664842 | 25.26638709 | 24.78516508 | 27.09898672 | 24.26033324 | 24.7433992 | 25.67594498 |
| Fumarylacetoacetase | P16930;P16930-2 | FAH | Extracellular or secreted | -0.866226974 | 1.624317887 | 27.9926967 | 28.19746486 | 28.04853626 | 28.26780732 | 28.4313746 | 27.3742633 | 27.47552837 | 27.28647908 | 26.79538316 |
| Acylpyruvase FAHD1, mitochondrial | Q6P587;Q6P587-2;Q6P587-3 | FAHD1 | Other | -0.274981006 | 0.631651772 | 27.32380011 | 27.63346412 | 27.66309126 | 27.67458528 | 27.85655917 | 26.76445858 | 27.35539882 | 27.33517715 | 26.94246028 |
| Fumarylacetoacetate hydrolase doma | Q96GK7 | FAHD2A | Other | -0.227298039 | 0.331072538 | 27.15170502 | 27.31484085 | 27.20880947 | 27.57210274 | 27.70246367 | 26.14664527 | 27.09083445 | 27.19422722 | 26.58132772 |
| Fas apoptotic inhibitory molecule 1 | Q9NVQ4;Q9NVQ4-3;Q9NVQ4 | FAIM | Other | 0.816410483 | 0.490018292 | 22.60806166 | 22.91590271 | 20.91039577 | 22.89888476 | 22.8787205 | 20.88427488 | 23.55987557 | 23.63422611 | 21.8032499 |
| RNMT-activating mini protein | Q9BTL3 | FAM103A1 | Other | 1.297114634 | 0.590934754 | 22.8828345 | 21.04957703 | 20.51368187 | 22.05133974 | 21.15437867 | 21.76223871 | 24.14615969 | 21.29263805 | 23.15957143 |
| Protein FAM107B | Q9H098;Q9H098-2 | FAM107B | Extracellular or secreted | -0.489235185 | 1.126186185 | 26.02227383 | 24.487078 | 24.74026252 | 24.89541257 | 25.06496769 | 25.0665698 | 24.69826973 | 24.43198823 | 24.54031868 |
| Protein NOXP20 | Q8IWE2;Q8IWE2-2 | FAM114A1 | Other | -1.21244073 | 0.576807512 | 25.81264581 | 26.02538224 | 26.14893502 | 25.46410372 | 25.11267112 | 25.30326682 | 25.50902761 | 24.91067932 | 22.87647324 |
| Protein FAM114A2 | Q9NRY5 | FAM114A2 | Other | -0.487844328 | 0.684977883 | 24.40405634 | 24.71528713 | 24.6696338 | 23.84278077 | 23.66046864 | 23.30507758 | 23.96815982 | 23.72536043 | 23.1415989 |
| Protein FAM118A | Q9NWS6 | FAM118A | Other | -0.31405596 | 0.150386753 | 20.23134159 | 21.54266522 | 19.36576709 | 21.33575083 | 22.76194231 | 20.06789122 | 19.54217415 | 20.52487453 | 21.64346257 |
| Constitutive coactivator of PPAR-gam | Q9NZB2;Q9NZB2-6;Q9NZB2- | FAM120A | Extracellular or secreted | 0.306034529 | 0.679290776 | 28.50725907 | 28.53863991 | 28.52924871 | 28.22747522 | 28.09249884 | 29.52686335 | 28.96587234 | 28.93072299 | 28.73250082 |
| Constitutive coactivator of peroxisom | Q96EK7;Q96EK7-2 | FAM120B | Other | 0.388424145 | 0.722944797 | 22.39861323 | 22.53229028 | 22.1550529 | 20.90545047 | 22.03015656 | 21.93635757 | 22.51130386 | 22.4993817 | 22.13354738 |
| Niban-like protein 1 | Q96TA1-2;Q96TA1 | FAM129B | Extracellular or secreted | 0.519596252 | 1.804222769 | 30.02194568 | 30.20314832 | 30.0916921 | 29.30612992 | 29.51095994 | 29.66378688 | 30.36045212 | 30.40750461 | 30.18966344 |
| Protein FAM134C | Q86VR2 | FAM134C | Other | -0.345462557 | 0.24632825 | 21.26613232 | 21.01780885 | 21.23864731 | 22.78337284 | 20.85048904 | 22.79284906 | 21.95206058 | 20.49289129 | 21.49331016 |
| Protein FAM136A | Q96C01 | FAM136A | Other | 0.963559331 | 0.849011869 | 26.41349713 | 26.26508175 | 26.20267672 | 26.38314237 | 25.93992996 | 25.63632989 | 27.57823528 | 27.47374922 | 26.25902241 |
| Protein FAM160B1 | Q5W0V3-2;Q5W0V3 | FAM160B1 | Other | -0.477257872 | 1.057934433 | 22.34815965 | 22.29139894 | 21.91159387 | 21.98842951 | 21.45903932 | 22.44765938 | 21.48212452 | 21.37856581 | 21.93067639 |
| Soluble lamin-associated protein of 7 | Q9Y6X4 | FAM169A | Other | 0.428677434 | 0.913282982 | 26.26180426 | 26.05287414 | 26.04626874 | 25.64960134 | 25.13617884 | 25.29130056 | 26.37729527 | 26.25961513 | 25.86813584 |
| Protein FAM171A1 | Q5VUB5 | FAM171A1 | Extracellular or secreted | 0.386125993 | 0.359040689 | 20.95048755 | 20.91502738 | 21.28234867 | 21.40036039 | 21.34856272 | 21.80096202 | 20.88680697 | 22.17621212 | 21.94423326 |
| BRISC complex subunit Abro1 | Q15018 | FAM175B | Other | -0.164553061 | 0.241787262 | 28.31125017 | 28.10019055 | 27.96845648 | 27.39214733 | 27.09758097 | 27.32843019 | 27.64883371 | 27.79259342 | 27.16394153 |
| Protein FAM188A | Q9H8M7;Q9H8M7-2 | FAM188A | Other | 0.063008551 | 0.017168562 | 23.05690558 | 22.77378312 | 22.15087651 | 22.69108663 | 21.10698727 | 22.34980789 | 23.49692203 | 23.59605141 | 20.16077572 |
| Protein FAM192A | Q9GZU8 | FAM192A | Other | 1.406118532 | 1.677585656 | 21.4532029 | 24.59291601 | 22.72657971 | 23.5049431 | 23.66719905 | 23.1306755 | 24.43217944 | 24.88292323 | 24.44101106 |
| Glycosaminoglycan xylosylkinase | O75063 | FAM20B | Other | 0.078350812 | 0.037000304 | 23.6042202 | 22.98107736 | 23.35514674 | 23.40950489 | 23.53076732 | 25.86161987 | 23.48922372 | 23.10523971 | 25.0117572 |
| Redox-regulatory protein FAM213A | Q9BRX8-2;Q9BRX8 | FAM213A | Extracellular or secreted | 0.281539952 | 0.125635956 | 21.90608152 | 21.94520948 | 22.44813849 | 21.6501033 | 22.18653305 | 24.5803216 | 22.59627398 | 21.61786739 | 23.98867221 |
| Prostamide/prostaglandin F synthase | Q8TBF2-4;Q8TBF2-2;Q8TBF2- | FAM213B | Extracellular or secreted | -0.164152199 | 0.187883512 | 22.3250946 | 23.55274132 | 23.43504452 | 24.35702787 | 24.51291982 | 23.81382033 | 23.69551123 | 23.53694437 | 23.27341203 |
| WASH complex subunit FAM21A;WAS | Q641Q2;Q641Q2-2;Q9Y4E1-3 | FAM21A;FAM21C | Extracellular or secreted | 0.485933702 | 2.470631449 | 25.88227026 | 25.84455368 | 25.83344913 | 25.82708932 | 25.9853533 | 25.327077 | 26.29468707 | 26.26158909 | 26.2514213 |
| Protein FAM3C | Q92520 | FAM3C | Extracellular or secreted | 0.089010461 | 0.040741845 | 28.08679159 | 28.16590702 | 28.25651415 | 29.56440023 | 29.59464629 | 30.64821658 | 28.33883633 | 28.74887205 | 30.33756093 |
| Protein FAM45A;Protein FAM45B | Q8TCE6-2;Q8TCE6;Q6NSW5 | FAM45A;FAM45E | Other | -0.191525624 | 0.793001156 | 24.81792361 | 25.53067804 | 25.325774 | 25.01847969 | 25.37265361 | 25.16399909 | 24.97963034 | 24.92836403 | 25.13218277 |
| Protein FAM49B | Q9NUQ9;Q9NUQ9-2 | FAM49B | Extracellular or secreted | 0.040844517 | 0.140780787 | 29.53998974 | 29.67264786 | 29.60791356 | 29.19171187 | 29.31557519 | 28.99878779 | 29.43971988 | 29.45430701 | 29.39181967 |
| Protein FAM50A | Q14320 | FAM50A | Other | 0.015316746 | 0.026338487 | 27.713125 | 27.45175506 | 27.44782015 | 28.05154718 | 27.75238575 | 26.65908066 | 27.44600648 | 27.54980722 | 27.58799343 |
| Protein FAM53C | Q9NYF3 | FAM53C | Other | -0.453596911 | 0.211728284 | 21.44682686 | 20.3688914 | 20.86483904 | 23.63067595 | 23.74447747 | 22.19032577 | 20.49994526 | 21.61041596 | 22.65186629 |
| Protein FAM65A | Q6ZS17-2;Q6ZS17;Q6ZS17-3; | FAM65A | Extracellular or secreted | 0.524756816 | 0.588852371 | 21.65202216 | 21.54751123 | 21.63826285 | 22.33780589 | 20.81511119 | 22.54842026 | 21.86849855 | 22.90675851 | 22.06858018 |
| Protein FAM83B | Q5T0W9 | FAM83B | Other | -0.986974407 | 1.434110225 | 25.09188374 | 24.72364303 | 24.53848451 | 23.27639707 | 24.41531987 | 25.66291476 | 23.23015358 | 23.88678836 | 23.77645633 |
| Protein FAM83D | Q9H4H8-2;Q9H4H8 | FAM83D | Other | -1.249808943 | 1.768829826 | 23.66957974 | 23.80783993 | 23.40457614 | 22.4110061 | 22.53900542 | 22.10402408 | 21.24473773 | 22.0915529 | 21.88229825 |
| Protein FAM83H | Q6ZRV2 | FAM83H | Other | 0.28147435 | 0.285385116 | 23.90563894 | 23.38811191 | 23.74386141 | 22.29868798 | 22.69521385 | 24.64355897 | 23.32457283 | 24.10349592 | 23.75389083 |
| Protein FAM84B | Q96KN1 | FAM84B | Extracellular or secreted | 0.044595317 | 0.177189117 | 28.22706202 | 28.21715574 | 28.1797808 | 28.08623481 | 27.99647297 | 28.08182359 | 28.07055706 | 28.34363026 | 28.11386359 |
| Protein FAM91A1 | Q658Y4 | FAM91A1 | Other | -0.072604875 | 0.097823279 | 25.73497645 | 25.70237119 | 25.72595844 | 25.46494503 | 25.59676234 | 25.3880462 | 25.74188313 | 25.80690632 | 25.03992575 |

| | | | | | | | | | | | | | | |
|---|---|---|---|---|---|---|---|---|---|---|---|---|---|---|
| MIP18 family protein FAM96A | Q9H5X1;Q9H5X1-2 | FAM96A | Extracellular or secreted | -0.036878341 | 0.057757694 | 24.27923423 | 24.48971373 | 24.1266799 | 23.82590131 | 24.20179848 | 23.14074117 | 23.99317198 | 23.76445858 | 24.16376882 |
| Protein FAM98A | Q8NCA5-2;Q8NCA5 | FAM98A | Other | -0.281188456 | 0.437701272 | 26.03090586 | 25.93186159 | 25.73182138 | 24.59286213 | 24.74627279 | 25.18763569 | 25.06361068 | 25.34888103 | 24.85462264 |
| Protein FAM98B | Q52LJ0-2;Q52LJ0 | FAM98B | Other | -0.510415258 | 1.078559894 | 26.95338139 | 26.88576488 | 26.88063654 | 27.14763535 | 27.16729563 | 27.196105 | 26.77567799 | 26.61133014 | 26.19715549 |
| Protein FAM98C | Q17RN3 | FAM98C | Other | -0.681545517 | 0.372595069 | 20.48369757 | 23.53527345 | 21.1777261 | 22.42125146 | 22.58055197 | 22.20681383 | 21.93052224 | 22.0935852 | 20.1339132 |
| Fanconi anemia group D2 protein | Q9BXW9;Q9BXW9-1;Q9BXW | FANCD2 | Other | 0.512019807 | 0.650813586 | 25.09856514 | 25.12356542 | 25.12856896 | 23.40909764 | 24.93651933 | 26.02407203 | 25.77399435 | 25.36282379 | 25.25943554 |
| Fanconi anemia group I protein | Q9NVI1;Q9NVI1-2;Q9NVI1-1 | FANCI | Other | -1.48418166 | 3.013399808 | 27.56608924 | 28.01850091 | 27.76306584 | 27.94949048 | 27.23306476 | 28.34451942 | 26.34830699 | 26.02941109 | 26.60710226 |
| FERM, RhoGEF and pleckstrin domain | Q9Y4F1;Q9Y4F1-2 | FARP1 | Extracellular or secreted | 0.629209384 | 0.468423454 | 22.9728466 | 23.2474101 | 23.23828286 | 21.45726129 | 22.21791748 | 23.16035648 | 23.45951467 | 24.12809693 | 22.44705396 |
| FERM, RhoGEF and pleckstrin domain | O94887-3;O94887-2;O94887 | FARP2 | Other | 1.452029873 | 1.166979248 | 21.85629016 | 23.48505195 | 20.49113892 | 23.77334046 | 24.10892831 | 24.40314625 | 24.13899294 | 24.75501224 | 24.52103246 |
| Phenylalanine--tRNA ligase alpha subu | Q9Y285;Q9Y285-2 | FARSA | Other | 0.052471823 | 0.304965939 | 28.83222409 | 29.03270385 | 28.92229657 | 28.9673849 | 28.99749626 | 29.14021751 | 28.93413046 | 29.0401453 | 29.1293013 |
| Phenylalanine--tRNA ligase beta subu | Q9NSD9;Q9NSD9-2 | FARSB | Other | 0.075180626 | 0.555401311 | 29.55213665 | 29.62198566 | 29.56025105 | 29.72982974 | 29.65875728 | 29.89850655 | 29.67736052 | 29.79893998 | 29.75997484 |
| Fatty acid synthase;[Acyl-carrier-prote | P49327 | FASN | Extracellular or secreted | 1.661241671 | 7.212728941 | 33.22201947 | 33.3782642 | 33.27280939 | 33.03290414 | 33.0928107 | 33.25729564 | 34.83257685 | 34.89025721 | 34.88894272 |
| FAST kinase domain-containing protei | Q9NYY8;Q9NYY8-2 | FASTKD2 | Other | -4.528651737 | 1.41147158 | 27.70021607 | 27.93477725 | 28.00635228 | 27.50786395 | 24.29942884 | 28.77196886 | 21.30687739 | 25.16434443 | 22.05312659 |
| Protocadherin Fat 1;Protocadherin Fa | Q14517 | FAT1 | Extracellular or secreted | -0.601542031 | 0.288876289 | 25.04247464 | 24.53273022 | 24.66828167 | 25.66703659 | 25.80734863 | 27.32868698 | 25.10809082 | 23.5798039 | 26.03075856 |
| 40S ribosomal protein S30 | P62861 | FAU | Extracellular or secreted | 0.827432865 | 0.436387347 | 24.93445108 | 25.02736643 | 24.82284192 | 25.93447357 | 25.48888675 | 26.26753053 | 26.91900314 | 26.98304943 | 24.81802117 |
| rRNA 2-O-methyltransferase fibrillarin | P22087 | FBL | Extracellular or secreted | -0.849498118 | 1.831466598 | 26.71528713 | 26.82578003 | 26.66641365 | 27.43029813 | 27.34036533 | 28.06732915 | 26.05506923 | 26.54153033 | 26.3776428 |
| Filamin-binding LIM protein 1 | Q8WUP2-3;Q8WUP2 | FBLIM1 | Other | 1.397442103 | 1.222681903 | 22.077772 | 21.47349851 | 21.31613184 | 24.40717231 | 24.08245314 | 23.94601314 | 24.22831963 | 24.26441969 | 24.35110746 |
| Fibulin-1 | P23142;P23142-2;P23142-3 | FBLN1 | Extracellular or secreted | -0.069004396 | 0.036070229 | 29.03320289 | 28.976591 | 29.00616498 | 30.0126525 | 30.00985283 | 31.50213586 | 29.12262778 | 29.22814069 | 30.71251837 |
| Fibulin-1 | P23142-4 | FBLN1 | Extracellular or secreted | 0.108852908 | 0.055697663 | 26.66668453 | 27.24852886 | 26.81518939 | 27.81592228 | 27.77417034 | 29.0585053 | 27.15624559 | 27.05289486 | 28.80691861 |
| Fibrillin-2 | P35556;P35556-2 | FBN2 | Extracellular or secreted | -1.464056153 | 1.060464767 | 22.40541918 | 22.07311901 | 22.90634147 | 21.83888229 | 23.26399008 | 25.93127549 | 20.82252104 | 21.78923264 | 22.20559163 |
| Fibrosin-1-like protein | Q9HCM7 | FBRSL1 | Other | -5.862722917 | 1.885316887 | 27.62268393 | 27.24816689 | 27.1937574 | 27.19272324 | 26.3832083 | 27.27550931 | 19.89288692 | 21.48027911 | 22.49668976 |
| F-box/LRR-repeat protein 18 | Q96ME1-4;Q96ME1-2;Q96M | FBXL18 | Other | 1.02139412 | 0.773396375 | 23.62488825 | 23.66828167 | 23.73138119 | 21.87817063 | 19.8432781 | 22.87511488 | 23.39322809 | 23.97596269 | 23.50554894 |
| F-box only protein 2 | Q9UK22 | FBXO2 | Extracellular or secreted | 0.323291318 | 1.461056361 | 28.87572423 | 29.05250624 | 28.90512245 | 28.89466694 | 28.94715088 | 28.27550931 | 29.19507253 | 29.15711344 | 29.09302801 |
| F-box only protein 22 | Q8NEZ5;Q8NEZ5-3 | FBXO22 | Other | -0.403802627 | 0.383021273 | 26.63975581 | 26.72416367 | 26.79290366 | 26.76913343 | 26.93404607 | 25.99088058 | 26.64003173 | 26.61764883 | 25.45635316 |
| F-box only protein 3 | Q9UK99-2;Q9UK99;Q9UK99- | FBXO3 | Other | -0.075316681 | 0.153619358 | 23.50154566 | 23.35783332 | 23.91843454 | 23.75246232 | 23.7971905 | 24.5455127 | 23.58989432 | 23.8811035 | 23.73954166 |
| F-BAR domain only protein 2 | Q0JRZ9-2;Q0JRZ9-3;Q0JRZ9 | FCHO2 | Extracellular or secreted | 0.218779167 | 0.216305799 | 21.1427054 | 21.70965096 | 22.60125302 | 21.41022981 | 21.95546261 | 21.05331342 | 21.88364116 | 21.30507758 | 22.40392636 |
| Squalene synthase | P37268;P37268-2;P37268-5;I | FDFT1 | Other | 2.876328014 | 0.992854637 | 22.85308107 | 23.14437639 | 21.80926215 | 21.92062274 | 22.79143947 | 23.81274372 | 24.59912013 | 24.49118274 | 27.70444394 |
| Farnesyl pyrophosphate synthase | P14324-2;P14324 | FDPS | Other | 2.401483686 | 2.860011268 | 31.6658548 | 31.84587895 | 31.79470947 | 31.45606131 | 31.50441088 | 30.97664562 | 33.85480319 | 33.61313643 | 34.35829196 |
| Adrenodoxin, mitochondrial | P10109 | FDX1 | Other | 0.882604557 | 0.966607627 | 23.60602922 | 23.58003401 | 23.51086942 | 24.06488549 | 24.11306872 | 24.93988513 | 25.15722911 | 25.25010234 | 24.14786821 |
| Adrenodoxin-like protein, mitochondr | Q6P4F2-2;Q6P4F2 | FDX1L | Other | 1.070904434 | 0.511531637 | 21.60068607 | 21.62484364 | 21.78927095 | 20.87495973 | 21.58011838 | 19.78883649 | 22.69858768 | 23.39453701 | 20.74894624 |
| NADPH:adrenodoxin oxidoreductase, | P22570;P22570-7;P22570-3;I | FDXR | Other | -1.476535996 | 0.421856141 | 22.41330666 | 22.53718142 | 23.42149679 | 24.35178146 | 24.76086002 | 28.77708998 | 22.34670525 | 21.1935807 | 25.16096422 |
| Flap endonuclease 1 | P39748;P39748-2 | FEN1 | Extracellular or secreted | -0.145974045 | 0.374790276 | 29.39523598 | 29.52151927 | 29.44114566 | 28.74156172 | 28.80372014 | 29.56703277 | 29.20569043 | 29.1463248 | 28.9451704 |
| Fermitin family homolog 1 | Q9BQL6;Q9BQL6-2;Q9BQL6-4 | FERMT1 | Extracellular or secreted | 1.550543635 | 1.758244879 | 23.05538373 | 22.61831559 | 23.67422158 | 24.20538281 | 24.44777287 | 25.27877357 | 24.94904513 | 25.36148576 | 25.98102509 |
| Fermitin family homolog 2 | Q96AC1;Q96AC1-3;Q96AC1-2 | FERMT2 | Extracellular or secreted | 0.494402423 | 0.628920913 | 28.17369467 | 28.14058321 | 28.24993061 | 28.28096039 | 28.29653489 | 27.38154256 | 28.79680694 | 28.95900465 | 27.98901884 |
| FYVE, RhoGEF and PH domain-contain | Q96M96 | FGD4 | Other | 1.513430908 | 1.774264579 | 23.14063196 | 22.11162626 | 20.31662197 | 22.04835854 | 21.93863911 | 23.27611304 | 23.31948249 | 23.66741564 | 23.96939003 |
| Fibroblast growth factor-binding prot | Q14512 | FGFBP1 | Extracellular or secreted | 0.505161666 | 0.243177918 | 26.77868857 | 26.66438044 | 26.60427676 | 25.76825016 | 25.84234919 | 29.74307171 | 26.76471167 | 26.85839825 | 28.59288351 |
| Fibroblast growth factor receptor 4 | P22455-2;P22455;P22455-3 | FGFR4 | Extracellular or secreted | 0.192885207 | 0.041713666 | 21.37409786 | 22.39848276 | 22.78588345 | 22.29004686 | 23.23334642 | 23.32649422 | 22.59299895 | 20.30833941 | 25.38149305 |
| Fumarate hydratase, mitochondrial | P07954-2;P07954 | FH | Extracellular or secreted | -0.112907469 | 0.422447741 | 31.14479433 | 31.1492234 | 31.16632614 | 31.30427042 | 31.32012407 | 31.02379715 | 31.24343039 | 31.07080014 | 30.90131482 |
| Bis(5-adenosyl)-triphosphatase | P49789 | FHIT | Extracellular or secreted | 0.039384777 | 0.026251782 | 24.13782106 | 24.33585863 | 24.3457041 | 24.12028598 | 23.95801455 | 21.65086037 | 23.2424903 | 24.23396074 | 23.91597563 |
| Four and a half LIM domains protein 1 | Q13642-1;Q13642-5;Q13642 | FHL1 | Extracellular or secreted | -0.587249628 | 0.740134449 | 25.95792602 | 26.06880566 | 25.86910129 | 26.36829677 | 26.20254595 | 26.51419984 | 25.12336807 | 25.45735558 | 26.14796523 |
| Four and a half LIM domains protein 2 | Q14192 | FHL2 | Other | 0.25794756 | 0.353958546 | 26.78006632 | 26.81249893 | 26.99291276 | 28.05092475 | 28.06321978 | 28.46419723 | 27.68311263 | 27.77442173 | 27.89821821 |
| Four and a half LIM domains protein 3 | Q13643 | FHL3 | Other | -0.154088156 | 0.214269311 | 25.41544882 | 24.88404192 | 23.67098466 | 24.68013714 | 25.45613379 | 24.65730973 | 24.7029788 | 24.79009669 | 24.42668807 |
| FH1/FH2 domain-containing protein 1 | Q9Y613 | FHOD1 | Other | 1.7234918 | 1.359194742 | 23.34394364 | 22.97344233 | 23.52492234 | 19.66361196 | 21.16949439 | 23.39479865 | 24.32848155 | 24.11314823 | 23.76395229 |
| Acidic fibroblast growth factor intrace | O43427-2;O43427 | FIBP | Other | 0.981999353 | 1.458564416 | 22.85654255 | 22.8501647 | 21.37094152 | 21.84563818 | 20.9224014 | 22.74759477 | 23.11873454 | 23.18519485 | 22.93871024 |
| Filamin A-interacting protein 1-like | Q4L180-2;Q4L180;Q4L180-5; | FILIP1L | Other | 0.040402535 | 0.011795544 | 22.26269027 | 22.6495119 | 22.61855478 | 22.85225721 | 22.68146749 | 22.54170203 | 22.96372232 | 20.65828627 | 24.30229084 |
| Pre-mRNA 3-end-processing factor FII | Q6UN15-5;Q6UN15;Q6UN15 | FIP1L1 | Other | -0.767799453 | 0.765093813 | 26.59077959 | 26.62268393 | 26.53612921 | 27.15267229 | 27.18933354 | 26.8467309 | 25.69386423 | 25.65744603 | 26.81445612 |
| Mitochondrial fission 1 protein | Q9Y3D6 | FIS1 | Other | -0.909394478 | 1.290451139 | 25.78511518 | 25.93761969 | 26.01556903 | 25.94966858 | 25.68335362 | 25.56072116 | 24.61138643 | 25.38089883 | 24.74555493 |
| Peptidyl-prolyl cis-trans isomerase FK | Q96AY3 | FKBP10 | Extracellular or secreted | -0.471410533 | 1.194053972 | 29.03852403 | 28.76663996 | 28.71734796 | 28.6670569 | 28.68612208 | 29.08572846 | 28.46649392 | 28.53856592 | 28.06141825 |
| Peptidyl-prolyl cis-trans isomerase FK | Q9NWM8 | FKBP14 | Extracellular or secreted | -1.408324649 | 0.462984227 | 25.15946364 | 24.80321511 | 25.03887982 | 24.30243031 | 24.31077363 | 23.95313713 | 24.43154198 | 24.2350571 | 20.89237679 |
| FK506-binding protein 15 | Q5T1M5;Q5T1M5-2 | FKBP15 | Other | -0.055211929 | 0.270484714 | 26.13200622 | 26.08523206 | 26.10319575 | 25.8288332 | 26.09295243 | 25.67849935 | 26.00354571 | 25.90513393 | 25.88604408 |
| Peptidyl-prolyl cis-trans isomerase FK | P62942 | FKBP1A | Extracellular or secreted | -0.304763741 | 0.212133491 | 30.42221892 | 30.3936618 | 30.36452798 | 30.61876926 | 30.62130453 | 28.84643203 | 30.39315033 | 30.3699446 | 28.9560711 |
| Peptidyl-prolyl cis-trans isomerase FK | P26885 | FKBP2 | Extracellular or secreted | -0.733504416 | 1.695035101 | 29.07747525 | 29.08266124 | 29.19204123 | 29.50982803 | 29.48087194 | 28.21456429 | 28.52211819 | 28.49281832 | 28.06327122 |
| Peptidyl-prolyl cis-trans isomerase FK | Q00688 | FKBP3 | Other | 0.280401383 | 0.53472528 | 29.15571499 | 29.13584625 | 29.05587085 | 28.86970731 | 28.93977307 | 28.5852377 | 29.46382708 | 29.40645072 | 28.84200144 |
| Peptidyl-prolyl cis-trans isomerase FK | Q02790 | FKBP4 | Extracellular or secreted | -0.101308974 | 0.172519167 | 34.2547641 | 34.27009605 | 34.21306398 | 34.29440279 | 34.28043501 | 33.7691965 | 34.1453563 | 34.38400288 | 33.70769312 |
| Peptidyl-prolyl cis-trans isomerase FK | Q13451;Q13451-2 | FKBP5 | Extracellular or secreted | 0.798283563 | 1.177834064 | 27.90437608 | 28.18669158 | 28.08273736 | 28.60218941 | 28.40263407 | 27.33406906 | 29.19903414 | 29.08889024 | 28.36327509 |
| Peptidyl-prolyl cis-trans isomerase FK | Q14318;Q14318-2;Q14318-3 | FKBP8 | Other | 0.286567516 | 0.11652849 | 22.91936237 | 22.1455115 | 20.48387513 | 20.80121772 | 22.82925747 | 26.51463629 | 22.77583369 | 22.58712648 | 23.34367261 |
| Peptidyl-prolyl cis-trans isomerase FK | O95302;O95302-3 | FKBP9 | Extracellular or secreted | -0.501169408 | 4.444564629 | 28.88463006 | 28.78053571 | 28.73156246 | 28.6577186 | 28.70064603 | 28.86106371 | 28.2729762 | 28.26660156 | 28.2649923 |
| FAD synthase;Molybdenum cofactor b | Q8NFF5-2;Q8NFF5;Q8NFF5-3 | FLAD1 | Extracellular or secreted | 0.019270912 | 0.018455811 | 26.59405893 | 27.15904012 | 27.06650821 | 27.07132454 | 27.05909868 | 26.70596032 | 27.43999722 | 27.08952176 | 26.35628915 |
| Protein flightless-1 homolog | Q13045;Q13045-3 | FLII | Other | 0.071878803 | 0.209259784 | 29.75840303 | 29.90426122 | 29.84988026 | 29.51611774 | 29.57304235 | 29.49642304 | 29.72139561 | 29.58175869 | 29.96154592 |
| Protein flightless-1 homolog | Q13045-2 | FLII | Other | -0.719240771 | 0.692544292 | 22.48155803 | 22.79411915 | 22.28217896 | 23.14934948 | 22.69227167 | 23.60203325 | 22.43941394 | 22.581253 | 21.32236602 |
| Filamin-A | P21333-2;P21333 | FLNA | Extracellular or secreted | 0.279483714 | 2.357026969 | 34.85122336 | 34.79459328 | 34.7564776 | 34.58093703 | 34.62363914 | 34.46969686 | 34.95770293 | 34.9891373 | 34.92989455 |
| Filamin-B | O75369-2;O75369-9;O75369 | FLNB | Extracellular or secreted | 0.604437237 | 2.185703571 | 27.35623877 | 27.33056872 | 27.29482727 | 26.86301525 | 27.09607329 | 26.97519742 | 27.62540396 | 27.6896585 | 27.95620961 |
| Filamin-B | O75369-8 | FLNB | Extracellular or secreted | 0.571868369 | 2.092246286 | 34.21306398 | 34.23976537 | 34.18164488 | 33.709543 | 33.75483628 | 33.51276915 | 34.41481753 | 34.42191808 | 34.68468083 |
| Flotillin-1 | O75955;O75955-2 | FLOT1 | Extracellular or secreted | -0.793388872 | 2.244042168 | 26.94770827 | 26.74740014 | 27.17512341 | 26.56497072 | 26.6191892 | 27.73382642 | 26.23839585 | 26.22285901 | 26.05268761 |
| Flotillin-2 | Q14254 | FLOT2 | Extracellular or secreted | -0.344355122 | 0.566208267 | 25.54968974 | 25.53724068 | 25.90547827 | 25.5142751 | 25.542121 | 26.80427423 | 25.08654866 | 25.57608825 | 25.73083724 |
| Leucine-rich repeat transmembrane p | Q9NZU0 | FLRT3 | Extracellular or secreted | 2.409111 | 1.605082172 | 21.8546249 | 22.98641705 | 22.87582733 | 20.77032168 | 20.65870374 | 21.89057411 | 24.12328912 | 23.33224311 | 25.29003516 |
| Receptor-type tyrosine-protein kinase | P36888-2;P36888 | FLT3 | Extracellular or secreted | 0.251028993 | 0.161287955 | 20.83768952 | 21.05530097 | 19.29031747 | 20.47960843 | 21.55327986 | 22.6614183 | 21.45998986 | 20.47129814 | 21.76060626 |
| Vascular endothelial growth factor re | P35916-3;P35916-1;P35916 | FLT4 | Extracellular or secreted | 0.02258894 | 0.016142465 | 21.71816493 | 21.75401321 | 21.87252459 | 22.39016056 | 23.41222146 | 22.80422006 | 23.04866514 | 21.77019768 | 22.2245564 |
| FLYWCH family member 2 | Q96CP2 | FLYWCH2 | Other | -0.919341022 | 3.03198358 | 27.46013989 | 27.29963845 | 27.13916864 | 27.53704809 | 27.27621956 | 26.58835093 | 26.23308306 | 26.48325376 | 26.17592289 |
| Fragile X mental retardation protein 1 | Q06787-6;Q06787-2;Q06787 | FMR1 | Other | -1.272524393 | 1.78307984 | 25.41583562 | 25.10569531 | 25.31672347 | 25.29193285 | 25.19238463 | 24.38317534 | 23.79520973 | 24.3069556 | 23.4331351 |
| Fibronectin;Anastellin;Ugl-Y1;Ugl-Y2;U | P02751-17;P02751-3;P02751 | FN1 | Extracellular or secreted | -0.786893575 | 1.558864416 | 31.42607419 | 31.39983622 | 31.37119171 | 31.86956759 | 31.86978819 | 30.19999361 | 30.42001127 | 30.45630616 | 30.8312276 |
| Fructosamine-3-kinase | Q9H479 | FN3K | Other | -0.157645781 | 0.04780856 | 23.71009257 | 24.24756946 | 20.32500754 | 22.27984993 | 23.46338667 | 23.69381107 | 23.70377096 | 23.79639852 | 20.8867518 |
| Ketosamine-3-kinase | Q9HA64 | FN3KRP | Other | 0.446916896 | 0.35641936 | 24.57611709 | 24.92134339 | 25.0232261 | 21.8202521 | 24.73288246 | 25.15826975 | 24.83388338 | 25.07610446 | 24.54680829 |
| Formin-binding protein 1 | Q96RU3-4;Q96RU3-3;Q96RU | FNBP1 | Extracellular or secreted | 0.188815695 | 0.205275205 | 24.92985435 | 24.77419548 | 24.78361736 | 24.92917712 | 24.63638521 | 25.22898012 | 24.96521181 | 24.56714885 | 25.67519125 |
| Formin-binding protein 1-like | Q5T0N5-3;Q5T0N5-4;Q5T0N5 | FNBP1L | Extracellular or secreted | 0.321736051 | 0.382809472 | 25.47116167 | 25.8887635 | 25.19411448 | 24.94815403 | 24.57709732 | 24.44764677 | 25.88697437 | 25.42723176 | 24.9144709 |
| Formin-binding protein 4 | Q8N3X1;Q8N3X1-2 | FNBP4 | Other | -1.067648053 | 2.193767218 | 26.24843837 | 26.56668446 | 26.36896278 | 26.38753684 | 26.53212082 | 26.42827058 | 25.19809278 | 25.26792338 | 25.59704662 |
| Protein farnesyltransferase/geranylge | P49354;P49354-2 | FNTA | Other | 0.089260713 | 0.205951009 | 27.58863687 | 27.4912898 | 27.47181287 | 27.47745974 | 27.53638122 | 26.81970296 | 27.59818316 | 27.62442813 | 27.23781258 |
| Protein farnesyltransferase subunit be | P49356;P49356-2 | FNTB | Other | 0.372475266 | 0.502789081 | 24.95322596 | 25.36269004 | 25.07152914 | 24.816313 | 25.1271524 | 24.08472536 | 25.55224295 | 25.53356215 | 24.73943865 |
| Focadhesin | Q5VW36 | FOCAD | Extracellular or secreted | -0.107256165 | 0.146799569 | 25.5133416 | 24.86514121 | 25.14534354 | 25.04034391 | 23.96719248 | 24.87436455 | 25.0295164 | 24.91857102 | 24.43300771 |
| Forkhead box protein K1 | P85037;P85037-2 | FOXK1 | Other | 0.305429003 | 0.146331131 | 25.17020259 | 25.67624098 | 25.39116412 | 21.27143588 | 23.05229609 | 24.59690449 | 24.72925603 | 25.03073751 | 23.73541554 |
| Fucose-1-phosphate guanylyltransfera | O14772 | FPGT | Other | 0.485893485 | 0.743435099 | 21.37941221 | 22.13814928 | 21.8199514 | 21.5868746 | 21.93211517 | 22.78075154 | 22.50212865 | 22.79392077 | 21.98025814 |
| Extracellular matrix protein FRAS1 | Q86XX4-2;Q86XX4;Q86XX4-5 | FRAS1 | Extracellular or secreted | 0.756563929 | 0.561665796 | 26.24259933 | 26.14874111 | 25.21811709 | 27.00731511 | 26.89458022 | 28.43209181 | 26.75595457 | 27.22683241 | 28.25862713 |
| FRAS1-related extracellular matrix pro | Q5SZK8-2;Q5SZK8 | FREM2 | Extracellular or secreted | 1.828495849 | 0.820428504 | 25.86947788 | 20.11425654 | 22.18722036 | 24.29642111 | 24.53290853 | 26.88425158 | 25.46811886 | 25.15007853 | 26.80955815 |
| FERM domain-containing protein 8 | Q9BZ67;Q9BZ67-2;Q9BZ67-3 | FRMD8 | Other | 1.228539716 | 0.891805182 | 26.04297553 | 23.50724395 | 22.26922258 | 25.45481686 | 26.34118014 | 25.94986891 | 26.19932907 | 26.08310271 | 26.18584136 |
| Protein furry homolog-like | O94915 | FRYL | Other | 0.931029027 | 0.779233305 | 21.19418213 | 20.65219727 | 21.58012605 | 20.87087038 | 22.33928767 | 22.86561322 | 23.16723437 | 22.70773574 | 21.66925533 |
| Fascin | Q16658 | FSCN1 | Extracellular or secreted | 0.642507835 | 1.129620416 | 31.98813855 | 32.00051475 | 31.91231509 | 31.92722974 | 31.91755278 | 31.76236104 | 32.32549524 | 32.41573692 | 32.94034732 |
| Fibronectin type III and SPRY domain- | Q9BTV5 | FSD1 | Other | -1.059845419 | 2.14186638 | 24.97022423 | 25.10940665 | 24.62984262 | 25.00260181 | 23.91433403 | 24.92338516 | 23.77041952 | 23.97491307 | 23.35002841 |
| Follistatin-related protein 1 | Q12841;Q12841-2 | FSTL1 | Extracellular or secreted | -0.178667493 | 0.085902983 | 27.7854395 | 27.83575157 | 27.73182138 | 29.41128821 | 29.59421914 | 30.63105427 | 27.91570216 | 28.29509011 | 29.74799228 |
| Follistatin-related protein 3 | O95633-2;O95633 | FSTL3 | Extracellular or secreted | 0.991931635 | 0.542891179 | 24.44922221 | 24.75694717 | 24.56651027 | 26.46704614 | 26.68338039 | 27.40251209 | 26.11060183 | 26.03676472 | 27.9912375 |
| Ferritin heavy chain;Ferritin heavy cha | P02794 | FTH1 | Extracellular or secreted | 1.154901272 | 0.535540453 | 28.97306346 | 28.88852546 | 28.88875769 | 29.66006735 | 29.78247423 | 26.42753549 | 30.72858155 | 30.71136999 | 28.33496412 |
| Ferritin light chain | P02792 | FTL | Extracellular or secreted | 1.049933874 | 0.559624983 | 25.0462271 | 25.50893697 | 25.40132433 | 26.70833057 | 26.81286611 | 23.78384012 | 27.32851579 | 27.2348927 | 25.21715574 |
| Alpha-ketoglutarate-dependent dioxy | Q9C0B1 | FTO | Other | -0.212795895 | 0.313790494 | 28.81099866 | 28.97232404 | 28.92073597 | 28.93041282 | 28.8375572 | 27.96057966 | 28.86552473 | 28.61999374 | 28.09239802 |
| Putative tRNA (cytidine(32)/guanosin | Q9UET6-2;Q9UET6 | FTSJ1 | Other | 1.103427785 | 0.820927439 | 23.41686658 | 20.84502756 | 19.75105246 | 21.65576986 | 23.89605962 | 22.7980414 | 23.15407949 | 23.16120263 | 23.17640997 |
| pre-rRNA processing protein FTSJ3 | Q8IY81 | FTSJ3 | Other | -1.946768382 | 3.062054781 | 26.12149194 | 26.4619515 | 26.32793359 | 25.12648298 | 25.32851579 | 26.39979345 | 23.59092233 | 24.13610059 | 24.31575656 |
| Far upstream element-binding protein | Q96AE4 | FUBP1 | Other | 0.300738997 | 0.443231684 | 30.5788471 | 30.6114005 | 30.49686109 | 30.65770156 | 30.69975289 | 31.00470595 | 31.23609102 | 31.22595957 | 30.46480095 |
| Far upstream element-binding protein | Q96AE4-2 | FUBP1 | Other | 1.222772866 | 1.900712151 | 24.09030952 | 22.67906261 | 24.01137332 | 23.50748593 | 23.93280064 | 25.11965294 | 25.3860077 | 25.11251205 | 24.84014133 |
| Far upstream element-binding protein | Q96I24;Q96I24-2 | FUBP3 | Other | 0.041907601 | 0.065082506 | 26.24394332 | 26.16913229 | 26.42252371 | 26.79327586 | 26.87326179 | 27.56456377 | 26.73750586 | 26.55937966 | 26.86218765 |
| Tissue alpha-L-fucosidase | P04066 | FUCA1 | Extracellular or secreted | -0.965525156 | 2.479413589 | 27.42157704 | 27.35304436 | 27.33534755 | 27.37335081 | 27.25412771 | 26.22221414 | 26.24993965 | 26.12506436 | 26.20825133 |
| Plasma alpha-L-fucosidase | Q9BTY2 | FUCA2 | Extracellular or secreted | 0.147007027 | 0.249233578 | 27.08861228 | 27.32354245 | 27.19882344 | 27.48069863 | 27.69842871 | 26.54738209 | 27.19779291 | 27.19008749 | 27.72188448 |
| L-fucose kinase | Q8N0W3;Q8N0W3-2 | FUK | Other | 0.339646489 | 0.203148754 | 24.19790537 | 23.80960722 | 23.47060327 | 23.34421462 | 20.36919496 | 22.24013332 | 23.55262407 | 23.61728451 | 22.56486027 |
| Furin | P09958 | FURIN | Extracellular or secreted | -0.09264294 | 0.02699285 | 22.02378087 | 21.77776216 | 21.60424147 | 23.22486547 | 23.1932799 | 24.44764677 | 20.777669 | 22.74951921 | 24.33067129 |
| RNA-binding protein FUS | P35637-2;P35637 | FUS | Other | 0.218699181 | 1.512684181 | 31.78483913 | 31.69718619 | 31.71608073 | 31.95521898 | 31.98644417 | 31.93540267 | 32.09565841 | 32.1398639 | 31.95816117 |
| Frataxin, mitochondrial;Frataxin inter | Q16595;Q16595-2;Q16595-3 | FXN | Extracellular or secreted | -0.268417526 | 0.291770324 | 26.55404508 | 26.4031625 | 26.55889816 | 27.07357349 | 27.30147122 | 25.22482868 | 26.47963522 | 26.51882853 | 25.75427323 |
| Fragile X mental retardation syndrom | P51114;P51114-2;P51114-3 | FXR1 | Extracellular or secreted | -0.214722916 | 1.251638339 | 27.97771035 | 28.049731 | 28.08024851 | 28.2673162 | 28.23954347 | 28.56499978 | 27.9714198 | 27.97792866 | 27.99625745 |
| Fragile X mental retardation syndrom | P51116 | FXR2 | Extracellular or secreted | -0.327212985 | 1.060572738 | 26.21547183 | 25.60416363 | 25.31572201 | 25.70627657 | 25.17283675 | 25.86589635 | 25.29530035 | 25.34627917 | 25.31696509 |
| FXYD domain-containing ion transpor | Q14802;Q14802-2;Q14802-3 | FXYD3 | Extracellular or secreted | 2.203377453 | 3.219456391 | 24.88105681 | 25.111 | 25.1806346 | 24.65670984 | 24.77504999 | 26.60526632 | 27.42133627 | 27.44947412 | 27.34418075 |
| FYVE and coiled-coil domain-containin | Q9BQS8;Q9BQS8-4 | FYCO1 | Other | 0.712239545 | 3.066463689 | 26.00388879 | 26.31193439 | 26.82335227 | 26.45472275 | 26.25239619 | 26.2905977 | 27.19055852 | 26.97891064 | 27.03569552 |
| Frizzled-6 | O60353-2;O60353 | FZD6 | Extracellular or secreted | 0.157559407 | 0.080353087 | 22.91654064 | 21.29579446 | 21.97502567 | 22.58473965 | 22.70790625 | 21.47170509 | 23.01028086 | 22.84837002 | 21.08988322 |
| Frizzled-7;Frizzled-2 | O75084;Q14332 | FZD7;FZD2 | Extracellular or secreted | 1.428700672 | 1.793635569 | 22.26464876 | 21.35250903 | 20.83312949 | 20.46050614 | 22.23279953 | 22.98046735 | 22.81037237 | 22.99101038 | 23.54674942 |
| Ras GTPase-activating protein-binding | Q13283 | G3BP1 | Extracellular or secreted | -0.301857349 | 1.245255812 | 31.0471975 | 31.11010894 | 31.01790386 | 31.07788267 | 31.13436582 | 30.99671539 | 30.61999374 | 30.76477493 | 30.90174638 |
| Ras GTPase-activating protein-binding | Q9UN86-2;Q9UN86 | G3BP2 | Other | -0.538572135 | 1.325735672 | 30.84273582 | 30.78900534 | 30.6718553 | 30.52747871 | 30.52037688 | 30.83906016 | 29.91342122 | 30.15375971 | 30.41235877 |
| Glucose-6-phosphate 1-dehydrogenas | P11413;P11413-2;P11413-3 | G6PD | Extracellular or secreted | 0.184376654 | 0.362208461 | 31.19289957 | 31.31516487 | 31.29285443 | 30.30007503 | 30.16410942 | 31.09200729 | 30.98134907 | 31.16997331 | 31.08036289 |
| Lysosomal alpha-glucosidase;76 kDa l | P10253 | GAA | Extracellular or secreted | -1.12523496 | 3.055659714 | 29.36605085 | 29.29335886 | 29.26680259 | 29.21430952 | 29.29524342 | 28.23968003 | 27.99210239 | 27.80986477 | 28.16005059 |
| Gamma-aminobutyric acid receptor-a | Q9H0R8;O95166;Q9H0R8-2 | GABARAPL1;GAB | Extracellular or secreted | 0.109701905 | 0.242805989 | 25.12990556 | 24.56720689 | 24.78401693 | 25.29775088 | 25.42120784 | 24.71565371 | 24.93359593 | 25.3265628 | 25.02681789 |
| Gamma-aminobutyric acid receptor-a | P60520 | GABARAPL2 | Other | 0.495514947 | 1.414796378 | 26.05438621 | 26.15923264 | 26.03212579 | 26.44988338 | 26.7652177 | 25.5530051 | 26.85412324 | 26.61554568 | 26.52380134 |
| GA-binding protein alpha chain | Q06546 | GABPA | Other | 0.348786882 | 0.533081708 | 26.0626847 | 26.24454226 | 26.34034835 | 26.23690075 | 26.46268486 | 25.71905327 | 26.84062159 | 26.70107586 | 26.0377703 |
| Growth arrest and DNA damage-induc | Q8TAE8 | GADD45GIP1 | Other | -1.351173573 | 1.342656241 | 22.61085724 | 22.79637871 | 22.96469199 | 22.6371042 | 23.44385158 | 24.73262373 | 22.16327745 | 22.29734508 | 21.07861047 |
| Cyclin-G-associated kinase | O14976-2;O14976 | GAK | Other | -0.374792715 | 0.470138971 | 20.58930003 | 21.80774168 | 21.36618345 | 22.28567191 | 22.38882139 | 21.77912459 | 21.55925623 | 20.84567381 | 21.57911334 |
| UDP-glucose 4-epimerase | Q14376;Q14376-2 | GALE | Extracellular or secreted | 0.793388377 | 3.142763847 | 25.53587716 | 25.48649503 | 25.5030937 | 25.15309769 | 25.22596876 | 25.73559631 | 26.15296235 | 26.14285338 | 26.40441372 |
| Galactokinase | P51570;P51570-2 | GALK1 | Extracellular or secreted | -0.182227657 | 0.34816987 | 25.85031257 | 25.45013518 | 25.61610682 | 25.69930281 | 25.80378171 | 26.30445104 | 25.70646103 | 25.85457509 | 25.25432598 |
| N-acetylgalactosamine kinase | Q01415;Q01415-2 | GALK2 | Other | 0.127339241 | 0.419921886 | 26.14751891 | 26.10797113 | 26.2642228 | 26.05626891 | 25.56569713 | 25.82708932 | 26.27143638 | 26.12324965 | 25.97171579 |
| N-acetylgalactosamine-6-sulfatase | P34059 | GALNS | Extracellular or secreted | -0.197409064 | 0.223072177 | 25.57181351 | 26.46295003 | 26.13828993 | 26.21741463 | 26.4048522 | 25.29098432 | 26.15069836 | 25.24919825 | 26.0510285 |
| Polypeptide N-acetylgalactosaminyltr | Q8IXK2;Q8IXK2-2 | GALNT12 | Extracellular or secreted | -1.31720056 | 0.356834363 | 23.77082276 | 20.3695512 | 23.85424216 | 23.32251134 | 23.83296648 | 25.90616671 | 20.13383751 | 21.80840963 | 24.63428151 |
| Polypeptide N-acetylgalactosaminyltr | Q10471;Q10471-2 | GALNT2 | Extracellular or secreted | -1.625093847 | 1.413695819 | 25.6292034 | 25.510628 | 25.31806915 | 25.86961907 | 25.80749604 | 27.31388998 | 23.61717239 | 24.12083966 | 25.11115924 |
| Polypeptide N-acetylgalactosaminyltr | Q14435 | GALNT3 | Extracellular or secreted | -1.211569565 | 0.877465119 | 25.69893205 | 25.66836283 | 25.34709064 | 25.27990723 | 25.22864991 | 25.92628405 | 23.49289471 | 24.21103986 | 25.23597009 |
| N-acetylgalactosaminyltransferase 7 | Q86SF2 | GALNT7 | Extracellular or secreted | -1.358238172 | 0.360701776 | 24.96049128 | 24.67427547 | 24.80582455 | 25.38419695 | 25.10293554 | 25.48943812 | 26.04685156 | 21.19077043 | 23.89624444 |
| Galactose-1-phosphate uridylyltransfe | P07902 | GALT | Other | -0.917865491 | 0.427158231 | 23.8754899 | 24.12470949 | 23.900673 | 23.08441719 | 23.57669377 | 21.63588219 | 23.25537095 | 23.22793787 | 20.86202748 |
| Guanidinoacetate N-methyltransferas | Q14353;Q14353-2 | GAMT | Extracellular or secreted | -0.302574965 | 0.430990171 | 23.84555896 | 23.70355976 | 23.87126531 | 23.30702511 | 23.28305556 | 22.75309512 | 23.11197108 | 23.60342804 | 22.7586559 |
| Gigaxonin | Q9H2C0 | GAN | Other | -0.158393256 | 0.306024439 | 26.44167601 | 26.36919582 | 26.1339667 | 26.47382662 | 26.60583147 | 25.44430113 | 26.23706492 | 26.2061098 | 25.81604439 |
| Neutral alpha-glucosidase AB | Q14697 | GANAB | Extracellular or secreted | 0.152609495 | 0.449648913 | 32.31497587 | 32.26483575 | 32.22256655 | 32.21509105 | 32.16299978 | 32.92048398 | 32.43397774 | 32.3746157 | 32.69971153 |
| Neutral alpha-glucosidase AB | Q14697-2 | GANAB | Extracellular or secreted | -0.010842675 | 0.008860916 | 23.33292598 | 23.25436202 | 23.19242226 | 23.42470349 | 23.50263857 | 25.49240578 | 23.30033692 | 23.80055159 | 23.96631252 |
| Neutral alpha-glucosidase C | Q8TET4 | GANC | Other | -1.100386026 | 0.431706633 | 22.88496419 | 23.98745833 | 23.9767494 | 23.42483161 | 23.77685788 | 24.06043933 | 23.28107539 | 23.78546444 | 20.68795246 |
| Glyceraldehyde-3-phosphate dehydro | P04406 | GAPDH | Extracellular or secreted | -0.134542528 | 0.340610312 | 35.66004715 | 35.8937642 | 35.87737815 | 35.94800691 | 36.0437026 | 35.50613253 | 35.58947313 | 35.51112022 | 35.96029483 |
| Glyceraldehyde-3-phosphate dehydro | P04406-2 | GAPDH | Extracellular or secreted | -0.195343611 | 0.165537125 | 27.68130394 | 28.41175289 | 28.20508446 | 27.01137332 | 26.80156332 | 28.55906598 | 27.09224678 | 27.43069692 | 28.22609742 |
| GTPase-activating protein and VPS9 d | Q14C86-3;Q14C86-5;Q14C86 | GAPVD1 | Other | -0.148838216 | 1.709297822 | 27.31941358 | 27.50574578 | 27.32345655 | 27.29096675 | 27.35976122 | 27.54863191 | 27.21141126 | 27.23744792 | 27.27861407 |
| H/ACA ribonucleoprotein complex sul | Q9NY12-2;Q9NY12 | GAR1 | Other | -0.472770019 | 1.494389733 | 25.84954923 | 25.88992408 | 25.78179292 | 26.66112134 | 26.56310944 | 26.4129321 | 25.76499 | 25.76118984 | 25.63472465 |
| Glycine--tRNA ligase | P41250 | GARS | Extracellular or secreted | 0.060602946 | 0.081563855 | 31.4971943 | 31.47633986 | 31.43370421 | 31.59690449 | 31.4576276 | 30.52458613 | 31.61258731 | 31.58718873 | 30.97521109 |
| Trifunctional purine biosynthetic prot | P22102 | GART | Extracellular or secreted | 0.4888362 | 0.563573537 | 31.77768574 | 31.86908952 | 31.85504463 | 31.93985712 | 31.90662836 | 31.58785935 | 32.60928821 | 32.6688793 | 31.65642344 |
| Transcriptional repressor p66-alpha | Q86YP4-2;Q86YP4;Q86YP4-3 | GATAD2A | Other | -1.298989561 | 0.719299315 | 25.22618932 | 24.13092682 | 24.29452933 | 23.97517555 | 21.42854521 | 24.17409486 | 21.27161453 | 23.46388553 | 22.9822618 |
| Transcriptional repressor p66-beta | Q8WXI9 | GATAD2B | Other | -1.044247602 | 3.163255855 | 26.19734299 | 26.14074897 | 26.32455566 | 25.47935769 | 25.72660817 | 26.50659342 | 25.12407839 | 25.02495972 | 24.90582253 |
| Glycine amidinotransferase, mitochor | P50440-3;P50440-2;P50440 | GATM | Extracellular or secreted | 1.80729439 | 2.649706534 | 24.04092913 | 23.72598444 | 22.86186244 | 23.8622113 | 24.18978596 | 22.10813004 | 25.55256544 | 25.37016084 | 24.89360853 |
| GATS-like protein 2 | A6NHX0 | GATSL2 | Other | 0.132905301 | 0.108983365 | 23.85519318 | 23.3295084 | 23.6623172 | 22.96449811 | 20.86068316 | 23.28531529 | 23.15386309 | 23.18814148 | 23.035469 |
| Glucosylceramidase | P04062-4;P04062-2;P04062;I | GBA | Extracellular or secreted | 0.033269202 | 0.033601335 | 26.71082829 | 26.84337997 | 26.82213686 | 27.3770801 | 27.33543275 | 27.32663137 | 26.75887951 | 26.84852285 | 27.70014991 |
| 1,4-alpha-glucan-branching enzyme | Q04446 | GBE1 | Extracellular or secreted | 1.037262318 | 1.230819787 | 27.38079977 | 27.7269589 | 27.28815259 | 25.90357187 | 25.07034209 | 27.12474892 | 27.50279029 | 28.01950869 | 27.83677504 |

| | | | | | | | | | | | | | | |
|---|---|---|---|---|---|---|---|---|---|---|---|---|---|---|
| Golgi-specific brefeldin A-resistance g | Q92538-3;Q92538-2;Q92538 | GBF1 | Other | -0.502660205 | 0.209988974 | 23.6027487 | 23.13194737 | 22.8752649 | 22.80459427 | 22.71066016 | 22.55966556 | 23.67033641 | 22.85502204 | 20.80910141 |
| Interferon-induced guanylate-binding | P32456;P32455 | GBP2;GBP1 | Other | -0.405504484 | 0.164916387 | 23.02421579 | 23.20883923 | 22.59167574 | 22.38784905 | 21.86098086 | 21.42290862 | 22.49989259 | 20.37533685 | 23.15649175 |
| Grancalcin | P28676 | GCA | Extracellular or secreted | 0.433365621 | 1.279952321 | 26.18044491 | 26.15238218 | 26.26606534 | 25.72616639 | 25.69731546 | 26.28050913 | 26.27412331 | 26.72168895 | 26.4557263 |
| GRIP and coiled-coil domain-containir | Q8IWJ2 | GCC2 | Other | -0.944618835 | 0.981761717 | 25.70640833 | 25.65471753 | 25.28496244 | 25.41438459 | 25.2371014 | 25.39705335 | 24.69646669 | 24.98093796 | 23.83605267 |
| GC-rich sequence DNA-binding factor | P16383;P16383-4;P16383-2;F | GCFC2 | Other | -0.180467444 | 0.147230834 | 23.47432188 | 23.44569083 | 23.18865464 | 23.51605009 | 23.13260641 | 22.97340729 | 23.62455363 | 23.42457536 | 22.27483425 |
| GTP cyclohydrolase 1 feedback regula | P30047;P30047-2 | GCHFR | Other | -1.797834391 | 3.274961439 | 25.00058325 | 24.55929213 | 24.58857969 | 25.80520954 | 25.80928828 | 24.44934817 | 23.15924804 | 23.02013423 | 23.53326509 |
| Glutamate--cysteine ligase catalytic su | P48506 | GCLC | Other | -0.114186546 | 0.180057997 | 28.76398393 | 28.74919183 | 28.65870622 | 28.43968025 | 28.34688782 | 27.92976407 | 28.47390401 | 28.65075201 | 27.9768914 |
| Glutamate--cysteine ligase regulatory | P48507;P48507-2 | GCLM | Other | 0.65530482 | 0.518843007 | 27.58641938 | 27.43061717 | 27.49159563 | 26.690458 | 26.54632258 | 24.22493905 | 27.41134883 | 27.74188313 | 26.7978584 |
| Translational activator GCN1 | Q92616 | GCN1L1 | Other | 0.158938592 | 0.306241078 | 26.67949299 | 26.99020976 | 26.97421291 | 26.70332212 | 26.93977307 | 27.56026927 | 27.20890247 | 27.38187256 | 26.80968081 |
| Glycine cleavage system H protein, mi | P23434 | GCSH | Extracellular or secreted | -0.685625921 | 0.581088039 | 30.67118043 | 30.57027598 | 30.75771978 | 31.46187736 | 31.59454841 | 29.54124598 | 30.50861968 | 30.51671891 | 29.21620762 |
| Guanine deaminase | Q9Y2T3;Q9Y2T3-3;Q9Y2T3-2 | GDA | Extracellular or secreted | -0.56396244 | 0.30443615 | 30.03091638 | 30.14321356 | 30.02762481 | 30.23020398 | 30.31083862 | 26.67276587 | 29.60406463 | 29.44403251 | 27.96779714 |
| Growth/differentiation factor 15 | Q99988 | GDF15 | Extracellular or secreted | -1.448991398 | 0.812718484 | 24.81254789 | 24.50233507 | 24.1562263 | 24.96811586 | 25.77089835 | 22.8836016 | 22.4812131 | 22.28394878 | 24.43472646 |
| Rab GDP dissociation inhibitor alpha | P31150 | GDI1 | Other | -0.228447378 | 0.73527543 | 31.49552747 | 31.53802005 | 31.44623332 | 31.53593647 | 31.63758965 | 30.65497354 | 31.14473356 | 31.15749898 | 31.16656558 |
| Rab GDP dissociation inhibitor beta | P50395;P50395-2 | GDI2 | Extracellular or secreted | 0.106409928 | 0.597933583 | 33.59989959 | 33.78068041 | 33.69239245 | 33.48063123 | 33.53310725 | 33.33989831 | 33.69509219 | 33.76337062 | 33.5740716 |
| GDP-D-glucose phosphorylase 1 | Q6ZNW5 | GDPGP1 | Other | 0.490416579 | 0.304705053 | 22.76935038 | 23.11908297 | 23.24453863 | 22.00563846 | 20.59077538 | 22.34537926 | 23.55894194 | 23.1381024 | 21.81158793 |
| Gem-associated protein 4 | P57678 | GEMIN4 | Extracellular or secreted | 1.262477994 | 1.433269962 | 22.82609534 | 23.54580725 | 22.70678771 | 23.1226574 | 22.95709465 | 23.85109935 | 24.79525928 | 24.71507761 | 23.78186794 |
| Gem-associated protein 5 | Q8TEQ6 | GEMIN5 | Other | 0.116409279 | 0.843681723 | 29.10109277 | 29.31211622 | 29.21196818 | 28.92025326 | 28.90512245 | 29.12885899 | 29.21708638 | 29.24974981 | 29.17209757 |
| Gem-associated protein 6 | Q8WXD5 | GEMIN6 | Other | -0.091875529 | 0.117643619 | 24.7188443 | 24.74237153 | 24.47661025 | 25.97696785 | 25.78745863 | 24.47004465 | 24.98076367 | 25.07398201 | 24.75577634 |
| Gem-associated protein 7 | Q9H840 | GEMIN7 | Other | 1.094444864 | 1.641205146 | 23.26404217 | 21.23030182 | 23.02304839 | 23.48287983 | 23.19767882 | 23.22504206 | 23.95544487 | 23.91688683 | 24.12249943 |
| Golgi to ER traffic protein 4 homolog | Q7L5D6-2;Q7L5D6 | GET4 | Other | -1.15407996 | 1.297018993 | 23.16784687 | 23.51086942 | 23.1517592 | 23.83845916 | 23.97219803 | 22.24645342 | 21.49341297 | 22.34980583 | 22.63833437 |
| FAD-linked sulfhydryl oxidase ALR | P55789;P55789-2 | GFER | Extracellular or secreted | -0.284012265 | 0.507747066 | 24.01350465 | 24.17424729 | 24.26642284 | 24.611499 | 24.89587478 | 23.27526064 | 24.10852956 | 23.68292517 | 23.97491307 |
| Elongation factor G, mitochondrial | Q96RP9;Q96RP9-2 | GFM1 | Other | 0.233002068 | 0.184916146 | 23.62176215 | 24.41956934 | 24.03607297 | 24.1521307 | 23.12012774 | 25.9856573 | 24.57894065 | 23.89300669 | 24.89471898 |
| Ribosome-releasing factor 2, mitocho | Q969S9-5;Q969S9-4;Q969S9- | GFM2 | Other | -3.012614851 | 2.965803759 | 25.73688682 | 26.23182045 | 26.34201144 | 27.04228676 | 27.06568681 | 24.14507139 | 23.72640029 | 22.91999877 | 22.59763822 |
| Glutamine--fructose-6-phosphate ami | Q06210-2;Q06210 | GFPT1 | Extracellular or secreted | -0.094597972 | 0.162930078 | 29.86736715 | 29.95850692 | 30.00034685 | 30.01385069 | 30.0507691 | 29.63299633 | 30.04934152 | 29.99846501 | 29.43031807 |
| ADP-ribosylation factor-binding prote | Q9UJY5;Q9UJY5-4;Q9UJY5-3; | GGA1 | Other | -0.080813278 | 0.239373122 | 26.10850962 | 26.46323074 | 26.61076714 | 26.73969616 | 26.97552544 | 26.58075288 | 26.49739419 | 26.37658342 | 26.62282355 |
| Gamma-glutamylaminecyclotransfera | Q9BVM4 | GGACT | Extracellular or secreted | -1.48418523 | 3.170401651 | 21.07982518 | 22.42403708 | 22.27062042 | 22.21865665 | 22.3515119 | 22.34152374 | 20.54234163 | 20.67542656 | 20.67276361 |
| Gamma-glutamylcyclotransferase | O75223;O75223-3 | GGCT | Extracellular or secreted | -0.372405214 | 1.503480662 | 30.98073644 | 31.0663799 | 30.88925106 | 31.13356992 | 31.10017801 | 30.30639071 | 30.56447291 | 30.47807723 | 30.57848723 |
| Gamma-glutamyl hydrolase | Q92820 | GGH | Extracellular or secreted | -0.763291668 | 3.003470694 | 31.65484554 | 31.65407729 | 31.57637122 | 32.12724835 | 32.13412097 | 31.40922566 | 30.986851 | 30.9158446 | 31.08537391 |
| Geranylgeranyl pyrophosphate syntha | O95749;O95749-2 | GGPS1 | Other | -0.402368379 | 1.047278872 | 24.61357996 | 24.3607828 | 24.82366811 | 24.67379041 | 24.44120108 | 24.21711875 | 24.24212682 | 24.29207332 | 23.82376528 |
| Glucose-induced degradation protein | Q8IVV7 | GID4 | Other | -0.172902112 | 0.287503455 | 24.59599452 | 24.66784872 | 24.1978304 | 25.07496199 | 25.13186889 | 23.62008466 | 24.4665017 | 24.18290895 | 24.4761776 |
| Glucose-induced degradation protein | Q9NWU2 | GID8 | Other | 0.487683834 | 1.347620653 | 26.00568862 | 26.06578951 | 26.14308702 | 26.78519003 | 26.99905671 | 25.98317994 | 26.83653429 | 26.68043228 | 26.93708084 |
| PERQ amino acid-rich with GYF domai | Q6Y7W6-4;Q6Y7W6-5;Q6Y7V | GIGYF2 | Other | 0.560224305 | 3.357134654 | 25.2791988 | 25.2534606 | 25.16184845 | 25.66193681 | 25.21959485 | 25.27252243 | 25.97191309 | 25.81714293 | 25.81589785 |
| DNA replication complex GINS protein | Q14691 | GINS1 | Other | 0.377262338 | 0.270753045 | 25.48437597 | 25.57839365 | 25.49780531 | 24.99890612 | 25.08440097 | 22.27634027 | 25.46013989 | 25.45129916 | 24.68045911 |
| DNA replication complex GINS protein | Q9Y248 | GINS2 | Other | -0.18887962 | 0.355655351 | 26.16215588 | 26.39827725 | 26.04780855 | 25.78994749 | 25.65261306 | 25.49646494 | 25.87110077 | 25.94955727 | 25.38633669 |
| DNA replication complex GINS protein | Q9BRX5;Q9BRX5-3;Q9BRX5-2 | GINS3 | Other | -0.099985896 | 0.160116359 | 27.3691126 | 27.04853626 | 26.97344671 | 26.58820794 | 26.68137097 | 26.4207743 | 26.90827588 | 26.96080057 | 26.37169025 |
| DNA replication complex GINS protein | Q9BRT9 | GINS4 | Other | -0.279723519 | 0.947933262 | 26.74611899 | 26.5197285 | 26.31402832 | 26.93427109 | 26.77843793 | 26.07040351 | 26.41204375 | 26.29281056 | 26.13746931 |
| PDZ domain-containing protein GIPC1 | O14908;O14908-2 | GIPC1 | Extracellular or secreted | 0.959528311 | 1.286259442 | 26.79463975 | 25.65957068 | 25.63729771 | 27.16652966 | 26.85388537 | 27.94156509 | 27.36436099 | 27.56993208 | 27.97103601 |
| PDZ domain-containing protein GIPC2 | Q8TF65 | GIPC2 | Extracellular or secreted | 0.940141805 | 0.386343154 | 19.80567032 | 21.59773433 | 21.34114959 | 22.85245255 | 22.79976145 | 21.32339127 | 23.77585378 | 23.05538373 | 20.84926766 |
| ARF GTPase-activating protein GIT1 | Q9Y2X7;Q9Y2X7-3 | GIT1 | Other | -0.111215997 | 0.206890444 | 25.92333982 | 25.78551433 | 25.62279563 | 25.41535211 | 25.36489528 | 25.55224295 | 25.32906352 | 25.86372425 | 25.30563429 |
| Alpha-galactosidase A | P06280 | GLA | Extracellular or secreted | -0.482303348 | 1.60229062 | 27.16432525 | 27.17997058 | 27.0204098 | 27.7416903 | 27.52850371 | 27.49609917 | 26.82553744 | 26.70464182 | 27.08841009 |
| Beta-galactosidase | P16278-3;P16278;P16278-2 | GLB1 | Extracellular or secreted | -0.529688508 | 2.215363169 | 28.58785041 | 28.49636589 | 28.40876017 | 28.16676907 | 28.26941344 | 28.45673698 | 27.88896086 | 27.71194429 | 28.0029773 |
| Golgi apparatus protein 1 | Q92896-2;Q92896;Q92896-3 | GLG1 | Extracellular or secreted | -0.203966084 | 0.399933956 | 27.26936885 | 27.17245528 | 27.07694034 | 28.16408543 | 28.1447214 | 28.03228346 | 27.36310796 | 27.37301885 | 27.58190232 |
| Golgi-associated plant pathogenesis-r | Q9H4G4 | GLIPR2 | Extracellular or secreted | 0.044129475 | 0.057656187 | 23.08849906 | 23.60263544 | 23.56325494 | 22.83483827 | 22.7234868 | 23.55718975 | 23.55730663 | 23.40015189 | 22.85988204 |
| Glomulin | Q92990;Q92990-2 | GLMN | Other | 2.368916084 | 1.734559246 | 24.0752885 | 20.93665118 | 21.04484778 | 22.69230049 | 21.2878106 | 23.15045821 | 24.3648619 | 25.53165988 | 23.80390486 |
| Lactoylglutathione lyase | Q04760;Q04760-2 | GLO1 | Extracellular or secreted | 0.315295403 | 0.928274904 | 30.78986046 | 30.76374655 | 30.71513981 | 31.22750965 | 31.5420398 | 30.76121203 | 31.16200938 | 31.4978605 | 31.18577048 |
| Glyoxalase domain-containing protein | Q9HC38-2;Q9HC38;Q9HC38- | GLOD4 | Extracellular or secreted | -0.341292429 | 1.26525466 | 31.49013278 | 31.43400261 | 31.34667651 | 31.35620728 | 31.45933878 | 30.61052076 | 30.92379883 | 30.9658173 | 30.93494593 |
| Glutaredoxin-3 | O76003 | GLRX3 | Extracellular or secreted | 0.520182528 | 1.319170302 | 29.75697262 | 29.6378919 | 29.59281224 | 29.80728721 | 29.57704688 | 29.70370166 | 30.21593001 | 30.41679411 | 29.96567972 |
| Glutaminase kidney isoform, mitocho | O94925-3;O94925 | GLS | Extracellular or secreted | 0.989049085 | 0.338375719 | 22.09239197 | 20.71280237 | 21.23087185 | 23.93274028 | 23.36733049 | 26.89504269 | 23.34407914 | 23.02441872 | 25.71423923 |
| Glycolipid transfer protein | Q9NZD2 | GLTP | Extracellular or secreted | -0.466831296 | 0.656223139 | 25.60416363 | 25.75608187 | 25.68565435 | 25.27976557 | 25.31962031 | 25.50391258 | 25.09168202 | 25.50884632 | 24.57357693 |
| Glutamate dehydrogenase 1, mitocho | P00367;P00367-3;P00367-2;F | GLUD1;GLUD2 | Extracellular or secreted | -0.032288542 | 0.024772863 | 29.08580442 | 29.15310736 | 29.08547521 | 29.54316517 | 29.65357343 | 30.6602544 | 29.13985171 | 29.13711748 | 30.21685518 |
| Glutamine synthetase | P15104 | GLUL | Extracellular or secreted | 0.700849477 | 0.936678523 | 27.26149942 | 27.26749481 | 27.20853043 | 26.08790451 | 25.95418051 | 26.66125728 | 27.79041989 | 27.63844442 | 26.89411759 |
| Putative oxidoreductase GLYR1 | Q49A26-5;Q49A26-4;Q49A26 | GLYR1 | Other | -1.522925202 | 1.870436962 | 26.01820375 | 25.79071817 | 26.07657343 | 23.80508651 | 24.34346931 | 26.12506436 | 23.72108917 | 24.03825189 | 23.75144109 |
| Ganglioside GM2 activator;Gangliosid | P17900 | GM2A | Extracellular or secreted | 0.024771929 | 0.016269137 | 26.52507174 | 26.1982989 | 26.45211654 | 26.56648124 | 26.99948688 | 26.64265043 | 26.15319436 | 26.08660939 | 27.52656491 |
| GDP-mannose 4,6 dehydratase | O60547-2;O60547 | GMDS | Extracellular or secreted | 0.267794401 | 1.504302311 | 28.33883633 | 28.05894391 | 28.20036786 | 27.88436805 | 27.98315819 | 28.37289435 | 28.29635984 | 28.49365832 | 28.43264938 |
| Glia maturation factor beta | P60983 | GMFB | Other | 0.051809591 | 0.068409581 | 29.45548332 | 29.53934272 | 29.43089627 | 29.58906567 | 29.61380826 | 28.6677337 | 29.5746491 | 29.72535068 | 29.00359396 |
| GEM-interacting protein | Q9P107-2;Q9P107 | GMIP | Other | -1.454441024 | 0.829368541 | 23.40639395 | 22.95011373 | 22.74669309 | 22.56334805 | 22.37925355 | 22.71628193 | 21.66136059 | 22.26565053 | 20.09070796 |
| Geminin | O75496 | GMNN | Other | 1.648695402 | 3.233084238 | 21.70470004 | 22.33567389 | 22.9186241 | 21.39706715 | 21.77759281 | 22.62353497 | 24.00478898 | 23.77786129 | 23.54203241 |
| Mannose-1-phosphate guanyltransfer | Q96IJ6;Q96IJ6-2 | GMPPA | Extracellular or secreted | 0.523861934 | 0.691470918 | 25.52145188 | 24.06661086 | 25.23538584 | 25.53768503 | 25.34844208 | 25.64132789 | 25.3357905 | 25.65839979 | 26.25284731 |
| Mannose-1-phosphate guanyltransfer | Q9Y5P6;Q9Y5P6-2 | GMPPB | Extracellular or secreted | -0.039523603 | 0.121990478 | 27.8008848 | 27.86632094 | 27.82941403 | 27.70252972 | 27.76471167 | 27.09707859 | 27.59576694 | 27.6414795 | 27.67465262 |
| GMP reductase 1 | P36959 | GMPR | Other | 1.860514913 | 1.087304954 | 24.30632986 | 24.59331817 | 24.35056803 | 23.10765194 | 23.33660789 | 25.63392136 | 26.22064682 | 27.14520747 | 24.87988907 |
| GMP reductase 2 | Q9P2T1;Q9P2T1-2;Q9P2T1-3 | GMPR2 | Other | -0.402860137 | 0.611453656 | 27.13428013 | 27.07561494 | 26.86950141 | 26.87900096 | 26.88296986 | 26.16892196 | 26.61231487 | 26.74393844 | 25.94031091 |
| GMP synthase [glutamine-hydrolyzing | P49915;P49915-2 | GMPS | Other | 0.059834851 | 0.058679083 | 30.83921037 | 30.96767348 | 30.90210591 | 30.85912433 | 30.81711243 | 30.47014553 | 31.20833274 | 31.18547513 | 30.21338271 |
| Guanine nucleotide-binding protein su | P29992 | GNA11 | Extracellular or secreted | 0.594779408 | 3.279395847 | 26.510628 | 26.5489993 | 26.61456315 | 26.6707416 | 26.79686881 | 26.59605141 | 27.1231707 | 27.22287743 | 27.30721624 |
| Guanine nucleotide-binding protein su | Q14344;Q14344-2 | GNA13 | Extracellular or secreted | 0.794702808 | 2.543466831 | 25.03355474 | 24.63073149 | 24.97478182 | 24.58250541 | 24.98689439 | 25.27792273 | 25.63718713 | 25.56048794 | 25.92962864 |
| Guanine nucleotide-binding protein G | P63096;P63096-2 | GNAI1 | Extracellular or secreted | 0.627556233 | 0.955915301 | 23.64932723 | 23.76425609 | 23.85176658 | 22.73849062 | 23.23888051 | 25.07724602 | 24.20895827 | 24.53783311 | 24.29586084 |
| Guanine nucleotide-binding protein G | P04899;P04899-6;P04899-3;F | GNAI2 | Extracellular or secreted | 0.407965065 | 1.727355052 | 29.24215863 | 29.23098267 | 29.46835973 | 29.10694346 | 29.15590796 | 29.58572123 | 29.53411898 | 29.83805335 | 29.74675971 |
| Guanine nucleotide-binding protein G | P08754 | GNAI3 | Extracellular or secreted | 0.774533826 | 2.728536033 | 26.58132772 | 26.54377361 | 26.88739281 | 26.44720534 | 26.73104448 | 26.93483348 | 27.27799365 | 27.50468554 | 27.603711 |
| Guanine nucleotide-binding protein G | P50148 | GNAQ | Extracellular or secreted | 1.16739507 | 2.344080263 | 24.80661137 | 24.16744877 | 24.56836717 | 24.50227436 | 23.90389361 | 23.90692362 | 25.25670309 | 25.83306302 | 25.34017854 |
| Guanine nucleotide-binding protein G | P63092;P63092-4;Q5JWF2;P6 | GNAS | Extracellular or secreted | 0.925545556 | 4.286110137 | 27.69928957 | 27.78668621 | 27.9280703 | 28.00683377 | 28.08648792 | 27.48975966 | 28.71610527 | 28.72656919 | 28.83252592 |
| Guanine nucleotide-binding protein G | P62873;P62873-2 | GNB1 | Extracellular or secreted | 0.51000959 | 3.213192996 | 30.25653664 | 30.30083874 | 30.38071721 | 30.16554768 | 30.13148625 | 29.97678217 | 30.69062451 | 30.81352373 | 30.63183488 |
| Guanine nucleotide-binding protein G | P62879 | GNB2 | Extracellular or secreted | 0.362841919 | 2.531806682 | 28.16920876 | 28.04557113 | 28.12514321 | 27.78344251 | 27.74746416 | 27.93590141 | 28.28011969 | 28.39386633 | 28.31790532 |
| Guanine nucleotide-binding protein su | P63244 | GNB2L1 | Extracellular or secreted | 0.364977614 | 1.533958784 | 30.96051061 | 30.98393284 | 30.96856634 | 31.466066 | 31.45938764 | 31.38267663 | 31.69029148 | 31.59987742 | 31.41533397 |
| Guanine nucleotide-binding protein su | Q9HAV0 | GNB4 | Extracellular or secreted | 0.891443481 | 1.96082665 | 25.31028826 | 25.36205457 | 25.30994147 | 24.90944377 | 25.03531797 | 24.01945567 | 25.61624707 | 25.88560199 | 26.14573224 |
| Bifunctional UDP-N-acetylglucosamin | Q9Y223-5;Q9Y223-3;Q9Y223 | GNE | Other | 1.449672374 | 0.981054836 | 24.58267768 | 23.82434815 | 24.04793333 | 25.19734299 | 23.32429813 | 20.40407983 | 24.39335903 | 24.99360391 | 25.65239424 |
| Guanine nucleotide-binding protein G | P50151;Q9H1X3-3 | GNG10 | Extracellular or secreted | 0.774417604 | 0.440930453 | 21.62283472 | 21.57427014 | 21.4391349 | 21.9428981 | 21.58114522 | 22.29661715 | 21.89304373 | 21.81846984 | 23.84018936 |
| Guanine nucleotide-binding protein G | Q9UBI6 | GNG12 | Extracellular or secreted | 0.79707214 | 1.489966861 | 27.39631851 | 27.33270408 | 27.51141246 | 27.52492234 | 27.88081167 | 27.36744715 | 28.15373555 | 28.07841723 | 28.66587174 |
| Guanine nucleotide-binding protein-li | P36915;P36915-2 | GNL1 | Extracellular or secreted | -0.413395945 | 0.821964543 | 26.96763226 | 26.89272883 | 27.02844182 | 26.49743988 | 26.61694812 | 27.63027324 | 26.70990858 | 26.70767256 | 26.1589631 |
| Guanine nucleotide-binding protein-li | Q9BVP2-2;Q9BVP2 | GNL3 | Extracellular or secreted | -1.071581897 | 0.921059375 | 21.06358958 | 22.82811169 | 22.6462537 | 22.75594948 | 22.96944273 | 24.85338571 | 21.31978289 | 22.46844208 | 21.55539578 |
| Glucosamine-6-phosphate isomerase | P46926;P46926-2 | GNPDA1 | Extracellular or secreted | -0.152958939 | 0.566347767 | 30.60310964 | 30.74607093 | 30.6378919 | 30.68979178 | 30.770221 | 29.98549445 | 30.37565168 | 30.49609917 | 30.38566219 |
| Glucosamine-6-phosphate isomerase | Q8TDQ7;Q8TDQ7-3;Q8TDQ7 | GNPDA2 | Other | -0.496170051 | 1.271475662 | 27.37318484 | 27.40332506 | 27.38038695 | 27.51050728 | 27.70701424 | 26.34360485 | 26.92840922 | 26.68511962 | 26.75697262 |
| Glucosamine 6-phosphate N-acetyltra | Q96EK6 | GNPNAT1 | Other | 0.037060332 | 0.096615379 | 29.38391278 | 29.43646699 | 29.39911299 | 29.12391071 | 29.07083852 | 29.39145096 | 29.45364002 | 29.45949513 | 29.10089232 |
| N-acetylglucosamine-6-sulfatase | P15586;P15586-2 | GNS | Extracellular or secreted | -0.470713469 | 1.203233419 | 29.54255647 | 29.57616395 | 29.49155741 | 29.74005016 | 29.87329113 | 28.63336018 | 28.88997628 | 28.88826416 | 29.2381088 |
| Golgin subfamily A member 1 | Q92805 | GOLGA1 | Other | 1.00620076 | 1.353662894 | 24.41989076 | 24.36105063 | 24.53552125 | 25.13080902 | 24.71214114 | 24.95920922 | 25.39237665 | 25.45826341 | 26.22727323 |
| Golgin subfamily A member 2 | Q08379;Q08379-2 | GOLGA2 | Other | 0.099650831 | 0.067600379 | 25.50448855 | 25.07834597 | 25.76040322 | 25.08025868 | 24.41338414 | 26.93685626 | 25.67651002 | 24.83994918 | 26.16936171 |
| Golgin subfamily A member 3 | Q08378;Q08378-2;Q08378-4 | GOLGA3 | Other | -0.196543692 | 0.691932182 | 26.39045908 | 26.21761801 | 26.40020076 | 26.20351708 | 26.09786223 | 26.20232175 | 26.09430228 | 25.85043181 | 26.22162429 |
| Golgin subfamily A member 4 | Q13439-3;Q13439-4;Q13439 | GOLGA4 | Extracellular or secreted | -0.641568333 | 1.250673006 | 28.02949534 | 28.11565046 | 27.99015565 | 27.82274469 | 27.83647409 | 27.96422046 | 27.30652109 | 27.026438 | 27.62170626 |
| Golgin subfamily A member 5 | Q8TBA6-2;Q8TBA6 | GOLGA5 | Other | -1.194988354 | 1.721822539 | 28.56070658 | 28.63985928 | 28.52234273 | 27.75957808 | 28.2714631 | 27.87502111 | 26.66193681 | 27.04124254 | 27.52634103 |
| Golgin subfamily A member 7 | Q7Z5G4-3;Q7Z5G4 | GOLGA7 | Extracellular or secreted | -0.59516667 | 0.307075563 | 23.29151135 | 23.4241909 | 23.19380627 | 21.26200724 | 23.60692501 | 23.7365514 | 21.93708027 | 21.69741875 | 23.83749705 |
| Golgin subfamily B member 1 | Q14789-4;Q14789-3;Q14789 | GOLGB1 | Other | -1.882353436 | 2.052009222 | 30.02722931 | 29.92011125 | 29.80052999 | 29.46446991 | 29.5480439 | 29.86350273 | 27.68177308 | 27.61175225 | 28.37135791 |
| Golgi integral membrane protein 4 | O00461 | GOLIM4 | Other | -0.726345988 | 0.289326051 | 23.80479119 | 21.20400987 | 21.47669677 | 23.75847292 | 23.4891012 | 25.7370674 | 21.04139713 | 22.85014562 | 23.66448895 |
| Golgi membrane protein 1 | Q8NBJ4-2;Q8NBJ4 | GOLM1 | Extracellular or secreted | -0.599700795 | 1.00383006 | 24.86905421 | 25.58792192 | 25.55511357 | 26.48646434 | 26.70754092 | 25.88192033 | 25.3182071 | 24.95215966 | 25.4745385 |
| Golgi phosphoprotein 3 | Q9H4A6 | GOLPH3 | Extracellular or secreted | 0.324038384 | 0.532330925 | 25.44253049 | 25.26549315 | 25.29687617 | 25.60975318 | 25.76273645 | 25.90944377 | 25.90877979 | 26.25021079 | 25.45654117 |
| Golgi-associated PDZ and coiled-coil n | Q9HD26-2;Q9HD26;Q9HD26- | GOPC | Extracellular or secreted | 0.971118237 | 0.654899926 | 24.16966754 | 23.38942548 | 23.68442418 | 24.5785952 | 24.98498406 | 23.74026252 | 25.59003714 | 25.66876858 | 23.92822847 |
| Golgi reassembly-stacking protein 1 | Q9BQQ3;Q9BQQ3-2;Q9BQQ3 | GORASP1 | Other | 0.633488303 | 2.573819868 | 23.69720939 | 23.4241909 | 23.17362224 | 22.90873915 | 23.30632986 | 22.95224855 | 23.84680262 | 23.95331478 | 23.83151756 |
| Golgi reassembly-stacking protein 2 | Q9H8Y8-2;Q9H8Y8;Q9H8Y8-3 | GORASP2 | Other | -0.305751127 | 1.115122127 | 28.08734815 | 27.89012128 | 28.08101086 | 27.37567239 | 27.75995897 | 27.66064544 | 27.67080912 | 27.49251275 | 27.3468033 |
| Golgi SNAP receptor complex membe | O95249;O95249-2 | GOSR1 | Other | -1.676490684 | 1.995488338 | 26.7006791 | 25.67745126 | 26.11328735 | 26.10795119 | 26.12758539 | 26.13173154 | 23.97342481 | 24.55438181 | 24.87206424 |
| Aspartate aminotransferase, cytoplas | P17174;P17174-2 | GOT1 | Extracellular or secreted | 0.083351497 | 0.250018877 | 31.38468484 | 31.43101587 | 31.34244387 | 31.52971414 | 31.62392253 | 30.82297257 | 31.42492369 | 31.31613649 | 31.57637122 |
| Aspartate aminotransferase, mitocho | P00505;P00505-2 | GOT2 | Extracellular or secreted | -0.574719794 | 0.933294688 | 32.32795928 | 32.52887626 | 32.44265407 | 33.08236304 | 33.11732674 | 31.90659252 | 32.26282144 | 32.1847956 | 31.53110953 |
| GPALPP motifs-containing protein 1 | Q8IXQ4;Q8IXQ4-2 | GPALPP1 | Other | 0.168964647 | 0.273632374 | 22.43398191 | 23.4354261 | 23.27255803 | 23.71397713 | 23.24347851 | 22.67551432 | 23.03024077 | 23.26613684 | 23.59798433 |
| G patch domain-containing protein 4 | Q5T3I0;Q5T3I0-3 | GPATCH4 | Other | -1.024893291 | 1.346762481 | 24.11044253 | 24.03178936 | 23.87229913 | 24.18124145 | 24.30897 | 24.24016244 | 23.13201015 | 22.65863949 | 23.50712294 |
| G patch domain-containing protein 8 | Q9UKJ3-2;Q9UKJ3 | GPATCH8 | Other | -0.865941082 | 0.792214506 | 24.02055816 | 24.17538995 | 24.08966319 | 21.3927101 | 21.77410497 | 23.76729057 | 22.14500918 | 22.22934692 | 22.63767913 |
| Glypican-1;Secreted glypican-1 | P35052 | GPC1 | Extracellular or secreted | 1.620608067 | 2.55697667 | 26.83460684 | 26.78643695 | 26.90242222 | 27.23196689 | 27.4985359 | 28.77523842 | 28.81329436 | 29.04468561 | 29.01844785 |
| Glypican-3;Secreted glypican-3 | P51654-2;P51654;P51654-3 | GPC3 | Extracellular or secreted | -0.45967844 | 0.294703648 | 21.80376694 | 21.93272207 | 22.5639325 | 22.57156472 | 21.88636045 | 25.40645883 | 21.76372946 | 22.03251259 | 22.90712539 |
| Glypican-4;Secreted glypican-4 | O75487;O75487-2 | GPC4 | Extracellular or secreted | 3.141776278 | 4.30881896 | 25.97556917 | 25.31350945 | 25.19208357 | 25.20717165 | 24.78001625 | 26.76192532 | 28.63813366 | 29.04895194 | 28.35338095 |
| Glycerol-3-phosphate dehydrogenase | Q8N335 | GPD1L | Extracellular or secreted | 0.164600081 | 0.227369688 | 26.44234065 | 26.386238 | 26.74791227 | 25.89291407 | 25.05299848 | 26.91501826 | 26.57286893 | 26.27341203 | 26.36623015 |
| Glycerol-3-phosphate dehydrogenase | P43304;P43304-2 | GPD2 | Extracellular or secreted | -1.52207956 | 0.456654039 | 21.91454947 | 19.90100828 | 20.95766038 | 19.70514299 | 19.90744817 | 27.62505553 | 19.71112247 | 18.89829434 | 21.82977692 |
| Gephyrin;Molybdopterin adenylyltran | Q9NQX3;Q9NQX3-2 | GPHN | Extracellular or secreted | -0.257943021 | 0.918131402 | 26.77918973 | 26.85685596 | 26.92546932 | 26.8754899 | 27.0342267 | 26.66193681 | 26.70596032 | 26.70464182 | 26.382153 |
| Glucose-6-phosphate isomerase | P06744 | GPI | Extracellular or secreted | -0.540686215 | 0.949059204 | 34.57339454 | 34.64409557 | 34.6463507 | 34.67624476 | 34.80024646 | 33.54869162 | 34.18785098 | 34.11303145 | 33.52157075 |
| G patch domain and KOW motifs-cont | Q92917 | GPKOW | Other | 0.13044502 | 0.852970231 | 26.23066666 | 26.23633513 | 26.13475014 | 26.22237999 | 26.49926611 | 26.5785952 | 26.4171081 | 26.42636816 | 26.49885541 |
| GPN-loop GTPase 1 | Q9HCN4;Q9HCN4-5;Q9HCN4 | GPN1 | Other | 0.86363956 | 0.807712948 | 24.79986024 | 24.22729159 | 24.34996094 | 21.90686853 | 22.02619815 | 23.06115437 | 24.36766376 | 24.42911707 | 23.97980475 |
| GPN-loop GTPase 3 | Q9UHW5;Q9UHW5-2;Q9UHV | GPN3 | Other | 0.691746936 | 0.431211223 | 23.32017143 | 22.5239658 | 22.76087017 | 23.26255713 | 23.73995362 | 21.8716743 | 23.80055159 | 24.52205083 | 22.49223462 |
| Protein GPR107 | Q5VW38-3;Q5VW38-2;Q5VW | GPR107 | Other | -0.73713348 | 0.220336123 | 23.19711048 | 23.25537095 | 23.45035547 | 22.54210328 | 22.58879697 | 22.72719008 | 20.06395754 | 22.40273001 | 24.20237562 |
| G-protein coupled receptor 126 | Q86SQ4-2;Q86SQ4;Q86SQ4-4 | GPR126 | Extracellular or secreted | 1.866732075 | 1.40281201 | 23.69636056 | 24.07234723 | 23.49130509 | 25.92483533 | 25.57934356 | 25.87309748 | 26.1447603 | 26.12230194 | 27.65177862 |
| G-protein coupled receptor 56;GPR56 | Q9Y653-2;Q9Y653-3;Q9Y653 | GPR56 | Extracellular or secreted | 1.691652999 | 4.351489193 | 25.87298011 | 25.93319069 | 26.10701333 | 26.21410104 | 26.07236767 | 26.92852217 | 27.78275541 | 27.87373115 | 27.98255993 |
| Retinoic acid-induced protein 3 | Q8NFJ5 | GPRC5A | Extracellular or secreted | 1.042759725 | 2.199313112 | 25.90462874 | 25.93997478 | 25.95619853 | 26.47769133 | 26.46699948 | 27.42093488 | 27.30199444 | 27.40632907 | 27.50316954 |
| G protein-regulated inducer of neurite | Q7Z2K8-2;Q7Z2K8 | GPRIN1 | Extracellular or secreted | 0.993971865 | 0.576410356 | 24.52456371 | 21.01078775 | 24.25255861 | 23.99308558 | 24.11862366 | 20.06616018 | 24.00496039 | 23.81166632 | 24.14817864 |
| COP9 signalosome complex subunit 1 | Q13098-5;Q13098;Q13098-7 | GPS1 | Other | 0.045011263 | 0.068359056 | 29.29848086 | 29.4031625 | 29.27115138 | 29.00284321 | 28.91122241 | 28.54984394 | 29.28142477 | 29.33255471 | 28.73940646 |
| Glutathione peroxidase 1 | P07203 | GPX1 | Extracellular or secreted | 1.925539421 | 3.735510825 | 23.72837395 | 22.89727899 | 23.68570781 | 23.93075683 | 23.14198861 | 24.49374993 | 25.67946614 | 25.66985002 | 25.36623015 |
| Phospholipid hydroperoxide glutathio | P36969-2;P36969 | GPX4 | Extracellular or secreted | 0.492350195 | 1.317709197 | 27.78269293 | 27.53430454 | 27.54045172 | 27.9264311 | 27.83701575 | 28.30691215 | 28.30534726 | 28.56699649 | 28.06861093 |
| Growth factor receptor-bound protein | P62993;P62993-2 | GRB2 | Extracellular or secreted | -0.234312495 | 0.536959843 | 29.14065634 | 29.21680894 | 29.11433534 | 29.44766253 | 29.51715084 | 29.13814342 | 28.8630448 | 28.8628084 | 29.35858803 |
| Grainyhead-like protein 2 homolog | Q6ISB3-2;Q6ISB3 | GRHL2 | Extracellular or secreted | 0.115996551 | 0.050276117 | 24.57530934 | 25.37032716 | 25.75536887 | 21.68656869 | 24.47525004 | 22.78926912 | 24.34665115 | 25.02593133 | 23.30145377 |
| Glyoxylate reductase/hydroxypyruvat | Q9UBQ7 | GRHPR | Extracellular or secreted | -0.427017451 | 0.375595603 | 29.5984849 | 29.72601693 | 29.56951566 | 29.49844079 | 29.49440245 | 28.2178953 | 29.19952565 | 29.44774134 | 28.12405866 |
| GRIP1-associated protein 1 | Q4V328;Q4V328-3;Q4V328-2 | GRIPAP1 | Extracellular or secreted | 0.322940794 | 0.580168134 | 25.58740694 | 25.53023153 | 25.30834514 | 25.43182887 | 25.45979605 | 25.09611352 | 25.42978752 | 25.60944322 | 26.13645267 |
| Granulins;Acrogranin;Paragranulin;Gr | P28799;P28799-3;P28799-2 | GRN | Extracellular or secreted | -0.25824632 | 1.14805212 | 27.89700657 | 27.8251128 | 27.84200144 | 28.26887826 | 28.06095463 | 28.34798611 | 27.67942587 | 27.73427879 | 27.93252629 |
| GrpE protein homolog 1, mitochondri | Q9HAV7 | GRPEL1 | Other | 0.589241154 | 1.101927674 | 27.27825958 | 26.67976142 | 26.55584496 | 27.27115138 | 25.92712092 | 27.69491374 | 27.48131474 | 27.68872519 | 27.30120953 |
| G-rich sequence factor 1 | Q12849-5;Q12849 | GRSF1 | Other | 0.504391392 | 0.753480608 | 25.64806568 | 25.49664779 | 25.95002471 | 26.46123337 | 26.58017781 | 26.70569671 | 26.75085355 | 27.00185055 | 26.18139312 |
| Glutamate-rich WD repeat-containing | Q9BQ67 | GRWD1 | Other | 0.901835039 | 1.574069558 | 27.53608473 | 27.70490562 | 27.73776372 | 28.23781258 | 28.17849914 | 26.86561322 | 28.90118533 | 28.74947957 | 28.18517972 |
| Glycogen synthase kinase-3 alpha | P49840 | GSK3A | Other | 0.027403265 | 0.079078876 | 25.5919496 | 25.82211254 | 25.7762053 | 25.43631607 | 25.5180331 | 25.3782549 | 25.75513962 | 25.66332204 | 25.42518388 |
| Glycogen synthase kinase-3 beta | P49841;P49841-2 | GSK3B | Extracellular or secreted | 0.313186211 | 0.86162827 | 23.26083571 | 23.04244959 | 23.09294034 | 23.49313911 | 23.24142868 | 23.73871738 | 23.48259233 | 23.49326129 | 23.89846042 |
| Gelsolin | P06396;P06396-4;P06396-3 | GSN | Extracellular or secreted | 0.661625428 | 1.273503012 | 32.77033917 | 32.7018195 | 32.66756876 | 32.2067328 | 32.39504185 | 33.54489762 | 33.12827826 | 33.24019201 | 33.75960585 |
| Gelsolin | P06396-2 | GSN | Extracellular or secreted | -0.532386175 | 1.054928955 | 26.3413668 | 26.11511463 | 26.02903189 | 26.17096661 | 26.93337081 | 27.23388762 | 26.04857784 | 25.60161575 | 26.16451707 |
| Eukaryotic peptide chain release facto | P15170-2;P15170-3;P15170 | GSPT1 | Other | 0.220266957 | 0.584908175 | 30.43658612 | 30.43946229 | 30.38977009 | 31.04316072 | 31.19219413 | 30.51163866 | 31.05329633 | 30.92528538 | 30.68862516 |
| Glutathione reductase, mitochondrial | P00390-2;P00390;P00390-4;F | GSR | Extracellular or secreted | -0.316394698 | 0.447860867 | 31.64894511 | 31.68018075 | 31.62618763 | 31.25400377 | 31.37647992 | 30.71898144 | 31.32972438 | 31.31958585 | 30.55389499 |
| Glutathione synthetase | P48637;P48637-2 | GSS | Extracellular or secreted | -0.137899957 | 0.204185973 | 31.40420864 | 31.53908383 | 31.4576276 | 31.54310062 | 31.70779596 | 30.28173428 | 31.39468419 | 31.30067512 | 30.85771628 |
| Glutathione S-transferase kappa 1 | Q9Y2Q3;Q9Y2Q3-3;Q9Y2Q3- | GSTK1 | Extracellular or secreted | -0.345264878 | 0.321568162 | 26.51323616 | 26.84900032 | 26.99312878 | 26.17163479 | 26.38368623 | 27.25601917 | 26.71174741 | 26.7598955 | 25.57591519 |
| Glutathione S-transferase Mu 3 | P21266 | GSTM3 | Extracellular or secreted | -0.63572248 | 0.708173404 | 27.12878526 | 27.38277968 | 27.56797568 | 27.99129157 | 28.13780152 | 27.8303211 | 27.23407041 | 27.51080907 | 26.36743048 |
| Glutathione S-transferase P | P09211 | GSTP1 | Extracellular or secreted | -0.32872695 | 0.975109527 | 34.33863713 | 34.40597166 | 34.29522425 | 34.13208408 | 34.31749836 | 33.57688925 | 33.96344204 | 33.97160138 | 33.61192809 |
| Glutathione S-transferase theta-1 | P30711 | GSTT1 | Extracellular or secreted | -0.568445124 | 0.679489499 | 25.92723398 | 25.81709413 | 25.9099015 | 24.99360391 | 25.20534552 | 24.51279929 | 25.32302699 | 24.88734632 | 24.26728048 |
| Maleylacetoacetate isomerase | O43708-2;O43708 | GSTZ1 | Other | 1.547438096 | 1.038558033 | 21.91125451 | 20.81093439 | 21.14090825 | 23.23306842 | 23.20157416 | 23.97867066 | 24.50893697 | 24.44056759 | 22.83101492 |
| Transcription initiation factor IIA subu | P52655;P52655-2 | GTF2A1 | Other | -0.325549771 | 0.443717536 | 24.43891924 | 24.53296796 | 24.17592289 | 25.39018024 | 25.46176419 | 24.27980099 | 24.24212682 | 24.07985193 | 24.84114969 |
| Transcription initiation factor IIA subu | P52657 | GTF2A2 | Other | -2.408623937 | 1.203488398 | 24.60286194 | 24.52725124 | 23.97289918 | 25.77248493 | 26.11432045 | 25.03061124 | 21.6024138 | 22.14820612 | 24.03372276 |
| General transcription factor IIE subun | P29083 | GTF2E1 | Other | 1.251226268 | 2.131979365 | 23.63588725 | 24.06035686 | 23.24243215 | 23.98172196 | 23.56523227 | 25.11905921 | 25.55713131 | 25.12293382 | 24.87595853 |
| General transcription factor IIF subun | P35269 | GTF2F1 | Other | -0.538265669 | 1.01214813 | 27.17359937 | 27.55457211 | 27.26552892 | 27.3655295 | 27.40624796 | 27.58541681 | 26.81347786 | 26.53743325 | 27.20973922 |

| | | | | | | | | | | | | | | |
|---|---|---|---|---|---|---|---|---|---|---|---|---|---|---|
| General transcription factor IIF subun | P13984 | GTF2F2 | Other | -0.597070708 | 2.485670794 | 27.43674496 | 26.88203698 | 26.91216517 | 27.24399778 | 27.38401575 | 26.79897088 | 26.40228438 | 26.47843217 | 26.65703708 |
| General transcription factor II-I | P78347-2;P78347-4;P78347-3 | GTF2I | Other | 0.212228987 | 0.76095559 | 28.1325162 | 28.03280892 | 28.12824446 | 28.22209434 | 28.19305237 | 28.91760968 | 28.43555327 | 28.46816549 | 28.54613119 |
| General transcription factor 3C polype | Q12789-3;Q12789 | GTF3C1 | Other | -0.014092406 | 0.005845585 | 25.15634205 | 25.19516642 | 24.92279561 | 24.57991896 | 24.81885012 | 26.44688995 | 26.21695229 | 25.49950944 | 23.8012426 |
| General transcription factor 3C polype | Q8WUA4-2;Q8WUA4 | GTF3C2 | Other | -0.442471411 | 0.255873934 | 22.46368601 | 22.38313578 | 22.6922792 | 22.67333755 | 22.77277688 | 23.26899423 | 23.01021256 | 22.7651671 | 21.02431093 |
| General transcription factor 3C polype | Q9Y5Q9 | GTF3C3 | Other | 0.287338935 | 0.455728165 | 24.06693926 | 24.52366676 | 24.57085862 | 23.96031451 | 23.54333113 | 25.3833072 | 24.80720121 | 24.67438324 | 24.4046411 |
| General transcription factor 3C polype | Q9UKN8 | GTF3C4 | Other | -0.382449453 | 0.227779006 | 24.88553217 | 25.20198538 | 25.0248752 | 28.40993373 | 28.30024963 | 26.37257058 | 26.13359441 | 25.93562045 | 25.88101012 |
| General transcription factor 3C polype | Q9Y5Q8-2;Q9Y5Q8;Q9Y5Q8- | GTF3C5 | Other | 0.619296689 | 0.825794083 | 24.91406025 | 24.15421859 | 24.36111759 | 24.1375084 | 24.51731232 | 25.02462162 | 24.98784861 | 25.69572361 | 24.72873723 |
| GTP-binding protein 1 | O00178 | GTPBP1 | Other | -0.55704885 | 1.238105261 | 24.64311854 | 24.33333554 | 24.27298509 | 24.28383274 | 24.15930965 | 24.30910882 | 23.93804613 | 23.45261931 | 23.93903321 |
| Nucleolar GTP-binding protein 1 | Q9BZE4;Q9BZE4-2;Q9BZE4-3 | GTPBP4 | Other | -0.128444949 | 0.094446946 | 25.05775678 | 24.80607048 | 25.61178039 | 26.10675381 | 26.29438909 | 27.73699002 | 25.53356215 | 25.57328799 | 26.3146853 |
| Guanylate kinase | Q16774;Q16774-2;Q16774-3 | GUK1 | Other | -0.024452792 | 0.033353849 | 25.01877679 | 25.04264163 | 25.3458056 | 25.07361434 | 24.82861533 | 24.68142459 | 25.21356349 | 25.17816667 | 24.53035061 |
| Beta-glucuronidase | P08236;P08236-3;P08236-2 | GUSB | Extracellular or secreted | -1.218903579 | 3.228275994 | 27.9678521 | 28.39341632 | 28.21168975 | 27.96620249 | 28.09083445 | 27.12622694 | 26.87759756 | 26.71986276 | 26.62393997 |
| Glycogenin-1 | P46976-2;P46976 | GYG1 | Extracellular or secreted | 0.037646657 | 0.051370104 | 28.70731052 | 28.70120809 | 28.59291914 | 28.39009822 | 28.57141572 | 27.53304225 | 28.58064507 | 28.68271089 | 28.09758097 |
| Glycogen [starch] synthase, muscle | P13807;P13807-2 | GYS1 | Other | -0.001561494 | 0.001331458 | 27.09174254 | 27.11048235 | 26.95726184 | 26.71711912 | 26.64030761 | 26.80476657 | 27.16786984 | 27.30947316 | 26.17881254 |
| Histone H1.0;Histone H1.0, N-termina | P07305;P07305-2 | H1F0 | Other | -0.562710944 | 0.22694122 | 29.72710501 | 29.35123386 | 29.38059337 | 29.75816472 | 29.18368993 | 30.38740948 | 29.89403083 | 30.01518086 | 27.29675367 |
| Histone H1x | Q92522 | H1FX | Other | 0.540203333 | 0.175742944 | 26.10755217 | 24.4759303 | 24.41628676 | 25.18978596 | 25.42828656 | 26.2367548 | 26.58118403 | 27.20825133 | 23.75847292 |
| Histone H2A.V;Histone H2A.Z | Q71UI9;P0C0S5;Q71UI9-4;Q7 | H2AFV;H2AFZ | Extracellular or secreted | 0.606873276 | 0.34404961 | 29.20165361 | 28.78874093 | 28.90219218 | 28.37239622 | 28.236399 | 29.69702044 | 29.93719312 | 30.28934048 | 28.19328742 |
| Histone H2AX | P16104 | H2AFX | Extracellular or secreted | 0.579412816 | 0.198902943 | 27.08121409 | 26.60201238 | 27.41796116 | 27.29210843 | 27.25601917 | 27.69119049 | 28.91153673 | 28.75779925 | 25.73915533 |
| Core histone macro-H2A.1 | O75367-3;O75367;O75367-2 | H2AFY | Extracellular or secreted | -0.477484769 | 0.222537845 | 30.54912786 | 30.51568549 | 30.456502 | 29.15518419 | 29.27399887 | 30.04126866 | 30.1265716 | 30.41316621 | 28.02369142 |
| Core histone macro-H2A.2 | Q9P0M6 | H2AFY2 | Extracellular or secreted | -0.599345211 | 0.272174281 | 27.95620961 | 28.03957022 | 28.14194769 | 27.39623684 | 27.62791021 | 28.92493158 | 28.20634272 | 28.21669332 | 25.82233139 |
| Histone H3.3 | P84243 | H3F3A | Extracellular or secreted | 0.584393336 | 0.524364055 | 28.04489401 | 28.85064045 | 29.08195054 | 30.05982072 | 30.11842564 | 30.64443944 | 30.63079397 | 30.0450503 | 29.47742114 |
| GDH/6PGL endoplasmic bifunctional p | O95479 | H6PD | Extracellular or secreted | -3.012776406 | 3.995302316 | 24.21170831 | 24.65965234 | 24.58674864 | 22.54694556 | 24.5473968 | 24.47302146 | 21.4311686 | 20.90017762 | 21.14306112 |
| Very-long-chain (3R)-3-hydroxyacyl-Co | Q9P035 | HACD3 | Other | -3.670545322 | 2.308859094 | 29.61988883 | 29.59272315 | 29.47094067 | 27.6807676 | 27.72351284 | 29.07014757 | 24.9191623 | 24.42091882 | 26.22727323 |
| Hydroxyacyl-coenzyme A dehydrogen | Q16836;Q16836-3;Q16836-2 | HADH | Other | -0.48339183 | 0.478193477 | 30.24070382 | 30.47913793 | 30.42171748 | 29.8935825 | 29.9960419 | 29.90210591 | 30.12632542 | 29.97746471 | 28.91267914 |
| Trifunctional enzyme subunit alpha, m | P40939 | HADHA | Other | -0.981260633 | 0.411971765 | 26.45331033 | 26.62282355 | 27.15325235 | 27.26060244 | 27.40494963 | 29.85583179 | 25.87342609 | 25.39073786 | 28.1674392 |
| Trifunctional enzyme subunit beta, mi | P55084-2;P55084 | HADHB | Extracellular or secreted | -0.751222895 | 0.317704708 | 25.1141417 | 25.1467812 | 25.90593727 | 25.55765721 | 25.82672575 | 28.43328635 | 23.64735214 | 25.77851313 | 26.31273079 |
| Hydroxyacylglutathione hydrolase, mi | Q16775-2;Q16775 | HAGH | Extracellular or secreted | 0.093514971 | 0.160921953 | 27.1175342 | 27.48830452 | 27.46302801 | 27.70714593 | 27.70338814 | 26.55920459 | 27.65587777 | 27.45708938 | 27.18688046 |
| Histidine ammonia-lyase | P42357-3;P42357-2;P42357 | HAL | Other | -2.506664219 | 1.211188469 | 24.30890058 | 24.30521678 | 24.42937252 | 25.74773305 | 25.79842711 | 22.61530027 | 23.53243554 | 21.71052016 | 20.83952679 |
| Hyaluronan and proteoglycan link pro | Q96S86 | HAPLN3 | Extracellular or secreted | 0.795028171 | 0.844378436 | 22.57103061 | 20.07343185 | 20.69882775 | 22.68822896 | 22.43308415 | 22.21765128 | 22.80130182 | 22.49514163 | 22.42976837 |
| Histidine--tRNA ligase, cytoplasmic | P12081;P12081-4;P12081-2;P | HARS | Other | 0.095901987 | 0.236211279 | 29.66453303 | 29.54502655 | 29.62427124 | 29.93213201 | 29.95656967 | 29.40831477 | 29.91755278 | 29.91598703 | 29.5195898 |
| Histone acetyltransferase type B catal | O14929;O14929-2 | HAT1 | Other | -0.042390539 | 0.0424201 | 28.63266706 | 28.63100221 | 28.49193962 | 28.14904165 | 28.26709291 | 28.02289816 | 28.73463412 | 28.52114482 | 27.71437026 |
| HAUS augmin-like complex subunit 1 | Q96CS2;Q96CS2-2 | HAUS1 | Other | 0.139101901 | 0.090737003 | 22.92030778 | 22.30005757 | 22.03358835 | 21.04037197 | 22.06228925 | 21.98827104 | 22.32248384 | 22.95027395 | 21.31699089 |
| HAUS augmin-like complex subunit 4 | Q9H6D7;Q9H6D7-2;Q9H6D7- | HAUS4 | Other | 0.32298714 | 0.403009643 | 21.4601216 | 22.70750406 | 22.76694698 | 22.37074828 | 22.01273774 | 22.53817661 | 23.09982549 | 22.65075201 | 22.14650155 |
| HAUS augmin-like complex subunit 5 | O94927;O94927-2 | HAUS5 | Other | -1.185116927 | 0.902299663 | 22.95904114 | 22.60580322 | 22.59757522 | 22.49638987 | 22.11610278 | 23.41699519 | 22.44422529 | 21.33718028 | 20.75919736 |
| HAUS augmin-like complex subunit 6 | Q7Z4H7-3;Q7Z4H7;Q7Z4H7-2 | HAUS6 | Other | 0.405387488 | 0.44774737 | 22.00237004 | 20.40924388 | 22.1229412 | 22.19213626 | 22.55321115 | 21.72563087 | 22.13238676 | 22.75131849 | 21.8352239 |
| Hemoglobin subunit beta;LVV-hemor | P68871;P02042 | HBB;HBD | Extracellular or secreted | -2.05106618 | 1.445171522 | 24.15050469 | 22.4571113 | 25.46970317 | 22.25635038 | 23.00207812 | 21.67042286 | 20.52382917 | 20.68850066 | 22.13755689 |
| HBS1-like protein | Q9Y450;Q9Y450-4 | HBS1L | Extracellular or secreted | 1.405913893 | 3.950784152 | 23.7234868 | 23.66600724 | 23.54698487 | 24.12076058 | 24.27575794 | 24.04184828 | 25.10713309 | 25.48348443 | 25.31454701 |
| Host cell factor 1;HCF N-terminal chai | P51610;P51610-2 | HCFC1 | Other | 0.315909256 | 0.252040297 | 25.14094395 | 25.00774283 | 25.10089232 | 24.50682038 | 24.72754326 | 22.04071986 | 24.98620002 | 24.78246798 | 24.44139107 |
| Host cell factor 1;HCF N-terminal chai | P51610-4 | HCFC1 | Other | -0.373615888 | 1.883846315 | 30.03708428 | 30.09320436 | 30.06650821 | 29.78956511 | 29.78398884 | 29.98549445 | 29.68885856 | 29.62580456 | 29.44241184 |
| Histone deacetylase 1 | Q13547 | HDAC1 | Other | -1.288860401 | 1.892732167 | 26.91854828 | 27.04614382 | 26.86407862 | 26.43190855 | 26.63131452 | 27.52155671 | 25.98189612 | 25.16042573 | 25.69787219 |
| Histone deacetylase 2 | Q92769;Q92769-3 | HDAC2 | Other | -0.294886748 | 1.234668397 | 30.49304746 | 30.41457815 | 30.466066 | 30.02630607 | 29.94293561 | 30.66771678 | 29.94321515 | 30.13552827 | 30.04192137 |
| Histone deacetylase 6 | Q9UBN7;Q9UBN7-2 | HDAC6 | Extracellular or secreted | 0.168527832 | 0.072984263 | 25.17153936 | 25.82220981 | 25.55294649 | 25.37851951 | 25.82014136 | 24.92791212 | 25.71088083 | 26.86820651 | 24.26313048 |
| HD domain-containing protein 2 | Q7Z4H3 | HDDC2 | Extracellular or secreted | 0.694555952 | 0.513000297 | 26.86396051 | 26.94871103 | 26.94692786 | 27.17883153 | 27.16901756 | 25.87560707 | 28.08258511 | 27.99091303 | 26.5016975 |
| Guanosine-3,5-bis(diphosphate) 3-pyr | Q8N4P3;Q8N4P3-2 | HDDC3 | Other | 1.125001471 | 0.856552158 | 24.18993673 | 24.45293345 | 23.89559747 | 24.1155314 | 24.26155322 | 21.37573787 | 25.65706435 | 24.78601312 | 24.07757202 |
| Hepatoma-derived growth factor | P51858;P51858-2;P51858-3 | HDGF | Extracellular or secreted | -0.400478563 | 0.912910922 | 32.76641486 | 32.83982983 | 32.76538765 | 32.46657658 | 32.56317764 | 31.6454276 | 32.32605806 | 32.13778931 | 31.85812402 |
| Hepatoma-derived growth factor-rela | Q7Z4V5 | HDGFRP2 | Other | -0.338904045 | 1.42284015 | 29.39719619 | 29.32390746 | 29.34690895 | 29.21664707 | 29.3567425 | 29.22736046 | 29.10749231 | 28.83023041 | 28.97994645 |
| Hepatoma-derived growth factor-rela | Q9Y3E1 | HDGFRP3 | Other | -0.291467872 | 0.488651658 | 28.32048995 | 28.04218237 | 27.99781925 | 27.6114005 | 27.76856568 | 27.10589508 | 27.67215889 | 27.76344581 | 27.1131681 |
| Pseudouridine-5-phosphatase | Q08623;Q08623-3;Q08623-4 | HDHD1 | Extracellular or secreted | -0.180867208 | 0.237952704 | 27.6921222 | 27.33636953 | 27.66871449 | 27.48331528 | 28.39652267 | 26.68859182 | 27.54671999 | 27.59612252 | 26.94737386 |
| Haloacid dehalogenase-like hydrolase | Q9H0R4;Q9H0R4-2 | HDHD2 | Extracellular or secreted | 0.227136755 | 0.552153658 | 26.10848968 | 25.98226615 | 26.10027077 | 26.04259988 | 26.0523974 | 25.17611318 | 26.26246753 | 25.88994728 | 26.26006399 |
| Haloacid dehalogenase-like hydrolase | Q9BSH5 | HDHD3 | Other | -0.75862147 | 2.011331702 | 27.80888338 | 27.76667155 | 27.84355968 | 27.61161156 | 27.59227761 | 27.54436336 | 26.89041124 | 27.14928398 | 26.76812394 |
| Vigilin | Q00341;Q00341-2 | HDLBP | Extracellular or secreted | -0.002187858 | 0.006404895 | 27.64313231 | 27.62107741 | 27.59583806 | 27.74258997 | 27.82389887 | 28.01307864 | 27.84583409 | 27.81188676 | 27.5555232 |
| HEAT repeat-containing protein 1;HEA | Q9H583 | HEATR1 | Other | -2.189881576 | 1.768989512 | 26.50560951 | 28.15923264 | 26.38525076 | 25.7252304 | 25.79644803 | 29.61896175 | 25.39679211 | 24.50845345 | 24.62047625 |
| HEAT repeat-containing protein 3 | Q7Z4Q2;Q7Z4Q2-2 | HEATR3 | Other | 0.434635592 | 0.569913914 | 24.41034628 | 24.48867227 | 23.76425609 | 24.73371008 | 25.13076975 | 25.28637332 | 25.48815126 | 25.14883807 | 24.57398134 |
| HEAT repeat-containing protein 5A | Q86XA9-2;Q86XA9 | HEATR5A | Extracellular or secreted | -0.097843027 | 0.211056973 | 21.35051408 | 21.3552543 | 21.46642702 | 22.07187812 | 22.26324069 | 21.19808528 | 21.47679564 | 21.60132104 | 21.481054 |
| HEAT repeat-containing protein 5B | Q9P2D3-3;Q9P2D3 | HEATR5B | Extracellular or secreted | 0.130598953 | 0.294439682 | 23.79669556 | 24.10421609 | 23.9013177 | 23.18617773 | 23.25316481 | 23.69391738 | 23.99619279 | 23.75348283 | 23.60986588 |
| HEAT repeat-containing protein 6 | Q6AI08 | HEATR6 | Other | 0.102183209 | 0.168284229 | 22.14675251 | 21.66357721 | 21.03977212 | 21.09639379 | 21.76494922 | 21.45733679 | 21.32086003 | 21.8567325 | 21.71334791 |
| Heme-binding protein 1 | Q9NRV9 | HEBP1 | Extracellular or secreted | -0.311906615 | 1.365856968 | 28.18172484 | 27.97475994 | 28.06301401 | 28.56161723 | 28.5673955 | 28.20899547 | 27.8050742 | 28.05045775 | 27.9875017 |
| Heme-binding protein 2 | Q9Y5Z4;Q9Y5Z4-2 | HEBP2 | Extracellular or secreted | -0.150774383 | 0.18896806 | 29.12361475 | 29.10206953 | 29.04160811 | 28.73628042 | 29.03354424 | 27.91279333 | 28.90239347 | 28.92911503 | 28.19112354 |
| E3 ubiquitin-protein ligase HECTD1 | Q9ULT8 | HECTD1 | Other | 0.718940833 | 1.033208226 | 24.46363612 | 25.02631135 | 24.59086524 | 25.142386 | 25.43157386 | 24.52438437 | 25.90423825 | 25.79377197 | 25.04839075 |
| E3 ubiquitin-protein ligase HECTD3 | Q5T447 | HECTD3 | Other | -0.156743 | 0.223446243 | 25.15151148 | 25.42062976 | 25.59875109 | 25.53978642 | 25.6056902 | 24.92827366 | 25.62123115 | 25.24190869 | 24.78895246 |
| E3 ubiquitin-protein ligase HERC2 | O95714 | HERC2 | Other | 1.053998792 | 0.196939992 | 27.16863509 | 22.47984411 | 27.31337105 | 23.05050495 | 22.87918054 | 21.81151137 | 26.86749971 | 26.99744242 | 21.64857781 |
| Probable E3 ubiquitin-protein ligase H | Q5GLZ8-6;Q5GLZ8-3;Q5GLZ8 | HERC4 | Other | -1.067693036 | 0.419605739 | 23.4761776 | 23.16160246 | 23.08147824 | 23.2992192 | 23.41738178 | 23.43135069 | 22.93392905 | 23.44392189 | 20.35267493 |
| Beta-hexosaminidase subunit alpha | P06865 | HEXA | Extracellular or secreted | -0.833845035 | 1.401381343 | 29.04780855 | 29.00696749 | 28.89510049 | 28.84738821 | 29.13075012 | 28.34201992 | 27.83370246 | 27.81885012 | 28.4809297 |
| Beta-hexosaminidase subunit beta;Be | P07686 | HEXB | Extracellular or secreted | -0.741068807 | 1.710135871 | 30.13968097 | 30.34720472 | 30.291867 | 30.31397645 | 30.45718725 | 29.5062567 | 29.38784084 | 29.17507581 | 29.74196347 |
| Protein HEXIM1 | O94992 | HEXIM1 | Other | 1.721050397 | 2.818514036 | 22.47223489 | 23.59866592 | 24.16852033 | 24.0126525 | 24.09707859 | 24.27958848 | 25.4096991 | 25.19940396 | 25.86841848 |
| Protein HGH1 homolog | Q9BTY7 | HGH1 | Other | 1.032370429 | 0.578786142 | 22.74888484 | 22.76401305 | 23.03979191 | 22.67009864 | 23.47370278 | 23.2408903 | 24.68869852 | 24.70588124 | 22.67122228 |
| Hepatocyte growth factor-regulated t | O14964;O14964-2 | HGS | Extracellular or secreted | -0.446083658 | 0.578317113 | 27.80427423 | 27.53845491 | 27.49716575 | 27.64512693 | 27.38722457 | 27.62644876 | 27.15759535 | 26.62100752 | 27.63249373 |
| 3-hydroxyisobutyrate dehydrogenase | P31937 | HIBADH | Extracellular or secreted | -0.933635165 | 1.682325764 | 28.76831327 | 28.66366814 | 28.63186956 | 28.24349849 | 28.26145458 | 27.85839825 | 27.57744312 | 27.76382568 | 27.07142685 |
| 3-hydroxyisobutyryl-CoA hydrolase, m | Q6NVY1;Q6NVY1-2 | HIBCH | Extracellular or secreted | -0.602031618 | 2.126736771 | 28.48864929 | 28.71852425 | 28.69866053 | 28.46478148 | 28.68194059 | 28.56354589 | 28.14952627 | 28.00372798 | 27.84870192 |
| Protein HID1 | Q8IV36-2;Q8IV36;Q8IV36-3 | HID1 | Extracellular or secreted | -0.118294543 | 0.231025829 | 26.70239762 | 26.54927845 | 26.64732469 | 25.84658745 | 25.90712998 | 26.34793544 | 26.34906669 | 26.35967745 | 25.93669904 |
| Hypoxia-inducible factor 1-alpha inhib | Q9NWT6 | HIF1AN | Other | -0.183853843 | 0.33201305 | 24.49978313 | 24.33408611 | 24.68586819 | 23.80183464 | 24.36151923 | 24.46413489 | 24.28079229 | 24.43764998 | 23.80360929 |
| Histidine triad nucleotide-binding pro | P49773 | HINT1 | Extracellular or secreted | 0.470421 | 0.472651937 | 30.16422933 | 29.89598164 | 29.81069229 | 29.39856268 | 29.59905272 | 28.70295899 | 30.2405901 | 30.5929904 | 29.36342132 |
| Histidine triad nucleotide-binding pro | Q9BX68 | HINT2 | Extracellular or secreted | 0.876858304 | 0.774696558 | 26.99528721 | 26.86584916 | 26.93494593 | 26.02601579 | 26.43841167 | 25.60314502 | 27.85851682 | 27.71679215 | 26.48709333 |
| Huntingtin-interacting protein 1 | O00291-3;O00291-4;O00291 | HIP1 | Other | -0.611084833 | 0.970765711 | 27.59754396 | 27.36703048 | 27.49785098 | 26.95914287 | 27.50483705 | 27.20247122 | 27.07224499 | 26.87572423 | 26.28321456 |
| Huntingtin-interacting protein 1-relat | O75146 | HIP1R | Other | -0.193137133 | 0.970541747 | 25.14821744 | 24.89587478 | 25.09297259 | 25.30016233 | 25.1049359 | 25.24318067 | 24.92827366 | 25.07410454 | 24.81088225 |
| HIRA-interacting protein 3 | Q9BW71 | HIRIP3 | Other | -0.962876507 | 0.807063445 | 27.2496142 | 26.68016398 | 26.80488963 | 26.22923689 | 26.38784905 | 26.01968896 | 24.88804347 | 25.46176419 | 26.44728417 |
| Histone H1.2 | P16403 | HIST1H1C | Other | -0.465442682 | 0.138147968 | 31.83251083 | 31.89504269 | 31.93680713 | 32.1068998 | 32.25375024 | 32.6985736 | 32.64093245 | 32.97865839 | 29.34587326 |
| Histone H1.3 | P16402 | HIST1H1D | Other | -0.37498268 | 0.202633591 | 22.43618896 | 20.66982572 | 22.51605009 | 23.46438421 | 23.5480439 | 23.43377185 | 22.74652908 | 22.97954312 | 21.18311213 |
| Histone H2A type 1-C;Histone H2A typ | Q93077;Q7L7L0;P04908 | HIST1H2AC;HIST1 | Extracellular or secreted | -0.16082497 | 0.093554186 | 33.52075154 | 33.61137852 | 33.69695828 | 33.69913234 | 33.77490081 | 33.77696061 | 34.12038919 | 34.06751828 | 32.36965865 |
| Histone H2B type 1-M;Histone H2B ty | Q99879;Q99877;Q93079;Q5 | HIST1H2BM;HIST | Extracellular or secreted | 0.044143969 | 0.021179707 | 33.23034715 | 33.38406723 | 33.3843246 | 33.40349776 | 33.51241597 | 34.01080333 | 34.19190009 | 34.1711671 | 32.23209273 |
| Histone H3.1;Histone H3.1t;Histone H | P68431;Q16695;Q6NXT2 | HIST1H3A;HIST3H | Extracellular or secreted | 0.241286557 | 0.181365813 | 31.98451015 | 31.36801986 | 31.87763412 | 32.34651801 | 32.04813996 | 34.80947574 | 32.58477193 | 32.43571719 | 32.92051948 |
| Histone H4 | P62805 | HIST1H4A | Extracellular or secreted | -0.02452005 | 0.010557513 | 34.00252805 | 34.08962594 | 34.07789858 | 34.32328206 | 34.26723806 | 34.43954898 | 34.83012272 | 34.95878345 | 32.73759451 |
| Histone H2B type 2-E;Histone H2B typ | Q16778;P33778;P23527;P06 | HIST2H2BE;HIST1 | Extracellular or secreted | -0.047553885 | 0.023947287 | 29.62119971 | 30.16326977 | 30.12669467 | 30.36442361 | 30.63270172 | 30.22481029 | 30.86407862 | 30.64727322 | 28.91253639 |
| Histone H3.2 | Q71DI3 | HIST2H3A | Extracellular or secreted | -0.547695992 | 0.282909365 | 27.15817343 | 27.96427555 | 27.70971142 | 29.09032971 | 29.31399806 | 30.92980639 | 27.06537866 | 28.36013812 | 29.01454253 |
| Transcription factor HIVEP3 | Q5T1R4-2;Q5T1R4 | HIVEP3 | Other | 1.905318144 | 0.670840522 | 26.93111766 | 21.95909467 | 21.21950502 | 25.4691442 | 24.99929332 | 27.28453891 | 27.52065788 | 27.39754303 | 24.72910041 |
| Hexokinase-1 | P19367;P19367-4;P19367-2;P | HK1 | Other | -0.238928315 | 0.492889769 | 29.76998465 | 29.97650906 | 29.97336459 | 29.32738115 | 29.43061717 | 30.30584734 | 29.50170889 | 29.31507847 | 29.85827967 |
| Hexokinase-2 | P52789 | HK2 | Other | 0.591498798 | 1.450670668 | 24.76283781 | 24.70804108 | 24.8915705 | 25.49747034 | 25.59214927 | 25.85514564 | 25.67659072 | 26.02002822 | 25.73148477 |
| HLA class I histocompatibility antigen, | P01892;P30459;P10314 | HLA-A | Extracellular or secreted | -0.616324252 | 0.223475552 | 23.72588045 | 23.43084045 | 23.77354169 | 20.82252792 | 21.94688772 | 28.26181323 | 22.53295607 | 22.77962559 | 23.81919132 |
| HLA class II histocompatibility antigen | P01903 | HLA-DRA | Extracellular or secreted | 3.434510988 | 1.37305179 | 25.33735676 | 24.5860614 | 25.27394553 | 21.5005735 | 23.99653762 | 22.98705916 | 28.19802719 | 28.46349581 | 25.48277694 |
| HLA class II histocompatibility antigen | P01911;Q5Y7A7;P01912 | HLA-DRB1 | Extracellular or secreted | 4.110000347 | 2.175956301 | 26.01763047 | 24.85333811 | 25.6960952 | 23.57056914 | 23.50663882 | 20.57518015 | 28.37720424 | 29.17176362 | 26.89075912 |
| HLA class II histocompatibility antigen | Q30134 | HLA-DRB1 | Extracellular or secreted | 0.943252971 | 0.489671468 | 22.8383021 | 20.94873039 | 22.10377732 | 21.7371627 | 21.33707003 | 20.12084856 | 22.97605012 | 23.35110746 | 21.04554688 |
| HLA class II histocompatibility antigen | Q95IE3 | HLA-DRB1 | Extracellular or secreted | 2.310964071 | 1.435352212 | 23.08513073 | 23.4537499 | 23.77172964 | 21.46643211 | 23.11938991 | 21.5895738 | 24.64784617 | 26.26205526 | 24.26599383 |
| HLA class II histocompatibility antigen | Q29974;P04229;P13760;Q30 | HLA-DRB1;HLA-D | Extracellular or secreted | 1.545212475 | 0.873456148 | 22.04112972 | 22.16938845 | 21.35799436 | 21.921968 | 22.47560208 | 22.47882229 | 24.16660628 | 24.37358313 | 22.31790046 |
| Biotin--protein ligase;Biotin--[methyl | P50747 | HLCS | Other | 1.549242344 | 2.128014086 | 25.96684056 | 26.36738049 | 25.96223571 | 25.19460298 | 24.08204701 | 26.47297499 | 27.24463298 | 27.53051433 | 26.89562058 |
| Porphobilinogen deaminase | P08397-2;P08397;P08397-4;P | HMBS | Other | 0.08190085 | 0.060028905 | 27.44284683 | 27.40446245 | 27.37658342 | 27.42710385 | 27.26642284 | 25.54813212 | 27.45646283 | 27.59135045 | 26.43066502 |
| Embryonic stem cell-specific 5-hydrox | Q96FZ2 | HMCES | Other | -0.335749339 | 0.688818668 | 22.41191941 | 21.31997856 | 21.0289771 | 21.7718199 | 21.21667482 | 21.14824071 | 21.3626566 | 21.00963183 | 21.06926881 |
| High mobility group protein 20A | Q9NP66 | HMG20A | Other | 1.181902911 | 1.181149669 | 23.60975318 | 20.99590543 | 22.89092991 | 24.07422706 | 24.15298169 | 24.14086596 | 24.47116167 | 24.68072737 | 24.3261513 |
| High mobility group protein HMG-I/HI | P17096;P17096-3 | HMGA1 | Other | 0.014721481 | 0.021729239 | 29.6881916 | 29.59553576 | 29.67133231 | 30.68369829 | 30.59938975 | 30.32334918 | 30.1319768 | 29.90612656 | 30.28680953 |
| High mobility group protein HMG-I/HI | P17096-2 | HMGA1 | Other | -1.675010362 | 0.483184935 | 26.29356496 | 24.99937935 | 25.54185523 | 26.66749685 | 23.87173532 | 26.96035871 | 23.00783691 | 22.4878263 | 26.64650092 |
| High mobility group protein B1;Putati | P09429;B2RPK0 | HMGB1;HMGB1P | Extracellular or secreted | -0.473779744 | 2.350155184 | 33.38817967 | 33.57745212 | 33.52554319 | 33.5241424 | 33.51911171 | 33.14306147 | 33.10155619 | 32.96873798 | 32.84711188 |
| High mobility group protein B2 | P26583 | HMGB2 | Extracellular or secreted | -0.672754565 | 3.065551607 | 31.03950485 | 31.3795816 | 31.31980117 | 31.29542955 | 31.22745227 | 30.68403285 | 30.59076175 | 30.44657843 | 30.41729727 |
| High mobility group protein B3 | O15347 | HMGB3 | Other | -0.466424451 | 0.765672702 | 30.31094694 | 30.32356393 | 30.08385341 | 30.40375169 | 30.37710079 | 29.3572041 | 29.93100491 | 29.88019862 | 29.21773355 |
| Hydroxymethylglutaryl-CoA lyase, mit | P35914;P35914-3;P35914-2 | HMGCL | Extracellular or secreted | -0.043599608 | 0.027453834 | 25.7696127 | 26.27460323 | 25.87963204 | 23.88688137 | 23.9125992 | 23.98207027 | 25.15907862 | 25.23735674 | 24.32546522 |
| Hydroxymethylglutaryl-CoA synthase, | Q01581 | HMGCS1 | Extracellular or secreted | 3.967638446 | 5.879763021 | 29.91484723 | 29.95961275 | 29.99185919 | 29.19598771 | 29.13452495 | 29.19232348 | 33.57531204 | 33.44923796 | 33.57294299 |
| Non-histone chromosomal protein HM | P05114 | HMGN1 | Other | 0.269064212 | 0.389728843 | 30.1753376 | 29.70701424 | 29.97158425 | 30.76477493 | 30.86274929 | 29.33138053 | 30.53556574 | 30.63521264 | 30.04283468 |
| Non-histone chromosomal protein HM | P05204 | HMGN2 | Other | -0.634859381 | 0.235769964 | 21.13075405 | 21.8120582 | 22.89626292 | 23.54462868 | 21.23899405 | 21.86332196 | 20.36292409 | 23.31824158 | 20.65726611 |
| High mobility group nucleosome-bind | Q15651 | HMGN3 | Other | 1.352364103 | 1.22539708 | 23.07589235 | 23.39610615 | 23.56127489 | 24.89291407 | 25.37653374 | 22.43389917 | 25.41734959 | 25.61436657 | 24.39368634 |
| HMG domain-containing protein 4 | Q9UGU5 | HMGXB4 | Other | 0.320510035 | 0.295992027 | 20.73274077 | 20.94610063 | 21.19291885 | 22.16043173 | 22.55041539 | 21.21531259 | 21.11254386 | 22.12347859 | 22.12446763 |
| Hyaluronan mediated motility recepto | O75330-4;O75330-2;O75330 | HMMR | Extracellular or secreted | -1.249566073 | 0.65072724 | 24.38857179 | 24.44985191 | 24.30180261 | 23.5535618 | 23.55379614 | 23.23037349 | 23.37861872 | 21.20267507 | 23.40898686 |
| Heme oxygenase 1 | P09601 | HMOX1 | Extracellular or secreted | -2.295283104 | 1.662259978 | 23.51376325 | 24.02411431 | 24.52915933 | 23.45913941 | 23.01000762 | 21.43137051 | 19.98400418 | 21.45256909 | 21.66135463 |
| Heme oxygenase 2 | P30519;P30519-2 | HMOX2 | Extracellular or secreted | -0.625863812 | 0.555917271 | 26.62212535 | 26.71934056 | 26.54526228 | 26.62421894 | 26.75849834 | 26.43042575 | 25.6546629 | 25.48259233 | 26.83508894 |
| Serpin-like protein HMSD | A8MTL9 | HMSD | Extracellular or secreted | -0.160502869 | 0.145224763 | 22.19825875 | 22.99748066 | 21.28922484 | 22.87851599 | 21.78360751 | 21.53285233 | 21.48725453 | 21.84961126 | 22.52159565 |
| Hematological and neurological expre | Q9UK76;Q9UK76-3;Q9UK76- | HN1 | Extracellular or secreted | 1.794843459 | 1.927229895 | 27.49571806 | 27.34375732 | 27.55691213 | 27.84535557 | 27.99339876 | 25.09635485 | 29.20279813 | 29.4213764 | 28.42610419 |
| Hematological and neurological expre | Q9H910;Q9H910-3;Q9H910-2 | HN1L | Extracellular or secreted | 1.234119487 | 1.535673588 | 27.55215498 | 27.05682704 | 27.27604203 | 27.35632274 | 27.43595061 | 27.08911762 | 28.76030803 | 28.79686881 | 28.02838912 |
| Heterogeneous nuclear ribonucleopro | Q13151 | HNRNPA0 | Other | 0.614355121 | 0.920151466 | 25.37669932 | 25.1965553 | 24.71429164 | 26.55131903 | 26.54939597 | 26.4172208 | 26.37323463 | 26.41393286 | 26.4586389 |
| Heterogeneous nuclear ribonucleopro | P09651-2;P09651;P09651-3; | HNRNPA1;HNRNP | Extracellular or secreted | -0.239779689 | 1.268904482 | 34.24090281 | 34.18814585 | 34.05956289 | 34.10524571 | 33.88494659 | 33.61817852 | 33.70016645 | 33.81840828 | 33.81057738 |
| Heterogeneous nuclear ribonucleopro | P22626;P22626-2 | HNRNPA2B1 | Other | 0.061539026 | 0.10780744 | 34.52839892 | 34.49857394 | 34.49589672 | 34.68682129 | 34.7675696 | 33.88703686 | 34.681438 | 34.72451134 | 34.2108164 |
| Heterogeneous nuclear ribonucleopro | P51991 | HNRNPA3 | Other | 0.349464588 | 0.866700305 | 30.42462341 | 30.21083091 | 30.25157931 | 30.61429986 | 30.47169661 | 30.8501337 | 30.60443585 | 30.73408493 | 31.12145489 |
| Heterogeneous nuclear ribonucleopro | P51991-2 | HNRNPA3 | Other | 0.731761002 | 0.979849791 | 24.1178314 | 24.21541628 | 23.75796452 | 24.08804609 | 23.94386906 | 24.46101474 | 24.6510806 | 25.37238792 | 24.46388553 |
| Heterogeneous nuclear ribonucleopro | Q99729-3;Q99729-4;Q99729 | HNRNPAB | Other | -0.147578382 | 0.329191243 | 32.17521861 | 32.39030839 | 32.23549198 | 32.16347973 | 32.35696808 | 31.33665683 | 32.12333723 | 32.00678028 | 31.75620915 |
| Heterogeneous nuclear ribonucleopro | P07910;P07910-3 | HNRNPC | Extracellular or secreted | -0.914751065 | 0.436432783 | 23.51231706 | 24.15383217 | 23.08626518 | 23.80734863 | 24.19385138 | 23.16968283 | 24.16146407 | 22.61961461 | 21.43631675 |
| Heterogeneous nuclear ribonucleopro | P07910-2;P07910-4 | HNRNPC | Extracellular or secreted | -0.820359988 | 1.275944856 | 32.19128241 | 32.28002011 | 32.23352196 | 32.08989423 | 31.96935572 | 32.16542788 | 31.40441169 | 31.65548544 | 30.94377407 |
| Heterogeneous nuclear ribonucleopro | Q14103-3;Q14103;Q14103-4 | HNRNPD | Extracellular or secreted | -0.356654471 | 1.553709583 | 32.5774071 | 32.25030907 | 32.13980293 | 32.31221902 | 32.21112112 | 32.12654083 | 31.91712592 | 32.06313619 | 31.75847451 |
| Heterogeneous nuclear ribonucleopro | O14979-3;O14979-2;O14979 | HNRNPDL | Extracellular or secreted | 0.391782558 | 1.936508148 | 29.65729269 | 29.72557821 | 29.45028073 | 29.5644184 | 29.56387314 | 29.26019862 | 29.98196143 | 30.02591022 | 29.77829692 |
| Heterogeneous nuclear ribonucleopro | P52597 | HNRNPF | Other | 0.312596025 | 2.061233996 | 31.3516867 | 31.24881158 | 31.19178246 | 31.18930997 | 31.29137303 | 31.32109236 | 31.67581377 | 31.52070471 | 31.53829763 |
| Heterogeneous nuclear ribonucleopro | P31943 | HNRNPH1 | Other | 0.395741528 | 0.574767596 | 30.96375208 | 30.9699389 | 30.88954119 | 31.35903856 | 31.3661968 | 31.01424987 | 31.77615665 | 31.72647175 | 30.96595488 |
| Heterogeneous nuclear ribonucleopro | P55795 | HNRNPH2 | Other | 0.4471482 | 1.303382663 | 27.39680844 | 27.39746143 | 27.44679532 | 26.98315819 | 26.79711627 | 27.71914469 | 27.93083576 | 27.77026827 | 27.51058274 |
| Heterogeneous nuclear ribonucleopro | P31942-2;P31942;P31942-3;P | HNRNPH3 | Other | 0.456193966 | 0.978170526 | 28.21507369 | 28.25011138 | 28.22582171 | 28.26239584 | 28.28991208 | 27.42374223 | 28.80245725 | 28.68120339 | 28.21844972 |
| Heterogeneous nuclear ribonucleopro | P61978-2;P61978 | HNRNPK | Extracellular or secreted | 0.736570184 | 1.642828495 | 32.11337306 | 31.97124162 | 32.08007693 | 32.91787283 | 32.88476468 | 32.38131048 | 33.26598154 | 33.32428847 | 32.79376034 |
| Heterogeneous nuclear ribonucleopro | P61978-3 | HNRNPK | Extracellular or secreted | 0.775706344 | 2.601118402 | 25.05163015 | 24.9100388 | 24.862164 | 25.4809297 | 25.52814597 | 25.54763214 | 26.07371648 | 26.13657001 | 25.80710292 |
| Heterogeneous nuclear ribonucleopro | P14866;P14866-2 | HNRNPL | Extracellular or secreted | 0.07601391 | 0.375090071 | 31.49624206 | 31.49995226 | 31.44519748 | 31.27269817 | 31.31148839 | 31.87063349 | 31.56896327 | 31.55750567 | 31.54967872 |
| Heterogeneous nuclear ribonucleopro | Q8WVV9-5;Q8WVV9-4;Q8W | HNRNPLL | Other | -0.968830472 | 3.000694875 | 26.05426199 | 26.13848524 | 26.24822119 | 26.47275813 | 26.2030316 | 27.09073352 | 25.33762898 | 25.29056255 | 25.56906289 |
| Heterogeneous nuclear ribonucleopro | P52272;P52272-2 | HNRNPM | Extracellular or secreted | 0.126224688 | 0.16258548 | 29.1261777 | 28.94726238 | 29.16286176 | 28.89775675 | 28.72591945 | 30.65241133 | 29.45910422 | 29.4569915 | 29.21877303 |
| Heterogeneous nuclear ribonucleopro | O43390;O43390-3 | HNRNPR | Other | -0.543995773 | 1.200363871 | 31.53862142 | 31.44544418 | 31.48678079 | 31.24564194 | 31.26801934 | 31.17194253 | 30.90748817 | 31.03865484 | 30.50009477 |
| Heterogeneous nuclear ribonucleopro | O43390-2;O43390-4 | HNRNPR | Other | -0.662723713 | 1.032273151 | 24.47648665 | 24.46070235 | 24.37855258 | 23.51821323 | 23.32868698 | 23.84488885 | 23.71156364 | 23.41956934 | 22.88446121 |
| Heterogeneous nuclear ribonucleopro | Q00839-2 | HNRNPU | Other | -0.894140957 | 2.344941017 | 31.73068826 | 31.72712125 | 31.69573522 | 31.10674383 | 31.08556385 | 32.3540276 | 30.61307054 | 30.69228852 | 30.86215808 |
| Heterogeneous nuclear ribonucleopro | Q9BUJ2-2;Q9BUJ2;Q9BUJ2-4 | HNRNPUL1 | Other | 1.539440652 | 3.01889603 | 28.76979553 | 28.78861649 | 28.84260096 | 28.64388921 | 28.73502166 | 29.59699332 | 30.33383455 | 30.21117915 | 30.76176684 |
| Heterogeneous nuclear ribonucleopro | Q1KMD3 | HNRNPUL2 | Other | -1.168026052 | 2.491278766 | 30.87752443 | 30.91598703 | 30.86274929 | 30.52561319 | 30.54489762 | 30.98983094 | 29.47707367 | 29.48298846 | 29.89416097 |
| Homer protein homolog 1 | Q86YM7-2;Q86YM7 | HOMER1 | Extracellular or secreted | -0.156495099 | 0.13689247 | 21.65070819 | 21.64882275 | 21.54403678 | 22.29759697 | 21.38671704 | 24.00601914 | 21.75046002 | 21.8045155 | 22.24248962 |
| Homer protein homolog 2 | Q9NSB8-2;Q9NSB8 | HOMER2 | Extracellular or secreted | 0.022580168 | 0.021338119 | 21.59208003 | 23.12929245 | 23.07703409 | 23.53183832 | 23.29165187 | 22.92965573 | 23.33551793 | 22.54205604 | 22.96594278 |
| Homer protein homolog 3 | Q9NSC5-2;Q9NSC5;Q9NSC5-4 | HOMER3 | Extracellular or secreted | 1.968657843 | 1.559265794 | 20.24856965 | 23.90948955 | 22.61241256 | 23.44988893 | 24.24836598 | 21.26376335 | 24.75108348 | 24.68971182 | 24.33142324 |
| Protein Hook homolog 1 | Q9UJC3-2;Q9UJC3 | HOOK1 | Other | 0.348515128 | 0.885018414 | 26.29584333 | 26.3478172 | 26.40803127 | 26.96455099 | 27.00077664 | 26.45429917 | 27.20778605 | 26.84457763 | 26.728841 |
| Protein Hook homolog 3 | Q86VS8 | HOOK3 | Other | -0.392144744 | 0.586101902 | 23.82929304 | 24.20329303 | 23.64680302 | 23.66155631 | 23.51592982 | 23.30354554 | 23.39597545 | 23.68592164 | 22.82187906 |
| Heterochromatin protein 1-binding pr | Q5SSJ5;Q5SSJ5-2;Q5SSJ5-3 | HP1BP3 | Other | -0.843562056 | 0.463747279 | 27.59569581 | 27.47359441 | 27.81720394 | 27.05082098 | 27.17073745 | 27.3691126 | 27.11852465 | 27.38492152 | 25.20445025 |

| | | | | | | | | | | | | | | |
|---|---|---|---|---|---|---|---|---|---|---|---|---|---|---|
| Hippocalcin-like protein 1 | P37235 | HPCAL1 | Extracellular or secreted | 0.927265858 | 1.239734024 | 24.7073566 | 24.37914777 | 25.03971662 | 25.17588483 | 25.24405223 | 24.95491265 | 25.5245936 | 25.61298972 | 26.3947496 |
| Hypoxanthine-guanine phosphoribosy | P00492 | HPRT1 | Extracellular or secreted | -0.324073158 | 1.329772374 | 31.6353424 | 31.56151708 | 31.54107054 | 31.46236503 | 31.53732586 | 31.0292057 | 31.16512833 | 31.27907921 | 30.96698629 |
| GTPase HRas;GTPase HRas, N-termina | P01112;P01112-2 | HRAS | Extracellular or secreted | 1.836788656 | 1.408006812 | 21.03911837 | 20.60354601 | 23.70967199 | 23.54167802 | 19.70452142 | 21.39195081 | 23.46425956 | 23.54686715 | 23.49448257 |
| Ribonuclease UK114 | P52758 | HRSP12 | Extracellular or secreted | -0.834595538 | 1.067091736 | 28.63640596 | 28.84361958 | 28.73489249 | 29.05886652 | 29.06440243 | 27.932301 | 28.08957227 | 28.25858221 | 27.28330289 |
| HCLS1-binding protein 3 | Q53T59 | HS1BP3 | Other | 0.384021131 | 0.464332233 | 25.13551359 | 25.04272511 | 24.94686095 | 25.1034559 | 25.18317406 | 24.41415874 | 25.37921388 | 25.88115019 | 24.8046435 |
| Heparan-sulfate 6-O-sulfotransferase | Q96MM7-2;Q96MM7-3;Q96 | HS6ST2 | Extracellular or secreted | -0.014842551 | 0.014917204 | 24.66492291 | 24.49210012 | 24.30931703 | 25.64174132 | 25.86587276 | 25.54105761 | 24.7293079 | 25.06931799 | 25.41435233 |
| Heat shock factor-binding protein 1 | O75506 | HSBP1 | Other | 1.609646713 | 1.479671649 | 21.39843057 | 20.38869003 | 21.53269455 | 22.55234264 | 23.77384348 | 21.88513182 | 23.94055735 | 22.782308 | 23.87164133 |
| 3-hydroxyacyl-CoA dehydrogenase typ | Q99714;Q99714-2 | HSD17B10 | Extracellular or secreted | -0.146857349 | 0.360668309 | 28.49144273 | 28.62062307 | 28.8248701 | 28.79937239 | 28.74704793 | 29.57038457 | 28.8049204 | 28.76030803 | 28.52106991 |
| Very-long-chain 3-oxoacyl-CoA reduct | Q53GQ0 | HSD17B12 | Extracellular or secreted | 0.438877655 | 0.091292604 | 21.48561708 | 24.24901736 | 24.68437067 | 25.08594115 | 24.77001616 | 28.28065072 | 23.39532179 | 24.13045556 | 28.06866218 |
| Estradiol 17-beta-dehydrogenase 8 | Q92506 | HSD17B8 | Extracellular or secreted | -0.869760687 | 1.63374126 | 24.89305299 | 24.26563622 | 24.4778457 | 24.26011784 | 23.77394406 | 24.25212545 | 23.63632989 | 23.66871449 | 23.04703469 |
| Hydroxysteroid dehydrogenase-like p | Q6YN16;Q6YN16-2 | HSDL2 | Other | -1.029426671 | 3.852659517 | 27.96675257 | 28.35565086 | 28.39998897 | 27.86366518 | 27.93224467 | 28.46291104 | 27.29333693 | 27.05837627 | 27.05061343 |
| Heat shock factor protein 1 | Q00613-2;Q00613 | HSF1 | Other | 0.018288462 | 0.010226658 | 22.55647948 | 22.06219457 | 22.99884588 | 21.67788155 | 22.06956585 | 21.54298298 | 22.75056225 | 22.64306348 | 21.11521482 |
| Heat shock protein HSP 90-alpha | P07900;P07900-2 | HSP90AA1 | Extracellular or secreted | -0.207868931 | 0.32099926 | 35.31638604 | 35.45779269 | 35.45240207 | 35.51159154 | 35.6261169 | 34.61258731 | 35.37907157 | 35.30034783 | 34.68541208 |
| Heat shock protein HSP 90-alpha A2 | Q14568 | HSP90AA2P | Other | -0.334739055 | 0.158999236 | 23.42290862 | 24.5234873 | 24.15692063 | 24.87979561 | 25.21141126 | 22.66930366 | 23.2282609 | 25.15668923 | 23.04274625 |
| Heat shock protein HSP 90-beta | P08238 | HSP90AB1 | Extracellular or secreted | 0.163005438 | 0.479784102 | 35.48102236 | 35.51382818 | 35.42664284 | 35.61659107 | 35.66044568 | 35.31847514 | 35.73932597 | 35.84120387 | 35.41698911 |
| Putative heat shock protein HSP 90-be | Q58FF8 | HSP90AB2P | Other | -0.462181222 | 1.127559226 | 29.8023648 | 29.80513572 | 29.6256304 | 29.46888405 | 29.80627334 | 28.66129127 | 29.31189975 | 28.90839042 | 28.97795595 |
| Putative heat shock protein HSP 90-be | Q58FF6 | HSP90AB4P | Other | 0.742921507 | 0.854431251 | 30.54591033 | 30.8264621 | 31.04322592 | 30.25766094 | 29.9283245 | 30.19084106 | 31.54977051 | 31.53658502 | 30.53862142 |
| Endoplasmin | P14625 | HSP90B1 | Extracellular or secreted | -0.341061823 | 2.215250745 | 34.11884331 | 34.09238542 | 34.12840119 | 33.9182106 | 33.95912905 | 34.45831231 | 33.72973657 | 33.77519524 | 33.80952366 |
| Heat shock 70 kDa protein 13 | P48723 | HSPA13 | Extracellular or secreted | 0.488101689 | 1.547351949 | 23.30452067 | 24.20642657 | 23.82026312 | 24.2137118 | 24.35991199 | 24.14980727 | 24.6159385 | 24.51002429 | 24.36566298 |
| Heat shock 70 kDa protein 14 | Q0VDF9 | HSPA14 | Other | 0.020638956 | 0.032639912 | 26.4531376 | 26.54153033 | 26.16146407 | 26.59789911 | 26.66166503 | 26.18181013 | 26.65430781 | 26.63117572 | 26.07518647 |
| Heat shock 70 kDa protein 1B;Heat sh | P0DMV9;P0DMV8;P0DMV8-2 | HSPA1B;HSPA1A | Extracellular or secreted | -0.114649271 | 0.27690768 | 34.08480391 | 34.06181735 | 34.03714975 | 34.1978427 | 34.20259382 | 33.36958066 | 33.96799978 | 34.01538028 | 33.64956622 |
| Heat shock-related 70 kDa protein 2 | P54652 | HSPA2 | Extracellular or secreted | 0.211586874 | 0.687923501 | 26.7693857 | 26.96697254 | 26.82456666 | 26.89238143 | 27.14423517 | 27.01211965 | 27.09516793 | 27.37152409 | 26.97289918 |
| Heat shock 70 kDa protein 4 | P34932 | HSPA4 | Extracellular or secreted | -0.122716343 | 0.284313452 | 33.20966951 | 33.17073446 | 33.08241063 | 33.25124069 | 33.31810472 | 32.65072206 | 33.18342078 | 33.07956841 | 32.71030281 |
| Heat shock 70 kDa protein 4L | O95757 | HSPA4L | Other | 0.016421618 | 0.023047447 | 26.04626874 | 25.81438278 | 25.77240942 | 26.68618888 | 26.56883102 | 26.10659409 | 26.23604311 | 26.46962555 | 25.84093367 |
| 78 kDa glucose-regulated protein | P11021 | HSPA5 | Extracellular or secreted | -0.499065269 | 4.414401983 | 33.7178382 | 33.71241587 | 33.70192275 | 33.48627732 | 33.52740879 | 33.69249638 | 33.11842564 | 33.16013176 | 33.14342645 |
| Heat shock 70 kDa protein 6;Putative | P17066;P48741 | HSPA6;HSPA7 | Extracellular or secreted | 0.737152204 | 3.074681017 | 31.69253795 | 31.85801283 | 31.98461201 | 31.35400132 | 31.56674252 | 31.25231272 | 32.43522042 | 32.25453886 | 32.37580701 |
| Heat shock cognate 71 kDa protein | P11142;P11142-2 | HSPA8 | Extracellular or secreted | 0.448427496 | 1.401136512 | 34.85531387 | 34.89012128 | 34.83775455 | 34.9883756 | 35.04233732 | 34.33969925 | 35.40590828 | 35.38817967 | 35.02799549 |
| Stress-70 protein, mitochondrial | P38646 | HSPA9 | Extracellular or secreted | -0.233404255 | 0.657893945 | 30.69544485 | 30.49046755 | 30.57235577 | 31.09961384 | 31.07259001 | 31.5180331 | 30.65112852 | 30.78939409 | 30.58351718 |
| Heat shock protein beta-1 | P04792 | HSPB1 | Extracellular or secreted | 0.603989408 | 0.481865698 | 27.03296652 | 26.02424115 | 25.66424479 | 26.31127616 | 26.51392887 | 27.27985411 | 27.45646283 | 27.61757878 | 26.15118242 |
| Intraflagellar transport protein 25 hor | Q9Y547 | HSPB11 | Extracellular or secreted | -0.119341958 | 0.127961435 | 27.73563504 | 27.76040322 | 27.60016995 | 27.51863347 | 27.86260151 | 27.06702135 | 27.8312276 | 27.72552946 | 26.85744934 |
| Hsp70-binding protein 1 | Q9NZL4;Q9NZL4-3;Q9NZL4-2 | HSPBP1 | Other | 0.469278425 | 0.857433863 | 27.83623329 | 28.30155843 | 28.11187559 | 27.99593411 | 27.85126618 | 27.78612532 | 28.66013537 | 28.65512713 | 28.03406924 |
| 60 kDa heat shock protein, mitochond | P10809 | HSPD1 | Extracellular or secreted | -0.227645813 | 0.76559605 | 33.31378729 | 33.31770051 | 33.4108436 | 33.80914029 | 33.92927732 | 33.91117239 | 33.50320272 | 33.44097246 | 33.21884808 |
| 10 kDa heat shock protein, mitochond | P61604 | HSPE1 | Extracellular or secreted | 0.225393175 | 0.659320178 | 32.23203553 | 32.29532006 | 32.21572755 | 32.12105341 | 31.91338555 | 32.06493558 | 32.47351216 | 32.51852561 | 32.10537061 |
| Basement membrane-specific hepara | P98160 | HSPG2 | Extracellular or secreted | 1.911316632 | 1.155909717 | 26.27668103 | 25.58523054 | 26.26373942 | 26.35309485 | 25.90455984 | 30.37056829 | 28.19446208 | 28.1753614 | 29.7410634 |
| Heat shock protein 105 kDa | Q92598;Q92598-2;Q92598-4 | HSPH1 | Extracellular or secreted | 0.473367763 | 0.992606412 | 31.54871458 | 31.57844224 | 31.4953845 | 32.12903102 | 32.22943059 | 31.41099519 | 32.42557408 | 32.34543446 | 31.84509381 |
| HIV Tat-specific factor 1 | O43719 | HTATSF1 | Other | 0.12344017 | 0.344806301 | 29.01297211 | 28.96801695 | 28.98736616 | 28.87105963 | 28.7945468 | 28.40279668 | 28.77307002 | 28.93724925 | 29.17838041 |
| Serine protease HTRA1 | Q92743 | HTRA1 | Extracellular or secreted | 1.41901367 | 0.968553887 | 24.54067342 | 23.42636816 | 24.71947113 | 26.93449607 | 26.42616019 | 27.16949552 | 26.56353134 | 26.4458171 | 27.85602481 |
| Serine protease HTRA2, mitochondria | O43464-3;O43464;O43464-2 | HTRA2 | Extracellular or secreted | 0.290823056 | 0.16693921 | 26.71214114 | 25.89809132 | 25.82070135 | 25.60803345 | 25.53883969 | 24.43669731 | 26.52777324 | 26.53226947 | 24.8196786 |
| Huntingtin | P42858 | HTT | Other | 0.73701794 | 0.729345427 | 23.15744501 | 23.84009329 | 23.62878637 | 21.46195692 | 21.48784698 | 23.78785714 | 23.83778575 | 23.81049005 | 23.24477088 |
| E3 ubiquitin-protein ligase HUWE1 | Q7Z6Z7;Q7Z6Z7-2;Q7Z6Z7-3 | HUWE1 | Extracellular or secreted | 0.173550011 | 0.27457113 | 31.06881591 | 31.18429313 | 31.10224478 | 30.67244555 | 30.72362676 | 30.70734344 | 31.35026452 | 31.26153304 | 30.63823725 |
| Hypoxia up-regulated protein 1 | Q9Y4L1;Q9Y4L1-2 | HYOU1 | Extracellular or secreted | 0.128218723 | 0.284973084 | 28.9762359 | 28.89527388 | 29.04132089 | 28.95939165 | 29.02588382 | 29.67741093 | 29.12250437 | 29.04624792 | 29.50366241 |
| Huntingtin-interacting protein K | Q9NX55 | HYPK | Other | -0.162085722 | 0.258312927 | 26.1960487 | 26.48608068 | 26.6345031 | 26.69431601 | 26.87829943 | 25.63707655 | 26.53826248 | 25.92172928 | 26.31691332 |
| Isoamyl acetate-hydrolyzing esterase | Q2TAA2;Q2TAA2-2 | IAH1 | Extracellular or secreted | -0.1814195 | 0.168833472 | 28.26767339 | 28.27927851 | 28.26602064 | 27.09285164 | 27.17025991 | 26.52121223 | 27.74400262 | 27.63747737 | 26.87290967 |
| Isoleucine--tRNA ligase, cytoplasmic | P41252 | IARS | Extracellular or secreted | -0.272692991 | 0.548861681 | 30.58817219 | 30.76817127 | 30.65215486 | 30.61219182 | 30.59503773 | 30.67016754 | 30.14976851 | 30.22710794 | 30.74799228 |
| Isoleucine--tRNA ligase, mitochondria | Q9NSE4 | IARS2 | Other | 0.053638928 | 0.046199318 | 28.00227989 | 28.29561565 | 28.27799365 | 28.28859266 | 28.43987837 | 30.04061565 | 28.38096487 | 28.30090418 | 29.15153568 |
| Putative transferase CAF17, mitochon | Q5T440 | IBA57 | Other | 0.752203885 | 0.69480385 | 23.29347726 | 22.95780206 | 23.37424672 | 21.27197776 | 22.37790421 | 24.48406861 | 23.10998046 | 24.07341004 | 23.95295946 |
| ICOS ligand | O75144;O75144-2;O75144-3 | ICOSLG | Extracellular or secreted | 0.486987754 | 0.277199436 | 21.43862741 | 20.55615513 | 21.89843763 | 21.33605648 | 21.99213417 | 22.26416194 | 20.95737699 | 22.12249943 | 23.12387322 |
| Insulin-degrading enzyme | P14735;P14735-2 | IDE | Extracellular or secreted | 0.745745665 | 4.144003533 | 28.03112678 | 28.09093538 | 28.07760258 | 27.88756712 | 27.7362159 | 27.82116379 | 28.79380297 | 28.62902271 | 28.6367171 |
| Isocitrate dehydrogenase [NADP] cyto | O75874 | IDH1 | Extracellular or secreted | -0.039772811 | 0.109256754 | 31.9847478 | 32.05086638 | 31.95722724 | 31.51031863 | 31.70968678 | 31.23380764 | 31.71001536 | 31.74659155 | 31.64740189 |
| Isocitrate dehydrogenase [NADP], mit | P48735;P48735-2 | IDH2 | Extracellular or secreted | -0.480429882 | 0.373784179 | 26.76572355 | 26.97825606 | 27.20732062 | 28.05966603 | 27.85750867 | 29.24079479 | 26.91877573 | 26.68404957 | 28.01051991 |
| Isocitrate dehydrogenase [NAD] subu | P50213;P50213-2 | IDH3A | Other | 0.193400144 | 0.217764083 | 26.73918109 | 26.11447932 | 26.31644727 | 27.51788296 | 27.59184976 | 27.75837125 | 26.89608272 | 27.10029082 | 27.60293272 |
| Isocitrate dehydrogenase [NAD] subu | O43837-2;O43837 | IDH3B | Other | -0.579507245 | 1.071385201 | 24.99196188 | 25.22390861 | 25.65168283 | 25.56264374 | 26.07879385 | 26.51207589 | 24.89129236 | 24.93992996 | 25.44078935 |
| Isopentenyl-diphosphate Delta-isome | Q13907;Q13907-2 | IDI1 | Other | 2.576510307 | 4.769345384 | 27.94703938 | 27.89000528 | 27.73343857 | 27.78450376 | 27.87220518 | 26.90000497 | 30.2724201 | 30.25867206 | 30.26203734 |
| Iduronate 2-sulfatase;Iduronate 2-sulf | P22304;P22304-3;P22304-2 | IDS | Extracellular or secreted | 0.698046937 | 0.319460933 | 21.93402672 | 20.68890243 | 23.76273645 | 20.91880666 | 24.38231794 | 23.77525098 | 22.08421436 | 23.41493293 | 24.32601411 |
| Gamma-interferon-inducible protein | Q16666-3;Q16666-6;Q16666 | IFI16 | Other | -2.779291565 | 1.288079383 | 24.91857102 | 25.76425609 | 25.35127599 | 23.35366698 | 23.64030761 | 28.52738549 | 21.61557935 | 21.71606706 | 24.10821049 |
| Gamma-interferon-inducible lysosom | P13284 | IFI30 | Extracellular or secreted | -0.503943999 | 1.540090237 | 27.58956577 | 27.3948477 | 27.47243279 | 27.69205567 | 27.89162844 | 27.00345992 | 27.03443662 | 26.77718407 | 27.19854246 |
| Interferon-induced 35 kDa protein | P80217;P80217-2 | IFI35 | Other | 0.342092584 | 0.313046913 | 25.99761469 | 25.69659935 | 25.80368319 | 24.54445181 | 23.90177802 | 23.49265026 | 25.12601026 | 25.53466076 | 25.08399539 |
| Interferon-induced protein with tetra | Q5T764;P09914-2;P09914 | IFIT1B;IFIT1 | Other | 0.337686114 | 1.331600631 | 23.9685993 | 23.88892604 | 23.88529942 | 23.82667727 | 23.79936622 | 24.29768092 | 24.08691304 | 24.39348997 | 24.36592992 |
| Interferon-induced transmembrane p | P13164;Q01629;Q01628 | IFITM1;IFITM2;IF | Extracellular or secreted | 1.202873102 | 0.993009232 | 23.70017638 | 23.61582627 | 23.66524829 | 21.01398716 | 22.93944503 | 25.57019273 | 24.55326883 | 24.62856391 | 24.6792245 |
| Intraflagellar transport protein 27 hor | Q9BW83-2;Q9BW83 | IFT27 | Other | -0.177793786 | 0.26827149 | 23.19413703 | 23.44240393 | 23.43377185 | 23.03286566 | 23.4820999 | 23.52850371 | 23.17700338 | 23.58474326 | 22.76176304 |
| Immunoglobulin-binding protein 1 | P78318 | IGBP1 | Other | 0.507691398 | 0.623684651 | 24.60376759 | 25.74388709 | 25.74773305 | 25.07512525 | 25.06328151 | 24.62036438 | 25.05693038 | 25.85526448 | 26.03795877 |
| Insulin-like growth factor 1 receptor;I | P08069 | IGF1R | Extracellular or secreted | 0.798745407 | 2.635366753 | 24.10677378 | 24.0480165 | 24.06693926 | 24.15437313 | 23.81889887 | 24.67341303 | 24.75552169 | 24.94458411 | 25.13033772 |
| Insulin-like growth factor II;Insulin-like | P01344;P01344-2;P01344-3 | IGF2 | Extracellular or secreted | -1.864029254 | 1.062593642 | 22.78367243 | 23.46587923 | 22.91807052 | 24.25673908 | 25.56313854 | 20.43659525 | 20.65739024 | 22.56609214 | 20.89647738 |
| Insulin-like growth factor 2 mRNA-bin | Q9NZI8;Q9NZI8-2 | IGF2BP1 | Other | 0.157258262 | 0.102036211 | 23.40028221 | 23.74242293 | 24.34692162 | 24.43096803 | 23.52169149 | 25.05283269 | 23.39270418 | 24.27383884 | 25.05279124 |
| Insulin-like growth factor 2 mRNA-bin | Q9Y6M1-5;Q9Y6M1-6;Q9Y6M | IGF2BP2 | Other | 0.091817506 | 0.05905811 | 23.8931919 | 24.55221362 | 24.4169954 | 23.42636816 | 23.74847542 | 26.47171986 | 23.85471775 | 24.82478514 | 24.85043181 |
| Insulin-like growth factor 2 mRNA-bin | O00425;O00425-2 | IGF2BP3 | Other | 0.053515601 | 0.070492012 | 23.26570775 | 23.35056803 | 23.23788915 | 22.28243352 | 22.04596891 | 23.15670465 | 22.84747184 | 23.2237908 | 22.75892017 |
| Cation-independent mannose-6-phos | P11717 | IGF2R | Extracellular or secreted | -0.183564428 | 0.20177452 | 26.46488273 | 26.55080549 | 26.52049304 | 27.25124069 | 27.36961186 | 27.39067227 | 26.44249885 | 26.45043413 | 27.33022677 |
| Insulin-like growth factor-binding prot | P18065 | IGFBP2 | Extracellular or secreted | 0.796479006 | 0.430256525 | 29.29622854 | 29.23500687 | 29.22501723 | 30.86104891 | 30.84041149 | 32.32503947 | 30.3959101 | 30.55178839 | 32.33311478 |
| Insulin-like growth factor-binding prot | P17936;P17936-2 | IGFBP3 | Extracellular or secreted | -1.252270136 | 1.372845159 | 27.21066837 | 27.20125632 | 27.09345625 | 28.1729321 | 28.28881265 | 29.05302438 | 26.31430498 | 26.16905581 | 27.26990384 |
| Insulin-like growth factor-binding prot | P22692;P22692-2 | IGFBP4 | Extracellular or secreted | 1.01422773 | 0.496808506 | 27.82638633 | 27.68565435 | 27.68224206 | 29.86503794 | 30.01358451 | 31.55439828 | 29.32375717 | 29.61719341 | 31.41538435 |
| Insulin-like growth factor-binding prot | P24593 | IGFBP5 | Extracellular or secreted | -3.127945826 | 0.933246679 | 25.51177437 | 25.53424517 | 25.8446734 | 24.80847837 | 24.33196987 | 28.61118941 | 20.61857411 | 22.29662806 | 25.02212564 |
| Insulin-like growth factor-binding prot | P24592 | IGFBP6 | Extracellular or secreted | -1.558615144 | 0.427282536 | 21.09061643 | 22.74400518 | 23.16914758 | 24.14024189 | 24.865094 | 28.77369888 | 20.18773385 | 23.64812055 | 23.87970215 |
| Insulin-like growth factor-binding prot | Q8WX77 | IGFBPL1 | Extracellular or secreted | 1.118745615 | 0.63691186 | 22.56437188 | 22.86938374 | 22.84166799 | 26.73402031 | 26.6597612 | 25.06389864 | 25.13969561 | 25.56743902 | 26.0156541 |
| Immunoglobulin J chain | P01591 | IGJ | Extracellular or secreted | 1.102898668 | 0.525270174 | 21.02649839 | 21.20724272 | 22.01571409 | 22.52160011 | 20.53265386 | 21.83986079 | 23.14758878 | 23.66957974 | 21.06331247 |
| Immunoglobulin superfamily member | Q969P0;Q969P0-3 | IGSF8 | Extracellular or secreted | 0.295809645 | 0.273555398 | 28.07907879 | 28.0001319 | 27.82213686 | 28.29065921 | 28.23301904 | 28.96801695 | 27.93893229 | 28.39827725 | 29.24674078 |
| Protein Red | Q13123 | IK | Extracellular or secreted | -0.186435016 | 0.36668936 | 23.78026662 | 23.12508802 | 23.46961002 | 23.36813021 | 22.9588465 | 23.45951467 | 22.98959061 | 23.00595412 | 23.52587824 |
| Elongator complex protein 1 | O95163 | IKBKAP | Other | 0.429744409 | 1.145762401 | 28.47123146 | 28.41856444 | 28.48987447 | 28.50324538 | 28.19854246 | 28.70994144 | 28.98125379 | 29.08365056 | 28.62002871 |
| NF-kappa-B essential modulator | Q9Y6K9;Q9Y6K9-2 | IKBKG | Other | -2.004726426 | 1.636262014 | 25.17077564 | 25.26241376 | 25.55015963 | 25.88005261 | 25.92669124 | 25.83511304 | 24.25031924 | 23.60817449 | 22.93992996 |
| Interleukin-18 | Q14116;Q14116-2 | IL18 | Extracellular or secreted | 1.318488327 | 0.910395348 | 26.55978807 | 26.65921679 | 26.69471453 | 26.29804816 | 26.46265366 | 25.51255819 | 28.26740551 | 28.17840416 | 26.60314502 |
| Interleukin-1 receptor accessory prote | Q9NPH3-2;Q9NPH3;Q9NPH3 | IL1RAP | Extracellular or secreted | 0.732471415 | 0.479779389 | 25.06138734 | 25.56182841 | 25.1330064 | 26.32673422 | 26.37237131 | 26.4943147 | 26.06387807 | 25.9127134 | 27.69564397 |
| Interleukin-1 receptor antagonist prot | P18510-2;P18510-4;P18510; | IL1RN | Extracellular or secreted | 0.160980714 | 0.50270547 | 23.67949299 | 23.35595324 | 24.13508298 | 23.77243459 | 23.63765702 | 23.27497639 | 23.95544487 | 23.75999705 | 23.69529882 |
| Interleukin-27 receptor subunit alpha | Q6UWB1 | IL27RA | Extracellular or secreted | -0.011038524 | 0.012214782 | 21.64552827 | 21.67581579 | 22.01089547 | 23.03106577 | 22.67846174 | 21.72575567 | 21.97722122 | 21.88185034 | 22.49157423 |
| Interleukin enhancer-binding factor 2 | Q12905 | ILF2 | Other | -0.184181108 | 0.294846123 | 31.64525579 | 31.80305033 | 31.69780758 | 30.95220133 | 30.82289661 | 31.47522877 | 31.384479 | 31.43549366 | 30.82570423 |
| Interleukin enhancer-binding factor 3 | Q12906;Q12906-5;Q12906-2 | ILF3 | Other | 0.004617317 | 0.003485349 | 27.84493673 | 27.96350406 | 27.85090865 | 26.15609126 | 26.3145643 | 26.98609149 | 27.62400972 | 27.31172656 | 26.63616392 |
| Interleukin enhancer-binding factor 3 | Q12906-7;Q12906-4;Q12906 | ILF3 | Other | -0.181707851 | 0.300297957 | 32.82498387 | 32.90465602 | 32.84871684 | 32.03515144 | 32.08885234 | 32.42337157 | 32.50052221 | 32.55885165 | 31.95836863 |
| Integrin-linked protein kinase | Q13418-2;Q13418;Q13418-3 | ILK | Extracellular or secreted | 0.80669425 | 0.968919317 | 22.42180494 | 22.54177253 | 22.90277181 | 22.3305482 | 22.40278205 | 22.49071772 | 23.51508765 | 23.72192358 | 22.72827014 |
| Integrin-linked kinase-associated serin | Q9H0C8 | ILKAP | Other | -0.034524524 | 0.048195071 | 27.54170755 | 27.56666994 | 27.64821658 | 28.02870527 | 28.04166032 | 28.05491403 | 28.11386359 | 27.85072985 | 27.37276983 |
| Acetolactate synthase-like protein | A1L0T0 | ILVBL | Extracellular or secreted | -1.096902935 | 0.495487875 | 23.71313811 | 22.90391199 | 24.15746043 | 23.90472061 | 20.5845669 | 25.89735285 | 22.6758266 | 21.09450649 | 23.51953356 |
| MICOS complex subunit MIC60 | Q16891-2;Q16891;Q16891-3 | IMMT | Other | -1.675290734 | 0.809093447 | 27.97481463 | 27.86679256 | 28.08263586 | 27.0868928 | 27.04364312 | 29.10931199 | 24.96322842 | 25.93012515 | 27.66281971 |
| Inositol monophosphatase 1 | P29218;P29218-3;P29218-2 | IMPA1 | Extracellular or secreted | 0.429111596 | 1.800324881 | 27.15749898 | 27.23872383 | 27.2459932 | 26.92263684 | 27.39247492 | 26.74637531 | 27.55193504 | 27.69444886 | 27.39280243 |
| Inositol monophosphatase 2 | O14732-2;O14732 | IMPA2 | Other | 0.272033814 | 0.457155539 | 25.75920979 | 26.08019768 | 26.03044286 | 25.51481685 | 25.41541659 | 25.65212066 | 26.03791689 | 26.3833072 | 25.62097956 |
| Protein IMPACT | Q9P2X3;Q9P2X3-2 | IMPACT | Other | 0.089005988 | 0.11194994 | 25.49655637 | 25.65239424 | 25.58860828 | 25.27081287 | 25.54819092 | 25.28326756 | 26.04730933 | 25.50763715 | 25.1319866 |
| Inositol monophosphatase 3 | Q9NX62 | IMPAD1 | Other | 0.287225534 | 0.160065063 | 21.36009345 | 20.48412277 | 20.88532143 | 22.339302 | 22.42652173 | 23.98276663 | 22.06753021 | 21.48605919 | 23.04715121 |
| Inosine-5-monophosphate dehydroge | P20839;P20839-3;P20839-7; | IMPDH1 | Other | 0.696919525 | 0.873689034 | 25.15143406 | 25.52169149 | 25.34126499 | 25.84433817 | 25.70395573 | 26.02013423 | 26.63588725 | 26.55990474 | 25.68637592 |
| Inosine-5-monophosphate dehydroge | P12268 | IMPDH2 | Extracellular or secreted | 0.920895441 | 2.42445253 | 29.66351546 | 29.68784133 | 29.68692354 | 30.38566219 | 30.46168224 | 30.6995047 | 31.0175057 | 30.87862101 | 31.15912435 |
| Immortalization up-regulated protein | Q9GZP8-2;Q9GZP8 | IMUP | Other | 0.688522819 | 0.967789312 | 27.67976142 | 27.82662878 | 27.72974872 | 27.99808835 | 27.66824109 | 27.06002695 | 28.37683178 | 28.80519724 | 27.8647871 |
| Inverted formin-2 | Q27J81-2;Q27J81 | INF2 | Other | -0.297025762 | 1.14449993 | 26.37790752 | 26.3510569 | 26.35576846 | 26.13731295 | 26.24401593 | 26.31767248 | 26.08086859 | 25.81169081 | 26.10823043 |
| Inhibin beta A chain | P08476 | INHBA | Extracellular or secreted | 0.521622629 | 0.412248005 | 25.90306614 | 25.80570157 | 25.49750079 | 26.37464472 | 26.0248752 | 25.81787483 | 25.91202808 | 25.99098875 | 27.37368269 |
| SOSS complex subunit C | Q9NRY2;Q9NRY2-2 | INIP | Other | 0.170605619 | 0.098132492 | 23.48873355 | 23.23476482 | 23.65392529 | 23.45525597 | 23.8781123 | 23.21784354 | 23.81440723 | 24.57548247 | 22.5862449 |
| Inositol polyphosphate 1-phosphatase | P49441 | INPP1 | Other | 0.124741154 | 0.20106589 | 21.74201167 | 21.48143485 | 21.54415698 | 21.57885809 | 22.33067377 | 22.26732572 | 22.18860937 | 22.07643071 | 21.58141392 |
| Type II inositol 1,4,5-trisphosphate 5- | P32019-3;P32019-4;P32019-2 | INPP5B | Extracellular or secreted | -0.69909101 | 0.402530359 | 23.72004548 | 21.56589385 | 21.487456 | 23.61031658 | 23.49911402 | 22.58012605 | 21.84976321 | 23.17636432 | 21.10807543 |
| Phosphatidylinositol 4,5-bisphosphate | Q15735-3;Q15735-2;Q15735 | INPP5J | Extracellular or secreted | -2.773414792 | 1.357560923 | 22.05391778 | 22.2371301 | 25.2579621 | 26.04901428 | 25.40269098 | 25.58373953 | 23.11561078 | 21.36285215 | 20.49352009 |
| Phosphatidylinositol 3,4,5-trisphospha | O15357-2;O15357 | INPPL1 | Extracellular or secreted | -0.233581638 | 0.140968591 | 22.34559583 | 23.02943215 | 23.24597508 | 22.59450321 | 22.39079033 | 20.78052695 | 23.00645073 | 22.21602348 | 21.27019265 |
| Integrator complex subunit 2 | Q9H0H0 | INTS2 | Other | 1.120436273 | 0.866869948 | 19.44679658 | 21.0129082 | 20.85703637 | 21.75324111 | 21.39446926 | 24.12083966 | 22.80943059 | 22.42611219 | 22.41841163 |
| Integrator complex subunit 3 | Q68E01-2;Q68E01;Q68E01-3 | INTS3 | Extracellular or secreted | 0.281434226 | 1.168367295 | 26.75582727 | 27.06455661 | 27.01073331 | 26.82213686 | 26.98869387 | 27.43872892 | 27.41425554 | 27.13486763 | 27.33551793 |
| Integrator complex subunit 7 | Q9NVH2-4;Q9NVH2-3;Q9NV | INTS7 | Other | 0.17634494 | 0.156222602 | 21.28818877 | 23.17697296 | 22.91327512 | 23.27241565 | 22.53194911 | 23.74375871 | 22.51679554 | 23.41944075 | 23.05607868 |
| Integrator complex subunit 9 | Q9NV88-2;Q9NV88 | INTS9 | Other | 0.68055093 | 0.584067915 | 23.27383884 | 22.67231677 | 20.04625607 | 23.03525084 | 21.03721753 | 22.0606607 | 22.91402374 | 22.887997 | 22.30240242 |
| Importin-11 | Q9UI26;Q9UI26-2 | IPO11 | Extracellular or secreted | 0.284313407 | 0.749811008 | 26.56101262 | 26.84959695 | 26.85281446 | 26.6191892 | 26.00058325 | 26.4774443 | 27.05444832 | 26.8793516 | 26.59946069 |
| Importin-4 | Q8TEX9;Q8TEX9-2 | IPO4 | Other | 0.520204026 | 0.653969928 | 27.1969492 | 27.46029615 | 27.28912057 | 27.5673955 | 27.37343379 | 27.10938672 | 28.18593585 | 28.12055296 | 27.25241424 |
| Importin-5 | O00410;O00410-3;O00410-2 | IPO5 | Other | 0.394840034 | 0.638783216 | 30.99711934 | 31.19466168 | 31.11091788 | 30.99104823 | 30.97903334 | 30.83266172 | 31.62736234 | 31.66449912 | 30.94537974 |
| Importin-7 | O95373 | IPO7 | Other | 0.225609765 | 0.690316619 | 29.98793533 | 30.20384843 | 30.07239834 | 29.86639464 | 30.03078486 | 30.13075012 | 30.5098469 | 30.22905807 | 30.08398018 |
| Importin-8 | O15397;O15397-2 | IPO8 | Other | 0.395594914 | 0.687608732 | 24.53011243 | 24.38455928 | 24.5920637 | 24.66258885 | 24.32505342 | 25.08598165 | 25.17649367 | 25.24702612 | 24.55344462 |
| Importin-9 | Q96P70 | IPO9 | Extracellular or secreted | 0.378387848 | 0.42546904 | 29.42219886 | 29.65505887 | 29.62891845 | 28.81796629 | 29.11872266 | 28.97642712 | 29.89879483 | 30.05180645 | 28.9942084 |
| Ras GTPase-activating-like protein IQ( | P46940 | IQGAP1 | Extracellular or secreted | -0.274881419 | 0.924839218 | 33.40578151 | 33.48435773 | 33.42166732 | 32.73628446 | 32.80179842 | 32.82505972 | 32.85262845 | 32.84602845 | 32.81417341 |
| Ras GTPase-activating-like protein IQ( | Q13576;Q13576-3;Q13576-2 | IQGAP2 | Extracellular or secreted | -0.721750702 | 1.227477369 | 28.87273358 | 28.25394744 | 28.25151159 | 27.16269373 | 27.41005508 | 27.40055909 | 27.16585911 | 26.87911785 | 27.46552119 |
| Ras GTPase-activating-like protein IQ( | Q86VI3 | IQGAP3 | Extracellular or secreted | 1.411092901 | 1.797703649 | 21.2585177 | 20.7395496 | 21.45659324 | 20.02979564 | 20.25056656 | 22.05999396 | 21.80660083 | 22.57724717 | 22.74693906 |
| Iron-responsive element-binding prot | P48200 | IREB2 | Other | -0.965873507 | 1.461769855 | 25.44401671 | 25.5085139 | 25.52572892 | 25.17249343 | 24.65277715 | 25.04143056 | 24.4777222 | 24.52641566 | 23.77072196 |
| Interferon regulatory factor 2-binding | Q8IU81 | IRF2BP1 | Other | -4.081546472 | 0.968719284 | 27.17464732 | 27.52566919 | 27.71973223 | 28.14107067 | 26.68271089 | 28.72439139 | 21.80762378 | 22.37111192 | 26.56073573 |
| Interferon regulatory factor 2-binding | Q7Z5L9-2;Q7Z5L9 | IRF2BP2 | Other | 0.577531459 | 0.598729268 | 26.44504352 | 25.74023678 | 26.02599468 | 25.95238187 | 25.93138822 | 26.13766474 | 26.22414789 | 26.27597101 | 27.34883039 |
| Interferon regulatory factor 3 | Q14653;Q14653-4 | IRF3 | Other | -0.261194294 | 0.432974638 | 26.80476657 | 26.67962721 | 26.78656159 | 26.78892758 | 26.69365157 | 26.68150502 | 26.74124025 | 26.66695535 | 26.02574129 |
| Interferon regulatory factor 6 | O14896;O14896-2 | IRF6 | Extracellular or secreted | 0.444462974 | 0.479234458 | 23.79143875 | 24.11441975 | 23.95588825 | 23.81440723 | 24.65741877 | 25.31181316 | 24.31948249 | 24.4732073 | 25.36339208 |
| Immunity-related GTPase family Q pro | Q8WZA9 | IRGQ | Other | 0.184608464 | 0.353733926 | 28.30734654 | 28.27963275 | 28.37807295 | 27.98196143 | 27.77007919 | 27.59967351 | 28.39439799 | 28.43041778 | 27.88739281 |
| Iron-sulfur cluster assembly enzyme IS | Q9H1K1-2;Q9H1K1 | ISCU | Other | 0.876937219 | 0.287520736 | 24.1733325 | 21.7759056 | 20.75871025 | 21.86015245 | 20.82068035 | 22.11524966 | 23.78496546 | 23.95491265 | 20.64294895 |
| Ubiquitin-like protein ISG15 | P05161 | ISG15 | Extracellular or secreted | -0.22688663 | 0.046114466 | 28.66648137 | 28.57437843 | 28.52062042 | 27.986851 | 28.03149492 | 27.48446817 | 29.5861867 | 29.5383809 | 24.82691967 |
| Interferon-stimulated gene 20 kDa pro | Q96AZ6 | ISG20 | Other | 0.191457207 | 0.09805408 | 23.1398361 | 23.06761227 | 23.15017927 | 24.99929332 | 24.70498475 | 27.1054955 | 24.90994727 | 24.80193329 | 23.94619166 |
| Isochorismatase domain-containing p | Q96CN7 | ISOC1 | Extracellular or secreted | -0.267265464 | 1.016588363 | 28.77818724 | 29.2302727 | 29.05935659 | 28.55997765 | 28.40758565 | 28.58168687 | 28.46929173 | 28.56361862 | 28.47382662 |
| Isochorismatase domain-containing p | Q96AB3;Q96AB3-2;Q96AB3-3 | ISOC2 | Other | 0.592608307 | 1.434728413 | 27.65102584 | 27.60660819 | 27.43332616 | 27.46933055 | 27.58226134 | 27.94937915 | 28.21803392 | 28.46072579 | 27.94503083 |
| IST1 homolog | P53990-2;P53990-4;P53990-3 | IST1 | Extracellular or secreted | -0.275469869 | 0.44461309 | 27.89186017 | 28.11351589 | 28.1850379 | 28.12755587 | 28.30951653 | 27.4912898 | 28.00496039 | 27.93775437 | 27.29026372 |
| Pre-mRNA-splicing factor ISY1 homolo | Q9ULR0;Q9ULR0-2;Q9ULR0-1 | ISY1 | Other | 0.515478516 | 3.875616858 | 24.6646517 | 24.90747384 | 24.96318432 | 24.93741765 | 24.75445165 | 24.63150881 | 25.37567239 | 25.3191379 | 25.28096923 |
| Inositol-3-phosphate synthase 1 | Q9NPH2-3;Q9NPH2 | ISYNA1 | Other | 0.380163255 | 0.302124032 | 26.54941066 | 25.80299334 | 23.04301727 | 24.86910129 | 25.11119904 | 25.42005144 | 25.54701429 | 25.88079999 | 25.110562 |
| Inositol-3-phosphate synthase 1 | Q9NPH2-2 | ISYNA1 | Other | -0.678937386 | 1.620219399 | 32.98862955 | 33.09721362 | 32.99578926 | 32.48682873 | 32.49612299 | 32.30209252 | 32.17566477 | 32.24061853 | 31.73024287 |
| E3 ubiquitin-protein ligase Itchy homo | Q96J02-2;Q96J02;Q96J02-3 | ITCH | Extracellular or secreted | 0.535077228 | 2.336973035 | 25.76592584 | 25.67338607 | 25.90359486 | 26.15792296 | 26.04818285 | 25.83918033 | 26.53589199 | 26.26808406 | 26.4953521 |
| Integrin-alpha FG-GAP repeat-contain | Q969R8 | ITFG2 | Other | 0.410526302 | 0.787851308 | 26.12518263 | 25.99892763 | 25.90483543 | 25.98428876 | 26.06901061 | 25.80225385 | 26.73647398 | 26.37411403 | 26.06324036 |
| Protein ITFG3 | Q9H0X4;Q9H0X4-2 | ITFG3 | Extracellular or secreted | 0.435893313 | 0.971440681 | 24.95659736 | 24.56540661 | 24.94779743 | 24.4093754 | 24.29326676 | 24.87960868 | 24.864055 | 25.02512875 | 25.44452231 |
| Integrin alpha-3;Integrin alpha-3 heav | P26006;P26006-1 | ITGA3 | Extracellular or secreted | 1.197287113 | 1.918227361 | 29.13609081 | 29.1316089 | 29.24837051 | 28.92699656 | 28.78998479 | 29.13040646 | 30.13001361 | 30.03420046 | 30.60937628 |
| Integrin alpha-6;Integrin alpha-6 heav | P23229-5;P23229-3;P23229; | ITGA6 | Extracellular or secreted | 1.452758113 | 3.416341913 | 29.7737146 | 29.80274997 | 29.9147047 | 29.44251071 | 29.54912786 | 30.4137715 | 31.03773889 | 31.25248191 | 31.51634321 |
| Integrin alpha-V;Integrin alpha-V heav | P06756-3;P06756;P06756-2 | ITGAV | Extracellular or secreted | 0.330775374 | 0.948395321 | 25.44480663 | 25.53907643 | 25.69954111 | 25.04193181 | 25.46198271 | 24.82376528 | 25.66016938 | 25.87335568 | 25.46435305 |
| Integrin beta-1 | P05556;P05556-2;P05556-5; | ITGB1 | Extracellular or secreted | 0.98593655 | 1.551283648 | 30.05452595 | 29.85598027 | 29.99806144 | 29.81272843 | 29.96719248 | 30.7310121 | 30.69353526 | 30.98250553 | 31.49151918 |
| Integrin beta-4 | P16144 | ITGB4 | Extracellular or secreted | 1.433549013 | 1.788496201 | 30.6405662 | 30.51352232 | 30.54765788 | 30.14126561 | 30.18919213 | 31.14217904 | 31.58799343 | 31.80512803 | 32.49471716 |
| Integrin beta-4 | P16144-2;P16144-4;P16144-3 | ITGB4 | Extracellular or secreted | 2.229078441 | 1.455253474 | 20.19142369 | 22.98082083 | 24.0619229 | 21.2134456 | 22.70163558 | 24.96516777 | 24.40639395 | 24.50397322 | 25.83407634 |
| Integrin beta-6 | P18564;P18564-2 | ITGB6 | Extracellular or secreted | 0.905334155 | 2.851116478 | 25.91314727 | 25.90795512 | 26.17142483 | 25.12970908 | 25.46023365 | 25.91800226 | 26.5363071 | 26.73840815 | 26.69152331 |
| Integral membrane protein 2B;BRI2, n | Q9Y287-2;Q9Y287 | ITM2B | Extracellular or secreted | 1.361683842 | 1.188574001 | 22.36169324 | 20.65210972 | 20.75197222 | 22.46163931 | 22.1123279 | 23.78216799 | 23.296141 | 22.76068747 | 24.08917825 |
| Inosine triphosphate pyrophosphatas | Q9BY32;Q9BY32-2;Q9BY32-3 | ITPA | Other | 0.592282817 | 0.889379447 | 29.13312892 | 29.22393622 | 29.11681569 | 28.12356542 | 28.08967329 | 28.36640526 | 29.47136714 | 29.55039452 | 28.78184919 |

| | | | | | | | | | | | | | | |
|---|---|---|---|---|---|---|---|---|---|---|---|---|---|---|
| Inositol 1,4,5-trisphosphate receptor t | Q14573 | ITPR3 | Extracellular or secreted | -2.636511779 | 0.801933204 | 24.38850611 | 23.64669318 | 24.06693926 | 24.10557543 | 23.47237321 | 27.41530375 | 19.85517592 | 21.58096145 | 24.20202276 |
| Intersectin-1 | Q15811-6;Q15811-12;Q1581 | ITSN1 | Extracellular or secreted | -0.522959337 | 0.50540385 | 25.1201673 | 25.32711127 | 25.12679804 | 25.76240698 | 25.35675929 | 24.32457283 | 25.41918354 | 24.30535596 | 24.21549034 |
| Intersectin-2 | Q9NZM3-4;Q9NZM3-3;Q9NZ | ITSN2 | Extracellular or secreted | 0.285426583 | 0.203056532 | 23.08917825 | 20.53372734 | 21.25971206 | 23.68088829 | 22.65170995 | 20.831319 | 22.22446808 | 22.59898388 | 22.05609523 |
| Involucrin | P07476 | IVL | Extracellular or secreted | -0.100640046 | 0.045367188 | 26.7278029 | 25.54111671 | 25.98888886 | 26.90035054 | 27.16863509 | 25.34302871 | 25.06870318 | 26.07312398 | 27.39116412 |
| Protein IWS1 homolog | Q96ST2;Q96ST2-3;Q96ST2-2 | IWS1 | Other | -0.592503182 | 1.73709837 | 26.96322842 | 26.87537271 | 26.88098677 | 27.44931668 | 27.51126163 | 26.39543621 | 26.24298086 | 26.4633243 | 26.55398651 |
| Protein jagged-1 | P78504;P78504-2 | JAG1 | Extracellular or secreted | 0.039648793 | 0.02299834 | 28.99685006 | 29.03938719 | 28.87088331 | 28.88623019 | 28.85528231 | 29.79873918 | 28.56416397 | 28.60194156 | 30.17652696 |
| Protein jagged-2 | Q9Y219-2;Q9Y219 | JAG2 | Extracellular or secreted | 0.978415789 | 0.906579292 | 21.49949576 | 19.99378809 | 22.03397856 | 22.70336965 | 22.11357534 | 23.77052034 | 22.63087588 | 23.31465073 | 23.04708463 |
| Bifunctional arginine demethylase and | Q6NYC1-2;Q6NYC1;Q6NYC1- | JMJD6 | Extracellular or secreted | 0.285638051 | 0.572708832 | 24.97473806 | 25.12060239 | 24.99701163 | 25.42460739 | 25.47125472 | 25.14748009 | 25.75281958 | 25.55891276 | 25.11302897 |
| JmjC domain-containing protein 7 | P0C870 | JMJD7 | Other | -0.059681361 | 0.100422477 | 22.739212 | 23.16186382 | 23.55496725 | 22.65318184 | 22.36220176 | 22.84136087 | 22.60326955 | 23.07576181 | 22.79831833 |
| Protein JTB | O76095-2;O76095 | JTB | Extracellular or secreted | -0.024229005 | 0.043319815 | 22.56211383 | 21.99569263 | 22.31185473 | 22.2219045 | 22.47417332 | 23.3540707 | 22.44111241 | 22.51838134 | 22.42772409 |
| Junction plakoglobin | P14923 | JUP | Extracellular or secreted | 0.458380994 | 1.505764826 | 31.1787247 | 31.30029327 | 31.40557865 | 30.77854759 | 30.70586147 | 31.20617968 | 31.5925895 | 31.72833827 | 31.3418079 |
| Lysine--tRNA ligase | Q15046;Q15046-2 | KARS | Extracellular or secreted | 0.163641026 | 0.61733399 | 31.92941842 | 31.99954065 | 31.86986171 | 31.78826643 | 31.84460755 | 31.43226109 | 32.07431385 | 32.05965958 | 31.78892758 |
| Histone acetyltransferase KAT7 | O95251-2;O95251-4;O95251 | KAT7 | Other | -0.989655963 | 2.044742091 | 24.23527627 | 23.90985574 | 24.16951463 | 24.01707821 | 23.71994107 | 24.35937585 | 22.87297541 | 23.39636751 | 22.96721007 |
| Katanin p60 ATPase-containing subun | Q8IYT4-2;Q8IYT4 | KATNAL2 | Other | 0.177129755 | 0.334831982 | 29.16062769 | 29.04531074 | 29.17581346 | 28.50657829 | 28.66441435 | 28.19069979 | 29.01472874 | 29.20976246 | 28.67862023 |
| Kelch repeat and BTB domain-contain | Q8IY47 | KBTBD2 | Other | 1.292302235 | 1.14068714 | 23.40989328 | 23.75419676 | 23.31506551 | 21.9306246 | 20.63555282 | 20.76731766 | 23.35810171 | 23.55800771 | 23.86712261 |
| E3 ubiquitin-protein ligase KCMF1 | Q9P0J7 | KCMF1 | Other | 0.587689503 | 3.74004893 | 25.81031843 | 25.63953502 | 25.64138302 | 25.96602642 | 25.97615941 | 25.69381107 | 26.39984233 | 26.35999575 | 26.36684711 |
| Potassium channel subfamily K memb | O00180 | KCNK1 | Extracellular or secreted | 0.453456579 | 0.200605276 | 21.31779576 | 20.76220408 | 21.57109447 | 21.48803306 | 19.62181723 | 22.94154271 | 20.58646396 | 21.51050728 | 23.11464215 |
| Intermediate conductance calcium-ac | O15554 | KCNN4 | Extracellular or secreted | 1.930796752 | 2.271746578 | 23.14042914 | 22.35863832 | 22.3425066 | 22.75745593 | 22.64557224 | 22.60979826 | 24.13226124 | 24.62449786 | 24.96283142 |
| BTB/POZ domain-containing protein K | Q96CX2 | KCTD12 | Extracellular or secreted | -0.764783982 | 1.229437179 | 27.40096619 | 27.28550932 | 27.22967697 | 27.26847675 | 27.28347953 | 26.85922803 | 26.69590942 | 26.652531 | 26.02087603 |
| BTB/POZ domain-containing protein K | Q96SI1;Q96SI1-2 | KCTD15 | Other | 0.560188524 | 0.360849372 | 25.20258331 | 25.23286355 | 24.9817655 | 25.65236688 | 21.3818057 | 24.31056563 | 25.19651778 | 25.22703447 | 24.63798861 |
| BTB/POZ domain-containing protein K | Q9Y597-2;Q9Y597 | KCTD3 | Other | 0.465333596 | 0.47883508 | 22.30409041 | 22.37872455 | 22.58927716 | 21.63559945 | 21.09701712 | 23.58886558 | 22.18213608 | 22.93801023 | 23.07264162 |
| BTB/POZ domain-containing protein K | Q9NXV2 | KCTD5 | Other | -0.173891525 | 0.317571589 | 24.57438564 | 24.60969683 | 25.19539173 | 25.54164848 | 25.17047005 | 24.20687366 | 24.62282355 | 24.93723803 | 24.56749705 |
| BTB/POZ domain-containing protein K | Q7L273 | KCTD9 | Other | 0.957330355 | 0.461805327 | 21.86632495 | 23.1382743 | 22.17542041 | 23.44227736 | 22.78312774 | 21.44371164 | 24.54173709 | 23.82026312 | 21.93455906 |
| KDEL motif-containing protein 2 | Q7Z4H8;Q7Z4H8-2;Q7Z4H8-3 | KDELC2 | Extracellular or secreted | 0.11908434 | 0.207011268 | 26.64663825 | 26.82989787 | 26.89481147 | 26.30530377 | 26.28459186 | 26.94915648 | 26.58749278 | 26.5824767 | 27.14248339 |
| Keratinocyte differentiation factor 1 | Q8NAX2 | KDF1 | Other | 0.626191278 | 1.151044447 | 22.6599898 | 24.32471015 | 24.44031412 | 23.76060626 | 24.06134614 | 23.88855449 | 24.23666358 | 24.68522659 | 24.52444415 |
| Lysine-specific histone demethylase 1 | O60341;O60341-2 | KDM1A | Extracellular or secreted | -0.368250277 | 0.480139825 | 28.70454289 | 28.84493673 | 28.76673473 | 28.07989261 | 28.18016033 | 28.40462486 | 28.48933858 | 28.33492151 | 27.56143514 |
| Lysine-specific demethylase 2A | Q9Y2K7-4;Q9Y2K7-5;Q9Y2K7 | KDM2A | Other | -0.407257061 | 0.558765977 | 23.43147822 | 23.03654256 | 23.23971098 | 23.37039368 | 23.26184012 | 25.13025915 | 22.93212751 | 23.39335903 | 23.18785464 |
| Lysine-specific demethylase 3B | Q7LBC6;Q7LBC6-2 | KDM3B | Extracellular or secreted | -0.828512263 | 2.354880373 | 25.52139197 | 25.43501272 | 25.30962928 | 24.92206968 | 24.66784872 | 25.53703327 | 24.29642111 | 24.65441708 | 24.26011784 |
| Lysine-specific demethylase 5C | P41229-2;P41229;P41229-4;P | KDM5C | Other | 1.649325157 | 2.160261778 | 23.02678412 | 20.69481606 | 21.36694524 | 21.13029686 | 21.74408735 | 22.67128837 | 23.80252504 | 23.45713636 | 23.00542307 |
| 3-ketodihydrosphingosine reductase | Q06136;Q06136-2 | KDSR | Extracellular or secreted | -0.222702483 | 0.123361079 | 22.60575801 | 20.88889395 | 20.26843588 | 22.74810684 | 22.37398132 | 24.17203556 | 21.25839284 | 22.10065701 | 22.50144848 |
| Kelch-like ECH-associated protein 1 | Q14145 | KEAP1 | Other | -0.558871966 | 0.266270126 | 28.34201992 | 28.31328455 | 28.40600461 | 28.39042627 | 28.65632115 | 27.34578869 | 28.55435254 | 28.32504484 | 26.17090932 |
| KH domain-containing, RNA-binding, s | Q07666;Q07666-3;Q07666-2 | KHDRBS1 | Extracellular or secreted | 0.110374665 | 0.164414793 | 29.33496412 | 29.47018433 | 29.57394524 | 29.13141265 | 29.14379743 | 28.96402762 | 29.59217066 | 29.61852424 | 28.92959479 |
| KH domain-containing, RNA-binding, s | O75525-2;O75525 | KHDRBS3 | Other | 1.719663464 | 1.734348577 | 24.6290088 | 24.4804367 | 24.91205092 | 21.70444343 | 24.56372043 | 24.80970534 | 25.89194127 | 25.89099099 | 25.92574095 |
| Ketohexokinase | P50053-2 | KHK | Extracellular or secreted | -0.151480386 | 0.164528384 | 25.93064405 | 25.88154698 | 24.05523477 | 26.14829503 | 25.62218121 | 24.70144607 | 25.03896352 | 25.4189906 | 25.25727878 |
| Far upstream element-binding protein | Q92945 | KHSRP | Other | 0.039384615 | 0.075158389 | 30.54250112 | 30.55508431 | 30.50578364 | 30.80424345 | 30.92570983 | 31.2822868 | 30.96133891 | 30.93592951 | 30.52869 |
| Pumilio domain-containing protein KI | Q15397 | KIAA0020 | Other | 0.59935948 | 0.642144499 | 28.89154153 | 26.28077459 | 28.00179687 | 28.26843213 | 28.15325235 | 29.22492526 | 29.05188422 | 28.65441025 | 28.50214534 |
| Uncharacterized protein KIAA0040 | Q15053 | KIAA0040 | Other | 1.013326506 | 0.430606747 | 22.38115855 | 19.36397216 | 23.37710309 | 21.79581087 | 23.27227321 | 21.85212866 | 22.17630344 | 22.22764414 | 24.6572552 |
| WASH complex subunit strumpellin | Q12768 | KIAA0196 | Other | 0.635646075 | 0.715342744 | 27.06773944 | 26.99183217 | 27.04395595 | 25.37904859 | 25.71696218 | 26.32856715 | 27.4143362 | 27.30921293 | 26.44744184 |
| Dyslexia-associated protein KIAA0319 | Q8IZA0-2;Q8IZA0 | KIAA0319L | Extracellular or secreted | 0.854071523 | 0.653135962 | 23.01721008 | 21.067494 | 22.3620178 | 21.5308017 | 21.60943274 | 23.14424685 | 22.11463564 | 23.02582153 | 23.78735899 |
| WASH complex subunit 7 | Q2M389;Q2M389-2 | KIAA1033 | Other | 0.086000258 | 0.161314844 | 26.64320113 | 26.53669246 | 26.75327878 | 26.5135675 | 26.05647565 | 26.52548998 | 26.84911967 | 26.65799111 | 26.26524275 |
| Uncharacterized protein KIAA1143 | Q96AT1 | KIAA1143 | Other | 1.183033538 | 2.577183605 | 24.75338081 | 24.8534809 | 24.85790015 | 25.05237667 | 25.01469682 | 24.08480644 | 26.08198608 | 26.14612083 | 25.62931459 |
| Sickle tail protein homolog | Q5T5P2;Q5T5P2-9;Q5T5P2-2 | KIAA1217 | Other | -1.567498535 | 0.826143875 | 25.19377619 | 25.28210118 | 25.49600771 | 24.15120178 | 24.18895641 | 26.17790064 | 22.06323855 | 24.15452765 | 24.32471015 |
| KIF1-binding protein | Q96EK5 | KIAA1279 | Other | 0.332314122 | 1.518083869 | 26.18848487 | 26.04257901 | 26.17327531 | 26.5220658 | 26.74509326 | 26.5464845 | 26.64608886 | 26.68391576 | 26.77592911 |
| Protein virilizer homolog | Q69YN4-3;Q69YN4;Q69YN4-1 | KIAA1429 | Other | -0.936272364 | 1.609969899 | 27.5068809 | 27.88232857 | 27.71371499 | 26.11495583 | 26.49040252 | 27.58276381 | 26.16645305 | 26.27731976 | 26.39293342 |
| LisH domain and HEAT repeat-contain | Q9P260;Q9P260-2 | KIAA1468 | Other | -0.409636454 | 0.604669912 | 22.93662716 | 22.7476972 | 23.00170025 | 22.40101177 | 22.66072975 | 21.89696501 | 22.41833442 | 22.47869754 | 21.69642424 |
| Uncharacterized protein KIAA1522 | Q9P206;Q9P206-3;Q9P206-2 | KIAA1522 | Other | -0.390943259 | 0.532491461 | 23.59274809 | 23.79471411 | 23.86230591 | 23.9294029 | 23.93355091 | 21.88246471 | 23.22710059 | 22.9554094 | 23.14225355 |
| Shootin-1 | A0MZ66;A0MZ66-3;A0MZ66- | KIAA1598 | Other | 0.252575036 | 0.382361356 | 27.92914326 | 28.01850091 | 28.04082464 | 27.70325611 | 27.71102529 | 27.3058256 | 28.36869641 | 28.17512341 | 27.56819319 |
| Kinesin-like protein KIF11 | P52732 | KIF11 | Other | -0.286848761 | 0.80005669 | 25.14351525 | 24.73086314 | 24.79312699 | 25.35796752 | 24.72961908 | 25.41277062 | 24.81083323 | 24.50324538 | 24.90930642 |
| Kinesin-like protein KIF13B | Q9NQT8 | KIF13B | Other | 0.138694217 | 0.048319719 | 21.36749829 | 21.13365076 | 20.40621584 | 23.55051195 | 21.5313542 | 22.8304154 | 20.27787869 | 22.49541005 | 23.05261714 |
| Kinesin-like protein KIF15 | Q9NS87-3;Q9NS87-2;Q9NS87 | KIF15 | Other | 1.093046076 | 1.17915338 | 20.95859076 | 19.74535924 | 22.21746565 | 20.55962435 | 21.98290656 | 21.33265287 | 22.81846984 | 22.14650155 | 21.71246655 |
| Kinesin-like protein KIF1A | Q12756;Q12756-3;Q12756-2 | KIF1A | Other | 0.974751581 | 1.967135139 | 26.67640241 | 26.54427492 | 26.44415893 | 26.87337915 | 26.80833106 | 27.03758181 | 27.93168128 | 27.79494955 | 27.38968805 |
| Kinesin-like protein KIF21A | Q7Z4S6-6;Q7Z4S6-5;Q7Z4S6- | KIF21A | Other | 0.592586172 | 1.715361785 | 25.94360083 | 25.86924252 | 25.97208845 | 25.03401675 | 25.11509478 | 25.82679847 | 26.16392234 | 26.36340879 | 26.13084829 |
| Kinesin-like protein KIF23 | Q02241-2;Q02241;Q02241-3 | KIF23 | Extracellular or secreted | 0.077473681 | 0.089455459 | 26.25778231 | 25.87175882 | 26.37270342 | 26.02976913 | 26.24932486 | 27.509979 | 26.04403935 | 26.64705015 | 26.6869903 |
| Kinesin-like protein KIF2A | O00139-2;O00139-1;O00139 | KIF2A | Other | 0.609107803 | 0.492334551 | 24.55215498 | 24.14382661 | 24.29270526 | 24.04934671 | 21.96596038 | 26.01473938 | 24.44783591 | 24.75918438 | 25.12966978 |
| Kinesin-like protein KIF2C | Q99661;Q99661-2 | KIF2C | Other | -0.999450358 | 0.578551676 | 23.69890556 | 23.72108917 | 23.79223346 | 23.98346266 | 24.23586056 | 27.59313292 | 23.77494949 | 22.48561708 | 24.25342453 |
| Chromosome-associated kinesin KIF4/ | O95239;O95239-2;Q2VIQ3 | KIF4A;KIF4B | Other | 0.004778724 | 0.00307561 | 23.24978417 | 23.64019727 | 23.85661853 | 23.35501228 | 23.78256796 | 25.0453524 | 24.67427547 | 23.27668103 | 23.52814597 |
| Kinesin-1 heavy chain | P33176 | KIF5B | Extracellular or secreted | -0.524656649 | 3.827785607 | 31.32785224 | 31.20705293 | 31.12453941 | 31.35688939 | 31.34614811 | 31.00403618 | 30.72817606 | 30.75597048 | 30.62514264 |
| Kinesin-associated protein 3 | Q92845-4;Q92845-2;Q92845 | KIFAP3 | Extracellular or secreted | -1.1401228 | 0.600702051 | 25.64229236 | 23.0143563 | 25.62204154 | 21.13141028 | 21.12842498 | 21.91812513 | 22.28728965 | 22.08909741 | 21.43156984 |
| Kinesin-like protein KIFC1 | Q9BW19 | KIFC1 | Other | -1.178800564 | 1.053595015 | 28.26874444 | 28.16298178 | 28.19497863 | 27.63478004 | 27.86366518 | 28.28969226 | 26.76167175 | 26.27456769 | 27.63478004 |
| DNA/RNA-binding protein KIN17 | O60870-2;O60870 | KIN | Other | -1.36085821 | 2.283273158 | 23.53409671 | 23.51520799 | 23.33932919 | 22.47180273 | 23.28611941 | 23.24500309 | 21.58235785 | 22.28005596 | 21.75079113 |
| Kin of IRRE-like protein 1 | Q96J84;Q96J84-2;Q96J84-3 | KIRREL | Extracellular or secreted | 0.842557331 | 1.950179783 | 23.58497259 | 23.13089541 | 23.49240578 | 24.16507321 | 24.08099054 | 23.05953195 | 24.28736012 | 24.73096677 | 24.26627985 |
| Mast/stem cell growth factor recepto | P10721-2;P10721;P10721-3 | KIT | Extracellular or secreted | -0.355820958 | 0.408839375 | 26.06523484 | 26.29393325 | 26.33992379 | 26.20314365 | 25.79936622 | 25.6040222 | 25.94277346 | 26.0897844 | 25.05279124 |
| Kinesin light chain 1 | Q07866-8 | KLC1 | Other | -0.28281116 | 1.48966937 | 29.15197108 | 29.22347597 | 29.15385633 | 29.33947361 | 29.43603007 | 29.05765351 | 28.99372267 | 29.03849786 | 28.80057627 |
| Kinesin light chain 2 | Q9H0B6;Q9H0B6-2 | KLC2 | Other | 0.711887696 | 0.64772383 | 23.98893219 | 24.627785 | 23.84018936 | 25.542121 | 25.45782522 | 26.18576576 | 25.2850683 | 25.29691117 | 26.37499288 |
| Kinesin light chain 3 | Q6P597-2;Q6P597;Q6P597-3 | KLC3 | Other | -0.547641084 | 1.089456197 | 26.56531944 | 26.77643123 | 26.96179428 | 26.69816373 | 26.90942088 | 26.43572812 | 26.28824061 | 25.81056359 | 26.43170137 |
| Kallikrein-10 | O43240 | KLK10 | Extracellular or secreted | 0.368875639 | 0.220148412 | 27.62463729 | 27.46629943 | 27.53919479 | 28.53667764 | 28.72224288 | 30.70347065 | 28.22094198 | 28.50650263 | 29.67544365 |
| Kallikrein-6 | Q92876;Q92876-2 | KLK6 | Extracellular or secreted | 0.197948125 | 0.069930609 | 30.40496992 | 30.28427414 | 30.21777977 | 30.8359774 | 30.83575157 | 32.97026469 | 30.1279248 | 30.38030437 | 32.86012396 |
| Kallikrein-8 | O60259;O60259-2;O60259-3 | KLK8 | Extracellular or secreted | -0.700476225 | 0.263460596 | 26.9942084 | 27.2828612 | 27.37060986 | 26.7811926 | 26.88611388 | 29.04936749 | 25.62803541 | 25.86919545 | 28.58351718 |
| Lysine-rich nucleolar protein 1 | Q1ED39 | KNOP1 | Other | -3.098364282 | 1.046963712 | 23.65578226 | 24.41319043 | 24.21963177 | 23.76506591 | 24.66887676 | 25.10924722 | 19.69299121 | 20.73817795 | 23.18963517 |
| Kinetochore-associated protein 1 | P50748 | KNTC1 | Extracellular or secreted | -0.226358582 | 0.20137443 | 25.96088893 | 26.22377975 | 25.92492592 | 25.7893008 | 26.9234305 | 25.26724475 | 26.35184884 | 25.9134897 | 25.10037104 |
| Importin subunit alpha-5;Importin sub | P52294 | KPNA1 | Other | 0.162084143 | 1.02785377 | 27.69955434 | 27.50218329 | 27.50771275 | 27.36686378 | 27.58756431 | 27.7269589 | 27.69185606 | 27.84050154 | 27.64931353 |
| Importin subunit alpha-1 | P52292 | KPNA2 | Other | -0.081788326 | 0.129044985 | 28.90624127 | 28.91735359 | 28.83827882 | 29.82656817 | 29.85768663 | 29.67461895 | 29.40803127 | 29.3553148 | 29.00166267 |
| Importin subunit alpha-4 | O00505 | KPNA3 | Other | 0.036677828 | 0.035914911 | 27.78606299 | 27.84236118 | 27.80698005 | 27.0857791 | 27.28806456 | 27.33304545 | 27.96080057 | 27.7679977 | 26.95238187 |
| Importin subunit alpha-3 | O00629 | KPNA4 | Extracellular or secreted | -0.007903078 | 0.008586736 | 29.77591342 | 29.88978776 | 29.8455799 | 29.27719558 | 29.35142344 | 29.16492461 | 29.81476169 | 29.80358158 | 29.01035984 |
| Importin subunit alpha-7;Importin sub | O60684;O15131 | KPNA6;KPNA5 | Other | 0.224813037 | 0.448932644 | 24.64207197 | 24.73773793 | 24.53094588 | 24.52725124 | 24.31423582 | 25.32374859 | 25.07924159 | 25.06685717 | 24.56633607 |
| Importin subunit beta-1 | Q14974;Q14974-2 | KPNB1 | Extracellular or secreted | -0.176571873 | 0.636086661 | 32.64032916 | 32.71896104 | 32.67615017 | 32.30348131 | 32.26150502 | 32.43064209 | 32.47435863 | 32.37306035 | 32.13839979 |
| Kaptin | Q9Y664;Q9Y664-2 | KPTN | Other | 0.957329083 | 1.873898966 | 23.67055252 | 23.71889655 | 23.35930882 | 23.51183468 | 21.99577056 | 23.31299039 | 24.36091672 | 24.21467545 | 24.08107183 |
| GTPase KRas;GTPase KRas, N-termina | P01116-2;P01116 | KRAS | Extracellular or secreted | 0.586708186 | 0.957482721 | 26.76357245 | 26.39473325 | 26.63588725 | 26.79017128 | 27.37293585 | 26.91843454 | 27.39599179 | 26.99139971 | 27.81060037 |
| Protein KRI1 homolog | Q8N9T8 | KRI1 | Other | 0.103158076 | 0.10645639 | 23.64008691 | 24.04410191 | 23.59240594 | 23.64757173 | 23.77896423 | 23.82454238 | 23.60896405 | 24.50008717 | 23.46425956 |
| KRR1 small subunit processome comp | Q13601-2;Q13601 | KRR1 | Extracellular or secreted | 0.284987146 | 0.183321325 | 22.76251343 | 22.55166227 | 22.8707011 | 22.97508806 | 23.1013012 | 24.01928598 | 23.91442528 | 23.80774168 | 22.27307049 |
| Keratin, type I cytoskeletal 18 | P05783 | KRT18 | Extracellular or secreted | 0.518466279 | 0.586382527 | 31.61977516 | 31.64362262 | 31.63663932 | 30.61841922 | 30.57920687 | 32.2399076 | 31.31154252 | 32.07718234 | 32.33545937 |
| Keratinocyte-associated protein 3 | Q53RY4-2;Q53RY4 | KRTCAP3 | Other | 2.102240617 | 2.731746963 | 23.36198767 | 20.82313193 | 22.91000219 | 23.11432442 | 22.34078569 | 22.11793308 | 24.63727007 | 24.7020805 | 24.30145377 |
| Kinectin | Q86UP2;Q86UP2-4;Q86UP2- | KTN1 | Extracellular or secreted | -0.254733736 | 0.747009718 | 31.73658688 | 31.6169744 | 31.61175225 | 31.54116288 | 31.57474834 | 30.66957636 | 31.23968003 | 31.11228584 | 31.25923348 |
| Neural cell adhesion molecule L1 | P32004-3;P32004-2;P32004 | L1CAM | Extracellular or secreted | 0.044022024 | 0.045591245 | 27.56928024 | 27.74976726 | 27.64547055 | 28.39304802 | 28.33039775 | 27.86985436 | 27.66621046 | 27.69484734 | 28.54991736 |
| Beta-lactamase-like protein 2 | Q53H82 | LACTB2 | Other | 0.202169109 | 0.372905339 | 28.94363436 | 29.03010077 | 28.91607248 | 29.7532469 | 29.81392122 | 28.73521538 | 29.50455295 | 29.58098643 | 29.11706349 |
| Ladinin-1 | O00515 | LAD1 | Extracellular or secreted | 1.093216445 | 1.924198669 | 27.64072133 | 27.40795026 | 27.48285386 | 27.75008684 | 27.91735359 | 27.38418048 | 28.36101716 | 28.7376348 | 28.97257056 |
| EKC/KEOPS complex subunit LAGE3 | Q14657 | LAGE3 | Other | 0.295542516 | 0.428394305 | 24.85062256 | 24.77881388 | 24.80129195 | 25.08147824 | 24.3934245 | 24.89684494 | 24.80710292 | 25.61076714 | 24.86999552 |
| Laminin subunit alpha-1 | P25391 | LAMA1 | Extracellular or secreted | 2.827915116 | 0.582286191 | 22.71009257 | 22.80801676 | 23.91360383 | 24.22140303 | 24.22486547 | 30.76335083 | 25.64669318 | 25.99757163 | 31.16014679 |
| Laminin subunit alpha-3 | Q16787;Q16787-3 | LAMA3 | Extracellular or secreted | 0.206044547 | 0.192493129 | 28.92620486 | 28.83827882 | 28.88907695 | 29.53194612 | 29.70640504 | 30.62078035 | 29.27646363 | 29.35913284 | 30.23888324 |
| Laminin subunit alpha-5 | O15230 | LAMA5 | Extracellular or secreted | 0.804226691 | 1.289729955 | 29.28387689 | 29.35053853 | 29.17940117 | 29.25601917 | 29.31898281 | 31.15888367 | 30.23031852 | 30.37637641 | 30.57983626 |
| Laminin subunit beta-1 | P07942 | LAMB1 | Extracellular or secreted | 0.934943572 | 0.97744359 | 29.13525916 | 29.10462103 | 29.13021005 | 30.09509246 | 30.10811575 | 31.15249098 | 30.34952654 | 30.40059981 | 31.41759908 |
| Laminin subunit beta-2 | P55268 | LAMB2 | Extracellular or secreted | -0.308721055 | 0.244159421 | 27.53244784 | 27.8150672 | 27.51638078 | 27.24136321 | 27.25682904 | 29.7168085 | 27.13310442 | 27.25988446 | 28.22029625 |
| Laminin subunit beta-3 | Q13751 | LAMB3 | Extracellular or secreted | -1.205510469 | 0.394366212 | 26.51746251 | 27.07877349 | 27.10389606 | 27.96537697 | 27.50937502 | 29.54773142 | 25.23191198 | 25.33347204 | 28.67939231 |
| Laminin subunit gamma-1 | P11047 | LAMC1 | Extracellular or secreted | 0.737495758 | 0.899268524 | 29.25788569 | 29.17021215 | 29.05209159 | 29.85734552 | 29.83121249 | 31.20040294 | 30.13332494 | 30.27052774 | 30.99320978 |
| Laminin subunit gamma-2 | Q13753;Q13753-2 | LAMC2 | Extracellular or secreted | 0.073897415 | 0.018557192 | 26.51279929 | 26.77178 | 26.55382543 | 26.59420134 | 26.54955755 | 29.59981537 | 26.23297331 | 25.75236023 | 29.5273482 |
| Lysosome-associated membrane glyco | P11279;P11279-2 | LAMP1 | Extracellular or secreted | 0.350393466 | 1.040968689 | 26.72103701 | 26.96256669 | 27.02569905 | 27.12760506 | 27.03559065 | 27.2337048 | 27.28700777 | 27.19591733 | 27.62135693 |
| Lysosome-associated membrane glyco | P13473-2;P13473;P13473-3 | LAMP2 | Extracellular or secreted | -0.219333983 | 0.595169003 | 26.80783993 | 26.9647713 | 26.87420036 | 26.83773764 | 27.28586204 | 26.30175029 | 26.74419514 | 26.39290067 | 26.74098301 |
| Ragulator complex protein LAMTOR1 | Q6IAA8 | LAMTOR1 | Extracellular or secreted | 0.916741843 | 1.878982627 | 25.6978987 | 25.30291833 | 25.10629456 | 24.4221387 | 24.25154771 | 25.30344103 | 25.83470327 | 25.71999328 | 26.23764849 |
| Ragulator complex protein LAMTOR2 | Q9Y2Q5;Q9Y2Q5-2 | LAMTOR2 | Extracellular or secreted | 0.115036774 | 0.152666182 | 25.58600411 | 25.13041628 | 25.62861953 | 24.22971364 | 24.56755507 | 25.04859862 | 25.02605802 | 25.50282064 | 24.91168529 |
| Ragulator complex protein LAMTOR3 | Q9UHA4;Q9UHA4-2 | LAMTOR3 | Extracellular or secreted | -0.095907749 | 0.17494697 | 27.36327509 | 26.9647713 | 26.77906445 | 26.30234315 | 26.2385234 | 26.65908066 | 26.36986142 | 26.70938277 | 26.78656159 |
| LanC-like protein 1 | O43813 | LANCL1 | Extracellular or secreted | -1.839796585 | 0.530828754 | 26.59562468 | 26.94547741 | 27.02432571 | 26.18584136 | 26.00298803 | 26.58175869 | 25.96192672 | 26.08921867 | 22.0974728 |
| LanC-like protein 2 | Q9NS86 | LANCL2 | Extracellular or secreted | -0.309202348 | 1.342288084 | 22.54271734 | 22.30824792 | 22.41302252 | 22.3778248 | 21.97868812 | 22.71885475 | 22.14954371 | 21.92918616 | 22.16334081 |
| Cytosol aminopeptidase | P28838-2;P28838 | LAP3 | Extracellular or secreted | -0.179665227 | 0.116849775 | 31.08036289 | 30.99408698 | 30.98046408 | 31.62575231 | 31.72995936 | 30.0510285 | 31.42337157 | 31.35005371 | 29.91840611 |
| La-related protein 1 | Q6PKG0;Q6PKG0-3 | LARP1 | Other | 0.515459188 | 1.302286103 | 27.45935832 | 27.88797377 | 27.90987862 | 28.21974254 | 28.24049912 | 28.00651279 | 28.37563096 | 28.26432125 | 28.76840793 |
| La-related protein 4B | Q92615 | LARP4B | Other | -0.535455228 | 0.140954015 | 23.5597589 | 23.46188906 | 23.89078231 | 21.67931228 | 21.21480039 | 23.62142681 | 23.50251717 | 23.76951181 | 19.8355902 |
| La-related protein 7 | Q4G0J3;Q4G0J3-3;Q4G0J3-2 | LARP7 | Other | -0.180063267 | 0.340153177 | 25.82645907 | 25.6773706 | 25.62463729 | 25.56150798 | 26.03019025 | 25.81166632 | 25.67581041 | 25.849239 | 25.20067654 |
| Leucine--tRNA ligase, cytoplasmic | Q9P2J5;Q9P2J5-2 | LARS | Other | -0.173799003 | 0.494460629 | 28.77464165 | 28.83294838 | 28.80983411 | 29.14996228 | 29.40515257 | 28.86002773 | 28.68922526 | 28.66891733 | 29.03674376 |
| Ribosomal biogenesis protein LAS1L | Q9Y4W2-3;Q9Y4W2-2;Q9Y4V | LAS1L | Other | 0.421053404 | 0.284659346 | 20.75245282 | 23.10298359 | 22.79857543 | 23.01140745 | 23.37398132 | 25.38376862 | 23.6676322 | 23.35877245 | 23.44834018 |
| LIM and SH3 domain protein 1 | Q14847;Q14847-2;Q14847-3 | LASP1 | Extracellular or secreted | 0.477962853 | 0.643633225 | 29.44740636 | 29.30214701 | 29.33877259 | 29.30046784 | 28.89440675 | 28.86455098 | 29.92309042 | 29.98155322 | 29.10312069 |
| Leucine carboxyl methyltransferase 1 | Q9UIC8;Q9UIC8-2;Q9UIC8-3 | LCMT1 | Other | 0.153962815 | 0.248265107 | 26.40631285 | 26.26606534 | 26.21574953 | 26.73272723 | 26.48268464 | 25.68482544 | 26.49419261 | 26.77555241 | 26.08632593 |
| Neutrophil gelatinase-associated lipo | P80188;P80188-2 | LCN2 | Extracellular or secreted | 1.706518827 | 1.59975015 | 31.58131874 | 31.8239596 | 31.73008087 | 33.39368737 | 33.52880641 | 32.81090292 | 33.78993817 | 33.76327167 | 35.0007246 |
| Plastin-2 | P13796 | LCP1 | Extracellular or secreted | 1.44273071 | 2.308233459 | 30.41790082 | 30.40547722 | 30.36045212 | 29.74115987 | 29.75160389 | 29.41013597 | 31.27469875 | 31.19601117 | 31.90084715 |
| L-lactate dehydrogenase A chain | P00338;P00338-3;P00338-5;P | LDHA | Extracellular or secreted | 0.100377093 | 0.316835852 | 34.55693496 | 34.68206621 | 34.73670783 | 34.74900197 | 34.72907812 | 34.07295729 | 34.62744932 | 34.58592489 | 34.85113026 |
| L-lactate dehydrogenase B chain | P07195 | LDHB | Extracellular or secreted | 0.178038281 | 1.093717022 | 34.45157331 | 34.53357143 | 34.43122763 | 34.69078059 | 34.52653459 | 34.13934553 | 34.57807103 | 34.61434375 | 34.72821661 |
| Low-density lipoprotein receptor | P01130-3;P01130-4;P01130-5 | LDLR | Extracellular or secreted | 2.16034288 | 1.429770445 | 27.93049742 | 27.82462736 | 27.73576414 | 27.85910952 | 27.97983746 | 27.99058849 | 29.65165889 | 29.57437843 | 30.91520352 |
| LEM domain-containing protein 2 | Q8NC56 | LEMD2 | Other | -1.121639636 | 1.62963199 | 24.36646363 | 23.66698243 | 23.63045378 | 22.29608497 | 22.79580425 | 22.70634271 | 22.26399008 | 22.42149679 | 21.6806601 |
| Inner nuclear membrane protein Man | Q9Y2U8 | LEMD3 | Other | -2.246995272 | 1.319724791 | 27.75021465 | 28.94310334 | 27.72643927 | 27.55924836 | 28.46384656 | 27.22820951 | 24.77821231 | 25.47367182 | 26.8426609 |
| RNA polymerase-associated protein L | Q8WVC0;Q8WVC0-2 | LEO1 | Other | -0.215780769 | 0.432820127 | 27.30929968 | 27.20060172 | 27.20508446 | 26.96774218 | 27.01414344 | 26.93010259 | 27.14111941 | 27.00131369 | 26.52371162 |
| Prolyl 3-hydroxylase 1 | Q32P28-4;Q32P28;Q32P28-3 | LEPRE1 | Extracellular or secreted | 0.339213453 | 0.450103557 | 24.85794759 | 24.82352235 | 25.23235124 | 25.1882396 | 24.93629466 | 26.35203413 | 25.82871217 | 25.08958238 | 25.79454061 |
| LETM1 and EF-hand domain-containin | O95202 | LETM1 | Other | -0.305131028 | 0.239821129 | 23.40807987 | 22.80463366 | 24.04776695 | 23.88604408 | 23.98693778 | 26.36639693 | 23.56755507 | 23.72463209 | 24.04234939 |
| Galectin-1 | P09382 | LGALS1 | Extracellular or secreted | 0.440821359 | 1.081176379 | 31.7819117 | 31.6114005 | 31.70285169 | 31.47460039 | 31.59294586 | 31.72285358 | 31.85318639 | 32.05656222 | 32.35599733 |
| Galectin-3 | P17931 | LGALS3 | Extracellular or secreted | -0.021635979 | 0.018497039 | 30.38987264 | 30.46635778 | 30.47846303 | 30.11073124 | 30.139559 | 29.40375169 | 30.42382236 | 30.54498972 | 29.46064768 |
| Galectin-3-binding protein | Q08380 | LGALS3BP | Extracellular or secreted | -0.214134192 | 0.191477597 | 30.17092842 | 30.25225632 | 30.15617325 | 31.17753715 | 31.32860139 | 31.28051798 | 30.10412109 | 30.2178953 | 31.21858829 |
| Galectin-7 | P47929 | LGALS7 | Extracellular or secreted | 0.768312953 | 0.6175553 | 31.71124689 | 31.81478461 | 31.88551035 | 30.1016188 | 30.28040736 | 32.45432172 | 31.79048204 | 31.63741691 | 33.00098477 |
| Galectin-related protein | Q3ZCW2 | LGALSL | Other | 1.057181023 | 3.418745804 | 26.27175694 | 26.07381861 | 26.15865499 | 25.99498522 | 25.53895807 | 25.65415566 | 26.85174275 | 26.96862127 | 27.19732425 |
| Leucine-rich repeat LGI family membe | Q8N145-2;Q8N145 | LGI3 | Extracellular or secreted | -1.250626976 | 0.810961818 | 19.13771164 | 21.00254451 | 22.83418646 | 22.42036644 | 19.86900336 | 21.65416574 | 20.76656594 | 19.02258455 | 19.91795766 |
| Legumain | Q99538;Q99538-2;Q99538-3 | LGMN | Extracellular or secreted | 1.366141019 | 1.369737135 | 27.44110607 | 27.43332616 | 27.48968311 | 27.92569568 | 27.91940102 | 27.80119326 | 28.76350913 | 28.65615063 | 29.68396594 |
| Phospholysine phosphohistidine inorg | Q9H008;Q9H008-2 | LHPP | Other | -0.304461155 | 0.607121656 | 29.11145279 | 29.05115819 | 29.00109889 | 28.3273169 | 28.5393797 | 28.44679532 | 28.3032581 | 28.22237078 | 28.79958855 |
| Leukemia inhibitory factor | P15018 | LIF | Extracellular or secreted | 2.694373515 | 1.03426467 | 20.49031085 | 22.42704453 | 21.25269265 | 21.49954483 | 21.45214491 | 21.06722774 | 24.5785952 | 22.24598057 | 25.35302753 |
| DNA ligase 1 | P18858-3;P18858;P18858-2 | LIG1 | Other | -0.207286723 | 0.247600182 | 27.44837169 | 27.55405972 | 27.6125258 | 27.45873276 | 27.23708316 | 26.95415832 | 27.54008215 | 27.38919569 | 26.58132772 |
| LIM domain and actin-binding protein | Q9UHB6-4;Q9UHB6 | LIMA1 | Extracellular or secreted | 0.7889946 | 0.755182979 | 27.75193903 | 27.74033973 | 27.77373031 | 27.73963178 | 27.65601421 | 28.02115146 | 28.25601917 | 28.11495583 | 29.33741206 |
| LIM and senescent cell antigen-like-co | P48059-2;P48059;P48059-4; | LIMS1 | Extracellular or secreted | 0.749859886 | 1.606759313 | 24.46276286 | 24.94561136 | 24.52838448 | 24.14312595 | 24.0988462 | 25.17047005 | 25.21297009 | 25.64809312 | 25.0631169 |
| Protein lin-7 homolog C | Q9NUP9 | LIN7C | Extracellular or secreted | 0.037415437 | 0.222853178 | 28.32122142 | 28.3671555 | 28.34823945 | 28.24717103 | 28.37546526 | 28.63968861 | 28.37592091 | 28.50062668 | 28.38516845 |
| Leucine-rich repeat and immunoglobu | Q96FE5-2;Q96FE5 | LINGO1 | Extracellular or secreted | 1.708612189 | 1.739255283 | 21.58494386 | 22.14736247 | 21.45023951 | 21.10862327 | 20.45618102 | 23.24723623 | 22.80581471 | 23.26713757 | 24.05017747 |
| Lysosomal acid lipase/cholesteryl este | P38571-2;P38571 | LIPA | Extracellular or secreted | 0.424227015 | 0.182196376 | 23.62566872 | 23.46704312 | 22.93738172 | 22.27181787 | 21.34868037 | 22.09468873 | 24.20045205 | 23.39100019 | 21.55386908 |
| Lethal(2) giant larvae protein homolo | Q15334 | LLGL1 | Extracellular or secreted | 0.338501914 | 0.593268369 | 25.91428841 | 26.15261428 | 26.11838603 | 25.70028223 | 25.95562226 | 26.21713724 | 26.56248362 | 26.56906289 | 25.91312444 |
| Lethal(2) giant larvae protein homolo | Q6P1M3-2;Q6P1M3 | LLGL2 | Extracellular or secreted | 0.48264727 | 1.315055609 | 26.61315839 | 26.44412733 | 26.49686109 | 26.04820364 | 26.64487947 | 26.5145159 | 26.53775907 | 26.81531156 | 26.97574408 |
| Protein LLP homolog | Q9BRT6 | LLPH | Other | 0.911470457 | 0.271953826 | 19.71802215 | 21.12016694 | 22.41665737 | 23.33183324 | 22.57528625 | 24.28291421 | 25.306008 | 21.40379637 | 22.74644707 |
| Protein ERGIC-53 | P49257 | LMAN1 | Extracellular or secreted | 0.239012737 | 0.143297185 | 26.28848706 | 26.38715882 | 26.39304802 | 26.34209624 | 25.90341098 | 28.20452488 | 26.15645267 | 26.49932695 | 27.84062159 |
| Vesicular integral-membrane protein | Q12907 | LMAN2 | Extracellular or secreted | 0.763674856 | 0.812890157 | 28.73366483 | 28.63488387 | 28.70747509 | 29.73681265 | 30.0292057 | 30.92684108 | 29.84072662 | 30.04869216 | 30.7860474 |
| VIP36-like protein | Q9H0V9;Q9H0V9-2;Q9H0V9- | LMAN2L | Other | 0.719463765 | 1.001844667 | 20.93341367 | 21.1378502 | 21.61481587 | 22.74398465 | 21.89610657 | 23.06606882 | 22.54096896 | 22.49047291 | 22.87306931 |
| Probable lysosomal cobalamin transp | Q9NUN5-4;Q9NUN5-2;Q9NU | LMBRD1 | Extracellular or secreted | 0.387183306 | 0.316207827 | 24.59046553 | 24.70935647 | 24.51827327 | 25.14755772 | 25.29715613 | 22.01783689 | 24.45963973 | 25.04776695 | 24.79446624 |
| LIM and cysteine-rich domains protei | Q9NZU5;Q9NZU5-2 | LMCD1 | Extracellular or secreted | 1.016975392 | 0.94166253 | 25.407302 | 24.95863413 | 24.38738893 | 24.93674396 | 24.83470327 | 25.02690229 | 25.412383 | 25.68528006 | 26.72910041 |

| | | | | | | | | | | | | | | |
|---|---|---|---|---|---|---|---|---|---|---|---|---|---|---|
| Prelamin-A/C;Lamin-A/C | P02545;P02545-6;P02545-3;F | LMNA | Other | 0.084076594 | 0.213500285 | 34.51523902 | 34.4697575 | 34.45910666 | 33.84931358 | 33.86931019 | 33.98320237 | 34.23948087 | 34.14338083 | 34.44233274 |
| Prelamin-A/C;Lamin-A/C | P02545-2 | LMNA | Other | -0.343855219 | 0.864921428 | 30.40527432 | 30.23876938 | 30.29219622 | 29.86381285 | 30.00597767 | 30.35405388 | 29.64046277 | 29.75957808 | 30.14843566 |
| Lamin-B1 | P20700 | LMNB1 | Other | -0.408502537 | 0.701585103 | 31.52565986 | 31.54999996 | 31.52864343 | 30.79660584 | 30.93072299 | 30.80616576 | 30.56092519 | 30.5666518 | 31.21581432 |
| Lamin-B2 | Q03252 | LMNB2 | Other | -0.855921139 | 1.431189452 | 30.354159 | 30.15882349 | 30.1207754 | 30.1316089 | 29.91242218 | 30.25754855 | 29.0019579 | 29.23594271 | 29.66200474 |
| LIM domain only protein 7 | Q8WWI1-3;Q8WWI1-5;Q8W\ | LMO7 | Other | 0.744651459 | 0.700593689 | 26.77240942 | 26.57629012 | 26.72819228 | 27.14782941 | 26.84026141 | 25.92881581 | 26.9097642 | 27.10439608 | 28.21669332 |
| Leucyl-cystinyl aminopeptidase;Leucy | Q9UIQ6-3;Q9UIQ6-2;Q9UIQ6 | LNPEP | Extracellular or secreted | 0.884208664 | 0.668276328 | 24.56952652 | 24.34719204 | 24.76905774 | 22.54752625 | 21.90943331 | 25.89219611 | 24.61588239 | 24.81772848 | 25.23648112 |
| Lon protease homolog, mitochondrial | P36776-2;P36776;P36776-3 | LONP1 | Other | -0.297626846 | 0.48371058 | 26.85447998 | 26.46024927 | 27.01531381 | 27.17645563 | 27.70898828 | 28.20307829 | 26.78656159 | 26.85519318 | 27.17464732 |
| Lysyl oxidase homolog 2 | Q9Y4K0 | LOXL2 | Extracellular or secreted | -0.489786807 | 0.740570827 | 27.78238049 | 27.56499978 | 27.56572618 | 28.94773614 | 28.93685626 | 29.12679312 | 27.64464572 | 27.86938374 | 27.9788561 |
| Lysophosphatidic acid receptor 2 | Q9HBW0 | LPAR2 | Extracellular or secreted | 1.009699755 | 1.321406587 | 22.20886899 | 22.44476768 | 20.57357693 | 22.66993537 | 22.3332647 | 22.43303319 | 22.58792764 | 23.31851743 | 23.45437762 |
| Latrophilin-1 | O94910-2;O94910 | LPHN1 | Extracellular or secreted | -0.245619495 | 0.247473917 | 23.75633642 | 23.32044691 | 23.34948858 | 22.02485514 | 22.88688137 | 23.43758649 | 22.42956407 | 22.74482645 | 23.47654845 |
| Latrophilin-3 | Q9HAR2-3;Q9HAR2-2;Q9HAR | LPHN3 | Extracellular or secreted | 0.263537916 | 0.138129484 | 21.32352108 | 21.18494468 | 22.30055355 | 23.83188127 | 22.09459584 | 22.97381016 | 23.13109958 | 23.11684841 | 21.39731905 |
| Lipoma-preferred partner | Q93052 | LPP | Extracellular or secreted | 0.183257811 | 0.438500579 | 28.2998131 | 28.2634708 | 28.20233109 | 28.43567249 | 28.43241045 | 28.43276884 | 28.61558075 | 28.74528564 | 28.22214042 |
| Lipopolysaccharide-responsive and be | P50851;P50851-2 | LRBA | Extracellular or secreted | -0.18158647 | 0.741091154 | 27.78830533 | 27.6602374 | 27.57094546 | 27.46154564 | 27.70385015 | 27.70457587 | 27.64553926 | 27.32628848 | 27.42814277 |
| Leucine-rich alpha-2-glycoprotein | P02750 | LRG1 | Extracellular or secreted | 0.020415555 | 0.011158847 | 28.64378602 | 28.27466543 | 28.34557722 | 29.28396517 | 29.44764283 | 29.56281839 | 28.274621 | 28.55266804 | 30.01318515 |
| Prolow-density lipoprotein receptor-r | Q07954 | LRP1 | Extracellular or secreted | -3.729752323 | 0.968869475 | 28.42802293 | 28.33266141 | 28.99816907 | 28.08719639 | 27.84888097 | 28.82662878 | 26.67060655 | 25.29056255 | 22.11035371 |
| Low-density lipoprotein receptor-rela | O75581 | LRP6 | Extracellular or secreted | 1.044925805 | 0.787261819 | 21.97655123 | 24.1766078 | 22.23823757 | 20.19912327 | 23.93201934 | 23.93553054 | 23.70988229 | 23.94699477 | 23.70693523 |
| Low-density lipoprotein receptor-rela | Q14114-2;Q14114-3;Q14114 | LRP8 | Extracellular or secreted | 0.726146378 | 0.759529488 | 22.6925451 | 22.61508538 | 22.3772993 | 22.93310475 | 22.21555668 | 23.31658538 | 22.99989543 | 23.10460003 | 24.14903196 |
| Alpha-2-macroglobulin receptor-asso | P30533 | LRPAP1 | Extracellular or secreted | -0.429724931 | 1.100860813 | 26.0013996 | 25.85317152 | 25.32199551 | 26.04090823 | 26.17274139 | 26.33682919 | 25.26277217 | 25.55970056 | 25.7518752 |
| Leucine-rich PPR motif-containing pro | P42704 | LRPPRC | Other | -0.420052824 | 0.355742178 | 29.22570681 | 29.46964107 | 29.63611205 | 30.62296315 | 30.70074523 | 31.97336459 | 29.58362477 | 29.53359928 | 30.43688393 |
| Leucine-rich repeat-containing protei | Q9BTT6;Q9BTT6-2 | LRRC1 | Other | 0.633788996 | 2.408523064 | 22.38490177 | 22.97839137 | 23.07255985 | 22.58047113 | 22.59443487 | 22.37822182 | 23.46301242 | 23.26685172 | 23.16599325 |
| Leucine-rich repeat-containing protei | Q5VZK9-2;Q5VZK9 | LRRC16A | Extracellular or secreted | 0.230980497 | 0.348196983 | 27.67276587 | 27.8469102 | 27.7549995 | 27.28550932 | 27.12494608 | 27.55501115 | 27.93724925 | 28.07612485 | 27.29963845 |
| Leucine-rich repeat-containing protei | Q8ND23;Q8ND23-2 | LRRC16B | Other | -0.125416096 | 0.07285475 | 22.12067801 | 24.54786745 | 24.10501586 | 24.67750502 | 24.96829168 | 23.69731546 | 23.55683906 | 23.34678639 | 24.778463 |
| Leucine-rich repeat-containing protei | Q9H9A6 | LRRC40 | Other | -0.758538298 | 1.178961328 | 26.63256306 | 26.200901 | 26.24216318 | 25.72210605 | 25.77921478 | 25.41464266 | 25.51126163 | 25.46587923 | 24.7430396 |
| Leucine-rich repeat-containing protei | Q8N1G4 | LRRC47 | Other | -0.594696829 | 0.843627386 | 27.87367249 | 27.95132607 | 27.88535761 | 27.65901258 | 27.72090658 | 27.04416446 | 27.42958003 | 27.29649113 | 26.55705825 |
| Leucine-rich repeat-containing protei | Q9NT99 | LRRC4B | Extracellular or secreted | -1.399491695 | 0.672745726 | 21.7980414 | 21.86936491 | 22.17581632 | 22.81762435 | 21.92326868 | 22.27727718 | 20.65378259 | 19.43489044 | 22.1435483 |
| Leucine-rich repeat-containing protei | Q8N9N7 | LRRC57 | Extracellular or secreted | 0.2631366 | 0.449876276 | 25.05266688 | 24.48732338 | 25.01920113 | 24.2505361 | 24.47611578 | 24.33939715 | 24.90476654 | 25.20705991 | 24.49020357 |
| Leucine-rich repeat-containing protei | Q96AG4 | LRRC59 | Other | -0.966970498 | 1.943858965 | 28.62292825 | 28.73094734 | 28.77853193 | 27.60413535 | 27.64278813 | 28.7559864 | 27.07439041 | 27.48123774 | 27.61111904 |
| Volume-regulated anion channel subu | Q8IWT6 | LRRC8A | Extracellular or secreted | 0.939045724 | 0.891326061 | 22.44748283 | 21.81711901 | 21.47202237 | 22.77885321 | 21.92199859 | 22.00716965 | 22.45686072 | 23.79887202 | 22.78372725 |
| Leucine-rich repeat flightless-interact | Q32MZ4-3;Q32MZ4-2;Q32M | LRRFIP1 | Extracellular or secreted | 0.175046685 | 0.431501775 | 31.27258695 | 31.29279959 | 31.29712552 | 31.09748051 | 31.05251919 | 30.56219974 | 31.3589338 | 31.44825352 | 31.00530848 |
| Leucine-rich repeat flightless-interact | Q32MZ4-4 | LRRFIP1 | Extracellular or secreted | -0.169452936 | 0.50368563 | 23.97718627 | 24.65567309 | 24.55015963 | 24.27951763 | 24.18842827 | 23.68816492 | 24.13194737 | 24.05837627 | 23.97088246 |
| Leucine-rich repeat flightless-interact | Q9Y608-4;Q9Y608;Q9Y608-2 | LRRFIP2 | Other | 0.42827108 | 1.001399708 | 27.20825133 | 27.30164564 | 27.59534013 | 27.15102754 | 26.98413662 | 26.40057538 | 27.34984286 | 27.42598418 | 27.82947451 |
| Leucine-rich repeat serine/threonine- | Q38SD2 | LRRK1 | Other | 0.523644231 | 0.113984223 | 26.39213095 | 22.94590879 | 21.48262522 | 25.73143298 | 26.42104728 | 25.72907447 | 27.11326748 | 22.50791451 | 26.30086055 |
| E3 ubiquitin-protein ligase LRSAM1 | Q6UWE0-2;Q6UWE0 | LRSAM1 | Extracellular or secreted | 1.292584994 | 1.885208699 | 24.62839703 | 24.22501263 | 24.33326729 | 23.7696127 | 24.3073726 | 22.34029475 | 25.29792576 | 25.52220053 | 24.8596072 |
| Leucine-rich repeat and WD repeat-co | Q9UFC0 | LRWD1 | Other | -0.013948643 | 0.018135504 | 23.97875793 | 24.83585997 | 24.26713757 | 23.15616457 | 24.3959101 | 24.27951763 | 24.39165581 | 23.98936539 | 24.03380676 |
| U6 snRNA-associated Sm-like protein | O15116 | LSM1 | Other | 0.535863493 | 0.836362938 | 24.28827582 | 24.06644662 | 24.17637953 | 23.92270489 | 23.98927876 | 23.9685993 | 24.70988229 | 24.95242631 | 24.15112434 |
| Protein LSM12 homolog | Q3MHD2;Q3MHD2-2 | LSM12 | Other | 2.56720211 | 2.617532705 | 22.7858236 | 23.3937518 | 23.12227824 | 21.04693128 | 20.86265606 | 20.91252015 | 24.29635109 | 24.4429101 | 25.02432571 |
| Protein LSM14 homolog A | Q8ND56-3;Q8ND56-2;Q8ND5 | LSM14A | Other | -0.636385994 | 0.96214267 | 26.01301472 | 26.30699035 | 26.04809968 | 26.75824416 | 26.89504269 | 26.49041782 | 25.32628848 | 25.81051456 | 26.20994368 |
| Protein LSM14 homolog B | Q9BX40-2;Q9BX40 | LSM14B | Other | -0.182428361 | 0.44348425 | 24.97548171 | 24.40710747 | 24.89254356 | 24.43637961 | 23.9942084 | 24.22354042 | 24.25277514 | 24.16023336 | 24.504337 |
| U6 snRNA-associated Sm-like protein | Q9Y333 | LSM2 | Other | 0.894485326 | 2.490156774 | 27.31717217 | 27.65423951 | 27.25205325 | 26.75110903 | 26.77517562 | 27.31267886 | 28.16739135 | 28.23086818 | 27.81641066 |
| U6 snRNA-associated Sm-like protein | P62310 | LSM3 | Other | 2.476994124 | 1.275306116 | 19.13836104 | 24.89161685 | 22.15179954 | 24.88152364 | 25.33429075 | 24.21963177 | 26.33924423 | 26.1635769 | 25.23677305 |
| U6 snRNA-associated Sm-like protein | Q9Y4Z0 | LSM4 | Other | 0.019048249 | 0.02054439 | 28.11152741 | 28.08618418 | 27.99652685 | 26.86147789 | 26.98793533 | 27.28727204 | 27.84481704 | 27.71855691 | 27.15923264 |
| U6 snRNA-associated Sm-like protein | Q9Y4Y9;Q9Y4Y9-2 | LSM5 | Other | 1.837602833 | 1.080679861 | 25.06533757 | 25.11557109 | 21.47861425 | 24.59565313 | 24.61042923 | 20.54546433 | 25.84388311 | 25.68597509 | 24.6884851 |
| U6 snRNA-associated Sm-like protein | P62312 | LSM6 | Extracellular or secreted | 0.392277261 | 0.784687768 | 27.16115649 | 27.13232008 | 27.06630291 | 26.16257849 | 26.06972771 | 26.93943681 | 27.17340875 | 27.38796406 | 26.88122022 |
| U6 snRNA-associated Sm-like protein | Q9UK45 | LSM7 | Other | 0.246265917 | 0.395886643 | 26.86502318 | 26.6443019 | 26.55755497 | 26.11713783 | 26.6389277 | 26.89087506 | 26.41230223 | 27.08101086 | 27.10239497 |
| U6 snRNA-associated Sm-like protein | O95777 | LSM8 | Other | 0.644550843 | 0.818633954 | 28.06476217 | 28.14734422 | 28.18872066 | 28.03454157 | 27.84320024 | 26.68284482 | 28.83656438 | 28.68755771 | 27.89023728 |
| Lipolysis-stimulated lipoprotein recep | Q86X29-6;Q86X29-5;Q86X29 | LSR | Extracellular or secreted | 1.215113526 | 2.664786022 | 27.50415512 | 27.49990475 | 27.77918973 | 28.45018239 | 28.39145096 | 28.66922153 | 29.03676996 | 29.21842662 | 29.53719624 |
| Lanosterol synthase | P48449;P48449-2;P48449-3 | LSS | Other | 3.720145253 | 1.339099027 | 22.26327379 | 20.91473549 | 21.45858885 | 22.23221651 | 23.48162167 | 24.43370819 | 25.19027592 | 25.1279789 | 28.23425318 |
| Leukotriene A-4 hydrolase | P09960;P09960-4;P09960-2;F | LTA4H | Extracellular or secreted | -0.029885006 | 0.130073167 | 33.34572525 | 33.45928991 | 33.37852261 | 33.33989831 | 33.38676734 | 32.93119869 | 33.35389619 | 33.21971369 | 33.25743615 |
| Latent-transforming growth factor be | Q14766-3;Q14766-2;Q14766 | LTBP1 | Extracellular or secreted | -1.455556242 | 0.532572685 | 21.24631839 | 20.44960006 | 20.47803093 | 20.86476349 | 23.36212148 | 27.59398772 | 19.93890762 | 20.76891208 | 21.92292261 |
| Latent-transforming growth factor be | Q14767 | LTBP2 | Extracellular or secreted | 0.31531727 | 0.05939741 | 22.15192948 | 22.26330244 | 22.65530186 | 24.11806913 | 23.63444771 | 28.81280492 | 22.66689577 | 22.79717071 | 27.2998131 |
| Latent-transforming growth factor be | Q9NS15-2;Q9NS15 | LTBP3 | Extracellular or secreted | 1.103368366 | 0.197717928 | 21.82010727 | 21.66916881 | 21.24986303 | 25.55923377 | 24.91054209 | 27.97716443 | 22.93189314 | 23.38218599 | 28.58906567 |
| Latent-transforming growth factor be | Q8N2S1-2;Q8N2S1-3;Q8N2S1 | LTBP4 | Extracellular or secreted | -1.936576274 | 0.432737332 | 28.91484723 | 26.82347376 | 25.49905317 | 27.88983127 | 28.36952866 | 30.95650044 | 24.69136357 | 24.27916336 | 29.44636151 |
| E3 ubiquitin-protein ligase listerin | O94822;O94822-3 | LTN1 | Other | 0.994077922 | 1.175537267 | 22.25300606 | 21.47679564 | 22.95283507 | 22.8286347 | 20.84760326 | 22.76455982 | 23.36599664 | 23.56313854 | 22.61481587 |
| Protein LTV1 homolog | Q96GA3 | LTV1 | Other | 2.311680335 | 2.746585027 | 21.36484891 | 21.62258107 | 21.03315515 | 22.41318183 | 21.36118846 | 22.96574906 | 23.69019155 | 23.9421694 | 24.68303229 |
| Putative RNA-binding protein Luc7-lik | Q9NQ29-2;Q9NQ29;Q9NQ29 | LUC7L | Other | -1.170953436 | 3.813744838 | 26.29502003 | 25.85095633 | 26.34764826 | 25.64325619 | 26.10866913 | 26.55367897 | 25.02047338 | 25.05407564 | 24.81220512 |
| Putative RNA-binding protein Luc7-lik | Q9Y383 | LUC7L2 | Other | -0.917914935 | 3.929110796 | 30.06650821 | 29.89937121 | 29.84714922 | 29.73259785 | 29.93550806 | 30.38966754 | 29.0431216 | 29.03241485 | 29.1061198 |
| Luc7-like protein 3 | O95232 | LUC7L3 | Other | -0.750090671 | 2.350345896 | 28.37260379 | 28.22527011 | 28.28224261 | 28.306608 | 28.33509195 | 28.87147096 | 27.53815885 | 27.52962107 | 27.87859177 |
| Leucine zipper protein 1 | Q86V48-2;Q86V48-3;Q86V48 | LUZP1 | Extracellular or secreted | -2.708814503 | 0.857375301 | 24.21148553 | 23.65261306 | 23.58508724 | 24.05977947 | 23.52468327 | 23.39296616 | 20.05029039 | 19.73498014 | 23.30159332 |
| Latexin | Q9BS40 | LXN | Extracellular or secreted | -2.540095228 | 1.06606592 | 24.65250365 | 24.40795026 | 24.05647565 | 23.47444567 | 23.57207382 | 23.71893718 | 23.0623018 | 20.96669531 | 20.29191032 |
| Cell growth-regulating nucleolar prote | Q9NX58 | LYAR | Other | -0.323117658 | 0.280624572 | 27.82541613 | 28.10254515 | 28.11976176 | 27.85007407 | 27.83526969 | 28.91436255 | 27.41586785 | 27.33577347 | 28.60272038 |
| Tyrosine-protein kinase Lyn | P07948;P07948-2 | LYN | Extracellular or secreted | 0.714375979 | 1.049473097 | 25.50181897 | 25.39568134 | 25.37726217 | 24.31409749 | 24.40905163 | 24.77087315 | 26.05506923 | 25.78920129 | 25.1832498 |
| Ly6/PLAUR domain-containing protein | O95274 | LYPD3 | Extracellular or secreted | 0.930331007 | 0.883465708 | 27.04291293 | 27.19215884 | 27.25862713 | 27.30086055 | 27.55171507 | 28.58928002 | 28.00640578 | 28.01998582 | 29.23237869 |
| Acyl-protein thioesterase 1 | O75608-2;O75608 | LYPLA1 | Extracellular or secreted | 0.572670606 | 1.041844098 | 26.57715496 | 26.60286194 | 26.57095993 | 25.71800811 | 25.96327253 | 27.24417929 | 26.93842758 | 26.7444518 | 27.37335081 |
| Acyl-protein thioesterase 2 | O95372 | LYPLA2 | Extracellular or secreted | -0.155456931 | 0.509496686 | 27.39312987 | 27.4376976 | 27.40405634 | 26.94178894 | 27.11306872 | 27.78206798 | 27.1419964 | 27.34206232 | 27.08547521 |
| LYR motif-containing protein 2 | Q9NU23 | LYRM2 | Other | 0.818533433 | 0.375361143 | 22.49792101 | 22.10722091 | 20.98726767 | 22.33146545 | 21.22287715 | 21.09911131 | 23.30033692 | 23.36639693 | 20.9117982 |
| Protein LZIC | Q8WZA0;Q8WZA0-2 | LZIC | Other | -0.168169997 | 0.255246576 | 25.81142134 | 26.22855817 | 25.94453943 | 26.94960179 | 27.07673651 | 25.65665529 | 26.29084374 | 26.1419964 | 25.89640614 |
| Leucine zipper transcription factor-lik | Q9NQ48;Q9NQ48-2;Q9NQ48 | LZTFL1 | Extracellular or secreted | 0.083822679 | 0.143607759 | 26.98988506 | 26.9538254 | 26.98587441 | 26.84816464 | 27.15769171 | 26.45134633 | 27.07744978 | 27.24934294 | 26.6180691 |
| Cation-dependent mannose-6-phosph | P20645 | M6PR | Extracellular or secreted | -0.81853879 | 0.924144924 | 27.8237167 | 27.344858 | 27.41860465 | 27.19112354 | 27.08253435 | 27.59270533 | 26.16303938 | 26.36866311 | 27.23945242 |
| O-acetyl-ADP-ribose deacetylase MAC | Q9BQ69 | MACROD1 | Other | -2.691945617 | 0.93094812 | 21.59111445 | 26.71030281 | 26.28312623 | 21.78170265 | 23.88613714 | 25.32885815 | 20.68418844 | 20.30139127 | 23.72920416 |
| Mitotic spindle assembly checkpoint p | Q9Y6D9-3;Q9Y6D9 | MAD1L1 | Extracellular or secreted | -0.228941958 | 0.523493377 | 26.19458419 | 26.06076914 | 26.11735584 | 25.1437877 | 25.17283675 | 25.62488825 | 25.47932684 | 25.41879764 | 25.57216058 |
| Mitotic spindle assembly checkpoint p | Q13257 | MAD2L1 | Other | 0.094936159 | 0.080075323 | 26.68725735 | 26.6945817 | 26.61596655 | 26.54126444 | 26.40072194 | 26.85067025 | 26.950492 | 27.26257505 | 25.96697254 |
| Macrophage erythroblast attacher | Q7L5Y9;Q7L5Y9-3;Q7L5Y9-2; | MAEA | Extracellular or secreted | 0.311010939 | 0.984835887 | 24.40503081 | 24.61290538 | 24.46013989 | 24.79644803 | 25.13241814 | 25.28050913 | 25.23589707 | 24.97285536 | 25.06800607 |
| Repressor of RNA polymerase III trans | Q9H063 | MAF1 | Extracellular or secreted | 0.386429073 | 0.451649356 | 21.89280492 | 23.16235568 | 22.86566985 | 24.19477456 | 22.70256671 | 22.08313343 | 23.50421575 | 22.84632442 | 23.25939962 |
| Melanoma-associated antigen D1 | Q9Y5V3;Q9Y5V3-2 | MAGED1 | Extracellular or secreted | 0.881855227 | 3.641920299 | 22.06177462 | 21.64820835 | 21.8196786 | 21.8499548 | 21.77106148 | 22.33970976 | 22.84331767 | 22.77958551 | 22.7678563 |
| Melanoma-associated antigen D2 | Q9UNF1-2;Q9UNF1 | MAGED2 | Other | -1.029405499 | 1.811408813 | 27.52223047 | 27.21706327 | 27.33577347 | 25.68506614 | 26.32390317 | 26.73569959 | 25.73329633 | 25.812401 | 25.77595422 |
| Protein mago nashi homolog | P61326 | MAGOH | Other | 0.323081482 | 0.973599402 | 29.77220489 | 30.22481029 | 30.09295243 | 29.29230594 | 29.34764826 | 29.65727565 | 30.0640683 | 30.18364261 | 29.91513227 |
| Protein MAK16 homolog | Q9BXY0 | MAK16 | Other | -2.038393757 | 1.635592736 | 24.97110181 | 25.30966397 | 25.11052218 | 23.95251518 | 23.41673775 | 23.50567008 | 21.79253137 | 23.28093385 | 21.944459 |
| Endoplasmic reticulum mannosyl-olig | Q9UKM7 | MAN1B1 | Extracellular or secreted | -0.60948431 | 0.18502005 | 23.12357331 | 23.48689393 | 22.52778815 | 22.71736685 | 21.85145277 | 24.76724005 | 21.11693209 | 21.57371213 | 24.71806038 |
| Alpha-mannosidase 2 | Q16706 | MAN2A1 | Extracellular or secreted | 0.274210838 | 0.122594334 | 21.5451138 | 22.65094918 | 21.19685757 | 23.21658602 | 23.05842582 | 24.97832153 | 22.63962334 | 22.24071452 | 24.26542161 |
| Lysosomal alpha-mannosidase;Lysoso | O00754-2;O00754 | MAN2B1 | Extracellular or secreted | -0.612335017 | 2.001210272 | 26.75327878 | 26.84002124 | 26.73052632 | 27.58756431 | 27.44671645 | 26.84110168 | 26.3782549 | 26.31934467 | 26.56499978 |
| Alpha-mannosidase 2C1 | Q9NTJ4-2;Q9NTJ4;Q9NTJ4-4; | MAN2C1 | Other | -0.247272058 | 0.417895144 | 24.97910696 | 25.32728263 | 25.09880605 | 24.88729984 | 25.17058465 | 23.8305508 | 24.79758633 | 24.76951181 | 24.33790115 |
| Beta-mannosidase | O00462 | MANBA | Extracellular or secreted | -0.050457744 | 0.038388225 | 24.58520188 | 24.88613714 | 24.69795171 | 24.65255836 | 25.02047338 | 24.50542779 | 24.45418933 | 24.08901657 | 25.479296 |
| Mesencephalic astrocyte-derived neu | P55145 | MANF | Extracellular or secreted | -0.567552115 | 1.800543477 | 29.24297632 | 29.24336229 | 29.20759989 | 29.21282634 | 29.35902809 | 28.5590295 | 28.73676427 | 28.6322857 | 28.34070489 |
| MANSC domain-containing protein 1 | Q9H8J5;Q9H8J5-2 | MANSC1 | Extracellular or secreted | 2.485469838 | 0.398206929 | 21.71408037 | 23.37504261 | 23.52874215 | 26.46229482 | 21.36014144 | 23.06732347 | 26.58434185 | 29.27697382 | 21.34890628 |
| Microtubule-associated protein 1B;M | P46821 | MAP1B | Extracellular or secreted | -1.345780795 | 1.526073265 | 27.59391651 | 27.46723276 | 27.56623444 | 26.7521305 | 26.79141391 | 26.66057744 | 26.0464561 | 25.15344565 | 26.17850864 |
| Microtubule-associated proteins 1A/1 | Q9GZQ8;A6NCE7 | MAP1LC3B;MAP1 | Other | 0.664800558 | 1.894274026 | 26.67815007 | 26.89758367 | 26.77756034 | 26.47325376 | 26.79253137 | 25.70601303 | 27.36168654 | 27.18129833 | 27.11396292 |
| Microtubule-associated protein 1S;M | Q66K74-2;Q66K74 | MAP1S | Other | -0.242320905 | 0.783053934 | 27.03359675 | 27.28621467 | 27.3316795 | 27.1191186 | 27.18110872 | 27.39042627 | 27.16739135 | 26.76002244 | 27.01669575 |
| Dual specificity mitogen-activated pro | Q02750;Q02750-2 | MAP2K1 | Extracellular or secreted | 0.221095032 | 0.409841343 | 28.16734349 | 28.05222118 | 28.29513392 | 28.96411027 | 29.42726373 | 29.05835047 | 28.91287897 | 28.9616563 | 28.77096134 |
| Dual specificity mitogen-activated pro | P36507 | MAP2K2 | Extracellular or secreted | 0.920094766 | 1.208584348 | 23.17783983 | 23.05860748 | 21.88981734 | 23.44253049 | 23.19778541 | 23.4563845 | 23.41066976 | 24.04984523 | 24.41125184 |
| Dual specificity mitogen-activated pro | P46734-2;P46734;P46734-3 | MAP2K3 | Other | 0.090335215 | 0.06812074 | 23.63267399 | 23.9655201 | 24.03397475 | 23.97324962 | 23.71554898 | 23.45801303 | 24.7160202 | 23.66893085 | 23.27554483 |
| Microtubule-associated protein 4 | P27816;P27816-6;P27816-2 | MAP4 | Extracellular or secreted | 0.236018667 | 0.59232643 | 31.57673161 | 31.52733888 | 31.52196849 | 31.43251001 | 31.52159415 | 30.94726238 | 31.80339682 | 31.80654992 | 31.36181202 |
| Microtubule-associated protein 4 | P27816-5 | MAP4 | Extracellular or secreted | -0.310667435 | 0.156560022 | 21.75049417 | 22.64647345 | 22.32124551 | 23.37570553 | 23.33660789 | 22.75782249 | 22.74453906 | 21.09319767 | 23.32443549 |
| Ensconsin | Q14244-2;Q14244;Q14244-3 | MAP7 | Extracellular or secreted | -0.760790057 | 1.424470748 | 26.59448612 | 26.18710707 | 26.13854383 | 25.76769469 | 25.53124342 | 24.68602854 | 25.05017747 | 25.10377603 | 25.01622816 |
| MAP7 domain-containing protein 1 | Q3KQU3-3;Q3KQU3-2;Q3KQU | MAP7D1 | Other | -2.066145696 | 1.088362809 | 24.96106563 | 24.65894451 | 24.86031788 | 24.55718975 | 24.84627659 | 24.6597612 | 23.78406687 | 21.58265471 | 22.70661911 |
| MAP7 domain-containing protein 2 | Q96T17-5;Q96T17-4;Q96T17 | MAP7D2 | Other | -0.634852584 | 1.09679306 | 23.683889 | 23.77645633 | 23.30103505 | 22.70484759 | 23.4867712 | 22.30167704 | 22.59498151 | 22.89827588 | 22.22952296 |
| Mitogen-activated protein kinase 1 | P28482;P28482-2 | MAPK1 | Extracellular or secreted | -0.1630957 | 0.429260279 | 30.14952627 | 30.23728835 | 30.18919213 | 29.79584759 | 29.86604084 | 30.13980293 | 29.85397458 | 30.15966573 | 29.68592164 |
| Mitogen-activated protein kinase 12 | P53778-2;P53778 | MAPK12 | Other | -0.133868628 | 0.043473335 | 24.77384348 | 25.11584886 | 24.87051298 | 23.96974133 | 23.48763006 | 19.69329274 | 24.14895441 | 24.2550107 | 22.14986373 |
| Mitogen-activated protein kinase 14 | Q16539;Q16539-3;Q16539-5 | MAPK14 | Extracellular or secreted | -0.010318278 | 0.014125956 | 27.84828405 | 27.77975331 | 27.83569135 | 27.93893229 | 27.87876715 | 27.94357847 | 28.02416716 | 28.14554762 | 27.41183369 |
| Mitogen-activated protein kinase 15 | Q8TD08-2;Q8TD08-3;Q8TD08 | MAPK15 | Extracellular or secreted | 0.371274011 | 0.435668614 | 26.60710226 | 26.41781634 | 26.53552125 | 26.17056555 | 26.09297259 | 27.0558967 | 27.11555125 | 27.21706327 | 26.22114486 |
| MAPK-interacting and spindle-stabiliz | Q8NDC0 | MAPK1IP1L | Other | -0.004488759 | 0.001116485 | 23.07483136 | 23.13617884 | 22.96382814 | 24.32319883 | 22.90458426 | 22.04412975 | 24.27653906 | 24.29487984 | 20.63849041 |
| Mitogen-activated protein kinase 3 | P27361;P27361-2;P27361-3 | MAPK3 | Extracellular or secreted | -0.201080762 | 0.71797594 | 26.23083155 | 26.21957638 | 26.28330289 | 25.86858333 | 26.02660685 | 26.03080064 | 25.9263293 | 26.08425903 | 25.7160202 |
| Mitogen-activated protein kinase 9 | P45984-2;P45984;P45984-3;F | MAPK9 | Other | -0.122623562 | 0.208576217 | 25.76108836 | 25.61512468 | 25.69365157 | 25.43526712 | 25.70672449 | 25.00187202 | 25.51294995 | 25.68557416 | 25.04046933 |
| MAP kinase-activated protein kinase 3 | Q16644 | MAPKAPK3 | Other | -0.10367146 | 0.266035802 | 26.01695073 | 25.99031798 | 25.90974131 | 25.92241001 | 26.35845388 | 25.6019274 | 25.9730087 | 25.99780848 | 25.6180691 |
| Microtubule-associated protein RP/EB | Q15691 | MAPRE1 | Extracellular or secreted | 0.372589551 | 1.663855619 | 31.16026703 | 30.89684782 | 30.89627043 | 30.88990378 | 30.99084542 | 31.14995017 | 31.21071481 | 31.41341844 | 31.48567772 |
| Microtubule-associated protein RP/EB | Q15555-4;Q15555-5;Q15555 | MAPRE2 | Other | -0.760808573 | 0.551548999 | 22.09013181 | 22.48234613 | 22.61337762 | 21.91928331 | 22.12069954 | 21.45202035 | 21.74644707 | 21.99696855 | 20.31308804 |
| Microtubule-associated protein tau | P10636-2;P10636-6;P10636-3 | MAPT | Extracellular or secreted | -0.349347408 | 1.214863405 | 24.46631499 | 24.06570735 | 24.1547594 | 24.78979828 | 24.47642484 | 23.81528713 | 24.05606214 | 23.8657076 | 23.91433403 |
| Microtubule-associated protein tau | P10636-4;P10636-5;P10636-1 | MAPT | Extracellular or secreted | -0.160029576 | 0.790863498 | 27.0557933 | 26.96664257 | 27.05372358 | 27.28612652 | 27.32173753 | 26.73969616 | 26.80857656 | 26.98544016 | 26.93775437 |
| Myristoylated alanine-rich C-kinase su | P29966 | MARCKS | Extracellular or secreted | 0.836415416 | 2.445645354 | 29.14102193 | 28.78718459 | 28.97051499 | 28.51450085 | 28.57855921 | 29.42676404 | 29.56632518 | 29.67841875 | 29.97377513 |
| MARCKS-related protein | P49006 | MARCKSL1 | Extracellular or secreted | 0.924483079 | 0.258668377 | 25.48040589 | 22.22445426 | 24.67222634 | 24.18465019 | 22.99229692 | 21.10769982 | 25.5341561 | 25.64902566 | 21.92113419 |
| Serine/threonine-protein kinase MAR | Q7KZI7-15;Q7KZI7-5;Q7KZI7- | MARK2 | Extracellular or secreted | 0.16504864 | 0.261406581 | 26.59974444 | 26.41005508 | 26.55625438 | 25.39054108 | 25.26083571 | 26.3952891 | 26.13155494 | 26.37951137 | 26.29043951 |
| MAP/microtubule affinity-regulating k | P27448-8;P27448-3;P27448-4 | MARK3 | Extracellular or secreted | 0.510400487 | 0.869481205 | 23.42495972 | 23.22070955 | 22.98659065 | 23.50724395 | 22.8800386 | 23.75999705 | 23.35420525 | 24.03010603 | 24.03665993 |
| Methionine--tRNA ligase, cytoplasmic | P56192;P56192-2 | MARS | Extracellular or secreted | -0.212110711 | 0.878204043 | 29.25595166 | 29.23011234 | 29.18397382 | 29.02593661 | 29.07370116 | 29.15487044 | 29.02902135 | 29.03826236 | 28.75865717 |
| MARVEL domain-containing protein 3 | Q96A59;Q96A59-2 | MARVELD3 | Other | 1.683007691 | 2.013839115 | 22.00958788 | 21.30781478 | 21.11148495 | 20.33708432 | 22.33401135 | 23.30256976 | 23.18910727 | 23.51833331 | 23.542859 |
| S-adenosylmethionine synthase isofor | P31153;P31153-2 | MAT2A | Other | -0.042710571 | 0.12050754 | 31.07335641 | 31.21864602 | 31.13301865 | 31.14272682 | 31.20856531 | 30.60063078 | 31.08765165 | 31.13430461 | 30.83838402 |
| Methionine adenosyltransferase 2 sul | Q9NZL9;Q9NZL9-2;Q9NZL9-4 | MAT2B | Extracellular or secreted | 0.118447719 | 0.419131058 | 30.60249032 | 30.71317418 | 30.60875965 | 30.27907921 | 30.40557865 | 30.12275119 | 30.6855374 | 30.628623 | 30.40709936 |
| Matrilin-2 | O00339-3;O00339-2;O00339 | MATN2 | Extracellular or secreted | -1.37434497 | 0.387324668 | 22.52439479 | 19.74641306 | 22.55595025 | 23.40846865 | 24.69975289 | 26.86985436 | 20.25352896 | 21.34385349 | 24.18199964 |
| Matrin-3 | P43243;P43243-2 | MATR3 | Extracellular or secreted | -0.343998666 | 0.960941599 | 29.41520299 | 29.8555645 | 29.5030937 | 29.14282418 | 29.06676481 | 30.1016188 | 29.12489679 | 29.34335494 | 29.04228676 |
| Mitochondrial antiviral-signaling prot | Q7Z434;Q7Z434-4 | MAVS | Other | -0.854796987 | 3.146023606 | 26.61090791 | 26.60328653 | 26.58376822 | 26.65880835 | 26.36861316 | 26.17155845 | 25.47657935 | 25.68297873 | 25.77452227 |
| Protein max | P61244-3;P61244-4;P61244-2 | MAX | Other | -0.075451426 | 0.018337998 | 23.12867908 | 20.93236183 | 21.73888227 | 24.56040048 | 23.77605466 | 25.27533169 | 21.8092343 | 22.1765165 | 25.49374993 |
| Myoglobin | P02144 | MB | Extracellular or secreted | -2.49605408 | 0.78044877 | 26.13259465 | 26.5016064 | 26.49229881 | 25.92915455 | 26.31736196 | 24.55981724 | 24.67109268 | 24.70798844 | 21.09917344 |
| Methyl-CpG-binding domain protein 2 | Q9UBB5 | MBD2 | Other | -1.614253433 | 0.802536149 | 23.53919479 | 23.32058463 | 23.62678294 | 22.76409407 | 22.80688725 | 23.0347641 | 20.08665755 | 22.18043732 | 22.43629871 |
| Methyl-CpG-binding domain protein 3 | O95983-2;O95983 | MBD3 | Other | -0.932517038 | 2.018647284 | 25.68757773 | 25.97574408 | 25.85922803 | 25.56340042 | 25.33630142 | 25.74342488 | 24.65600057 | 24.56226525 | 25.06702135 |
| Metallo-beta-lactamase domain-cont | A4D2B0 | MBLAC1 | Other | 0.930815185 | 0.539499957 | 22.92117593 | 22.36728023 | 20.42242174 | 21.30326317 | 21.85143513 | 22.37500288 | 23.89235826 | 22.85448949 | 21.66588734 |
| Muscleblind-like protein 1 | Q9NR56-3;Q9NR56-4;Q9NR5 | MBNL1 | Extracellular or secreted | -0.146290744 | 0.130109589 | 23.10722091 | 23.17610557 | 23.40145455 | 23.06305928 | 23.08207951 | 22.00939216 | 23.24784467 | 23.11227341 | 22.12066567 |
| Lysophospholipid acyltransferase 7 | Q96N66;Q96N66-3;Q96N66-1 | MBOAT7 | Other | -0.67766137 | 0.287999701 | 22.57937181 | 21.82734842 | 24.06184052 | 23.14119349 | 22.00140205 | 26.55628362 | 22.34378854 | 21.73764511 | 23.9693022 |
| Myelin basic protein | P02686-2;P02686 | MBP | Extracellular or secreted | 2.217464107 | 1.85492874 | 21.30547561 | 24.66101258 | 24.72416367 | 22.51807406 | 24.43657024 | 22.15586252 | 25.26878013 | 25.65512713 | 25.62906441 |
| Membrane-bound transcription facto | Q14703 | MBTPS1 | Other | 1.639727828 | 0.870818106 | 21.94128306 | 20.6222647 | 20.97062048 | 24.72666013 | 20.75126646 | 25.10749231 | 23.48443744 | 23.51412457 | 24.98041505 |
| Cell surface glycoprotein MUC18 | P43121;P43121-2 | MCAM | Extracellular or secreted | -1.187479117 | 2.257080636 | 25.67903653 | 25.88776465 | 25.89640614 | 26.20780466 | 26.36724716 | 25.90435311 | 25.11481686 | 24.75669273 | 24.53735919 |
| Malonyl-CoA-acyl carrier protein tran | Q8IVS2 | MCAT | Other | -1.438921259 | 0.475337904 | 23.00235287 | 22.9613218 | 23.01005886 | 23.84536753 | 22.9648506 | 24.05490368 | 23.12298909 | 22.89613355 | 19.58354125 |
| Methylcrotonoyl-CoA carboxylase bet | Q9HCC0-2;Q9HCC0 | MCCC2 | Other | -0.387713066 | 0.170069263 | 24.66871449 | 20.77751016 | 24.42809483 | 24.40490091 | 25.02947428 | 25.69766013 | 24.76167175 | 23.6736826 | 22.90468386 |
| DNA replication licensing factor MCM | P49736 | MCM2 | Other | 0.53119844 | 0.813835363 | 29.62366094 | 29.55687559 | 29.51045069 | 29.20827459 | 29.26869983 | 29.13072557 | 30.2190501 | 30.09848482 | 29.42540401 |
| DNA replication licensing factor MCM | P25205;P25205-2 | MCM3 | Other | 0.336421829 | 0.435040744 | 29.539694 | 29.55771563 | 29.47173536 | 29.10636945 | 29.12344208 | 30.07813725 | 30.23740233 | 29.93578904 | 29.274621 |
| DNA replication licensing factor MCM | P33991 | MCM4 | Other | 0.336815479 | 0.917107651 | 29.55738698 | 29.59793462 | 29.47519978 | 29.43233079 | 29.50094953 | 29.6567303 | 30.07902792 | 29.92053722 | 29.62114729 |
| DNA replication licensing factor MCM | P33992 | MCM5 | Other | 0.028236503 | 0.071953448 | 28.55603506 | 28.55786168 | 28.52798198 | 28.12015742 | 28.16106036 | 28.77313292 | 28.59281224 | 28.54966036 | 28.29035162 |
| DNA replication licensing factor MCM | Q14566 | MCM6 | Other | 0.289553806 | 0.972850492 | 30.52962107 | 30.57181351 | 30.53278223 | 30.28669939 | 30.35971934 | 30.49628969 | 30.89951528 | 30.82547678 | 30.53213196 |
| DNA replication licensing factor MCM | P33993;P33993-3 | MCM7 | Other | 0.549483987 | 1.10987067 | 28.56230893 | 28.5953757 | 28.69716961 | 28.25200812 | 28.41090424 | 28.99824979 | 29.41908708 | 29.21131842 | 28.77605466 |
| Mini-chromosome maintenance comp | Q9BTE3-2;Q9BTE3;Q9BTE3-3 | MCMBP | Extracellular or secreted | 0.470454338 | 0.720782987 | 27.9306666 | 28.04598765 | 28.05940817 | 27.74464427 | 27.88308643 | 28.02601579 | 28.79401996 | 28.50282822 | 27.95941929 |
| Malignant T-cell-amplified sequence 1 | Q9ULC4;Q9ULC4-3;Q9ULC4-2 | MCTS1 | Extracellular or secreted | 0.040217278 | 0.103177228 | 28.46719388 | 28.55512089 | 28.54285163 | 27.93713698 | 27.86082698 | 28.35737192 | 28.42438314 | 28.36436099 | 28.19215884 |
| Mediator of DNA damage checkpoint | Q14676;Q14676-2;Q14676-3 | MDC1 | Other | -0.960974598 | 1.300956192 | 29.30849704 | 29.29177919 | 29.22239381 | 29.29320535 | 29.33310944 | 29.05260988 | 28.53400763 | 28.51069591 | 27.82317003 |
| Malate dehydrogenase, cytoplasmic | P40925;P40925-3;P40925-2 | MDH1 | Extracellular or secreted | -0.140770901 | 0.523603891 | 32.62897058 | 32.68916693 | 32.59916803 | 32.73212474 | 33.02082041 | 32.14029065 | 32.57713694 | 32.47522877 | 32.43059224 |
| Malate dehydrogenase, mitochondria | P40926;P40926-2 | MDH2 | Extracellular or secreted | -0.333673278 | 0.686931759 | 33.44997585 | 33.35639096 | 33.31810472 | 33.55670661 | 33.71845076 | 32.90849064 | 33.16044737 | 33.30130222 | 32.69129034 |
| Midkine | P21741 | MDK | Extracellular or secreted | -1.030838238 | 0.507028248 | 28.82341302 | 28.83282769 | 28.78147403 | 29.01059994 | 29.24240853 | 32.13827772 | 27.66383777 | 27.77812457 | 29.88002341 |
| Midasin | Q9NU22 | MDN1 | Other | -0.587208534 | 0.551349324 | 23.39152471 | 23.54014129 | 23.13832118 | 21.44700363 | 24.31056563 | 24.69434258 | 22.63274055 | 22.66042511 | 23.20615825 |

| | | | | | | | | | | | | | | |
|---|---|---|---|---|---|---|---|---|---|---|---|---|---|---|
| Magnesium-dependent phosphatase | Q86V88;Q86V88-3;Q86V88-2 | MDP1 | Extracellular or secreted | -0.031441467 | 0.046555624 | 23.31506551 | 23.25738669 | 23.22682139 | 23.20127502 | 23.29502003 | 22.66819509 | 22.76937056 | 23.47283559 | 23.14535132 |
| NADP-dependent malic enzyme | P48163;P48163-2 | ME1 | Other | 0.830962809 | 2.351016317 | 31.0487571 | 30.98359313 | 31.00350013 | 30.67404644 | 30.63009962 | 29.95684658 | 31.43574202 | 31.43688393 | 31.76868398 |
| NAD-dependent malic enzyme, mitoc | P23368;P23368-2 | ME2 | Other | -0.519245564 | 2.123676155 | 27.7693857 | 27.76850258 | 27.77184296 | 28.03627215 | 28.12084954 | 28.34528111 | 27.36861316 | 27.35152875 | 27.62818842 |
| Male-enhanced antigen 1 | Q16626 | MEA1 | Other | 1.52529685 | 2.218579572 | 21.02220833 | 21.61415071 | 20.8760893 | 23.21861373 | 21.96107663 | 22.10587458 | 23.29558062 | 23.62978705 | 23.04952952 |
| Methyl-CpG-binding protein 2 | P51608;P51608-2 | MECP2 | Extracellular or secreted | 0.031681998 | 0.036562489 | 26.04470645 | 24.53789234 | 25.00684447 | 25.0232684 | 24.72889289 | 25.89947494 | 25.4687093 | 25.32299262 | 24.92388382 |
| Mediator of RNA polymerase II transc | Q15648;Q15648-3 | MED1 | Other | -1.269379691 | 0.83917934 | 26.08111248 | 26.18754131 | 24.8644329 | 23.88483381 | 23.46276286 | 25.39842405 | 24.3561548 | 22.58856825 | 24.18669158 |
| Mediator of RNA polymerase II transc | Q9P086 | MED11 | Other | 0.098911648 | 0.269726558 | 22.31727569 | 22.30810901 | 22.28968347 | 22.02635037 | 22.25557265 | 22.17237135 | 22.32391347 | 22.55613742 | 22.10136533 |
| Mediator of RNA polymerase II transc | Q93074-3;Q93074;Q93074-2 | MED12 | Other | -1.620539376 | 0.919121978 | 23.5005735 | 23.61144272 | 24.02013423 | 23.62332604 | 21.22994139 | 24.78935056 | 21.62611371 | 20.79558029 | 23.10407209 |
| Mediator of RNA polymerase II transc | O60244 | MED14 | Other | -0.625404412 | 0.982570653 | 25.40665345 | 25.67317038 | 24.73407201 | 25.44180263 | 25.13914912 | 25.51406436 | 24.52354712 | 24.37835413 | 25.17634149 |
| Mediator of RNA polymerase II transc | Q96RN5-3;Q96RN5-2;Q96RN | MED15 | Other | -0.47680407 | 0.545222971 | 25.70453629 | 25.42374223 | 24.91095375 | 24.95020274 | 24.85471775 | 25.02580464 | 24.96974133 | 25.02917937 | 24.00564579 |
| Mediator of RNA polymerase II transc | Q9NVC6 | MED17 | Other | -0.374327639 | 0.547366786 | 23.23037349 | 23.92732442 | 23.55613742 | 23.18122628 | 23.32457283 | 23.48652572 | 23.43288032 | 22.5721895 | 23.22502734 |
| Mediator of RNA polymerase II transc | Q9BUE0 | MED18 | Other | -0.065464244 | 0.085284813 | 24.25190883 | 24.26813761 | 24.51074872 | 23.25336682 | 22.85241215 | 23.4986272 | 23.6070458 | 23.72650423 | 23.7876579 |
| Mediator of RNA polymerase II transc | Q15528-2;Q15528 | MED22 | Other | -2.435911468 | 1.251577526 | 25.21248778 | 25.31565292 | 25.25911223 | 25.51409447 | 25.46519421 | 25.65061507 | 23.59479932 | 23.56302213 | 21.74302249 |
| Mediator of RNA polymerase II transc | Q9ULK4-4;Q9ULK4-5;Q9ULK4 | MED23 | Other | -0.23915198 | 0.241566963 | 23.37980879 | 23.35823588 | 23.58966577 | 23.57276776 | 23.55285856 | 24.41982648 | 23.29165187 | 24.04893116 | 22.87854266 |
| Mediator of RNA polymerase II transc | Q6P2C8;Q6P2C8-2 | MED27 | Other | -0.281965197 | 0.421335429 | 22.05453449 | 22.42008358 | 23.31782771 | 22.25596704 | 22.76824007 | 23.42393453 | 22.06141783 | 22.70066323 | 22.51231706 |
| Mediator of RNA polymerase II transc | Q9NX70;Q9NX70-2 | MED29 | Other | 0.640419768 | 2.208939288 | 24.59536858 | 23.56987415 | 24.21504591 | 24.35521397 | 24.48363818 | 24.25472243 | 24.82599833 | 24.98080725 | 24.85138534 |
| Mediator of RNA polymerase II transc | Q96HR3-2;Q96HR3 | MED30 | Other | 0.15874471 | 0.160278615 | 22.84532925 | 22.83360355 | 23.0318903 | 20.80610721 | 22.01441233 | 23.03959113 | 22.83155621 | 22.40795026 | 22.52219454 |
| Mediator of RNA polymerase II transc | Q9Y3C7 | MED31 | Other | -0.889568568 | 1.755330894 | 23.47047915 | 23.27852546 | 23.16568663 | 22.92927647 | 23.20986562 | 24.10101259 | 22.08669767 | 22.61936834 | 22.70265125 |
| Mediator of RNA polymerase II transc | Q9NPJ6;Q9NPJ6-2 | MED4 | Other | -0.849059212 | 2.621755313 | 24.64855946 | 24.65534554 | 24.50784883 | 24.14335955 | 24.3461777 | 24.82764662 | 23.5659295 | 23.5513337 | 23.90002801 |
| Mediator of RNA polymerase II transc | O43513 | MED7 | Other | -0.806903719 | 0.360676429 | 23.27582896 | 23.01816553 | 23.14119349 | 20.74514878 | 22.05665387 | 23.64921758 | 22.77371938 | 20.25243418 | 22.49623939 |
| Mediator of RNA polymerase II transc | Q96G25-3;Q96G25;Q96G25-2 | MED8 | Other | -1.154618385 | 0.871834794 | 23.75694717 | 23.11784725 | 23.26313048 | 24.04198863 | 21.98450857 | 21.90392579 | 20.85526588 | 22.24452411 | 22.4705288 |
| Myocyte-specific enhancer factor 2C;I | Q06413-6;Q06413-5;Q06413 | MEF2C;MEF2D;M | Other | -0.700534889 | 1.383682198 | 24.86499957 | 24.77966566 | 24.76486349 | 24.34435009 | 23.68912526 | 24.76106299 | 23.64427439 | 24.21408251 | 23.64207197 |
| Protein MEMO1 | Q9Y316;Q9Y316-3;Q9Y316-2 | MEMO1 | Other | 0.490116923 | 3.344847652 | 27.41304512 | 27.46060863 | 27.47691921 | 27.25655914 | 27.14530467 | 27.61526502 | 27.95748327 | 27.82662878 | 27.87008962 |
| 7SK snRNA methylphosphate capping | Q7L2J0 | MEPCE | Other | -0.773983984 | 0.368074567 | 24.2327172 | 21.39490325 | 23.15949805 | 23.39218009 | 23.447962 | 24.06964578 | 23.87932822 | 21.28797766 | 22.35919535 |
| LDLR chaperone MESD | Q14696 | MESDC2 | Extracellular or secreted | -0.027093615 | 0.051859621 | 27.7153526 | 28.08997629 | 27.84834375 | 27.71881818 | 28.03044286 | 27.19535418 | 27.79550702 | 27.92823977 | 27.49411629 |
| Hepatocyte growth factor receptor | P08581;P08581-2;P08581-3 | MET | Extracellular or secreted | 2.530564186 | 0.507947393 | 23.12461484 | 20.60059533 | 22.94933017 | 23.68163906 | 21.84842733 | 28.16729563 | 23.93454106 | 24.23286355 | 29.61023913 |
| Methionine aminopeptidase 1 | P53582 | METAP1 | Other | -0.031004979 | 0.016991321 | 28.7391167 | 28.88069491 | 28.70757383 | 28.5657988 | 28.69158987 | 27.45951467 | 28.95999959 | 29.08420834 | 27.38492152 |
| Methionine aminopeptidase 2 | P50579;P50579-3 | METAP2 | Extracellular or secreted | 0.537015421 | 0.685371504 | 30.26796355 | 30.18872066 | 30.17307511 | 30.40334538 | 30.36369284 | 29.57213889 | 30.85378129 | 31.12293628 | 30.11879691 |
| tRNA (guanine-N(7)-)-methyltransfera | Q9UBP6 | METTL1 | Other | 0.238475395 | 1.305312643 | 24.59331817 | 24.74046841 | 24.71858304 | 24.70603939 | 24.38370271 | 24.25616318 | 24.81279268 | 24.90016625 | 24.70160471 |
| Methyltransferase-like protein 13 | Q8N6R0-3;Q8N6R0-4;Q8N6R | METTL13 | Other | -0.073720491 | 0.033172752 | 22.57600172 | 21.55772732 | 21.17537636 | 22.70659803 | 21.1715526 | 24.03724665 | 22.70746193 | 22.7525644 | 20.93106354 |
| Methyltransferase-like protein 16 | Q86W50;Q86W50-2 | METTL16 | Other | -0.41811551 | 0.76621108 | 22.79171871 | 22.57929752 | 22.41593875 | 22.91639486 | 22.92059855 | 23.60873851 | 22.84921036 | 22.21035624 | 22.30243031 |
| N6-adenosine-methyltransferase 70 k | Q86U44 | METTL3 | Other | -0.339385007 | 0.420862902 | 26.88692787 | 26.36592992 | 26.28402697 | 26.36379308 | 26.5903513 | 26.32393752 | 26.39285155 | 26.46931502 | 25.52716174 |
| Microfibrillar-associated protein 1 | P55081 | MFAP1 | Extracellular or secreted | 1.086062112 | 1.000325984 | 25.43682437 | 25.33019257 | 25.29270526 | 25.36138536 | 25.27756807 | 24.49801844 | 25.97088246 | 25.80311655 | 27.08253435 |
| Lactadherin;Lactadherin short form;M | Q08431;Q08431-3;Q08431-4 | MFGE8 | Extracellular or secreted | 1.169492737 | 0.732650963 | 23.96446285 | 22.4341255 | 21.21099198 | 23.75276855 | 24.32106655 | 26.56867159 | 24.61475971 | 24.65053291 | 25.36922911 |
| Melanotransferrin | P08582 | MFI2 | Extracellular or secreted | 0.073327992 | 0.085297588 | 28.18238806 | 28.10084221 | 28.20373176 | 28.71109096 | 28.77947155 | 28.78749599 | 28.18768288 | 28.35594484 | 29.05886652 |
| Alpha-1,3-mannosyl-glycoprotein 2-be | P26572 | MGAT1 | Extracellular or secreted | -0.113292608 | 0.321376823 | 23.64812055 | 23.62544577 | 23.50796977 | 23.57161101 | 23.68602854 | 24.18412047 | 23.41841163 | 23.55894194 | 23.79441666 |
| Protein O-GlcNAcase | O60502;O60502-4;O60502-2 | MGEA5 | Other | -0.295341256 | 0.539342093 | 27.50907293 | 27.50861968 | 27.30486877 | 27.59085096 | 27.58627619 | 27.27533169 | 27.39173774 | 27.35800107 | 26.75174753 |
| Mitochondrial genome maintenance e | Q9BQP7 | MGME1 | Other | 1.090563758 | 0.55493542 | 24.9294029 | 24.79337509 | 24.95522314 | 20.72513473 | 20.8885692 | 24.6845847 | 24.9729868 | 24.90136374 | 23.88548562 |
| Matrix Gla protein | P08493;P08493-2 | MGP | Extracellular or secreted | 3.953498426 | 2.362760413 | 23.53338392 | 23.67443712 | 23.55367897 | 25.40366637 | 25.25237814 | 24.42046914 | 27.60766669 | 27.83960085 | 29.33223457 |
| E3 ubiquitin-protein ligase MGRN1 | O60291-4;O60291;O60291-3 | MGRN1 | Extracellular or secreted | 0.589257881 | 0.895744942 | 22.6432617 | 22.1875197 | 23.46176419 | 21.55157523 | 22.8124528 | 23.45513053 | 23.14634617 | 23.57888308 | 23.09839647 |
| Microsomal glutathione S-transferase | P10620;P10620-2 | MGST1 | Other | 0.541620209 | 0.135833881 | 23.52862294 | 22.30114672 | 23.64383418 | 22.94326099 | 22.89506119 | 27.64437067 | 22.51881353 | 23.87942171 | 26.70477373 |
| Melanoma-derived growth regulatory | Q16674 | MIA | Extracellular or secreted | 3.980755422 | 1.733967508 | 24.12115595 | 23.77595422 | 23.37583807 | 25.19576718 | 25.1074524 | 24.64008691 | 27.42149679 | 27.858991 | 29.76990585 |
| MHC class I polypeptide-related seque | Q29983 | MICA | Extracellular or secreted | 1.66218877 | 1.417640351 | 21.99241346 | 22.52035059 | 24.20463681 | 21.17641817 | 24.5941159 | 21.67827389 | 24.71324302 | 23.7329342 | 24.6234935 |
| Protein-methionine sulfoxide oxidase | Q7RTP6;Q7RTP6-2;Q7RTP6-4 | MICAL3 | Extracellular or secreted | -0.886370691 | 0.479808533 | 23.99524407 | 24.65758232 | 25.26549315 | 23.10939071 | 23.30326682 | 23.78366731 | 22.17200224 | 22.75746751 | 24.46874037 |
| MICAL-like protein 1 | Q8N3F8 | MICALL1 | Extracellular or secreted | 0.666695342 | 0.944964652 | 24.10597499 | 25.12048375 | 25.46007738 | 25.09695799 | 26.21905933 | 24.81254789 | 25.90246823 | 25.36586319 | 26.13930528 |
| MHC class I polypeptide-related seque | Q29980;Q29980-2 | MICB | Extracellular or secreted | 3.926447605 | 0.840636095 | 22.96924949 | 21.26990464 | 20.10152135 | 25.10992468 | 20.21430396 | 23.77515049 | 24.46463349 | 24.28369147 | 29.75104516 |
| Migration and invasion enhancer 1 | Q9BRT3 | MIEN1 | Extracellular or secreted | 0.025381895 | 0.022852878 | 24.84904806 | 24.85186187 | 24.76754316 | 24.50809071 | 24.46276286 | 24.78191796 | 24.77183037 | 25.30504278 | 24.10988484 |
| Mesoderm induction early response p | Q8N108-16;Q8N108-14;Q8N1 | MIER1 | Extracellular or secreted | 0.209798932 | 0.281471727 | 26.45494234 | 26.40798266 | 26.53130292 | 25.13171192 | 26.40026592 | 25.09735994 | 26.20290086 | 26.44726841 | 25.99101038 |
| Macrophage migration inhibitory fact | P14174 | MIF | Extracellular or secreted | 0.240858893 | 0.577384394 | 32.55545006 | 32.81214697 | 32.7591693 | 32.28060094 | 32.33151401 | 31.69909096 | 32.84121793 | 32.63272339 | 32.46762147 |
| MIF4G domain-containing protein | A9UHW6;A9UHW6-2;A9UHW | MIF4GD | Other | 0.36300099 | 1.075693781 | 23.08453074 | 23.09082638 | 23.173546 | 22.35431288 | 22.4299471 | 22.52377442 | 23.13918035 | 23.30089545 | 22.97739593 |
| Bifunctional lysine-specific demethyla | Q8IUF8;Q8IUF8-4;Q8IUF8-3 | MINA | Other | -0.294844491 | 0.100632233 | 25.01784282 | 25.02356453 | 22.97533301 | 25.34292702 | 24.6065376 | 25.10972546 | 25.12750667 | 25.52886136 | 22.49706372 |
| Misshapen-like kinase 1 | Q8N4C8-4;Q8N4C8;Q8N4C8- | MINK1 | Extracellular or secreted | 1.064991631 | 2.274194578 | 26.30666015 | 25.03363875 | 26.16184845 | 25.36813021 | 25.69962053 | 26.08723686 | 26.75149216 | 26.61329892 | 27.15875128 |
| Multiple inositol polyphosphate phos | Q9UNW1;Q9UNW1-4;Q9UNW | MINPP1 | Extracellular or secreted | -0.134854626 | 0.260278075 | 25.55109896 | 25.43090424 | 25.48566621 | 25.01648323 | 25.20093841 | 25.865094 | 25.06056302 | 25.19091639 | 25.61904924 |
| WD repeat-containing protein mio | Q9NXC5-2;Q9NXC5 | MIOS | Other | 0.883194757 | 0.658731765 | 24.37218862 | 24.04042753 | 24.11949464 | 20.66571052 | 22.06915419 | 24.32717982 | 24.10301561 | 24.18495281 | 24.1586935 |
| Mitochondrial intermediate peptidase | Q99797 | MIPEP | Other | 0.32905938 | 0.23035281 | 24.4994182 | 23.95464646 | 24.2511865 | 23.80232782 | 21.39524498 | 24.98962525 | 24.10197439 | 24.62945357 | 23.70197479 |
| Protein MIS12 homolog | Q9H081 | MIS12 | Other | -0.36962702 | 0.311372322 | 24.68260375 | 23.21083184 | 24.274621 | 23.64185154 | 23.85329052 | 22.93725599 | 24.1052557 | 22.70998744 | 23.37610313 |
| Mitotic interactor and substrate of PL | Q8IVT2 | MISP | Extracellular or secreted | 3.450964542 | 1.627826131 | 21.14680247 | 20.90001466 | 21.61362236 | 20.77005597 | 22.26497155 | 22.58666848 | 24.28926132 | 24.30785895 | 26.3968411 |
| MIT domain-containing protein 1 | Q8WV92 | MITD1 | Extracellular or secreted | 0.727386927 | 0.353117899 | 20.97992413 | 23.28329396 | 21.45523002 | 21.81776751 | 20.87781418 | 22.57493993 | 23.82570726 | 21.27147973 | 22.57945866 |
| Antigen KI-67 | P46013;P46013-2 | MKI67 | Other | -0.954607593 | 1.478817654 | 28.38590899 | 28.56793943 | 28.43686407 | 28.25835756 | 28.2127104 | 28.51863347 | 27.34722584 | 27.15131792 | 27.82784042 |
| MKL/myocardin-like protein 2 | Q9ULH7-4;Q9ULH7;Q9ULH7- | MKL2 | Other | 0.60976372 | 1.536910042 | 22.10392807 | 22.11387197 | 20.95887007 | 22.33845893 | 21.81324612 | 22.14388639 | 22.50273567 | 22.64706663 | 22.41561963 |
| Muskelin | Q9UL63;Q9UL63-2 | MKLN1 | Other | 0.648439021 | 0.456298592 | 23.15205331 | 23.86834783 | 23.35675929 | 23.74919183 | 24.5923489 | 22.02550466 | 25.10801103 | 23.68592164 | 23.5234873 |
| Probable E3 ubiquitin-protein ligase m | Q13064 | MKRN3 | Other | 1.013634051 | 1.617833833 | 21.21926847 | 20.41963617 | 21.77100418 | 22.36711716 | 21.86638698 | 21.58652229 | 22.87700772 | 22.13608494 | 22.64277711 |
| Putative E3 ubiquitin-protein ligase m | Q13434 | MKRN4P | Other | -2.691979302 | 3.041233292 | 24.59428678 | 24.68586819 | 24.75948919 | 24.86159621 | 25.02309916 | 22.31155881 | 21.69813475 | 21.22681749 | 22.11705902 |
| Malectin | Q14165 | MLEC | Extracellular or secreted | -1.031834916 | 0.762649806 | 22.96511491 | 23.0664959 | 22.51113493 | 24.50415512 | 24.2128959 | 25.85436107 | 23.27056339 | 23.24862657 | 21.94238421 |
| Myeloid leukemia factor 2 | Q15773 | MLF2 | Other | 1.419131814 | 1.371108988 | 22.11718143 | 21.46970938 | 21.79217388 | 21.54561874 | 21.89026279 | 23.19332502 | 23.74437481 | 23.88306312 | 22.63409314 |
| DNA mismatch repair protein Mlh1 | P40692-2;P40692-3;P40692 | MLH1 | Other | -0.299110081 | 0.771197827 | 24.13414301 | 24.98119934 | 24.61941312 | 24.21563845 | 24.69853469 | 24.86773535 | 24.03724665 | 24.34523034 | 24.47852475 |
| Protein ENL | Q03111 | MLLT1 | Other | 0.478730436 | 0.919273986 | 23.17674475 | 23.11791065 | 23.29473965 | 22.22878934 | 22.61662532 | 22.76872467 | 23.62131501 | 22.97095266 | 23.44569083 |
| Afadin | P55196;P55196-5;P55196-6 | MLLT4 | Extracellular or secreted | 0.652838544 | 0.906660666 | 23.01896352 | 22.9555513 | 23.02345455 | 23.97009255 | 22.86486788 | 23.11033101 | 24.35110746 | 23.65075201 | 23.42828656 |
| Afadin | P55196-1;P55196-2 | MLLT4 | Extracellular or secreted | 0.196749305 | 0.69124049 | 29.01974728 | 29.12368874 | 29.14525607 | 28.95999959 | 28.89044024 | 28.41155088 | 29.23920201 | 29.17231221 | 28.9540751 |
| Target of rapamycin complex subunit | Q9BVC4-4;Q9BVC4;Q9BVC4-3 | MLST8 | Other | 0.697396461 | 1.081032975 | 23.9693022 | 23.98154777 | 23.61739662 | 24.05092475 | 24.08237192 | 22.1339933 | 24.1967804 | 24.63261853 | 24.18055873 |
| Matrix metalloproteinase-15 | P51511 | MMP15 | Extracellular or secreted | -0.157956271 | 0.13438831 | 25.58806494 | 25.72327847 | 25.34948858 | 24.93210948 | 24.15189852 | 25.46824319 | 24.69938225 | 24.66779459 | 25.76549594 |
| 72 kDa type IV collagenase;PEX | P08253-2;P08253-3;P08253 | MMP2 | Extracellular or secreted | 0.77725606 | 0.281861236 | 21.00037745 | 22.78680085 | 23.74139457 | 23.02820151 | 24.34516265 | 29.10439608 | 24.86537723 | 25.04389339 | 24.42566411 |
| Matrilysin | P09237 | MMP7 | Extracellular or secreted | 0.074661287 | 0.048983757 | 23.38390043 | 22.46946098 | 23.19239216 | 22.04346791 | 21.7926108 | 24.92193353 | 22.7521356 | 23.57288338 | 22.80084778 |
| MMS19 nucleotide excision repair pro | Q96T76;Q96T76-8;Q96T76-9 | MMS19 | Other | 0.461925545 | 0.35457759 | 22.81213656 | 23.48333065 | 23.87633333 | 23.38271373 | 24.75669273 | 24.26212697 | 24.99907822 | 24.25212545 | 23.42123994 |
| MOB kinase activator 1B;MOB kinase | Q7L9L4;Q9H8S9;Q7L9L4-2;Q9 | MOB1B;MOB1A | Extracellular or secreted | 0.068136341 | 0.267432253 | 27.28771238 | 27.57845124 | 27.44126441 | 27.33892132 | 27.33227726 | 27.24789536 | 27.58096847 | 27.45622781 | 27.28047373 |
| MOB-like protein phocein | Q9Y3A3;Q9Y3A3-2;Q9Y3A3-3 | MOB4 | Other | 0.013022415 | 0.036049906 | 27.63075926 | 27.50134824 | 27.49701343 | 27.0744925 | 27.10749231 | 26.924337 | 27.37525809 | 27.29928908 | 27.23224144 |
| Adenylyltransferase and sulfurtransfe | O95396 | MOCS3 | Other | 1.378579869 | 1.022676483 | 24.12620725 | 23.88241021 | 23.77152816 | 21.1111971 | 22.47677092 | 24.40353636 | 24.78166787 | 25.43303956 | 23.80685717 |
| Mannosyl-oligosaccharide glucosidase | Q13724-2;Q13724 | MOGS | Extracellular or secreted | 0.640672085 | 0.354118499 | 21.91127499 | 23.65184703 | 23.72286172 | 23.73520893 | 20.90624877 | 26.21397131 | 23.56010889 | 24.11996949 | 24.31264425 |
| Protein MON2 homolog | Q7Z3U7-2;Q7Z3U7-6;Q7Z3U7 | MON2 | Extracellular or secreted | 0.511123906 | 0.52278426 | 26.52032817 | 26.91056497 | 26.99194026 | 26.67006621 | 26.36059863 | 26.57279666 | 27.67357478 | 27.41771978 | 26.45522461 |
| MORC family CW-type zinc finger prot | Q9Y6X9-2;Q9Y6X9 | MORC2 | Other | -1.606031757 | 1.644277842 | 24.02538224 | 23.96622449 | 23.58840813 | 22.17510801 | 21.68616225 | 24.26327379 | 22.11286198 | 21.0401696 | 21.8811526 |
| MORC family CW-type zinc finger prot | Q14149 | MORC3 | Other | 1.408184481 | 2.32400317 | 23.04111301 | 23.18246952 | 23.0197441 | 22.3972854 | 21.47933479 | 23.50445824 | 24.36813021 | 24.3413668 | 23.82725896 |
| Mortality factor 4-like protein 1 | Q9UBU8-2;Q9UBU8;Q9UBU8 | MORF4L1 | Other | -0.757877144 | 2.280081951 | 27.19947885 | 27.25574912 | 27.39917412 | 26.52017828 | 26.69922337 | 26.48951468 | 25.99224289 | 26.20884667 | 26.30693822 |
| Mortality factor 4-like protein 2 | Q15014 | MORF4L2 | Extracellular or secreted | -0.073934553 | 0.103198505 | 25.0248752 | 25.96736841 | 25.63693831 | 26.08841009 | 26.3961715 | 25.301977 | 25.43472646 | 25.57577096 | 25.97556917 |
| Motile sperm domain-containing prot | Q8NHP6;Q8NHP6-2 | MOSPD2 | Other | -0.913188548 | 0.451878722 | 23.08914591 | 23.84028542 | 23.89282145 | 22.53931314 | 22.32676658 | 24.48989744 | 22.29409457 | 21.23275869 | 23.82269608 |
| Putative helicase MOV-10 | Q9HCE1;Q9HCE1-2 | MOV10 | Extracellular or secreted | -0.360514134 | 0.842643214 | 27.23196689 | 27.42726373 | 27.25051803 | 26.53482399 | 26.19580472 | 27.08253435 | 26.58046538 | 26.76205209 | 26.437396 |
| 6.8 kDa mitochondrial proteolipid | P56378;P56378-2 | MP68 | Other | 0.211583171 | 0.100462476 | 22.7050797 | 22.69395991 | 22.6593148 | 20.79220918 | 21.5917192 | 24.24981309 | 21.57046141 | 22.79574481 | 23.61459123 |
| DNA-3-methyladenine glycosylase | P29372-5;P29372-4;P29372-2 | MPG | Other | -0.209424708 | 0.086082565 | 23.89245091 | 23.69551123 | 23.62800759 | 23.61522609 | 21.4327408 | 22.34462702 | 23.62444208 | 23.65534554 | 21.39622008 |
| M-phase phosphoprotein 8 | Q99549;Q99549-2 | MPHOSPH8 | Extracellular or secreted | -1.366516955 | 1.163354361 | 23.59479932 | 24.06406316 | 24.61419805 | 24.30459029 | 24.63167532 | 23.72484022 | 23.23529088 | 23.20371123 | 21.9285302 |
| Mannose-6-phosphate isomerase | P34949;P34949-2 | MPI | Extracellular or secreted | -0.049786552 | 0.034505807 | 28.02263365 | 28.11391326 | 28.00875815 | 27.2300436 | 27.45935832 | 26.11360531 | 27.9256391 | 27.71056558 | 26.68859182 |
| M-phase-specific PLK1-interacting pro | Q8TAP9 | MPLKIP | Other | -0.356002972 | 1.203468116 | 23.77152816 | 24.1084498 | 24.04076195 | 24.06521429 | 24.1027754 | 24.8583271 | 23.81704532 | 23.82259884 | 23.76587527 |
| MAGUK p55 subfamily member 5 | Q8N3R9-2;Q8N3R9 | MPP5 | Extracellular or secreted | -0.308044919 | 0.165868936 | 22.96877506 | 22.94640587 | 22.8899844 | 22.49620182 | 22.84963035 | 20.99910049 | 22.90126245 | 22.74847542 | 21.00117637 |
| MAGUK p55 subfamily member 6 | Q9NZW5 | MPP6 | Extracellular or secreted | 0.981813839 | 2.384209087 | 24.61739662 | 23.96692855 | 24.82609534 | 24.97897608 | 25.48928498 | 24.64746194 | 25.81391816 | 25.75133892 | 25.64325619 |
| Myosin phosphatase Rho-interacting | Q6WCQ1-3;Q6WCQ1;Q6WCQ | MPRIP | Other | -0.778963673 | 0.670977542 | 26.81665479 | 26.81555588 | 26.94223653 | 26.99679619 | 26.9888022 | 26.8606494 | 25.58442788 | 25.80793817 | 26.98109043 |
| 3-mercaptopyruvate sulfurtransferase | P25325;P25325-2 | MPST | Extracellular or secreted | 2.253715809 | 1.861303306 | 24.79610141 | 25.04489401 | 24.60240891 | 24.34238452 | 21.74566861 | 25.53910602 | 26.986851 | 27.0342267 | 25.77535147 |
| Myelin protein zero-like protein 1 | O95297-2;O95297;O95297-4 | MPZL1 | Extracellular or secreted | 1.763805507 | 3.06486114 | 24.25717085 | 23.72951536 | 24.06109888 | 24.54238673 | 24.91392334 | 25.46338667 | 25.96380168 | 26.21232078 | 26.59903497 |
| C-type mannose receptor 2 | Q9UBG0 | MRC2 | Other | -3.131673091 | 3.739987379 | 26.39512564 | 26.32122142 | 26.34820567 | 26.13010201 | 25.75539434 | 27.77108732 | 22.84435254 | 23.36852991 | 23.75266648 |
| Double-strand break repair protein M | P49959;P49959-3;P49959-2 | MRE11A | Other | -0.206253931 | 0.347696019 | 26.90655668 | 27.08375199 | 27.15044659 | 26.81127434 | 26.62533428 | 26.33367676 | 26.89307614 | 26.73815041 | 26.20553197 |
| MRG/MORF4L-binding protein | Q9NV56 | MRGBP | Other | -0.222827367 | 0.589845461 | 26.43077666 | 26.89850655 | 26.6389277 | 26.71489425 | 26.20187324 | 26.56935268 | 26.42321326 | 26.08626518 | 26.549205 |
| Methylthioribose-1-phosphate isomer | Q9BV20;Q9BV20-2 | MRI1 | Other | 0.310596838 | 1.291603653 | 27.50127231 | 27.69245481 | 27.56521774 | 27.44584866 | 27.06773944 | 27.27106231 | 27.71181304 | 27.89792982 | 27.59384529 |
| 39S ribosomal protein L1, mitochondr | Q9BYD6 | MRPL1 | Other | 0.178240623 | 0.163683821 | 25.47299048 | 25.76867925 | 25.57758718 | 26.16641473 | 26.1718829 | 26.50883121 | 26.58175869 | 26.37904859 | 25.40710747 |
| 39S ribosomal protein L12, mitochond | P52815 | MRPL12 | Other | 0.62031352 | 1.728746463 | 27.49235994 | 27.39345723 | 27.39762463 | 27.62233484 | 27.37451207 | 26.76091076 | 27.93246997 | 28.21001803 | 27.7390523 |
| 39S ribosomal protein L19, mitochond | P49406 | MRPL19 | Other | -0.300452813 | 0.163271689 | 19.64762209 | 21.13408033 | 20.83202776 | 22.99499484 | 21.46017488 | 24.39420989 | 21.14339069 | 21.57005951 | 21.61674625 |
| 39S ribosomal protein L37, mitochond | Q9BZE1 | MRPL37 | Other | 0.330233631 | 0.152407945 | 22.06100339 | 21.39231907 | 19.96746281 | 23.16923935 | 21.77722245 | 25.06093402 | 21.67964157 | 23.04436877 | 22.9807811 |
| 39S ribosomal protein L39, mitochond | Q9NYK5;Q9NYK5-2 | MRPL39 | Other | -0.512049166 | 0.803894111 | 22.65129963 | 23.20271034 | 23.22071956 | 21.8049961 | 23.26699466 | 24.05390998 | 22.61335514 | 22.51475065 | 22.43606185 |
| 39S ribosomal protein L4, mitochondr | Q9BYD3;Q9BYD3-2 | MRPL4 | Other | -0.843279268 | 0.323833338 | 21.18095534 | 19.6322391 | 19.81917182 | 21.93501497 | 22.60292619 | 26.19747424 | 20.04313411 | 21.29311237 | 21.81780654 |
| 39S ribosomal protein L44, mitochond | Q9H9J2 | MRPL44 | Other | -0.633547737 | 0.297298509 | 20.39544652 | 20.37909487 | 20.74098301 | 21.99565813 | 22.03318503 | 24.74072573 | 20.31617103 | 20.69667893 | 22.22905349 |
| 39S ribosomal protein L46, mitochond | Q9H2W6 | MRPL46 | Other | 1.400630207 | 1.072655681 | 24.02816778 | 24.16146407 | 21.9915525 | 24.01648323 | 21.09556591 | 25.08966319 | 24.98897552 | 25.21774741 | 24.18661603 |
| 39S ribosomal protein L53, mitochond | Q96EL3 | MRPL53 | Other | -0.75063843 | 1.015029824 | 20.6416555 | 22.10734863 | 21.94892041 | 21.55795369 | 21.98099024 | 22.11283017 | 21.35772596 | 20.42852011 | 21.13668797 |
| 28S ribosomal protein S22, mitochond | P82650 | MRPS22 | Extracellular or secreted | -0.818170136 | 0.504243744 | 22.50064455 | 21.88045289 | 21.12559257 | 23.3524322 | 23.05669063 | 26.01235413 | 22.08592494 | 21.71101366 | 22.71263447 |
| 28S ribosomal protein S25, mitochond | P82663 | MRPS25 | Other | -1.008839326 | 0.669302811 | 20.34754352 | 21.54832072 | 21.90470161 | 21.43588387 | 23.17501742 | 24.53824767 | 20.31694438 | 21.42658966 | 21.70480538 |
| 28S ribosomal protein S27, mitochond | Q92552;Q92552-2 | MRPS27 | Other | 0.02546662 | 0.030678778 | 23.62678294 | 23.9294029 | 23.9255146 | 24.08488752 | 24.35561725 | 25.54049606 | 24.29080859 | 24.14794583 | 24.36899608 |
| 28S ribosomal protein S35, mitochond | P82673 | MRPS35 | Other | 0.205656981 | 0.115815535 | 22.25988805 | 21.0686609 | 22.50334245 | 22.28262824 | 20.99186793 | 24.7971905 | 22.2741802 | 22.07512525 | 23.21945452 |
| 28S ribosomal protein S36, mitochond | P82909 | MRPS36 | Other | 0.162752329 | 0.445254884 | 21.57090934 | 22.12407905 | 21.48700278 | 21.94909858 | 22.04292715 | 22.21400837 | 22.18537361 | 22.1542495 | 21.84264652 |
| 28S ribosomal protein S9, mitochondr | P82933 | MRPS9 | Other | -0.479174591 | 0.148222557 | 22.44777752 | 21.59498908 | 21.33032828 | 21.88254496 | 23.39165581 | 25.45920196 | 20.20458666 | 22.90947124 | 23.50166714 |
| Ribosome-recycling factor, mitochond | Q96E11-3;Q96E11;Q96E11-8 | MRRF | Other | 0.215029996 | 0.248845575 | 25.76531888 | 26.11919777 | 26.16612737 | 25.98383229 | 25.92011693 | 26.62589163 | 26.53225461 | 26.69285384 | 25.71022398 |
| mRNA turnover protein 4 homolog | Q9UKD2 | MRTO4 | Other | 0.54288294 | 3.198506369 | 28.10948636 | 28.23539497 | 28.12991538 | 28.287228 | 28.52432458 | 28.57924284 | 28.89844889 | 28.78431653 | 28.87867947 |
| DNA mismatch repair protein Msh2 | P43246 | MSH2 | Other | 0.141544718 | 0.393699831 | 28.73350322 | 28.7409187 | 28.80808552 | 28.83282769 | 28.81372249 | 29.40594377 | 29.19422722 | 29.09040544 | 28.80750218 |
| DNA mismatch repair protein Msh6 | P52701;P52701-3;P52701-4;P | MSH6 | Extracellular or secreted | 0.130453276 | 0.208300146 | 27.14014435 | 27.05548303 | 27.36277364 | 27.67249613 | 27.56245451 | 28.61757878 | 27.65819546 | 27.5588106 | 27.87981898 |
| RNA-binding protein Musashi homolo | Q96DH6-2;Q96DH6 | MSI2 | Other | 0.97477569 | 0.954468717 | 23.21834624 | 21.65186289 | 21.24534177 | 20.0719795 | 21.72256302 | 23.31506551 | 22.83373865 | 22.72951536 | 22.97365253 |
| Mesothelin;Megakaryocyte-potentiat | Q13421-4;Q13421-3;Q13421 | MSLN | Extracellular or secreted | 2.290554717 | 0.99154523 | 22.4228573 | 22.69538379 | 22.79689356 | 23.77936509 | 24.06142855 | 26.56531944 | 25.01788529 | 25.22721814 | 27.78718459 |
| Moesin | P26038 | MSN | Extracellular or secreted | 0.238575139 | 1.022029501 | 33.30999891 | 33.34347672 | 33.27932833 | 32.95497641 | 32.93726329 | 32.63146631 | 33.34744235 | 33.25743615 | 33.3391019 |
| Mitochondrial peptide methionine su | Q9UJ68-2;Q9UJ68-3;Q9UJ68 | MSRA | Extracellular or secreted | 0.709372617 | 2.572868023 | 24.2857386 | 23.79124001 | 24.10229485 | 24.52450393 | 24.43695142 | 23.7194189 | 24.81577573 | 24.9378217 | 24.80459427 |
| Macrophage-stimulating protein rece | Q04912-2;Q04912;Q04912-7 | MST1R | Extracellular or secreted | 2.608982453 | 2.409959526 | 23.26255713 | 20.73229751 | 23.14982277 | 23.24250484 | 23.60331484 | 24.85219535 | 25.79761107 | 25.32185792 | 26.12882459 |
| Metallothionein-1X | P80297 | MT1X | Other | -2.136814261 | 0.511989959 | 27.57765921 | 27.18347698 | 27.25880682 | 27.27479871 | 26.97792866 | 25.15390946 | 26.33660789 | 26.36130168 | 21.60493756 |
| Metallothionein-2;Metallothionein-1G | P02795;P13640-2;P13640 | MT2A;MT1G | Other | -1.062154235 | 2.029016791 | 24.90122562 | 24.88702087 | 25.30344103 | 24.65135438 | 25.03825189 | 23.75887951 | 23.98633024 | 23.75796452 | 23.33932919 |
| Metastasis-associated protein MTA1 | Q13330;Q13330-3;Q13330-2 | MTA1 | Other | -1.242354173 | 3.04997796 | 28.05667203 | 28.01035984 | 28.00362076 | 27.42141653 | 27.46256006 | 27.90558156 | 26.8086993 | 26.54980722 | 26.34453635 |
| Metastasis-associated protein MTA2 | O94776;O94776-2 | MTA2 | Other | -0.758272899 | 2.265594509 | 28.5403039 | 28.40397511 | 28.37629361 | 27.77379318 | 27.91433403 | 28.6075256 | 27.31682703 | 27.48684791 | 27.72961908 |
| Metastasis-associated protein MTA3 | Q9BTC8-2;Q9BTC8 | MTA3 | Other | -1.490375219 | 4.15268659 | 24.98463645 | 25.2556951 | 25.15641921 | 24.96802794 | 25.18725812 | 24.23447248 | 23.38297752 | 23.60512499 | 23.43402648 |
| S-methyl-5-thioadenosine phosphoryl | Q13126;Q13126-4;Q13126-3 | MTAP | Extracellular or secreted | 0.055371492 | 0.082040522 | 30.83062333 | 30.79884731 | 30.74101517 | 30.01145331 | 29.99738858 | 30.07813725 | 30.51314578 | 30.73739304 | 30.14430812 |
| Mitochondrial carrier homolog 2 | Q9Y6C9 | MTCH2 | Extracellular or secreted | -0.929582549 | 0.230045715 | 22.25179328 | 23.01389647 | 24.06915406 | 21.96465674 | 22.3950864 | 29.16583516 | 21.98262739 | 21.44839059 | 25.21044543 |
| Cytochrome c oxidase subunit 2 | P00403 | MT-CO2 | Other | -0.572709376 | 0.264941843 | 24.43536251 | 24.30653847 | 24.11569014 | 24.41086381 | 24.06529648 | 28.02548785 | 23.44075767 | 23.99196188 | 25.52877196 |
| Protein LYRIC | Q86UE4 | MTDH | Extracellular or secreted | -0.722917072 | 3.713856805 | 30.09471504 | 30.13833876 | 30.16266972 | 30.16985388 | 30.13148625 | 29.62757978 | 29.28978019 | 29.31094694 | 29.39284336 |
| C-1-tetrahydrofolate synthase, cytopl | P11586 | MTHFD1 | Extracellular or secreted | 0.367378463 | 2.244667716 | 29.95698501 | 29.89339444 | 29.96182187 | 29.81773763 | 29.82470322 | 30.33127374 | 30.37751455 | 30.38966754 | 30.22791125 |
| Monofunctional C1-tetrahydrofolate s | Q6UB35 | MTHFD1L | Other | 1.673536922 | 0.888197843 | 20.59039252 | 20.63539973 | 21.50166357 | 21.96967932 | 22.5174051 | 26.09667655 | 22.96209888 | 24.68586819 | 24.0282521 |
| Bifunctional methylenetetrahydrofola | P13995;P13995-2 | MTHFD2 | Extracellular or secreted | 0.669258436 | 1.316428278 | 26.41335186 | 26.60158741 | 26.67828442 | 27.62644876 | 27.59391651 | 26.91934419 | 27.84619288 | 27.78743372 | 27.29061528 |
| 5-formyltetrahydrofolate cyclo-ligase | P49914-2;P49914 | MTHFS | Other | 0.897956176 | 0.866031883 | 23.91952605 | 23.8120582 | 24.14429353 | 23.43987044 | 21.77189391 | 24.22191924 | 24.57045333 | 25.00727233 | 23.77092355 |
| Myotubularin | Q13496;Q13496-2 | MTM1 | Extracellular or secreted | 0.02299886 | 0.019248181 | 21.02452017 | 22.50011149 | 22.73131903 | 22.94719102 | 24.04668507 | 22.38574446 | 22.51277518 | 22.652438 | 22.72156902 |
| Myotubularin-related protein 1 | Q13613-2;Q13613 | MTMR1 | Extracellular or secreted | 0.418386473 | 1.127888868 | 22.88373437 | 23.04486901 | 23.22423254 | 23.02681789 | 22.07564758 | 23.11197108 | 23.47852475 | 23.34177397 | 23.11849693 |
| Myotubularin-related protein 12 | Q9C0I1-3;Q9C0I1-2;Q9C0I1 | MTMR12 | Other | -0.180574159 | 0.222692816 | 23.63644053 | 23.82259884 | 23.84143766 | 22.43999722 | 22.42100875 | 23.90756553 | 23.44707919 | 23.10376803 | 22.94195457 |
| Myotubularin-related protein 6 | Q9Y217;Q9Y217-2 | MTMR6 | Other | 0.275641484 | 0.177671094 | 23.23861037 | 20.23094503 | 19.24631362 | 21.8718481 | 21.5786861 | 22.31869367 | 21.60724903 | 21.5666264 | 21.89559747 |
| Myotubularin-related protein 9 | Q96QG7-2;Q96QG7 | MTMR9 | Other | 0.430266262 | 0.486344315 | 23.91451652 | 24.15607196 | 23.20123014 | 21.73154367 | 23.31658538 | 23.98311469 | 23.48615742 | 24.09756088 | 23.85861167 |
| Myotrophin | P58546 | MTPN | Extracellular or secreted | 0.639928476 | 1.106813115 | 26.04136789 | 25.60893586 | 25.99744242 | 26.53577336 | 26.86076779 | 25.80508651 | 27.21808013 | 26.54794097 | 26.57845124 |
| Methionine synthase | Q99707;Q99707-2 | MTR | Other | 1.251434969 | 0.346503252 | 24.64537434 | 25.10557543 | 25.12911948 | 26.4650852 | 25.77356684 | 22.94701261 | 27.86502318 | 27.28559751 | 23.63655117 |

| | | | | | | | | | | | | | | |
|---|---|---|---|---|---|---|---|---|---|---|---|---|---|---|
| Metaxin-2 | O75431;O75431-2 | MTX2 | Extracellular or secreted | 0.509141543 | 0.639555096 | 20.94840174 | 20.97232817 | 22.3034898 | 22.60777763 | 22.40171494 | 23.16123339 | 22.54597218 | 22.68093121 | 22.49799408 |
| Mucin-1;Mucin-1 subunit alpha;Mucir | P15941-7;P15941-17;P15941 | MUC1 | Extracellular or secreted | 1.013360619 | 0.415761248 | 25.1655333 | 25.40200782 | 25.64449445 | 20.12438382 | 21.99693704 | 24.28220725 | 25.33231141 | 25.43609363 | 23.57945866 |
| Methylmalonyl-CoA mutase, mitochor | P22033 | MUT | Other | 1.665087041 | 1.72544736 | 23.1523271 | 21.47313323 | 21.91546943 | 21.32613716 | 22.485381 | 24.48842711 | 24.11505508 | 24.2278057 | 24.07283785 |
| Multivesicular body subunit 12A | Q96EY5;Q96EY5-2;Q96EY5-3 | MVB12A | Extracellular or secreted | -0.424936531 | 0.171392036 | 23.80025533 | 23.26813761 | 23.73675782 | 24.29670116 | 24.29600093 | 24.67362869 | 21.86823229 | 24.51044692 | 24.38225197 |
| Diphosphomevalonate decarboxylase | P53602 | MVD | Other | 2.302822829 | 1.973252355 | 27.24354388 | 27.38319182 | 27.2843624 | 27.18480151 | 27.33526236 | 27.30990678 | 29.39415264 | 29.2985901 | 30.08626012 |
| Mevalonate kinase | Q03426 | MVK | Extracellular or secreted | 3.815099069 | 2.819076221 | 24.10117294 | 21.21687958 | 23.66719905 | 22.83276298 | 19.98472988 | 21.404406 | 25.46235723 | 26.76825016 | 25.81826503 |
| Major vault protein | Q14764 | MVP | Extracellular or secreted | 0.390313399 | 0.536442876 | 31.77482228 | 31.78916086 | 31.83171838 | 30.23180672 | 30.25191785 | 30.52859686 | 31.40319298 | 31.51976786 | 31.45199082 |
| Matrix-remodeling-associated protein | P84157-2;P84157-3;P84157 | MXRA7 | Extracellular or secreted | -0.746380933 | 1.635405855 | 24.05953195 | 23.63666179 | 24.26577927 | 24.59263405 | 24.90476654 | 23.39584475 | 23.36212148 | 23.46824319 | 23.35810171 |
| Myeloid-associated differentiation ma | Q96S97 | MYADM | Extracellular or secreted | 1.054707933 | 1.608824418 | 26.16729563 | 26.22776899 | 26.76610282 | 26.36329181 | 26.42279636 | 27.26382895 | 27.22857652 | 27.47961981 | 28.06146975 |
| Myb-binding protein 1A | Q9BQG0;Q9BQG0-2 | MYBBP1A | Other | -0.71086058 | 0.553713913 | 25.25241424 | 25.9330781 | 25.86024682 | 26.26056655 | 26.18307938 | 27.95160399 | 25.57048229 | 24.73169192 | 26.2857386 |
| C-Myc-binding protein | Q99417 | MYCBP | Other | 1.938778078 | 3.12374262 | 24.90912327 | 25.08005532 | 25.34377426 | 24.34428236 | 25.39506025 | 23.7858635 | 27.03674376 | 26.34695543 | 26.86171452 |
| Myeloid-derived growth factor | Q969H8 | MYDGF | Extracellular or secreted | -0.301167934 | 0.349555849 | 28.50324538 | 28.17868909 | 28.11207451 | 29.11383876 | 29.51943982 | 27.54200288 | 28.44730782 | 28.39929638 | 27.73453722 |
| Myelin expression factor 2 | Q9P2K5-4;Q9P2K5-2;Q9P2K5 | MYEF2 | Other | -0.42793172 | 1.305564887 | 23.87229913 | 23.89965932 | 24.3937518 | 24.17714029 | 24.10533564 | 24.8916632 | 23.98006634 | 23.70271466 | 23.70334853 |
| Myosin-10 | P35580;P35580-5;P35580-2;P | MYH10 | Extracellular or secreted | -0.677104989 | 1.34105365 | 29.54323893 | 29.4969563 | 29.5458183 | 28.32422945 | 28.40774771 | 29.33415433 | 28.58265615 | 28.57390913 | 28.13819226 |
| Myosin-14 | Q7Z406-6;Q7Z406-2;Q7Z406; | MYH14 | Extracellular or secreted | -0.144634243 | 0.304885123 | 29.85796835 | 29.90626995 | 29.9608558 | 29.29797822 | 29.1815116 | 29.86627672 | 29.3155536 | 29.48908971 | 29.79688428 |
| Myosin-9 | P35579;P35579-2 | MYH9 | Extracellular or secreted | -0.333453666 | 0.926311935 | 33.57880212 | 33.55773389 | 33.62036525 | 32.84204266 | 32.92707075 | 33.38097518 | 32.85632541 | 32.87334981 | 33.22345871 |
| Myosin regulatory light chain 12A;My | P19105;O14950;P24844 | MYL12A;MYL12B | Extracellular or secreted | 0.613848253 | 0.976767963 | 29.91099376 | 30.10086727 | 30.17152504 | 29.70046414 | 29.76312917 | 30.07775536 | 30.43061717 | 30.2521435 | 31.02115146 |
| Myosin light polypeptide 6 | P60660-2;P60660 | MYL6 | Extracellular or secreted | -0.147014858 | 0.084092842 | 30.42902127 | 30.21962715 | 30.44065076 | 29.45552251 | 29.68021429 | 28.15131792 | 29.80605818 | 30.32742398 | 28.61365021 |
| Myosin light chain kinase, smooth mu | Q15746-9;Q15746-11;Q1574 | MYLK | Extracellular or secreted | -0.305925188 | 0.753592482 | 25.72473616 | 26.11729639 | 25.95805881 | 26.23213163 | 26.2054201 | 24.98910549 | 25.51285955 | 25.65212066 | 25.53061851 |
| Unconventional myosin-XVIIIa | Q92614-4;Q92614;Q92614-3 | MYO18A | Other | -0.793262756 | 1.923828978 | 29.60192386 | 29.62973495 | 29.57023978 | 28.69786557 | 28.71636698 | 28.57942269 | 28.26033324 | 28.30912617 | 28.44852923 |
| Unconventional myosin-Ib | O43795-2;O43795 | MYO1B | Extracellular or secreted | 0.421165073 | 1.015368795 | 25.58135645 | 26.49408577 | 26.00757175 | 26.05469672 | 26.4798973 | 26.99539505 | 26.5429623 | 26.71502522 | 26.81200922 |
| Unconventional myosin-Ic | O00159-3;O00159;O00159-2 | MYO1C | Extracellular or secreted | 0.736325614 | 2.532145396 | 28.31665442 | 28.40592349 | 28.3889084 | 28.05976916 | 28.2510149 | 29.09541948 | 29.06237077 | 29.19999361 | 29.20545739 |
| Unconventional myosin-Id | O94832 | MYO1D | Extracellular or secreted | 1.633830018 | 2.777090463 | 25.76357245 | 26.00343848 | 26.09311367 | 25.16518825 | 24.74319373 | 26.70028223 | 27.05889231 | 27.46699948 | 27.60999265 |
| Unconventional myosin-Ie | Q12965 | MYO1E | Extracellular or secreted | 1.746346975 | 1.325913628 | 22.21871576 | 23.95748327 | 23.13207292 | 21.65979485 | 21.71918493 | 24.3415704 | 23.84258897 | 24.38337313 | 25.52748989 |
| Unconventional myosin-Vb | Q9ULV0;Q9ULV0-2 | MYO5B | Extracellular or secreted | 0.076758161 | 0.056689355 | 21.51177227 | 22.7807472 | 22.66553023 | 22.05986197 | 22.13345331 | 22.8438352 | 21.65645679 | 22.51431004 | 23.05710775 |
| Unconventional myosin-VI | Q9UM54-6;Q9UM54-5;Q9UM | MYO6 | Extracellular or secreted | -0.59990846 | 1.041326954 | 28.29224011 | 28.31760348 | 28.35274978 | 27.64051448 | 27.68063348 | 28.64632924 | 27.22848478 | 27.45873276 | 27.97809237 |
| Myoferlin | Q9NZM1-6;Q9NZM1;Q9NZM | MYOF | Extracellular or secreted | -0.438565477 | 0.323659131 | 29.69728563 | 29.51284449 | 29.83686531 | 29.49817446 | 29.54447392 | 30.69610847 | 28.76284414 | 28.97858339 | 30.33575217 |
| Mitotic-spindle organizing protein 1 | Q08AG7 | MZT1 | Other | 0.672795652 | 0.891911779 | 21.8915302 | 21.66443472 | 22.12342652 | 21.40154474 | 23.07502859 | 20.722398 | 22.65634979 | 22.75594948 | 22.04526907 |
| Protein-lysine N-methyltransferase N6 | Q8WVE0 | N6AMT2 | Extracellular or secreted | 0.708137327 | 0.484754924 | 22.82410701 | 21.2613153 | 22.42875934 | 21.59080814 | 22.73480072 | 21.26132366 | 23.46263806 | 23.03897189 | 21.67335911 |
| N-alpha-acetyltransferase 10 | P41227;P41227-2 | NAA10 | Other | 0.394413708 | 2.867218244 | 28.18470694 | 28.11654306 | 28.16700844 | 28.49910641 | 28.40133247 | 28.16158901 | 28.63481465 | 28.67367586 | 28.63989378 |
| N-alpha-acetyltransferase 11 | Q9BSU3 | NAA11 | Other | 0.046773184 | 0.005497889 | 28.48366124 | 28.56132588 | 28.63256306 | 27.34451942 | 21.56924271 | 21.38340494 | 29.09032971 | 21.39119092 | 27.64615754 |
| N-alpha-acetyltransferase 15, NatA au | Q9BXJ9;Q9BXJ9-4 | NAA15 | Other | 0.273300237 | 1.775787912 | 29.1654758 | 29.29684117 | 29.2124785 | 29.39368225 | 29.5290625 | 29.23345339 | 29.66527879 | 29.60241952 | 29.4676992 |
| N-alpha-acetyltransferase 20 | P61599;P61599-2 | NAA20 | Other | 0.02505378 | 0.030855513 | 24.97469431 | 25.28690204 | 25.29382804 | 25.09345625 | 25.24488699 | 25.01179984 | 25.64721488 | 25.10477597 | 24.77595422 |
| N-alpha-acetyltransferase 25, NatB au | Q14CX7;Q14CX7-2 | NAA25 | Other | 0.285341954 | 0.533108616 | 28.24172689 | 28.29346849 | 28.27546491 | 27.98016442 | 28.01632382 | 27.63547214 | 28.544695 | 28.57499189 | 27.95764931 |
| N-alpha-acetyltransferase 35, NatC au | Q5VZE5 | NAA35 | Other | -0.374948395 | 0.191531945 | 23.05170482 | 23.20167885 | 23.50336671 | 23.64207197 | 22.25168096 | 23.42123994 | 23.55578647 | 23.41157512 | 21.44366486 |
| N-alpha-acetyltransferase 38, NatC au | Q9BRA0;Q9BRA0-2 | NAA38 | Extracellular or secreted | -0.408598685 | 0.498545688 | 22.24733766 | 21.55604384 | 22.1857393 | 22.89786982 | 22.8045259 | 22.53509403 | 22.07368788 | 21.40436824 | 22.40945309 |
| N-alpha-acetyltransferase 50 | Q9GZZ1;Q9GZZ1-2 | NAA50 | Extracellular or secreted | 0.298738116 | 2.171056602 | 27.34604241 | 27.46676617 | 27.32568823 | 27.37310185 | 27.41651227 | 27.74419514 | 27.66458389 | 27.8237167 | 27.74406679 |
| N-acylethanolamine-hydrolyzing acid | Q02083-2;Q02083;Q02083-3 | NAAA | Extracellular or secreted | -0.449326844 | 1.144008862 | 22.50438549 | 22.92223305 | 22.95537392 | 22.73520893 | 22.89803595 | 21.93646541 | 22.31746892 | 22.36876967 | 21.94163225 |
| SOSS complex subunit B1 | Q9BQ15;Q9BQ15-2 | NABP2 | Other | 0.69491214 | 2.211768678 | 23.34881351 | 23.68976513 | 23.58909425 | 22.93874318 | 23.16654499 | 23.95287061 | 24.15977158 | 24.30236058 | 23.9655201 |
| Nascent polypeptide-associated comp | Q13765;E9PAV3-2;E9PAV3 | NACA | Extracellular or secreted | 0.495742436 | 1.403757926 | 29.86118206 | 29.99819598 | 29.93705277 | 30.3878203 | 30.55022937 | 29.60434747 | 30.58324816 | 30.75477656 | 30.31861655 |
| Nucleus accumbens-associated protei | Q96RE7 | NACC1 | Other | 0.059814911 | 0.117971725 | 25.4373325 | 25.51261847 | 25.40509575 | 24.94176656 | 24.70361256 | 25.98089439 | 25.45422071 | 25.39021305 | 25.32567108 |
| NAD kinase | O95544;O95544-2 | NADK | Other | 1.210882001 | 1.275957393 | 24.62939798 | 24.50324538 | 24.59923366 | 22.63721212 | 21.93775197 | 24.13202584 | 24.68725735 | 24.8520048 | 25.31281733 |
| NAD kinase 2, mitochondrial | Q4G0N4-2;Q4G0N4;Q4G0N4 | NADK2 | Other | 0.471028458 | 0.345216721 | 23.96728045 | 21.14763078 | 24.041347 | 24.2250862 | 24.28961312 | 25.16449789 | 24.01060527 | 24.58135645 | 24.23885136 |
| Glutamine-dependent NAD(+) synthet | Q6IA69-2;Q6IA69 | NADSYN1 | Other | -0.028817796 | 0.021061892 | 24.13930528 | 24.31851743 | 24.53913561 | 21.92716271 | 23.77786129 | 25.13907103 | 24.03515013 | 23.70998744 | 24.08893572 |
| NEDD8-activating enzyme E1 regulato | Q13564-2;Q13564;Q13564-4 | NAE1 | Extracellular or secreted | -0.097959545 | 0.230923451 | 30.02458992 | 30.10711812 | 30.13528362 | 30.01464894 | 30.02736116 | 29.71608891 | 30.10099255 | 30.00128685 | 29.61638731 |
| H/ACA ribonucleoprotein complex no | Q96HR8-2;Q96HR8 | NAF1 | Other | -0.647314253 | 0.291770019 | 23.56964241 | 23.61324271 | 24.14148976 | 23.46687505 | 23.84114969 | 23.27113357 | 24.39027866 | 21.57267696 | 23.04686821 |
| Alpha-N-acetylgalactosaminidase | P17050 | NAGA | Extracellular or secreted | -1.184073396 | 3.727592057 | 27.45560089 | 27.39811412 | 27.3711086 | 27.57289783 | 27.72260119 | 26.72351284 | 26.29388064 | 26.14551847 | 26.13029844 |
| N-acetyl-D-glucosamine kinase | Q9UJ70;Q9UJ70-2 | NAGK | Extracellular or secreted | 0.569141272 | 1.603119583 | 26.8003294 | 27.00356715 | 26.86407862 | 26.14084646 | 25.88070659 | 26.64099707 | 27.19168834 | 27.17378996 | 27.00720816 |
| Alpha-N-acetylglucosaminidase;Alpha | P54802 | NAGLU | Extracellular or secreted | 0.281062989 | 0.651033055 | 23.66133884 | 23.836149 | 23.91122812 | 24.39695539 | 24.35366698 | 23.85167128 | 24.04760056 | 24.22088664 | 24.58020657 |
| Nicotinamide phosphoribosyltransfera | P43490 | NAMPT | Extracellular or secreted | 0.184676808 | 0.486470576 | 31.5975262 | 31.82293459 | 31.80301182 | 31.23071934 | 31.17676472 | 30.86665993 | 31.68055804 | 31.67181313 | 31.45046756 |
| Sialic acid synthase | Q9NR45 | NANS | Extracellular or secreted | -0.145768438 | 0.411977896 | 30.59779257 | 30.64091091 | 30.5388064 | 30.11582406 | 30.11321779 | 30.00718142 | 30.25124069 | 30.33255471 | 29.98576586 |
| Nucleosome assembly protein 1-like 1 | P55209-2;P55209 | NAP1L1 | Other | -0.108097176 | 0.120287863 | 31.37477115 | 31.25743615 | 31.30791105 | 31.55114665 | 31.62091142 | 30.56447291 | 31.42006149 | 31.48812252 | 30.60584913 |
| Nucleosome assembly protein 1-like 1 | P55209-3 | NAP1L1 | Other | 1.255435189 | 1.407292957 | 22.96128647 | 23.30702511 | 23.29235421 | 21.86778242 | 23.87586113 | 20.88710476 | 23.95411393 | 23.90370977 | 24.00418891 |
| Nucleosome assembly protein 1-like 4 | Q99733;Q99733-2 | NAP1L4 | Other | 0.322508191 | 1.337340265 | 28.85528231 | 28.73104448 | 28.68885856 | 28.256919 | 28.38964703 | 28.2566941 | 28.92524293 | 28.93685626 | 28.69464812 |
| Alpha-soluble NSF attachment protein | P54920 | NAPA | Extracellular or secreted | 0.392892304 | 1.537034737 | 28.07846813 | 28.07892615 | 28.16312578 | 28.55244818 | 28.50275236 | 28.38997518 | 28.60335729 | 28.57621802 | 28.8819495 |
| Gamma-soluble NSF attachment prote | Q99747;Q99747-2 | NAPG | Extracellular or secreted | 0.46569784 | 2.088464762 | 26.47602304 | 26.53146654 | 26.58362477 | 26.34573794 | 26.73143298 | 26.84720897 | 26.94558904 | 26.99366869 | 27.21558291 |
| Nicotinate phosphoribosyltransferase | Q6XQN6;Q6XQN6-2;Q6XQN6 | NAPRT | Extracellular or secreted | -0.132459336 | 0.490014 | 29.27801582 | 29.29651301 | 29.2811152 | 28.71227236 | 28.74868014 | 28.85507434 | 28.97443175 | 28.83692549 | 28.87710019 |
| Asparagine--tRNA ligase, cytoplasmic | O43776 | NARS | Extracellular or secreted | 0.13239258 | 1.024792173 | 30.31710746 | 30.34804945 | 30.31516487 | 30.34857716 | 30.36650949 | 29.95753862 | 30.38525076 | 30.42382236 | 30.41457815 |
| Probable asparagine--tRNA ligase, mit | Q96I59;Q96I59-2 | NARS2 | Other | 1.020470501 | 0.361961993 | 20.58673719 | 21.81731629 | 21.09384307 | 21.90267982 | 21.58870549 | 26.50184933 | 22.25951456 | 22.59636995 | 24.95109258 |
| Nuclear autoantigenic sperm protein | P49321;P49321-3;P49321-4 | NASP | Other | 0.402871626 | 1.173406689 | 32.47128477 | 32.52288053 | 32.44963154 | 32.58174523 | 32.63116271 | 31.82175683 | 32.99322665 | 32.88767243 | 32.56694661 |
| Arylamine N-acetyltransferase 1 | P18440 | NAT1 | Other | 0.556668305 | 0.431341225 | 21.30709909 | 21.71193877 | 23.44808807 | 24.34014458 | 24.52462349 | 22.6871227 | 23.29684117 | 23.90279021 | 23.47988188 |
| N-acetyltransferase 10 | Q9H0A0-2;Q9H0A0 | NAT10 | Other | -0.838061472 | 0.285520506 | 24.38699443 | 24.14491585 | 24.92718876 | 26.88180367 | 26.67640241 | 28.90210591 | 24.26205526 | 24.16369205 | 27.01977378 |
| Neurobeachin-like protein 2 | Q6ZNJ1-2;Q6ZNJ1-3;Q6ZNJ1 | NBEAL2 | Other | 0.379028523 | 0.573001421 | 24.31112022 | 23.58003401 | 23.88827578 | 23.2093302 | 24.50645723 | 24.59057974 | 23.99524407 | 24.72510035 | 24.45963973 |
| Nibrin | O60934 | NBN | Other | -0.179852371 | 0.158535593 | 24.19625512 | 24.21955792 | 24.1049359 | 23.94975762 | 21.86179682 | 24.523906 | 23.69996465 | 23.28065072 | 23.90793221 |
| Neural cell adhesion molecule 1 | P13591-4;P13591-3;P13591-1 | NCAM1 | Extracellular or secreted | -0.33669655 | 0.535760896 | 24.22339311 | 24.25169217 | 24.36419398 | 23.90380169 | 23.88408852 | 23.14923358 | 23.37318484 | 24.06151095 | 23.44341608 |
| Condensin complex subunit 1 | Q15021 | NCAPD2 | Other | 0.754052802 | 0.959369352 | 27.86702832 | 28.15846238 | 28.0619332 | 28.09164167 | 28.12080012 | 28.32979921 | 29.24163598 | 29.02635885 | 28.30899603 |
| Condensin-2 complex subunit D3 | P42695 | NCAPD3 | Other | -1.662589075 | 0.933052656 | 27.51705696 | 28.2312803 | 30.45218725 | 28.17255066 | 28.07193824 | 27.67397907 | 27.65027267 | 25.3565914 | 27.06486494 |
| Condensin complex subunit 3 | Q9BPX3 | NCAPG | Other | 0.746786669 | 1.261565457 | 26.39204904 | 26.43785631 | 26.80131662 | 26.64814799 | 26.76736635 | 27.04113808 | 27.68972515 | 27.59583806 | 26.998734 |
| Condensin complex subunit 2 | Q15003;Q15003-2 | NCAPH | Other | 0.937541473 | 1.024915822 | 26.49664779 | 26.75072579 | 26.62156654 | 27.02422001 | 27.22296953 | 26.84050154 | 28.20578363 | 27.97547078 | 27.10968562 |
| Nuclear cap-binding protein subunit 1 | Q09161 | NCBP1 | Other | 0.205092676 | 0.666987369 | 28.9584516 | 29.11391326 | 29.06422252 | 28.82853665 | 28.91846298 | 29.57260876 | 29.39229066 | 29.37276983 | 29.07831542 |
| Nuclear cap-binding protein subunit 2 | P52298;P52298-3;P52298-2 | NCBP2 | Other | 0.628293275 | 1.507905879 | 25.28065072 | 25.13918816 | 24.96934612 | 25.15892459 | 25.20612843 | 26.07461499 | 25.89052721 | 25.68829834 | 26.22048077 |
| Neurochondrin | Q9UBB6-2;Q9UBB6;Q9UBB6- | NCDN | Other | 1.215754374 | 1.0051993 | 23.30284862 | 23.00176896 | 23.29221377 | 21.31001398 | 21.6034849 | 22.69338571 | 24.10181413 | 24.31548018 | 22.83182678 |
| Neutrophil cytosol factor 2 | P19878-2;P19878-4;P19878-3 | NCF2 | Extracellular or secreted | -2.034741141 | 0.635208966 | 25.57533819 | 25.26760196 | 25.21830189 | 24.52450393 | 25.6027487 | 24.78511518 | 20.85597444 | 23.56523227 | 24.9613748 |
| Cytoplasmic protein NCK1 | P16333;P16333-2 | NCK1 | Extracellular or secreted | -0.239792949 | 0.451513168 | 25.67801571 | 25.50902761 | 25.36034746 | 25.94681634 | 24.87239308 | 24.69646669 | 25.36289066 | 25.07316485 | 24.87609909 |
| Cytoplasmic protein NCK2 | O43639 | NCK2 | Other | -0.182308657 | 0.557167207 | 24.34218104 | 24.34448555 | 24.0212362 | 24.10181413 | 23.9593419 | 23.99083731 | 23.88585795 | 23.76819967 | 24.17896447 |
| Nck-associated protein 1 | Q9Y2A7;Q9Y2A7-2 | NCKAP1 | Extracellular or secreted | 0.23866002 | 0.475207602 | 29.33653979 | 29.51286332 | 29.4884386 | 28.68153853 | 28.47316859 | 29.32268325 | 29.56230893 | 29.46285255 | 29.09843462 |
| NCK-interacting protein with SH3 dom | Q9NZQ3-3;Q9NZQ3;Q9NZQ3 | NCKIPSD | Other | 0.129015448 | 0.237465874 | 24.85932283 | 25.35806816 | 25.25587515 | 24.93678888 | 24.73075951 | 24.45694843 | 25.29872993 | 25.15038848 | 24.73680943 |
| Nucleolin | P19338 | NCL | Extracellular or secreted | 0.475678019 | 2.459997848 | 34.48579766 | 34.67029419 | 34.63925124 | 34.89102722 | 34.96288208 | 34.33004507 | 35.18334683 | 35.17849469 | 35.05484127 |
| Nuclear receptor coactivator 5 | Q9HCD5 | NCOA5 | Extracellular or secreted | 0.163357237 | 0.457968455 | 25.34607623 | 25.70316368 | 25.65711888 | 25.23859629 | 25.3807007 | 25.94715088 | 25.80131662 | 25.84610918 | 25.47904925 |
| Nuclear receptor corepressor 1 | O75376;O75376-3;O75376-2 | NCOR1 | Other | -1.946445445 | 1.129153197 | 24.81592228 | 27.68163906 | 24.81753333 | 22.18696567 | 22.60467267 | 22.46413892 | 22.41951791 | 21.97802471 | 22.04855701 |
| Neuronal calcium sensor 1 | P62166-2;P62166 | NCS1 | Extracellular or secreted | -0.23437861 | 0.39820087 | 23.13032986 | 22.57110175 | 22.97901971 | 23.18811129 | 23.17657737 | 23.07711561 | 23.06337204 | 22.93590815 | 22.35871177 |
| Nicastrin | Q92542-2;Q92542 | NCSTN | Extracellular or secreted | 0.839493378 | 2.104566409 | 24.46811886 | 24.98150422 | 24.69354523 | 23.95907653 | 24.36719716 | 25.06467995 | 25.37026064 | 25.21515703 | 25.70012345 |
| Kinetochore protein NDC80 homolog | O14777 | NDC80 | Other | -0.079685804 | 0.059712561 | 25.61879726 | 25.33715256 | 25.45998361 | 25.13613971 | 25.64265043 | 25.56917882 | 25.67882168 | 25.93265019 | 24.53142192 |
| Nuclear distribution protein nudE hon | Q9NXR1;Q9NXR1-1 | NDE1 | Other | 0.582101484 | 2.046095522 | 24.32388599 | 24.50863479 | 24.93836027 | 24.92944805 | 24.96626851 | 25.12518263 | 25.32481314 | 25.24524978 | 25.57213166 |
| Melanoma-associated antigen G1 | Q96MG7 | NDNL2 | Other | -0.638955076 | 1.377598843 | 23.37278643 | 23.25695498 | 23.37252077 | 22.984857 | 22.16686665 | 22.2581922 | 22.46774583 | 22.05049304 | 22.27098492 |
| Protein NDRG1 | Q92597;Q92597-3;Q92597-2 | NDRG1 | Extracellular or secreted | -0.56017314 | 1.058301219 | 27.94424897 | 27.82850638 | 27.8588132 | 26.65744603 | 26.62072793 | 27.36411047 | 26.65935291 | 26.71594168 | 27.08111248 |
| Protein NDRG2 | Q9UN36-2;Q9UN36-6;Q9UN3 | NDRG2 | Extracellular or secreted | -1.158769982 | 1.202149469 | 28.02939002 | 28.15378386 | 28.05196198 | 26.75798994 | 27.09476537 | 26.73143298 | 26.74740014 | 26.61610682 | 25.56984518 |
| Protein NDRG3 | Q9UGV2-2;Q9UGV2;Q9UGV2 | NDRG3 | Extracellular or secreted | 0.057609382 | 0.109290798 | 26.68351425 | 26.74791227 | 27.06599489 | 26.86903067 | 26.70556489 | 26.75887951 | 26.87443491 | 27.15701704 | 26.55682444 |
| NADH dehydrogenase [ubiquinone] 1 | O95299;O95299-2 | NDUFA10 | Extracellular or secreted | -0.742564825 | 0.339720543 | 23.46401022 | 21.46996822 | 23.73851123 | 23.72307011 | 23.68067372 | 26.80304263 | 23.86287341 | 21.83410287 | 23.5149673 |
| NADH dehydrogenase [ubiquinone] 1 | O43678 | NDUFA2 | Other | -0.266481611 | 0.238521253 | 24.1931746 | 24.13304561 | 24.27895076 | 24.11449918 | 24.68431716 | 24.50033035 | 24.34698923 | 24.54704372 | 23.25868104 |
| NADH dehydrogenase [ubiquinone] 1 | Q16718;Q16718-2 | NDUFA5 | Other | 0.866256678 | 0.500591265 | 24.19023824 | 23.73840815 | 23.24569951 | 22.9933966 | 21.67269907 | 27.15605267 | 24.44670068 | 24.44758372 | 25.20273276 |
| NADH dehydrogenase [ubiquinone] 1 | O95182 | NDUFA7 | Other | -0.464424361 | 0.262566386 | 22.61101489 | 23.4910933 | 21.35767551 | 22.68767383 | 22.8725058 | 26.38457575 | 23.21356349 | 22.52049304 | 22.57493993 |
| NADH dehydrogenase [ubiquinone] 1 | P51970 | NDUFA8 | Other | 0.329779941 | 0.925246107 | 27.39304802 | 27.36059863 | 27.32465866 | 27.36260644 | 27.30024963 | 27.40178003 | 27.86254239 | 27.7568454 | 27.44142273 |
| NADH dehydrogenase [ubiquinone] 1 | Q16795 | NDUFA9 | Other | -0.942723567 | 0.391018598 | 21.16714244 | 20.35820905 | 20.36943547 | 20.25707012 | 21.56718368 | 26.62533428 | 21.32231437 | 20.19219648 | 20.82950597 |
| Acyl carrier protein, mitochondrial | O14561 | NDUFAB1 | Other | -0.336150687 | 0.298016068 | 26.43760236 | 27.11018363 | 27.23214993 | 26.17016438 | 26.83569135 | 25.52127214 | 26.4146104 | 26.74201167 | 25.48845776 |
| NADH dehydrogenase [ubiquinone] 1 | O43674-2;O43674 | NDUFB5 | Extracellular or secreted | -7.476940377 | 2.814917159 | 29.54342332 | 29.64072133 | 29.76971672 | 29.30136219 | 29.4165928 | 24.14211329 | 22.99979725 | 20.01310947 | 20.46323698 |
| NADH dehydrogenase [ubiquinone] 1 | O95298-2;O95298 | NDUFC2 | Other | -0.696535919 | 0.356541262 | 21.18168125 | 20.27849892 | 21.38676621 | 22.40142868 | 22.52376376 | 24.79827877 | 20.32289638 | 22.17280354 | 21.69990113 |
| NADH-ubiquinone oxidoreductase 75 | P28331-4;P28331-5;P28331;P | NDUFS1 | Other | -0.174290408 | 0.223559865 | 25.53341363 | 25.22073906 | 25.36872971 | 26.31101625 | 26.03290348 | 27.12099782 | 25.96258875 | 25.74699029 | 25.56144971 |
| NADH dehydrogenase [ubiquinone] ir | O75489 | NDUFS3 | Other | -1.509435545 | 0.403541165 | 23.19315956 | 22.89469122 | 21.98770119 | 20.80427915 | 24.54874949 | 26.69005831 | 19.22587866 | 22.78680085 | 23.51833331 |
| NADH dehydrogenase [ubiquinone] ir | O43920 | NDUFS5 | Other | -0.908994362 | 1.499739232 | 24.48129934 | 24.67648311 | 24.29817055 | 25.99416523 | 25.91677296 | 25.05080023 | 24.47010673 | 24.06529648 | 23.94645942 |
| NADH dehydrogenase [ubiquinone] ir | O75380 | NDUFS6 | Other | 1.352866964 | 0.990358845 | 21.59903975 | 23.74847542 | 24.12242044 | 24.44094772 | 23.00849455 | 25.5266246 | 25.24416114 | 25.79800678 | 24.23943421 |
| NADH dehydrogenase [ubiquinone] fl | P49821-2;P49821 | NDUFV1 | Extracellular or secreted | -2.94645729 | 3.927885747 | 27.77762304 | 27.16154095 | 27.67215889 | 26.52019327 | 26.0078925 | 27.85186187 | 24.52557959 | 23.79074303 | 24.33994077 |
| NADH dehydrogenase [ubiquinone] fl | P19404 | NDUFV2 | Other | -0.424262132 | 0.421154546 | 25.40454366 | 25.42207452 | 25.59647801 | 26.93111766 | 26.71004 | 25.31299039 | 25.63184181 | 25.94246028 | 24.84153364 |
| Nebulette | O76041-2 | NEBL | Extracellular or secreted | 1.174289374 | 1.287972516 | 23.78456616 | 23.99455371 | 23.86966613 | 23.49179439 | 21.79031504 | 22.38490451 | 24.74914067 | 24.67804258 | 23.75358484 |
| Adaptin ear-binding coat-associated p | Q8NC96-2;Q8NC96 | NECAP1 | Extracellular or secreted | 0.492697024 | 2.382903547 | 23.57899821 | 23.60342804 | 23.71565371 | 23.97105794 | 24.06389864 | 23.44088437 | 24.30646894 | 24.26327379 | 24.0953088 |
| Adaptin ear-binding coat-associated p | Q9NVZ3;Q9NVZ3-2;Q9NVZ3- | NECAP2 | Extracellular or secreted | 0.07577051 | 0.058794272 | 23.67217238 | 24.34867845 | 24.4984446 | 25.10880869 | 24.49625159 | 23.87829943 | 24.64943688 | 24.92691741 | 23.65228481 |
| Protein NEDD1 | Q8NHV4;Q8NHV4-3;Q8NHV4 | NEDD1 | Extracellular or secreted | 1.185524875 | 0.948860766 | 20.42298298 | 23.78486565 | 23.2404973 | 23.87642702 | 24.56482539 | 24.041347 | 24.85528825 | 24.50275995 | 24.16399909 |
| E3 ubiquitin-protein ligase NEDD4 | P46934-4 | NEDD4 | Extracellular or secreted | 0.422660884 | 0.156184326 | 24.89795289 | 23.62042031 | 24.70234477 | 23.19067659 | 22.32557392 | 22.70326403 | 24.99334477 | 24.77525098 | 22.21950316 |
| E3 ubiquitin-protein ligase NEDD4-like | Q96PU5-3;Q96PU5-2;Q96PU5 | NEDD4L | Extracellular or secreted | 0.689010618 | 0.492660648 | 23.85671351 | 24.01341946 | 23.88399533 | 24.48652572 | 24.0180976 | 20.41862508 | 24.25781827 | 24.34252017 | 23.80538177 |
| NEDD8 | Q15843 | NEDD8 | Extracellular or secreted | -0.013422586 | 0.008183442 | 26.85150449 | 25.63502923 | 25.79419353 | 25.89866799 | 26.04720531 | 24.52019327 | 26.10433608 | 26.32765954 | 24.90113353 |
| Serine/threonine-protein kinase Nek9 | Q8TD19 | NEK9 | Other | -0.139232429 | 0.244768106 | 27.10908777 | 27.05837627 | 27.07132454 | 27.313025 | 27.28374445 | 26.52103246 | 27.17016438 | 27.04228676 | 26.54814682 |
| Negative elongation factor A | Q9H3P2;Q9H3P2-7 | NELFA | Other | -1.018495164 | 0.804293947 | 23.17148591 | 23.18545197 | 23.2642765 | 24.78885292 | 24.09006718 | 23.57623244 | 23.63599792 | 22.15293889 | 22.19376116 |
| Negative elongation factor B | Q8WX92 | NELFB | Other | -0.357933274 | 0.259735396 | 26.31979255 | 26.57945866 | 26.4108153 | 25.57351914 | 25.19053968 | 26.33803721 | 26.42165729 | 25.91615792 | 24.79446624 |
| Negative elongation factor C/D | Q8IXH7-4;Q8IXH7 | NELFCD | Other | 0.064037914 | 0.06120365 | 24.77610487 | 24.25508276 | 24.2882054 | 23.99731319 | 23.93292046 | 25.33973694 | 24.51611022 | 24.99088058 | 23.97980475 |
| Negative elongation factor E | P18615;P18615-3;P18615-4 | NELFE | Other | 0.080014095 | 0.203290118 | 25.758295 | 25.48339217 | 25.63375511 | 25.37842028 | 25.70556489 | 24.74252573 | 25.48977497 | 25.57363471 | 25.5276092 |
| Neudesin | Q9UMX5 | NENF | Extracellular or secreted | 0.755741846 | 0.380482269 | 25.00791388 | 24.81122533 | 24.94427132 | 24.05854143 | 24.78710985 | 23.57876794 | 25.71166865 | 26.29846775 | 23.851004 |
| Neogenin | Q92859-3;Q92859-2;Q92859 | NEO1 | Extracellular or secreted | 0.5193587 | 0.453019196 | 25.5149673 | 25.248945 | 25.41483618 | 26.57068494 | 26.49526059 | 26.40010301 | 26.09085464 | 25.99838969 | 27.29123029 |
| Sialidase-1 | Q99519 | NEU1 | Extracellular or secreted | 2.459668871 | 4.009250803 | 24.83672689 | 24.26820902 | 24.42847826 | 24.91675019 | 24.91761537 | 24.35110746 | 27.24008963 | 26.77152816 | 27.22683241 |
| Merlin | P35240-4;P35240-6;P35240-5 | NF2 | Extracellular or secreted | 0.734595947 | 0.505641045 | 24.97967394 | 24.95943034 | 24.57172672 | 25.33915926 | 22.87144702 | 25.57750075 | 24.69811073 | 25.23143595 | 26.42371018 |
| Nuclear factor of activated T-cells, cyt | Q13469-3;Q13469-4;Q13469 | NFATC2 | Extracellular or secreted | 0.122153484 | 0.11511675 | 24.01631319 | 24.46201392 | 24.15003978 | 25.30263948 | 25.13339844 | 25.11747475 | 24.56313854 | 24.45287062 | 25.44139107 |
| Nuclear factor 1 X-type | Q14938-6;Q14938-4;Q14938 | NFIX | Other | -0.694154169 | 1.385477391 | 24.9122794 | 24.41673775 | 24.59172138 | 24.61543342 | 24.81474947 | 26.13893437 | 24.10797113 | 24.24517723 | 24.30931703 |
| Nuclear factor NF-kappa-B p105 subu | P19838;P19838-2;P19838-3 | NFKB1 | Other | -1.064741119 | 0.286835402 | 24.88063654 | 25.14821744 | 24.63433691 | 24.88259678 | 24.80518493 | 25.0252555 | 25.19163187 | 25.19696796 | 21.10529086 |
| Nuclear factor NF-kappa-B p100 subu | Q00653-4;Q00653;Q00653-3 | NFKB2 | Extracellular or secreted | 2.094247907 | 1.754858008 | 19.88810398 | 23.74765623 | 21.08203173 | 21.881981 | 22.89996887 | 23.4693616 | 24.67044447 | 23.84747184 | 24.24937911 |
| NF-kappa-B inhibitor beta | Q15653-2;Q15653 | NFKBIB | Other | -0.65867139 | 0.14962616 | 24.77545194 | 25.33476811 | 25.32704273 | 25.58457124 | 25.83691947 | 20.99697897 | 25.25396546 | 25.56191579 | 21.1359708 |
| NFU1 iron-sulfur cluster scaffold hom | Q9UMS0-3;Q9UMS0;Q9UMS | NFU1 | Other | 0.112193906 | 0.119281088 | 26.53953501 | 26.58448523 | 26.49993516 | 26.9293126 | 26.94402551 | 25.9868727 | 26.95526749 | 27.03349173 | 26.0899056 |
| Nuclear transcription factor Y subunit | P23511-2;P23511 | NFYA | Other | -0.20724635 | 0.20150819 | 24.67330519 | 23.79491238 | 24.4994182 | 25.04543573 | 24.89485772 | 24.33871734 | 24.08115312 | 23.90756553 | 25.01286558 |
| Nuclear transcription factor Y subunit | P25208 | NFYB | Other | 0.182405066 | 0.885506565 | 24.90944377 | 25.25378517 | 24.73974765 | 24.76703794 | 24.59792751 | 24.98975516 | 24.97197885 | 25.06138734 | 25.14269761 |
| Nuclear transcription factor Y subunit | Q13952-6;Q13952-7;Q13952 | NFYC | Other | -0.213843223 | 0.463715953 | 26.17938218 | 26.35972771 | 26.17264603 | 25.92651028 | 25.77109992 | 25.87218169 | 26.07400243 | 25.92020782 | 25.50503399 |
| Neuroguidin | Q8NEJ9-2;Q8NEJ9 | NGDN | Other | -2.752749165 | 2.806423528 | 22.98182646 | 22.55138064 | 23.22336365 | 23.68656293 | 25.66782166 | 23.4806216 | 20.96091544 | 21.3300284 | 20.24659714 |
| Tumor necrosis factor receptor super | P08138 | NGFR | Extracellular or secreted | -1.279329297 | 0.789003339 | 25.30657324 | 25.9312304 | 25.99763623 | 23.5653485 | 23.70809372 | 27.48968311 | 23.43834821 | 23.61166784 | 25.11127865 |
| Peptide-N(4)-(N-acetyl-beta-glucosam | Q96IV0-3;Q96IV0-2;Q96IV0;Q | NGLY1 | Extracellular or secreted | -0.367274878 | 0.951212485 | 23.11359339 | 22.99603759 | 22.83124693 | 22.8645557 | 22.96261081 | 22.72267414 | 22.28418587 | 22.57466281 | 22.78468596 |
| Non-homologous end-joining factor 1 | Q9H9Q4;Q9H9Q4-2 | NHEJ1 | Other | -0.142143368 | 0.167112819 | 27.60759615 | 27.40632907 | 27.38327423 | 26.51370302 | 26.45040267 | 26.44303661 | 26.92773131 | 27.17512341 | 26.37288604 |
| NHL repeat-containing protein 2 | Q8NBF2;Q8NBF2-2 | NHLRC2 | Other | 0.1721736 | 0.299374567 | 26.89677566 | 27.18697488 | 27.0857791 | 26.76597641 | 26.77115031 | 26.50289649 | 27.22563786 | 27.26060244 | 26.63505692 |
| NHL repeat-containing protein 3 | Q5JS37 | NHLRC3 | Extracellular or secreted | -0.378904528 | 0.539524521 | 24.36699715 | 24.74529846 | 24.50700193 | 24.74134313 | 24.71444887 | 24.32189232 | 24.11576951 | 24.68094193 | 23.76506591 |
| H/ACA ribonucleoprotein complex sub | Q9NX24 | NHP2 | Other | -1.274785901 | 0.27437795 | 24.52151178 | 24.17577064 | 24.50191006 | 24.26191184 | 24.05258397 | 24.46768365 | 24.39040987 | 19.67992555 | 25.09599284 |
| NHP2-like protein 1;NHP2-like protein | P55769 | NHP2L1 | Other | -0.847148396 | 3.405305242 | 28.72669911 | 28.72689396 | 28.72299197 | 28.24485978 | 28.46248205 | 29.09007728 | 27.75525425 | 27.76641881 | 27.92388382 |
| NIF3-like protein 1 | Q9GZT8-2;Q9GZT8;Q9GZT8-3 | NIF3L1 | Other | 0.075721236 | 0.127014552 | 29.29867749 | 29.34534457 | 29.27390997 | 29.3830888 | 29.49604201 | 28.40068123 | 29.48314226 | 29.36638442 | 28.97650906 |
| MKI67 FHA domain-interacting nucleo | Q9BYG3 | NIFK | Other | -0.4483191 | 0.501864277 | 26.73350322 | 26.69418315 | 26.84110168 | 27.51705696 | 26.99085894 | 26.85424216 | 26.98043684 | 26.64787361 | 25.8422053 |
| 60S ribosome subunit biogenesis prot | Q9Y221;Q9Y221-2 | NIP7 | Other | 0.636548332 | 1.2428059 | 25.27213084 | 25.83460684 | 25.81328213 | 26.20340506 | 26.39334266 | 26.88215363 | 26.78867871 | 26.93280785 | 26.38761901 |
| Protein NipSnap homolog 1 | Q9BPW8 | NIPSNAP1 | Extracellular or secreted | -2.609173854 | 1.11127313 | 24.13508298 | 23.88557871 | 24.31319804 | 24.18412047 | 24.11973208 | 27.07152914 | 20.97370039 | 21.25482333 | 23.79857543 |
| Protein NipSnap homolog 3A | Q9UFN0 | NIPSNAP3A | Other | -0.735266556 | 2.165809761 | 29.54449234 | 29.74784827 | 29.69114056 | 29.36778039 | 29.26470155 | 28.73747363 | 28.81741744 | 28.72487925 | 28.42862201 |
| Nischarin | Q9Y2I1-4;Q9Y2I1-3;Q9Y2I1 | NISCH | Extracellular or secreted | 0.066552869 | 0.062826463 | 23.53919479 | 23.60036848 | 23.91360383 | 23.3413668 | 23.70977715 | 21.7465088 | 23.28559751 | 23.75521604 | 23.08425498 |
| Nitrilase homolog 1 | Q86X76-2;Q86X76-4;Q86X76 | NIT1 | Extracellular or secreted | -0.324583449 | 1.024813049 | 28.3726453 | 28.12962065 | 28.08648792 | 27.66526185 | 27.88663721 | 27.90535202 | 27.5744867 | 27.92297704 | 27.55178839 |

| | | | | | | | | | | | | | | |
|---|---|---|---|---|---|---|---|---|---|---|---|---|---|---|
| Omega-amidase NIT2 | Q9NQR4 | NIT2 | Extracellular or secreted | -0.050977054 | 0.108386137 | 29.95379766 | 29.81801202 | 29.80604281 | 29.2269013 | 29.25766094 | 29.00784974 | 29.52260464 | 29.53326509 | 29.32633135 |
| Notchless protein homolog 1 | Q9NVX2 | NLE1 | Other | -0.117596673 | 0.270965934 | 27.6480794 | 27.66648137 | 27.5168316 | 27.7619887 | 27.90173199 | 27.68905859 | 27.71992802 | 27.75053414 | 27.26883366 |
| Neuroligin-2 | Q8NFZ4 | NLGN2 | Extracellular or secreted | -0.062104275 | 0.056891862 | 23.55602044 | 23.51713207 | 23.06457306 | 23.87661437 | 24.23352196 | 24.150892 | 23.44960006 | 23.25102754 | 24.31243652 |
| Neurolysin, mitochondrial | Q9BYT8 | NLN | Extracellular or secreted | 0.562815438 | 1.21343249 | 27.88925106 | 28.12912931 | 27.98000095 | 27.70708009 | 27.80931282 | 27.32079119 | 28.60431212 | 28.48346905 | 28.01844785 |
| NACHT, LRR and PYD domains-contair | P59047 | NLRP5 | Extracellular or secreted | -2.68699869 | 1.193686298 | 24.31319804 | 24.31125884 | 24.48234613 | 24.96009352 | 24.67319734 | 24.99416523 | 20.46068632 | 22.6184418 | 22.72700536 |
| 60S ribosomal export protein NMD3 | Q96D46 | NMD3 | Other | 0.341710287 | 0.431203291 | 25.68319296 | 25.76192532 | 25.66711782 | 26.03950747 | 26.1091675 | 25.61158342 | 26.50700193 | 26.39234389 | 25.56203229 |
| Nucleoside diphosphate kinase A | P15531;P15531-2 | NME1 | Extracellular or secreted | -0.131291423 | 0.172584268 | 31.21251329 | 30.73586096 | 30.71350197 | 30.78448815 | 29.85297812 | 30.79559991 | 30.198238 | 30.48096821 | 30.97439072 |
| Nucleoside diphosphate kinase B | P22392-2;P22392 | NME2 | Other | 0.089865345 | 0.442904563 | 33.85285166 | 34.15043448 | 34.07495178 | 33.8459537 | 33.85563875 | 33.83798946 | 34.03813151 | 33.90194413 | 34.13843031 |
| Putative nucleoside diphosphate kina | O60361 | NME2P1 | Other | 0.838579778 | 2.22976861 | 25.06796506 | 25.35598684 | 25.35245514 | 24.42783916 | 24.3423167 | 24.70872524 | 25.6517923 | 25.93235733 | 25.55923377 |
| Nucleoside diphosphate kinase 7 | Q9Y5B8-2;Q9Y5B8 | NME7 | Other | -0.201695305 | 0.218298011 | 23.24336229 | 23.06895733 | 22.86593409 | 21.36641996 | 21.82256112 | 22.24295543 | 22.57387736 | 22.28517416 | 21.84095767 |
| N-myc-interactor | Q13287 | NMI | Other | -0.392821419 | 0.39422424 | 25.98328869 | 25.85481285 | 25.79399516 | 24.20493526 | 25.07058777 | 23.44644828 | 24.82094476 | 24.69959406 | 24.47803093 |
| Nicotinamide/nicotinic acid mononuc | Q9HAN9 | NMNAT1 | Other | -0.955336417 | 1.253581765 | 23.47010673 | 23.48123774 | 23.93256008 | 22.87884198 | 23.185759 | 23.40184511 | 22.21698238 | 22.99917287 | 22.09301082 |
| NmrA-like family domain-containing p | Q9HBL8 | NMRAL1 | Other | 0.521481582 | 0.557861723 | 24.66459745 | 24.72291382 | 24.75276855 | 22.2935483 | 23.1204937 | 23.40054281 | 24.26735193 | 24.49161092 | 24.28291421 |
| Glycylpeptide N-tetradecanoyltransfe | P30419;P30419-2 | NMT1 | Extracellular or secreted | 0.149557473 | 0.768473681 | 29.23247019 | 29.25383476 | 29.2957689 | 28.89752597 | 28.86446242 | 28.90589711 | 29.27373215 | 29.27699599 | 29.12292394 |
| Glycylpeptide N-tetradecanoyltransfe | O60551 | NMT2 | Extracellular or secreted | -0.848302534 | 0.734283758 | 23.24605746 | 23.64152084 | 22.16441349 | 20.9908068 | 23.23527627 | 22.70619752 | 21.26606546 | 21.58923634 | 22.59192678 |
| Nicotinamide N-methyltransferase | P40261 | NNMT | Other | 1.754533635 | 2.355709451 | 22.81092457 | 20.516007 | 20.77666357 | 21.89786631 | 22.25325608 | 22.18988575 | 23.65850875 | 23.44151772 | 23.38587609 |
| Nucleolar complex protein 2 homolog | Q9Y3T9 | NOC2L | Other | -0.446787768 | 0.414436424 | 21.15696691 | 21.29684152 | 22.99536237 | 22.80822795 | 21.03611227 | 23.16518058 | 22.10533564 | 21.25776861 | 21.52587824 |
| Nucleolar complex protein 4 homolog | Q9BVI4 | NOC4L | Other | -0.279170852 | 0.155153878 | 20.79233277 | 20.72642108 | 21.22817281 | 20.9643042 | 20.85991994 | 24.4221387 | 20.63520088 | 21.89777134 | 21.12615997 |
| Nucleolar protein 10 | Q9BSC4-2;Q9BSC4-4;Q9BSC4 | NOL10 | Other | -2.617055609 | 1.57834695 | 24.45016665 | 24.58072413 | 24.38317534 | 23.91570216 | 23.60093555 | 23.68110284 | 20.91514566 | 21.02685216 | 22.51273869 |
| Nucleolar protein 11 | Q9H8H0 | NOL11 | Other | -0.948762371 | 2.469158777 | 26.84314032 | 27.02305685 | 26.92104822 | 26.58305085 | 26.50585177 | 27.04978292 | 25.80609507 | 26.12036509 | 25.6902182 |
| Nucleolar protein 3 | O60936;O60936-3;O60936-1 | NOL3 | Other | 0.429129603 | 0.69602668 | 26.85661853 | 26.87630991 | 26.9647713 | 26.77793652 | 26.83569135 | 26.07408412 | 27.13408424 | 27.58003401 | 26.76597641 |
| Nucleolar protein 4-like | Q96MY1-2;Q96MY1 | NOL4L | Other | -0.796846745 | 1.362846526 | 23.28305556 | 23.52480281 | 23.40782063 | 23.48959124 | 22.78560412 | 23.95535618 | 22.24382713 | 23.01908233 | 22.56966558 |
| Nucleolar protein 6 | Q9H6R4-4;Q9H6R4;Q9H6R4- | NOL6 | Other | -1.097600943 | 2.452203728 | 24.101734 | 24.59826836 | 24.63128676 | 23.95029175 | 24.27319858 | 25.26377523 | 23.10833813 | 23.35957692 | 23.64855946 |
| Nucleolar protein 7 | Q9UMY1-2;Q9UMY1 | NOL7 | Other | -0.849169959 | 0.405173811 | 26.1843286 | 21.63726613 | 21.46337955 | 22.65414388 | 22.38202763 | 21.63338374 | 21.07041693 | 21.4135391 | 22.94579887 |
| Nucleolar and coiled-body phosphopr | Q14978-2;Q14978;Q14978-3 | NOLC1 | Other | -0.837543779 | 1.680595816 | 28.79110336 | 28.70441096 | 28.88861255 | 29.64754771 | 29.56378225 | 28.53129549 | 27.8588132 | 28.18796599 | 28.50396564 |
| Nodal modulator 1;Nodal modulator 2 | Q15155;Q5JPE7-2;P69849;Q5 | NOMO1;NOMO2 | Other | 0.128494328 | 0.074002753 | 23.76395229 | 24.50112042 | 24.64790105 | 25.07422706 | 25.52151178 | 26.57816328 | 24.42540801 | 24.96710451 | 26.03640841 |
| Non-POU domain-containing octamer | Q15233;Q15233-2 | NONO | Other | 1.005358111 | 1.80057409 | 28.34747931 | 28.22006556 | 28.18565235 | 29.22540803 | 29.25178245 | 29.83969094 | 29.73254934 | 29.68904192 | 30.1295224 |
| H/ACA ribonucleoprotein complex sul | Q9NPE3 | NOP10 | Other | 1.007020327 | 1.425700623 | 24.10900804 | 23.91861652 | 23.74088011 | 23.71680523 | 24.2680662 | 26.04149322 | 25.318414 | 25.22427671 | 25.37580494 |
| Nucleolar protein 14 | P78316-2;P78316 | NOP14 | Other | 0.202950699 | 0.2328326 | 22.04851867 | 22.1537858 | 21.20430845 | 22.60240891 | 22.78634222 | 23.74129169 | 22.59343216 | 22.58247096 | 22.70127685 |
| Nucleolar protein 16 | Q9Y3C1;Q9Y3C1-3;Q9Y3C1-2 | NOP16 | Other | 1.593062721 | 1.381284774 | 21.68588785 | 23.2292394 | 23.84258897 | 24.314789 | 24.43784044 | 25.1063345 | 25.60351293 | 25.90030447 | 24.58371084 |
| Probable 28S rRNA (cytosine(4447)-C( | P46087;P46087-4;P46087-2;P | NOP2 | Other | -0.445320465 | 0.887246194 | 27.00056176 | 27.10269532 | 26.94570066 | 27.90873399 | 28.00913204 | 28.31527286 | 27.18290895 | 27.02178687 | 27.10039109 |
| Nucleolar protein 56 | O00567 | NOP56 | Other | -1.482799333 | 4.142636425 | 28.81726494 | 28.71692295 | 28.74765623 | 28.5667788 | 28.42466345 | 29.50865746 | 27.19102939 | 27.32156551 | 27.42997902 |
| Nucleolar protein 58 | Q9Y2X3 | NOP58 | Other | -1.206502847 | 2.679252134 | 28.36394343 | 28.91837767 | 28.88436805 | 28.55709478 | 28.34129892 | 29.71191148 | 27.56005056 | 27.57491973 | 27.63401834 |
| Nitric oxide synthase-interacting prot | Q9Y314 | NOSIP | Other | 0.638482666 | 1.662000703 | 23.23622563 | 23.16792342 | 23.56150798 | 24.04385168 | 23.67992245 | 24.36899608 | 24.52294879 | 24.22088664 | 24.20082618 |
| Neurogenic locus notch homolog prot | P46531 | NOTCH1 | Extracellular or secreted | -0.496038259 | 0.93030326 | 23.50639669 | 23.88874028 | 23.7365514 | 22.93671701 | 23.98489225 | 24.20642657 | 22.8332947 | 23.4732073 | 23.33524532 |
| Neurogenic locus notch homolog prot | Q04721 | NOTCH2 | Extracellular or secreted | -0.093618018 | 0.067020224 | 23.87661437 | 24.44170766 | 23.95180406 | 25.67882168 | 25.16227115 | 24.41402967 | 23.76172246 | 24.60229562 | 25.11775216 |
| Neurogenic locus notch homolog prot | Q9UM47 | NOTCH3 | Extracellular or secreted | -0.15723123 | 0.135247134 | 27.51960854 | 27.43738012 | 27.42766336 | 28.06208765 | 27.99620357 | 29.12684234 | 27.39165581 | 27.47041709 | 28.4511262 |
| Protein NOV homolog | P48745 | NOV | Extracellular or secreted | -0.497847283 | 0.201880523 | 26.93303306 | 26.66193681 | 26.54014129 | 27.90086874 | 28.24947858 | 30.27564251 | 26.4157067 | 26.58405506 | 28.78724688 |
| RNA-binding protein Nova-2 | Q9UNW9 | NOVA2 | Other | 0.164053894 | 0.082514091 | 21.83599486 | 21.95776664 | 21.6100011 | 22.29375789 | 20.2616625 | 22.24472733 | 23.03618096 | 20.94226239 | 21.61567349 |
| Niemann-Pick C1 protein | O15118;O15118-2 | NPC1 | Extracellular or secreted | 0.572865597 | 0.298615721 | 23.11878206 | 23.16226347 | 24.40125922 | 23.5162906 | 23.43173325 | 23.76698741 | 23.53480916 | 23.34840831 | 25.53403733 |
| Epididymal secretory protein E1 | P61916;P61916-2 | NPC2 | Extracellular or secreted | 0.704220424 | 1.051256685 | 30.2657301 | 30.20978569 | 30.13833876 | 30.35562985 | 29.97418555 | 30.21581432 | 30.62531686 | 30.7019847 | 31.36510185 |
| Probable aminopeptidase NPEPL1 | Q8NDH3;Q8NDH3-5;Q8NDH3 | NPEPL1 | Other | -0.556236821 | 1.463043981 | 27.48684791 | 27.48715468 | 27.48223841 | 27.03107419 | 27.27923423 | 26.36334194 | 26.85376642 | 26.49797277 | 26.54449603 |
| Puromycin-sensitive aminopeptidase | P55786;P55786-2 | NPEPPS | Extracellular or secreted | -0.177824574 | 0.419226231 | 32.1266639 | 32.15864295 | 32.11306251 | 32.22308465 | 32.26765665 | 31.48697254 | 32.07278164 | 31.97514274 | 31.60664349 |
| Nuclear protein localization protein 4 | Q8TAT6-2;Q8TAT6 | NPLOC4 | Other | -0.081903243 | 0.147867628 | 27.84511625 | 27.68545385 | 27.53356215 | 27.21863448 | 27.65710525 | 27.24227223 | 27.70859369 | 27.52320311 | 27.11356557 |
| Nucleophosmin | P06748-2;P06748;P06748-3 | NPM1 | Other | 0.138568924 | 0.789648514 | 34.37431772 | 34.36658765 | 34.34049533 | 34.25074673 | 34.39093615 | 34.19527791 | 34.44183824 | 34.58250811 | 34.35054116 |
| Nucleoplasmin-3 | O75607 | NPM3 | Other | 0.391694133 | 0.984739755 | 28.88212447 | 28.90959255 | 28.94315925 | 29.11944022 | 29.17750151 | 28.60844243 | 29.59869786 | 29.32798069 | 29.06853406 |
| Neuroplastin | Q9Y639-1;Q9Y639;Q9Y639-3 | NPTN | Extracellular or secreted | -0.159717503 | 0.382038821 | 28.11083079 | 28.26927966 | 28.36269004 | 28.37335081 | 28.16509238 | 29.14214251 | 28.110333 | 28.44375594 | 28.17845165 |
| NAD(P)H dehydrogenase [quinone] 1 | P15559-3;P15559-2;P15559 | NQO1 | Extracellular or secreted | -0.482931967 | 1.01599695 | 27.80051456 | 28.09843462 | 27.96212536 | 27.14150925 | 27.26391847 | 27.97601734 | 27.26588655 | 27.50415512 | 26.90242222 |
| Ribosyldihydronicotinamide dehydrog | P16083 | NQO2 | Extracellular or secreted | -0.336256969 | 0.274495247 | 28.06640556 | 27.98473422 | 27.91718284 | 28.27155215 | 28.30508628 | 26.43790392 | 27.84762715 | 27.94218059 | 26.69285384 |
| Nuclear receptor 2C2-associated prot | Q86WQ0;Q86WQ0-2 | NR2C2AP | Extracellular or secreted | 0.299231346 | 0.308360006 | 25.73122579 | 25.55537692 | 25.56066286 | 26.88506663 | 26.52273932 | 26.78743372 | 25.92689479 | 26.49106038 | 27.00099148 |
| Nuclear receptor-binding protein | Q9UHY1 | NRBP1 | Other | 0.330817099 | 0.325758582 | 26.57643429 | 26.75378884 | 26.2998131 | 26.70226549 | 26.66220853 | 24.91250784 | 26.84816464 | 27.06722656 | 26.03056914 |
| Nardilysin | O43847;O43847-2 | NRD1 | Other | 0.757443298 | 2.098011447 | 29.40157661 | 29.45314938 | 29.29155966 | 29.34112922 | 29.49449783 | 29.36896695 | 30.2831704 | 30.18577048 | 29.97882884 |
| Nuclear respiratory factor 1 | Q16656;Q16656-4;Q16656-2 | NRF1 | Extracellular or secreted | -0.099440993 | 0.261921535 | 24.12573444 | 24.29607097 | 24.32608271 | 24.67357478 | 24.80906744 | 24.95579958 | 24.52862294 | 24.26856599 | 24.49765306 |
| Neuropilin-1 | O14786-3;O14786-2;O14786 | NRP1 | Extracellular or secreted | 0.769865458 | 0.163000393 | 23.13777417 | 24.06480328 | 23.63931421 | 24.33012416 | 24.90177802 | 25.74719523 | 24.52915933 | 22.55763114 | 28.13330044 |
| Sterol-4-alpha-carboxylate 3-dehydro | Q15738 | NSDHL | Other | 2.478658032 | 1.29066547 | 20.71112153 | 22.64489047 | 22.93142429 | 22.93716617 | 22.3294247 | 25.74334784 | 24.45098467 | 24.83248367 | 26.80119326 |
| Vesicle-fusing ATPase | P46459;P46459-2 | NSF | Extracellular or secreted | 0.149181231 | 0.104714324 | 27.58541681 | 27.76452186 | 27.84481704 | 27.17054645 | 27.34798611 | 27.75659093 | 28.21590687 | 28.18801317 | 26.77856326 |
| NSFL1 cofactor p47 | Q9UNZ2;Q9UNZ2-5;Q9UNZ2- | NSFL1C | Extracellular or secreted | -0.023352022 | 0.033465278 | 30.94761074 | 30.90712998 | 30.86562796 | 31.02933736 | 31.00356715 | 29.9838649 | 30.91741051 | 30.96450967 | 30.4165928 |
| Kinetochore-associated protein NSL1 | Q96IY1;Q96IY1-2 | NSL1 | Other | -0.409772312 | 0.779389757 | 24.78979828 | 24.8110783 | 25.04051114 | 24.28679632 | 24.11996949 | 23.39858714 | 24.14507139 | 23.98875887 | 23.86022314 |
| Protein FAN | Q92636;Q92636-2 | NSMAF | Other | 0.242895122 | 1.200933016 | 25.24524978 | 25.43920466 | 25.27305626 | 24.78276791 | 25.25147548 | 25.1264042 | 25.51307047 | 25.30747683 | 25.46721721 |
| Non-structural maintenance of chrom | Q8WV22 | NSMCE1 | Other | -0.401247382 | 0.23762234 | 23.42764738 | 23.05695517 | 23.51580954 | 22.8302026 | 21.4722818 | 23.41286751 | 23.00766585 | 23.23062269 | 21.41585132 |
| E3 SUMO-protein ligase NSE2 | Q96MF7 | NSMCE2 | Other | -0.453348839 | 0.976361811 | 22.81993192 | 23.36319153 | 23.54108716 | 23.1196846 | 23.89161685 | 23.9413636 | 22.70788315 | 23.26026145 | 23.01024671 |
| Non-structural maintenance of chrom | Q9NXX6;Q9NXX6-2 | NSMCE4A | Other | 0.273538472 | 0.233365437 | 21.8499548 | 23.08723686 | 22.63701573 | 22.20699286 | 22.25865229 | 21.02911195 | 22.98330608 | 21.75107327 | 22.62071831 |
| Nuclear speckle splicing regulatory pr | Q9H0G5 | NSRP1 | Other | -1.180392749 | 1.998168683 | 24.74780986 | 24.29207332 | 23.64427439 | 24.09571122 | 25.2155644 | 23.48480618 | 23.3626566 | 23.15132567 | 22.68495917 |
| tRNA (cytosine(34)-C(5))-methyltransf | Q08J23;Q08J23-2;Q08J23-3 | NSUN2 | Other | 0.271394225 | 0.500695916 | 27.61554568 | 27.35413798 | 27.65628705 | 28.28841665 | 28.2383594 | 28.82310927 | 28.46960226 | 28.25322613 | 28.07928229 |
| 5(3)-deoxyribonucleotidase, cytosolic | Q8TCD5 | NT5C | Extracellular or secreted | 0.291329289 | 0.751856795 | 27.2429082 | 27.33364264 | 27.37724562 | 26.50370032 | 26.5380256 | 26.76698741 | 27.40299993 | 27.2950025 | 27.05724033 |
| Cytosolic purine 5-nucleotidase | P49902;P49902-2 | NT5C2 | Other | -0.125638988 | 0.769542681 | 27.78450376 | 27.86632094 | 27.85661853 | 27.50370032 | 27.62380046 | 27.77843793 | 27.5676131 | 27.54229816 | 27.71986276 |
| 5-nucleotidase domain-containing pro | Q5TFE4;Q5TFE4-2 | NT5DC1 | Other | 0.323017888 | 0.979672298 | 30.28250775 | 30.25248191 | 30.18482515 | 29.66108735 | 29.60794882 | 29.66600724 | 30.40821353 | 30.33426091 | 30.05400834 |
| 5-nucleotidase domain-containing pro | Q86UY8;Q86UY8-2 | NT5DC3 | Extracellular or secreted | 0.150140842 | 0.310010583 | 26.36653033 | 26.34129892 | 26.22929191 | 26.87701241 | 26.70094362 | 27.09254924 | 26.46950134 | 26.82128546 | 26.96344893 |
| Protein N-terminal asparagine amidol | Q96AB6 | NTAN1 | Other | 0.271659468 | 0.237428249 | 24.9983036 | 24.57968883 | 24.73055223 | 22.22467415 | 24.27106231 | 24.59741609 | 24.70144607 | 24.74072573 | 24.0736552 |
| N-terminal Xaa-Pro-Lys N-methyltrans | Q9BV86;Q9BV86-2 | NTMT1 | Other | 0.551459707 | 1.061357676 | 22.76048444 | 22.12641208 | 22.80341222 | 23.35877245 | 22.91890763 | 24.01077599 | 23.51123147 | 23.52695288 | 23.60557718 |
| Netrin-4 | Q9HB63-3;Q9HB63-2;Q9HB6 | NTN4 | Extracellular or secreted | -0.588367871 | 0.270182718 | 23.05074212 | 23.68313941 | 23.67001217 | 25.12028598 | 24.85105167 | 26.51429015 | 23.01469682 | 23.37225505 | 25.29270526 |
| Cancer-related nucleoside-triphospha | Q9BSD7 | NTPCR | Extracellular or secreted | -0.048640192 | 0.028407692 | 22.56581332 | 22.29134272 | 22.66055568 | 21.49180358 | 22.60909413 | 22.89356225 | 21.33679419 | 22.78947357 | 22.98389751 |
| NEDD8 ultimate buster 1 | Q9Y5A7-2;Q9Y5A7 | NUB1 | Other | 0.594814941 | 1.453655098 | 25.15197592 | 23.81068616 | 23.98728483 | 24.69975289 | 24.49460464 | 24.81938625 | 25.04168121 | 25.01000762 | 25.21460134 |
| Cytosolic Fe-S cluster assembly factor | P53384-2;P53384 | NUBP1 | Extracellular or secreted | 0.788528719 | 0.994824947 | 26.77693316 | 26.73156246 | 26.78331761 | 26.62672725 | 26.63159207 | 25.66884972 | 27.73873026 | 27.50854412 | 26.7278029 |
| Cytosolic Fe-S cluster assembly factor | Q9Y5Y2 | NUBP2 | Other | 0.388135574 | 1.489015671 | 24.23191198 | 23.88362252 | 24.22383498 | 24.33932919 | 24.46867823 | 24.71649127 | 24.7678967 | 24.80651304 | 24.52193106 |
| Nucleobindin-1 | Q02818 | NUCB1 | Extracellular or secreted | 0.66451071 | 0.441830031 | 27.12386138 | 27.3672805 | 27.44640095 | 28.51694428 | 28.53682583 | 28.48550499 | 28.00517461 | 28.03779124 | 29.68897525 |
| Nucleobindin-2;Nesfatin-1 | P80303-2;P80303 | NUCB2 | Extracellular or secreted | -0.66996867 | 0.713596556 | 21.81577573 | 21.53965077 | 23.40810795 | 23.20009279 | 23.14960573 | 23.50215293 | 21.8164009 | 21.78400694 | 22.6973791 |
| Nuclear ubiquitous casein and cyclin-c | Q9H1E3;Q9H1E3-2 | NUCKS1 | Other | -0.649708759 | 2.137141058 | 30.25462333 | 30.28217631 | 30.27619737 | 30.66906944 | 30.63105427 | 29.62627468 | 29.48828536 | 29.68065025 | 29.75163581 |
| Nuclear migration protein nudC | Q9Y266 | NUDC | Other | 0.004770941 | 0.008378668 | 30.67438324 | 30.62931807 | 30.74975127 | 30.90784627 | 30.99212941 | 30.17200216 | 30.83815857 | 30.84183653 | 30.39703294 |
| NudC domain-containing protein 1 | Q96RS6;Q96RS6-2;Q96RS6-3 | NUDCD1 | Other | 0.100013533 | 0.120621784 | 29.71202632 | 29.81471586 | 29.76483819 | 29.38595012 | 29.37403523 | 29.26952491 | 29.89174431 | 29.96209777 | 29.10674383 |
| NudC domain-containing protein 2 | Q8WVJ2 | NUDCD2 | Extracellular or secreted | 1.610003993 | 1.936735531 | 25.44909624 | 25.01576043 | 25.09534905 | 24.72150644 | 25.36111759 | 25.29659615 | 26.94837686 | 27.07694034 | 26.27440773 |
| NudC domain-containing protein 3 | Q8IVD9 | NUDCD3 | Other | 0.373010649 | 0.406444129 | 24.28029672 | 23.35528119 | 24.4602649 | 23.30075584 | 23.55695597 | 23.60263544 | 24.74411814 | 24.0293479 | 23.62366094 |
| 7,8-dihydro-8-oxoguanine triphospha | P36639-4;P36639-3;P36639-2 | NUDT1 | Extracellular or secreted | -0.529431332 | 0.84918776 | 28.34435009 | 28.52260464 | 28.358043 | 28.71891614 | 28.84613309 | 27.30765054 | 28.00185055 | 28.07106876 | 27.38763544 |
| Peroxisomal NADH pyrophosphatase | Q9BQG2 | NUDT12 | Other | -0.080527585 | 0.096836322 | 21.60630038 | 21.58536236 | 20.8450612 | 21.82714264 | 22.24138504 | 20.75993575 | 21.53416797 | 21.01113442 | 21.64570854 |
| Uridine diphosphate glucose pyropho | O95848 | NUDT14 | Extracellular or secreted | -0.728271052 | 0.494997631 | 28.04702845 | 27.83358183 | 27.80556628 | 27.6768059 | 27.60731394 | 26.15065963 | 27.58527353 | 27.051236 | 25.73915533 |
| Probable 8-oxo-dGTP diphosphatase N | Q9NV35 | NUDT15 | Other | 1.056623402 | 1.152661997 | 24.17934421 | 25.28940205 | 24.75541981 | 24.73701581 | 24.82546465 | 23.56975828 | 25.7134528 | 26.17580871 | 24.95881111 |
| Protein syndesmos | Q9BRJ7 | NUDT16L1 | Other | 0.181281786 | 0.683188961 | 24.22530689 | 24.34834077 | 24.41163977 | 24.51231706 | 24.23498403 | 24.38844041 | 24.72322638 | 24.37431306 | 24.50682038 |
| Nucleoside diphosphate-linked moiety | A8MXV4 | NUDT19 | Extracellular or secreted | 0.11345546 | 0.187670072 | 27.30678181 | 27.27079505 | 27.30504278 | 27.22131083 | 27.01329166 | 26.98010994 | 27.42694395 | 27.57737108 | 26.88471738 |
| Bis(5-nucleosyl)-tetraphosphatase [as | P50583 | NUDT2 | Other | -0.639392696 | 0.526268571 | 26.50092294 | 26.31629189 | 26.48396101 | 26.16541829 | 26.83653429 | 25.46948582 | 26.22574817 | 25.99677465 | 24.74560622 |
| Cleavage and polyadenylation specific | O43809 | NUDT21 | Other | -0.16675065 | 0.358010434 | 29.47647506 | 29.65647459 | 29.64677556 | 28.74701591 | 28.70421305 | 29.65759935 | 29.27728428 | 29.03165266 | 29.13508787 |
| Diphosphoinositol polyphosphate pho | O95989 | NUDT3 | Extracellular or secreted | 0.13538465 | 0.939795964 | 27.924337 | 28.11570006 | 28.08679159 | 27.80839244 | 27.86614404 | 28.10629456 | 28.09219637 | 28.21572176 | 28.05206567 |
| Diphosphoinositol polyphosphate pho | Q9NZJ9;A0A024RBG1;Q9NZJ9 | NUDT4 | Other | 0.104702342 | 0.184122699 | 23.23918653 | 23.59877948 | 23.58026409 | 23.85614357 | 23.7365514 | 23.69890556 | 23.92460884 | 23.33960104 | 23.90481247 |
| ADP-sugar pyrophosphatase | Q9UKK9 | NUDT5 | Extracellular or secreted | -0.048540959 | 0.097060056 | 31.22181786 | 31.03039023 | 31.0822044 | 31.46441148 | 31.44351884 | 30.26237343 | 30.99530069 | 31.08493059 | 31.02650396 |
| ADP-ribose pyrophosphatase, mitoch | Q9BW91-2;Q9BW91 | NUDT9 | Extracellular or secreted | -1.897089898 | 0.713872639 | 24.87366076 | 24.96450692 | 24.8316142 | 25.91501826 | 25.89580546 | 22.9728466 | 25.0380844 | 21.81046923 | 22.18690277 |
| Kinetochore protein Nuf2 | Q9BZD4 | NUF2 | Other | 0.394300903 | 0.406484813 | 29.71130434 | 29.9411173 | 30.10387106 | 29.08527259 | 29.09416131 | 30.35089677 | 30.50975254 | 30.39468419 | 29.42177766 |
| Nuclear fragile X mental retardation-i | Q7Z417 | NUFIP2 | Other | 1.417775419 | 2.338288735 | 25.15861647 | 25.17436159 | 24.64586905 | 24.72951536 | 24.87211122 | 25.24695365 | 26.75136446 | 26.251638 | 26.16403747 |
| Nuclear mitotic apparatus protein 1 | Q14980-2 | NUMA1 | Extracellular or secreted | -1.9179354 | 4.432442417 | 32.20748935 | 32.168659 | 32.11566906 | 31.22974572 | 31.34931562 | 31.7510292 | 29.80019052 | 29.82341302 | 30.03354424 |
| Nuclear pore complex protein Nup107 | P57740;P57740-3;P57740-2 | NUP107 | Other | 0.182589602 | 0.073533598 | 23.72286172 | 24.08796519 | 23.85033642 | 22.46672573 | 22.64337181 | 27.07856993 | 23.23584596 | 24.00598837 | 25.23084987 |
| Nuclear pore complex protein Nup133 | Q8WUM0 | NUP133 | Other | -1.075937455 | 1.163843123 | 25.08253435 | 23.63632989 | 24.98641705 | 24.3607828 | 24.32402339 | 26.88623019 | 23.69614827 | 23.54674942 | 24.16744877 |
| Nuclear pore complex protein Nup153 | P49790;P49790-3;P49790-2 | NUP153 | Other | -1.259330545 | 5.741071452 | 26.95471301 | 26.75429872 | 26.80833106 | 26.77391891 | 27.02823102 | 27.22719977 | 25.62103548 | 25.74614463 | 25.62817451 |
| Nuclear pore complex protein Nup155 | O75694;O75694-2 | NUP155 | Extracellular or secreted | -1.567057449 | 0.650834117 | 22.73966526 | 23.35299387 | 23.15618 | 23.85310011 | 24.13977366 | 27.5341561 | 20.99204728 | 22.71822766 | 23.97648721 |
| Nuclear pore complex protein Nup160 | Q12769 | NUP160 | Other | -0.030776233 | 0.022027356 | 24.74103446 | 24.81811872 | 24.83855534 | 23.73686102 | 23.74990788 | 26.82493078 | 24.62912001 | 24.64207197 | 24.99118343 |
| Nucleoporin NUP188 homolog | Q5SRE5;Q5SRE5-2 | NUP188 | Other | 0.118404145 | 0.063420095 | 22.55933881 | 23.30131421 | 23.78835512 | 24.75669273 | 22.26719474 | 26.2602435 | 24.08383313 | 23.41956934 | 24.31837952 |
| Nuclear pore complex protein Nup205 | Q92621 | NUP205 | Other | 0.144059718 | 0.157553216 | 25.37049346 | 25.117118 | 25.18737141 | 25.60124735 | 25.3684966 | 27.08253435 | 25.51310059 | 25.65357001 | 26.12913913 |
| Nuclear pore complex protein Nup214 | P35658-2;P35658-4;P35658;P | NUP214 | Extracellular or secreted | -0.680450751 | 1.360450284 | 26.75951459 | 26.71947113 | 26.74393844 | 26.64581409 | 26.48703198 | 26.87981898 | 25.88518303 | 25.83436573 | 26.35689359 |
| Nucleoporin Nup37 | Q8NFH4 | NUP37 | Other | 0.506081855 | 1.772368739 | 27.75015075 | 27.91775193 | 27.77297567 | 28.29140596 | 28.43272902 | 28.2756869 | 28.59718873 | 28.74733611 | 28.39407084 |
| Nucleoporin Nup43 | Q8NFH3;Q8NFH3-2 | NUP43 | Other | 0.093348829 | 0.126067163 | 26.20232175 | 26.28091615 | 26.22787913 | 26.50830232 | 26.52962107 | 26.02491746 | 26.67626788 | 26.59377407 | 25.89698348 |
| Nuclear pore complex protein Nup50 | Q9UKX7-2;Q9UKX7 | NUP50 | Other | 0.151376652 | 1.31085854 | 27.49113686 | 27.35009587 | 27.41417488 | 27.35304436 | 27.48069863 | 27.68739085 | 27.68237603 | 27.60618458 | 27.55384007 |
| Nucleoporin p54 | Q7Z3B4-3;Q7Z3B4;Q7Z3B4-2 | NUP54 | Other | -0.226755963 | 0.603574922 | 26.46396346 | 26.41015215 | 26.57310015 | 26.21447165 | 26.38226846 | 27.25511878 | 26.27854318 | 26.17350407 | 26.5172222 |
| Nuclear pore glycoprotein p62 | P37198 | NUP62 | Other | -0.498146545 | 1.649432775 | 26.39879913 | 25.93526075 | 26.25122263 | 26.06665191 | 26.25243228 | 26.71607255 | 25.55733585 | 25.81198473 | 25.94645942 |
| Nuclear pore complex protein Nup85 | Q9BW27-3;Q9BW27;Q9BW2 | NUP85 | Extracellular or secreted | 0.160799919 | 0.215484616 | 24.31610196 | 25.05631026 | 24.98019712 | 25.44960006 | 25.63017601 | 26.19118003 | 25.66003333 | 25.47830872 | 25.15584042 |
| Nuclear pore complex protein Nup88 | Q99567 | NUP88 | Other | -1.02907506 | 2.088417457 | 26.03426869 | 25.9174788 | 25.95451334 | 25.27202402 | 25.07189734 | 26.7951354 | 24.6916298 | 24.70128742 | 25.04251639 |
| Nuclear pore complex protein Nup93 | Q8N1F7;Q8N1F7-2 | NUP93 | Other | -0.216306797 | 0.793171054 | 27.40365012 | 27.61006308 | 27.54325737 | 27.50157603 | 27.50703218 | 28.24929773 | 27.49686109 | 27.44474345 | 27.31691332 |
| Nuclear pore complex protein Nup98- | P52948-6;P52948-2;P52948-5 | NUP98 | Other | -0.47600483 | 0.326655747 | 26.37376565 | 26.42149679 | 26.19723049 | 26.28586204 | 25.15085328 | 29.01685512 | 25.86374788 | 25.66478731 | 26.766482 |
| Nucleoporin p58/p45 | Q9BVL2-2;Q9BVL2-3;Q9BVL2 | NUPL1 | Other | -0.18612161 | 0.156681107 | 24.78291785 | 25.0258891 | 24.5961083 | 24.98884553 | 24.29782083 | 26.3770801 | 24.22479189 | 24.84234919 | 25.40882494 |
| Nucleolar and spindle-associated prot | Q9BXS6-2;Q9BXS6;Q9BXS6-7 | NUSAP1 | Other | -1.172004984 | 1.718322779 | 26.70819899 | 26.5144858 | 26.42034063 | 27.01254594 | 26.81958115 | 26.9237705 | 25.12545856 | 25.80562777 | 25.75236023 |
| Nuclear transport factor 2 | P61970 | NUTF2 | Extracellular or secreted | 0.1355409 | 0.352272776 | 31.16991359 | 31.33308811 | 31.1919589 | 31.13968097 | 31.25141001 | 30.55380346 | 31.43355499 | 31.26807513 | 31.0249201 |
| Nuclear RNA export factor 1 | Q9UBU9;Q9UBU9-2 | NXF1 | Other | -0.739332263 | 1.56283497 | 24.33285771 | 25.22927357 | 24.94775285 | 25.04988676 | 25.21992713 | 25.91474461 | 24.64471447 | 24.08820788 | 24.39630217 |
| Nucleoredoxin | Q6DKJ4;Q6DKJ4-3 | NXN | Other | -0.051030087 | 0.498356994 | 26.48871823 | 26.57873915 | 26.74560622 | 26.51389876 | 26.55404508 | 26.67060655 | 26.56172646 | 26.49047903 | 26.57051124 |
| NTF2-related export protein 1;NTF2-r | Q9UKK6;Q9NPJ8;Q9NPJ8-2;Q | NXT1;NXT2 | Other | 0.252436447 | 0.168060682 | 26.25353273 | 26.2831439 | 26.05548303 | 26.28434475 | 26.41262528 | 22.89385846 | 26.0252555 | 26.1365309 | 25.68701701 |
| O-acetyl-ADP-ribose deacetylase 1 | Q9Y530 | OARD1 | Other | 0.394865035 | 0.29526781 | 22.1200328 | 22.62495516 | 22.38624458 | 22.22090231 | 21.80340528 | 24.25031924 | 22.12669565 | 23.28008429 | 23.48074485 |
| Ornithine aminotransferase, mitochor | P04181;P04181-2 | OAT | Other | -0.650419062 | 0.417369233 | 29.71035208 | 29.83633865 | 29.84882129 | 30.77407606 | 30.88474649 | 31.06535298 | 29.20053156 | 29.05442244 | 30.85363259 |
| Obscurin-like protein 1 | O75147-2;O75147-1;O75147 | OBSL1 | Other | 0.918689343 | 1.235729602 | 23.5395498 | 23.48419156 | 21.86730761 | 22.93044758 | 21.7531913 | 23.94815403 | 23.68281803 | 23.88082334 | 23.9538476 |
| OCIA domain-containing protein 2 | Q56VL3 | OCIAD2 | Other | -2.063552872 | 0.636207873 | 21.27729354 | 19.91763416 | 24.19203722 | 20.80609413 | 22.31087606 | 26.90046571 | 19.10331342 | 22.69917486 | 19.70905351 |
| Occludin | Q16625-4;Q16625;Q16625-2 | OCLN | Extracellular or secreted | 0.447732402 | 0.917862676 | 25.40142199 | 25.83716015 | 25.85854053 | 25.86891296 | 26.08208763 | 25.69455513 | 26.04855705 | 26.05310209 | 26.61287727 |
| Inositol polyphosphate 5-phosphatase | Q01968-2;Q01968 | OCRL | Extracellular or secreted | 0.113875791 | 0.085847319 | 24.92479004 | 25.08172202 | 24.92678171 | 24.70197479 | 24.98719808 | 22.08576088 | 24.568889 | 24.51231706 | 24.61453507 |
| Oral-facial-digital syndrome 1 protein | O75665-3;O75665 | OFD1 | Other | -0.085492252 | 0.076197416 | 22.09061841 | 21.78628267 | 22.27515888 | 21.20035338 | 19.68337503 | 21.70582372 | 21.19219648 | 21.33041143 | 21.59172138 |
| 2-oxoglutarate dehydrogenase, mitoc | Q02218;Q02218-2 | OGDH | Other | 0.189980266 | 0.344314899 | 26.59562468 | 26.55178839 | 26.81848447 | 26.95426927 | 26.68592164 | 28.04379954 | 27.0964755 | 27.05713702 | 27.24127228 |
| Prolyl 3-hydroxylase OGFOD1 | Q8N543;Q8N543-2 | OGFOD1 | Other | 0.098579266 | 0.192247854 | 28.31116354 | 28.14928398 | 28.10819054 | 28.48730805 | 28.53136986 | 27.76458513 | 28.51867098 | 28.44580921 | 28.00720816 |
| Opioid growth factor receptor | Q9NZT2-2;Q9NZT2 | OGFR | Other | -0.152158415 | 0.642950181 | 28.21326682 | 28.36686378 | 28.36344221 | 28.31799155 | 28.30412896 | 28.02284526 | 28.22080363 | 28.18181961 | 27.93517081 |
| UDP-N-acetylglucosamine--peptide N- | O15294;O15294-3;O15294-2 | OGT | Extracellular or secreted | 0.108590212 | 0.180111071 | 25.65616428 | 25.14887685 | 25.55616666 | 26.47559021 | 26.34250321 | 25.93676642 | 26.11277053 | 25.72634831 | 26.04468561 |
| Obg-like ATPase 1 | Q9NTK5;Q9NTK5-3;Q9NTK5-2 | OLA1 | Extracellular or secreted | 0.161613788 | 1.132722566 | 30.66999866 | 30.88278042 | 30.85348387 | 30.57362026 | 30.57199428 | 30.56683323 | 30.87151502 | 30.91591582 | 30.75676588 |
| Olfactomedin-4 | Q6UX06 | OLFM4 | Extracellular or secreted | 0.413738419 | 1.129498519 | 22.47313297 | 22.9551433 | 22.76247288 | 22.24043823 | 21.87140562 | 22.04159766 | 22.84412263 | 22.61862926 | 22.9505587 |
| Olfactomedin-like protein 2A | Q68BL7-2;Q68BL7;Q68BL7-3 | OLFML2A | Extracellular or secreted | -0.082933418 | 0.032146527 | 22.78208799 | 23.13436239 | 22.18818795 | 23.17680561 | 24.41441685 | 25.05254251 | 22.48081879 | 22.79419849 | 24.85038411 |
| Dynamin-like 120 kDa protein, mitoch | O60313;O60313-2;O60313-1 | OPA1 | Other | -0.386261045 | 0.807631614 | 27.55413293 | 27.59526898 | 27.72201482 | 27.70021607 | 27.34214712 | 27.72760818 | 26.93314565 | 27.17426634 | 27.55449892 |
| 5-oxoprolinase | O14841 | OPLAH | Other | 1.160798434 | 1.248441487 | 23.39191797 | 21.37832633 | 23.54297705 | 24.04937376 | 21.88630944 | 21.5956703 | 23.75480841 | 24.01631319 | 23.63356112 |
| Optineurin | Q96CV9-3;Q96CV9-2;Q96CV9 | OPTN | Extracellular or secreted | 0.676994848 | 0.368098734 | 21.66914545 | 21.57144899 | 20.92882484 | 22.52036607 | 21.68029711 | 21.13340668 | 21.47243898 | 21.69451263 | 23.6157775 |
| Origin recognition complex subunit 2 | Q13416 | ORC2 | Other | 0.2400441 | 0.311930513 | 22.69644547 | 24.12604966 | 24.15274965 | 23.52300863 | 22.6776233 | 24.50639669 | 23.85367125 | 23.82434815 | 23.88324961 |
| Origin recognition complex subunit 6 | Q9Y5N6 | ORC6 | Other | -0.440088496 | 0.494850028 | 21.27612851 | 23.35440974 | 20.96366882 | 20.93070269 | 21.12886784 | 21.66660401 | 20.97161933 | 20.93833335 | 21.42997264 |
| Oxysterol-binding protein 1 | P22059 | OSBP | Other | 0.151139307 | 0.474625961 | 27.98239673 | 27.79234518 | 27.74791227 | 27.2634708 | 27.21808013 | 27.76452186 | 27.82796153 | 27.63878964 | 27.87103025 |
| Oxysterol-binding protein-related pro | Q9BXB4 | OSBPL11 | Other | 0.132752783 | 0.134472911 | 25.19381379 | 25.17272232 | 25.3514445 | 24.74847542 | 24.77188073 | 25.60218233 | 24.99545974 | 24.90214617 | 25.92091197 |
| Oxysterol-binding protein-related pro | Q9BXW6;Q9BXW6-4;Q9BXW | OSBPL1A | Extracellular or secreted | 0.36790547 | 0.759327439 | 27.14004681 | 27.45740255 | 27.26051271 | 26.24878218 | 26.28346187 | 26.80328903 | 27.19629264 | 27.466066 | 27.03810534 |
| Oxysterol-binding protein-related pro | Q9H1P3-2;Q9H1P3 | OSBPL2 | Other | 0.464964997 | 0.889005404 | 22.75124795 | 22.60855736 | 22.02644645 | 23.11936798 | 22.45250253 | 22.07372057 | 23.26470603 | 23.03845286 | 22.60765753 |

| | | | | | | | | | | | | | | |
|---|---|---|---|---|---|---|---|---|---|---|---|---|---|---|
| Oxysterol-binding protein-related pro | Q9H4L5-2;Q9H4L5;Q9H4L5-6 | OSBPL3 | Extracellular or secreted | 1.380164623 | 3.086641331 | 22.84284357 | 22.33248898 | 22.00744344 | 22.44085903 | 21.80401357 | 22.90921485 | 23.98398446 | 23.47766045 | 23.84728067 |
| Oxysterol-binding protein-related pro | Q9BZF1-3;Q9BZF1-2;Q9BZF1 | OSBPL8 | Other | 0.440276796 | 0.166830229 | 24.68821829 | 24.72977465 | 24.94895605 | 20.14309366 | 20.72123782 | 25.24604759 | 24.58164378 | 23.29782083 | 23.6800298 |
| Oxysterol-binding protein-related pro | Q96SU4-7;Q96SU4-2;Q96SU4 | OSBPL9 | Other | 0.646839836 | 0.778724469 | 23.50845345 | 23.40807987 | 23.59889304 | 21.39329548 | 23.55309301 | 24.06324036 | 23.71471088 | 23.77826245 | 24.21007378 |
| Probable tRNA N6-adenosine threony | Q9NPF4 | OSGEP | Other | -0.053090576 | 0.04100128 | 28.12957153 | 28.12322004 | 27.89261304 | 27.66864687 | 27.83303889 | 27.4374595 | 28.18238806 | 28.21974254 | 26.9808726 |
| Oncostatin-M-specific receptor subun | Q99650-2;Q99650 | OSMR | Extracellular or secreted | 0.503180831 | 0.35098154 | 23.45412656 | 21.35351044 | 23.14626847 | 24.43338983 | 24.62628164 | 24.63964542 | 23.85642857 | 23.6884851 | 24.79124001 |
| Osteoclast-stimulating factor 1 | Q92882 | OSTF1 | Extracellular or secreted | 0.778893545 | 1.44453948 | 26.85602481 | 26.86726403 | 27.08588038 | 26.77077236 | 26.7206457 | 26.17157753 | 27.77121329 | 27.67976142 | 27.12178833 |
| Ubiquitin thioesterase OTUB1 | Q96FW1 | OTUB1 | Extracellular or secreted | -0.1366916 | 0.15495526 | 31.05050964 | 30.97199529 | 30.92195622 | 30.60293272 | 30.6462434 | 30.06843155 | 30.78939409 | 30.92719441 | 30.0043711 |
| OTU domain-containing protein 6B | Q8N6M0;Q8N6M0-2 | OTUD6B | Other | 0.402136905 | 0.855012067 | 25.37145762 | 25.33671003 | 25.36202112 | 25.75513962 | 25.37102549 | 25.11151746 | 25.89896777 | 26.02795694 | 25.43342167 |
| OTU domain-containing protein 7B | Q6GQQ9-2;Q6GQQ9 | OTUD7B | Other | -0.021549421 | 0.01266341 | 21.99289547 | 22.47137257 | 22.41635119 | 21.70064815 | 22.56316182 | 22.51732433 | 23.10032291 | 22.35511985 | 21.31078575 |
| Ubiquitin thioesterase otulin | Q96BN8 | OTULIN | Other | 0.402779658 | 0.90912462 | 25.22342994 | 25.14059296 | 25.1510469 | 25.33476811 | 25.70846214 | 25.2180062 | 25.82082306 | 25.91358101 | 25.36208803 |
| Ovarian cancer-associated gene 2 pro | Q8WZ82 | OVCA2 | Other | 0.418347525 | 0.653061399 | 25.80424961 | 25.65973399 | 25.52951682 | 25.46288764 | 25.56418578 | 25.32501909 | 26.19289252 | 26.24465113 | 25.49029539 |
| Succinyl-CoA:3-ketoacid coenzyme A | P55809 | OXCT1 | Extracellular or secreted | -0.640949064 | 3.006127027 | 28.99037209 | 29.14879929 | 29.05674954 | 29.35653263 | 29.58102236 | 28.91803639 | 28.5413383 | 28.42918093 | 28.63238972 |
| Oxidation resistance protein 1 | Q8N573-8;Q8N573;Q8N573- | OXR1 | Other | -1.254786957 | 1.328193162 | 28.08278811 | 28.08339695 | 28.00998627 | 28.63668253 | 29.28711789 | 28.25822276 | 27.13192775 | 26.52628132 | 27.75652731 |
| 3-oxoacyl-[acyl-carrier-protein] synth | Q9NWU1;Q9NWU1-2 | OXSM | Other | -0.25482048 | 0.650192604 | 25.99150783 | 26.60087886 | 25.80176065 | 25.74142028 | 25.82425102 | 25.89444145 | 25.63267399 | 25.98324519 | 25.54674942 |
| Serine/threonine-protein kinase OSR1 | O95747 | OXSR1 | Extracellular or secreted | -0.239776148 | 0.758476461 | 27.43817368 | 27.33722063 | 27.45104757 | 26.81567802 | 26.90735923 | 26.73337391 | 27.00934564 | 26.89145462 | 26.72129782 |
| Prolyl 4-hydroxylase subunit alpha-1 | P13674-3;P13674-2;P13674 | P4HA1 | Extracellular or secreted | -0.612827531 | 2.177415093 | 27.44758372 | 27.44529617 | 27.45826341 | 27.18999327 | 27.45450313 | 28.11018363 | 26.77994113 | 26.83942064 | 27.0950673 |
| Prolyl 4-hydroxylase subunit alpha-2 | O15460-2;O15460 | P4HA2 | Extracellular or secreted | -1.030219579 | 2.286567633 | 24.83079255 | 24.33388145 | 24.59559623 | 24.61907723 | 24.84536753 | 24.49661732 | 23.83422104 | 23.58279251 | 23.35299387 |
| Protein disulfide-isomerase | P07237 | P4HB | Extracellular or secreted | -0.223817705 | 2.746050955 | 31.97705523 | 31.96949296 | 31.90806109 | 31.84344737 | 31.99006772 | 32.11250335 | 31.70989215 | 31.79587081 | 31.72309778 |
| Transmembrane prolyl 4-hydroxylase | Q9NXG6-2;Q9NXG6;Q9NXG6 | P4HTM | Other | -0.13785881 | 0.198847811 | 23.58920857 | 23.30396353 | 23.19300912 | 22.41537144 | 22.80833106 | 23.11047439 | 22.89119037 | 23.31699961 | 22.58841265 |
| Proliferation-associated protein 2G4 | Q9UQ80;Q9UQ80-2 | PA2G4 | Extracellular or secreted | 0.613331011 | 2.333939651 | 30.33734825 | 30.39130755 | 30.40141386 | 30.85934653 | 30.86695465 | 30.74903194 | 31.13491657 | 31.10280793 | 31.40496992 |
| Proteasomal ATPase-associated facto | Q9BRP4-2;Q9BRP4;Q9BRP4-3 | PAAF1 | Other | -0.437972296 | 1.446617039 | 26.29864254 | 26.3005813 | 26.26897639 | 26.16739135 | 26.31637822 | 25.57447227 | 25.6970768 | 25.91845729 | 25.53377006 |
| Polyadenylate-binding protein 1 | P11940;P11940-2 | PABPC1 | Extracellular or secreted | 0.615879759 | 3.790762768 | 31.40232911 | 31.52323116 | 31.48375733 | 31.64203064 | 31.65744603 | 31.58557798 | 32.24402614 | 32.16656558 | 32.08423369 |
| Polyadenylate-binding protein 4 | Q13310;Q13310-3;Q13310-2 | PABPC4 | Other | 0.666940952 | 2.057884573 | 28.34006815 | 28.5218562 | 28.52757192 | 28.2953967 | 28.37335081 | 28.84008128 | 29.32109236 | 29.19884685 | 28.93004617 |
| Polyadenylate-binding protein 2 | Q86U42-2;Q86U42;Q92843-2 | PABPN1 | Other | -0.203675588 | 0.449838601 | 28.60179991 | 28.94408138 | 28.79076167 | 28.62383534 | 28.75468101 | 28.13662868 | 28.14150925 | 28.68789137 | 28.4854666 |
| Phosphofurin acidic cluster sorting pr | Q86VP3-4;Q86VP3;Q86VP3-3 | PACS2 | Extracellular or secreted | -0.181816215 | 0.354446691 | 22.67042286 | 23.22091615 | 23.21599386 | 21.87243553 | 22.73728888 | 22.33399061 | 22.58701199 | 22.50390045 | 22.38916286 |
| Protein kinase C and casein kinase sub | Q9BY11 | PACSIN1 | Extracellular or secreted | -2.344969842 | 1.519355654 | 27.84792578 | 29.31189975 | 28.51750756 | 28.22448832 | 27.55134837 | 27.65273613 | 26.22690589 | 24.76415483 | 26.52698272 |
| Protein kinase C and casein kinase sub | Q9UNF0;Q9UNF0-2 | PACSIN2 | Extracellular or secreted | 0.351559996 | 1.77996952 | 29.10941163 | 29.1369953 | 29.0626281 | 28.79686881 | 29.11924231 | 28.73927768 | 29.45399317 | 29.38007725 | 29.20282148 |
| Protein kinase C and casein kinase sub | Q9UKS6 | PACSIN3 | Extracellular or secreted | 0.534989659 | 3.138919217 | 26.89203394 | 27.01690824 | 27.05040585 | 26.73660301 | 26.59718873 | 27.11306872 | 27.36952866 | 27.45630616 | 27.48223841 |
| Protein-arginine deiminase type-2 | Q9Y2J8;Q9Y2J8-2 | PADI2 | Extracellular or secreted | -0.697832821 | 3.099161527 | 28.63082868 | 28.69879298 | 28.6559801 | 28.50846856 | 28.49331474 | 28.03548578 | 27.83279751 | 27.78000373 | 27.80513572 |
| Protein-arginine deiminase type-3 | Q9ULW8 | PADI3 | Other | 0.276922356 | 0.118195682 | 20.93774687 | 22.5698278 | 22.67803183 | 23.40377769 | 23.26544633 | 24.23330253 | 22.32961789 | 22.48937074 | 24.55584496 |
| RNA polymerase II-associated factor 1 | Q8N7H5;Q8N7H5-3;Q8N7H5 | PAF1 | Other | -2.98E-05 | 7.05E-05 | 26.90793221 | 26.84314032 | 26.79476368 | 26.24988541 | 26.46216998 | 26.8931919 | 26.78369229 | 26.517087 | 26.77467307 |
| Platelet-activating factor acetylhydrol | P43034 | PAFAH1B1 | Extracellular or secreted | -0.279990823 | 1.265431986 | 31.04895194 | 31.20565547 | 30.98725772 | 31.12342974 | 31.3225973 | 30.55682079 | 30.82084436 | 30.80562777 | 30.65591188 |
| Platelet-activating factor acetylhydrol | P68402;P68402-3;P68402-2;P | PAFAH1B2 | Extracellular or secreted | -0.153131653 | 0.243481051 | 29.27921208 | 29.47086311 | 29.32905069 | 28.87038363 | 28.89755482 | 28.27150762 | 29.22841597 | 28.68471845 | 28.6867566 |
| Platelet-activating factor acetylhydrol | Q15102 | PAFAH1B3 | Extracellular or secreted | -0.128036818 | 0.267413437 | 30.00383518 | 30.16098826 | 29.99212941 | 29.79468623 | 29.87207304 | 29.57069218 | 29.76657678 | 30.06162425 | 29.48489067 |
| Multifunctional protein ADE2;Phosph | P22234;P22234-2 | PAICS | Extracellular or secreted | 0.047143252 | 0.171458877 | 31.96154592 | 31.94342477 | 31.94642595 | 32.03209951 | 31.94555416 | 31.95355485 | 32.11185072 | 32.10224478 | 31.81863683 |
| Polyadenylate-binding protein-interac | Q9H074;Q9H074-2;Q9H074-3 | PAIP1 | Other | 0.257151127 | 0.69072354 | 28.79485662 | 28.83007927 | 29.10614477 | 28.74454804 | 28.74178672 | 28.66475341 | 29.23226432 | 29.19399233 | 28.78628115 |
| Polyadenylate-binding protein-interac | Q9BPZ3 | PAIP2 | Other | 0.949696146 | 0.389290205 | 23.62756238 | 23.41484377 | 20.74694769 | 24.78820574 | 25.86410224 | 24.41319043 | 24.62522279 | 26.17548514 | 23.47580664 |
| Serine/threonine-protein kinase PAK 1 | Q13153;Q13153-2 | PAK1 | Extracellular or secreted | -0.116920579 | 0.250311937 | 26.36961186 | 26.4428152 | 26.40885733 | 25.61028841 | 25.7523347 | 26.60625519 | 26.10645432 | 26.2290168 | 25.90884849 |
| p21-activated protein kinase-interacti | Q9NWT1 | PAK1IP1 | Extracellular or secreted | -1.465567256 | 4.80626454 | 27.1178314 | 27.21141126 | 27.07081293 | 26.64663825 | 26.80513572 | 27.47374922 | 25.44950561 | 25.65206594 | 25.66451608 |
| Serine/threonine-protein kinase PAK 2 | Q13177 | PAK2 | Extracellular or secreted | 0.807721512 | 1.017092238 | 28.61477375 | 28.32513064 | 28.47518045 | 28.28520062 | 28.17650318 | 28.43364453 | 29.38582673 | 29.55373023 | 28.63882415 |
| Serine/threonine-protein kinase PAK 4 | O96013;O96013-2;O96013-3 | PAK4 | Other | -0.260250837 | 0.581743584 | 25.99016648 | 26.17761556 | 26.09057201 | 25.67675211 | 25.74975447 | 25.27575794 | 25.38284563 | 25.44154938 | 25.87516176 |
| Paladin | Q9ULE6 | PALD1 | Other | 0.237511594 | 0.848718901 | 23.70282032 | 24.1220254 | 24.35602043 | 24.70377096 | 24.25205325 | 24.23169229 | 24.55625438 | 24.41531987 | 24.42515186 |
| Paralemmin-1 | O75781-2;O75781 | PALM | Extracellular or secreted | -0.322022167 | 0.5886719 | 26.46971869 | 25.99642987 | 26.19398293 | 26.00787112 | 25.9789979 | 24.90044268 | 25.72811441 | 25.76298983 | 25.31655085 |
| Paralemmin-3 | A6NDB9 | PALM3 | Extracellular or secreted | -0.753393107 | 0.32451324 | 24.18842827 | 24.3312182 | 24.29382804 | 23.55227227 | 24.06241709 | 23.21182711 | 24.30966397 | 23.74519586 | 21.50495633 |
| Peptidyl-glycine alpha-amidating mon | P19021-6;P19021;P19021-5;P | PAM | Extracellular or secreted | 0.654611302 | 0.773754267 | 23.78935056 | 24.33114985 | 23.3626566 | 25.20523364 | 24.91524626 | 25.41638341 | 24.82259884 | 24.97753568 | 25.67370955 |
| Pantothenate kinase 3 | Q9H999 | PANK3 | Other | 1.168218457 | 1.172717945 | 22.84838913 | 22.48640297 | 22.47152142 | 20.18220186 | 22.71659672 | 23.88203698 | 23.29712114 | 23.83142091 | 23.66968786 |
| Pantothenate kinase 4 | Q9NVE7 | PANK4 | Other | 0.238667528 | 0.137839458 | 24.56022553 | 25.43634784 | 25.51487703 | 25.71931445 | 26.86076779 | 23.6846382 | 25.86818295 | 26.14538242 | 24.59052263 |
| Poly(A) polymerase alpha | P51003 | PAPOLA | Other | 0.321711662 | 0.392957363 | 28.11619601 | 28.26450023 | 28.2071344 | 27.89110691 | 27.85239778 | 28.05842788 | 28.73969616 | 28.64388921 | 27.77643123 |
| Bifunctional 3-phosphoadenosine 5-p | O43252 | PAPSS1 | Other | 0.116223462 | 0.575161613 | 27.91689822 | 27.77705862 | 27.77045732 | 27.98891053 | 28.0973298 | 27.77045732 | 27.95277066 | 27.91090802 | 28.14554762 |
| Protein deglycase DJ-1 | Q99497 | PARK7 | Extracellular or secreted | -0.021529496 | 0.035439284 | 32.60961846 | 32.537673 | 32.56905384 | 32.9035611 | 32.91609384 | 31.64568527 | 32.6049439 | 32.5967268 | 32.32458356 |
| Poly(A)-specific ribonuclease PARN | O95453-2;O95453;O95453-3 | PARN | Other | 0.190088854 | 0.715558703 | 27.86342888 | 27.91917367 | 27.76755579 | 27.87513832 | 27.87859177 | 28.08603229 | 28.22637309 | 28.14913859 | 27.88971524 |
| Poly [ADP-ribose] polymerase 1 | P09874 | PARP1 | Other | -1.072065371 | 1.04874518 | 32.64501953 | 32.71210834 | 32.67383589 | 32.01571258 | 32.06204261 | 32.30310021 | 31.68678999 | 31.72012378 | 30.58279969 |
| Poly [ADP-ribose] polymerase 4 | Q9UKK3 | PARP4 | Extracellular or secreted | 0.938453319 | 1.974175397 | 27.08810676 | 27.25565909 | 27.01126667 | 25.93172636 | 25.89970541 | 26.7050375 | 27.61365021 | 27.70358616 | 27.44387448 |
| Alpha-parvin | Q9NVD7 | PARVA | Other | 0.687943425 | 3.315426419 | 24.49118274 | 24.39113134 | 24.12801824 | 24.63809912 | 24.19633017 | 24.05299848 | 25.12888357 | 24.95681889 | 24.92700786 |
| Beta-parvin | Q9HBI1;Q9HBI1-3;Q9HBI1-2 | PARVB | Extracellular or secreted | -0.775036358 | 0.488772441 | 21.94252739 | 21.66262144 | 21.78636217 | 21.9006646 | 21.99823216 | 22.65360884 | 21.79137913 | 21.83968974 | 20.01583035 |
| Protein PAT1 homolog 1 | Q86TB9-4;Q86TB9;Q86TB9-2 | PATL1 | Other | -0.333081594 | 0.422923875 | 23.30033692 | 23.27653906 | 23.62053218 | 23.79044475 | 23.2641333 | 23.71502522 | 23.61795704 | 22.61006871 | 23.25623518 |
| PRKC apoptosis WT1 regulator protei | Q96IZ0 | PAWR | Extracellular or secreted | 2.730767741 | 2.562888346 | 23.88595102 | 24.33905729 | 24.52540037 | 25.5621779 | 25.92768611 | 22.98697249 | 27.63041212 | 27.65799111 | 26.51752258 |
| PAX3- and PAX7-binding protein 1 | Q9Y5B6-3;Q9Y5B6-4;Q9Y5B6 | PAXBP1 | Other | -0.156320785 | 0.201539541 | 22.70878837 | 22.80183464 | 22.6743078 | 23.42751951 | 23.25270296 | 21.73022646 | 22.3532047 | 22.49211235 | 22.98341047 |
| Protein PBDC1 | Q9BVG4 | PBDC1 | Other | 0.716656918 | 1.362715796 | 27.66857925 | 27.59184976 | 27.56906289 | 27.66166503 | 27.64677556 | 26.94815403 | 28.40722095 | 28.44919859 | 27.83659448 |
| Lymphokine-activated killer T-cell-orig | Q96KB5;Q96KB5-2 | PBK | Other | 0.510293174 | 0.549214444 | 26.78069214 | 26.67559508 | 26.40655614 | 26.89458022 | 26.52465338 | 26.51933859 | 27.52551985 | 27.46583254 | 26.4402349 |
| Pterin-4-alpha-carbinolamine dehydra | P61457 | PCBD1 | Extracellular or secreted | 0.747050736 | 0.997092875 | 27.45560089 | 27.29824049 | 27.28744819 | 26.83532993 | 26.91467619 | 27.96922533 | 27.76818705 | 27.73466642 | 28.61855925 |
| Poly(rC)-binding protein 1 | Q15365 | PCBP1 | Extracellular or secreted | 0.444175611 | 0.699218982 | 30.95982 | 30.91142244 | 31.01178651 | 31.12145489 | 31.14892047 | 31.34678217 | 31.74226069 | 31.79277182 | 31.04758756 |
| Poly(rC)-binding protein 2 | Q15366-3;Q15366-2 | PCBP2 | Extracellular or secreted | 0.391201274 | 0.754606633 | 28.84332006 | 29.01315852 | 29.08871336 | 29.38518903 | 29.35053853 | 29.92309042 | 29.85681144 | 29.84093667 | 29.27786067 |
| Propionyl-CoA carboxylase beta chain | P05166;P05166-2 | PCCB | Extracellular or secreted | -1.031416766 | 2.052561274 | 23.31105091 | 22.98946934 | 23.08704258 | 24.18260591 | 23.28700777 | 24.45173935 | 22.47822848 | 22.51681958 | 22.56515957 |
| Protocadherin-1 | Q08174;Q08174-2 | PCDH1 | Extracellular or secreted | 2.447613224 | 2.328301489 | 24.42707187 | 25.2570989 | 25.27091978 | 27.27896848 | 27.16134874 | 27.06825215 | 28.16542788 | 28.23731114 | 29.17193061 |
| Pre-mRNA cleavage complex 2 protei | O94913 | PCF11 | Extracellular or secreted | -2.320297601 | 0.633164701 | 26.76331916 | 26.43466284 | 26.83520944 | 26.18184804 | 25.94263924 | 26.30949051 | 25.11521387 | 25.78937544 | 21.3681025 |
| PCI domain-containing protein 2 | Q5JVF3-3;Q5JVF3;Q5JVF3-4;C | PCID2 | Other | 0.54476015 | 1.389951524 | 24.4297556 | 24.85638107 | 25.04343453 | 25.01346206 | 24.28057992 | 25.63558285 | 25.56392404 | 25.32680278 | 25.37315164 |
| Phosphorylated CTD-interacting facto | Q9H4Z3 | PCIF1 | Extracellular or secreted | 0.309255212 | 0.366274884 | 22.79554665 | 22.85249065 | 23.11762535 | 22.1217054 | 23.944316 | 22.65208783 | 23.57623244 | 22.70176332 | 23.39165581 |
| Phosphoenolpyruvate carboxykinase | Q16822;Q16822-3;Q16822-2 | PCK2 | Extracellular or secreted | -0.639908085 | 0.274665593 | 27.08911762 | 26.99291276 | 26.78444135 | 26.15176307 | 26.33971995 | 24.26942235 | 26.57830727 | 26.28968347 | 24.02597356 |
| Pericentriolar material 1 protein | Q15154-5;Q15154-4;Q15154 | PCM1 | Other | 0.087252628 | 0.093823024 | 27.02231616 | 27.50248682 | 28.31055697 | 28.1325162 | 28.38562105 | 28.84035146 | 27.80328903 | 28.03852403 | 28.51686916 |
| Protein-L-isoaspartate(D-aspartate) O | P22061;P22061-2 | PCMT1 | Extracellular or secreted | -0.037650749 | 0.122227328 | 30.15942514 | 30.05607765 | 30.07367563 | 29.87781693 | 29.67623425 | 29.9141344 | 29.83116718 | 30.09911216 | 29.83545042 |
| Proliferating cell nuclear antigen | P12004 | PCNA | Extracellular or secreted | 0.372000347 | 2.444603917 | 32.98437433 | 33.02290478 | 32.91787283 | 32.64785217 | 32.57893705 | 32.75455757 | 33.22216346 | 33.19326685 | 33.1538201 |
| PEST proteolytic signal-containing nuc | Q8WW12;Q8WW12-2 | PCNP | Other | 0.03448208 | 0.064895987 | 28.54892583 | 28.46084294 | 28.40515257 | 28.5363071 | 28.44007645 | 28.14214251 | 28.56685137 | 28.67640241 | 28.12691617 |
| Procollagen C-endopeptidase enhanc | Q9UKZ9 | PCOLCE2 | Extracellular or secreted | 1.51877993 | 1.922464499 | 23.86947788 | 24.67728995 | 23.94520948 | 24.99937935 | 25.26888718 | 26.25212545 | 26.24245396 | 25.96582831 | 26.85424216 |
| ProSAAS;KEP;Big SAAS;Little SAAS;Big | Q9UHG2 | PCSK1N | Extracellular or secreted | 2.730579963 | 1.070072301 | 21.69612387 | 20.41788533 | 21.19127404 | 23.3710145 | 21.0970047 | 23.02631135 | 23.93012515 | 23.23593358 | 26.42548805 |
| Proprotein convertase subtilisin/kexir | P29122-2;P29122;P29122-8;P | PCSK6 | Extracellular or secreted | 0.970522635 | 1.219018395 | 24.32827609 | 25.79335028 | 24.21348933 | 24.46207635 | 25.10589508 | 26.6988261 | 26.01678075 | 26.26902992 | 25.92671386 |
| Phosphatidylcholine transfer protein | Q9UKL6 | PCTP | Other | -2.06671942 | 0.781173406 | 23.50312404 | 23.4299471 | 23.68656293 | 24.07193824 | 24.32120421 | 23.98693778 | 23.71397713 | 20.87076895 | 20.71495281 |
| Prenylcysteine oxidase 1 | Q9UHG3-2;Q9UHG3 | PCYOX1 | Extracellular or secreted | -0.087982778 | 0.029911852 | 22.29574876 | 23.49789666 | 23.57588635 | 23.17771819 | 23.23072529 | 27.31949972 | 23.13233969 | 22.97732605 | 25.17512341 |
| Choline-phosphate cytidylyltransferas | P49585 | PCYT1A | Other | 0.803802323 | 1.837546594 | 25.97259247 | 26.45150355 | 26.18735253 | 25.32546522 | 25.4994182 | 26.75021465 | 26.91456214 | 26.71240357 | 26.87771456 |
| Ethanolamine-phosphate cytidylyltran | Q99447-3;Q99447;Q99447-4 | PCYT2 | Other | 2.890095567 | 1.440244487 | 22.4542772 | 23.75654003 | 23.9735124 | 22.55810116 | 24.13555274 | 23.78166787 | 25.63624691 | 25.63749119 | 27.7263743 |
| 28 kDa heat- and acid-stable phospho | Q13442 | PDAP1 | Other | 0.294158541 | 0.419542501 | 29.34199872 | 29.2775725 | 29.32551669 | 29.55462699 | 29.77581925 | 28.10819054 | 29.72053154 | 29.7359255 | 29.11788093 |
| Programmed cell death protein 10 | Q9BUL8 | PDCD10 | Extracellular or secreted | 0.76467728 | 1.791229792 | 26.93562045 | 27.0517546 | 27.11812855 | 26.17314184 | 26.59135045 | 27.35237095 | 27.48523626 | 27.94832115 | 27.47165785 |
| Protein RRP5 homolog | Q14690 | PDCD11 | Other | -1.375565733 | 0.825994256 | 22.12999986 | 22.98254035 | 23.37159055 | 21.52353516 | 21.38700758 | 26.33377911 | 21.72322815 | 22.18440657 | 20.82989439 |
| Programmed cell death protein 2-like | Q9BRP1 | PDCD2L | Other | 1.150532946 | 2.428024086 | 24.43090424 | 24.10421609 | 24.11068147 | 24.80168665 | 24.83238709 | 23.25218321 | 25.36402695 | 25.58325175 | 25.27034951 |
| Programmed cell death protein 4 | Q53EL6-2;Q53EL6 | PDCD4 | Other | -0.216707421 | 0.087009734 | 25.37650062 | 25.00748621 | 20.47788263 | 24.78770771 | 24.89933664 | 24.93449607 | 24.64179643 | 24.4137715 | 23.03601475 |
| Programmed cell death protein 5 | O14737;O14737-2 | PDCD5 | Extracellular or secreted | 0.192029175 | 0.174018579 | 29.4160894 | 29.47556702 | 29.45067403 | 29.75867305 | 29.70673437 | 28.45273712 | 29.80658064 | 30.04179085 | 28.85795353 |
| Programmed cell death protein 6 | O75340;O75340-2 | PDCD6 | Extracellular or secreted | -0.023306558 | 0.071558916 | 27.79903265 | 27.95986144 | 27.96207019 | 27.83713609 | 27.59106505 | 27.85863538 | 27.64209952 | 27.81035521 | 27.981526 |
| Programmed cell death 6-interacting | Q8WUM4 | PDCD6IP | Extracellular or secreted | 0.042375396 | 0.158826669 | 31.41855439 | 31.4412941 | 31.41452774 | 31.43380368 | 31.43071685 | 31.79559991 | 31.4182025 | 31.49046755 | 31.68570447 |
| Programmed cell death 6-interacting | Q8WUM4-2 | PDCD6IP | Extracellular or secreted | 0.286715054 | 0.320641192 | 23.4020946 | 21.36554933 | 21.03568353 | 22.03575487 | 21.49415431 | 22.18014189 | 22.2806297 | 22.44649876 | 21.88970596 |
| Phosducin-like protein 3 | Q9H2J4 | PDCL3 | Other | 1.23210177 | 4.518363541 | 25.58448523 | 25.59877948 | 25.69075104 | 25.86485793 | 26.22048077 | 26.13657001 | 27.12780183 | 27.15643849 | 26.96002722 |
| Parkinson disease 7 domain-containin | Q8NB37;Q8NB37-3;Q8NB37- | PDDC1 | Extracellular or secreted | -0.317544887 | 1.407175664 | 25.18271956 | 25.25078907 | 25.04785014 | 25.22714467 | 24.78740881 | 24.55162707 | 24.65850875 | 24.6792782 | 24.73334805 |
| 2,5-phosphodiesterase 12 | Q6L8Q7;Q6L8Q7-2 | PDE12 | Other | 0.411601654 | 1.022787559 | 25.94612472 | 26.01075465 | 25.85276684 | 26.16817599 | 26.09196443 | 25.75322777 | 26.61400141 | 26.43310325 | 26.09920749 |
| Retinal rod rhodopsin-sensitive cGMP | O43924 | PDE6D | Other | -0.696794039 | 0.784980316 | 25.61860125 | 25.26538584 | 25.94605777 | 26.56809169 | 26.74098301 | 25.68843174 | 25.26391847 | 24.69683809 | 25.86263698 |
| Platelet-derived growth factor recept | Q15198 | PDGFRL | Extracellular or secreted | -0.051155042 | 0.050683504 | 24.37828798 | 23.90848205 | 23.98554874 | 24.52408541 | 24.57299899 | 25.62382837 | 24.19430238 | 24.20448756 | 24.94436069 |
| Pyruvate dehydrogenase E1 compone | P08559-4;P08559-3;P08559;P | PDHA1 | Extracellular or secreted | 0.221707653 | 0.129195587 | 25.00530313 | 25.16169471 | 25.15784588 | 26.19208357 | 26.34192664 | 27.90178953 | 25.6917629 | 25.76022554 | 27.09345625 |
| Pyruvate dehydrogenase E1 compone | P11177-2;P11177;P11177-3 | PDHB | Extracellular or secreted | -0.009573328 | 0.005256727 | 25.47701189 | 25.35736353 | 25.53489819 | 26.59548241 | 26.46156126 | 27.55464529 | 25.62692216 | 25.71463228 | 27.12020686 |
| Protein disulfide-isomerase A3 | P30101 | PDIA3 | Extracellular or secreted | -0.679429288 | 3.669959658 | 33.81859874 | 33.916521 | 33.83064221 | 33.93059964 | 34.04807499 | 33.83977352 | 33.28705732 | 33.12671005 | 33.24004981 |
| Protein disulfide-isomerase A4 | P13667 | PDIA4 | Extracellular or secreted | -0.681580323 | 3.607769874 | 33.89066491 | 33.88812623 | 33.80510881 | 33.4480073 | 33.54754298 | 33.81488009 | 33.05467149 | 33.07014437 | 33.02760833 |
| Protein disulfide-isomerase A5 | Q14554 | PDIA5 | Extracellular or secreted | -0.498918684 | 0.841159953 | 27.79054418 | 27.80771712 | 27.73660301 | 27.27355432 | 27.40324378 | 27.41538435 | 27.37691456 | 27.21048259 | 26.62937018 |
| Protein disulfide-isomerase A6 | Q15084-3;Q15084;Q15084-4 | PDIA6 | Extracellular or secreted | -0.373669546 | 2.849788918 | 31.05323159 | 31.29800007 | 31.26338125 | 31.03662587 | 31.11545203 | 31.27758359 | 30.86791205 | 30.82061615 | 30.71260036 |
| PDZ and LIM domain protein 1 | O00151 | PDLIM1 | Other | 0.50253606 | 1.712559119 | 28.29241565 | 28.30482526 | 28.25385729 | 27.92671386 | 27.88826416 | 28.32053299 | 28.76667155 | 28.78827421 | 28.44596703 |
| PDZ and LIM domain protein 2 | Q96JY6;Q96JY6-4;Q96JY6-3;C | PDLIM2 | Extracellular or secreted | 0.144297412 | 0.420570334 | 25.0055173 | 24.87159433 | 25.17649367 | 25.08946115 | 24.65239424 | 24.24016244 | 25.04435209 | 25.0398421 | 24.86650961 |
| PDZ and LIM domain protein 4 | P50479;P50479-2 | PDLIM4 | Other | 0.269000883 | 0.383327151 | 23.27511852 | 23.02370834 | 23.26570775 | 23.77239509 | 22.78576375 | 23.0311331 | 23.60036848 | 23.81215615 | 22.97139129 |
| PDZ and LIM domain protein 5 | Q96HC4;Q96HC4-4;Q96HC4- | PDLIM5 | Extracellular or secreted | 1.014982505 | 5.393110884 | 26.68992505 | 26.76471167 | 26.81567802 | 27.08972379 | 26.96939003 | 26.6699311 | 27.77906445 | 27.84445791 | 27.92110499 |
| PDZ and LIM domain protein 7 | Q9NR12;Q9NR12-2 | PDLIM7 | Other | 0.870227615 | 2.047321329 | 26.54671999 | 26.41599676 | 26.43658612 | 26.95881111 | 26.03783313 | 26.93370848 | 27.67505661 | 27.4708825 | 27.12957153 |
| [Pyruvate dehydrogenase [acetyl-tran | Q9P0J1;Q9P0J1-2 | PDP1 | Other | 0.742058373 | 1.051743639 | 23.06114834 | 23.31368243 | 23.35689359 | 23.27838366 | 22.99007556 | 25.22912685 | 24.40911639 | 24.42034063 | 24.01137332 |
| Pyruvate dehydrogenase phosphatase | Q8NCN5 | PDPR | Other | -0.479047798 | 1.297972345 | 26.12453202 | 26.18478259 | 26.06081037 | 25.67230728 | 25.56072116 | 26.77391891 | 25.45387546 | 25.78176791 | 25.5157494 |
| p53 and DNA damage-regulated prote | Q9NUG6 | PDRG1 | Other | 2.123118223 | 1.928099929 | 20.98209456 | 21.22718295 | 20.77522478 | 22.26124881 | 23.83518534 | 23.75389083 | 24.00821317 | 24.16959109 | 24.60896405 |
| Sister chromatid cohesion protein PDS | Q29RF7 | PDS5A | Extracellular or secreted | -0.702796618 | 2.385214396 | 28.54229816 | 28.5552672 | 28.29079102 | 27.74117594 | 27.82140712 | 28.39917412 | 27.4909839 | 27.67512393 | 27.40055909 |
| Sister chromatid cohesion protein PDS | Q9NTI5-2;Q9NTI5 | PDS5B | Other | -1.599984894 | 2.659970002 | 27.22462632 | 27.37393155 | 27.46372965 | 27.31120686 | 27.12534031 | 27.68585482 | 25.924337 | 25.91029047 | 25.45776261 |
| Pyridoxal-dependent decarboxylase d | Q6P996-4;Q6P996-5;Q6P996 | PDXDC1 | Other | -0.218688364 | 1.192462521 | 26.83146923 | 26.96069012 | 27.18688046 | 27.1931934 | 27.2141937 | 27.46232603 | 26.84361958 | 26.9497131 | 26.9749787 |
| Pyridoxal kinase | O00764;O00764-3 | PDXK | Extracellular or secreted | -0.25513075 | 0.582156422 | 29.61696564 | 29.57528048 | 29.56097984 | 29.11587366 | 29.01672231 | 28.95532292 | 28.94134121 | 28.85546054 | 29.35837843 |
| Pyridoxal phosphate phosphatase | Q96GD0 | PDXP | Extracellular or secreted | -0.183930334 | 0.344354961 | 23.58886558 | 23.89633684 | 23.78845469 | 24.06488549 | 24.19880471 | 23.76587527 | 24.06587166 | 23.41196296 | 23.62198566 |
| PDZ domain-containing protein 11 | Q5EBL8;Q5EBL8-2 | PDZD11 | Extracellular or secreted | 0.328143245 | 1.422940016 | 24.25781827 | 24.55637133 | 24.15050469 | 24.36673041 | 24.20791634 | 23.73117399 | 24.42027637 | 24.58078163 | 24.61862926 |
| Astrocytic phosphoprotein PEA-15 | Q15121;Q15121-2 | PEA15 | Other | 0.257953989 | 0.148781858 | 26.40480349 | 26.58491526 | 26.77843793 | 26.41076679 | 26.50373065 | 23.11379494 | 26.54218005 | 26.58233313 | 25.54757331 |
| Phosphatidylethanolamine-binding pr | P30086 | PEBP1 | Extracellular or secreted | -0.39076406 | 1.171861136 | 31.95168041 | 31.89327147 | 31.85708592 | 32.47853536 | 32.62731885 | 31.68925026 | 31.56524498 | 31.63460696 | 31.87642702 |
| Peflin | Q9UBV8 | PEF1 | Extracellular or secreted | 0.008196369 | 0.003872545 | 22.05876542 | 23.58543113 | 23.68228225 | 24.84790189 | 22.38989807 | 25.4266241 | 22.83341053 | 24.41873331 | 23.7678967 |
| Retrotransposon-derived protein PEG | Q86TG7-2;Q86TG7 | PEG10 | Other | 0.471365994 | 0.290218285 | 23.17456756 | 22.08410897 | 22.06578951 | 21.42555959 | 20.64426482 | 22.01034969 | 22.78859408 | 23.05137292 | 21.27645104 |
| Protein pelota homolog | Q9BRX2 | PELO | Other | 1.827351015 | 1.055701679 | 23.04725108 | 20.69323774 | 22.62936463 | 23.05216934 | 21.72963251 | 21.69102271 | 25.01405828 | 24.16023336 | 22.72910041 |
| Proline-, glutamic acid- and leucine-ri | Q8IZL8 | PELP1 | Other | -0.640701134 | 1.497901835 | 26.2464282 | 26.26726262 | 26.25628918 | 25.62329813 | 25.8073732 | 27.153349 | 25.71232485 | 25.63245213 | 25.4101198 |
| Xaa-Pro dipeptidase | P12955;P12955-2;P12955-3 | PEPD | Extracellular or secreted | -0.385028203 | 0.982143377 | 30.75748137 | 30.73884298 | 30.7175931 | 30.98535873 | 31.03786977 | 29.83930049 | 30.4182025 | 30.38133626 | 30.08359984 |
| p53 apoptosis effector related to PMP | Q96FX8 | PERP | Extracellular or secreted | 1.234735343 | 1.194495306 | 22.30309955 | 22.78358739 | 22.38637616 | 22.31928953 | 22.33298059 | 25.1973055 | 24.59531167 | 24.2327172 | 23.53749653 |
| Pescadillo homolog | O00541-2;O00541 | PES1 | Other | -1.504330599 | 0.460891597 | 23.19775542 | 25.56409854 | 25.63939702 | 24.07267433 | 24.61442274 | 26.40036365 | 21.06745691 | 23.96128647 | 25.20262067 |
| Peroxisomal membrane protein PEX14 | O75381-2;O75381 | PEX14 | Other | -0.773965622 | 1.116882523 | 23.22134403 | 22.960615 | 22.65414242 | 21.92145077 | 22.39113134 | 21.55421785 | 21.21549034 | 22.08203784 | 21.73202566 |
| Peroxisomal biogenesis factor 19 | P40855;P40855-5 | PEX19 | Extracellular or secreted | 1.220317006 | 0.694457085 | 25.88504335 | 25.97274583 | 26.29755849 | 26.15703632 | 26.11899983 | 21.85777117 | 27.01786405 | 27.18101391 | 25.60665055 |
| Peroxisomal targeting signal 2 recept | O00628-2;O00628 | PEX7 | Other | 0.339921316 | 0.388547675 | 21.92001695 | 21.77366241 | 21.82150931 | 20.64910983 | 20.75596742 | 20.79209442 | 22.17563361 | 21.47159584 | 21.22871467 |
| Phosphoribosylformylglycinamidine s | O15067 | PFAS | Extracellular or secreted | 0.081013321 | 0.162056466 | 30.09270045 | 30.14175285 | 30.05336108 | 30.31570474 | 30.37056829 | 29.66900184 | 30.39437755 | 30.29032964 | 29.87987739 |
| Prefoldin subunit 1 | O60925 | PFDN1 | Other | 0.649531943 | 2.444733084 | 27.52738549 | 27.27417665 | 27.21595314 | 26.74791227 | 26.79637376 | 27.34815501 | 27.80421267 | 27.77133924 | 27.82802208 |
| Prefoldin subunit 2 | Q9UHV9 | PFDN2 | Extracellular or secreted | 0.791992663 | 2.135276737 | 28.50771275 | 28.77542683 | 28.59331104 | 28.66397346 | 28.67788134 | 28.22453432 | 29.49111774 | 29.4898362 | 29.11644391 |
| Prefoldin subunit 4 | Q9NQP4 | PFDN4 | Other | 0.530740865 | 0.931496109 | 26.71934056 | 26.70727761 | 26.79476368 | 27.1841394 | 27.04103361 | 25.99468317 | 27.17407581 | 27.68471845 | 26.95404735 |
| Prefoldin subunit 5 | Q99471;Q99471-3 | PFDN5 | Other | 0.294441828 | 0.918047837 | 28.76458513 | 28.79318282 | 28.87882561 | 28.67556143 | 28.69055791 | 28.38038695 | 29.15918451 | 29.06576383 | 28.74992706 |
| Prefoldin subunit 6 | O15212 | PFDN6 | Other | 0.21551129 | 0.491110354 | 28.83563112 | 28.90821861 | 28.97702792 | 28.30238673 | 28.42237934 | 28.0986856 | 28.94533787 | 28.93623849 | 28.53712216 |
| 6-phosphofructo-2-kinase/fructose-2, | O60825-2;O60825 | PFKFB2 | Other | 0.646640602 | 0.43947947 | 23.98953864 | 23.81616649 | 23.88706737 | 24.53883969 | 24.3426558 | 24.72374717 | 23.89836815 | 24.87422382 | 25.81633742 |
| ATP-dependent 6-phosphofructokinas | P17858;P17858-2 | PFKL | Extracellular or secreted | 0.348986385 | 0.942620457 | 28.24907163 | 28.26123038 | 28.2429082 | 27.71679215 | 27.6881916 | 28.8181492 | 28.35795913 | 28.51163866 | 28.66553294 |
| ATP-dependent 6-phosphofructokinas | P08237;P08237-3;P08237-2 | PFKM | Extracellular or secreted | 0.219483888 | 0.266844125 | 28.29017582 | 28.18980481 | 28.32864418 | 28.24399778 | 28.2488455 | 29.49566088 | 28.42846228 | 28.42197825 | 29.20657562 |
| ATP-dependent 6-phosphofructokinas | Q01813;Q01813-2 | PFKP | Extracellular or secreted | 0.168036007 | 0.260758696 | 30.30486877 | 30.37015253 | 30.44173932 | 30.12755587 | 29.92238165 | 31.1759324 | 30.30007503 | 30.43430094 | 30.94104732 |
| Profilin-1 | P07737 | PFN1 | Extracellular or secreted | 0.221694548 | 1.130487856 | 33.80087323 | 33.75851422 | 33.7861448 | 33.61642675 | 33.61785022 | 33.38509644 | 33.96025167 | 33.95350282 | 33.73378199 |
| Profilin-2 | P35080-2 | PFN2 | Extracellular or secreted | -0.347110079 | 0.371391506 | 28.8591984 | 28.79287264 | 28.86888353 | 28.1729321 | 28.30864893 | 28.2783039 | 28.5021074 | 28.58391165 | 27.51307047 |
| Phosphoglycerate mutase 1 | P18669 | PGAM1 | Extracellular or secreted | -0.385256134 | 1.522925242 | 32.72837882 | 32.73711093 | 32.56855563 | 32.51580296 | 32.60518682 | 32.09320436 | 32.28051798 | 32.21679738 | 31.97103601 |
| Serine/threonine-protein phosphatas | Q96HS1;Q96HS1-2 | PGAM5 | Extracellular or secreted | -3.653850562 | 2.377261965 | 24.7658247 | 25.17714029 | 24.83571544 | 25.28383274 | 21.45797864 | 26.85221916 | 21.25902479 | 20.1474146 | 21.81836441 |
| 6-phosphogluconate dehydrogenase, | P52209-2;P52209 | PGD | Extracellular or secreted | 0.84000678 | 1.801021461 | 31.78593052 | 31.87708556 | 31.8207683 | 31.22101115 | 31.17444494 | 31.89590944 | 32.28231442 | 32.30456962 | 32.82071124 |
| Geranylgeranyl transferase type-1 sub | P53609;P53609-2 | PGGT1B | Other | -0.67370868 | 0.163923552 | 25.65477215 | 25.47299048 | 24.91884395 | 22.26724031 | 21.88323303 | 25.48520554 | 20.92172966 | 24.97521929 | 24.92306774 |
| Phosphoglycerate kinase 1 | P00558;P00558-2 | PGK1 | Extracellular or secreted | -0.21707221 | 0.543200423 | 34.6698192 | 34.63547214 | 34.66325883 | 34.46447234 | 34.47647265 | 33.87325813 | 34.31257068 | 34.44022995 | 33.98735938 |

| | | | | | | | | | | | | | | |
|---|---|---|---|---|---|---|---|---|---|---|---|---|---|---|
| 6-phosphogluconolactonase | O95336 | PGLS | Extracellular or secreted | 0.288405382 | 0.477902187 | 30.503416 | 30.37906512 | 30.35688939 | 29.61911922 | 29.74275057 | 29.54932987 | 30.44628262 | 30.57965646 | 29.91456214 |
| Phosphoglucomutase-1 | P36871;P36871-2;P36871-3 | PGM1 | Extracellular or secreted | -0.040313852 | 0.03755669 | 32.72403353 | 32.88414603 | 32.85110233 | 32.02890941 | 32.06875186 | 31.91737494 | 32.67625527 | 32.68789554 | 31.75206668 |
| Phosphoglucomutase-2 | Q96G03 | PGM2 | Extracellular or secreted | -0.254827522 | 0.636448939 | 31.47720881 | 31.48769139 | 31.42327138 | 31.05394363 | 31.09917488 | 30.83364215 | 31.16943578 | 31.06483924 | 30.68870852 |
| Glucose 1,6-bisphosphate synthase | Q6PCE3 | PGM2L1 | Other | -0.770667383 | 0.572553167 | 26.91136529 | 27.06435103 | 27.03485638 | 25.4592645 | 27.32868698 | 25.92408777 | 26.20523364 | 26.48465255 | 24.85941763 |
| Phosphoacetylglucosamine mutase | O95394;O95394-3;O95394-4 | PGM3 | Other | -0.135332307 | 0.218912473 | 29.07106876 | 29.16173315 | 29.07732244 | 28.82974668 | 28.77184296 | 27.96686256 | 28.89203394 | 28.79779657 | 28.34346084 |
| Phosphoglycolate phosphatase | A6NDG6 | PGP | Other | 0.889738976 | 3.080536316 | 26.19651778 | 26.25062645 | 26.01476066 | 26.56450562 | 26.91193667 | 26.42694395 | 27.18262485 | 27.20293822 | 27.46629943 |
| Membrane-associated progesterone r | O00264;O00264-2 | PGRMC1 | Extracellular or secreted | 0.434691457 | 0.252164728 | 25.77512537 | 25.76182389 | 25.82716203 | 23.97403781 | 24.23008026 | 28.05594841 | 25.67626788 | 25.63156431 | 26.80833106 |
| Membrane-associated progesterone r | O15173;O15173-2 | PGRMC2 | Other | -2.144188448 | 0.647864193 | 25.58256284 | 26.10383605 | 25.74481105 | 26.01718443 | 26.03033761 | 26.84637225 | 21.46792986 | 24.72437188 | 25.53768503 |
| Phosphorylated adapter RNA export p | Q9H814 | PHAX | Other | 1.453457694 | 3.874292359 | 23.88846159 | 24.08788428 | 23.95898806 | 24.2286866 | 23.85804248 | 23.93111766 | 25.58313695 | 25.29659615 | 25.45723031 |
| Prohibitin | P35232 | PHB | Extracellular or secreted | -0.528304677 | 0.260496461 | 26.27401664 | 26.60045355 | 27.10639441 | 27.27408776 | 27.16307778 | 29.90426122 | 25.96331663 | 26.54169278 | 28.07122224 |
| Prohibitin-2 | Q99623;Q99623-2 | PHB2 | Extracellular or secreted | -0.614892269 | 0.28162907 | 26.68792474 | 26.40460862 | 27.263023 | 27.40843626 | 27.23908817 | 30.01757207 | 25.90444499 | 26.49277249 | 28.26843213 |
| PHD finger protein 10 | Q8WUB8-3;Q8WUB8-2;Q8W | PHF10 | Other | -0.650407368 | 0.712134951 | 23.85452753 | 22.89733439 | 23.49521484 | 21.91775564 | 23.67120068 | 24.75862541 | 22.34621829 | 22.88091673 | 23.11897211 |
| PHD finger protein 14 | O94880 | PHF14 | Other | -0.466063274 | 1.19233228 | 24.02774607 | 23.67443712 | 24.27845456 | 24.49258915 | 24.32745396 | 24.93809101 | 23.62187391 | 23.80931282 | 24.04000939 |
| Lysine-specific demethylase PHF2 | O75151 | PHF2 | Other | -2.740786977 | 2.629884224 | 23.82909944 | 23.66503138 | 23.95916499 | 23.97596269 | 23.32292387 | 23.23876391 | 20.42039929 | 21.28562303 | 21.0670899 |
| PHD finger protein 3 | Q92576-2;Q92576 | PHF3 | Other | 0.551634659 | 0.368535639 | 24.54692601 | 24.41680217 | 23.66209984 | 25.03502425 | 25.00727233 | 21.77996803 | 23.858991 | 24.72686798 | 25.29309131 |
| PHD finger-like domain-containing pro | Q7RTV0 | PHF5A | Other | -0.064445534 | 0.119865619 | 27.08506993 | 27.14335955 | 27.03705809 | 27.30120953 | 27.19901073 | 27.39607348 | 26.95659736 | 26.94380201 | 27.48715468 |
| PHD finger protein 6 | Q8IWS0-3;Q8IWS0;Q8IWS0-5 | PHF6 | Other | -0.473592016 | 1.08698607 | 26.31388998 | 25.89182541 | 25.66768633 | 26.11684047 | 26.15191787 | 27.2115041 | 25.78994749 | 25.66936347 | 25.79674507 |
| D-3-phosphoglycerate dehydrogenase | O43175 | PHGDH | Extracellular or secreted | -0.086191263 | 0.117014 | 30.45316901 | 30.52215562 | 30.61289484 | 30.7673032 | 30.82259276 | 30.96746736 | 30.22699314 | 30.52598649 | 31.06123797 |
| PH-interacting protein | Q8WWQ0 | PHIP | Extracellular or secreted | -1.625403856 | 0.626141219 | 24.15699776 | 25.25659513 | 25.37252077 | 22.08381328 | 22.23472825 | 25.31737921 | 23.24904631 | 23.57022169 | 20.51553764 |
| Phosphorylase b kinase regulatory sub | P46019 | PHKA2 | Extracellular or secreted | 0.605864265 | 0.485041671 | 20.55629434 | 22.40859822 | 22.21225769 | 22.76220147 | 22.05304062 | 21.84003565 | 22.57498611 | 23.35245514 | 21.80636554 |
| Phosphorylase b kinase regulatory sub | Q93100-4;Q93100-2;Q93100 | PHKB | Extracellular or secreted | 0.647347205 | 1.39417707 | 23.07848442 | 23.52766885 | 23.6935984 | 22.59791615 | 23.37092575 | 23.85367125 | 24.22169803 | 24.10908777 | 23.67238822 |
| Phosphorylase b kinase gamma cataly | P15735-2;P15735 | PHKG2 | Other | 1.65114107 | 1.728722265 | 23.77605466 | 22.59732515 | 23.00336556 | 21.86928085 | 20.56463346 | 23.0988462 | 24.13672645 | 24.62739539 | 23.6440543 |
| Pleckstrin homology-like domain fami | Q8WV24 | PHLDA1 | Other | 0.742756389 | 0.635004185 | 21.87346364 | 22.14317414 | 19.72893027 | 21.7778048 | 23.47022322 | 21.19937591 | 22.17038599 | 22.18213608 | 22.9722331 |
| PH domain leucine-rich repeat-contai | O60346 | PHLPP1 | Extracellular or secreted | 0.062329624 | 0.065864203 | 21.548773 | 21.46468334 | 21.43568043 | 21.8194837 | 21.92922228 | 21.3634957 | 21.96049128 | 21.95571092 | 21.05145591 |
| 14 kDa phosphohistidine phosphatase | Q9NRX4;Q9NRX4-2 | PHPT1 | Extracellular or secreted | 0.889061718 | 1.20943475 | 27.79804387 | 26.97322772 | 27.0050675 | 26.48048293 | 26.44192924 | 27.03663897 | 27.97836518 | 28.25696398 | 27.29955111 |
| Phytanoyl-CoA dioxygenase, peroxiso | O14832-2;O14832 | PHYH | Other | -1.351989722 | 1.899996994 | 24.53984557 | 24.26212697 | 24.29045709 | 23.97237335 | 23.61851724 | 23.85097225 | 23.21813927 | 22.4667047 | 22.5263331 |
| Elafin | P19957 | PI3 | Extracellular or secreted | 2.455577152 | 2.79608599 | 29.47496779 | 29.14019312 | 29.25685153 | 28.37019411 | 28.5354545 | 28.97117309 | 31.04856225 | 31.27553151 | 31.91705477 |
| Phosphatidylinositol 4-kinase alpha | P42356 | PI4KA | Other | 0.868056408 | 1.087297317 | 20.79416339 | 22.06775996 | 22.46916284 | 21.62346769 | 21.88893325 | 23.5414417 | 22.55646489 | 22.97431795 | 23.26585079 |
| Phosphatidylinositol 4-kinase beta | Q9UBF8-3;Q9UBF8-2;Q9UBF8 | PI4KB | Extracellular or secreted | -0.705031004 | 0.800610162 | 20.96754431 | 21.30401926 | 22.89417913 | 21.84982028 | 23.06724515 | 22.41564745 | 21.65210972 | 21.65399088 | 20.82803419 |
| Phosphatidylinositol-binding clathrin a | Q13492-3;Q13492-2;Q13492 | PICALM | Extracellular or secreted | 0.224460049 | 0.782831497 | 28.15976196 | 27.76477493 | 27.86325163 | 27.86053101 | 27.62282355 | 28.51818321 | 28.21465692 | 28.2707505 | 28.08263586 |
| PIH1 domain-containing protein 1 | Q9NWS0 | PIH1D1 | Other | 1.452672495 | 0.896210498 | 23.19150387 | 23.20672465 | 24.69077768 | 23.19044925 | 20.04375281 | 25.07439041 | 25.21804316 | 24.99187541 | 23.84689825 |
| Phosphatidylinositol 4,5-bisphosphate | P42336 | PIK3CA | Extracellular or secreted | 0.752467291 | 2.137234779 | 22.06969494 | 21.32262967 | 21.77258518 | 22.60764204 | 21.8792263 | 22.00959766 | 22.75680469 | 22.84113049 | 22.49015459 |
| Phosphatidylinositol 3-kinase regulato | O00459 | PIK3R2 | Other | 0.308251401 | 0.146842545 | 21.56175959 | 26.09641517 | 26.1911612 | 26.00859788 | 25.62005668 | 26.61694812 | 25.54792627 | 26.06773944 | 25.35655782 |
| Phosphoinositide 3-kinase regulatory | Q99570 | PIK3R4 | Other | 0.485849264 | 0.644103237 | 23.07200369 | 23.35568446 | 23.40184511 | 21.53574932 | 23.06109751 | 23.6907244 | 23.48554338 | 23.84957309 | 23.18098357 |
| Peptidyl-prolyl cis-trans isomerase NI | Q13526 | PIN1 | Other | 0.500127289 | 1.180420042 | 26.19134949 | 25.60331484 | 25.99776542 | 26.18090013 | 26.42922883 | 27.00592415 | 26.63075926 | 26.98923545 | 26.58462859 |
| Peptidyl-prolyl cis-trans isomerase NI | Q9Y237;Q9Y237-2 | PIN4 | Extracellular or secreted | -0.099970785 | 0.122963341 | 28.80928215 | 28.84323019 | 28.7210044 | 28.94634228 | 28.83077442 | 28.39398904 | 28.94159308 | 28.91743896 | 28.11336685 |
| Prolactin-inducible protein | P12273 | PIP | Extracellular or secreted | -1.529073006 | 1.492307204 | 22.34253373 | 21.627785 | 22.51869349 | 24.29017582 | 24.44347932 | 21.56917882 | 20.87117129 | 21.53999502 | 21.39753776 |
| Phosphatidylinositol 5-phosphate 4-ki | Q8TBX8-3;Q8TBX8 | PIP4K2C | Extracellular or secreted | 0.33397858 | 0.916723103 | 25.22188238 | 25.48323838 | 25.36185384 | 25.24495955 | 25.72020208 | 25.26678025 | 25.93898835 | 25.7558018 | 25.45660383 |
| Pirin | O00625 | PIR | Other | 0.84206424 | 2.848466251 | 27.34087464 | 27.46847626 | 27.3220815 | 27.53200188 | 27.40511198 | 26.56425848 | 28.18702209 | 28.18711651 | 27.96845648 |
| PITH domain-containing protein 1 | Q9GZP4-2;Q9GZP4 | PITHD1 | Other | -0.275127308 | 0.252759312 | 26.18198069 | 25.35779977 | 25.17577064 | 25.98602637 | 26.59077959 | 25.82021442 | 25.17161571 | 25.24778673 | 26.31150138 |
| Phosphatidylinositol transfer protein a | Q00169 | PITPNA | Extracellular or secreted | -0.068984814 | 0.173225157 | 27.75919708 | 27.85620295 | 27.83056288 | 27.45144067 | 27.49411629 | 27.28082768 | 27.60907681 | 27.70668497 | 27.31345755 |
| Phosphatidylinositol transfer protein | P48739;P48739-2;P48739-3 | PITPNB | Extracellular or secreted | 0.255325774 | 0.935323307 | 29.80611966 | 29.72269889 | 29.66966083 | 29.27001527 | 29.25943554 | 29.43603007 | 29.74401866 | 29.96801695 | 29.63592183 |
| Presequence protease, mitochondrial | Q5JRX3;Q5JRX3-2;Q5JRX3-3 | PITRM1 | Other | -0.063300555 | 0.076686602 | 28.41021686 | 28.60918252 | 28.61708829 | 28.5987156 | 28.50017077 | 28.92462016 | 28.7891453 | 28.83915029 | 28.01179984 |
| cAMP-dependent protein kinase inhib | Q9C010;Q9C010-2 | PKIB | Other | 0.525471301 | 0.263288222 | 22.91219027 | 22.43612991 | 19.93057787 | 22.43670016 | 20.82445764 | 21.91571264 | 22.63506799 | 23.16361528 | 21.00561488 |
| Pyruvate kinase PKM | P14618;P14618-3 | PKM | Extracellular or secreted | 0.071405037 | 0.324701064 | 33.70594384 | 33.78292722 | 33.73721169 | 33.56841973 | 33.65725435 | 33.66246322 | 33.85117681 | 33.82202286 | 33.59812546 |
| Serine/threonine-protein kinase N1 | Q16512;Q16512-2 | PKN1 | Extracellular or secreted | 0.101319746 | 0.142837026 | 24.20291954 | 24.30910882 | 23.91369513 | 24.38159206 | 24.04551905 | 23.91342122 | 24.1052557 | 23.89217295 | 24.6896585 |
| Plakophilin-2 | Q99959-2;Q99959 | PKP2 | Extracellular or secreted | 0.205667502 | 0.122026823 | 24.89578235 | 25.27813549 | 22.36741218 | 22.41589877 | 24.56639414 | 24.93606996 | 24.46344904 | 23.64107978 | 24.74232013 |
| Plakophilin-3 | Q9Y446;Q9Y446-2 | PKP3 | Extracellular or secreted | 0.080154631 | 0.082710689 | 27.49678492 | 27.68885856 | 27.51990843 | 25.95358121 | 26.81421162 | 27.6759988 | 27.40681562 | 26.85447998 | 27.55384007 |
| Group XV phospholipase A2 | Q8NCC3;Q8NCC3-2 | PLA2G15 | Extracellular or secreted | 0.103660961 | 0.461645785 | 25.03967479 | 25.28107539 | 25.25313595 | 25.11537265 | 25.04330937 | 24.90439905 | 25.14654817 | 25.11672152 | 25.3661968 |
| Phospholipase A-2-activating protein | Q9Y263 | PLAA | Extracellular or secreted | -0.011699557 | 0.012043999 | 28.40952916 | 28.50967704 | 28.47649438 | 28.74432348 | 28.83424516 | 28.38640248 | 28.79711627 | 28.89113589 | 27.95698501 |
| Tissue-type plasminogen activator;Tis | P00750;P00750-3;P00750-4 | PLAT | Extracellular or secreted | 1.702710859 | 0.802655691 | 25.91195953 | 26.18482042 | 26.11559093 | 25.26660156 | 25.34702304 | 28.98980388 | 27.17997058 | 27.21808013 | 29.61798155 |
| Urokinase plasminogen activator surfa | Q03405;Q03405-2;Q03405-3 | PLAUR | Extracellular or secreted | 0.554523292 | 0.338755093 | 26.37326783 | 26.17943913 | 26.53528392 | 26.71187867 | 26.72832205 | 27.69258783 | 26.66438044 | 26.65335134 | 28.45622781 |
| Putative phospholipase B-like 2;Putat | Q8NHP8;Q8NHP8-2 | PLBD2 | Extracellular or secreted | 0.315918457 | 0.492569548 | 26.35240463 | 26.33914226 | 26.2684946 | 26.87150033 | 27.01275904 | 25.31240189 | 26.49486401 | 26.52211071 | 27.00913204 |
| 1-phosphatidylinositol 4,5-bisphospha | Q01970;Q01970-2 | PLCB3 | Extracellular or secreted | -0.384546927 | 2.257875747 | 30.05866012 | 30.16722384 | 30.13577288 | 29.78359863 | 29.69047465 | 29.75692491 | 29.60151657 | 29.56900855 | 29.47216161 |
| 1-phosphatidylinositol 4,5-bisphospha | Q15147;Q15147-5;Q15147-4 | PLCB4 | Other | -0.829523928 | 0.893701872 | 24.26549315 | 24.32347373 | 24.31610196 | 22.84806432 | 23.02863994 | 23.25666711 | 23.03508299 | 23.32649422 | 22.1690711 |
| 1-phosphatidylinositol 4,5-bisphospha | P51178;P51178-2 | PLCD1 | Extracellular or secreted | -0.765276181 | 1.645055032 | 23.4898362 | 22.17554224 | 22.58070113 | 23.72426528 | 23.29050874 | 23.12237304 | 22.1911725 | 22.22843707 | 22.47617521 |
| 1-phosphatidylinositol 4,5-bisphospha | P19174;P19174-2 | PLCG1 | Extracellular or secreted | -0.406247984 | 1.204388261 | 26.57428458 | 26.62811887 | 26.57344691 | 26.73711899 | 26.71384607 | 26.50033035 | 26.28256077 | 26.37565582 | 25.98661234 |
| 1-phosphatidylinositol 4,5-bisphospha | P16885 | PLCG2 | Extracellular or secreted | -0.839484497 | 0.363992823 | 24.20545739 | 24.12573444 | 24.60325823 | 23.51773282 | 21.65406084 | 23.57692438 | 23.41351327 | 23.78186794 | 21.12774935 |
| Phospholipase D1 | Q13393-2;Q13393;Q13393-4 | PLD1 | Extracellular or secreted | 3.890916546 | 1.816531112 | 20.85680177 | 20.09920177 | 21.4478439 | 22.61862249 | 22.42774432 | 25.28535057 | 25.69226856 | 27.55340067 | 24.79486282 |
| Phospholipase D3 | Q8IV08 | PLD3 | Extracellular or secreted | -0.07951107 | 0.289040386 | 24.62483248 | 24.85195716 | 25.18517972 | 25.02635357 | 24.99005825 | 24.57732786 | 24.7159155 | 24.75378884 | 24.91961697 |
| Plectin | Q15149-2;Q15149-7;Q15149 | PLEC | Extracellular or secreted | -0.561022241 | 0.857504884 | 23.65348801 | 24.20366643 | 24.15344565 | 23.490326 | 24.03975845 | 23.9990352 | 23.29949871 | 22.9621695 | 23.82512494 |
| Plectin | Q15149-3 | PLEC | Extracellular or secreted | 0.29108143 | 0.176794696 | 24.28602073 | 22.42470349 | 23.62477672 | 20.67988712 | 20.71565014 | 23.54533594 | 22.80785958 | 22.78931075 | 22.91426103 |
| Plectin | Q15149-4 | PLEC | Extracellular or secreted | -1.067302515 | 1.083089904 | 33.86193632 | 33.89428387 | 33.8614742 | 32.75372111 | 32.79445772 | 33.93420781 | 32.03311754 | 32.10543306 | 33.20958237 |
| Pleckstrin homology domain-containi | Q9HAU0-5;Q9HAU0-8;Q9HAU | PLEKHA5 | Other | -0.587982187 | 0.387194956 | 23.64008691 | 23.74221732 | 23.86985436 | 21.94909441 | 23.1260339 | 23.8406696 | 23.35796752 | 23.27923423 | 21.68282994 |
| Perilipin-3 | O60664-4;O60664;O60664-3 | PLIN3 | Other | 0.409587302 | 1.643145599 | 28.51945857 | 28.43511607 | 28.46964107 | 28.52230531 | 28.47734393 | 29.24678607 | 28.96611996 | 29.00295048 | 29.09501698 |
| Serine/threonine-protein kinase PLK1 | P53350 | PLK1 | Other | -0.32725643 | 0.625360593 | 23.54616064 | 22.72129782 | 23.06289464 | 23.39636751 | 24.01358984 | 24.08285916 | 23.02270983 | 23.40418631 | 23.00291937 |
| Procollagen-lysine,2-oxoglutarate 5-d | Q02809;Q02809-2 | PLOD1 | Extracellular or secreted | -0.164728447 | 0.202398432 | 27.36794698 | 27.38993417 | 27.46598819 | 27.88092841 | 27.86059021 | 28.49174853 | 27.26669091 | 27.37509234 | 28.09259965 |
| Procollagen-lysine,2-oxoglutarate 5-d | O60568 | PLOD3 | Extracellular or secreted | 0.394754099 | 0.641709655 | 30.25146645 | 30.26539478 | 30.18659714 | 30.2147727 | 30.3215225 | 30.81360018 | 30.5202832 | 30.48366124 | 31.20699473 |
| Proteolipid protein 2 | Q04941;Q04941-2 | PLP2 | Extracellular or secreted | 3.306360809 | 2.055292494 | 22.16217523 | 21.12635297 | 21.73647329 | 20.38263049 | 22.21417418 | 26.12534031 | 25.9097642 | 24.8381225 | 26.04476897 |
| Pleiotropic regulator 1 | O43660;O43660-2 | PLRG1 | Other | -1.101472393 | 1.806163343 | 29.38329484 | 29.42961994 | 29.34502726 | 29.72201482 | 29.80093107 | 29.28995604 | 28.27626394 | 28.13437806 | 28.7703628 |
| Plastin-1 | Q14651 | PLS1 | Extracellular or secreted | 0.109766264 | 0.130173158 | 30.08765165 | 30.02221032 | 30.03721522 | 29.47988959 | 29.46113577 | 29.16916097 | 29.96306302 | 30.18541605 | 29.30945148 |
| Plastin-3 | P13797;P13797-2;P13797-3 | PLS3 | Other | 0.374308006 | 1.474992543 | 30.24240853 | 30.31915514 | 30.2997476 | 30.3243153 | 30.27941136 | 30.28482569 | 30.53398907 | 30.78830533 | 30.67556143 |
| Phospholipid scramblase 1 | O15162-2;O15162 | PLSCR1 | Extracellular or secreted | -0.324389134 | 0.294926202 | 24.31817261 | 24.40073823 | 24.76612811 | 24.3965635 | 24.19798034 | 23.20097581 | 24.32333629 | 24.2128959 | 23.1308797 |
| Plexin domain-containing protein 2 | Q6UX71;Q6UX71-2 | PLXDC2 | Extracellular or secreted | -0.778036455 | 0.643867165 | 25.74352761 | 25.78409184 | 25.84160562 | 25.68757773 | 25.96344893 | 26.36516236 | 24.50306336 | 24.84330808 | 26.01222623 |
| Plexin-A1 | Q9UIW2 | PLXNA1 | Extracellular or secreted | 1.179582263 | 1.281088628 | 23.34285922 | 23.02501043 | 23.46749709 | 22.61557986 | 20.6521817 | 23.84239715 | 23.88576488 | 24.07806088 | 24.04768376 |
| Plexin-A3 | P51805 | PLXNA3 | Extracellular or secreted | -0.14679538 | 0.093193663 | 23.02003246 | 23.73696422 | 23.17377472 | 20.87167109 | 22.66078806 | 22.98260998 | 23.08906507 | 22.91152073 | 21.78194833 |
| Plexin-B1 | O43157-2;O43157 | PLXNB1 | Extracellular or secreted | 0.192468422 | 0.532026125 | 25.39322809 | 25.52548998 | 25.83809845 | 25.89015608 | 25.86646244 | 25.99841121 | 25.7184524 | 25.94966858 | 26.16520742 |
| Plexin-B2 | O15031 | PLXNB2 | Extracellular or secreted | -0.147319578 | 0.461177021 | 27.63311763 | 27.60724338 | 27.53051433 | 28.15716164 | 27.97465055 | 28.20573703 | 27.63193893 | 27.61617695 | 27.86413768 |
| Peptidase M20 domain-containing pro | Q8IYS1 | PM20D2 | Extracellular or secreted | 0.655504413 | 0.706231548 | 28.93429921 | 28.84074163 | 28.83267681 | 28.47452303 | 28.53945366 | 27.74432348 | 29.44260958 | 29.68809153 | 28.51882103 |
| Polyamine-modulated factor 1 | Q6P1K2;Q6P1K2-4;Q6P1K2-2 | PMF1 | Other | -0.429649836 | 0.531538206 | 23.68378194 | 23.95118154 | 23.69476765 | 23.17092078 | 24.32264887 | 23.29080859 | 23.37901552 | 23.7040877 | 22.68500196 |
| Phosphomannomutase 2 | O15305 | PMM2 | Extracellular or secreted | 0.309423858 | 1.242249436 | 28.34861936 | 28.27564251 | 28.34726809 | 28.53549158 | 28.40645072 | 28.33611411 | 28.62076288 | 28.86088616 | 28.57141572 |
| Mitochondrial-processing peptidase s | Q10713;Q10713-2 | PMPCA | Extracellular or secreted | 0.215126127 | 0.338345469 | 25.27980099 | 24.79466454 | 25.48173046 | 25.24977693 | 26.06899012 | 26.16255928 | 25.49481824 | 26.05593807 | 25.61338324 |
| Mitochondrial-processing peptidase s | O75439 | PMPCB | Other | 0.101667316 | 0.108989815 | 26.00315964 | 26.1956921 | 26.16920876 | 26.63768465 | 26.45849811 | 26.34769895 | 26.70305804 | 26.72142821 | 25.78648681 |
| Phosphomevalonate kinase | Q15126 | PMVK | Extracellular or secreted | 0.99205583 | 1.359029319 | 24.666224 | 24.42123994 | 24.52384619 | 23.71261348 | 23.57600172 | 25.90391659 | 25.61675186 | 25.73249435 | 25.02884225 |
| Bifunctional polynucleotide phosphat | Q96T60;Q96T60-2 | PNKP | Other | -0.174774219 | 0.137786844 | 24.66806521 | 23.19772543 | 22.24683709 | 21.83076394 | 21.98975361 | 23.51460619 | 22.91887124 | 22.37798362 | 22.90269822 |
| Pinin | Q9H307 | PNN | Extracellular or secreted | -1.070279389 | 2.059684451 | 28.17959099 | 27.86673362 | 28.02543505 | 27.13868053 | 27.15624559 | 28.00404958 | 26.42500776 | 26.50567008 | 27.04385168 |
| RNA-binding protein PNO1 | Q9NRX1 | PNO1 | Other | 1.117349915 | 1.765398853 | 23.58634779 | 23.26642284 | 23.15174372 | 22.20415246 | 21.69559091 | 23.60964047 | 24.08310271 | 24.10237495 | 23.92352118 |
| Purine nucleoside phosphorylase | P00491 | PNP | Extracellular or secreted | 0.30283259 | 1.235904993 | 32.25673346 | 32.494741 | 32.38339785 | 32.21827071 | 32.21997328 | 31.63248506 | 32.51493345 | 32.41704571 | 32.57931928 |
| Pyridoxine-5-phosphate oxidase | Q9NVS9;Q9NVS9-3;Q9NVS9- | PNPO | Extracellular or secreted | 0.203503103 | 1.721147092 | 27.64065238 | 27.72507434 | 27.57629012 | 27.52976999 | 27.62672725 | 27.64917645 | 27.77379318 | 27.79742554 | 27.91313585 |
| Polyribonucleotide nucleotidyltransfe | Q8TCS8 | PNPT1 | Other | -0.357602031 | 0.567642315 | 26.66125728 | 26.68525332 | 26.76306584 | 27.60526632 | 27.41134883 | 28.26579715 | 26.77379318 | 26.69856119 | 27.15083391 |
| Podocalyxin | O00592-2;O00592 | PODXL | Extracellular or secreted | 0.32329806 | 1.746473029 | 26.12255867 | 25.94752993 | 25.88019278 | 25.46970317 | 25.78206798 | 26.04456055 | 26.24147233 | 26.17371373 | 26.17801466 |
| Podocalyxin-like protein 2 | Q9NZ53 | PODXL2 | Extracellular or secreted | 0.364471816 | 0.181010828 | 20.0653951 | 19.7602164 | 20.20752915 | 21.10559141 | 24.05609938 | 20.72117264 | 20.64869111 | 21.18978596 | 22.21294041 |
| GDP-fucose protein O-fucosyltransfer | Q9H488 | POFUT1 | Extracellular or secreted | -0.280283557 | 1.443095533 | 28.04092913 | 28.13868053 | 28.13574842 | 27.66132525 | 27.7619887 | 28.10998444 | 27.72142821 | 27.79153812 | 27.57051124 |
| GDP-fucose protein O-fucosyltransfer | Q9Y2G5-2;Q9Y2G5-1;Q9Y2G5 | POFUT2 | Extracellular or secreted | -1.603041982 | 1.173180823 | 23.90517984 | 23.53433423 | 23.72317429 | 23.18286062 | 22.57857749 | 24.28171217 | 20.99754994 | 22.75397242 | 22.04227102 |
| Pogo transposable element with KRAB | Q9P215 | POGK | Other | 0.876185126 | 1.580879466 | 23.19709548 | 22.22550602 | 21.29554299 | 22.69538874 | 22.81187093 | 23.01295933 | 23.46861609 | 23.30479915 | 23.47432188 |
| Protein O-glucosyltransferase 1 | Q8NBL1 | POGLUT1 | Extracellular or secreted | 0.046556634 | 0.121344665 | 25.24572126 | 25.44104273 | 24.91437966 | 25.48431451 | 25.42965984 | 25.542121 | 25.44521722 | 25.53857332 | 25.18449887 |
| Pogo transposable element with ZNF | Q7Z3K3-5;Q7Z3K3-7;Q7Z3K3 | POGZ | Other | -0.633769046 | 0.537484861 | 22.66758889 | 22.92603968 | 22.3667971 | 22.77868356 | 23.39401358 | 24.24372546 | 22.63276273 | 21.59589542 | 23.05845885 |
| DNA polymerase alpha catalytic subun | P09884 | POLA1 | Other | 0.199502784 | 0.542643622 | 27.81799678 | 27.45159787 | 27.5724642 | 27.8148228 | 27.71443577 | 28.02284526 | 27.9942084 | 28.13050466 | 27.67087664 |
| DNA polymerase alpha subunit B | Q14181;Q14181-2 | POLA2 | Other | 0.785344369 | 1.603868213 | 24.29452933 | 24.6258359 | 25.17607512 | 24.49405524 | 25.44287847 | 25.43523532 | 26.02648022 | 25.63378282 | 25.43007476 |
| DNA polymerase delta catalytic subun | P28340 | POLD1 | Other | -0.55794792 | 2.362661906 | 25.79934151 | 25.53960896 | 25.57954498 | 26.02559346 | 25.87759756 | 26.2274385 | 25.43033003 | 25.26223452 | 25.15815416 |
| DNA polymerase delta subunit 2 | P49005 | POLD2 | Other | -0.13803551 | 0.183613887 | 26.12662083 | 26.18574686 | 26.15686278 | 26.15682422 | 25.98233144 | 25.78054197 | 26.37320144 | 25.94246028 | 25.4646958 |
| DNA polymerase delta subunit 3 | Q15054-3;Q15054-2;Q15054 | POLD3 | Other | -0.390184407 | 0.616039186 | 24.12589206 | 24.27334088 | 24.05034357 | 25.04042753 | 25.64823029 | 25.39584475 | 24.19520397 | 24.35494504 | 24.5463373 |
| Polymerase delta-interacting protein | Q9Y2S7 | POLDIP2 | Other | -0.318084253 | 0.637182508 | 25.84227725 | 25.90244523 | 26.02045219 | 26.24974077 | 26.49574855 | 27.07398201 | 26.15060153 | 26.03176833 | 25.65570038 |
| Polymerase delta-interacting protein | Q9BY77-2;Q9BY77 | POLDIP3 | Other | -0.514881149 | 2.013147418 | 23.58221826 | 23.8316142 | 23.26956503 | 23.41028157 | 23.23748074 | 24.01665325 | 23.17081384 | 23.01857307 | 22.93987617 |
| DNA polymerase epsilon subunit 3 | Q9NRF9 | POLE3 | Other | 0.456786061 | 0.665578624 | 26.68859182 | 26.90563894 | 27.07019876 | 26.52460855 | 26.50728932 | 25.60189907 | 27.22637309 | 27.27506522 | 26.5180331 |
| DNA-directed RNA polymerase I subu | O95602 | POLR1A | Other | 0.284742858 | 0.667126567 | 26.20155546 | 25.91738774 | 25.97795049 | 25.73817619 | 25.76913343 | 26.91547423 | 26.30876174 | 26.22455272 | 26.58075288 |
| DNA-directed RNA polymerase I subu | Q9H9Y6;Q9H9Y6-3;Q9H9Y6-2 | POLR1B | Other | 0.191857156 | 0.481270547 | 24.88091673 | 25.18389339 | 25.48222302 | 25.67022834 | 24.86994847 | 25.89471898 | 25.41641563 | 25.55438181 | 25.59573849 |
| DNA-directed RNA polymerases I and | O15160-2;O15160 | POLR1C | Other | 0.282607805 | 0.575748582 | 27.08628543 | 26.92603515 | 27.04603971 | 26.98966856 | 26.80501268 | 27.29465202 | 27.51472657 | 27.47289755 | 26.93404607 |
| DNA-directed RNA polymerase I subu | Q9GZS1-2;Q9GZS1 | POLR1E | Other | 0.250819323 | 0.067403546 | 22.31665499 | 22.49707765 | 21.09082389 | 20.90061696 | 21.21539153 | 24.2775326 | 23.86994847 | 20.05096894 | 22.98058937 |
| DNA-directed RNA polymerase II subu | P24928 | POLR2A | Other | -0.184450068 | 0.579402732 | 27.96069012 | 27.79779657 | 27.90821861 | 27.54576307 | 27.53103515 | 28.35573486 | 27.75754495 | 27.74483671 | 27.49388732 |
| DNA-directed RNA polymerase II subu | P30876 | POLR2B | Other | -0.09079381 | 0.218097979 | 27.58527353 | 27.69743478 | 27.5824767 | 27.12405866 | 27.13848524 | 27.80974213 | 27.53066316 | 27.49944861 | 27.16624232 |
| DNA-directed RNA polymerase II subu | P19387 | POLR2C | Other | 0.089676586 | 0.115011554 | 25.88185034 | 25.52817579 | 25.44616428 | 25.27433663 | 25.09365773 | 26.06804709 | 25.89774521 | 25.8548604 | 25.16254008 |
| DNA-directed RNA polymerase II subu | O15514 | POLR2D | Other | -0.287784637 | 0.318474394 | 26.26243168 | 26.6896585 | 26.45560089 | 25.91868475 | 26.16939994 | 24.92370251 | 26.11366492 | 26.01109602 | 25.22162429 |
| DNA-directed RNA polymerases I, II, a | P19388 | POLR2E | Other | -0.126447944 | 0.701340466 | 27.74470842 | 28.07270499 | 27.73724796 | 27.91136529 | 27.99937935 | 28.22697018 | 27.8312276 | 27.89116487 | 27.7444518 |
| DNA-directed RNA polymerase II subu | P62487 | POLR2G | Other | 0.221659841 | 0.553862283 | 26.92150229 | 27.45120482 | 27.39460242 | 26.77655673 | 26.46653282 | 27.23598834 | 27.32448699 | 27.40438124 | 27.05930501 |
| DNA-directed RNA polymerases I, II, a | P52434;P52434-3;P52434-4 | POLR2H | Other | 0.622737077 | 0.628265969 | 28.78203673 | 28.85061064 | 28.84077163 | 28.31272213 | 28.69155659 | 27.05299848 | 29.34036533 | 29.45840032 | 28.33479368 |
| DNA-directed RNA polymerase II subu | P52435;Q9H1A7;Q9GZM3 | POLR2J;POLR2J3; | Other | 1.078753303 | 1.367453124 | 22.40874519 | 21.66758616 | 19.82046551 | 21.93231091 | 22.45259566 | 21.23193715 | 22.85351897 | 22.722424 | 22.41713723 |
| DNA-directed RNA polymerases I, II, a | P62875 | POLR2L | Other | 0.024075662 | 0.037349653 | 24.68747094 | 24.51532832 | 24.63649585 | 24.00273056 | 24.30612123 | 24.82682271 | 24.43319879 | 24.86768822 | 24.25882478 |
| DNA-directed RNA polymerase III sub | O14802 | POLR3A | Other | -0.127104611 | 0.442314 | 25.26867306 | 25.10809082 | 25.12159074 | 24.53901725 | 24.97408159 | 25.20635204 | 25.05262543 | 24.86334616 | 24.81161733 |
| DNA-directed RNA polymerase III sub | Q9BUI4 | POLR3C | Other | 0.181762655 | 0.314463687 | 26.27602427 | 26.34172308 | 26.43676084 | 25.83699168 | 25.93152348 | 26.41247992 | 26.64636358 | 26.5323438 | 25.98433223 |
| DNA-directed RNA polymerase III sub | P05423 | POLR3D | Other | 0.492289885 | 1.748993875 | 22.70838783 | 23.58026409 | 23.45022959 | 23.04421867 | 23.57876794 | 23.27440773 | 23.7234868 | 23.66166503 | 23.90985574 |
| DNA-directed RNA polymerase III sub | Q9H1D9 | POLR3F | Other | 0.457947416 | 0.476782237 | 23.57045333 | 23.45838859 | 20.98632483 | 23.17535949 | 22.84714683 | 23.87464597 | 23.56697472 | 23.32429813 | 23.43872892 |
| DNA-directed RNA polymerase III sub | Q9Y535 | POLR3H | Other | 0.249400759 | 0.069808412 | 21.56453469 | 22.33369041 | 22.0153606 | 21.91672458 | 21.5687187 | 21.2036515 | 21.5561975 | 20.26245782 | 24.23088719 |
| Ribonucleases P/MRP protein subunit | Q99575 | POP1 | Extracellular or secreted | 0.631307998 | 1.318684313 | 25.58098284 | 25.25655914 | 25.31188244 | 25.18476368 | 24.35971096 | 26.00966599 | 25.96276524 | 26.13471098 | 25.64823029 |
| Ribonuclease P protein subunit p29 | O95707 | POP4 | Other | 0.506413003 | 0.463427776 | 22.91404985 | 21.65989892 | 20.94549528 | 22.61089227 | 21.16844299 | 23.14659478 | 23.11044253 | 22.61299535 | 22.01848818 |
| Ribonuclease P protein subunit p20 | O75817 | POP7 | Other | 0.484891103 | 0.407491218 | 23.71722374 | 21.28344215 | 21.36305563 | 23.88343607 | 23.29963845 | 23.99636522 | 23.32868698 | 23.50093814 | 23.39662882 |
| NADPH--cytochrome P450 reductase | P16435 | POR | Other | -1.33267665 | 0.40866141 | 24.60450301 | 24.55098158 | 24.86145423 | 23.14892338 | 23.74170315 | 25.7509813 | 20.67784626 | 24.16982043 | 24.48357668 |
| POU domain, class 2, transcription fac | P14859-2;P14859-4;P14859-5 | POU2F1 | Other | -1.899488359 | 1.295687457 | 25.01720567 | 24.87890744 | 24.50112042 | 24.96820377 | 25.20187324 | 24.36285722 | 23.20160407 | 21.9787815 | 23.58623324 |
| Inorganic pyrophosphatase | Q15181 | PPA1 | Extracellular or secreted | 0.537798205 | 1.906688633 | 31.54452919 | 31.46951493 | 31.39744103 | 31.37528917 | 31.4213162 | 31.81741744 | 32.02617413 | 32.22460907 | 31.87536539 |
| Inorganic pyrophosphatase 2, mitoch | Q9H2U2;Q9H2U2-2;Q9H2U2 | PPA2 | Extracellular or secreted | -0.083747065 | 0.655311131 | 28.85486634 | 28.67033641 | 28.68244301 | 28.94826545 | 28.95163178 | 29.01446272 | 28.77354169 | 28.77737222 | 28.75884775 |
| Suppressor of SWI4 1 homolog | Q9NQ55-2;Q9NQ55;Q9NQ55 | PPAN | Other | -0.640055741 | 0.8984889 | 22.17816842 | 23.69413 | 23.54179616 | 23.09746443 | 23.1485976 | 24.73407201 | 22.54997169 | 22.90394876 | 22.82302664 |
| Lipid phosphate phosphohydrolase 2 | O43688;O43688-2;O43688-3 | PPAP2C | Extracellular or secreted | 1.709713001 | 2.33928981 | 24.26756625 | 24.60489885 | 25.0115866 | 24.03405874 | 24.08342739 | 26.11493598 | 26.01830988 | 26.2938105 | 26.87525552 |
| Amidophosphoribosyltransferase | Q06203 | PPAT | Other | 0.255293437 | 1.00995232 | 30.02221032 | 30.0126525 | 30.02815196 | 30.03301905 | 30.04583147 | 29.56940701 | 30.29197674 | 30.3146248 | 30.01491492 |
| Phosphopantothenoylcysteine decarb | Q96CD2;Q96CD2-2 | PPCDC | Other | 0.181880965 | 0.467456675 | 25.94817631 | 25.72881506 | 25.71460608 | 25.2546864 | 25.25850134 | 24.95996091 | 25.69173628 | 25.79659656 | 25.48968311 |
| Phosphopantothenate--cysteine ligase | Q9HAB8;Q9HAB8-2 | PPCS | Extracellular or secreted | 0.594358482 | 2.209344352 | 27.17674094 | 27.63186956 | 27.65717341 | 26.83243538 | 27.05568988 | 27.45974916 | 27.81262133 | 27.9281268 | 27.94915648 |
| Liprin-alpha-1 | Q13136-2;Q13136 | PPFIA1 | Extracellular or secreted | -0.220612465 | 0.344330378 | 24.36566298 | 24.53029107 | 24.71392471 | 25.0117572 | 25.22960364 | 24.27653906 | 24.11727657 | 24.41821859 | 24.86655677 |
| Liprin-beta-1 | Q86W92;Q86W92-3;Q86W92 | PPFIBP1 | Extracellular or secreted | 0.090195086 | 0.210273244 | 26.31994756 | 26.36011299 | 25.93573284 | 25.77323984 | 25.57649196 | 26.60413535 | 26.26740551 | 26.07541092 | 26.21259909 |
| Periphilin-1 | Q8NEY8-6;Q8NEY8-9;Q8NEY8 | PPHLN1 | Other | 1.020745545 | 1.040709488 | 23.07076956 | 20.39498312 | 22.30703216 | 23.31437414 | 23.23437014 | 22.07394933 | 23.09873378 | 23.22823154 | 23.93301054 |
| Peptidyl-prolyl cis-trans isomerase A;P | P62937 | PPIA | Extracellular or secreted | 0.367061188 | 2.286426966 | 33.60730512 | 33.68614295 | 33.65170593 | 33.28512895 | 33.31351702 | 33.44393868 | 33.75463721 | 33.92193849 | 33.91847719 |
| Peptidyl-prolyl cis-trans isomerase B | P23284 | PPIB | Extracellular or secreted | -0.18460588 | 2.094946021 | 32.08850487 | 32.20995995 | 32.18047455 | 31.9433549 | 31.97137868 | 32.12694078 | 31.86030899 | 31.91067218 | 31.93550806 |
| Peptidyl-prolyl cis-trans isomerase C | P45877 | PPIC | Extracellular or secreted | 0.228693741 | 0.857549222 | 24.47382662 | 24.7678462 | 24.94127404 | 24.47171986 | 24.42252371 | 24.83518534 | 24.73412371 | 25.06076914 | 24.84737626 |
| Peptidyl-prolyl cis-trans isomerase D | Q08752 | PPID | Other | 0.317244583 | 0.607854626 | 25.34411301 | 25.23702844 | 25.20906986 | 24.83349739 | 24.69561742 | 25.78808125 | 25.84016534 | 25.45211654 | 25.21315555 |
| Peptidyl-prolyl cis-trans isomerase E | Q9UNP9;Q9UNP9-2;Q9UNP9 | PPIE | Other | -0.401017328 | 0.850147352 | 28.8482542 | 28.83866954 | 28.90069603 | 29.015686 | 29.03052179 | 27.90207715 | 28.54447392 | 28.48604231 | 28.03438415 |

| | | | | | | | | | | | | | |
|---|---|---|---|---|---|---|---|---|---|---|---|---|---|
| Peptidyl-prolyl cis-trans isomerase F, | P30405;P30405-2 | PPIF | Other | 0.122599475 | 0.05159505 | 27.91347829 | 27.63975581 | 27.28709586 | 27.88227026 | 27.7852524 | 25.41486843 | 28.16959109 | 28.12548812 | 26.03407974 |
| Peptidyl-prolyl cis-trans isomerase G | Q13427;Q13427-2 | PPIG | Other | -1.12456036 | 0.257357176 | 26.73789263 | 22.22031101 | 21.93019316 | 26.02555122 | 25.23786727 | 25.54651394 | 22.7308301 | 26.36862981 | 21.37602362 |
| Peptidyl-prolyl cis-trans isomerase H | O43447;O43447-2 | PPIH | Other | 0.174614601 | 0.305201191 | 28.77291276 | 28.60629049 | 28.46092103 | 28.18125093 | 28.18191437 | 28.50922398 | 28.70928416 | 28.93027182 | 28.24054461 |
| Peptidyl-prolyl cis-trans isomerase-lik | Q9Y3C6 | PPIL1 | Extracellular or secreted | 0.225753483 | 0.804696753 | 30.37803159 | 30.38535362 | 30.30465121 | 29.92903035 | 30.14284852 | 30.02960065 | 30.61412431 | 30.39846075 | 30.24943337 |
| Peptidyl-prolyl cis-trans isomerase-lik | Q9H2H8;Q9H2H8-2 | PPIL3 | Other | 0.257474442 | 0.505367228 | 27.02411431 | 26.83412457 | 26.78618765 | 27.17245528 | 27.30617339 | 26.00470327 | 27.31129349 | 27.18895641 | 26.83605267 |
| Peptidyl-prolyl cis-trans isomerase-lik | Q8WUA2 | PPIL4 | Other | 0.020816634 | 0.066836789 | 27.2514213 | 27.07989261 | 27.13398629 | 27.2337048 | 27.09254924 | 27.25862713 | 27.39329356 | 27.0844618 | 27.10978523 |
| Periplakin | O60437 | PPL | Extracellular or secreted | 0.641713985 | 1.203892035 | 28.75388446 | 28.67865381 | 28.62167133 | 27.94731812 | 27.90827588 | 28.47622396 | 28.76957486 | 28.90454836 | 29.44403251 |
| Protein phosphatase 1A | P35813;P35813-3;P35813-2 | PPM1A | Other | -0.0639322 | 0.084402887 | 25.09587215 | 25.16813772 | 24.92129798 | 25.91499546 | 25.30535596 | 25.38284563 | 24.93836027 | 25.67965405 | 25.08444153 |
| Protein phosphatase 1B | O75688;O75688-4;O75688-2 | PPM1B | Other | 0.046500367 | 0.074913542 | 27.55691213 | 27.59754396 | 27.48123774 | 27.48400712 | 27.52522113 | 27.31639548 | 27.68772455 | 27.79346192 | 27.13897341 |
| Protein phosphatase 1F | P49593;P49593-2 | PPM1F | Other | 0.039868918 | 0.08679523 | 26.20849322 | 26.51586968 | 26.54352289 | 26.4034876 | 26.65867217 | 25.99509308 | 26.65307794 | 26.42649614 | 26.20260199 |
| Protein phosphatase 1G | O15355 | PPM1G | Other | 0.156544151 | 0.360484648 | 30.53834389 | 30.60831904 | 30.59814765 | 30.80747147 | 30.95345078 | 30.5829791 | 31.11352831 | 30.83680513 | 30.56365498 |
| Protein phosphatase methylesterase | Q9Y570;Q9Y570-4;Q9Y570-2 | PPME1 | Other | -0.259776694 | 1.565709648 | 28.0981836 | 28.23731114 | 28.15894384 | 27.98912715 | 27.90793221 | 28.3312951 | 27.88681162 | 27.95221522 | 27.7430396 |
| Protoporphyrinogen oxidase | P50336 | PPOX | Extracellular or secreted | 0.130722801 | 0.082776394 | 21.11171643 | 20.71246655 | 22.28282939 | 21.68478799 | 21.20386267 | 24.11592822 | 21.60127569 | 21.82849912 | 22.51818922 |
| Serine/threonine-protein phosphatas | P62136;P62136-2;P62136-3 | PPP1CA | Extracellular or secreted | 0.062527999 | 0.087846159 | 31.40988317 | 31.30677095 | 31.3843246 | 30.9509508 | 30.93044102 | 30.82843072 | 31.48370929 | 31.35147609 | 30.75779925 |
| Serine/threonine-protein phosphatas | P62140 | PPP1CB | Extracellular or secreted | -0.077199719 | 0.337275781 | 28.96162871 | 28.88028621 | 29.00645929 | 28.63519534 | 28.61445784 | 28.71374776 | 28.80882202 | 28.77665084 | 28.58881555 |
| Serine/threonine-protein phosphatas | P36873;P36873-2 | PPP1CC | Other | 0.127586208 | 0.064544327 | 26.85269542 | 26.92320379 | 27.05734364 | 26.7278029 | 26.90173199 | 25.95555573 | 27.45442469 | 27.54406852 | 25.59343216 |
| Serine/threonine-protein phosphatas | Q96QC0 | PPP1R10 | Other | -1.061181017 | 0.800940728 | 25.4092459 | 25.43641139 | 25.66901198 | 24.87792514 | 25.10313571 | 26.67384432 | 23.79699255 | 24.13625708 | 25.46799453 |
| Protein phosphatase 1 regulatory sub | O60927 | PPP1R11 | Other | 0.455644001 | 1.157579068 | 24.85214771 | 25.36138536 | 25.15271097 | 25.26549315 | 25.51965353 | 24.39734718 | 25.76777045 | 25.55663445 | 25.31689606 |
| Protein phosphatase 1 regulatory sub | O14974;O14974-3;O14974-2 | PPP1R12A | Other | -0.159038043 | 0.434752725 | 27.82535547 | 27.69252132 | 27.91871887 | 28.42758344 | 28.44632207 | 27.58355305 | 27.9370247 | 27.71574534 | 27.81714293 |
| Protein phosphatase 1 regulatory sub | Q9BZL4-5;Q9BZL4-3;Q9BZL4- | PPP1R12C | Other | 1.147458785 | 1.477020376 | 19.92000149 | 21.41622232 | 21.44303661 | 22.49084011 | 21.64024141 | 22.42054296 | 23.04942981 | 22.72336181 | 22.33502718 |
| RelA-associated inhibitor | Q8WUF5 | PPP1R13L | Extracellular or secreted | 0.138907822 | 0.110196666 | 22.96606603 | 23.3267685 | 23.25199548 | 22.86608269 | 21.22016213 | 24.01324906 | 22.83678467 | 22.77687796 | 23.62522279 |
| Protein phosphatase 1 regulatory sub | Q96C90 | PPP1R14B | Other | 0.277589655 | 0.666105535 | 29.53504656 | 29.44864737 | 29.44116545 | 29.63612934 | 29.23742512 | 28.73508623 | 29.67910703 | 29.83251083 | 29.33790115 |
| Protein phosphatase 1 regulatory sub | Q8TAE6 | PPP1R14C | Other | -0.620380789 | 1.02589305 | 25.43628429 | 25.00568862 | 24.61397332 | 24.57536705 | 24.85405188 | 23.92976407 | 23.8865093 | 24.56459284 | 23.89532011 |
| Protein phosphatase 1 regulatory sub | Q9UD71;Q9UD71-2 | PPP1R1B | Other | 1.246121791 | 3.145782333 | 28.63509153 | 28.97828334 | 28.84469734 | 28.26203734 | 27.98728483 | 27.83833894 | 29.68276112 | 29.81605965 | 29.51241126 |
| Protein phosphatase inhibitor 2;Prote | P41236;Q6NXS1 | PPP1R2;PPP1R2P | Other | 1.12265896 | 3.605326954 | 27.62156654 | 27.57195813 | 27.56848315 | 27.38459222 | 27.67263101 | 27.44300498 | 28.65095739 | 28.55504773 | 28.79308977 |
| Protein phosphatase 1 regulatory sub | Q6ZMI0;Q6ZMI0-3;Q6ZMI0-5 | PPP1R21 | Other | -0.188280833 | 0.219554079 | 25.22989696 | 24.96741239 | 24.8298253 | 23.35555005 | 23.68816492 | 24.29977817 | 24.36218838 | 24.37391496 | 23.88436805 |
| Protein phosphatase 1 regulatory sub | O95685 | PPP1R3D | Other | 0.440719056 | 1.11632164 | 22.25985933 | 22.11714972 | 22.15390946 | 22.24982756 | 22.28070735 | 22.05316416 | 22.56087856 | 22.42695674 | 22.89163065 |
| Protein phosphatase 1 regulatory sub | Q15435;Q15435-3;Q15435-2 | PPP1R7 | Extracellular or secreted | 0.073052174 | 0.108622249 | 29.44507905 | 29.32089876 | 29.40961009 | 28.98266873 | 28.99563765 | 28.39370271 | 29.37058907 | 29.32868698 | 28.79367896 |
| Nuclear inhibitor of protein phosphat | Q12972;Q12972-2 | PPP1R8 | Other | 0.366479661 | 1.354564653 | 25.3033365 | 25.75133892 | 26.12455174 | 25.66660327 | 25.42687999 | 25.98674253 | 25.94755222 | 26.21077983 | 26.0708334 |
| Neurabin-1 | Q9ULJ8;Q9ULJ8-4;Q9ULJ8-5; | PPP1R9A | Other | -0.984328185 | 0.415250627 | 23.3273169 | 23.07613709 | 23.98546188 | 22.40603057 | 22.72652502 | 22.47055363 | 23.16097347 | 20.26479244 | 22.61726208 |
| Neurabin-2 | Q96SB3 | PPP1R9B | Extracellular or secreted | -0.423214039 | 1.105730498 | 24.83475149 | 24.40412133 | 24.71560135 | 24.82992205 | 24.56028385 | 24.55683906 | 24.16813772 | 24.01137332 | 24.5016064 |
| Serine/threonine-protein phosphatas | P67775;P67775-2 | PPP2CA | Extracellular or secreted | -0.074628957 | 0.171244387 | 30.30083874 | 30.28107097 | 30.16578725 | 30.12730986 | 30.23409326 | 29.73259785 | 30.17878405 | 30.2291727 | 29.78900534 |
| Serine/threonine-protein phosphatas | P62714 | PPP2CB | Extracellular or secreted | 0.720365389 | 0.357127752 | 25.16096422 | 24.73474393 | 24.62220915 | 25.04506072 | 25.37188962 | 20.49782927 | 25.20880947 | 25.57735668 | 24.09127848 |
| Serine/threonine-protein phosphatas | P30153 | PPP2R1A | Extracellular or secreted | 0.284823309 | 0.587309145 | 31.67037862 | 31.87265287 | 31.89868673 | 31.63352475 | 31.58794874 | 31.56483629 | 32.15662534 | 32.19125299 | 31.62060559 |
| Serine/threonine-protein phosphatas | P30154;P30154-2;P30154-5; | PPP2R1B | Extracellular or secreted | 0.612065915 | 0.162100942 | 21.96093582 | 22.04746075 | 20.04794323 | 21.13044763 | 23.32127304 | 22.44514216 | 23.62332604 | 23.81098028 | 19.87849274 |
| Serine/threonine-protein phosphatas | P63151;P63151-2 | PPP2R2A | Other | 0.157011309 | 0.587440062 | 28.99285875 | 29.13726409 | 29.01294548 | 28.97921056 | 28.98563016 | 29.03666517 | 29.23194401 | 29.325774 | 28.98560302 |
| Serine/threonine-protein phosphatas | Q15257;Q15257-2;Q15257-3 | PPP2R4 | Extracellular or secreted | -0.029710937 | 0.047763626 | 30.45424817 | 30.39447977 | 30.38226434 | 29.89230904 | 29.81245303 | 29.92973586 | 30.33127374 | 30.2456986 | 29.76663996 |
| Serine/threonine-protein phosphatas | Q13362-4;Q13362-3;Q13362 | PPP2R5C | Other | 0.336558692 | 0.621911279 | 24.7745474 | 24.82948661 | 24.67551432 | 23.96578428 | 25.09034991 | 24.71319057 | 25.27774541 | 25.12502494 | 24.63134227 |
| Serine/threonine-protein phosphatas | Q14738;Q14738-2;Q14738-3 | PPP2R5D | Other | -0.0151902 | 0.011358512 | 28.46645503 | 28.263023 | 28.3320638 | 27.19676164 | 26.83737674 | 27.23927031 | 28.11768281 | 27.97125533 | 27.03296652 |
| Serine/threonine-protein phosphatas | Q08209-2;Q08209;Q08209-3 | PPP3CA;PPP3CB | Extracellular or secreted | 0.43352203 | 2.806498621 | 28.14666469 | 28.23849607 | 28.16336575 | 28.00308456 | 28.0720405 | 28.08588038 | 28.56015992 | 28.6293007 | 28.46587145 |
| Serine/threonine-protein phosphatas | P48454-2;P48454;P48454-3 | PPP3CC | Extracellular or secreted | 0.142885838 | 0.138738053 | 22.71580032 | 22.47753694 | 22.96215184 | 21.45048276 | 22.47338636 | 22.50076237 | 22.72173588 | 23.03049337 | 21.96648855 |
| Calcineurin subunit B type 1 | P63098 | PPP3R1 | Extracellular or secreted | 0.60565532 | 0.807731522 | 24.28644383 | 24.11624559 | 24.11894044 | 24.35002841 | 24.5835387 | 23.65970677 | 25.35302753 | 24.61778893 | 24.40360137 |
| Serine/threonine-protein phosphatas | P60510 | PPP4C | Extracellular or secreted | 0.061732943 | 0.117845396 | 29.45173542 | 29.49004668 | 29.36319153 | 29.33696535 | 29.33234129 | 28.75385259 | 29.52719903 | 29.48341139 | 29.03865484 |
| Serine/threonine-protein phosphatas | Q8TF05-2;Q8TF05 | PPP4R1 | Other | 0.851541037 | 2.365259955 | 27.06578951 | 27.21456429 | 27.10039109 | 27.76894421 | 27.89469585 | 26.92206968 | 28.41771978 | 28.08370127 | 28.03642937 |
| Serine/threonine-protein phosphatas | Q9NY27;Q9NY27-2;Q9NY27- | PPP4R2 | Other | 0.108758472 | 1.190084384 | 28.08861228 | 28.08283886 | 28.11793046 | 28.17126256 | 28.18115612 | 27.88605572 | 28.16959109 | 28.24263567 | 28.17797666 |
| Serine/threonine-protein phosphatas | P53041 | PPP5C | Other | 0.211443271 | 0.659149256 | 28.81555588 | 28.74393844 | 28.79810569 | 29.34720472 | 29.42956008 | 28.7604984 | 29.25444312 | 29.30765054 | 29.01966775 |
| Serine/threonine-protein phosphatas | O00743;O00743-3;O00743-2 | PPP6C | Other | -0.218682795 | 0.568626792 | 28.25033731 | 28.34006815 | 28.1559562 | 28.23772142 | 28.09315398 | 27.58856539 | 28.03836703 | 28.01775788 | 27.62072793 |
| Serine/threonine-protein phosphatas | Q9UPN7 | PPP6R1 | Other | -0.442833602 | 0.112948824 | 24.81685007 | 24.84388311 | 24.49692203 | 24.43822128 | 20.74335569 | 24.79069332 | 24.93098236 | 24.68859182 | 21.11688777 |
| Serine/threonine-protein phosphatas | O75170-6;O75170-2;O75170 | PPP6R2 | Other | 1.044260807 | 1.402606708 | 22.36988804 | 23.51544864 | 23.66307768 | 21.70180309 | 21.55147405 | 22.9005256 | 23.92614829 | 23.75796452 | 23.29977817 |
| Serine/threonine-protein phosphatas | Q5H9R7-6;Q5H9R7-2;Q5H9R | PPP6R3 | Extracellular or secreted | 0.056872025 | 0.083715895 | 28.90408892 | 28.83927045 | 28.73072065 | 28.62428867 | 28.81552534 | 28.18574686 | 28.9560711 | 28.94477956 | 28.31958585 |
| Palmitoyl-protein thioesterase 1 | P50897 | PPT1 | Extracellular or secreted | -0.134027254 | 0.281494723 | 27.79457779 | 27.91330708 | 27.82935353 | 28.61407164 | 28.68341386 | 27.83502869 | 28.14894471 | 27.79358595 | 27.99026387 |
| Lysosomal thioesterase PPT2 | Q9UMR5-2;Q9UMR5;Q9UMF | PPT2 | Extracellular or secreted | -0.207711458 | 0.158303419 | 21.69782758 | 22.11626146 | 22.56799598 | 23.19562452 | 22.66871449 | 21.40919409 | 22.50767949 | 22.44601197 | 21.25098323 |
| Peptidylprolyl isomerase domain and | Q96BP3;Q96BP3-2 | PPWD1 | Other | 0.176677421 | 0.340459374 | 26.74046841 | 27.09365773 | 27.03433167 | 26.58132772 | 25.71476327 | 26.26817331 | 26.88227026 | 26.67047148 | 26.69365157 |
| Polyglutamine-binding protein 1 | O60828-2;O60828;O60828-5 | PQBP1 | Other | 1.555816628 | 3.01898736 | 25.13985171 | 25.98122112 | 24.69178952 | 26.22764047 | 25.96868718 | 25.58204594 | 27.26552892 | 27.056517 | 27.14102193 |
| Melanoma antigen preferentially expr | P78395 | PRAME | Extracellular or secreted | 0.232083146 | 0.063363812 | 21.98613924 | 21.77048001 | 21.84942037 | 26.65717341 | 26.7931518 | 21.51349822 | 22.75295225 | 23.00288505 | 25.22534367 |
| Protein regulator of cytokinesis 1 | O43663-2;O43663-4;O43663 | PRC1 | Extracellular or secreted | -0.025578713 | 0.013346163 | 22.22374201 | 21.96419842 | 24.19430238 | 24.0123115 | 23.99714088 | 21.93955339 | 22.88048707 | 22.32310166 | 23.88529942 |
| Proline-rich protein PRCC | Q92733 | PRCC | Other | -0.131951528 | 0.453879339 | 25.31679251 | 25.68142459 | 25.64971097 | 25.39011463 | 25.58960863 | 25.12825429 | 25.13946142 | 25.43332616 | 25.40931065 |
| Lysosomal Pro-X carboxypeptidase | P42785;P42785-2 | PRCP | Extracellular or secreted | -0.604010132 | 0.87058495 | 25.82716203 | 25.95584392 | 25.82361953 | 26.68939191 | 27.041347 | 24.73018941 | 25.15957912 | 25.56857013 | 25.49359725 |
| Peroxiredoxin-1 | Q06830 | PRDX1 | Extracellular or secreted | 0.012652747 | 0.024844374 | 34.60039817 | 34.62804715 | 34.56354134 | 34.29645557 | 34.35073873 | 34.07447336 | 34.47954754 | 34.6539492 | 34.16128866 |
| Peroxiredoxin-2 | P32119 | PRDX2 | Extracellular or secreted | 0.066811502 | 0.203239918 | 33.02227647 | 33.03896548 | 32.91393831 | 32.67724289 | 32.48606149 | 32.51667195 | 32.92298412 | 32.92717675 | 32.67785194 |
| Thioredoxin-dependent peroxide redu | P30048-2;P30048 | PRDX3 | Extracellular or secreted | -0.083723268 | 0.166706348 | 31.12145489 | 31.136812 | 31.09458921 | 30.92046624 | 30.89929918 | 30.84116169 | 31.19190009 | 30.9635454 | 30.60027631 |
| Peroxiredoxin-4 | Q13162 | PRDX4 | Extracellular or secreted | -0.090164134 | 0.250824169 | 30.72720241 | 30.90784627 | 30.76461677 | 30.93086396 | 30.89821821 | 31.22561488 | 31.01464894 | 30.84415856 | 30.59788135 |
| Peroxiredoxin-5, mitochondrial | P30044-2;P30044;P30044-3; | PRDX5 | Extracellular or secreted | 0.858356817 | 2.841147958 | 29.89687669 | 29.98644417 | 30.03577416 | 29.50720235 | 29.41713628 | 30.08080761 | 30.49495554 | 30.68118663 | 30.86104891 |
| Peroxiredoxin-6 | P30041 | PRDX6 | Extracellular or secreted | 0.173064086 | 0.648522362 | 32.57580792 | 32.73622396 | 32.6898959 | 32.35660085 | 32.37479705 | 32.20317749 | 32.75037041 | 32.75918915 | 32.47788429 |
| Prolargin | P51888 | PRELP | Extracellular or secreted | -0.214051999 | 0.055767111 | 23.56906289 | 23.35218571 | 23.99963742 | 24.52282909 | 24.54968974 | 24.43980705 | 25.18082427 | 25.00774283 | 21.38588286 |
| Prolyl endopeptidase | P48147 | PREP | Other | -0.107155208 | 0.245363585 | 30.22676353 | 30.36087069 | 30.29712552 | 30.17152504 | 30.16758276 | 29.54447392 | 30.18269588 | 30.11359041 | 29.76641881 |
| Prolyl endopeptidase-like | Q4J6C6-4;Q4J6C6;Q4J6C6-2; | PREPL | Other | -0.268705565 | 0.411212936 | 27.08669037 | 26.99096711 | 26.88285328 | 27.93809101 | 28.19356942 | 26.83134842 | 27.33056872 | 27.06568681 | 26.75938759 |
| DNA primase small subunit | P49642 | PRIM1 | Other | 0.445672014 | 1.158352988 | 24.32127304 | 24.41538435 | 24.60127569 | 25.2178953 | 24.95650874 | 25.3421132 | 25.03170524 | 25.32597981 | 25.40655614 |
| DNA primase large subunit | P49643 | PRIM2 | Other | 0.440155643 | 0.428310295 | 24.54008215 | 24.24901736 | 23.99963742 | 23.86031788 | 21.83734614 | 24.79585378 | 24.57149528 | 24.34224887 | 24.04785014 |
| 5-AMP-activated protein kinase cataly | Q13131;Q13131-2 | PRKAA1 | Extracellular or secreted | -0.199830203 | 1.436438417 | 25.43615718 | 25.3667971 | 25.48667915 | 25.11565046 | 25.26180426 | 25.6030601 | 25.12376273 | 25.19977837 | 25.21204242 |
| 5-AMP-activated protein kinase subur | Q9Y478 | PRKAB1 | Other | 1.508365975 | 1.187948667 | 23.34353708 | 23.40820948 | 19.93206786 | 21.78394211 | 20.7223734 | 23.60093555 | 23.96146312 | 23.39754303 | 23.56162451 |
| cAMP-dependent protein kinase catal | P17612-2;P17612 | PRKACA | Extracellular or secreted | 0.069220788 | 0.125076555 | 27.84302048 | 28.29517772 | 28.10564536 | 27.31259232 | 27.12553739 | 27.87008962 | 27.66458389 | 27.99636522 | 27.82274469 |
| cAMP-dependent protein kinase catal | P22694-4;P22694-3;P22694; | PRKACB | Extracellular or secreted | 0.337976884 | 0.466236041 | 22.32319228 | 23.32223625 | 23.2268067 | 21.61945311 | 22.16927935 | 23.35097262 | 22.96130414 | 23.27042081 | 22.78817586 |
| cAMP-dependent protein kinase catal | P22612 | PRKACG | Other | 0.723892814 | 0.440223321 | 25.40031478 | 25.89400191 | 26.263023 | 22.00787639 | 22.74592383 | 24.34969104 | 25.39260593 | 25.26977902 | 24.83970896 |
| 5-AMP-activated protein kinase subur | P54619-2;P54619;P54619-3 | PRKAG1 | Extracellular or secreted | -0.14738152 | 0.32067403 | 24.1447603 | 24.74026252 | 24.70640833 | 23.81274372 | 24.2907383 | 24.8046435 | 24.29789078 | 24.06537866 | 24.44436433 |
| cAMP-dependent protein kinase type | P10644;P10644-2 | PRKAR1A | Extracellular or secreted | 0.318086722 | 1.264361931 | 28.845206 | 28.80992608 | 28.71623613 | 29.14804769 | 28.95900465 | 28.54730854 | 29.13606635 | 29.33095332 | 29.00010503 |
| cAMP-dependent protein kinase type | P13861-2;P13861 | PRKAR2A | Extracellular or secreted | -0.204045399 | 1.015652633 | 28.16274174 | 28.12760506 | 28.05915026 | 27.68157204 | 27.73292126 | 27.81347786 | 27.7693857 | 27.60392319 | 27.80328903 |
| cAMP-dependent protein kinase type | P31323 | PRKAR2B | Extracellular or secreted | -1.37748736 | 3.123738315 | 26.78755827 | 26.62296315 | 26.50133306 | 26.05131898 | 26.00496039 | 25.91335274 | 25.22273006 | 24.86613225 | 24.7194189 |
| Protein kinase C delta-binding protein | Q969G5 | PRKCDBP | Extracellular or secreted | -2.583824808 | 2.641650334 | 24.50506428 | 24.72645226 | 24.6671449 | 26.1702217 | 24.86716975 | 22.79495203 | 22.61578916 | 22.00601735 | 21.49222153 |
| Protein kinase C iota type | P41743 | PRKCI | Extracellular or secreted | 0.666047299 | 1.43644239 | 25.93660919 | 25.65537284 | 25.92565041 | 26.39532179 | 26.32457283 | 26.080523 | 26.62268393 | 27.05982072 | 26.47466227 |
| Glucosidase 2 subunit beta | P14314-2;P14314 | PRKCSH | Extracellular or secreted | -0.172459173 | 0.867754389 | 32.94006722 | 32.85515604 | 32.85446883 | 33.12235008 | 33.29447126 | 33.3381722 | 32.86870327 | 32.82232684 | 32.9939352 |
| DNA-dependent protein kinase cataly | P78527;P78527-2 | PRKDC | Other | 0.049464347 | 0.066423809 | 30.93522702 | 31.1319768 | 31.18163007 | 31.05355529 | 31.12774034 | 32.16701442 | 31.55896566 | 31.44702202 | 30.94097733 |
| Interferon-inducible double-stranded | O75569-3;O75569-2;O75569 | PRKRA | Other | 0.475241729 | 0.46366305 | 22.82462007 | 22.817748 | 20.744035 | 21.33914067 | 23.26360663 | 22.95888189 | 22.42744278 | 22.82749157 | 23.14480696 |
| 52 kDa repressor of the inhibitor of th | O43422 | PRKRIR | Other | 0.656102561 | 0.71501312 | 20.97858365 | 23.42854215 | 22.99020591 | 23.17057581 | 22.26978043 | 22.15005656 | 23.50893697 | 23.32402339 | 22.62921958 |
| Protein arginine N-methyltransferase | Q99873-3;Q99873-2;Q99873 | PRMT1 | Other | 0.428370871 | 1.230857872 | 31.82831722 | 32.03213236 | 31.9836271 | 31.81329436 | 31.94775007 | 31.99320978 | 32.5171884 | 32.44119514 | 32.12589452 |
| Protein arginine N-methyltransferase | O60678;O60678-2 | PRMT3 | Other | 0.385504066 | 0.574340717 | 26.17803366 | 26.03743518 | 26.33686323 | 26.64567668 | 27.06393977 | 26.20806523 | 27.07826452 | 27.01393054 | 26.29932402 |
| Protein arginine N-methyltransferase | O14744;O14744-5;O14744-2 | PRMT5 | Other | 0.052214809 | 0.086185862 | 29.74915985 | 29.8132026 | 29.77975331 | 29.82414174 | 29.90540941 | 29.3826354 | 30.03052179 | 29.94293561 | 29.41033819 |
| Protein arginine N-methyltransferase | Q9NVM4-4;Q9NVM4-3;Q9NV | PRMT7 | Other | 0.58543406 | 0.441274704 | 22.17692315 | 22.99717535 | 23.19191044 | 24.40509575 | 22.38828344 | 23.27554483 | 24.06233474 | 24.18903185 | 22.72240207 |
| Major prion protein | P04156-2;P04156 | PRNP | Extracellular or secreted | 1.626071288 | 1.113839975 | 24.12115595 | 24.03027445 | 23.83103426 | 25.09948843 | 25.37759316 | 25.46957897 | 25.66451608 | 25.78406687 | 27.39419353 |
| Endothelial protein C receptor | Q9UNN8 | PROCR | Extracellular or secreted | -0.195267746 | 0.050860765 | 24.89082868 | 24.68142459 | 24.56214878 | 25.11048235 | 25.44591179 | 27.00227989 | 24.14817864 | 23.62310273 | 27.48945343 |
| Proline synthase co-transcribed bacte | O94903 | PROSC | Extracellular or secreted | 0.483334172 | 0.627065935 | 26.48538983 | 26.22694263 | 26.24519536 | 26.20359176 | 26.32957683 | 27.21057548 | 26.66641365 | 26.61596655 | 27.51825826 |
| Pre-mRNA-processing factor 19 | Q9UMS4 | PRPF19 | Other | -0.640232507 | 1.549071455 | 31.77301498 | 31.93199117 | 31.8170743 | 30.95199298 | 31.10262024 | 31.41427571 | 31.00463899 | 31.01378415 | 30.55636403 |
| U4/U6 small nuclear ribonucleoprotei | O43395 | PRPF3 | Other | -0.902349189 | 0.309886051 | 22.71933534 | 23.96507967 | 24.60235227 | 23.64163108 | 24.44094772 | 23.52874215 | 20.8643359 | 24.45167647 | 23.42598418 |
| U4/U6 small nuclear ribonucleoprotei | Q8WWY3-2;Q8WWY3-4;Q8W | PRPF31 | Other | -1.891609714 | 0.910101661 | 25.43647493 | 25.61700419 | 25.80173598 | 25.35346508 | 25.40859822 | 26.41808988 | 22.42137915 | 23.89402505 | 25.0274508 |
| Pre-mRNA-splicing factor 38A | Q8NAV1 | PRPF38A | Other | -0.376351298 | 0.641917685 | 22.76038291 | 22.51053143 | 22.36292409 | 22.4539508 | 22.40639395 | 24.07054682 | 22.40870187 | 22.23908453 | 22.50552471 |
| Pre-mRNA-splicing factor 38B | Q5VTL8;Q5VTL8-2 | PRPF38B | Other | -0.427604079 | 0.26013076 | 22.19932548 | 21.36416746 | 23.66535674 | 23.01592202 | 23.42764738 | 24.61155528 | 21.65579912 | 23.14067877 | 23.06269706 |
| Pre-mRNA-processing factor 39 | Q86UA1 | PRPF39 | Other | 0.014977396 | 0.003585001 | 23.73748007 | 24.52808634 | 24.44088437 | 23.51508765 | 23.00908504 | 23.10616674 | 25.35669214 | 24.62995376 | 21.2266814 |
| U4/U6 small nuclear ribonucleoprotei | O43172-2;O43172 | PRPF4 | Other | 0.685944068 | 1.610403582 | 27.46154564 | 27.32508774 | 27.43682437 | 27.60293272 | 27.60540763 | 27.48784469 | 28.35077034 | 28.24861933 | 27.91826392 |
| Pre-mRNA-processing factor 40 homo | O75400-2;O75400-3;O75400 | PRPF40A | Other | -1.081151745 | 1.165618839 | 26.23297331 | 26.11527342 | 26.24083572 | 25.96558615 | 26.12528118 | 27.459671 | 25.20583023 | 24.69381107 | 25.92671386 |
| Serine/threonine-protein kinase PRP4 | Q13523 | PRPF4B | Other | -0.888956544 | 1.749668598 | 26.26745909 | 26.39752671 | 26.5247281 | 25.56546472 | 25.71387228 | 26.66234437 | 25.0517546 | 25.16852033 | 25.67855308 |
| Pre-mRNA-processing factor 6 | O94906;O94906-2 | PRPF6 | Other | -0.117801288 | 0.125971856 | 24.12328912 | 23.63599792 | 23.68142459 | 24.25673908 | 23.89170955 | 25.71947113 | 23.83961286 | 24.08440097 | 24.376898 |
| Pre-mRNA-processing-splicing factor 8 | Q6P2Q9 | PRPF8 | Other | -0.475138519 | 1.218491132 | 30.61315839 | 30.73940646 | 30.67673866 | 29.87857716 | 29.76724005 | 30.90834747 | 29.99954065 | 29.95864519 | 29.90813269 |
| Ribose-phosphate pyrophosphokinase | P60891;P60891-2 | PRPS1 | Other | 0.222761668 | 0.699281537 | 26.69829623 | 26.58276381 | 26.40556648 | 26.73815041 | 26.61694812 | 27.17150118 | 26.70332212 | 27.04176475 | 27.02981125 |
| Ribose-phosphate pyrophosphokinase | P11908;P11908-2 | PRPS2 | Extracellular or secreted | -0.101862007 | 0.517803668 | 28.96005484 | 29.11830186 | 29.21368399 | 29.40782873 | 29.32023169 | 29.3550627 | 29.18276691 | 29.19140597 | 29.00782301 |
| Phosphoribosyl pyrophosphate synth | Q14558;Q14558-2 | PRPSAP1 | Other | 0.325303394 | 0.856330698 | 26.10511579 | 26.04903506 | 26.10653419 | 26.20685504 | 26.22943861 | 27.06896963 | 26.41675385 | 26.62950915 | 26.81262133 |
| Phosphoribosyl pyrophosphate synth | O60256;O60256-4;O60256-3 | PRPSAP2 | Other | 0.355036701 | 1.326869534 | 27.9276183 | 27.91723976 | 27.89775675 | 27.76502796 | 27.68899192 | 28.32113538 | 28.11063169 | 28.46275506 | 28.25060838 |
| Proline-rich protein 36 | Q9H6K5-2;Q9H6K5 | PRR36 | Other | -0.380274761 | 0.535224973 | 25.34989347 | 24.99533035 | 24.98228792 | 25.62945357 | 24.7990203 | 23.53919479 | 24.77851313 | 24.48191519 | 24.24633759 |
| Protein PRRC1 | Q96M27;Q96M27-3;Q96M27 | PRRC1 | Other | -0.926025451 | 0.81219445 | 25.69399712 | 25.8364139 | 26.07532931 | 25.85666602 | 25.87939834 | 25.87537271 | 24.84555896 | 24.27696494 | 25.70798844 |
| Protein PRRC2A | P48634;P48634-3;P48634-2; | PRRC2A | Extracellular or secreted | -0.763846812 | 4.230236393 | 27.16873072 | 27.13721522 | 27.04447717 | 26.88518303 | 27.01797022 | 26.91079367 | 26.16714247 | 26.33246508 | 26.29103703 |
| Protein PRRC2C | Q9Y520-3;Q9Y520-4;Q9Y520 | PRRC2C | Other | -0.516540508 | 1.633613309 | 29.11485657 | 29.23582861 | 29.10507082 | 29.09224678 | 29.12659621 | 28.88075329 | 28.50827965 | 28.43730074 | 28.78247423 |
| Brain-specific serine protease 4 | Q9GZN4 | PRSS22 | Extracellular or secreted | 0.184328852 | 0.104105576 | 25.76761892 | 26.01968896 | 25.57888308 | 25.59804114 | 26.78755827 | 26.36289066 | 25.55987557 | 26.05564851 | 27.49480298 |
| Serine protease 23 | O95084;O95084-2 | PRSS23 | Extracellular or secreted | 1.859978787 | 1.50617009 | 21.31881183 | 21.6264485 | 21.69077816 | 20.20634496 | 22.82472743 | 23.00646785 | 23.48123774 | 22.91128299 | 24.524205 |
| Prostasin;Prostasin light chain;Prostas | Q16651;Q16651-2 | PRSS8 | Extracellular or secreted | 0.058514298 | 0.039637635 | 26.36213308 | 26.33097895 | 26.28162375 | 26.2747454 | 25.9437573 | 26.5130554 | 25.84359562 | 25.89082868 | 27.27426553 |
| Phosphoribosyltransferase domain-co | Q9NRG1;Q9NRG1-2 | PRTFDC1 | Other | -0.208251862 | 0.516189967 | 27.72231754 | 27.77410749 | 27.72123262 | 27.08658915 | 27.21632328 | 27.25961513 | 27.43991799 | 27.33073966 | 26.99517936 |
| Protein prune homolog | Q86TP1;Q86TP1-3;Q86TP1-2 | PRUNE | Other | -0.036114554 | 0.025335157 | 25.71381986 | 25.84957309 | 25.96386781 | 25.52939767 | 25.37321803 | 24.67546048 | 25.94033331 | 25.87220518 | 24.63178632 |
| Prosaposin;Saposin-A;Saposin-B-Val;S | P07602;P07602-2;P07602-3 | PSAP | Extracellular or secreted | 0.474784249 | 1.507271581 | 27.97710983 | 28.03951792 | 28.1639895 | 28.55482823 | 28.82532514 | 27.76875495 | 28.63013434 | 28.69335248 | 28.76562872 |
| Phosphoserine aminotransferase | Q9Y617;Q9Y617-2 | PSAT1 | Extracellular or secreted | -0.062594126 | 0.082924638 | 32.23842774 | 32.19650958 | 32.13213006 | 32.5192289 | 32.61467284 | 30.87122124 | 32.07249418 | 32.16973443 | 31.85608419 |
| PC4 and SFRS1-interacting protein | O75475;O75475-3 | PSIP1 | Other | -0.699229231 | 0.744801982 | 29.60272038 | 29.51487703 | 29.51636199 | 29.60309195 | 29.65458097 | 28.85129597 | 29.1330064 | 29.09717908 | 28.04359098 |
| Proteasome subunit alpha type-1 | P25786;P25786-2 | PSMA1 | Other | -0.144076336 | 0.35720391 | 32.19372216 | 32.13461062 | 32.08359984 | 32.39424976 | 32.55270468 | 31.40709936 | 31.95823033 | 31.88015482 | 32.11237906 |
| Proteasome subunit alpha type-2 | P25787 | PSMA2 | Other | -0.161143412 | 0.362396621 | 31.4112478 | 31.57925184 | 31.53973097 | 31.83815857 | 31.85775336 | 30.70281042 | 31.47745009 | 31.33260806 | 31.17098809 |
| Proteasome subunit alpha type-3 | P25788-2;P25788 | PSMA3 | Extracellular or secreted | -0.242039917 | 0.694800054 | 31.17676472 | 31.27269817 | 31.19148833 | 31.47788429 | 31.51615532 | 30.55169673 | 30.90791788 | 30.81688363 | 31.14242252 |
| Proteasome subunit alpha type-4 | P25789;P25789-2 | PSMA4 | Extracellular or secreted | -0.062174293 | 0.104136208 | 31.46173102 | 31.73606264 | 31.7165714 | 31.86104891 | 31.96447525 | 31.28421897 | 31.48610946 | 31.36718675 | 31.97223501 |
| Proteasome subunit alpha type-5 | P28066;P28066-2 | PSMA5 | Extracellular or secreted | -0.063548582 | 0.124934388 | 31.80096963 | 31.83303889 | 31.67261415 | 32.00138752 | 32.09823381 | 31.13943702 | 31.49709911 | 31.64271928 | 31.94237638 |
| Proteasome subunit alpha type-6 | P60900;P60900-2;P60900-3 | PSMA6 | Extracellular or secreted | -0.069819536 | 0.132048073 | 32.15840219 | 32.08007693 | 32.0313766 | 32.42071407 | 32.57533459 | 31.45302179 | 32.01913086 | 31.85545312 | 32.2754205 |
| Proteasome subunit alpha type-7 | O14818 | PSMA7 | Extracellular or secreted | -0.073814494 | 0.258166397 | 32.22463782 | 32.23956623 | 32.20841996 | 32.47101822 | 32.50033225 | 31.75059004 | 32.22426406 | 32.08160141 | 32.16997331 |
| Proteasome subunit alpha type-7 | O14818-2 | PSMA7 | Extracellular or secreted | -2.471945097 | 1.74919048 | 24.47969689 | 24.58336654 | 24.28982416 | 20.34533954 | 21.78895965 | 24.01000762 | 21.02114913 | 20.55877322 | 20.75283957 |
| Proteasome subunit beta type-1 | P20618 | PSMB1 | Extracellular or secreted | -0.128511592 | 0.37354856 | 31.86097494 | 32.07899612 | 31.93526215 | 32.22354502 | 32.27641925 | 31.49257005 | 31.80328133 | 31.713461 | 32.03160666 |
| Proteasome subunit beta type-10 | P40306 | PSMB10 | Other | -3.202812777 | 1.061625708 | 27.5506147 | 27.12908016 | 27.39827725 | 26.39407902 | 26.65184703 | 24.65458097 | 24.49075444 | 24.59280511 | 21.19724169 |
| Proteasome subunit beta type-2 | P49721 | PSMB2 | Extracellular or secreted | -0.009590642 | 0.016787156 | 30.96787957 | 31.03688784 | 30.9564312 | 31.27630832 | 31.52229595 | 30.41205586 | 30.92980639 | 30.86901596 | 31.2583351 |
| Proteasome subunit beta type-3 | P49720 | PSMB3 | Extracellular or secreted | 0.096530667 | 0.22683057 | 30.79343091 | 31.10243252 | 31.01112002 | 31.26584185 | 31.36515401 | 30.22837009 | 31.06702135 | 31.09829657 | 31.00744878 |
| Proteasome subunit beta type-4 | P28070 | PSMB4 | Extracellular or secreted | -0.020348544 | 0.048051417 | 31.26108464 | 30.99745588 | 30.97309084 | 31.5129104 | 31.41048983 | 30.66932292 | 31.06580235 | 31.02015808 | 31.2651712 |
| Proteasome subunit beta type-5 | P28074;P28074-3 | PSMB5 | Extracellular or secreted | 0.010590223 | 0.029660344 | 31.85143747 | 31.84800789 | 31.8370684 | 32.13341681 | 32.32680814 | 31.69303669 | 32.02221032 | 31.80478196 | 32.04966609 |
| Proteasome subunit beta type-6 | P28072 | PSMB6 | Extracellular or secreted | -0.030559385 | 0.0444044 | 30.73747363 | 30.69494694 | 30.72476543 | 31.33228793 | 31.39222923 | 29.97541611 | 31.01305201 | 30.65129963 | 30.67252985 |
| Proteasome subunit beta type-7 | Q99436 | PSMB7 | Extracellular or secreted | -0.054160739 | 0.124849335 | 31.2301467 | 31.52009583 | 31.32377864 | 31.6240969 | 31.73755421 | 30.68678899 | 31.36196885 | 31.19513121 | 31.34164886 |
| Proteasome subunit beta type-8 | P28062-2;P28062 | PSMB8 | Extracellular or secreted | -1.476003152 | 2.336723396 | 29.26537243 | 29.34994829 | 29.1348921 | 28.89694884 | 29.1017941 | 27.27230885 | 27.45607111 | 27.49197784 | 27.13457391 |
| Proteasome subunit beta type-9 | P28065-2;P28065 | PSMB9 | Extracellular or secreted | -2.374233658 | 2.979885733 | 28.43069692 | 28.7652177 | 28.58981576 | 28.23187537 | 28.37467788 | 26.64677556 | 26.22086819 | 25.89050402 | 25.28545641 |
| 26S protease regulatory subunit 4 | P62191;P62191-2 | PSMC1 | Extracellular or secreted | 0.241577615 | 0.815376659 | 30.33287478 | 30.33968598 | 30.20467835 | 30.18647907 | 30.32620418 | 30.40740331 | 30.7169393 | 30.54480552 | 30.33170085 |
| 26S protease regulatory subunit 7 | P35998;P35998-2 | PSMC2 | Other | 0.268169341 | 0.965196303 | 31.07546193 | 31.02154863 | 30.99206186 | 30.8195507 | 30.90059527 | 31.00504072 | 31.34493205 | 31.33671003 | 31.0299955 |
| 26S protease regulatory subunit 6A | P17980 | PSMC3 | Other | 0.437111003 | 0.734029279 | 30.18810073 | 30.20920469 | 30.14746553 | 29.85765697 | 29.89266566 | 29.78891203 | 30.71464865 | 30.61754375 | 29.99118343 |
| 26S protease regulatory subunit 6B | P43686;P43686-2 | PSMC4 | Other | 0.281384403 | 0.973599227 | 30.41165189 | 30.36723883 | 30.45845899 | 30.18919213 | 30.28372237 | 30.3119214 | 30.82403549 | 30.57109017 | 30.46012035 |
| 26S protease regulatory subunit 8 | P62195;P62195-2 | PSMC5 | Extracellular or secreted | 0.272111968 | 0.935150156 | 30.75660684 | 30.71095963 | 30.68085142 | 30.38946241 | 30.5792968 | 30.50379508 | 31.05975627 | 30.89352464 | 30.67354108 |
| 26S protease regulatory subunit 10B | P62333 | PSMC6 | Extracellular or secreted | 0.291334898 | 0.758964003 | 30.03065333 | 30.02366498 | 29.95240965 | 29.29876488 | 30.39805295 | 30.17343258 | 30.56392768 | 30.60523099 | 30.14333521 |
| 26S proteasome non-ATPase regulato | Q99460;Q99460-2 | PSMD1 | Extracellular or secreted | 0.200322134 | 0.568079803 | 31.16590702 | 31.13019777 | 31.11929179 | 30.64306348 | 30.75803761 | 31.02458992 | 31.33356801 | 31.26896747 | 30.91897471 |

| | | | | | | | | | | | | | | |
|---|---|---|---|---|---|---|---|---|---|---|---|---|---|---|
| 26S proteasome non-ATPase regulato | O75832;O75832-2 | PSMD10 | Other | -0.037707729 | 0.136779329 | 27.71875287 | 27.90173199 | 27.67330519 | 27.4954131 | 27.60137488 | 27.61133014 | 27.47010673 | 27.65935291 | 27.75837125 |
| 26S proteasome non-ATPase regulato | O00231;O00231-2 | PSMD11 | Extracellular or secreted | 0.430417819 | 0.974723643 | 30.01531381 | 30.0567237 | 29.99859951 | 29.81613597 | 29.88716036 | 29.93789465 | 30.56165364 | 30.5188773 | 30.06663652 |
| 26S proteasome non-ATPase regulato | O00232;O00232-2 | PSMD12 | Extracellular or secreted | 0.170827502 | 1.192097575 | 30.16470887 | 30.16014679 | 30.09345625 | 29.86406386 | 29.87324713 | 30.06162425 | 30.27453214 | 30.24070382 | 30.10587011 |
| 26S proteasome non-ATPase regulato | Q9UNM6;Q9UNM6-2 | PSMD13 | Extracellular or secreted | 0.184970661 | 0.753119798 | 30.39560372 | 30.5042688 | 30.36484103 | 29.93480536 | 29.95988907 | 30.28769037 | 30.51314578 | 30.48778721 | 30.27752817 |
| 26S proteasome non-ATPase regulato | O00487 | PSMD14 | Extracellular or secreted | 0.211100769 | 1.040066621 | 29.53504656 | 29.57690276 | 29.57897303 | 29.00378158 | 29.13383935 | 29.38899049 | 29.59782808 | 29.60895348 | 29.53528763 |
| 26S proteasome non-ATPase regulato | Q13200;Q13200-3;Q13200-2 | PSMD2 | Extracellular or secreted | 0.216377154 | 0.938485023 | 31.85804989 | 32.0550757 | 32.01976053 | 31.73416571 | 31.8685377 | 31.91723265 | 32.25467964 | 32.16581719 | 31.95504572 |
| 26S proteasome non-ATPase regulato | O43242;O43242-2 | PSMD3 | Extracellular or secreted | 0.001977132 | 0.006441675 | 30.56538118 | 30.61307054 | 30.55435254 | 30.44124462 | 30.34212592 | 30.56001411 | 30.64289139 | 30.55837271 | 30.34276175 |
| 26S proteasome non-ATPase regulato | P55036;P55036-2 | PSMD4 | Other | 1.024031516 | 3.141742435 | 28.74733611 | 29.00613823 | 28.98666116 | 28.65536601 | 28.78197422 | 28.65867217 | 29.95004697 | 29.88820609 | 29.65191545 |
| 26S proteasome non-ATPase regulato | Q16401;Q16401-2 | PSMD5 | Other | 0.007096378 | 0.013030577 | 30.1016188 | 30.21326682 | 30.17569451 | 29.74111164 | 29.76317667 | 29.51111079 | 30.00088406 | 30.11929179 | 29.6541029 |
| 26S proteasome non-ATPase regulato | Q15008;Q15008-4;Q15008-3 | PSMD6 | Extracellular or secreted | 0.198471586 | 0.705255596 | 30.11916808 | 30.38730676 | 30.32442261 | 29.69488054 | 29.77621157 | 30.07724602 | 30.3876149 | 30.2808498 | 30.11656785 |
| 26S proteasome non-ATPase regulato | P51665 | PSMD7 | Extracellular or secreted | -0.091007873 | 0.243152368 | 29.45138171 | 29.61694812 | 29.62392253 | 29.12324472 | 29.18876781 | 29.08271199 | 29.47720881 | 29.1806346 | 29.11262141 |
| 26S proteasome non-ATPase regulato | P48556 | PSMD8 | Extracellular or secreted | 0.159752346 | 0.585361875 | 29.17009273 | 29.37328857 | 29.34216832 | 29.11572486 | 29.1281707 | 29.10079209 | 29.5575148 | 29.33462321 | 29.20223766 |
| 26S proteasome non-ATPase regulato | O00233;O00233-2 | PSMD9 | Other | 0.205273253 | 0.255780109 | 28.26369465 | 27.94960179 | 27.9256391 | 28.0204098 | 28.11252199 | 27.47707367 | 28.43264938 | 28.46244305 | 27.59519783 |
| Proteasome activator complex subuni | Q06323;Q06323-3;Q06323-2 | PSME1 | Extracellular or secreted | -0.476497332 | 0.797416237 | 30.82213686 | 30.85087885 | 30.78744929 | 30.15978601 | 30.2178953 | 29.9845441 | 30.10024069 | 30.3223824 | 29.55923012 |
| Proteasome activator complex subuni | Q9UL46 | PSME2 | Extracellular or secreted | -0.538976985 | 0.694376726 | 30.93677204 | 31.08916814 | 31.08663976 | 30.46878697 | 30.52309092 | 29.75252931 | 30.39988714 | 30.38514788 | 29.5265276 |
| Proteasome activator complex subuni | P61289;P61289-3;P61289-2 | PSME3 | Other | 0.020310342 | 0.0224154 | 29.87579746 | 30.02696559 | 29.95629271 | 29.78478454 | 29.74132062 | 29.72344774 | 30.14843566 | 30.16038726 | 29.30641244 |
| Proteasome activator complex subuni | Q14997;Q14997-2 | PSME4 | Other | 0.834035363 | 1.445357934 | 27.11812855 | 27.09928777 | 27.06804709 | 25.48600393 | 26.38477334 | 27.44426953 | 27.39892957 | 27.63796098 | 27.76547065 |
| Proteasome inhibitor PI31 subunit | Q92530 | PSMF1 | Other | 0.037002624 | 0.048152167 | 27.78188045 | 27.43991799 | 27.50809071 | 27.71456678 | 27.69398383 | 27.52043309 | 27.76925957 | 27.99539505 | 27.17578967 |
| Proteasome assembly chaperone 1 | O95456-2;O95456 | PSMG1 | Other | 0.604490386 | 1.425448106 | 26.43095208 | 26.86466904 | 26.67222634 | 26.52463843 | 26.58792192 | 26.71881818 | 27.31112022 | 27.44758372 | 26.95438022 |
| Proteasome assembly chaperone 2 | Q969U7-2;Q969U7 | PSMG2 | Other | 0.016047248 | 0.041308101 | 25.92309042 | 25.8508133 | 25.9425945 | 26.07412496 | 26.109845 | 25.76017477 | 26.19415206 | 25.83990114 | 25.84441001 |
| Proteasome assembly chaperone 3 | Q9BT73 | PSMG3 | Other | 1.756658701 | 1.375373413 | 24.48246923 | 24.55952553 | 24.35602043 | 21.57497863 | 22.07504076 | 21.69971162 | 25.20474874 | 25.2934071 | 24.14569337 |
| Proteasome assembly chaperone 4 | Q5JS54;Q5JS54-2;Q5JS54-3 | PSMG4 | Other | 0.448001753 | 1.079591612 | 24.57270994 | 24.4790184 | 24.34658353 | 24.4171242 | 24.57375026 | 23.81645949 | 25.13394711 | 24.68827166 | 24.62460941 |
| Paraspeckle component 1 | Q8WXF1;Q8WXF1-2 | PSPC1 | Other | 0.800219492 | 1.527299364 | 29.65215486 | 29.50296097 | 29.47807723 | 29.65092316 | 29.72266632 | 29.39637976 | 30.48615742 | 30.57181351 | 30.0442687 |
| Phosphoserine phosphatase | P78330 | PSPH | Other | -0.062669465 | 0.266683744 | 27.47877159 | 27.43324655 | 27.36536263 | 27.59135045 | 27.44419053 | 26.93055382 | 27.27453214 | 27.35093891 | 27.30825833 |
| Polypyrimidine tract-binding protein 1 | P26599;P26599-2;P26599-3 | PTBP1 | Extracellular or secreted | 0.144516216 | 0.197316884 | 31.6306638 | 31.82244081 | 31.79857697 | 31.73137633 | 31.71141102 | 31.55545006 | 32.08910499 | 32.13439642 | 31.33500673 |
| Polypyrimidine tract-binding protein 3 | O95758-1;O95758-6;O95758 | PTBP3 | Other | 1.18685701 | 0.780725298 | 22.04654948 | 24.09772161 | 20.55422631 | 23.44871826 | 23.77384348 | 25.48747673 | 24.96811586 | 24.3098721 | 23.986851 |
| Phosphotriesterase-related protein | Q96BW5-2;Q96BW5 | PTER | Extracellular or secreted | -1.27483557 | 0.914275927 | 25.03073751 | 26.02648022 | 25.23348539 | 26.10337586 | 26.85459887 | 23.207827 | 24.34068791 | 24.85290968 | 23.21014812 |
| Prostaglandin E synthase 2;Prostaglan | Q9H7Z7 | PTGES2 | Other | 0.356938509 | 0.968877014 | 25.50124193 | 25.81892324 | 25.38436166 | 25.60902043 | 25.64935465 | 26.30528637 | 25.88760198 | 25.99204835 | 26.32525933 |
| Prostaglandin E synthase 3 | Q15185-4;Q15185;Q15185-3 | PTGES3 | Extracellular or secreted | 1.159789053 | 2.088686799 | 29.56176288 | 29.58496901 | 29.58199208 | 29.67256357 | 29.74275057 | 29.54336801 | 30.84093667 | 30.94935132 | 30.53278223 |
| Prostaglandin F2 receptor negative re | Q9P2B2 | PTGFRN | Other | 0.293871979 | 0.346807361 | 28.41747836 | 28.36340043 | 28.3841393 | 28.85682628 | 28.99490972 | 28.72129782 | 28.5615444 | 28.652531 | 29.53656649 |
| Prostaglandin reductase 2 | Q8N8N7;Q8N8N7-2 | PTGR2 | Extracellular or secreted | -0.288185754 | 1.697627891 | 27.71920998 | 27.73013757 | 27.81433388 | 27.68304568 | 27.86555422 | 27.27248684 | 27.37094237 | 27.40713989 | 27.39974456 |
| Focal adhesion kinase 1 | Q05397-7;Q05397;Q05397-5 | PTK2 | Extracellular or secreted | -0.583286341 | 1.634169319 | 26.38949113 | 26.3827467 | 26.45729295 | 26.40579368 | 26.77806189 | 26.62198566 | 26.00545305 | 25.67718239 | 26.08519153 |
| Inactive tyrosine-protein kinase 7 | Q13308;Q13308-6;Q13308-4 | PTK7 | Extracellular or secreted | 0.48913217 | 1.099292107 | 30.37534096 | 30.34561951 | 30.33255471 | 29.9542138 | 29.88133693 | 30.54287007 | 30.54167063 | 30.5706379 | 31.07105597 |
| Prothymosin alpha;Prothymosin alpha | P06454-2;P06454 | PTMA | Extracellular or secreted | 0.076033264 | 0.092362258 | 33.70614975 | 33.41664313 | 33.49321452 | 33.36306617 | 32.96318363 | 32.82390267 | 33.31972043 | 33.83450889 | 32.95695041 |
| Parathymosin | P20962 | PTMS | Other | 0.320930524 | 0.241953857 | 27.03485638 | 27.98304943 | 28.00023938 | 28.7765881 | 28.15557024 | 25.46559904 | 28.20862345 | 28.07571694 | 27.38640248 |
| Protein tyrosine phosphatase type IVA | Q93096 | PTP4A1 | Extracellular or secreted | 1.146635929 | 0.944384295 | 22.28647203 | 22.50091383 | 21.55496725 | 22.69884199 | 20.51278284 | 23.12940252 | 22.7755926 | 24.17012617 | 22.83587924 |
| Tyrosine-protein phosphatase non-rec | P18031 | PTPN1 | Extracellular or secreted | 0.192454734 | 0.191452368 | 25.55832892 | 25.59291914 | 25.63242439 | 25.95889958 | 25.87208773 | 27.80704149 | 26.01501598 | 26.15611055 | 26.61708829 |
| Tyrosine-protein phosphatase non-rec | Q06124;Q06124-2;Q06124-3 | PTPN11 | Other | -0.120245593 | 0.361181492 | 28.45998361 | 28.48273848 | 28.50793954 | 28.35808493 | 28.46559904 | 28.02178687 | 28.42322128 | 28.32615988 | 28.0379483 |
| Tyrosine-protein phosphatase non-rec | Q05209-2;Q05209-3;Q05209 | PTPN12 | Extracellular or secreted | 0.445870093 | 0.396213996 | 24.33817326 | 24.06365182 | 24.7025033 | 24.95681889 | 24.80168665 | 21.93310246 | 24.20471143 | 24.75959078 | 24.77127627 |
| Tyrosine-protein phosphatase non-rec | P17706-3;P17706-2;P17706;P | PTPN2 | Extracellular or secreted | 0.530530561 | 1.096261659 | 24.11092037 | 24.35977797 | 23.8634407 | 23.35971096 | 24.31610196 | 24.77615508 | 24.70424604 | 24.3595099 | 24.92088926 |
| Tyrosine-protein phosphatase non-rec | Q9H3S7 | PTPN23 | Extracellular or secreted | -0.098871804 | 0.62022416 | 28.54399479 | 28.56048794 | 28.63311763 | 28.3164818 | 28.77514421 | 28.3235854 | 28.4432817 | 28.44509879 | 28.39140999 |
| Tyrosine-protein phosphatase non-rec | P29350-2;P29350;P29350-3;P | PTPN6 | Extracellular or secreted | -0.959677696 | 1.044621325 | 27.4604524 | 26.81102929 | 27.04166032 | 25.74599082 | 26.21863448 | 26.62992598 | 25.96690655 | 26.12062217 | 24.98728483 |
| Receptor-type tyrosine-protein phosp | P10586-2;P10586 | PTPRF | Extracellular or secreted | 0.001009217 | 0.001507205 | 30.90806109 | 30.96368319 | 30.86850091 | 30.70610857 | 30.61841922 | 30.64142782 | 30.53519491 | 30.63823725 | 31.18269588 |
| Receptor-type tyrosine-protein phosp | P23470-2;P23470 | PTPRG | Extracellular or secreted | 0.165566324 | 0.195963027 | 23.09276295 | 23.18001232 | 23.70967199 | 23.96305199 | 23.51340184 | 23.72795867 | 23.34989347 | 23.47975856 | 24.26047682 |
| Receptor-type tyrosine-protein phosp | Q12913;Q12913-2 | PTPRJ | Extracellular or secreted | 0.74420273 | 0.877044242 | 23.22507149 | 22.7919951 | 21.09945113 | 22.06084871 | 22.64561622 | 20.70116192 | 23.05536718 | 22.79713112 | 22.64218217 |
| Receptor-type tyrosine-protein phosp | Q15262;Q15262-2;Q15262-3 | PTPRK | Extracellular or secreted | 0.877759733 | 0.79764139 | 27.01403699 | 27.08415764 | 26.9375299 | 27.3095599 | 27.4194086 | 27.94937915 | 27.64958763 | 27.84846315 | 28.9922645 |
| Receptor-type tyrosine-protein phosp | Q13332-6;Q13332-2;Q13332 | PTPRS | Extracellular or secreted | -0.51869334 | 0.871516734 | 27.85620295 | 27.84840345 | 27.75786281 | 28.14053446 | 28.04452928 | 28.95543379 | 27.35749777 | 27.36009624 | 28.02780934 |
| Receptor-type tyrosine-protein phosp | P23471-3;P23471-2;P23471 | PTPRZ1 | Extracellular or secreted | 0.056869998 | 0.008128199 | 21.36027996 | 21.65041894 | 22.51121156 | 22.75712704 | 21.83691942 | 29.27464322 | 21.17840226 | 21.54113444 | 27.14637336 |
| Polymerase I and transcript release fa | Q6NZI2;Q6NZI2-2 | PTRF | Extracellular or secreted | -0.568360023 | 1.226664467 | 26.41932823 | 25.94966858 | 26.00889703 | 25.69914392 | 25.74044267 | 27.20545739 | 25.55765721 | 25.49371939 | 25.75501224 |
| Peptidyl-tRNA hydrolase 2, mitochond | Q9Y3E5 | PTRH2 | Other | 0.051498338 | 0.027214011 | 22.39252077 | 23.07967293 | 23.43580758 | 23.31755173 | 23.36051491 | 25.93287542 | 22.99043918 | 23.49326129 | 24.43026622 |
| Putative peptidyl-tRNA hydrolase PTR | Q6GMV3 | PTRHD1 | Extracellular or secreted | 0.831401801 | 1.279542609 | 27.92529953 | 27.80974213 | 27.7691965 | 27.63304832 | 27.54222434 | 27.55354715 | 28.42225903 | 28.95820266 | 28.2302727 |
| Transcriptional activator protein Pur-a | Q00577 | PURA | Other | 0.107078733 | 0.116417616 | 26.01703572 | 26.31925852 | 26.1940581 | 25.90253723 | 26.10569531 | 25.23260742 | 26.41862073 | 26.29330184 | 25.49490977 |
| Transcriptional activator protein Pur-b | Q96QR8 | PURB | Other | -0.074793071 | 0.229634805 | 27.11545203 | 26.8358118 | 26.72533443 | 26.52047805 | 26.3116746 | 26.63270172 | 26.49734851 | 26.73879467 | 26.61020392 |
| tRNA pseudouridine synthase A, mito | Q9Y606-2;Q9Y606 | PUS1 | Other | 0.248604955 | 0.318830635 | 25.75447713 | 26.43548969 | 26.46964107 | 25.0661181 | 24.85438485 | 25.93059893 | 26.1800275 | 26.19296775 | 25.62817451 |
| Pseudouridylate synthase 7 homolog | Q96PZ0;Q96PZ0-2 | PUS7 | Other | -0.012949565 | 0.025066743 | 28.93373662 | 29.18971057 | 29.16837208 | 28.99663459 | 29.07513036 | 28.99601495 | 29.29605346 | 29.09624927 | 28.74864816 |
| Poliovirus receptor | P15151-3;P15151-2;P15151;P | PVR | Extracellular or secreted | 1.055174311 | 1.258690492 | 26.41141349 | 26.73143298 | 26.78780733 | 26.77303857 | 26.88878672 | 27.72676406 | 27.70556489 | 27.54877888 | 28.57080073 |
| Nectin-1 | Q15223;Q15223-3;Q15223-2 | PVRL1 | Extracellular or secreted | -0.33813491 | 0.238529611 | 27.13261426 | 27.10669392 | 27.03254622 | 27.23635338 | 27.34654971 | 28.48393027 | 26.54442233 | 26.57629012 | 28.0342267 |
| Nectin-2 | Q92692-2;Q92692 | PVRL2 | Extracellular or secreted | 0.333553447 | 1.388243103 | 27.31319804 | 27.22655684 | 27.06804709 | 27.43928394 | 27.54532121 | 27.32920042 | 27.47606941 | 27.6978987 | 27.78749599 |
| Nectin-4;Processed poliovirus recepto | Q96NY8 | PVRL4 | Extracellular or secreted | 0.255446639 | 0.544243749 | 28.34053512 | 28.31168326 | 28.31967198 | 27.54671999 | 27.80882202 | 28.50801512 | 28.14705303 | 28.36030559 | 28.67670504 |
| Periodic tryptophan protein 1 homolo | Q13610 | PWP1 | Other | 0.417414499 | 1.005295698 | 27.89394407 | 27.42678403 | 27.47923432 | 28.15879942 | 28.16552372 | 27.28973623 | 28.30103505 | 28.29544049 | 27.86277884 |
| Periodic tryptophan protein 2 homolo | Q15269 | PWP2 | Other | -0.346418244 | 0.229511166 | 23.24043907 | 23.45888918 | 23.46699948 | 21.2101585 | 22.73638895 | 25.40940777 | 22.5213744 | 23.40236571 | 22.79814663 |
| Peroxidasin homolog | Q92626 | PXDN | Extracellular or secreted | 1.539997207 | 2.03656494 | 28.2606473 | 27.95249297 | 28.12030576 | 28.90970698 | 28.89460913 | 30.53083056 | 30.16830034 | 30.19201771 | 30.59396992 |
| Paxillin | P49023-2;P49023;P49023-3;P | PXN | Extracellular or secreted | 0.754957652 | 0.873635553 | 23.4759303 | 23.40431626 | 23.2992192 | 21.67012755 | 23.74786107 | 23.59479932 | 23.59946069 | 23.70756725 | 24.55397187 |
| Pyrroline-5-carboxylate reductase 1, r | P32322;P32322-3;P32322-2 | PYCR1 | Other | -0.425255556 | 0.444046362 | 26.11642408 | 26.19561701 | 26.07838669 | 26.81457836 | 26.58777889 | 27.84565467 | 25.94561136 | 25.78424164 | 26.81360018 |
| Pyrroline-5-carboxylate reductase 2 | Q96C36 | PYCR2 | Other | -0.271332339 | 0.310257068 | 25.98672083 | 25.90774888 | 25.98302768 | 26.01222623 | 26.01695073 | 27.89226561 | 25.68837838 | 26.0458523 | 26.35124229 |
| Pyrroline-5-carboxylate reductase 3 | Q53H96-2;Q53H96 | PYCRL | Other | -0.109958796 | 0.257587665 | 26.91273624 | 27.10169393 | 26.90127166 | 26.72168895 | 26.92535613 | 26.70859369 | 26.95438022 | 26.88180367 | 26.46961002 |
| Glycogen phosphorylase, brain form | P11216 | PYGB | Extracellular or secreted | 0.839095841 | 2.45533357 | 32.12730986 | 32.26829827 | 32.21821296 | 31.57145188 | 31.58669675 | 31.7888887 | 32.63456369 | 32.64950198 | 33.01365106 |
| Glycogen phosphorylase, liver form | P06737-2;P06737 | PYGL | Extracellular or secreted | 0.44372833 | 1.455237678 | 31.09270045 | 31.34862992 | 31.30975503 | 30.53120251 | 30.61736859 | 30.57650637 | 31.35888141 | 31.29263506 | 31.41774996 |
| Pygopus homolog 2 | Q9BRQ0 | PYGO2 | Other | 1.188392915 | 0.655465141 | 23.54049606 | 21.47587847 | 23.74724646 | 21.36441245 | 21.94124201 | 22.60159308 | 24.37145762 | 24.26391847 | 22.26523692 |
| Pyridine nucleotide-disulfide oxidored | Q8WU10-2;Q8WU10 | PYROXD1 | Other | 0.163523174 | 0.166572008 | 21.96395157 | 22.20937483 | 22.00305667 | 23.47204915 | 22.45500121 | 22.09979078 | 22.00477184 | 23.01570088 | 22.57170891 |
| Glutamine--tRNA ligase | P47897;P47897-2 | QARS | Other | -0.226522329 | 0.77555244 | 30.18989904 | 30.1639895 | 30.15013181 | 29.90598316 | 29.9616839 | 30.31516487 | 29.74570238 | 29.80854588 | 30.1096109 |
| Dihydropteridine reductase | P09417;P09417-2 | QDPR | Extracellular or secreted | -0.044586559 | 0.153431399 | 29.71782186 | 29.69875987 | 29.59088665 | 29.40458426 | 29.49016148 | 29.32723122 | 29.54944004 | 29.60995744 | 29.32156551 |
| Glutaminyl-peptide cyclotransferase | Q16769;Q16769-2 | QPCT | Extracellular or secreted | -0.957254707 | 0.938747934 | 27.68945856 | 27.57513619 | 27.48031343 | 29.08144268 | 29.10564536 | 29.83248065 | 27.2023778 | 27.13946142 | 28.16863509 |
| Glutamine-rich protein 1 | Q2TAL8 | QRICH1 | Other | 0.145670545 | 0.17110191 | 24.9871547 | 25.28771238 | 25.12218343 | 25.06792404 | 24.73985064 | 24.70487924 | 25.47351699 | 25.36936225 | 24.54898461 |
| Sulfhydryl oxidase 1 | O00391;O00391-2 | QSOX1 | Extracellular or secreted | 0.340678682 | 0.1787255 | 27.69371803 | 27.95454662 | 27.88518303 | 28.77461023 | 28.93404607 | 30.00356715 | 28.10938672 | 28.42973964 | 30.10574525 |
| Sulfhydryl oxidase 2 | Q6ZRP7 | QSOX2 | Extracellular or secreted | 0.361325749 | 0.479687876 | 25.28517416 | 25.37404767 | 25.30768527 | 25.98520127 | 25.99772236 | 25.89312245 | 25.75105794 | 25.69386423 | 26.56053167 |
| Queuine tRNA-ribosyltransferase | Q9BXR0-2;Q9BXR0 | QTRT1 | Other | 0.647576759 | 1.064580654 | 23.37252077 | 23.52826523 | 22.11456348 | 23.59821156 | 23.18092289 | 24.19648026 | 24.24074476 | 24.01529254 | 23.68217507 |
| Queuine tRNA-ribosyltransferase subu | Q9H974-2;Q9H974-3;Q9H974 | QTRTD1 | Other | 0.681518031 | 0.994109932 | 24.95131496 | 24.58640506 | 24.45757476 | 24.79758633 | 24.82216118 | 24.76880542 | 25.61051372 | 25.70880416 | 24.91716008 |
| Ras-related protein Rab-10 | P61026 | RAB10 | Extracellular or secreted | 0.291789464 | 0.46230221 | 27.33756093 | 27.32954262 | 27.50642696 | 27.11852465 | 27.33671003 | 28.12863779 | 27.47328473 | 27.57686673 | 28.20391842 |
| Ras-related protein Rab-11B | Q15907;Q15907-2 | RAB11B | Extracellular or secreted | 0.237505559 | 0.972892905 | 30.11570006 | 30.0555606 | 29.99685006 | 30.02366498 | 29.90555287 | 30.32999163 | 30.16050748 | 30.28162375 | 30.48404555 |
| Rab11 family-interacting protein 1 | Q6WKZ4-2;Q6WKZ4-3;Q6WK | RAB11FIP1 | Other | 1.159135527 | 0.56722161 | 21.45235868 | 21.44299568 | 21.21655642 | 22.26882295 | 20.65677365 | 22.46473319 | 21.21047151 | 23.27511852 | 23.74293684 |
| Ras-related protein Rab-12 | Q6IQ22 | RAB12 | Other | -0.316188856 | 0.3345081 | 26.19338143 | 26.11352583 | 26.48134554 | 26.28252543 | 26.60921775 | 27.18536879 | 25.54203241 | 26.22954863 | 26.71253477 |
| Ras-related protein Rab-13 | P51153 | RAB13 | Extracellular or secreted | 1.092477471 | 0.897435678 | 23.48406861 | 23.40772677 | 21.05398244 | 21.20519995 | 21.3094922 | 24.22479189 | 23.0756639 | 23.84517608 | 23.69922337 |
| Ras-related protein Rab-14 | P61106 | RAB14 | Extracellular or secreted | 0.353245505 | 0.663203642 | 28.77385605 | 28.76303417 | 28.80113157 | 28.90020657 | 28.8937416 | 29.72997557 | 29.06134099 | 29.22271624 | 29.70665204 |
| Ras-related protein Rab-18 | Q9NP72;Q9NP72-2 | RAB18 | Extracellular or secreted | 0.171648069 | 0.186176926 | 25.2461926 | 24.93030566 | 25.16741049 | 25.39545255 | 25.2281361 | 26.14176259 | 25.25039153 | 25.19238463 | 26.12679804 |
| Ras-related protein Rab-1A | P62820;P62820-2;P62820-3 | RAB1A | Extracellular or secreted | -0.211604253 | 0.650671486 | 29.50227815 | 29.70990858 | 29.79532122 | 29.51596741 | 29.40829452 | 29.80883736 | 29.31529446 | 29.263023 | 29.65717341 |
| Ras-related protein Rab-1B;Putative R | Q9H0U4;Q92928 | RAB1B;RAB1C | Extracellular or secreted | -0.299773562 | 0.769494175 | 27.89261304 | 27.92082364 | 27.91330708 | 27.80390486 | 27.96295274 | 27.69032478 | 27.3661134 | 27.47583756 | 27.85049142 |
| Ras-related protein Rab-21 | Q9UL25 | RAB21 | Extracellular or secreted | -0.068710862 | 0.082313785 | 26.82662878 | 26.98782693 | 27.1890507 | 27.03946563 | 26.99905671 | 27.52492234 | 26.97103601 | 26.59263405 | 27.51367291 |
| Ras-related protein Rab-22A | Q9UL26 | RAB22A | Extracellular or secreted | 0.424086582 | 0.868798963 | 25.29466954 | 24.79019614 | 24.99196188 | 25.45707372 | 25.22342994 | 25.39106576 | 25.29551056 | 25.60348463 | 25.94746305 |
| Ras-related protein Rab-23 | Q9ULC3 | RAB23 | Extracellular or secreted | 1.389862207 | 1.567878158 | 23.44303661 | 23.1954443 | 23.38469102 | 20.99330743 | 23.83990114 | 24.04443548 | 24.36979487 | 24.33837732 | 24.91182242 |
| Ras-related protein Rab-25 | P57735 | RAB25 | Extracellular or secreted | -1.541263107 | 0.564666996 | 25.13120166 | 25.21248778 | 25.69032478 | 24.99062095 | 22.83509637 | 26.17159662 | 21.51266388 | 23.83711202 | 25.04209886 |
| Ras-related protein Rab-27B | O00194 | RAB27B | Extracellular or secreted | 0.188240637 | 0.116630869 | 23.10272735 | 22.90205414 | 23.02025294 | 22.39643258 | 21.63319183 | 22.71255051 | 22.06223591 | 22.5633946 | 23.82269608 |
| Ras-related protein Rab-2A | P61019;P61019-2 | RAB2A | Extracellular or secreted | 0.009543316 | 0.022084635 | 28.16202139 | 28.19774605 | 28.23031852 | 28.29307377 | 28.28537703 | 28.60392319 | 28.09687758 | 28.28590612 | 28.53207621 |
| Ras-related protein Rab-2B | Q8WUD1 | RAB2B | Extracellular or secreted | 1.619249889 | 0.65168697 | 21.85285255 | 21.83834374 | 21.64601193 | 21.36826346 | 21.48878021 | 22.53606694 | 22.80565729 | 25.24350757 | 22.17374422 |
| Ras-related protein Rab-35 | Q15286;Q15286-2 | RAB35 | Extracellular or secreted | 1.350137666 | 2.723341138 | 24.87422382 | 24.82255021 | 24.94046774 | 25.04797492 | 25.00598837 | 25.94415959 | 26.19953501 | 26.376898 | 26.79166231 |
| Ras-related protein Rab-3D | O95716 | RAB3D | Extracellular or secreted | 0.111235742 | 0.329437601 | 25.6027487 | 25.96507967 | 25.87027779 | 25.86752327 | 26.27788727 | 26.51559903 | 26.22354042 | 26.0342267 | 26.12549798 |
| Rab3 GTPase-activating protein cataly | Q15042-4;Q15042;Q15042-3 | RAB3GAP1 | Extracellular or secreted | -0.79131503 | 1.230854058 | 27.5734758 | 27.36678043 | 27.46185785 | 26.98196143 | 26.99258867 | 27.05631026 | 26.3889987 | 26.881687 | 26.07185643 |
| Rab3 GTPase-activating protein non-c | Q9H2M9 | RAB3GAP2 | Extracellular or secreted | -0.578409382 | 0.777749471 | 24.59320417 | 24.80040347 | 24.72977465 | 23.96463911 | 24.84866611 | 24.5574235 | 24.32189232 | 24.25616318 | 23.43377185 |
| Ras-related protein Rab-4A | P20338 | RAB4A | Extracellular or secreted | 0.873573837 | 0.61405689 | 24.37980879 | 24.28827582 | 24.25263079 | 23.75389083 | 21.08734298 | 25.35454157 | 24.37232149 | 24.39329356 | 25.41335186 |
| Ras-related protein Rab-5A | P20339-2;P20339 | RAB5A | Extracellular or secreted | 0.592616033 | 0.709527077 | 24.13836806 | 24.17691211 | 24.45644717 | 23.82687119 | 21.84594303 | 24.19670537 | 24.27419442 | 24.33046614 | 24.49381099 |
| Ras-related protein Rab-5B | P61020;P61020-2 | RAB5B | Extracellular or secreted | 0.501583826 | 0.498118231 | 23.417253 | 24.19324981 | 24.23695548 | 23.78206798 | 22.21782875 | 24.65184703 | 23.82997043 | 24.08958238 | 24.8347997 |
| Ras-related protein Rab-5C | P51148;P51148-2 | RAB5C | Extracellular or secreted | 0.131266032 | 0.179418053 | 27.98343004 | 28.22655684 | 28.28387689 | 28.462209 | 28.371441 | 28.48496747 | 28.1075422 | 28.26087159 | 28.93162492 |
| Ras-related protein Rab-6A | P20340-2;P20340;P20340-4 | RAB6A | Extracellular or secreted | -0.175710046 | 0.202330335 | 29.11589846 | 29.20487464 | 29.15754717 | 29.46600764 | 29.45071336 | 29.67456845 | 28.7597368 | 28.98457127 | 29.76336666 |
| Ras-related protein Rab-6B | Q9NRW1;Q9NRW1-2 | RAB6B | Other | -1.340788478 | 0.554856861 | 23.56395313 | 23.40379637 | 23.38020527 | 23.0392397 | 23.50118118 | 23.57137955 | 22.8239596 | 23.1291352 | 20.25441736 |
| Ras-related protein Rab-7a | P51149 | RAB7A | Extracellular or secreted | 0.507884672 | 2.260086665 | 29.41191449 | 29.28420794 | 29.29353427 | 29.60448887 | 29.56743177 | 29.91641422 | 29.86528873 | 30.07584442 | 30.12151664 |
| Ras-related protein Rab-8A | P61006;P61006-2 | RAB8A | Extracellular or secreted | 0.85425845 | 2.242233996 | 26.08371142 | 25.52220053 | 25.81369803 | 25.82776775 | 25.6988261 | 26.86112289 | 26.69139019 | 26.95060324 | 26.82444527 |
| Prenylated Rab acceptor protein 1 | Q9UI14 | RABAC1 | Extracellular or secreted | -0.813204118 | 0.283876212 | 23.02472306 | 23.6073845 | 23.49716575 | 23.0069814 | 23.5659295 | 21.94885342 | 20.25388623 | 22.95027395 | 23.68174627 |
| Rab GTPase-binding effector protein 1 | Q15276;Q15276-2 | RABEP1 | Other | 0.522651621 | 0.405357034 | 25.45500507 | 25.89101417 | 25.07422706 | 24.92288633 | 25.25889665 | 23.59991466 | 25.99440264 | 26.01209833 | 24.66242586 |
| Rab GTPase-binding effector protein 2 | Q9H5N1;Q9H5N1-2 | RABEP2 | Other | 0.744717706 | 0.699949452 | 24.36185384 | 24.16077193 | 24.08715591 | 24.13500467 | 24.38324127 | 21.54589298 | 24.83200069 | 24.75913358 | 23.97997915 |
| Rab9 effector protein with kelch moti | Q7Z6M1;Q7Z6M1-2 | RABEPK | Extracellular or secreted | 0.473049172 | 1.606937593 | 26.50301785 | 25.82738012 | 26.03942379 | 26.28483892 | 26.42809483 | 25.96607044 | 26.56083775 | 26.86749971 | 26.51522303 |
| Rab GTPase-activating protein 1 | Q9Y3P9;Q9Y3P9-2 | RABGAP1 | Other | 0.462179995 | 1.545456672 | 28.61967897 | 28.68006335 | 28.67394538 | 28.47243279 | 28.63325623 | 28.79829113 | 29.27070595 | 29.13052921 | 28.92413875 |
| Rab GTPase-activating protein 1-like | Q5R372-4;Q5R372;Q5R372-3 | RABGAP1L | Other | 0.050038213 | 0.06977326 | 25.09285164 | 24.35339777 | 24.83793008 | 25.34996094 | 25.44284683 | 24.36125148 | 25.17558031 | 24.62527853 | 25.06837517 |
| Rab GTPase-activating protein 1-like;R | Q5R372-7;Q5R372-8;Q5R372 | RABGAP1L | Other | 0.275907128 | 0.866491189 | 22.54250481 | 22.40901277 | 22.45460353 | 22.24745356 | 22.7603423 | 23.02155816 | 22.77088323 | 23.06943272 | 22.70514299 |
| Geranylgeranyl transferase type-2 sub | Q92696 | RABGGTA | Other | 0.384357461 | 1.272068693 | 27.65574132 | 27.44640095 | 27.39304802 | 26.85162362 | 27.06476217 | 26.97781951 | 27.75729061 | 27.67445059 | 27.41602898 |
| Geranylgeranyl transferase type-2 sub | P53611 | RABGGTB | Other | 0.943739531 | 2.873485805 | 25.89887554 | 25.97973935 | 25.86273156 | 26.24890882 | 25.93957132 | 25.64460446 | 26.90735923 | 27.03359675 | 26.67747814 |
| Guanine nucleotide exchange factor M | P47224 | RABIF | Other | 1.722988539 | 2.887942493 | 24.11418142 | 23.37278643 | 23.7427313 | 22.85261968 | 22.49450129 | 24.16484311 | 25.08180328 | 25.2898945 | 25.16809945 |
| Rab-like protein 6 | Q3YEC7;Q3YEC7-2;Q3YEC7-3 | RABL6 | Other | -0.244037432 | 0.472345726 | 26.80180998 | 27.08911762 | 26.97059727 | 26.12573444 | 25.85493171 | 26.23024521 | 26.130377 | 26.1790784 | 26.49465041 |
| Ras-related C3 botulinum toxin substr | P63000-2;P63000;P15153;P6 | RAC1;RAC2;RAC3 | Extracellular or secreted | 0.910320615 | 1.274651511 | 28.19544806 | 28.29911437 | 28.31484085 | 27.65744603 | 27.61083753 | 29.15872721 | 28.75496766 | 28.94008123 | 29.65411998 |
| Rac GTPase-activating protein 1 | Q9H0H5 | RACGAP1 | Extracellular or secreted | -0.039122909 | 0.058863249 | 24.93880893 | 24.98259257 | 24.80316583 | 24.61284915 | 24.35534841 | 25.50306336 | 24.50809071 | 24.87525552 | 25.09719918 |
| Cell cycle checkpoint protein RAD1 | O60671-2;O60671;O60671-3 | RAD1 | Other | 0.376945517 | 0.702508312 | 23.4891012 | 23.74386141 | 23.92759569 | 23.53017198 | 23.95730613 | 23.12748305 | 24.201948 | 24.21741463 | 23.59923366 |
| Double-strand-break repair protein ra | O60216 | RAD21 | Other | -0.324568814 | 0.481056078 | 29.50584041 | 29.48029417 | 29.50737251 | 28.88471738 | 28.98525014 | 28.63941082 | 29.11671656 | 29.05700787 | 28.35401184 |
| UV excision repair protein RAD23 hom | P54725-3;P54725;P54725-2 | RAD23A | Other | 0.420362097 | 1.495645985 | 30.6539492 | 30.54333113 | 30.47033951 | 30.61324622 | 30.65999932 | 29.97773764 | 30.98983094 | 31.03262503 | 30.69793183 |
| UV excision repair protein RAD23 hom | P54727;P54727-2 | RAD23B | Other | 0.344537439 | 0.608022119 | 30.89258409 | 30.88103055 | 30.78986046 | 30.5420398 | 30.61815664 | 30.41104571 | 31.23888324 | 31.26074824 | 30.60133946 |
| DNA repair protein RAD50 | Q92878-3;Q92878;Q92878-2 | RAD50 | Other | -0.084445141 | 0.435824814 | 27.16892196 | 27.14773238 | 27.31605015 | 27.27914565 | 27.20069525 | 27.64485197 | 27.2756869 | 27.22591362 | 27.12376273 |
| mRNA export factor | P78406 | RAE1 | Other | -0.356523101 | 2.039840986 | 28.57455888 | 28.53411898 | 28.61645742 | 28.74947957 | 28.83731658 | 28.35934233 | 28.33790115 | 28.30011868 | 28.12804775 |
| Ras-related protein Ral-A | P11233 | RALA | Extracellular or secreted | 1.052439889 | 2.574775791 | 27.65880835 | 27.62554331 | 27.54738209 | 26.70252972 | 26.55841651 | 27.54318361 | 28.33985585 | 28.47371052 | 28.1616851 |
| Ras-related protein Ral-B | P11234;P11234-2;P11234-3 | RALB | Extracellular or secreted | 0.703889723 | 2.685869477 | 24.28616178 | 24.20806523 | 24.43084045 | 23.56790317 | 24.22905349 | 24.37331762 | 24.75062358 | 24.90504209 | 25.00367437 |
| RalA-binding protein 1 | Q15311 | RALBP1 | Other | -0.148792997 | 0.532462501 | 24.75654003 | 24.3990437 | 24.3027092 | 24.17843265 | 24.24176325 | 24.26556469 | 24.07651226 | 24.157152 | 24.39198351 |
| RNA-binding protein Raly | Q9UKM9-2;Q9UKM9 | RALY | Other | -0.884067489 | 0.399615444 | 27.62002871 | 27.73666752 | 27.68438405 | 27.17674094 | 27.2950025 | 27.95243742 | 27.52312832 | 27.52357703 | 25.03372276 |
| GTP-binding nuclear protein Ran | P62826 | RAN | Extracellular or secreted | -0.032414643 | 0.102532372 | 31.3089418 | 31.60924417 | 31.45537553 | 30.93845563 | 31.02769071 | 30.98996625 | 31.22302709 | 31.20244786 | 31.14211816 |
| Ran-specific GTPase-activating protein | P43487-2;P43487 | RANBP1 | Other | 0.704216059 | 1.109913774 | 30.67429905 | 30.60116232 | 30.72630933 | 31.08948388 | 31.19560059 | 29.97432233 | 31.63222502 | 31.68608032 | 30.92493158 |
| Ran-binding protein 10 | Q6VN20-3;Q6VN20;Q6VN20- | RANBP10 | Other | -0.068571195 | 0.060840228 | 25.38175707 | 25.63478004 | 25.23480135 | 25.84412263 | 25.63303446 | 25.63225796 | 26.05266688 | 25.60690465 | 24.81509164 |
| E3 SUMO-protein ligase RanBP2 | P49792 | RANBP2 | Other | -1.046623637 | 1.357381538 | 28.31777597 | 28.2318296 | 28.17421871 | 27.92840922 | 28.00629877 | 28.52854097 | 26.75340632 | 27.06146975 | 27.63878964 |
| Ran-binding protein 3 | Q9H6Z4-3;Q9H6Z4 | RANBP3 | Other | 0.108401724 | 0.328626041 | 29.31921975 | 29.34697233 | 29.36773874 | 29.22798009 | 29.40084407 | 29.21113272 | 29.60084342 | 29.4719485 | 29.1893571 |
| Ran-binding protein 9 | Q96S59;Q96S59-2;Q96S59-3 | RANBP9 | Other | 0.730279015 | 1.149873921 | 24.61952506 | 24.59388802 | 24.55490872 | 24.3405521 | 25.13953949 | 24.56651027 | 25.46699948 | 25.71609873 | 24.91520067 |
| Ran GTPase-activating protein 1 | P46060 | RANGAP1 | Other | 0.493426131 | 1.163738986 | 28.71845893 | 28.81286611 | 28.80593522 | 28.92422372 | 28.98535873 | 28.97350145 | 29.57614593 | 29.43499682 | 29.07930773 |
| Ran guanine nucleotide release factor | Q9HD47-4;Q9HD47-2;Q9HD4 | RANGRF | Extracellular or secreted | -0.156682616 | 0.209898839 | 22.84237797 | 22.37642113 | 22.72953611 | 23.95162622 | 24.01188513 | 23.00166589 | 23.19829515 | 22.98224439 | 22.80616884 |
| Ras-related protein Rap-1A | P62834 | RAP1A | Extracellular or secreted | 0.90633544 | 1.272067673 | 22.23733485 | 20.66364576 | 22.32251134 | 23.06955386 | 21.16283111 | 21.55888232 | 22.50932367 | 22.75517528 | 22.96188699 |
| Ras-related protein Rap-1b;Ras-relate | P61224;P61224-3;P61224-2;/ | RAP1B | Extracellular or secreted | 0.905641822 | 1.937268117 | 27.96400007 | 27.91222228 | 27.97311821 | 28.08430973 | 28.10838998 | 28.68435061 | 28.8232004 | 28.89512939 | 29.36179111 |
| Rap1 GTPase-GDP dissociation stimula | P52306-4;P52306;P52306-5;P | RAP1GDS1 | Extracellular or secreted | 0.264558029 | 0.282739043 | 25.50112042 | 26.43987044 | 26.51304034 | 25.14856657 | 24.88548562 | 25.57919968 | 26.40960199 | 25.9611098 | 25.45660383 |
| Ras-related protein Rap-2b;Ras-relate | P61225;P10114 | RAP2B;RAP2A | Extracellular or secreted | 0.479819402 | 0.945814892 | 25.36014649 | 25.32178913 | 25.42460739 | 25.7411888 | 25.19197066 | 26.15485596 | 25.8480691 | 25.79014642 | 26.39852191 |
| Ras-associated and pleckstrin homolo | Q70E73;Q70E73-5;Q70E73-2 | RAPH1 | Extracellular or secreted | 0.421821632 | 0.18039117 | 20.79661194 | 23.00855514 | 23.27951763 | 21.40172607 | 20.77303082 | 23.66654909 | 23.26685172 | 23.31837952 | 21.14322901 |
| Retinoic acid receptor responder prot | P49788;P49788-2 | RARRES1 | Extracellular or secreted | 1.188289216 | 1.54986359 | 30.29438909 | 30.24909424 | 30.21731755 | 31.31996263 | 31.47682267 | 32.58884235 | 31.91812172 | 32.23834231 | 32.48161788 |

| | | | | | | | | | | | | | | |
|---|---|---|---|---|---|---|---|---|---|---|---|---|---|---|
| Arginine--tRNA ligase, cytoplasmic | P54136 | RARS | Extracellular or secreted | -0.375837605 | 1.244815924 | 31.09440044 | 31.16871877 | 31.18157084 | 31.09502956 | 31.11947732 | 30.84460755 | 30.81772238 | 30.8195507 | 30.48711634 |
| Ras association domain-containing pr | O75901 | RASSF9 | Extracellular or secreted | -5.775040152 | 3.295492466 | 26.97377513 | 27.77077236 | 27.81207044 | 26.85816108 | 26.86573119 | 26.52978488 | 20.62597853 | 21.89270292 | 21.56134564 |
| Ribonucleoprotein PTB-binding 1 | Q8IY67;Q8IY67-2 | RAVER1 | Other | -0.977230762 | 1.724015905 | 25.212228 | 24.91131957 | 25.36379308 | 25.7677957 | 25.76655782 | 25.60766669 | 24.73727375 | 24.05125674 | 24.59445765 |
| Histone-binding protein RBBP4 | Q09028-3;Q09028;Q09028-2 | RBBP4 | Other | -0.309248749 | 0.819086147 | 30.76168759 | 30.86599661 | 30.65411998 | 29.81735644 | 29.94642595 | 30.34540802 | 30.14698023 | 30.14685887 | 29.97391195 |
| Retinoblastoma-binding protein 5 | Q15291;Q15291-2 | RBBP5 | Other | -0.056084274 | 0.302864465 | 27.90058088 | 28.02638523 | 28.03033761 | 27.94837686 | 27.80119326 | 27.85424216 | 27.93128677 | 27.93797881 | 27.7430396 |
| E3 ubiquitin-protein ligase RBBP6 | Q7Z6E9-2;Q7Z6E9;Q7Z6E9-4 | RBBP6 | Other | -1.508968428 | 1.874171227 | 25.81394262 | 25.89330764 | 25.61040107 | 26.14330115 | 24.24742458 | 25.55265339 | 23.45525597 | 24.16023336 | 24.48812061 |
| Histone-binding protein RBBP7 | Q16576;Q16576-2 | RBBP7 | Other | 0.065262536 | 0.127336769 | 31.91712592 | 32.02115146 | 31.93863089 | 31.1288467 | 31.2366613 | 31.28300477 | 31.81734119 | 31.65667916 | 31.48447778 |
| Putative hydrolase RBBP9 | O75884;O75884-2 | RBBP9 | Extracellular or secreted | -0.236263712 | 0.702597083 | 29.28270658 | 29.42682401 | 29.32761671 | 28.87105963 | 29.03262503 | 28.40681562 | 28.82647726 | 28.91013604 | 28.72841936 |
| RNA binding protein fox-1 homolog 2; | O43251-9;Q9NWB1-3;O4325 | RBFOX2;RBFOX1; | Other | -0.815883799 | 0.296692087 | 24.40535548 | 24.33599489 | 24.33032936 | 23.07380227 | 23.82561022 | 24.07152914 | 24.49185554 | 23.87351996 | 21.20828378 |
| Ribokinase | Q9H477-2;Q9H477 | RBKS | Extracellular or secreted | 0.009233616 | 0.012976873 | 24.52731091 | 24.39675946 | 24.32147949 | 23.66068624 | 23.47630123 | 23.10061165 | 23.93714821 | 23.88781112 | 23.944316 |
| RNA-binding protein 10 | P98175-2;P98175;P98175-5;F | RBM10 | Other | -0.373868093 | 1.509221713 | 27.98869387 | 28.10474598 | 27.97678217 | 28.06363125 | 27.9712005 | 27.51705696 | 27.64883371 | 27.66593949 | 27.37467788 |
| RNA-binding protein 12 | Q9NTZ6 | RBM12 | Other | -0.273312767 | 0.808190921 | 27.94480748 | 27.92936904 | 28.0992376 | 28.43880822 | 28.25223375 | 28.26279904 | 28.04390381 | 27.97486932 | 27.62491613 |
| RNA-binding protein 12B | Q8IXT5 | RBM12B | Other | -0.110288226 | 0.132876661 | 23.11454684 | 22.94304632 | 23.57114805 | 22.50246528 | 23.26337126 | 22.33041158 | 23.09754481 | 23.07370422 | 22.36038095 |
| RNA-binding protein 14 | Q96PK6;Q96PK6-2 | RBM14 | Other | -0.27334033 | 0.687461324 | 23.89901388 | 23.80429885 | 23.57207382 | 24.59741609 | 24.57103229 | 24.44493298 | 24.00829867 | 23.78396699 | 23.8320973 |
| RNA-binding protein 14 | Q96PK6-5 | RBM14 | Other | -0.518211988 | 0.537174249 | 24.79297811 | 24.69927633 | 24.74149744 | 24.4652565 | 25.01763047 | 27.03191553 | 24.84771078 | 24.13992975 | 24.83200069 |
| Putative RNA-binding protein 15 | Q96T37-4;Q96T37-2;Q96T37 | RBM15 | Other | -1.620319136 | 1.846688933 | 25.80375708 | 25.62153859 | 25.49987435 | 25.04239114 | 25.1176729 | 25.59115068 | 23.33619925 | 23.85946502 | 24.28157069 |
| Splicing factor 45 | Q96I25 | RBM17 | Other | 0.636668002 | 2.381841907 | 25.60487058 | 25.0709562 | 25.20627751 | 25.78321768 | 25.24724348 | 25.78686066 | 26.11996949 | 25.92998976 | 26.20975781 |
| RNA-binding protein 25 | P49756;P49756-2;P49756-3 | RBM25 | Other | -1.038254922 | 2.614156147 | 28.53675174 | 28.49548934 | 28.49381099 | 28.20349841 | 28.09602301 | 28.97374776 | 27.32817335 | 27.2514213 | 27.70530122 |
| RNA-binding protein 26 | Q5T8P6-3;Q5T8P6-2;Q5T8P6 | RBM26 | Other | -1.440968382 | 1.80061369 | 26.10029082 | 25.80955815 | 25.73280484 | 25.91182242 | 25.70218622 | 26.12492637 | 24.33735676 | 24.13766474 | 24.89286776 |
| RNA-binding protein 28 | Q9NW13;Q9NW13-2 | RBM28 | Other | -1.703697717 | 0.597704711 | 23.76860354 | 24.41654448 | 24.47753694 | 24.27084851 | 24.1958798 | 25.93862948 | 20.7975339 | 23.10910372 | 24.5162906 |
| RNA-binding protein 3 | P98179 | RBM3 | Extracellular or secreted | 1.081857944 | 1.242618138 | 28.01094668 | 28.00356715 | 28.24054461 | 30.27519846 | 30.00088406 | 28.2979345 | 29.75748137 | 29.6203434 | 30.2822868 |
| RNA-binding protein 38 | Q9H0Z9 | RBM38 | Other | -0.100321538 | 0.054593478 | 25.20452488 | 25.2671733 | 25.16645305 | 25.10677378 | 25.34709064 | 24.43942662 | 25.46653282 | 25.67951983 | 23.81870387 |
| RNA-binding protein 39 | Q14498-3;Q14498-2;Q14498 | RBM39 | Other | -1.096331913 | 2.189491922 | 27.25574912 | 27.41739788 | 27.27799365 | 27.03684854 | 26.98239673 | 27.39345723 | 26.29705115 | 26.25780029 | 25.83807439 |
| RNA-binding protein 4 | Q9BWF3;Q9BWF3-4;Q9BWF3 | RBM4 | Other | 1.789697032 | 2.324254223 | 26.60794882 | 26.14914828 | 26.32392034 | 25.10940665 | 26.63089809 | 27.80396643 | 28.59331104 | 28.34975852 | 27.73866584 |
| RNA-binding protein 4B | Q9BQ04 | RBM4B | Other | 1.281777368 | 0.537375361 | 21.22244534 | 22.60730112 | 19.48023618 | 20.8837 | 22.03167117 | 22.62390232 | 23.55578647 | 23.73675782 | 20.97741588 |
| RNA-binding protein 8A | Q9Y5S9-2;Q9Y5S9 | RBM8A | Other | 0.146275137 | 0.598276396 | 30.24320337 | 30.35940517 | 30.21280315 | 30.61016871 | 30.67219261 | 30.00958591 | 30.44529617 | 30.43827285 | 30.60893586 |
| RNA-binding motif, single-stranded-in | P29558-2;P29558 | RBMS1 | Other | 0.025413428 | 0.020543742 | 22.3557651 | 22.22152104 | 22.24074476 | 20.92516059 | 21.00644271 | 22.82061372 | 22.42890636 | 21.96548328 | 21.4669746 |
| RNA-binding motif protein, X chromos | P38159;P38159-2 | RBMX | Extracellular or secreted | -0.064207366 | 0.161217937 | 31.65283868 | 31.42096499 | 31.33346138 | 31.94677452 | 31.64224588 | 31.19829656 | 31.25490486 | 31.57704688 | 31.57271717 |
| Retinol-binding protein 1 | P09455;P09455-2;P09455-3 | RBP1 | Other | -0.333944361 | 0.365688318 | 29.47645575 | 29.54076578 | 29.48947258 | 28.69175624 | 28.64835374 | 29.66399042 | 28.61473865 | 28.57250034 | 29.56632518 |
| Retinol-binding protein 4;Plasma retir | P02753 | RBP4 | Extracellular or secreted | -2.969409297 | 1.340262781 | 23.30284862 | 22.37729527 | 22.48308459 | 27.99480184 | 27.73705451 | 22.65298498 | 22.41162684 | 20.67155583 | 21.28262435 |
| Recombining binding protein suppres | Q06330-5;Q06330-4;Q06330 | RBPJ | Other | -0.25784164 | 0.276165742 | 26.30940377 | 25.99735627 | 26.03705809 | 26.11680082 | 26.07928229 | 26.12585265 | 26.16470887 | 26.22295111 | 25.17169205 |
| E3 ubiquitin-protein ligase RBX1;E3 ub | P62877 | RBX1 | Other | 0.722488639 | 0.74998808 | 27.77912709 | 27.74188313 | 27.68096875 | 27.23434456 | 27.36461146 | 27.65013569 | 28.60713754 | 28.7096457 | 27.57621802 |
| Regulator of chromosome condensati | P18754;P18754-2 | RCC1 | Other | -0.273565538 | 1.013406045 | 28.06080006 | 28.14981696 | 27.99636522 | 28.62512522 | 28.66315914 | 28.68475188 | 28.13222201 | 28.10938672 | 28.02770389 |
| Protein RCC2 | Q9P258 | RCC2 | Extracellular or secreted | -0.453411904 | 0.758601533 | 31.75907006 | 31.92039525 | 31.81310319 | 31.62270139 | 31.66581246 | 31.16692467 | 31.50597288 | 31.35405388 | 30.75374103 |
| RCC1 domain-containing protein 1 | A6NED2 | RCCD1 | Extracellular or secreted | 0.622323218 | 1.268487471 | 21.77410497 | 21.79962313 | 21.76376998 | 21.57030387 | 22.30929749 | 23.09811243 | 22.71357866 | 22.42349861 | 22.88749832 |
| RING finger and CHY zinc finger doma | Q96PM5-6;Q96PM5-2;Q96PM | RCHY1 | Other | 1.507270034 | 1.150150293 | 23.72889289 | 20.55973029 | 21.21716192 | 24.07659381 | 21.87051901 | 24.13390792 | 23.60354123 | 24.50845345 | 24.20321834 |
| Reticulocalbin-1 | Q15293;Q15293-2 | RCN1 | Extracellular or secreted | 0.089330097 | 0.237431229 | 30.17712128 | 30.51210981 | 30.54637042 | 30.51962729 | 30.72167266 | 29.91384917 | 30.6223523 | 30.53473125 | 30.30628205 |
| Reticulocalbin-2 | Q14257;Q14257-2 | RCN2 | Extracellular or secreted | 1.050346807 | 1.933350227 | 26.6416862 | 26.5472497 | 26.76002244 | 27.18999327 | 27.2525947 | 28.09672681 | 27.91176528 | 27.96834661 | 28.51506509 |
| Reticulocalbin-3 | Q96D15 | RCN3 | Extracellular or secreted | -1.002356841 | 0.436429076 | 24.04785014 | 24.71093336 | 24.21867143 | 25.78324266 | 25.35168038 | 22.74886437 | 24.55578647 | 24.05531752 | 21.81244666 |
| REST corepressor 1 | Q9UKL0 | RCOR1 | Extracellular or secreted | 0.046094183 | 0.054675536 | 27.44134357 | 27.42061369 | 27.44900175 | 27.2337048 | 27.17417108 | 26.91695515 | 27.57036647 | 27.59875109 | 26.78706001 |
| Retinol dehydrogenase 11 | Q8TC12;Q8TC12-2;Q8TC12-3 | RDH11 | Extracellular or secreted | -0.484747874 | 0.242733731 | 22.70682986 | 22.88063654 | 23.49313911 | 22.84806432 | 22.21320748 | 26.51609519 | 22.11617529 | 22.88382758 | 23.87473976 |
| Radixin | P35241;P35241-5;P35241-4 | RDX | Extracellular or secreted | 0.029423136 | 0.228511298 | 29.92549762 | 29.70724469 | 29.75168369 | 29.81060037 | 29.75155601 | 29.65836573 | 29.79864649 | 29.857479 | 29.73461797 |
| ATP-dependent DNA helicase Q1 | P46063 | RECQL | Other | -0.03583231 | 0.064752761 | 28.15537723 | 28.21955792 | 28.16221352 | 27.90798949 | 28.15170502 | 28.51814568 | 28.40202409 | 28.22273927 | 27.82523414 |
| Receptor expression-enhancing prote | Q00765 | REEP5 | Extracellular or secreted | 0.334372775 | 0.212237732 | 26.05382714 | 25.89737594 | 25.59454307 | 25.13453474 | 25.03942379 | 25.36044794 | 24.77193109 | 26.1574026 | 26.61386094 |
| Receptor expression-enhancing prote | Q96HR9-2;Q96HR9 | REEP6 | Other | -0.428969395 | 0.414262826 | 23.59605141 | 23.52778815 | 24.78681082 | 24.34245235 | 23.38758615 | 24.70001758 | 23.41905492 | 23.13522392 | 24.3291662 |
| Transcription factor p65 | Q04206;Q04206-2;Q04206-3 | RELA | Other | -0.205878641 | 0.560767914 | 27.23288184 | 26.76673473 | 26.77291276 | 26.82989787 | 26.47551291 | 26.97563477 | 26.89434892 | 26.54114626 | 26.47365633 |
| RELT-like protein 1 | Q8IUW5 | RELL1 | Extracellular or secreted | 1.857891097 | 2.36521238 | 22.52426111 | 21.12085487 | 22.10380004 | 21.67607272 | 21.31807801 | 23.77577178 | 23.81274372 | 23.73200259 | 24.28834624 |
| Replication initiator 1 | Q9BWE0;Q9BWE0-4 | REPIN1 | Extracellular or secreted | 0.119519897 | 0.081745031 | 24.59985792 | 24.43262549 | 24.18895641 | 24.48744606 | 24.32532797 | 24.75938759 | 24.57415463 | 23.75643823 | 25.42476755 |
| RalBP1-associated Eps domain-contai | Q96D71-3;Q96D71;Q96D71-2 | REPS1 | Extracellular or secreted | 0.262505073 | 0.366570458 | 25.92567305 | 25.21089128 | 25.15367758 | 24.30486877 | 24.03975845 | 25.48833517 | 25.45550683 | 25.09365773 | 25.2999528 |
| RalBP1-associated Eps domain-contai | Q8NFH8-4;Q8NFH8;Q8NFH8- | REPS2 | Extracellular or secreted | -0.112544376 | 0.085344441 | 23.10298359 | 23.58416979 | 24.00864062 | 21.75769046 | 22.30564428 | 23.56267285 | 22.48328144 | 23.56859912 | 22.7713871 |
| Protein RER1 | O15258 | RER1 | Extracellular or secreted | 0.09535216 | 0.031507386 | 20.4137477 | 20.6466924 | 22.54500593 | 21.45340077 | 21.17017576 | 26.4171886 | 22.1643521 | 21.33886693 | 23.10594303 |
| Oligoribonuclease, mitochondrial | Q9Y3B8-3;Q9Y3B8-2;Q9Y3B8 | REXO2 | Other | -0.064881233 | 0.169621587 | 27.60066622 | 27.3691126 | 27.71960168 | 27.12287459 | 27.76825016 | 27.00206524 | 27.43173325 | 27.22048077 | 27.44442752 |
| Replication factor C subunit 1 | P35251-2;P35251 | RFC1 | Extracellular or secreted | -1.424228212 | 1.555934881 | 23.31174388 | 24.26018965 | 23.15810793 | 23.53599578 | 23.53765541 | 26.09087482 | 22.68497726 | 22.0796241 | 22.90999773 |
| Replication factor C subunit 2 | P35250;P35250-2 | RFC2 | Other | 0.459597647 | 0.267508539 | 23.70292598 | 21.31379586 | 23.42764738 | 23.49338347 | 20.75851372 | 25.33422254 | 23.19634518 | 23.67217238 | 23.52551985 |
| Replication factor C subunit 3 | P40938;P40938-2 | RFC3 | Other | -0.639736513 | 0.430570633 | 23.16433676 | 22.98020584 | 22.92801155 | 23.70956682 | 23.2185384 | 25.89145462 | 22.31423582 | 22.81129394 | 23.9013177 |
| Replication factor C subunit 4 | P35249;P35249-2 | RFC4 | Other | -0.237138874 | 0.458803256 | 24.86759397 | 24.64509943 | 24.77329015 | 25.08828877 | 25.19779291 | 25.67669831 | 24.60817449 | 24.65758232 | 25.14720834 |
| MHC class II regulatory factor RFX1 | P22670 | RFX1 | Other | 0.054806539 | 0.099165843 | 25.93660919 | 25.77818724 | 25.92608041 | 25.31620556 | 25.20586751 | 25.76103762 | 25.94042293 | 25.76835114 | 25.41763931 |
| RanBP2-like and GRIP domain-contain | A6NKT7;Q7Z3J3;Q99666;O14 | RGPD3;RGPD4;R | Other | -1.042106053 | 0.619385269 | 23.78835512 | 23.26026145 | 23.60602922 | 22.30923488 | 22.16974302 | 23.96031451 | 23.34716215 | 21.1624042 | 21.91108458 |
| Regulator of G-protein signaling 10 | O43665-3;O43665-2;O43665 | RGS10 | Extracellular or secreted | -0.032443367 | 0.071800531 | 25.8976529 | 25.04485233 | 25.35460882 | 25.1790784 | 25.632147 | 24.95433584 | 25.30851874 | 25.42758344 | 25.19790537 |
| Inactive rhomboid protein 2 | Q6PJF5-2;Q6PJF5 | RHBDF2 | Extracellular or secreted | 0.91815524 | 1.139576644 | 23.03360515 | 23.0322267 | 23.02546673 | 22.94851054 | 23.1205233 | 23.36906266 | 23.50663882 | 24.05977947 | 24.45274497 |
| GTP-binding protein Rheb | Q15382 | RHEB | Extracellular or secreted | 0.225909283 | 0.275550065 | 24.79857543 | 24.87384847 | 25.03825189 | 25.05001136 | 24.36579646 | 25.73531224 | 24.66177375 | 25.41331957 | 25.53353245 |
| Rho-related GTP-binding protein RhoB | P62745 | RHOB | Extracellular or secreted | 1.19924088 | 1.21375144 | 26.26189391 | 26.20884667 | 26.63339483 | 26.40749651 | 26.49570281 | 27.16614653 | 27.18366627 | 27.60349878 | 28.39729821 |
| Rho-related GTP-binding protein RhoC | P08134;P61586 | RHOC;RHOA | Extracellular or secreted | 0.357730026 | 0.568019284 | 26.54820563 | 26.97344671 | 26.86289705 | 26.62463729 | 26.82043356 | 27.16939994 | 26.82031182 | 27.11277053 | 27.63961782 |
| Rho-related GTP-binding protein RhoG | P84095 | RHOG | Extracellular or secreted | 0.514162756 | 0.973924784 | 24.50403385 | 23.39204904 | 24.13531788 | 23.47741341 | 23.54545378 | 24.82118813 | 24.23717436 | 24.64790105 | 24.5951409 |
| Rhophilin-2 | Q8IUC4;Q8IUC4-2 | RHPN2 | Other | 0.943629355 | 1.571381251 | 24.89017928 | 24.49801844 | 24.72296592 | 24.44499615 | 24.5387805 | 24.97853975 | 25.91499546 | 25.71936668 | 25.23326595 |
| Synembryn-A | Q9NPQ8-4;Q9NPQ8;Q9NPQ8 | RIC8A | Extracellular or secreted | 1.041185212 | 0.911860453 | 24.51797304 | 21.28154042 | 23.90930642 | 24.79466454 | 24.09055182 | 25.0126525 | 25.15611055 | 25.02960065 | 24.7411888 |
| Telomere-associated protein RIF1 | Q5UIP0-2;Q5UIP0 | RIF1 | Extracellular or secreted | -0.959098486 | 1.661739038 | 27.55596195 | 27.59106505 | 27.59860913 | 27.76281247 | 27.39737982 | 27.51382348 | 26.53549158 | 26.36614676 | 26.93089215 |
| E3 ubiquitin-protein ligase RING1 | Q06587;Q06587-2 | RING1 | Other | -0.237651522 | 0.590967073 | 25.96214743 | 25.53038038 | 25.56935268 | 25.37032716 | 25.52375648 | 26.25013849 | 25.70714593 | 25.42560009 | 25.25735072 |
| Serine/threonine-protein kinase RIO1 | Q9BRS2 | RIOK1 | Other | -2.981959013 | 0.879645594 | 22.02869893 | 24.34550108 | 24.192648 | 24.94793117 | 24.67809633 | 25.88462423 | 23.81479836 | 19.44682545 | 20.83124902 |
| Receptor-interacting serine/threonine | Q13546;Q13546-2 | RIPK1 | Extracellular or secreted | -0.096867986 | 0.201071594 | 24.10381604 | 24.36666372 | 24.70197479 | 24.83465506 | 24.31271348 | 24.16583995 | 24.14771298 | 24.19865486 | 24.60585972 |
| Regulator of microtubule dynamics pr | Q96DB5;Q96DB5-2;Q96DB5- | RMDN1 | Extracellular or secreted | 0.085132343 | 0.117792411 | 27.38475688 | 27.49006582 | 27.33475106 | 27.03810534 | 27.10149357 | 26.44594336 | 27.48015932 | 27.37914777 | 26.79364796 |
| Regulator of microtubule dynamics pr | Q96TC7-2;Q96TC7 | RMDN3 | Other | -0.841091105 | 4.154241849 | 27.30903941 | 27.16806119 | 27.55946719 | 27.11465803 | 27.39214733 | 27.00217257 | 26.36258972 | 26.45862326 | 26.42828656 |
| RecQ-mediated genome instability pr | Q9H9A7 | RMI1 | Other | -0.115084074 | 0.071931797 | 21.97812948 | 22.45986482 | 22.24286823 | 20.92361771 | 22.49164762 | 22.44464629 | 21.94005546 | 22.84318346 | 21.14189593 |
| Protein RMD5 homolog A | Q9H871 | RMND5A | Other | -0.193481199 | 0.504923634 | 23.5414417 | 23.67476037 | 23.38112994 | 23.58164378 | 23.6896585 | 22.61827078 | 23.33688025 | 23.18398423 | 23.14214446 |
| Ribonuclease 4 | P34096 | RNASE4 | Extracellular or secreted | -0.755520895 | 0.567904437 | 27.31544563 | 27.08334622 | 26.95349241 | 26.91911683 | 27.15942514 | 28.57708291 | 26.14584883 | 26.01007167 | 27.58147139 |
| Ribonuclease H2 subunit A | O75792 | RNASEH2A | Other | 0.422342718 | 0.498280048 | 24.20784189 | 24.55613742 | 24.48111453 | 23.76759367 | 22.4014162 | 23.70998744 | 24.71670058 | 24.09071333 | 24.02165981 |
| Ribonuclease H2 subunit B | Q5TBB1-2;Q5TBB1 | RNASEH2B | Other | 0.40221588 | 1.03204316 | 23.25868104 | 22.83010587 | 22.72878911 | 22.61524257 | 23.86003365 | 23.17299693 | 23.46288764 | 23.34678639 | 23.62989819 |
| Ribonuclease H2 subunit C | Q8TDP1 | RNASEH2C | Other | 0.998799873 | 1.117196042 | 24.41319043 | 24.83407634 | 24.69253463 | 22.06990198 | 23.99005825 | 23.86580198 | 25.25684704 | 25.19004981 | 24.48228458 |
| Ribonuclease T2 | O00584 | RNASET2 | Extracellular or secreted | -0.183462989 | 0.299122599 | 27.93853975 | 27.9511037 | 27.98174373 | 28.18872066 | 28.31760348 | 28.06074853 | 27.65717341 | 27.6747873 | 28.33688025 |
| RING finger protein 113A | O15541 | RNF113A | Other | 1.934700741 | 1.876170745 | 22.50338479 | 22.4440785 | 21.29165325 | 23.89879124 | 21.13208704 | 24.26484917 | 24.4429101 | 24.29970831 | 24.82890582 |
| E3 ubiquitin-protein ligase RNF114 | Q9Y508;Q9Y508-2 | RNF114 | Other | 0.400294223 | 0.849317575 | 27.0232684 | 27.14918705 | 26.92886098 | 27.32783083 | 27.45002503 | 26.89006329 | 27.66444826 | 27.75193903 | 27.16911317 |
| E3 ubiquitin-protein ligase RNF14 | Q9UBS8 | RNF14 | Other | 1.100001008 | 0.530992387 | 20.34662477 | 21.34952664 | 21.2517036 | 21.09057318 | 23.06518675 | 21.28771814 | 23.19479833 | 23.36225528 | 20.93861595 |
| E3 ubiquitin-protein ligase RNF149 | Q8NC42 | RNF149 | Extracellular or secreted | 0.81430677 | 0.614080113 | 21.02905944 | 18.74015711 | 22.25482445 | 21.12418285 | 21.99829353 | 22.96862465 | 21.82791793 | 22.22999228 | 22.44258111 |
| E3 ubiquitin-protein ligase RING2 | Q99496;Q99496-2 | RNF2 | Other | -0.479004443 | 0.171360364 | 20.56959462 | 24.02106672 | 23.93472102 | 23.92088926 | 23.16622316 | 21.43009597 | 24.05622756 | 21.83462098 | 21.19343352 |
| E3 ubiquitin-protein ligase BRE1A | Q5VTR2 | RNF20 | Other | -0.023427041 | 0.030240045 | 24.97906333 | 25.34140073 | 25.17016438 | 24.97044367 | 25.63092586 | 24.18101391 | 24.88282997 | 25.37202251 | 24.81137234 |
| E3 ubiquitin-protein ligase RNF213 | Q63HN8;Q63HN8-4 | RNF213 | Other | 1.305651704 | 1.03239219 | 23.30187237 | 24.1418795 | 24.32594551 | 24.01750304 | 24.36192076 | 24.14693654 | 24.43198823 | 25.75018909 | 25.88280665 |
| RING finger protein 214 | Q8ND24 | RNF214 | Other | 1.584233234 | 0.903194143 | 21.20944175 | 21.58815646 | 24.39355543 | 24.5588252 | 24.76212814 | 24.3514445 | 24.56232349 | 26.25515481 | 24.36699715 |
| E3 ubiquitin-protein ligase RNF220 | Q5VTB9-3;Q5VTB9 | RNF220 | Extracellular or secreted | -0.250363741 | 0.13554136 | 21.27820291 | 22.63632989 | 21.52712598 | 23.50324538 | 23.59080814 | 22.79286892 | 21.19284034 | 22.85266209 | 22.86769696 |
| E3 ubiquitin-protein ligase RNF25 | Q96BH1 | RNF25 | Other | 0.289193322 | 0.163134529 | 24.55724819 | 24.51929359 | 24.53872131 | 24.04000939 | 24.27341203 | 22.95633149 | 25.03531797 | 25.02559346 | 23.24917654 |
| E3 ubiquitin-protein ligase BRE1B | O75150-4;O75150;O75150-3 | RNF40 | Other | 0.085335957 | 0.165499413 | 25.86105187 | 25.49859676 | 25.66979597 | 25.9883905 | 25.81689888 | 25.70725127 | 26.16200217 | 25.8010452 | 25.56395313 |
| RING-box protein 2 | Q9UBF6;Q9UBF6-4 | RNF7 | Other | 0.217474037 | 0.068014897 | 20.5987902 | 22.92582243 | 23.04914726 | 22.44184382 | 21.23872923 | 22.95960722 | 23.98988506 | 20.56465954 | 22.70484759 |
| mRNA-capping enzyme;Polynucleotid | O60942-3;O60942-4;O60942 | RNGTT | Other | -2.57569058 | 1.115385324 | 20.6945989 | 19.26718602 | 23.91661353 | 25.11147766 | 25.56935268 | 23.93606996 | 21.51399408 | 20.81591246 | 19.19067109 |
| Ribonuclease inhibitor | P13489 | RNH1 | Extracellular or secreted | 0.23814157 | 0.622756547 | 30.77878255 | 30.64718743 | 30.64013519 | 30.79428341 | 30.95004697 | 30.4112478 | 31.08853646 | 31.08594368 | 30.65078624 |
| mRNA cap guanine-N7 methyltransfer | O43148;O43148-2 | RNMT | Other | 0.318682462 | 0.715197576 | 26.77504999 | 26.92422372 | 26.83701575 | 26.8878576 | 26.96928024 | 26.51014505 | 27.35363334 | 27.25448818 | 26.79971205 |
| Aminopeptidase B | Q9H4A4 | RNPEP | Extracellular or secreted | -0.173524673 | 0.285226542 | 31.37710079 | 31.44623332 | 31.39631851 | 31.14613055 | 31.19946715 | 30.5025627 | 31.26919047 | 31.1320381 | 30.61210391 |
| RNA-binding protein with serine-rich ( | Q15287-3;Q15287-2;Q15287 | RNPS1 | Other | -1.599964863 | 0.370814892 | 25.63295128 | 25.69946168 | 25.50039114 | 25.98048042 | 26.23229634 | 27.59484202 | 21.50179659 | 25.0097087 | 27.00881157 |
| Roundabout homolog 1 | Q9Y6N7-6;Q9Y6N7-5;Q9Y6N7 | ROBO1 | Extracellular or secreted | -0.352518761 | 0.645709784 | 23.32882391 | 22.53471418 | 23.7177467 | 24.12904085 | 23.95695179 | 23.79233277 | 22.9882388 | 23.29235421 | 23.39165581 |
| Rho-associated protein kinase 1 | Q13464 | ROCK1 | Extracellular or secreted | -0.273130864 | 1.254638804 | 28.83026064 | 28.79426791 | 28.64687854 | 28.34832388 | 28.50646479 | 28.13975415 | 28.27386552 | 28.33402642 | 28.20569043 |
| Rho-associated protein kinase 2 | O75116 | ROCK2 | Extracellular or secreted | 0.004362601 | 0.01362988 | 27.61897925 | 27.49121333 | 27.48369968 | 27.46224801 | 27.79847655 | 27.20871646 | 27.5896372 | 27.5869204 | 27.36819684 |
| Tyrosine-protein kinase transmembra | Q01974 | ROR2 | Extracellular or secreted | 0.653688462 | 0.482427996 | 21.72434065 | 21.9477796 | 21.99982436 | 22.6215301 | 21.66031067 | 22.71845762 | 22.15875513 | 22.35646379 | 23.78196797 |
| Protein XRP2 | O75695 | RP2 | Extracellular or secreted | 0.38099633 | 1.000049384 | 26.10423609 | 26.25672109 | 25.93914533 | 25.24096307 | 25.57464549 | 25.90903168 | 26.00921748 | 26.19415206 | 26.45199082 |
| Replication protein A 70 kDa DNA-bin | P27694 | RPA1 | Other | -0.068485857 | 0.069003849 | 29.65742899 | 29.64955337 | 29.55780691 | 29.10317073 | 28.91367801 | 29.74840824 | 29.69761705 | 29.65739492 | 28.75455359 |
| Replication protein A 32 kDa subunit | P15927;P15927-2;P15927-3 | RPA2 | Other | -0.232503555 | 0.342825453 | 27.87929316 | 28.28224261 | 28.13148625 | 27.38352145 | 27.61883926 | 28.11083079 | 27.95825799 | 27.85067025 | 27.19666786 |
| Replication protein A 14 kDa subunit | P35244 | RPA3 | Other | 0.20078154 | 0.404775239 | 28.0781627 | 28.04478981 | 28.02448424 | 27.19309938 | 27.37409744 | 27.83887989 | 28.03002182 | 28.18158268 | 27.66749685 |
| RPA-interacting protein | Q86UA6;Q86UA6-8;Q86UA6- | RPAIN | Other | 0.114843548 | 0.034920619 | 23.61042923 | 23.17136374 | 23.83132426 | 23.82745281 | 21.9647938 | 22.43930567 | 24.07675689 | 24.46220119 | 21.22790731 |
| RNA polymerase II-associated protein | Q9BWH6-2;Q9BWH6 | RPAP1 | Other | 0.267636271 | 0.13714179 | 23.6516281 | 23.76151958 | 23.7163866 | 23.6686063 | 23.62867514 | 22.66014217 | 24.60783598 | 24.26549315 | 22.47305863 |
| RNA polymerase II-associated protein | Q9H6T3;Q9H6T3-2;Q9H6T3-3 | RPAP3 | Other | 0.537657441 | 1.69667685 | 26.15992552 | 26.27815322 | 26.03395376 | 26.88016942 | 26.87712946 | 26.80833106 | 27.02453708 | 26.92048043 | 27.18678602 |
| Ribulose-phosphate 3-epimerase | Q96AT9;Q96AT9-4;Q96AT9-2 | RPE | Extracellular or secreted | -0.149093211 | 0.161545539 | 28.2135913 | 28.36214657 | 28.2115041 | 28.14909012 | 28.28661127 | 27.28153532 | 28.25628918 | 28.21187538 | 27.33679514 |
| Ribosome production factor 2 homolo | Q9H7B2 | RPF2 | Other | -0.199072304 | 0.2321596 | 23.36639693 | 23.10771578 | 23.23277574 | 23.22051767 | 24.10237495 | 24.90370977 | 23.34583943 | 23.18850373 | 23.83518534 |
| Rabphilin-3A | Q9Y2J0-2;Q9Y2J0 | RPH3A | Other | -4.951591969 | 1.355999789 | 29.27399887 | 28.56292754 | 28.70908692 | 28.45454235 | 27.48914715 | 29.43485767 | 23.02779668 | 25.96666457 | 22.11304309 |
| Ribose-5-phosphate isomerase | P49247 | RPIA | Other | 0.077509701 | 0.092596844 | 27.84523591 | 28.16925656 | 28.07902792 | 28.06141825 | 28.20293822 | 27.53571404 | 28.3648619 | 28.29193285 | 27.52252981 |
| 60S ribosomal protein L10 | P27635 | RPL10 | Other | 1.331181283 | 1.737595715 | 27.60179991 | 27.58333784 | 27.80452042 | 28.21210738 | 27.96030346 | 29.33573088 | 29.82832479 | 29.56145335 | 28.85266566 |
| 60S ribosomal protein L10a | P62906 | RPL10A | Extracellular or secreted | 2.059177555 | 4.945306087 | 29.56743177 | 29.57719098 | 29.65913171 | 29.23256168 | 29.40859822 | 29.95698501 | 31.54609438 | 31.50497907 | 31.82740889 |
| 60S ribosomal protein L11 | P62913-2;P62913 | RPL11 | Extracellular or secreted | 0.70911697 | 1.444101353 | 27.16317378 | 27.09978938 | 27.2754205 | 27.77498718 | 27.90408892 | 28.70895541 | 28.31259232 | 28.28537703 | 28.49258915 |
| 60S ribosomal protein L12 | P30050;P30050-2 | RPL12 | Extracellular or secreted | 1.470668033 | 2.952957047 | 30.20489796 | 30.32130745 | 30.11644391 | 29.58838668 | 29.65466632 | 30.38339785 | 31.7156308 | 31.64108323 | 31.18984015 |
| 60S ribosomal protein L13 | P26373;P26373-2 | RPL13 | Other | 1.408293325 | 2.022254679 | 27.61519485 | 27.4104595 | 27.62860562 | 28.22149523 | 28.46824319 | 29.38933931 | 29.74832825 | 29.80325823 | 29.03996234 |
| 60S ribosomal protein L13a;Putative 6 | P40429;Q6NVV1 | RPL13A;RPL13AP | Other | 1.683967633 | 2.508158309 | 26.48193058 | 26.12780183 | 26.46514749 | 26.40820948 | 26.29743604 | 28.25277514 | 28.60367563 | 28.51912109 | 27.94575646 |
| 60S ribosomal protein L14 | P50914 | RPL14 | Extracellular or secreted | 1.337648754 | 1.638667268 | 26.30005757 | 27.28241937 | 27.2210342 | 27.23041014 | 27.16633811 | 28.9327797 | 29.22237078 | 28.62320742 | 28.23388762 |
| 60S ribosomal protein L15 | P61313;P61313-2 | RPL15 | Extracellular or secreted | 1.202305434 | 2.333761756 | 27.33116694 | 27.44031412 | 27.65123119 | 27.53727031 | 27.73000797 | 29.03294026 | 29.11354073 | 28.92719441 | 28.92764655 |
| 60S ribosomal protein L17 | P18621;P18621-3;P18621-2 | RPL17 | Other | 1.129353082 | 2.205861367 | 27.52424984 | 27.49944861 | 27.59149313 | 27.45513053 | 27.86531823 | 28.85931691 | 29.16314978 | 29.05006846 | 28.57231963 |
| 60S ribosomal protein L18 | Q07020;Q07020-2 | RPL18 | Other | 1.3291653 | 1.821568589 | 27.43809435 | 27.69677178 | 27.85554965 | 27.33866634 | 27.4326892 | 29.74695187 | 29.4703589 | 29.24013514 | 29.03136345 |
| 60S ribosomal protein L18a | Q02543 | RPL18A | Other | 1.476782816 | 2.160962993 | 25.69834922 | 25.97521929 | 26.20560654 | 26.4445697 | 26.31970643 | 28.05878912 | 28.03580038 | 27.84750769 | 27.89816054 |
| 60S ribosomal protein L19 | P84098 | RPL19 | Other | 0.672314021 | 1.438169072 | 28.22912685 | 27.7876828 | 27.89081709 | 28.22218649 | 28.3393037 | 29.1173608 | 29.16122859 | 29.03899491 | 28.60995744 |
| 60S ribosomal protein L21 | P46778 | RPL21 | Other | 1.742825008 | 1.831603337 | 27.33296011 | 27.73834372 | 27.98571159 | 27.08598165 | 27.37260379 | 29.06504476 | 29.96195983 | 29.75975267 | 28.79708534 |
| 60S ribosomal protein L22 | P35268 | RPL22 | Extracellular or secreted | 1.374446974 | 1.620529063 | 27.4826231 | 27.58240492 | 27.81659377 | 27.99112935 | 27.85869465 | 30.1054955 | 29.24592522 | 29.25788569 | 30.03800065 |
| 60S ribosomal protein L23 | P62829 | RPL23 | Extracellular or secreted | 0.804577604 | 1.595363605 | 27.15508766 | 27.26686959 | 27.18082427 | 27.8275376 | 27.87812399 | 28.79352393 | 28.56125304 | 28.54022999 | 28.36323331 |
| 60S ribosomal protein L23a | P62750 | RPL23A | Extracellular or secreted | 0.943290767 | 1.435128204 | 28.27208633 | 28.02194568 | 28.08009599 | 27.99215643 | 28.02511818 | 30.0713757 | 29.23911094 | 29.39231113 | 29.42983939 |
| 60S ribosomal protein L24 | P83731 | RPL24 | Extracellular or secreted | 1.084537445 | 2.003827403 | 27.12465033 | 27.31984422 | 27.42237934 | 27.33278943 | 27.0517546 | 28.87584139 | 28.56601663 | 28.43698318 | 28.81424218 |
| 60S ribosomal protein L26;60S riboso | P61254;Q9UNX3 | RPL26;RPL26L1 | Extracellular or secreted | 1.118289215 | 1.034236599 | 26.66722613 | 25.46979631 | 25.06291111 | 26.74931972 | 26.73453722 | 28.71911204 | 27.25178245 | 27.84888097 | 27.95565549 |
| 60S ribosomal protein L27 | P61353 | RPL27 | Extracellular or secreted | 1.32213389 | 2.275305149 | 27.86944257 | 27.93747377 | 28.03427918 | 27.91342122 | 27.94268398 | 29.67799891 | 29.69975289 | 29.47500645 | 29.47929215 |
| 60S ribosomal protein L27a | P46776 | RPL27A | Other | 1.35013426 | 2.037712175 | 27.17388525 | 26.94391376 | 27.20060172 | 27.20321834 | 27.30947316 | 28.91467619 | 29.04252161 | 29.03041654 | 28.35034884 |
| 60S ribosomal protein L28 | P46779-4;P46779-5;P46779;P | RPL28 | Extracellular or secreted | 1.878728263 | 1.769646165 | 27.84433817 | 26.90058088 | 27.69537848 | 27.64663825 | 25.28817019 | 29.36308707 | 29.53743695 | 29.48064086 | 28.9872035 |
| 60S ribosomal protein L29 | P47914 | RPL29 | Other | 0.715105404 | 1.282497231 | 26.90712998 | 26.92580885 | 26.86041261 | 26.83207315 | 27.09295243 | 28.19267622 | 28.07423727 | 28.03233602 | 27.44426953 |
| 60S ribosomal protein L3 | P39023 | RPL3 | Extracellular or secreted | 0.920456677 | 1.610529924 | 29.12895729 | 29.27050547 | 29.45448352 | 29.22151828 | 29.38974958 | 30.52579985 | 30.770221 | 30.46021802 | 30.026438 |
| 60S ribosomal protein L30 | P62888 | RPL30 | Extracellular or secreted | 1.023181528 | 2.736058316 | 28.19821458 | 28.36256464 | 28.42193813 | 28.87557778 | 28.86844203 | 29.42628418 | 29.82160479 | 29.56806631 | 29.75638415 |
| 60S ribosomal protein L31 | P62899;P62899-3;P62899-2 | RPL31 | Extracellular or secreted | 1.070213605 | 1.683176338 | 27.1969492 | 27.24127228 | 27.58892275 | 28.81154996 | 28.6241492 | 28.83707592 | 29.19891709 | 29.09186862 | 29.06981477 |
| 60S ribosomal protein L32 | P62910 | RPL32 | Other | 1.333003164 | 2.132304285 | 26.72286172 | 26.80415111 | 26.92694002 | 27.18650269 | 27.33432485 | 28.81219288 | 28.75779925 | 28.8003294 | 28.33436748 |
| 60S ribosomal protein L34 | P49207 | RPL34 | Extracellular or secreted | 1.134434766 | 1.911114992 | 26.24452411 | 25.95684104 | 26.00172172 | 26.23735674 | 26.14487696 | 27.91803639 | 27.60314502 | 27.69358511 | 27.35825265 |
| 60S ribosomal protein L35 | P42766 | RPL35 | Other | 1.115195605 | 1.480496949 | 26.37157394 | 26.25264883 | 26.48871823 | 26.5122417 | 26.10041114 | 28.61684299 | 27.96570723 | 27.77240942 | 27.77868857 |
| 60S ribosomal protein L35a | P18077 | RPL35A | Extracellular or secreted | 0.367253238 | 0.370554887 | 28.09893653 | 27.86985436 | 28.03883796 | 27.34747931 | 27.41288366 | 28.29043951 | 28.85355823 | 28.17887901 | 27.59853814 |
| 60S ribosomal protein L36 | Q9Y3U8 | RPL36 | Other | 1.707764812 | 1.802654508 | 25.85474153 | 26.05471741 | 26.08885486 | 26.69988525 | 26.79637376 | 28.27635269 | 28.76065701 | 28.61484394 | 27.63325623 |
| 60S ribosomal protein L36a-like;60S ri | Q969Q0;P83881 | RPL36AL;RPL36A | Other | 1.097430624 | 1.870438182 | 26.33183324 | 26.2590763 | 26.56409854 | 26.48709333 | 26.22280375 | 28.02738753 | 27.95082569 | 27.90351441 | 27.38409811 |
| 60S ribosomal protein L37a | P61513 | RPL37A | Extracellular or secreted | 1.145046216 | 1.456120051 | 25.20500986 | 25.32625419 | 25.4563845 | 25.38122898 | 25.44846621 | 27.76008591 | 27.11555125 | 26.80415111 | 26.80415111 |

| | | | | | | | | | | | | | | |
|---|---|---|---|---|---|---|---|---|---|---|---|---|---|---|
| 60S ribosomal protein L38 | P63173 | RPL38 | Other | 1.29167481 | 2.374807136 | 25.19708048 | 24.88115019 | 25.15421859 | 25.55920459 | 25.59032275 | 25.48802864 | 26.79897088 | 26.75837125 | 26.25268492 |
| Putative 60S ribosomal protein L39-lik | Q59GN2;P62891 | RPL39P5;RPL39 | Other | 1.105104747 | 1.736869448 | 24.130377 | 24.27632607 | 24.17500916 | 24.74688782 | 25.03590523 | 25.01814006 | 26.13422137 | 25.52546011 | 25.34695543 |
| 60S ribosomal protein L4 | P36578 | RPL4 | Extracellular or secreted | 1.570485708 | 2.232232654 | 27.92970765 | 27.88843837 | 28.07244945 | 28.04150367 | 28.03506621 | 29.95227077 | 30.23683234 | 29.98005544 | 29.4542874 |
| 60S ribosomal protein L5 | P46777 | RPL5 | Extracellular or secreted | 1.715155032 | 2.679645167 | 28.52301611 | 28.5993188 | 28.63322158 | 29.30838854 | 29.34709909 | 30.46645503 | 31.20034447 | 30.86311867 | 30.52075154 |
| 60S ribosomal protein L6 | Q02878 | RPL6 | Other | 0.327326738 | 0.626720282 | 29.82470322 | 30.0483024 | 30.0768639 | 29.25948044 | 29.20699473 | 30.59637137 | 30.37813498 | 30.18032634 | 29.92987692 |
| 60S ribosomal protein L7 | P18124 | RPL7 | Extracellular or secreted | 1.260698189 | 2.089746593 | 28.53923177 | 28.60378173 | 28.75238575 | 28.74400262 | 28.83866954 | 30.52393216 | 30.47547039 | 30.15834199 | 30.14928398 |
| 60S ribosomal protein L7a | P62424 | RPL7A | Extracellular or secreted | 1.151431242 | 2.069185873 | 28.32671708 | 28.34971634 | 28.5768307 | 28.62822319 | 28.65201806 | 30.17152504 | 30.11433534 | 30.00316501 | 29.68930858 |
| 60S ribosomal protein L7-like 1 | Q6DKI1 | RPL7L1 | Other | -0.547332829 | 0.347888294 | 22.90650678 | 21.19604168 | 21.87781284 | 21.71962779 | 22.00758031 | 25.03883796 | 21.76222954 | 21.33200416 | 22.63697149 |
| 60S ribosomal protein L8 | P62917 | RPL8 | Other | 1.46709186 | 2.259653263 | 28.31695646 | 28.17512341 | 28.42057354 | 28.39603264 | 28.43618896 | 30.16758276 | 30.33511325 | 30.32098481 | 29.70140641 |
| 60S ribosomal protein L9 | P32969 | RPL9 | Other | 1.06276713 | 1.551178612 | 26.94737386 | 26.77605466 | 27.03107419 | 27.43165356 | 27.38459222 | 29.12117078 | 28.36164472 | 28.52316571 | 28.64945059 |
| 60S acidic ribosomal protein P0;60S a | P05388;P05388-2;Q8NHW5 | RPLP0;RPLP0P6 | Extracellular or secreted | 0.911205071 | 3.54766009 | 30.85860574 | 31.04492006 | 31.00041401 | 31.17051064 | 31.13839979 | 31.08366324 | 32.02015808 | 32.0299955 | 31.83171838 |
| 60S acidic ribosomal protein P1 | P05386 | RPLP1 | Extracellular or secreted | 0.197185775 | 0.209105937 | 31.10599496 | 31.29032964 | 31.26963637 | 30.95740024 | 30.92174346 | 29.46423618 | 31.36895654 | 31.1822815 | 30.54498972 |
| 60S acidic ribosomal protein P2 | P05387 | RPLP2 | Extracellular or secreted | 0.172097978 | 0.353968609 | 32.38257357 | 32.44953316 | 32.52708249 | 31.82559051 | 31.89388622 | 31.39166605 | 32.23953778 | 32.44013092 | 32.07179123 |
| Dolichyl-diphosphooligosaccharide--p | P04843 | RPN1 | Other | -1.203824777 | 0.560816334 | 27.18517972 | 27.34375732 | 27.88460095 | 27.83719625 | 27.90466319 | 30.10998444 | 25.62761804 | 26.52273932 | 28.37085925 |
| Dolichyl-diphosphooligosaccharide--p | P04844-2;P04844 | RPN2 | Extracellular or secreted | -0.841096267 | 0.548465821 | 27.05196198 | 26.9480426 | 27.32817335 | 27.15141471 | 27.13016094 | 29.52660222 | 26.10273536 | 26.15690135 | 27.7852524 |
| Ribonuclease P protein subunit p25 | Q9BUL9 | RPP25 | Other | 0.73131753 | 1.95682589 | 23.49252803 | 23.24091941 | 23.30869232 | 23.87652069 | 23.1815751 | 24.38817762 | 24.18903185 | 24.29305622 | 24.45607111 |
| Ribonuclease P protein subunit p30 | P78346;P78346-2 | RPP30 | Other | 0.630337758 | 2.33143195 | 26.68404957 | 26.92082113 | 26.30610384 | 26.5584603 | 26.36203785 | 26.92331715 | 27.44655871 | 27.12218343 | 27.19966606 |
| Ribonuclease P protein subunit p40 | O75818-2;O75818 | RPP40 | Other | 0.615785331 | 2.71767012 | 24.52766885 | 24.84603742 | 24.80582455 | 24.87164133 | 24.57605941 | 25.04318419 | 25.41857248 | 25.24448782 | 25.51950356 |
| Regulation of nuclear pre-mRNA dom | Q96P16;Q96P16-3 | RPRD1A | Other | -0.105015637 | 0.111387229 | 26.26733407 | 26.16088731 | 25.98128645 | 25.86499957 | 25.75169646 | 25.47243279 | 26.08642717 | 26.15098881 | 25.19685543 |
| Regulation of nuclear pre-mRNA dom | Q9NQG5 | RPRD1B | Other | -0.432871275 | 1.468700236 | 29.59441493 | 29.81537264 | 29.79632734 | 29.43603007 | 29.36041026 | 28.94013725 | 29.18146421 | 29.12220812 | 28.8690601 |
| Regulation of nuclear pre-mRNA dom | Q5VT52-5;Q5VT52-3;Q5VT52 | RPRD2 | Other | -3.251730288 | 1.40084226 | 24.50476129 | 23.49289471 | 24.27717784 | 22.6376144 | 22.03895437 | 27.36202112 | 19.10794196 | 22.12738859 | 21.16619045 |
| 40S ribosomal protein S10;Putative 40 | P46783;Q9NQ39 | RPS10;RPS10P5 | Extracellular or secreted | 1.265181554 | 1.981011846 | 26.03881704 | 26.37908165 | 26.46424398 | 26.15855869 | 25.77326499 | 27.57852322 | 27.47861732 | 27.3831094 | 28.13006272 |
| 40S ribosomal protein S11 | P62280 | RPS11 | Extracellular or secreted | 0.97050329 | 2.134674152 | 28.9704327 | 28.74233298 | 28.9428797 | 28.87108902 | 28.84750769 | 30.09043068 | 30.25236912 | 30.14758683 | 29.7438903 |
| 40S ribosomal protein S12 | P25398 | RPS12 | Other | 0.556668201 | 2.991931882 | 30.88467372 | 30.71366584 | 30.80678036 | 31.05219526 | 31.10886352 | 30.54397636 | 31.37134752 | 31.40932684 | 31.44440778 |
| 40S ribosomal protein S13 | P62277 | RPS13 | Extracellular or secreted | 1.268418168 | 2.280753076 | 27.91245073 | 28.22356803 | 28.15634205 | 27.8522787 | 27.98928959 | 29.34625381 | 29.82022051 | 29.588619 | 29.13650645 |
| 40S ribosomal protein S14 | P62263 | RPS14 | Extracellular or secreted | 0.58223312 | 1.226887072 | 28.20041463 | 27.86076779 | 27.67929163 | 27.79618808 | 27.95510117 | 28.99275072 | 28.57416185 | 28.97983746 | 28.43495706 |
| 40S ribosomal protein S15 | P62841 | RPS15 | Other | 1.334541126 | 2.165591958 | 26.91991242 | 26.83653429 | 26.90322707 | 27.01051991 | 26.34319819 | 28.54093202 | 28.38409811 | 28.37587949 | 28.52080773 |
| 40S ribosomal protein S15a | P62244 | RPS15A | Extracellular or secreted | 1.226827156 | 2.315183442 | 27.49205426 | 27.84559485 | 28.02104553 | 27.72442392 | 27.61862926 | 29.10307066 | 29.41884588 | 29.32519499 | 28.83884984 |
| 40S ribosomal protein S16 | P62249 | RPS16 | Extracellular or secreted | 1.189935315 | 2.369752732 | 27.93432734 | 28.4034876 | 28.33964351 | 28.6907577 | 28.65679848 | 29.72300825 | 29.90956394 | 29.75608187 | 29.77817158 |
| 40S ribosomal protein S17 | P08708 | RPS17 | Extracellular or secreted | 1.283565313 | 2.284698924 | 27.77102433 | 27.61750872 | 27.81353902 | 27.74271845 | 27.47961981 | 29.31155335 | 29.29476155 | 29.26948032 | 29.15443591 |
| 40S ribosomal protein S18 | P62269 | RPS18 | Extracellular or secreted | 1.138244333 | 2.056116543 | 28.41465879 | 28.67909024 | 28.76960639 | 28.97065212 | 29.04954926 | 30.32195252 | 30.16710418 | 30.18399748 | 30.166386 |
| 40S ribosomal protein S19 | P39019 | RPS19 | Extracellular or secreted | 1.197103184 | 1.877019376 | 27.29079102 | 27.25520885 | 27.14287286 | 26.83134842 | 27.44015568 | 29.08866282 | 28.46804893 | 28.80974213 | 28.83803832 |
| 40S ribosomal protein S2 | P15880 | RPS2 | Extracellular or secreted | 0.811388018 | 2.137897323 | 29.45352228 | 29.44931668 | 29.45199082 | 29.30391129 | 29.34718359 | 30.53825137 | 30.57983626 | 30.38071721 | 30.2456986 |
| 40S ribosomal protein S20 | P60866;P60866-2 | RPS20 | Extracellular or secreted | 0.992659142 | 1.86833462 | 27.9434108 | 28.24377085 | 28.28718395 | 27.66905254 | 27.91302169 | 29.51312696 | 29.23210417 | 29.22172569 | 29.30893095 |
| 40S ribosomal protein S21 | P63220 | RPS21 | Other | 0.868688729 | 1.106585606 | 29.08669037 | 28.82462736 | 28.81259073 | 29.44166018 | 29.50884632 | 28.97059727 | 30.27164119 | 30.26326931 | 29.3936618 |
| 40S ribosomal protein S23 | P62266 | RPS23 | Other | 1.230957096 | 1.806864486 | 25.83059915 | 26.20176109 | 26.90035054 | 26.10015044 | 25.96470521 | 27.12129431 | 28.067842 | 27.49975272 | 27.18470694 |
| 40S ribosomal protein S24 | P62847-2;P62847-3;P62847;F | RPS24 | Other | 0.388267408 | 0.316715767 | 26.81763091 | 26.99851881 | 27.00859788 | 26.62644876 | 26.79352393 | 27.49579429 | 27.98277751 | 27.57520834 | 26.47707367 |
| 40S ribosomal protein S25 | P62851 | RPS25 | Extracellular or secreted | 1.105768208 | 2.807379996 | 27.93719312 | 27.77875123 | 27.69351865 | 28.04082464 | 27.84439804 | 28.95662506 | 29.24138595 | 29.1519469 | 29.04962715 |
| 40S ribosomal protein S26;Putative 40 | P62854;Q5JNZ5 | RPS26;RPS26P11 | Extracellular or secreted | 0.895145386 | 0.974940052 | 27.13133905 | 27.10928708 | 27.11107962 | 26.94681634 | 26.9138777 | 28.04603971 | 28.5030937 | 28.40685615 | 27.40470606 |
| 40S ribosomal protein S27;40S riboso | P42677;Q71UM5 | RPS27;RPS27L | Other | 1.372757452 | 2.071298759 | 27.29473965 | 27.14228861 | 27.23086818 | 27.90109899 | 27.94754108 | 29.25171474 | 29.37349602 | 29.35338095 | 28.77552102 |
| Ubiquitin-40S ribosomal protein S27a | P62979 | RPS27A | Extracellular or secreted | 0.288905725 | 0.418174868 | 32.25013962 | 32.25417277 | 32.27733417 | 32.66362997 | 32.61003667 | 32.32418116 | 33.18362782 | 32.55880604 | 32.31403049 |
| 40S ribosomal protein S28 | P62857 | RPS28 | Extracellular or secreted | 0.83483111 | 1.021774698 | 28.5978636 | 28.30942979 | 28.45485607 | 29.09174254 | 28.98544016 | 28.13858289 | 29.49499367 | 29.9362104 | 28.86224678 |
| 40S ribosomal protein S29 | P62273;P62273-2 | RPS29 | Extracellular or secreted | 1.104197082 | 1.124535666 | 24.78780733 | 25.86648603 | 26.12957153 | 25.45688578 | 25.73990213 | 28.29928908 | 26.85993889 | 27.20359176 | 27.38903153 |
| 40S ribosomal protein S3 | P23396;P23396-2 | RPS3 | Extracellular or secreted | 1.104789099 | 2.25666471 | 29.77588203 | 29.88564853 | 29.90037934 | 30.33159409 | 30.50559437 | 31.33122034 | 31.43663576 | 31.39304802 | 31.34984286 |
| 40S ribosomal protein S3a | P61247 | RPS3A | Extracellular or secreted | 1.174010852 | 2.345260916 | 29.55784342 | 29.44383497 | 29.58645517 | 29.71010571 | 29.80080767 | 31.03872024 | 31.19301711 | 31.17134608 | 30.72655295 |
| 40S ribosomal protein S4, X isoform;4 | P62701;Q8TD47 | RPS4X;RPS4Y2 | Extracellular or secreted | 1.060232452 | 1.891323558 | 28.81142747 | 29.03351799 | 29.14559621 | 29.14816894 | 29.14729569 | 30.23054757 | 30.66593949 | 30.33766726 | 29.93536754 |
| 40S ribosomal protein S5;40S ribosom | P46782 | RPS5 | Extracellular or secreted | 1.178629925 | 2.313255264 | 28.0089184 | 28.00720816 | 28.07958748 | 28.36723883 | 28.25318104 | 29.46573525 | 29.76146569 | 29.66707382 | 29.19828485 |
| 40S ribosomal protein S6 | P62753 | RPS6 | Other | 1.34259793 | 1.258773302 | 26.66709075 | 26.8046435 | 26.65485407 | 27.371441 | 26.87619279 | 28.56325494 | 29.02477483 | 28.78596948 | 27.685788 |
| Ribosomal protein S6 kinase alpha-3 | P51812 | RPS6KA3 | Other | -0.294935733 | 0.64472873 | 26.33097895 | 26.42969176 | 26.50595774 | 26.11547187 | 26.09986963 | 26.89017928 | 26.35171407 | 26.17169205 | 25.77786129 |
| 40S ribosomal protein S7 | P62081 | RPS7 | Extracellular or secreted | 1.158349192 | 2.262275879 | 28.75512688 | 28.51837084 | 28.66393954 | 28.46801007 | 28.48393027 | 30.10474598 | 30.09005203 | 30.14065634 | 29.74140099 |
| 40S ribosomal protein S8 | P62241 | RPS8 | Extracellular or secreted | 1.143007242 | 1.864292194 | 28.84155163 | 28.82089 | 28.96653256 | 29.262239 | 29.39294569 | 30.4016173 | 30.71112379 | 30.58441354 | 29.97637249 |
| 40S ribosomal protein S9 | P46781 | RPS9 | Extracellular or secreted | 1.32948458 | 2.56788969 | 28.52062042 | 28.70285994 | 28.81311084 | 28.69062451 | 28.85090865 | 30.20466479 | 30.55032112 | 30.29362197 | 30.03590523 |
| 40S ribosomal protein SA | P08865 | RPSA | Extracellular or secreted | 0.940728037 | 4.350000546 | 30.26091645 | 30.33521975 | 30.24308985 | 30.44628262 | 30.65906364 | 30.75342226 | 31.43559301 | 31.46138952 | 31.27419887 |
| RNA pseudouridylate synthase domai | Q8IZ73;Q8IZ73-2 | RPUSD2 | Other | 0.213682018 | 0.266045226 | 27.20899547 | 27.46497618 | 27.36394343 | 27.18470694 | 27.20050818 | 27.15923264 | 27.90426122 | 27.60907681 | 26.91888944 |
| Cell differentiation protein RCD1 hom | Q92600;Q92600-2;Q92600-3 | RQCD1 | Extracellular or secreted | -0.275311329 | 0.594876751 | 26.03747708 | 25.92045772 | 26.04936749 | 25.27973016 | 24.82977692 | 26.05335072 | 25.4875074 | 25.34309651 | 25.42854215 |
| Ras-related GTP-binding protein A;Ra | Q7L523;Q5VZM2-2;Q5VZM2 | RRAGA;RRAGB | Other | 1.189815011 | 0.788627213 | 22.56978145 | 23.51207589 | 23.3937518 | 22.66858466 | 24.03296652 | 24.35890656 | 23.61694812 | 24.59263405 | 25.6278963 |
| Ras-related GTP-binding protein C | Q9HB90 | RRAGC | Other | 2.134769285 | 3.054786624 | 23.93246997 | 24.15429586 | 23.67841875 | 24.03263029 | 23.44960006 | 24.21170831 | 26.41425554 | 25.822769 | 25.89684494 |
| Ras-related protein R-Ras | P10301 | RRAS | Extracellular or secreted | 0.859608032 | 2.391024748 | 24.86240051 | 24.9225688 | 25.17926827 | 24.90269822 | 24.77841286 | 25.11957379 | 25.8522668 | 25.99036127 | 25.61865726 |
| Ras-related protein R-Ras2 | P62070;P62070-4;P62070-3;F | RRAS2 | Extracellular or secreted | 0.707454496 | 1.920470468 | 26.64223727 | 26.69842871 | 26.89110691 | 26.65075201 | 26.78706001 | 26.73414956 | 27.26936885 | 27.42862201 | 27.62623986 |
| Ribosome-binding protein 1 | Q9P2E9;Q9P2E9-3 | RRBP1 | Other | -1.354618057 | 2.547408635 | 30.7250906 | 30.64443944 | 30.67581377 | 30.36515401 | 30.54600236 | 30.09005203 | 29.31574792 | 29.30873571 | 28.8349383 |
| Ribonucleoside-diphosphate reductas | P23921 | RRM1 | Extracellular or secreted | -0.150652672 | 1.223981558 | 28.57632616 | 28.56946133 | 28.57520834 | 28.78066086 | 28.64079027 | 28.73792485 | 28.52163159 | 28.56241812 | 28.40417818 |
| Ribonucleoside-diphosphate reductas | P31350;P31350-2 | RRM2 | Other | 0.22512522 | 0.520823324 | 27.08172202 | 26.91649964 | 26.97147462 | 26.45834165 | 26.02320494 | 26.36359259 | 26.81066165 | 27.06814962 | 26.70398212 |
| Ribonucleoside-diphosphate reductas | Q7LG56;Q7LG56-6;Q7LG56-2 | RRM2B | Extracellular or secreted | 0.520298379 | 0.616637553 | 26.34883039 | 26.54241625 | 26.33296011 | 25.49054023 | 26.22870495 | 24.01256726 | 26.33403495 | 26.33372794 | 26.37114184 |
| RNA polymerase I-specific transcriptio | Q9NYV6-2;Q9NYV6-3;Q9NYV | RRN3 | Other | 0.482424219 | 0.373508381 | 19.09546209 | 21.88725335 | 22.26759482 | 22.21871101 | 21.94788041 | 21.62395113 | 22.47305863 | 21.9990352 | 21.49560523 |
| Ribosomal RNA processing protein 1 h | P56182 | RRP1 | Other | -0.757328229 | 0.382055486 | 21.04370421 | 19.33695205 | 21.8273365 | 22.5980069 | 23.59491549 | 24.8488571 | 20.78649788 | 22.03489835 | 21.53150521 |
| RRP12-like protein | Q5JTH9-2;Q5JTH9-3;Q5JTH9 | RRP12 | Other | 0.838760953 | 0.52949321 | 22.14840786 | 20.78883945 | 21.48123774 | 22.69791205 | 20.63218584 | 25.40204036 | 23.03101527 | 23.01379427 | 23.04678497 |
| RRP15-like protein | Q9Y3B9 | RRP15 | Other | 0.66016578 | 1.005959916 | 25.00598837 | 23.74540106 | 23.60127569 | 25.02106672 | 24.23629863 | 25.55098158 | 25.24890882 | 25.33796918 | 24.97412536 |
| U3 small nucleolar RNA-interacting pr | O43818 | RRP9 | Other | -0.622439979 | 0.984097824 | 26.92479004 | 26.54197335 | 26.77001616 | 26.71554898 | 26.59789911 | 27.34621153 | 26.06467995 | 25.85414703 | 26.66207267 |
| Ribosome biogenesis regulatory prote | Q15050 | RRS1 | Other | -0.367675487 | 0.371129183 | 23.55555245 | 23.54262288 | 23.36452798 | 24.62689431 | 24.51068836 | 25.62061608 | 24.12636481 | 23.34624535 | 24.03481441 |
| Remodeling and spacing factor 1 | Q96T23-2;Q96T23;Q96T23-3 | RSF1 | Extracellular or secreted | -0.303032133 | 1.089180595 | 28.94717876 | 28.7831927 | 28.74239723 | 29.01786405 | 28.8119786 | 28.66566847 | 28.62348653 | 28.63792644 | 28.31363054 |
| Ribosomal L1 domain-containing prot | O76021 | RSL1D1 | Other | -0.172845601 | 0.205097028 | 26.99959441 | 27.22609742 | 27.49823153 | 28.27346539 | 28.30504278 | 29.09549493 | 27.8048281 | 27.79470172 | 27.58089661 |
| RING finger and SPRY domain-contain | Q96DX4 | RSPRY1 | Extracellular or secreted | -0.413686309 | 0.422897737 | 22.95303053 | 22.57223577 | 22.75831025 | 22.25535227 | 21.69042359 | 21.74429374 | 21.27375838 | 21.99012751 | 22.48187825 |
| Ras suppressor protein 1 | Q15404;Q15404-2 | RSU1 | Extracellular or secreted | 0.502270073 | 1.405287023 | 28.8485527 | 28.85043181 | 28.77127627 | 28.02469558 | 28.16226155 | 28.23434456 | 29.10231988 | 29.10739254 | 28.74287903 |
| RNA 3-terminal phosphate cyclase | O00442;O00442-2 | RTCA | Other | 0.889447401 | 1.501500802 | 23.71229861 | 23.82919624 | 23.96578428 | 23.49789666 | 22.00667977 | 23.85823223 | 24.53100539 | 24.15977158 | 24.41260913 |
| tRNA-splicing ligase RtcB homolog | Q9Y3I0 | RTCB | Other | -0.223337937 | 0.745687694 | 29.61812162 | 29.47134776 | 29.55362039 | 29.25750359 | 29.46608546 | 29.70330562 | 29.42582416 | 29.37923041 | 29.05992384 |
| RNA polymerase-associated protein R | Q92541 | RTF1 | Extracellular or secreted | -0.659571909 | 1.421808052 | 28.42065385 | 28.43185278 | 28.36156106 | 28.47452303 | 28.46629943 | 27.829535 | 27.88547398 | 27.80002076 | 27.3280021 |
| Protein RTF2 homolog | Q9BY42 | RTFDC1 | Other | 0.784373108 | 2.386142424 | 26.06878516 | 26.33526236 | 26.58175869 | 27.05878912 | 26.79872373 | 27.09526856 | 27.45755911 | 27.31138011 | 27.55347391 |
| Rhotekin | Q9BST9;Q9BST9-3;Q9BST9-2 | RTKN | Other | -0.366858618 | 0.275689629 | 21.99464738 | 20.870461 | 22.29552457 | 23.46948582 | 23.41415874 | 22.66338177 | 22.67478191 | 21.37347693 | 22.20499494 |
| Reticulon-3 | O95197-3;O95197-6;O95197 | RTN3 | Extracellular or secreted | -0.680148852 | 0.707038343 | 27.68445094 | 27.80630407 | 27.64629491 | 26.54220958 | 26.61820916 | 28.64416435 | 26.54071775 | 26.41977827 | 27.46987392 |
| Reticulon-4 | Q9NQC3-2;Q9NQC3;Q9NQC3 | RTN4 | Extracellular or secreted | -0.280638765 | 0.229069472 | 29.18243543 | 29.08894077 | 29.1325162 | 28.34583098 | 28.60109146 | 30.02986389 | 28.56019637 | 28.2115041 | 29.5767226 |
| Reticulon-4 receptor | Q9BZR6 | RTN4R | Extracellular or secreted | 1.493974569 | 0.770322286 | 21.71687013 | 21.83286864 | 23.48566621 | 23.56894696 | 23.98953864 | 26.24163598 | 24.31008019 | 24.28277285 | 26.30683395 |
| Reticulon-4 receptor-like 2 | Q86UN3;Q86UN3-2 | RTN4RL2 | Extracellular or secreted | -1.269310196 | 0.601565282 | 24.01554777 | 24.45619647 | 21.8177895 | 24.90733631 | 24.53278965 | 21.77107225 | 21.84792953 | 21.28637123 | 23.80813463 |
| RUN and FYVE domain-containing pro | Q96T51;Q96T51-2 | RUFY1 | Other | 0.13374671 | 0.29103393 | 26.77278694 | 26.80783993 | 26.65553662 | 26.78256796 | 26.82092042 | 26.81408935 | 27.11674134 | 27.03580038 | 26.57556902 |
| RUN and FYVE domain-containing pro | Q8WXA3;Q8WXA3-3;Q8WXA | RUFY2 | Other | -0.277597952 | 0.337052649 | 20.20975373 | 21.12459509 | 21.36284414 | 21.76871838 | 22.71571463 | 21.63820963 | 21.05377743 | 21.1318218 | 21.39152471 |
| RuvB-like 1 | Q9Y265;Q9Y265-2 | RUVBL1 | Extracellular or secreted | 0.219299449 | 0.520024757 | 30.98318538 | 30.9510203 | 30.91299314 | 30.97828334 | 30.92648766 | 30.53547304 | 31.26198131 | 31.25749235 | 30.78214612 |
| RuvB-like 2 | Q9Y230;Q9Y230-2 | RUVBL2 | Extracellular or secreted | 0.2362032 | 0.703553202 | 31.01531381 | 30.83996119 | 30.80131662 | 30.87078045 | 30.94013725 | 30.84161161 | 31.28206582 | 31.21025033 | 30.87085393 |
| RWD domain-containing protein 1 | Q9H446;Q9H446-2 | RWDD1 | Other | 0.542552843 | 1.091105537 | 25.74255143 | 25.79956384 | 25.81972732 | 25.62878637 | 25.86563681 | 24.71014513 | 26.48670984 | 26.04936749 | 25.87478666 |
| RWD domain-containing protein 4 | Q6NW29;Q6NW29-2 | RWDD4 | Other | 0.973803793 | 1.60763756 | 23.33742482 | 23.12664839 | 22.85134721 | 21.87883743 | 23.55086419 | 22.43721089 | 23.99774389 | 24.10701333 | 23.40782063 |
| Retinoic acid receptor RXR-alpha | P19793-2;P19793 | RXRA | Other | -0.248171148 | 0.089338943 | 21.05992797 | 21.28737421 | 21.45035547 | 22.32381886 | 23.00351759 | 22.27250108 | 21.4676502 | 20.19402223 | 23.29256171 |
| RING1 and YY1-binding protein | Q8N488 | RYBP | Other | -0.387631955 | 0.203652247 | 22.05602849 | 22.14371692 | 20.7601733 | 23.73665462 | 24.00324544 | 23.24895947 | 21.93617782 | 23.35460882 | 21.52070661 |
| Protein S100-A10 | P60903 | S100A10 | Extracellular or secreted | 0.722102102 | 0.714865204 | 27.57029408 | 27.40169866 | 27.57000448 | 28.50703218 | 28.47235531 | 28.46135048 | 28.49925851 | 28.18749412 | 29.47092128 |
| Protein S100-A11;Protein S100-A11, N | P31949 | S100A11 | Extracellular or secreted | 0.098218998 | 0.216245402 | 33.02083696 | 32.84316653 | 33.12827826 | 33.39151242 | 33.51406342 | 32.82549574 | 33.3985372 | 32.95023821 | 33.30755826 |
| Protein S100-A13 | Q99584 | S100A13 | Extracellular or secreted | 1.198995259 | 1.829784168 | 24.73918109 | 24.41828294 | 24.11036287 | 25.03468849 | 24.85576349 | 24.52790743 | 25.81685007 | 25.41512641 | 26.20810246 |
| Protein S100-A14 | Q9HCY8 | S100A14 | Extracellular or secreted | 1.698160919 | 1.86295195 | 28.47576027 | 27.94307539 | 28.52193106 | 28.48193058 | 28.28956035 | 29.18288528 | 30.06496769 | 29.71912837 | 30.75795816 |
| Protein S100-A16 | Q96FQ6 | S100A16 | Extracellular or secreted | 1.947543148 | 2.427371066 | 29.2236831 | 29.2425221 | 29.47487111 | 29.06141825 | 29.10009027 | 30.30432482 | 30.96567972 | 31.21569862 | 31.86470594 |
| Protein S100-A2 | P29034 | S100A2 | Other | 1.344029373 | 2.139508766 | 28.13481868 | 28.4107021 | 28.68962518 | 28.06157275 | 28.7348279 | 27.84128168 | 30.04113808 | 29.63813366 | 29.28923053 |
| Protein S100-A3 | P33764 | S100A3 | Other | 2.569669363 | 2.425429579 | 22.59439353 | 22.99937726 | 20.25501688 | 21.16684839 | 22.17613892 | 22.06368569 | 23.82220981 | 24.36833007 | 25.14619854 |
| Protein S100-A4 | P26447 | S100A4 | Extracellular or secreted | 1.005135364 | 4.284518666 | 27.66519407 | 27.7881186 | 27.91062215 | 27.7050375 | 27.73776372 | 27.25430795 | 28.73343857 | 28.63982479 | 28.67266472 |
| Protein S100-A6 | P06703 | S100A6 | Extracellular or secreted | 0.446399281 | 1.069479895 | 32.49008495 | 32.07076816 | 32.59845828 | 32.53843641 | 32.80501268 | 33.29021977 | 33.35757118 | 33.00540888 | 32.8727079 |
| Protein S100-A7;Protein S100-A7A | P31151;Q86SG5 | S100A7;S100A7A | Extracellular or secreted | -1.761194555 | 1.451043661 | 22.50784883 | 20.68292158 | 23.41118718 | 25.36769709 | 22.77223321 | 22.69742152 | 21.01685724 | 21.37561239 | 21.04360141 |
| Protein S100-A8;Protein S100-A8, N-t | P05109 | S100A8 | Extracellular or secreted | -2.277535219 | 1.692347273 | 25.20869786 | 24.911731 | 25.17016438 | 25.0274508 | 25.19681791 | 25.27134733 | 22.60498931 | 22.41619654 | 23.53931314 |
| Protein S100-A9 | P06702 | S100A9 | Extracellular or secreted | -1.683218014 | 3.181867194 | 27.21900392 | 25.58923715 | 26.10409609 | 25.94254976 | 26.13018059 | 26.56083775 | 24.5017886 | 24.62153859 | 24.59997139 |
| Protein S100-P | P25815 | S100P | Extracellular or secreted | 1.51889089 | 1.011505404 | 24.2472797 | 23.65348801 | 24.1655333 | 25.75409479 | 25.92091197 | 23.71156364 | 25.76233094 | 25.29974324 | 27.2210342 |
| Protein SAAL1 | Q96ER3 | SAAL1 | Extracellular or secreted | 0.373974003 | 0.952564774 | 26.11888105 | 26.14466307 | 26.16894108 | 26.41464266 | 26.12403894 | 25.81753333 | 26.6416862 | 26.67047148 | 26.20411438 |
| Phosphatidylinositide phosphatase SA | Q9NTJ5;Q9NTJ5-2 | SACM1L | Extracellular or secreted | -0.62761358 | 0.322581974 | 24.03582135 | 24.12966978 | 24.38883452 | 23.6793856 | 22.44894506 | 26.33050033 | 22.28743058 | 24.15491389 | 24.18139312 |
| SUMO-activating enzyme subunit 1;SU | Q9UBE0;Q9UBE0-2;Q9UBE0- | SAE1 | Other | 0.193331227 | 0.382622636 | 30.23534932 | 30.27974344 | 30.26091645 | 30.54655441 | 30.49581335 | 30.19565925 | 30.7369094 | 30.69585965 | 30.15424274 |
| Scaffold attachment factor B1 | Q15424;Q15424-4;Q15424-3 | SAFB | Other | -0.302854383 | 0.527348166 | 30.23797209 | 30.21222336 | 30.15713754 | 29.89994737 | 30.02141625 | 28.75327878 | 29.32229643 | 29.62022103 | 29.78990708 |
| Scaffold attachment factor B2 | Q14151 | SAFB2 | Extracellular or secreted | -1.150624133 | 1.994742151 | 26.8854158 | 27.32860139 | 27.23644462 | 26.6007371 | 26.93943681 | 26.13475014 | 25.3086576 | 25.7517986 | 26.05036433 |
| Protein Smaug homolog 2 | Q5PRF9 | SAMD4B | Other | 0.656596345 | 0.707779712 | 22.72643731 | 21.37888879 | 22.18639351 | 21.1044144 | 23.56050562 | 20.83321447 | 22.4093754 | 22.87096442 | 22.58437626 |
| Deoxynucleoside triphosphate tripho | Q9Y3Z3-4;Q9Y3Z3;Q9Y3Z3-3 | SAMHD1 | Extracellular or secreted | -0.217023922 | 0.144192132 | 25.43787218 | 25.22890674 | 25.0398421 | 25.81401599 | 25.81186227 | 26.64567668 | 25.93346087 | 25.94272872 | 24.46182663 |
| Histone deacetylase complex subunit | Q9H0E3;Q9H0E3-3;Q9H0E3-2 | SAP130 | Other | -0.759403632 | 0.217698878 | 24.14615969 | 24.34000871 | 23.92333982 | 21.59105621 | 21.64019345 | 23.02639579 | 23.93030566 | 19.98477157 | 23.1402887 |
| Histone deacetylase complex subunit | O00422 | SAP18 | Other | 0.208838168 | 0.455029061 | 26.68645607 | 26.29997026 | 26.46106159 | 26.87701241 | 26.63561052 | 26.84445791 | 26.92161579 | 27.08790451 | 26.51927859 |
| SAP30-binding protein | Q9UHR5;Q9UHR5-2 | SAP30BP | Other | 0.342972001 | 0.788329201 | 26.14298968 | 26.17081384 | 25.99748549 | 26.41610954 | 26.55057066 | 26.13825086 | 26.70977715 | 26.24668189 | 26.780567 |
| GTP-binding protein SAR1a | Q9NR31;Q9NR31-2 | SAR1A | Extracellular or secreted | -0.636926792 | 0.309983127 | 25.21044543 | 25.16518825 | 25.30305773 | 25.60631168 | 25.450324 | 26.97475994 | 23.96437471 | 24.5501009 | 26.42978752 |
| SAP domain-containing ribonucleopro | P82979 | SARNP | Other | 0.328931853 | 0.538196789 | 28.27967703 | 28.26901207 | 28.23022688 | 29.08765165 | 29.0370057 | 28.23986209 | 28.6574801 | 28.61792902 | 29.28310415 |
| Serine--tRNA ligase, cytoplasmic | P49591 | SARS | Extracellular or secreted | 0.309694969 | 0.996586194 | 28.97448645 | 28.79451582 | 28.77548962 | 28.958175 | 29.01881923 | 28.81274372 | 29.30534726 | 29.32322031 | 28.96763226 |
| Serine--tRNA ligase, mitochondrial | Q9NP81;Q9NP81-2 | SARS2 | Other | -0.227383423 | 0.063857725 | 20.64684679 | 23.67454488 | 21.95773172 | 24.23870561 | 24.03363875 | 24.79272994 | 24.18971057 | 20.91224101 | 23.887997 |
| U4/U6.U5 tri-snRNP-associated prote | O43290 | SART1 | Other | 0.085892361 | 0.073006082 | 25.5411758 | 25.79810569 | 25.75832042 | 25.84112568 | 25.91877573 | 26.46123337 | 25.7163866 | 25.4875994 | 26.71305943 |
| Squamous cell carcinoma antigen rec | Q15020;Q15020-4 | SART3 | Other | -0.40169586 | 0.804864807 | 28.31557519 | 28.51273148 | 28.37085925 | 27.84014133 | 27.84619288 | 28.01844785 | 28.01520745 | 27.85459887 | 27.3770801 |
| Ribosome maturation protein SBDS | Q9Y3A5 | SBDS | Other | 0.310053648 | 0.692513505 | 27.09607329 | 27.00088406 | 27.06291111 | 27.30138399 | 27.51352232 | 26.46642391 | 27.46295003 | 27.60801935 | 27.07979091 |
| Myotubularin-related protein 5 | O95248;O95248-5;O95248-4 | SBF1 | Other | 0.000121285 | 0.000132717 | 23.39506025 | 24.05697171 | 23.94360083 | 23.25882478 | 23.12126664 | 23.15486754 | 23.97989196 | 23.26570775 | 23.22006002 |
| Protein strawberry notch homolog 1 | A3KN83-3;A3KN83-2;A3KN83 | SBNO1 | Other | 0.021111276 | 0.01931579 | 25.47994354 | 24.75582727 | 24.69731546 | 25.01278035 | 24.87084217 | 22.89953026 | 24.76663364 | 24.63660648 | 24.51821323 |
| Suprabasin | Q6UWP8 | SBSN | Extracellular or secreted | -0.575316484 | 0.104231227 | 25.12198589 | 23.42123994 | 23.36212148 | 25.46995153 | 23.46238844 | 25.75348283 | 22.51096597 | 21.57658513 | 27.48208451 |
| Splicing factor, arginine/serine-rich 19 | Q9H7N4 | SCAF1 | Other | -1.359766895 | 0.845764327 | 24.99066422 | 24.77389377 | 24.57582866 | 25.18563345 | 24.75113458 | 24.67868739 | 23.87154733 | 24.21430489 | 22.31276812 |
| Protein SCAF11 | Q99590-2;Q99590 | SCAF11 | Other | -1.143048553 | 4.137002667 | 27.65266776 | 27.86820651 | 27.68096875 | 27.27710688 | 27.09828401 | 27.49442153 | 26.45751214 | 26.32839595 | 26.32077397 |
| Splicing factor, arginine/serine-rich 15 | O95104-3;O95104;O95104-2 | SCAF4 | Other | -1.211875167 | 3.432563085 | 26.54251957 | 26.54339013 | 26.58505858 | 26.10115289 | 26.52264953 | 26.2622883 | 25.3188277 | 25.01150129 | 25.31257501 |
| Protein SCAF8 | Q9UPN6;Q9UPN6-2 | SCAF8 | Other | -0.550318578 | 0.82055983 | 24.73897502 | 24.15197592 | 24.3357905 | 23.86994847 | 23.9910969 | 24.22971364 | 23.72723163 | 24.08415764 | 23.19640522 |
| Protein SCAI | Q8N9R8;Q8N9R8-2 | SCAI | Other | -0.552599846 | 0.656311349 | 23.54521808 | 24.05233521 | 24.41982648 | 24.17843265 | 24.02605802 | 24.93539565 | 23.22655684 | 23.42649614 | 24.26778053 |
| Secretory carrier-associated membra | O15126;O15126-2 | SCAMP1 | Other | -0.766546385 | 1.14486163 | 21.92798763 | 22.87713882 | 22.18513434 | 22.90944935 | 21.53829667 | 23.49935737 | 21.94685203 | 21.90712539 | 21.31506551 |
| Secretory carrier-associated membra | O14828;O14828-2 | SCAMP3 | Extracellular or secreted | -0.3322905 | 1.790596335 | 25.75197732 | 25.33091059 | 25.6500672 | 25.64759918 | 25.50993371 | 25.48704732 | 25.24832979 | 25.10057155 | 25.34299482 |
| Secretory carrier-associated membra | Q969E2-2;Q969E2 | SCAMP4 | Other | 1.019341925 | 1.327280697 | 21.1389282 | 23.12697665 | 21.18848912 | 22.27692377 | 22.47427926 | 23.42123994 | 23.18210576 | 23.18803581 | 23.50130268 |
| Lysosome membrane protein 2 | Q14108;Q14108-2 | SCARB2 | Extracellular or secreted | -0.917975979 | 0.888067653 | 25.64325619 | 25.5438031 | 25.73068179 | 25.75832042 | 26.02495972 | 26.41420714 | 24.50373065 | 24.59252 | 25.7074356 |
| Saccharopine dehydrogenase-like oxi | Q8NBX0 | SCCPDH | Other | -0.146436623 | 0.057360279 | 20.92411949 | 20.97833899 | 22.51677116 | 21.22330278 | 21.20448756 | 25.07655304 | 20.71498331 | 22.81944527 | 21.98804805 |
| Sec1 family domain-containing protei | Q8WVM8;Q8WVM8-2;Q8WV | SCFD1 | Extracellular or secreted | -0.102176871 | 0.154844764 | 28.02474842 | 28.18683324 | 28.09149035 | 27.51840836 | 27.67134918 | 27.70418007 | 27.97508806 | 27.98315819 | 27.33372794 |
| Uteroglobin | P11684 | SCGB1A1 | Extracellular or secreted | 0.788104335 | 0.196230602 | 19.96048842 | 21.75790885 | 22.08120407 | 22.15946889 | 22.67255158 | 23.88390214 | 21.2250307 | 21.77366241 | 25.62338187 |
| Selenocysteine lyase | Q96I15;Q96I15-2 | SCLY | Other | -0.276089116 | 0.425221705 | 27.49533685 | 27.62079783 | 27.54325737 | 26.58949434 | 26.81787483 | 27.11008404 | 27.17588483 | 27.10429609 | 26.47997437 |
| Protein SCO1 homolog, mitochondrial | O75880 | SCO1 | Other | 0.519815371 | 0.993139786 | 27.13633532 | 27.21113272 | 27.21613822 | 27.01403699 | 27.36986142 | 27.1814879 | 27.79259342 | 27.98294067 | 27.34840831 |
| Non-specific lipid-transfer protein | P22307-7;P22307-8;P22307;F | SCP2 | Extracellular or secreted | -0.423817882 | 0.638624259 | 27.22582171 | 27.34274056 | 27.29008791 | 26.41346485 | 26.21421224 | 27.66593949 | 26.67855308 | 26.92829626 | 26.1978304 |
| Retinoid-inducible serine carboxypept | Q9HB40;Q9HB40-2 | SCPEP1 | Extracellular or secreted | -1.564401493 | 0.728380885 | 24.0873178 | 23.93651933 | 24.15984855 | 24.69280064 | 24.76597641 | 25.66046864 | 21.35669393 | 23.87839299 | 23.72317429 |
| Protein scribble homolog | Q14160;Q14160-3;Q14160-2 | SCRIB | Extracellular or secreted | 0.470070816 | 1.911294679 | 27.00421035 | 26.77253527 | 27.09798275 | 26.79773474 | 26.97825606 | 27.22866826 | 27.48031343 | 27.59142179 | 27.27817094 |
| Secernin-1 | Q12765;Q12765-2;Q12765-3 | SCRN1 | Other | -0.017550286 | 0.015812168 | 30.61184017 | 30.50918622 | 30.45797 | 30.36348398 | 30.51662499 | 29.19121769 | 30.58091458 | 30.52775833 | 29.66383777 |
| Secernin-2 | Q96FV2-2;Q96FV2 | SCRN2 | Extracellular or secreted | -0.727563937 | 1.60050719 | 26.84326015 | 26.8460135 | 26.81421162 | 27.11107962 | 27.15643849 | 26.11330723 | 26.28098693 | 26.23896066 | 25.73951591 |
| Secernin-3 | Q0VDG4-2;Q0VDG4 | SCRN3 | Other | -0.24336481 | 0.544506027 | 22.34983949 | 23.43555327 | 23.09489017 | 23.33265287 | 23.44909624 | 23.11805328 | 23.11214612 | 22.67646159 | 22.87134052 |

| | | | | | | | | | | | | | |
|---|---|---|---|---|---|---|---|---|---|---|---|---|---|
| N-terminal kinase-like protein | Q96KG9-3;Q96KG9-4;Q96KG9 | SCYL1 | Other | -0.397673498 | 0.518944157 | 25.9111824 | 25.70898828 | 25.70268824 | 24.47116167 | 25.31049629 | 25.69559088 | 24.64691286 | 25.56560998 | 24.99451055 |
| SCY1-like protein 2 | Q6P3W7 | SCYL2 | Other | -0.367896641 | 0.89590326 | 27.29867749 | 27.41256067 | 27.65020419 | 27.03590523 | 26.62561298 | 27.42862201 | 26.98163487 | 26.58777889 | 27.05268761 |
| Syndecan-1 | P18827 | SDC1 | Extracellular or secreted | 1.966888604 | 2.078366423 | 25.24561247 | 25.16054114 | 25.06155215 | 24.79372236 | 25.1104027 | 27.68511962 | 27.07398201 | 27.26508175 | 28.09007728 |
| Syndecan-4 | P31431;P31431-2 | SDC4 | Extracellular or secreted | 2.481492247 | 2.315657713 | 27.02580464 | 26.67519125 | 26.91421996 | 25.86837138 | 25.18929583 | 28.7153526 | 28.67141667 | 29.07556394 | 29.89161396 |
| Syntenin-1 | O00560;O00560-3 | SDCBP | Extracellular or secreted | 0.344928974 | 0.341495156 | 28.62209043 | 28.5811481 | 28.68749097 | 29.5938809 | 29.69433262 | 30.55288787 | 29.24907163 | 29.45526773 | 30.196363 |
| Syntenin-1 | O00560-2 | SDCBP | Extracellular or secreted | 0.408202031 | 0.257643556 | 23.38218599 | 23.35514674 | 23.42290862 | 24.8404295 | 24.87759756 | 26.30504278 | 24.13953949 | 24.58106907 | 25.59565313 |
| Protein SDE2 homolog | Q6IQ49-3;Q6IQ49-2;Q6IQ49 | SDE2 | Other | 0.321415438 | 0.849448804 | 25.38883452 | 25.26956503 | 25.25975878 | 24.79773474 | 25.38867032 | 26.18338232 | 25.66684703 | 25.68549395 | 25.75587819 |
| Stromal cell-derived factor 2 | Q99470 | SDF2 | Extracellular or secreted | -0.515689262 | 2.104945715 | 25.73897502 | 25.58906567 | 25.62209741 | 25.89682184 | 25.90738215 | 26.12518263 | 25.12234144 | 25.34024647 | 25.43010667 |
| Stromal cell-derived factor 2-like prot | Q9HCN8 | SDF2L1 | Extracellular or secreted | -0.684239784 | 1.371770511 | 25.57577096 | 25.50121155 | 25.58046538 | 26.14914828 | 26.10171397 | 25.8948346 | 24.80705377 | 25.08212825 | 25.459671 |
| 45 kDa calcium-binding protein | Q9BRK5;Q9BRK5-6;Q9BRK5-2 | SDF4 | Extracellular or secreted | 0.143063032 | 0.205528985 | 27.94553323 | 27.95293725 | 27.89064317 | 28.09918742 | 28.34091707 | 27.20265804 | 27.78062957 | 27.87976056 | 28.48473705 |
| Succinate dehydrogenase [ubiquinone | P31040;P31040-3 | SDHA | Other | -0.322985584 | 0.967532532 | 29.13447599 | 29.23972554 | 29.22660277 | 29.32808773 | 29.36015905 | 30.22481029 | 29.15863091 | 29.06612324 | 29.06321978 |
| Succinate dehydrogenase assembly fa | Q9NX18 | SDHAF2 | Other | 1.247287739 | 1.488173287 | 20.80899873 | 20.83576362 | 20.68183204 | 22.4085464 | 22.95068332 | 22.61627512 | 22.8086453 | 22.99719258 | 23.08707496 |
| Succinate dehydrogenase [ubiquinone | P21912 | SDHB | Extracellular or secreted | -1.027973845 | 0.723262939 | 23.4781544 | 23.72035867 | 23.62555725 | 23.81870387 | 23.54002301 | 27.67283329 | 23.20498002 | 23.19347543 | 23.44543826 |
| Protein sidekick-2 | Q58EX2-2;Q58EX2-3;Q58EX2 | SDK2 | Extracellular or secreted | -0.675189301 | 0.355201532 | 24.29228399 | 21.19814525 | 22.50550048 | 21.66001157 | 22.34272361 | 22.47610341 | 20.70860062 | 21.54172613 | 22.9614895 |
| Serine dehydratase-like | Q96GA7 | SDSL | Extracellular or secreted | -0.852700916 | 1.855252339 | 26.03456256 | 26.00562438 | 26.27490532 | 25.57007689 | 25.74239723 | 25.38413106 | 25.22000096 | 25.08050267 | 24.64724233 |
| Protein SEC13 homolog | P55735;P55735-2;P55735-4;P | SEC13 | Extracellular or secreted | -0.327453218 | 4.742102705 | 29.36308707 | 29.30725967 | 29.24843837 | 29.32622418 | 29.36663454 | 29.20104594 | 29.00546912 | 28.97289918 | 28.94561694 |
| SEC14-like protein 2 | O76054;O76054-5;O76054-4 | SEC14L2 | Extracellular or secreted | 0.285208623 | 0.73155193 | 26.08196577 | 26.50663882 | 26.12060239 | 25.95398077 | 25.95422489 | 26.59988629 | 26.26992167 | 26.43636373 | 26.75798994 |
| Protein transport protein Sec16A | O15027-2;O15027-4;O15027 | SEC16A | Extracellular or secreted | -0.794715432 | 1.167383652 | 26.849239 | 26.75276855 | 26.94010924 | 26.66776753 | 26.13071085 | 27.20629614 | 25.71172116 | 25.69455513 | 26.48302306 |
| Vesicle-trafficking protein SEC22b | O75396 | SEC22B | Other | -1.273942719 | 0.843944629 | 26.47787657 | 26.11793046 | 26.32194392 | 25.75276855 | 26.02000702 | 27.09858521 | 24.2831969 | 24.64784617 | 26.14168464 |
| Protein transport protein Sec23A | Q15436;Q15436-2 | SEC23A | Other | -0.398582739 | 0.997737342 | 28.05940817 | 27.97032298 | 27.70391614 | 27.3699446 | 27.60236643 | 27.93404607 | 27.30921293 | 27.66837636 | 27.14666469 |
| Protein transport protein Sec23B | Q15437 | SEC23B | Other | -0.069332346 | 0.231731313 | 29.2017938 | 29.38710129 | 29.29554997 | 28.96886842 | 29.01542016 | 29.02989022 | 29.1672717 | 29.18368993 | 28.89035326 |
| SEC23-interacting protein | Q9Y6Y8;Q9Y6Y8-2 | SEC23IP | Other | -0.69643959 | 2.127420274 | 27.19741799 | 26.96146312 | 27.093557 | 27.43189262 | 27.50271443 | 27.03621974 | 26.54652866 | 26.28612652 | 26.6896585 |
| Protein transport protein Sec24A | O95486 | SEC24A | Other | 0.133776772 | 0.069080499 | 24.45431486 | 25.28316157 | 24.89258988 | 24.14094395 | 21.50328341 | 23.79649754 | 24.18396909 | 24.86872461 | 23.38403222 |
| Protein transport protein Sec24B | O95487-2;O95487;O95487-3 | SEC24B | Other | 0.042388068 | 0.093608605 | 24.39434075 | 24.67211841 | 24.24814879 | 24.56168277 | 24.12573444 | 24.83903613 | 24.46475811 | 24.74857778 | 24.33435895 |
| Protein transport protein Sec24C | P53992 | SEC24C | Other | -0.322370643 | 1.625717838 | 28.25110522 | 28.35896523 | 28.35317059 | 28.05216934 | 28.01786405 | 28.16389355 | 27.90299716 | 27.99361471 | 27.73486019 |
| Protein transport protein Sec24D | O94855;O94855-2 | SEC24D | Other | -1.323138676 | 0.934380624 | 24.40982856 | 24.5368851 | 24.32361117 | 24.36011299 | 24.19760546 | 24.31209024 | 23.48836582 | 23.57484757 | 22.03743735 |
| Protein transport protein Sec31A | O94979-4;O94979;O94979-6 | SEC31A | Extracellular or secreted | -0.143847209 | 0.603313837 | 28.61820916 | 28.66217456 | 28.59027991 | 28.65229165 | 28.7079029 | 28.55211833 | 28.54932987 | 28.60240183 | 28.30821493 |
| Protein transport protein Sec61 subur | P60468 | SEC61B | Other | 0.542030097 | 0.386150946 | 24.82565874 | 23.88864739 | 24.50500369 | 24.93400107 | 22.10883389 | 26.52285902 | 24.75547075 | 24.97438798 | 25.28873346 |
| Translocation protein SEC63 homolog | Q9UGP8 | SEC63 | Other | -0.549463739 | 0.216601023 | 21.75333616 | 21.43995782 | 22.03533587 | 22.00324152 | 22.74217464 | 25.59559623 | 20.64782606 | 22.15363119 | 23.33497265 |
| Secreted and transmembrane protein | Q8WVN6 | SECTM1 | Extracellular or secreted | 0.438725278 | 0.172268548 | 22.33235239 | 23.25561712 | 22.38779647 | 21.44264338 | 21.3792016 | 25.25789019 | 22.30674705 | 22.38046952 | 24.65670984 |
| Nucleoporin SEH1 | Q96EE3;Q96EE3-1 | SEH1L | Other | 0.204216034 | 0.731978772 | 27.04447717 | 27.13428013 | 27.06177874 | 27.27008212 | 27.33022677 | 27.48323838 | 27.34950545 | 27.63360269 | 27.29158161 |
| Selenium-binding protein 1 | Q13228;Q13228-4;Q13228-2 | SELENBP1 | Extracellular or secreted | -2.485220122 | 1.591251408 | 26.82747703 | 25.77140222 | 25.44912773 | 26.49200841 | 26.75964157 | 27.18857919 | 23.24884369 | 23.53409671 | 25.0055173 |
| Selenoprotein M | Q8WWX9 | SELM | Extracellular or secreted | -0.315028839 | 0.250835553 | 23.2641333 | 22.78306777 | 22.94472707 | 23.20040715 | 24.06266412 | 22.23128213 | 22.95812078 | 23.3848886 | 21.95504487 |
| Semaphorin-3F | Q13275;Q13275-2 | SEMA3F | Extracellular or secreted | -0.409937339 | 0.215182807 | 24.32175473 | 22.99327565 | 24.06332266 | 24.06718552 | 22.69962583 | 26.91067932 | 23.89522764 | 22.90529004 | 24.49759215 |
| Semaphorin-4B | Q9NPR2-2;Q9NPR2 | SEMA4B | Extracellular or secreted | -0.292808662 | 0.337309776 | 28.69102405 | 28.70902116 | 28.66955947 | 28.48146873 | 28.44683475 | 29.8247639 | 28.12558665 | 28.32225345 | 29.08506993 |
| Semaphorin-4C | Q9C0C4 | SEMA4C | Extracellular or secreted | -0.491769869 | 0.991439025 | 23.57727023 | 24.04768376 | 23.62176215 | 24.48830452 | 23.53290853 | 24.06915406 | 23.06500058 | 23.41299669 | 23.71523475 |
| Semaphorin-4D | Q92854;Q92854-2 | SEMA4D | Extracellular or secreted | 0.058249129 | 0.056826766 | 26.72403353 | 26.73130349 | 26.67465262 | 26.19253513 | 26.20968345 | 26.85638107 | 26.12581325 | 26.51933859 | 27.2238902 |
| Semaphorin-6B | Q9H3T3-3;Q9H3T3 | SEMA6B | Extracellular or secreted | 0.049281022 | 0.007763366 | 23.43198823 | 21.01270029 | 20.3384519 | 25.71740679 | 26.08565755 | 26.76395229 | 20.59372278 | 24.7317437 | 26.49745511 |
| Semaphorin-7A | O75326-2;O75326 | SEMA7A | Extracellular or secreted | -0.792156246 | 0.384133891 | 23.63644053 | 24.09715898 | 23.49057084 | 23.05811199 | 23.20557671 | 23.26856599 | 22.24643908 | 21.59284146 | 24.16246325 |
| Sentrin-specific protease 8 | Q96LD8 | SENP8 | Other | 0.353585405 | 0.350482495 | 22.34009023 | 21.23141676 | 22.34813812 | 23.64520825 | 21.08347291 | 22.87938554 | 22.91402374 | 22.61389465 | 22.29669372 |
| 15 kDa selenoprotein | O60613;O60613-2 | SEP15 | Extracellular or secreted | -0.791255563 | 0.395497736 | 24.06685717 | 23.53587716 | 24.00650209 | 24.52001338 | 24.54604285 | 25.25519084 | 24.23644462 | 24.31451244 | 22.04051799 |
| Selenide, water dikinase 1 | P49903;P49903-2;P49903-4;P | SEPHS1 | Extracellular or secreted | -0.251956815 | 0.593231744 | 29.99023682 | 30.05594841 | 29.99631133 | 29.53017942 | 29.55307103 | 29.21382302 | 29.55092655 | 29.6674461 | 29.19554192 |
| Selenide, water dikinase 2 | Q99611 | SEPHS2 | Other | -0.598558341 | 1.204796158 | 25.23815892 | 25.80333831 | 25.741446 | 25.76802295 | 25.95060324 | 25.33244801 | 25.42101517 | 24.8727688 | 24.82754972 |
| O-phosphoseryl-tRNA(Sec) selenium t | Q9HD40-3;Q9HD40 | SEPSECS | Other | 0.546823817 | 0.362729219 | 24.02369142 | 24.55496725 | 23.59718873 | 21.49302097 | 20.91231686 | 21.46159257 | 23.40366637 | 23.21184196 | 23.04635202 |
| Septin-10 | Q9P0V9;Q9P0V9-2;Q9P0V9-3 | SEPT10 | Other | 0.762693388 | 0.621203473 | 25.36182038 | 25.32240819 | 25.70308445 | 25.67494889 | 25.44950561 | 25.12856896 | 26.68016398 | 26.65321465 | 25.27486978 |
| Septin-11 | Q9NVA2;Q9NVA2-2 | SEPT11 | Extracellular or secreted | 0.222549179 | 0.346598342 | 30.80585836 | 30.97644078 | 30.86274929 | 30.37440842 | 30.30606472 | 30.36452798 | 31.04465955 | 31.08739874 | 30.38061401 |
| Septin-2 | Q15019;Q15019-3;Q15019-2 | SEPT2 | Extracellular or secreted | 0.343478457 | 0.705560776 | 30.75342226 | 30.88030081 | 30.82000742 | 30.20408172 | 30.12644852 | 30.25664911 | 31.08714579 | 30.96326977 | 30.50047472 |
| Neuronal-specific septin-3 | Q9UH03-2;Q9UH03 | SEPT3 | Other | 0.006478748 | 0.013621759 | 26.3121595 | 26.56118747 | 26.4767184 | 25.79815514 | 25.81531156 | 26.08790451 | 26.13555274 | 26.35591965 | 26.05368216 |
| Septin-6 | Q14141-2;Q14141-4;Q14141 | SEPT6 | Other | -0.158862161 | 0.353682022 | 25.14000779 | 25.44588023 | 25.08082793 | 25.02170217 | 24.58371084 | 24.23454557 | 24.91063358 | 24.72702384 | 24.63909336 |
| Septin-7 | Q16181;Q16181-2 | SEPT7 | Extracellular or secreted | 0.207064759 | 0.419480185 | 30.52775833 | 30.43628826 | 30.4546404 | 30.14831443 | 30.1679416 | 30.25721133 | 30.67337259 | 30.76927534 | 30.17462351 |
| Septin-8 | Q92599-2;Q92599-4;Q92599 | SEPT8 | Other | -0.232694979 | 0.213241641 | 26.66654909 | 26.68029814 | 26.50636642 | 25.74411814 | 25.9542027 | 25.28517416 | 26.14854718 | 26.35878921 | 25.212933 |
| Septin-9 | Q9UHD8-7;Q9UHD8-2;Q9UH | SEPT9 | Extracellular or secreted | 0.396235854 | 1.154365027 | 29.13628642 | 28.98408228 | 29.05649116 | 28.7412724 | 28.78041056 | 28.64581409 | 29.46767977 | 29.36342132 | 29.02978493 |
| Selenoprotein W | P63302 | SEPW1 | Other | 1.043695182 | 1.010894546 | 21.5250357 | 23.71753754 | 23.82269608 | 24.88413511 | 24.81186227 | 24.47605396 | 24.89730669 | 25.12947327 | 24.72296592 |
| Plasminogen activator inhibitor 1 RNA | Q8NC51;Q8NC51-2 | SERBP1 | Extracellular or secreted | 0.562079667 | 1.019273474 | 26.57142295 | 26.72559447 | 26.77228356 | 27.52969553 | 27.69318628 | 27.05961446 | 27.47931142 | 27.35892332 | 28.02390288 |
| Plasminogen activator inhibitor 1 RNA | Q8NC51-3;Q8NC51-4 | SERBP1 | Extracellular or secreted | 0.374330902 | 1.747528048 | 31.04849729 | 30.70165428 | 30.90634164 | 31.15870313 | 31.17384952 | 30.62888369 | 31.19101762 | 31.29510106 | 31.4458388 |
| Small EDRK-rich factor 2 | P84101-4;P84101;P84101-3;P | SERF2 | Other | 1.480354743 | 0.336979473 | 23.95046975 | 22.15184168 | 22.01222275 | 24.16169471 | 24.64548429 | 22.7404569 | 26.41552941 | 26.33705045 | 21.51956941 |
| Secretion-regulating guanine nucleoti | Q9UGK8 | SERGEF | Other | 0.24713377 | 0.079301583 | 20.95697371 | 22.78452622 | 20.31558648 | 22.38415796 | 23.59696134 | 23.22371716 | 20.60888605 | 23.03323545 | 23.73024125 |
| Alpha-1-antichymotrypsin;Alpha-1-an | P01011 | SERPINA3 | Extracellular or secreted | 2.819148078 | 1.892423489 | 28.00287003 | 27.95565549 | 27.83557089 | 27.50846856 | 27.55939425 | 26.94100532 | 29.9158446 | 30.14940513 | 31.29367677 |
| Leukocyte elastase inhibitor | P30740;P30740-2 | SERPINB1 | Extracellular or secreted | 0.753512473 | 1.656622695 | 32.08369493 | 32.22504022 | 32.12866237 | 31.36290946 | 31.42842235 | 30.79405096 | 32.59323534 | 32.41629078 | 32.26240144 |
| Plasminogen activator inhibitor 2 | P05120 | SERPINB2 | Extracellular or secreted | 1.225215816 | 1.389208749 | 28.11545203 | 28.24154506 | 28.14734422 | 28.95365891 | 28.93379289 | 26.53403733 | 29.68754102 | 29.70115851 | 28.74986314 |
| Serpin B3 | P29508;P29508-2 | SERPINB3 | Extracellular or secreted | -0.878645923 | 1.072183148 | 30.59477085 | 30.69536188 | 30.69818029 | 29.65652574 | 29.56824756 | 30.21835733 | 29.69297021 | 29.66076443 | 28.72604942 |
| Serpin B5 | P36952 | SERPINB5 | Extracellular or secreted | -0.129716097 | 0.334324155 | 32.54802552 | 32.59410344 | 32.55240695 | 31.80789366 | 31.85162362 | 32.22897209 | 32.06984037 | 32.28991757 | 32.0426064 |
| Serpin B6 | P35237 | SERPINB6 | Extracellular or secreted | -0.156196031 | 0.196081795 | 31.72305709 | 31.82312446 | 31.72975683 | 31.93385619 | 32.01375088 | 30.43658612 | 31.76840793 | 31.52444602 | 31.06862374 |
| Serpin B8 | P50452;P50452-2 | SERPINB8 | Extracellular or secreted | 0.827264553 | 1.711117902 | 26.84409868 | 26.77768574 | 26.72546445 | 27.02474842 | 27.21456429 | 26.65430781 | 27.59077959 | 27.48569691 | 28.02575185 |
| Serpin B9 | P50453 | SERPINB9 | Extracellular or secreted | -0.35523851 | 1.216444922 | 29.70913623 | 29.78336445 | 29.66839327 | 29.27517626 | 29.50058869 | 28.83171083 | 29.02202508 | 29.10372101 | 29.19272324 |
| Plasminogen activator inhibitor 1 | P05121-2;P05121 | SERPINE1 | Extracellular or secreted | -0.517734383 | 0.441751831 | 24.0053888 | 22.96234604 | 22.42777524 | 21.73399885 | 21.02418598 | 21.16066423 | 22.19005734 | 21.38184286 | 21.53207621 |
| Glia-derived nexin | P07093-2;P07093-3;P07093 | SERPINE2 | Extracellular or secreted | 1.425705013 | 1.77288787 | 26.45073302 | 26.75327878 | 26.47054121 | 27.47158033 | 27.48515947 | 28.26530535 | 28.21766422 | 28.32808773 | 29.17966217 |
| Pigment epithelium-derived factor | P36955 | SERPINF1 | Extracellular or secreted | -1.346746028 | 1.132630538 | 24.45287062 | 24.03632456 | 22.89602265 | 23.01706121 | 22.88569042 | 25.91620349 | 23.22514505 | 22.50242005 | 21.83428328 |
| Serpin H1 | P50454 | SERPINH1 | Extracellular or secreted | -0.290219329 | 0.19266839 | 32.10555794 | 32.33063282 | 32.26930196 | 30.54323893 | 30.66356636 | 31.28499111 | 31.7792915 | 31.6499302 | 30.29876488 |
| Protein SET | Q01105 | SET | Other | -0.103181966 | 0.353031549 | 33.69072857 | 33.89482593 | 33.7467517 | 33.81564367 | 33.81535737 | 33.65693483 | 33.81009851 | 33.45598785 | 33.73448877 |
| Protein SET | Q01105-2;Q01105-3;Q01105 | SET | Other | 0.608012156 | 0.239049041 | 29.81269783 | 28.34430776 | 28.38734785 | 30.82835506 | 27.9534369 | 31.38797432 | 28.53493528 | 30.32377864 | 31.3223824 |
| Histone-lysine N-methyltransferase SE | Q9BYW2 | SETD2 | Other | -0.371290167 | 0.427643348 | 22.51057972 | 22.49592235 | 21.23756126 | 22.89977423 | 22.62740653 | 22.7543395 | 21.4875997 | 22.20892851 | 22.45239308 |
| Histone-lysine N-methyltransferase se | Q86TU7;Q86TU7-3;Q86TU7-2 | SETD3 | Other | 0.100547472 | 0.484893102 | 27.0372676 | 26.95127048 | 27.07683843 | 27.10499587 | 27.19751173 | 27.39852191 | 27.14092445 | 27.18072944 | 27.36319153 |
| Histone-lysine N-methyltransferase SE | Q8WTS6 | SETD7 | Other | 0.0430526 | 0.058511035 | 25.54830853 | 25.10445607 | 25.28011969 | 25.81894761 | 26.13741068 | 24.48934624 | 25.2535688 | 25.42537599 | 25.63950742 |
| Seizure 6-like protein 2 | Q6UXD5-6;Q6UXD5-3;Q6UXD | SEZ6L2 | Extracellular or secreted | -0.92162366 | 1.870331428 | 26.14495474 | 26.04491485 | 25.95850139 | 26.2298603 | 26.21576804 | 26.32311291 | 25.04651855 | 25.13590495 | 25.51126163 |
| Splicing factor 1 | Q15637-4;Q15637-6;Q15637 | SF1 | Other | 0.001699385 | 0.003101619 | 29.52428721 | 29.42894142 | 29.46822376 | 29.73657075 | 29.96609244 | 29.59300822 | 29.88913499 | 29.62552589 | 29.34899918 |
| Splicing factor 1 | Q15637-5 | SF1 | Other | 0.206138449 | 0.208969468 | 23.95668598 | 24.22081285 | 23.85090865 | 24.17226451 | 22.26315785 | 23.79342471 | 24.37762625 | 23.88166366 | 23.48775271 |
| Splicing factor 3A subunit 1 | Q15459;Q15459-2 | SF3A1 | Other | -0.109325953 | 0.434411683 | 31.06400403 | 31.13314117 | 31.1369953 | 30.97062469 | 31.02650396 | 30.72378948 | 30.97514274 | 31.00021251 | 30.72419621 |
| Splicing factor 3A subunit 2 | Q15428 | SF3A2 | Other | -0.078418053 | 0.290284664 | 28.58244081 | 28.89330764 | 28.78852314 | 28.65812735 | 28.7019021 | 28.33811374 | 28.73114161 | 28.56026927 | 28.45454235 |
| Splicing factor 3A subunit 3 | Q12874 | SF3A3 | Other | 0.312287605 | 2.388311847 | 29.83419994 | 29.8255526 | 29.83423008 | 29.74925577 | 29.84523591 | 29.43215156 | 30.10674383 | 30.02867893 | 30.06175299 |
| Splicing factor 3B subunit 1 | O75533 | SF3B1 | Other | 0.897993765 | 2.667521638 | 30.45904558 | 30.86798567 | 30.79149154 | 29.87437628 | 29.98304943 | 30.50843078 | 31.39447977 | 31.35326525 | 31.18842591 |
| Splicing factor 3B subunit 2 | Q13435 | SF3B2 | Other | 0.232626781 | 1.413514867 | 31.07680021 | 31.12774034 | 30.98352518 | 30.98874804 | 31.0089184 | 30.56174467 | 31.18364261 | 31.1322833 | 31.25569285 |
| Splicing factor 3B subunit 3 | Q15393 | SF3B3 | Other | -0.033425777 | 0.113807814 | 32.19521923 | 32.28165138 | 32.20585935 | 31.77120542 | 31.85333514 | 31.74258194 | 32.03223091 | 32.00530848 | 31.88710951 |
| Splicing factor 3B subunit 4 | Q15427 | SF3B4 | Other | -0.738155695 | 1.697089947 | 28.65765046 | 29.02799384 | 29.04468561 | 29.02983757 | 29.23242444 | 28.55244818 | 28.50206945 | 28.07963834 | 27.97634517 |
| Splicing factor 3B subunit 5 | Q9BWJ5 | SF3B5 | Other | 0.699550729 | 0.488595562 | 25.05353716 | 23.89106054 | 24.46506962 | 25.02812561 | 25.59781388 | 24.174933 | 26.19112354 | 24.38126199 | 25.63153656 |
| Splicing factor 3B subunit 6 | Q9Y3B4 | SF3B6 | Other | 0.011909632 | 0.030451711 | 26.94692786 | 26.82735589 | 26.80661137 | 27.24154506 | 27.33543275 | 27.31691332 | 26.97092633 | 27.03621974 | 27.26597595 |
| 14-3-3 protein sigma | P31947;P31947-2 | SFN | Extracellular or secreted | 1.418220129 | 1.156050925 | 31.63922967 | 31.69780758 | 31.90232159 | 31.64667258 | 31.79246155 | 32.15915443 | 32.78980217 | 32.83552571 | 34.04815621 |
| Splicing factor, proline- and glutamine | P23246;P23246-2 | SFPQ | Other | -0.419017185 | 0.477760588 | 31.16566747 | 30.89309061 | 30.96368319 | 32.25064791 | 32.11916808 | 31.912672 | 30.73448877 | 30.94349464 | 31.71742968 |
| Secreted frizzled-related protein 1 | Q8N474 | SFRP1 | Extracellular or secreted | 0.200375225 | 0.066208276 | 25.13402547 | 25.29953365 | 25.15618772 | 26.51695179 | 26.5308566 | 29.03123198 | 25.21600867 | 26.09412103 | 28.12538958 |
| Splicing factor, suppressor of white-ap | Q12872;Q12872-2 | SFSWAP | Other | -1.132260529 | 0.641408135 | 23.37597061 | 23.52516138 | 23.60127569 | 21.86473043 | 21.85550812 | 24.42527994 | 22.94990007 | 22.26699649 | 20.71028493 |
| Vesicle transport protein SFT2B | O95562 | SFT2D2 | Extracellular or secreted | 1.738998315 | 0.444351886 | 22.55431826 | 22.34829828 | 21.82207221 | 25.11723693 | 21.89411149 | 21.90509269 | 21.47064855 | 25.26162495 | 26.30528637 |
| Sideroflexin-1 | Q9H9B4 | SFXN1 | Other | -0.958953504 | 0.300047798 | 22.52551985 | 21.81624463 | 22.46789506 | 21.71279763 | 21.7940398 | 27.50922398 | 20.13473292 | 22.56943381 | 23.33183324 |
| Epsilon-sarcoglycan | O43556;O43556-3;O43556-4 | SGCE | Extracellular or secreted | 0.089715844 | 0.060088402 | 25.62953694 | 25.55537692 | 25.4256321 | 25.75110903 | 26.09914728 | 26.66180093 | 25.52554972 | 25.43163762 | 26.87326179 |
| N-sulphoglucosamine sulphohydrolas | P51688 | SGSH | Extracellular or secreted | -0.850559973 | 0.797428979 | 23.9680719 | 24.34948858 | 24.63599792 | 24.12864762 | 24.37006104 | 22.2038369 | 23.39427532 | 23.54226863 | 22.33982811 |
| Small glutamine-rich tetratricopeptide | O43765 | SGTA | Other | 0.059403383 | 0.148754101 | 28.23548628 | 28.46228702 | 28.4503004 | 28.87707093 | 29.02408788 | 28.31639548 | 28.74970333 | 28.64385481 | 28.467466 |
| SH2 domain-containing protein 4A | Q9H788-2;Q9H788 | SH2D4A | Other | 0.507018124 | 1.431585692 | 24.61470355 | 25.20288218 | 25.02867366 | 25.72327847 | 24.63627457 | 24.63394907 | 25.58591818 | 25.40060795 | 25.454409 |
| SH3 domain-binding glutamic acid-ric | O75368 | SH3BGRL | Extracellular or secreted | 0.133203264 | 0.15101925 | 23.00581709 | 22.74978515 | 22.35968415 | 23.14276771 | 23.12803398 | 22.08883869 | 22.77776098 | 23.39759765 | 22.46171456 |
| SH3 domain-binding glutamic acid-ric | Q9UJC5 | SH3BGRL2 | Extracellular or secreted | -0.256608156 | 0.23862013 | 26.23229634 | 26.22690589 | 26.03397475 | 25.3807007 | 25.24818499 | 24.34624535 | 25.67028237 | 25.59477085 | 24.69927633 |
| SH3 domain-binding glutamic acid-ric | Q9H299 | SH3BGRL3 | Extracellular or secreted | 0.16039394 | 0.138101553 | 29.37920975 | 28.7901402 | 28.71558171 | 28.68645607 | 28.89879483 | 27.69842871 | 29.19917459 | 29.23512102 | 28.13119184 |
| SH3 domain-binding protein 1 | Q9Y3L3-2;Q9Y3L3 | SH3BP1 | Other | -0.712992891 | 0.814474482 | 22.33142324 | 22.76510639 | 22.01994766 | 21.88211164 | 22.36412717 | 20.81607851 | 20.98070499 | 21.96055688 | 21.00915676 |
| SH3 domain-binding protein 5-like | Q7L8J4;Q7L8J4-2 | SH3BP5L | Other | 1.651430379 | 3.046109975 | 22.49633543 | 23.34746241 | 22.96657656 | 23.778463 | 23.8247366 | 23.04204875 | 25.04222413 | 25.11489628 | 24.5249821 |
| Endophilin-A2 | Q99961;Q99961-2 | SH3GL1 | Other | 0.738683029 | 0.676232734 | 24.28961312 | 24.00358859 | 24.05523477 | 23.80508651 | 21.15525791 | 24.23987119 | 24.36499544 | 24.65326933 | 23.97211036 |
| Endophilin-B2 | Q9NR46;Q9NR46-2 | SH3GLB2 | Other | -0.01331102 | 0.03566453 | 27.40356886 | 27.62651839 | 27.59690449 | 27.23434456 | 27.16038726 | 27.41828294 | 27.59184976 | 27.33031226 | 27.25790817 |
| SH3 domain-containing kinase-binding | Q96B97-3 | SH3KBP1 | Extracellular or secreted | 1.315558877 | 2.726457571 | 26.45838859 | 26.30171541 | 26.23976196 | 27.00345992 | 26.62393997 | 25.36759712 | 27.59676234 | 27.67660417 | 27.6707416 |
| SH3 and PX domain-containing protei | A1X283 | SH3PXD2B | Extracellular or secreted | 0.031212317 | 0.070181934 | 24.29600093 | 24.29305622 | 24.52581852 | 24.13969561 | 24.24270835 | 24.64861432 | 24.60291856 | 24.41828294 | 24.14538242 |
| SHC-transforming protein 1 | P29353-2;P29353-7;P29353;P | SHC1 | Extracellular or secreted | 0.122015842 | 0.121999403 | 24.24314434 | 23.45412656 | 24.09474524 | 23.49850546 | 24.00376014 | 23.01014425 | 23.75877788 | 24.40743167 | 23.35205097 |
| Protein shisa-2 homolog | Q6UWI4 | SHISA2 | Other | 2.692404374 | 1.589485004 | 21.68411826 | 22.98154777 | 21.91412753 | 23.50978273 | 21.12444381 | 23.60308841 | 24.52121223 | 24.5318978 | 26.43265735 |
| Serine hydroxymethyltransferase, cyt | P34896-2;P34896;P34896-3;P | SHMT1 | Extracellular or secreted | 0.115393544 | 0.437121203 | 29.55418783 | 29.45075268 | 29.51591103 | 28.98239673 | 29.03601007 | 29.2583351 | 29.30862723 | 29.40851724 | 29.52783288 |
| Serine hydroxymethyltransferase, mit | P34897-3;P34897;P34897-2 | SHMT2 | Extracellular or secreted | -0.1702641 | 0.191673834 | 29.76411052 | 29.89094752 | 29.86800039 | 30.5549014 | 30.67682271 | 30.97705523 | 29.8723373 | 29.82092042 | 30.66186887 |
| Leucine-rich repeat protein SHOC-2 | Q9UQ13;Q9UQ13-2 | SHOC2 | Other | 0.400650555 | 0.106628311 | 24.50815117 | 20.70310093 | 25.40840386 | 22.23059756 | 20.88297546 | 21.47000297 | 24.28757148 | 23.75368684 | 20.76230932 |
| Sedoheptulokinase | Q9UHJ6 | SHPK | Other | -0.052200152 | 0.072457388 | 27.2131741 | 27.30138399 | 27.12396003 | 26.95670813 | 27.1061947 | 26.26359616 | 27.20489796 | 27.07683843 | 26.54417172 |
| Protein Shroom3 | Q8TF72 | SHROOM3 | Extracellular or secreted | 1.067092037 | 2.10446464 | 24.47975856 | 24.18343912 | 24.6494917 | 25.00598837 | 25.51583961 | 24.90407743 | 25.82859112 | 25.54527701 | 26.19670537 |
| Sialate O-acetylesterase | Q9HAT2;Q9HAT2-2 | SIAE | Extracellular or secreted | -1.109707492 | 3.154982101 | 28.24581192 | 28.28135844 | 28.21377668 | 28.45485607 | 28.51020543 | 27.53304225 | 26.98272312 | 26.97574408 | 27.33193572 |
| Paired amphipathic helix protein Sin3 | Q96ST3 | SIN3A | Extracellular or secreted | -0.918207721 | 0.582530168 | 26.71685755 | 27.10978523 | 26.91843454 | 24.10445607 | 24.778864 | 25.55546469 | 25.43860203 | 25.51108062 | 23.88762522 |
| Signal-regulatory protein beta-1 isofo | Q5TFQ8;P78324-4;P78324;P7 | SIRPB1;SIRPA | Extracellular or secreted | 0.412417785 | 0.444175077 | 24.65796386 | 24.55385472 | 24.77349138 | 25.37195607 | 24.86353522 | 25.00812766 | 25.06952286 | 24.81641066 | 25.96578428 |
| NAD-dependent protein deacylase sir | Q9NXA8-2;Q9NXA8;Q9NXA8- | SIRT5 | Other | -0.074880461 | 0.100614994 | 24.58066663 | 24.30194212 | 24.36859651 | 24.02021902 | 23.63278491 | 23.98971186 | 24.49899233 | 23.72515237 | 23.99817445 |
| Helicase SKI2W | Q15477 | SKIV2L | Other | 0.045107985 | 0.132544023 | 25.87342609 | 25.68011031 | 25.83646206 | 25.32687134 | 25.65447171 | 25.61197732 | 25.84143766 | 25.76865401 | 25.5168917 |
| Superkiller viralicidic activity 2-like 2 | P42285 | SKIV2L2 | Other | -0.123524972 | 0.171395146 | 28.7284518 | 28.897324 | 28.81558641 | 28.67296814 | 28.86917779 | 28.26758411 | 28.837106 | 28.82174163 | 28.09612358 |
| S-phase kinase-associated protein 1 | P63208;P63208-2 | SKP1 | Extracellular or secreted | -0.074086887 | 0.183637541 | 30.25146645 | 30.07737338 | 30.04895194 | 30.3483661 | 30.5388064 | 29.88021322 | 30.12878526 | 30.32130745 | 29.90023536 |
| Solute carrier family 12 member 2 | P55011-3;P55011 | SLC12A2 | Extracellular or secreted | 2.450578696 | 1.672575451 | 24.20702266 | 23.68495917 | 24.14623739 | 20.7756321 | 20.23968367 | 23.96102146 | 25.12257842 | 25.3862051 | 25.35023079 |
| Solute carrier family 12 member 7 | Q9Y666 | SLC12A7 | Extracellular or secreted | 2.530162781 | 2.531316904 | 20.67918339 | 23.82454238 | 20.76142054 | 22.30556634 | 22.65602122 | 21.34875716 | 24.18139312 | 24.36706382 | 24.82977692 |
| Monocarboxylate transporter 1 | P53985;P53985-2 | SLC16A1 | Extracellular or secreted | 1.770800936 | 3.149980345 | 26.30923028 | 26.62240467 | 26.74316804 | 25.74085438 | 26.31430498 | 26.49658685 | 27.84535557 | 28.12730986 | 28.45301197 |
| Monocarboxylate transporter 4 | O15427 | SLC16A3 | Extracellular or secreted | 1.462568245 | 2.474664977 | 25.35396978 | 25.3959101 | 25.65965234 | 24.15468216 | 24.33388145 | 25.88583469 | 26.42042095 | 26.4397912 | 26.91945785 |
| Folate transporter 1 | P41440;P41440-3;P41440-2 | SLC19A1 | Extracellular or secreted | 3.535436716 | 3.053298313 | 21.21249637 | 20.62434983 | 22.29877824 | 21.23532938 | 21.08350702 | 23.9199806 | 24.64107978 | 25.18737141 | 25.96507967 |
| Neutral amino acid transporter A | P43007;P43007-2 | SLC1A4 | Extracellular or secreted | 1.644040361 | 1.544941175 | 25.50130268 | 24.9383154 | 25.55353251 | 25.38343905 | 25.27020691 | 26.20692954 | 26.64361402 | 26.87384847 | 27.84152164 |
| Neutral amino acid transporter B(0) | Q15758;Q15758-3;Q15758-2 | SLC1A5 | Extracellular or secreted | 1.190898083 | 2.13952595 | 27.12898187 | 27.42037276 | 27.44379546 | 27.3359438 | 27.32997025 | 27.52178133 | 28.42422294 | 28.39701253 | 28.8418815 |
| Sodium-dependent phosphate transp | Q8WUM9 | SLC20A1 | Extracellular or secreted | 0.878993098 | 1.1075481 | 22.62631341 | 23.350978 | 22.06320208 | 22.9817493 | 22.69672333 | 21.16968133 | 22.94581673 | 23.25270296 | 23.88278333 |
| Mitochondrial 2-oxoglutarate/malate | Q02978;Q02978-2 | SLC25A11 | Extracellular or secreted | -0.702551214 | 0.221850039 | 22.22349623 | 22.11286198 | 22.80903799 | 22.31274117 | 22.21865665 | 27.62058811 | 21.7149414 | 21.51151444 | 24.31458158 |
| Calcium-binding mitochondrial carrier | Q9UJS0;Q9UJS0-2 | SLC25A13 | Extracellular or secreted | -1.749407898 | 0.844261293 | 24.38692867 | 24.25486657 | 24.18737141 | 24.04959599 | 23.76941092 | 27.52110737 | 22.83803591 | 21.63900501 | 24.35937585 |
| Calcium-binding mitochondrial carrier | Q6NUK1-2;Q6NUK1 | SLC25A24 | Other | -1.294323618 | 0.542412157 | 24.43949003 | 23.40392636 | 23.46861609 | 23.54215053 | 23.53978642 | 25.70126098 | 20.91176552 | 23.67066057 | 23.58221826 |
| Phosphate carrier protein, mitochond | Q00325-2;Q00325 | SLC25A3 | Extracellular or secreted | 0.199023859 | 0.058703772 | 26.51583961 | 27.04124254 | 27.46645503 | 26.73363251 | 26.47159584 | 31.10199442 | 26.62282355 | 27.03485638 | 29.60477163 |
| ADP/ATP translocase 2;ADP/ATP trans | P05141 | SLC25A5 | Extracellular or secreted | 0.946242106 | 0.272155762 | 23.68110284 | 24.33565422 | 22.70212279 | 25.59346065 | 22.03387938 | 29.85716751 | 24.91241646 | 24.63561052 | 27.39239303 |
| ADP/ATP translocase 3;ADP/ATP trans | P12236;P12235 | SLC25A6;SLC25A4 | Other | 0.366498819 | 0.129000447 | 27.10948636 | 27.05755022 | 27.64718743 | 26.76850258 | 26.59163579 | 31.15454455 | 27.04301727 | 27.57852322 | 29.64240943 |
| Sulfate transporter | P50443 | SLC26A2 | Extracellular or secreted | 2.600930732 | 3.222357764 | 22.00573842 | 22.17903126 | 21.60307508 | 22.02837762 | 20.11727107 | 20.96904624 | 24.2298603 | 24.45920196 | 23.56499978 |
| Equilibrative nucleoside transporter 1 | Q99808;Q99808-2 | SLC29A1 | Extracellular or secreted | 2.158679806 | 4.078075482 | 24.95566658 | 25.46911314 | 25.33258459 | 25.85447998 | 25.69861417 | 26.3757718 | 27.52245498 | 27.72916525 | 28.06753431 |
| Equilibrative nucleoside transporter 2 | Q14542;Q14542-3 | SLC29A2 | Extracellular or secreted | 2.683546652 | 2.501857005 | 21.01175058 | 22.5091545 | 21.04604249 | 23.0404108 | 21.48519983 | 23.27895076 | 24.00966599 | 24.90573074 | 25.32099771 |
| Solute carrier family 2, facilitated gluc | P11166 | SLC2A1 | Extracellular or secreted | 0.897892389 | 0.989702842 | 27.06424822 | 27.45222653 | 27.47877159 | 26.16109881 | 26.48136094 | 27.459671 | 27.66343063 | 27.47243279 | 28.6065023 |
| Zinc transporter 1 | Q9Y6M5 | SLC30A1 | Extracellular or secreted | 1.094306988 | 1.002287031 | 21.40786352 | 23.98953864 | 24.38640248 | 24.56412762 | 25.16315458 | 24.51195529 | 25.00238721 | 25.1694764 | 25.12257842 |
| High affinity copper uptake protein 1 | O15431 | SLC31A1 | Extracellular or secreted | 2.286661365 | 1.497515517 | 23.57634779 | 23.60681996 | 23.57161101 | 23.95358121 | 21.53233701 | 20.6075217 | 24.53492786 | 24.50573064 | 26.24343493 |
| Adenosine 3-phospho 5-phosphosulfa | Q8TB61-5;Q8TB61-4;Q8TB61 | SLC35B2 | Other | -0.696045705 | 0.267663931 | 21.06553169 | 23.00523459 | 22.46368601 | 23.50990352 | 20.92491635 | 24.11751438 | 21.36586461 | 20.5714185 | 23.51797304 |
| Solute carrier family 35 member F6 | Q8N357 | SLC35F6 | Extracellular or secreted | -0.571492906 | 0.150485682 | 24.26363198 | 21.36240352 | 21.21080212 | 24.47035502 | 21.69463959 | 25.5251315 | 21.88731216 | 20.85621082 | 24.80548017 |
| Sodium-coupled neutral amino acid tr | Q9H2H9 | SLC38A1 | Extracellular or secreted | 4.328892376 | 2.703704401 | 22.04496903 | 22.12738859 | 21.03556968 | 22.36273685 | 21.99082 | 22.27042081 | 25.99072913 | 25.8385313 | 27.07336918 |
| Zinc transporter ZIP1 | Q9NY26-2;Q9NY26 | SLC39A1 | Extracellular or secreted | 0.684658616 | 0.206459267 | 20.3691585 | 20.63597961 | 22.89085729 | 22.78839018 | 21.31498095 | 21.0944733 | 23.57230517 | 23.10749231 | 19.92109829 |

| | | | | | | | | | | | | | | |
|---|---|---|---|---|---|---|---|---|---|---|---|---|---|---|
| Zinc transporter ZIP10 | Q9ULF5 | SLC39A10 | Extracellular or secreted | 1.685461776 | 1.14543774 | 24.3743794 | 25.28563278 | 25.14720834 | 22.70895672 | 22.74067427 | 25.31447786 | 25.96921435 | 24.97941229 | 26.89342338 |
| Zinc transporter ZIP14 | Q15043-2;Q15043-3;Q15043 | SLC39A14 | Extracellular or secreted | 2.117742998 | 3.829425052 | 22.8847593 | 22.73434082 | 23.78346749 | 24.07724602 | 24.05870656 | 23.37052672 | 25.56717787 | 25.89041124 | 25.35016333 |
| Zinc transporter ZIP6 | Q13433;Q13433-2 | SLC39A6 | Extracellular or secreted | 1.496173923 | 1.19381493 | 23.23625483 | 23.31741372 | 23.42495972 | 23.13026701 | 23.57819208 | 23.889576 | 24.07773499 | 25.26795908 | 25.43115937 |
| Zinc transporter SLC39A7 | Q92504 | SLC39A7 | Other | -0.259029267 | 0.07108938 | 22.43209021 | 21.92078025 | 22.3528592 | 22.05278295 | 20.79590331 | 26.34861092 | 21.25390056 | 21.49243023 | 24.42809483 |
| Zinc transporter ZIP9 | Q9NUM3-3;Q9NUM3-2;Q9NU | SLC39A9 | Other | -0.371415582 | 0.691591817 | 22.17936862 | 22.59609692 | 21.92317658 | 22.73708804 | 22.81019583 | 23.70819899 | 22.1947833 | 22.32487493 | 22.34315752 |
| 4F2 cell-surface antigen heavy chain | P08195-2;P08195-3;P08195;F | SLC3A2 | Extracellular or secreted | 1.008540766 | 3.495356495 | 30.36765543 | 30.26673558 | 30.44519748 | 30.70528474 | 30.89987536 | 30.63512613 | 31.38710129 | 31.59214392 | 31.70631446 |
| Choline transporter-like protein 1 | Q8WWI5-3;Q8WWI5-2;Q8WV | SLC44A1 | Extracellular or secreted | 0.856757858 | 1.504333062 | 26.96024822 | 27.1580771 | 27.07132454 | 26.87900096 | 26.95193742 | 27.73885908 | 27.77718407 | 27.76547065 | 28.40734252 |
| Choline transporter-like protein 2 | Q8IWA5;Q8IWA5-3;Q8IWA5- | SLC44A2 | Extracellular or secreted | 0.787750064 | 2.00923126 | 27.81763091 | 27.92167253 | 27.90936365 | 27.14170413 | 27.19281729 | 27.59981537 | 28.10699337 | 28.44217452 | 28.60558424 |
| Kanadaptin | Q9BWU0 | SLC4A1AP | Extracellular or secreted | 0.566596357 | 1.542007888 | 27.30312743 | 26.93134313 | 27.00528171 | 27.09858521 | 27.14374878 | 26.81200922 | 27.77001616 | 27.72318732 | 27.35363334 |
| Anion exchange protein 2 | P04920-2;P04920-3;P04920 | SLC4A2 | Extracellular or secreted | 2.323006155 | 2.918798581 | 24.32964526 | 22.51674746 | 23.89078231 | 24.75414577 | 25.05084174 | 23.91588448 | 26.27458546 | 26.44627473 | 26.47718178 |
| Sodium bicarbonate cotransporter 3 | Q9Y6M7-6;Q9Y6M7-13;Q9Y6 | SLC4A7 | Extracellular or secreted | 1.792839619 | 1.246476653 | 23.73727375 | 24.15930965 | 24.19835511 | 22.30710851 | 23.34421462 | 23.36693047 | 24.4427203 | 25.19053968 | 26.30185493 |
| Sodium-dependent multivitamin trans | Q9Y289 | SLC5A6 | Extracellular or secreted | 1.708923234 | 1.597098154 | 23.51086942 | 23.76547065 | 23.98494061 | 22.98250553 | 24.28355018 | 23.80282083 | 24.70382375 | 25.67704794 | 25.91097662 |
| Sodium- and chloride-dependent GAB | P48066-2;P48066 | SLC6A11 | Extracellular or secreted | 0.562873301 | 0.891651123 | 21.92078363 | 21.20287471 | 21.82534574 | 21.77632547 | 21.2026906 | 22.10677378 | 22.09029337 | 21.90569402 | 22.71002949 |
| Sodium- and chloride-dependent neu | Q9UN76 | SLC6A14 | Extracellular or secreted | 3.141400083 | 2.41324253 | 19.51511497 | 23.07996583 | 23.23873476 | 22.85527375 | 22.88324961 | 24.12612846 | 25.42403068 | 25.77024306 | 26.0791602 |
| Sodium- and chloride-dependent taur | P31641-2;P31641 | SLC6A6 | Extracellular or secreted | 3.469354507 | 4.797869791 | 21.84903852 | 21.51127973 | 21.25141769 | 20.8825834 | 22.03412704 | 22.44614535 | 24.75067469 | 25.24524978 | 25.39943492 |
| Sodium- and chloride-dependent crea | P48029;P48029-4 | SLC6A8 | Extracellular or secreted | 3.209938737 | 1.903945449 | 21.25394662 | 21.53431834 | 20.3748788 | 21.60451322 | 23.12130668 | 22.70927232 | 24.46818103 | 24.2756869 | 26.18506627 |
| High affinity cationic amino acid trans | P30825 | SLC7A1 | Extracellular or secreted | 2.106579714 | 4.279412053 | 23.9624343 | 23.99619279 | 24.02993759 | 24.24474185 | 23.98111222 | 24.21852362 | 26.09758097 | 26.34243539 | 26.09619397 |
| Large neutral amino acids transporter | Q01650 | SLC7A5 | Extracellular or secreted | 1.42079339 | 2.183913922 | 27.11674134 | 26.9612423 | 27.15189852 | 27.64842232 | 27.43053741 | 28.28603836 | 28.58301497 | 28.64835374 | 29.32845159 |
| Y+L amino acid transporter 2 | Q92536 | SLC7A6 | Extracellular or secreted | 2.169175603 | 3.035949655 | 22.79225333 | 22.85163315 | 21.15180777 | 20.89264025 | 21.70586528 | 21.9492411 | 24.48701664 | 23.82454238 | 23.86768822 |
| Sodium/hydrogen exchanger 1 | P19634;P19634-2 | SLC9A1 | Extracellular or secreted | 0.292971818 | 0.147027352 | 20.7914195 | 21.6743752 | 22.45244336 | 21.65219227 | 22.40049679 | 21.40973931 | 22.62086773 | 20.69259441 | 22.75578653 |
| Na(+)/H(+) exchange regulatory cofac | O14745;O14745-2 | SLC9A3R1 | Extracellular or secreted | 0.148636423 | 0.741658771 | 29.56219974 | 29.55055966 | 29.58367857 | 29.53519491 | 29.51222286 | 29.60960525 | 29.61050316 | 29.65758232 | 29.85455428 |
| Na(+)/H(+) exchange regulatory cofac | Q15599-2;Q15599 | SLC9A3R2 | Extracellular or secreted | 0.587757124 | 0.348024807 | 24.490326 | 20.6492541 | 23.66264317 | 25.15093073 | 25.19340023 | 24.50827209 | 24.14833382 | 24.75327878 | 24.68907192 |
| Solute carrier organic anion transport | Q9UIG8-2;Q9UIG8 | SLCO3A1 | Extracellular or secreted | 1.101801946 | 1.344271467 | 23.90187007 | 22.26899423 | 22.48145949 | 22.99543387 | 21.36067565 | 23.97997915 | 23.61616292 | 24.10221474 | 24.08123441 |
| SRA stem-loop-interacting RNA-bindir | Q9GZT3-2;Q9GZT3 | SLIRP | Other | -0.522200083 | 0.491387543 | 25.56903391 | 25.88296986 | 26.68538701 | 26.54220958 | 26.77756034 | 28.36790534 | 26.0935167 | 25.62959253 | 26.62282355 |
| Slit homolog 1 protein | O75093-2;O75093 | SLIT1 | Extracellular or secreted | -6.975731808 | 6.233158324 | 28.3223824 | 28.47823156 | 28.53359928 | 28.03433167 | 28.67367586 | 26.61217424 | 21.03985883 | 21.36121131 | 20.99893194 |
| SLIT and NTRK-like protein 6 | Q9H5Y7 | SLITRK6 | Extracellular or secreted | 0.036015189 | 0.010367634 | 22.2651068 | 22.74865967 | 22.43374639 | 23.38086581 | 21.62502207 | 21.76464662 | 20.37435849 | 22.89133781 | 23.95137294 |
| STE20-like serine/threonine-protein k | Q9H2G2-2;Q9H2G2 | SLK | Extracellular or secreted | 0.043628859 | 0.057219915 | 26.27097323 | 26.33258459 | 26.13520043 | 26.16392234 | 26.14275603 | 25.45791913 | 25.93089215 | 26.57873915 | 25.87293315 |
| Antileukoproteinase | P03973 | SLPI | Extracellular or secreted | -0.458093293 | 0.679658287 | 29.95726184 | 29.61866426 | 29.56065193 | 29.57828927 | 29.66117232 | 31.10218219 | 29.22837009 | 29.24780484 | 29.8886561 |
| SAFB-like transcription modulator | Q9NWH9 | SLTM | Extracellular or secreted | -1.4461838 | 2.93356703 | 28.07142685 | 27.96306302 | 27.92903035 | 27.14540185 | 27.40624796 | 27.27132951 | 25.94433835 | 26.51316085 | 26.09719918 |
| Mothers against decapentaplegic hon | Q13485 | SMAD4 | Other | 0.101625464 | 0.328338168 | 24.33722063 | 24.19415206 | 23.9680719 | 24.41060507 | 24.71229861 | 24.38357089 | 24.26549315 | 24.5068809 | 24.53546192 |
| Small acidic protein | O00193 | SMAP | Extracellular or secreted | 0.41452471 | 0.83270747 | 26.79587855 | 27.37633501 | 26.99711934 | 27.51344702 | 27.49982874 | 26.05100775 | 27.25898648 | 27.60865392 | 27.49274193 |
| Stromal membrane-associated protei | Q8IYB5-3;Q8IYB5-2;Q8IYB5 | SMAP1 | Extracellular or secreted | -0.134997403 | 0.656916941 | 26.3824169 | 26.64030761 | 26.62491613 | 26.83207315 | 26.75264096 | 26.23750262 | 26.37230488 | 26.49393312 | 26.46369848 |
| Stromal membrane-associated protei | Q8WU79-2;Q8WU79;Q8WU7 | SMAP2 | Other | 0.420378659 | 0.55009908 | 22.78948987 | 22.4736037 | 20.68913793 | 21.83023927 | 21.73326529 | 22.54373231 | 22.58136795 | 22.63043156 | 22.07907065 |
| Transcription activator BRG1 | P51532-5;P51532-2;P51532-3 | SMARCA4 | Extracellular or secreted | -0.634657206 | 1.226726387 | 27.33730571 | 27.32199551 | 27.29684117 | 27.34832388 | 27.07999431 | 27.81726494 | 27.01361113 | 26.82238002 | 26.36089998 |
| SWI/SNF-related matrix-associated ac | O60264 | SMARCA5 | Other | -0.684241206 | 1.436372359 | 27.89411759 | 27.8086993 | 27.79890909 | 26.62058811 | 26.79995902 | 27.91096518 | 26.7854395 | 26.66709075 | 26.91136529 |
| SWI/SNF-related matrix-associated ac | Q12824-2;Q12824 | SMARCB1 | Other | -0.472024191 | 0.58537992 | 25.6614204 | 25.75760853 | 25.46877144 | 25.08094989 | 24.63494617 | 25.34989347 | 25.43399465 | 24.71858304 | 24.40814468 |
| SWI/SNF complex subunit SMARCC1 | Q92922 | SMARCC1 | Other | -0.621288396 | 0.950197316 | 29.70175341 | 29.69183942 | 29.68667313 | 29.40653182 | 29.39776742 | 29.6086363 | 29.18914499 | 29.20643589 | 28.48715468 |
| SWI/SNF complex subunit SMARCC2 | Q8TAQ2;Q8TAQ2-2;Q8TAQ2- | SMARCC2 | Other | -1.307080249 | 3.318609352 | 28.50627562 | 28.37873447 | 28.29447674 | 28.24413392 | 28.21155051 | 27.98201585 | 27.00399598 | 27.11763327 | 26.76572355 |
| SWI/SNF-related matrix-associated ac | Q96GM5;Q96GM5-2 | SMARCD1 | Other | -0.975339717 | 0.458522844 | 23.19648026 | 23.0391058 | 23.06175814 | 21.95466827 | 21.82469115 | 22.98730218 | 20.55103417 | 23.24401997 | 21.31092961 |
| SWI/SNF-related matrix-associated ac | Q92925-3;Q92925;Q92925-2 | SMARCD2 | Other | -0.635357605 | 1.838774658 | 27.29509011 | 27.42518388 | 27.41320657 | 26.78006632 | 26.83026064 | 27.33739079 | 26.8046435 | 26.38725745 | 26.4426254 |
| SWI/SNF-related matrix-associated ac | Q969G3;Q969G3-2;Q969G3-4 | SMARCE1 | Other | -0.269470871 | 1.238132319 | 28.00233355 | 27.92654421 | 27.88465917 | 27.74432348 | 27.51058274 | 27.50937502 | 27.55581572 | 27.58448523 | 27.34019552 |
| Structural maintenance of chromosor | Q14683 | SMC1A | Other | -0.599990042 | 0.864329468 | 31.10562038 | 31.13087283 | 31.0457013 | 30.4690782 | 30.61342188 | 30.14455124 | 30.38164569 | 30.4546404 | 29.61836671 |
| Structural maintenance of chromosor | O95347;O95347-2 | SMC2 | Extracellular or secreted | 0.422953048 | 0.796180301 | 28.45885008 | 28.47630123 | 28.54874214 | 27.75066191 | 27.97454115 | 28.52954661 | 29.01108002 | 28.79952679 | 28.32757389 |
| Structural maintenance of chromosor | Q9UQE7 | SMC3 | Extracellular or secreted | -0.603636407 | 0.802005364 | 30.99138619 | 31.03420046 | 31.00262864 | 29.98712216 | 30.14515887 | 30.02907403 | 30.17521861 | 30.2443381 | 29.36431924 |
| Structural maintenance of chromosor | Q9NTJ3;Q9NTJ3-2 | SMC4 | Other | 0.128328999 | 0.24017457 | 29.51271265 | 29.36836338 | 29.40037586 | 29.15561849 | 29.20969275 | 29.56935268 | 29.80525875 | 29.53378491 | 29.15400124 |
| Structural maintenance of chromosor | Q8IY18 | SMC5 | Other | -0.382293391 | 0.250310824 | 24.7926803 | 24.60665055 | 24.78511518 | 23.67745126 | 23.37278643 | 24.22861321 | 24.23425318 | 24.53747768 | 22.81303742 |
| Structural maintenance of chromosor | Q96SB8;Q96SB8-2 | SMC6 | Other | -0.294789657 | 0.616826521 | 25.37716286 | 25.04839075 | 24.74493934 | 25.21044543 | 24.52019327 | 24.8447931 | 24.44512248 | 25.00812766 | 24.53534326 |
| Structural maintenance of chromosor | A6NHR9-2;A6NHR9;A6NHR9- | SMCHD1 | Other | -0.715896158 | 0.954472348 | 27.5597589 | 27.64787361 | 27.80279618 | 27.95947457 | 27.93882014 | 28.19295834 | 27.45544412 | 27.36753046 | 26.58017781 |
| Serine/threonine-protein phosphatas | Q6IN85;Q6IN85-2;Q6IN85-5;( | SMEK1 | Other | 0.115011658 | 0.12020244 | 28.07948576 | 28.13138812 | 28.02120442 | 27.31691332 | 27.223522 | 27.36527918 | 28.2341618 | 27.94670481 | 27.23306476 |
| Serine/threonine-protein phosphatas | Q5MIZ7-3;Q5MIZ7-2;Q5MIZ7 | SMEK2 | Other | 0.24360948 | 0.094334465 | 23.26083571 | 22.92255972 | 22.91252611 | 23.09879802 | 23.06892454 | 22.71571655 | 24.27497639 | 23.92034413 | 21.52518824 |
| Serine/threonine-protein kinase SMG | Q96Q15-3;Q96Q15-2;Q96Q1 | SMG1 | Other | -0.040930538 | 0.064964166 | 22.94345775 | 23.27696494 | 23.54498235 | 22.90628142 | 23.34489185 | 22.19622509 | 23.09998597 | 23.1142132 | 22.76941092 |
| Protein SMG8 | Q8ND04;Q8ND04-2;Q8ND04- | SMG8 | Other | -0.155110777 | 0.174148888 | 22.61103741 | 22.51667533 | 22.53478541 | 22.49655637 | 22.54769097 | 20.76714072 | 21.73537422 | 22.34913758 | 22.18709898 |
| Protein SMG9 | Q9H0W8-2;Q9H0W8 | SMG9 | Other | 0.443463958 | 0.953912752 | 22.91402374 | 22.78952967 | 23.83142091 | 23.29670116 | 23.40496586 | 23.39963049 | 23.95898806 | 23.80656221 | 23.38297752 |
| Survival of motor neuron-related-splic | O75940 | SMNDC1 | Other | 2.624695448 | 2.631873593 | 21.57634995 | 23.05731884 | 23.4474576 | 21.83594618 | 24.25803401 | 24.52933809 | 25.44199254 | 25.58431318 | 26.20000296 |
| SPARC-related modular calcium-bindi | Q9H4F8;Q9H4F8-2 | SMOC1 | Extracellular or secreted | 2.753021552 | 1.373206251 | 25.02723986 | 24.76248302 | 25.21941021 | 27.24081753 | 27.21428636 | 27.33193572 | 28.14554762 | 28.09179297 | 30.41981041 |
| Sphingomyelin phosphodiesterase 4 | Q9NXE4-8;Q9NXE4-6;Q9NXE4 | SMPD4 | Other | -3.331726102 | 1.729485265 | 26.44561191 | 25.67648311 | 26.97661831 | 26.07060824 | 21.53335591 | 25.65512713 | 22.6933219 | 22.82257939 | 20.66782271 |
| Spermine synthase | P52788;P52788-2 | SMS | Extracellular or secreted | 0.142335769 | 0.302217807 | 30.5042688 | 30.52971414 | 30.51474538 | 30.29340271 | 30.42532397 | 29.85838343 | 30.67657054 | 30.63374125 | 30.17961472 |
| Smoothelin | P53814-5;P53814;P53814-6;F | SMTN | Other | -3.39617212 | 0.880520022 | 26.55337137 | 26.97967394 | 27.08618418 | 26.46174858 | 27.39933713 | 27.61421209 | 23.31437414 | 21.38413979 | 26.16023336 |
| WD40 repeat-containing protein SMU | Q2TAY7;Q2TAY7-2 | SMU1 | Other | -0.811704712 | 0.864149601 | 28.02464275 | 28.0753089 | 28.00485326 | 27.86183281 | 27.96708251 | 27.5122417 | 27.53148141 | 27.34451942 | 26.41186601 |
| N-lysine methyltransferase SMYD2 | Q9NRG4;Q9NRG4-2 | SMYD2 | Other | 1.436434121 | 1.317169539 | 21.67504045 | 21.94966858 | 21.60792626 | 22.52704836 | 21.5313546 | 21.23847861 | 23.67831129 | 23.48505195 | 22.41069755 |
| SET and MYND domain-containing pro | Q6GMV2 | SMYD5 | Other | 0.58975859 | 0.846801241 | 27.44071015 | 27.47622396 | 27.42542402 | 26.6367171 | 27.20499121 | 26.82359523 | 28.08425903 | 27.96366941 | 27.22517816 |
| Synaptosomal-associated protein 23 | O00161;O00161-2 | SNAP23 | Extracellular or secreted | 0.234864793 | 0.106432342 | 24.74811708 | 23.99576163 | 24.32889238 | 24.32443549 | 24.65883558 | 24.56639414 | 24.6889119 | 25.94775285 | 23.37914777 |
| Synaptosomal-associated protein 29 | O95721 | SNAP29 | Extracellular or secreted | 3.103395519 | 1.805617917 | 23.03360515 | 21.88676976 | 22.25755934 | 23.45035547 | 22.29731709 | 23.12218343 | 25.47481697 | 26.83857938 | 25.02068532 |
| SNARE-associated protein Snapin | O95295 | SNAPIN | Other | 0.231751325 | 1.009572683 | 23.85205244 | 23.41118718 | 23.68110284 | 23.3095252 | 23.10157372 | 23.53278965 | 23.79808096 | 23.71617724 | 23.62511128 |
| Staphylococcal nuclease domain-cont | Q7KZF4 | SND1 | Extracellular or secreted | -0.103302688 | 0.632612889 | 31.72525315 | 31.7997352 | 31.74286298 | 31.65667916 | 31.71046705 | 31.44588812 | 31.66636285 | 31.5938809 | 31.47029102 |
| Vacuolar-sorting protein SNF8 | Q96H20-2;Q96H20 | SNF8 | Extracellular or secreted | 0.291222151 | 0.370466606 | 23.32347373 | 23.21701518 | 22.82735589 | 23.36594396 | 21.33284034 | 23.42866993 | 23.16214051 | 23.37464472 | 23.08453074 |
| U5 small nuclear ribonucleoprotein 2( | O75643 | SNRNP200 | Other | -0.194442062 | 0.529000373 | 31.08815732 | 31.16206942 | 31.13858289 | 30.55581572 | 30.61981888 | 30.91070791 | 30.86090096 | 30.82631056 | 30.46703837 |
| U5 small nuclear ribonucleoprotein 4( | Q96DI7;Q96DI7-2 | SNRNP40 | Other | -0.136509395 | 0.56953246 | 27.34028043 | 27.54347864 | 27.4778457 | 27.15624559 | 27.38195505 | 27.88965723 | 27.32199551 | 27.24090849 | 27.42229913 |
| U1 small nuclear ribonucleoprotein 7( | P08621;P08621-2;P08621-3 | SNRNP70 | Other | 2.018924236 | 3.788163638 | 28.51833331 | 28.28850466 | 28.25525388 | 28.59843166 | 28.54325737 | 29.83791805 | 30.72647175 | 30.45012338 | 30.90102704 |
| U1 small nuclear ribonucleoprotein A | P09012 | SNRPA | Other | 0.56197175 | 2.409021067 | 29.66232739 | 30.11817807 | 30.03524979 | 30.24263567 | 29.91056497 | 30.31861655 | 30.49781292 | 30.56001411 | 30.77187443 |
| U2 small nuclear ribonucleoprotein A | P09661 | SNRPA1 | Other | 1.148940385 | 5.585039203 | 28.06840592 | 27.98695947 | 28.14860536 | 28.28387689 | 28.30573865 | 28.26700359 | 29.32457283 | 29.40865895 | 29.24388432 |
| Small nuclear ribonucleoprotein-asso | P14678-2;P63162;P14678;P6 | SNRPB;SNRPN | Extracellular or secreted | -0.082772894 | 0.361128925 | 30.11879691 | 30.44282706 | 30.35751875 | 30.23089108 | 30.16482873 | 29.89426218 | 30.12065183 | 30.0115866 | 30.22400524 |
| U2 small nuclear ribonucleoprotein B | P08579 | SNRPB2 | Other | 0.528939141 | 0.958888192 | 27.13065194 | 27.48738472 | 27.65123119 | 28.3382838 | 27.90190458 | 27.25115038 | 28.26239584 | 27.75970505 | 28.44501984 |
| U1 small nuclear ribonucleoprotein C | P09234 | SNRPC | Other | 1.557435473 | 1.659206608 | 26.29810061 | 24.99031798 | 25.71850466 | 25.89307614 | 25.50172787 | 27.98391925 | 27.13692198 | 27.51480181 | 28.21340589 |
| Small nuclear ribonucleoprotein Sm D | P62314 | SNRPD1 | Other | 0.213546339 | 0.1141149 | 30.86178845 | 31.10193182 | 31.2414996 | 30.42961994 | 30.38853895 | 29.12846572 | 31.4245233 | 31.22716533 | 29.56487262 |
| Small nuclear ribonucleoprotein Sm D | P62316;P62316-2 | SNRPD2 | Extracellular or secreted | 0.269775693 | 0.955179877 | 31.24756719 | 31.22319975 | 31.33681642 | 30.64022141 | 30.56746804 | 31.01285227 | 31.21552506 | 31.21829958 | 31.38956498 |
| Small nuclear ribonucleoprotein Sm D | P62318-2;P62318 | SNRPD3 | Extracellular or secreted | -0.119570052 | 0.127867678 | 30.9664363 | 30.88867061 | 30.91840611 | 30.88816253 | 30.87562172 | 29.83261646 | 30.86695465 | 30.92811267 | 30.03117938 |
| Small nuclear ribonucleoprotein E | P62304 | SNRPE | Extracellular or secreted | -0.323308483 | 2.119797756 | 29.20764644 | 29.13672645 | 29.08626012 | 29.30619512 | 29.35386465 | 28.87903018 | 28.74294326 | 28.85355823 | 28.91843454 |
| Small nuclear ribonucleoprotein F | P62306 | SNRPF | Other | 1.160723957 | 4.550753687 | 28.8842807 | 29.11152741 | 29.10931199 | 28.70869235 | 28.45603193 | 28.99509848 | 29.95240965 | 30.14479433 | 30.01743932 |
| Small nuclear ribonucleoprotein G;Pu | P62308;A8MWD9 | SNRPG;SNRPGP1 | Other | 0.333490799 | 1.035255347 | 28.39464331 | 28.57268103 | 28.27595326 | 27.82207606 | 27.62407946 | 28.27226435 | 28.6514365 | 28.47591485 | 28.35396978 |
| Beta-2-syntrophin | Q13425;Q13425-2 | SNTB2 | Extracellular or secreted | 0.483211796 | 1.204753087 | 24.31125884 | 24.35843712 | 24.39917412 | 23.9850275 | 24.14647048 | 23.89374739 | 24.92048043 | 24.39296616 | 24.68324652 |
| Snurportin-1 | O95149 | SNUPN | Extracellular or secreted | 2.02516452 | 1.647573683 | 21.88756062 | 24.152827 | 21.50550793 | 23.41866898 | 23.15407949 | 23.03321663 | 25.46970317 | 25.22769555 | 23.95402516 |
| SNW domain-containing protein 1 | Q13573 | SNW1 | Other | 0.230838796 | 0.627219751 | 26.642375 | 26.51150294 | 26.4932002 | 26.30547774 | 26.1998158 | 26.86124123 | 26.46270046 | 26.84888097 | 26.88774141 |
| Sorting nexin-1 | Q13596;Q13596-2;Q13596-3 | SNX1 | Other | 0.026083102 | 0.183553635 | 27.53200188 | 27.54605758 | 27.42886158 | 27.40786924 | 27.72968391 | 27.46816549 | 27.56848315 | 27.4867712 | 27.57931479 |
| Sorting nexin-12 | Q9UMY4-2;Q9UMY4;Q9UMY | SNX12 | Extracellular or secreted | -0.162634924 | 0.224774527 | 27.34163826 | 27.44134357 | 27.35077034 | 27.69955434 | 27.48907056 | 27.25069873 | 27.39982604 | 27.62992598 | 26.76888112 |
| Sorting nexin-15 | Q9NRS6-2;Q9NRS6 | SNX15 | Other | 0.257847158 | 1.024257093 | 25.72463209 | 25.6978987 | 25.58803634 | 25.78324266 | 25.7394644 | 25.60314502 | 25.91428841 | 25.80626719 | 26.12119549 |
| Sorting nexin-16 | P57768-2;P57768 | SNX16 | Other | -0.175066352 | 0.096575623 | 21.74579084 | 22.60494408 | 22.53805818 | 23.21954315 | 23.66492291 | 20.84670526 | 22.91533746 | 22.60645853 | 21.26298717 |
| Sorting nexin-17 | Q15036-2;Q15036 | SNX17 | Other | 1.677719534 | 1.778165218 | 21.68883581 | 20.81010465 | 21.23828635 | 22.06989157 | 21.84861836 | 23.21437901 | 23.93012515 | 23.75674362 | 22.78134771 |
| Sorting nexin-2 | O60749-2;O60749 | SNX2 | Extracellular or secreted | -0.218071512 | 0.399853302 | 26.91285042 | 26.95548922 | 26.84493673 | 27.30129677 | 27.41691489 | 26.84959695 | 27.09123812 | 26.95582175 | 26.43926808 |
| Sorting nexin-27 | Q96L92-3;Q96L92;Q96L92-2 | SNX27 | Extracellular or secreted | -0.044006171 | 0.08472752 | 27.98706794 | 28.08420834 | 28.0210985 | 27.87220518 | 27.72299197 | 27.75779925 | 28.04941942 | 28.01073331 | 27.53051433 |
| Sorting nexin-3 | O60493;O60493-2;O60493-4 | SNX3 | Extracellular or secreted | 0.464792231 | 0.577728201 | 24.08269676 | 23.73912958 | 24.07030114 | 24.69147006 | 25.66936347 | 23.37729527 | 25.03002182 | 24.80503729 | 24.37444573 |
| Sorting nexin-30 | Q5VWJ9 | SNX30 | Other | 0.54718824 | 0.656597778 | 21.7831178 | 22.95861643 | 22.87145333 | 20.92227877 | 22.65265682 | 22.2219045 | 23.18879045 | 22.83335262 | 22.32443549 |
| Sorting nexin-4 | O95219;O95219-2 | SNX4 | Extracellular or secreted | 0.069300352 | 0.038723172 | 23.08326506 | 23.15344565 | 22.32409177 | 21.71799464 | 20.80098359 | 23.32484747 | 22.88270872 | 22.98274923 | 21.5447572 |
| Sorting nexin-5 | Q9Y5X3 | SNX5 | Extracellular or secreted | -0.147804895 | 0.55126619 | 26.35937585 | 26.41212453 | 26.46215438 | 26.39185244 | 26.5357437 | 26.46238844 | 26.39055747 | 26.39196713 | 26.08588038 |
| Sorting nexin-6;Sorting nexin-6, N-ter | Q9UNH7;Q9UNH7-2 | SNX6 | Other | -0.300605785 | 0.606719479 | 26.64938206 | 26.75824416 | 26.79104124 | 26.19578595 | 26.57150975 | 26.29410859 | 26.44142273 | 26.41929608 | 25.86749971 |
| Sorting nexin-8 | Q9Y5X2 | SNX8 | Other | 1.013122751 | 1.593914043 | 23.64427439 | 22.30777646 | 23.35904066 | 23.78506527 | 23.97342481 | 24.71486806 | 24.72411162 | 24.55952553 | 24.64795593 |
| Sorting nexin-9 | Q9Y5X1 | SNX9 | Extracellular or secreted | 0.383341499 | 1.101915521 | 28.36952866 | 28.77360457 | 28.56150798 | 28.50047472 | 28.37355824 | 27.85566845 | 28.92512972 | 28.90813269 | 28.5339334 |
| Sterol O-acyltransferase 1 | P35610;P35610-3;P35610-2 | SOAT1 | Other | 0.064669019 | 0.033257635 | 21.76094324 | 21.40691291 | 20.49351993 | 22.15517646 | 20.59803811 | 24.10269532 | 21.11286198 | 21.8284118 | 22.51137626 |
| Superoxide dismutase [Cu-Zn] | P00441 | SOD1 | Extracellular or secreted | -0.272927732 | 0.406527422 | 33.79100629 | 33.8243011 | 33.91197595 | 33.54846196 | 33.67535111 | 32.53834389 | 33.52950472 | 33.43477318 | 32.86165906 |
| Superoxide dismutase [Mn], mitochor | P04179;P04179-4;P04179-2;F | SOD2 | Extracellular or secreted | -0.831452884 | 1.648136323 | 30.52878313 | 30.63996276 | 30.50748593 | 31.09496666 | 31.28190006 | 29.84826912 | 30.18506154 | 29.65713933 | 29.61412431 |
| Protein SON | P18583-2;P18583-10;P18583 | SON | Other | -3.306645248 | 0.880809326 | 26.1306323 | 26.26056655 | 26.68110284 | 26.1147573 | 24.9656522 | 26.73298594 | 20.11371359 | 24.13657001 | 24.27262922 |
| Vinexin | O60504-2;O60504 | SORBS3 | Other | 0.06556492 | 0.124756365 | 26.52315823 | 26.11751438 | 26.18574686 | 26.02766172 | 25.86775891 | 26.37419696 | 26.49888584 | 26.30596473 | 25.93986272 |
| Sorbitol dehydrogenase | Q00796 | SORD | Extracellular or secreted | 0.100683216 | 0.06952753 | 30.96547331 | 31.04641707 | 30.97978295 | 30.52037688 | 30.30628205 | 29.61745617 | 31.05141753 | 31.17765595 | 29.79087039 |
| Sortilin-related receptor | Q92673 | SORL1 | Extracellular or secreted | 0.076413538 | 0.04901506 | 23.1677397 | 23.0788427 | 20.36929586 | 22.88310042 | 21.68177996 | 23.97000475 | 22.747943 | 22.31713766 | 22.73954166 |
| Sortilin | Q99523;Q99523-2 | SORT1 | Extracellular or secreted | -0.587343082 | 0.24048306 | 23.65567309 | 23.5162906 | 23.46238844 | 23.0163642 | 22.61108902 | 25.06249944 | 21.93015997 | 22.30276413 | 24.66719905 |
| Son of sevenless homolog 1 | Q07889 | SOS1 | Extracellular or secreted | 1.265967515 | 0.974910032 | 23.96710451 | 23.97421291 | 23.99636522 | 21.52353248 | 21.28180335 | 24.37364951 | 23.94824316 | 24.21734066 | 25.1906527 |
| Sclerostin domain-containing protein | Q6X4U4;Q6X4U4-2 | SOSTDC1 | Extracellular or secreted | -1.867975603 | 1.036614292 | 21.43714431 | 21.18392367 | 21.53673099 | 25.32378294 | 25.4607336 | 25.2879237 | 21.50649355 | 21.07084159 | 21.93385766 |
| Transcription factor SOX-9 | P48436 | SOX9 | Extracellular or secreted | 1.567850922 | 2.125101611 | 25.78254297 | 24.9272792 | 24.96013772 | 25.93898835 | 25.8316142 | 23.95278176 | 27.26660156 | 26.77743493 | 26.35618839 |
| Transcription factor Sp1 | P08047-3;P08047;P08047-2 | SP1 | Other | -0.625719055 | 0.661654044 | 24.30047657 | 24.63682771 | 24.74026252 | 24.70392934 | 24.86679256 | 24.06693926 | 24.21992713 | 24.3515119 | 23.20901778 |
| Transcription factor Sp3 | Q02447-4;Q02447-3;Q02447 | SP3 | Extracellular or secreted | 0.565956832 | 0.630917191 | 24.12099782 | 24.22920021 | 24.25335239 | 21.95743713 | 23.5712638 | 22.31892935 | 24.04735094 | 23.88371573 | 23.99239418 |
| Sperm-associated antigen 7 | O75391 | SPAG7 | Other | 0.31886734 | 0.336994883 | 26.84541539 | 26.86242416 | 26.84481704 | 26.52468327 | 26.76509121 | 25.95850139 | 27.19460298 | 27.38747111 | 26.27499416 |
| C-Jun-amino-terminal kinase-interacti | O60271-4;O60271-2;O60271 | SPAG9 | Extracellular or secreted | 0.259600221 | 1.397256148 | 27.60066622 | 27.75429872 | 27.68825831 | 27.32251134 | 27.5132211 | 27.31492726 | 27.90529463 | 27.79606427 | 27.67438324 |
| SPARC | P09486 | SPARC | Extracellular or secreted | -0.063650052 | 0.026665341 | 23.78706001 | 22.45033029 | 23.71722374 | 25.12257842 | 25.49628207 | 26.65444439 | 23.99507151 | 23.87408307 | 25.55385472 |
| Spastin | Q9UBP0-4;Q9UBP0-3;Q9UBP | SPAST | Extracellular or secreted | -0.5716365 | 1.709785317 | 23.27298509 | 23.52229034 | 23.43504452 | 23.2454529 | 23.06877697 | 23.5143654 | 22.86368645 | 22.53271833 | 22.91814333 |
| Spermatogenesis-associated protein 3 | Q96N06;Q96N06-2 | SPATA33 | Other | 0.251707998 | 0.228159063 | 21.26665971 | 20.62390649 | 21.42207149 | 21.85551143 | 22.92767218 | 21.71693518 | 22.13097394 | 22.26570775 | 21.26482054 |
| Spermatogenesis-associated protein 5 | Q8NB90-2;Q8NB90;Q8NB90- | SPATA5 | Other | -2.054433986 | 0.917024612 | 22.32790434 | 22.22873254 | 23.99296848 | 21.40052422 | 21.27293374 | 22.67195358 | 19.89671377 | 18.99203106 | 21.89546166 |
| Kinetochore protein Spc24 | Q8NBT2;Q8NBT2-2 | SPC24 | Other | 1.74375849 | 1.308927679 | 24.32560246 | 23.80242643 | 24.10069185 | 21.33137507 | 20.97372013 | 21.00774903 | 23.92795732 | 24.306608 | 24.76749264 |
| Protein Spindly | Q96EA4-3;Q96EA4;Q96EA4-2 | SPDL1 | Other | -0.453873436 | 0.160436264 | 24.4230369 | 23.56767112 | 23.79382157 | 24.66741564 | 24.21660082 | 23.9738627 | 23.77926488 | 25.28753626 | 21.89278292 |
| Cytospin-B | Q5M775-5;Q5M775-3;Q5M7 | SPECC1 | Other | -0.12940661 | 0.144179947 | 24.7321579 | 24.71103843 | 25.12206491 | 25.32570539 | 25.31900004 | 24.54132353 | 24.56412762 | 24.50784883 | 25.41544882 |
| Cytospin-A | Q69YQ0-2;Q69YQ0 | SPECC1L | Other | 0.730974843 | 1.376974053 | 24.07545173 | 24.64251272 | 23.9709702 | 24.76819967 | 25.37937916 | 23.53670728 | 25.01090401 | 25.23480135 | 25.13382955 |
| Msx2-interacting protein | Q96T58 | SPEN | Extracellular or secreted | -0.382164312 | 0.704311876 | 23.55180306 | 23.1371644 | 24.22875999 | 23.77001616 | 23.92714354 | 23.04458556 | 23.21396389 | 22.91630373 | 23.55297579 |
| Spartin | Q8N0X7 | SPG20 | Extracellular or secreted | -1.008643564 | 0.787703824 | 22.56132151 | 22.19955374 | 22.19385138 | 21.48583911 | 21.92538783 | 21.82108432 | 20.08843069 | 21.70691415 | 21.2722434 |
| Spindlin-1 | Q9Y657 | SPIN1 | Other | -0.886512369 | 2.182793806 | 26.86088616 | 26.89284461 | 26.81665479 | 26.09214595 | 25.88103347 | 25.87342609 | 25.69771315 | 25.36312467 | 25.48812061 |
| Kunitz-type protease inhibitor 1 | O43278 | SPINT1 | Extracellular or secreted | 0.602307327 | 0.731387878 | 29.52056422 | 29.51145016 | 29.34904138 | 29.71146846 | 29.83445615 | 29.74648744 | 29.88672442 | 29.92351551 | 30.83341596 |
| Kunitz-type protease inhibitor 1 | O43278-2 | SPINT1 | Extracellular or secreted | -0.769201253 | 0.3238287 | 23.50348803 | 23.20685876 | 23.30938642 | 24.80183464 | 20.80024793 | 22.38785258 | 20.72229056 | 22.49357892 | 23.48136094 |
| Kunitz-type protease inhibitor 2 | O43291;O43291-2 | SPINT2 | Extracellular or secreted | 1.555047884 | 1.252178877 | 21.36168483 | 23.75776111 | 24.06587166 | 24.94436069 | 25.27639707 | 25.51977349 | 25.52288894 | 26.10199442 | 25.50318471 |
| Sepiapterin reductase | P35270 | SPR | Extracellular or secreted | -0.363744292 | 0.38825039 | 25.71109096 | 25.86422035 | 25.96721447 | 26.08119377 | 25.64094193 | 25.41773588 | 25.75631097 | 25.77580356 | 24.71785127 |
| SPRY domain-containing protein 4 | Q8WW59 | SPRYD4 | Other | 0.202237858 | 0.163918979 | 25.2727004 | 24.85495548 | 25.24278102 | 24.40008672 | 24.23717436 | 24.19061503 | 25.18914499 | 25.4171564 | 24.09956869 |
| SPRY domain-containing protein 7 | Q5W111 | SPRYD7 | Other | 1.138770181 | 0.903911325 | 22.55667537 | 22.56230266 | 20.96568473 | 21.31569467 | 21.59488742 | 22.99870818 | 23.13544314 | 22.30100713 | 23.97683678 |
| Spectrin alpha chain, non-erythrocytic | Q13813;Q13813-3 | SPTAN1 | Extracellular or secreted | 0.267973821 | 1.47165046 | 27.18886212 | 27.48669449 | 27.44978895 | 27.08790451 | 27.15344565 | 27.2115041 | 27.39419353 | 27.59370284 | 27.60512499 |
| Spectrin alpha chain, non-erythrocytic | Q13813-2 | SPTAN1 | Extracellular or secreted | -0.061694226 | 0.289119579 | 33.89599969 | 33.87096413 | 33.80385868 | 33.67682271 | 33.76890084 | 33.32696882 | 33.65288141 | 33.62723187 | 33.70656148 |
| Spectrin beta chain, non-erythrocytic | Q01082;Q01082-2 | SPTBN1 | Extracellular or secreted | 0.191347863 | 0.663641502 | 33.46878697 | 33.48014969 | 33.41752363 | 32.83872213 | 32.91783728 | 32.84946273 | 33.36541478 | 33.41928301 | 33.27558701 |
| Spectrin beta chain, non-erythrocytic | Q01082-3 | SPTBN1 | Extracellular or secreted | 0.202016338 | 0.050720244 | 21.11583901 | 24.56389496 | 24.46313718 | 23.35076468 | 24.51881353 | 22.94100719 | 25.12226244 | 21.27539522 | 24.68511962 |
| Spectrin beta chain, non-erythrocytic | O15020;O15020-2 | SPTBN2 | Extracellular or secreted | -0.73214924 | 1.881039716 | 29.04419052 | 29.01278568 | 28.89044024 | 27.90242222 | 28.15397709 | 28.37608656 | 27.9264311 | 27.78375472 | 27.78331761 |
| Serine palmitoyltransferase 1 | O15269 | SPTLC1 | Other | 0.167553741 | 0.093729162 | 21.2253334 | 22.41296021 | 22.42370377 | 20.72083188 | 21.04878864 | 24.79233277 | 22.5774033 | 21.82224872 | 22.41498453 |
| Sulfide:quinone oxidoreductase, mito | Q9Y6N5 | SQRDL | Other | 0.699662375 | 0.20580971 | 21.99345265 | 22.80185437 | 22.80329396 | 21.00641649 | 22.61172583 | 27.27355432 | 23.16974399 | 22.43402648 | 25.74036547 |
| Sequestosome-1 | Q13501;Q13501-2 | SQSTM1 | Extracellular or secreted | 0.960536322 | 2.730351807 | 26.8364139 | 26.61133014 | 26.67868739 | 27.33176491 | 27.50370032 | 26.55786168 | 27.87255747 | 27.92405378 | 27.84487688 |
| Steroid receptor RNA activator 1 | Q9HD15 | SRA1 | Extracellular or secreted | 0.886755448 | 0.713768901 | 22.97326714 | 19.95755988 | 23.32443549 | 23.63444771 | 23.70038807 | 23.40197528 | 23.51833331 | 23.69338571 | 23.94458411 |
| S1 RNA-binding domain-containing pr | Q8N5C6;Q8N5C6-2 | SRBD1 | Other | -0.840518139 | 0.691896608 | 22.53278965 | 22.43025346 | 22.79062372 | 22.15502201 | 21.6202049 | 21.95155508 | 22.33021229 | 20.86335285 | 21.02510486 |
| Proto-oncogene tyrosine-protein kina | P12931;P12931-2 | SRC | Extracellular or secreted | 0.988662844 | 2.194015299 | 24.69099076 | 24.17012617 | 24.60794882 | 24.53029107 | 24.27440773 | 25.12616785 | 25.5464845 | 25.84570252 | 25.27376772 |
| Splicing regulatory glutamine/lysine-r | Q8WXA9-2;Q8WXA9 | SREK1 | Other | -0.990174041 | 1.396866207 | 25.45331033 | 25.31513463 | 25.24067198 | 25.28104001 | 25.36312467 | 25.04697642 | 24.57502075 | 23.85290968 | 24.45167647 |
| SLIT-ROBO Rho GTPase-activating pro | Q7Z6B7-2;Q7Z6B7 | SRGAP1 | Other | 0.78659547 | 0.525937649 | 22.91917911 | 19.3732189 | 21.3929491 | 23.31035761 | 22.98099024 | 22.64495646 | 22.97042612 | 22.26158397 | 23.43860203 |
| SLIT-ROBO Rho GTPase-activating pro | O75044;P0DJJ0 | SRGAP2;SRGAP2( | Other | 0.549413567 | 1.148071756 | 25.0611401 | 24.91328425 | 24.41763931 | 23.75470649 | 23.87060704 | 24.21859753 | 25.12427564 | 24.98567902 | 24.6562734 |

| | | | | | | | | | | | | | |
|---|---|---|---|---|---|---|---|---|---|---|---|---|---|
| Sorcin | P30626-2;P30626;P30626-3 | SRI | Extracellular or secreted | 0.294797352 | 2.761837484 | 29.14797493 | 29.27208633 | 29.28275076 | 29.44496062 | 29.41912727 | 29.33524106 | 29.64244387 | 29.6547858 | 29.53823287 |
| Spermidine synthase | P19623 | SRM | Other | 0.60054569 | 0.785572092 | 27.12415729 | 27.3468033 | 27.2429082 | 27.51795803 | 27.52065788 | 26.42572813 | 28.10938672 | 28.07198937 | 27.20936739 |
| Signal recognition particle 14 kDa pro | P37108 | SRP14 | Extracellular or secreted | -0.101281666 | 0.099097868 | 28.8519512 | 28.94265602 | 28.83481778 | 28.67653692 | 28.56325494 | 29.36454885 | 29.25885174 | 28.92093468 | 28.13325143 |
| Signal recognition particle 19 kDa pro | P09132;P09132-2 | SRP19 | Other | 0.512992768 | 1.206695548 | 25.53599578 | 25.04314246 | 25.05486229 | 25.47351699 | 25.34871222 | 25.78294284 | 25.71111722 | 25.73262373 | 26.21482364 |
| Signal recognition particle 54 kDa pro | P61011-2;P61011 | SRP54 | Other | -0.354588514 | 1.159789121 | 24.46656393 | 24.39152471 | 24.29368774 | 24.50742544 | 24.2188931 | 24.92732442 | 23.96472724 | 24.03271435 | 24.34150254 |
| Signal recognition particle subunit SRI | Q9UHB9;Q9UHB9-4;Q9UHB9 | SRP68 | Other | 0.32970059 | 0.793832102 | 26.53590681 | 26.57873915 | 26.35630594 | 26.47010673 | 26.54395056 | 27.64402679 | 26.92138879 | 26.97213228 | 27.16009869 |
| Signal recognition particle subunit SRI | O76094;O76094-2 | SRP72 | Extracellular or secreted | 0.001782085 | 0.004915078 | 27.00442468 | 27.21474955 | 27.23999862 | 27.1125717 | 27.33193572 | 27.63221635 | 27.09557039 | 27.30678181 | 27.37094237 |
| Signal recognition particle 9 kDa prote | P49458 | SRP9 | Extracellular or secreted | 0.737269622 | 1.185689848 | 28.15320402 | 28.08962278 | 28.12504465 | 28.1199596 | 28.14972007 | 28.5322992 | 29.06108343 | 29.22699314 | 28.50865746 |
| SRSF protein kinase 1 | Q96SB4-4;Q96SB4;Q96SB4-3 | SRPK1 | Extracellular or secreted | -0.290895121 | 1.208283739 | 27.46513192 | 27.18839054 | 27.22324579 | 27.86431482 | 27.56870058 | 27.65982924 | 27.05185829 | 27.28982416 | 27.27043863 |
| SRSF protein kinase 2;SRSF protein kir | P78362;P78362-2 | SRPK2 | Other | -0.916973954 | 1.261496629 | 24.54968974 | 25.00444612 | 24.47283559 | 24.75908277 | 24.45663516 | 24.77001616 | 23.82045791 | 23.28446831 | 24.15050469 |
| Signal recognition particle receptor su | P08240;P08240-2 | SRPR | Extracellular or secreted | -1.619190562 | 0.751842897 | 22.04736758 | 23.91716008 | 23.85880134 | 23.52193106 | 23.61121756 | 24.35346508 | 21.26367496 | 20.9633872 | 23.57033751 |
| Signal recognition particle receptor su | Q9Y5M8 | SRPRB | Other | -0.080276157 | 0.034530207 | 21.99853415 | 22.73398929 | 22.90453687 | 21.96677017 | 20.7482233 | 25.39796729 | 21.98457561 | 22.11705457 | 23.53255188 |
| Sushi repeat-containing protein SRPX | P78539;P78539-4;P78539-3;F | SRPX | Extracellular or secreted | 2.661498337 | 1.478129342 | 20.57453619 | 23.2157083 | 20.67829196 | 24.77339077 | 25.46142074 | 25.3261513 | 25.70242404 | 26.08535364 | 26.21146696 |
| Serine racemase | Q9GZT4 | SRR | Extracellular or secreted | -0.432030621 | 0.600273668 | 22.58634779 | 22.25885353 | 22.07861471 | 23.28304982 | 23.07934885 | 22.10380022 | 22.39322809 | 22.38345223 | 21.62223528 |
| Serine/arginine repetitive matrix prot | Q8IYB3-2;Q8IYB3 | SRRM1 | Other | -1.314804628 | 1.65489855 | 27.84517608 | 27.89902541 | 27.77020525 | 27.74156172 | 27.33568829 | 28.39374362 | 26.22493905 | 26.26033324 | 27.06301401 |
| Serine/arginine repetitive matrix prot | Q9UQ35;Q9UQ35-2 | SRRM2 | Other | -0.298218311 | 1.109786777 | 29.46480095 | 29.38769706 | 29.36220929 | 29.55862817 | 29.606061 | 30.07851903 | 29.10051642 | 29.32096329 | 29.4128231 |
| Serrate RNA effector molecule homol | Q9BXP5-2;Q9BXP5;Q9BXP5-4 | SRRT | Other | 0.084598261 | 0.126992019 | 28.46637723 | 28.37840375 | 28.49132803 | 28.95551693 | 29.03878564 | 29.14816894 | 28.58448523 | 28.67841875 | 29.23018107 |
| Serine/arginine-rich splicing factor 1 | Q07955;Q07955-2 | SRSF1 | Extracellular or secreted | 0.584471532 | 0.472998707 | 30.57172311 | 30.44025471 | 30.53889888 | 30.35216045 | 30.4504184 | 31.09697809 | 30.81184084 | 30.58262026 | 32.08417031 |
| Serine/arginine-rich splicing factor 10 | O75494-5;O75494-4;O75494 | SRSF10 | Other | -0.595081251 | 0.203267314 | 27.20275144 | 26.88959921 | 26.93988513 | 26.11705854 | 26.43882408 | 26.58419847 | 24.44278357 | 25.84244511 | 28.015686 |
| Serine/arginine-rich splicing factor 11 | Q05519-2;Q05519 | SRSF11 | Other | -0.712753391 | 1.859202115 | 27.51058274 | 27.53207621 | 27.44939541 | 27.57801928 | 27.5363071 | 27.79921798 | 27.0431216 | 26.87876715 | 26.64265043 |
| Serine/arginine-rich splicing factor 2 | Q01130-2;Q01130 | SRSF2 | Extracellular or secreted | 0.50950289 | 0.319527504 | 29.06655954 | 29.07017316 | 29.17376614 | 29.72668287 | 29.72888964 | 30.01398377 | 29.4056598 | 29.37088003 | 31.1419964 |
| Serine/arginine-rich splicing factor 3 | P84103-2;P84103 | SRSF3 | Other | 0.13558526 | 0.065772385 | 28.54126444 | 28.46248205 | 28.62348653 | 28.65935291 | 28.76135472 | 30.397849 | 28.26297821 | 28.56659737 | 30.30007503 |
| Serine/arginine-rich splicing factor 4 | Q08170 | SRSF4 | Other | 0.382953595 | 0.213387105 | 25.00979411 | 24.97657461 | 24.91615792 | 25.63167532 | 25.56590046 | 25.12872628 | 24.89508893 | 25.00611682 | 26.86206938 |
| Serine/arginine-rich splicing factor 5 | Q13243;Q13243-2 | SRSF5 | Other | 1.102316554 | 0.248851961 | 23.44972599 | 24.02892654 | 24.18767345 | 24.39434075 | 24.39204904 | 27.27852546 | 25.44126441 | 23.15364665 | 28.57765921 |
| Serine/arginine-rich splicing factor 6 | Q13247-3;Q13247;Q13247-2 | SRSF6 | Other | 0.633532373 | 0.556020069 | 28.96919788 | 28.88675349 | 28.81881965 | 28.43440037 | 28.59583806 | 29.81306495 | 29.23778979 | 29.01454253 | 30.407302 |
| Serine/arginine-rich splicing factor 7 | Q16629-3;Q16629-2;Q16629 | SRSF7 | Extracellular or secreted | 0.041525784 | 0.01532564 | 28.4169954 | 28.53552866 | 28.62114729 | 28.62397484 | 28.59405893 | 29.71158333 | 28.05310209 | 27.86720511 | 30.45591439 |
| Serine/arginine-rich splicing factor 9 | Q13242 | SRSF9 | Other | 0.52535625 | 0.960367495 | 27.03862868 | 27.00109889 | 26.98141712 | 26.98652555 | 27.08810676 | 26.60696111 | 27.53185319 | 27.09908707 | 27.79649754 |
| Sulfiredoxin-1 | Q9BYN0 | SRXN1 | Other | -1.094868569 | 0.440031119 | 29.17612269 | 29.07940946 | 28.46025709 | 28.93809101 | 28.46392449 | 30.01371761 | 29.37368269 | 28.23530366 | 26.17216912 |
| Lupus La protein | P05455 | SSB | Other | 0.32264094 | 0.431082668 | 31.89983936 | 31.94174697 | 31.93919158 | 31.7165714 | 31.76754 | 31.06445382 | 32.35429039 | 32.28377756 | 31.49452644 |
| Single-stranded DNA-binding protein, | Q04837 | SSBP1 | Extracellular or secreted | -0.465203922 | 0.693478821 | 31.7326302 | 31.70111718 | 31.54830118 | 32.09071459 | 32.18695128 | 31.00798336 | 31.59303494 | 31.38843631 | 30.75676588 |
| Protein phosphatase Slingshot homol | Q8TE77;Q8TE77-3;Q8TE77-2 | SSH3 | Other | 1.231373843 | 0.586634516 | 25.73277897 | 25.52596783 | 25.29274036 | 24.07005541 | 23.71250853 | 24.82230708 | 27.0115866 | 26.76293916 | 24.49777486 |
| Translocon-associated protein subuni | P43307-2;P43307 | SSR1 | Other | -1.01875667 | 0.398602128 | 20.95812411 | 21.7781788 | 23.12120339 | 24.16292417 | 23.32498477 | 25.70725127 | 21.70391456 | 20.95428642 | 23.81186227 |
| Translocon-associated protein subuni | P51571 | SSR4 | Extracellular or secreted | -1.005023987 | 0.423542277 | 22.70248217 | 23.25753057 | 23.73799575 | 24.75943839 | 23.6896585 | 26.77630571 | 22.35145798 | 22.29398235 | 24.80119326 |
| FACT complex subunit SSRP1 | Q08945 | SSRP1 | Other | 0.126833466 | 0.209561769 | 31.37710079 | 31.25479225 | 31.31581269 | 30.83243538 | 30.85749383 | 30.94761074 | 31.4276234 | 31.43136464 | 30.8141352 |
| Sjoegren syndrome/scleroderma auto | O60232 | SSSCA1 | Other | 1.383530879 | 1.327327044 | 26.44903325 | 26.38030437 | 26.36178693 | 26.39822831 | 26.54108716 | 26.41184985 | 28.15923264 | 28.08253435 | 27.17997058 |
| RNA polymerase II subunit A C-termin | Q9NP77 | SSU72 | Other | 0.261964829 | 0.48566353 | 24.85257637 | 25.05212787 | 24.77786129 | 25.58523054 | 26.15143406 | 24.6896585 | 25.30552992 | 25.47233982 | 25.56246907 |
| Hsc70-interacting protein;Putative pr | P50502;Q8IZP2;Q8NFI4 | ST13;ST13P4;ST1 | Extracellular or secreted | -0.169971938 | 0.581483363 | 31.57375568 | 31.56547198 | 31.52645297 | 31.7774898 | 31.8138295 | 30.98868033 | 31.49319065 | 31.36708257 | 31.25265109 |
| Suppressor of tumorigenicity 14 prote | Q9Y5Y6 | ST14 | Extracellular or secreted | 0.307583074 | 0.493870646 | 27.60349878 | 27.50286615 | 27.49197784 | 27.61855925 | 27.27949992 | 28.58509441 | 27.63913477 | 27.96493651 | 28.35942612 |
| Cohesin subunit SA-1 | Q8WVM7-2;Q8WVM7 | STAG1 | Other | -0.06362878 | 0.030765735 | 26.22376134 | 26.5073347 | 26.26047682 | 24.96411027 | 25.40584236 | 25.6053511 | 26.42564811 | 26.36074932 | 24.50615453 |
| Cohesin subunit SA-2 | Q8N3U4-2;Q8N3U4 | STAG2 | Extracellular or secreted | -0.35566512 | 0.572071494 | 29.29548429 | 29.36915421 | 29.31954279 | 28.31842262 | 28.37459498 | 28.79736369 | 28.80433578 | 28.72556196 | 28.14038818 |
| Signal transducing adapter molecule 1 | Q92783-2;Q92783 | STAM | Other | 0.037423809 | 0.178434826 | 26.97136498 | 27.00238721 | 26.96631252 | 27.25250447 | 27.17597998 | 26.78618765 | 26.97158425 | 27.12218343 | 27.09587215 |
| Signal transducing adapter molecule 2 | O75886;O75886-2 | STAM2 | Other | 1.230677517 | 1.121098041 | 24.13202584 | 24.31782771 | 23.80074905 | 20.96459456 | 22.15641685 | 22.54247128 | 24.37517522 | 24.47611578 | 23.7977842 |
| STAM-binding protein | O95630;O95630-2 | STAMBP | Extracellular or secreted | -0.044809729 | 0.063408388 | 26.52880176 | 26.89896777 | 26.75722701 | 26.65621884 | 26.49719621 | 26.30248261 | 26.83918033 | 26.72325243 | 26.12358515 |
| PCTP-like protein | Q9Y365 | STARD10 | Other | -0.99123467 | 3.227211264 | 25.47187487 | 25.36872971 | 25.7518752 | 25.1277428 | 25.01767294 | 25.32340501 | 24.55625438 | 24.22479189 | 24.27589999 |
| Signal transducer and activator of tra | P42224;P42224-2 | STAT1 | Other | 0.303503155 | 0.312291494 | 26.86679256 | 26.78456616 | 26.93999719 | 26.51618538 | 26.49414682 | 26.58591818 | 27.41530375 | 27.29911437 | 26.2898945 |
| Signal transducer and activator of tra | P40763-3;P40763;P40763-2 | STAT3 | Extracellular or secreted | 0.631235494 | 0.909184772 | 27.4326892 | 27.63242439 | 27.6063258 | 26.64498945 | 26.95814734 | 27.57022169 | 28.28135844 | 28.1242066 | 27.41054037 |
| Signal transducer and activator of tra | P51692 | STAT5B | Other | -0.287033711 | 0.252267834 | 24.1973055 | 23.81860636 | 23.72837395 | 23.3321065 | 24.22236156 | 23.57172672 | 24.33278943 | 23.14085036 | 23.10049937 |
| Double-stranded RNA-binding protein | O95793-2;O95793;O95793-3 | STAU1 | Extracellular or secreted | 0.482116645 | 1.917510623 | 25.99920729 | 25.90662549 | 25.96276524 | 26.24027164 | 26.00893976 | 26.73569959 | 26.68123691 | 26.50327571 | 26.68859182 |
| Starch-binding domain-containing pro | O95210 | STBD1 | Extracellular or secreted | -0.636483737 | 3.038423918 | 22.86710375 | 23.06574021 | 22.92547838 | 22.85689326 | 23.04677037 | 22.85521219 | 22.18425671 | 22.37254734 | 22.34234383 |
| Stanniocalcin-1 | P52823;P52823-2 | STC1 | Extracellular or secreted | -0.75411714 | 0.407932859 | 27.10539559 | 26.55417685 | 26.43245027 | 27.01531381 | 27.00870473 | 29.42830253 | 25.84050154 | 25.87703582 | 27.79228312 |
| Stanniocalcin-2 | O76061 | STC2 | Extracellular or secreted | -0.061984356 | 0.03663583 | 27.99048029 | 27.89232352 | 27.94391376 | 29.17350407 | 29.44112587 | 29.24361198 | 28.05631026 | 28.0824836 | 29.51773282 |
| Metalloreductase STEAP4 | Q687X5;Q687X5-2 | STEAP4 | Extracellular or secreted | 1.355161031 | 2.235247287 | 22.81876476 | 22.05391948 | 21.52171177 | 21.91301211 | 21.9166248 | 23.31423582 | 23.31105091 | 23.51749255 | 24.00607401 |
| Stress-induced-phosphoprotein 1 | P31948;P31948-2;P31948-3 | STIP1 | Other | 0.078678062 | 0.235232466 | 32.82464253 | 32.72805439 | 32.71454631 | 33.17541197 | 33.24516022 | 32.49897807 | 32.99809508 | 33.02349976 | 32.80783609 |
| Serine/threonine-protein kinase 10 | O94804 | STK10 | Extracellular or secreted | 0.859389515 | 1.929548271 | 22.56923658 | 21.73540398 | 21.29459355 | 21.51825799 | 21.63302891 | 21.49701952 | 22.88879601 | 22.44821413 | 22.36492867 |
| Serine/threonine-protein kinase 24;Se | Q9Y6E0-2;Q9Y6E0 | STK24 | Extracellular or secreted | 0.379178558 | 1.176059521 | 26.58892275 | 26.46450886 | 26.32070512 | 26.78818084 | 26.52911464 | 25.95006922 | 26.92150229 | 26.95560007 | 26.58118403 |
| Serine/threonine-protein kinase 26 | Q9P289;Q9P289-2;Q9P289-3 | STK26 | Extracellular or secreted | 0.229466734 | 1.157912517 | 25.39568134 | 25.22659358 | 25.67028237 | 25.82648332 | 25.74036547 | 25.79864958 | 25.81102929 | 25.8891582 | 25.81724054 |
| Serine/threonine-protein kinase 3;Ser | Q13188;Q13188-2 | STK3 | Other | 0.956035406 | 1.946229426 | 24.19249751 | 24.85842196 | 25.83742486 | 24.90389361 | 24.89684494 | 24.57686673 | 26.16748706 | 25.67201047 | 25.66158349 |
| Serine/threonine-protein kinase 38 | Q15208 | STK38 | Other | 0.402462955 | 1.067174935 | 24.23110631 | 23.5574235 | 24.25637917 | 24.38462515 | 24.32051577 | 23.82152877 | 24.72061961 | 24.52832485 | 24.24423374 |
| STE20/SPS1-related proline-alanine-ri | Q9UEW8-2;Q9UEW8 | STK39 | Extracellular or secreted | 0.990487386 | 2.04228391 | 23.63754647 | 21.97646681 | 22.5980525 | 22.7398844 | 23.04786678 | 23.447962 | 23.76547065 | 24.11107962 | 23.81880137 |
| Stathmin | P16949;P16949-2 | STMN1 | Extracellular or secreted | 1.358038325 | 1.824980745 | 27.96069012 | 27.94625861 | 28.1705942 | 27.98836883 | 28.14923552 | 27.87619279 | 29.56527222 | 29.56619814 | 28.98831465 |
| Stathmin-2 | Q93045;Q93045-2 | STMN2 | Other | 2.617939969 | 1.670989437 | 21.27400476 | 26.09101612 | 26.05587602 | 25.36629686 | 25.74555493 | 26.47739797 | 27.73453722 | 27.91826392 | 27.7060921 |
| Erythrocyte band 7 integral membran | P27105 | STOM | Extracellular or secreted | 0.821227619 | 1.613823689 | 25.07312398 | 25.27127608 | 25.73096677 | 24.15614914 | 25.34340153 | 25.92755048 | 25.89061998 | 26.00006742 | 26.32422945 |
| Stomatin-like protein 2, mitochondria | Q9UJZ1;Q9UJZ1-2 | STOML2 | Extracellular or secreted | -1.403674793 | 0.706320782 | 23.74108591 | 23.40275603 | 24.01971017 | 24.17058465 | 24.26191184 | 27.3529602 | 21.77579351 | 23.20845228 | 24.27923423 |
| Stonin-1 | Q9Y6Q2;Q9Y6Q2-3;Q9Y6Q2- | STON1 | Other | -1.421188978 | 0.954975775 | 24.55666368 | 23.62019655 | 23.6741138 | 20.0936439 | 23.17489172 | 24.14328169 | 21.08777108 | 21.7231618 | 22.55689586 |
| Stonin-2 | Q8WXE9;Q8WXE9-3 | STON2 | Other | -4.636601539 | 3.154394606 | 25.94578994 | 26.51395898 | 26.72273146 | 25.47685743 | 25.92915455 | 25.09051144 | 21.87087938 | 20.57633437 | 21.48248353 |
| Serine-threonine kinase receptor-ass | Q9Y3F4;Q9Y3F4-2 | STRAP | Other | 0.604019033 | 1.127627021 | 30.5243059 | 30.46041334 | 30.37772139 | 30.57758718 | 30.67909024 | 30.04465955 | 31.24932033 | 31.24615181 | 30.64847375 |
| Spermatid perinuclear RNA-binding p | Q96SI9-2;Q96SI9 | STRBP | Other | -0.910022253 | 1.647676745 | 29.2785033 | 29.31658969 | 29.38650526 | 28.08217902 | 28.06224209 | 29.57071027 | 27.99102119 | 28.10284546 | 28.0244314 |
| Striatin | O43815;O43815-2 | STRN | Extracellular or secreted | -0.338458206 | 1.019945586 | 28.18536879 | 28.26789659 | 28.1773114 | 27.52230531 | 27.47212286 | 27.75193903 | 27.61715837 | 27.68865851 | 27.3672805 |
| Striatin-4 | Q9NRL3;Q9NRL3-3 | STRN4 | Other | 0.936768054 | 1.213441411 | 24.70239762 | 24.7776105 | 24.05017747 | 25.31958585 | 23.75725245 | 23.96252256 | 25.65176494 | 25.67052551 | 24.77278694 |
| Dolichyl-diphosphooligosaccharide--p | Q8TCJ2 | STT3B | Other | -2.029097736 | 0.68484224 | 23.18498306 | 23.01161219 | 23.62511128 | 24.11965294 | 23.87229913 | 26.70661911 | 20.57619896 | 21.30808923 | 24.28855746 |
| E3 ubiquitin-protein ligase CHIP | Q9UNE7-2;Q9UNE7 | STUB1 | Extracellular or secreted | 0.952884613 | 1.336548041 | 23.67626788 | 23.78086732 | 24.54904338 | 23.66990407 | 24.37285284 | 24.9294932 | 24.5791709 | 25.48882547 | 25.27987182 |
| Syntaxin-10 | O60499;O60499-2 | STX10 | Other | 0.346957647 | 1.05052077 | 23.66448895 | 23.38179007 | 23.2831969 | 23.48345367 | 22.96135713 | 22.83155621 | 23.55835812 | 23.45073302 | 23.83470327 |
| Syntaxin-12 | Q86Y82 | STX12 | Extracellular or secreted | -0.201037856 | 0.349523358 | 25.99062095 | 26.11612658 | 26.01599431 | 26.15531932 | 26.29409106 | 26.05885104 | 25.64809312 | 25.73619009 | 26.32810485 |
| Syntaxin-3 | Q13277-2;Q13277-3;Q13277 | STX3 | Extracellular or secreted | 0.58963955 | 1.339040495 | 24.1155314 | 23.98641705 | 23.88874028 | 23.92705309 | 23.75041913 | 24.37696422 | 24.30570386 | 24.61296161 | 24.87281576 |
| Syntaxin-4 | Q12846;Q12846-2 | STX4 | Extracellular or secreted | 1.682245017 | 1.478922031 | 26.13791876 | 25.44827715 | 26.16482393 | 23.94207989 | 22.55725988 | 24.28206582 | 26.80353539 | 26.26155322 | 26.24785915 |
| Syntaxin-6 | O43752 | STX6 | Extracellular or secreted | -0.0159229 | 0.018240579 | 24.77011701 | 24.20784189 | 24.34921859 | 24.91479022 | 24.30883117 | 23.22081285 | 24.57068494 | 24.29165187 | 23.97570036 |
| Syntaxin-7 | O15400-2;O15400 | STX7 | Extracellular or secreted | -0.331922935 | 0.901282539 | 26.78693542 | 26.59177844 | 26.66451608 | 26.62728406 | 26.36609672 | 26.42828656 | 25.99379824 | 26.26230622 | 26.48057538 |
| Syntaxin-8 | Q9UNK0 | STX8 | Extracellular or secreted | -0.65972843 | 0.9383135 | 24.90081117 | 24.98511439 | 24.53249243 | 24.23914281 | 24.77293792 | 23.93606996 | 24.24016244 | 24.09980944 | 23.36412717 |
| Syntaxin-binding protein 1 | P61764;P61764-2 | STXBP1 | Extracellular or secreted | 0.184998319 | 0.944328002 | 26.92354384 | 27.16844381 | 27.15759535 | 26.87067758 | 26.87970215 | 26.69245481 | 27.07673651 | 27.05724033 | 27.26722689 |
| Syntaxin-binding protein 2 | Q15833-2;Q15833;Q15833-3 | STXBP2 | Extracellular or secreted | 0.063470147 | 0.103168029 | 28.39631851 | 28.43479803 | 28.41816228 | 27.79054418 | 28.00982614 | 27.6289532 | 28.33845383 | 28.35695234 | 27.83430544 |
| Syntaxin-binding protein 3 | O00186 | STXBP3 | Extracellular or secreted | 0.336532272 | 0.991246151 | 25.54297705 | 25.67516432 | 26.0274719 | 25.81320872 | 26.18789993 | 25.43256178 | 25.90955822 | 26.10013038 | 26.33955007 |
| Activated RNA polymerase II transcrip | P53999 | SUB1 | Extracellular or secreted | 0.358074415 | 0.384999142 | 30.31656811 | 30.15255142 | 30.09672681 | 30.57704688 | 30.56392768 | 30.40385324 | 30.96285623 | 31.14370013 | 30.02300396 |
| Succinyl-CoA ligase [ADP-forming] sub | Q9P2R7-2;Q9P2R7 | SUCLA2 | Extracellular or secreted | -0.248893321 | 0.990327724 | 26.98032788 | 26.962677 | 27.0986856 | 27.28020821 | 27.20004039 | 27.7351831 | 27.08436042 | 26.90873399 | 26.88878672 |
| Succinyl-CoA ligase [ADP/GDP-formin | P53597 | SUCLG1 | Extracellular or secreted | -1.988186035 | 3.070594307 | 28.78998479 | 28.32555958 | 28.30151482 | 27.87695388 | 27.85459887 | 27.90690069 | 25.99948688 | 25.98252729 | 26.58118403 |
| Succinyl-CoA ligase [GDP-forming] sub | Q96I99;Q96I99-2 | SUCLG2 | Extracellular or secreted | -1.576056608 | 2.509509476 | 29.86555422 | 29.81046247 | 29.79439187 | 29.853469 | 29.914277 | 29.47763343 | 28.05377536 | 28.07050588 | 28.50544293 |
| Sin3 histone deacetylase corepressor | Q9H7L9 | SUDS3 | Extracellular or secreted | -0.865308734 | 1.365125942 | 24.37570553 | 24.48142253 | 24.17752052 | 23.46550562 | 23.56406946 | 24.91551982 | 23.10463202 | 23.01597305 | 23.77334046 |
| SURP and G-patch domain-containing | Q8IWZ8 | SUGP1 | Other | 0.013611796 | 0.041744382 | 24.35400343 | 24.30201187 | 24.21415664 | 23.98207027 | 24.56209053 | 24.05299848 | 24.11020355 | 24.32347373 | 24.34082372 |
| Suppressor of G2 allele of SKP1 homo | Q9Y2Z0-2;Q9Y2Z0 | SUGT1 | Other | 0.62392656 | 1.738392948 | 27.47173536 | 27.3661134 | 27.41916746 | 27.73576414 | 27.59903497 | 27.40908401 | 28.22921855 | 28.25619918 | 27.88681162 |
| Extracellular sulfatase Sulf-2 | Q8IWU5;Q8IWU5-2 | SULF2 | Extracellular or secreted | 0.31566626 | 0.393872478 | 27.16595492 | 26.89006329 | 27.08294035 | 27.13133905 | 27.17797666 | 29.09660116 | 27.67134918 | 27.62114729 | 27.92694002 |
| Sulfotransferase 1A4;Sulfotransferase | P0DMN0;P0DMM9;P0DMM9 | SULT1A4;SULT1A | Other | 0.701225573 | 0.158842425 | 21.83402577 | 22.68933858 | 23.25537095 | 21.75534308 | 21.26907159 | 23.1661312 | 24.53931314 | 24.5676131 | 19.98139107 |
| Sulfatase-modifying factor 1 | Q8NBK3-2;Q8NBK3-5;Q8NBK | SUMF1 | Extracellular or secreted | 0.488141077 | 0.976512283 | 24.04743415 | 23.59798433 | 23.50373065 | 23.5920637 | 23.34786787 | 24.72223636 | 24.33067129 | 24.54339013 | 23.99602034 |
| Sulfatase-modifying factor 2 | Q8NBJ7;Q8NBJ7-2;Q8NBJ7-3 | SUMF2 | Extracellular or secreted | -0.42805874 | 0.97254473 | 27.34528111 | 27.40819328 | 27.38038695 | 27.54635202 | 27.2590763 | 26.97311821 | 26.9139918 | 26.58892275 | 27.16911317 |
| Small ubiquitin-related modifier 1 | P63165 | SUMO1 | Extracellular or secreted | -0.481132716 | 0.486239323 | 29.60803698 | 29.67754535 | 29.80077682 | 29.38580616 | 29.47278137 | 28.42658411 | 29.10694346 | 29.42852218 | 28.2069016 |
| Small ubiquitin-related modifier 2 | P61956-2;P61956 | SUMO2 | Other | 0.230931387 | 0.261735139 | 30.25900894 | 29.85332026 | 29.99361471 | 30.12423126 | 30.33564571 | 28.95737256 | 30.24229494 | 30.59663795 | 29.61545798 |
| Small ubiquitin-related modifier 3 | P55854;P55854-2 | SUMO3 | Extracellular or secreted | 0.424820328 | 0.505060525 | 23.699647 | 23.59012283 | 23.99101038 | 23.51713207 | 23.87229913 | 21.57280755 | 23.80970534 | 23.99101038 | 23.59525475 |
| Small ubiquitin-related modifier 4 | Q6EEV6 | SUMO4 | Other | -5.578260569 | 1.72960598 | 29.28363407 | 29.00402278 | 29.01310527 | 24.57253648 | 27.38508615 | 26.26520697 | 20.0665038 | 24.14693654 | 21.81357381 |
| Sulfite oxidase, mitochondrial | P51687 | SUOX | Other | -1.568569553 | 0.795931585 | 25.59812635 | 25.27148981 | 25.24822119 | 24.70429882 | 25.01303603 | 24.65801836 | 24.34536572 | 24.13703928 | 22.05848162 |
| FACT complex subunit SPT16 | Q9Y5B9 | SUPT16H | Other | -0.253131652 | 0.245250315 | 32.17456399 | 32.22983166 | 32.12460104 | 31.66033942 | 31.68386558 | 31.60443585 | 32.07934588 | 31.97182404 | 30.92825389 |
| Transcription elongation factor SPT4 | P63272 | SUPT4H1 | Other | 0.185186524 | 0.219596369 | 25.14914828 | 25.45070156 | 25.37663309 | 25.45016665 | 25.52190111 | 24.77635592 | 25.77989105 | 25.7517986 | 24.88632323 |
| Transcription elongation factor SPT5 | O00267-2;O00267 | SUPT5H | Other | 0.280602918 | 0.642844092 | 29.33022677 | 29.27081732 | 29.20043802 | 29.47376857 | 29.53543596 | 29.18269588 | 29.7727083 | 29.78620324 | 29.27958848 |
| Transcription elongation factor SPT6 | Q7KZ85 | SUPT6H | Other | 0.190932161 | 0.545674309 | 28.38730676 | 28.51585465 | 28.51077135 | 28.26718223 | 28.25313595 | 28.76622923 | 28.84242113 | 28.68217507 | 28.39844036 |
| Surfeit locus protein 4 | O15260-2;O15260;O15260-3 | SURF4 | Other | 0.104333748 | 0.030169864 | 23.95464646 | 23.73624171 | 23.72941164 | 22.74094185 | 22.93261414 | 28.12099782 | 23.34095951 | 23.55039452 | 26.02907403 |
| Sushi domain-containing protein 5 | O60279 | SUSD5 | Extracellular or secreted | -0.941480814 | 0.76917918 | 22.7692293 | 22.01981195 | 22.41173027 | 22.54210328 | 22.38210682 | 21.51841883 | 20.52307391 | 22.14249205 | 21.33169183 |
| Supervillin | O95425-2;O95425-4;O95425 | SVIL | Extracellular or secreted | -0.562931318 | 0.745542414 | 24.38007312 | 24.91428841 | 25.01750304 | 24.75536887 | 25.42165729 | 25.2643123 | 23.94752993 | 24.30124443 | 24.93903321 |
| Switch-associated protein 70 | Q9UH65 | SWAP70 | Extracellular or secreted | 0.132855637 | 0.211121155 | 26.62338187 | 25.60856933 | 25.89555125 | 25.24070837 | 26.77580356 | 25.7746228 | 26.16495816 | 26.17816667 | 26.01476066 |
| Synapse-associated protein 1 | Q96A49 | SYAP1 | Extracellular or secreted | 0.322072107 | 0.343182048 | 25.31762073 | 25.54901399 | 25.5791709 | 25.76617867 | 25.46407256 | 24.72343472 | 26.35173092 | 25.63098139 | 25.1832498 |
| Symplekin | Q92797;Q92797-2 | SYMPK | Extracellular or secreted | 0.055377107 | 0.130553157 | 27.01818252 | 26.97836518 | 26.97059727 | 26.80919014 | 26.58935147 | 27.29867749 | 27.11068147 | 27.13613971 | 26.75149216 |
| Heterogeneous nuclear ribonucleopro | O60506;O60506-2 | SYNCRIP | Other | -0.105783881 | 0.202841457 | 31.0251842 | 31.17390907 | 30.99974224 | 31.02696559 | 31.06297542 | 30.75716342 | 30.98807081 | 31.16566747 | 30.55188005 |
| Heterogeneous nuclear ribonucleopro | O60506-3;O60506-4;O60506 | SYNCRIP | Other | -0.188623926 | 0.118401883 | 25.77964061 | 25.53549158 | 25.46827427 | 24.5666264 | 24.78775752 | 23.49692203 | 24.96925828 | 25.43093614 | 23.85129001 |
| Nesprin-1 | Q8NF91-4;Q8NF91;Q8NF91-1 | SYNE1 | Extracellular or secreted | -2.621598593 | 0.773584312 | 25.14176259 | 24.76202673 | 25.10697341 | 27.26588655 | 27.65812735 | 20.32060313 | 24.77650653 | 20.62356955 | 21.86281802 |
| Nesprin-2 | Q8WXH0;Q8WXH0-2;Q8WXH | SYNE2 | Extracellular or secreted | -2.022751183 | 1.729179853 | 29.07823906 | 28.33415433 | 28.61729852 | 28.41651227 | 28.19037013 | 30.70734344 | 26.47075841 | 26.37489342 | 27.7580535 |
| Synaptogyrin-3 | O43761 | SYNGR3 | Other | -0.388710043 | 0.330488106 | 21.88866597 | 21.8450612 | 22.02634513 | 22.08388181 | 22.02282834 | 22.26633705 | 21.69604212 | 22.35511985 | 20.84926764 |
| Synaptophysin-like protein 1 | Q16563-2;Q16563 | SYPL1 | Extracellular or secreted | 0.472189805 | 0.475429924 | 24.46929949 | 25.21500887 | 25.32642565 | 24.30347587 | 23.12407839 | 25.93786659 | 24.92020782 | 25.07091527 | 25.61352376 |
| SUZ domain-containing protein 1 | Q7Z422-2;Q7Z422-4;Q7Z422- | SZRD1 | Other | 0.587573166 | 0.549720727 | 25.70192192 | 26.17365655 | 26.12103735 | 26.95260405 | 26.8959672 | 24.49203898 | 27.08162045 | 26.86325163 | 25.98646045 |
| TGF-beta-activated kinase 1 and MAP | Q15750;Q15750-2 | TAB1 | Other | -0.178392646 | 0.541823802 | 26.08415764 | 26.10171397 | 25.93064405 | 26.27074159 | 26.53894327 | 25.82767085 | 26.03674376 | 26.07689957 | 25.72811441 |
| Transforming acidic coiled-coil-contai | O75410;O75410-2;O75410-6 | TACC1 | Other | -1.374443726 | 0.338545756 | 26.46291884 | 26.90827588 | 26.86360611 | 22.72407809 | 21.21112329 | 25.92415575 | 25.93775437 | 21.22275475 | 23.76323869 |
| Transforming acidic coiled-coil-contai | O95359-3;O95359;O95359-2 | TACC2 | Other | 2.747062085 | 3.323549059 | 23.4374595 | 22.76832084 | 22.79481325 | 23.79788313 | 22.29647712 | 23.35299387 | 26.06136674 | 26.0977417 | 25.30605167 |
| Transforming acidic coiled-coil-contai | Q9Y6A5 | TACC3 | Other | -0.029675415 | 0.042982248 | 25.51821323 | 25.66985002 | 25.60424848 | 26.45727729 | 26.43012263 | 25.862164 | 25.77122588 | 25.68354102 | 26.22714467 |
| Tumor-associated calcium signal trans | P09758 | TACSTD2 | Extracellular or secreted | 0.987416009 | 1.625243318 | 29.7167431 | 29.57891906 | 29.72427754 | 29.20697145 | 29.3157911 | 29.92860688 | 30.27963275 | 30.43678467 | 30.98148517 |
| Transcription initiation factor TFIID su | Q12962 | TAF10 | Other | 0.250685803 | 0.493932119 | 23.0142371 | 22.95665053 | 22.78466599 | 23.65294123 | 23.53670728 | 23.35837004 | 23.53017198 | 23.13794611 | 23.73572541 |
| TATA-binding protein-associated facto | Q92804-2;Q92804 | TAF15 | Other | 0.429549586 | 1.304586053 | 27.8277193 | 27.9237705 | 27.89954409 | 28.09778188 | 28.1892864 | 27.69052461 | 28.487078 | 28.49800322 | 28.11788093 |
| Transcription initiation factor TFIID su | P49848-4;P49848-2;P49848;F | TAF6 | Other | -0.983304576 | 0.673091206 | 22.53945514 | 22.57614016 | 22.36407372 | 22.337303 | 22.79920434 | 22.46655149 | 20.59670375 | 21.5052824 | 22.48946404 |
| Transcription initiation factor TFIID su | Q15545 | TAF7 | Other | -0.20169737 | 0.130290776 | 22.80701445 | 23.69498014 | 23.85262399 | 21.96199634 | 23.10752424 | 22.18765203 | 22.96082708 | 21.82607582 | 23.41390059 |
| Transgelin | Q01995 | TAGLN | Other | 0.61638644 | 0.285106291 | 21.69033253 | 25.30549513 | 25.78446631 | 27.43785631 | 27.61294755 | 25.59607985 | 25.87600539 | 26.1375084 | 26.54923438 |
| Transgelin-2 | P37802;P37802-2 | TAGLN2 | Extracellular or secreted | -0.024863452 | 0.083828351 | 33.04412211 | 33.09139261 | 33.14067158 | 33.33471377 | 33.25855974 | 32.86789365 | 32.93759654 | 33.14873869 | 33.20775115 |
| Transaldolase | P37837 | TALDO1 | Extracellular or secreted | -0.279539923 | 1.501822022 | 32.09106153 | 32.14315272 | 32.05759541 | 31.9620288 | 32.06702135 | 31.82304852 | 31.86082698 | 31.75509504 | 31.61741238 |
| Protein TANC1 | Q9C0D5-2;Q9C0D5 | TANC1 | Extracellular or secreted | -1.405166615 | 0.399854511 | 22.9908631 | 21.46132982 | 22.49006788 | 22.07111363 | 22.73712598 | 21.86268427 | 18.90580417 | 20.3147616 | 23.37052672 |
| Serine/threonine-protein kinase TAO1 | Q7L7X3;Q7L7X3-3 | TAOK1 | Extracellular or secreted | 0.354596336 | 0.592920304 | 25.44673222 | 25.4452488 | 25.85524072 | 24.76890636 | 25.68461145 | 25.54716143 | 26.16186767 | 25.86947788 | 25.40639395 |
| RISC-loading complex subunit TARBP2 | Q15633-2;Q15633 | TARBP2 | Other | -0.351252135 | 0.425028414 | 21.64540632 | 23.12407862 | 22.96560325 | 22.98181385 | 22.20253417 | 21.45658503 | 21.66899575 | 22.03577102 | 22.42948745 |
| TAR DNA-binding protein 43 | Q13148 | TARDBP | Other | -0.121545013 | 0.172350246 | 27.22361406 | 27.35388568 | 27.63885867 | 27.96212536 | 27.93915654 | 27.73246847 | 27.83683522 | 27.65328299 | 27.07030114 |
| Threonine--tRNA ligase, cytoplasmic | P26639;P26639-2 | TARS | Extracellular or secreted | 0.431252318 | 3.166124705 | 30.5556329 | 30.748952 | 30.69760048 | 30.58772525 | 30.55581572 | 30.29723486 | 31.07909151 | 30.98338927 | 30.95275677 |
| Putative deoxyribonuclease TATDN1 | Q6P1N9;Q6P1N9-2 | TATDN1 | Other | -0.339810388 | 0.483316335 | 28.98207027 | 29.05297258 | 29.12067654 | 28.90859085 | 29.14019312 | 28.42778323 | 29.03094268 | 28.64739332 | 28.11837613 |
| Putative deoxyribonuclease TATDN3 | Q17R31-5;Q17R31-4;Q17R31 | TATDN3 | Other | -0.763993857 | 0.321046868 | 23.94190085 | 23.51400414 | 23.74499065 | 23.83248367 | 23.93526075 | 21.86698534 | 23.66209984 | 23.48000519 | 20.98372609 |
| Tax1-binding protein 3 | O14907 | TAX1BP3 | Extracellular or secreted | 0.54809952 | 1.182681516 | 26.94022128 | 26.79550702 | 26.75047025 | 27.60583147 | 27.56601663 | 26.41125184 | 27.64931353 | 27.77190591 | 27.25772837 |
| TBC1 domain family member 1 | Q86TI0-2;Q86TI0 | TBC1D1 | Other | -1.40861628 | 0.797894753 | 23.12469371 | 23.61840522 | 23.06272999 | 23.05910903 | 21.62070918 | 22.25614878 | 22.6871609 | 21.03836568 | 20.41952254 |
| TBC1 domain family member 10B | Q4KMP7 | TBC1D10B | Extracellular or secreted | -0.235437763 | 0.848645164 | 26.46199832 | 26.47477056 | 26.18629487 | 26.11370465 | 26.54685243 | 25.67788134 | 26.05337143 | 26.03250418 | 25.93856219 |
| TBC1 domain family member 13 | Q9NVG8-2;Q9NVG8 | TBC1D13 | Other | 0.309685336 | 0.765181066 | 25.4778457 | 25.32488179 | 25.2045995 | 25.56633607 | 25.55946719 | 25.46715501 | 25.81841132 | 25.44603804 | 25.96474927 |
| TBC1 domain family member 15 | Q8TC07-2;Q8TC07;Q8TC07-3 | TBC1D15 | Extracellular or secreted | -0.030132993 | 0.026769593 | 25.31510007 | 25.47336216 | 25.26549315 | 25.14074897 | 25.32625419 | 25.22070217 | 25.75491033 | 25.45177078 | 24.57375026 |
| TBC1 domain family member 2A | Q9BYX2-4;Q9BYX2-3;Q9BYX2 | TBC1D2 | Other | -0.411608961 | 0.74068643 | 22.4078439 | 21.62947131 | 21.82563312 | 22.68124645 | 20.9548744 | 22.47092593 | 21.60521544 | 21.56096901 | 21.58398623 |
| TBC1 domain family member 22A | Q8WUA7-3;Q8WUA7-4;Q8W | TBC1D22A | Other | 0.371401232 | 0.137204869 | 23.26828042 | 21.23765578 | 22.98379316 | 24.68142459 | 21.85207283 | 24.97307441 | 22.06582237 | 23.72077614 | 24.82575577 |
| TBC1 domain family member 23 | Q9NUY8-2;Q9NUY8 | TBC1D23 | Other | -0.169158983 | 0.593358147 | 26.21183825 | 26.2326989 | 26.54732325 | 26.01960413 | 26.09148026 | 25.5686571 | 25.92739224 | 25.93865192 | 25.96227984 |
| TBC1 domain family member 24 | Q9ULP9-2;Q9ULP9 | TBC1D24 | Other | 0.136753872 | 0.544582074 | 25.79168715 | 26.23108799 | 26.10407609 | 25.87977225 | 26.37515865 | 26.03235704 | 26.05212787 | 26.27951763 | 26.28568569 |
| TBC1 domain family member 4 | O60343-2;O60343-3;O60343 | TBC1D4 | Extracellular or secreted | 0.322657793 | 0.581124512 | 26.44748913 | 26.17693112 | 26.35045001 | 25.57951621 | 25.43587115 | 25.94717318 | 26.603711 | 26.38437813 | 25.94859965 |

| | | | | | | | | | | | | | | |
|---|---|---|---|---|---|---|---|---|---|---|---|---|---|---|
| TBC1 domain family member 5 | Q92609;Q92609-2 | TBC1D5 | Extracellular or secreted | -2.082085393 | 0.473702436 | 24.48376117 | 22.52497626 | 24.95060324 | 21.43441946 | 24.35608761 | 24.87979561 | 19.7429294 | 20.34039139 | 24.98524471 |
| TBC1 domain family member 9B | Q66K14-2;Q66K14 | TBC1D9B | Other | -0.626887064 | 1.1370482 | 26.58290734 | 26.40632907 | 26.58677727 | 26.4840225 | 26.58046538 | 26.34267275 | 25.99798072 | 26.11346621 | 25.49947903 |
| Tubulin-specific chaperone A | O75347;O75347-2 | TBCA | Extracellular or secreted | 0.377614662 | 0.484546772 | 30.57009499 | 30.65693483 | 30.60805461 | 30.4161901 | 30.48979793 | 29.23924754 | 31.00021251 | 30.94816796 | 30.17462351 |
| Tubulin-folding cofactor B | Q99426;Q99426-2 | TBCB | Other | 0.191996263 | 0.928027403 | 28.53482399 | 28.46053051 | 28.4768806 | 28.60907681 | 28.65095739 | 28.39860345 | 28.63460696 | 28.87255747 | 28.63426074 |
| Tubulin-specific chaperone C | Q15814 | TBCC | Other | -0.017705554 | 0.017300429 | 24.32573969 | 24.55642981 | 24.48086809 | 24.30514718 | 24.0220833 | 24.0220833 | 23.77625551 | 24.84852285 | 24.17828067 |
| Tubulin-specific chaperone D | Q9BTW9;Q9BTW9-4;Q9BTW9 | TBCD | Extracellular or secreted | -0.023468892 | 0.051397942 | 27.03160008 | 26.8844845 | 27.01307864 | 26.17558031 | 26.3541548 | 26.96002722 | 26.62142681 | 26.72559447 | 26.79203483 |
| Tubulin-specific chaperone E | Q15813;Q15813-2 | TBCE | Other | -0.110292636 | 0.207030681 | 26.12835263 | 26.0987659 | 26.02221032 | 26.72234061 | 26.7284518 | 26.65662802 | 25.9703559 | 26.41828294 | 26.4588579 |
| F-box-like/WD repeat-containing prot | Q9BZK7 | TBL1XR1 | Other | -0.527495218 | 2.188265611 | 29.76728741 | 29.83764742 | 29.74070965 | 29.76796614 | 29.68907526 | 29.83249574 | 29.12765426 | 29.27739514 | 29.33005576 |
| Transducin beta-like protein 3 | Q12788 | TBL3 | Other | -1.616351953 | 1.825827924 | 22.78644194 | 23.09910313 | 23.08441719 | 23.29193285 | 23.42098305 | 25.40080336 | 21.51234119 | 22.51014505 | 21.67029866 |
| TRPM8 channel-associated factor 1 | Q9Y4C2-2;Q9Y4C2 | TCAF1 | Extracellular or secreted | 1.506586645 | 0.589500781 | 20.91833875 | 21.64924776 | 20.49201048 | 20.3079869 | 20.76647609 | 24.1586935 | 23.49692203 | 21.01754834 | 24.15166631 |
| Transcription elongation factor A prot | P23193;P23193-2 | TCEA1 | Other | 0.050993882 | 0.16669804 | 28.09164167 | 28.01499471 | 27.99145377 | 28.08931971 | 28.23662709 | 28.27684074 | 28.34367261 | 27.99139971 | 28.16834817 |
| Transcription elongation factor A prot | Q15170-2;Q15170 | TCEAL1 | Other | -1.111785801 | 0.412179219 | 22.38031097 | 21.28591322 | 21.08796614 | 23.47741341 | 23.42021211 | 22.05473811 | 21.7003669 | 19.23846236 | 22.57909031 |
| Transcription elongation factor A prot | Q969E4;Q6IPX3-2;Q6IPX3 | TCEAL3;TCEAL6 | Other | -0.656112926 | 1.537966114 | 27.54178139 | 27.4157067 | 27.39844036 | 26.86266062 | 26.66016938 | 26.18332553 | 26.40516068 | 26.25290143 | 26.4046411 |
| Transcription elongation factor A prot | Q96EI5;Q96EI5-2 | TCEAL4 | Other | 0.282630633 | 0.598737542 | 26.37610313 | 26.6628876 | 26.80550478 | 26.30810207 | 26.46260686 | 25.99998143 | 26.92705309 | 26.85780525 | 26.37062649 |
| Transcription elongation factor B poly | Q15369;Q15369-2 | TCEB1 | Other | 0.619339334 | 0.58230634 | 28.72035214 | 28.05263579 | 28.13618862 | 28.0796892 | 28.52492234 | 27.38426283 | 29.32309143 | 29.0015553 | 27.98239673 |
| Transcription elongation factor B poly | Q15370;Q15370-2 | TCEB2 | Extracellular or secreted | 0.799021942 | 1.026050811 | 28.690458 | 28.44312358 | 28.34663425 | 27.97568943 | 28.61911922 | 28.12538958 | 29.43320675 | 29.47732463 | 28.58674148 |
| Transcription elongation regulator 1 | O14776-2;O14776 | TCERG1 | Other | -0.707790517 | 2.379532635 | 28.58917285 | 28.5151403 | 28.5526314 | 29.30343232 | 29.15097913 | 28.89726629 | 28.03033761 | 28.02770389 | 28.32289809 |
| Transcription factor 20 | Q9UGU0-2;Q9UGU0 | TCF20 | Other | -0.128135397 | 0.099511488 | 22.86325163 | 22.62433051 | 22.73694358 | 22.28827582 | 22.36879631 | 22.33188789 | 22.50564585 | 21.62307339 | 23.09361744 |
| Transcription factor 25 | Q9BQ70 | TCF25 | Other | 0.082591948 | 0.116556036 | 22.10523971 | 21.88769959 | 22.32366614 | 20.58693943 | 22.21157465 | 21.89983124 | 21.97053144 | 21.85289064 | 21.93182913 |
| Transcobalamin-1 | P20061 | TCN1 | Extracellular or secreted | 2.527317181 | 4.20094148 | 27.27941136 | 27.27533169 | 27.31768973 | 28.42081445 | 28.43531482 | 27.84577429 | 30.03629835 | 30.20419835 | 30.628623 |
| Treacle protein | Q13428-3;Q13428;Q13428-4 | TCOF1 | Other | -1.008424868 | 2.770575153 | 29.71774016 | 29.71610527 | 29.69228852 | 29.54419752 | 29.71979749 | 29.51335289 | 28.52604247 | 28.80036026 | 28.60006358 |
| T-complex protein 1 subunit alpha | P17987 | TCP1 | Extracellular or secreted | 0.056049162 | 0.05853648 | 32.31175904 | 32.3746157 | 32.2994201 | 32.16083803 | 32.2059176 | 31.48308459 | 32.51246306 | 32.46557958 | 31.60792238 |
| Tyrosyl-DNA phosphodiesterase 1 | Q9NUW8;Q9NUW8-2 | TDP1 | Extracellular or secreted | -0.149558093 | 0.235460631 | 25.39227837 | 25.37921388 | 25.60266376 | 25.32680278 | 25.30204675 | 24.72567247 | 25.38228496 | 25.32123862 | 24.71214114 |
| Tyrosyl-DNA phosphodiesterase 2 | O95551-3;O95551;O95551-2 | TDP2 | Extracellular or secreted | 0.666899768 | 0.261559197 | 23.62889759 | 23.7234868 | 23.64515442 | 21.52467863 | 22.15281014 | 20.79575365 | 23.82561022 | 24.35904066 | 21.55143904 |
| Transcriptional enhancer factor TEF-5 | Q99594 | TEAD3 | Other | -0.918724248 | 0.461837778 | 22.94569173 | 22.79918833 | 22.87327844 | 22.89374686 | 20.04460276 | 23.06503346 | 20.11864142 | 22.42680322 | 22.0091534 |
| Very-long-chain enoyl-CoA reductase | Q9NZ01 | TECR | Other | -2.131909851 | 0.520703607 | 24.97395026 | 24.80911652 | 25.27944679 | 25.61551761 | 25.17455209 | 29.06293683 | 23.0560456 | 21.37897922 | 26.62700568 |
| Telomeric repeat-binding factor 2-inte | Q9NYB0 | TERF2IP | Other | 0.351489603 | 1.033873413 | 27.66627819 | 27.40576122 | 27.50627562 | 27.71626884 | 27.50582149 | 26.73569959 | 27.92852217 | 27.82377743 | 27.57022169 |
| Testin | Q9UGI8-2;Q9UGI8 | TES | Extracellular or secreted | -0.193007671 | 0.98138692 | 28.17398053 | 28.00308456 | 27.86561322 | 28.19187656 | 28.40628852 | 28.04551905 | 27.87050122 | 27.83863949 | 28.0550175 |
| Serotransferrin | P02787 | TF | Extracellular or secreted | 13.58074234 | 5.018303028 | 18.92181189 | 17.65431436 | 19.24571981 | 19.71724466 | 19.37026596 | 17.5068809 | 31.59321308 | 32.10942409 | 33.24770865 |
| Transcription factor A, mitochondrial | Q00059;Q00059-2 | TFAM | Extracellular or secreted | -0.135307551 | 0.098565824 | 24.4619515 | 24.19505375 | 24.15182112 | 25.79056904 | 26.20769297 | 26.9922645 | 24.94628092 | 25.27997806 | 25.26749481 |
| Transcription factor AP-2-alpha;Trans | P05549-5;P05549-6;P05549;( | TFAP2A;TFAP2E;T | Other | -0.017437988 | 0.027608806 | 22.61362492 | 21.70897776 | 22.23680954 | 22.58301704 | 21.91056954 | 22.08637858 | 22.47449518 | 21.94884914 | 22.09403041 |
| Transcription factor AP-4 | Q01664 | TFAP4 | Other | 0.854723327 | 2.022536999 | 21.43527991 | 20.91298284 | 20.8283069 | 21.5689962 | 21.24591296 | 22.19229978 | 22.3508108 | 21.95716441 | 22.34808407 |
| Dimethyladenosine transferase 2, mit | Q9H5Q4 | TFB2M | Other | 0.460526323 | 0.364591764 | 22.03280317 | 21.65175105 | 22.09924756 | 23.69636056 | 23.60433333 | 24.7976358 | 23.74108591 | 23.56313854 | 23.01842026 |
| Alpha-globin transcription factor CP2 | Q12800-3;Q12800;Q12800-2 | TFCP2 | Other | -1.701284989 | 0.574401426 | 24.42098305 | 24.45907685 | 24.46301242 | 25.225932 | 25.00466042 | 24.2365906 | 24.14841141 | 20.69919127 | 23.95367001 |
| Protein TFG | Q92734-2;Q92734;Q92734-4 | TFG | Extracellular or secreted | 0.139829555 | 0.217859043 | 29.58421639 | 29.51410576 | 29.48344983 | 29.52807516 | 29.6367171 | 29.08846064 | 29.81785959 | 29.85510405 | 29.16403747 |
| Tuftelin-interacting protein 11 | Q9UBB9-2;Q9UBB9 | TFIP11 | Extracellular or secreted | -0.547819668 | 0.712787151 | 22.97330218 | 22.61983846 | 23.19320469 | 20.89795697 | 23.30985748 | 22.32317134 | 21.88959457 | 22.05824414 | 22.06736785 |
| Transferrin receptor protein 1;Transfe | P02786 | TFRC | Extracellular or secreted | -0.789663281 | 1.341150421 | 30.66084942 | 30.90282471 | 30.85623264 | 31.55439828 | 31.62261412 | 32.05407305 | 30.261477 | 30.27442106 | 30.92060821 |
| Transforming growth factor beta-1;La | P01137 | TGFB1 | Extracellular or secreted | 1.377358649 | 0.51252358 | 21.18597991 | 19.45942802 | 22.39777859 | 25.10029082 | 24.57767361 | 26.50500369 | 24.12589206 | 23.79916856 | 25.82009266 |
| Protein-glutamine gamma-glutamyltra | P21980;P21980-2 | TGM2 | Extracellular or secreted | -1.99147825 | 2.038019814 | 25.78049191 | 26.20918144 | 25.42300483 | 25.63956262 | 25.27810003 | 24.50657829 | 23.95571092 | 23.67745126 | 22.81086264 |
| Protein-glutamine gamma-glutamyltra | Q08188 | TGM3 | Extracellular or secreted | -1.061363668 | 2.179596774 | 22.49279693 | 22.41841163 | 22.81642043 | 23.47258772 | 23.84412263 | 22.53167178 | 21.68637058 | 21.78384713 | 22.13369686 |
| Trans-Golgi network integral membra | O43493-4;O43493-2;O43493 | TGOLN2 | Extracellular or secreted | -0.854623094 | 2.594372532 | 24.82274469 | 24.65053291 | 25.3148927 | 25.17607512 | 24.82934144 | 25.52270939 | 23.9703559 | 24.27923423 | 24.34468872 |
| Thrombospondin-1 | P07996;P07996-2 | THBS1 | Extracellular or secreted | 0.575426123 | 0.305679378 | 30.49667065 | 30.38679303 | 30.36233472 | 31.37326783 | 31.43146427 | 31.0140503 | 30.71513981 | 30.75023063 | 32.79319832 |
| Thrombospondin-4 | P35443 | THBS4 | Extracellular or secreted | -2.694120169 | 1.982526602 | 24.71298074 | 24.68045911 | 24.68437067 | 25.33350616 | 25.69956758 | 22.98072882 | 21.42921432 | 22.98626684 | 21.54796487 |
| Probable tRNA(His) guanylyltransferas | Q9NWX6 | THG1L | Other | 0.548588139 | 1.5062862 | 23.66470595 | 24.2740522 | 24.42136837 | 24.4170598 | 24.46662616 | 24.447962 | 25.05845885 | 24.85419459 | 24.57899821 |
| Threonine synthase-like 1 | Q8IYQ7 | THNSL1 | Other | -0.342458814 | 0.279697245 | 22.72250738 | 22.67635398 | 22.35940361 | 19.89767114 | 22.44788635 | 23.29361758 | 22.1795872 | 21.73314114 | 21.75861524 |
| THO complex subunit 1 | Q96FV9 | THOC1 | Extracellular or secreted | -0.997733314 | 2.775019152 | 26.89365482 | 26.96046919 | 27.18980481 | 26.26332752 | 26.46855395 | 26.17207372 | 25.53023153 | 25.87991244 | 25.57059809 |
| THO complex subunit 2 | Q8NI27 | THOC2 | Other | -2.046835336 | 1.680808997 | 26.91718284 | 26.88378563 | 26.80919014 | 26.22589524 | 26.22685078 | 26.23551367 | 23.86787671 | 25.17569451 | 24.46513192 |
| THO complex subunit 3 | Q96J01;Q96J01-2 | THOC3 | Extracellular or secreted | 0.296174568 | 0.420501599 | 28.57589356 | 28.43161371 | 28.39030326 | 28.4929329 | 28.50074063 | 28.13609081 | 28.92362885 | 29.03642937 | 28.19225293 |
| THO complex subunit 5 homolog | Q13769 | THOC5 | Other | -1.318839269 | 1.45366326 | 25.54318361 | 25.98604808 | 26.02462162 | 24.90269822 | 26.04680994 | 26.42350181 | 24.36666372 | 23.98893219 | 25.15131792 |
| THO complex subunit 6 homolog | Q86W42;Q86W42-3;Q86W42 | THOC6 | Other | -0.128785301 | 0.141925233 | 24.54333113 | 25.32865274 | 25.39868499 | 24.10701333 | 24.3454334 | 25.44110607 | 25.21652681 | 24.48486763 | 24.49436049 |
| Thimet oligopeptidase | P52888 | THOP1 | Other | 0.154556334 | 0.333851742 | 29.21442533 | 29.13726409 | 29.08377734 | 29.34276175 | 29.46892288 | 28.42221892 | 29.47593417 | 29.29524342 | 29.02717657 |
| Thyroid hormone receptor-associated | Q9Y2W1 | THRAP3 | Extracellular or secreted | -1.181883247 | 1.131299886 | 29.64742763 | 29.41354959 | 29.43932357 | 29.40051838 | 29.26008643 | 30.04335631 | 27.87067758 | 28.10769184 | 29.07811179 |
| Thrombospondin type-1 domain-cont | Q6ZMP0;Q6ZMP0-2;Q6ZMP0 | THSD4 | Extracellular or secreted | -0.829024039 | 0.378450973 | 26.31817261 | 22.45337313 | 22.52193106 | 21.90067945 | 24.04809968 | 24.19219648 | 21.89182078 | 22.2570989 | 24.08123441 |
| Thiamine-triphosphatase | Q9BU02 | THTPA | Other | 0.228099612 | 0.260597399 | 22.63158651 | 22.01947264 | 22.4626131 | 22.4318147 | 21.82178708 | 22.98848297 | 22.42193332 | 23.20563637 | 22.23460765 |
| THUMP domain-containing protein 1 | Q9NXG2 | THUMPD1 | Other | -0.188788812 | 0.341017793 | 30.45100826 | 30.51323993 | 30.50946941 | 29.87075106 | 29.81880442 | 30.17783413 | 30.2521435 | 30.19260568 | 29.65943798 |
| THUMP domain-containing protein 3 | Q9BV44 | THUMPD3 | Other | 1.304123768 | 1.239814559 | 23.14514915 | 23.51617035 | 23.89892165 | 24.7490895 | 24.7658247 | 24.41660891 | 26.19811152 | 24.9730306 | 24.98711132 |
| Thymocyte nuclear protein 1 | Q9P016;Q9P016-2 | THYN1 | Other | -0.585895508 | 2.795355491 | 25.73184727 | 25.83233879 | 25.64770896 | 25.16323137 | 25.48203833 | 25.68279125 | 24.88035628 | 25.10393607 | 25.02799911 |
| Nucleolysin TIA-1 isoform p40 | P31483;P31483-3 | TIA1 | Other | 0.018809067 | 0.036780715 | 26.84337997 | 26.77064635 | 26.63561052 | 26.51825826 | 26.42644815 | 26.91923051 | 26.85483662 | 26.85649981 | 26.40187766 |
| Nucleolysin TIAR | Q01085 | TIAL1 | Extracellular or secreted | -0.266065094 | 0.141886257 | 25.87065407 | 25.65880835 | 25.60399391 | 22.82110132 | 22.78844752 | 24.91693237 | 24.89790674 | 23.53883969 | 24.59502705 |
| Nucleolysin TIAR | Q01085-2 | TIAL1 | Extracellular or secreted | 0.122411148 | 0.773866608 | 28.03391176 | 28.03228346 | 27.94268398 | 28.00736858 | 27.99204835 | 28.20195735 | 28.1175342 | 28.27666329 | 28.0781627 |
| TIR domain-containing adapter molec | Q86XR7-2;Q9Y3B3-2;Q9Y3B3 | TICAM2;TMED7 | Extracellular or secreted | -0.251279601 | 0.154323613 | 23.62142681 | 23.67508354 | 23.75450262 | 21.07330694 | 22.20710165 | 25.34715824 | 22.81624463 | 22.9162126 | 23.35299387 |
| Fructose-2,6-bisphosphatase TIGAR | Q9NQ88 | TIGAR | Other | -0.008450924 | 0.013132276 | 27.15296235 | 27.41021686 | 27.6772765 | 27.52917423 | 27.45928014 | 27.17436159 | 27.66898494 | 27.47854018 | 27.02875796 |
| Mitochondrial import inner membran | P62072 | TIMM10 | Other | 0.407667446 | 0.289853839 | 24.10829026 | 23.74529846 | 24.00350281 | 25.29045709 | 25.64270551 | 21.73416497 | 24.80321511 | 24.38040346 | 24.30159332 |
| Mitochondrial import inner membran | Q9Y5L4 | TIMM13 | Other | 0.243219378 | 0.737384759 | 28.04291293 | 27.46396346 | 27.51073363 | 27.80341222 | 27.84020137 | 27.87267488 | 27.90454836 | 28.24299903 | 27.84906 |
| Mitochondrial import inner membran | O43615 | TIMM44 | Other | -0.36104491 | 0.756945486 | 25.24626509 | 25.10121302 | 25.19636769 | 24.88497351 | 24.60433333 | 25.90813842 | 24.72858155 | 25.09172236 | 24.56720689 |
| Mitochondrial import inner membran | Q3ZCQ8;Q3ZCQ8-2;Q3ZCQ8- | TIMM50 | Other | -0.113448293 | 0.138596374 | 23.61268045 | 24.05829369 | 24.28291421 | 23.36506221 | 23.08509831 | 24.90053481 | 23.63533375 | 23.59855234 | 24.07806088 |
| Mitochondrial import inner membran | O60220 | TIMM8A | Other | 0.204835362 | 0.183661479 | 24.80148931 | 24.50796977 | 23.81274372 | 24.94471814 | 25.20538281 | 23.76131667 | 25.39972827 | 24.28135844 | 24.45022959 |
| Mitochondrial import inner membran | Q9Y5J7 | TIMM9 | Other | -0.313553383 | 0.332325915 | 26.7649647 | 26.07647149 | 26.49879456 | 26.98457127 | 27.00998627 | 26.30110485 | 26.83376277 | 26.37754351 | 25.66598014 |
| Metalloproteinase inhibitor 1 | P01033 | TIMP1 | Extracellular or secreted | 0.902030107 | 0.601979563 | 30.19037013 | 29.89387176 | 29.79549154 | 31.60341919 | 31.55586142 | 33.12840119 | 31.28201057 | 31.66033942 | 32.84744795 |
| Metalloproteinase inhibitor 2 | P16035 | TIMP2 | Extracellular or secreted | -0.063824761 | 0.033660358 | 25.96622449 | 25.7166221 | 26.03298753 | 27.07265389 | 27.06845718 | 27.9808726 | 26.00187202 | 26.09071333 | 27.63484926 |
| Tubulointerstitial nephritis antigen-lik | Q9GZM7;Q9GZM7-3 | TINAGL1 | Extracellular or secreted | 1.116191649 | 0.854806697 | 27.01382408 | 27.18574686 | 27.27364324 | 28.34218951 | 28.40186139 | 29.07183598 | 28.42977954 | 28.56946133 | 29.9938846 |
| TIMELESS-interacting protein | Q9BVW5 | TIPIN | Other | 1.328283759 | 2.194373053 | 22.43055975 | 22.74847542 | 22.92411949 | 22.84781589 | 22.2434204 | 20.97479046 | 23.63300673 | 23.73044858 | 23.70598667 |
| TIP41-like protein | O75663;O75663-2 | TIPRL | Other | -0.078581311 | 0.118368366 | 29.92365718 | 29.98617289 | 29.87381914 | 29.65429073 | 29.56621629 | 29.17195446 | 29.79895543 | 29.84522096 | 29.20813502 |
| Tight junction protein ZO-1 | Q07157-2;Q07157 | TJP1 | Extracellular or secreted | 0.474396213 | 1.234605452 | 29.16065173 | 29.22232471 | 29.20106932 | 28.92886098 | 28.93275154 | 29.21463376 | 29.49944861 | 29.39844036 | 29.85544569 |
| Tight junction protein ZO-2 | Q9UDY2-3;Q9UDY2;Q9UDY2- | TJP2 | Extracellular or secreted | 0.604916422 | 1.876272768 | 27.97257056 | 27.98912715 | 27.85400431 | 27.71979749 | 27.77448456 | 27.30686871 | 28.51795803 | 28.47289755 | 28.13232008 |
| Tight junction protein ZO-3 | O95049-5;O95049;O95049-4 | TJP3 | Other | -0.568302215 | 0.398008547 | 23.84114969 | 23.38429578 | 23.41673775 | 21.41738562 | 22.2405992 | 22.49196561 | 22.91763358 | 22.47956122 | 21.29396538 |
| Thymidine kinase, cytosolic | P04183 | TK1 | Other | 0.546274396 | 0.811410041 | 25.79096669 | 25.36719716 | 25.65529094 | 25.15375488 | 24.89864493 | 25.30966397 | 26.30943847 | 25.99369028 | 25.42345373 |
| Transketolase | P29401;P29401-2 | TKT | Extracellular or secreted | -0.541211344 | 0.976793824 | 33.21022129 | 33.28719496 | 33.19640698 | 33.36814999 | 33.40641524 | 32.48896047 | 32.82147175 | 32.84331632 | 32.19025237 |
| TLD domain-containing protein 1 | Q6P9B6 | TLDC1 | Other | 0.409896172 | 1.203268206 | 26.38465808 | 26.51174421 | 26.54833794 | 25.99438106 | 25.75865082 | 25.55727741 | 26.5072137 | 26.4540481 | 26.64595148 |
| Transducin-like enhancer protein 1;Tr | Q04724;Q04727-2;Q04727-4 | TLE1;TLE4 | Other | 0.656032571 | 0.541295271 | 21.36649103 | 23.84345186 | 21.05002786 | 23.71250853 | 23.64493446 | 23.13951098 | 23.71313811 | 23.67422158 | 22.95920037 |
| Transducin-like enhancer protein 3 | Q04726-2;Q04726-7;Q04726 | TLE3 | Other | -0.619688736 | 0.775559019 | 25.87832282 | 25.86050733 | 25.5034577 | 25.61882526 | 25.75488485 | 25.92390648 | 25.56107091 | 25.28778282 | 24.56203229 |
| Talin-1 | Q9Y490 | TLN1 | Extracellular or secreted | -0.321364585 | 0.278276047 | 32.04391033 | 31.93364517 | 31.93026476 | 31.96574851 | 32.05827304 | 31.14710157 | 31.93079348 | 31.94980353 | 30.69478094 |
| Toll-like receptor 5 | O60602 | TLR5 | Extracellular or secreted | -1.622333421 | 0.693370775 | 23.84153364 | 24.41021686 | 24.22486547 | 25.74742575 | 24.0986856 | 23.34177397 | 21.09847201 | 24.19580472 | 22.67097365 |
| Transmembrane 9 superfamily memb | Q99805 | TM9SF2 | Extracellular or secreted | -1.216088702 | 1.534916508 | 25.59097942 | 25.58677727 | 25.44325798 | 25.94008683 | 25.83826679 | 26.1279789 | 24.28340888 | 24.26892287 | 25.06307574 |
| Transmembrane 9 superfamily memb | Q9HD45 | TM9SF3 | Other | -1.409253815 | 0.611232485 | 23.89513517 | 24.41525538 | 24.10996452 | 23.8624951 | 24.058624 | 23.83942064 | 23.2649923 | 20.90441431 | 23.69327935 |
| Transmembrane 9 superfamily memb | Q92544 | TM9SF4 | Other | -1.482305283 | 1.851629011 | 25.04685156 | 24.96000511 | 24.93979548 | 24.90894009 | 24.99606345 | 25.79753686 | 23.35863832 | 23.36212148 | 24.15692063 |
| Translation machinery-associated pro | Q96EY4 | TMA16 | Other | -0.042937065 | 0.021843657 | 21.32498291 | 23.0632239 | 22.93788455 | 23.22100469 | 23.22911218 | 23.67042903 | 23.37914777 | 23.46687505 | 21.74848461 |
| Translation machinery-associated pro | Q9Y2S6 | TMA7 | Other | -1.566716032 | 0.359088692 | 28.65707117 | 24.56470912 | 24.01273774 | 25.84606134 | 22.30500227 | 26.3933263 | 20.65123392 | 25.04481065 | 25.49326129 |
| Protein lifeguard 3 | Q969X1 | TMBIM1 | Extracellular or secreted | 2.153976366 | 2.74727358 | 22.72442392 | 22.47550981 | 22.66561697 | 23.06138093 | 21.89701437 | 24.46948582 | 24.64091435 | 24.94975762 | 25.51797304 |
| Transmembrane and coiled-coil doma | Q9UM00-2;Q9UM00 | TMCO1 | Other | 0.78026457 | 0.482466221 | 24.35070291 | 23.69763362 | 24.6741138 | 21.8796737 | 20.85055681 | 25.28132306 | 23.9461024 | 24.16353851 | 24.59815475 |
| Transmembrane emp24 domain-cont | P49755 | TMED10 | Extracellular or secreted | -0.788012602 | 0.584452865 | 24.18412047 | 24.38640248 | 24.58319435 | 24.82987368 | 25.41341643 | 26.81860636 | 23.64306348 | 23.91734221 | 25.18336339 |
| Transmembrane emp24 domain-cont | Q15363 | TMED2 | Extracellular or secreted | 0.246932028 | 0.084727325 | 24.60597272 | 24.67943929 | 24.6863492 | 24.20791634 | 20.0118858 | 26.86985436 | 23.65512713 | 24.16936171 | 25.44701611 |
| Protein TMED8 | Q6PL24 | TMED8 | Other | 0.386837271 | 0.472227912 | 26.29813558 | 25.95766038 | 26.24648257 | 25.51517791 | 26.12390084 | 25.49378046 | 26.59377407 | 26.6676322 | 25.71667442 |
| Transmembrane emp24 domain-cont | Q9BVK6 | TMED9 | Extracellular or secreted | -1.180517571 | 0.971147805 | 24.69710332 | 24.77600444 | 25.01771541 | 25.16840556 | 25.14988478 | 26.62867514 | 23.64976578 | 23.53112441 | 24.99645142 |
| Transmembrane protein 132A | Q24JP5;Q24JP5-2;Q24JP5-4 | TMEM132A | Extracellular or secreted | 0.89065871 | 1.402488182 | 26.97103601 | 27.41183369 | 27.31855191 | 27.92897389 | 27.99139971 | 28.57596567 | 28.33964351 | 28.36565464 | 29.06555842 |
| Transmembrane protein 2 | Q9UHN6-2;Q9UHN6 | TMEM2 | Extracellular or secreted | 1.410003369 | 0.748881511 | 21.6806308 | 21.36602333 | 21.16021213 | 24.94882241 | 24.54751448 | 23.14320382 | 23.19583475 | 24.0873178 | 25.37006104 |
| Transmembrane protein 205 | Q6UW68 | TMEM205 | Extracellular or secreted | 1.239151472 | 0.399838122 | 21.56840811 | 20.55741872 | 20.18753975 | 24.0129082 | 24.23454557 | 27.45904558 | 23.29963845 | 23.62544577 | 25.80230316 |
| Transmembrane protein 33 | P57088 | TMEM33 | Extracellular or secreted | -0.030589946 | 0.012405136 | 23.63189731 | 23.98737158 | 24.34982599 | 23.14382661 | 23.38679714 | 27.277905 | 23.48480618 | 23.97491307 | 25.33732273 |
| Transmembrane protein 43 | Q9BTV4 | TMEM43 | Other | -0.935906993 | 0.210809124 | 23.77293792 | 24.10317573 | 25.17588483 | 23.61223049 | 22.7453395 | 28.20251793 | 21.28748694 | 23.30131421 | 26.40952107 |
| CSC1-like protein 1 | O94886 | TMEM63A | Extracellular or secreted | -0.299732493 | 0.261038476 | 22.07646288 | 21.12066567 | 21.04466893 | 22.8283515 | 22.52650776 | 21.30229084 | 22.11690261 | 20.91612146 | 21.51725224 |
| Transmembrane protein 87A | Q8NBN3-3;Q8NBN3 | TMEM87A | Extracellular or secreted | 0.013800098 | 0.017433454 | 23.73138119 | 24.08180328 | 24.2158606 | 24.28249008 | 24.2545062 | 25.27398109 | 24.31118953 | 24.03389076 | 24.61633121 |
| TATA element modulatory factor | P82094;P82094-2 | TMF1 | Other | -0.533921072 | 1.750471015 | 24.59588073 | 24.64322866 | 24.7663809 | 25.08756061 | 25.25241424 | 25.07312398 | 24.15676636 | 24.397086 | 24.55367897 |
| Tropomodulin-1 | P28289-2;P28289 | TMOD1 | Other | -0.363885364 | 0.764048587 | 24.53118392 | 24.41170441 | 24.37643438 | 24.13539617 | 23.70746193 | 24.29698116 | 24.078631 | 24.04168121 | 23.51761269 |
| Tropomodulin-3 | Q9NYL9 | TMOD3 | Other | 0.376939237 | 1.038961461 | 29.10714307 | 28.97836518 | 29.03194181 | 29.11421121 | 29.34165947 | 28.94910081 | 29.24918468 | 29.42698393 | 29.71585988 |
| Lamina-associated polypeptide 2, isof | P42166 | TMPO | Other | -1.173876813 | 2.975485868 | 31.34810223 | 31.26634464 | 31.28554239 | 31.46036451 | 31.60743742 | 31.09024137 | 30.33052598 | 30.21697079 | 29.95988907 |
| Lamina-associated polypeptide 2, isof | P42167;P42167-2;P42167-3 | TMPO | Other | -0.915456717 | 5.008247724 | 27.8117643 | 27.91718284 | 27.99377665 | 28.00688726 | 28.03286146 | 27.68096875 | 27.0480165 | 27.03044286 | 26.89689112 |
| Thymosin beta-10 | P63313 | TMSB10 | Other | 0.371688458 | 0.076771398 | 26.68284482 | 25.57192921 | 25.65706435 | 26.50754642 | 26.6673615 | 22.34482629 | 27.26356035 | 27.64663825 | 22.92065307 |
| Thymosin beta-4;Hematopoietic syste | P62328 | TMSB4X | Extracellular or secreted | -0.794044422 | 0.141251839 | 28.24798588 | 27.68919192 | 27.70140641 | 28.15947326 | 28.15518419 | 22.36733847 | 27.97486932 | 28.16633811 | 22.63694937 |
| Thioredoxin-related transmembrane p | Q9H3N1 | TMX1 | Other | -2.02452252 | 0.618937242 | 24.55221362 | 25.77467307 | 24.9791942 | 23.90223819 | 23.89550502 | 26.03386976 | 20.42508517 | 23.4801285 | 24.59006571 |
| Thioredoxin-related transmembrane p | Q9Y320-2;Q9Y320 | TMX2 | Extracellular or secreted | -2.812115957 | 2.150888922 | 21.29887892 | 22.57981279 | 18.12120972 | 22.38624714 | 21.93528852 | 21.34104097 | 18.08441395 | 18.39527472 | 18.91520249 |
| Protein disulfide-isomerase TMX3 | Q96JJ7 | TMX3 | Extracellular or secreted | -0.505860389 | 0.545632482 | 24.31989589 | 23.82463949 | 24.20179848 | 23.65534554 | 23.67852621 | 25.56981621 | 23.52945724 | 23.27540274 | 24.30256976 |
| Thioredoxin-related transmembrane p | Q9H1E5 | TMX4 | Other | -0.591784252 | 0.559522694 | 22.96470961 | 22.66309941 | 22.7994848 | 20.08887922 | 22.13564667 | 23.14201978 | 21.74613018 | 21.38853373 | 21.98690307 |
| Tenascin | P24821;P24821-3;P24821-5;P | TNC | Extracellular or secreted | 3.229857604 | 0.898520804 | 25.53978642 | 25.15448902 | 25.33036356 | 22.58943719 | 23.72077614 | 31.43151409 | 27.49975272 | 27.63478004 | 31.43822327 |
| Tumor necrosis factor alpha-induced p | Q03169 | TNFAIP2 | Extracellular or secreted | 0.381665589 | 0.39906981 | 25.88736957 | 25.91645408 | 26.33022677 | 25.9992288 | 26.69657282 | 26.84218132 | 26.32421228 | 26.29678867 | 27.3600125 |
| Tumor necrosis factor alpha-induced p | O95379-3;O95379-2;O95379 | TNFAIP8 | Other | 0.151942348 | 0.271764761 | 26.45956157 | 25.94683865 | 26.65758232 | 27.04364312 | 26.75085355 | 26.87455217 | 26.44010815 | 26.94100532 | 26.94122926 |
| Tumor necrosis factor receptor supert | O00220 | TNFRSF10A | Extracellular or secreted | 2.213788816 | 2.273701605 | 21.90428425 | 24.14063196 | 23.84966853 | 24.6371042 | 24.7830178 | 23.12653812 | 25.41235069 | 25.89762982 | 26.55200836 |
| Tumor necrosis factor receptor supert | O14763-2;O14763 | TNFRSF10B | Extracellular or secreted | 2.226999101 | 2.446293501 | 23.45324752 | 23.47345506 | 23.36305782 | 23.56836717 | 24.10789134 | 20.93890954 | 25.27600652 | 25.18563345 | 25.67182156 |
| 182 kDa tankyrase-1-binding protein | Q9C0C2 | TNKS1BP1 | Extracellular or secreted | -0.08886403 | 0.259071276 | 29.06791891 | 28.84547522 | 28.80826968 | 28.68398267 | 28.80029854 | 28.26972553 | 28.52436195 | 28.61610682 | 28.83077442 |
| Transportin-1 | Q92973-2;Q92973;Q92973-3 | TNPO1 | Extracellular or secreted | 0.130037359 | 0.207726703 | 28.60399391 | 28.64602017 | 28.64337318 | 28.47417487 | 28.54107977 | 28.33836882 | 28.9714198 | 28.80347381 | 28.23872383 |
| Transportin-2 | O14787-2;O14787 | TNPO2 | Other | 0.158183951 | 0.292659804 | 29.09629955 | 29.35053853 | 29.18491971 | 28.56656108 | 28.66438044 | 28.85854646 | 29.34161705 | 29.23544062 | 28.75811706 |
| Transportin-3 | Q9Y5L0-3;Q9Y5L0-1;( | TNPO3 | Other | 0.306310102 | 1.251191021 | 28.35333888 | 28.48362281 | 28.50861968 | 27.99991693 | 28.05227301 | 28.80538177 | 28.64027313 | 28.70305804 | 28.67717567 |
| Tensin-3 | Q68CZ2-2;Q68CZ2 | TNS3 | Other | -0.238212718 | 0.086113273 | 23.52384619 | 22.40174097 | 23.16154095 | 20.45124677 | 22.8006248 | 23.44594336 | 23.2300656 | 23.24890158 | 20.69886619 |
| Target of EGR1 protein 1 | Q96GM8-2;Q96GM8 | TOE1 | Other | 0.791844683 | 1.122248943 | 22.55349316 | 21.43775012 | 22.49059123 | 23.33701641 | 23.28135844 | 23.90903168 | 23.53694437 | 23.63311763 | 23.71009257 |
| Toll-interacting protein | Q9H0E2;Q9H0E2-2 | TOLLIP | Other | 1.019036121 | 4.675117166 | 23.80616884 | 23.57403911 | 23.78316771 | 23.73933563 | 24.10821049 | 23.49399418 | 24.79367276 | 24.67201047 | 24.84388311 |
| Target of Myb protein 1 | O60784;O60784-2;O60784-4;( | TOM1 | Extracellular or secreted | -0.147848672 | 0.288745319 | 23.73272723 | 23.8772699 | 23.7717467 | 23.90260623 | 23.85052719 | 22.59150299 | 23.55250681 | 23.38099788 | 23.45913941 |
| TOM1-like protein 2 | Q6ZVM7;Q6ZVM7-5;Q6ZVM7 | TOM1L2 | Extracellular or secreted | 1.300333361 | 2.093957981 | 26.02007063 | 25.99987393 | 26.52766885 | 26.12486722 | 24.28665533 | 25.71748524 | 27.25078907 | 26.83797819 | 27.15054343 |
| Mitochondrial import receptor subun | Q15388 | TOMM20 | Other | -0.062836017 | 0.024243563 | 23.06803888 | 23.23131143 | 23.50020876 | 20.59884822 | 22.16174244 | 23.2831969 | 22.96643575 | 21.15765101 | 23.60907681 |
| Mitochondrial import receptor subun | Q9NS69 | TOMM22 | Other | -1.317818678 | 0.881253489 | 25.75447713 | 26.22306162 | 26.36024698 | 26.03841937 | 26.07324658 | 26.83881979 | 23.92651028 | 24.92306774 | 25.84110168 |
| Mitochondrial import receptor subun | Q15785 | TOMM34 | Other | 0.230626648 | 0.311231946 | 23.06032388 | 23.15169728 | 23.56406946 | 23.7665831 | 23.46799453 | 24.87970215 | 23.57242083 | 23.92424638 | 24.14039793 |
| Mitochondrial import receptor subun | O96008;O96008-2 | TOMM40 | Extracellular or secreted | 0.189024265 | 0.05148807 | 21.4135391 | 21.09636957 | 18.99681232 | 21.65226739 | 23.24130476 | 27.19891709 | 21.61443737 | 22.06098348 | 23.69125706 |
| Mitochondrial import receptor subun | O94826 | TOMM70A | Extracellular or secreted | -0.999611707 | 3.789063091 | 27.32276919 | 27.38434519 | 27.48761473 | 27.27861407 | 27.50104447 | 28.04390381 | 26.52202089 | 26.42908514 | 26.55920459 |
| DNA topoisomerase 1 | P11387 | TOP1 | Other | 0.157936019 | 0.09035552 | 29.83764742 | 29.84233121 | 29.80379711 | 29.68560423 | 29.79215898 | 30.55096324 | 30.69718619 | 30.60805461 | 28.92481835 |
| DNA topoisomerase 2-alpha | P11388;P11388-2;P11388-3;( | TOP2A | Other | -1.560078528 | 3.457681559 | 30.15617325 | 30.16470887 | 30.21326682 | 29.68902525 | 29.63618121 | 31.03845861 | 28.7104342 | 28.60349878 | 28.45473844 |
| DNA topoisomerase 2-beta | Q02880-2;Q02880 | TOP2B | Other | -1.649807472 | 2.176046993 | 27.58218954 | 27.62846657 | 27.3569943 | 27.25799806 | 27.13907103 | 27.74727207 | 25.86582557 | 26.13282995 | 25.40791785 |
| DNA topoisomerase 3-alpha | Q13472-3;Q13472-2;Q13472 | TOP3A | Other | -0.950953396 | 0.506845259 | 24.50263857 | 24.52981466 | 24.46270046 | 23.97438798 | 23.91186812 | 24.21474955 | 24.20955332 | 23.82211254 | 21.91355362 |
| Torsin-1A-interacting protein 1 | Q5JTV8-3;Q5JTV8;Q5JTV8-2 | TOR1AIP1 | Other | -0.680709333 | 1.191156682 | 26.86608507 | 26.69032478 | 26.82590131 | 26.79389597 | 26.73582868 | 28.07127339 | 25.9943379 | 26.25028309 | 26.70490562 |
| Torsin-1A-interacting protein 2 | Q8NFQ8 | TOR1AIP2 | Other | -0.900296494 | 1.72251336 | 25.42178568 | 26.08048234 | 25.95876686 | 25.67968089 | 25.28580913 | 24.84704167 | 24.30236058 | 24.64438442 | 24.98914881 |
| Torsin-1B | O14657 | TOR1B | Extracellular or secreted | 1.680794453 | 1.253060308 | 19.8972259 | 23.25565909 | 23.16038726 | 21.12459193 | 23.3340179 | 23.85186187 | 23.84306842 | 23.67465262 | 24.83653429 |
| TOX high mobility group box family m | O94842-3;O94842-2;O94842 | TOX4 | Other | -0.348068945 | 0.417737106 | 24.76228024 | 24.51376325 | 24.31402832 | 25.37272002 | 25.29119515 | 25.05911931 | 23.94449474 | 24.747144 | 24.92070758 |
| Cellular tumor antigen p53 | P04637-4;P04637;P04637-7;( | TP53 | Other | 0.327021962 | 0.230942625 | 26.07159051 | 26.10858938 | 26.05885104 | 26.27517182 | 26.33290891 | 27.5791709 | 26.26926182 | 26.22466311 | 27.70028223 |
| Tumor suppressor p53-binding protei | Q12888;Q12888-2;Q12888-3 | TP53BP1 | Other | -0.551841005 | 1.696605195 | 28.6896585 | 28.77746628 | 28.7756466 | 28.14146053 | 28.174933 | 28.36168654 | 28.08810676 | 28.0240086 | 27.69278735 |
| Quinone oxidoreductase PIG3 | Q53FA7 | TP53I3 | Extracellular or secreted | 0.396321524 | 1.053960234 | 26.77994113 | 27.09412173 | 26.86938374 | 26.5791709 | 26.65129963 | 25.84997866 | 27.23022688 | 26.97377513 | 26.86691045 |
| TP53-regulating kinase | Q96S44 | TP53RK | Other | -0.402065928 | 0.899551307 | 24.1346914 | 24.66828167 | 24.41034628 | 24.4252159 | 25.04626874 | 25.40103131 | 24.05183756 | 24.50621507 | 24.27866724 |

| | | | | | | | | | | | | | | |
|---|---|---|---|---|---|---|---|---|---|---|---|---|---|---|
| Trophoblast glycoprotein | Q13641 | TPBG | Extracellular or secreted | 0.617494765 | 0.889353378 | 25.25475847 | 25.76994052 | 25.74458012 | 25.60588797 | 25.68471845 | 25.37550668 | 25.81994653 | 26.05924312 | 26.69099076 |
| Tumor protein D52 | P55327-2;P55327-4;P55327;F | TPD52 | Extracellular or secreted | 0.143138272 | 0.354408433 | 28.37343379 | 28.53978642 | 28.63502231 | 28.64972467 | 29.11237285 | 27.96636753 | 28.57484757 | 28.63609475 | 28.85682628 |
| Tumor protein D53 | Q16890-4;Q16890-3;Q16890 | TPD52L1 | Other | 0.78001983 | 0.481900694 | 21.2360212 | 24.20478604 | 21.71020669 | 24.3273169 | 24.49381099 | 24.4646958 | 24.63350569 | 24.57467436 | 23.35029825 |
| Tumor protein D54 | O43399;O43399-5;O43399-7 | TPD52L2 | Other | 0.613284814 | 1.966871633 | 29.24893596 | 29.1716682 | 29.15588384 | 28.92555421 | 28.94912865 | 29.21988099 | 29.70005067 | 29.56105269 | 29.914277 |
| Tumor protein D54 | O43399-2;O43399-4;O43399 | TPD52L2 | Other | -0.178748614 | 0.027712193 | 25.97622497 | 21.81875012 | 20.76768302 | 26.13625708 | 25.90524872 | 26.51998339 | 26.09490628 | 26.32798497 | 20.60293656 |
| Triosephosphate isomerase | P60174-1;P60174;P60174-4 | TPI1 | Extracellular or secreted | -0.262470028 | 0.652048485 | 35.24278471 | 35.35264719 | 35.32244284 | 35.24593939 | 35.37974942 | 34.27981261 | 34.97954448 | 34.95280884 | 34.69192468 |
| Tropomyosin alpha-3 chain | P06753-2;P06753-3;P06753-5 | TPM3 | Extracellular or secreted | -0.025619323 | 0.053824573 | 33.72882479 | 33.76742161 | 33.78156001 | 34.31108232 | 34.42047566 | 33.4431977 | 33.84277328 | 33.79110336 | 34.01554644 |
| Tropomyosin alpha-3 chain | P06753-6 | TPM3 | Extracellular or secreted | 1.18087381 | 1.173865951 | 21.4239459 | 22.67912424 | 21.2284863 | 24.20000296 | 23.97910696 | 22.82526083 | 23.85005022 | 24.01000762 | 23.85052719 |
| Tropomyosin alpha-4 chain | P67936 | TPM4 | Extracellular or secreted | 0.52697073 | 2.344125852 | 31.18435225 | 31.21239732 | 31.21506212 | 31.69673033 | 31.72817606 | 31.10230736 | 31.96605806 | 31.79838384 | 31.88598301 |
| Tropomyosin alpha-4 chain | P67936-2 | TPM4 | Extracellular or secreted | 0.751395999 | 2.33839447 | 27.40072194 | 27.70457587 | 27.66471951 | 27.41522314 | 27.57188582 | 27.7602763 | 28.41078296 | 28.13794806 | 28.46415827 |
| Thiopurine S-methyltransferase | P51580 | TPMT | Extracellular or secreted | 0.400572518 | 0.606948427 | 27.90890574 | 26.88727659 | 26.62030844 | 26.26153529 | 25.95233743 | 27.07643071 | 26.89099099 | 27.3569943 | 27.30712936 |
| Tripeptidyl-peptidase 1 | O14773;O14773-2 | TPP1 | Extracellular or secreted | 0.473934967 | 0.514614367 | 24.64801081 | 25.05415847 | 25.55821212 | 25.60887948 | 25.98676422 | 26.68204109 | 25.5912363 | 25.97009255 | 26.62950915 |
| Tripeptidyl-peptidase 2 | P29144 | TPP2 | Other | 0.075471829 | 0.22966337 | 29.50780725 | 29.6383581 | 29.56657922 | 29.24284007 | 29.30384599 | 29.32270473 | 29.60059533 | 29.62844918 | 29.28843865 |
| Tubulin polymerization-promoting pro | O94811 | TPPP | Other | -0.622536718 | 0.50486494 | 22.96657656 | 24.41892628 | 22.16257081 | 24.19370101 | 23.74960104 | 21.48802251 | 22.84435254 | 22.8686681 | 21.90906832 |
| Nucleoprotein TPR | P12270 | TPR | Other | -0.827199804 | 2.13845549 | 31.47392336 | 31.43604 | 31.44618401 | 31.44114566 | 31.56483629 | 31.35499967 | 30.48289232 | 30.60071938 | 30.79335338 |
| EKC/KEOPS complex subunit TPRKB | Q9Y3C4;Q9Y3C4-3;Q9Y3C4-2 | TPRKB | Other | 0.094826855 | 0.149793661 | 25.44015568 | 25.40616685 | 25.49524534 | 25.57065599 | 25.64883371 | 24.66822756 | 25.57340357 | 25.72530842 | 25.10041114 |
| Translationally-controlled tumor prot | P13693;P13693-2 | TPT1 | Extracellular or secreted | 0.093028644 | 0.145457428 | 31.23837079 | 31.21777977 | 31.18736433 | 31.72959478 | 31.93019426 | 30.61500186 | 31.52234273 | 31.6087156 | 31.10718049 |
| Targeting protein for Xklp2 | Q9ULW0;Q9ULW0-2 | TPX2 | Other | -0.806599271 | 0.33017755 | 23.63511229 | 23.94064696 | 24.07414538 | 24.23205841 | 24.49911402 | 22.96557302 | 22.29184629 | 22.07903752 | 24.88264342 |
| Transformer-2 protein homolog alpha | Q13595-4;Q13595-3;Q13595 | TRA2A | Other | -0.087338927 | 0.053098407 | 24.9850275 | 25.07324658 | 25.25093361 | 25.32858427 | 25.55581572 | 27.02854721 | 25.04776695 | 24.9103591 | 26.39093461 |
| Transformer-2 protein homolog beta | P62995;P62995-3 | TRA2B | Other | -0.307245937 | 0.161436658 | 27.50741032 | 27.69909096 | 27.41956934 | 27.82493078 | 27.95199298 | 29.2490038 | 26.96366941 | 27.03443662 | 28.90615524 |
| Tumor necrosis factor receptor type 1 | Q15628-2;Q15628 | TRADD | Extracellular or secreted | -0.705755654 | 0.935616952 | 22.85110073 | 20.69308239 | 22.21949177 | 22.95921265 | 22.38108818 | 22.98526875 | 21.32256635 | 21.78755827 | 21.81723065 |
| TNF receptor-associated factor 6 | Q9Y4K3 | TRAF6 | Extracellular or secreted | -0.089949817 | 0.417294378 | 23.72920416 | 24.09980944 | 24.19663034 | 24.25939962 | 24.40041251 | 24.10980515 | 24.03229398 | 24.08334622 | 24.01214096 |
| TRAF-type zinc finger domain-contain | O14545 | TRAFD1 | Other | -0.126033452 | 0.123161501 | 22.35065502 | 21.66344692 | 21.46298746 | 22.36663704 | 21.66062096 | 21.79066349 | 21.1431981 | 22.27309896 | 21.85310803 |
| Heat shock protein 75 kDa, mitochon | Q12931;Q12931-2 | TRAP1 | Extracellular or secreted | -0.06538644 | 0.117775201 | 29.37434623 | 29.61101348 | 29.6371664 | 30.03813151 | 30.00651279 | 30.69162314 | 29.82239522 | 29.97350145 | 29.68734079 |
| Trafficking protein particle complex su | Q9Y5R8 | TRAPPC1 | Other | -0.320080889 | 0.625774163 | 22.70474207 | 21.9921694 | 22.31387614 | 21.42429916 | 22.26192618 | 22.18474098 | 22.16671353 | 21.65796386 | 21.65595691 |
| Trafficking protein particle complex su | P48553 | TRAPPC10 | Other | -0.403120442 | 0.550506193 | 21.31257501 | 21.24400141 | 22.09767339 | 22.04484212 | 22.4233426 | 20.69776284 | 21.23648841 | 21.60715871 | 20.85709023 |
| Trafficking protein particle complex su | Q9UL33-2;Q9UL33 | TRAPPC2L | Other | 1.422006362 | 1.196387476 | 23.76091076 | 21.56977934 | 21.47792057 | 20.68818593 | 20.87451576 | 23.80725035 | 23.88455437 | 22.8927844 | 23.57796167 |
| Trafficking protein particle complex su | O43617;O43617-2 | TRAPPC3 | Other | -0.016026899 | 0.018145548 | 26.83617309 | 26.81982475 | 26.94603545 | 26.64814799 | 26.82177204 | 26.19946013 | 26.93258261 | 26.98522299 | 26.16982043 |
| Trafficking protein particle complex su | Q96Q05-3;Q96Q05;Q96Q05- | TRAPPC9 | Other | 1.104118822 | 0.827959299 | 22.00422321 | 22.19165446 | 20.68859182 | 21.84388708 | 21.97598551 | 21.54184341 | 22.91741506 | 23.63876202 | 21.87927213 |
| Tripartite motif-containing protein 16 | O95361;O95361-2 | TRIM16 | Other | 1.526327774 | 1.486921571 | 27.10439608 | 26.6896585 | 27.25718884 | 26.48660244 | 26.58434185 | 23.7716289 | 27.88820609 | 27.98809791 | 27.64958763 |
| Tripartite motif-containing protein 2 | Q9C040;Q9C040-2 | TRIM2 | Other | 1.239571972 | 0.81087033 | 24.8200196 | 24.7815178 | 24.83547451 | 21.00632301 | 21.39262312 | 24.0944231 | 25.02009183 | 24.59017995 | 24.57363471 |
| E3 ubiquitin-protein ligase TRIM23 | P36406-3;P36406-2;P36406 | TRIM23 | Extracellular or secreted | 0.279872525 | 0.213219891 | 20.33766417 | 21.35197012 | 21.51758866 | 21.30000169 | 22.145569 | 21.23637163 | 21.45457843 | 20.9129098 | 22.41671198 |
| Transcription intermediary factor 1-al | O15164-2;O15164 | TRIM24 | Extracellular or secreted | 1.462029436 | 1.463612678 | 22.70807267 | 20.5342961 | 22.33409976 | 21.72617969 | 20.73181001 | 23.32896084 | 23.95491265 | 23.26770911 | 22.84517608 |
| E3 ubiquitin/ISG15 ligase TRIM25 | Q14258 | TRIM25 | Other | -0.255196465 | 1.010111054 | 28.14622768 | 28.22131083 | 28.22609742 | 27.57174119 | 27.78749599 | 28.27510964 | 27.75951459 | 27.89942884 | 27.68945856 |
| Transcription intermediary factor 1-be | Q13263;Q13263-2 | TRIM28 | Extracellular or secreted | 0.488918424 | 0.664263868 | 30.96133891 | 30.93297676 | 30.87122124 | 31.22020398 | 31.24937685 | 31.05930501 | 31.80850752 | 31.82376983 | 30.9816893 |
| Tripartite motif-containing protein 29 | Q14134-2;Q14134 | TRIM29 | Other | 0.560534822 | 0.32315129 | 27.43093614 | 26.86147789 | 26.75595457 | 26.08814721 | 24.57033751 | 28.5896372 | 26.61386094 | 26.9763998 | 28.23958899 |
| E3 ubiquitin-protein ligase TRIM33 | Q9UPN9-2;Q9UPN9 | TRIM33 | Other | -0.325068292 | 0.540853901 | 25.98183081 | 26.52288894 | 26.23690075 | 26.92286365 | 26.62212535 | 27.18224597 | 26.31027092 | 25.86992494 | 26.579027 |
| Tripartite motif-containing protein 47 | Q96LD4;Q96LD4-2 | TRIM47 | Other | 0.688848692 | 1.098522962 | 24.34374038 | 25.00881157 | 25.32048134 | 24.92854476 | 25.17641758 | 24.28086307 | 25.99828207 | 25.4120599 | 25.18563345 |
| E3 ubiquitin-protein ligase TRIM56 | Q9BRZ2 | TRIM56 | Other | -0.858926147 | 1.667687823 | 21.35906747 | 22.24106493 | 22.48438827 | 20.98820706 | 22.38083274 | 21.497117 | 20.86499013 | 21.24266474 | 20.79090542 |
| Tripartite motif-containing protein 65 | Q6PJ69 | TRIM65 | Other | 2.631978821 | 1.614781183 | 21.80986229 | 24.00041132 | 20.46353408 | 21.9086624 | 22.27642616 | 23.13408033 | 25.92938033 | 24.98819545 | 23.77484898 |
| TRIO and F-actin-binding protein | Q9H2D6-6;Q9H2D6-5;Q9H2D | TRIOBP | Other | -1.1641098 | 0.580768848 | 24.3098721 | 24.14421572 | 23.82347376 | 23.36426079 | 23.61313028 | 23.7427313 | 23.08589253 | 21.19692493 | 23.7236951 |
| Cdc42-interacting protein 4 | Q15642-2;Q15642-4;Q15642 | TRIP10 | Extracellular or secreted | -0.286988383 | 0.868031762 | 29.95670813 | 29.89032426 | 29.77525412 | 29.65157336 | 29.65758232 | 28.93086396 | 29.40659264 | 29.45757868 | 29.20601661 |
| Thyroid receptor-interacting protein 1 | Q15643 | TRIP11 | Other | -1.459675667 | 0.497374126 | 26.47861732 | 26.25747661 | 26.39785308 | 25.49118274 | 25.45346733 | 26.01854336 | 22.84529096 | 24.18767345 | 26.63657882 |
| E3 ubiquitin-protein ligase TRIP12 | Q14669-3;Q14669;Q14669-2 | TRIP12 | Other | -1.633870099 | 2.802286551 | 25.33759495 | 25.47051018 | 24.98115578 | 25.51728228 | 23.67766631 | 24.77107472 | 23.44442752 | 23.38139403 | 23.15021026 |
| Pachytene checkpoint protein 2 homo | Q15645 | TRIP13 | Other | 0.318243133 | 0.578856514 | 27.03653417 | 27.10739254 | 27.10239497 | 27.26284384 | 27.48354593 | 27.43928394 | 27.59455731 | 27.89942884 | 27.17674094 |
| Thyroid receptor-interacting protein 6 | Q15654 | TRIP6 | Extracellular or secreted | 0.283941128 | 0.474072569 | 27.38566219 | 27.43991799 | 27.22361406 | 27.77912709 | 27.81011001 | 27.63360269 | 28.12075069 | 27.97978295 | 27.38730676 |
| tRNA (guanine(26)-N(2))-dimethyltran | Q9NXH9;Q9NXH9-2 | TRMT1 | Other | -0.091000805 | 0.700746836 | 27.23635338 | 27.05465532 | 27.11406224 | 27.08780337 | 27.40226811 | 27.32585976 | 27.13897341 | 27.14180156 | 27.0567237 |
| Mitochondrial ribonuclease P protein | Q7L0Y3 | TRMT10C | Other | -0.113877329 | 0.052210314 | 21.52773674 | 20.80990736 | 21.36062207 | 21.91776907 | 22.06391509 | 23.94440538 | 21.23456018 | 21.1926179 | 23.04336778 |
| tRNA (guanine(10)-N2)-methyltransfe | Q7Z4G4-3;Q7Z4G4-2;Q7Z4G4 | TRMT11 | Other | 0.536576415 | 2.184377222 | 25.85483662 | 25.61697616 | 25.8080364 | 26.26218075 | 26.09615374 | 25.81567802 | 26.3787014 | 26.62477672 | 26.33318197 |
| Multifunctional methyltransferase sub | Q9UI30;Q9UI30-2 | TRMT112 | Extracellular or secreted | 0.856187858 | 1.243276218 | 26.53863251 | 26.30935173 | 26.46943924 | 26.67626788 | 26.90529463 | 26.93865192 | 27.74675971 | 27.73995362 | 27.0006692 |
| TRMT1-like protein | Q7Z2T5;Q7Z2T5-2 | TRMT1L | Other | 0.214986036 | 0.102913942 | 21.48248897 | 22.5481932 | 20.2443499 | 21.34383352 | 22.43752213 | 20.05284927 | 21.77603457 | 20.40872778 | 22.51481425 |
| tRNA (adenine(58)-N(1))-methyltransf | Q9UJA5;Q9UJA5-4;Q9UJA5-2 | TRMT6 | Other | 0.462776586 | 0.97146105 | 28.86345842 | 29.15221292 | 29.06504476 | 29.07722055 | 29.14116815 | 28.65270194 | 29.70279391 | 29.55851869 | 29.10292053 |
| tRNA (adenine(58)-N(1))-methyltransf | Q96FX7 | TRMT61A | Other | 0.520490135 | 1.11973477 | 27.8003294 | 28.01515427 | 27.97700062 | 27.78450376 | 27.95776 | 27.21965023 | 28.55822672 | 28.41804162 | 27.96240121 |
| CCA tRNA nucleotidyltransferase 1, m | Q96Q11-2;Q96Q11 | TRNT1 | Other | 0.417220966 | 1.337643351 | 27.20927442 | 27.50597288 | 27.37517522 | 27.60272038 | 27.5867057 | 27.04666425 | 27.96554211 | 27.88016942 | 27.56920779 |
| 60 kDa SS-A/Ro ribonucleoprotein | P10155;P10155-3;P10155-5;F | TROVE2 | Other | 0.107427864 | 0.291705242 | 29.46610491 | 29.52469821 | 29.43620882 | 29.05982072 | 29.1486296 | 28.62432354 | 29.44029432 | 29.3000532 | 29.21182897 |
| Transient receptor potential cation ch | Q9Y5S1 | TRPV2 | Extracellular or secreted | -4.097764034 | 1.651159136 | 27.22315371 | 26.80931282 | 27.05755022 | 27.14860536 | 27.33355734 | 26.79476368 | 23.4458171 | 23.76314184 | 21.68122053 |
| TSC22 domain family protein 2;TSC22 | O75157-2;O75157;Q99576-4 | TSC22D2;TSC22D | Other | 0.582828346 | 1.486237639 | 21.55290546 | 22.29810061 | 22.11920965 | 21.96303435 | 21.78076722 | 21.44283741 | 22.40155871 | 22.71973223 | 22.20562145 |
| tRNA-splicing endonuclease subunit S | Q8WW01;Q8WW01-2 | TSEN15 | Other | 0.078488846 | 0.201515096 | 24.82832479 | 24.40593971 | 24.41015215 | 24.49996556 | 24.51316085 | 24.55502578 | 24.82186933 | 24.64982058 | 24.37006104 |
| tRNA-splicing endonuclease subunit S | Q9BSV6 | TSEN34 | Other | -0.08399781 | 0.21613261 | 21.38571606 | 21.13019316 | 21.20036224 | 22.14883965 | 21.37386182 | 21.43822128 | 21.44288479 | 21.2927193 | 21.35099959 |
| Elongation factor Ts, mitochondrial | P43897;P43897-2 | TSFM | Other | -0.078617753 | 0.065517325 | 24.66990407 | 24.76946137 | 25.22066527 | 26.21515703 | 25.88278333 | 27.37790752 | 25.48637227 | 25.63328395 | 25.71242981 |
| Tumor susceptibility gene 101 protein | Q99816;Q99816-2 | TSG101 | Extracellular or secreted | 0.359367124 | 0.726932803 | 26.30244774 | 26.27132951 | 26.22624445 | 26.62366094 | 26.03768653 | 27.19713674 | 26.59163579 | 26.67694037 | 27.13877816 |
| Tsukushin | Q8WUA8 | TSKU | Extracellular or secreted | 0.901634617 | 0.99084542 | 22.02738765 | 22.20558656 | 23.39884805 | 23.85747307 | 24.00881157 | 24.25515481 | 23.73520893 | 24.15081455 | 24.69551123 |
| Translin | Q15631 | TSN | Other | -0.082464639 | 0.102185255 | 30.47111515 | 30.45199082 | 30.4504184 | 29.8678826 | 30.10011534 | 29.30628205 | 30.23260742 | 30.26762875 | 29.57627209 |
| Translin-associated protein X | Q99598 | TSNAX | Other | 0.124183704 | 0.184133351 | 29.87781693 | 29.96719248 | 29.94600756 | 29.47123146 | 29.46855395 | 28.74576648 | 29.85976121 | 29.91013604 | 29.34093829 |
| Tetraspanin-1 | O60635 | TSPAN1 | Extracellular or secreted | 1.610374741 | 1.165002073 | 24.83557089 | 25.26402589 | 25.15611055 | 24.24807639 | 24.73034492 | 25.23191198 | 26.06278762 | 26.00403886 | 27.49731805 |
| Tetraspanin-13 | O95857 | TSPAN13 | Extracellular or secreted | 1.291910175 | 1.169570543 | 21.09181814 | 21.742611 | 21.18022459 | 21.6017053 | 22.99545868 | 24.56621992 | 23.11050625 | 23.68163906 | 23.67260403 |
| Tetraspanin-14 | Q8NG11;Q8NG11-2;Q8NG11 | TSPAN14 | Extracellular or secreted | 0.651280744 | 0.456522286 | 23.24071565 | 23.49448257 | 24.48333065 | 24.58755001 | 22.31273647 | 25.38702731 | 23.84057357 | 24.46992049 | 25.39626951 |
| Tetraspanin-15 | O95858 | TSPAN15 | Extracellular or secreted | 1.108635631 | 0.753694988 | 23.94975762 | 20.59407786 | 21.59516007 | 24.19843005 | 24.10605489 | 24.55789088 | 23.74960104 | 24.36044794 | 24.7165436 |
| Tetraspanin-6 | O43657 | TSPAN6 | Extracellular or secreted | 1.00928341 | 0.492177405 | 23.26799479 | 24.41176905 | 25.13891484 | 25.53255188 | 25.33145741 | 27.86767644 | 25.53139217 | 25.63956262 | 27.63207765 |
| Translocator protein | P30536 | TSPO | Extracellular or secreted | -0.36931666 | 0.188876079 | 21.29365866 | 22.09535242 | 21.80710501 | 22.33723424 | 21.90697865 | 24.90903168 | 20.98788641 | 22.18824715 | 22.89059679 |
| Pre-rRNA-processing protein TSR1 hor | Q2NL82 | TSR1 | Other | 0.886454404 | 2.114009915 | 25.75207944 | 25.45597708 | 25.51376325 | 25.22217728 | 24.96622449 | 25.28217189 | 26.33203819 | 26.46254446 | 25.96097728 |
| Pre-rRNA-processing protein TSR2 hor | Q969E8 | TSR2 | Other | 0.654526606 | 1.208301671 | 24.68072737 | 24.87323832 | 24.69959406 | 25.2168968 | 25.37348357 | 24.08245314 | 25.74411814 | 25.60851293 | 25.07414538 |
| Protein TSSC1 | Q53HC9 | TSSC1 | Other | 0.875081551 | 1.305134276 | 24.48769139 | 24.60529458 | 24.36038095 | 23.86787671 | 23.99049111 | 23.65698255 | 25.2719172 | 25.31091228 | 24.52677383 |
| Protein TSSC4 | Q9Y5U2-2;Q9Y5U2 | TSSC4 | Other | 0.623930259 | 0.66681051 | 23.82997043 | 23.58932289 | 23.54686715 | 23.75378884 | 24.1800275 | 22.30774146 | 24.62042031 | 24.36686378 | 23.48836582 |
| Thiosulfate sulfurtransferase | Q16762 | TST | Extracellular or secreted | -0.278218836 | 0.273726133 | 22.47734839 | 21.52032457 | 20.36070097 | 22.68775603 | 21.95940957 | 22.16885694 | 21.65250365 | 21.97526303 | 21.12477504 |
| GDP-L-fucose synthase | Q13630 | TSTA3 | Extracellular or secreted | -0.281696905 | 2.610310813 | 28.89821821 | 29.07237278 | 29.03519734 | 28.79906354 | 29.03485638 | 28.77796786 | 28.65030692 | 28.63183488 | 28.68160555 |
| Thiosulfate sulfurtransferase/rhodane | Q8NFU3;Q8NFU3-3;Q8NFU3- | TSTD1 | Other | 1.044935115 | 2.104942806 | 26.2293836 | 26.14720834 | 26.13803598 | 26.16351931 | 26.36180365 | 26.5234873 | 27.33304545 | 27.53400763 | 27.04947136 |
| Tetratricopeptide repeat protein 1 | Q99614 | TTC1 | Other | 0.384068238 | 0.604798386 | 27.14404064 | 27.17436159 | 27.20050818 | 27.37252077 | 27.27967703 | 26.82650756 | 27.81292729 | 27.77096134 | 27.06712396 |
| Tetratricopeptide repeat protein 19, r | Q6DKK2 | TTC19 | Extracellular or secreted | 0.183834553 | 0.201200199 | 25.1091675 | 25.44047255 | 25.192347 | 24.47475509 | 24.70345415 | 23.88231691 | 25.05366144 | 25.41638341 | 24.48271541 |
| Tetratricopeptide repeat protein 27 | Q6P3X3 | TTC27 | Other | 2.385920653 | 1.019813026 | 24.81587343 | 25.25143936 | 24.71156364 | 22.16551797 | 22.16158708 | 22.53400169 | 27.15131792 | 26.61680794 | 24.20962768 |
| Tetratricopeptide repeat protein 37 | Q6PGP7 | TTC37 | Other | -0.055698098 | 0.069469528 | 25.4176715 | 25.77502487 | 25.80646388 | 25.65523634 | 25.35056803 | 25.45456588 | 25.94277346 | 25.56354589 | 25.05635161 |
| Tetratricopeptide repeat protein 38 | Q5R3I4 | TTC38 | Extracellular or secreted | 2.630154142 | 1.511676238 | 21.58210338 | 21.91261747 | 23.31575656 | 23.32512206 | 21.99141726 | 20.71813775 | 24.60732806 | 25.97526303 | 23.73044858 |
| Tetratricopeptide repeat protein 39C | Q8N584-2;Q8N584 | TTC39C | Other | 0.141259918 | 0.188511282 | 23.76192532 | 23.6212032 | 23.78446631 | 23.57427014 | 23.61459123 | 23.77434631 | 24.04959599 | 24.13617884 | 23.30340619 |
| Tetratricopeptide repeat protein 4 | O95801 | TTC4 | Other | 0.732701159 | 0.960434952 | 24.38040346 | 24.37099224 | 24.48345367 | 24.75154324 | 24.67691348 | 23.69699723 | 25.38452635 | 25.45299627 | 24.54073253 |
| Tetratricopeptide repeat protein 5 | Q8N0Z6 | TTC5 | Other | 1.250016948 | 0.814693787 | 24.43377185 | 23.85993889 | 24.05986197 | 23.90884849 | 21.49899635 | 24.85704587 | 25.1958798 | 25.89691421 | 23.96648855 |
| Tetratricopeptide repeat protein 9C | Q8N5M4;Q8N5M4-2 | TTC9C | Other | 0.034053104 | 0.053401365 | 23.14050715 | 23.2727004 | 22.46629682 | 23.49947903 | 23.30758106 | 22.30426998 | 23.13447207 | 23.13619449 | 22.82690997 |
| TELO2-interacting protein 1 homolog | O43156 | TTI1 | Other | 0.126432242 | 0.059332231 | 20.96803674 | 20.96845868 | 20.78779737 | 22.61781704 | 22.2906294 | 21.1007234 | 22.2599455 | 22.20021255 | 20.28586998 |
| Tubulin--tyrosine ligase-like protein 1 | Q14166 | TTLL12 | Other | 0.327138763 | 0.620885368 | 29.23683234 | 29.13244266 | 29.00262864 | 29.18113242 | 29.10517076 | 29.03244112 | 29.59473526 | 29.68430044 | 29.04770456 |
| Titin | Q8WZ42-5;Q8WZ42-11;Q8W | TTN | Extracellular or secreted | -0.865244074 | 0.437681235 | 26.70780418 | 25.87088919 | 26.96840155 | 28.25408264 | 28.0171207 | 26.27104449 | 24.71889655 | 26.66898494 | 27.06105767 |
| Protein tweety homolog 1 | Q9H313-3;Q9H313;Q9H313-2 | TTYH1 | Extracellular or secreted | 1.074591491 | 1.741196751 | 27.094262 | 26.95315934 | 27.01424987 | 27.33039775 | 27.57809128 | 28.10314571 | 28.07765351 | 28.31704275 | 28.86573119 |
| Protein tweety homolog 2 | Q9BSA4 | TTYH2 | Extracellular or secreted | 1.403234427 | 0.996212335 | 22.72388255 | 22.64717645 | 22.07929857 | 19.79445775 | 23.18745013 | 24.91223371 | 23.55285856 | 23.85880134 | 24.47029296 |
| Protein tweety homolog 3 | Q9C0H2-3;Q9C0H2-2;Q9C0H | TTYH3 | Extracellular or secreted | 0.625097174 | 0.6806144 | 21.99455371 | 21.90954565 | 21.89230267 | 21.85831839 | 21.82559081 | 21.45287062 | 21.98853353 | 22.22696835 | 23.12638057 |
| Tubulin alpha-1A chain;Tubulin alpha- | Q71U36;Q71U36-2;P0DPH8;F | TUBA1A;TUBA3E | Extracellular or secreted | 0.307773991 | 0.656324927 | 28.32281216 | 28.26445549 | 28.21502739 | 28.96600991 | 29.08311793 | 28.35481057 | 29.16036321 | 28.68782465 | 28.67825083 |
| Tubulin alpha-1B chain | P68363;P68363-2 | TUBA1B | Extracellular or secreted | 0.473036257 | 1.277843104 | 33.64667258 | 33.7380175 | 33.66712459 | 33.60476721 | 33.73368099 | 33.57801478 | 34.27162728 | 34.23670406 | 33.89491625 |
| Tubulin alpha-1C chain | Q9BQE3 | TUBA1C | Other | 0.317348073 | 0.290933246 | 27.95532292 | 28.13877816 | 27.99959441 | 28.37683178 | 28.44407202 | 26.86088616 | 28.63778828 | 28.66132525 | 27.54067342 |
| Tubulin alpha-4A chain | P68366-2;P68366 | TUBA4A | Extracellular or secreted | -0.20562012 | 0.404673234 | 27.44671645 | 27.51668134 | 27.5501009 | 26.68952521 | 26.48048293 | 27.72864642 | 27.15981007 | 26.89538945 | 27.03401675 |
| Tubulin alpha chain-like 3 | A6NHL2-2;A6NHL2 | TUBAL3 | Other | 1.60904383 | 1.987116811 | 23.90113353 | 24.21111415 | 24.45744952 | 23.64030761 | 22.24999831 | 25.20761851 | 25.63153656 | 25.52716174 | 25.502244 |
| Tubulin beta chain | P07437 | TUBB | Extracellular or secreted | 0.930576974 | 1.084895524 | 31.65493088 | 31.74169833 | 31.67265629 | 31.19601117 | 31.32157626 | 31.30850789 | 32.76505167 | 32.67301449 | 31.80135517 |
| Tubulin beta-2A chain | Q13885 | TUBB2A | Extracellular or secreted | 0.452219616 | 0.781888834 | 27.99647297 | 27.96394496 | 27.87378981 | 28.08304183 | 28.09803297 | 28.44193715 | 28.8321939 | 28.65263357 | 28.10044122 |
| Tubulin beta-3 chain | Q13509;Q13509-2 | TUBB3 | Extracellular or secreted | 0.741221562 | 2.052155385 | 27.8722639 | 28.19952565 | 28.15238218 | 28.0021189 | 27.5590295 | 28.90615524 | 28.86894239 | 28.99212941 | 28.70833057 |
| Tubulin beta-4A chain | P04350 | TUBB4A | Extracellular or secreted | -0.942308739 | 2.350764665 | 25.12313122 | 24.91665908 | 25.02778825 | 26.05880976 | 25.82529481 | 25.75755767 | 24.59531167 | 24.47951188 | 24.45287062 |
| Tubulin beta-4B chain | P68371 | TUBB4B | Extracellular or secreted | 0.550940038 | 0.926326935 | 33.93192075 | 34.03575778 | 34.04839985 | 33.74424728 | 33.67944684 | 33.83648538 | 34.74279272 | 34.53918786 | 34.00896847 |
| Tubulin beta-6 chain | Q9BUF5 | TUBB6 | Extracellular or secreted | 0.668494998 | 1.930988825 | 28.02310974 | 28.18697488 | 28.25966002 | 28.33355734 | 28.41013597 | 29.23628494 | 29.06373409 | 29.1265716 | 29.04004075 |
| Tubulin beta-8 chain | Q3ZCM7 | TUBB8 | Extracellular or secreted | 0.788207963 | 2.938342871 | 30.35541982 | 30.43251001 | 30.29865565 | 30.25900894 | 30.25788569 | 31.06804709 | 31.26561834 | 31.27702926 | 31.15773989 |
| Tubulin gamma-1 chain;Tubulin gamm | P23258;Q9NRH3 | TUBG1;TUBG2 | Other | -0.379885811 | 0.945379627 | 27.47274264 | 27.03380676 | 27.16930435 | 26.51395898 | 26.28102231 | 27.11336685 | 26.69232178 | 26.61428231 | 26.34583943 |
| Gamma-tubulin complex component | Q9BSJ2;Q9BSJ2-4;Q9BSJ2-3 | TUBGCP2 | Other | -0.483120599 | 1.151204975 | 25.81303742 | 25.80845382 | 25.88378563 | 24.64311854 | 25.09039029 | 25.41573893 | 25.01499471 | 25.09430228 | 24.76860354 |
| Gamma-tubulin complex component | Q96CW5-2;Q96CW5;Q96CW | TUBGCP3 | Other | -0.924973504 | 0.408957187 | 22.90530841 | 24.57536705 | 22.03049337 | 23.36064886 | 23.57276776 | 22.77806189 | 20.78149453 | 22.37724231 | 23.67766631 |
| Gamma-tubulin complex component | Q96RT8-2;Q96RT8 | TUBGCP5 | Other | 1.226180865 | 0.502438367 | 21.53515293 | 21.89632888 | 19.89987889 | 21.72589198 | 21.58020867 | 19.95976884 | 22.87451465 | 23.57450114 | 20.5281419 |
| Elongation factor Tu, mitochondrial | P49411 | TUFM | Extracellular or secreted | -0.212851462 | 0.214182352 | 28.50967704 | 28.64701583 | 29.02059525 | 29.38152193 | 29.42440317 | 31.11035789 | 29.16125262 | 28.96303545 | 29.2839431 |
| Speckle targeted PIP5K1A-regulated p | Q9H6E5 | TUT1 | Extracellular or secreted | 0.950463367 | 0.80919347 | 20.31288201 | 21.54674942 | 21.31659442 | 22.11987454 | 22.2128217 | 20.36937576 | 22.71655407 | 22.69230049 | 21.38168447 |
| Twinfilin-1 | Q12792;Q12792-3;Q12792-4 | TWF1 | Extracellular or secreted | -0.134470783 | 0.190206438 | 29.70786999 | 29.78261483 | 29.70056336 | 28.92241001 | 28.85001444 | 29.09458921 | 29.36580063 | 29.46703837 | 28.79277957 |
| Twinfilin-2 | Q6IBS0 | TWF2 | Extracellular or secreted | 0.023176196 | 0.0644526 | 29.53267077 | 29.59553576 | 29.50485598 | 29.21268721 | 29.05982072 | 28.97768306 | 29.31546722 | 29.46663005 | 29.22905807 |
| DNA-directed RNA polymerase I subu | Q3B726 | TWISTNB | Other | 0.077482074 | 0.099878853 | 23.95712897 | 23.77986601 | 24.06258178 | 24.49167208 | 23.07564758 | 24.62377256 | 24.00633087 | 24.40704262 | 23.81440723 |
| Twisted gastrulation protein homolog | Q9GZX9 | TWSG1 | Extracellular or secreted | 0.346858266 | 0.483888237 | 25.26820902 | 24.83754517 | 24.89749135 | 25.7796907 | 26.12998414 | 26.60002813 | 25.67446406 | 25.89834509 | 26.2242399 |
| Alpha-taxilin | P40222 | TXLNA | Extracellular or secreted | 0.626314445 | 1.330059873 | 27.09978938 | 27.03977936 | 26.93460855 | 27.79563087 | 28.04874412 | 26.46587923 | 27.89035326 | 27.90959255 | 27.77121329 |
| Gamma-taxilin | Q9NUQ3;Q9NUQ3-2 | TXLNG | Other | -0.08910475 | 0.169318014 | 28.84870192 | 28.83674495 | 28.79983554 | 29.52466085 | 29.50322642 | 28.63727007 | 29.08534858 | 29.0462219 | 28.67633515 |
| Thioredoxin | P10599;P10599-2 | TXN | Extracellular or secreted | 0.310520477 | 0.929377488 | 32.24024889 | 32.33697599 | 32.36674396 | 32.38422166 | 32.52470288 | 32.03025867 | 32.7959095 | 32.70479021 | 32.37243774 |
| Thioredoxin, mitochondrial | Q99757 | TXN2 | Other | -0.236752408 | 0.091890804 | 27.11406224 | 26.56468005 | 26.80685717 | 27.38812834 | 27.84935833 | 24.87173532 | 27.2649923 | 27.38919569 | 24.9329655 |
| Thioredoxin domain-containing prote | O95881 | TXNDC12 | Extracellular or secreted | -0.421831915 | 0.774450032 | 29.61629966 | 29.61196326 | 29.71148487 | 29.86489039 | 29.93170945 | 28.85575755 | 29.3746157 | 29.38463338 | 28.77130776 |
| Thioredoxin domain-containing prote | Q9BRA2 | TXNDC17 | Extracellular or secreted | -0.029209106 | 0.024274429 | 30.22366009 | 30.01637696 | 30.11557605 | 30.22008863 | 30.4648983 | 28.85233824 | 30.35468448 | 30.20209751 | 29.30205983 |
| Thioredoxin domain-containing prote | Q8NBS9;Q8NBS9-2 | TXNDC5 | Extracellular or secreted | -0.288003962 | 0.832714855 | 31.74169833 | 31.69540337 | 31.65961662 | 31.53949988 | 31.59027098 | 31.35431667 | 31.46153589 | 31.41960952 | 31.04524563 |
| Thioredoxin domain-containing prote | O14530-2;O14530 | TXNDC9 | Other | 0.919807392 | 3.546348006 | 24.34577177 | 24.38508615 | 23.87004257 | 24.42675205 | 24.22934692 | 24.62884198 | 25.22258267 | 25.09671676 | 25.38304346 |
| Thioredoxin-like protein 1 | O43396 | TXNL1 | Extracellular or secreted | 0.487637848 | 1.533511119 | 29.14367581 | 29.05418953 | 29.03393801 | 29.09998998 | 29.03230974 | 29.29570322 | 29.72227546 | 29.68093523 | 29.389606 |
| Thioredoxin-like protein 4A | P83876 | TXNL4A | Other | 0.35057789 | 0.354260051 | 25.62257223 | 25.3547097 | 25.39633484 | 24.84474522 | 26.03414272 | 24.24517723 | 25.76688634 | 26.06759585 | 24.96609244 |
| Thioredoxin reductase 1, cytoplasmic | Q16881-4;Q16881-3;Q16881 | TXNRD1 | Extracellular or secreted | -0.307005636 | 0.49291697 | 32.55944439 | 32.70221182 | 32.60372869 | 32.8634695 | 32.96456131 | 31.62823188 | 32.51608486 | 32.37534096 | 31.84838107 |
| Thioredoxin reductase 2, mitochondri | Q9NNW7-2;Q9NNW7;Q9NN\ | TXNRD2 | Extracellular or secreted | -0.688054743 | 2.637386433 | 27.03506621 | 26.97213228 | 26.84505641 | 27.4221387 | 27.57701085 | 26.87537271 | 26.38773404 | 26.43955344 | 26.47193688 |
| Thioredoxin reductase 3 | Q86VQ6 | TXNRD3 | Other | 0.076486453 | 0.120979331 | 26.52173641 | 26.61652753 | 26.61554568 | 26.82335227 | 26.61428231 | 25.44297335 | 26.62603093 | 26.65034116 | 26.27029604 |
| Thymidylate synthase | P04818;P04818-2 | TYMS | Other | 2.222003392 | 2.643639902 | 24.26219867 | 24.27433663 | 24.5473968 | 24.71240357 | 25.36669707 | 26.01930719 | 26.60935867 | 27.63740828 | 27.0104132 |
| Tyrosine-protein kinase receptor TYR | Q06418 | TYRO3 | Extracellular or secreted | -0.027560042 | 0.014212944 | 24.08123441 | 23.24959612 | 23.50118118 | 23.79995902 | 24.00855514 | 24.51032618 | 23.44101106 | 23.06332266 | 24.98841218 |
| Splicing factor U2AF 35 kDa subunit | P0DN76;Q01081;Q01081-2;Q | U2AF1 | Other | -0.359523486 | 0.39411625 | 26.48753806 | 26.76660837 | 26.82723473 | 27.80636554 | 27.86832427 | 28.11500546 | 26.68123691 | 26.66220853 | 27.51352232 |
| Splicing factor U2AF 65 kDa subunit | P26368-2;P26368 | U2AF2 | Other | -0.96375199 | 1.317318713 | 30.32324179 | 30.41316621 | 30.28140266 | 30.33319477 | 30.39407084 | 30.31267886 | 29.02725568 | 29.32021017 | 29.79015574 |
| U2 snRNP-associated SURP motif-con | O15042-2;O15042;O15042-3 | U2SURP | Other | -0.722517767 | 1.478356073 | 27.87987739 | 28.06249944 | 27.74348909 | 26.61175225 | 27.26481339 | 28.28020821 | 26.86254239 | 26.91843454 | 26.97278964 |
| Uveal autoantigen with coiled-coil do | Q9BZF9-2;Q9BZF9 | UACA | Extracellular or secreted | -2.697092726 | 1.603925955 | 27.56376407 | 27.45442469 | 27.6073845 | 25.71311189 | 25.94156509 | 26.80205659 | 24.74734891 | 22.93740982 | 24.7651165 |
| UDP-N-acetylhexosamine pyrophosph | Q16222-3;Q16222;Q16222-2 | UAP1 | Extracellular or secreted | 0.437446308 | 1.893986314 | 27.9264311 | 28.05553474 | 28.04676832 | 27.7509813 | 27.86177367 | 27.61498432 | 28.33819877 | 28.44994634 | 28.15243054 |
| Ubiquitin-like modifier-activating enz | P22314-2;P22314 | UBA1 | Extracellular or secreted | -0.020845246 | 0.033278312 | 32.97456167 | 33.01411683 | 32.93828034 | 32.69565227 | 32.75244557 | 32.31753879 | 32.94875976 | 32.93468236 | 32.40031987 |
| SUMO-activating enzyme subunit 2 | Q9UBT2;Q9UBT2-2 | UBA2 | Other | 0.229891116 | 0.578022688 | 31.305793 | 31.24547194 | 31.19689059 | 30.74575045 | 30.91783728 | 30.94816796 | 31.42612419 | 31.45002503 | 30.99347974 |
| NEDD8-activating enzyme E1 catalytic | Q8TBC4;Q8TBC4-2 | UBA3 | Other | 0.060524808 | 0.112501527 | 29.31505687 | 29.43125503 | 29.40108829 | 29.10279542 | 29.12346675 | 28.94514249 | 29.43473839 | 29.47109576 | 28.9351427 |
| Ubiquitin-like modifier-activating enz | Q9GZZ9;Q9GZZ9-2 | UBA5 | Other | 1.318937528 | 0.841996054 | 25.8748101 | 26.22910851 | 26.24178143 | 25.60070875 | 26.16031031 | 23.55027708 | 27.58727816 | 27.46013989 | 25.73789263 |
| Ubiquitin-like modifier-activating enz | A0AVT1;A0AVT1-2 | UBA6 | Other | 0.273805385 | 1.202500598 | 29.57547886 | 29.55636403 | 29.52878313 | 29.53045852 | 29.49411629 | 29.50531042 | 29.80742541 | 29.93339895 | 29.67584742 |
| Ubiquitin-associated domain-containi | Q9BSL1 | UBAC1 | Extracellular or secreted | 0.688785413 | 1.270708543 | 21.04500237 | 20.69139835 | 21.81660681 | 21.67663376 | 20.68146366 | 21.95858103 | 21.64412033 | 22.20332291 | 22.15375599 |
| Ubiquitin-associated protein 2 | Q5T6F2 | UBAP2 | Extracellular or secreted | 0.388555801 | 0.617005975 | 25.08403596 | 25.04055294 | 25.28996483 | 26.19882344 | 25.47580664 | 24.35756489 | 25.87375462 | 25.72961908 | 25.28566805 |
| Ubiquitin-associated protein 2-like | Q14157-4;Q14157-1;Q14157 | UBAP2L | Other | 0.65330886 | 2.023644628 | 27.95831331 | 27.99167001 | 28.02865258 | 28.50684308 | 28.53241068 | 27.90890574 | 28.96380719 | 28.88980226 | 28.56971483 |
| Ubiquitin-associated and SH3 domain | Q8TF42 | UBASH3B | Other | 0.730086386 | 2.405125775 | 25.14682004 | 24.55759879 | 24.63039823 | 24.82313357 | 24.56714885 | 25.37178994 | 25.61268045 | 25.38988495 | 25.73613846 |

| | | | | | | | | | | | | | | |
|---|---|---|---|---|---|---|---|---|---|---|---|---|---|---|
| Ubiquitin-conjugating enzyme E2 A | P49459;P49459-3;P49459-2 | UBE2A | Other | -0.258138912 | 0.315896265 | 27.07785721 | 26.87103025 | 26.95160399 | 27.41151047 | 27.37658342 | 27.22674056 | 27.2043383 | 27.18196174 | 26.29694616 |
| Ubiquitin-conjugating enzyme E2 C | O00762;O00762-2;O00762-3 | UBE2C | Other | 1.134422789 | 2.184192292 | 25.27742618 | 25.17550417 | 25.47855561 | 26.18638934 | 25.84541539 | 26.20711578 | 27.05662035 | 26.98576586 | 26.44608539 |
| Ubiquitin-conjugating enzyme E2 D3;U | P61077;P61077-2;P61077-3;P | UBE2D3;UBE2D2 | Extracellular or secreted | 0.848267071 | 1.303330958 | 26.03915183 | 27.68545385 | 27.69637383 | 27.83098592 | 27.7064215 | 28.17131029 | 28.47150281 | 28.62933544 | 28.00881157 |
| Ubiquitin-conjugating enzyme E2 E1 | P51965-2;P51965 | UBE2E1 | Other | -0.135809677 | 0.222605692 | 23.94619166 | 23.98927876 | 23.84928673 | 23.58152885 | 23.98250553 | 24.21386009 | 24.0146117 | 23.99222127 | 23.36706382 |
| Ubiquitin-conjugating enzyme E2 E2;U | Q96LR5;P51965-3 | UBE2E2;UBE2E1 | Other | 0.013318852 | 0.020150071 | 26.60851293 | 26.84254102 | 26.92580885 | 26.66492291 | 26.51121638 | 27.0333867 | 26.92082113 | 27.025171 | 26.38715882 |
| Ubiquitin-conjugating enzyme E2 G1;U | P62253 | UBE2G1 | Extracellular or secreted | 0.076075846 | 0.05809084 | 27.42189802 | 27.58978005 | 27.44379546 | 27.29535291 | 27.63055098 | 26.56656834 | 27.71934056 | 27.89336551 | 26.58949434 |
| Ubiquitin-conjugating enzyme E2 G2 | P60604-2;P60604 | UBE2G2 | Other | -0.252744696 | 0.28217041 | 26.13151569 | 26.06566627 | 26.17495204 | 25.92100281 | 26.2163973 | 26.12259816 | 26.11901962 | 26.23832295 | 25.20048947 |
| Ubiquitin-conjugating enzyme E2 H | P62256;P62256-2 | UBE2H | Other | 0.501486052 | 0.714922781 | 26.4517865 | 26.05063419 | 25.72533443 | 26.70938277 | 26.74111164 | 26.37278643 | 26.96741239 | 27.23187537 | 26.33068838 |
| SUMO-conjugating enzyme UBC9 | P63279 | UBE2I | Other | 0.017139348 | 0.084649489 | 30.08309256 | 30.22124168 | 30.06548138 | 29.98739327 | 30.09219637 | 29.84864223 | 30.15581148 | 30.07099202 | 29.9736383 |
| Ubiquitin-conjugating enzyme E2 J1 | Q9Y385 | UBE2J1 | Other | -0.603103706 | 0.919523853 | 21.9050329 | 22.52197897 | 22.24321699 | 21.69869894 | 21.91818702 | 23.34975852 | 21.44101106 | 21.44561506 | 22.12249943 |
| Ubiquitin-conjugating enzyme E2 K | P61086;P61086-2;P61086-3 | UBE2K | Extracellular or secreted | 0.116971526 | 0.263593266 | 29.90870536 | 29.9808726 | 29.83340088 | 29.89836239 | 29.83624835 | 29.36936225 | 29.96897825 | 30.15436347 | 29.64104877 |
| Ubiquitin-conjugating enzyme E2 L3 | P68036;P68036-3;P68036-2;A | UBE2L3 | Extracellular or secreted | 0.214174915 | 0.303371064 | 30.51953356 | 30.68269415 | 30.58450315 | 30.37409744 | 30.20139655 | 29.98766433 | 30.62653579 | 31.03367551 | 30.15725803 |
| Ubiquitin/ISG15-conjugating enzyme | O14933-2;O14933 | UBE2L6 | Other | -2.576939261 | 1.19372201 | 26.0092602 | 26.04699723 | 25.55505505 | 24.9497131 | 24.47951188 | 25.31893111 | 23.56883102 | 23.57888308 | 21.3012024 |
| NEDD8-conjugating enzyme Ubc12 | P61081 | UBE2M | Extracellular or secreted | -0.296921068 | 0.764308361 | 28.65693483 | 28.69282059 | 28.69032478 | 28.60830142 | 28.69710332 | 28.63325623 | 28.44015568 | 28.56837442 | 28.09007728 |
| Ubiquitin-conjugating enzyme E2 N | P61088 | UBE2N | Extracellular or secreted | 0.056411935 | 0.123463951 | 30.68478532 | 30.75358165 | 30.79482564 | 31.04882205 | 30.94363436 | 30.36557122 | 30.83379292 | 31.05193606 | 30.57911694 |
| E2/E3 hybrid ubiquitin-protein ligase U | Q9C0C9 | UBE2O | Other | 0.078073025 | 0.139777969 | 28.07653265 | 28.04176475 | 27.94536578 | 28.23840496 | 28.26270945 | 27.94586806 | 28.33717809 | 28.36623849 | 27.78612532 |
| Ubiquitin-conjugating enzyme E2 Q1 | Q7Z7E8;Q7Z7E8-2 | UBE2Q1 | Other | 0.483128407 | 1.242634494 | 26.14254181 | 25.78399196 | 26.10073195 | 25.9916592 | 25.77296309 | 25.96066803 | 26.59348915 | 26.57386581 | 26.15830828 |
| Ubiquitin-conjugating enzyme E2 R2 | Q712K3 | UBE2R2 | Other | -0.227686186 | 0.33080234 | 25.74811708 | 25.45409518 | 25.68005664 | 26.18082427 | 25.83610084 | 24.55783247 | 25.46354258 | 25.60385246 | 24.97805963 |
| Ubiquitin-conjugating enzyme E2 S | Q16763 | UBE2S | Other | 0.261180528 | 0.310659222 | 25.62747889 | 24.91848004 | 25.0238183 | 26.30765054 | 26.16365367 | 26.57960252 | 25.99942237 | 26.42334151 | 25.67111968 |
| Ubiquitin-conjugating enzyme E2 T | Q9NPD8 | UBE2T | Other | 1.167268002 | 2.384248114 | 24.34285922 | 24.79664606 | 24.29375789 | 23.43923637 | 23.93418109 | 24.28418587 | 25.07981125 | 25.68656293 | 25.28086307 |
| Ubiquitin-conjugating enzyme E2 vari | Q13404;Q13404-7;Q13404-2 | UBE2V1 | Extracellular or secreted | -0.039417191 | 0.072812995 | 28.49342928 | 28.53385915 | 28.02865258 | 28.77467307 | 28.86056061 | 28.3375184 | 28.47363311 | 28.71607255 | 28.20638931 |
| Ubiquitin-conjugating enzyme E2 vari | Q15819 | UBE2V2 | Extracellular or secreted | 0.167040297 | 0.318702488 | 30.306608 | 30.46801007 | 30.2999659 | 30.18777726 | 30.07405859 | 30.0937081 | 30.4165928 | 30.73141679 | 30.06817525 |
| Ubiquitin-conjugating enzyme E2 Z | Q9H832;Q9H832-2 | UBE2Z | Other | 0.26362809 | 0.680910003 | 27.57751515 | 27.42878173 | 27.39982604 | 26.90046571 | 26.57571327 | 27.06013006 | 27.55002749 | 27.50960154 | 27.20247122 |
| Ubiquitin-protein ligase E3A | Q05086-3;Q05086;Q05086-2 | UBE3A | Other | -0.288153156 | 1.718505021 | 29.1809665 | 29.1597379 | 29.02390288 | 29.17769159 | 29.1305783 | 28.99142674 | 28.8424511 | 28.91359242 | 28.71164896 |
| Ubiquitin conjugation factor E4 A | Q14139;Q14139-2 | UBE4A | Other | -0.171417955 | 0.329917988 | 25.17459018 | 25.66269749 | 25.62770152 | 24.77218286 | 25.11247228 | 25.48406861 | 25.31247114 | 25.28711348 | 24.80301799 |
| Ubiquitin conjugation factor E4 B | O95155-2;O95155;O95155-4 | UBE4B | Other | -0.308740166 | 1.030748585 | 23.4379674 | 23.73624171 | 23.19472319 | 23.27767448 | 23.92614829 | 23.32278638 | 23.33960104 | 23.19709548 | 22.98485371 |
| Ubiquitin domain-containing protein U | O14562 | UBFD1 | Other | 0.017019739 | 0.019642166 | 27.53504656 | 27.39009822 | 27.34112922 | 27.33296011 | 27.26794123 | 26.90552418 | 27.35009587 | 27.74957547 | 26.83773764 |
| Ubiquitin-like protein 4A | P11441 | UBL4A | Other | -0.070195442 | 0.058986373 | 25.77072196 | 25.63555517 | 25.50221365 | 25.26072805 | 25.32013699 | 27.36527918 | 25.44351093 | 25.48661778 | 26.28660246 |
| Ubiquitin-like protein 5 | Q9BZL1 | UBL5 | Other | 2.081143146 | 1.047770424 | 25.15491389 | 24.91383206 | 24.98350615 | 24.86561322 | 24.37378224 | 20.67796734 | 26.99248062 | 26.94904513 | 24.78671113 |
| Ubiquitin-like domain-containing CTD | Q8WVY7 | UBLCP1 | Other | 0.379711333 | 0.528858297 | 27.41924785 | 27.5339334 | 27.55749654 | 27.0744925 | 27.05682704 | 26.62254431 | 27.75913358 | 27.94648173 | 27.06578951 |
| Upstream-binding protein 1 | Q9NZI7-4;Q9NZI7 | UBP1 | Other | -0.039055358 | 0.043008336 | 22.17000555 | 20.92449649 | 22.40195852 | 21.93782366 | 22.15524998 | 22.97163462 | 21.74443641 | 22.27699333 | 22.14198861 |
| Ubiquilin-1 | Q9UMX0;Q9UMX0-2;Q9UMX | UBQLN1 | Extracellular or secreted | 0.123468998 | 0.12486425 | 30.19436814 | 30.20092905 | 30.1779529 | 30.35783332 | 30.33639082 | 29.42740361 | 30.62078035 | 30.49762261 | 29.59944296 |
| Ubiquilin-2 | Q9UHD9 | UBQLN2 | Extracellular or secreted | 0.076971708 | 0.354525858 | 27.10609483 | 26.92286365 | 26.98598296 | 27.22949362 | 26.71332169 | 26.96818179 | 26.99959441 | 27.18697488 | 27.00731511 |
| Ubiquilin-4 | Q9NRR5 | UBQLN4 | Other | -0.147246431 | 0.277579199 | 27.68418337 | 27.78450376 | 27.85055103 | 28.03307158 | 28.26758411 | 27.47969689 | 27.83098592 | 27.92823977 | 27.34883039 |
| E3 ubiquitin-protein ligase UBR1 | Q8IWV7;Q8IWV7-2 | UBR1 | Other | -0.211842051 | 0.240635347 | 25.18767345 | 25.39888066 | 25.60935867 | 25.00564579 | 24.29487984 | 24.6462537 | 25.25925593 | 24.8939325 | 24.28263147 |
| E3 ubiquitin-protein ligase UBR2 | Q8IWV8-4;Q8IWV8;Q8IWV8- | UBR2 | Extracellular or secreted | 0.631102727 | 0.858907372 | 23.96754431 | 23.69381107 | 24.39335903 | 21.85042542 | 23.57939203 | 23.77595422 | 24.21141126 | 24.15367758 | 24.15846238 |
| E3 ubiquitin-protein ligase UBR3 | Q6ZT12;Q6ZT12-4 | UBR3 | Extracellular or secreted | 0.626118964 | 1.092598733 | 21.34435009 | 22.0873836 | 21.20018274 | 22.70636617 | 22.36324501 | 22.90551041 | 22.69140617 | 22.7830078 | 22.70746193 |
| E3 ubiquitin-protein ligase UBR4 | Q5T4S7-2;Q5T4S7-3;Q5T4S7; | UBR4 | Other | 0.111081259 | 0.520385425 | 29.96815432 | 30.01929659 | 30.03012709 | 29.94237638 | 29.98834174 | 29.63985928 | 30.15689653 | 30.06817525 | 29.9022497 |
| E3 ubiquitin-protein ligase UBR5 | O95071-2;O95071 | UBR5 | Other | 0.89257648 | 1.486869996 | 25.67039044 | 26.016802 | 25.68981844 | 24.87539615 | 24.49606868 | 26.52290391 | 26.65908066 | 26.22846643 | 26.42587216 |
| Putative E3 ubiquitin-protein ligase U | Q8N806 | UBR7 | Other | 0.260864583 | 1.141101308 | 26.58233313 | 26.37982532 | 26.30778948 | 26.63713184 | 26.67989562 | 26.60907681 | 26.88471738 | 26.87923473 | 26.61666774 |
| Ubiquitin domain-containing protein 2 | Q8WUN7;Q9HAC8 | UBTD2;UBTD1 | Other | 1.421988816 | 2.550603395 | 23.27980099 | 22.83978583 | 23.93669904 | 23.54132353 | 21.97799974 | 23.24398688 | 24.30277891 | 24.54079164 | 24.8321939 |
| Nucleolar transcription factor 1 | P17480-2;P17480 | UBTF | Other | -0.926876137 | 1.07166037 | 26.34903294 | 27.41643173 | 26.91147958 | 26.61400141 | 26.67895597 | 26.96046919 | 26.40086849 | 25.28665533 | 25.99703317 |
| UBX domain-containing protein 1 | Q04323;Q04323-2 | UBXN1 | Other | 0.313596988 | 0.628455879 | 27.90747384 | 27.63975581 | 28.03988392 | 28.2233839 | 28.19975965 | 27.19666786 | 28.16916097 | 28.48439134 | 27.89070115 |
| UBX domain-containing protein 4 | Q92575 | UBXN4 | Other | -1.296442359 | 1.162591723 | 26.72585446 | 26.42476755 | 26.54639618 | 26.18173432 | 26.01860703 | 25.64273305 | 25.25637917 | 25.46572358 | 24.15861647 |
| UBX domain-containing protein 7 | O94888 | UBXN7 | Other | 0.43189818 | 0.421781253 | 27.15701704 | 27.12947327 | 26.84050154 | 27.07907879 | 27.09083445 | 26.63062041 | 27.849239 | 27.75837125 | 26.65184703 |
| Ubiquitin carboxyl-terminal hydrolase | P15374 | UCHL3 | Extracellular or secreted | 0.582861241 | 1.423140581 | 29.57212081 | 29.53943517 | 29.53782571 | 29.72791648 | 29.82020529 | 28.70196818 | 30.20990186 | 30.22008863 | 29.76832905 |
| Ubiquitin carboxyl-terminal hydrolase | Q9Y5K5-2;Q9Y5K5-4;Q9Y5K5 | UCHL5 | Other | -0.057376275 | 0.100014089 | 28.13486763 | 27.97339197 | 28.07127339 | 28.30273534 | 28.39521555 | 27.79457779 | 28.24671813 | 28.22379816 | 27.69338571 |
| Uridine-cytidine kinase 1 | Q9HA47-3;Q9HA47;Q9HA47- | UCK1 | Other | -0.051772011 | 0.023891499 | 23.83263813 | 23.2085267 | 21.91361508 | 20.5796934 | 22.36599971 | 23.04391841 | 23.08940457 | 21.32040474 | 22.90707036 |
| Uridine-cytidine kinase 2 | Q9BZX2 | UCK2 | Other | 0.721794431 | 1.01152817 | 24.42130416 | 24.2641333 | 24.378751 | 25.16211746 | 25.09244843 | 24.88952959 | 25.6951926 | 25.7020805 | 24.87225215 |
| Ubiquitin-conjugating enzyme E2 vari | Q8IX04-6;Q8IX04;Q8IX04-4;Q | UEVLD | Extracellular or secreted | 0.19715612 | 0.10776362 | 23.71366256 | 23.94672712 | 24.26119451 | 23.72525641 | 21.26798924 | 24.9992503 | 23.84047753 | 24.61610682 | 23.09192409 |
| Ubiquitin-fold modifier-conjugating er | Q9Y3C8 | UFC1 | Extracellular or secreted | 0.035503368 | 0.041383203 | 27.1723599 | 27.18470694 | 27.15286567 | 26.96113188 | 27.11733603 | 26.13627664 | 27.29096675 | 27.16211746 | 26.51576443 |
| Ubiquitin fusion degradation protein 1 | Q92890;Q92890-1;Q92890-3 | UFD1L | Other | -0.143217194 | 0.401653522 | 26.91113667 | 27.02981125 | 26.91934419 | 26.35972771 | 26.39078705 | 26.22492066 | 26.39037707 | 26.47892585 | 26.61890926 |
| E3 UFM1-protein ligase 1 | O94874;O94874-2;O94874-3 | UFL1 | Other | -0.231010951 | 0.41562246 | 26.10976531 | 25.95677459 | 25.98648215 | 24.61262422 | 25.52540037 | 25.84249307 | 25.38122898 | 25.30803261 | 25.6344754 |
| Ubiquitin-fold modifier 1 | P61960;P61960-2 | UFM1 | Extracellular or secreted | 0.295924648 | 0.220480243 | 27.2279342 | 27.61238518 | 27.57974637 | 27.75824416 | 27.35716215 | 25.84637225 | 27.95798135 | 27.9644959 | 26.65621884 |
| UDP-glucose 6-dehydrogenase | O60701;O60701-2;O60701-3 | UGDH | Extracellular or secreted | -0.587889059 | 2.17860852 | 29.71522166 | 29.80102361 | 29.81430331 | 29.15472561 | 29.06056817 | 29.54235351 | 28.97374776 | 28.93803491 | 28.86864808 |
| UDP-glucose:glycoprotein glucosyltra | Q9NYU2-2;Q9NYU2 | UGGT1 | Extracellular or secreted | -0.352617803 | 1.947209296 | 29.2662441 | 29.32911486 | 29.31943512 | 29.17928251 | 29.28942843 | 29.76599221 | 29.07986719 | 28.88869964 | 29.04832838 |
| UTP--glucose-1-phosphate uridylyltra | Q16851 | UGP2 | Extracellular or secreted | -0.131601358 | 0.224857041 | 25.45898302 | 25.7648888 | 25.83860343 | 25.74262852 | 25.77878882 | 24.3688629 | 25.32858427 | 25.41985862 | 25.33313078 |
| UTP--glucose-1-phosphate uridylyltra | Q16851-2 | UGP2 | Extracellular or secreted | 0.004376544 | 0.006147073 | 31.4125102 | 31.43574202 | 31.38391278 | 31.18441137 | 31.1669845 | 30.50758044 | 31.44282706 | 31.29214135 | 30.82373188 |
| E3 ubiquitin-protein ligase UHRF1 | Q96T88;Q96T88-2 | UHRF1 | Extracellular or secreted | -1.216481015 | 0.97654408 | 25.9276635 | 25.92818328 | 25.99187541 | 25.52811616 | 25.4592645 | 27.02601579 | 24.44941115 | 24.13680467 | 25.69490046 |
| UHRF1-binding protein 1-like | A0JNW5 | UHRF1BP1L | Other | -0.099901657 | 0.094165346 | 22.92611209 | 22.07444467 | 22.60220499 | 21.63815841 | 21.23931016 | 23.089744 | 22.42544643 | 22.39058043 | 21.66925533 |
| BRCA1-A complex subunit RAP80 | Q96RL1-2;Q96RL1-3;Q96RL1 | UIMC1 | Other | 0.924323534 | 1.2045503 | 20.82896974 | 22.27554483 | 20.27825698 | 21.3695942 | 22.64223709 | 22.34720195 | 22.49572721 | 22.46530633 | 22.68283946 |
| NKG2D ligand 2;Retinoic acid early tra | Q9BZM5;Q6H3X3-2;Q5VY80; | ULBP2;RAET1G;R | Extracellular or secreted | 0.72793554 | 0.607949336 | 25.97039979 | 26.29628106 | 26.2113927 | 25.95504572 | 25.9400196 | 27.84505641 | 26.74419514 | 26.57116252 | 27.9775466 |
| NKG2D ligand 3 | Q9BZM4 | ULBP3 | Extracellular or secreted | 1.071443074 | 1.588066382 | 20.59270513 | 22.46316085 | 20.90753594 | 20.5700644 | 21.06360213 | 21.5068809 | 22.37928 | 22.58570234 | 21.80132156 |
| Uridine 5-monophosphate synthase;O | P11172;P11172-3;P11172-2 | UMPS | Other | -0.059061201 | 0.087445277 | 27.55457211 | 27.7353768 | 27.54811741 | 27.62777109 | 27.70911979 | 28.20922793 | 27.31708589 | 28.04926361 | 27.64855946 |
| Protein unc-119 homolog B;Protein un | A6NIH7;Q13432-2;Q13432 | UNC119B;UNC11 | Other | 0.147581017 | 0.101995365 | 22.28957091 | 23.03960786 | 21.71544168 | 23.64801081 | 23.65938014 | 21.59989426 | 23.26642284 | 23.08527664 | 22.0669964 |
| Protein unc-45 homolog A | Q9H3U1-2;Q9H3U1 | UNC45A | Other | 0.07309005 | 0.138960209 | 28.68368156 | 28.62620504 | 28.6416173 | 28.82935353 | 28.75216241 | 28.49830763 | 28.94885025 | 28.89709315 | 28.38899049 |
| Uracil-DNA glycosylase | P13051-2;P13051 | UNG | Other | -0.011996269 | 0.007051073 | 26.16987776 | 26.12847064 | 26.10777164 | 23.46898889 | 24.34441783 | 25.50078621 | 25.478895 | 25.73298594 | 24.61228674 |
| Regulator of nonsense transcripts 1 | Q92900-2;Q92900 | UPF1 | Other | 0.164939873 | 0.424399863 | 28.78612532 | 28.90635597 | 28.90236472 | 28.38894945 | 28.42590417 | 29.50508322 | 29.04934152 | 29.00522816 | 28.89764136 |
| Regulator of nonsense transcripts 2 | Q9HAU5;Q9HAU5-2 | UPF2 | Other | -0.817394158 | 2.159057886 | 26.13860242 | 25.98945202 | 25.92788952 | 26.03655513 | 25.5133416 | 26.03615684 | 25.01554777 | 24.9693022 | 25.38396632 |
| Regulator of nonsense transcripts 3B | Q9BZI7-2;Q9BZI7 | UPF3B | Other | -1.1922346 | 0.462873625 | 24.12589206 | 24.28128768 | 23.89133872 | 23.70535396 | 23.55837554 | 23.86533003 | 20.79566689 | 23.95002471 | 23.3913936 |
| Ubiquinol-cytochrome-c reductase co | Q9NVA1-4;Q9NVA1-2;Q9NVA | UQCC1 | Other | -1.111426184 | 0.829640297 | 19.52327379 | 22.53602867 | 20.80723122 | 22.16776327 | 22.57497482 | 22.11235296 | 21.40768649 | 19.94934798 | 20.16949934 |
| Cytochrome b-c1 complex subunit 9 | Q9UDW1;Q9UDW1-2 | UQCR10 | Extracellular or secreted | -0.499371262 | 0.149511804 | 21.95681003 | 20.44239921 | 22.1455379 | 21.76312157 | 21.93800376 | 26.15392879 | 19.95460908 | 22.72014043 | 23.02703733 |
| Cytochrome b-c1 complex subunit 7 | P14927;P14927-2 | UQCRB | Other | 0.25707246 | 0.134521968 | 21.59009221 | 21.64061145 | 22.73092532 | 22.41766506 | 21.18907133 | 25.31879323 | 22.55517795 | 22.17678026 | 23.48283848 |
| Cytochrome b-c1 complex subunit 1, r | P31930 | UQCRC1 | Other | -0.424580975 | 0.569358542 | 27.22738341 | 27.11852465 | 27.50869523 | 27.84613309 | 27.97978295 | 29.28162375 | 27.24063559 | 27.27896848 | 27.68772455 |
| Cytochrome b-c1 complex subunit 2, r | P22695 | UQCRC2 | Extracellular or secreted | -0.25238459 | 0.051092219 | 23.0359891 | 23.65600057 | 23.8997515 | 23.64119006 | 22.735495 | 28.71672675 | 22.86912012 | 22.35116139 | 26.86514121 |
| Cytochrome b-c1 complex subunit Rie | P47985;P0C7P4 | UQCRFS1;UQCRF | Other | -0.256357142 | 0.329007758 | 26.19537296 | 25.96115397 | 25.98457127 | 26.06472106 | 26.13756703 | 27.03349173 | 26.39839143 | 26.04990753 | 25.47106862 |
| Cytochrome b-c1 complex subunit 6, r | P07919 | UQCRH | Other | -0.60043099 | 0.875006809 | 27.72858155 | 28.02342705 | 28.19347543 | 28.43077666 | 28.57819928 | 26.35247198 | 27.10219471 | 27.43920466 | 27.31077363 |
| Unconventional prefoldin RPB5 intera | O94763-2;O94763-3;O94763 | URI1 | Other | -0.050235665 | 0.043076263 | 24.45268214 | 24.54521808 | 23.94127404 | 24.88063654 | 24.56935268 | 22.40097317 | 24.27277158 | 23.7298265 | 24.24176325 |
| Uroporphyrinogen decarboxylase | P06132 | UROD | Other | 0.151866593 | 0.293780244 | 29.10512079 | 29.16851555 | 29.0742628 | 28.19309938 | 28.27017126 | 28.73046153 | 29.04241724 | 29.02709746 | 28.65690074 |
| Uroporphyrinogen-III synthase | P10746 | UROS | Other | 0.964414301 | 0.29163597 | 24.28637332 | 23.94261687 | 23.68645607 | 22.38419117 | 20.13780284 | 21.88164914 | 24.79897088 | 24.85908582 | 21.39473092 |
| General vesicular transport factor p11 | O60763;O60763-2 | USO1 | Other | -0.053249186 | 0.122044273 | 30.05452595 | 30.19095876 | 30.13332494 | 29.66292155 | 29.70665204 | 29.60891824 | 30.03564309 | 29.86599661 | 29.61726348 |
| Ubiquitin carboxyl-terminal hydrolase | Q14694;Q14694-3;Q14694-2 | USP10 | Other | 0.770197908 | 1.308679732 | 26.90816133 | 27.11882165 | 27.21706327 | 27.30486877 | 27.48508267 | 27.00206524 | 28.10474598 | 28.19455601 | 27.52932319 |
| Ubiquitin carboxyl-terminal hydrolase | P51784 | USP11 | Extracellular or secreted | 1.107366512 | 2.659672449 | 26.01507981 | 26.07814234 | 25.94270635 | 26.20761851 | 26.11406224 | 24.84598958 | 26.82602258 | 26.95004697 | 27.14782941 |
| Ubiquitin carboxyl-terminal hydrolase | P54578;P54578-2;P54578-3 | USP14 | Extracellular or secreted | 0.144667232 | 0.367053123 | 32.47860768 | 32.61221379 | 32.47563949 | 32.45329168 | 32.58542128 | 32.04775006 | 32.72313848 | 32.72900719 | 32.30831801 |
| Ubiquitin carboxyl-terminal hydrolase | Q9Y4E8-2;Q9Y4E8-3;Q9Y4E8 | USP15 | Other | 0.240243275 | 0.894504097 | 27.77931499 | 27.83291821 | 27.7509813 | 28.37542383 | 28.43432083 | 27.78849203 | 28.28876865 | 28.19826143 | 28.21442533 |
| Ubiquitin carboxyl-terminal hydrolase | O94966-7;O94966-6;O94966 | USP19 | Other | 0.156633444 | 0.127346916 | 24.35184884 | 23.76759367 | 23.6586177 | 23.71826947 | 24.42104728 | 23.36279035 | 24.49588576 | 24.37974271 | 23.23435552 |
| Ubiquitin carboxyl-terminal hydrolase | Q9UPU5 | USP24 | Other | -0.015562468 | 0.026380534 | 28.2994201 | 28.38574446 | 28.39427532 | 28.22121863 | 28.2828612 | 28.02981125 | 28.48861098 | 28.37563096 | 27.89573613 |
| Ubiquitin carboxyl-terminal hydrolase | Q9UHP3;Q9UHP3-1;Q9UHP3 | USP25 | Other | 0.559981751 | 0.914576633 | 24.46569244 | 24.52061293 | 24.47091352 | 24.35245514 | 22.67572635 | 23.77896423 | 24.78860404 | 24.50573064 | 24.51779288 |
| Ubiquitin carboxyl-terminal hydrolase | Q96RU2-2;Q96RU2;Q96RU2- | USP28 | Other | -0.150524102 | 0.406352626 | 23.29642111 | 23.43872892 | 23.32044691 | 23.17125683 | 23.04455221 | 24.13437806 | 23.16933112 | 23.26126626 | 23.32072234 |
| Ubiquitin carboxyl-terminal hydrolase | Q8NFA0 | USP32 | Other | 0.660775681 | 0.916397255 | 22.29484129 | 20.89074521 | 20.6858575 | 21.31606051 | 21.0295164 | 22.62432 | 22.29274738 | 22.34388944 | 21.76636067 |
| Ubiquitin carboxyl-terminal hydrolase | Q70CQ2-3;Q70CQ2-2;Q70CQ | USP34 | Other | -0.043231465 | 0.030618706 | 22.5130065 | 21.95025555 | 20.16249392 | 21.06535239 | 21.94320971 | 21.0264154 | 21.80289969 | 20.72708618 | 21.67068646 |
| U4/U6.U5 tri-snRNP-associated prote | Q53GS9;Q53GS9-3;Q53GS9-2 | USP39 | Other | -0.738354677 | 2.20268646 | 29.13716635 | 29.14206946 | 29.02894234 | 28.37579665 | 28.36189566 | 29.26985926 | 28.14477002 | 28.09541948 | 28.20261133 |
| Ubiquitin carboxyl-terminal hydrolase | Q96K76-4;Q96K76;Q96K76-2 | USP47 | Other | 2.299449675 | 2.132198483 | 21.23483031 | 23.25753057 | 22.67471727 | 23.0953088 | 22.75548304 | 24.06972771 | 25.64965615 | 25.43936321 | 24.35312851 |
| Ubiquitin carboxyl-terminal hydrolase | P45974-2;P45974 | USP5 | Other | 0.31033738 | 0.637490572 | 30.58244081 | 30.49743227 | 30.47314923 | 30.26785196 | 30.40466546 | 30.27097323 | 30.94551928 | 30.87518227 | 30.35856707 |
| Ubiquitin carboxyl-terminal hydrolase | Q93009-3;Q93009 | USP7 | Other | -0.212703384 | 0.993076623 | 31.67623425 | 31.72045 | 31.60302118 | 31.31651417 | 31.27425442 | 31.56215424 | 31.43524526 | 31.33394115 | 31.16901756 |
| Ubiquitin carboxyl-terminal hydrolase | P40818;P40818-2 | USP8 | Extracellular or secreted | 0.081283297 | 0.079678264 | 24.67502968 | 24.40204036 | 24.11941548 | 24.47004465 | 24.94547741 | 25.5167114 | 25.14250286 | 24.98567902 | 24.1800275 |
| Probable ubiquitin carboxyl-terminal h | Q93008-1;Q93008 | USP9X | Other | 0.407848087 | 3.387339988 | 28.47823156 | 28.60342804 | 28.49552747 | 28.44505931 | 28.19093522 | 28.38306819 | 28.87908863 | 28.85798318 | 28.78459736 |
| U3 small nucleolar RNA-associated pr | Q8TED0-3;Q8TED0;Q8TED0-2 | UTP15 | Other | -1.427020961 | 2.146951103 | 25.62544577 | 26.04370569 | 25.91394616 | 25.25078907 | 25.54751448 | 26.32651137 | 24.11314823 | 24.14133384 | 24.81841132 |
| U3 small nucleolar RNA-associated pr | Q9Y5J1 | UTP18 | Other | 0.861913876 | 0.598918807 | 22.64376813 | 23.43657024 | 24.0950673 | 21.55735127 | 21.63838885 | 22.99208294 | 22.71543098 | 23.55250681 | 24.4994182 |
| Small subunit processome componen | O75691 | UTP20 | Other | -2.353015369 | 0.84653163 | 22.95666826 | 23.31021891 | 22.62611451 | 21.36465164 | 21.96873991 | 29.18177223 | 20.46313718 | 20.32080496 | 22.86109448 |
| Something about silencing protein 10 | Q9NQZ2 | UTP3 | Other | -1.545389796 | 3.557320128 | 24.22449752 | 25.45186509 | 25.33036356 | 24.60907681 | 24.66161067 | 25.14557677 | 23.41015215 | 23.53931314 | 23.12586054 |
| Utrophin | P46939;P46939-2 | UTRN | Extracellular or secreted | 0.281802479 | 0.430519432 | 25.23352196 | 25.69763362 | 25.44161269 | 24.20717165 | 24.2719172 | 24.28750103 | 24.95091466 | 25.19839258 | 25.26577927 |
| Urotensin-2 | O95399;O95399-2 | UTS2 | Extracellular or secreted | 1.889214142 | 0.55508666 | 22.59435929 | 29.67418117 | 29.72142821 | 22.21630605 | 29.61647495 | 30.06033625 | 29.18072944 | 29.12430522 | 29.30415072 |
| UDP-glucuronic acid decarboxylase 1 | Q8NBZ7-3;Q8NBZ7;Q8NBZ7-1 | UXS1 | Extracellular or secreted | 0.743582709 | 0.845405814 | 23.34678639 | 21.32479821 | 23.44632207 | 23.92822847 | 23.91879847 | 24.06274645 | 24.2128217 | 23.81509164 | 24.21667482 |
| Protein UXT | Q9UBK9;Q9UBK9-2 | UXT | Other | 0.15006477 | 0.154915544 | 24.2859502 | 24.13186889 | 23.57807688 | 24.98672083 | 22.83770877 | 24.94721778 | 24.32786509 | 23.97867066 | 24.52743023 |
| Protein VAC14 homolog | Q08AM6 | VAC14 | Other | -0.172320079 | 0.415438984 | 27.16853946 | 27.55508431 | 27.49663256 | 27.0480165 | 27.12780183 | 27.36603 | 26.97366566 | 26.96917043 | 27.421256 |
| Vesicle-associated membrane protein | Q15836 | VAMP3 | Extracellular or secreted | 0.124648494 | 0.384434694 | 26.31522103 | 26.09139955 | 26.27515405 | 26.26381104 | 26.03120042 | 25.55806611 | 26.3157911 | 26.27831276 | 26.04726772 |
| Vesicle-associated membrane protein | P51809;P51809-2;P51809-3 | VAMP7 | Extracellular or secreted | -0.235560786 | 0.873660957 | 25.84878548 | 25.91230224 | 25.75340632 | 25.13868053 | 25.27009995 | 25.70012345 | 25.38063465 | 25.40548533 | 25.31889664 |
| Vesicle-associated membrane protein | Q9BV40 | VAMP8 | Extracellular or secreted | 0.432932923 | 0.783857343 | 24.4388558 | 24.65387064 | 24.09119776 | 23.96631252 | 24.10637444 | 25.66364779 | 24.98307119 | 25.0442687 | 24.73158835 |
| Vang-like protein 1 | Q8TAA9-2;Q8TAA9 | VANGL1 | Extracellular or secreted | 2.290851743 | 1.248791389 | 24.19249751 | 21.10924096 | 22.03313868 | 24.92782172 | 19.45526751 | 24.96525586 | 24.92497121 | 25.18914499 | 25.10005015 |
| Vesicle-associated membrane protein | Q9P0L0 | VAPA | Extracellular or secreted | -0.410240604 | 0.528624514 | 30.62226501 | 30.34783831 | 30.29701616 | 29.09368291 | 28.96743987 | 30.88678255 | 29.44069035 | 29.51868974 | 29.91741051 |
| Vesicle-associated membrane protein | O95292 | VAPB | Other | -0.431310849 | 0.514784256 | 29.41757896 | 29.08542456 | 29.16228556 | 27.70034838 | 27.94475164 | 29.68135421 | 28.08749991 | 28.3273169 | 28.7871223 |
| Valine--tRNA ligase | P26640 | VARS | Other | 0.365312703 | 1.304322884 | 30.45757868 | 30.42462341 | 30.36942465 | 30.53899136 | 30.64941632 | 31.10886352 | 30.77893917 | 30.96333868 | 31.12810922 |
| Vasorin | Q6EMK4 | VASN | Extracellular or secreted | 0.129413429 | 0.076280349 | 23.1129574 | 23.02820151 | 22.98507963 | 24.31347485 | 23.14645494 | 24.13108388 | 23.29305622 | 22.84676437 | 24.6070458 |
| Vasodilator-stimulated phosphoprote | P50552 | VASP | Extracellular or secreted | -0.174299875 | 0.548236697 | 29.11807903 | 29.22763588 | 29.21071481 | 28.66488901 | 28.71872021 | 28.91547423 | 28.93382102 | 28.87929316 | 28.59174278 |
| Synaptic vesicle membrane protein VA | Q99536;Q99536-3;Q99536-2 | VAT1 | Extracellular or secreted | 0.412237552 | 1.420422554 | 28.97530677 | 29.08245822 | 29.0125193 | 28.44011607 | 28.60349878 | 28.23676393 | 29.05721451 | 29.0684828 | 29.28634688 |
| Guanine nucleotide exchange factor V | P52735-3;P52735-2;P52735 | VAV2 | Extracellular or secreted | 0.088098741 | 0.103162869 | 27.71894879 | 27.83683522 | 28.12903102 | 27.97880157 | 28.06321978 | 28.18200912 | 27.77385605 | 27.80593522 | 28.6389277 |
| Prefoldin subunit 3 | P61758 | VBP1 | Other | 0.309904805 | 1.976618902 | 29.33766726 | 29.49510807 | 29.48064086 | 29.38189318 | 29.38673137 | 29.17951981 | 29.73866584 | 29.75144428 | 29.57038457 |
| Vinculin | P18206-2;P18206 | VCL | Extracellular or secreted | 0.731921153 | 1.975972149 | 24.73241671 | 25.12750667 | 24.65064246 | 24.47537375 | 24.40944014 | 23.98259257 | 25.21989021 | 25.11024338 | 25.55461602 |
| Transitional endoplasmic reticulum AT | P55072 | VCP | Extracellular or secreted | 0.034865335 | 0.031967704 | 34.85726208 | 34.8766923 | 34.81416386 | 34.12455482 | 34.23242158 | 33.60531931 | 34.76549633 | 34.74359542 | 33.85071123 |
| Deubiquitinating protein VCIP135 | Q96JH7 | VCPIP1 | Other | 0.173898026 | 0.19916634 | 20.75946668 | 22.57968883 | 22.55910537 | 23.01120768 | 21.56072886 | 22.44245456 | 22.40629014 | 22.28717691 | 22.28455303 |
| Voltage-dependent anion-selective ch | P21796 | VDAC1 | Extracellular or secreted | -0.097122642 | 0.026568285 | 23.24123954 | 24.8997515 | 24.68944523 | 23.27440773 | 23.2397838 | 28.96543202 | 23.20445771 | 24.47215386 | 26.18705042 |
| Voltage-dependent anion-selective ch | P45880-2;P45880;P45880-1 | VDAC2 | Extracellular or secreted | -0.751845732 | 0.244391004 | 24.57525163 | 24.47264969 | 25.23516669 | 25.27444328 | 25.19917927 | 28.51307047 | 23.06496769 | 24.67793509 | 26.63644053 |
| Voltage-dependent anion-selective ch | Q9Y277;Q9Y277-2 | VDAC3 | Extracellular or secreted | -1.62644368 | 0.570136189 | 22.42869548 | 21.58936175 | 24.70255614 | 23.39756987 | 24.29852019 | 26.63048155 | 20.1939327 | 22.79171106 | 23.6586177 |
| Neurosecretory protein VGF;Neuroen | O15240 | VGF | Extracellular or secreted | 2.91682342 | 0.902560844 | 21.90245229 | 22.69452806 | 23.10343172 | 21.8189668 | 21.58825272 | 21.9297928 | 23.58279251 | 24.28827582 | 27.39811412 |
| Villin-like protein | O15195-2;O15195 | VILL | Other | 0.292367404 | 0.328111909 | 22.72158214 | 21.19434565 | 20.88707913 | 22.45512696 | 21.8494735 | 20.81391452 | 21.91630373 | 21.63582084 | 22.2857386 |
| Vimentin | P08670 | VIM | Extracellular or secreted | 4.400110888 | 1.401632547 | 30.82000742 | 23.90545532 | 23.97211036 | 22.76826026 | 22.46806913 | 24.78311774 | 27.52080773 | 28.68009689 | 31.35793817 |
| Protein VPRBP | Q9Y4B6-2;Q9Y4B6;Q9Y4B6-3 | VPRBP | Other | 0.931088317 | 1.170074651 | 25.99105364 | 26.30243031 | 26.6345031 | 25.50824186 | 25.49436049 | 26.13811412 | 27.36302439 | 27.11674134 | 26.34785098 |
| Vacuolar protein sorting-associated p | Q9H269;Q9H269-2 | VPS16 | Other | 0.660923402 | 0.31999854 | 23.76445858 | 20.66473542 | 23.69115054 | 23.49203898 | 23.57853762 | 22.42897077 | 24.5438031 | 23.99403571 | 22.25487735 |
| Vacuolar protein sorting-associated p | Q9P253 | VPS18 | Other | -0.706285947 | 0.846682247 | 22.90442662 | 22.4927236 | 22.63858528 | 23.9097642 | 22.46555544 | 22.15610283 | 22.18046768 | 22.52152376 | 21.46272971 |
| Vacuolar protein sorting-associated p | O75436;O75436-2 | VPS26A | Extracellular or secreted | 0.423378574 | 1.226750468 | 29.29900516 | 29.73293743 | 29.5891014 | 29.28951638 | 29.18108502 | 29.64903937 | 29.82429351 | 29.68212483 | 30.13405976 |
| Vacuolar protein sorting-associated p | Q4G0F5 | VPS26B | Other | 0.066255115 | 0.181370688 | 26.09160132 | 26.54206194 | 26.44189759 | 26.12208467 | 26.74021104 | 26.41565835 | 26.57773123 | 26.53731475 | 26.26047682 |
| Vacuolar protein sorting-associated p | Q9UK41;Q9UK41-2 | VPS28 | Extracellular or secreted | 0.710926546 | 0.888346138 | 24.86258969 | 24.80163732 | 24.58583224 | 25.50754642 | 25.44063096 | 26.26515331 | 25.58106907 | 25.74124025 | 26.54216529 |
| Vacuolar protein sorting-associated p | Q9UBQ0;Q9UBQ0-2 | VPS29 | Extracellular or secreted | 0.300968648 | 0.938107897 | 28.79628092 | 28.85993889 | 28.7950115 | 28.17302744 | 28.02627969 | 28.38701909 | 28.80338142 | 28.94648173 | 28.67182156 |
| Vacuolar protein sorting-associated p | Q9H267;Q9H267-2 | VPS33B | Other | -0.589499384 | 0.843489391 | 24.58147139 | 24.51875351 | 24.8547653 | 24.25313595 | 24.43657024 | 24.76915866 | 23.52145188 | 24.43447196 | 23.98250553 |
| Vacuolar protein sorting-associated p | Q96QK1 | VPS35 | Extracellular or secreted | 0.584751892 | 2.4279829 | 29.97841973 | 30.25327123 | 30.29559376 | 29.73085019 | 29.58659833 | 30.21604569 | 30.68277786 | 30.56964241 | 30.53222488 |
| Vacuolar protein-sorting-associated p | Q86VN1-2;Q86VN1 | VPS36 | Extracellular or secreted | -1.037263903 | 0.348539905 | 24.57120593 | 23.83016391 | 23.98058937 | 23.53670728 | 22.82886028 | 22.41439906 | 20.30288118 | 23.77806189 | 23.38822814 |
| Vacuolar protein sorting-associated p | Q9H9H4 | VPS37B | Extracellular or secreted | 0.562953106 | 1.421954357 | 25.36636357 | 25.39073786 | 25.30542555 | 25.62123115 | 25.78880315 | 26.1915001 | 26.02812561 | 26.01771541 | 26.47504899 |
| Vacuolar protein sorting-associated p | A5D8V6 | VPS37C | Extracellular or secreted | 1.204837274 | 1.089473108 | 22.89568991 | 22.28878273 | 22.98959061 | 23.27326973 | 23.05143931 | 22.0687697 | 23.48296154 | 23.62075589 | 24.79456539 |
| Vacuolar protein sorting-associated p | P49754-2;P49754-3;P49754 | VPS41 | Other | 0.538979893 | 0.824263616 | 22.13814596 | 22.77374289 | 22.8569794 | 22.74943737 | 22.82164554 | 21.02593501 | 22.87640828 | 23.22298058 | 22.7004939 |
| Vacuolar protein sorting-associated p | Q9NRW7;Q9NRW7-2 | VPS45 | Other | 0.066660374 | 0.10259295 | 26.02307801 | 26.3148927 | 26.31032293 | 25.23786727 | 25.25472243 | 26.02863151 | 26.11026329 | 26.00081961 | 25.67365565 |
| Vacuolar protein sorting-associated p | Q9UN37 | VPS4A | Extracellular or secreted | 0.13015721 | 0.200657805 | 24.15073709 | 24.67567583 | 24.60161575 | 24.19174481 | 23.92352118 | 24.27625506 | 24.0792009 | 24.8239596 | 24.397086 |
| Vacuolar protein sorting-associated p | O75351 | VPS4B | Extracellular or secreted | 0.067111727 | 0.215661574 | 29.26001911 | 29.32098481 | 29.3021906 | 29.32272622 | 29.3402592 | 29.85698949 | 29.51551632 | 29.58281763 | 29.30458594 |

| | | | | | | | | | | | | | | |
|---|---|---|---|---|---|---|---|---|---|---|---|---|---|---|
| Vacuolar protein sorting-associated pı | Q9UID3-2;Q9UID3 | VPS51 | Extracellular or secreted | -0.57167221 | 0.298595894 | 22.6953413 | 23.90747384 | 23.82036052 | 23.88753226 | 24.05796331 | 23.3431304 | 23.77906445 | 23.70725127 | 21.65456846 |
| Vacuolar protein sorting-associated pı | Q8N1B4-2;Q8N1B4 | VPS52 | Other | 1.42392643 | 1.534342059 | 23.47072738 | 23.22671852 | 21.4630424 | 20.95360542 | 23.13732456 | 21.15020945 | 23.53076732 | 23.58198849 | 23.85983733 |
| Vacuolar protein sorting-associated pı | Q5VIR6-3;Q5VIR6;Q5VIR6-4;( | VPS53 | Other | -0.11978698 | 1.49249259 | 24.29130056 | 24.11569014 | 24.04635202 | 24.18465019 | 24.23753909 | 24.05109075 | 24.02976913 | 24.01894654 | 24.05523477 |
| Serine/threonine-protein kinase VRK1 | Q99986 | VRK1 | Other | -0.374482075 | 0.407447546 | 23.42265202 | 21.54019685 | 23.27866724 | 23.59400196 | 23.99584787 | 24.27540274 | 22.92842729 | 23.15197592 | 22.84953491 |
| Visinin-like protein 1 | P62760 | VSNL1 | Other | 0.246644514 | 0.16004852 | 24.33101313 | 24.89078231 | 24.80666053 | 25.81580015 | 25.22059147 | 27.0115866 | 25.32926887 | 24.99662382 | 26.45225796 |
| Vacuolar protein sorting-associated pı | Q9NP79 | VTA1 | Extracellular or secreted | -0.016261752 | 0.031293616 | 28.21660082 | 28.09672681 | 28.0880562 | 28.17317044 | 28.04702845 | 27.77931499 | 28.13565058 | 28.27124045 | 27.74477257 |
| Vesicle transport through interaction | Q9UEU0 | VTI1B | Extracellular or secreted | -0.49237376 | 0.963695575 | 24.61211798 | 24.52808634 | 24.63389366 | 24.43358086 | 24.49112156 | 24.34374038 | 24.28806456 | 24.09224678 | 23.66383777 |
| von Willebrand factor A domain-conta | Q6PCB0 | VWA1 | Extracellular or secreted | 1.678973247 | 1.037407632 | 21.09242789 | 22.75694421 | 21.67356977 | 23.27611304 | 23.407302 | 24.57588635 | 24.00084109 | 23.73469226 | 25.69250802 |
| von Willebrand factor A domain-conta | Q96SY0-4;Q96SY0-3;Q96SY0 | VWA9 | Other | -0.076818642 | 0.20564624 | 25.97605012 | 26.07677728 | 26.15971384 | 25.96252256 | 25.67341303 | 26.53593647 | 25.98450608 | 26.15073709 | 25.82650756 |
| WW domain-containing adapter prote | Q9BTA9-5;Q9BTA9-2;Q9BTA9 | WAC | Other | -0.045642406 | 0.026397284 | 25.06332266 | 24.93750745 | 24.36833007 | 21.74051433 | 23.52664379 | 24.56877305 | 24.36459477 | 23.37888327 | 24.22214042 |
| Wings apart-like protein homolog | Q7Z5K2;Q7Z5K2-2;Q7Z5K2-3 | WAPAL | Other | -0.978733304 | 0.754976302 | 26.68351425 | 23.19914931 | 23.33538163 | 24.50233507 | 22.46438421 | 24.72504833 | 23.00937554 | 23.03860356 | 23.47072738 |
| Tryptophan--tRNA ligase, cytoplasmic | P23381;P23381-2 | WARS | Extracellular or secreted | -0.160551421 | 0.37044791 | 30.97514274 | 30.99980943 | 30.92287782 | 30.69627432 | 30.68219182 | 29.94907297 | 30.65693483 | 30.63122777 | 30.34286769 |
| Wiskott-Aldrich syndrome protein fan | Q9Y6W5;Q9Y6W5-2 | WASF2 | Extracellular or secreted | -0.063540159 | 0.15100837 | 26.61932915 | 26.85376642 | 26.83279751 | 26.34516265 | 26.59120776 | 26.07428832 | 26.57816328 | 26.61006308 | 26.27942908 |
| Wiskott-Aldrich syndrome protein fan | Q9UPY6;Q9UPY6-2 | WASF3 | Extracellular or secreted | 0.756024669 | 0.914497336 | 21.31853774 | 20.99216329 | 22.71792116 | 22.0284376 | 21.28362051 | 23.29824049 | 22.37116511 | 22.62207506 | 23.09429422 |
| Putative WAS protein family homolog | C4AMC7;Q6VEQ5;A8K0Z3 | WASH3P;WASH2 | Other | 0.658912877 | 0.800167564 | 25.52258967 | 25.17466637 | 25.17306558 | 24.45951467 | 22.85399479 | 24.22589524 | 25.14615969 | 25.33517715 | 25.20026495 |
| WAS protein family homolog 6 | Q9NQA3 | WASH6P | Other | -0.01298465 | 0.008210486 | 22.31780011 | 20.9760023 | 21.56275824 | 22.09852586 | 22.4033018 | 20.7734409 | 22.13905541 | 22.07776758 | 20.81013766 |
| Neural Wiskott-Aldrich syndrome prot | O00401 | WASL | Extracellular or secreted | -0.041406387 | 0.069128453 | 25.69256123 | 25.34458714 | 25.62343769 | 26.24655505 | 26.44531195 | 25.44404832 | 25.98563559 | 25.67133568 | 25.61706026 |
| WW domain-binding protein 11 | Q9Y2W2 | WBP11 | Other | 0.065576979 | 0.037643996 | 27.22692426 | 27.27550931 | 27.05734364 | 27.80722578 | 27.52611712 | 27.98304943 | 26.84361958 | 27.17169205 | 28.61950407 |
| WW domain-binding protein 2 | Q969T9;Q969T9-2 | WBP2 | Other | 1.541435163 | 3.464189727 | 24.72484022 | 25.28637332 | 24.51328135 | 25.18933354 | 24.93539565 | 24.45475412 | 26.53537292 | 26.53504656 | 26.10587511 |

| | | | | | | | | | | | | | | |
|---|---|---|---|---|---|---|---|---|---|---|---|---|---|---|
| WW domain-binding protein 4 | O75554-2;O75554 | WBP4 | Extracellular or secreted | 0.672557455 | 0.827198604 | 22.92339423 | 22.9376152 | 23.02860622 | 23.49118274 | 21.68996634 | 24.27042081 | 24.07398201 | 23.82220981 | 23.29207332 |
| Probable 18S rRNA (guanine-N(7))-me | O43709;O43709-3 | WBSCR22 | Other | 0.298004054 | 0.317026822 | 21.05911987 | 21.41016189 | 21.56825423 | 22.56497653 | 22.21744421 | 22.93854873 | 21.9447052 | 22.04343453 | 22.78512516 |
| WD repeat and FYVE domain-containi | Q8IWB7 | WDFY1 | Other | 0.079737715 | 0.58446172 | 26.6945817 | 26.58305085 | 26.5923489 | 26.40186139 | 26.5317491 | 26.75366134 | 26.74188313 | 26.67465262 | 26.60130403 |
| WD repeat and HMG-box DNA-bindin | O75717;O75717-2 | WDHD1 | Other | 0.180921241 | 0.694563691 | 29.02448424 | 29.04377347 | 28.98937081 | 28.85257637 | 28.94720663 | 28.78979828 | 29.2423631 | 29.19992342 | 28.9240821 |
| WD repeat-containing protein 1 | O75083;O75083-3 | WDR1 | Extracellular or secreted | -0.149886461 | 0.433050644 | 32.78534595 | 32.8216238 | 32.75908991 | 32.46329117 | 32.42514886 | 32.22825539 | 32.5280146 | 32.56233623 | 32.20136734 |
| WD repeat-containing protein 11 | Q9BZH6 | WDR11 | Other | -0.584062454 | 1.445526606 | 25.45556954 | 25.68107602 | 25.11798989 | 25.22140303 | 25.2992192 | 25.25860916 | 24.99761469 | 24.48511339 | 24.78201798 |
| Ribosome biogenesis protein WDR12 | Q9GZL7 | WDR12 | Other | -0.165785206 | 0.321452076 | 29.83223918 | 29.82880899 | 29.77602327 | 29.91513227 | 30.035512 | 29.28151321 | 29.75605004 | 29.82527964 | 29.25592916 |
| WD repeat-containing protein 13 | Q9H1Z4-2;Q9H1Z4 | WDR13 | Other | -0.151610358 | 0.340524349 | 24.61329892 | 24.40872778 | 24.46326193 | 23.8624951 | 23.56685862 | 24.06266412 | 23.8345104 | 24.01784282 | 24.18146895 |
| WD repeat-containing protein 18 | Q9BV38 | WDR18 | Other | 0.278117098 | 0.4680539 | 25.18646491 | 25.1287656 | 25.37318484 | 25.48440671 | 25.29789078 | 26.77228356 | 25.83942064 | 25.97322772 | 25.64320113 |
| WD repeat-containing protein 20 | Q8TBZ3-4;Q8TBZ3-5;Q8TBZ3 | WDR20 | Other | 0.605497848 | 1.791701466 | 23.64416435 | 24.19813026 | 24.11949464 | 23.87492733 | 23.26527852 | 24.07585972 | 24.22831963 | 24.58743555 | 24.58966577 |
| WD repeat-containing protein 26 | Q9H7D7-2;Q9H7D7 | WDR26 | Other | -0.20470552 | 0.481432044 | 26.89735285 | 27.24099945 | 27.25106006 | 27.00388879 | 27.08405624 | 26.85721202 | 27.153349 | 26.789052 | 26.61076714 |
| WD repeat-containing protein 3 | Q9UNX4 | WDR3 | Other | -2.061924713 | 3.91868595 | 25.4683675 | 25.52007334 | 26.04199446 | 24.76233094 | 25.11350596 | 26.14726658 | 23.23718895 | 23.35420525 | 23.74960104 |
| pre-mRNA 3 end processing protein W | Q9C0J8 | WDR33 | Other | -0.867258762 | 2.475882072 | 25.80037878 | 25.57801928 | 25.44347932 | 25.67667142 | 25.81333106 | 25.85566845 | 24.61565792 | 24.91355818 | 24.95278176 |
| WD repeat-containing protein 36 | Q8NI36 | WDR36 | Other | -1.485490499 | 0.629661328 | 21.77518203 | 23.56755507 | 23.26327379 | 23.99377665 | 23.81147034 | 25.52246995 | 22.39558329 | 20.51955437 | 23.59525475 |
| tRNA (guanine-N(7)-)-methyltransfera | P57081-2;P57081;P57081-3 | WDR4 | Other | 0.298283386 | 0.632225831 | 23.73974765 | 23.94770827 | 24.12801824 | 23.81665479 | 22.74638549 | 23.99144296 | 23.91086228 | 24.22831963 | 23.94064696 |
| WD repeat-containing protein 43 | Q15061 | WDR43 | Other | -0.811969433 | 2.872292723 | 27.55756957 | 27.77391891 | 27.74072573 | 27.35984498 | 27.44986765 | 28.28498009 | 26.72598444 | 26.92013965 | 27.00142108 |
| WD repeat-containing protein 44 | Q5JSH3;Q5JSH3-2;Q5JSH3-4 | WDR44 | Other | 0.638995449 | 0.614448959 | 27.38870316 | 27.25412771 | 27.13437806 | 26.6056902 | 26.63436461 | 25.84018936 | 27.83394368 | 27.84535557 | 26.66641365 |
| WD repeat domain phosphoinositide- | Q5MNZ6 | WDR45B | Other | 0.328841447 | 0.498422373 | 24.73252022 | 24.38482274 | 24.22464472 | 23.80370782 | 23.6946614 | 23.70524848 | 24.66084942 | 24.64542932 | 23.9530483 |
| WD repeat-containing protein 46 | O15213 | WDR46 | Other | -1.556660279 | 1.244708597 | 21.48870708 | 23.38548117 | 23.40444621 | 24.01128801 | 22.06467166 | 25.74601645 | 21.38701794 | 21.65443206 | 22.33887445 |
| WD repeat-containing protein 48 | Q8TAF3-3;Q8TAF3;Q8TAF3-4 | WDR48 | Other | 0.004273665 | 0.011820674 | 27.53926876 | 27.68036521 | 27.53356215 | 27.38706019 | 27.49327656 | 27.420694 | 27.37260379 | 27.44245139 | 27.72487925 |
| WD repeat-containing protein 5 | P61964 | WDR5 | Other | -0.387747444 | 3.933905969 | 30.10412109 | 30.0457013 | 29.97473259 | 29.9564312 | 29.96526687 | 29.89463803 | 29.56770375 | 29.57805528 | 29.66144418 |
| WD repeat-containing protein 55 | Q9H6Y2;Q9H6Y2-2 | WDR55 | Other | 0.177425033 | 0.158677249 | 24.14335955 | 24.09143991 | 23.21270778 | 24.27703591 | 24.55432326 | 23.75052136 | 24.89550502 | 23.95393638 | 23.69752758 |
| WD repeat-containing protein 6 | Q9NNW5 | WDR6 | Other | -1.106703563 | 0.328391564 | 24.04785014 | 24.25601917 | 24.0993279 | 24.74827066 | 24.66768633 | 23.82842164 | 24.4171886 | 24.41628676 | 20.67020188 |
| WD repeat-containing protein 61;WD | Q9GZS3 | WDR61 | Other | 0.395221345 | 0.709056147 | 27.32680278 | 27.34401139 | 27.26695892 | 27.08040101 | 27.21178257 | 26.75059802 | 27.86555422 | 27.66275181 | 27.14763535 |
| WD repeat-containing protein 64 | B1ANS9-2;B1ANS9 | WDR64 | Other | -1.164164186 | 0.626910669 | 25.27070595 | 24.86499957 | 24.07234723 | 27.66424479 | 28.15735441 | 25.69237499 | 23.78236799 | 24.74709277 | 25.83906016 |
| WD repeat-containing protein 75 | Q8IWA0 | WDR75 | Other | -1.484508168 | 2.703220712 | 25.46104598 | 24.82113946 | 26.04253726 | 24.52486258 | 24.28206582 | 25.04959599 | 23.40691291 | 23.35769911 | 23.87248702 |
| Methylosome protein 50 | Q9BQA1;Q9BQA1-2 | WDR77 | Other | 0.102703467 | 0.290642342 | 29.30938642 | 29.23505253 | 29.14654331 | 29.53930574 | 29.38156318 | 28.52924871 | 29.30732482 | 29.23139475 | 29.33994077 |
| WD repeat-containing protein 82 | Q6UXN9 | WDR82 | Other | 0.449588442 | 1.211462342 | 27.8754899 | 27.83394368 | 27.90466319 | 27.39941862 | 26.90357187 | 28.17250297 | 28.07229611 | 28.25178245 | 28.06948189 |
| WD repeat domain-containing protei | Q9BRX9 | WDR83 | Other | -1.341038391 | 2.723014173 | 23.96578428 | 24.5168316 | 24.51038655 | 24.11131846 | 24.10613478 | 23.70059973 | 23.08882252 | 22.82189852 | 22.52169149 |
| WD repeat-containing protein 89 | Q96FK6 | WDR89 | Other | 0.039859475 | 0.040399281 | 24.15930965 | 24.48946875 | 24.11457861 | 23.27554483 | 23.21939544 | 23.21533322 | 24.10725284 | 23.95215966 | 23.29698116 |
| WD repeat-containing protein 92 | Q96MX6;Q96MX6-2 | WDR92 | Other | 0.098217505 | 0.279272361 | 26.89226561 | 26.7509813 | 26.65785486 | 26.63325623 | 26.34229974 | 26.40184511 | 26.79723999 | 26.85150449 | 26.48515947 |
| WD repeat, SAM and U-box domain-c | Q8N9V3-2;Q8N9V3 | WDSUB1 | Other | 1.182827003 | 1.003848861 | 24.17417108 | 23.78825553 | 24.15599479 | 21.42486659 | 21.01642674 | 23.58233313 | 24.46531878 | 24.22758539 | 23.92660076 |
| Wolframin | O76024 | WFS1 | Other | -2.308579605 | 1.372701274 | 23.61020392 | 23.46513192 | 23.35380157 | 21.55099076 | 22.68242361 | 23.24684496 | 19.94343424 | 20.18561964 | 21.89990567 |
| Partner of Y14 and mago | Q9BRP8-2;Q9BRP8 | WIBG | Other | 2.418472974 | 5.119068813 | 25.58689177 | 25.30903941 | 25.52058295 | 26.16915141 | 26.11312835 | 25.09796267 | 28.08714579 | 28.22075751 | 27.8458939 |
| WAS/WASL-interacting protein family | Q8TF74;Q8TF74-2 | WIPF2 | Other | -0.595154885 | 0.148648489 | 23.94601314 | 23.88725335 | 24.25198104 | 23.84038148 | 23.81088225 | 22.45231393 | 20.34599739 | 24.46438421 | 24.49856633 |
| WD repeat domain phosphoinositide- | Q9Y4P8-2;Q9Y4P8-4;Q9Y4P8 | WIPI2 | Other | 0.840900481 | 1.371139533 | 23.78166787 | 24.13672645 | 25.29722612 | 23.66242586 | 24.54510022 | 23.10381604 | 24.77399435 | 25.04918051 | 24.96300788 |
| Protein Wiz | O95785-2;O95785-4;O95785 | WIZ | Extracellular or secreted | 0.121391065 | 0.059047913 | 24.87305053 | 25.11115924 | 25.83696761 | 21.53592863 | 26.04857784 | 25.6747873 | 25.16254008 | 25.25111425 | 24.49075444 |
| Serine/threonine-protein kinase WNK | Q9H4A3-2;Q9H4A3;Q9H4A3- | WNK1 | Extracellular or secreted | -0.313420997 | 1.780062235 | 27.59960257 | 27.36636357 | 27.41578728 | 27.59583806 | 27.71220675 | 27.1806346 | 27.13133905 | 27.28118154 | 27.08243284 |
| Serine/threonine-protein kinase WNK | Q9Y3S1-2;Q9Y3S1-4;Q9Y3S1; | WNK2 | Extracellular or secreted | -0.535378515 | 0.342608824 | 23.30033692 | 23.37583807 | 23.17285963 | 23.92279561 | 23.46911314 | 24.00178614 | 23.28658483 | 21.8927844 | 23.83585997 |
| Telomerase Cajal body protein 1 | Q9BUR4 | WRAP53 | Other | 0.141327772 | 0.224972108 | 25.76749264 | 25.90474357 | 25.82310927 | 25.89705274 | 26.12407839 | 25.27408776 | 26.16689355 | 26.13478931 | 25.51758265 |
| ATPase WRNIP1 | Q96S55-2;Q96S55;Q96S55-3 | WRNIP1 | Other | 0.507655349 | 1.756247045 | 26.14709186 | 26.54651394 | 26.65908066 | 26.22716304 | 26.33459764 | 26.04589395 | 27.00474613 | 26.6285361 | 26.86985436 |
| Pre-mRNA-splicing regulator WTAP | Q15007;Q15007-2 | WTAP | Other | -1.085566623 | 4.169316832 | 27.50982803 | 27.55032112 | 27.53808483 | 27.02664906 | 26.97869248 | 27.25187272 | 26.15323302 | 26.36831342 | 26.14947781 |
| NEDD4-like E3 ubiquitin-protein ligase | O00308-2;O00308;O00308-4 | WWP2 | Extracellular or secreted | 0.291479195 | 0.14554239 | 23.21010352 | 20.38204307 | 20.64526438 | 23.06977687 | 23.16260461 | 21.32003367 | 22.72456964 | 22.85029826 | 21.19448275 |
| Pre-mRNA-splicing factor SYF1 | Q9HCS7 | XAB2 | Other | 0.340707709 | 0.3043227 | 25.52516138 | 22.77950536 | 23.78286787 | 24.85262399 | 24.83518534 | 25.65313263 | 25.10029082 | 24.98849886 | 24.64757173 |
| E3 ubiquitin-protein ligase XIAP | P98170 | XIAP | Other | 0.819420253 | 0.3881575 | 22.69920218 | 22.96726285 | 22.53207621 | 23.6722803 | 22.9906296 | 21.15018648 | 24.24219953 | 24.28376211 | 21.93811794 |
| Xaa-Pro aminopeptidase 1 | Q9NQW7-3;Q9NQW7;Q9NQ\ | XPNPEP1 | Extracellular or secreted | 0.348615718 | 0.610282838 | 28.63526455 | 28.71243637 | 28.61887426 | 28.58763584 | 28.65659393 | 27.40178003 | 29.02883698 | 28.88302815 | 28.44027451 |
| Probable Xaa-Pro aminopeptidase 3 | Q9NQH7-4;Q9NQH7;Q9NQH | XPNPEP3 | Extracellular or secreted | 0.334318375 | 0.238918075 | 25.19238463 | 25.10665399 | 25.0499283 | 24.66041423 | 24.50415512 | 24.60715871 | 25.4468584 | 25.89659091 | 24.2198533 |
| Exportin-1 | O14980 | XPO1 | Other | -0.053607994 | 0.061003529 | 31.39110265 | 31.73008087 | 31.64125553 | 30.84647686 | 30.90684336 | 31.2301467 | 31.60681996 | 31.37316409 | 30.73214496 |
| Exportin-4 | Q9C0E2 | XPO4 | Other | -0.052649417 | 0.073323107 | 27.5497338 | 27.88145363 | 27.80452042 | 27.14520747 | 27.57297009 | 27.10089232 | 27.80741005 | 27.48753806 | 27.0744925 |
| Exportin-5 | Q9HAV4 | XPO5 | Other | 0.284623234 | 0.70105067 | 28.61470355 | 28.8363537 | 28.73120637 | 28.74679173 | 28.77953417 | 29.11107962 | 29.29533101 | 29.18527426 | 28.783099 |
| Exportin-7 | Q9UIA9 | XPO7 | Other | -0.21863436 | 0.375355706 | 30.27974344 | 30.407302 | 30.3271027 | 29.72219401 | 29.74971931 | 29.8766612 | 30.08474056 | 30.00691401 | 29.43380368 |
| Exportin-T | O43592 | XPOT | Other | 0.381365432 | 0.935689209 | 29.73566732 | 29.95698501 | 29.92436532 | 29.44177889 | 29.38847737 | 29.53636269 | 30.19952565 | 30.20583023 | 29.73055871 |
| DNA repair protein XRCC1 | P18887 | XRCC1 | Other | 0.651051539 | 1.716356993 | 26.19383258 | 26.39706967 | 26.39990751 | 25.28887424 | 25.69117717 | 25.87946846 | 26.80980345 | 26.66356636 | 26.40494963 |
| DNA repair protein XRCC4 | Q13426;Q13426-3;Q13426-2 | XRCC4 | Other | -0.443608026 | 0.592409516 | 24.5869204 | 24.69842871 | 24.73753165 | 24.42668807 | 24.86438567 | 23.96868718 | 24.45243079 | 24.31934467 | 23.53872131 |
| X-ray repair cross-complementing pro | P13010 | XRCC5 | Extracellular or secreted | -0.23891008 | 0.36391873 | 33.1342281 | 33.18482515 | 33.11858034 | 32.80276152 | 32.87840176 | 32.6123676 | 33.01954515 | 32.88974062 | 32.23956623 |
| X-ray repair cross-complementing pro | P12956;P12956-2 | XRCC6 | Other | -0.394506438 | 0.555607324 | 33.03698606 | 33.01947887 | 32.98835867 | 32.55513003 | 32.65710525 | 32.4811848 | 32.7144849 | 32.61269715 | 31.85842048 |
| 5-3 exoribonuclease 2 | Q9H0D6;Q9H0D6-2 | XRN2 | Other | -0.926627342 | 2.560775164 | 29.80562777 | 29.74810428 | 29.67796532 | 28.82341302 | 28.90667136 | 29.59022636 | 28.39046728 | 28.50335913 | 28.60229562 |
| Xylulose kinase | O75191;O75191-2 | XYLB | Extracellular or secreted | 0.552001402 | 1.293662897 | 24.30938642 | 24.33626737 | 24.75200285 | 24.50039114 | 24.74601645 | 24.73334805 | 25.42451129 | 25.02677568 | 24.89342338 |
| YY1-associated factor 2 | Q8IY57-3;Q8IY57;Q8IY57-5 | YAF2 | Other | 0.069661902 | 0.110626399 | 22.06782559 | 22.66804356 | 22.56395313 | 22.4741238 | 22.83596622 | 23.29389818 | 23.02956695 | 22.65554208 | 22.47578191 |
| Transcriptional coactivator YAP1 | P46937-5;P46937-3;P46937-( | YAP1 | Other | 1.341622484 | 0.800995557 | 23.09603307 | 19.43277392 | 22.78388708 | 24.25277514 | 24.69773966 | 23.00437447 | 24.96062384 | 23.71187867 | 23.98615661 |
| Tyrosine--tRNA ligase, cytoplasmic;Ty | P54577 | YARS | Extracellular or secreted | 0.473767372 | 0.711015998 | 28.76376238 | 28.87276293 | 28.86496416 | 28.8363537 | 28.80335063 | 28.40157661 | 29.47448435 | 29.49228352 | 28.72591945 |
| Nuclease-sensitive element-binding p | P67809 | YBX1 | Extracellular or secreted | 0.64539382 | 1.895499002 | 31.48106448 | 31.49290425 | 31.52654625 | 31.8991551 | 31.84370943 | 32.23349339 | 32.49688489 | 32.54153217 | 32.13620084 |
| Y-box-binding protein 3 | P16989;P16989-3 | YBX3 | Other | 1.468202138 | 3.343717086 | 28.65765046 | 28.25714387 | 28.38339785 | 29.16966276 | 29.18160638 | 29.49615633 | 30.36723883 | 30.42582416 | 30.18435225 |
| Y-box-binding protein 3 | P16989-2 | YBX3 | Other | 1.62897073 | 2.031302294 | 24.09079408 | 23.38811191 | 23.43402648 | 25.60189907 | 24.69773966 | 25.58545979 | 26.15848164 | 26.23485616 | 25.89258988 |
| YEATS domain-containing protein 4 | O95619 | YEATS4 | Other | -0.229164128 | 0.432843651 | 24.30959459 | 24.79332547 | 24.34712444 | 23.51508765 | 24.03573746 | 23.60388075 | 23.91095375 | 24.07993329 | 23.62399577 |
| Tyrosine-protein kinase Yes | P07947 | YES1 | Extracellular or secreted | 1.045983185 | 3.64401785 | 27.63733917 | 27.56710532 | 27.71246917 | 27.55171507 | 27.7498951 | 28.33568829 | 28.71695564 | 28.98163487 | 28.7164651 |
| Protein YIPF3;Protein YIPF3, 36 kDa fo | Q9GZM5 | YIPF3 | Extracellular or secreted | -0.38596345 | 0.652782398 | 21.98733833 | 23.22418837 | 23.20953844 | 22.29007736 | 22.36113098 | 21.76320264 | 22.00199225 | 21.896189 | 22.36166647 |
| Synaptobrevin homolog YKT6 | O15498;O15498-2 | YKT6 | Extracellular or secreted | -0.041417945 | 0.105504162 | 26.77102433 | 26.69272084 | 26.46757483 | 26.77693316 | 26.83593224 | 26.29403846 | 26.55815372 | 26.81078421 | 26.42592017 |
| YLP motif-containing protein 1 | P49750;P49750-1;P49750-3 | YLPM1 | Other | -0.255470366 | 1.365242377 | 27.58498692 | 27.3426558 | 27.23388762 | 27.59434374 | 27.57830727 | 27.06270529 | 27.12494608 | 27.07091527 | 27.23617087 |
| ATP-dependent zinc metalloprotease | Q96TA2-3;Q96TA2-2;Q96TA2 | YME1L1 | Other | 0.062829826 | 0.055793513 | 27.07673651 | 27.03149492 | 27.22903514 | 27.97951041 | 27.78699772 | 26.60540763 | 27.59213501 | 27.74868014 | 26.70226549 |
| YTH domain-containing protein 1 | Q96MU7-2;Q96MU7 | YTHDC1 | Other | -1.607628306 | 1.773208959 | 24.40021705 | 24.74046841 | 24.78511518 | 25.73451138 | 25.00581709 | 23.87323832 | 22.74632405 | 22.87505862 | 23.82541613 |
| YTH domain-containing family protein | Q9BYJ9 | YTHDF1 | Other | -0.11809184 | 0.070375124 | 22.12691617 | 22.17508533 | 22.39068538 | 23.2431734 | 22.06354662 | 21.63436944 | 22.65890094 | 22.70182677 | 21.10188495 |
| YTH domain-containing family protein | Q9Y5A9;Q9Y5A9-2 | YTHDF2 | Other | -0.506824162 | 0.924721677 | 27.04478981 | 26.74355329 | 26.46724831 | 27.28665533 | 27.59541127 | 26.78805635 | 26.80279618 | 26.54470237 | 26.09488615 |
| YTH domain-containing family protein | Q7Z739 | YTHDF3 | Other | -0.200827084 | 1.570602562 | 26.4732073 | 26.9509369 | 26.7050375 | 26.59818316 | 26.68939191 | 26.61133014 | 26.44714226 | 26.49242106 | 26.47199887 |
| 14-3-3 protein beta/alpha;14-3-3 prot | P31946-2;P31946 | YWHAB | Extracellular or secreted | 1.457297829 | 3.171074556 | 31.36531048 | 31.4432224 | 31.45238366 | 31.14534111 | 31.52248304 | 32.39695641 | 32.96914984 | 32.76633587 | 33.29925633 |
| 14-3-3 protein epsilon | P62258;P62258-2 | YWHAE | Extracellular or secreted | 0.332157955 | 1.275005052 | 34.38071721 | 34.45280093 | 34.36208646 | 34.17771534 | 34.24168427 | 33.57068313 | 34.53258486 | 34.54455222 | 34.51218046 |
| 14-3-3 protein gamma;14-3-3 protein | P61981 | YWHAG | Extracellular or secreted | 0.169748238 | 0.396598968 | 31.3969309 | 31.25338395 | 31.28454994 | 31.15617325 | 31.35126546 | 30.83394368 | 31.47614669 | 31.59143071 | 31.07979091 |
| 14-3-3 protein eta | Q04917 | YWHAH | Extracellular or secreted | 1.163648164 | 4.84311953 | 30.45110654 | 30.20174707 | 30.23797209 | 30.00142108 | 30.08296572 | 30.43757855 | 31.37673865 | 31.41624044 | 31.40436093 |
| 14-3-3 protein theta | P27348 | YWHAQ | Extracellular or secreted | 0.960736674 | 1.386216605 | 31.58982469 | 31.63711457 | 31.57294299 | 31.35662708 | 31.45792109 | 31.52234273 | 32.69438659 | 32.69751763 | 32.05869237 |
| 14-3-3 protein zeta/delta | P63104;P63104-2 | YWHAZ | Extracellular or secreted | 1.092306945 | 4.331453447 | 33.53519491 | 33.46708697 | 33.37528917 | 33.2019369 | 33.48531784 | 33.60410441 | 34.43936321 | 34.51253371 | 34.65948902 |
| Transcriptional repressor protein YY1 | P25490 | YY1 | Other | -0.105813968 | 0.303186802 | 24.42930866 | 23.87239308 | 24.07977057 | 23.77976584 | 24.13680467 | 24.69885258 | 24.09353685 | 24.10261523 | 23.98485371 |
| Mitogen-activated protein kinase kina | Q9NYL2-2;Q9NYL2-3;Q9NYL2 | ZAK | Other | -0.239512657 | 0.45783176 | 24.22751195 | 24.28714872 | 24.25551502 | 23.64877887 | 24.13029844 | 23.66177375 | 23.66795697 | 23.5535618 | 24.16545663 |
| Zinc finger and BTB domain-containin | O15156;O15156-2 | ZBTB7B | Other | -0.262816967 | 0.174277157 | 23.78346749 | 21.60449194 | 22.25755934 | 22.54453435 | 22.88908062 | 23.98450608 | 21.71338987 | 22.85602147 | 23.17395767 |
| Zinc finger CCCH domain-containing p | O75152 | ZC3H11A | Other | -0.757296659 | 0.33081794 | 23.22711529 | 23.07439041 | 22.38318852 | 22.82590131 | 23.60345911 | 24.00487469 | 23.18537635 | 20.74678271 | 23.35541562 |
| Zinc finger CCCH domain-containing p | Q5T200;Q5T200-2 | ZC3H13 | Other | -1.67741653 | 2.361396897 | 26.92965122 | 26.7573542 | 26.73182138 | 26.70925129 | 26.51270888 | 26.05655834 | 25.08820788 | 24.51520799 | 25.21300719 |
| Zinc finger CCCH domain-containing p | Q6PJT7-5;Q6PJT7-9;Q6PJT7-2 | ZC3H14 | Other | -0.753897037 | 1.372487039 | 25.92689479 | 25.93127549 | 25.9053635 | 25.78511518 | 25.60002813 | 24.7508791 | 24.67233426 | 24.70065264 | 25.31510007 |
| Zinc finger CCCH domain-containing p | Q8WU90;Q8WU90-2 | ZC3H15 | Other | -0.53358215 | 0.678727945 | 25.90692362 | 25.63994896 | 25.72197572 | 26.12803792 | 26.20998086 | 26.49730282 | 25.42008358 | 24.99683928 | 26.03441563 |
| Zinc finger CCCH domain-containing p | Q86VM9;Q86VM9-2 | ZC3H18 | Other | -1.049716976 | 1.423567934 | 28.05025014 | 28.25660413 | 28.27012669 | 28.25484856 | 28.18271956 | 27.82183285 | 26.82031182 | 26.88634649 | 27.56238172 |
| Zinc finger CCCH domain-containing p | Q9UPT8 | ZC3H4 | Other | 0.220512234 | 0.379662306 | 26.09444323 | 25.9682697 | 25.91458495 | 26.47422903 | 26.02799911 | 26.36941217 | 26.19439632 | 26.10679374 | 26.78481574 |
| Zinc finger CCCH-type antiviral protei | Q7Z2W4;Q7Z2W4-2;Q7Z2W4 | ZC3HAV1 | Other | 0.783083 | 0.97091904 | 24.01214096 | 24.81279268 | 25.05858271 | 25.73402031 | 25.41341643 | 26.62839703 | 25.72161074 | 26.47478603 | 25.98252729 |
| Zinc finger CCCH-type antiviral protei | Q96H79 | ZC3HAV1L | Other | 2.23899109 | 1.608752281 | 20.6354738 | 21.93623491 | 21.63572529 | 24.58629051 | 24.55250681 | 21.07526338 | 24.41189833 | 24.75521604 | 24.76060626 |
| Nuclear-interacting partner of ALK | Q86WB0;Q86WB0-2;Q86WB( | ZC3HC1 | Other | 1.771064703 | 1.762507594 | 23.77263595 | 24.314789 | 24.9791942 | 24.6164434 | 25.25695498 | 23.21038597 | 26.01337686 | 26.84098167 | 25.53403733 |
| Zinc finger CCHC domain-containing p | Q6NZY4;Q6NZY4-2 | ZCCHC8 | Other | 0.391071879 | 0.878653685 | 23.46425956 | 23.61672382 | 23.08487131 | 23.32072234 | 23.64504444 | 23.19189538 | 23.91743327 | 23.98058937 | 23.43695142 |
| Probable palmitoyltransferase ZDHHC | Q5W0Z9-4;Q5W0Z9-3;Q5W0 | ZDHHC20 | Extracellular or secreted | 2.143770845 | 3.617131782 | 21.41405702 | 20.952399 | 20.4396924 | 21.88837402 | 21.9202095 | 21.45432815 | 23.46425956 | 23.51653107 | 23.48505195 |
| Palmitoyltransferase ZDHHC5 | Q9C0B5-2;Q9C0B5 | ZDHHC5 | Extracellular or secreted | 1.273897531 | 3.98790356 | 25.15842386 | 24.67206444 | 24.74668284 | 25.10485594 | 24.84090966 | 24.50239577 | 25.92143419 | 26.23890601 | 26.17401864 |
| AN1-type zinc finger protein 1 | Q8TCF1;Q8TCF1-2;Q8TCF1-4; | ZFAND1 | Other | 0.75644402 | 1.087576838 | 23.2209014 | 23.05619448 | 22.94177551 | 24.01724816 | 24.25573111 | 22.74326565 | 24.28411525 | 24.47109964 | 23.63167532 |
| AN1-type zinc finger protein 5 | O76080 | ZFAND5 | Other | 0.827220011 | 0.786172889 | 21.09569584 | 21.90389796 | 22.99891473 | 23.29193285 | 23.44480663 | 22.40792433 | 23.68528006 | 23.743656 | 22.62431013 |
| AN1-type zinc finger protein 6 | Q6FIF0-2;Q6FIF0 | ZFAND6 | Other | 0.016645104 | 0.008939767 | 25.75521604 | 26.14024189 | 26.01288689 | 25.73678363 | 26.21602718 | 25.40373137 | 26.57600172 | 26.34450248 | 24.76187461 |
| E3 ubiquitin-protein ligase ZFP91 | Q96JP5-2;Q96JP5 | ZFP91 | Other | -0.743994614 | 1.722946242 | 24.95548922 | 24.93215455 | 24.98446262 | 24.88609061 | 25.5535618 | 25.08492806 | 24.27703591 | 24.07422706 | 24.61509661 |
| Zinc finger protein-like 1 | O95159 | ZFPL1 | Other | -0.770741339 | 2.982494487 | 27.19648026 | 27.05320569 | 26.97858339 | 26.42027637 | 26.52853352 | 26.747144 | 26.00262327 | 25.99681774 | 26.15044659 |
| Zinc finger RNA-binding protein | Q96KR1 | ZFR | Other | -0.761946558 | 2.359441543 | 28.11182585 | 28.21122558 | 28.1793537 | 27.5593213 | 27.91416292 | 28.75808528 | 27.31414937 | 27.32130745 | 27.44569083 |
| Abscission/NoCut checkpoint regulato | Q96K21-2;Q96K21;Q96K21-4 | ZFYVE19 | Other | 1.641519981 | 2.329822432 | 21.61547826 | 21.43903627 | 20.48513576 | 21.84907477 | 22.77523235 | 22.66809315 | 23.41931215 | 23.5721895 | 23.34908357 |
| Zinc finger FYVE domain-containing pr | Q9BQ24;Q9BQ24-2 | ZFYVE21 | Extracellular or secreted | 0.609492065 | 0.380865503 | 21.87500527 | 20.74139115 | 21.65555167 | 22.16506231 | 21.45288199 | 22.81135274 | 21.24260254 | 22.81614695 | 23.12034927 |
| Zymogen granule protein 16 homolog | Q96DA0 | ZG16B | Extracellular or secreted | -0.929572195 | 0.398452459 | 24.78971805 | 22.59859026 | 21.79158593 | 23.05055522 | 21.80758975 | 22.73914631 | 21.04806641 | 20.99403571 | 23.55777405 |
| CAAX prenyl protease 1 homolog | O75844 | ZMPSTE24 | Extracellular or secreted | -0.514156024 | 0.095854698 | 20.52945724 | 23.12275218 | 22.98163487 | 21.89559747 | 21.94356506 | 27.23963451 | 20.11629319 | 21.57644006 | 25.62111934 |
| Zinc finger MYM-type protein 2 | Q9UBW7 | ZMYM2 | Other | 0.876620789 | 0.607585686 | 20.36623843 | 20.90239572 | 20.73538179 | 22.48029074 | 21.5650048 | 24.50754642 | 22.5412999 | 22.19675826 | 23.17023316 |
| Zinc finger MYM-type protein 3 | Q14202-3;Q14202-2;Q14202 | ZMYM3 | Other | 0.652406246 | 0.687759496 | 20.98386744 | 23.22343562 | 20.80695547 | 20.98592414 | 21.23992793 | 20.57424901 | 21.52406149 | 22.39013431 | 21.95020274 |
| Protein kinase C-binding protein 1 | Q9ULU4-17;Q9ULU4-8;Q9ULU | ZMYND8 | Other | -0.869475964 | 0.918698972 | 24.25738669 | 23.99057767 | 23.87633333 | 23.77936509 | 23.43224317 | 24.80099585 | 22.62546806 | 22.98193095 | 23.85262399 |
| Zinc finger protein 185 | O15231-3;O15231-2 | ZNF185 | Extracellular or secreted | -0.325235439 | 0.397580633 | 26.23222314 | 26.00695145 | 26.06782149 | 25.9127134 | 25.93952649 | 25.41622232 | 25.44215078 | 25.17797666 | 26.19189538 |
| BUB3-interacting and GLEBS motif-co | O43670-2 | ZNF207 | Extracellular or secreted | -0.377644647 | 0.844022434 | 28.33573088 | 28.33972845 | 28.32225345 | 28.38112994 | 28.21405471 | 28.10699337 | 27.77580356 | 28.23498403 | 27.70622387 |
| Zinc finger protein 217 | O75362 | ZNF217 | Other | -0.427466721 | 0.941346864 | 22.07551702 | 22.20999944 | 22.57544784 | 22.59897553 | 22.38345543 | 22.07747423 | 22.20123014 | 21.88198235 | 21.59482209 |
| DBIRD complex subunit ZNF326 | Q5BKZ1;Q5BKZ1-3 | ZNF326 | Other | -1.266554805 | 2.352065897 | 27.75557262 | 28.01547333 | 28.10594502 | 27.1525756 | 27.16250166 | 26.52260464 | 26.24517723 | 26.45723031 | 25.85526448 |
| Zinc finger protein 428 | Q96B54 | ZNF428 | Other | -0.052999153 | 0.057436002 | 25.9789979 | 26.08050267 | 25.84184552 | 26.15174372 | 26.07598209 | 25.17405675 | 26.08679159 | 26.12813627 | 25.27763901 |
| Zinc finger protein 483 | Q8TF39-2;Q8TF39 | ZNF483 | Other | -0.952699496 | 0.418268494 | 22.49382321 | 20.48315841 | 20.18131728 | 24.89901388 | 21.7856808 | 26.17216912 | 21.99056605 | 21.38784182 | 21.771075 |
| Zinc finger protein 511 | Q8NB15-2;Q8NB15 | ZNF511 | Extracellular or secreted | 0.057852196 | 0.074750947 | 21.37443247 | 21.04919713 | 22.08608697 | 21.18861671 | 21.50913597 | 22.13877426 | 21.85991994 | 21.21193106 | 21.77482734 |
| Zinc finger protein 622 | Q969S3 | ZNF622 | Other | -2.431407604 | 0.774230324 | 24.64047311 | 24.2073951 | 23.82599833 | 24.62215328 | 24.88716036 | 23.56732296 | 19.56863396 | 23.36024698 | 22.65214782 |
| Zinc finger protein 638 | Q14966;Q14966-3;Q14966-5 | ZNF638 | Other | 0.190347293 | 0.055963439 | 28.88518303 | 29.7156308 | 28.97514274 | 22.09701076 | 27.83104634 | 28.66610885 | 28.51126163 | 27.83086506 | 27.31397645 |
| Zinc finger protein 687 | Q8N1G0-2;Q8N1G0 | ZNF687 | Other | -1.399809957 | 0.593884702 | 24.03489835 | 23.89254356 | 24.01826743 | 24.45356153 | 24.52963596 | 24.05382714 | 21.01216083 | 23.38297752 | 23.89679875 |
| Zinc finger protein 706 | Q9Y5V0 | ZNF706 | Other | 0.980003212 | 0.582912854 | 25.12561621 | 24.34252017 | 24.45393824 | 25.8586828 | 25.95193742 | 23.51291982 | 26.75786281 | 26.15458559 | 24.65036855 |
| Zinc finger protein 787 | Q6DD87 | ZNF787 | Other | -0.271387164 | 0.327728652 | 22.81493523 | 22.09188319 | 23.2598306 | 22.07172552 | 23.39923932 | 24.13578756 | 22.68752434 | 22.91990788 | 22.465107 |
| Putative uncharacterized zinc finger p | B7Z6K7 | ZNF814 | Other | -2.057557048 | 2.183178831 | 23.45813823 | 23.34502726 | 20.95890949 | 23.55004217 | 23.96675257 | 24.12943396 | 20.9805938 | 20.87373365 | 21.67715324 |
| Zinc finger protein 830 | Q96NB3 | ZNF830 | Other | -0.317574262 | 0.270640551 | 25.20966486 | 25.30577343 | 24.66084942 | 25.76248302 | 25.86719332 | 24.04826601 | 25.2360066 | 25.17276046 | 24.06562518 |
| Zinc finger protein ZPR1 | O75312 | ZPR1 | Other | 0.528729907 | 0.704893601 | 26.73647398 | 26.6800298 | 26.57017825 | 27.2266487 | 27.25718884 | 26.25736871 | 27.59063684 | 27.61645742 | 26.7430396 |
| Zinc finger Ran-binding domain-conta | O95218-2;O95218 | ZRANB2 | Other | -0.133573513 | 0.435089959 | 27.54251957 | 27.49434523 | 27.55735046 | 27.78855426 | 27.80790133 | 27.56499978 | 27.30382422 | 27.69544485 | 27.4778457 |
| Centromere/kinetochore protein zw1 | O43264;O43264-2 | ZW10 | Other | 0.127146343 | 0.211154719 | 26.14782941 | 26.17289396 | 26.43828475 | 25.58817934 | 25.83144507 | 25.19591733 | 26.09466471 | 26.24720725 | 25.726842 |
| Protein zwilch homolog | Q9H900;Q9H900-2 | ZWILCH | Other | 0.110646098 | 0.268035929 | 24.79441666 | 24.8676411 | 24.69837572 | 24.57605941 | 24.79243207 | 24.17721634 | 24.84278077 | 24.9371033 | 24.50512488 |
| Protein zyg-11 homolog B | Q9C0D3 | ZYG11B | Other | 0.263180992 | 0.668616154 | 22.80797746 | 23.03135188 | 22.9384231 | 23.13186889 | 23.56767112 | 22.4742476 | 23.4891012 | 23.19499365 | 23.08121815 |
| Zyxin | Q15942;Q15942-2 | ZYX | Extracellular or secreted | 1.155419973 | 3.970640383 | 27.96845648 | 27.73988926 | 27.9383154 | 28.4273037 | 28.53552866 | 28.274621 | 29.44164039 | 29.27930065 | 29.18737613 |
| Zinc finger ZZ-type and EF-hand doma | O43149 | ZZEF1 | Other | 0.513666418 | 0.250147022 | 25.09960882 | 25.05548303 | 25.08371142 | 23.3836368 | 25.32068791 | 24.43459921 | 25.79325105 | 26.1248475 | 23.8117643 |
| | P0DPI2;A0A0B4J2D5;P0DPI2-2;A0A0B4J2D5-2 | | NA | -0.374188482 | 0.93014589 | 30.0353809 | 29.82935353 | 29.80134746 | 29.74270239 | 29.61750872 | 29.80863793 | 29.47163847 | 29.67075848 | 29.15250307 |
| | P09493-3;P09493-8;P09493-9;P09493-10;P094 | | NA | 0.668920817 | 0.559304763 | 27.75244956 | 27.18999327 | 27.33381322 | 27.86466904 | 27.60151657 | 27.69112391 | 27.98576586 | 27.61463336 | 29.12314602 |
| | P09493-5 | | NA | 1.383053538 | 0.966104845 | 21.07863365 | 24.92347584 | 24.93553054 | 23.83113093 | 24.36245595 | 25.050219 | 25.24641008 | 24.77776098 | 26.21571251 |
| | P0DP25;P0DP24;P0DP23 | | NA | -1.069764581 | 1.188137569 | 30.36859235 | 30.50483705 | 30.86791205 | 30.29559376 | 30.2721976 | 27.8248701 | 29.12465033 | 29.09582186 | 28.63723551 |
| | P0DPB6 | | NA | 0.638862777 | 1.249342669 | 24.3850203 | 24.6937579 | 24.29607097 | 24.55285856 | 24.4523051 | 25.02407203 | 25.27070595 | 25.50418543 | 24.84373937 |
| | Q13491-4 | | NA | -0.131711836 | 0.098215242 | 23.86938374 | 23.7329342 | 23.62232088 | 22.43824633 | 23.43173325 | 21.02796316 | 23.24055553 | 22.88541114 | 22.54018859 |
| | Q6ZT62-2;Q6ZT62 | | NA | -0.314874027 | 0.596546636 | 25.67777383 | 24.86924252 | 25.11779178 | 25.67489503 | 25.48200755 | 24.59741609 | 24.639811 | 24.89919832 | 25.225932 |
| | Q86XK3-2 | | NA | 0.659422302 | 0.20070023 | 21.24796436 | 20.46081941 | 22.29568775 | 21.87057785 | 22.32430456 | 21.89779851 | 20.08851866 | 23.91177671 | 23.02654776 |
| | Q9UHC1-2 | | NA | 1.492671021 | 2.491813143 | 25.31972366 | 23.94752993 | 23.86721689 | 24.69343889 | 23.42358683 | 23.91442528 | 25.50109004 | 26.07093574 | 25.48894802 |
| | Q9UHX1-6 | | NA | -0.279943515 | 1.828872062 | 30.30791105 | 30.41094466 | 30.41518284 | 30.66610885 | 30.6853703 | 30.60514266 | 30.28482569 | 30.29570322 | 30.12497072 |
| | Q9UPN3-2;Q9UPN3-3 | | NA | 0.189529467 | 0.735478174 | 30.56310944 | 30.45189259 | 30.46051098 | 29.9139918 | 29.92309042 | 30.22492526 | 30.40628852 | 30.37927173 | 30.55178839 |
| | Q9UPN3-5 | | NA | -0.156227661 | 0.124051657 | 23.28996483 | 23.17148591 | 22.2541671 | 23.13472273 | 23.82405674 | 23.34109528 | 22.21129218 | 23.42046914 | 23.407302 |
